# Reduced NCOR2 expression accelerates androgen deprivation therapy failure in prostate cancer

**DOI:** 10.1101/2020.07.02.182758

**Authors:** Mark D Long, Prashant K Singh, Gerard Llimos, Justine J Jacobi, Aryn M Rowsam, Spencer R Rosario, Jason Kirk, Hayley C Affronti, Moray J Campbell, Dominic J Smiraglia

## Abstract

NCOR2 is frequently and significantly mutated in late stage androgen deprivation therapy resistant prostate cancer (ADT-RPCa). NCOR2 has been characterized as a transcriptional corepressor and has mechanistic links to DNA methylation, but its global functions and overall contributions to PCa progression remain enigmatic. In the current study, we utilize immunohistochemical staining of samples from over 700 PCa patients and reveal associations of reduced NCOR2 expression with correlates of aggressive primary PCa and recurrence in patients who received adjuvant androgen deprivation therapy. We mapped the dihydrotestosterone (DHT) dependent and independent effects of NCOR2 on the transcriptome, cistrome and DNA methylome in androgen sensitive (AS) and ADT-RPCa cells using the isogenic LNCaP and LNCaP-C4-2 (C4-2) cell models. Transcriptional profiling identified androgen dependent and independent regulatory roles of NCOR2, the latter of which was enhanced in the ADT-RPCa state and included neuronal differentiation. Interestingly, reduced expression of NCOR2 resulted in a striking global DNA hypermethylation pattern that significantly enriched at enhancer regions. ChIP-seq revealed that NCOR2 was more clearly associated with promoters in AS LNCaP cells, which was modestly enhanced by DHT treatment. However, in ADT-RPCa C4-2 cells, the NCOR2 cistrome was larger and more distal. Motif analyses and integration of large-scale public cistrome data revealed strong enrichment for FOXA1 in mediating NCOR2 binding, and included additional factors such as AR, E2F, TET2, MED1 and MBD2. Utilizing the CWR22 xenograft model, we demonstrate a direct role for NCOR2 in PCa progression as reduced NCOR2 expression attenuated the impact of ADT, and significantly accelerated recurrence of disease. Transcriptomic analyses in recurrent CWR22 tumors indicated NCOR2-dependent gene expression profiles during ADT that were enriched for neuroendocrine associated genes and also associated with worse survival in human patients with ADT-RPCa. DNA methylation profiles in CWR22 tumors with reduced NCOR2 expression recapitulated the hypermethylation observed *in vitro*, and further revealed that hypermethylation patterns are commonly associated with ADT-RPCa disease, which was also confirmed in human samples. These studies reveal robust roles for NCOR2 in regulating the PCa transcriptome and epigenome and underscore recent mutational studies linking NCOR2 loss of function to PCa disease progression.

## Introduction

Prostate Cancer (PCa) patients with advanced disease receive androgen deprivation therapy (ADT) but frequently experience treatment failure leading to ADT-resistant PCa (ADT-RPCa); mortality rates for ADT-RPCa remain stubbornly high. Recent clinical successes of the LATITUDE ^1^ and STAMPEDE ^2^ clinical trials suggests that ADT-combination therapies can deliver significant survival benefits and that next generation androgen receptor (AR) antagonists ^3,4^ may enhance ADT efficacy. Given these successes, there is enthusiasm to identify mechanisms that either augment or synergize with ADT to improve duration of ADT benefits, and to limit ADT failure.

ADT-RPCa drivers include genetic, epigenetic and metabolic oncogenic processes, which amplify and re-direct androgen receptor (AR) signaling (reviewed in ^5^). As a result AR signaling devolves from its normal control of cell growth and differentiation to favor growth promoting pathways ^6,7^. In this manner AR signaling is “re-wired”, often associated with distorted histone modifications and DNA methylation patterns that impact AR access to enhancer regions ^8,9^. These observations fit with the emerging concept of oncogenic enhancer addiction ^10,11^ acting as a driver of lineage plasticity ^12,13^. Understanding the mechanisms that reconfigure AR genomic interactions have potential to be exploited to augment ADT.

Alterations to a number of AR interacting proteins, such as pioneer factors and co-regulators, contribute to these changes in AR signaling. Of these co-regulators, Nuclear Receptor Corepressor 2/Silencing Mediator For Retinoid And Thyroid Hormone Receptors (NCOR2) has been identified as frequently altered in PCa and other cancers ^14,15^. Notably, NCOR2 is among the top 5 mutated corepressors in the recent SU2C study of 444 men with advanced PCa ^16,17^. The current study addresses the role of NCOR2 in determining the effectiveness of ADT.

NCOR2 binds nuclear hormone receptors (NR) ^18^, and allosterically interacts with histone deacetylases ^19,20^ to promote repressive histone marks such as H3K9me3. This can lead to direct epigenetic silencing, or acts as an histone mark that recruits the CpG methylation machinery and promotes increased DNA methylation ^21^. Similarly, NCOR2 interacts with KAISO ^22^, and with the lncRNA, SHARP, in both cases to trigger DNA methylation ^23^. Therefore, in PCa, we and others have reasoned that expression and mutation changes in NCOR2 disrupt its ability to regulate the epigenome and control gene expression. In this manner, altered NCOR2 functions may induce an onco-epigenomc that re-wires AR-genomic interactions and impacts the duration and success of ADT^14^.

There remain significant ambiguities on NCOR2’s functions. Whilst both NCOR2 and HDAC3 knockout mice are embryonically lethal ^24,25^, mice with mutant NCOR2 that cannot recruit HDACs are viable ^26^, suggesting HDAC-independent roles for NCOR2. Furthermore NCOR2 significantly accumulates at open chromatin and actively transcribed genomic regions ^27^, and can actively enhance transcription by estrogen receptor beta ^28^ and AR ^29^. This suggests that NCOR2 is not an obligate co-repressor. It also remains unclear where NCOR2 interacts with the genome, given the diversity of interacting transcription factors ^30,31^ and the presence of a MYB-like SANT domain that may allow for direct binding to histone tails ^32^.

Indeed, establishing NCOR2-dependent cistrome-transcriptome is obscured by the distribution of enhancers across the genome and their complex relationship to genes. On average, every human gene is regulated by ∼ 6 enhancers ^33^; the median distance between gene and enhancer is 158 kb^34^; and genomic topology (e.g. topologically associated domains (TADs)), impacts these relationships in a non-predictable manner (reviewed in ^35^).

Furthermore there are significant ambiguities concerning the impact of NCOR2 on CpG methylation patterns. Clearly, in various cancers elevated DNA methylation at promoter regions of high CpG density results in transcriptional silencing (reviewed in ^36^), but in low density CpG regions, for example within enhancers, DNA methylation may selectively recruit transcription factor binding ^37^, and participate in allelic-specific gene regulation ^38^. Therefore, it is unclear what are the genomic contexts and consequences of NCOR2 interactions with CpG methylation.

Thus, it is unknown how and where NCOR2 is recruited to the genome, what effects it may confer to gene expression, and what functional relationships exist between NCOR2-genomic interactions and DNA methylation. Furthermore, it is still unclear to what extent these relationships impact transcriptional signals that are essential for the success of ADT in PCa patients.

We sought to address these ambiguities and define how NCOR2 drives the onco-epigenome and determines androgen dependent and independent transcriptional signaling in progression to ADT-RPCa. Specifically, we exploited a large tissue microarray spanning over 700 PCa patients with extensive clinical follow-up to identify relationships between NCOR2 expression and biochemical recurrence. We then utilized isogenic models of androgen sensitive (LNCaP) and ADT-resistant (C4-2) PCa cells to map the global regulatory functions of NCOR2 with regards to its cistrome (ChIP-seq), transcriptome (RNA-seq, miRNA-seq), and DNA methylome (EPIC array). Finally, the CWR22 xenograft model of PCa progression was utilized to examine the role of NCOR2 in determining response to ADT *in vivo*, and identified molecular features compared to those observed in human patients with ADT-RPCa.

## Methods

### Cell culture and materials

LNCaP cells were derived from a 50-year old male with PCa who responded briefly to androgen deprivation therapy ^39^ and serve as a model for androgen sensitive PCa. The C4-2 variant was derived in vivo from LNCaP using multiple rounds of selection in castrated mice and has metastatic potential, thus serving as an isogenic ADT resistant cell line model of aggressive PCa ^40^.

All cells were maintained at 37°C and 5.0% CO2 using a cell culture incubator with UV contamination control (Sanyo). LNCaP and LNCaP-C4-2 cells were regularly maintained in RPMI 1640 Medium containing 10% FBS. All media was supplemented with 100 U/mL Penicillin-Streptomycin. Dihydrotestosterone (DHT, D-073-1ML, Sigma-Aldrich) was kept as 10mM EtOH stocks, and diluted to 1000x stocks prior to treatments. Prior to androgen treatment, cells were serum starved using charcoal stripped FBS (10%) for 72 hours.

Cell lines were authenticated by STR profiling and confirmed mycoplasma free by RT-PCR in the RPCCC Genomics Shared Resource.

### Stable knockdown of NCOR2

Knockdown of NCOR2 in LNCaP and C4-2 cells was achieved by stable selection after transduction with lentiviral shRNA constructs targeting *NCOR2*. Two targeting constructs (V2LHS-251658 (shNCOR2-A), V2LHS-196739 (shNCOR2-B)) and one non-silencing control construct were selected from the V2LHS pGIPZ based lentiviral shRNA library (Thermo Fisher Scientific). Viral packaging and cellular infection (RPCCC shRNA Resource) yielded pGIPZ containing cells, which were maintained in media supplemented with puromycin (2µg/mL). For xenograft studies, NCOR2 targeting shRNA (V2LHS-251658) or non-targeting shRNA control were introduced to digested CWR22 tissue under puromycin selection for 24 hours prior to implantation.

### RT-qPCR

Total RNA was isolated via TRIzol® reagent (Thermo Fisher Scientific) for candidate mRNA detection by use of the AllPrep DNA/RNA/miRNA Universal Kit (Qiagen), following manufacturer’s protocols. Complementary DNA (cDNA) was prepared using iScriptTM cDNA Synthesis Kit (Bio-Rad), following manufacturer’s protocols. Relative gene expression was subsequently quantified via Applied Biosystems 7300 Real-Time PCR System (Applied Biosystems), for both TaqMan and SYBR Green (Thermo Fisher Scientific) applications. All targets were detected using either pre-designed TaqMan Gene Expression Assays (Thermo Fisher Scientific; *AR, NCOR2*, miR-96-5p, miR-125b-5p, miR-193b-3p, miR-10a-5p, miR126-3p, miR-200a-3p, miR-22-3p, let-7e-5p, FOXA1, GAPDH, KRT8), pre-designed PrimeTime qPCR primers (IDT; *KRT18, DENND1B, TMPRSS2, HERC3, CXCR7, FOS, KLK3*) or custom designed qPCR primers (IDT) using a final primer concentration of 500nM. All primers for use with SYBR Green application were tested for specificity by melting curve analysis with subsequent product visualization on agarose gel. All RT-qPCR experiments were performed in biological triplicates, with at least technical duplicates. Fold changes were determined using the 2-ΔΔCt method as the difference between experimental group and respective control group. Significance of experimental comparisons was performed using Student’s t-test.

### Immunoblotting

Total cellular protein was isolated from exponentially growing cells for determination of target protein expression. Cells were harvested, then washed in ice cold PBS before lysing in ice cold RIPA buffer (50mM Tris-HCl pH 7.4, 150mM NaCl, 1% v/v Triton X-100, 1mM EDTA pH 8.0, 0.5% w/v sodium deoxychlorate, 0.1% w/v SDS) containing 1x cOmplete Mini Protease Inhibitor Tablets (Roche). Protein concentrations were quantified using the DC Protein Assay (Bio-Rad), following manufacturer’s protocols. Equal amounts of proteins (30-60µg) were separated via SDS polyacrylamide gel electrophoresis (SDS-PAGE) using precast 10% polyacrylamide gels (Mini-Protean TGX, Bio-Rad). Proteins were transferred onto polyvinylidene fluoride (PVDF) membrane (Roche) for 80V for 1.5 hours. Post transfer, membranes were blocked with 5% non-fat dry milk (NFDM) for 1 hour at room temperature. Blocked membranes were probed with primary antibody against NCOR2 (ab24551, Abcam; 62370, Cell Signaling), FOXA1 (ab23738, Abcam), AR (PG-21, Millipore), TET1 (GTX124207, GeneTex), DNMT1 (NB100-392, Novus Biologicals), IgG (2729S, Cell Signaling), TBP (8515S, Cell Signaling), GAPDH (2118, Cell Signaling), β–Actin (ab8227, Abcam), H3K9me3 (13969, Cell Signaling), H3K27me3 (9733, Cell Signaling), H3 (9715, Cell Signaling), either overnight at 4°C or for 3 hours at room temperature. Primary antibody was detected after probing for 1 hour with HRP-linked rabbit anti-mouse IgG (P0161, Dako) or goat anti-rabbit IgG (P0448, Dako) secondary antibody at room temperature using ECL Western Blotting substrate (Pierce). Signal and quantification was performed using the ChemiDoc XRS+ system (Bio-Rad).

### Cell viability

Bioluminescent detection of cellular ATP as a measure of cell viability was undertaken using ViaLight® Plus Kit (Lonza Inc.) reagents. Cells were plated at optimal seeding density to ensure exponential growth (4×10^3^ cells per well) in 96-well, white-walled plates. Wells were dosed with agents to a final volume of 100 µl. Dosing occurred at the beginning of the experiment, and cells were incubated for up to 120 hours. Luminescence was detected with Synergy™2 multi-mode microplate reader (BioTek® Instruments). Each experiment was performed in at least triplicate wells in triplicate experiments.

### RNA-sequencing

RNA was extracted from LNCaP and C4-2 cells in the presence of DHT (10nM, 6hr) or EtOH. RNA was extracted at the indicated time points in the CWR22 xenograft model. To profile global transcriptional patterns, a minimum of biological triplicate samples (cell line studies; n = 3, xenograft studies; n = 5-6) per experimental condition were analyzed by RNA-seq. Sequencing was performed at the RPCCC Genomics Shared Resource core facility, and sequencing libraries prepared with the TruSeq Stranded Total RNA kit (Illumina Inc), from 1ug total RNA. Quasi-alignment of raw sequence reads to the human transcriptome (hg19) and subsequent transcript abundance estimation was performed via *Salmon* ^41^. For CWR22 samples, alignments were first filtered to remove and reads aligning to mouse (GRCm38). Transcript abundance estimates were normalized and differentially expressed genes (DEGs) were identified using a standard *DESeq2* ^42^ pipeline. For cell line studies, final DEG determination was called after combining samples from multiple NCOR2 targeting shRNA into a single group. Transcriptional regulator analysis on DEGs was performed using LISA ^43^.

### Small RNA-seq

Cell lines were treated as RNA-Seq. Sequencing was performed at the RPCCC Genomics Shared Resource core facility. Sequencing libraries were prepared with the TruSeq Small RNA kit (Illumina Inc), from 1ug total RNA. Following manufacturer’s protocols, ligation of 5’ and 3’ RNA adapters to the mature miRNAs 5’-phosphate and 3’-hydroxyl groups, respectively was undertaken. Following cDNA synthesis, the cDNA was amplified with 11-13 PCR cycles using a universal primer and a primer containing one of 48 index sequences, which allowed pooling of libraries and multiplex sequencing. Prior to pooling, each individual sample’s amplified cDNA construct was visualized on a DNA-HS Bioanalyzer DNA chip (Agilent Technologies) for mature miRNA and other small RNA products (140-150bp). Successful constructs were purified using a Pippen prep (Sage Inc.), using 125 – 160 bp product size settings with separation on a 3% agarose gel. The purified samples were validated for size, purity and concentration using a DNA-HS Bioanalyzer chip. Validated libraries were pooled at equal molar to a final concentration of 10nM in Tris-HCI 10 mM, pH 8.5, before 50 cycle sequencing on a MiSeq (Illumina, Inc.). Fastq files were aligned to the genome (hg19) using *bowtie2*. Expression counts were called against the miRbase consensus miRnome using *featureCounts* and A standard *DESeq2* ^42^ pipeline determined differentially expressed miRNA.

### DNA methylation profiling

Cell lines were treated as for RNA-Seq. DNA was extracted from all samples using the AllPrep DNA/RNA/miRNA Universal Kit (Qiagen), following manufacturer’s protocol. DNA methylation profiles were obtained using the Infinium MethylationEPIC BeadChip (EPIC array) platform ^44^, performed in the RPCCC Genomics Shared Resource. Data processing and quantification was accomplished using *ChAMP* ^45^. Briefly, detectible beta values for all probed CpG sites were initially compiled and filtered to remove those associated with multiple alignments and known SNPs, leaving reliable information for 791,398 CpG sites. To adjust for probe design bias (Infinium Type-I, Type-II), a beta-mixture quantile normalization method (BMIQ) was employed ^46^. Additionally, cross-array batch effect was corrected using *ComBat* ^47^. Differentially methylated Positions (DMPs) were determined using *ChAMP* and subsequently differentially methylated regions (DMR) identified using *DMRcate* ^48^.

### Chromatin Immunoprecipitation

ChIP was performed in LNCaP and C4-2 cells in the presence of DHT (10nM, 1hr) or EtOH in triplicate independent experiments. Briefly, approximately 20×10^6^ cells were crosslinked with 1% formaldehyde solution, quenched with glycine (0.125 M) and harvested in cold PBS. Sonication of crosslinked chromatin was performed using a Bioruptor® UCD-200™Sonicator (Diagenode) with optimized cycles for each cell type. Immunoprecipitation of sonicated material was performed with antibodies against NCOR2 (ab24551, Abcam) or IgG (sc-2027x, santa cruz) for 16 hours, and antibody/bead complexes isolated with Magna ChIPTM Protein A+G magnetic beads (Millipore). Complexes were washed, reverse crosslinked, and treated sequentially with RNase and proteinase K prior to DNA isolation. Sequencing (75bp single end, 49.1×10^6^, 50.9×10^6^ average reads/sample in LNCaP, C4-2 respectively) was performed at the RPCCC Genomics Shared Resource core facility. The NCOR2 cistrome was analyzed with *Rsubread/csaw* ^49^, along with TF motif analyses (*MotifDb*). Peak density plots were performed using the annotatePeaks.pl tool available from the *HOMER* (Hypergeometric Optimization of Motif EnRichment) suite v4.10. In order to find potential transcription factor binding enrichment within NCOR2 cistromes, we utilized *GIGGLE* ^50^ to query the complete human transcription factor ChIP-seq dataset collection (10,361 and 10,031 datasets across 1,111 transcription factors and 75 histone marks, respectively) in Cistrome DB ^51^. Prostate specific filtering limited analysis to 681 datasets across 74 TFs and 238 datasets across 19 HMs. For each query dataset, we determined the overlap of each NCOR2 cistrome. Putative co-enriched factors were identified by assessment of the number of time a given factor was observed in the top 200 most enriched datasets relative to the total number of datasets for that factor in the complete Cistrome DB (> 1.2 FC enrichment over background). For prostate specific analysis, overlaps across datasets were averaged for each factor.

### CWR22 model of PCa progression

All animal experiments were carried out at the Department of Laboratory Animal Research at RPCCC in accordance with an Institutional Animal Care and Use Committee approved protocol. Male Athymic Nude Balb/c mice were purchased from Harlan at approximately 2 months of age. Mice were allowed to reach approximately 3 months of age at which point they were surgically castrated and implanted with silastic tubing containing 12.5 mg of testosterone for sustained release 2 weeks prior to xenograft implantation. 1×10^6^ CWR22 cells in a 1:1 mix of media to matrigel were injected subcutaneously on the right flank as previously described ^52^. Total initial cohort size was 65 xenografts per group (shCTL, shNCOR2; V2LHS-251658), with 50 animals per group designated for completion to recurrence. Tumor volumes were calculated from caliper measurements using the formula (length^2^ x width x 0.5234). Once tumors reached approximately 0.3 cm^3^ in size (two consecutive measurements > 0.3 cm^3^), androgen withdrawal was achieved by removal of the silastic tubing and tumor volumes were followed for a maximum of 336 days. Mice designated in recurrence group were sacrificed once tumors reached approximately 1.0 cm^3^, or if mice presented with ascites or were otherwise required by veterinary staff. At the time of sacrifice body and tumor weight were taken. Additionally, serum, tumor and in some cases liver, spleen and/or pancreas tissues with possible metastases were obtained and immediately flash frozen and kept at -80°C. A tumor regression response to androgen withdrawal was defined as tumors that achieved a 40% loss in tumor size relative to size at withdrawal. A tumor was considered to be recurrent following androgen withdrawal once the primary subcutaneous tumor had reached a size that was 200% that of the original size of the tumor at withdrawal. Presence of shRNA targeting construct in recurrent tumors was verified by Sanger sequencing for select animals (n=10 per group).

Approximately 30-50 milligrams of flash frozen tumor tissue was homogenized in 1 mL of TRIzol (for RNA) and 1 mL of Szak’s RIPA buffer containing 1x Protease Inhibitor (Mini Tablets) (for Protein) using a Polytron PT 2100 tissue homogenizer. Approximately 10 ug of extracted RNA was then DNase treated using the TURBO DNA-free™kit. RNA from tissue samples utilized for RNA-seq was isolated using the AllPrep DNA/RNA/miRNA Universal Kit (Qiagen) as per manufacturer’s protocols.

### Candidate Drug Screen

Drug screening was performed at the Small Molecule Screening Shared Resource (SMSSR) at RPCCC.

### Tissue Microarray

The RPCCC prostate adenocarcinoma tissue microarray and associated de-identified clinical information was made available through the RPCCC Pathology Resource Network (PRN) and Data Bank and Bio-Repository (DBBR) core facilities. This collection includes tissue (3 distinct core samples from tumor and matching normal tissue) for 707 patients that underwent radical prostatectomy (RP) at RPCCC between 1993 and 2005. De-identified clinical annotations include patient characteristics (BMI, race, age, PSA), pathological information (Gleason sum, TNM), adjuvant therapy (ADT, radiation), and outcomes post-RP (biochemical recurrence, metastases, death) with maximum follow-up time of 18.6 years (mean = 8.8 years). Patients were considered to have received adjuvant ADT if given prior to surgery, or at any point post-surgery but before biochemical recurrence (BCR). Optimization and staining of NCOR2 (HPA001928, Sigma-Aldrich) and NCOR1 (HPA051168, Sigma-Aldrich) were performed by the RPCCC PRN. Quality assessment, nuclear identification and staining quantification (H-score) was performed using Aperio Nuclear v9 algorithm. Tissue cores were filtered for those with at least 20 detectable epithelial nuclei, and each individual core was pathologically examined to ensure tumor or normal involvement. Only patients with 2 or more cores that passed these criteria were retained for further analysis (564 patients available for NCOR2 analysis, and 463 for NCOR1). Univariate and multivariate linear regression was applied to examine relationships between extraneous clinical variables with NCORs protein expression. BCR survival separated on staining quantification (median cut-off) was assessed by Cox proportional hazards regression within clinical sub-groups to limit confounding variables, and statistical differences deemed by log-rank test.

### Functional annotation of gene sets

Pathway enrichment analysis and gene set enrichment analysis (GSEA) were performed using gene sets from the Molecular signatures database (MSigDB). Specifically, gene sets were compiled to assess enrichment of all BROAD Hallmark pathways, curated pathways (KEGG, BioCarta, Canonical, Reactome, Chemical/Genetic perturbations), and GO terms (Biological Processes). GSEA was implemented using the *clusterProfiler* and *fgsea* packages in R. Master regulator analysis (MRA) was performed on select gene sets using *iRegulon* implemented in Cytoscape.

### Data analyses and integration

All analyses were undertaken using the R platform for statistical computing (version 3.6.1) and the indicated library packages implemented in Bioconductor.

## Results

### NCOR2 expression is associated with ADT recurrence in human PCa

A 707 patient tissue microarray compiled from men who underwent radical prostatectomy (RP) at RPCCC was examined for NCOR2 and NCOR1 expression by immunohistochemistry (Figure 1A). Patient clinical follow up post-surgery was maintained for maximum 18 years (mean follow-up = 8.8 years) (Table S1). Notably, a patient subset (n = 136) received adjuvant ADT prior to (n=126) or following RP (n=10). Univariate regression analyses identified significant associations of NCOR2 expression (H-score) in patients with; race (decreased in African Americans), BMI (decreased in overweight/obese); pre-surgical PSA (decreased with pre-surgical PSA > 4 ng/mL); and adjuvant ADT. Multivariate regression identified additional association with Gleason sum (decreased with Gleason sum 8+) (Table S2). NCOR1 expression significantly associated with elevated BMI, pre-surgical PSA and adjuvant ADT (Table S3). Neither age nor pathologic stage associated with expression of NCOR2 or NCOR1.

**Figure 1.**
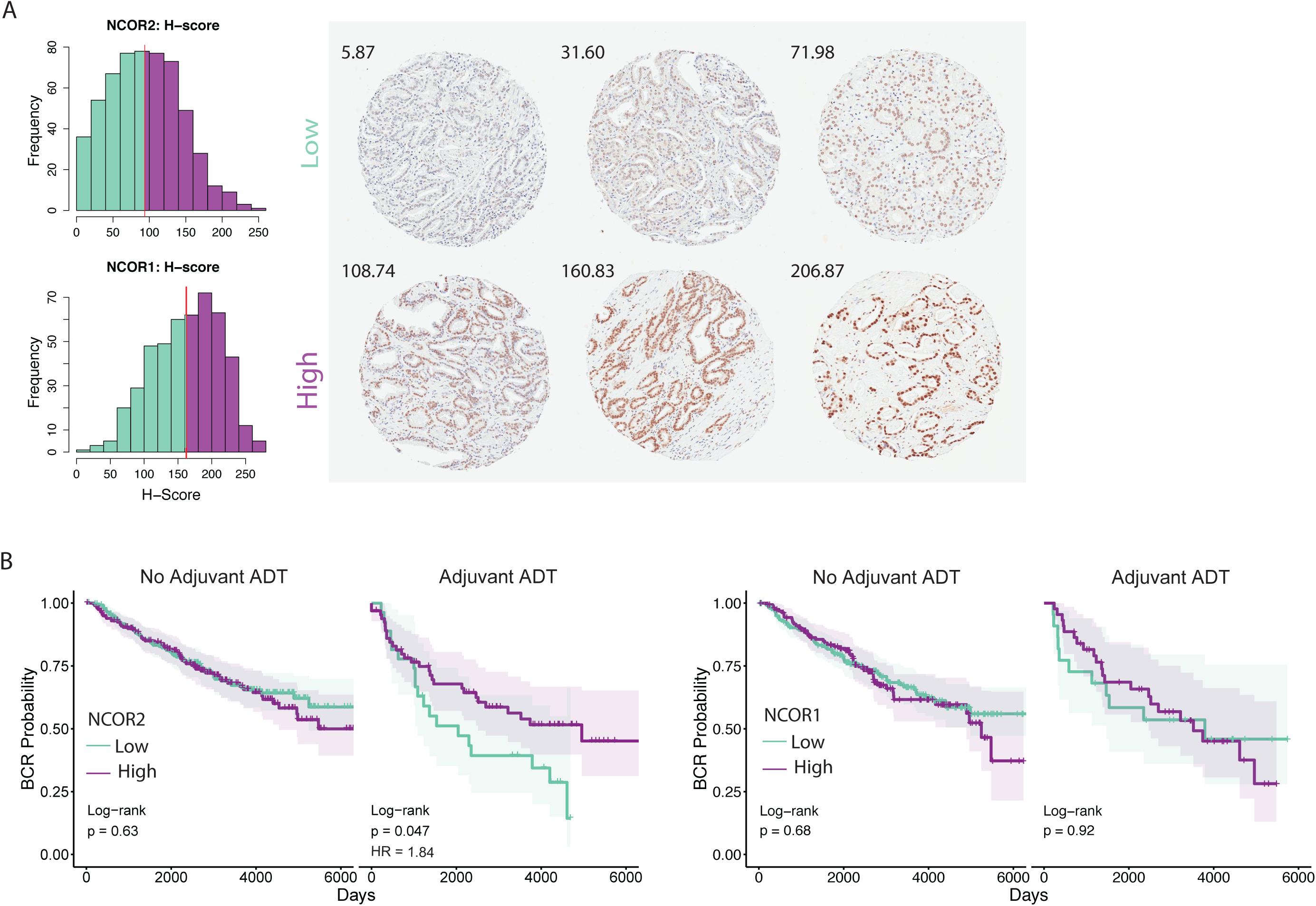
Legend: **A)** Distribution of protein expression (H-score) determined by IHC in the RPCCC PCa TMA (left). Red line indicates median expression, separating low expressing (green) and high expressing (purple) tumors. Representative tissue cores from six individual patients showing low and high nuclear staining of NCOR2 (right). H-scores for each individual core are shown. **B)** BCR survival assessment of patients with high and low expression of NCOR2 (left) and NCOR1 (right). For each, survival was examined within sub-cohorts of patients that did or did not receive adjuvant ADT with RP. Significance of Cox proportional hazards regression is shown (log-rank test), as well as hazards ratio (HR) for significant shifts in survival.

Univariate and multivariate Cox proportional hazards regression revealed relationships between clinical variables and time to BCR (Table S4). Gleason sum and pathologic stage were identified as significant indications of reduced BCR survival (Figure S1A). Patients receiving adjuvant ADT also had reduced survival but had a significantly skewed distribution of Gleason sum (Figure S1B), consistent with the fact that patients diagnosed with aggressive primary disease are more likely to be given adjuvant ADT.

Following normalization for either Gleason sum, pathologic stage, race or BMI NCOR2 levels (median cut-off) did not stratify survival of patients (Figure S1C-D). Strikingly, reduced NCOR2 (and not NCOR1) was significantly associated with worse BCR survival in patients receiving adjuvant ADT (Figure 1B). No such relationships were observed in patients who received surgery without ADT. These observations strongly support the concept that reduced NCOR2 dampens response to ADT in PCa patients.

### Reduced NCOR2 impacts DHT-dependent and independent transcriptomes

Given the relationships between reduced NCOR2 and ADT, we next established the impact of stable lentiviral-mediated shRNA knockdown of NCOR2 on DHT-dependent (10nM, 6hr) and independent transcriptomes (Figure S2A-D) in the isogenic LNCaP and LNCaP-C4-2 (C4-2) cell lines. Parental LNCaP and C4-2 cell growth responses to R1881 and Enzalutamide were as predicted (Figure S2E) and DHT exposure did not alter NCOR2 expression in the control or knockdown clones (Figure S2F).

Similarity and principal component analyses revealed that experimental conditions explained the majority of variation in expression (Figure S3A-B). Differentially expressed genes (DEGs; FDR < 0.05, FC > 1.2) were determined as either DHT-dependent in CTL cells (Figure 1A; magenta), DHT-dependent in NCOR2 knockdown cells (green), or NCOR2-dependent (orange). There were more DHT-dependent DEGs in LNCaP (1,396 total - 917 upregulated, 479 downregulated) than in C4-2 (700 total - 423 upregulated, 277 downregulated) cells (Figure 2A-B). Conversely, the NCOR2 dependent transcriptome was strikingly larger in C4-2 (2,138 total - 1331 upregulated, 807 downregulated) than LNCaP (444 total - 252 up, 192 down) cells.

**Figure 2:**
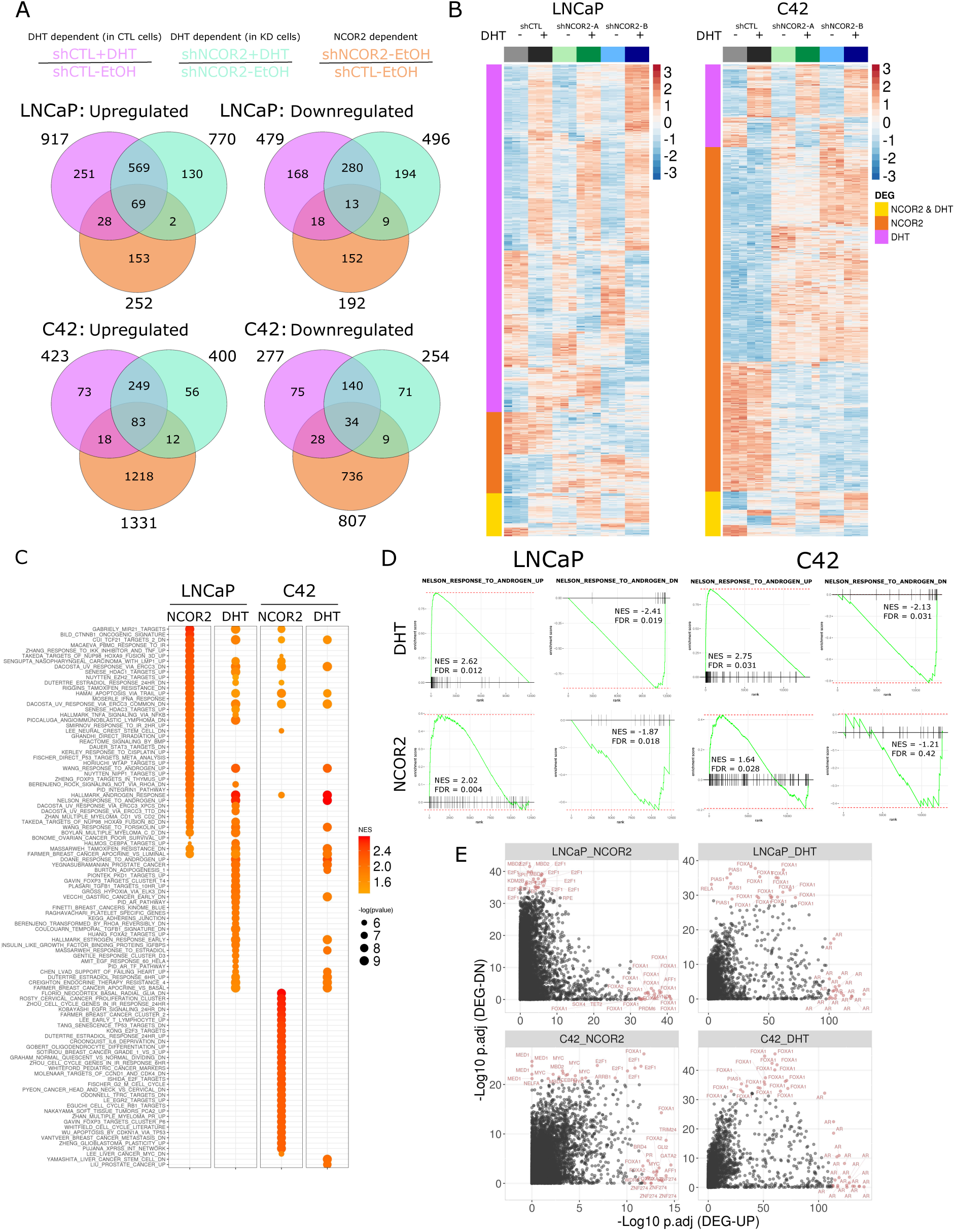
Identifying NCOR2 and DHT dependent gene expression patterns in LNCaP and C4-2 cells. **A)** Venn diagram depicting DEGs determined in each comparison. Gene sets were considered either NCOR2 dependent (shNCOR2-EtOH / shCTL-EtOH, orange), DHT dependent in shCTL cells (shCTL+DHT / shCTL-EtOH, magenta), or DHT dependent in shNCOR2 cells (shNCOR2+DHT / shNCOR2-EtOH, green). **B)** Heatmap representing all DEGs determined as either DHT or NCOR2 dependent in LNCaP (left) and C4-2 (right). **C)** The top 30 most significant upregulated pathways from GSEA analysis of NCOR2 and DHT dependent gene expression changes. **D)** GSEA enrichment plots of the NELSON_RESPONSE_TO_ANDROGEN up and down pathways. **E)** LISA analysis of DHT and NCOR2 dependent DEGs. Each point represents a single ChIP-seq dataset queried from the CistromeDB. The top 20 most significant enrichments for both up and downregulated genes is highlighted.

Within a given cell background, the majority of DHT-dependent genes were consistent between CTL (magenta) and NCOR2 knockdown cells (green) (Figure 2A, Figure S4A-B). For instance, the DHT dependent transcriptome in CTL and NCOR2 knockdown cells was similar both in the number of DEGs detected (931 and 506 shared DEGs in LNCaP and C4-2 cells, respectively), and in the magnitude of DHT associated induction (r = 0.90 and 0.80 in LNCaP and C4-2 cells, respectively). Thus, while loss of NCOR2 expression did not alter a majority of the DHT capacity (genes affected) or sensitivity (magnitude of change), there were nevertheless an additional 335 and 148 genes that gained DHT regulation, and conversely 465 and 194 genes lost DHT regulation, suggesting a skewed transcriptional response to DHT stimulation.

Gene set enrichment analysis (GSEA) was applied to DEGs using a comprehensive set of pathways compiled from MSigDB (Figure 2C, Figure S5, Table S5). In LNCaP cells, DHT stimulation resulted in strong enrichment for androgen-associated pathways (e.g. NELSON_RESPONSE_TO_ANDROGEN, Figure 2D), as expected. Interestingly, NCOR2 knockdown, independently of DHT treatment, also significantly enriched for androgen response pathways. This suggested NCOR2 levels alone control gene expression in a similar manner to DHT exposure. Indeed, there was a strong degree of commonality between NCOR2- and DHT-dependent pathways in LNCaP cells, including gene sets regulated by HDAC activity (e.g. SENESE_HDAC3_TARGETS). However, distinct NCOR2 associated enrichments were also observed, including pathways regulated by STAT, NFKB and FOXP3 (e.g. ZHENG_BOUND_BY_FOXP3) that were not induced by DHT, suggesting roles for NCOR2 in mediating other transcription factor signaling responses.

C4-2 cells displayed similar DHT regulated enrichment of androgen associated pathways (Figure S4A-B), and overall pathway enrichment patterns were similar to that observed in LNCaP cells. NCOR2 knockdown in C4-2 cells also enriched for androgen response pathways, although the overall concordance between DHT and NCOR2 regulated pathways was weaker in C42 than in LNCaP cells. NCOR2 enrichment patterns were more distinct between cell types as only 6% of NCOR2 regulated genes in C4-2 were also observed in LNCaP (Figure S4A). Unique enrichments to the NCOR2 dependent transcriptome in C4-2 cells were observed for pathways involving neuronal differentiation (e.g. GOBERT_OLIGODENDROCYTE_DIFFERENTIATION), RB1/E2F (e.g. ISHIDA_E2F_TARGETS) and TP53 regulation (e.g. TANG_SENESCENCE_TP53_TARGETS).

Transcriptional master regulators (MR) of DEGs were inferred using LISA ^43^ (Figure 2E). DHT-dependent DEGs enriched strongly for AR as well as the forkhead box factor FOXA1, which has characterized AR pioneering function, in both cell lines. NCOR2-dependent regulators included FOXA1 but were more diverse, including E2F and the methyl binding factor MBD2. Cell line specific regulation was also observed, including TET2 regulation in LNCaP and MYC regulation in C4-2. iRegulon analysis largely confirmed these observations (Table S6), including common FOXA1 enrichment. In C4-2 cells, several factors associated with neuroendorine differentiation were enriched, including FOXM1 and MYCN.

Small RNA-seq (Figure S6, Table S7) also supported a greater impact of NCOR2 in C4-2; 15 miRNA were significantly regulated including regulation of several known tumor-suppressors (e.g. miR-10a) and oncomirs (e.g. let-7e). Another six were also regulated by DHT treatment, confirmed by RT-qPCR (Figure S6A-C). Upregulated Let-7e and miR-200a seed sequences were significantly enriched in NCOR2-dependent downregulated DEGs in C4-2 cells suggesting that NCOR2-dependent transcriptome reflects direct events amplified by altered miRNA-gene networks. This was not evident in LNCaP cells (Figure S6B-C).

In total, these analyses strongly support the concept that reduced NCOR2 levels alter the DHT response in two ways; by (1) skewing the capacity of the DHT regulated transcriptome and by (2) mimicking a subset of the androgen transcriptional responses, even in the absence of DHT. In C4-2 cells, the impact of reduced NCOR2 alone was significantly more extensive than in LNCaP cells and enriched for cell-cycle and neuronal differentiation responses, as well as genes associated with endocrine therapy resistance and putative regulation by neuroendocrine transcription factors.

### NCOR2 loss induces hypermethylation at enhancer regions

Given the links between NCOR2 and DNA methylation machinery, we examined changes in DNA methylation following NCOR2 knockdown and DHT treatment. Again, experimental conditions largely explained the variance in DNA methylation levels (Figure S7A-B). Differential methylation analyses (FDR < 0.05, 10% change in methylation) revealed a substantial shift towards hypermethylation events following NCOR2 knockdown. In LNCaP cells differentially methylated positions (DMP) were readily identified (25,843 total (DMPs); 98% hypermethylated) and was even more striking in C4-2 (184,663 total DMPs; 82% hypermethylated) (Figure 3A). Notably, C4-2 DMPs largely encompassed LNCaP DMPs (50%) (Figure 3B). DHT treatment had no significant effect on DNA methylation in either cell line (Figure S7C).

**Figure 3:**
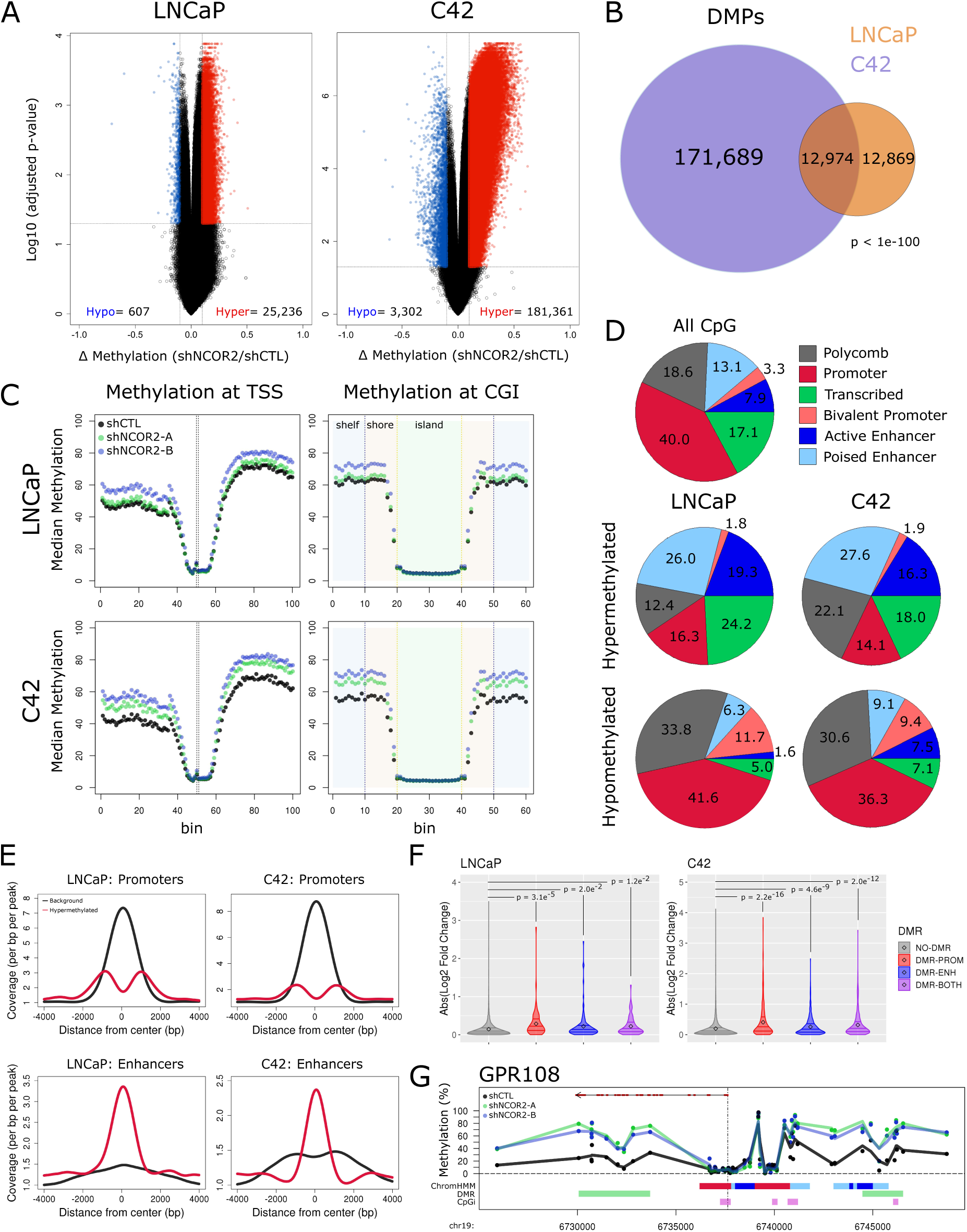
NCOR2 loss results in a hypermethylation phenotype in LNCaP and C4-2 cells. **A)** Volcano plots representing methylation changes identified upon NCOR2 knockdown in LNCaP (left) and C4-2 (right) cells. Determined DMPs are shown in blue (hypomethylated) and red (hypermethylated). **B)** Venn diagram of DMPs identified in each cell. **C)** Binning analysis depicting the median methylation calculated for genomic regions relative to TSS (left) and CpG island (right) loci. For TSS, each bin represents 100bp, with the TSS centered at bin 50. For CpG islands, shore (orange) and shelf (blue) bins represent 200bp, while islands (green) are variable depending on genomic length but centered on bin 30. **D)** Relative proportions of all CpGs (top) or DMPs (middle, bottom) that annotate to ChromHMM regions. **E)** Peak centered densities of non-DMP CpG sites (black) or hypermethylated DMPs (red) centered at ChromHMM promoter or enhancer regions. **F)** Comparison of NCOR2 knockdown associated expression changes of genes annotated with DMR-promoter and/or DMR-enhancers relative to genes not annotated. Distributions are compared by KS-test. **G)** Representative genomic view of an NCOR2 dependent gene locus with annotated enhancer hypermethylation (*GPR108*). Tracks are as follows starting from top; RefSeq gene (exons (red) and introns (black)); Methylation detected in shCTL (black) or shNCOR2 C4-2 cells (green, blue); ChromHMM regions (color code is same as shown in **D**); Determined DMR regions (green); CpG island regions (purple).

The NCOR2-dependent global distribution shifts in DMPs were most pronounced at genomic sites distal to both transcriptional start sites (TSS) and CpG islands (CGI) (Figure 3C). We exploited ChromHMM defined chromatin states in LNCaP cells ^53-55^ to reveal that NCOR2 dependent hypermethylated DMPs (hyper-DMP) were observed at a lower than expected rate at active promoter loci in both LNCaP and C4-2 cells and conversely enriched in poised and active enhancer regions (21% background, 45% hyper-DMP (LNCaP), 44% hyper-DMP (C4-2)). (Figure 3D-E, Figure S8A). Although fewer, hypomethylated DMPs (hypo-DMP) were enriched at Polycomb and bivalent promoter regions.

Differentially methylated regions (DMRs; FDR < 0.05, Δ regional methylation > 5%) ^48^ were identified, revealing 929 and 7,845 DMRs in LNCaP and C4-2 cells respectively, with >99% of these being hyper-DMRs (Figure S8B). In both LNCaP and C4-2 a substantial proportion of DMRs mapped to at least one enhancer region (37%, 29% respectively), with fewer mapping to promoter regions (30%, 21%). DMRs were annotated to gene proximal enhancers or promoters (enhancer region within +/- 20kb of TSS; promoter region within +/- 2kb of TSS) (Figure S8C). In both cell lines, genes with DMR associated enhancers or promoters were significantly more affected at the expression level by NCOR2 knockdown than genes without (Figure 3F). DMR-enhancers tended to be distinct from promoter regions and exclusive of CpG islands, as shown at representative loci for GPR108, TRIP10 and TLE6 (Figure 3G, Figure S8D).

GSEA of genes associated with NCOR2-dependent DMR-enhancers did not enrich for DHT regulated genes (Figure S8E, Table S8), but rather for pathways associated with cancer and endocrine therapy resistance (e.g. CREIGHTON_ENDOCRINE_THERAPY_RESISTANCE). Reflecting the DEG functional annotations, DMR pathways unique to LNCaP were strongly enriched in interferon response, while those unique to C4-2 included neuronal (e.g. KEGG_AXON_GUIDANCE) and P53 pathways (e.g. HALLMARK_P53_PATHWAY).

### The NCOR2 cistrome is regulated by DHT and associates with FOXA1

The NCOR2 cistromes in LNCaP and C4-2 cells were largely distinct to cell line and DHT stimulation (Figure 4A). In total, there were 1257 and 1865 robust NCOR2 binding peaks (FDR < 0.1) in untreated LNCaP and C4-2 cells, respectively. DHT exposure redistributed and increased the number of peaks to 1469 and 2068, respectively.

**Figure 4:**
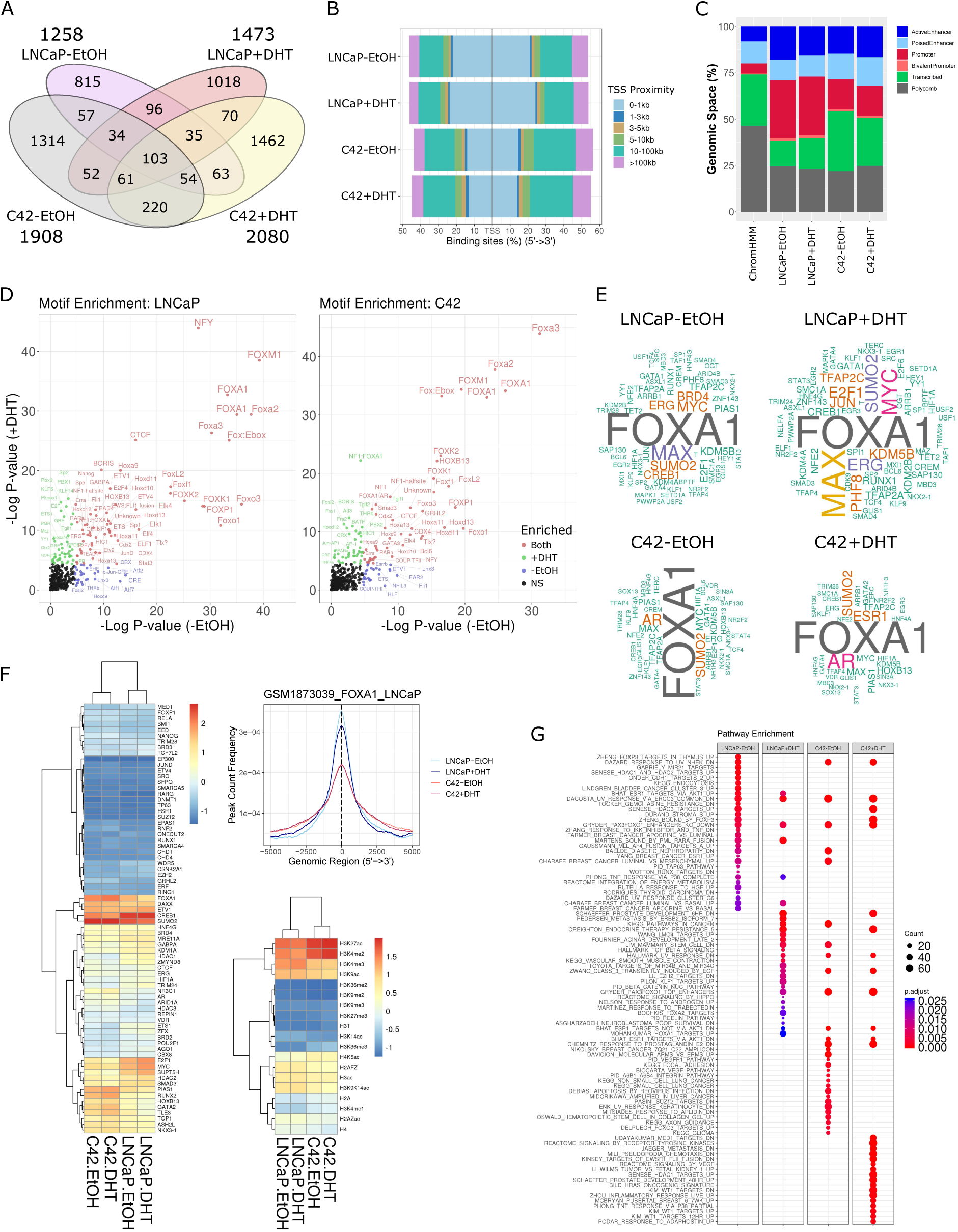
Defining the NCOR2 cistromes in LNCaP and C4-2 cells. **A)** Venn diagram of significant peaks identified in each condition. **B)** Proportional annotations of determined NCOR2 cistromes to TSS regions. **C)** Relative genomic proportions of all chromHMM regions (left) or NCOR2 cistromes that annotate to chromHMM regions. **D)** Motif analysis of NCOR2 cistromes in LNCaP (left) and C4-2 (right) cells. **E)** Word cloud depicting the frequencies of factors observed in the top 200 most overlapping datasets form the complete CistromeDB with each NCOR2 cistrome. Factors with observed proportions near background (<1.2x enrichment) were removed. **F)** Heatmap depicting the mean normalized overlaps observed for each factor with each NCOR2 cistrome within the prostate subset of the CistromeDB (left). Peak centered densities of NCOR2 cistromes (LNCaP) against a publicly available FOXA1 ChIP-seq dataset derived in LNCaP cells ^70^ (right). Mean normalized heatmap for histone mark datasets available within the prostate subset of the CistromeDB. **G)** The top 30 most significantly enriched pathways from functional enrichment analysis of genes annotated to NCOR2 cistromes.

In LNCaP cells, almost half of peaks (45%) were within 3kb of a TSS, which was modestly increased with DHT treatment (50%) (Figure 4B, Figure S9A-B). By contrast, in C4-2 cells the NCOR2 cistrome was skewed towards more distal regions, with the highest proportions (∼40%) of peaks falling between 10-100kb of the closest TSS in basal and DHT treated conditions. Compared to the background distribution of ChromHMM states, NCOR2 binding was enriched in enhancers and diminished in Polycomb associated regions across conditions. NCOR2 binding was also enriched in promoter regions, particularly in LNCaP cells (Figure 4C).

In both LNCaP and C4-2 cells, NCOR2 peaks were enriched for several NR motifs, including PPARs, RARs, THR, ERRa, GR and PGR elements. Notably, NR motifs were more evident upon DHT exposure, although some orphan NR (COUP-TFII, EAR2) enrichments were observed in basal conditions (Figure 4D, Figure S9C). Interestingly, in neither cell type nor in any condition were motifs for the AR half-site significantly enriched.

NRs only represented a small proportion of significant motif enrichments. Although the genomic sites of NCOR2 binding were largely unique across cell lines and upon DHT exposure, the motif analyses revealed similar enrichment patterns across conditions. For instance, Forkhead box (FOX) transcription factors (e.g. FOXA1, FOXO1) were significantly enriched in both LNCaP and C4-2 cells, regardless of DHT status, suggesting common interactions across cellular contexts. Similarly, ETS family members (e.g. ELK1, ETS1) as well as CTCF and BORIS (CTCFL) were significantly enriched across conditions. Meanwhile, other motifs were most enriched only in the absence of DHT such as the basic leucine zipper (bZIP) domain motifs (e.g. JUNB, CREB) in LNCaP cells.

Motifs were annotated to larger transcription factor families, and family-wide enrichments tested to gauge what types of factors were enriched (Figure S9D). FOX factors were the most highly enriched family across all conditions, suggesting an important role for these factors in the NCOR2 regulatory function. Other higher than expected enrichments were observed for ETS, E2F and bZIP factors. Notably, NR factors were enriched only at slightly higher than expected levels.

NCOR2 cistromes were queried for transcription factor binding against the complete CistromeDB collection (> 10,000 total ChIP-seq datasets, across > 1100 factors) using GIGGLE ^50^ (Figure 4E, Figure S9E-F, Table S9). This approach provided strong evidence for co-accumulation of NCOR2 with copious factors, but consistent with motif analysis, overlaps were strongly enriched for FOXA1 binding across conditions. Other commonly enriched factors included AR, MYC/MAX, E2F1, SUMO2, CREB1 and KMD5B. Filtering for prostate specific datasets confirmed strong overlap of NCOR2 cistromes with FOXA1, CREB1 and SUMO2 as well as DAXX and ETV1 (Figure 4F). Notably, enrichment patterns were cell line dependent and included enrichments more dominant in LNCaP (CREB1, E2F1, MYC, HDAC1) or C4-2 (AR, FOXA1, SUMO2, RUNX2). Similarly, prostate histone mark datasets were interrogated, revealing an increased H3K27ac and H3K4me2 and decreased H3K4me3 enrichments in C4-2 cells relative to LNCaP, consistent with a movement towards distal enhancer regions. Notably, enrichment for repressive marks H3K9me3 and H3K27me3 were relatively weak.

GSEA analyses of NCOR2 peak annotated genes reflected previous findings (Figure 4G). Androgen regulated pathways were enriched, particularly in LNCaP cells treated with DHT but were either reduced or not significant in C4-2 cells. HDAC activity and FOX function were commonly enriched amongst all conditions, including genes shown to be regulated by FOXO occupied. In C4-2 cells, several pathways involved stem cell or neuronal signaling (e.g. KEGG_AXON_GUIDANCE) were also enriched.

These data suggest NCOR2 bind sites are divergent dependent on cell types and DHT exposure but are commonly enriched for open chromatin and enhancer regions that contain a diverse set transcription factor binding elements, particularly those of FOX family members. Whilst AR motifs are not prominent in the motif analyses, there is pronounced overlap with AR-dependent ChIP-Seq data sets, as well as clear overlap with FOXA1 ChIP-Seq and functional pathways in both cell types.

### Integration of NCOR2 dependent transcriptomes, DNA methylomes, and cistromes

NCOR2-dependent omic integration was undertaken to define how the genomic context impacted NCOR2 peak and gene expression (peak:gene) relationships by first examining the extent of overlaps between gene sets identified across omic data types. NCOR2-dependent gene sets identified at the levels of expression, DNA-enhancer methylation and genomic occupancy shared significant overlap in both LNCaP and C4-2 cells. For instance, 257 genes were found to have both NCOR2 dependent gene expression and had a proximal DMR-enhancer in C4-2 cells (Fig 5A, which was significantly higher than expected by chance (OR = 1.78, p-val = 3.08e-^13^) and substantiated previous analysis integrating DMRs with expression (Figure 3F). Additionally, within each omic level, strong overlaps were observed between cell lines suggesting a core NCOR2 regulatory function that is conserved in the androgen sensitive and ADT-R contexts (Fig 5B).

**Figure 5:**
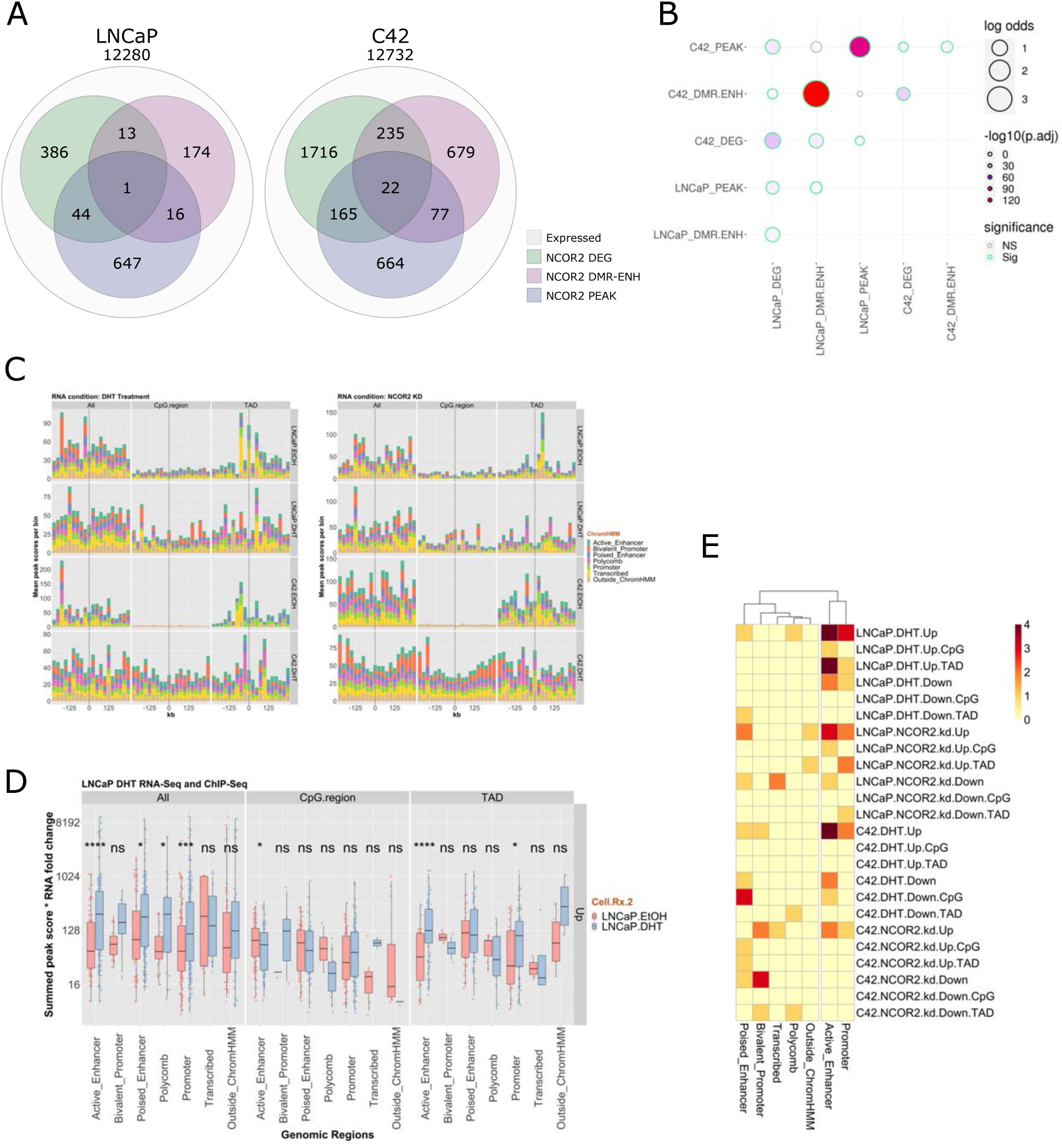
Integrative analyses of NCOR2-dependent transcriptome, methylome and cistrome. **A)** Venn diagram of gene sets (expressed genes) linked to NCOR2 regulatory function by expression, enhancer DNA methylation, or cistrome analyses. **B)** Assessment of gene set overlaps from **A**. Bubble plot depicts the log odds ratio (size) and significance (color) of each respective overlap. **C)** The spatial distribution of mean score of NCOR2 binding in ChromHMM regions in LNCaP and C42. Basal and DHT treated (10 nM, 1h) NCOR2 peaks were overlapped to the indicated ChromHMM regions derived in LNCaP cells, the NCOR2-dependent differentially methylated regions (CpG region) identified in the same cell type, or with TAD regions identified in LNCaP cells. **D)** The sum of peak scores within 250 kb of a given gene in either LNCaP or C42 were calculated and multiplied by the absolute fold change of the gene to derive the Summed Expression Score (SEC). The changes in SEC is shown for basal and DHT-induced NCOR2 binding in LNCaP cells using the LNCaP RNA-Seq following DHT treatment. A two-way t-test was used to test difference between SEC between basal and DHT-induced NCOR2 binding. **E)** The significance of difference between SEC was determined for the indicated both ChIP-Seq data sets (LNCaP or C42 treated with DHT) and DHT- and NCOR2-dependent RNA-Seq. Color represents -log10(p-values), with the darker color being more significant.

NCOR2 cistromes were next annotated by three levels of genomic feature; ChromHMM regions and TADs both defined in LNCaP ^56^, and the respective NCOR2-dependent DMRs. Given that the median distance for enhancer to TSS is 158 kb^34^, these annotated NCOR2 cistromes were related to NCOR2 or DHT regulated genes within a 250 kb window.

The number of NCOR2 peak:gene relationships differed between LNCaP and C4-2 by annotation and DHT treatment (Figure S10, Table S10). For example, peak:gene relationships were broadly constant in Active or Poised Enhancers, but increased with DHT treatment only in LNCaP cells. In C4-2 cells there were many more NCOR2 peak:gene relationships outside of ChromHMM classification. These peak:gene relationships were resolved further by overlapping with TAD (Table S11) and DMR (Table S12) regions. Within TADs, LNCaP peak:gene relationships at Promoter regions were approximately twice as frequent as in C4-2. Interestingly, whilst there were almost ten times as many DMRs in C4-2 than LNCaP cells, the enrichment of peak:gene relationships overlapping with DMRs were broadly consistent, with 6238 peak:gene DMR-promoter relationships in LNCaP and 7092 in C4-2.

Restricting peak:gene relationships to genes expressed in the respective cell background, we and addressed the impact of NCOR2 peak distance from target gene. Specifically, we summarized NCOR2 peak:gene relationships into 25kb bins up and downstream from target genes, and calculated changes in mean ChIP-Seq peak score for genomic annotation (e.g. ChromHMM state, contained within TAD or overlapped with DMR) (Figure 5C). This revealed a number of prominent genomic spatial relationships including in LNCaP cells, higher NCOR2 peak scores in Bivalent promoters associated upstream of genes regulated by DHT treatment (Figure 5C, left panel). In both cell types the mean significance of basal NCOR2 binding overlapped with DMR was generally low, but NCOR2 proximal binding was notable in proximal regions associated in TADs, most clearly in Transcribed regions. DHT treatment modestly increased the mean NCOR2 peak DMR-dependent score notably in Bivalent Promoters, and also in Transcribed regions enriched in TAD regions notably downstream of target genes. In C4-2 cells higher significance of basal upstream NCOR2 binding in Bivalent Promoters, low significance in DMRs but enhanced by DHT treatment, and TAD regions being enriched for proximal Transcribed regions were all apparent.

Considering genes regulated by NCOR2 knockdown (Figure 5C, right panel) revealed in LNCaP high basal NCOR2 binding in Bivalent Promoters, and within TADs, and proximal downstream Poised Enhancers. In C4-2 cells, some of the spatial relationships were more emphasized, with the basal NCOR2 peaks within TADS on average being more significant in proximal regions (including in Active Enhancers).

From these classifications we built on the BETA method ^57^ to test the NCOR2-dependent cistrome-transcriptome relationships. Specifically, the sum was calculated of NCOR2 peak scores within 250 kb of each regulated by either DHT or NCOR2 knockdown, multiplied by the absolute gene-specific fold change to define the Summed Expression Score (SEC); the SEC was used to test how NCOR2 binding in defined genomic features related to changes in gene expression (Figure 5D,E). For example, the SEC at Active Enhancers in LNCaP cells treated with DHT was significantly greater (p < 0.0001) in DHT-treated NCOR2 ChIP-Seq (Figure 5D). This is reversed with NCOR2 binding within DMRs, whereas it is sustained within TADs at Active Enhancers.

The clustering of significant differences in SEC are summarized in Figure 5E and revealed the NCOR2 binding in Active Enhancers and Promoters were highly similar. Perhaps surprisingly they reveal a mixed function coregulator impact of NCOR2. Thus, the SEC score for DHT-treated NCOR2 is significantly higher at Active Enhancers for both up and down-regulated genes in LNCaP. In C4-2 cells this was more pronounced for up-regulated genes suggesting its function at these sites is more of a coactivator function for DHT-regulated genes. Also reflecting the distribution of NCOR2 binding there are a number of significant relationships in C4-2 between Polycomb and Bivalent Promoter binding, suggesting these are more impactful.

### Reduced NCOR2 expression limits the effectiveness of androgen deprivation and accelerates disease progression in vivo

Next we sought to assess the impact of reduced NCOR2 expression on PCa progression *in vivo* in the CWR22 xenograft model, which recapitulates the tumor-impact of androgen withdrawal and tumor-recurrence. Mice were inoculated with CWR22-shCTL or CWR22-shNCOR2 tissue (n = 65 per group), with 100 animals (n = 50 per group) designated for follow up to either recurrence or end of study (Figure S11A-B). GFP detection of construct and reduced NCOR2 levels were confirmed at all stages of disease (Figure 6A-B,). Expression of androgen regulated genes (i.e. *TMPRSS2, HERC3*) conformed repression by androgen withdrawal, but re-expression in recurrence concomitant with increased *AR* expression (Figure S11C) ^58^.

**Figure 6:**
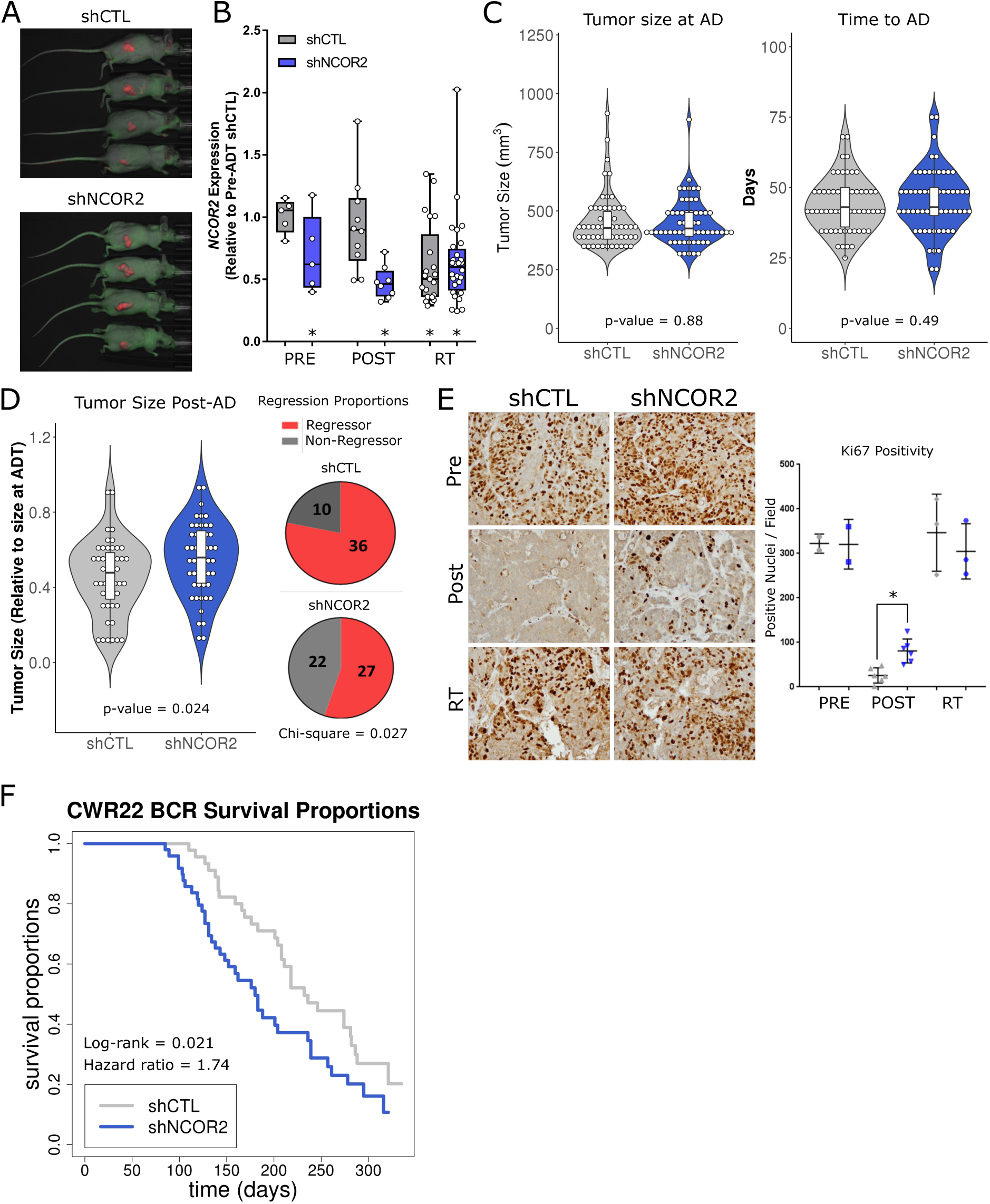
NCOR2 loss alters response to ADT in the CWR22 model of PCa progression. **A)** Representative *In vivo* imaging showing fluorescent (GFP) detection of xenograft tumors at point of recurrence > 300 days post androgen withdrawal. **B)** Relative NCOR2 expression in select tumors pre-ADT (n=5,5), post-ADT (n = 10,10), and in RT (n = 22,28). Asterisk represents significant difference (p < 0.05) between respective distribution and that of shCTL pre-ADT tumors. **C)** Violin plots showing the distribution of tumor sizes at time of ADT (left) and time to reach ADT (right). **D**) Violin plots showing the distribution of maximum regression post androgen withdrawal for each tumor (left), and overall proportions of tumors that reached 40% regression (right). **E)** Representative IHC (left) and quantification (right) of Ki67 staining in select tumors pre-ADT (n=2,2), post-ADT (n = 6,6) and in RT (n = 3,3). Asterisk represent p < 0.05. **F)** Kaplan-Meier representation of the biochemical recurrence (BCR) survival proportions in shCTL and shNCOR2 tumors post-ADT.

Reduced NCOR2 expression neither impacted androgen-stimulated tumor size nor growth rate prior to androgen deprivation (pre-AD) (Figure 6C). By contrast, the initial response to androgen deprivation (post-AD) was significantly reduced in shNCOR2 tumors (Figure 6D). A total of 36 out of 46 (78%) of control tumors reached regression, as defined as a 40% reduction in tumor size post-AD (mean regression = 54%). However, only 27 out of 49 shNCOR2 tumors (55%) reached regression (mean maximum regression = 45%) (Chi-square = 0.027). Furthermore, Ki-67 staining was starkly reduced 1 week following AD in control tumors, but not as clearly in shNCOR2 tumors, indicative of a dampened AD-response (Figure 6E).

Tumor recurrence rates in control tumors were similar to those previously reported (50% recurrence at 232 days post-AD) ^59^ (Figure 6F). However, shNCOR2 tumors recurred at a significantly faster rate (50% recurrence at 180 days, Log-rank = 0.021, Hazard ratio = 1.74). Interestingly, *NCOR2* expression in control recurrent tumors (RT) was also significantly reduced relative to pre and post-AD control tumors (Figure 6B). This suggests that reduced NCOR2 expression is common event in tumor-recurrence following AD.

### NCOR2 knockdown accelerates the molecular features of ADT resistance in vivo

Reflecting the lack of phenotypic impact of NCOR2 knockdown in tumors prior to AD, the RT displayed the greatest NCOR2-dependent transcriptomic and DNA methylation changes (Figure S12A-B). NCOR2-dependent transcriptional changes were not observed in pre-AD tumors, but significant expression changes were identified post-AD (96 total DEGs; 50 upregulated, 46 downregulated) and RT (529; 225 up, 304 down). In control tumors, the acute response to AD suppressed proliferative pathways (Figure S12C), corroborating Ki67 expression patterns (Figure 6E). Gene sets associated with ADT resistant phenotypes were observed ^60^, including transient depletion of AR and FGFR signaling post-ADT that was re-established in RT. These patterns were not affected by NCOR2 status. However, RT-shNCOR2 displayed increased enrichment for neuroendocrine signaling relative to RT-shCTL (Figure 7A). This observation is typified by increased levels of *SYP, CHGB* and *NTN1* and reduced *AR* mRNA, and elevated SYP protein in RT-shNCOR2 tumors (Figure 7B-C).

**Figure 7:**
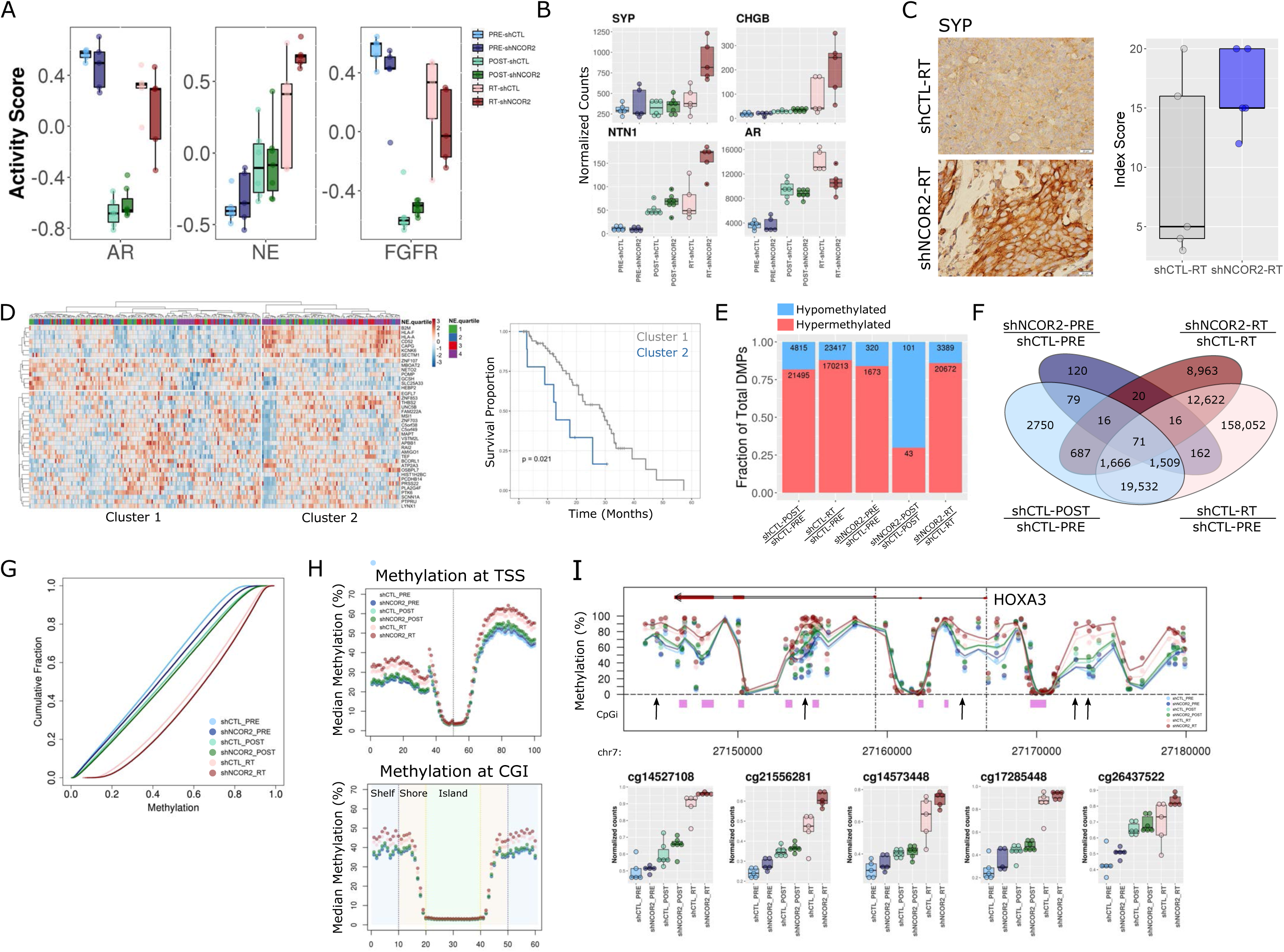
NCOR2 loss accelerates the molecular features of ADT resistance during PCa progression *in vivo*. **A)** GSVA analysis of established pathways associated with ADT-R in prostate cancer. **B)** Gene expression profiles for *SYP, CHGB, NTN1* and *AR* in CWR22 groups. **C)** IHC showing representative examples of SYP expression in shCTL-RT and shNCOR2-RT samples (left). Quantification of IHC in RT is shown (right). **D)** Heatmap of scaled expression depicting the 41 most significant DEGs in NCOR2-low relative to NCOR2-high tumors within the SU2C cohort that were also identified in RT-shNCOR2 (left). Column annotations show grouped neuroendocrine scores (NE.quariles) for each tumor. Kaplan-meier curves showing overall survival proportions between major tumor clusters identified. **E)** Proportion of DMPs in each comparison found to be relatively hypermethylated (red) or hypomethylated (blue). **F)** Total number of NCOR2 and stage specific DMPs determined prior to androgen withdrawal (PRE) and at recurrence (RT). **G)** Cumulative distribution plots showing the total methylation levels observed within each CWR22 group. **H)** Binning analysis depicting the median methylation calculated for genomic regions relative to TSS (top) and CpG island (bottom) loci. For TSS, each bin represents 100bp, with the TSS centered at bin 50. For CpG islands, shore (orange) and shelf (blue) bins represent 200bp, while islands (green) are variable depending on genomic length but centered on bin 30. **I)** Representative genomic view of the HOXA3 locus, showing the average methylation levels determined within each CWR22 group. CpG island regions are shown (purple). Arrows depict example DMPs which are depicted in order below.

We investigated how NCOR2-dependent RT expression associated with outcome in patients receiving ADT ^16^. Tumors (n = 270) were categorized by NCOR2-high or NCOR2-low expression (quartile) and DEGs identified. To identify the consequences of reduced NCOR2 levels, these DEGs were overlapped with NCOR2-dependent RT associated genes, revealing a 242 gene signature that was further filtered to reveal those most strongly associated with NCOR2 status (> 1.5 Z-scores > 20% tumors). Expression of these 41 genes separated patients into two major tumor clusters that were significantly associated with neuroendocrine score ^16^ (Chi-square = 0.014) (Figure 7D, Figure S12D). Furthermore, univariate Cox proportional hazards regression revealed tumor cluster membership significantly associated with risk of death following ADT (Log-rank = 0.07, Hazard ratio = 2.47).

Broad hypermethylation was detected in post-AD and RT relative to pre-AD (Figure 7E). Most (87%) methylation changes observed post-ADT were also observed in RT, suggesting that DNA methylation patterns observed in recurrence are progressive from events observed in acute response (Figure 7F). Similar to the impact of NCOR2 knockdown on DNA methylation patterns in cell line, NCOR2 knockdown *in vivo* also associated with hypermethylation both in primary and recurrent tumors. Notably, NCOR2-associated DMPs strongly overlapped with stage specific DMPs. Considering hypermethylation events observed in RT, shNCOR2 tumors were significantly more hypermethylated than shCTL counterparts within each stage of disease (Figure 7G). Thus, even prior to the stress of ADT, loss of NCOR2 leads to subtle increases in methylation at loci that gain high level methylation upon recurrence. Binning analyses revealed progressive and NCOR2-dependent hypermethylation was enriched at regions distal to TSS and CpG island loci (Figure 7H), and ChromHMM annotations indicated enrichment for RT and NCOR2 associated DMPs at enhancer regions (Figure S13B). For example, progressive hypermethylation was observed broadly at the *HOXA3* locus (Figure 7I). Gene-annotated DMR analysis for shNCOR2-RT relative to shCTL-RT revealed 32 genes that were also differentially expressed in the same samples, including several associated with neuronal development (e.g. *CHGA, NTN1*).

Intriguingly, the hypermethylation phenotype associated with ADT tumor recurrence was observed in a pilot study of primary and unpaired ADT-RPCa human tumor samples (local recurrences) (Figure S13A). DMP analysis revealed ADT-RPCa associated hypermethylation events (76% of 31,593 total DMPs hypermethylated). Genomic annotation identified ADT-RPCa DMP enrichment at enhancer regions, reflecting the hypermethylation associated with CWR22 recurrence, and with NCOR2 loss *in vitro* and *in vivo* (Figure S13B). This suggests that enhancer hypermethylation is a general phenomenon associated with ADT-RPCa.

## Discussion

Altered transcription co-factors are drivers of a range of hormone-responsive cancers ^14,15^. Of these multiple co-regulators, altered NCOR2 has consistently been reported in late stage ADT-RPCa ^16,61^, and furthermore it and other coregulators have been implicated in resistance to hormone based therapies ^15,62^. To date ambiguity existed over whether gain or loss of NCOR2 functions were a PCa driver ^14,15^, The current study aimed to address this ambiguity with a multi-omic integrative genomic approach exploiting *in vitro, in vivo* and *in silico* resources covering the emergence of the advanced PCa.

In the first instance we exploited a large TMA of patient tumor samples and revealed that reduced NCOR2, but not NCOR1, significantly associated with various factors known to reflect an underlying aggressive disease. Notably, reduced NCOR1 was more apparent in men of African ancestry, with elevated BMI and higher levels of PSA. Intriguingly, low levels of NCOR2 in the tumors of patients who received adjuvant ADT was significantly associated with shorter time to BCR.

At the transcriptomic level, NCOR2 knockdown significantly overlapped with DHT-responsive genes in both LNCaP and C4-2. Indeed, reduced NCOR2 expression alone significantly enriched for gene signatures that included androgen responses. It therefore seems that NCOR2 both directly regulates AR responses, but also regulates genes downstream of the AR and in this manner altered NCOR2 expression can phenocopy androgen actions. These gene expression patterns were associated with both up and down-regulated genes, and although we cannot exclude indirect effects, it may suggest that NCOR2 functions are involved in changing expression levels of genes in both a positive and negative direction. The up-regulation of genes following reduced NCOR2 expression fit with a model involving a loss of allosteric interactions with HDACs. Down-regulated genes fit with HDAC-independent modes of action as reported in transgenic mice where HDAC3 recruitment is impeded ^26^. Furthermore, NCOR function has been implicated in gene activation in several contexts including in association with nuclear receptors ^28,29,63^, further indicating complex and poorly understood regulatory roles.

Surprisingly, reduced NCOR2 expression resulted in a profound increase in CpG methylation. This was unexpected given the extensive literature linking repressive histone modifications that recruit the CpG methylation machinery ^64^. Nonetheless it is clear that reduced NCOR2 expression exerts a large-scale impact on gene expression and DNA methylation. Although there was overlap between genes altered in expression and DNA methylation, there was no widespread and significant inverse correlation and suggests a more complex relationship between changes in CpG methylation and genes expression. large-scale impact of NCOR2 knockdown on gene expression. It is notable that among the ChIP-seq enrichments associated with differentially expressed genes upon NCOR2 knockdown were MBD2 in both LNCaP and C4-2, TET2 in LNCaP, and MED1 in C4-2 (Fig 1E). All three are known to interact with DNA methylation at the level of reading and erasing and therefore support a role for NCOR2 to impact DNA methylation. The methylation phenotype associated with NCOR2 loss suggests an active regulatory function of NCOR2 that dictates methylation levels of enhancer regions. However, CpG methylation is not exclusively associated with gene repression. For example, CpG methylation at enhancer regions can both repress and attract different classes of TFs, including NRs ^37^, and therefore the status of CpG at enhancer sites will impact which TFs are recruited and regulate gene expression.

NCOR2 cistrome analyses also supported this diversity of transcriptional responses and revealed evolution and expansion of binding choices in the C4-2 compared to LNCaP cells. We applied several levels of integration of transcriptomic, DNA methylome and cistrome data to define how the spatial distribution of NCOR2 binding was associated with genomic features (e.g. ChromHMM) and gene expression patterns. From these studies we revealed NCOR2 cistrome-transcriptome relationships became more distal in C4-2 than LNCaP and there was a fairly even distribution of relationships where NCOR2 functioned as a canonical co-repressor. For example, in LNCaP cells, sites of NCOR2 binding that overlapped with CpG methylation were significantly up-regulated when NCOR2 was knocked-down. By contrast, in other circumstances its function was as a co-activator. For example, in C4-2 cells genes where NCOR2 was bound at Bivalent Enhancers were significantly reduced in expression when NCOR2 was reduced.

Together these studies suggest that NCOR2 may be part of a large complex that includes components of the DNA methylation regulatory system at enhancer regions governed by FOX family members, a subset of which may regulate lineage choice under the selective pressure of ADT in PCa cells. Interestingly, recurrent tumors in the CWR22 model from the shCTL group exhibited reduced expression of NCOR2, similar to the shNCOR2 recurrent tumors. Yet, only the shNCOR2 recurrent tumors exhibited increased neuroendocrine characteristics. While both groups of recurrent tumors end up with reduced NCOR2 expression, the shCTL tumors had normal NCOR2 expression levels at the time of ADT. These observations suggest that the timing of NCOR2 reduction relative to the selective pressure of ADT is important in altering the potential for lineage plasticity.

GSEA and MRA, DMR-enhancer identification, and assessment of NCOR2 cistromes all strongly implicated FOX factor function overlap with the NCOR2 dependent epigenome. FOXA1 is strongly linked to AR function in PCa cells through its actions as a pioneering factor^65,66^. FOXA1 has the capacity to bind nucleosomal DNA and shape global chromatin patterns in a manner that ultimately dictates lineage specific enhancer landscapes and occupancy^67^. Its pioneering function is also linked to the neuronal differentiation program driven by N-Myc^68^, suggesting it as a key component in governing lineage plasticity in PCa cells, which is increasingly recognized as a contributor to ADT-RPCa ^13,69^. Indeed, our studies demonstrated that NCOR2 loss significantly reduced response to ADT in a large cohort of CWR22 xenografts and led to recurrent tumors with more neuroendocrine characteristics. These PDX results align with our cell line findings that altered NCOR2 expression impacted both androgen response gene sets, as well as genes with N-Myc MR enrichment and gene sets associated with neuronal pathways. These findings suggest that levels of NCOR2 play a role in regulating the ability of FOXA1 to regulate the chromatin landscape of enhancers dictating cell lineage. ^13,69^.

Combined these observations in CWR22 and in human tumors of reduced NCOR2 expression associating with decreased maximum regression and faster time to recurrence, suggest that the reduction in NCOR2 makes some cells resistant to the effects of androgen depletion. However, RNA-seq from the 7-days post androgen withdrawal showed no evidence of NCOR2 reduction associating with expression of androgen regulated genes (Fig S10C). This might be attributed to the fact that despite a significant increase in Ki-67 positive cells, the remain a minority of the cell population contributing to the RNA-seq data.

## Supplementary Figure Legends

**Figure S1: A)** Associations of Gleason sum (left), pathologic stage (middle) and adjuvant ADT (right) with BCR survival. Overall log-rank test is shown, and significance of individual comparisons from univariate regression analysis are noted. **B)** Distribution of Gleason scores between patients that did or did not receive adjuvant ADT. Significance was determined by chi-square test. **C)** BCR survival assessment of patients with high and low expression of NCOR2 (left) and NCOR2 (right). NCOR associated survival was compared within sub-cohorts of patients stratified by Gleason sum and **D)** pathologic stage.

**Figure S2: A)** NCOR2 gene expression in LNCaP-shCTL and LNCaP-shNCOR2 cells. **B)** NCOR2 protein expression (below) and quantification (above) in LNCaP-shCTL and LNCaP-shNCOR2 cells. **C)** NCOR2 gene expression in C4-2-shCTL and C4-2-shNCOR2 cells. **D)** NCOR2 protein expression (below) and quantification (above) in C4-2-shCTL and C4-2-shNCOR2 cells. **E)** Dose response of LNCaP (blue) and C4-2 (red) to enzalutamide (left) and R1881 (right). **F)** NCOR2 gene expression in shCTL and shNCOR2 cell lines exposed to DHT.

**Figure S3: A)** Distance matrices for LNCaP (left) and C4-2 (right) samples based on total gene expression profiles. **B)** Principal component analyses for LNCaP (left) and C4-2 (right) samples based on total gene expression profiles.

**Figure S4: A)** Venn diagram of DHT mediated DEGs (top) and NCOR2 mediated DEGs (bottom) between cell lines. **B)** Scatterplot of DHT induced fold changes in shCTL cells and shNCOR2 cells in LNCaP (top) and C4-2 (bottom). Pearson correlation is shown. **C)** Scatterplot of gene expression change induced by NCOR2 knockdown (y-axis) or DHT exposure (x-axis) in LNCaP (top) and C4-2 (bottom). Red genes are those found to be both NCOR2 and DHT dependent.

**Figure S5:** Pathway enrichment maps for the top 50 significantly up (red) and down (blue) regulated pathways for both NCOR2 associated DEGs (left column) and DHT associated DEGs (right column).

**Figure S6: A)** Expression profile for miR-10a-5p determined by RT-qPCR (top) and small RNA-seq (middle). Summary of RT-qPCR validation shown in table (bottom). **B)** Expression of candidate miRNA (miR-200a-3p, let-7e-5p) determined as significantly upregulated upon NCOR2 knockdown. **C)** Network of all genes downregulated upon NCOR2 knockdown, showing subsets of genes containing miR-200a-3p and/or let-7e-5p target sequences.

**Figure S7: A)** Distance matrix for LNCaP (left) and C4-2 (right) samples based on total DNA methylation profiles. **B)** Principal component analyses for LNCaP (left) and C4-2 (right) samples based on total DNA methylation profiles. **C)** Volcano plots representing methylation changes identified upon DHT exposure in LNCaP (top) and C4-2 (bottom) cells. Determined DMPs are shown in blue (hypomethylated) and red (hypermethylated).

**Figure S8: A)** Peak centered densities of non-DMP CpG sites (black) or hypermethylated DMPs (red) centered at chromHMM regions. **B)** Venn diagram of DMRs identified in each cell. **C)** Venn diagram of annotated DMR-enhancer and DMR-promoter genes identified in LNCaP (top) and C4-2 (bottom). **D)** Representative genomic view of an NCOR2 dependent gene locus with annotated enhancer hypermethylation (*TRIP10, TLE6*). **E)** The top 30 most significant upregulated pathways from GSEA analysis DMR-enhancer associated genes in both LNCaP (left column) and C4-2 (right column)

**Figure S9: A)** Proportional annotations of determined NCOR2 cistromes to gene regions. **B)** Density plot of NCOR2 binding around TSS loci. **C)** Motif enrichments, comparing cistromes in untreated (left) and DHT treated (right) cells between cell lines. **D)** TF family motif enrichment analysis in LNCaP (left) and C4-2 (right). **E)** Scatterplot representing total overlaps identified between LNCaP (left) and C4-2 (right) cistromes and all transcription factor datasets available in the CistromeDB. The top 200 enriched datasets are shown in red. **F)** The top 20 most frequently observed factors within the top 200 enriched datasets for each NCOR2 cistrome. The observed frequencies (blue) were compared to the background frequencies (red), and factors occurring at rates similar to background (< 1.2x enrichment) were removed from subsequent analyses.

**Figure S10:** The number of peak:gene relationships observed for each NCOR2 cistrome within specific genomic subsets.

**Figure S11: A)** Absolute tumor growth and **B)** androgen withdrawal normalized growth of all tumors in the study. **C)** RT-qPCR of select androgen regulated genes at different stages of disease progression in shCTL and shNCOR2 tumors.

**Figure S12: A)** Volcano plots depicting NCOR2-dependent DEGs (red = upregulated, blue = downregulated) determined at different stages of disease. **B)** Volcano plots depicting NCOR2-dependent DMPs (red = upregulated, blue = downregulated) determined at different stages of disease. **C)** GSEA summary of the top enriched pathways associated with DEGs identified in post-AD and RT relative to pre-AD tumors, and NCOR2-dependent DEGs at each stage of disease.

**Figure S13: A)** Volcano plots representing methylation changes identified in human PCa samples (androgen independent (AI) relative to androgen dependent (AD)). Determined DMPs are shown in blue (hypomethylated) and red (hypermethylated). **B)** Relative enrichment of DMPs within chromHMM categories for respective comparisons.

## Acknowledgements

Shared Resources

Genomics Shared Resource (sequencing)

Gene Modulation Services (shRNA cell lines)

Pathology Network Shared Resource (TMA)

Translational Imaging Shared Resource (in vivo imaging)

Small Molecule Screening Shared Resource (drug screen)

Barbara Foster / Bryan Gillard / Ellen Karasik (Mouse work)

