## Supplemental figures for "Reduced NCOR2 expression accelerates androgen deprivation therapy failure in prostate cancer"

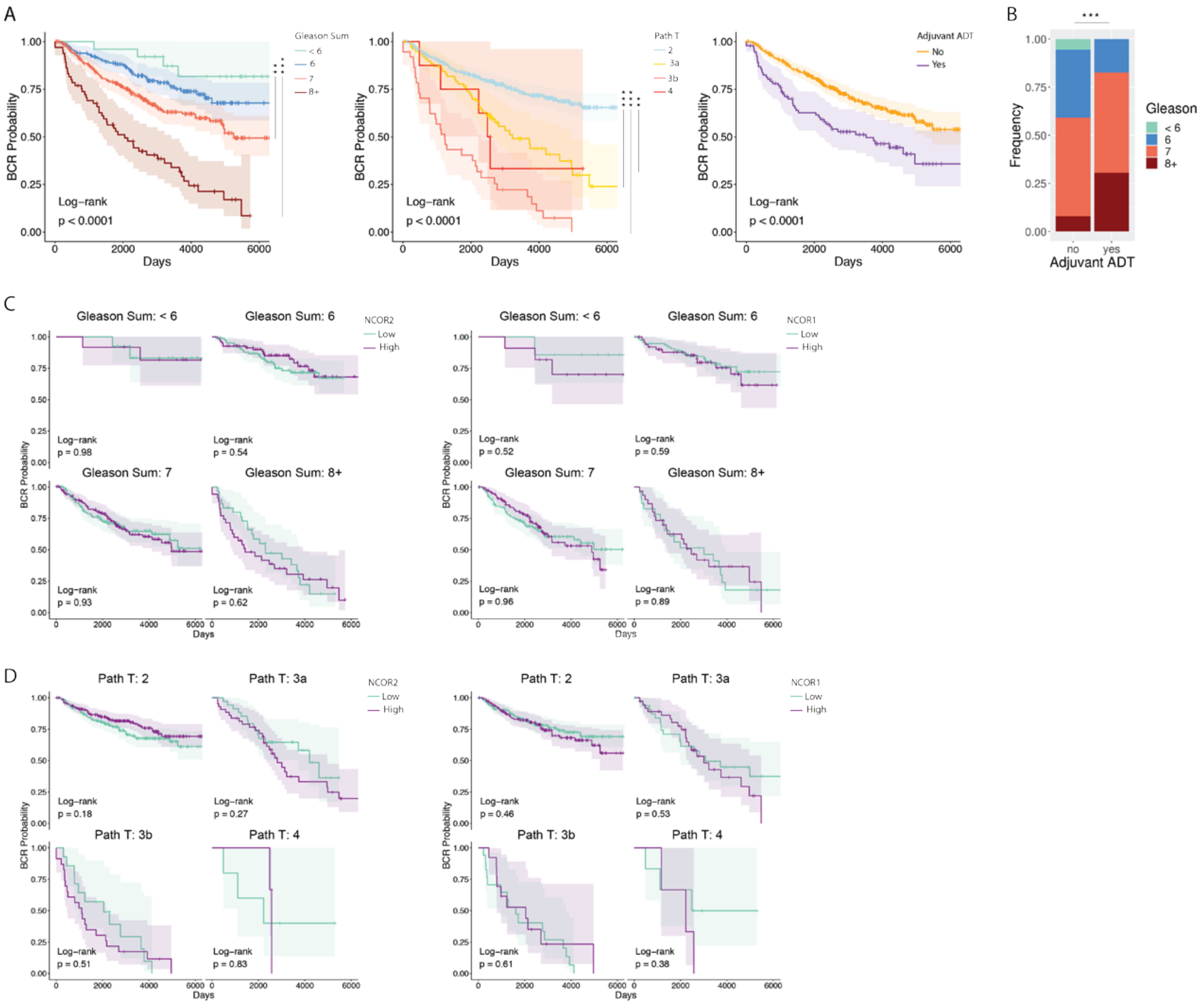

A
BCR Probability
Days
Gleason Sum
< 6
6
7
8+
Log−rank
p < 0.0001
Path T
2
3a
3b
4
Adjuvant ADT
No
Yes
B
Frequency
Gleason
no
yes
Adjuvant ADT
C
Gleason Sum: < 6
Gleason Sum: 6
Gleason Sum: 7
Gleason Sum: 8+
NCOR2
Low
High
NCOR1
p = 0.98
p = 0.54
p = 0.93
p = 0.62
p = 0.52
p = 0.59
p = 0.96
p = 0.89
D
Path T: 2
Path T: 3a
Path T: 3b
Path T: 4
p = 0.18
p = 0.27
p = 0.51
p = 0.83
p = 0.46
p = 0.53
p = 0.61
p = 0.38


Figure S1

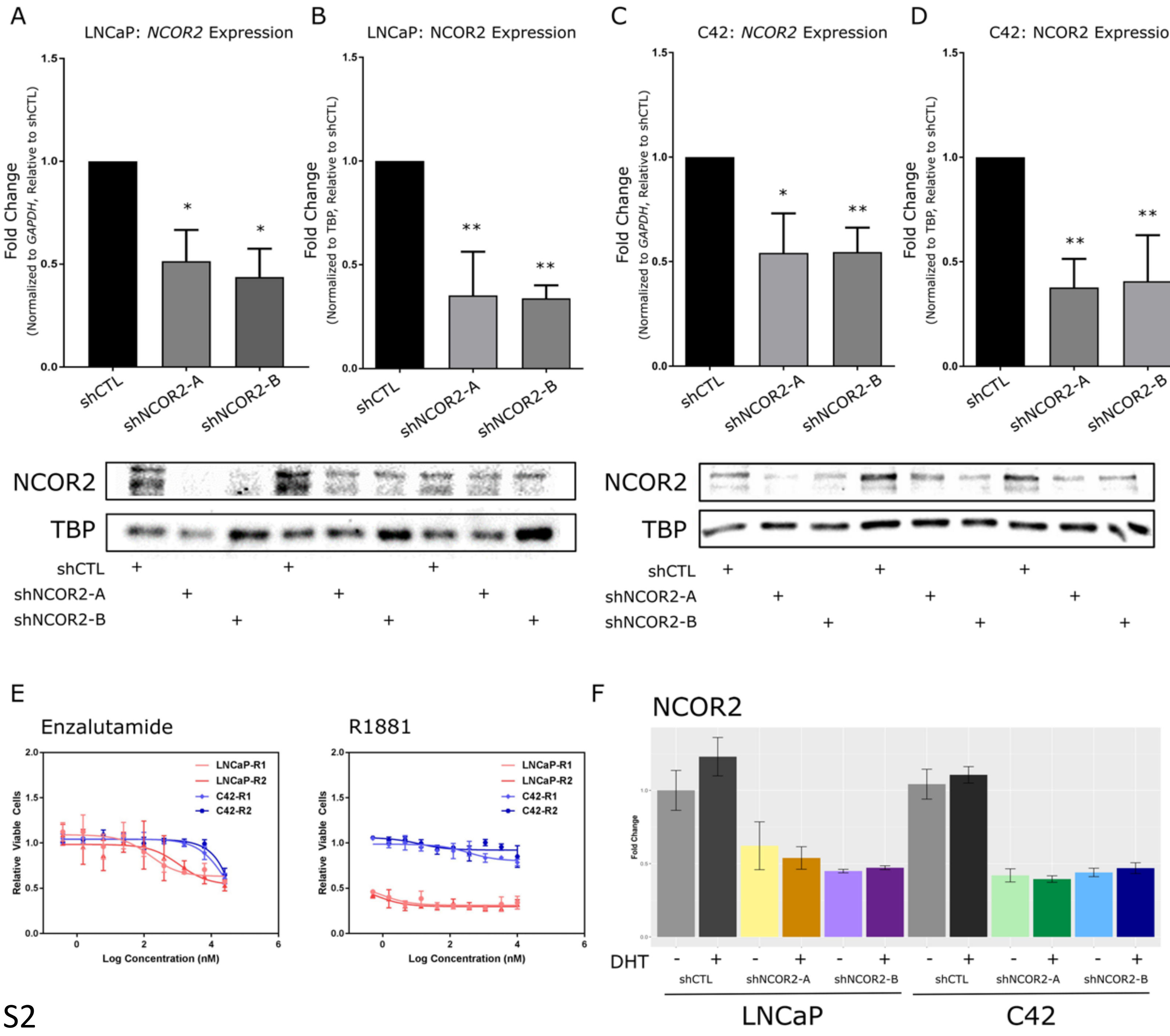
A
LNCaP: NCOR2 Expression
Fold Change
(Normalized to GAPDH, Relative to shCTL)
shCTL
shNCOR2-A
shNCOR2-B
B
LNCaP: NCOR2 Expression
(Normalized to TBP, Relative to shCTL)
C
C42: NCOR2 Expression
D
C42: NCOR2 Expression
NCOR2
TBP
E
Enzalutamide
R1881
LNCaP-R1
LNCaP-R2
C42-R1
C42-R2
Relative Viable Cells
Log Concentration (nM)
F
NCOR2
Fold Change
DHT
LNCaP
C42

Figure S2

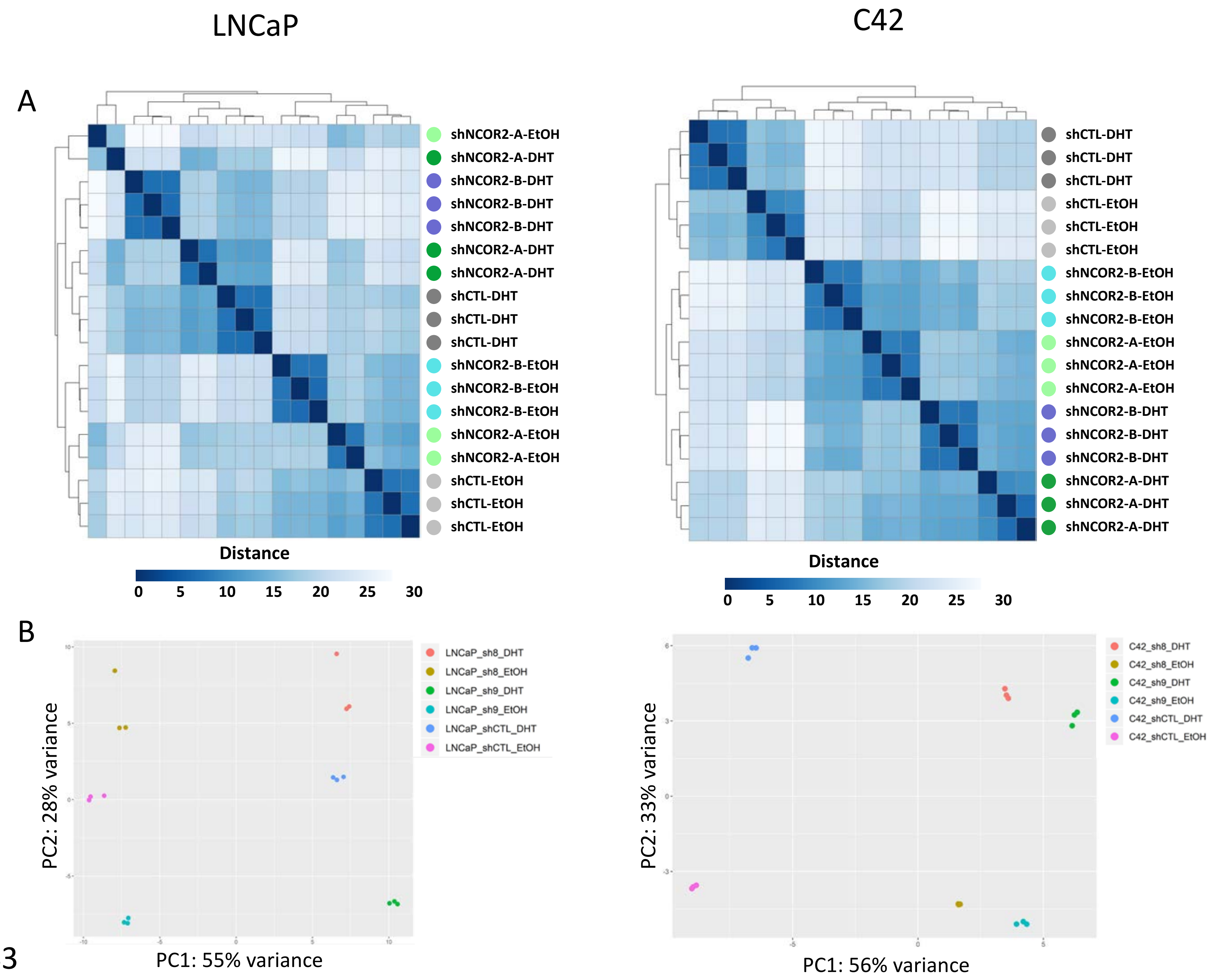

LNCaP
C42
A
shNCOR2-A-EtOH
shNCOR2-A-DHT
shNCOR2-B-DHT
shNCOR2-B-DHT
shNCOR2-B-DHT
shNCOR2-A-DHT
shNCOR2-A-DHT
shCTL-DHT
shCTL-DHT
shCTL-DHT
shNCOR2-B-EtOH
shNCOR2-B-EtOH
shNCOR2-B-EtOH
shNCOR2-A-EtOH
shNCOR2-A-EtOH
shCTL-EtOH
shCTL-EtOH
shCTL-EtOH
Distance
0 5 10 15 20 25 30
shCTL-DHT
shCTL-DHT
shCTL-DHT
shCTL-EtOH
shCTL-EtOH
shCTL-EtOH
shNCOR2-B-EtOH
shNCOR2-B-EtOH
shNCOR2-B-EtOH
shNCOR2-A-EtOH
shNCOR2-A-EtOH
shNCOR2-A-EtOH
shNCOR2-B-DHT
shNCOR2-B-DHT
shNCOR2-B-DHT
shNCOR2-A-DHT
shNCOR2-A-DHT
shNCOR2-A-DHT
Distance
0 5 10 15 20 25 30
B
LNCaP_sh8_DHT
LNCaP_sh8_EtOH
LNCaP_sh9_DHT
LNCaP_sh9_EtOH
LNCaP_shCTL_DHT
LNCaP_shCTL_EtOH
PC2: 28% variance
PC1: 55% variance
C42_sh8_DHT
C42_sh8_EtOH
C42_sh9_DHT
C42_sh9_EtOH
C42_shCTL_DHT
C42_shCTL_EtOH
PC2: 33% variance
PC1: 56% variance


Figure S3

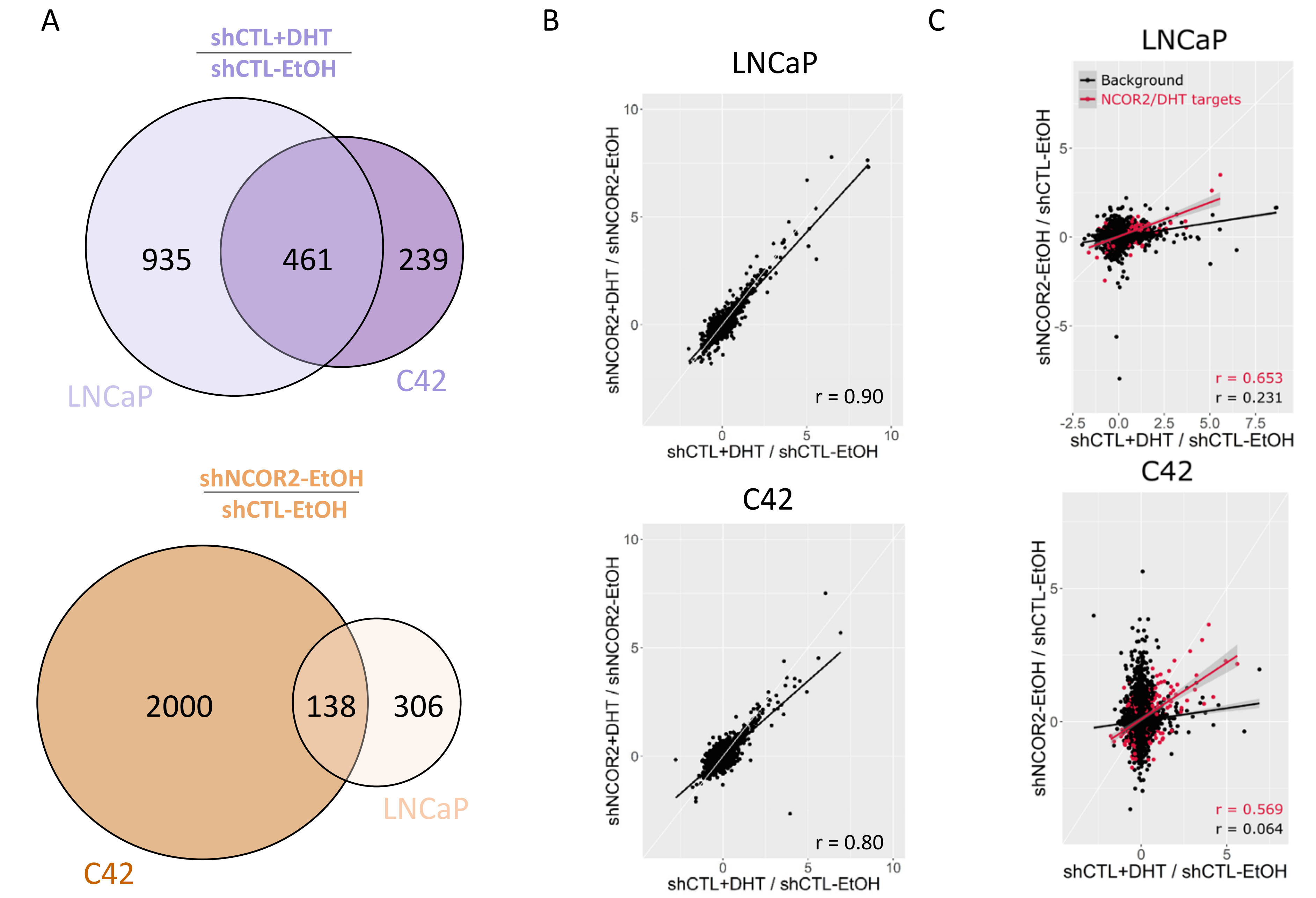
A
shCTL+DHT
shCTL-EtOH
935
461
239
LNCaP
C42
shNCOR2-EtOH
shCTL-EtOH
2000
138
306
LNCaP
C42
B
LNCaP
shNCOR2+DHT / shNCOR2-EtOH
shCTL+DHT / shCTL-EtOH
r = 0.90
C42
shNCOR2+DHT / shNCOR2-EtOH
shCTL+DHT / shCTL-EtOH
r = 0.80
C
LNCaP
Background
NCOR2/DHT targets
shNCOR2-EtOH / shCTL-EtOH
shCTL+DHT / shCTL-EtOH
r = 0.653
r = 0.231
C42
shNCOR2-EtOH / shCTL-EtOH
shCTL+DHT / shCTL-EtOH
r = 0.569
r = 0.064

Figure S4

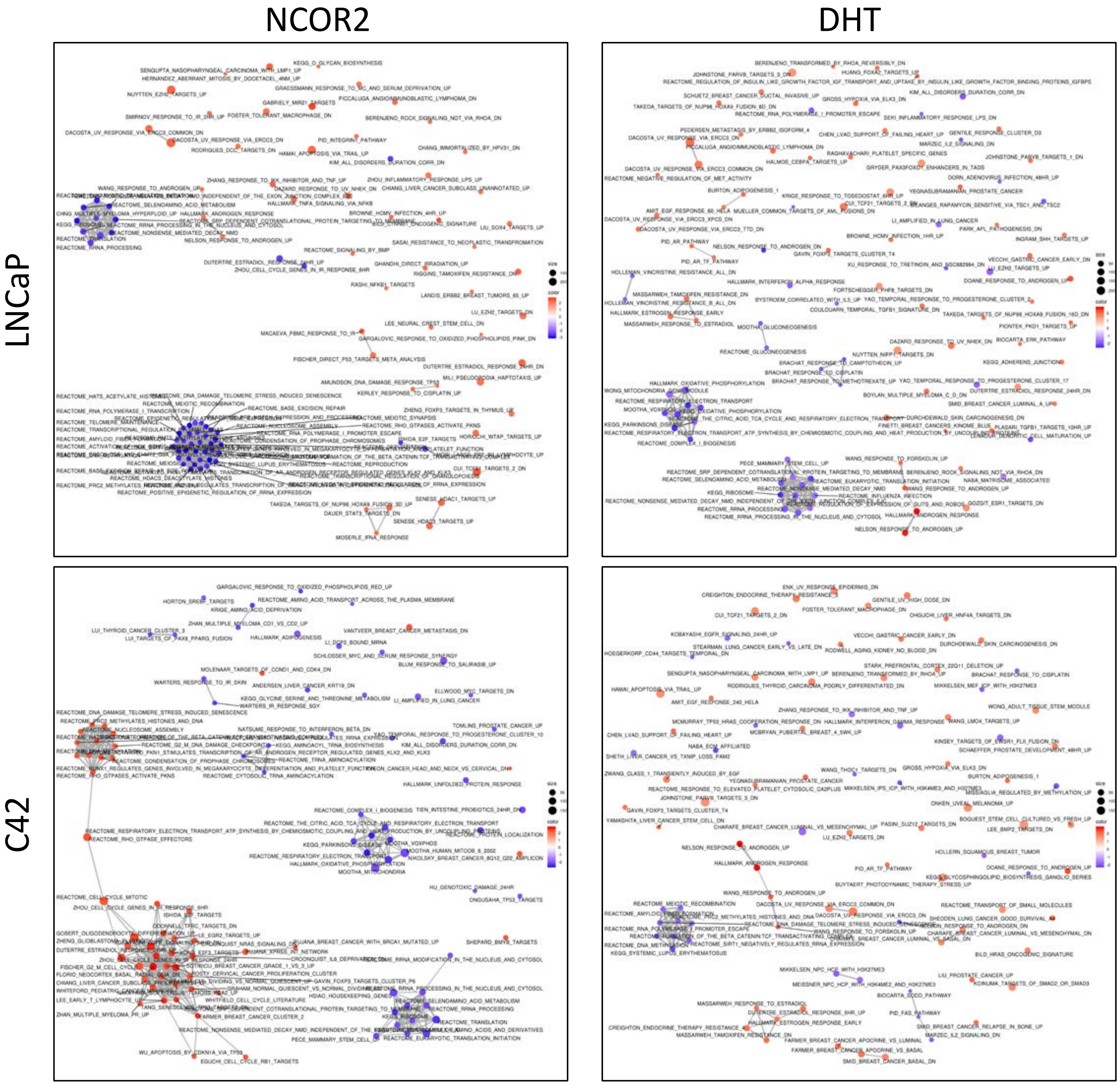


Figure S5

A

miR-10a-5p: qRT-PCR

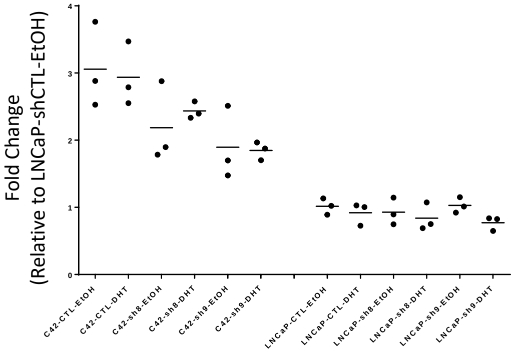


miR-10a-5p: Small RNA-seq

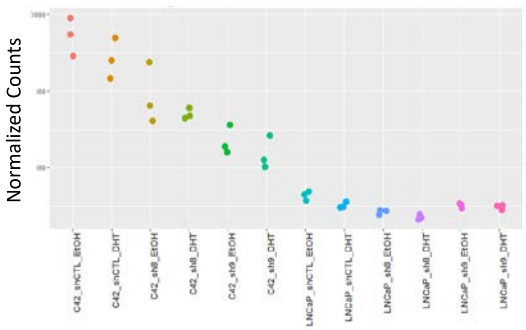


| | miRNA-seq | miRNA-RTqPCR | PCR-pval |
|---|---|---|---|
| **miR-96-5p** | UP | UP | 0.0011 |
| **miR-125b-5p** | DOWN | DOWN | 0.0020 |
| **miR-193b-3p** | DOWN | DOWN | 0.017 |
| **miR-10a-5p** | DOWN | DOWN | 0.046 |
| **let-7e-5p** | UP | UP | 0.044 |
| **miR-126-3p** | UP | UP | 0.097 |
| **miR-200a-3p** | UP | UP | 0.023 |
| **miR-22-3p** | DOWN | NS | 0.79 |

B

C4-2

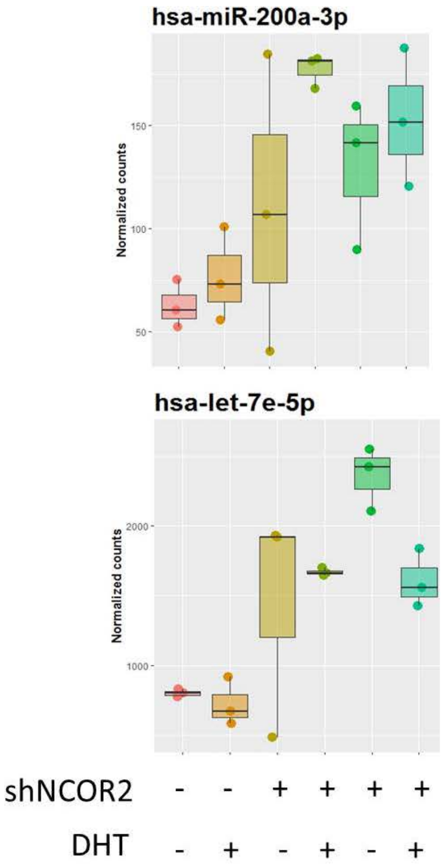


C

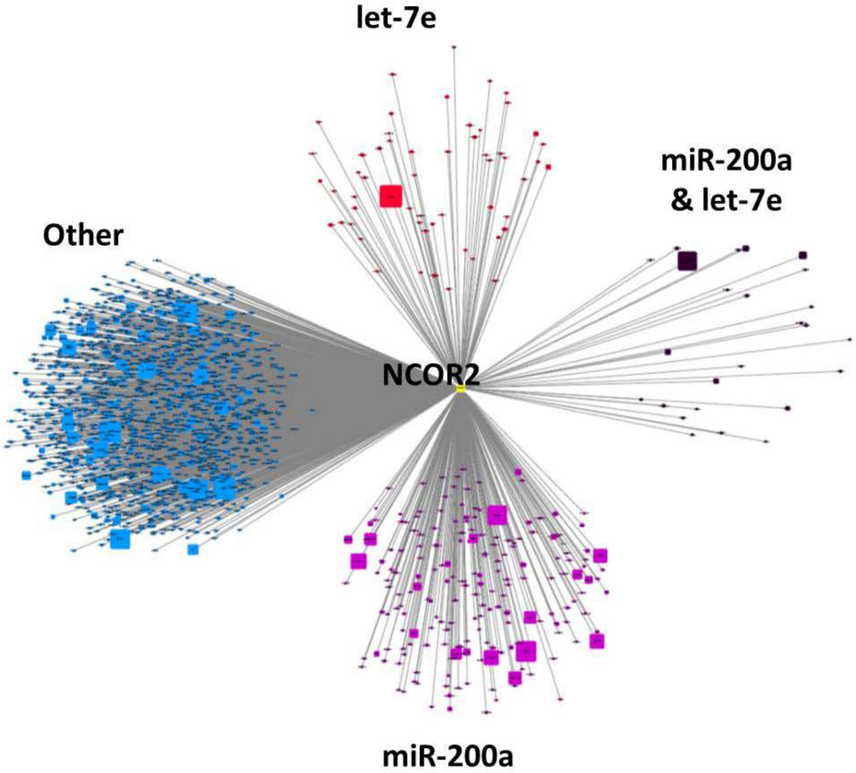


Figure S6

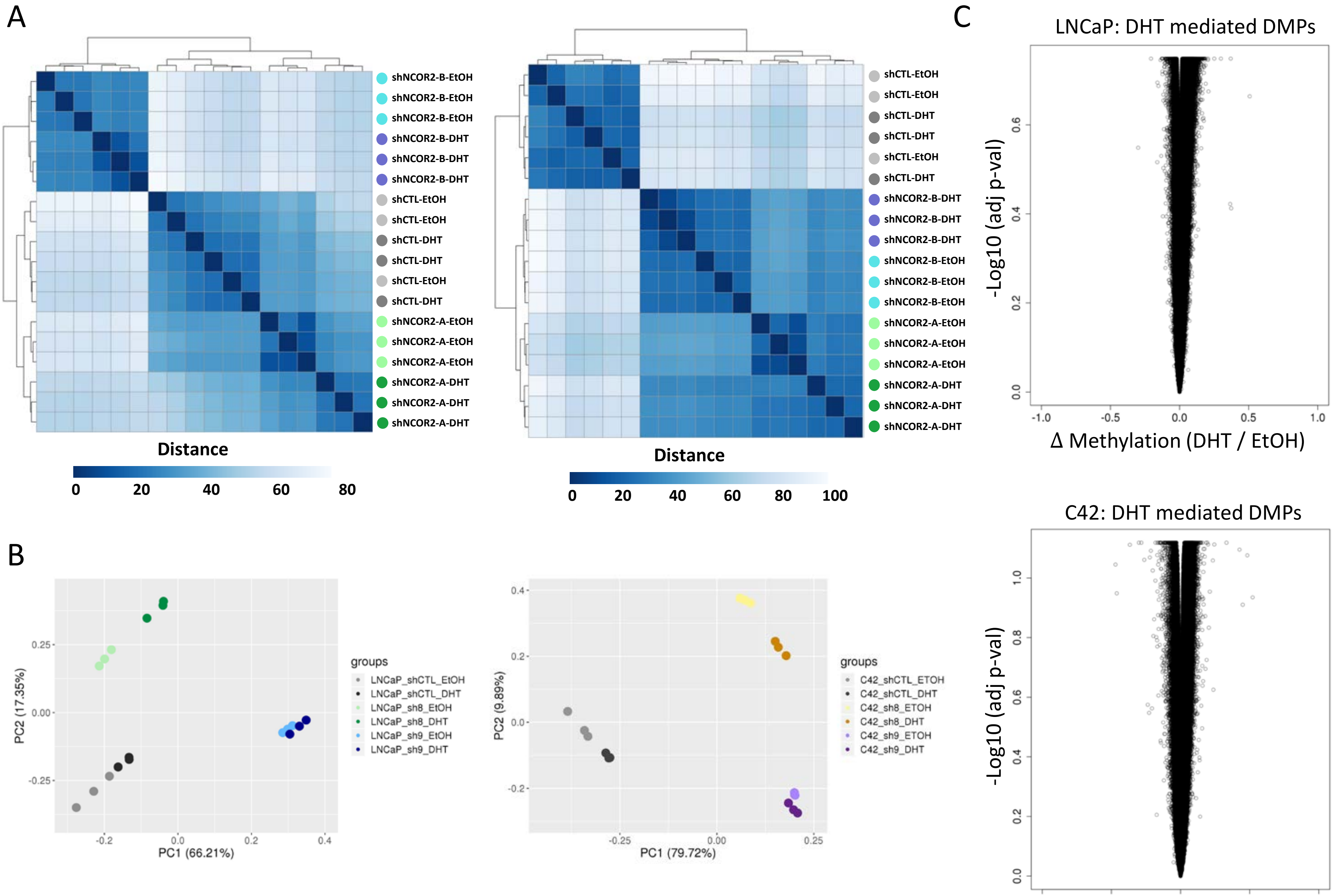
A
shNCOR2-B-EtOH
shNCOR2-B-EtOH
shNCOR2-B-EtOH
shNCOR2-B-DHT
shNCOR2-B-DHT
shNCOR2-B-DHT
shCTL-EtOH
shCTL-EtOH
shCTL-DHT
shCTL-DHT
shCTL-EtOH
shCTL-DHT
shNCOR2-A-EtOH
shNCOR2-A-EtOH
shNCOR2-A-EtOH
shNCOR2-A-DHT
shNCOR2-A-DHT
shNCOR2-A-DHT
Distance
0 20 40 60 80
shCTL-EtOH
shCTL-EtOH
shCTL-DHT
shCTL-DHT
shCTL-EtOH
shCTL-DHT
shNCOR2-B-DHT
shNCOR2-B-DHT
shNCOR2-B-DHT
shNCOR2-B-EtOH
shNCOR2-B-EtOH
shNCOR2-B-EtOH
shNCOR2-A-EtOH
shNCOR2-A-EtOH
shNCOR2-A-EtOH
shNCOR2-A-DHT
shNCOR2-A-DHT
shNCOR2-A-DHT
Distance
0 20 40 60 80 100
B
PC2 (17.35%)
PC1 (66.21%)
groups
LNCaP_shCTL_EtOH
LNCaP_shCTL_DHT
LNCaP_sh8_EtOH
LNCaP_sh8_DHT
LNCaP_sh9_EtOH
LNCaP_sh9_DHT
PC2 (9.89%)
PC1 (79.72%)
groups
C42_shCTL_ETOH
C42_shCTL_DHT
C42_sh8_ETOH
C42_sh8_DHT
C42_sh9_ETOH
C42_sh9_DHT
C
LNCaP: DHT mediated DMPs
-Log10 (adj p-val)
Δ Methylation (DHT / EtOH)
C42: DHT mediated DMPs
-Log10 (adj p-val)
Δ Methylation (DHT / EtOH)

Figure S7

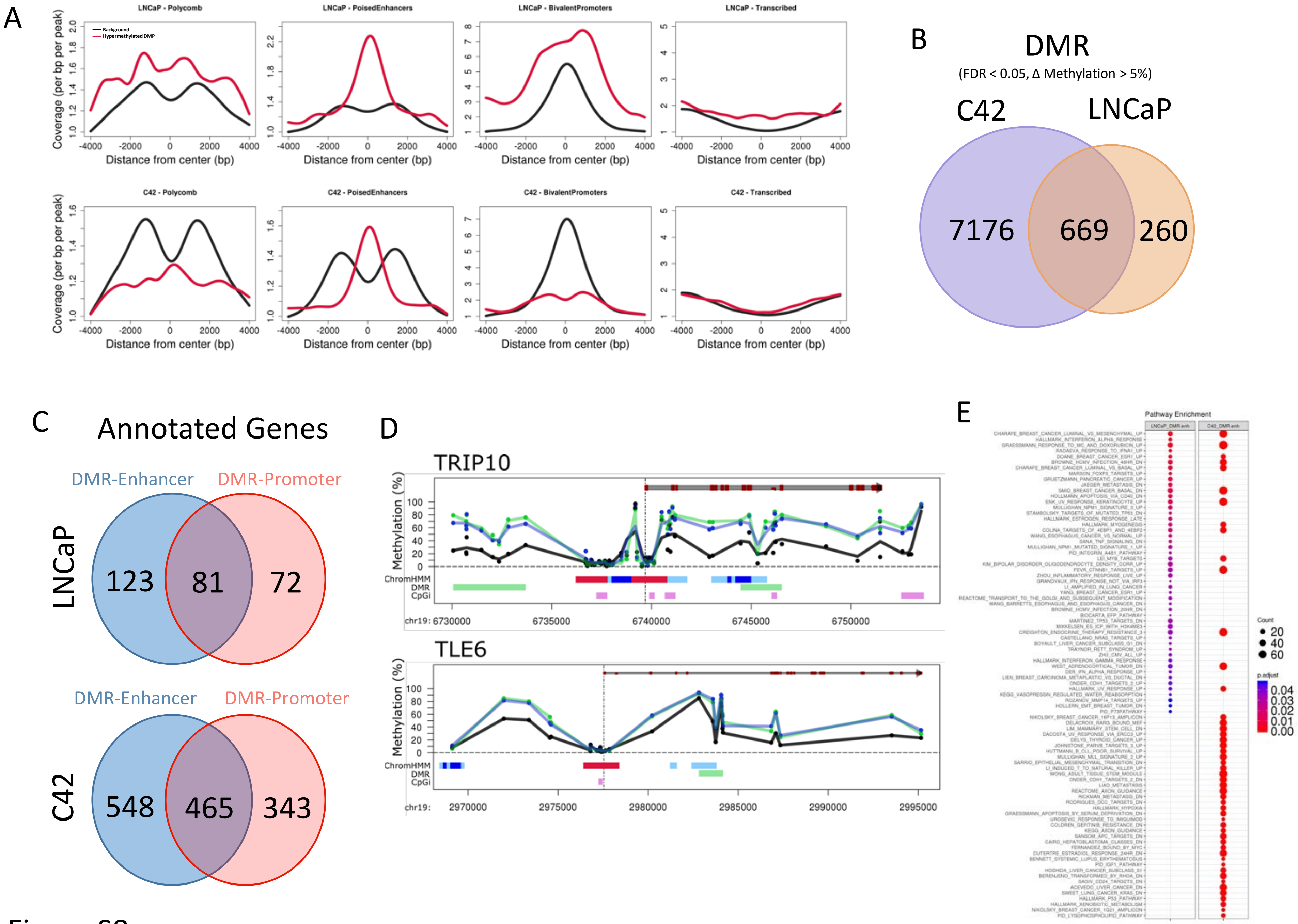

A
LNCaP - Polycomb
LNCaP - PoisedEnhancers
LNCaP - BivalentPromoters
LNCaP - Transcribed
Background
Hypermethylated DMP
Coverage (per bp per peak)
Distance from center (bp)
C42 - Polycomb
C42 - PoisedEnhancers
C42 - BivalentPromoters
C42 - Transcribed
B
DMR
(FDR < 0.05, Δ Methylation > 5%)
C42
LNCaP
7176
669
260
C
Annotated Genes
DMR-Enhancer
DMR-Promoter
LNCaP
123
81
72
DMR-Enhancer
DMR-Promoter
C42
548
465
343
D
TRIP10
Methylation (%)
ChromHMM
DMR
CpGi
chr19: 6730000
6735000
6740000
6745000
6750000
TLE6
chr19:
2970000
2975000
2980000
2985000
2990000
2995000
E
Pathway Enrichment
Count
20
40
60
p.adjust
0.04
0.03
0.02
0.01
0.00


Figure S8

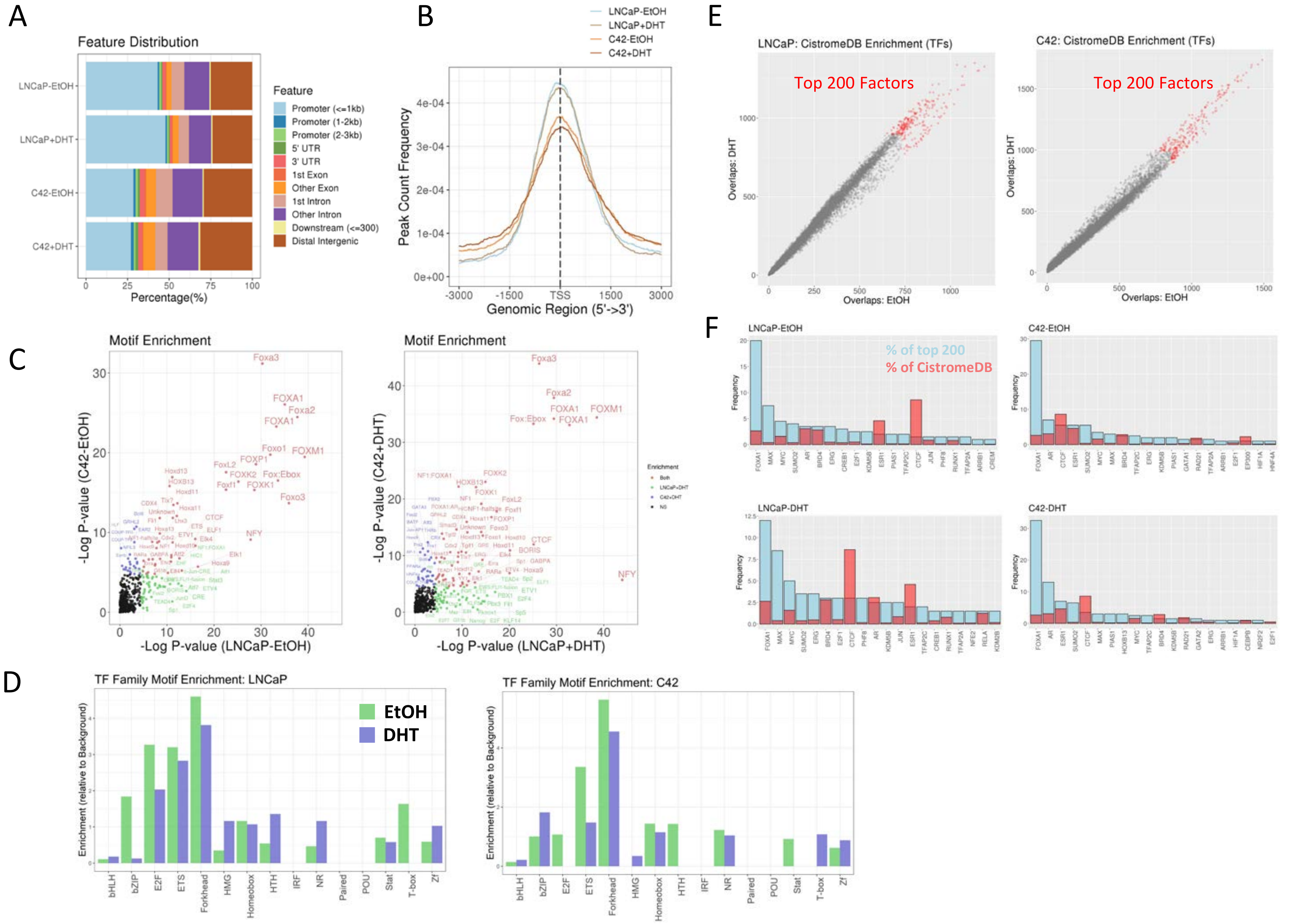
A
Feature Distribution
LNCaP-EtOH
LNCaP+DHT
C42-EtOH
C42+DHT
Percentage(%)
Feature
Promoter (<=1kb)
Promoter (1-2kb)
Promoter (2-3kb)
5' UTR
3' UTR
1st Exon
Other Exon
1st Intron
Other Intron
Downstream (<=300)
Distal Intergenic
B
LNCaP-EtOH
LNCaP+DHT
C42-EtOH
C42+DHT
Peak Count Frequency
Genomic Region (5'->3')
E
LNCaP: CistromeDB Enrichment (TFs)
C42: CistromeDB Enrichment (TFs)
Top 200 Factors
Overlaps: DHT
Overlaps: EtOH
C
Motif Enrichment
-Log P-value (C42-EtOH)
-Log P-value (LNCaP-EtOH)
-Log P-value (C42+DHT)
-Log P-value (LNCaP+DHT)
F
LNCaP-EtOH
C42-EtOH
LNCaP-DHT
C42-DHT
% of top 200
% of CistromeDB
Frequency
D
TF Family Motif Enrichment: LNCaP
TF Family Motif Enrichment: C42
EtOH
DHT
Enrichment (relative to Background)
bHLH
bZIP
E2F
ETS
Forkhead
HMG
Homeobox
HTH
IRF
NR
Paired
POU
Stat
T-box
Zf

Figure S9

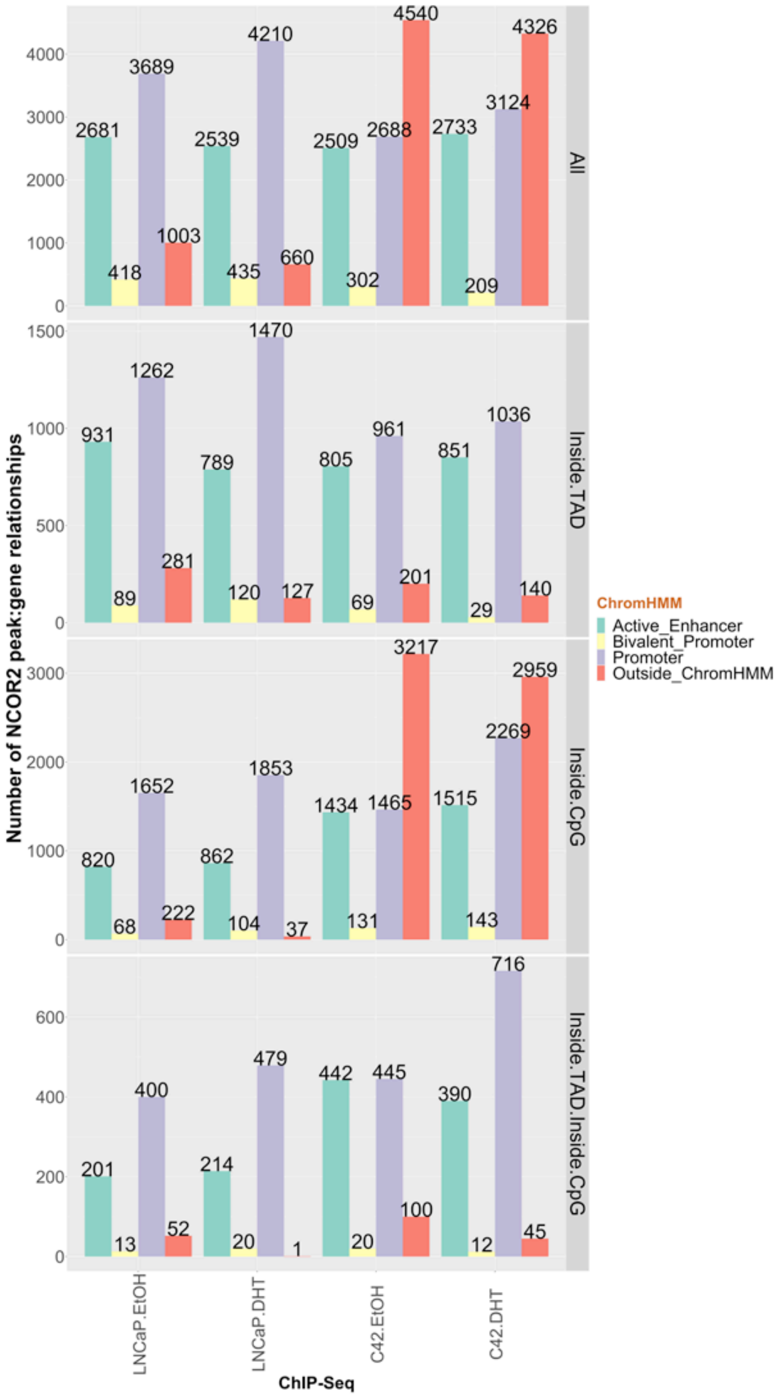
Number of NCOR2 peak:gene relationships
ChIP-Seq
LNCaP.EtOH
LNCaP.DHT
C42.EtOH
C42.DHT
All
Inside.TAD
Inside.CpG
Inside.TAD.Inside.CpG
ChromHMM
Active_Enhancer
Bivalent_Promoter
Promoter
Outside_ChromHMM
2681
418
3689
1003
2539
435
4210
660
2509
302
2688
4540
2733
209
3124
4326
931
89
1262
281
789
120
1470
127
805
69
961
201
851
29
1036
140
820
68
1652
222
862
104
1853
37
1434
131
1465
3217
1515
143
2269
2959
201
13
400
52
214
20
479
1
442
20
445
100
390
12
716
45

Figure S10

A

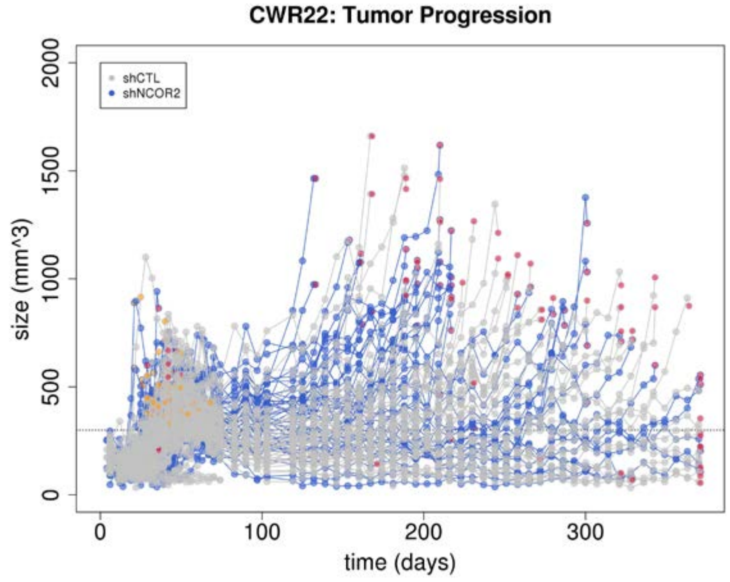


B

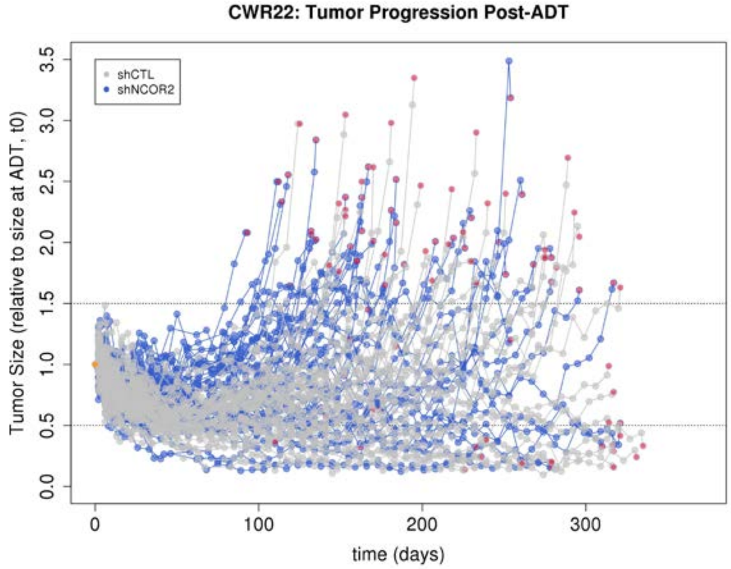


C

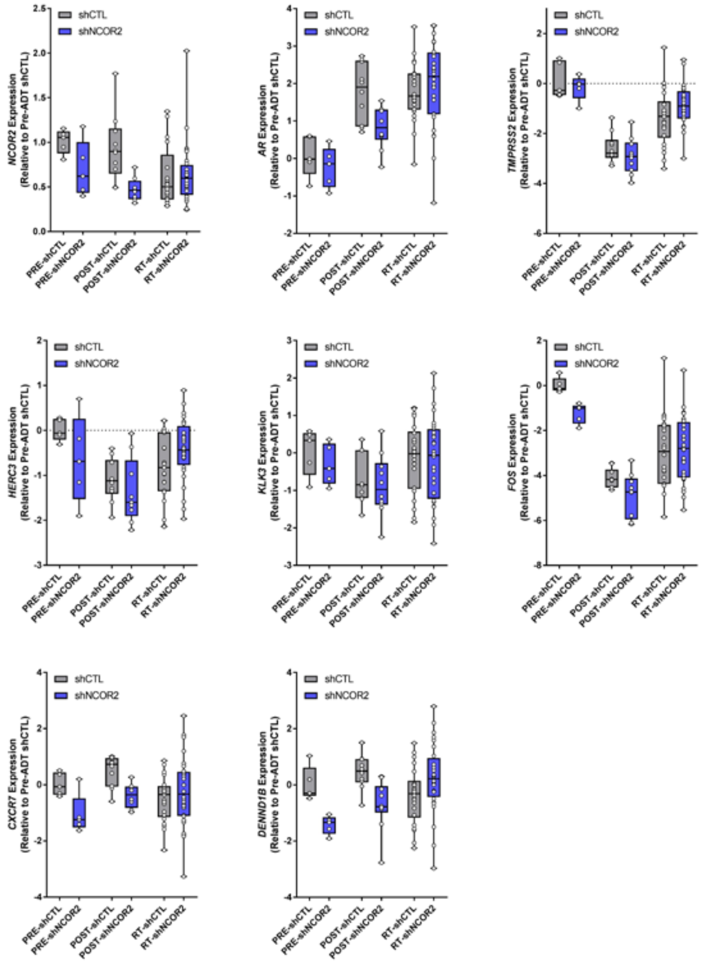


Figure S11

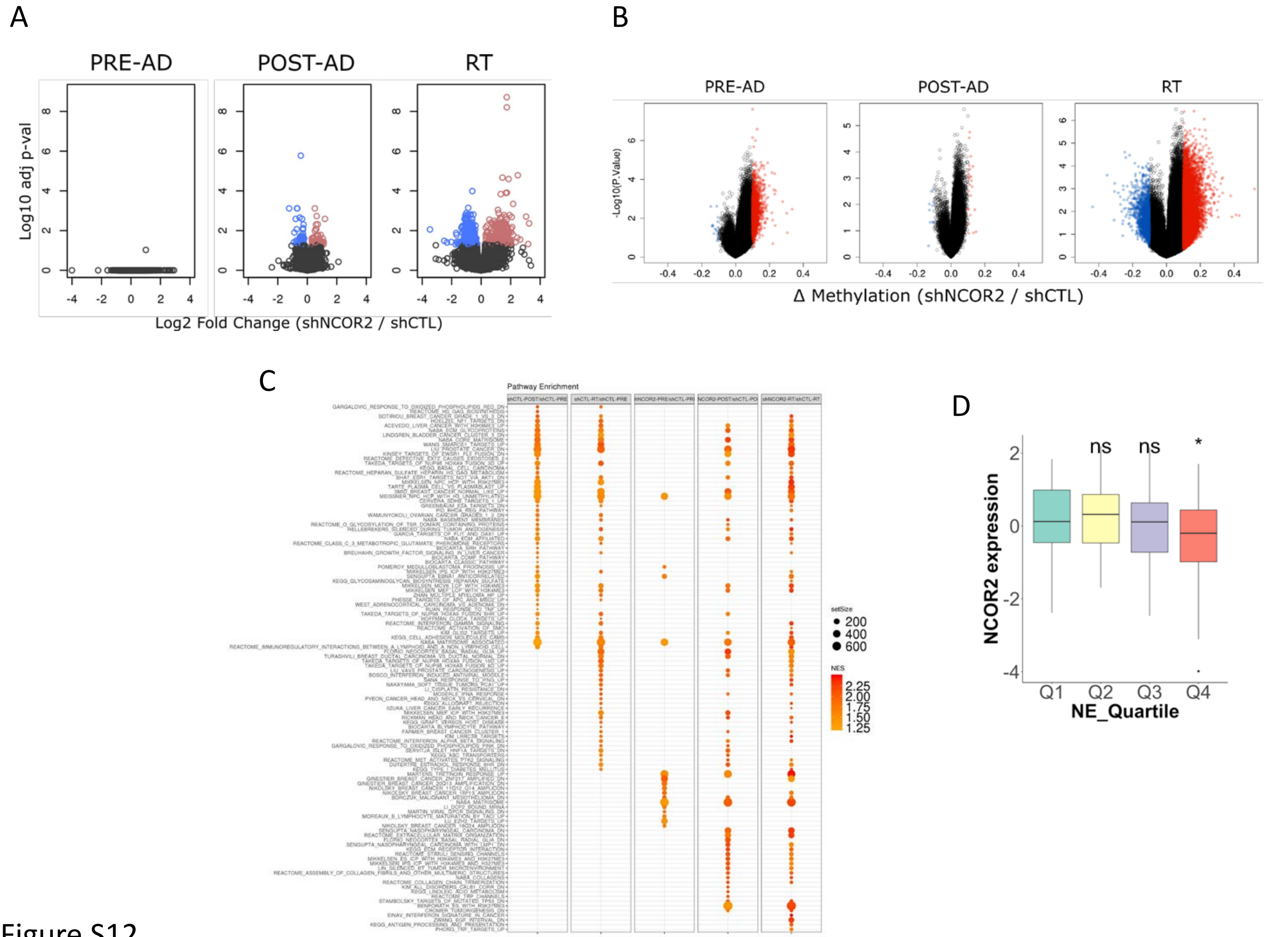
A
PRE-AD
POST-AD
RT
Log10 adj p-val
Log2 Fold Change (shNCOR2 / shCTL)
B
PRE-AD
POST-AD
RT
-Log10(P.Value)
Δ Methylation (shNCOR2 / shCTL)
C
Pathway Enrichment
setSize
200
400
600
NES
2.25
2.00
1.75
1.50
1.25
D
ns
ns
*
NCOR2 expression
Q1
Q2
Q3
Q4
NE_Quartile

Figure S12

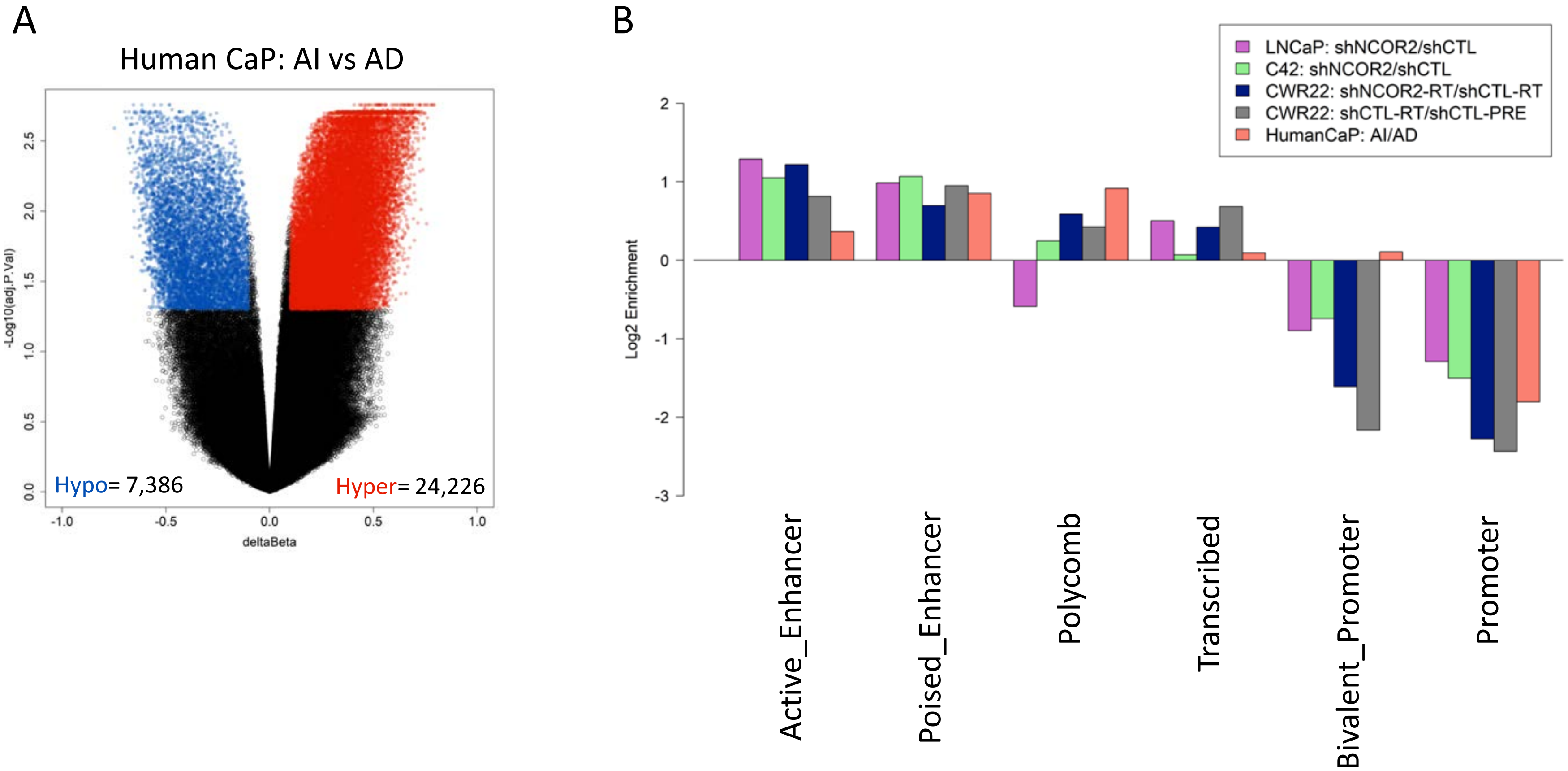

A
Human CaP: AI vs AD
-Log10(adj.P.Val)
deltaBeta
Hypo= 7,386
Hyper= 24,226
B
LNCaP: shNCOR2/shCTL
C42: shNCOR2/shCTL
CWR22: shNCOR2-RT/shCTL-RT
CWR22: shCTL-RT/shCTL-PRE
HumanCaP: AI/AD
Log2 Enrichment
Active_Enhancer
Poised_Enhancer
Polycomb
Transcribed
Bivalent_Promoter
Promoter


Figure S13
