## Supplemental tables for "Reduced NCOR2 expression accelerates androgen deprivation therapy failure in prostate cancer"

Table S1

| | Total | | | | | |
|---|---|---|---|---|---|---|
| NCOR2 | 564 | | | | | |
| NCOR1 | 463 | | | | | |
| **Hscore** | **Min** | **Q25** | **Median** | **Q75** | **Max** | |
| NCOR2 | 0 | 55.06 | 91.5 | 127.57 | 246 | |
| NCOR1 | 15.87 | 123.97 | 167.03 | 200.14 | 268.84 | |
| **BMI** | **Min** | **Q25** | **Median** | **Q75** | **Max** | **NA** |
| NCOR2 | 18.46 | 25.63 | 28.05 | 30.86 | 51.42 | 10 |
| NCOR1 | 18.46 | 25.77 | 28.06 | 31.12 | 65.01 | 10 |
| **Pre-RP PSA** | **Min** | **Q25** | **Median** | **Q75** | **Max** | **NA** |
| NCOR2 | 0 | 4.25 | 5.6 | 8 | 41.9 | 38 |
| NCOR1 | 0 | 4.47 | 5.9 | 8.5 | 35.7 | 34 |
| **Age at surgery** | **Min** | **Q25** | **Median** | **Q75** | **Max** | |
| NCOR2 | 39.18 | 55.04 | 60.08 | 65.74 | 74.62 | |
| NCOR1 | 39.18 | 55.32 | 59.91 | 65.67 | 74.3 | |
| **Gleason Sum** | **< 6** | **6** | **7** | **8+** | | |
| NCOR2 | 26 | 183 | 290 | 65 | | |
| NCOR1 | 19 | 140 | 251 | 53 | | |
| **PathT** | **2X** | **3a** | **3b** | **4** | | |
| NCOR2 | 437 | 81 | 38 | 8 | | |
| NCOR1 | 353 | 71 | 30 | 9 | | |
| **Adjuvant ADT** | **No** | **Yes** | | | | |
| NCOR2 | 472 | 92 | | | | |
| NCOR1 | 397 | 66 | | | | |
| **Race** | **White** | **Black** | **Other** | **NA** | | |
| NCOR2 | 520 | 34 | 6 | 4 | | |
| NCOR1 | 425 | 30 | 5 | 3 | | |

### Table S2

**NCOR2: H-score Linear Regression Analysis**

| | *Dependent variable: Hscore* | | | | | | | |
|---|---|---|---|---|---|---|---|---|
| | Hscore | | | | | | | |
| | Race: White | BMI: Normal (< | PSA: < 2.5 | Gleason: < 6 | Adjuvant ADT: | Age: < 55 | Path Stage: 2 | Multivariate |
| Race: Black | -16.750* | | | | | | | -17.854* |
| | -8.752 | | | | | | | -9.319 |
| Race: Other | -2.485 | | | | | | | -2.206 |
| | -20.302 | | | | | | | -20.13 |
| BMI: Overweight (> 25; < 30) | | -17.147*** | | | | | | -18.168*** |
| | | -5.683 | | | | | | -5.759 |
| BMI: Obese (> 30) | | -8.942 | | | | | | -11.351* |
| | | -6.103 | | | | | | -6.267 |
| PSA: > 2.5; < 4 | | | -6.068 | | | | | 4.01 |
| | | | -9.416 | | | | | -9.8 |
| PSA: > 4; < 10 | | | -16.000** | | | | | -3.384 |
| | | | -6.938 | | | | | -7.638 |
| PSA: > 10 | | | -22.786*** | | | | | -14.662 |
| | | | -8.675 | | | | | -9.184 |
| Gleason: 6 | | | | -14.707 | | | | -22.052** |
| | | | | -10.376 | | | | -10.44 |
| Gleason: 7 | | | | -7.589 | | | | -14.234 |
| | | | | -10.135 | | | | -10.222 |
| Gleason: 8+ | | | | -4.755 | | | | -23.934** |
| | | | | -11.488 | | | | -12.08 |
| Adjuvant ADT: TRUE | | | | | 26.655*** | | | 25.459*** |
| | | | | | -5.539 | | | -6.511 |
| Age: 55-65 | | | | | | -6.573 | | -7.361 |
| | | | | | | -5.208 | | -5.362 |
| Age: 65+ | | | | | | -3.909 | | -6.004 |
| | | | | | | -5.707 | | -5.949 |
| Path Stage: 3b | | | | | | | 5.502 | 7.796 |
| | | | | | | | -5.981 | -6.331 |
| Path Stage: 3a | | | | | | | 17.113** | 11.617 |
| | | | | | | | -8.363 | -8.994 |
| Path Stage: 4 | | | | | | | -13.101 | -8.877 |
| | | | | | | | -17.641 | -17.655 |
| Constant | 94.810*** | 105.006*** | 108.433*** | 102.896*** | 89.326*** | 97.848*** | 91.916*** | 127.774*** |
| | -2.168 | -4.847 | -6.411 | -9.709 | -2.237 | -4.204 | -2.365 | -12.792 |
| Observations | 560 | 554 | 526 | 564 | 564 | 564 | 564 | 517 |
| $R^2$ | 0.007 | 0.017 | 0.017 | 0.007 | 0.04 | 0.003 | 0.01 | 0.095 |
| Adjusted $R^2$ | 0.003 | 0.014 | 0.011 | 0.002 | 0.038 | -0.001 | 0.004 | 0.066 |
| Residual Std. Error | 49.444 (df = | 49.194 (df = | 49.246 (df = | 49.507 (df = | 48.604 (df = 562) | 49.568 (df = | 49.445 (df = | 47.975 (df = 500) |
| F Statistic | 1.834 (df = 2; | 4.862 (df = 2; | 2.972 (df = 3; | 1.331 (df = 3; | 23.157 (df = 1; | 0.800 (df = 2; | 1.801 (df = 3; | 3.265 (df = 16; |
| *Note:* | | | | | | | | *p<0.1;**p<0.05;***p<0.01 |

### Table S3

**NCOR1: H-score Linear Regression Analysis**

| | *Dependent variable: Hscore* | | | | | | | |
|---|---|---|---|---|---|---|---|---|
| | Race: White | BMI: Normal (< | PSA: < 2.5 | Gleason: < 6 | Adjuvant ADT: | Age: < 55 | Path Stage: 2 | Multivariate |
| Race: Black | 9.59 | | | | | | | 7.827 |
| | -9.506 | | | | | | | -10.417 |
| Race: Other | 2.922 | | | | | | | 2.35 |
| | -22.637 | | | | | | | -22.685 |
| BMI: Overweight (> 25; < 30) | | -20.209$^{***}$ | | | | | | -17.869$^{***}$ |
| | | -6.471 | | | | | | -6.761 |
| BMI: Obese (> 30) | | -15.125$^{**}$ | | | | | | -13.577$^{*}$ |
| | | -6.862 | | | | | | -7.189 |
| PSA: > 2.5; < 4 | | | -19.452$^{*}$ | | | | | -7.494 |
| | | | -11.235 | | | | | -12.325 |
| PSA: > 4; < 10 | | | -19.959$^{**}$ | | | | | -10.174 |
| | | | -8.618 | | | | | -9.808 |
| PSA: > 10 | | | -14.692 | | | | | -7.4 |
| | | | -10.16 | | | | | -11.214 |
| Gleason: 6 | | | | -10.829 | | | | -15.784 |
| | | | | -12.212 | | | | -12.651 |
| Gleason: 7 | | | | -0.345 | | | | -5.436 |
| | | | | -11.885 | | | | -12.397 |
| Gleason: 8+ | | | | 8.776 | | | | -0.857 |
| | | | | -13.356 | | | | -14.305 |
| Adjuvant ADT: TRUE | | | | | 21.804$^{***}$ | | | 15.657$^{**}$ |
| | | | | | -6.599 | | | -7.931 |
| Age: 55-65 | | | | | | -2.04 | | 0.381 |
| | | | | | | -5.857 | | -6.174 |
| Age: 65+ | | | | | | 4.716 | | 4.876 |
| | | | | | | -6.48 | | -6.972 |
| Path Stage: 3b | | | | | | | 0.128 | -1.836 |
| | | | | | | | -6.54 | -7.103 |
| Path Stage: 3a | | | | | | | -0.294 | -5.923 |
| | | | | | | | -9.562 | -10.322 |
| Path Stage: 4 | | | | | | | -16.846 | -27.051 |
| | | | | | | | -16.972 | -17.387 |
| Constant | 160.754$^{***}$ | 175.778$^{***}$ | 177.977$^{***}$ | 163.831$^{***}$ | 158.266$^{***}$ | 161.003$^{***}$ | 161.701$^{***}$ | 186.691$^{***}$ |
| | -2.441 | -5.536 | -8.094 | -11.459 | -2.491 | -4.786 | -2.676 | -16.643 |
| Observations | 460 | 453 | 429 | 463 | 463 | 463 | 463 | 421 |
| $R^2$ | 0.002 | 0.021 | 0.013 | 0.015 | 0.023 | 0.003 | 0.002 | 0.061 |
| Adjusted $R^2$ | -0.002 | 0.017 | 0.006 | 0.009 | 0.021 | -0.001 | -0.004 | 0.024 |
| Residual Std. Error | 50.322 (df = | 49.824 (df = | 49.894 (df = | 49.948 (df = | 49.639 (df = 461) | 50.196 (df = | 50.279 (df = | 49.483 (df = |
| F Statistic | 0.514 (df = 2; | 4.878 (df = 2; | 1.877 (df = 3; | 2.370 (df = 3; | 10.919 (df = 1; | 0.752 (df = 2; | 0.331 (df = 3; | 1.648 (df = 16; |
| *Note:* | | | | | | | | $^{*}$p<0.1; $^{**}$p<0.05; $^{***}$p<0.01 |

### Table S4

**BCR Cox Regression Analysis**

| | *Dependent variable: time to BCR* | | | | | | | |
|---|---|---|---|---|---|---|---|---|
| | Race: White | BMI: Normal | PSA: < 2.5 | Gleason: < 6 | Adjuvant ADT: | Age: < 55 | Path Stage: 2 | Multivariate |
| Race: Black | 0.24 | | | | | | | 0.152 |
| | -0.327 | | | | | | | -0.354 |
| Race: Other | 0.537 | | | | | | | -0.48 |
| | -0.713 | | | | | | | -0.751 |
| BMI: Overweight (> 25; < 30) | | 0.219 | | | | | | 0.336 |
| | | -0.344 | | | | | | -0.378 |
| BMI: Obese (> 30) | | 0.122 | | | | | | 0.274 |
| | | -0.255 | | | | | | -0.296 |
| PSA: > 2.5; < 4 | | 0.411 | | | | | | 0.383 |
| | | -0.295 | | | | | | -0.333 |
| PSA: > 4; < 10 | | | 0.037 | | | | | 0.026 |
| | | | -0.204 | | | | | -0.213 |
| PSA: > 10 | | | 0.012 | | | | | 0.098 |
| | | | -0.223 | | | | | -0.237 |
| Gleason: 6 | | | | 0.731 | | | | 0.712 |
| | | | | -0.527 | | | | -0.532 |
| Gleason: 7 | | | | 1.255$^{**}$ | | | | 0.998$^{*}$ |
| | | | | -0.513 | | | | -0.52 |
| Gleason: 8+ | | | | 2.218$^{***}$ | | | | 1.482$^{***}$ |
| | | | | -0.523 | | | | -0.546 |
| Adjuvant ADT: TRUE | | | | | 0.675$^{***}$ | | | 0.446$^{**}$ |
| | | | | | -0.169 | | | -0.2 |
| Age: 55-65 | | | | | | -0.124 | | -0.269 |
| | | | | | | -0.187 | | -0.194 |
| Age: 65+ | | | | | | -0.158 | | -0.431$^{**}$ |
| | | | | | | -0.203 | | -0.215 |
| Path Stage: 3b | | | | | | | 0.845$^{***}$ | 0.647$^{***}$ |
| | | | | | | | -0.185 | -0.203 |
| Path Stage: 3a | | | | | | | 1.839$^{***}$ | 1.389$^{***}$ |
| | | | | | | | -0.204 | -0.257 |
| Path Stage: 4 | | | | | | | 1.069$^{**}$ | 1.075$^{**}$ |
| | | | | | | | -0.459 | -0.488 |
| Observations | 521 | 513 | 515 | 521 | 521 | 521 | 521 | 508 |
| $R^2$ | 0.002 | 0.005 | 0.0001 | 0.099 | 0.027 | 0.001 | 0.127 | 0.171 |
| Max. Possible $R^2$ | 0.98 | 0.98 | 0.979 | 0.98 | 0.98 | 0.98 | 0.98 | 0.98 |
| Log Likelihood | -1,020.72 | -1,006.22 | -996.017 | -994.057 | -1,014.11 | -1,020.88 | -985.903 | -940.834 |
| Wald Test | 1.070 (df = 2) | 2.670 (df = 3) | 0.040 (df = 2) | 57.670$^{***}$ (df = 3) | 15.880$^{***}$ (df = 1) | 0.670 (df = 2) | 87.410$^{***}$ (df = 3) | 110.990$^{***}$ (df = 16) |
| LR Test | 0.961 (df = 2) | 2.554 (df = 3) | 0.040 (df = 2) | 54.294$^{***}$ (df = 3) | 14.190$^{***}$ (df = 1) | 0.659 (df = 2) | 70.602$^{***}$ (df = 3) | 95.471$^{***}$ (df = 16) |
| Score (Logrank) Test | 1.090 (df = 2) | 2.686 (df = 3) | 0.040 (df = 2) | 66.672$^{***}$ (df = 3) | 16.490$^{***}$ (df = 1) | 0.673 (df = 2) | 107.851$^{***}$ (df = 3) | 134.828$^{***}$ (df = 16) |

*Note:* *p<0.1;**p<0.05;***p<0.01

LNCaP_NCOR2

### Table S5

| | ID | Descriptior | setSize | enrichmen | NES | pvalue | p.adjust | qvalues | rank | leading_ed | core_enricl | Group |
|---|---|---|---|---|---|---|---|---|---|---|---|---|
| NUYTTEN_I | NUYTTEN_I | NUYTTEN_I | 729 | 0.451126 | 2.279387 | 0.000123 | 0.004767 | 0.003497 | 2658 | tags=38%, l | 619279/35 | LNCaP-NCOR2 |
| GOBERT_O | GOBERT_O | GOBERT_O | 714 | 0.338306 | 1.706807 | 0.000123 | 0.004767 | 0.003497 | 2799 | tags=32%, l | 79993/110 | LNCaP-NCOR2 |
| GEORGES_ | GEORGES_ | GEORGES_ | 682 | 0.334008 | 1.680825 | 0.000124 | 0.004767 | 0.003497 | 2928 | tags=32%, l | 5865/6598 | LNCaP-NCOR2 |
| DACOSTA_ | DACOSTA_ | DACOSTA_ | 668 | 0.456398 | 2.292646 | 0.000124 | 0.004767 | 0.003497 | 2953 | tags=45%, l | 11059/520 | LNCaP-NCOR2 |
| KRIGE_RES | KRIGE_RES | KRIGE_RES | 639 | 0.304294 | 1.524479 | 0.000125 | 0.004767 | 0.003497 | 2950 | tags=31%, l | 3685/8795, | LNCaP-NCOR2 |
| BUYTAERT_ | BUYTAERT_ | BUYTAERT_ | 633 | 0.387127 | 1.938429 | 0.000125 | 0.004767 | 0.003497 | 3520 | tags=46%, l | 4254/7107, | LNCaP-NCOR2 |
| ONKEN_UV | ONKEN_UV | ONKEN_UV | 571 | 0.329696 | 1.637867 | 0.000127 | 0.004767 | 0.003497 | 2704 | tags=33%, l | 65986/110 | LNCaP-NCOR2 |
| RODRIGUES | RODRIGUES | RODRIGUES | 550 | 0.339662 | 1.682469 | 0.000127 | 0.004767 | 0.003497 | 2713 | tags=33%, l | 5793/5908 | LNCaP-NCOR2 |
| RODRIGUES | RODRIGUES | RODRIGUES | 549 | 0.337517 | 1.671354 | 0.000127 | 0.004767 | 0.003497 | 2521 | tags=33%, l | 8795/6741, | LNCaP-NCOR2 |
| CUI_TCF21 | CUI_TCF21 | CUI_TCF21 | 555 | 0.480779 | 2.381946 | 0.000127 | 0.004767 | 0.003497 | 2907 | tags=45%, l | 5793/3572, | LNCaP-NCOR2 |
| SCHLOSSER | SCHLOSSER | SCHLOSSER | 524 | 0.368523 | 1.819657 | 0.000128 | 0.004767 | 0.003497 | 2747 | tags=33%, l | 5793/3572, | LNCaP-NCOR2 |
| KRIGE_RES | KRIGE_RES | KRIGE_RES | 513 | 0.328559 | 1.618693 | 0.000129 | 0.004767 | 0.003497 | 2887 | tags=32%, l | 3685/8795, | LNCaP-NCOR2 |
| CREIGHTON | CREIGHTON | CREIGHTON | 500 | 0.387877 | 1.907246 | 0.000129 | 0.004767 | 0.003497 | 2912 | tags=38%, l | 5205/5152, | LNCaP-NCOR2 |
| NUYTTEN_I | NUYTTEN_I | NUYTTEN_I | 503 | 0.395506 | 1.945184 | 0.000129 | 0.004767 | 0.003497 | 2537 | tags=34%, l | 4254/5716 | LNCaP-NCOR2 |
| HAMAI_AP | HAMAI_AP | HAMAI_AP | 494 | 0.457257 | 2.24577 | 0.00013 | 0.004767 | 0.003497 | 2989 | tags=44%, l | 11059/368 | LNCaP-NCOR2 |
| GOZGIT_ES | GOZGIT_ES | GOZGIT_ES | 471 | 0.390282 | 1.910104 | 0.00013 | 0.004767 | 0.003497 | 2754 | tags=39%, l | 5793/4254, | LNCaP-NCOR2 |
| WONG_AD | WONG_AD | WONG_AD | 435 | 0.363276 | 1.767697 | 0.000132 | 0.004767 | 0.003497 | 2621 | tags=33%, l | 4254/3915, | LNCaP-NCOR2 |
| FLECHNER_ | FLECHNER_ | FLECHNER_ | 417 | 0.367483 | 1.783077 | 0.000132 | 0.004767 | 0.003497 | 2339 | tags=32%, l | 3572/1105 | LNCaP-NCOR2 |
| NAKAMURA | NAKAMURA | NAKAMURA | 419 | 0.34415 | 1.670012 | 0.000132 | 0.004767 | 0.003497 | 2908 | tags=36%, l | 5205/6542, | LNCaP-NCOR2 |
| ENK_UV_R | ENK_UV_R | ENK_UV_R | 400 | 0.385164 | 1.861353 | 0.000133 | 0.004767 | 0.003497 | 3043 | tags=38%, l | 10565/559 | LNCaP-NCOR2 |
| MASSARWI | MASSARWI | MASSARWI | 381 | 0.40493 | 1.948437 | 0.000134 | 0.004767 | 0.003497 | 2733 | tags=39%, l | 619279/57 | LNCaP-NCOR2 |
| MILI_PSEU | MILI_PSEU | MILI_PSEU | 377 | 0.434034 | 2.08639 | 0.000134 | 0.004767 | 0.003497 | 2996 | tags=42%, l | 11059/233 | LNCaP-NCOR2 |
| DUTERTRE_ | DUTERTRE_ | DUTERTRE_ | 376 | 0.472887 | 2.271908 | 0.000134 | 0.004767 | 0.003497 | 2590 | tags=41%, l | 619279/36 | LNCaP-NCOR2 |
| MARTORIA | MARTORIA | MARTORIA | 361 | 0.353147 | 1.689129 | 0.000135 | 0.004767 | 0.003497 | 2169 | tags=28%, l | 5793/3572, | LNCaP-NCOR2 |
| DACOSTA_ | DACOSTA_ | DACOSTA_ | 362 | 0.460961 | 2.205187 | 0.000135 | 0.004767 | 0.003497 | 2953 | tags=44%, l | 10565/969 | LNCaP-NCOR2 |
| SHEN_SMA | SHEN_SMA | SHEN_SMA | 360 | 0.374262 | 1.78908 | 0.000136 | 0.004767 | 0.003497 | 2924 | tags=36%, l | 167227/67 | LNCaP-NCOR2 |
| SENESE_HD | SENESE_HD | SENESE_HD | 354 | 0.461194 | 2.20238 | 0.000136 | 0.004767 | 0.003497 | 1793 | tags=35%, l | 3572/4254, | LNCaP-NCOR2 |
| CREIGHTON | CREIGHTON | CREIGHTON | 342 | 0.411512 | 1.958126 | 0.000137 | 0.004767 | 0.003497 | 2720 | tags=39%, l | 619279/52 | LNCaP-NCOR2 |
| ZWANG_CL | ZWANG_CL | ZWANG_CL | 331 | 0.406132 | 1.925204 | 0.000138 | 0.004767 | 0.003497 | 2815 | tags=37%, l | 3572/7999 | LNCaP-NCOR2 |
| GRYDER_PA | GRYDER_PA | GRYDER_PA | 325 | 0.336915 | 1.594457 | 0.000138 | 0.004767 | 0.003497 | 2714 | tags=33%, l | 6542/2309 | LNCaP-NCOR2 |
| BAELDE_DI | BAELDE_DI | BAELDE_DI | 318 | 0.386675 | 1.825662 | 0.000138 | 0.004767 | 0.003497 | 2337 | tags=32%, l | 11059/368 | LNCaP-NCOR2 |
| ACEVEDO_ | ACEVEDO_ | ACEVEDO_ | 312 | 0.33857 | 1.595621 | 0.000138 | 0.004767 | 0.003497 | 2282 | tags=26%, l | 4193/9693, | LNCaP-NCOR2 |
| OSMAN_BL | OSMAN_BL | OSMAN_BL | 300 | 0.423336 | 1.986272 | 0.000139 | 0.004767 | 0.003497 | 2883 | tags=41%, l | 64393/289 | LNCaP-NCOR2 |
| SENESE_HD | SENESE_HD | SENESE_HD | 304 | 0.487725 | 2.29019 | 0.000139 | 0.004767 | 0.003497 | 1793 | tags=37%, l | 3572/4254, | LNCaP-NCOR2 |
| ZHENG_BO | ZHENG_BO | ZHENG_BO | 298 | 0.430935 | 2.019679 | 0.000139 | 0.004767 | 0.003497 | 1705 | tags=30%, l | 4193/3572, | LNCaP-NCOR2 |
| ONDER_CD | ONDER_CD | ONDER_CD | 275 | 0.360014 | 1.676558 | 0.00014 | 0.004767 | 0.003497 | 2529 | tags=32%, l | 1824/6653, | LNCaP-NCOR2 |
| PILON_KLF | PILON_KLF | PILON_KLF | 277 | 0.374922 | 1.747451 | 0.00014 | 0.004767 | 0.003497 | 3135 | tags=39%, l | 4193/4199, | LNCaP-NCOR2 |
| LU_EZH2_T | LU_EZH2_T | LU_EZH2_T | 274 | 0.43994 | 2.047666 | 0.00014 | 0.004767 | 0.003497 | 3106 | tags=45%, l | 3915/1672 | LNCaP-NCOR2 |
| FOSTER_TC | FOSTER_TC | FOSTER_TC | 270 | 0.43845 | 2.036546 | 0.00014 | 0.004767 | 0.003497 | 2825 | tags=40%, l | 3685/2309, | LNCaP-NCOR2 |
| WANG_LM | WANG_LM | WANG_LM | 267 | 0.380236 | 1.76602 | 0.00014 | 0.004767 | 0.003497 | 2571 | tags=33%, l | 57169/654 | LNCaP-NCOR2 |
| PHONG_TN | PHONG_TN | PHONG_TN | 255 | 0.369457 | 1.705667 | 0.000141 | 0.004767 | 0.003497 | 2047 | tags=29%, l | 65986/357 | LNCaP-NCOR2 |
| FULCHER_I | FULCHER_I | FULCHER_I | 254 | 0.418181 | 1.929806 | 0.000141 | 0.004767 | 0.003497 | 3033 | tags=43%, l | 11059/182 | LNCaP-NCOR2 |
| JOHNSTON | JOHNSTON | JOHNSTON | 250 | 0.419769 | 1.933168 | 0.000142 | 0.004767 | 0.003497 | 2591 | tags=39%, l | 5793/2581 | LNCaP-NCOR2 |
| SENGUPTA | SENGUPTA | SENGUPTA | 244 | 0.50609 | 2.325818 | 0.000142 | 0.004767 | 0.003497 | 2577 | tags=47%, l | 4193/6598 | LNCaP-NCOR2 |
| DOUGLAS_ | DOUGLAS_ | DOUGLAS_ | 234 | 0.425163 | 1.942094 | 0.000143 | 0.004767 | 0.003497 | 2117 | tags=33%, l | 9693/7107, | LNCaP-NCOR2 |
| DAZARD_R | DAZARD_R | DAZARD_R | 235 | 0.443961 | 2.028864 | 0.000143 | 0.004767 | 0.003497 | 2810 | tags=39%, l | 9693/2336 | LNCaP-NCOR2 |
| PASINI_SUZ | PASINI_SUZ | PASINI_SUZ | 220 | 0.398533 | 1.812034 | 0.000143 | 0.004767 | 0.003497 | 2642 | tags=35%, l | 3685/1287, | LNCaP-NCOR2 |
| FISCHER_D | FISCHER_D | FISCHER_D | 229 | 0.463215 | 2.111771 | 0.000143 | 0.004767 | 0.003497 | 1669 | tags=32%, l | 4193/8795, | LNCaP-NCOR2 |
| DEBIASI_AF | DEBIASI_AF | DEBIASI_AF | 228 | 0.38549 | 1.756684 | 0.000144 | 0.004767 | 0.003497 | 2872 | tags=41%, l | 3005/5472 | LNCaP-NCOR2 |
| ZHOU_INFL | ZHOU_INFL | ZHOU_INFL | 228 | 0.433428 | 1.97514 | 0.000144 | 0.004767 | 0.003497 | 1533 | tags=29%, l | 79993/854 | LNCaP-NCOR2 |
| GABRIELY_ | GABRIELY_ | GABRIELY_ | 231 | 0.576363 | 2.629642 | 0.000144 | 0.004767 | 0.003497 | 2976 | tags=58%, l | 10565/232 | LNCaP-NCOR2 |
| TORCHIA_T | TORCHIA_T | TORCHIA_T | 211 | 0.370417 | 1.673465 | 0.000145 | 0.004767 | 0.003497 | 2130 | tags=27%, l | 619279/57 | LNCaP-NCOR2 |
| DEURIG_T_ | DEURIG_T_ | DEURIG_T_ | 198 | 0.44456 | 1.992802 | 0.000146 | 0.004767 | 0.003497 | 2823 | tags=40%, l | 51635/710 | LNCaP-NCOR2 |
| HORIUCHI_ | HORIUCHI_ | HORIUCHI_ | 192 | 0.465302 | 2.080798 | 0.000146 | 0.004767 | 0.003497 | 2822 | tags=48%, l | 23092/576 | LNCaP-NCOR2 |
| DURCHDEV | DURCHDEV | DURCHDEV | 187 | 0.447387 | 1.993862 | 0.000146 | 0.004767 | 0.003497 | 1812 | tags=35%, l | 7107/5127 | LNCaP-NCOR2 |
| ZHOU_INFL | ZHOU_INFL | ZHOU_INFL | 183 | 0.457885 | 2.035012 | 0.000146 | 0.004767 | 0.003497 | 2833 | tags=38%, l | 5865/6598 | LNCaP-NCOR2 |
| CONCANNO | CONCANNO | CONCANNO | 180 | 0.411217 | 1.82446 | 0.000147 | 0.004767 | 0.003497 | 2769 | tags=37%, l | 4193/4254, | LNCaP-NCOR2 |
| PHONG_TN | PHONG_TN | PHONG_TN | 181 | 0.410982 | 1.823683 | 0.000147 | 0.004767 | 0.003497 | 2540 | tags=39%, l | 23362/230 | LNCaP-NCOR2 |
| CHEN_HOX | CHEN_HOX | CHEN_HOX | 174 | 0.40787 | 1.801184 | 0.000147 | 0.004767 | 0.003497 | 2958 | tags=36%, l | 8795/2309, | LNCaP-NCOR2 |
| LU_AGING_ | LU_AGING_ | LU_AGING_ | 177 | 0.398913 | 1.764477 | 0.000147 | 0.004767 | 0.003497 | 2301 | tags=36%, l | 3685/1287, | LNCaP-NCOR2 |
| RIGGINS_T | RIGGINS_T | RIGGINS_T | 168 | 0.514517 | 2.260077 | 0.000148 | 0.004767 | 0.003497 | 2801 | tags=51%, l | 3572/1056 | LNCaP-NCOR2 |
| WANG_SM | WANG_SM | WANG_SM | 167 | 0.415457 | 1.823313 | 0.000148 | 0.004767 | 0.003497 | 2379 | tags=39%, l | 5205/4199, | LNCaP-NCOR2 |
| LINSLEY_M | LINSLEY_M | LINSLEY_M | 159 | 0.427009 | 1.864133 | 0.000149 | 0.004767 | 0.003497 | 1994 | tags=28%, l | 5865/1105 | LNCaP-NCOR2 |
| ZHANG_RE | ZHANG_RE | ZHANG_RE | 148 | 0.544411 | 2.35156 | 0.00015 | 0.004767 | 0.003497 | 1988 | tags=43%, l | 79993/571 | LNCaP-NCOR2 |
| ZWANG_CL | ZWANG_CL | ZWANG_CL | 147 | 0.41883 | 1.806817 | 0.000151 | 0.004767 | 0.003497 | 2496 | tags=37%, l | 85463/969 | LNCaP-NCOR2 |

| | | | | | | | | | | | | |
|---|---|---|---|---|---|---|---|---|---|---|---|---|
| HALLMARK | HALLMARK | HALLMARK | 141 | 0.512585 | 2.200615 | 0.000151 | 0.004767 | 0.003497 | 3191 | tags=52%, l | 65986/357 | LNCaP-NCOR2 |
| ZHENG_FO | ZHENG_FO | ZHENG_FO | 141 | 0.493266 | 2.117677 | 0.000151 | 0.004767 | 0.003497 | 2440 | tags=45%, l | 51274/169 | LNCaP-NCOR2 |
| HALLMARK | HALLMARK | HALLMARK | 140 | 0.424009 | 1.817798 | 0.000151 | 0.004767 | 0.003497 | 1432 | tags=26%, l | 2309/8974 | LNCaP-NCOR2 |
| SENESE_HD | SENESE_HD | SENESE_HD | 138 | 0.452722 | 1.936607 | 0.000152 | 0.004767 | 0.003497 | 1794 | tags=32%, l | 5865/3572 | LNCaP-NCOR2 |
| NAGASHIM | NAGASHIM | NAGASHIM | 131 | 0.435113 | 1.847872 | 0.000153 | 0.004767 | 0.003497 | 3501 | tags=44%, l | 8795/2302 | LNCaP-NCOR2 |
| GRAESSMA | GRAESSMA | GRAESSMA | 120 | 0.491554 | 2.058328 | 0.000154 | 0.004767 | 0.003497 | 2672 | tags=42%, l | 28996/850 | LNCaP-NCOR2 |
| PHONG_TN | PHONG_TN | PHONG_TN | 115 | 0.473665 | 1.970932 | 0.000154 | 0.004767 | 0.003497 | 2301 | tags=39%, l | 3572/3685 | LNCaP-NCOR2 |
| LIU_SOX4_ | LIU_SOX4_ | LIU_SOX4_ | 107 | 0.509468 | 2.097857 | 0.000155 | 0.004767 | 0.003497 | 1808 | tags=36%, l | 59084/665 | LNCaP-NCOR2 |
| PICCALUGA | PICCALUGA | PICCALUGA | 100 | 0.530005 | 2.161907 | 0.000155 | 0.004767 | 0.003497 | 2940 | tags=49%, l | 65986/931 | LNCaP-NCOR2 |
| TAKEDA_TA | TAKEDA_TA | TAKEDA_TA | 104 | 0.552174 | 2.263579 | 0.000155 | 0.004767 | 0.003497 | 2587 | tags=52%, l | 57674/550 | LNCaP-NCOR2 |
| BROCKE_AI | BROCKE_AI | BROCKE_AI | 99 | 0.477867 | 1.945453 | 0.000155 | 0.004767 | 0.003497 | 2940 | tags=40%, l | 5793/3572 | LNCaP-NCOR2 |
| RODRIGUES | RODRIGUES | RODRIGUES | 95 | 0.511947 | 2.068242 | 0.000156 | 0.004767 | 0.003497 | 2587 | tags=42%, l | 5793/1078 | LNCaP-NCOR2 |
| HALLMARK | HALLMARK | HALLMARK | 93 | 0.519663 | 2.093026 | 0.000156 | 0.004767 | 0.003497 | 1061 | tags=32%, l | 65986/368 | LNCaP-NCOR2 |
| IKEDA_MIR | IKEDA_MIR | IKEDA_MIR | 91 | 0.501939 | 2.016204 | 0.000156 | 0.004767 | 0.003497 | 3191 | tags=57%, l | 6651/2803 | LNCaP-NCOR2 |
| RASHI_RES | RASHI_RES | RASHI_RES | 85 | 0.503087 | 1.997569 | 0.000157 | 0.004767 | 0.003497 | 2584 | tags=46%, l | 4193/1824 | LNCaP-NCOR2 |
| TAKEDA_TA | TAKEDA_TA | TAKEDA_TA | 82 | 0.490522 | 1.935812 | 0.000159 | 0.004767 | 0.003497 | 1620 | tags=33%, l | 79993/969 | LNCaP-NCOR2 |
| MACAEVA_ | MACAEVA_ | MACAEVA_ | 73 | 0.608796 | 2.363517 | 0.000159 | 0.004767 | 0.003497 | 2530 | tags=59%, l | 4193/8795 | LNCaP-NCOR2 |
| CHIANG_LI | CHIANG_LI | CHIANG_LI | 78 | 0.507322 | 1.990446 | 0.000159 | 0.004767 | 0.003497 | 2178 | tags=40%, l | 6542/2581 | LNCaP-NCOR2 |
| IVANOVSKA | IVANOVSKA | IVANOVSKA | 74 | 0.502997 | 1.957073 | 0.000159 | 0.004767 | 0.003497 | 1971 | tags=41%, l | 57169/590 | LNCaP-NCOR2 |
| SENESE_HD | SENESE_HD | SENESE_HD | 77 | 0.497645 | 1.948146 | 0.00016 | 0.004767 | 0.003497 | 3557 | tags=60%, l | 3572/8795 | LNCaP-NCOR2 |
| NELSON_RI | NELSON_RI | NELSON_RI | 75 | 0.522539 | 2.036673 | 0.00016 | 0.004767 | 0.003497 | 1488 | tags=36%, l | 65986/368 | LNCaP-NCOR2 |
| DUTERTRE_ | DUTERTRE_ | DUTERTRE_ | 68 | 0.499869 | 1.914748 | 0.00016 | 0.004767 | 0.003497 | 1905 | tags=41%, l | 619279/44 | LNCaP-NCOR2 |
| SEIDEN_ON | SEIDEN_ON | SEIDEN_ON | 68 | 0.503883 | 1.930124 | 0.00016 | 0.004767 | 0.003497 | 2800 | tags=49%, l | 25800/332 | LNCaP-NCOR2 |
| AZARE_NE | AZARE_NE | AZARE_NE | 67 | 0.504938 | 1.928322 | 0.000161 | 0.004767 | 0.003497 | 1342 | tags=34%, l | 6653/4747 | LNCaP-NCOR2 |
| LEE_NEURA | LEE_NEURA | LEE_NEURA | 67 | 0.571166 | 2.181241 | 0.000161 | 0.004767 | 0.003497 | 1394 | tags=34%, l | 1287/8795 | LNCaP-NCOR2 |
| GUO_TARG | GUO_TARG | GUO_TARG | 62 | 0.536655 | 2.020848 | 0.000162 | 0.004767 | 0.003497 | 2442 | tags=48%, l | 6542/3320 | LNCaP-NCOR2 |
| DACOSTA_ | DACOSTA_ | DACOSTA_ | 59 | 0.529018 | 1.972384 | 0.000163 | 0.004767 | 0.003497 | 2453 | tags=44%, l | 11059/520 | LNCaP-NCOR2 |
| KEGG_P53_ | KEGG_P53_ | KEGG_P53_ | 58 | 0.537039 | 1.996523 | 0.000163 | 0.004767 | 0.003497 | 1572 | tags=38%, l | 4193/8795 | LNCaP-NCOR2 |
| DACOSTA_ | DACOSTA_ | DACOSTA_ | 55 | 0.534322 | 1.96564 | 0.000164 | 0.004767 | 0.003497 | 3402 | tags=60%, l | 9732/6651 | LNCaP-NCOR2 |
| BILD_CTNN | BILD_CTNN | BILD_CTNN | 50 | 0.707809 | 2.554837 | 0.000165 | 0.004767 | 0.003497 | 1187 | tags=44%, l | 64393/665 | LNCaP-NCOR2 |
| GHANDHI_ | GHANDHI_ | GHANDHI_ | 51 | 0.59944 | 2.171444 | 0.000165 | 0.004767 | 0.003497 | 2145 | tags=51%, l | 4193/8546 | LNCaP-NCOR2 |
| CHIANG_LI | CHIANG_LI | CHIANG_LI | 48 | 0.587939 | 2.104415 | 0.000166 | 0.004767 | 0.003497 | 2179 | tags=56%, l | 9693/6653 | LNCaP-NCOR2 |
| SEKI_INFLA | SEKI_INFLA | SEKI_INFLA | 46 | 0.565118 | 2.008201 | 0.000166 | 0.004767 | 0.003497 | 1716 | tags=39%, l | 9180/9497 | LNCaP-NCOR2 |
| BOSCO_INT | BOSCO_INT | BOSCO_INT | 45 | 0.537701 | 1.901872 | 0.000167 | 0.004767 | 0.003497 | 2310 | tags=42%, l | 2803/5119 | LNCaP-NCOR2 |
| HALLMARK | HALLMARK | HALLMARK | 45 | 0.538079 | 1.90321 | 0.000167 | 0.004767 | 0.003497 | 1760 | tags=38%, l | 3572/9180 | LNCaP-NCOR2 |
| KIM_GERM | KIM_GERM | KIM_GERM | 45 | 0.553017 | 1.956045 | 0.000167 | 0.004767 | 0.003497 | 1371 | tags=33%, l | 5784/6439 | LNCaP-NCOR2 |
| ONDER_CD | ONDER_CD | ONDER_CD | 47 | 0.565684 | 2.013707 | 0.000167 | 0.004767 | 0.003497 | 1569 | tags=40%, l | 6653/4430 | LNCaP-NCOR2 |
| SMIRNOV_ | SMIRNOV_ | SMIRNOV_ | 40 | 0.621839 | 2.146459 | 0.000167 | 0.004767 | 0.003497 | 2548 | tags=52%, l | 4193/6598 | LNCaP-NCOR2 |
| DAUER_STA | DAUER_STA | DAUER_STA | 39 | 0.627794 | 2.15196 | 0.000168 | 0.004767 | 0.003497 | 1689 | tags=44%, l | 55008/511 | LNCaP-NCOR2 |
| PID_INTEG | PID_INTEG | PID_INTEG | 37 | 0.620996 | 2.098567 | 0.000169 | 0.004767 | 0.003497 | 1985 | tags=46%, l | 3685/3915 | LNCaP-NCOR2 |
| BERENJENO | BERENJENO | BERENJENO | 35 | 0.629127 | 2.104468 | 0.000169 | 0.004767 | 0.003497 | 1425 | tags=49%, l | 6741/3480 | LNCaP-NCOR2 |
| WANG_RES | WANG_RES | WANG_RES | 26 | 0.681111 | 2.118171 | 0.000172 | 0.004767 | 0.003497 | 2347 | tags=54%, l | 65986/514 | LNCaP-NCOR2 |
| MOSERLE_ | MOSERLE_ | MOSERLE_ | 22 | 0.746647 | 2.23115 | 0.000173 | 0.004767 | 0.003497 | 1689 | tags=68%, l | 51191/641 | LNCaP-NCOR2 |
| REACTOME | REACTOME | REACTOME | 21 | 0.734731 | 2.169145 | 0.000174 | 0.004767 | 0.003497 | 2132 | tags=62%, l | 657/93/73 | LNCaP-NCOR2 |
| GARGALOV | GARGALOV | GARGALOV | 17 | 0.726254 | 2.031601 | 0.000175 | 0.004767 | 0.003497 | 1398 | tags=47%, l | 619279/47 | LNCaP-NCOR2 |
| LANDIS_ER | LANDIS_ER | LANDIS_ER | 17 | 0.739823 | 2.069559 | 0.000175 | 0.004767 | 0.003497 | 1620 | tags=59%, l | 3709/1829 | LNCaP-NCOR2 |
| KEGG_O_G | KEGG_O_G | KEGG_O_G | 16 | 0.747608 | 2.060853 | 0.000175 | 0.004767 | 0.003497 | 1459 | tags=50%, l | 2650/5556 | LNCaP-NCOR2 |
| RASHI_NFK | RASHI_NFK | RASHI_NFK | 16 | 0.737592 | 2.033244 | 0.000175 | 0.004767 | 0.003497 | 1960 | tags=56%, l | 4193/4791 | LNCaP-NCOR2 |
| WU_HBX_T | WU_HBX_T | WU_HBX_T | 16 | 0.723465 | 1.9943 | 0.000175 | 0.004767 | 0.003497 | 990 | tags=44%, l | 5784/1316 | LNCaP-NCOR2 |
| AMUNDSO | AMUNDSO | AMUNDSO | 15 | 0.753784 | 2.04286 | 0.000175 | 0.004767 | 0.003497 | 1492 | tags=60%, l | 4193/7508 | LNCaP-NCOR2 |
| INGA_TP53 | INGA_TP53 | INGA_TP53 | 14 | 0.753243 | 2.003063 | 0.000176 | 0.004767 | 0.003497 | 1572 | tags=57%, l | 4193/8795 | LNCaP-NCOR2 |
| HERNANDE | HERNANDE | HERNANDE | 13 | 0.776508 | 2.022798 | 0.000176 | 0.004767 | 0.003497 | 1592 | tags=62%, l | 4982/4094 | LNCaP-NCOR2 |
| HOLLEMAN | HOLLEMAN | HOLLEMAN | 15 | -0.773727 | -2.18701 | 0.000233 | 0.004767 | 0.003497 | 1417 | tags=67%, l | 6130/6188 | LNCaP-NCOR2 |
| REACTOME | REACTOME | REACTOME | 18 | -0.758146 | -2.25388 | 0.000234 | 0.004767 | 0.003497 | 1406 | tags=50%, l | 8367/8354 | LNCaP-NCOR2 |
| EGUCHI_CE | EGUCHI_CE | EGUCHI_CE | 23 | -0.669944 | -2.128867 | 0.000236 | 0.004767 | 0.003497 | 3172 | tags=78%, l | 1062/1040 | LNCaP-NCOR2 |
| REACTOME | REACTOME | REACTOME | 24 | -0.665072 | -2.140144 | 0.000236 | 0.004767 | 0.003497 | 2481 | tags=58%, l | 4173/5422 | LNCaP-NCOR2 |
| REACTOME | REACTOME | REACTOME | 25 | -0.72998 | -2.376733 | 0.000237 | 0.004767 | 0.003497 | 808 | tags=56%, l | 10213/833 | LNCaP-NCOR2 |
| REACTOME | REACTOME | REACTOME | 26 | -0.67731 | -2.228979 | 0.000238 | 0.004767 | 0.003497 | 948 | tags=35%, l | 1457/8786 | LNCaP-NCOR2 |
| NIKOLSKY_ | NIKOLSKY_ | NIKOLSKY_ | 29 | -0.671968 | -2.271534 | 0.00024 | 0.004767 | 0.003497 | 2741 | tags=66%, l | 4580/2362 | LNCaP-NCOR2 |
| REACTOME | REACTOME | REACTOME | 29 | -0.686663 | -2.321211 | 0.00024 | 0.004767 | 0.003497 | 2481 | tags=66%, l | 4173/5982 | LNCaP-NCOR2 |
| KEGG_DNA | KEGG_DNA | KEGG_DNA | 33 | -0.670119 | -2.33392 | 0.000243 | 0.004767 | 0.003497 | 2481 | tags=67%, l | 4173/5982 | LNCaP-NCOR2 |
| BILANGES_ | BILANGES_ | BILANGES_ | 36 | -0.692815 | -2.466549 | 0.000245 | 0.004767 | 0.003497 | 1216 | tags=61%, l | 6138/6188 | LNCaP-NCOR2 |
| BURTON_A | BURTON_A | BURTON_A | 34 | -0.599372 | -2.102167 | 0.000245 | 0.004767 | 0.003497 | 1661 | tags=35%, l | 3835/5120 | LNCaP-NCOR2 |
| REACTOME | REACTOME | REACTOME | 38 | -0.802346 | -2.894056 | 0.000246 | 0.004767 | 0.003497 | 1140 | tags=61%, l | 8367/4595 | LNCaP-NCOR2 |
| POMEROY_ | POMEROY_ | POMEROY_ | 39 | -0.591155 | -2.141275 | 0.000247 | 0.004767 | 0.003497 | 1657 | tags=38%, l | 6633/6130 | LNCaP-NCOR2 |
| REACTOME | REACTOME | REACTOME | 40 | -0.816813 | -2.973875 | 0.000249 | 0.004767 | 0.003497 | 1853 | tags=75%, l | 8342/3021 | LNCaP-NCOR2 |

| | | | | | | | | | | | | |
|---|---|---|---|---|---|---|---|---|---|---|---|---|
| REACTOME | REACTOME | REACTOME | 47 | -0.77202 | -2.918138 | 0.00025 | 0.004767 | 0.003497 | 1417 | tags=57%, l | 8363/1059 | LNCaP-NCOR2 |
| REACTOME | REACTOME | REACTOME | 47 | -0.739069 | -2.793586 | 0.00025 | 0.004767 | 0.003497 | 1406 | tags=53%, l | 79172/301 | LNCaP-NCOR2 |
| REACTOME | REACTOME | REACTOME | 45 | -0.755422 | -2.821761 | 0.00025 | 0.004767 | 0.003497 | 1140 | tags=53%, l | 8367/4595 | LNCaP-NCOR2 |
| ISHIDA_E2F | ISHIDA_E2F | ISHIDA_E2F | 46 | -0.691988 | -2.596462 | 0.000251 | 0.004767 | 0.003497 | 1661 | tags=48%, l | 4176/2339 | LNCaP-NCOR2 |
| CHNG_MU | CHNG_MU | CHNG_MU | 42 | -0.704272 | -2.581253 | 0.000251 | 0.004767 | 0.003497 | 1794 | tags=64%, l | 1211/2587 | LNCaP-NCOR2 |
| REACTOME | REACTOME | REACTOME | 42 | -0.63731 | -2.335828 | 0.000251 | 0.004767 | 0.003497 | 2088 | tags=60%, l | 6189/6235 | LNCaP-NCOR2 |
| REACTOME | REACTOME | REACTOME | 42 | -0.640749 | -2.348432 | 0.000251 | 0.004767 | 0.003497 | 2141 | tags=55%, l | 51295/470 | LNCaP-NCOR2 |
| REACTOME | REACTOME | REACTOME | 42 | -0.812135 | -2.976584 | 0.000251 | 0.004767 | 0.003497 | 1417 | tags=67%, l | 8363/6908 | LNCaP-NCOR2 |
| KOINUMA_ | KOINUMA_ | KOINUMA_ | 606 | 0.299068 | 1.493035 | 0.000251 | 0.004767 | 0.003497 | 2636 | tags=30%, l | 3685/3915 | LNCaP-NCOR2 |
| REACTOME | REACTOME | REACTOME | 41 | -0.796497 | -2.903273 | 0.000252 | 0.004767 | 0.003497 | 1417 | tags=66%, l | 8363/3015 | LNCaP-NCOR2 |
| REACTOME | REACTOME | REACTOME | 48 | -0.752316 | -2.847698 | 0.000252 | 0.004767 | 0.003497 | 1417 | tags=67%, l | 8342/3021 | LNCaP-NCOR2 |
| ZHANG_RE | ZHANG_RE | ZHANG_RE | 52 | -0.591378 | -2.27794 | 0.000254 | 0.004767 | 0.003497 | 1987 | tags=50%, l | 4708/1025 | LNCaP-NCOR2 |
| REACTOME | REACTOME | REACTOME | 51 | -0.596565 | -2.287775 | 0.000255 | 0.004767 | 0.003497 | 1406 | tags=41%, l | 5599/8367 | LNCaP-NCOR2 |
| REACTOME | REACTOME | REACTOME | 51 | -0.738687 | -2.832801 | 0.000255 | 0.004767 | 0.003497 | 1488 | tags=59%, l | 3020/8363 | LNCaP-NCOR2 |
| REACTOME | REACTOME | REACTOME | 51 | -0.691469 | -2.651723 | 0.000255 | 0.004767 | 0.003497 | 1406 | tags=49%, l | 4001/3015 | LNCaP-NCOR2 |
| SONG_TAR | SONG_TAR | SONG_TAR | 50 | -0.583979 | -2.233742 | 0.000255 | 0.004767 | 0.003497 | 809 | tags=32%, l | 7374/6240 | LNCaP-NCOR2 |
| KAUFFMAN | KAUFFMAN | KAUFFMAN | 55 | -0.57473 | -2.241202 | 0.000256 | 0.004767 | 0.003497 | 2370 | tags=55%, l | 4173/9156 | LNCaP-NCOR2 |
| REACTOME | REACTOME | REACTOME | 55 | -0.656464 | -2.55993 | 0.000256 | 0.004767 | 0.003497 | 1416 | tags=51%, l | 8363/4001 | LNCaP-NCOR2 |
| REACTOME | REACTOME | REACTOME | 56 | -0.754208 | -2.955813 | 0.000257 | 0.004767 | 0.003497 | 1812 | tags=66%, l | 3021/1019 | LNCaP-NCOR2 |
| REACTOME | REACTOME | REACTOME | 56 | -0.508615 | -1.993309 | 0.000257 | 0.004767 | 0.003497 | 2901 | tags=54%, l | 5719/5695 | LNCaP-NCOR2 |
| REACTOME | REACTOME | REACTOME | 56 | -0.716976 | -2.809897 | 0.000257 | 0.004767 | 0.003497 | 1140 | tags=45%, l | 8367/8354 | LNCaP-NCOR2 |
| REN_BOUN | REN_BOUN | REN_BOUN | 56 | -0.571694 | -2.240521 | 0.000257 | 0.004767 | 0.003497 | 1522 | tags=41%, l | 10592/598 | LNCaP-NCOR2 |
| MORI_MAT | MORI_MAT | MORI_MAT | 57 | -0.548536 | -2.156702 | 0.000257 | 0.004767 | 0.003497 | 2366 | tags=51%, l | 79621/341 | LNCaP-NCOR2 |
| REACTOME | REACTOME | REACTOME | 57 | -0.689035 | -2.709106 | 0.000257 | 0.004767 | 0.003497 | 1417 | tags=49%, l | 8363/3015 | LNCaP-NCOR2 |
| KEGG_SYST | KEGG_SYST | KEGG_SYST | 58 | -0.764683 | -3.01282 | 0.000258 | 0.004767 | 0.003497 | 1417 | tags=60%, l | 8363/9555 | LNCaP-NCOR2 |
| REACTOME | REACTOME | REACTOME | 60 | -0.749725 | -2.973468 | 0.000259 | 0.004767 | 0.003497 | 1406 | tags=53%, l | 5984/3015 | LNCaP-NCOR2 |
| RHODES_U | RHODES_U | RHODES_U | 60 | -0.556261 | -2.206177 | 0.000259 | 0.004767 | 0.003497 | 3562 | tags=60%, l | 5111/1058 | LNCaP-NCOR2 |
| REACTOME | REACTOME | REACTOME | 59 | -0.696339 | -2.750498 | 0.000259 | 0.004767 | 0.003497 | 1812 | tags=58%, l | 3021/1019 | LNCaP-NCOR2 |
| CHICAS_RB | CHICAS_RB | CHICAS_RB | 61 | -0.497879 | -1.982727 | 0.00026 | 0.004767 | 0.003497 | 2801 | tags=54%, l | 1350/8372 | LNCaP-NCOR2 |
| MORI_PRE_ | MORI_PRE_ | MORI_PRE_ | 61 | -0.506292 | -2.016228 | 0.00026 | 0.004767 | 0.003497 | 2096 | tags=44%, l | 10797/262 | LNCaP-NCOR2 |
| REACTOME | REACTOME | REACTOME | 61 | -0.677792 | -2.699202 | 0.00026 | 0.004767 | 0.003497 | 1417 | tags=46%, l | 8363/3015 | LNCaP-NCOR2 |
| CROONQUI | CROONQUI | CROONQUI | 64 | -0.516935 | -2.074207 | 0.000263 | 0.004767 | 0.003497 | 1137 | tags=28%, l | 6192/8970 | LNCaP-NCOR2 |
| REACTOME | REACTOME | REACTOME | 64 | -0.723493 | -2.903025 | 0.000263 | 0.004767 | 0.003497 | 1417 | tags=56%, l | 3021/2965 | LNCaP-NCOR2 |
| SANA_RESI | SANA_RESI | SANA_RESI | 64 | -0.479095 | -1.922376 | 0.000263 | 0.004767 | 0.003497 | 2806 | tags=45%, l | 3155/9643 | LNCaP-NCOR2 |
| MOOTHA_ | MOOTHA_ | MOOTHA_ | 66 | -0.567946 | -2.292552 | 0.000265 | 0.004767 | 0.003497 | 3593 | tags=65%, l | 4702/1349 | LNCaP-NCOR2 |
| ZHOU_CELI | ZHOU_CELI | ZHOU_CELI | 66 | -0.658822 | -2.659383 | 0.000265 | 0.004767 | 0.003497 | 2549 | tags=62%, l | 4605/6118 | LNCaP-NCOR2 |
| REACTOME | REACTOME | REACTOME | 65 | -0.617919 | -2.484127 | 0.000265 | 0.004767 | 0.003497 | 1549 | tags=45%, l | 3020/8363 | LNCaP-NCOR2 |
| REACTOME | REACTOME | REACTOME | 65 | -0.666694 | -2.680211 | 0.000265 | 0.004767 | 0.003497 | 1812 | tags=52%, l | 3021/7531 | LNCaP-NCOR2 |
| CROONQUI | CROONQUI | CROONQUI | 67 | -0.575067 | -2.32682 | 0.000265 | 0.004767 | 0.003497 | 1657 | tags=33%, l | 3835/4176 | LNCaP-NCOR2 |
| GAVIN_FO | GAVIN_FO | GAVIN_FO | 67 | -0.557524 | -2.255841 | 0.000265 | 0.004767 | 0.003497 | 2389 | tags=43%, l | 4751/5740 | LNCaP-NCOR2 |
| NIKOLSKY_ | NIKOLSKY_ | NIKOLSKY_ | 67 | -0.51398 | -2.079652 | 0.000265 | 0.004767 | 0.003497 | 3452 | tags=58%, l | 10573/724 | LNCaP-NCOR2 |
| GRAHAM_I | GRAHAM_I | GRAHAM_I | 69 | -0.578308 | -2.351371 | 0.000266 | 0.004767 | 0.003497 | 2883 | tags=52%, l | 10403/395 | LNCaP-NCOR2 |
| REACTOME | REACTOME | REACTOME | 69 | -0.507543 | -2.063644 | 0.000266 | 0.004767 | 0.003497 | 2901 | tags=54%, l | 5427/5719 | LNCaP-NCOR2 |
| REACTOME | REACTOME | REACTOME | 69 | -0.686031 | -2.789366 | 0.000266 | 0.004767 | 0.003497 | 980 | tags=45%, l | 8970/8335 | LNCaP-NCOR2 |
| REACTOME | REACTOME | REACTOME | 70 | -0.535053 | -2.184388 | 0.000267 | 0.004767 | 0.003497 | 2271 | tags=49%, l | 5982/6396 | LNCaP-NCOR2 |
| MORI_IMM | MORI_IMM | MORI_IMM | 76 | -0.567886 | -2.354099 | 0.000267 | 0.004767 | 0.003497 | 1711 | tags=39%, l | 55388/417 | LNCaP-NCOR2 |
| REACTOME | REACTOME | REACTOME | 71 | -0.608882 | -2.490316 | 0.000267 | 0.004767 | 0.003497 | 1812 | tags=46%, l | 3021/1869 | LNCaP-NCOR2 |
| REACTOME | REACTOME | REACTOME | 76 | -0.563017 | -2.333916 | 0.000267 | 0.004767 | 0.003497 | 2141 | tags=46%, l | 51295/738 | LNCaP-NCOR2 |
| TAKAO_RES | TAKAO_RES | TAKAO_RES | 71 | -0.527497 | -2.157454 | 0.000267 | 0.004767 | 0.003497 | 1917 | tags=39%, l | 8848/1327 | LNCaP-NCOR2 |
| REACTOME | REACTOME | REACTOME | 72 | -0.70155 | -2.875903 | 0.000268 | 0.004767 | 0.003497 | 1443 | tags=49%, l | 8363/5984 | LNCaP-NCOR2 |
| REACTOME | REACTOME | REACTOME | 77 | -0.667477 | -2.772282 | 0.000268 | 0.004767 | 0.003497 | 1488 | tags=51%, l | 60/3020/8 | LNCaP-NCOR2 |
| REACTOME | REACTOME | REACTOME | 77 | -0.595258 | -2.472329 | 0.000268 | 0.004767 | 0.003497 | 1841 | tags=44%, l | 8342/3021 | LNCaP-NCOR2 |
| REACTOME | REACTOME | REACTOME | 77 | -0.6272 | -2.604998 | 0.000268 | 0.004767 | 0.003497 | 1417 | tags=44%, l | 8363/5438 | LNCaP-NCOR2 |
| KONG_E2F | KONG_E2F | KONG_E2F | 78 | -0.584649 | -2.434281 | 0.000268 | 0.004767 | 0.003497 | 2190 | tags=46%, l | 55166/650 | LNCaP-NCOR2 |
| REACTOME | REACTOME | REACTOME | 78 | -0.688838 | -2.868086 | 0.000268 | 0.004767 | 0.003497 | 1406 | tags=46%, l | 79172/598 | LNCaP-NCOR2 |
| REACTOME | REACTOME | REACTOME | 78 | -0.661359 | -2.753674 | 0.000268 | 0.004767 | 0.003497 | 1812 | tags=51%, l | 3021/1019 | LNCaP-NCOR2 |
| KEGG_RIBO | KEGG_RIBO | KEGG_RIBO | 73 | -0.716883 | -2.943678 | 0.000269 | 0.004767 | 0.003497 | 2121 | tags=73%, l | 6191/6189 | LNCaP-NCOR2 |
| MORI_LAR | MORI_LAR | MORI_LAR | 73 | -0.60648 | -2.490337 | 0.000269 | 0.004767 | 0.003497 | 2843 | tags=60%, l | 7153/5700 | LNCaP-NCOR2 |
| REACTOME | REACTOME | REACTOME | 79 | -0.528692 | -2.206886 | 0.00027 | 0.004767 | 0.003497 | 2141 | tags=44%, l | 51295/738 | LNCaP-NCOR2 |
| REACTOME | REACTOME | REACTOME | 80 | -0.615424 | -2.576952 | 0.00027 | 0.004767 | 0.003497 | 1499 | tags=45%, l | 3020/8363 | LNCaP-NCOR2 |
| REACTOME | REACTOME | REACTOME | 80 | -0.65741 | -2.752757 | 0.00027 | 0.004767 | 0.003497 | 1812 | tags=51%, l | 3021/2965 | LNCaP-NCOR2 |
| REACTOME | REACTOME | REACTOME | 81 | -0.651912 | -2.733576 | 0.000271 | 0.004767 | 0.003497 | 1417 | tags=53%, l | 5435/7959 | LNCaP-NCOR2 |
| REACTOME | REACTOME | REACTOME | 81 | -0.670891 | -2.81316 | 0.000271 | 0.004767 | 0.003497 | 2121 | tags=67%, l | 6191/6189 | LNCaP-NCOR2 |
| KEGG_OXID | KEGG_OXID | KEGG_OXID | 84 | -0.502187 | -2.120447 | 0.000274 | 0.004767 | 0.003497 | 2141 | tags=40%, l | 7385/4708 | LNCaP-NCOR2 |
| BURTON_A | BURTON_A | BURTON_A | 86 | -0.526998 | -2.23506 | 0.000274 | 0.004767 | 0.003497 | 1661 | tags=41%, l | 3835/5538 | LNCaP-NCOR2 |
| REACTOME | REACTOME | REACTOME | 88 | -0.582925 | -2.485118 | 0.000274 | 0.004767 | 0.003497 | 1468 | tags=39%, l | 60/8363/5 | LNCaP-NCOR2 |

| | | | | | | | | | | | | |
|---|---|---|---|---|---|---|---|---|---|---|---|---|
| CROONQUI | CROONQUI | CROONQUI | 85 | -0.530462 | -2.245623 | 0.000274 | 0.004767 | 0.003497 | 3562 | tags=56%, l | 2879/5111, | LNCaP-NCOR2 |
| REACTOME | REACTOME | REACTOME | 85 | -0.629157 | -2.663429 | 0.000274 | 0.004767 | 0.003497 | 1812 | tags=48%, l | 3021/1019, | LNCaP-NCOR2 |
| GRAESSMA | GRAESSMA | GRAESSMA | 331 | 0.322724 | 1.52982 | 0.000275 | 0.004767 | 0.003497 | 1841 | tags=24%, l | 5784/2899 | LNCaP-NCOR2 |
| JOHNSTON | JOHNSTON | JOHNSTON | 327 | 0.330625 | 1.565383 | 0.000275 | 0.004767 | 0.003497 | 2027 | tags=27%, l | 23092/553 | LNCaP-NCOR2 |
| REACTOME | REACTOME | REACTOME | 92 | -0.6728 | -2.885106 | 0.000278 | 0.004767 | 0.003497 | 2121 | tags=64%, l | 6191/6189, | LNCaP-NCOR2 |
| REACTOME | REACTOME | REACTOME | 92 | -0.59541 | -2.553244 | 0.000278 | 0.004767 | 0.003497 | 1531 | tags=55%, l | 5719/5695, | LNCaP-NCOR2 |
| REACTOME | REACTOME | REACTOME | 93 | -0.546595 | -2.34805 | 0.000278 | 0.004767 | 0.003497 | 1190 | tags=38%, l | 8363/5438, | LNCaP-NCOR2 |
| REACTOME | REACTOME | REACTOME | 93 | -0.507872 | -2.181705 | 0.000278 | 0.004767 | 0.003497 | 1856 | tags=44%, l | 8342/3021, | LNCaP-NCOR2 |
| ZHOU_CELI | ZHOU_CELI | ZHOU_CELI | 96 | -0.512049 | -2.209433 | 0.000279 | 0.004767 | 0.003497 | 2549 | tags=54%, l | 890/51002, | LNCaP-NCOR2 |
| RUTELLA_R | RUTELLA_R | RUTELLA_R | 272 | 0.345741 | 1.608433 | 0.00028 | 0.004767 | 0.003497 | 3536 | tags=43%, l | 3572/2336 | LNCaP-NCOR2 |
| CHARAFE_I | CHARAFE_I | CHARAFE_I | 280 | 0.346769 | 1.617334 | 0.00028 | 0.004767 | 0.003497 | 1878 | tags=26%, l | 5793/3685, | LNCaP-NCOR2 |
| CHEMNITZ_ | CHEMNITZ_ | CHEMNITZ_ | 105 | -0.473488 | -2.074947 | 0.000281 | 0.004767 | 0.003497 | 2069 | tags=35%, l | 10797/105 | LNCaP-NCOR2 |
| RUIZ_TNC_ | RUIZ_TNC_ | RUIZ_TNC_ | 105 | -0.461053 | -2.020455 | 0.000281 | 0.004767 | 0.003497 | 2346 | tags=42%, l | 57405/450 | LNCaP-NCOR2 |
| MISSIAGLIA | MISSIAGLIA | MISSIAGLIA | 97 | -0.513061 | -2.21466 | 0.000281 | 0.004767 | 0.003497 | 1977 | tags=40%, l | 10535/590 | LNCaP-NCOR2 |
| REACTOME | REACTOME | REACTOME | 97 | -0.620823 | -2.679824 | 0.000281 | 0.004767 | 0.003497 | 2121 | tags=60%, l | 6191/6189, | LNCaP-NCOR2 |
| REACTOME | REACTOME | REACTOME | 97 | -0.662232 | -2.858568 | 0.000281 | 0.004767 | 0.003497 | 2487 | tags=66%, l | 6224/6166, | LNCaP-NCOR2 |
| REACTOME | REACTOME | REACTOME | 98 | -0.647671 | -2.801949 | 0.000281 | 0.004767 | 0.003497 | 2487 | tags=63%, l | 6224/6166, | LNCaP-NCOR2 |
| WHITEFOR | WHITEFOR | WHITEFOR | 99 | -0.561667 | -2.432197 | 0.000281 | 0.004767 | 0.003497 | 3172 | tags=58%, l | 22974/157 | LNCaP-NCOR2 |
| REACTOME | REACTOME | REACTOME | 103 | -0.496881 | -2.17031 | 0.000282 | 0.004767 | 0.003497 | 1134 | tags=32%, l | 8367/2237, | LNCaP-NCOR2 |
| REACTOME | REACTOME | REACTOME | 107 | -0.532733 | -2.341013 | 0.000282 | 0.004767 | 0.003497 | 1812 | tags=37%, l | 3021/8480, | LNCaP-NCOR2 |
| REACTOME | REACTOME | REACTOME | 106 | -0.474259 | -2.079031 | 0.000283 | 0.004767 | 0.003497 | 3172 | tags=54%, l | 64785/646 | LNCaP-NCOR2 |
| REACTOME | REACTOME | REACTOME | 110 | -0.540706 | -2.384041 | 0.000284 | 0.004767 | 0.003497 | 1422 | tags=38%, l | 3020/8363, | LNCaP-NCOR2 |
| PECE_MAM | PECE_MAM | PECE_MAM | 102 | -0.542175 | -2.360593 | 0.000284 | 0.004767 | 0.003497 | 2450 | tags=50%, l | 6166/4502, | LNCaP-NCOR2 |
| REACTOME | REACTOME | REACTOME | 112 | -0.564549 | -2.497422 | 0.000285 | 0.004767 | 0.003497 | 1488 | tags=40%, l | 60/3020/83 | LNCaP-NCOR2 |
| KIM_ALL_D | KIM_ALL_D | KIM_ALL_D | 113 | -0.56127 | -2.485967 | 0.000285 | 0.004767 | 0.003497 | 2289 | tags=49%, l | 8887/8185 | LNCaP-NCOR2 |
| ROSTY_CEF | ROSTY_CEF | ROSTY_CEF | 121 | -0.527746 | -2.365554 | 0.000286 | 0.004767 | 0.003497 | 2366 | tags=45%, l | 4173/5514 | LNCaP-NCOR2 |
| REACTOME | REACTOME | REACTOME | 117 | -0.450885 | -2.010264 | 0.000286 | 0.004767 | 0.003497 | 2349 | tags=42%, l | 2271/3418, | LNCaP-NCOR2 |
| REACTOME | REACTOME | REACTOME | 126 | -0.451039 | -2.033089 | 0.000288 | 0.004767 | 0.003497 | 2590 | tags=42%, l | 9031/322/( | LNCaP-NCOR2 |
| REACTOME | REACTOME | REACTOME | 128 | -0.473103 | -2.139823 | 0.000289 | 0.004767 | 0.003497 | 2952 | tags=50%, l | 7316/5719, | LNCaP-NCOR2 |
| REACTOME | REACTOME | REACTOME | 129 | -0.409755 | -1.856389 | 0.000289 | 0.004767 | 0.003497 | 2901 | tags=44%, l | 5427/5719, | LNCaP-NCOR2 |
| WINNEPEN | WINNEPEN | WINNEPEN | 129 | -0.418219 | -1.894734 | 0.000289 | 0.004767 | 0.003497 | 2366 | tags=36%, l | 4173/5740 | LNCaP-NCOR2 |
| HOFFMANI | HOFFMANI | HOFFMANI | 127 | -0.489451 | -2.208552 | 0.000289 | 0.004767 | 0.003497 | 2170 | tags=38%, l | 9493/6494 | LNCaP-NCOR2 |
| REACTOME | REACTOME | REACTOME | 132 | -0.529577 | -2.410686 | 0.00029 | 0.004767 | 0.003497 | 2133 | tags=51%, l | 6170/6191, | LNCaP-NCOR2 |
| SOTIRIOU_ | SOTIRIOU_ | SOTIRIOU_ | 132 | -0.507849 | -2.311781 | 0.00029 | 0.004767 | 0.003497 | 3562 | tags=55%, l | 23560/414 | LNCaP-NCOR2 |
| FLORIO_NE | FLORIO_NE | FLORIO_NE | 134 | -0.414593 | -1.889968 | 0.000291 | 0.004767 | 0.003497 | 2370 | tags=37%, l | 4751/5740 | LNCaP-NCOR2 |
| IVANOVA_I | IVANOVA_I | IVANOVA_I | 190 | 0.382483 | 1.708269 | 0.000292 | 0.004767 | 0.003497 | 2283 | tags=32%, l | 51274/657 | LNCaP-NCOR2 |
| SARRIO_EP | SARRIO_EP | SARRIO_EP | 137 | -0.486645 | -2.228573 | 0.000292 | 0.004767 | 0.003497 | 2346 | tags=41%, l | 4502/9156, | LNCaP-NCOR2 |
| DANG_REG | DANG_REG | DANG_REG | 174 | 0.386205 | 1.705512 | 0.000294 | 0.004767 | 0.003497 | 2089 | tags=30%, l | 4193/3915 | LNCaP-NCOR2 |
| IWANAGA_ | IWANAGA_ | IWANAGA_ | 173 | 0.37998 | 1.676168 | 0.000294 | 0.004767 | 0.003497 | 1487 | tags=23%, l | 4254/1672 | LNCaP-NCOR2 |
| REACTOME | REACTOME | REACTOME | 141 | -0.535299 | -2.459565 | 0.000296 | 0.004767 | 0.003497 | 2489 | tags=53%, l | 140801/56 | LNCaP-NCOR2 |
| LI_AMPLIFI | LI_AMPLIFI | LI_AMPLIFI | 142 | -0.417573 | -1.920273 | 0.000297 | 0.004767 | 0.003497 | 1689 | tags=39%, l | 3249/6337, | LNCaP-NCOR2 |
| YAO_TEMP | YAO_TEMP | YAO_TEMP | 147 | -0.489726 | -2.261039 | 0.000297 | 0.004767 | 0.003497 | 1958 | tags=48%, l | 64960/569 | LNCaP-NCOR2 |
| DAZARD_R | DAZARD_R | DAZARD_R | 143 | -0.48002 | -2.208469 | 0.000297 | 0.004767 | 0.003497 | 2033 | tags=36%, l | 8971/2067, | LNCaP-NCOR2 |
| GRAHAM_( | GRAHAM_( | GRAHAM_( | 143 | -0.443541 | -2.040636 | 0.000297 | 0.004767 | 0.003497 | 2913 | tags=45%, l | 5427/6496 | LNCaP-NCOR2 |
| REACTOME | REACTOME | REACTOME | 151 | -0.552054 | -2.561646 | 0.000298 | 0.004767 | 0.003497 | 2487 | tags=48%, l | 51202/622 | LNCaP-NCOR2 |
| REACTOME | REACTOME | REACTOME | 148 | -0.417365 | -1.928465 | 0.000298 | 0.004767 | 0.003497 | 1856 | tags=34%, l | 51433/834 | LNCaP-NCOR2 |
| PLASARI_T | PLASARI_T | PLASARI_T | 149 | 0.39228 | 1.696008 | 0.0003 | 0.004767 | 0.003497 | 2485 | tags=37%, l | 3572/4254, | LNCaP-NCOR2 |
| MANALO_H | MANALO_H | MANALO_H | 144 | 0.403403 | 1.736587 | 0.000301 | 0.004767 | 0.003497 | 3280 | tags=40%, l | 8795/8974, | LNCaP-NCOR2 |
| PAL_PRMT | PAL_PRMT | PAL_PRMT | 154 | -0.454077 | -2.111239 | 0.000301 | 0.004767 | 0.003497 | 2092 | tags=32%, l | 3832/1042 | LNCaP-NCOR2 |
| WANG_CLI | WANG_CLI | WANG_CLI | 136 | 0.415799 | 1.775094 | 0.000304 | 0.004767 | 0.003497 | 3303 | tags=47%, l | 55500/803 | LNCaP-NCOR2 |
| REACTOME | REACTOME | REACTOME | 156 | -0.550377 | -2.559729 | 0.000304 | 0.004767 | 0.003497 | 2487 | tags=48%, l | 51202/622 | LNCaP-NCOR2 |
| REACTOME | REACTOME | REACTOME | 159 | -0.475844 | -2.218874 | 0.000305 | 0.004767 | 0.003497 | 1422 | tags=31%, l | 3020/8363, | LNCaP-NCOR2 |
| KONDO_EZ | KONDO_EZ | KONDO_EZ | 123 | 0.439278 | 1.848056 | 0.000306 | 0.004767 | 0.003497 | 2440 | tags=34%, l | 5865/5205, | LNCaP-NCOR2 |
| HALLMARK | HALLMARK | HALLMARK | 125 | 0.430512 | 1.814492 | 0.000306 | 0.004767 | 0.003497 | 2574 | tags=34%, l | 57169/576 | LNCaP-NCOR2 |
| FISCHER_G | FISCHER_G | FISCHER_G | 161 | -0.466395 | -2.176295 | 0.000307 | 0.004767 | 0.003497 | 1747 | tags=37%, l | 1869/9401 | LNCaP-NCOR2 |
| REACTOME | REACTOME | REACTOME | 161 | -0.445122 | -2.077031 | 0.000307 | 0.004767 | 0.003497 | 1531 | tags=29%, l | 10213/262 | LNCaP-NCOR2 |
| ZHENG_GL | ZHENG_GL | ZHENG_GL | 164 | -0.377299 | -1.767968 | 0.000307 | 0.004767 | 0.003497 | 2386 | tags=38%, l | 4173/4502, | LNCaP-NCOR2 |
| REACTOME | REACTOME | REACTOME | 165 | -0.39503 | -1.852716 | 0.000307 | 0.004767 | 0.003497 | 2901 | tags=41%, l | 5719/1499, | LNCaP-NCOR2 |
| ELVIDGE_H | ELVIDGE_H | ELVIDGE_H | 117 | 0.43608 | 1.819149 | 0.000308 | 0.004767 | 0.003497 | 1539 | tags=27%, l | 1824/6653, | LNCaP-NCOR2 |
| BOYLAN_M | BOYLAN_M | BOYLAN_M | 122 | 0.418894 | 1.759074 | 0.000308 | 0.004767 | 0.003497 | 2577 | tags=38%, l | 79993/107 | LNCaP-NCOR2 |
| HALLMARK | HALLMARK | HALLMARK | 170 | -0.41968 | -1.980809 | 0.000308 | 0.004767 | 0.003497 | 2815 | tags=41%, l | 3326/5169 | LNCaP-NCOR2 |
| REACTOME | REACTOME | REACTOME | 170 | -0.41465 | -1.957068 | 0.000308 | 0.004767 | 0.003497 | 2540 | tags=37%, l | 1950/5709, | LNCaP-NCOR2 |
| FUJII_YBX1 | FUJII_YBX1 | FUJII_YBX1 | 168 | -0.421775 | -1.98543 | 0.00031 | 0.004767 | 0.003497 | 2550 | tags=42%, l | 4605/2340 | LNCaP-NCOR2 |
| SANSOM_A | SANSOM_A | SANSOM_A | 168 | -0.379229 | -1.785156 | 0.00031 | 0.004767 | 0.003497 | 1994 | tags=29%, l | 10535/850 | LNCaP-NCOR2 |
| LIN_NPAS4 | LIN_NPAS4 | LIN_NPAS4 | 103 | 0.45163 | 1.849336 | 0.00031 | 0.004767 | 0.003497 | 2888 | tags=37%, l | 5865/8012 | LNCaP-NCOR2 |
| GARGALOV | GARGALOV | GARGALOV | 97 | 0.450391 | 1.82732 | 0.00031 | 0.004767 | 0.003497 | 2408 | tags=35%, l | 4254/8546 | LNCaP-NCOR2 |

| | | | | | | | | | | | | |
|---|---|---|---|---|---|---|---|---|---|---|---|---|
| PID_P53_D | PID_P53_D | PID_P53_D | 105 | 0.446445 | 1.832052 | 0.000311 | 0.004767 | 0.003497 | 1906 | tags=35%, | 4193/8795, | LNCaP-NCOR2 |
| HALLMARK | HALLMARK | HALLMARK | 173 | -0.474776 | -2.244113 | 0.000312 | 0.004767 | 0.003497 | 1982 | tags=38%, | 10797/105 | LNCaP-NCOR2 |
| HALLMARK | HALLMARK | HALLMARK | 176 | -0.429928 | -2.034974 | 0.000313 | 0.004767 | 0.003497 | 3302 | tags=51%, | 2879/1666, | LNCaP-NCOR2 |
| HALLMARK | HALLMARK | HALLMARK | 175 | -0.379436 | -1.794443 | 0.000314 | 0.004767 | 0.003497 | 2985 | tags=40%, | 6558/7486, | LNCaP-NCOR2 |
| MARKEY_R | MARKEY_R | MARKEY_R | 180 | -0.393048 | -1.867805 | 0.000315 | 0.004767 | 0.003497 | 2366 | tags=41%, | 8270/4173, | LNCaP-NCOR2 |
| REACTOME | REACTOME | REACTOME | 183 | -0.46121 | -2.197957 | 0.000315 | 0.004767 | 0.003497 | 1422 | tags=29%, | 3020/8363, | LNCaP-NCOR2 |
| REACTOME | REACTOME | REACTOME | 179 | -0.4456 | -2.114236 | 0.000316 | 0.004767 | 0.003497 | 1488 | tags=27%, | 6944/60/8 | LNCaP-NCOR2 |
| REACTOME | REACTOME | REACTOME | 179 | -0.444259 | -2.107873 | 0.000316 | 0.004767 | 0.003497 | 2489 | tags=50%, | 5719/5695, | LNCaP-NCOR2 |
| LU_EZH2_T | LU_EZH2_T | LU_EZH2_T | 190 | -0.389971 | -1.868622 | 0.000317 | 0.004767 | 0.003497 | 3267 | tags=47%, | 6284/1054 | LNCaP-NCOR2 |
| WONG_MI | WONG_MI | WONG_MI | 190 | -0.422352 | -2.023779 | 0.000317 | 0.004767 | 0.003497 | 2349 | tags=38%, | 27349/483 | LNCaP-NCOR2 |
| DE_YY1_TA | DE_YY1_TA | DE_YY1_TA | 74 | 0.478965 | 1.863569 | 0.000319 | 0.004767 | 0.003497 | 3020 | tags=51%, | 3685/2899 | LNCaP-NCOR2 |
| TAKEDA_TA | TAKEDA_TA | TAKEDA_TA | 77 | 0.473804 | 1.854817 | 0.000319 | 0.004767 | 0.003497 | 2154 | tags=38%, | 57674/550 | LNCaP-NCOR2 |
| KOBAYASH | KOBAYASH | KOBAYASH | 211 | -0.495922 | -2.410925 | 0.000324 | 0.004767 | 0.003497 | 2366 | tags=43%, | 4173/5740 | LNCaP-NCOR2 |
| IZADPANAH | IZADPANAH | IZADPANAH | 60 | 0.518497 | 1.938561 | 0.000326 | 0.004767 | 0.003497 | 2636 | tags=40%, | 1290/1607 | LNCaP-NCOR2 |
| ALTEMEIER | ALTEMEIER | ALTEMEIER | 54 | 0.51867 | 1.901111 | 0.000329 | 0.004767 | 0.003497 | 2445 | tags=48%, | 9180/6413 | LNCaP-NCOR2 |
| BASAKI_YB | BASAKI_YB | BASAKI_YB | 216 | -0.401092 | -1.953512 | 0.000329 | 0.004767 | 0.003497 | 2366 | tags=33%, | 4173/7908 | LNCaP-NCOR2 |
| CHEBOTAE | CHEBOTAE | CHEBOTAE | 52 | 0.524364 | 1.907104 | 0.000329 | 0.004767 | 0.003497 | 1936 | tags=38%, | 5152/1078 | LNCaP-NCOR2 |
| GRADE_CO | GRADE_CO | GRADE_CO | 219 | -0.354608 | -1.73023 | 0.00033 | 0.004767 | 0.003497 | 2393 | tags=32%, | 6242/4173, | LNCaP-NCOR2 |
| GOLDRATH | GOLDRATH | GOLDRATH | 229 | -0.355161 | -1.746004 | 0.00033 | 0.004767 | 0.003497 | 1711 | tags=28%, | 2597/699/ | LNCaP-NCOR2 |
| SANA_TNF_ | SANA_TNF_ | SANA_TNF_ | 49 | 0.525408 | 1.888187 | 0.00033 | 0.004767 | 0.003497 | 3522 | tags=67%, | 3685/5462 | LNCaP-NCOR2 |
| REACTOME | REACTOME | REACTOME | 226 | -0.337076 | -1.652264 | 0.00033 | 0.004767 | 0.003497 | 1531 | tags=23%, | 10213/569 | LNCaP-NCOR2 |
| BIOCARTA_ | BIOCARTA_ | BIOCARTA_ | 48 | 0.529537 | 1.895377 | 0.000332 | 0.004767 | 0.003497 | 1534 | tags=31%, | 4193/5591, | LNCaP-NCOR2 |
| CHANG_IM | CHANG_IM | CHANG_IM | 42 | 0.585749 | 2.045542 | 0.000332 | 0.004767 | 0.003497 | 2536 | tags=43%, | 1824/93/8 | LNCaP-NCOR2 |
| PHONG_TN | PHONG_TN | PHONG_TN | 42 | 0.569074 | 1.98731 | 0.000332 | 0.004767 | 0.003497 | 2846 | tags=52%, | 8795/1316, | LNCaP-NCOR2 |
| LINDGREN_ | LINDGREN_ | LINDGREN_ | 247 | -0.356656 | -1.764549 | 0.000339 | 0.004767 | 0.003497 | 2366 | tags=34%, | 10313/163 | LNCaP-NCOR2 |
| MANALO_H | MANALO_H | MANALO_H | 244 | -0.318407 | -1.574482 | 0.000339 | 0.004767 | 0.003497 | 2346 | tags=37%, | 890/22974, | LNCaP-NCOR2 |
| BROWNE_I | BROWNE_I | BROWNE_I | 33 | 0.6239 | 2.056951 | 0.00034 | 0.004767 | 0.003497 | 2482 | tags=55%, | 3572/5781, | LNCaP-NCOR2 |
| KERLEY_RE | KERLEY_RE | KERLEY_RE | 32 | 0.653502 | 2.141631 | 0.00034 | 0.004767 | 0.003497 | 1668 | tags=47%, | 64393/474 | LNCaP-NCOR2 |
| WILCOX_RI | WILCOX_RI | WILCOX_RI | 30 | 0.620158 | 2.003824 | 0.00034 | 0.004767 | 0.003497 | 1921 | tags=37%, | 4193/3480, | LNCaP-NCOR2 |
| REACTOME | REACTOME | REACTOME | 251 | -0.369925 | -1.833915 | 0.00034 | 0.004767 | 0.003497 | 1902 | tags=26%, | 30851/834 | LNCaP-NCOR2 |
| REACTOME | REACTOME | REACTOME | 254 | -0.503003 | -2.497637 | 0.000341 | 0.004767 | 0.003497 | 2121 | tags=41%, | 65003/116 | LNCaP-NCOR2 |
| DACOSTA_ | DACOSTA_ | DACOSTA_ | 255 | -0.338161 | -1.679564 | 0.000342 | 0.004767 | 0.003497 | 1971 | tags=27%, | 573/5702/ | LNCaP-NCOR2 |
| SCHAEFFER | SCHAEFFER | SCHAEFFER | 29 | 0.622743 | 1.991112 | 0.000343 | 0.004767 | 0.003497 | 3611 | tags=69%, | 4430/8020 | LNCaP-NCOR2 |
| WARTERS_ | WARTERS_ | WARTERS_ | 29 | 0.596228 | 1.906336 | 0.000343 | 0.004767 | 0.003497 | 2530 | tags=55%, | 4193/7832, | LNCaP-NCOR2 |
| ZHAN_MUI | ZHAN_MUI | ZHAN_MUI | 29 | 0.602237 | 1.925549 | 0.000343 | 0.004767 | 0.003497 | 1403 | tags=34%, | 5865/1078 | LNCaP-NCOR2 |
| CUI_TCF21 | CUI_TCF21 | CUI_TCF21 | 257 | -0.389413 | -1.936627 | 0.000343 | 0.004767 | 0.003497 | 2227 | tags=34%, | 83451/666 | LNCaP-NCOR2 |
| DUTTA_AP | DUTTA_AP | DUTTA_AP | 25 | 0.649737 | 2.001051 | 0.000346 | 0.004767 | 0.003497 | 1170 | tags=40%, | 4193/8795, | LNCaP-NCOR2 |
| DUTERTRE_ | DUTERTRE_ | DUTERTRE_ | 262 | -0.497774 | -2.477601 | 0.000347 | 0.004767 | 0.003497 | 2125 | tags=40%, | 26271/949 | LNCaP-NCOR2 |
| LINDGREN_ | LINDGREN_ | LINDGREN_ | 270 | -0.310831 | -1.553337 | 0.000347 | 0.004767 | 0.003497 | 2985 | tags=37%, | 6558/7077, | LNCaP-NCOR2 |
| REACTOME | REACTOME | REACTOME | 267 | -0.416623 | -2.077503 | 0.000347 | 0.004767 | 0.003497 | 2601 | tags=42%, | 80198/508 | LNCaP-NCOR2 |
| REACTOME | REACTOME | REACTOME | 270 | -0.462368 | -2.310625 | 0.000347 | 0.004767 | 0.003497 | 2489 | tags=44%, | 140801/56 | LNCaP-NCOR2 |
| WANG_RES | WANG_RES | WANG_RES | 264 | -0.360503 | -1.795608 | 0.000348 | 0.004767 | 0.003497 | 2370 | tags=34%, | 4173/2734 | LNCaP-NCOR2 |
| BLUM_RES | BLUM_RES | BLUM_RES | 279 | -0.347034 | -1.739737 | 0.000351 | 0.004768 | 0.003498 | 2883 | tags=38%, | 57122/107 | LNCaP-NCOR2 |
| WONG_EM | WONG_EM | WONG_EM | 281 | -0.444129 | -2.228318 | 0.000352 | 0.004768 | 0.003498 | 2892 | tags=41%, | 5427/5712 | LNCaP-NCOR2 |
| FARMER_B | FARMER_B | FARMER_B | 12 | 0.77665 | 1.983518 | 0.000352 | 0.004768 | 0.003498 | 1689 | tags=75%, | 55008/511 | LNCaP-NCOR2 |
| SCHAVOLT_ | SCHAVOLT_ | SCHAVOLT_ | 12 | 0.789244 | 2.015683 | 0.000352 | 0.004768 | 0.003498 | 1668 | tags=67%, | 4193/5048 | LNCaP-NCOR2 |
| REACTOME | REACTOME | REACTOME | 11 | 0.754647 | 1.877678 | 0.000354 | 0.004778 | 0.003505 | 1121 | tags=55%, | 7706/6413 | LNCaP-NCOR2 |
| REACTOME | REACTOME | REACTOME | 303 | -0.343172 | -1.740552 | 0.000356 | 0.00478 | 0.003506 | 2509 | tags=32%, | 5709/9702, | LNCaP-NCOR2 |
| RHEIN_ALL | RHEIN_ALL | RHEIN_ALL | 300 | -0.383235 | -1.93883 | 0.000357 | 0.004785 | 0.00351 | 2003 | tags=29%, | 4190/708/ | LNCaP-NCOR2 |
| VECCHI_GA | VECCHI_GA | VECCHI_GA | 313 | -0.385246 | -1.959892 | 0.000362 | 0.004811 | 0.003529 | 2591 | tags=35%, | 10857/838 | LNCaP-NCOR2 |
| HSIAO_HO | HSIAO_HO | HSIAO_HO | 318 | -0.472371 | -2.408084 | 0.000362 | 0.004811 | 0.003529 | 2342 | tags=41%, | 4502/8692, | LNCaP-NCOR2 |
| REACTOME | REACTOME | REACTOME | 314 | -0.37118 | -1.888848 | 0.000363 | 0.004811 | 0.003529 | 2906 | tags=41%, | 7316/5719, | LNCaP-NCOR2 |
| REACTOME | REACTOME | REACTOME | 330 | -0.312902 | -1.601116 | 0.000366 | 0.004838 | 0.003549 | 1902 | tags=23%, | 30851/982 | LNCaP-NCOR2 |
| REACTOME | REACTOME | REACTOME | 334 | -0.289766 | -1.484588 | 0.000368 | 0.004859 | 0.003564 | 1858 | tags=24%, | 57332/514 | LNCaP-NCOR2 |
| PUJANA_BI | PUJANA_BI | PUJANA_BI | 343 | -0.286099 | -1.469728 | 0.000374 | 0.004914 | 0.003605 | 2346 | tags=29%, | 79084/690 | LNCaP-NCOR2 |
| NUYTTEN_ | NUYTTEN_ | NUYTTEN_ | 626 | 0.280399 | 1.402476 | 0.000376 | 0.004922 | 0.00361 | 2501 | tags=29%, | 3572/3915, | LNCaP-NCOR2 |
| MOOTHA_ | MOOTHA_ | MOOTHA_ | 352 | -0.375486 | -1.935296 | 0.000378 | 0.004933 | 0.003619 | 2547 | tags=36%, | 2356/8192, | LNCaP-NCOR2 |
| MOOTHA_ | MOOTHA_ | MOOTHA_ | 359 | -0.360422 | -1.860688 | 0.000379 | 0.004938 | 0.003622 | 2352 | tags=32%, | 23600/273 | LNCaP-NCOR2 |
| FORTSCHE | FORTSCHE | FORTSCHE | 538 | 0.30323 | 1.499649 | 0.000383 | 0.004975 | 0.003649 | 1993 | tags=24%, | 79993/167 | LNCaP-NCOR2 |
| STARK_PRE | STARK_PRE | STARK_PRE | 365 | -0.399018 | -2.061504 | 0.000385 | 0.004981 | 0.003654 | 2601 | tags=36%, | 9993/6497 | LNCaP-NCOR2 |
| SHEDDEN_ | SHEDDEN_ | SHEDDEN_ | 371 | -0.317703 | -1.643265 | 0.000386 | 0.004981 | 0.003654 | 2404 | tags=29%, | 5688/4751, | LNCaP-NCOR2 |
| ENK_UV_R | ENK_UV_R | ENK_UV_R | 383 | -0.431121 | -2.236485 | 0.000398 | 0.005114 | 0.003751 | 2543 | tags=40%, | 645/1537/ | LNCaP-NCOR2 |
| SPIELMAN_ | SPIELMAN_ | SPIELMAN_ | 391 | -0.37939 | -1.973979 | 0.0004 | 0.005118 | 0.003754 | 2230 | tags=34%, | 5757/5691, | LNCaP-NCOR2 |
| REACTOME | REACTOME | REACTOME | 402 | -0.284211 | -1.48429 | 0.000401 | 0.005118 | 0.003754 | 1594 | tags=23%, | 25873/559 | LNCaP-NCOR2 |
| REACTOME | REACTOME | REACTOME | 397 | -0.281437 | -1.466844 | 0.000402 | 0.005118 | 0.003754 | 1860 | tags=24%, | 7266/5733 | LNCaP-NCOR2 |
| FEVR_CTN | FEVR_CTN | FEVR_CTN | 406 | -0.31399 | -1.639835 | 0.000407 | 0.00517 | 0.003793 | 1615 | tags=22%, | 3835/6749, | LNCaP-NCOR2 |

| | | | | | | | | | | | |
|---|---|---|---|---|---|---|---|---|---|---|---|
| GOBERT_O | GOBERT_O | GOBERT_O | 418 | -0.364524 | -1.90851 | 0.000408 | 0.00517 | 0.003793 | 2370 | tags=33%, l | 4751/4173, | LNCaP-NCOR2 |
| REACTOME | REACTOME | REACTOME | 427 | -0.356061 | -1.869059 | 0.000413 | 0.005214 | 0.003825 | 2509 | tags=33%, l | 51343/570 | LNCaP-NCOR2 |
| CHARAFE_I | CHARAFE_I | CHARAFE_I | 280 | 0.34173 | 1.59383 | 0.00042 | 0.005286 | 0.003877 | 3551 | tags=45%, l | 3915/8546, | LNCaP-NCOR2 |
| ZHOU_INFL | ZHOU_INFL | ZHOU_INFL | 227 | 0.362413 | 1.652007 | 0.000429 | 0.005388 | 0.003953 | 2681 | tags=34%, l | 79993/425 | LNCaP-NCOR2 |
| CAIRO_HEP | CAIRO_HEP | CAIRO_HEP | 488 | -0.378966 | -2.013766 | 0.000439 | 0.005491 | 0.004028 | 2366 | tags=34%, l | 4173/7908 | LNCaP-NCOR2 |
| BROWNE_I | BROWNE_I | BROWNE_I | 176 | 0.366022 | 1.618991 | 0.000441 | 0.005491 | 0.004028 | 2935 | tags=40%, l | 6651/5531 | LNCaP-NCOR2 |
| REACTOME | REACTOME | REACTOME | 499 | -0.323423 | -1.722912 | 0.000442 | 0.005491 | 0.004028 | 2509 | tags=32%, l | 51343/570 | LNCaP-NCOR2 |
| MARSON_E | MARSON_E | MARSON_E | 500 | -0.336483 | -1.792441 | 0.000443 | 0.005491 | 0.004028 | 2857 | tags=33%, l | 10733/104 | LNCaP-NCOR2 |
| MASSARWI | MASSARWI | MASSARWI | 160 | 0.383841 | 1.67839 | 0.000444 | 0.005492 | 0.004029 | 1865 | tags=24%, l | 65986/579 | LNCaP-NCOR2 |
| CASORELLI_ | CASORELLI_ | CASORELLI_ | 510 | -0.303505 | -1.619374 | 0.000445 | 0.005492 | 0.004029 | 2091 | tags=25%, l | 3832/5451 | LNCaP-NCOR2 |
| KIM_BIPOL | KIM_BIPOL | KIM_BIPOL | 514 | -0.373428 | -1.992231 | 0.000451 | 0.005519 | 0.004048 | 2015 | tags=30%, l | 7385/6881, | LNCaP-NCOR2 |
| REACTOME | REACTOME | REACTOME | 514 | -0.295693 | -1.577515 | 0.000451 | 0.005519 | 0.004048 | 3055 | tags=35%, l | 11193/339 | LNCaP-NCOR2 |
| KIM_WT1_ | KIM_WT1_ | KIM_WT1_ | 150 | 0.385355 | 1.66667 | 0.000452 | 0.005519 | 0.004048 | 3561 | tags=49%, l | 8507/5472 | LNCaP-NCOR2 |
| OSWALD_H | OSWALD_H | OSWALD_H | 136 | 0.400546 | 1.709974 | 0.000455 | 0.005549 | 0.004071 | 2693 | tags=38%, l | 6741/9314, | LNCaP-NCOR2 |
| SMIRNOV_ | SMIRNOV_ | SMIRNOV_ | 120 | 0.409377 | 1.71422 | 0.000462 | 0.005563 | 0.004081 | 2394 | tags=32%, l | 4193/8795, | LNCaP-NCOR2 |
| WAMUNYO | WAMUNYO | WAMUNYO | 113 | 0.412335 | 1.711955 | 0.000462 | 0.005563 | 0.004081 | 1731 | tags=33%, l | 5205/1824, | LNCaP-NCOR2 |
| HOLLEMAN | HOLLEMAN | HOLLEMAN | 14 | -0.730694 | -2.020586 | 0.000463 | 0.005563 | 0.004081 | 1425 | tags=86%, l | 26173/516 | LNCaP-NCOR2 |
| RUIZ_TNC_ | RUIZ_TNC_ | RUIZ_TNC_ | 107 | 0.431831 | 1.778167 | 0.000464 | 0.005563 | 0.004081 | 1906 | tags=30%, l | 1287/1078 | LNCaP-NCOR2 |
| LUI_THYRO | LUI_THYRO | LUI_THYRO | 20 | -0.689441 | -2.116414 | 0.000464 | 0.005563 | 0.004081 | 1425 | tags=55%, l | 5714/3312, | LNCaP-NCOR2 |
| DURAND_S | DURAND_S | DURAND_S | 99 | 0.424296 | 1.727362 | 0.000465 | 0.005563 | 0.004081 | 2311 | tags=35%, l | 5793/2309 | LNCaP-NCOR2 |
| REACTOME | REACTOME | REACTOME | 16 | -0.718875 | -2.068055 | 0.000467 | 0.005563 | 0.004081 | 547 | tags=25%, l | 2771/2787, | LNCaP-NCOR2 |
| MARTENS_ | MARTENS_ | MARTENS_ | 563 | -0.428235 | -2.302961 | 0.000469 | 0.005563 | 0.004081 | 2113 | tags=32%, l | 65003/262 | LNCaP-NCOR2 |
| KIM_ALL_D | KIM_ALL_D | KIM_ALL_D | 569 | -0.419096 | -2.255798 | 0.00047 | 0.005563 | 0.004081 | 2658 | tags=39%, l | 6223/1097 | LNCaP-NCOR2 |
| LINDGREN_ | LINDGREN_ | LINDGREN_ | 89 | 0.445761 | 1.783905 | 0.00047 | 0.005563 | 0.004081 | 3583 | tags=48%, l | 3709/5617, | LNCaP-NCOR2 |
| DIRMEIER_ | DIRMEIER_ | DIRMEIER_ | 23 | -0.659799 | -2.096627 | 0.000472 | 0.005563 | 0.004081 | 2901 | tags=74%, l | 6223/7454, | LNCaP-NCOR2 |
| LUI_TARGE | LUI_TARGE | LUI_TARGE | 24 | -0.644019 | -2.072396 | 0.000472 | 0.005563 | 0.004081 | 1425 | tags=46%, l | 6385/3312, | LNCaP-NCOR2 |
| GINESTIER_ | GINESTIER_ | GINESTIER_ | 71 | 0.490622 | 1.894298 | 0.000479 | 0.005609 | 0.004115 | 2894 | tags=51%, l | 25800/332 | LNCaP-NCOR2 |
| HALLMARK | HALLMARK | HALLMARK | 69 | 0.476455 | 1.830974 | 0.00048 | 0.005609 | 0.004115 | 2574 | tags=35%, l | 55008/770 | LNCaP-NCOR2 |
| REACTOME | REACTOME | REACTOME | 29 | -0.602043 | -2.035159 | 0.00048 | 0.005609 | 0.004115 | 1471 | tags=45%, l | 5984/5665, | LNCaP-NCOR2 |
| PUJANA_CI | PUJANA_CI | PUJANA_CI | 599 | -0.291004 | -1.571755 | 0.000484 | 0.005633 | 0.004132 | 1841 | tags=22%, l | 90410/112 | LNCaP-NCOR2 |
| IRITANI_M | IRITANI_M | IRITANI_M | 33 | -0.602036 | -2.0968 | 0.000486 | 0.005647 | 0.004143 | 1652 | tags=52%, l | 191/9361/ | LNCaP-NCOR2 |
| MODY_HIP | MODY_HIP | MODY_HIP | 36 | -0.600355 | -2.137374 | 0.00049 | 0.005658 | 0.00415 | 1472 | tags=53%, l | 1019/6164, | LNCaP-NCOR2 |
| KYNG_WER | KYNG_WER | KYNG_WER | 57 | 0.496415 | 1.838889 | 0.000491 | 0.005658 | 0.00415 | 1913 | tags=40%, l | 8795/8766, | LNCaP-NCOR2 |
| REACTOME | REACTOME | REACTOME | 613 | -0.319923 | -1.729495 | 0.000492 | 0.005658 | 0.00415 | 1531 | tags=23%, l | 389/6138/ | LNCaP-NCOR2 |
| GAVIN_FO | GAVIN_FO | GAVIN_FO | 50 | 0.521508 | 1.882382 | 0.000494 | 0.005658 | 0.00415 | 2608 | tags=52%, l | 169200/28 | LNCaP-NCOR2 |
| BENPORAT | BENPORAT | BENPORAT | 628 | -0.293335 | -1.591311 | 0.000495 | 0.005658 | 0.00415 | 2008 | tags=24%, l | 9218/7385, | LNCaP-NCOR2 |
| PID_AVB3_ | PID_AVB3_ | PID_AVB3_ | 48 | 0.508225 | 1.819093 | 0.000498 | 0.005658 | 0.00415 | 1097 | tags=25%, l | 3685/1287, | LNCaP-NCOR2 |
| TIEN_INTES | TIEN_INTES | TIEN_INTES | 40 | -0.567632 | -2.066652 | 0.000499 | 0.005658 | 0.00415 | 2121 | tags=62%, l | 6189/2512, | LNCaP-NCOR2 |
| REACTOME | REACTOME | REACTOME | 47 | -0.541961 | -2.048541 | 0.000499 | 0.005658 | 0.00415 | 1501 | tags=43%, l | 7520/8363, | LNCaP-NCOR2 |
| FRASOR_RI | FRASOR_RI | FRASOR_RI | 46 | -0.553815 | -2.078011 | 0.000502 | 0.005658 | 0.00415 | 2263 | tags=48%, l | 4798/3832, | LNCaP-NCOR2 |
| KANG_DOX | KANG_DOX | KANG_DOX | 46 | -0.571792 | -2.145465 | 0.000502 | 0.005658 | 0.00415 | 1661 | tags=39%, l | 29028/598 | LNCaP-NCOR2 |
| BILANGES_ | BILANGES_ | BILANGES_ | 48 | -0.515375 | -1.950818 | 0.000504 | 0.005658 | 0.00415 | 2701 | tags=46%, l | 645/6227/ | LNCaP-NCOR2 |
| PUJANA_BI | PUJANA_BI | PUJANA_BI | 48 | -0.518433 | -1.962395 | 0.000504 | 0.005658 | 0.00415 | 2544 | tags=42%, l | 9702/4173, | LNCaP-NCOR2 |
| SASAI_RESI | SASAI_RESI | SASAI_RESI | 34 | 0.615926 | 2.046798 | 0.000507 | 0.005676 | 0.004164 | 1593 | tags=35%, l | 3572/4982, | LNCaP-NCOR2 |
| CHIARADO | CHIARADO | CHIARADO | 30 | 0.585028 | 1.890314 | 0.000509 | 0.005694 | 0.004177 | 1341 | tags=40%, l | 4124/9180, | LNCaP-NCOR2 |
| REACTOME | REACTOME | REACTOME | 55 | -0.498924 | -1.945594 | 0.000512 | 0.005712 | 0.00419 | 2901 | tags=55%, l | 5719/5695, | LNCaP-NCOR2 |
| YAO_TEMP | YAO_TEMP | YAO_TEMP | 60 | -0.481142 | -1.908249 | 0.000518 | 0.005747 | 0.004216 | 1742 | tags=37%, l | 27316/191 | LNCaP-NCOR2 |
| GARGALOV | GARGALOV | GARGALOV | 27 | 0.619204 | 1.944963 | 0.000518 | 0.005747 | 0.004216 | 1904 | tags=48%, l | 4254/9314, | LNCaP-NCOR2 |
| LOPEZ_MB | LOPEZ_MB | LOPEZ_MB | 678 | -0.315173 | -1.718323 | 0.000521 | 0.005756 | 0.004223 | 2838 | tags=35%, l | 823/7737/ | LNCaP-NCOR2 |
| ROETH_TER | ROETH_TER | ROETH_TER | 12 | 0.744744 | 1.902033 | 0.000529 | 0.005831 | 0.004278 | 1689 | tags=58%, l | 55008/511 | LNCaP-NCOR2 |
| REACTOME | REACTOME | REACTOME | 69 | -0.493563 | -2.0068 | 0.000533 | 0.005832 | 0.004278 | 1812 | tags=39%, l | 641/7531/ | LNCaP-NCOR2 |
| RIZ_ERYTH | RIZ_ERYTH | RIZ_ERYTH | 69 | -0.470949 | -1.914852 | 0.000533 | 0.005832 | 0.004278 | 2934 | tags=45%, l | 5427/6827, | LNCaP-NCOR2 |
| REACTOME | REACTOME | REACTOME | 70 | -0.458715 | -1.87273 | 0.000534 | 0.005832 | 0.004278 | 2369 | tags=36%, l | 26224/690 | LNCaP-NCOR2 |
| WU_HBX_T | WU_HBX_T | WU_HBX_T | 10 | 0.802277 | 1.939722 | 0.000535 | 0.005832 | 0.004278 | 702 | tags=60%, l | 5784/1316, | LNCaP-NCOR2 |
| LI_DCP2_B | LI_DCP2_B | LI_DCP2_B | 77 | -0.463217 | -1.923912 | 0.000536 | 0.005832 | 0.004278 | 2969 | tags=56%, l | 57798/551 | LNCaP-NCOR2 |
| REACTOME | REACTOME | REACTOME | 82 | -0.451608 | -1.900441 | 0.000541 | 0.005875 | 0.00431 | 2901 | tags=49%, l | 5719/5695, | LNCaP-NCOR2 |
| NUYTTEN_I | NUYTTEN_I | NUYTTEN_I | 744 | -0.243564 | -1.336117 | 0.00055 | 0.005957 | 0.00437 | 2162 | tags=24%, l | 10244/572 | LNCaP-NCOR2 |
| BASAKI_YB | BASAKI_YB | BASAKI_YB | 276 | 0.339627 | 1.581696 | 0.00056 | 0.006047 | 0.004436 | 2055 | tags=28%, l | 3572/5152, | LNCaP-NCOR2 |
| ACEVEDO_ | ACEVEDO_ | ACEVEDO_ | 105 | -0.397003 | -1.739771 | 0.000561 | 0.006047 | 0.004436 | 855 | tags=22%, l | 8335/8348, | LNCaP-NCOR2 |
| KAUFFMAN | KAUFFMAN | KAUFFMAN | 118 | -0.395185 | -1.766537 | 0.000569 | 0.006082 | 0.004462 | 2509 | tags=38%, l | 1950/5763, | LNCaP-NCOR2 |
| BENPORAT | BENPORAT | BENPORAT | 120 | -0.398897 | -1.785992 | 0.000571 | 0.006082 | 0.004462 | 2550 | tags=36%, l | 7272/4605, | LNCaP-NCOR2 |
| GARCIA_TA | GARCIA_TA | GARCIA_TA | 121 | -0.412365 | -1.848376 | 0.000571 | 0.006082 | 0.004462 | 2239 | tags=36%, l | 5757/6502, | LNCaP-NCOR2 |
| YAO_TEMP | YAO_TEMP | YAO_TEMP | 120 | -0.393534 | -1.761983 | 0.000571 | 0.006082 | 0.004462 | 3545 | tags=51%, l | 5096/1891, | LNCaP-NCOR2 |
| FARMER_B | FARMER_B | FARMER_B | 235 | 0.355289 | 1.623644 | 0.000573 | 0.006082 | 0.004462 | 2753 | tags=37%, l | 4193/3572, | LNCaP-NCOR2 |
| KEGG_PATI | KEGG_PATI | KEGG_PATI | 233 | 0.350682 | 1.601373 | 0.000573 | 0.006082 | 0.004462 | 2242 | tags=29%, l | 4193/4254, | LNCaP-NCOR2 |
| PROVENZA | PROVENZA | PROVENZA | 137 | -0.378131 | -1.731636 | 0.000584 | 0.006169 | 0.004526 | 2125 | tags=30%, l | 23644/615 | LNCaP-NCOR2 |

| | | | | | | | | | | | | |
|---|---|---|---|---|---|---|---|---|---|---|---|---|
| REACTOME | REACTOME | REACTOME | 137 | -0.389644 | -1.784361 | 0.000584 | 0.006169 | 0.004526 | 3105 | tags=47%, | 64682/598 | LNCaP-NCOR2 |
| CHIANG_LI | CHIANG_LI | CHIANG_LI | 155 | -0.364406 | -1.693802 | 0.000607 | 0.006387 | 0.004685 | 1987 | tags=26%, | 10535/708 | LNCaP-NCOR2 |
| TIEN_INTES | TIEN_INTES | TIEN_INTES | 73 | 0.474662 | 1.842768 | 0.000637 | 0.006692 | 0.004909 | 2861 | tags=42%, | 9693/2353 | LNCaP-NCOR2 |
| INGRAM_S | INGRAM_S | INGRAM_S | 71 | 0.465539 | 1.797454 | 0.000639 | 0.006697 | 0.004913 | 1131 | tags=27%, | 3480/5617 | LNCaP-NCOR2 |
| KAUFFMAN | KAUFFMAN | KAUFFMAN | 204 | -0.334176 | -1.61838 | 0.000641 | 0.006699 | 0.004914 | 2601 | tags=36%, | 80198/537 | LNCaP-NCOR2 |
| WARTERS_ | WARTERS_ | WARTERS_ | 50 | 0.512366 | 1.849384 | 0.000658 | 0.006849 | 0.005024 | 2303 | tags=44%, | 4193/5048 | LNCaP-NCOR2 |
| HINATA_NI | HINATA_NI | HINATA_NI | 51 | 0.509211 | 1.844594 | 0.000659 | 0.006849 | 0.005024 | 1056 | tags=27%, | 6542/2581 | LNCaP-NCOR2 |
| REACTOME | REACTOME | REACTOME | 233 | -0.326513 | -1.608904 | 0.000661 | 0.006858 | 0.005031 | 2489 | tags=32%, | 5692/6118 | LNCaP-NCOR2 |
| MUELLER_I | MUELLER_I | MUELLER_I | 237 | -0.312752 | -1.542299 | 0.000664 | 0.006872 | 0.005041 | 2346 | tags=38%, | 890/22974 | LNCaP-NCOR2 |
| BURTON_A | BURTON_A | BURTON_A | 40 | 0.545081 | 1.881506 | 0.000668 | 0.006891 | 0.005055 | 3772 | tags=60%, | 10565/290 | LNCaP-NCOR2 |
| HESS_TARG | HESS_TARG | HESS_TARG | 38 | 0.556465 | 1.893608 | 0.000673 | 0.006928 | 0.005082 | 2728 | tags=50%, | 7107/9314 | LNCaP-NCOR2 |
| BOYLAN_M | BOYLAN_M | BOYLAN_M | 36 | 0.568658 | 1.914583 | 0.000675 | 0.006937 | 0.005089 | 1745 | tags=39%, | 65986/798 | LNCaP-NCOR2 |
| NIKOLSKY_ | NIKOLSKY_ | NIKOLSKY_ | 30 | 0.578879 | 1.870445 | 0.000679 | 0.006959 | 0.005105 | 2370 | tags=43%, | 4193/1171 | LNCaP-NCOR2 |
| HORTON_S | HORTON_S | HORTON_S | 22 | 0.666442 | 1.991478 | 0.00069 | 0.007053 | 0.005174 | 1800 | tags=55%, | 4199/1344 | LNCaP-NCOR2 |
| HAN_SATB | HAN_SATB | HAN_SATB | 293 | 0.328552 | 1.537868 | 0.000698 | 0.007115 | 0.005219 | 2052 | tags=28%, | 1287/6653 | LNCaP-NCOR2 |
| REACTOME | REACTOME | REACTOME | 272 | 0.327997 | 1.525885 | 0.0007 | 0.007115 | 0.005219 | 2294 | tags=25%, | 11059/368 | LNCaP-NCOR2 |
| WANG_TU | WANG_TU | WANG_TU | 286 | -0.309587 | -1.556835 | 0.000708 | 0.007183 | 0.005269 | 2540 | tags=31%, | 1733/5709 | LNCaP-NCOR2 |
| WEST_ADR | WEST_ADR | WEST_ADR | 310 | -0.293125 | -1.489427 | 0.00072 | 0.007289 | 0.005347 | 2848 | tags=33%, | 80210/218 | LNCaP-NCOR2 |
| SCHAEFFER | SCHAEFFER | SCHAEFFER | 181 | 0.361796 | 1.605427 | 0.000734 | 0.007397 | 0.005426 | 2231 | tags=27%, | 5865/6192 | LNCaP-NCOR2 |
| MULLIGHA | MULLIGHA | MULLIGHA | 176 | 0.362396 | 1.602952 | 0.000734 | 0.007397 | 0.005426 | 2145 | tags=28%, | 3685/5784 | LNCaP-NCOR2 |
| RAMALHO_ | RAMALHO_ | RAMALHO_ | 165 | 0.373394 | 1.637416 | 0.000741 | 0.007444 | 0.005461 | 3069 | tags=37%, | 64393/116 | LNCaP-NCOR2 |
| LINDGREN_ | LINDGREN_ | LINDGREN_ | 157 | 0.373739 | 1.629285 | 0.000744 | 0.007458 | 0.005471 | 2060 | tags=27%, | 1287/6653 | LNCaP-NCOR2 |
| HALLMARK | HALLMARK | HALLMARK | 104 | 0.422002 | 1.729953 | 0.000775 | 0.007751 | 0.005686 | 2596 | tags=40%, | 56172/931 | LNCaP-NCOR2 |
| RAO_BOUN | RAO_BOUN | RAO_BOUN | 84 | 0.441592 | 1.750187 | 0.000787 | 0.007834 | 0.005747 | 1931 | tags=33%, | 23362/951 | LNCaP-NCOR2 |
| SHAFFER_I | SHAFFER_I | SHAFFER_I | 84 | 0.438096 | 1.736332 | 0.000787 | 0.007834 | 0.005747 | 1600 | tags=31%, | 4124/3320 | LNCaP-NCOR2 |
| BRUECKNE | BRUECKNE | BRUECKNE | 80 | 0.452651 | 1.780474 | 0.000794 | 0.007884 | 0.005783 | 2085 | tags=38%, | 23092/553 | LNCaP-NCOR2 |
| VECCHI_GA | VECCHI_GA | VECCHI_GA | 70 | 0.456973 | 1.761063 | 0.000799 | 0.0079 | 0.005795 | 1567 | tags=29%, | 57674/129 | LNCaP-NCOR2 |
| EPPERT_HS | EPPERT_HS | EPPERT_HS | 67 | 0.466456 | 1.781362 | 0.000803 | 0.0079 | 0.005795 | 3220 | tags=54%, | 9314/3404 | LNCaP-NCOR2 |
| GARGALOV | GARGALOV | GARGALOV | 65 | 0.492486 | 1.870809 | 0.000803 | 0.0079 | 0.005795 | 3601 | tags=55%, | 3842/7964 | LNCaP-NCOR2 |
| KIM_MYC_ | KIM_MYC_ | KIM_MYC_ | 65 | 0.479338 | 1.820864 | 0.000803 | 0.0079 | 0.005795 | 1871 | tags=34%, | 5865/7999 | LNCaP-NCOR2 |
| LE_EGR2_T | LE_EGR2_T | LE_EGR2_T | 84 | -0.435636 | -1.839441 | 0.000822 | 0.008065 | 0.005916 | 1661 | tags=35%, | 79004/383 | LNCaP-NCOR2 |
| MILI_PSEU | MILI_PSEU | MILI_PSEU | 49 | 0.503938 | 1.81103 | 0.000825 | 0.008078 | 0.005926 | 2018 | tags=45%, | 6741/2302 | LNCaP-NCOR2 |
| GOBERT_C | GOBERT_C | GOBERT_C | 34 | 0.557269 | 1.851873 | 0.000844 | 0.008227 | 0.006035 | 4311 | tags=74%, | 7586/9053 | LNCaP-NCOR2 |
| REACTOME | REACTOME | REACTOME | 36 | 0.553334 | 1.862991 | 0.000844 | 0.008227 | 0.006035 | 2024 | tags=44%, | 8795/6406 | LNCaP-NCOR2 |
| STARK_HYF | STARK_HYF | STARK_HYF | 33 | 0.570928 | 1.882307 | 0.000849 | 0.008252 | 0.006053 | 2833 | tags=52%, | 23171/228 | LNCaP-NCOR2 |
| JISON_SICK | JISON_SICK | JISON_SICK | 120 | -0.368032 | -1.647801 | 0.000857 | 0.008308 | 0.006095 | 1946 | tags=35%, | 55658/620 | LNCaP-NCOR2 |
| HALLMARK | HALLMARK | HALLMARK | 132 | -0.360393 | -1.640545 | 0.00087 | 0.008414 | 0.006172 | 2494 | tags=36%, | 6118/2959 | LNCaP-NCOR2 |
| MIYAGAWA | MIYAGAWA | MIYAGAWA | 164 | 0.36699 | 1.608067 | 0.00089 | 0.00859 | 0.006301 | 2823 | tags=35%, | 85463/710 | LNCaP-NCOR2 |
| REACTOME | REACTOME | REACTOME | 531 | 0.289181 | 1.428732 | 0.000896 | 0.008634 | 0.006333 | 2522 | tags=27%, | 3572/4254 | LNCaP-NCOR2 |
| CHANDRAN | CHANDRAN | CHANDRAN | 141 | 0.384126 | 1.64912 | 0.000905 | 0.008699 | 0.006381 | 2520 | tags=40%, | 116064/21 | LNCaP-NCOR2 |
| VERHAAK_ | VERHAAK_ | VERHAAK_ | 126 | 0.390752 | 1.649287 | 0.000918 | 0.008805 | 0.006459 | 1413 | tags=20%, | 1824/2581 | LNCaP-NCOR2 |
| REACTOME | REACTOME | REACTOME | 13 | -0.745409 | -2.018361 | 0.000925 | 0.008853 | 0.006494 | 547 | tags=31%, | 2771/2787 | LNCaP-NCOR2 |
| WHITFIELD | WHITFIELD | WHITFIELD | 99 | 0.412382 | 1.678858 | 0.00093 | 0.008879 | 0.006513 | 2723 | tags=35%, | 57169/111 | LNCaP-NCOR2 |
| KEGG_CYT | KEGG_CYT | KEGG_CYT | 85 | 0.440085 | 1.747412 | 0.000944 | 0.008957 | 0.00657 | 2073 | tags=33%, | 3572/4254 | LNCaP-NCOR2 |
| ABE_INNEF | ABE_INNEF | ABE_INNEF | 24 | -0.611597 | -1.968067 | 0.000945 | 0.008957 | 0.00657 | 1523 | tags=46%, | 81876/299 | LNCaP-NCOR2 |
| REACTOME | REACTOME | REACTOME | 24 | -0.619219 | -1.992591 | 0.000945 | 0.008957 | 0.00657 | 3018 | tags=71%, | 54107/511 | LNCaP-NCOR2 |
| NADLER_O | NADLER_O | NADLER_O | 32 | -0.583315 | -2.016575 | 0.000973 | 0.009199 | 0.006748 | 2270 | tags=44%, | 1537/5834 | LNCaP-NCOR2 |
| SESTO_RES | SESTO_RES | SESTO_RES | 43 | 0.523495 | 1.838138 | 0.000995 | 0.009356 | 0.006863 | 1546 | tags=35%, | 1525/7586 | LNCaP-NCOR2 |
| KEGG_ARG | KEGG_ARG | KEGG_ARG | 40 | -0.526927 | -1.91845 | 0.000997 | 0.009356 | 0.006863 | 974 | tags=28%, | 8659/445/ | LNCaP-NCOR2 |
| MOLENAAF | MOLENAAF | MOLENAAF | 40 | -0.54078 | -1.968886 | 0.000997 | 0.009356 | 0.006863 | 2550 | tags=48%, | 64222/574 | LNCaP-NCOR2 |
| REACTOME | REACTOME | REACTOME | 47 | -0.507061 | -1.916625 | 0.000999 | 0.009356 | 0.006863 | 3298 | tags=49%, | 10469/267 | LNCaP-NCOR2 |
| DAZARD_U | DAZARD_U | DAZARD_U | 44 | -0.523286 | -1.943159 | 0.001001 | 0.009356 | 0.006863 | 2811 | tags=48%, | 3326/1350 | LNCaP-NCOR2 |
| PID_IL6_7_ | PID_IL6_7_ | PID_IL6_7_ | 36 | 0.535358 | 1.802469 | 0.001013 | 0.009453 | 0.006935 | 2522 | tags=47%, | 3572/6774 | LNCaP-NCOR2 |
| KRIEG_HYP | KRIEG_HYP | KRIEG_HYP | 541 | 0.285459 | 1.412251 | 0.001022 | 0.009517 | 0.006981 | 2001 | tags=23%, | 5865/5784 | LNCaP-NCOR2 |
| GENTILE_U | GENTILE_U | GENTILE_U | 29 | 0.570729 | 1.824807 | 0.001028 | 0.009524 | 0.006986 | 2868 | tags=48%, | 9693/1105 | LNCaP-NCOR2 |
| ZHAN_MUI | ZHAN_MUI | ZHAN_MUI | 29 | 0.571039 | 1.825797 | 0.001028 | 0.009524 | 0.006986 | 2280 | tags=38%, | 9693/2581 | LNCaP-NCOR2 |
| BIOCARTA_ | BIOCARTA_ | BIOCARTA_ | 23 | 0.636827 | 1.923609 | 0.001041 | 0.009631 | 0.007065 | 1121 | tags=22%, | 4193/5591 | LNCaP-NCOR2 |
| NEWMAN_ | NEWMAN_ | NEWMAN_ | 14 | 0.711976 | 1.893322 | 0.001056 | 0.009733 | 0.00714 | 2621 | tags=57%, | 5865/6439 | LNCaP-NCOR2 |
| OSMAN_BL | OSMAN_BL | OSMAN_BL | 293 | -0.292455 | -1.477535 | 0.001057 | 0.009733 | 0.00714 | 2088 | tags=26%, | 1606/6189 | LNCaP-NCOR2 |
| SPIELMAN_ | SPIELMAN_ | SPIELMAN_ | 11 | 0.746806 | 1.858168 | 0.001063 | 0.009766 | 0.007164 | 2796 | tags=82%, | 4124/4189 | LNCaP-NCOR2 |
| DAZARD_U | DAZARD_U | DAZARD_U | 111 | 0.402015 | 1.663611 | 0.001081 | 0.009913 | 0.007272 | 1613 | tags=26%, | 2309/9732 | LNCaP-NCOR2 |
| REACTOME | REACTOME | REACTOME | 747 | -0.234513 | -1.287657 | 0.001089 | 0.009958 | 0.007305 | 1669 | tags=20%, | 207/1211/ | LNCaP-NCOR2 |
| REACTOME | REACTOME | REACTOME | 90 | -0.426653 | -1.82375 | 0.001106 | 0.01009 | 0.007402 | 3480 | tags=51%, | 10573/781 | LNCaP-NCOR2 |
| KEGG_HUN | KEGG_HUN | KEGG_HUN | 133 | -0.351215 | -1.60039 | 0.001166 | 0.010611 | 0.007784 | 2141 | tags=35%, | 5435/7385 | LNCaP-NCOR2 |
| REACTOME | REACTOME | REACTOME | 18 | -0.66466 | -1.975955 | 0.001168 | 0.010611 | 0.007784 | 2481 | tags=83%, | 57804/511 | LNCaP-NCOR2 |
| RUTELLA_F | RUTELLA_F | RUTELLA_F | 168 | 0.363835 | 1.598188 | 0.001182 | 0.010713 | 0.007859 | 3567 | tags=45%, | 3572/6653 | LNCaP-NCOR2 |

| | | | | | | | | | | | |
|---|---|---|---|---|---|---|---|---|---|---|---|
| PID_MAPK_ | PID_MAPK_ | PID_MAPK_ | 30 | 0.566104 | 1.829165 | 0.001189 | 0.010755 | 0.007889 | 2971 | tags=50%, | 5906/5606, | LNCaP-NCOR2 |
| PUJANA_X | PUJANA_X | PUJANA_X | 145 | -0.348382 | -1.60473 | 0.001192 | 0.010757 | 0.007891 | 2544 | tags=37%, | 10403/715 | LNCaP-NCOR2 |
| BROWNE_ | BROWNE_ | BROWNE_ | 26 | 0.602003 | 1.872156 | 0.001206 | 0.010867 | 0.007972 | 1536 | tags=38%, | 3572/3480, | LNCaP-NCOR2 |
| REACTOME | REACTOME | REACTOME | 23 | 0.631613 | 1.907861 | 0.001215 | 0.010905 | 0.007999 | 1985 | tags=48%, | 3685/3915, | LNCaP-NCOR2 |
| WANG_RES | WANG_RES | WANG_RES | 21 | 0.651696 | 1.924002 | 0.001216 | 0.010905 | 0.007999 | 2080 | tags=48%, | 51465/905 | LNCaP-NCOR2 |
| HALLMARK | HALLMARK | HALLMARK | 104 | 0.405606 | 1.662741 | 0.00124 | 0.0111 | 0.008143 | 1929 | tags=32%, | 3915/3480, | LNCaP-NCOR2 |
| PLASARI_T | PLASARI_T | PLASARI_T | 70 | 0.451346 | 1.73938 | 0.001279 | 0.011423 | 0.00838 | 2694 | tags=49%, | 2803/1007 | LNCaP-NCOR2 |
| RUTELLA_R | RUTELLA_R | RUTELLA_R | 233 | 0.338891 | 1.547529 | 0.00129 | 0.011497 | 0.008434 | 3567 | tags=41%, | 3572/2336 | LNCaP-NCOR2 |
| GUO_HEX_ | GUO_HEX_ | GUO_HEX_ | 59 | 0.475806 | 1.77399 | 0.001302 | 0.011576 | 0.008492 | 2353 | tags=37%, | 59084/348 | LNCaP-NCOR2 |
| KEGG_TGF_ | KEGG_TGF_ | KEGG_TGF_ | 58 | 0.484581 | 1.801501 | 0.001304 | 0.011576 | 0.008492 | 1739 | tags=34%, | 657/93/10 | LNCaP-NCOR2 |
| ACEVEDO_ | ACEVEDO_ | ACEVEDO_ | 64 | -0.447239 | -1.794554 | 0.001317 | 0.011662 | 0.008555 | 296 | tags=27%, | 51203/897 | LNCaP-NCOR2 |
| MULLIGHA | MULLIGHA | MULLIGHA | 168 | 0.357038 | 1.568332 | 0.001329 | 0.011716 | 0.008595 | 2215 | tags=28%, | 11059/128 | LNCaP-NCOR2 |
| PID_TAP63 | PID_TAP63 | PID_TAP63 | 46 | 0.508085 | 1.805528 | 0.00133 | 0.011716 | 0.008595 | 1837 | tags=41%, | 4193/1105 | LNCaP-NCOR2 |
| SCHEIDERE | SCHEIDERE | SCHEIDERE | 45 | 0.514305 | 1.819121 | 0.001332 | 0.011716 | 0.008595 | 2381 | tags=47%, | 2309/3320, | LNCaP-NCOR2 |
| TAVAZOIE_ | TAVAZOIE_ | TAVAZOIE_ | 47 | 0.501988 | 1.786963 | 0.001334 | 0.011716 | 0.008595 | 2547 | tags=40%, | 4193/7414, | LNCaP-NCOR2 |
| REACTOME | REACTOME | REACTOME | 74 | -0.421651 | -1.737745 | 0.00134 | 0.011717 | 0.008595 | 2901 | tags=46%, | 5427/1045 | LNCaP-NCOR2 |
| BRUNO_HE | BRUNO_HE | BRUNO_HE | 39 | 0.524901 | 1.79926 | 0.001343 | 0.011717 | 0.008595 | 2537 | tags=44%, | 25800/300 | LNCaP-NCOR2 |
| REACTOME | REACTOME | REACTOME | 73 | -0.434326 | -1.783437 | 0.001343 | 0.011717 | 0.008595 | 2901 | tags=48%, | 5719/5695, | LNCaP-NCOR2 |
| KEGG_PAR | KEGG_PAR | KEGG_PAR | 79 | -0.425215 | -1.774951 | 0.001349 | 0.011749 | 0.008619 | 3233 | tags=53%, | 4696/2708 | LNCaP-NCOR2 |
| THUM_SYS | THUM_SYS | THUM_SYS | 133 | 0.377287 | 1.605272 | 0.00137 | 0.011904 | 0.008733 | 2690 | tags=39%, | 9693/2336 | LNCaP-NCOR2 |
| HOLLERN_ | HOLLERN_ | HOLLERN_ | 16 | 0.674244 | 1.858618 | 0.001398 | 0.012097 | 0.008874 | 1790 | tags=38%, | 8974/5510 | LNCaP-NCOR2 |
| KANG_DOX | KANG_DOX | KANG_DOX | 16 | 0.671555 | 1.851204 | 0.001398 | 0.012097 | 0.008874 | 2787 | tags=69%, | 11059/820 | LNCaP-NCOR2 |
| WHITFIELD | WHITFIELD | WHITFIELD | 97 | -0.391964 | -1.691939 | 0.001405 | 0.012137 | 0.008903 | 2604 | tags=39%, | 83879/839 | LNCaP-NCOR2 |
| REACTOME | REACTOME | REACTOME | 26 | -0.61108 | -2.011021 | 0.001429 | 0.012315 | 0.009034 | 3018 | tags=73%, | 57804/541 | LNCaP-NCOR2 |
| NIKOLSKY_ | NIKOLSKY_ | NIKOLSKY_ | 33 | -0.559238 | -1.947739 | 0.001459 | 0.01255 | 0.009206 | 1868 | tags=39%, | 353/2175/ | LNCaP-NCOR2 |
| GHANDHI_ | GHANDHI_ | GHANDHI_ | 42 | 0.508391 | 1.775393 | 0.001496 | 0.012838 | 0.009418 | 2193 | tags=52%, | 85463/573 | LNCaP-NCOR2 |
| HECKER_IF | HECKER_IF | HECKER_IF | 47 | 0.496649 | 1.767958 | 0.001501 | 0.012857 | 0.009431 | 1689 | tags=38%, | 55008/151 | LNCaP-NCOR2 |
| WU_APOP | WU_APOP | WU_APOP | 41 | -0.526474 | -1.919025 | 0.00151 | 0.012906 | 0.009467 | 1661 | tags=37%, | 7846/2902 | LNCaP-NCOR2 |
| ZHAN_MU | ZHAN_MU | ZHAN_MU | 35 | 0.543309 | 1.817402 | 0.00152 | 0.012967 | 0.009512 | 1622 | tags=31%, | 5865/1171 | LNCaP-NCOR2 |
| VERNELL_R | VERNELL_R | VERNELL_R | 58 | -0.46346 | -1.826014 | 0.001551 | 0.013202 | 0.009685 | 2125 | tags=50%, | 26271/100 | LNCaP-NCOR2 |
| ELVIDGE_H | ELVIDGE_H | ELVIDGE_H | 96 | 0.412324 | 1.66861 | 0.001558 | 0.013238 | 0.009711 | 1539 | tags=26%, | 1824/2581 | LNCaP-NCOR2 |
| BOWIE_RES | BOWIE_RES | BOWIE_RES | 13 | 0.714191 | 1.860463 | 0.001585 | 0.013435 | 0.009855 | 2772 | tags=62%, | 3433/4938, | LNCaP-NCOR2 |
| CHIARADO | CHIARADO | CHIARADO | 71 | 0.44251 | 1.708536 | 0.001598 | 0.013523 | 0.00992 | 2084 | tags=37%, | 4124/9180, | LNCaP-NCOR2 |
| REACTOME | REACTOME | REACTOME | 21 | -0.633214 | -1.96448 | 0.001649 | 0.013928 | 0.010217 | 3782 | tags=76%, | 54107/511 | LNCaP-NCOR2 |
| REACTOME | REACTOME | REACTOME | 24 | -0.605576 | -1.94869 | 0.001654 | 0.013936 | 0.010223 | 1443 | tags=71%, | 54107/511 | LNCaP-NCOR2 |
| REACTOME | REACTOME | REACTOME | 22 | -0.621333 | -1.951205 | 0.001664 | 0.013997 | 0.010268 | 3926 | tags=82%, | 57804/541 | LNCaP-NCOR2 |
| KEGG_HYP | KEGG_HYP | KEGG_HYP | 40 | 0.508116 | 1.75391 | 0.001669 | 0.01401 | 0.010278 | 2503 | tags=42%, | 3685/488/ | LNCaP-NCOR2 |
| REACTOME | REACTOME | REACTOME | 38 | 0.525355 | 1.787743 | 0.001682 | 0.014093 | 0.010338 | 1642 | tags=34%, | 5781/3433, | LNCaP-NCOR2 |
| DANG_MY | DANG_MY | DANG_MY | 104 | -0.373916 | -1.636035 | 0.00169 | 0.014126 | 0.010362 | 1797 | tags=29%, | 6207/8409, | LNCaP-NCOR2 |
| CAIRO_HE | CAIRO_HE | CAIRO_HE | 114 | 0.390574 | 1.62275 | 0.001696 | 0.014153 | 0.010382 | 1588 | tags=27%, | 3572/9455, | LNCaP-NCOR2 |
| PUJANA_BI | PUJANA_BI | PUJANA_BI | 106 | -0.372385 | -1.632444 | 0.0017 | 0.014153 | 0.010382 | 2231 | tags=41%, | 7153/6929, | LNCaP-NCOR2 |
| ACEVEDO_ | ACEVEDO_ | ACEVEDO_ | 260 | -0.287898 | -1.431935 | 0.001722 | 0.014309 | 0.010497 | 2849 | tags=33%, | 23636/569 | LNCaP-NCOR2 |
| BURTON_A | BURTON_A | BURTON_A | 22 | 0.635916 | 1.90026 | 0.001725 | 0.014309 | 0.010497 | 1669 | tags=45%, | 4254/3005, | LNCaP-NCOR2 |
| ODONNELL | ODONNELL | ODONNELL | 188 | 0.345786 | 1.542642 | 0.001753 | 0.014507 | 0.010642 | 2327 | tags=32%, | 64393/575 | LNCaP-NCOR2 |
| JEON_SMA | JEON_SMA | JEON_SMA | 12 | 0.727274 | 1.857416 | 0.001762 | 0.014559 | 0.01068 | 1537 | tags=50%, | 2200/355/ | LNCaP-NCOR2 |
| REACTOME | REACTOME | REACTOME | 10 | -0.777341 | -1.938536 | 0.001821 | 0.014991 | 0.010997 | 1991 | tags=60%, | 9391/5701 | LNCaP-NCOR2 |
| REACTOME | REACTOME | REACTOME | 43 | 0.501414 | 1.760607 | 0.001825 | 0.014991 | 0.010997 | 2123 | tags=33%, | 2650/5556 | LNCaP-NCOR2 |
| BROWNE_ | BROWNE_ | BROWNE_ | 42 | 0.499224 | 1.743382 | 0.001828 | 0.014991 | 0.010997 | 2896 | tags=48%, | 23269/573 | LNCaP-NCOR2 |
| STAMBOLS | STAMBOLS | STAMBOLS | 46 | 0.496894 | 1.765762 | 0.001829 | 0.014991 | 0.010997 | 2531 | tags=48%, | 5140/1836, | LNCaP-NCOR2 |
| REACTOME | REACTOME | REACTOME | 11 | -0.770659 | -1.981917 | 0.001836 | 0.015016 | 0.011015 | 547 | tags=55%, | 2786/2788, | LNCaP-NCOR2 |
| GOLUB_AL | GOLUB_AL | GOLUB_AL | 20 | -0.641916 | -1.970523 | 0.001858 | 0.015138 | 0.011105 | 770 | tags=25%, | 896/3925/ | LNCaP-NCOR2 |
| REACTOME | REACTOME | REACTOME | 20 | -0.650576 | -1.997109 | 0.001858 | 0.015138 | 0.011105 | 547 | tags=20%, | 2771/2787, | LNCaP-NCOR2 |
| GAZDA_DI | GAZDA_DI | GAZDA_DI | 22 | 0.62636 | 1.871706 | 0.001898 | 0.015434 | 0.011322 | 1754 | tags=41%, | 10565/879 | LNCaP-NCOR2 |
| LEE_BMP2_ | LEE_BMP2_ | LEE_BMP2_ | 662 | 0.268237 | 1.346586 | 0.00199 | 0.016152 | 0.011849 | 2639 | tags=27%, | 5793/2581 | LNCaP-NCOR2 |
| HAMAI_AP | HAMAI_AP | HAMAI_AP | 122 | -0.358266 | -1.607635 | 0.001997 | 0.016178 | 0.011868 | 2543 | tags=30%, | 51343/646 | LNCaP-NCOR2 |
| TANG_SEN | TANG_SEN | TANG_SEN | 46 | -0.493862 | -1.853059 | 0.002006 | 0.016219 | 0.011898 | 3562 | tags=67%, | 11065/490 | LNCaP-NCOR2 |
| RODRIGUE | RODRIGUE | RODRIGUE | 355 | 0.299477 | 1.430517 | 0.002036 | 0.016428 | 0.012051 | 3108 | tags=36%, | 5793/6653, | LNCaP-NCOR2 |
| PURBEY_T | PURBEY_T | PURBEY_T | 258 | -0.285327 | -1.418611 | 0.002075 | 0.016707 | 0.012255 | 1507 | tags=19%, | 23397/537 | LNCaP-NCOR2 |
| BROWNE_ | BROWNE_ | BROWNE_ | 70 | 0.436613 | 1.682601 | 0.002078 | 0.016707 | 0.012255 | 2587 | tags=39%, | 5737/4026, | LNCaP-NCOR2 |
| AKL_HTLV1 | AKL_HTLV1 | AKL_HTLV1 | 17 | -0.652966 | -1.910811 | 0.002104 | 0.016879 | 0.012382 | 2191 | tags=53%, | 5867/7266, | LNCaP-NCOR2 |
| LEE_EARLY | LEE_EARLY | LEE_EARLY | 70 | -0.42534 | -1.736474 | 0.002135 | 0.017097 | 0.012542 | 1711 | tags=36%, | 4173/6509, | LNCaP-NCOR2 |
| TONKS_TA | TONKS_TA | TONKS_TA | 112 | 0.389277 | 1.613224 | 0.002157 | 0.017203 | 0.012619 | 2940 | tags=38%, | 8507/2302 | LNCaP-NCOR2 |
| TSAI_RESP | TSAI_RESP | TSAI_RESP | 112 | 0.390195 | 1.617028 | 0.002157 | 0.017203 | 0.012619 | 1714 | tags=30%, | 25816/931 | LNCaP-NCOR2 |
| MULLIGHA | MULLIGHA | MULLIGHA | 87 | 0.414046 | 1.651188 | 0.002198 | 0.017435 | 0.01279 | 2305 | tags=31%, | 11059/128 | LNCaP-NCOR2 |
| LINDGREN_ | LINDGREN_ | LINDGREN_ | 196 | 0.338896 | 1.516748 | 0.002198 | 0.017435 | 0.01279 | 1863 | tags=24%, | 6542/1078 | LNCaP-NCOR2 |
| BURTON_A | BURTON_A | BURTON_A | 35 | -0.529104 | -1.872154 | 0.002205 | 0.017435 | 0.01279 | 1657 | tags=43%, | 4176/1059 | LNCaP-NCOR2 |

| | | | | | | | | | | | | |
|---|---|---|---|---|---|---|---|---|---|---|---|---|
| YU_MYC_T | YU_MYC_T | YU_MYC_T | 35 | -0.536731 | -1.89914 | 0.002205 | 0.017435 | 0.01279 | 1277 | tags=34%, | 3151/7973 | LNCaP-NCOR2 |
| AMIT_EGF_ | AMIT_EGF_ | AMIT_EGF_ | 33 | 0.538458 | 1.775256 | 0.002207 | 0.017435 | 0.01279 | 2041 | tags=42%, | 1316/7832 | LNCaP-NCOR2 |
| REACTOME | REACTOME | REACTOME | 477 | 0.284577 | 1.393933 | 0.002211 | 0.017435 | 0.01279 | 2294 | tags=24%, | 11059/368 | LNCaP-NCOR2 |
| RAO_BOUN | RAO_BOUN | RAO_BOUN | 308 | 0.304778 | 1.433661 | 0.002224 | 0.017491 | 0.012831 | 2776 | tags=31%, | 9695/9901 | LNCaP-NCOR2 |
| REACTOME | REACTOME | REACTOME | 39 | -0.514068 | -1.862051 | 0.002226 | 0.017491 | 0.012831 | 3018 | tags=54%, | 7316/5427 | LNCaP-NCOR2 |
| COULOUAF | COULOUAF | COULOUAF | 77 | 0.426578 | 1.669938 | 0.002233 | 0.017514 | 0.012848 | 3510 | tags=52%, | 65065/801 | LNCaP-NCOR2 |
| REACTOME | REACTOME | REACTOME | 42 | -0.506103 | -1.854936 | 0.002258 | 0.017679 | 0.012969 | 1130 | tags=31%, | 8367/8385 | LNCaP-NCOR2 |
| SAKAI_CHR | SAKAI_CHR | SAKAI_CHR | 62 | 0.460694 | 1.734808 | 0.002265 | 0.017697 | 0.012982 | 2248 | tags=39%, | 3842/3320 | LNCaP-NCOR2 |
| STARK_PRE | STARK_PRE | STARK_PRE | 137 | 0.368951 | 1.575896 | 0.00228 | 0.017778 | 0.013041 | 3069 | tags=46%, | 3480/1955 | LNCaP-NCOR2 |
| BOQUEST_ | BOQUEST_ | BOQUEST_ | 249 | 0.327017 | 1.505792 | 0.002406 | 0.01873 | 0.01374 | 2728 | tags=33%, | 65986/128 | LNCaP-NCOR2 |
| MORI_EMU | MORI_EMU | MORI_EMU | 79 | -0.409433 | -1.709073 | 0.002428 | 0.018868 | 0.013841 | 2060 | tags=33%, | 6598/5496 | LNCaP-NCOR2 |
| PEDRIOLI_ | PEDRIOLI_ | PEDRIOLI_ | 221 | 0.328048 | 1.491809 | 0.002438 | 0.018904 | 0.013867 | 1638 | tags=24%, | 51635/182 | LNCaP-NCOR2 |
| YANAGIHA | YANAGIHA | YANAGIHA | 20 | 0.630415 | 1.832693 | 0.002458 | 0.019026 | 0.013957 | 3191 | tags=70%, | 3572/8821 | LNCaP-NCOR2 |
| BILBAN_B_ | BILBAN_B_ | BILBAN_B_ | 40 | 0.50019 | 1.726552 | 0.002504 | 0.019345 | 0.014191 | 1906 | tags=40%, | 3321/7150 | LNCaP-NCOR2 |
| MOOTHA_ | MOOTHA_ | MOOTHA_ | 325 | -0.272157 | -1.389836 | 0.002552 | 0.019681 | 0.014437 | 1914 | tags=25%, | 10797/551 | LNCaP-NCOR2 |
| MOOTHA_ | MOOTHA_ | MOOTHA_ | 15 | -0.68595 | -1.938901 | 0.002563 | 0.019729 | 0.014473 | 2782 | tags=67%, | 4967/2271 | LNCaP-NCOR2 |
| SANSOM_ | SANSOM_ | SANSOM_ | 121 | -0.35786 | -1.604061 | 0.00257 | 0.019747 | 0.014486 | 2314 | tags=33%, | 2049/2199 | LNCaP-NCOR2 |
| BHAT_ESR | BHAT_ESR | BHAT_ESR | 63 | 0.449002 | 1.695907 | 0.002582 | 0.019802 | 0.014526 | 2142 | tags=35%, | 5784/2336 | LNCaP-NCOR2 |
| BRACHAT_ | BRACHAT_ | BRACHAT_ | 17 | 0.651933 | 1.823699 | 0.002621 | 0.020062 | 0.014717 | 1492 | tags=47%, | 4193/677/ | LNCaP-NCOR2 |
| BROWNE_ | BROWNE_ | BROWNE_ | 46 | 0.484062 | 1.720163 | 0.00266 | 0.020293 | 0.014887 | 1689 | tags=37%, | 3433/4938 | LNCaP-NCOR2 |
| RODRIGUE | RODRIGUE | RODRIGUE | 46 | 0.482267 | 1.713782 | 0.00266 | 0.020293 | 0.014887 | 2896 | tags=50%, | 57674/289 | LNCaP-NCOR2 |
| TONKS_TAI | TONKS_TAI | TONKS_TAI | 30 | -0.547869 | -1.86616 | 0.002674 | 0.020363 | 0.014938 | 390 | tags=10%, | 4071/2770 | LNCaP-NCOR2 |
| SHAFFER_I | SHAFFER_I | SHAFFER_I | 67 | 0.434758 | 1.660309 | 0.00273 | 0.020714 | 0.015195 | 3674 | tags=51%, | 1595/6482 | LNCaP-NCOR2 |
| PEREZ_TP5 | PEREZ_TP5 | PEREZ_TP5 | 671 | 0.26277 | 1.320726 | 0.002731 | 0.020714 | 0.015195 | 2956 | tags=28%, | 4193/5716 | LNCaP-NCOR2 |
| BANDRES_ | BANDRES_ | BANDRES_ | 86 | -0.39548 | -1.677277 | 0.002741 | 0.020758 | 0.015227 | 1277 | tags=35%, | 2886/2512 | LNCaP-NCOR2 |
| BRUINS_U | BRUINS_U | BRUINS_U | 132 | 0.368768 | 1.567001 | 0.002747 | 0.020764 | 0.015232 | 2754 | tags=35%, | 3709/9180 | LNCaP-NCOR2 |
| BEGUM_TA | BEGUM_TA | BEGUM_TA | 28 | 0.560114 | 1.776036 | 0.002752 | 0.020764 | 0.015232 | 2663 | tags=50%, | 5793/4254 | LNCaP-NCOR2 |
| IVANOVA_ | IVANOVA_ | IVANOVA_ | 332 | 0.299619 | 1.419818 | 0.002757 | 0.020768 | 0.015235 | 2209 | tags=25%, | 65986/619 | LNCaP-NCOR2 |
| BOWIE_RE | BOWIE_RE | BOWIE_RE | 13 | 0.695946 | 1.812933 | 0.002817 | 0.021177 | 0.015535 | 2772 | tags=62%, | 3433/4938 | LNCaP-NCOR2 |
| ZHANG_RE | ZHANG_RE | ZHANG_RE | 11 | 0.728225 | 1.811937 | 0.002835 | 0.021274 | 0.015606 | 2301 | tags=82%, | 7832/5366 | LNCaP-NCOR2 |
| REACTOME | REACTOME | REACTOME | 59 | -0.442613 | -1.748295 | 0.002853 | 0.021357 | 0.015667 | 1341 | tags=22%, | 816/84254 | LNCaP-NCOR2 |
| MARSON_ | MARSON_ | MARSON_ | 654 | 0.264891 | 1.329505 | 0.002856 | 0.021357 | 0.015667 | 3083 | tags=30%, | 4193/8795 | LNCaP-NCOR2 |
| KEGG_JAK_ | KEGG_JAK_ | KEGG_JAK_ | 68 | 0.431477 | 1.65277 | 0.002885 | 0.021532 | 0.015795 | 3430 | tags=46%, | 3572/9180 | LNCaP-NCOR2 |
| LI_WILMS_ | LI_WILMS_ | LI_WILMS_ | 135 | -0.337024 | -1.536333 | 0.002939 | 0.021902 | 0.016066 | 2226 | tags=29%, | 440/9493/ | LNCaP-NCOR2 |
| GAVIN_FO | GAVIN_FO | GAVIN_FO | 55 | 0.473037 | 1.740188 | 0.002951 | 0.021951 | 0.016103 | 1995 | tags=36%, | 8507/2353 | LNCaP-NCOR2 |
| HERNANDE | HERNANDE | HERNANDE | 41 | 0.489852 | 1.701555 | 0.002986 | 0.022152 | 0.01625 | 2064 | tags=44%, | 10788/348 | LNCaP-NCOR2 |
| BUYTAERT_ | BUYTAERT_ | BUYTAERT_ | 465 | -0.251892 | -1.334944 | 0.002989 | 0.022152 | 0.01625 | 2924 | tags=31%, | 8214/4521 | LNCaP-NCOR2 |
| BILD_SRC_ | BILD_SRC_ | BILD_SRC_ | 45 | 0.485971 | 1.718899 | 0.002998 | 0.02218 | 0.01627 | 3136 | tags=53%, | 6653/4012 | LNCaP-NCOR2 |
| REACTOME | REACTOME | REACTOME | 18 | -0.633202 | -1.882435 | 0.003036 | 0.022421 | 0.016447 | 547 | tags=22%, | 2771/2787 | LNCaP-NCOR2 |
| MOREIRA_ | MOREIRA_ | MOREIRA_ | 19 | -0.625838 | -1.888545 | 0.00305 | 0.022487 | 0.016495 | 3800 | tags=84%, | 3150/2026 | LNCaP-NCOR2 |
| BURTON_A | BURTON_A | BURTON_A | 22 | 0.61687 | 1.843348 | 0.003106 | 0.022855 | 0.016765 | 2897 | tags=59%, | 7414/3157 | LNCaP-NCOR2 |
| DURAND_S | DURAND_S | DURAND_S | 163 | 0.352797 | 1.544268 | 0.00312 | 0.0229 | 0.016799 | 1812 | tags=24%, | 4254/5163 | LNCaP-NCOR2 |
| REACTOME | REACTOME | REACTOME | 29 | -0.550923 | -1.862352 | 0.003123 | 0.0229 | 0.016799 | 2481 | tags=55%, | 4173/5982 | LNCaP-NCOR2 |
| TANAKA_M | TANAKA_M | TANAKA_M | 46 | 0.480014 | 1.705777 | 0.003159 | 0.023103 | 0.016948 | 1716 | tags=33%, | 2074/9749 | LNCaP-NCOR2 |
| HALLMARK | HALLMARK | HALLMARK | 32 | -0.533896 | -1.84573 | 0.003161 | 0.023103 | 0.016948 | 1090 | tags=16%, | 10014/686 | LNCaP-NCOR2 |
| REACTOME | REACTOME | REACTOME | 14 | 0.678918 | 1.805415 | 0.003168 | 0.023114 | 0.016955 | 1432 | tags=43%, | 2309/9314 | LNCaP-NCOR2 |
| BROWNE_ | BROWNE_ | BROWNE_ | 76 | 0.420743 | 1.642763 | 0.003196 | 0.023277 | 0.017075 | 1913 | tags=38%, | 4254/7046 | LNCaP-NCOR2 |
| BENPORAT | BENPORAT | BENPORAT | 404 | -0.256322 | -1.338959 | 0.003232 | 0.023498 | 0.017237 | 2446 | tags=26%, | 29767/501 | LNCaP-NCOR2 |
| PASQUALU | PASQUALU | PASQUALU | 102 | 0.392987 | 1.609078 | 0.00324 | 0.023511 | 0.017247 | 1724 | tags=27%, | 1769/2581 | LNCaP-NCOR2 |
| QI_HYPOXI | QI_HYPOXI | QI_HYPOXI | 96 | 0.396199 | 1.603353 | 0.003272 | 0.023704 | 0.017388 | 2255 | tags=31%, | 2309/8974 | LNCaP-NCOR2 |
| REACTOME | REACTOME | REACTOME | 52 | 0.470996 | 1.713007 | 0.003294 | 0.02382 | 0.017474 | 3008 | tags=60%, | 51360/159 | LNCaP-NCOR2 |
| GAL_LEUKE | GAL_LEUKE | GAL_LEUKE | 85 | 0.410081 | 1.628278 | 0.003302 | 0.02384 | 0.017488 | 2246 | tags=29%, | 2309/3480 | LNCaP-NCOR2 |
| DURCHDEV | DURCHDEV | DURCHDEV | 52 | -0.456608 | -1.75882 | 0.003308 | 0.02384 | 0.017488 | 927 | tags=25%, | 3028/3742 | LNCaP-NCOR2 |
| AFFAR_YY1 | AFFAR_YY1 | AFFAR_YY1 | 164 | -0.322078 | -1.50921 | 0.003375 | 0.024283 | 0.017813 | 2346 | tags=32%, | 6950/5110 | LNCaP-NCOR2 |
| KIM_WT1_ | KIM_WT1_ | KIM_WT1_ | 122 | 0.377516 | 1.585316 | 0.003386 | 0.02432 | 0.01784 | 2928 | tags=37%, | 3572/1105 | LNCaP-NCOR2 |
| HU_ANGIO | HU_ANGIO | HU_ANGIO | 31 | -0.536879 | -1.842866 | 0.0034 | 0.024375 | 0.017881 | 2005 | tags=39%, | 6418/5689 | LNCaP-NCOR2 |
| SWEET_KR | SWEET_KR | SWEET_KR | 54 | 0.465425 | 1.705949 | 0.003451 | 0.024701 | 0.01812 | 1705 | tags=24%, | 3685/8974 | LNCaP-NCOR2 |
| WHITFIELD | WHITFIELD | WHITFIELD | 40 | -0.494524 | -1.800476 | 0.00349 | 0.02494 | 0.018295 | 1690 | tags=30%, | 699/9134/ | LNCaP-NCOR2 |
| REACTOME | REACTOME | REACTOME | 17 | -0.64569 | -1.889519 | 0.003506 | 0.025011 | 0.018347 | 547 | tags=41%, | 152/2786/ | LNCaP-NCOR2 |
| BIOCARTA_ | BIOCARTA_ | BIOCARTA_ | 13 | 0.688172 | 1.792682 | 0.003521 | 0.025074 | 0.018393 | 1387 | tags=46%, | 2309/3480 | LNCaP-NCOR2 |
| IWANAGA_ | IWANAGA_ | IWANAGA_ | 100 | 0.394484 | 1.609111 | 0.003564 | 0.025338 | 0.018587 | 3028 | tags=40%, | 1769/9069 | LNCaP-NCOR2 |
| SENESE_HD | SENESE_HD | SENESE_HD | 60 | 0.451363 | 1.687559 | 0.003582 | 0.02542 | 0.018647 | 1970 | tags=32%, | 6653/4747 | LNCaP-NCOR2 |
| BROWNE_ | BROWNE_ | BROWNE_ | 88 | 0.399783 | 1.596665 | 0.003619 | 0.025617 | 0.018792 | 2822 | tags=41%, | 3915/2899 | LNCaP-NCOR2 |
| MARTENS_ | MARTENS_ | MARTENS_ | 232 | -0.292815 | -1.442599 | 0.003622 | 0.025617 | 0.018792 | 2849 | tags=33%, | 1969/2728 | LNCaP-NCOR2 |
| REACTOME | REACTOME | REACTOME | 72 | 0.418644 | 1.620548 | 0.003671 | 0.025879 | 0.018984 | 1967 | tags=33%, | 9069/667/ | LNCaP-NCOR2 |
| REACTOME | REACTOME | REACTOME | 11 | -0.733279 | -1.885788 | 0.003671 | 0.025879 | 0.018984 | 1560 | tags=73%, | 4176/8429 | LNCaP-NCOR2 |

| | | | | | | | | | | | | |
|---|---|---|---|---|---|---|---|---|---|---|---|---|
| REACTOME | REACTOME | REACTOME | 37 | -0.506922 | -1.819183 | 0.003678 | 0.025884 | 0.018988 | 2481 | tags=49%, | 5427/6118, | LNCaP-NCOR2 |
| PRAMOON | PRAMOON | PRAMOON | 38 | -0.494091 | -1.782184 | 0.003696 | 0.025924 | 0.019017 | 2208 | tags=42%, | 3916/5101 | LNCaP-NCOR2 |
| ZHAN_MUI | ZHAN_MUI | ZHAN_MUI | 38 | -0.502885 | -1.813903 | 0.003696 | 0.025924 | 0.019017 | 3151 | tags=45%, | 2305/7153, | LNCaP-NCOR2 |
| GRESHOCK | GRESHOCK | GRESHOCK | 233 | 0.318878 | 1.456138 | 0.003727 | 0.026092 | 0.01914 | 2719 | tags=32%, | 23092/576 | LNCaP-NCOR2 |
| ENK_UV_R | ENK_UV_R | ENK_UV_R | 386 | 0.288454 | 1.389185 | 0.00374 | 0.026141 | 0.019176 | 2100 | tags=27%, | 3685/8795, | LNCaP-NCOR2 |
| HOEBEKE_I | HOEBEKE_I | HOEBEKE_I | 56 | 0.458776 | 1.693769 | 0.003764 | 0.026269 | 0.01927 | 2854 | tags=52%, | 23022/545, | LNCaP-NCOR2 |
| REACTOME | REACTOME | REACTOME | 53 | -0.444333 | -1.719068 | 0.003823 | 0.026546 | 0.019474 | 2901 | tags=49%, | 5719/5695, | LNCaP-NCOR2 |
| NIKOLSKY_ | NIKOLSKY_ | NIKOLSKY_ | 77 | 0.410877 | 1.608475 | 0.003828 | 0.026546 | 0.019474 | 2193 | tags=34%, | 65986/619 | LNCaP-NCOR2 |
| MANN_RES | MANN_RES | MANN_RES | 19 | 0.629353 | 1.81339 | 0.003833 | 0.026546 | 0.019474 | 573 | tags=26%, | 4193/1371, | LNCaP-NCOR2 |
| SANSOM_A | SANSOM_A | SANSOM_A | 84 | -0.388144 | -1.638909 | 0.003836 | 0.026546 | 0.019474 | 2638 | tags=38%, | 57590/838 | LNCaP-NCOR2 |
| GINESTIER_ | GINESTIER_ | GINESTIER_ | 47 | 0.469339 | 1.670742 | 0.003836 | 0.026546 | 0.019474 | 2577 | tags=47%, | 56987/402 | LNCaP-NCOR2 |
| LIU_PROST | LIU_PROST | LIU_PROST | 233 | 0.318448 | 1.454177 | 0.00387 | 0.026672 | 0.019566 | 2084 | tags=23%, | 4254/1577, | LNCaP-NCOR2 |
| REACTOME | REACTOME | REACTOME | 14 | 0.673213 | 1.790243 | 0.003873 | 0.026672 | 0.019566 | 1178 | tags=43%, | 3572/9180, | LNCaP-NCOR2 |
| REACTOME | REACTOME | REACTOME | 13 | 0.682926 | 1.779017 | 0.003873 | 0.026672 | 0.019566 | 2215 | tags=62%, | 7706/6413 | LNCaP-NCOR2 |
| REACTOME | REACTOME | REACTOME | 34 | 0.523552 | 1.739829 | 0.003883 | 0.026697 | 0.019584 | 1178 | tags=32%, | 4254/6774, | LNCaP-NCOR2 |
| FORTSCHE | FORTSCHE | FORTSCHE | 190 | 0.337754 | 1.508498 | 0.003944 | 0.02707 | 0.019858 | 2000 | tags=26%, | 11059/348 | LNCaP-NCOR2 |
| MITSIADES | MITSIADES | MITSIADES | 318 | 0.298956 | 1.411502 | 0.004004 | 0.027436 | 0.020126 | 3329 | tags=36%, | 9693/5140, | LNCaP-NCOR2 |
| SAGIV_CD2 | SAGIV_CD2 | SAGIV_CD2 | 15 | 0.663869 | 1.79918 | 0.004028 | 0.027521 | 0.020188 | 1729 | tags=40%, | 10565/285 | LNCaP-NCOR2 |
| MULLIGHA | MULLIGHA | MULLIGHA | 213 | 0.325088 | 1.471743 | 0.004029 | 0.027521 | 0.020188 | 2215 | tags=26%, | 11059/128 | LNCaP-NCOR2 |
| KOKKINAKI | KOKKINAKI | KOKKINAKI | 91 | 0.390928 | 1.570292 | 0.004064 | 0.027711 | 0.020328 | 1871 | tags=32%, | 3572/8795, | LNCaP-NCOR2 |
| PID_PI3KCI | PID_PI3KCI | PID_PI3KCI | 31 | 0.53834 | 1.751839 | 0.004079 | 0.027768 | 0.02037 | 1167 | tags=32%, | 5591/2309, | LNCaP-NCOR2 |
| BRUINS_U | BRUINS_U | BRUINS_U | 424 | 0.281405 | 1.366129 | 0.004098 | 0.027803 | 0.020396 | 2926 | tags=31%, | 116068/97 | LNCaP-NCOR2 |
| FLECHNER_ | FLECHNER_ | FLECHNER_ | 411 | 0.28217 | 1.36662 | 0.00411 | 0.027803 | 0.020396 | 1900 | tags=22%, | 11059/36/ | LNCaP-NCOR2 |
| GOTTWEIN | GOTTWEIN | GOTTWEIN | 51 | 0.46864 | 1.697628 | 0.004117 | 0.027803 | 0.020396 | 2481 | tags=39%, | 57674/553 | LNCaP-NCOR2 |
| PEDERSEN_ | PEDERSEN_ | PEDERSEN_ | 51 | 0.467795 | 1.694565 | 0.004117 | 0.027803 | 0.020396 | 1016 | tags=25%, | 5865/1824, | LNCaP-NCOR2 |
| AMIT_SERU | AMIT_SERU | AMIT_SERU | 52 | 0.463759 | 1.686684 | 0.004117 | 0.027803 | 0.020396 | 3564 | tags=46%, | 5205/9314, | LNCaP-NCOR2 |
| REACTOME | REACTOME | REACTOME | 320 | 0.298712 | 1.411494 | 0.004139 | 0.027902 | 0.020468 | 2835 | tags=31%, | 11059/425 | LNCaP-NCOR2 |
| IGARASHI_ | IGARASHI_ | IGARASHI_ | 76 | 0.412792 | 1.611717 | 0.004155 | 0.02797 | 0.020518 | 1522 | tags=26%, | 6653/9314, | LNCaP-NCOR2 |
| IVANOVA_I | IVANOVA_I | IVANOVA_I | 98 | 0.38784 | 1.575626 | 0.004188 | 0.028144 | 0.020646 | 2640 | tags=36%, | 85463/107 | LNCaP-NCOR2 |
| YAMASHIT | YAMASHIT | YAMASHIT | 39 | -0.488122 | -1.76807 | 0.004205 | 0.028212 | 0.020695 | 2088 | tags=49%, | 6189/1122 | LNCaP-NCOR2 |
| MILI_PSEU | MILI_PSEU | MILI_PSEU | 30 | 0.541253 | 1.748869 | 0.004245 | 0.028437 | 0.020861 | 2293 | tags=50%, | 6741/2302 | LNCaP-NCOR2 |
| YOSHIMUR | YOSHIMUR | YOSHIMUR | 243 | 0.319033 | 1.465655 | 0.004255 | 0.028456 | 0.020874 | 2508 | tags=30%, | 3572/1160 | LNCaP-NCOR2 |
| LEE_CALOR | LEE_CALOR | LEE_CALOR | 22 | -0.587767 | -1.845795 | 0.00428 | 0.028551 | 0.020944 | 2838 | tags=50%, | 5720/7086, | LNCaP-NCOR2 |
| MATTIOLI_ | MATTIOLI_ | MATTIOLI_ | 72 | -0.403884 | -1.655664 | 0.004283 | 0.028551 | 0.020944 | 1167 | tags=42%, | 23636/570 | LNCaP-NCOR2 |
| ASGHARZA | ASGHARZA | ASGHARZA | 28 | -0.548419 | -1.8393 | 0.004298 | 0.028608 | 0.020986 | 799 | tags=21%, | 55339/942 | LNCaP-NCOR2 |
| VALK_AML | VALK_AML | VALK_AML | 19 | 0.623226 | 1.795735 | 0.004355 | 0.028944 | 0.021232 | 2504 | tags=47%, | 3572/1287, | LNCaP-NCOR2 |
| WANG_RES | WANG_RES | WANG_RES | 16 | -0.651818 | -1.875146 | 0.00444 | 0.029461 | 0.021612 | 501 | tags=19%, | 3006/2770 | LNCaP-NCOR2 |
| REN_ALVE | REN_ALVE | REN_ALVE | 316 | 0.297513 | 1.403466 | 0.004563 | 0.030228 | 0.022174 | 2153 | tags=26%, | 3685/3915, | LNCaP-NCOR2 |
| REACTOME | REACTOME | REACTOME | 82 | 0.40393 | 1.594083 | 0.004599 | 0.030324 | 0.022244 | 2505 | tags=35%, | 1795/5906, | LNCaP-NCOR2 |
| HOOI_ST7_ | HOOI_ST7_ | HOOI_ST7_ | 53 | 0.458988 | 1.674995 | 0.004607 | 0.030324 | 0.022244 | 1967 | tags=42%, | 3572/2309 | LNCaP-NCOR2 |
| PID_FAK_P | PID_FAK_P | PID_FAK_P | 53 | 0.460778 | 1.68153 | 0.004607 | 0.030324 | 0.022244 | 2504 | tags=36%, | 3685/2309 | LNCaP-NCOR2 |
| REACTOME | REACTOME | REACTOME | 53 | 0.458608 | 1.67361 | 0.004607 | 0.030324 | 0.022244 | 1170 | tags=30%, | 7706/5119 | LNCaP-NCOR2 |
| NADERI_BF | NADERI_BF | NADERI_BF | 33 | -0.517254 | -1.801517 | 0.004621 | 0.030324 | 0.022244 | 2940 | tags=48%, | 55839/104 | LNCaP-NCOR2 |
| MUELLER_ | MUELLER_ | MUELLER_ | 12 | -0.707718 | -1.86916 | 0.004621 | 0.030324 | 0.022244 | 3264 | tags=83%, | 3728/6929, | LNCaP-NCOR2 |
| BROWNE_I | BROWNE_I | BROWNE_I | 76 | 0.409475 | 1.598766 | 0.004635 | 0.030366 | 0.022276 | 2769 | tags=38%, | 9848/9334, | LNCaP-NCOR2 |
| HALLMARK | HALLMARK | HALLMARK | 99 | 0.387345 | 1.576927 | 0.004652 | 0.030386 | 0.02229 | 1816 | tags=26%, | 6542/9180, | LNCaP-NCOR2 |
| HILLION_H | HILLION_H | HILLION_H | 58 | -0.434047 | -1.710129 | 0.004652 | 0.030386 | 0.02229 | 1436 | tags=26%, | 7415/381/ | LNCaP-NCOR2 |
| DAIRKEE_C | DAIRKEE_C | DAIRKEE_C | 86 | -0.383983 | -1.628518 | 0.00466 | 0.030389 | 0.022292 | 1775 | tags=33%, | 468/2597/ | LNCaP-NCOR2 |
| RUTELLA_R | RUTELLA_R | RUTELLA_R | 246 | 0.315346 | 1.450288 | 0.00468 | 0.030472 | 0.022353 | 3490 | tags=39%, | 4199/1078 | LNCaP-NCOR2 |
| HALLMARK | HALLMARK | HALLMARK | 162 | 0.343055 | 1.501642 | 0.004748 | 0.030838 | 0.022622 | 2548 | tags=32%, | 4193/1105 | LNCaP-NCOR2 |
| KEGG_MISI | KEGG_MISI | KEGG_MISI | 22 | -0.584601 | -1.835853 | 0.004755 | 0.030838 | 0.022622 | 3782 | tags=73%, | 4436/5111, | LNCaP-NCOR2 |
| BOYAULT_I | BOYAULT_I | BOYAULT_I | 12 | 0.703578 | 1.796896 | 0.004759 | 0.030838 | 0.022622 | 2165 | tags=67%, | 7052/6282, | LNCaP-NCOR2 |
| BERENJEN | BERENJEN | BERENJEN | 403 | -0.252503 | -1.319046 | 0.004839 | 0.031309 | 0.022968 | 1700 | tags=19%, | 1869/699/ | LNCaP-NCOR2 |
| FURUKAW | FURUKAW | FURUKAW | 56 | -0.437338 | -1.713968 | 0.004882 | 0.031539 | 0.023136 | 2149 | tags=36%, | 64236/383 | LNCaP-NCOR2 |
| SHARMA_F | SHARMA_F | SHARMA_F | 15 | 0.656342 | 1.77878 | 0.004904 | 0.031609 | 0.023187 | 2042 | tags=47%, | 5915/840/ | LNCaP-NCOR2 |
| NELSON_RI | NELSON_RI | NELSON_RI | 16 | -0.648353 | -1.865179 | 0.004908 | 0.031609 | 0.023187 | 1865 | tags=56%, | 590/6385/ | LNCaP-NCOR2 |
| TAVOR_CEI | TAVOR_CEI | TAVOR_CEI | 33 | 0.523656 | 1.726453 | 0.004924 | 0.031663 | 0.023227 | 1346 | tags=36%, | 7414/5774, | LNCaP-NCOR2 |
| SERVITJA_I | SERVITJA_I | SERVITJA_I | 73 | 0.412369 | 1.600931 | 0.004938 | 0.031706 | 0.023258 | 1994 | tags=29%, | 4747/9515, | LNCaP-NCOR2 |
| PID_INTEGI | PID_INTEGI | PID_INTEGI | 22 | 0.59736 | 1.785047 | 0.005003 | 0.032077 | 0.023531 | 866 | tags=27%, | 3685/3915, | LNCaP-NCOR2 |
| PID_S1P_S | PID_S1P_S | PID_S1P_S | 11 | -0.715987 | -1.841319 | 0.005048 | 0.032216 | 0.023633 | 302 | tags=18%, | 2770 | LNCaP-NCOR2 |
| REACTOME | REACTOME | REACTOME | 11 | -0.717924 | -1.8463 | 0.005048 | 0.032216 | 0.023633 | 1783 | tags=36%, | 2786/2788, | LNCaP-NCOR2 |
| RODRIGUE | RODRIGUE | RODRIGUE | 11 | -0.714876 | -1.83846 | 0.005048 | 0.032216 | 0.023633 | 542 | tags=36%, | 6228/5582 | LNCaP-NCOR2 |
| TOYOTA_T | TOYOTA_T | TOYOTA_T | 314 | -0.264784 | -1.347422 | 0.005076 | 0.032345 | 0.023727 | 2550 | tags=28%, | 4605/6470 | LNCaP-NCOR2 |
| LINDGREN_ | LINDGREN_ | LINDGREN_ | 97 | 0.38849 | 1.576176 | 0.005121 | 0.032581 | 0.023901 | 2806 | tags=38%, | 10788/230 | LNCaP-NCOR2 |
| REACTOME | REACTOME | REACTOME | 34 | -0.504165 | -1.768251 | 0.005148 | 0.032696 | 0.023984 | 1783 | tags=26%, | 23416/278 | LNCaP-NCOR2 |
| SEITZ_NEO | SEITZ_NEO | SEITZ_NEO | 46 | 0.468039 | 1.663224 | 0.005155 | 0.032696 | 0.023984 | 2080 | tags=39%, | 6653/5500 | LNCaP-NCOR2 |

| | | | | | | | | | | | | |
|---|---|---|---|---|---|---|---|---|---|---|---|---|
| MALONEY_ | MALONEY_ | MALONEY_ | 60 | -0.418984 | -1.661722 | 0.005181 | 0.032766 | 0.024036 | 2952 | tags=47%, | 1365/7114, | LNCaP-NCOR2 |
| NAKAYAMA | NAKAYAMA | NAKAYAMA | 60 | -0.418698 | -1.660588 | 0.005181 | 0.032766 | 0.024036 | 3562 | tags=52%, | 2026/1058, | LNCaP-NCOR2 |
| JOHNSTON | JOHNSTON | JOHNSTON | 38 | 0.495645 | 1.68664 | 0.005215 | 0.032931 | 0.024157 | 2834 | tags=53%, | 9749/4012, | LNCaP-NCOR2 |
| HALMOS_C | HALMOS_C | HALMOS_C | 36 | 0.506318 | 1.704696 | 0.005235 | 0.033003 | 0.02421 | 1761 | tags=36%, | 25816/195 | LNCaP-NCOR2 |
| DAZARD_R | DAZARD_R | DAZARD_R | 93 | 0.385537 | 1.552812 | 0.005303 | 0.033386 | 0.024491 | 3294 | tags=39%, | 9693/2581 | LNCaP-NCOR2 |
| SANA_RESI | SANA_RESI | SANA_RESI | 39 | 0.49272 | 1.688951 | 0.00537 | 0.033754 | 0.024761 | 2975 | tags=49%, | 57674/151 | LNCaP-NCOR2 |
| STEIN_ESRI | STEIN_ESRI | STEIN_ESRI | 35 | -0.504932 | -1.786626 | 0.005391 | 0.033834 | 0.02482 | 2125 | tags=46%, | 54892/553 | LNCaP-NCOR2 |
| LINDSTEDT | LINDSTEDT | LINDSTEDT | 37 | 0.504248 | 1.704036 | 0.005402 | 0.033852 | 0.024833 | 1311 | tags=32%, | 10957/309 | LNCaP-NCOR2 |
| REACTOME | REACTOME | REACTOME | 61 | -0.416912 | -1.660287 | 0.005466 | 0.034203 | 0.02509 | 1342 | tags=39%, | 2967/5435, | LNCaP-NCOR2 |
| REACTOME | REACTOME | REACTOME | 43 | -0.468762 | -1.726059 | 0.005537 | 0.03455 | 0.025344 | 1471 | tags=40%, | 5984/9100, | LNCaP-NCOR2 |
| CHANG_IM | CHANG_IM | CHANG_IM | 39 | 0.491797 | 1.685787 | 0.005538 | 0.03455 | 0.025344 | 1479 | tags=28%, | 9069/324/ | LNCaP-NCOR2 |
| GROSS_HYI | GROSS_HYI | GROSS_HYI | 110 | 0.376451 | 1.556715 | 0.005555 | 0.034603 | 0.025384 | 2584 | tags=35%, | 9314/2302 | LNCaP-NCOR2 |
| BIOCARTA_ | BIOCARTA_ | BIOCARTA_ | 17 | 0.633185 | 1.771255 | 0.00559 | 0.034774 | 0.025509 | 1714 | tags=41%, | 4193/7046, | LNCaP-NCOR2 |
| LEE_LIVER_ | LEE_LIVER_ | LEE_LIVER_ | 36 | -0.494087 | -1.759039 | 0.005637 | 0.035013 | 0.025684 | 1711 | tags=39%, | 353/25873, | LNCaP-NCOR2 |
| REACTOME | REACTOME | REACTOME | 240 | 0.311348 | 1.427177 | 0.005699 | 0.035343 | 0.025926 | 1904 | tags=22%, | 11059/576 | LNCaP-NCOR2 |
| LASTOWSK | LASTOWSK | LASTOWSK | 26 | -0.540716 | -1.779459 | 0.005716 | 0.035383 | 0.025956 | 2470 | tags=62%, | 26160/101 | LNCaP-NCOR2 |
| BROWNE_I | BROWNE_I | BROWNE_I | 57 | 0.44095 | 1.633426 | 0.005727 | 0.035383 | 0.025956 | 2908 | tags=39%, | 9532/1710 | LNCaP-NCOR2 |
| ZHAN_MUI | ZHAN_MUI | ZHAN_MUI | 28 | -0.538457 | -1.805889 | 0.005731 | 0.035383 | 0.025956 | 2218 | tags=43%, | 10036/590 | LNCaP-NCOR2 |
| SCIBETTA_I | SCIBETTA_I | SCIBETTA_I | 15 | -0.649257 | -1.835185 | 0.005825 | 0.035867 | 0.026311 | 2679 | tags=40%, | 8635/5449 | LNCaP-NCOR2 |
| MARTINEZ_ | MARTINEZ_ | MARTINEZ_ | 419 | 0.277946 | 1.348752 | 0.005826 | 0.035867 | 0.026311 | 2040 | tags=25%, | 5793/3572, | LNCaP-NCOR2 |
| KYNG_DNA | KYNG_DNA | KYNG_DNA | 40 | 0.484233 | 1.671471 | 0.005842 | 0.035912 | 0.026344 | 3263 | tags=42%, | 8795/6774, | LNCaP-NCOR2 |
| MENSE_HY | MENSE_HY | MENSE_HY | 74 | 0.402692 | 1.566805 | 0.005899 | 0.03621 | 0.026562 | 3686 | tags=49%, | 8974/9314, | LNCaP-NCOR2 |
| REACTOME | REACTOME | REACTOME | 96 | 0.382331 | 1.547233 | 0.005921 | 0.036289 | 0.026621 | 2897 | tags=44%, | 134429/51 | LNCaP-NCOR2 |
| MORI_EMU | MORI_EMU | MORI_EMU | 15 | 0.646134 | 1.751113 | 0.005954 | 0.036442 | 0.026732 | 921 | tags=40%, | 3572/3309, | LNCaP-NCOR2 |
| SCHMIDT_I | SCHMIDT_I | SCHMIDT_I | 20 | 0.609085 | 1.770683 | 0.005969 | 0.036478 | 0.026759 | 2410 | tags=60%, | 134429/15 | LNCaP-NCOR2 |
| GAL_LEUKE | GAL_LEUKE | GAL_LEUKE | 117 | -0.343439 | -1.531218 | 0.006 | 0.036539 | 0.026804 | 1769 | tags=27%, | 1869/699/ | LNCaP-NCOR2 |
| HALLMARK | HALLMARK | HALLMARK | 47 | 0.461786 | 1.643856 | 0.006004 | 0.036539 | 0.026804 | 2156 | tags=47%, | 28996/657, | LNCaP-NCOR2 |
| NUTT_GBM | NUTT_GBM | NUTT_GBM | 29 | -0.532244 | -1.799207 | 0.006005 | 0.036539 | 0.026804 | 2920 | tags=48%, | 7381/6224, | LNCaP-NCOR2 |
| RUTELLA_R | RUTELLA_R | RUTELLA_R | 181 | 0.332463 | 1.475263 | 0.00602 | 0.036573 | 0.026829 | 3490 | tags=41%, | 6653/1078 | LNCaP-NCOR2 |
| GAUSSMAN | GAUSSMAN | GAUSSMAN | 101 | 0.380175 | 1.55277 | 0.006042 | 0.036654 | 0.026888 | 2828 | tags=39%, | 6542/9180, | LNCaP-NCOR2 |
| LU_IL4_SIG | LU_IL4_SIG | LU_IL4_SIG | 51 | 0.455868 | 1.651363 | 0.006093 | 0.036908 | 0.027075 | 1268 | tags=29%, | 51105/945 | LNCaP-NCOR2 |
| FEVR_CTNI | FEVR_CTNI | FEVR_CTNI | 401 | 0.279743 | 1.35199 | 0.006119 | 0.037018 | 0.027155 | 2622 | tags=29%, | 5152/4199, | LNCaP-NCOR2 |
| REACTOME | REACTOME | REACTOME | 10 | -0.72401 | -1.805539 | 0.006146 | 0.037126 | 0.027234 | 1114 | tags=80%, | 5111/6118, | LNCaP-NCOR2 |
| REACTOME | REACTOME | REACTOME | 27 | -0.537909 | -1.788813 | 0.00617 | 0.037215 | 0.0273 | 1410 | tags=41%, | 4001/6175, | LNCaP-NCOR2 |
| REACTOME | REACTOME | REACTOME | 99 | -0.351402 | -1.521684 | 0.006192 | 0.037294 | 0.027358 | 2645 | tags=41%, | 5427/1045 | LNCaP-NCOR2 |
| SHAFFER_II | SHAFFER_II | SHAFFER_II | 56 | 0.44319 | 1.636224 | 0.006219 | 0.037405 | 0.027439 | 3674 | tags=50%, | 4124/5146 | LNCaP-NCOR2 |
| FUJII_YBX1 | FUJII_YBX1 | FUJII_YBX1 | 21 | 0.596215 | 1.760204 | 0.006252 | 0.037549 | 0.027545 | 2372 | tags=57%, | 1730/355/ | LNCaP-NCOR2 |
| RICKMAN_ | RICKMAN_ | RICKMAN_ | 242 | 0.309271 | 1.418139 | 0.006279 | 0.037659 | 0.027625 | 3415 | tags=42%, | 51635/897 | LNCaP-NCOR2 |
| REACTOME | REACTOME | REACTOME | 11 | 0.704632 | 1.753233 | 0.006378 | 0.038198 | 0.028021 | 1150 | tags=45%, | 7706/6413 | LNCaP-NCOR2 |
| BHAT_ESR1 | BHAT_ESR1 | BHAT_ESR1 | 68 | 0.418868 | 1.604473 | 0.00641 | 0.038333 | 0.02812 | 2558 | tags=37%, | 23362/758 | LNCaP-NCOR2 |
| JAZAG_TGF | JAZAG_TGF | JAZAG_TGF | 59 | -0.416765 | -1.646198 | 0.006483 | 0.038715 | 0.0284 | 2796 | tags=44%, | 7703/2279 | LNCaP-NCOR2 |
| REACTOME | REACTOME | REACTOME | 29 | 0.536909 | 1.716674 | 0.006508 | 0.038806 | 0.028467 | 1675 | tags=34%, | 4193/1030, | LNCaP-NCOR2 |
| MEISSNER_ | MEISSNER_ | MEISSNER_ | 68 | 0.417304 | 1.598484 | 0.006571 | 0.039124 | 0.0287 | 1715 | tags=29%, | 4747/5670 | LNCaP-NCOR2 |
| KEGG_BUT | KEGG_BUT | KEGG_BUT | 25 | -0.545483 | -1.776031 | 0.006641 | 0.039433 | 0.028927 | 2756 | tags=52%, | 3155/1162 | LNCaP-NCOR2 |
| TIEN_INTES | TIEN_INTES | TIEN_INTES | 25 | -0.544612 | -1.773195 | 0.006641 | 0.039433 | 0.028927 | 1236 | tags=40%, | 6194/2709 | LNCaP-NCOR2 |
| AMUNDSO | AMUNDSO | AMUNDSO | 165 | 0.337702 | 1.480897 | 0.006668 | 0.039533 | 0.029 | 2554 | tags=32%, | 3572/8795, | LNCaP-NCOR2 |
| REACTOME | REACTOME | REACTOME | 75 | -0.394353 | -1.628982 | 0.006688 | 0.039598 | 0.029048 | 2645 | tags=41%, | 5887/5081 | LNCaP-NCOR2 |
| MARKEY_R | MARKEY_R | MARKEY_R | 119 | 0.3659 | 1.530074 | 0.006769 | 0.039989 | 0.029334 | 2038 | tags=29%, | 57169/516 | LNCaP-NCOR2 |
| KEGG_HEM | KEGG_HEM | KEGG_HEM | 21 | 0.592421 | 1.749003 | 0.006773 | 0.039989 | 0.029334 | 2503 | tags=43%, | 4254/3684, | LNCaP-NCOR2 |
| MARTINEZ_ | MARTINEZ_ | MARTINEZ_ | 358 | 0.284885 | 1.361824 | 0.006785 | 0.040003 | 0.029345 | 2155 | tags=25%, | 5793/3572, | LNCaP-NCOR2 |
| BIOCARTA_ | BIOCARTA_ | BIOCARTA_ | 21 | 0.591426 | 1.746066 | 0.006947 | 0.040898 | 0.030002 | 1714 | tags=48%, | 657/324/7 | LNCaP-NCOR2 |
| NATSUME_ | NATSUME_ | NATSUME_ | 46 | 0.459537 | 1.63301 | 0.006984 | 0.041058 | 0.030118 | 2762 | tags=39%, | 667/840/7 | LNCaP-NCOR2 |
| BIOCARTA_ | BIOCARTA_ | BIOCARTA_ | 77 | 0.396538 | 1.552339 | 0.007019 | 0.041205 | 0.030227 | 2971 | tags=39%, | 9693/7046, | LNCaP-NCOR2 |
| ZWANG_EC | ZWANG_EC | ZWANG_EC | 22 | 0.582299 | 1.740041 | 0.007074 | 0.041471 | 0.030422 | 1102 | tags=36%, | 4199/8426 | LNCaP-NCOR2 |
| ZHU_CMV_ | ZHU_CMV_ | ZHU_CMV_ | 36 | 0.495954 | 1.669802 | 0.007092 | 0.04152 | 0.030458 | 2587 | tags=36%, | 3572/3480, | LNCaP-NCOR2 |
| SERVITJA_L | SERVITJA_L | SERVITJA_L | 88 | -0.36893 | -1.572818 | 0.007131 | 0.04169 | 0.030582 | 2366 | tags=35%, | 291/55143, | LNCaP-NCOR2 |
| IVANOVA_I | IVANOVA_I | IVANOVA_I | 328 | 0.28902 | 1.368541 | 0.007169 | 0.041855 | 0.030704 | 2027 | tags=23%, | 79993/516 | LNCaP-NCOR2 |
| KEGG_SMA | KEGG_SMA | KEGG_SMA | 70 | 0.412289 | 1.588861 | 0.007194 | 0.041924 | 0.030754 | 1985 | tags=33%, | 3685/3915, | LNCaP-NCOR2 |
| WINZEN_D | WINZEN_D | WINZEN_D | 56 | 0.437223 | 1.614196 | 0.007201 | 0.041924 | 0.030754 | 1952 | tags=34%, | 9069/5737, | LNCaP-NCOR2 |
| REACTOME | REACTOME | REACTOME | 25 | 0.55824 | 1.719259 | 0.007259 | 0.042201 | 0.030957 | 1459 | tags=36%, | 2650/5556 | LNCaP-NCOR2 |
| HU_GENOT | HU_GENOT | HU_GENOT | 24 | -0.553348 | -1.780626 | 0.007323 | 0.042359 | 0.031073 | 2314 | tags=46%, | 10797/468, | LNCaP-NCOR2 |
| LY_AGING_ | LY_AGING_ | LY_AGING_ | 24 | -0.552978 | -1.779434 | 0.007323 | 0.042359 | 0.031073 | 1277 | tags=38%, | 9088/580/ | LNCaP-NCOR2 |
| REACTOME | REACTOME | REACTOME | 24 | -0.553095 | -1.779811 | 0.007323 | 0.042359 | 0.031073 | 3718 | tags=71%, | 2805/230/ | LNCaP-NCOR2 |
| FIGUEROA_ | FIGUEROA_ | FIGUEROA_ | 60 | 0.429736 | 1.606699 | 0.007327 | 0.042359 | 0.031073 | 3044 | tags=38%, | 6653/9053, | LNCaP-NCOR2 |
| MATHEW_ | MATHEW_ | MATHEW_ | 11 | -0.69908 | -1.797837 | 0.007343 | 0.042394 | 0.031099 | 1592 | tags=82%, | 2175/2176, | LNCaP-NCOR2 |
| PEDERSEN_ | PEDERSEN_ | PEDERSEN_ | 12 | 0.684248 | 1.74753 | 0.007402 | 0.042636 | 0.031276 | 2439 | tags=67%, | 25816/850 | LNCaP-NCOR2 |

| | | | | | | | | | | | | |
|---|---|---|---|---|---|---|---|---|---|---|---|---|
| ACEVEDO_ | ACEVEDO_ | ACEVEDO_ | 102 | 0.377533 | 1.545803 | 0.007405 | 0.042636 | 0.031276 | 2231 | tags=34%, | 23269/562 | LNCaP-NCOR2 |
| HOLLERN_ | HOLLERN_ | HOLLERN_ | 55 | -0.419515 | -1.635931 | 0.00743 | 0.042663 | 0.031296 | 346 | tags=11%, | 30061/555 | LNCaP-NCOR2 |
| DELASERN | DELASERN | DELASERN | 35 | 0.50007 | 1.672764 | 0.007431 | 0.042663 | 0.031296 | 1753 | tags=34%, | 3709/1344 | LNCaP-NCOR2 |
| CAIRO_HEI | CAIRO_HEI | CAIRO_HEI | 157 | 0.338387 | 1.475167 | 0.00744 | 0.042663 | 0.031296 | 1800 | tags=22%, | 3915/1672 | LNCaP-NCOR2 |
| UDAYAKUN | UDAYAKUN | UDAYAKUN | 186 | 0.327931 | 1.460637 | 0.007471 | 0.042782 | 0.031383 | 1047 | tags=18%, | 4193/8974 | LNCaP-NCOR2 |
| CHIBA_RES | CHIBA_RES | CHIBA_RES | 17 | 0.621419 | 1.73834 | 0.007512 | 0.042956 | 0.031511 | 2307 | tags=47%, | 4193/3685 | LNCaP-NCOR2 |
| KEGG_RIG_ | KEGG_RIG_ | KEGG_RIG_ | 46 | 0.457436 | 1.625543 | 0.007649 | 0.043618 | 0.031997 | 3024 | tags=43%, | 7706/6413 | LNCaP-NCOR2 |
| REACTOME | REACTOME | REACTOME | 46 | 0.457479 | 1.625696 | 0.007649 | 0.043618 | 0.031997 | 1985 | tags=33%, | 3685/1287 | LNCaP-NCOR2 |
| NAKAYAMA | NAKAYAMA | NAKAYAMA | 32 | 0.51501 | 1.687772 | 0.00781 | 0.044423 | 0.032587 | 1020 | tags=28%, | 4193/6439 | LNCaP-NCOR2 |
| PID_PI3KCI | PID_PI3KCI | PID_PI3KCI | 30 | 0.524954 | 1.696205 | 0.007811 | 0.044423 | 0.032587 | 2529 | tags=50%, | 2309/5933 | LNCaP-NCOR2 |
| PECE_MAN | PECE_MAN | PECE_MAN | 90 | 0.380882 | 1.526803 | 0.007832 | 0.044481 | 0.03263 | 3069 | tags=41%, | 4254/5617 | LNCaP-NCOR2 |
| REACTOME | REACTOME | REACTOME | 86 | 0.382539 | 1.521687 | 0.007869 | 0.044629 | 0.032739 | 2156 | tags=34%, | 657/93/25 | LNCaP-NCOR2 |
| BORCZUK_ | BORCZUK_ | BORCZUK_ | 259 | 0.305205 | 1.411808 | 0.00788 | 0.044629 | 0.032739 | 1594 | tags=20%, | 3685/3915 | LNCaP-NCOR2 |
| NIKOLSKY_ | NIKOLSKY_ | NIKOLSKY_ | 37 | 0.488083 | 1.649407 | 0.007934 | 0.044823 | 0.032881 | 1622 | tags=30%, | 9069/1577 | LNCaP-NCOR2 |
| FRASOR_T | FRASOR_T | FRASOR_T | 34 | 0.501806 | 1.667564 | 0.007935 | 0.044823 | 0.032881 | 2073 | tags=35%, | 5793/7534 | LNCaP-NCOR2 |
| OKUMURA | OKUMURA | OKUMURA | 127 | 0.353567 | 1.49461 | 0.007947 | 0.044832 | 0.032887 | 2471 | tags=33%, | 4199/1013 | LNCaP-NCOR2 |
| GYORFFY_ | GYORFFY_ | GYORFFY_ | 25 | 0.556015 | 1.712408 | 0.008123 | 0.045761 | 0.033569 | 3377 | tags=52%, | 9749/8023 | LNCaP-NCOR2 |
| KORKOLA_ | KORKOLA_ | KORKOLA_ | 16 | -0.621967 | -1.789269 | 0.008179 | 0.046016 | 0.033756 | 2490 | tags=56%, | 51202/259 | LNCaP-NCOR2 |
| ZHANG_IN | ZHANG_IN | ZHANG_IN | 17 | 0.620407 | 1.735508 | 0.008211 | 0.046132 | 0.033841 | 2772 | tags=59%, | 3433/4938 | LNCaP-NCOR2 |
| KEGG_BAS | KEGG_BAS | KEGG_BAS | 33 | -0.494982 | -1.723946 | 0.008268 | 0.046392 | 0.034032 | 2289 | tags=39%, | 143/4350/ | LNCaP-NCOR2 |
| KEGG_UBI | KEGG_UBI | KEGG_UBI | 112 | 0.366895 | 1.52047 | 0.008318 | 0.046607 | 0.03419 | 2260 | tags=27%, | 4193/1105 | LNCaP-NCOR2 |
| REACTOME | REACTOME | REACTOME | 13 | -0.66193 | -1.792322 | 0.008329 | 0.046609 | 0.034191 | 2263 | tags=92%, | 57804/511 | LNCaP-NCOR2 |
| MCBRYAN_ | MCBRYAN_ | MCBRYAN_ | 134 | 0.347956 | 1.482066 | 0.008371 | 0.046781 | 0.034317 | 1995 | tags=31%, | 3572/2899 | LNCaP-NCOR2 |
| SCHLOSSER | SCHLOSSER | SCHLOSSER | 81 | -0.379897 | -1.592972 | 0.008403 | 0.046897 | 0.034402 | 1191 | tags=37%, | 6050/1209 | LNCaP-NCOR2 |
| BONOME_ | BONOME_ | BONOME_ | 22 | 0.571607 | 1.70809 | 0.008454 | 0.047086 | 0.034541 | 997 | tags=27%, | 23362/892 | LNCaP-NCOR2 |
| GRANDVAU | GRANDVAU | GRANDVAU | 12 | 0.674796 | 1.723389 | 0.00846 | 0.047086 | 0.034541 | 2772 | tags=83%, | 9334/2908 | LNCaP-NCOR2 |
| HALLMARK | HALLMARK | HALLMARK | 112 | 0.36656 | 1.519082 | 0.008472 | 0.047092 | 0.034545 | 1921 | tags=31%, | 56172/300 | LNCaP-NCOR2 |
| WHITFIELD | WHITFIELD | WHITFIELD | 115 | -0.33507 | -1.489191 | 0.008549 | 0.047459 | 0.034814 | 1764 | tags=32%, | 641/15757 | LNCaP-NCOR2 |
| GARGALOV | GARGALOV | GARGALOV | 12 | 0.674314 | 1.722159 | 0.008636 | 0.047875 | 0.03512 | 1663 | tags=42%, | 91947/964 | LNCaP-NCOR2 |
| WANG_RES | WANG_RES | WANG_RES | 211 | 0.315147 | 1.423765 | 0.008671 | 0.048004 | 0.035214 | 2904 | tags=32%, | 64393/665 | LNCaP-NCOR2 |
| SMID_BRE | SMID_BRE | SMID_BRE | 83 | -0.370975 | -1.563537 | 0.008703 | 0.048118 | 0.035298 | 1177 | tags=22%, | 23464/991 | LNCaP-NCOR2 |
| WENG_POI | WENG_POI | WENG_POI | 10 | 0.712518 | 1.722705 | 0.008736 | 0.048238 | 0.035386 | 2204 | tags=60%, | 1595/3157 | LNCaP-NCOR2 |
| HALLMARK | HALLMARK | HALLMARK | 183 | 0.324747 | 1.443297 | 0.008789 | 0.048465 | 0.035552 | 1774 | tags=23%, | 10565/973 | LNCaP-NCOR2 |
| KAMMING | KAMMING | KAMMING | 39 | -0.457876 | -1.658514 | 0.008904 | 0.048982 | 0.035932 | 3018 | tags=49%, | 5427/7153 | LNCaP-NCOR2 |
| HILLION_H | HILLION_H | HILLION_H | 52 | -0.422232 | -1.626405 | 0.008906 | 0.048982 | 0.035932 | 1868 | tags=31%, | 207/5694/ | LNCaP-NCOR2 |
| MEISSNER_ | MEISSNER_ | MEISSNER_ | 23 | -0.557263 | -1.770802 | 0.00896 | 0.049155 | 0.036059 | 1238 | tags=26%, | 3229/5930 | LNCaP-NCOR2 |
| GENTILE_U | GENTILE_U | GENTILE_U | 26 | 0.542771 | 1.687951 | 0.008961 | 0.049155 | 0.036059 | 3778 | tags=58%, | 9444/7357 | LNCaP-NCOR2 |
| BILANGES_ | BILANGES_ | BILANGES_ | 27 | -0.517403 | -1.720621 | 0.009018 | 0.049402 | 0.036239 | 1554 | tags=41%, | 6164/1059 | LNCaP-NCOR2 |
| SIMBULAN | SIMBULAN | SIMBULAN | 15 | -0.634236 | -1.792725 | 0.009087 | 0.049715 | 0.036469 | 1079 | tags=73%, | 5111/871/ | LNCaP-NCOR2 |

LNCaP_DHT

| | ID | Descriptior | setSize | enrichmen | NES | pvalue | p.adjust | qvalues | rank | leading_ed | core_enricl | Group |
|---|---|---|---|---|---|---|---|---|---|---|---|---|
| GEORGES_ | GEORGES_ | GEORGES_ | 670 | 0.454252 | 1.492523 | 0.0001 | 0.017103 | 0.014729 | 2117 | tags=26%, | 23052/260 | LNCaP+DHT |
| DACOSTA_ | DACOSTA_ | DACOSTA_ | 666 | 0.579595 | 1.904311 | 0.0001 | 0.017103 | 0.014729 | 1745 | tags=35%, | 26018/129 | LNCaP+DHT |
| BUYTAERT_ | BUYTAERT_ | BUYTAERT_ | 627 | 0.466449 | 1.531754 | 0.0001 | 0.017103 | 0.014729 | 1945 | tags=32%, | 3480/2597 | LNCaP+DHT |
| KOINUMA_ | KOINUMA_ | KOINUMA_ | 604 | 0.474635 | 1.558133 | 0.0001 | 0.017103 | 0.014729 | 1535 | tags=24%, | 129642/22 | LNCaP+DHT |
| CUI_TCF21_ | CUI_TCF21_ | CUI_TCF21_ | 538 | 0.546752 | 1.792197 | 0.0001 | 0.017103 | 0.014729 | 1307 | tags=27%, | 23052/397 | LNCaP+DHT |
| ONKEN_UV | ONKEN_UV | ONKEN_UV | 563 | 0.454064 | 1.488647 | 0.0001 | 0.017103 | 0.014729 | 1798 | tags=25%, | 65986/716 | LNCaP+DHT |
| RODRIGUES | RODRIGUES | RODRIGUES | 541 | 0.492872 | 1.615471 | 0.0001 | 0.017103 | 0.014729 | 1538 | tags=22%, | 2289/2601 | LNCaP+DHT |
| KRIGE_RES | KRIGE_RES | KRIGE_RES | 506 | 0.472508 | 1.547602 | 0.0001 | 0.017103 | 0.014729 | 1765 | tags=28%, | 54206/110 | LNCaP+DHT |
| CREIGHTON | CREIGHTON | CREIGHTON | 486 | 0.516281 | 1.689508 | 0.0001 | 0.017103 | 0.014729 | 1420 | tags=25%, | 7113/2181 | LNCaP+DHT |
| HAMAI_AP | HAMAI_AP | HAMAI_AP | 489 | 0.454951 | 1.488879 | 0.0001 | 0.017103 | 0.014729 | 2302 | tags=30%, | 7163/1005 | LNCaP+DHT |
| WONG_AD | WONG_AD | WONG_AD | 417 | 0.524717 | 1.713202 | 0.000101 | 0.017103 | 0.014729 | 979 | tags=20%, | 54206/584 | LNCaP+DHT |
| SMID_BRE | SMID_BRE | SMID_BRE | 394 | 0.508815 | 1.657811 | 0.000101 | 0.017103 | 0.014729 | 955 | tags=20%, | 26018/348 | LNCaP+DHT |
| ENK_UV_R | ENK_UV_R | ENK_UV_R | 378 | 0.490066 | 1.596144 | 0.000101 | 0.017103 | 0.014729 | 1634 | tags=28%, | 22936/102 | LNCaP+DHT |
| DACOSTA_ | DACOSTA_ | DACOSTA_ | 362 | 0.547871 | 1.782382 | 0.000101 | 0.017103 | 0.014729 | 2285 | tags=44%, | 2181/2597 | LNCaP+DHT |
| ZWANG_CI | ZWANG_CI | ZWANG_CI | 327 | 0.535248 | 1.736006 | 0.000102 | 0.017103 | 0.014729 | 1191 | tags=26%, | 57619/110 | LNCaP+DHT |
| BAELDE_DI | BAELDE_DI | BAELDE_DI | 310 | 0.509021 | 1.650059 | 0.000102 | 0.017103 | 0.014729 | 1324 | tags=22%, | 26018/218 | LNCaP+DHT |
| SENESE_HD | SENESE_HD | SENESE_HD | 295 | 0.527277 | 1.705912 | 0.000102 | 0.017103 | 0.014729 | 1239 | tags=26%, | 54206/102 | LNCaP+DHT |
| LU_EZH2_T | LU_EZH2_T | LU_EZH2_T | 270 | 0.551409 | 1.776163 | 0.000103 | 0.017103 | 0.014729 | 1499 | tags=29%, | 26018/229 | LNCaP+DHT |
| DAZARD_R | DAZARD_R | DAZARD_R | 235 | 0.564706 | 1.805903 | 0.000104 | 0.017103 | 0.014729 | 1741 | tags=36%, | 9732/2597 | LNCaP+DHT |
| DURCHDEV | DURCHDEV | DURCHDEV | 182 | 0.56626 | 1.784131 | 0.000107 | 0.017103 | 0.014729 | 1142 | tags=26%, | 22936/576 | LNCaP+DHT |
| HALLMARK | HALLMARK | HALLMARK | 143 | 0.605317 | 1.876355 | 0.00011 | 0.017103 | 0.014729 | 904 | tags=22%, | 23052/228 | LNCaP+DHT |
| DOANE_RE | DOANE_RE | DOANE_RE | 129 | 0.726839 | 2.235147 | 0.000111 | 0.017103 | 0.014729 | 911 | tags=31%, | 7113/2305 | LNCaP+DHT |
| VECCHI_GA | VECCHI_GA | VECCHI_GA | 129 | 0.636209 | 1.956448 | 0.000111 | 0.017103 | 0.014729 | 695 | tags=20%, | 2289/3977 | LNCaP+DHT |
| BOYLAN_M | BOYLAN_M | BOYLAN_M | 119 | 0.635198 | 1.940924 | 0.000112 | 0.017103 | 0.014729 | 924 | tags=21%, | 4094/3480 | LNCaP+DHT |
| GROSS_HY | GROSS_HY | GROSS_HY | 107 | 0.653214 | 1.976913 | 0.000114 | 0.017103 | 0.014729 | 927 | tags=24%, | 54206/182 | LNCaP+DHT |
| PLASARI_T | PLASARI_T | PLASARI_T | 104 | 0.661286 | 1.996154 | 0.000115 | 0.017103 | 0.014729 | 914 | tags=22%, | 23052/228 | LNCaP+DHT |
| HALLMARK | HALLMARK | HALLMARK | 93 | 0.906231 | 2.703627 | 0.000116 | 0.017103 | 0.014729 | 584 | tags=66%, | 7113/2289 | LNCaP+DHT |
| YEGNASUB | YEGNASUB | YEGNASUB | 91 | 0.683998 | 2.034465 | 0.000117 | 0.017103 | 0.014729 | 382 | tags=21%, | 7113/3817 | LNCaP+DHT |
| NELSON_RI | NELSON_RI | NELSON_RI | 74 | 0.913027 | 2.639653 | 0.000121 | 0.017103 | 0.014729 | 582 | tags=70%, | 7113/2289 | LNCaP+DHT |
| GAVIN_FO | GAVIN_FO | GAVIN_FO | 62 | 0.699446 | 1.971824 | 0.000125 | 0.017103 | 0.014729 | 487 | tags=21%, | 23052/260 | LNCaP+DHT |
| PID_AR_PA | PID_AR_PA | PID_AR_PA | 48 | 0.711719 | 1.938511 | 0.000131 | 0.017103 | 0.014729 | 584 | tags=19%, | 7113/3817 | LNCaP+DHT |
| HALMOS_C | HALMOS_C | HALMOS_C | 33 | 0.765812 | 1.972976 | 0.000139 | 0.017103 | 0.014729 | 412 | tags=27%, | 25803/258 | LNCaP+DHT |
| WANG_RES | WANG_RES | WANG_RES | 26 | 0.901935 | 2.234399 | 0.000144 | 0.017103 | 0.014729 | 560 | tags=58%, | 3817/4094 | LNCaP+DHT |
| BURTON_A | BURTON_A | BURTON_A | 23 | 0.831439 | 2.011763 | 0.000147 | 0.017103 | 0.014729 | 863 | tags=57%, | 54206/229 | LNCaP+DHT |
| PIONTEK_P | PIONTEK_P | PIONTEK_P | 21 | 0.83613 | 1.991225 | 0.000149 | 0.017103 | 0.014729 | 227 | tags=29%, | 54206/228 | LNCaP+DHT |
| WANG_RES | WANG_RES | WANG_RES | 21 | 0.844886 | 2.012078 | 0.000149 | 0.017103 | 0.014729 | 531 | tags=48%, | 3817/1064 | LNCaP+DHT |
| FINETTI_BR | FINETTI_BR | FINETTI_BR | 14 | 0.875079 | 1.928437 | 0.000161 | 0.017908 | 0.015422 | 116 | tags=21%, | 3480/2734 | LNCaP+DHT |
| KRIGE_RES | KRIGE_RES | KRIGE_RES | 634 | 0.434646 | 1.427596 | 0.0002 | 0.017946 | 0.015455 | 1765 | tags=24%, | 54206/102 | LNCaP+DHT |
| JOHNSTON | JOHNSTON | JOHNSTON | 632 | 0.441639 | 1.450479 | 0.0002 | 0.017946 | 0.015455 | 2351 | tags=29%, | 2289/5583 | LNCaP+DHT |
| SCHLOSSER | SCHLOSSER | SCHLOSSER | 520 | 0.45707 | 1.49752 | 0.000201 | 0.017946 | 0.015455 | 1791 | tags=24%, | 26018/716 | LNCaP+DHT |
| MILI_PSEU | MILI_PSEU | MILI_PSEU | 496 | 0.4562 | 1.493551 | 0.000201 | 0.017946 | 0.015455 | 1226 | tags=17%, | 23052/348 | LNCaP+DHT |
| CHARAFE_I | CHARAFE_I | CHARAFE_I | 264 | 0.541846 | 1.744389 | 0.000206 | 0.017946 | 0.015455 | 797 | tags=15%, | 10892/561 | LNCaP+DHT |
| FOSTER_TC | FOSTER_TC | FOSTER_TC | 267 | 0.519062 | 1.671122 | 0.000207 | 0.017946 | 0.015455 | 1027 | tags=18%, | 54206/230 | LNCaP+DHT |
| CHARAFE_I | CHARAFE_I | CHARAFE_I | 263 | 0.529907 | 1.704882 | 0.000207 | 0.017946 | 0.015455 | 1705 | tags=27%, | 22936/561 | LNCaP+DHT |
| PASINI_SU | PASINI_SU | PASINI_SU | 214 | 0.547215 | 1.743231 | 0.00021 | 0.017946 | 0.015455 | 924 | tags=23%, | 7113/5420 | LNCaP+DHT |
| CREIGHTON | CREIGHTON | CREIGHTON | 184 | 0.546527 | 1.724917 | 0.000213 | 0.017946 | 0.015455 | 1209 | tags=22%, | 2289/2601 | LNCaP+DHT |
| LINDGREN_ | LINDGREN_ | LINDGREN_ | 179 | 0.54769 | 1.723906 | 0.000214 | 0.017946 | 0.015455 | 1497 | tags=27%, | 26018/409 | LNCaP+DHT |
| DUTERTRE_ | DUTERTRE_ | DUTERTRE_ | 174 | 0.550705 | 1.72961 | 0.000214 | 0.017946 | 0.015455 | 904 | tags=16%, | 23052/228 | LNCaP+DHT |
| LU_AGING_ | LU_AGING_ | LU_AGING_ | 170 | 0.545179 | 1.710325 | 0.000215 | 0.017946 | 0.015455 | 1142 | tags=25%, | 26018/259 | LNCaP+DHT |
| MASSARWI | MASSARWI | MASSARWI | 153 | 0.607882 | 1.890926 | 0.000218 | 0.017946 | 0.015455 | 851 | tags=24%, | 2289/2601 | LNCaP+DHT |
| PICCALUGA | PICCALUGA | PICCALUGA | 98 | 0.651439 | 1.951687 | 0.000231 | 0.018412 | 0.015856 | 1486 | tags=42%, | 65986/259 | LNCaP+DHT |
| COULOUAR | COULOUAR | COULOUAR | 97 | 0.63195 | 1.889217 | 0.000232 | 0.018412 | 0.015856 | 558 | tags=23%, | 54206/218 | LNCaP+DHT |
| DACOSTA_ | DACOSTA_ | DACOSTA_ | 59 | 0.678083 | 1.897145 | 0.000252 | 0.019339 | 0.016655 | 1486 | tags=37%, | 3480/1089 | LNCaP+DHT |
| DACOSTA_ | DACOSTA_ | DACOSTA_ | 55 | 0.693234 | 1.926487 | 0.000253 | 0.019339 | 0.016655 | 1490 | tags=38%, | 10892/973 | LNCaP+DHT |
| RAGHAVAC | RAGHAVAC | RAGHAVAC | 48 | 0.688575 | 1.875476 | 0.000261 | 0.019588 | 0.016869 | 1245 | tags=31%, | 23052/495 | LNCaP+DHT |
| NELSON_RI | NELSON_RI | NELSON_RI | 16 | -0.895183 | -2.426948 | 0.000275 | 0.019862 | 0.017105 | 877 | tags=88%, | 6337/4609 | LNCaP+DHT |
| GRYDER_PA | GRYDER_PA | GRYDER_PA | 745 | 0.429209 | 1.412239 | 0.0003 | 0.019862 | 0.017105 | 1500 | tags=20%, | 54206/260 | LNCaP+DHT |
| NUYTTEN_I | NUYTTEN_I | NUYTTEN_I | 617 | 0.432309 | 1.41961 | 0.0003 | 0.019862 | 0.017105 | 2094 | tags=28%, | 7113/1296 | LNCaP+DHT |
| OSMAN_BL | OSMAN_BL | OSMAN_BL | 296 | 0.490935 | 1.588644 | 0.000307 | 0.019862 | 0.017105 | 2402 | tags=37%, | 10645/912 | LNCaP+DHT |
| BRACHAT_ | BRACHAT_ | BRACHAT_ | 22 | -0.694235 | -2.035846 | 0.00031 | 0.019862 | 0.017105 | 466 | tags=27%, | 1026/7832 | LNCaP+DHT |
| FARMER_B | FARMER_B | FARMER_B | 239 | 0.521016 | 1.668396 | 0.000312 | 0.019862 | 0.017105 | 927 | tags=18%, | 7113/2181 | LNCaP+DHT |
| SCHAEFFER | SCHAEFFER | SCHAEFFER | 237 | 0.528274 | 1.690109 | 0.000313 | 0.019862 | 0.017105 | 1110 | tags=19%, | 7113/5420 | LNCaP+DHT |
| GRESHOCK_ | GRESHOCK_ | GRESHOCK_ | 232 | 0.534965 | 1.710332 | 0.000313 | 0.019862 | 0.017105 | 1793 | tags=26%, | 7113/3977 | LNCaP+DHT |
| GABRIELY_ | GABRIELY_ | GABRIELY_ | 230 | 0.53416 | 1.707165 | 0.000313 | 0.019862 | 0.017105 | 2471 | tags=47%, | 3977/2332 | LNCaP+DHT |
| FARMER_B | FARMER_B | FARMER_B | 229 | 0.526177 | 1.681315 | 0.000313 | 0.019862 | 0.017105 | 935 | tags=19%, | 7113/2289 | LNCaP+DHT |

| | | | | | | | | | | | |
|---|---|---|---|---|---|---|---|---|---|---|---|
| SCHUETZ_E | SCHUETZ_E | SCHUETZ_E | 134 | 0.586472 | 1.809944 | 0.000331 | 0.020393 | 0.017562 | 972 | tags=24%, | 4094/1025 | LNCaP+DHT |
| STARK_PRE | STARK_PRE | STARK_PRE | 134 | 0.575069 | 1.774753 | 0.000331 | 0.020393 | 0.017562 | 905 | tags=21%, | 23052/584 | LNCaP+DHT |
| TAKEDA_T | TAKEDA_T | TAKEDA_T | 75 | 0.656063 | 1.901462 | 0.000362 | 0.021952 | 0.018905 | 1140 | tags=28%, | 4094/9732, | LNCaP+DHT |
| KEGG_ADH | KEGG_ADH | KEGG_ADH | 59 | 0.671534 | 1.878822 | 0.000379 | 0.022637 | 0.019495 | 1112 | tags=25%, | 3480/7414, | LNCaP+DHT |
| REACTOME | REACTOME | REACTOME | 42 | -0.665823 | -2.290528 | 0.000401 | 0.023063 | 0.019862 | 3372 | tags=76%, | 4728/4726, | LNCaP+DHT |
| FORTSCHE | FORTSCHE | FORTSCHE | 528 | 0.433183 | 1.419634 | 0.000401 | 0.023063 | 0.019862 | 1327 | tags=18%, | 22936/716 | LNCaP+DHT |
| GOZGIT_ES | GOZGIT_ES | GOZGIT_ES | 448 | 0.452589 | 1.479479 | 0.000403 | 0.023063 | 0.019862 | 1489 | tags=24%, | 4094/7997 | LNCaP+DHT |
| SENGUPTA | SENGUPTA | SENGUPTA | 235 | 0.515537 | 1.648663 | 0.000417 | 0.023556 | 0.020286 | 1695 | tags=26%, | 3977/6598 | LNCaP+DHT |
| SWEET_LU | SWEET_LU | SWEET_LU | 186 | 0.524686 | 1.657232 | 0.000425 | 0.023712 | 0.020421 | 1101 | tags=23%, | 3977/7704, | LNCaP+DHT |
| DORN_ADE | DORN_ADE | DORN_ADE | 10 | -0.78765 | -1.865484 | 0.000493 | 0.026964 | 0.023221 | 643 | tags=50%, | 4609/4791, | LNCaP+DHT |
| REACTOME | REACTOME | REACTOME | 54 | 0.675146 | 1.871864 | 0.000507 | 0.026964 | 0.023221 | 1038 | tags=20%, | 3817/354/ | LNCaP+DHT |
| ACEVEDO_ | ACEVEDO_ | ACEVEDO_ | 297 | 0.485843 | 1.572294 | 0.000512 | 0.026964 | 0.023221 | 1754 | tags=22%, | 54206/228 | LNCaP+DHT |
| XU_RESPO | XU_RESPO | XU_RESPO | 12 | -0.775928 | -1.928346 | 0.000513 | 0.026964 | 0.023221 | 833 | tags=58%, | 3276/1434, | LNCaP+DHT |
| BOQUEST_ | BOQUEST_ | BOQUEST_ | 234 | 0.499707 | 1.598221 | 0.000521 | 0.026964 | 0.023221 | 1297 | tags=25%, | 4094/6598 | LNCaP+DHT |
| LENAOUR_ | LENAOUR_ | LENAOUR_ | 66 | -0.547572 | -2.065548 | 0.000531 | 0.026964 | 0.023221 | 2057 | tags=44%, | 3988/9308, | LNCaP+DHT |
| MOOTHA_ | MOOTHA_ | MOOTHA_ | 66 | -0.582212 | -2.196213 | 0.000531 | 0.026964 | 0.023221 | 3255 | tags=71%, | 4726/6835, | LNCaP+DHT |
| CHEN_HOX | CHEN_HOX | CHEN_HOX | 173 | 0.534109 | 1.676649 | 0.000536 | 0.026964 | 0.023221 | 1977 | tags=35%, | 25976/235 | LNCaP+DHT |
| SHEDDEN_ | SHEDDEN_ | SHEDDEN_ | 117 | 0.574445 | 1.751639 | 0.000563 | 0.027567 | 0.023741 | 1386 | tags=27%, | 79098/770 | LNCaP+DHT |
| BRACHAT_ | BRACHAT_ | BRACHAT_ | 17 | -0.710739 | -1.952309 | 0.000564 | 0.027567 | 0.023741 | 649 | tags=41%, | 9249/3983, | LNCaP+DHT |
| KEGG_RIB | KEGG_RIB | KEGG_RIB | 73 | -0.617466 | -2.365844 | 0.000568 | 0.027567 | 0.023741 | 2390 | tags=59%, | 6133/6155, | LNCaP+DHT |
| REACTOME | REACTOME | REACTOME | 76 | -0.629188 | -2.436288 | 0.000586 | 0.027627 | 0.023793 | 3255 | tags=72%, | 6341/4726, | LNCaP+DHT |
| REACTOME | REACTOME | REACTOME | 78 | -0.620293 | -2.410494 | 0.000604 | 0.027627 | 0.023793 | 3255 | tags=71%, | 6341/4726, | LNCaP+DHT |
| MOOTHA_ | MOOTHA_ | MOOTHA_ | 21 | -0.664896 | -1.926719 | 0.000605 | 0.027627 | 0.023793 | 1210 | tags=57%, | 2820/2821, | LNCaP+DHT |
| KEGG_PAR | KEGG_PAR | KEGG_PAR | 79 | -0.454294 | -1.770295 | 0.000606 | 0.027627 | 0.023793 | 3070 | tags=62%, | 6390/1351, | LNCaP+DHT |
| SCHAEFFER | SCHAEFFER | SCHAEFFER | 335 | 0.477498 | 1.549944 | 0.00061 | 0.027627 | 0.023793 | 1677 | tags=24%, | 129642/79 | LNCaP+DHT |
| REACTOME | REACTOME | REACTOME | 81 | -0.525022 | -2.053895 | 0.000614 | 0.027627 | 0.023793 | 2390 | tags=56%, | 6133/6155, | LNCaP+DHT |
| CHEN_LVA | CHEN_LVA | CHEN_LVA | 63 | 0.636875 | 1.800362 | 0.000623 | 0.027627 | 0.023793 | 1058 | tags=35%, | 2289/7704, | LNCaP+DHT |
| KEGG_OXI | KEGG_OXI | KEGG_OXI | 83 | -0.547672 | -2.162768 | 0.000626 | 0.027627 | 0.023793 | 3428 | tags=66%, | 4700/9296, | LNCaP+DHT |
| LIM_MAM | LIM_MAM | LIM_MAM | 211 | 0.504614 | 1.607043 | 0.00063 | 0.027627 | 0.023793 | 1419 | tags=21%, | 3977/1089 | LNCaP+DHT |
| GENTILE_R | GENTILE_R | GENTILE_R | 48 | 0.67577 | 1.840597 | 0.000653 | 0.027906 | 0.024032 | 1491 | tags=44%, | 25976/258 | LNCaP+DHT |
| MASSARW | MASSARW | MASSARW | 44 | 0.693482 | 1.87053 | 0.00066 | 0.027906 | 0.024032 | 776 | tags=36%, | 26018/348 | LNCaP+DHT |
| ONDER_CD | ONDER_CD | ONDER_CD | 139 | 0.554295 | 1.714353 | 0.000661 | 0.027906 | 0.024032 | 1246 | tags=27%, | 2289/2601 | LNCaP+DHT |
| THUM_SYS | THUM_SYS | THUM_SYS | 133 | 0.553442 | 1.70767 | 0.000663 | 0.027906 | 0.024032 | 1372 | tags=26%, | 22936/849 | LNCaP+DHT |
| REACTOME | REACTOME | REACTOME | 92 | -0.564977 | -2.256241 | 0.000717 | 0.02915 | 0.025104 | 2390 | tags=54%, | 6133/6155, | LNCaP+DHT |
| REACTOME | REACTOME | REACTOME | 97 | -0.486213 | -1.971425 | 0.000721 | 0.02915 | 0.025104 | 2390 | tags=51%, | 6133/6155, | LNCaP+DHT |
| REACTOME | REACTOME | REACTOME | 97 | -0.498421 | -2.020924 | 0.000721 | 0.02915 | 0.025104 | 2390 | tags=51%, | 6133/5859, | LNCaP+DHT |
| REACTOME | REACTOME | REACTOME | 97 | -0.41803 | -1.694966 | 0.000721 | 0.02915 | 0.025104 | 2390 | tags=46%, | 6133/6155, | LNCaP+DHT |
| PECE_MA | PECE_MA | PECE_MA | 101 | -0.423829 | -1.731095 | 0.000759 | 0.030402 | 0.026182 | 2024 | tags=47%, | 30844/356 | LNCaP+DHT |
| LANDIS_ER | LANDIS_ER | LANDIS_ER | 115 | 0.566477 | 1.724905 | 0.000789 | 0.030907 | 0.026617 | 1063 | tags=23%, | 54206/229 | LNCaP+DHT |
| MIYAGAW | MIYAGAW | MIYAGAW | 112 | 0.567812 | 1.726466 | 0.000791 | 0.030907 | 0.026617 | 897 | tags=22%, | 22936/110 | LNCaP+DHT |
| BERENJEN | BERENJEN | BERENJEN | 15 | 0.842777 | 1.880968 | 0.000794 | 0.030907 | 0.026617 | 215 | tags=27%, | 54206/573 | LNCaP+DHT |
| SENESE_H | SENESE_H | SENESE_H | 342 | 0.476022 | 1.546079 | 0.000812 | 0.031027 | 0.02672 | 1434 | tags=27%, | 54206/102 | LNCaP+DHT |
| BERTUCCI_ | BERTUCCI_ | BERTUCCI_ | 97 | 0.576367 | 1.72305 | 0.000813 | 0.031027 | 0.02672 | 737 | tags=13%, | 23052/129 | LNCaP+DHT |
| HUANG_F | HUANG_F | HUANG_F | 29 | 0.742569 | 1.869709 | 0.000849 | 0.031674 | 0.027277 | 1386 | tags=41%, | 182/3248/ | LNCaP+DHT |
| INGRAM_S | INGRAM_S | INGRAM_S | 71 | 0.629349 | 1.81197 | 0.000852 | 0.031674 | 0.027277 | 930 | tags=23%, | 3480/5617 | LNCaP+DHT |
| BILANGES_ | BILANGES_ | BILANGES_ | 48 | -0.54189 | -1.918045 | 0.000853 | 0.031674 | 0.027277 | 2928 | tags=54%, | 6950/8483 | LNCaP+DHT |
| KIM_ALL_ | KIM_ALL_ | KIM_ALL_ | 112 | -0.468516 | -1.938171 | 0.00087 | 0.032014 | 0.02757 | 3351 | tags=55%, | 271/8417/ | LNCaP+DHT |
| HALLMARK | HALLMARK | HALLMARK | 138 | 0.538328 | 1.665149 | 0.00088 | 0.032014 | 0.02757 | 1555 | tags=28%, | 54206/716 | LNCaP+DHT |
| REACTOME | REACTOME | REACTOME | 116 | -0.544748 | -2.27026 | 0.000885 | 0.032014 | 0.02757 | 3255 | tags=62%, | 6341/4726, | LNCaP+DHT |
| GENTILE_U | GENTILE_U | GENTILE_U | 225 | 0.496287 | 1.584696 | 0.00094 | 0.033721 | 0.029041 | 1491 | tags=29%, | 25976/258 | LNCaP+DHT |
| HELLER_H | HELLER_H | HELLER_H | 178 | 0.522671 | 1.644529 | 0.000962 | 0.034078 | 0.029348 | 2094 | tags=33%, | 4094/9120, | LNCaP+DHT |
| MARZEC_I | MARZEC_I | MARZEC_I | 25 | -0.640845 | -1.939086 | 0.000967 | 0.034078 | 0.029348 | 52 | tags=32%, | 5608/901/ | LNCaP+DHT |
| PEDERSEN_ | PEDERSEN_ | PEDERSEN_ | 62 | 0.63221 | 1.782279 | 0.001001 | 0.034997 | 0.03014 | 512 | tags=23%, | 85414/229 | LNCaP+DHT |
| GAUSSMA | GAUSSMA | GAUSSMA | 114 | 0.560952 | 1.706786 | 0.001015 | 0.03509 | 0.030219 | 1526 | tags=26%, | 26018/283 | LNCaP+DHT |
| REN_ALVE | REN_ALVE | REN_ALVE | 309 | 0.466583 | 1.512119 | 0.001021 | 0.03509 | 0.030219 | 2050 | tags=29%, | 23052/229 | LNCaP+DHT |
| REACTOME | REACTOME | REACTOME | 132 | -0.395399 | -1.681665 | 0.00104 | 0.035373 | 0.030463 | 2390 | tags=44%, | 6133/6155, | LNCaP+DHT |
| HOLLEMA | HOLLEMA | HOLLEMA | 14 | -0.731507 | -1.904864 | 0.001059 | 0.035373 | 0.030463 | 2262 | tags=71%, | 6210/2340 | LNCaP+DHT |
| PID_AR_TF | PID_AR_TF | PID_AR_TF | 43 | 0.680049 | 1.829191 | 0.001059 | 0.035373 | 0.030463 | 419 | tags=19%, | 7113/3817, | LNCaP+DHT |
| GOZGIT_ES | GOZGIT_ES | GOZGIT_ES | 83 | 0.59691 | 1.75312 | 0.001071 | 0.035373 | 0.030463 | 894 | tags=22%, | 3915/2581 | LNCaP+DHT |
| JOHNSTON | JOHNSTON | JOHNSTON | 38 | 0.702626 | 1.85032 | 0.001084 | 0.035373 | 0.030463 | 1556 | tags=45%, | 3977/5734, | LNCaP+DHT |
| REACTOME | REACTOME | REACTOME | 17 | 0.807966 | 1.849478 | 0.001084 | 0.035373 | 0.030463 | 786 | tags=35%, | 26018/579 | LNCaP+DHT |
| LI_AMPLIFI | LI_AMPLIFI | LI_AMPLIFI | 139 | -0.394967 | -1.69757 | 0.001089 | 0.035373 | 0.030463 | 1851 | tags=39%, | 8140/1936, | LNCaP+DHT |
| REACTOME | REACTOME | REACTOME | 141 | -0.397887 | -1.710357 | 0.001119 | 0.035852 | 0.030875 | 2426 | tags=42%, | 9349/6133, | LNCaP+DHT |
| HALLMARK | HALLMARK | HALLMARK | 119 | 0.552208 | 1.687335 | 0.001121 | 0.035852 | 0.030875 | 930 | tags=17%, | 26018/348 | LNCaP+DHT |
| BROWNE_ | BROWNE_ | BROWNE_ | 25 | 0.754521 | 1.854202 | 0.00116 | 0.036786 | 0.03168 | 1476 | tags=44%, | 3480/3164, | LNCaP+DHT |
| YAO_TEMP | YAO_TEMP | YAO_TEMP | 146 | -0.413772 | -1.785474 | 0.001174 | 0.036949 | 0.031821 | 2534 | tags=45%, | 1163/2199 | LNCaP+DHT |

| | | | | | | | | | | | | |
|---|---|---|---|---|---|---|---|---|---|---|---|---|
| REACTOME | REACTOME | REACTOME | 151 | -0.457435 | -1.988925 | 0.001199 | 0.037259 | 0.032087 | 2390 | tags=45%, | 6133/6155, | LNCaP+DHT |
| TAKEDA_TA | TAKEDA_TA | TAKEDA_TA | 41 | 0.677447 | 1.806863 | 0.001202 | 0.037259 | 0.032087 | 1104 | tags=22%, | 4094/2293 | LNCaP+DHT |
| REACTOME | REACTOME | REACTOME | 156 | -0.459616 | -2.002324 | 0.00128 | 0.039406 | 0.033936 | 2390 | tags=46%, | 6133/6155, | LNCaP+DHT |
| MOHANKU | MOHANKU | MOHANKU | 72 | 0.613328 | 1.768343 | 0.001337 | 0.04085 | 0.03518 | 495 | tags=19%, | 4094/1105 | LNCaP+DHT |
| AMIT_EGF_ | AMIT_EGF_ | AMIT_EGF_ | 30 | 0.719284 | 1.821974 | 0.001404 | 0.042481 | 0.036585 | 364 | tags=30%, | 25976/235 | LNCaP+DHT |
| BRACHAT_ | BRACHAT_ | BRACHAT_ | 17 | -0.691863 | -1.900458 | 0.001411 | 0.042481 | 0.036585 | 649 | tags=35%, | 55766/102 | LNCaP+DHT |
| YAO_TEMP | YAO_TEMP | YAO_TEMP | 27 | 0.741272 | 1.845251 | 0.001433 | 0.042669 | 0.036747 | 1267 | tags=26%, | 54206/259 | LNCaP+DHT |
| MUELLER_ | MUELLER_ | MUELLER_ | 19 | -0.672271 | -1.90279 | 0.001448 | 0.042669 | 0.036747 | 826 | tags=21%, | 6788/6385, | LNCaP+DHT |
| WANG_LM | WANG_LM | WANG_LM | 260 | 0.480264 | 1.543919 | 0.001449 | 0.042669 | 0.036747 | 2131 | tags=32%, | 58480/847 | LNCaP+DHT |
| HADDAD_E | HADDAD_E | HADDAD_E | 172 | 0.513555 | 1.61267 | 0.001501 | 0.043696 | 0.037631 | 2309 | tags=40%, | 26018/659 | LNCaP+DHT |
| HALLMARK | HALLMARK | HALLMARK | 176 | -0.436911 | -1.934349 | 0.001517 | 0.043696 | 0.037631 | 3176 | tags=52%, | 2395/8802, | LNCaP+DHT |
| REACTOME | REACTOME | REACTOME | 63 | -0.494162 | -1.851869 | 0.001518 | 0.043696 | 0.037631 | 3782 | tags=63%, | 8363/3017, | LNCaP+DHT |
| SEKI_INFLA | SEKI_INFLA | SEKI_INFLA | 12 | -0.741133 | -1.841873 | 0.001539 | 0.043696 | 0.037631 | 971 | tags=42%, | 3983/8991, | LNCaP+DHT |
| HOLLEMAN | HOLLEMAN | HOLLEMAN | 13 | -0.736155 | -1.877482 | 0.001559 | 0.043696 | 0.037631 | 2262 | tags=69%, | 6210/2340 | LNCaP+DHT |
| HALLMARK | HALLMARK | HALLMARK | 65 | -0.486242 | -1.83347 | 0.001563 | 0.043696 | 0.037631 | 2858 | tags=54%, | 10561/924 | LNCaP+DHT |
| REACTOME | REACTOME | REACTOME | 24 | -0.632216 | -1.893907 | 0.001592 | 0.043696 | 0.037631 | 1210 | tags=50%, | 6576/2597, | LNCaP+DHT |
| BIOCARTA_ | BIOCARTA_ | BIOCARTA_ | 24 | 0.756923 | 1.844598 | 0.001603 | 0.043696 | 0.037631 | 407 | tags=17%, | 3817/3480, | LNCaP+DHT |
| NGUYEN_N | NGUYEN_N | NGUYEN_N | 64 | 0.612436 | 1.735939 | 0.001613 | 0.043696 | 0.037631 | 1341 | tags=30%, | 3480/2293 | LNCaP+DHT |
| SCHAEFFER | SCHAEFFER | SCHAEFFER | 63 | 0.611351 | 1.728207 | 0.00162 | 0.043696 | 0.037631 | 474 | tags=22%, | 54206/228 | LNCaP+DHT |
| CREIGHTON | CREIGHTON | CREIGHTON | 336 | 0.460515 | 1.495103 | 0.001626 | 0.043696 | 0.037631 | 1336 | tags=23%, | 7113/5848 | LNCaP+DHT |
| BYSTROEM | BYSTROEM | BYSTROEM | 26 | -0.595859 | -1.822602 | 0.001639 | 0.043696 | 0.037631 | 865 | tags=27%, | 6809/5441, | LNCaP+DHT |
| LU_EZH2_T | LU_EZH2_T | LU_EZH2_T | 185 | -0.406811 | -1.806475 | 0.001647 | 0.043696 | 0.037631 | 2995 | tags=50%, | 84287/126 | LNCaP+DHT |
| BERENJENC | BERENJENC | BERENJENC | 35 | 0.7044 | 1.82996 | 0.001649 | 0.043696 | 0.037631 | 780 | tags=34%, | 3480/7414, | LNCaP+DHT |
| PILON_KLF | PILON_KLF | PILON_KLF | 272 | 0.470197 | 1.515337 | 0.00165 | 0.043696 | 0.037631 | 1877 | tags=24%, | 129642/34 | LNCaP+DHT |
| NABA_MAT | NABA_MAT | NABA_MAT | 188 | -0.315363 | -1.404761 | 0.001653 | 0.043696 | 0.037631 | 2325 | tags=32%, | 4185/1027 | LNCaP+DHT |
| WONG_MI | WONG_MI | WONG_MI | 190 | -0.38062 | -1.697117 | 0.001669 | 0.043852 | 0.037765 | 3117 | tags=51%, | 10128/470 | LNCaP+DHT |
| SMID_BREA | SMID_BREA | SMID_BREA | 28 | 0.730693 | 1.827944 | 0.001714 | 0.044624 | 0.03843 | 873 | tags=39%, | 81563/770 | LNCaP+DHT |
| DUTERTRE_ | DUTERTRE_ | DUTERTRE_ | 367 | 0.449619 | 1.463188 | 0.00172 | 0.044624 | 0.03843 | 1006 | tags=20%, | 7113/5420 | LNCaP+DHT |
| DEURIG_T_ | DEURIG_T_ | DEURIG_T_ | 196 | 0.493596 | 1.564162 | 0.001909 | 0.049215 | 0.042384 | 1523 | tags=23%, | 4094/2181, | LNCaP+DHT |
| VECCHI_GA | VECCHI_GA | VECCHI_GA | 73 | 0.603661 | 1.74338 | 0.001941 | 0.04972 | 0.042819 | 1037 | tags=16%, | 7113/7163, | LNCaP+DHT |
| PARK_APL_ | PARK_APL_ | PARK_APL_ | 32 | 0.711222 | 1.818615 | 0.001953 | 0.04972 | 0.042819 | 923 | tags=25%, | 2289/2752, | LNCaP+DHT |

C42_NCOR2

| | ID | Descriptior | setSize | enrichmen | NES | pvalue | p.adjust | qvalues | rank | leading_ed | core_enricl | Group |
|---|---|---|---|---|---|---|---|---|---|---|---|---|
| GEORGES_ | GEORGES_ | GEORGES_ | 691 | 0.35951 | 1.751091 | 0.000147 | 0.00517 | 0.004001 | 2498 | tags=32%, | 93664/157 | C42-NCOR2 |
| DACOSTA_ | DACOSTA_ | DACOSTA_ | 674 | 0.331739 | 1.613191 | 0.000147 | 0.00517 | 0.004001 | 2365 | tags=29%, | 11073/556 | C42-NCOR2 |
| JOHNSTON | JOHNSTON | JOHNSTON | 647 | 0.312552 | 1.514525 | 0.000149 | 0.00517 | 0.004001 | 2340 | tags=28%, | 157570/10 | C42-NCOR2 |
| PATIL_LIVE | PATIL_LIVE | PATIL_LIVE | 526 | 0.375043 | 1.789181 | 0.000152 | 0.00517 | 0.004001 | 1755 | tags=27%, | 157570/41 | C42-NCOR2 |
| MARSON_E | MARSON_E | MARSON_E | 516 | 0.380871 | 1.814142 | 0.000152 | 0.00517 | 0.004001 | 2322 | tags=29%, | 157570/30 | C42-NCOR2 |
| MEISSNER_ | MEISSNER_ | MEISSNER_ | 542 | 0.32352 | 1.545432 | 0.000152 | 0.00517 | 0.004001 | 2323 | tags=29%, | 4885/7431 | C42-NCOR2 |
| REACTOME | REACTOME | REACTOME | 510 | 0.445968 | 2.121866 | 0.000152 | 0.00517 | 0.004001 | 2109 | tags=31%, | 157570/41 | C42-NCOR2 |
| BENPORAT | BENPORAT | BENPORAT | 500 | 0.395848 | 1.880522 | 0.000153 | 0.00517 | 0.004001 | 1872 | tags=28%, | 10769/417 | C42-NCOR2 |
| HAMAI_AP | HAMAI_AP | HAMAI_AP | 499 | 0.353444 | 1.67831 | 0.000153 | 0.00517 | 0.004001 | 2252 | tags=32%, | 54443/113 | C42-NCOR2 |
| GOZGIT_ES | GOZGIT_ES | GOZGIT_ES | 482 | 0.330907 | 1.567768 | 0.000153 | 0.00517 | 0.004001 | 2870 | tags=38%, | 7227/1076 | C42-NCOR2 |
| REACTOME | REACTOME | REACTOME | 436 | 0.464487 | 2.184718 | 0.000154 | 0.00517 | 0.004001 | 2872 | tags=40%, | 157570/41 | C42-NCOR2 |
| GOBERT_O | GOBERT_O | GOBERT_O | 431 | 0.504475 | 2.368166 | 0.000155 | 0.00517 | 0.004001 | 2370 | tags=42%, | 157570/74 | C42-NCOR2 |
| BERENJENC | BERENJENC | BERENJENC | 413 | 0.330937 | 1.546861 | 0.000156 | 0.00517 | 0.004001 | 2105 | tags=28%, | 10769/417 | C42-NCOR2 |
| ONKEN_UV | ONKEN_UV | ONKEN_UV | 396 | 0.342351 | 1.592636 | 0.000157 | 0.00517 | 0.004001 | 1846 | tags=28%, | 93664/556 | C42-NCOR2 |
| CHICAS_RB | CHICAS_RB | CHICAS_RB | 374 | 0.341005 | 1.579816 | 0.000157 | 0.00517 | 0.004001 | 1965 | tags=27%, | 4173/3070 | C42-NCOR2 |
| DACOSTA_ | DACOSTA_ | DACOSTA_ | 367 | 0.329256 | 1.52358 | 0.000158 | 0.00517 | 0.004001 | 1905 | tags=25%, | 11073/556 | C42-NCOR2 |
| REACTOME | REACTOME | REACTOME | 359 | 0.327769 | 1.513928 | 0.000158 | 0.00517 | 0.004001 | 2408 | tags=29%, | 2787/5567 | C42-NCOR2 |
| PUJANA_BI | PUJANA_BI | PUJANA_BI | 348 | 0.436999 | 2.013662 | 0.000158 | 0.00517 | 0.004001 | 2947 | tags=38%, | 4173/1107 | C42-NCOR2 |
| CREIGHTON | CREIGHTON | CREIGHTON | 350 | 0.344736 | 1.589118 | 0.000158 | 0.00517 | 0.004001 | 2296 | tags=32%, | 10769/113 | C42-NCOR2 |
| SHEDDEN_ | SHEDDEN_ | SHEDDEN_ | 368 | 0.450477 | 2.083833 | 0.000158 | 0.00517 | 0.004001 | 2673 | tags=39%, | 4173/5587 | C42-NCOR2 |
| REACTOME | REACTOME | REACTOME | 334 | 0.458383 | 2.104013 | 0.000158 | 0.00517 | 0.004001 | 2939 | tags=45%, | 3817/3012 | C42-NCOR2 |
| ZHANG_BR | ZHANG_BR | ZHANG_BR | 333 | 0.329049 | 1.509442 | 0.000158 | 0.00517 | 0.004001 | 2273 | tags=27%, | 157570/72 | C42-NCOR2 |
| GRYDER_P | GRYDER_P | GRYDER_P | 324 | 0.35501 | 1.625213 | 0.000159 | 0.00517 | 0.004001 | 2257 | tags=31%, | 7431/5444 | C42-NCOR2 |
| TOYOTA_T | TOYOTA_T | TOYOTA_T | 320 | 0.384526 | 1.756044 | 0.000159 | 0.00517 | 0.004001 | 2664 | tags=36%, | 157313/30 | C42-NCOR2 |
| REACTOME | REACTOME | REACTOME | 310 | 0.437653 | 1.993254 | 0.00016 | 0.00517 | 0.004001 | 2572 | tags=35%, | 3012/8335 | C42-NCOR2 |
| HELLER_HD | HELLER_HD | HELLER_HD | 309 | 0.354742 | 1.614834 | 0.00016 | 0.00517 | 0.004001 | 2534 | tags=33%, | 4885/1076 | C42-NCOR2 |
| BLUM_RES | BLUM_RES | BLUM_RES | 283 | 0.477159 | 2.152112 | 0.000161 | 0.00517 | 0.004001 | 1990 | tags=37%, | 11073/558 | C42-NCOR2 |
| LINDGREN_ | LINDGREN_ | LINDGREN_ | 277 | 0.369889 | 1.66446 | 0.000161 | 0.00517 | 0.004001 | 2911 | tags=40%, | 157313/29 | C42-NCOR2 |
| WANG_RES | WANG_RES | WANG_RES | 272 | 0.368576 | 1.654021 | 0.000162 | 0.00517 | 0.004001 | 1590 | tags=23%, | 157570/30 | C42-NCOR2 |
| BENPORAT | BENPORAT | BENPORAT | 266 | 0.374514 | 1.67657 | 0.000162 | 0.00517 | 0.004001 | 2176 | tags=30%, | 64849/101 | C42-NCOR2 |
| DUTERTRE_ | DUTERTRE_ | DUTERTRE_ | 269 | 0.545755 | 2.443919 | 0.000163 | 0.00517 | 0.004001 | 2137 | tags=43%, | 157570/41 | C42-NCOR2 |
| REACTOME | REACTOME | REACTOME | 269 | 0.363744 | 1.628868 | 0.000163 | 0.00517 | 0.004001 | 1912 | tags=25%, | 11073/301 | C42-NCOR2 |
| COLINA_TA | COLINA_TA | COLINA_TA | 250 | 0.352009 | 1.565309 | 0.000164 | 0.00517 | 0.004001 | 2082 | tags=30%, | 5236/9086 | C42-NCOR2 |
| LINDGREN_ | LINDGREN_ | LINDGREN_ | 253 | 0.385505 | 1.715751 | 0.000164 | 0.00517 | 0.004001 | 2054 | tags=32%, | 4173/2912 | C42-NCOR2 |
| CREIGHTON | CREIGHTON | CREIGHTON | 244 | 0.410828 | 1.822507 | 0.000164 | 0.00517 | 0.004001 | 4013 | tags=45%, | 4885/7419 | C42-NCOR2 |
| REACTOME | REACTOME | REACTOME | 233 | 0.391744 | 1.727616 | 0.000164 | 0.00517 | 0.004001 | 2211 | tags=28%, | 2787/3012 | C42-NCOR2 |
| RIGGI_EWI | RIGGI_EWI | RIGGI_EWI | 235 | 0.381691 | 1.684517 | 0.000164 | 0.00517 | 0.004001 | 2118 | tags=33%, | 93664/488 | C42-NCOR2 |
| REACTOME | REACTOME | REACTOME | 236 | 0.435727 | 1.923951 | 0.000164 | 0.00517 | 0.004001 | 2572 | tags=38%, | 4173/1107 | C42-NCOR2 |
| REACTOME | REACTOME | REACTOME | 252 | 0.512184 | 2.277269 | 0.000164 | 0.00517 | 0.004001 | 2939 | tags=49%, | 3817/3012 | C42-NCOR2 |
| HORIUCHI_ | HORIUCHI_ | HORIUCHI_ | 248 | 0.430673 | 1.912806 | 0.000164 | 0.00517 | 0.004001 | 1503 | tags=30%, | 5865/5444 | C42-NCOR2 |
| BASAKI_YB | BASAKI_YB | BASAKI_YB | 219 | 0.478383 | 2.099646 | 0.000164 | 0.00517 | 0.004001 | 1600 | tags=34%, | 10769/417 | C42-NCOR2 |
| GOLDRATH | GOLDRATH | GOLDRATH | 232 | 0.439554 | 1.937441 | 0.000165 | 0.00517 | 0.004001 | 2782 | tags=44%, | 4173/1107 | C42-NCOR2 |
| KOBAYASH | KOBAYASH | KOBAYASH | 209 | 0.600493 | 2.621094 | 0.000165 | 0.00517 | 0.004001 | 2764 | tags=55%, | 4173/5587 | C42-NCOR2 |
| FISCHER_G | FISCHER_G | FISCHER_G | 205 | 0.531437 | 2.313771 | 0.000165 | 0.00517 | 0.004001 | 2566 | tags=51%, | 54443/558 | C42-NCOR2 |
| MITSIADES | MITSIADES | MITSIADES | 205 | 0.396116 | 1.724613 | 0.000165 | 0.00517 | 0.004001 | 1642 | tags=29%, | 4173/1107 | C42-NCOR2 |
| YANG_BCL | YANG_BCL | YANG_BCL | 206 | 0.458429 | 1.997094 | 0.000165 | 0.00517 | 0.004001 | 2108 | tags=39%, | 10157/558 | C42-NCOR2 |
| SENGUPTA | SENGUPTA | SENGUPTA | 201 | 0.466896 | 2.026585 | 0.000165 | 0.00517 | 0.004001 | 2369 | tags=38%, | 157570/41 | C42-NCOR2 |
| HADDAD_E | HADDAD_E | HADDAD_E | 181 | 0.459293 | 1.968699 | 0.000167 | 0.00517 | 0.004001 | 1913 | tags=31%, | 8653/9086 | C42-NCOR2 |
| MARKEY_R | MARKEY_R | MARKEY_R | 182 | 0.436956 | 1.873498 | 0.000167 | 0.00517 | 0.004001 | 2322 | tags=38%, | 4173/1107 | C42-NCOR2 |
| REACTOME | REACTOME | REACTOME | 182 | 0.407311 | 1.746389 | 0.000167 | 0.00517 | 0.004001 | 1788 | tags=26%, | 7404/3012 | C42-NCOR2 |
| HALLMARK | HALLMARK | HALLMARK | 176 | 0.412652 | 1.763516 | 0.000167 | 0.00517 | 0.004001 | 2189 | tags=35%, | 55872/588 | C42-NCOR2 |
| HALLMARK | HALLMARK | HALLMARK | 175 | 0.506327 | 2.16229 | 0.000167 | 0.00517 | 0.004001 | 2782 | tags=45%, | 4173/3070 | C42-NCOR2 |
| FUJII_YBX1 | FUJII_YBX1 | FUJII_YBX1 | 174 | 0.48081 | 2.051477 | 0.000167 | 0.00517 | 0.004001 | 2397 | tags=43%, | 4173/3070 | C42-NCOR2 |
| ZHENG_GL | ZHENG_GL | ZHENG_GL | 172 | 0.516981 | 2.202884 | 0.000168 | 0.00517 | 0.004001 | 2263 | tags=42%, | 4173/1057 | C42-NCOR2 |
| REACTOME | REACTOME | REACTOME | 163 | 0.414862 | 1.75501 | 0.000169 | 0.00517 | 0.004001 | 2109 | tags=28%, | 2787/3012 | C42-NCOR2 |
| REACTOME | REACTOME | REACTOME | 165 | 0.40069 | 1.697109 | 0.000169 | 0.00517 | 0.004001 | 2015 | tags=28%, | 3012/8335 | C42-NCOR2 |
| CHICAS_RB | CHICAS_RB | CHICAS_RB | 161 | 0.438013 | 1.849596 | 0.000169 | 0.00517 | 0.004001 | 2324 | tags=40%, | 4173/5510 | C42-NCOR2 |
| REACTOME | REACTOME | REACTOME | 157 | 0.475232 | 1.99973 | 0.00017 | 0.00517 | 0.004001 | 2572 | tags=38%, | 5885/8461 | C42-NCOR2 |
| GRAHAM_ | GRAHAM_ | GRAHAM_ | 146 | 0.530918 | 2.211708 | 0.000171 | 0.00517 | 0.004001 | 1600 | tags=40%, | 4173/360/ | C42-NCOR2 |
| WHITFIELD | WHITFIELD | WHITFIELD | 146 | 0.440569 | 1.835331 | 0.000171 | 0.00517 | 0.004001 | 2680 | tags=44%, | 4173/5444 | C42-NCOR2 |
| REACTOME | REACTOME | REACTOME | 150 | 0.454483 | 1.89919 | 0.000171 | 0.00517 | 0.004001 | 2361 | tags=35%, | 3009/3012 | C42-NCOR2 |
| PUJANA_XI | PUJANA_XI | PUJANA_XI | 147 | 0.527294 | 2.196908 | 0.000171 | 0.00517 | 0.004001 | 2484 | tags=39%, | 4173/1107 | C42-NCOR2 |
| SARRIO_EP | SARRIO_EP | SARRIO_EP | 142 | 0.507943 | 2.106839 | 0.000171 | 0.00517 | 0.004001 | 1804 | tags=42%, | 4173/5444 | C42-NCOR2 |
| CHIANG_LI | CHIANG_LI | CHIANG_LI | 139 | 0.531214 | 2.197641 | 0.000171 | 0.00517 | 0.004001 | 2054 | tags=47%, | 54443/558 | C42-NCOR2 |
| FLORIO_NE | FLORIO_NE | FLORIO_NE | 137 | 0.661221 | 2.727831 | 0.000172 | 0.00517 | 0.004001 | 2353 | tags=64%, | 157570/54 | C42-NCOR2 |

| | | | | | | | | | | | | |
|---|---|---|---|---|---|---|---|---|---|---|---|---|
| SOTIRIOU_ | SOTIRIOU_ | SOTIRIOU_ | 134 | 0.57382 | 2.360405 | 0.000172 | 0.00517 | 0.004001 | 1692 | tags=49%, l | 4173/2912 | C42-NCOR2 |
| VILLANUEV | VILLANUEV | VILLANUEV | 134 | 0.476367 | 1.95953 | 0.000172 | 0.00517 | 0.004001 | 2667 | tags=41%, l | 54443/715 | C42-NCOR2 |
| BENPORAT | BENPORAT | BENPORAT | 122 | 0.458931 | 1.864313 | 0.000173 | 0.00517 | 0.004001 | 3315 | tags=54%, l | 54443/558 | C42-NCOR2 |
| FERREIRA_ | FERREIRA_ | FERREIRA_ | 122 | 0.471744 | 1.916364 | 0.000173 | 0.00517 | 0.004001 | 2263 | tags=37%, l | 81931/544 | C42-NCOR2 |
| SHEPARD_( | SHEPARD_( | SHEPARD_( | 122 | 0.493472 | 2.00463 | 0.000173 | 0.00517 | 0.004001 | 1693 | tags=34%, l | 4192/8555 | C42-NCOR2 |
| SHEPARD_I | SHEPARD_I | SHEPARD_I | 128 | 0.483515 | 1.975005 | 0.000173 | 0.00517 | 0.004001 | 1484 | tags=32%, l | 4192/4173 | C42-NCOR2 |
| REACTOME | REACTOME | REACTOME | 130 | 0.467094 | 1.911803 | 0.000173 | 0.00517 | 0.004001 | 1912 | tags=32%, l | 4173/1107 | C42-NCOR2 |
| WINNEPEN | WINNEPEN | WINNEPEN | 130 | 0.51659 | 2.114389 | 0.000173 | 0.00517 | 0.004001 | 2263 | tags=42%, l | 4173/5444 | C42-NCOR2 |
| HOFFMANI | HOFFMANI | HOFFMANI | 129 | 0.49987 | 2.044457 | 0.000173 | 0.00517 | 0.004001 | 1819 | tags=37%, l | 3012/5444 | C42-NCOR2 |
| REACTOME | REACTOME | REACTOME | 126 | 0.498542 | 2.030715 | 0.000173 | 0.00517 | 0.004001 | 1912 | tags=33%, l | 11073/834 | C42-NCOR2 |
| REACTOME | REACTOME | REACTOME | 116 | 0.479681 | 1.933369 | 0.000173 | 0.00517 | 0.004001 | 2938 | tags=47%, l | 84617/649 | C42-NCOR2 |
| GAL_LEUKE | GAL_LEUKE | GAL_LEUKE | 123 | 0.490332 | 1.991828 | 0.000173 | 0.00517 | 0.004001 | 2189 | tags=37%, l | 7544/8358 | C42-NCOR2 |
| ROSTY_CEF | ROSTY_CEF | ROSTY_CEF | 123 | 0.653394 | 2.65422 | 0.000173 | 0.00517 | 0.004001 | 2667 | tags=67%, l | 4173/5587 | C42-NCOR2 |
| WHITFIELD | WHITFIELD | WHITFIELD | 123 | 0.443652 | 1.802205 | 0.000173 | 0.00517 | 0.004001 | 2094 | tags=39%, l | 54443/157 | C42-NCOR2 |
| POOLA_IN\ | POOLA_IN\ | POOLA_IN\ | 113 | 0.474066 | 1.902698 | 0.000174 | 0.00517 | 0.004001 | 2312 | tags=44%, l | 4173/5587 | C42-NCOR2 |
| KEGG_CELL | KEGG_CELL | KEGG_CELL | 109 | 0.515365 | 2.058017 | 0.000174 | 0.00517 | 0.004001 | 1955 | tags=39%, l | 4173/8555 | C42-NCOR2 |
| REACTOME | REACTOME | REACTOME | 108 | 0.504026 | 2.00963 | 0.000174 | 0.00517 | 0.004001 | 2179 | tags=38%, l | 3012/8335 | C42-NCOR2 |
| PUJANA_BI | PUJANA_BI | PUJANA_BI | 107 | 0.537393 | 2.139354 | 0.000174 | 0.00517 | 0.004001 | 2747 | tags=42%, l | 4173/1107 | C42-NCOR2 |
| REACTOME | REACTOME | REACTOME | 103 | 0.509398 | 2.017164 | 0.000176 | 0.00517 | 0.004001 | 1912 | tags=33%, l | 11073/834 | C42-NCOR2 |
| CROONQUI | CROONQUI | CROONQUI | 87 | 0.637218 | 2.459445 | 0.000176 | 0.00517 | 0.004001 | 2782 | tags=60%, l | 10157/417 | C42-NCOR2 |
| REACTOME | REACTOME | REACTOME | 89 | 0.561629 | 2.174971 | 0.000176 | 0.00517 | 0.004001 | 2550 | tags=42%, l | 3012/8335 | C42-NCOR2 |
| BURTON_A | BURTON_A | BURTON_A | 90 | 0.513789 | 1.993391 | 0.000176 | 0.00517 | 0.004001 | 1872 | tags=46%, l | 4173/1107 | C42-NCOR2 |
| REACTOME | REACTOME | REACTOME | 88 | 0.515569 | 1.991518 | 0.000176 | 0.00517 | 0.004001 | 2432 | tags=36%, l | 3012/8335 | C42-NCOR2 |
| REACTOME | REACTOME | REACTOME | 101 | 0.521253 | 2.057949 | 0.000176 | 0.00517 | 0.004001 | 2938 | tags=50%, l | 5885/8461 | C42-NCOR2 |
| ZHOU_CELI | ZHOU_CELI | ZHOU_CELI | 98 | 0.672737 | 2.64442 | 0.000176 | 0.00517 | 0.004001 | 2263 | tags=59%, l | 11073/544 | C42-NCOR2 |
| REACTOME | REACTOME | REACTOME | 80 | 0.541741 | 2.058871 | 0.000176 | 0.00517 | 0.004001 | 2432 | tags=39%, l | 3012/8335 | C42-NCOR2 |
| LE_EGR2_T | LE_EGR2_T | LE_EGR2_T | 85 | 0.59144 | 2.270685 | 0.000177 | 0.00517 | 0.004001 | 2066 | tags=49%, l | 4173/3012 | C42-NCOR2 |
| NIKOLSKY_ | NIKOLSKY_ | NIKOLSKY_ | 93 | 0.508029 | 1.981899 | 0.000177 | 0.00517 | 0.004001 | 3446 | tags=53%, l | 7227/5885 | C42-NCOR2 |
| MISSIAGLIA | MISSIAGLIA | MISSIAGLIA | 99 | 0.508152 | 2.001121 | 0.000177 | 0.00517 | 0.004001 | 2004 | tags=43%, l | 23310/475 | C42-NCOR2 |
| VANTVEER_ | VANTVEER_ | VANTVEER_ | 99 | 0.56595 | 2.228733 | 0.000177 | 0.00517 | 0.004001 | 1974 | tags=46%, l | 5885/9134 | C42-NCOR2 |
| WHITEFOR | WHITEFOR | WHITEFOR | 99 | 0.596636 | 2.349575 | 0.000177 | 0.00517 | 0.004001 | 1902 | tags=47%, l | 4173/5587 | C42-NCOR2 |
| ODONNELL | ODONNELL | ODONNELL | 96 | 0.572463 | 2.244968 | 0.000177 | 0.00517 | 0.004001 | 3036 | tags=59%, l | 54443/307 | C42-NCOR2 |
| REACTOME | REACTOME | REACTOME | 95 | 0.527708 | 2.065756 | 0.000177 | 0.00517 | 0.004001 | 2361 | tags=42%, l | 3012/8335 | C42-NCOR2 |
| KONG_E2F | KONG_E2F | KONG_E2F | 78 | 0.647649 | 2.448383 | 0.000177 | 0.00517 | 0.004001 | 2664 | tags=62%, l | 55872/291 | C42-NCOR2 |
| REACTOME | REACTOME | REACTOME | 78 | 0.55253 | 2.088793 | 0.000177 | 0.00517 | 0.004001 | 2245 | tags=37%, l | 3012/8335 | C42-NCOR2 |
| REACTOME | REACTOME | REACTOME | 55 | 0.625492 | 2.227061 | 0.000178 | 0.00517 | 0.004001 | 1912 | tags=44%, l | 3009/3012 | C42-NCOR2 |
| REACTOME | REACTOME | REACTOME | 60 | 0.586859 | 2.124586 | 0.000178 | 0.00517 | 0.004001 | 1990 | tags=35%, l | 3012/8335 | C42-NCOR2 |
| LEE_EARLY | LEE_EARLY | LEE_EARLY | 75 | 0.674923 | 2.538791 | 0.000178 | 0.00517 | 0.004001 | 1902 | tags=61%, l | 4173/5587 | C42-NCOR2 |
| NIKOLSKY_ | NIKOLSKY_ | NIKOLSKY_ | 75 | 0.581776 | 2.18841 | 0.000178 | 0.00517 | 0.004001 | 1586 | tags=47%, l | 9134/2578 | C42-NCOR2 |
| REACTOME | REACTOME | REACTOME | 53 | 0.605908 | 2.143425 | 0.000178 | 0.00517 | 0.004001 | 2300 | tags=40%, l | 3012/8335 | C42-NCOR2 |
| PUJANA_BI | PUJANA_BI | PUJANA_BI | 48 | 0.629316 | 2.177347 | 0.000178 | 0.00517 | 0.004001 | 1366 | tags=33%, l | 4173/1107 | C42-NCOR2 |
| REACTOME | REACTOME | REACTOME | 48 | 0.681657 | 2.358437 | 0.000178 | 0.00517 | 0.004001 | 2109 | tags=54%, l | 3012/8335 | C42-NCOR2 |
| ISHIDA_E2F | ISHIDA_E2F | ISHIDA_E2F | 47 | 0.669902 | 2.304325 | 0.000178 | 0.00517 | 0.004001 | 2944 | tags=68%, l | 54443/715 | C42-NCOR2 |
| REACTOME | REACTOME | REACTOME | 47 | 0.653958 | 2.249479 | 0.000178 | 0.00517 | 0.004001 | 2245 | tags=49%, l | 3012/8335 | C42-NCOR2 |
| KEGG_SYST | KEGG_SYST | KEGG_SYST | 61 | 0.575951 | 2.091243 | 0.000178 | 0.00517 | 0.004001 | 2300 | tags=39%, l | 3012/8335 | C42-NCOR2 |
| REACTOME | REACTOME | REACTOME | 50 | 0.670454 | 2.341883 | 0.000178 | 0.00517 | 0.004001 | 2334 | tags=50%, l | 3012/8335 | C42-NCOR2 |
| SHEPARD_I | SHEPARD_I | SHEPARD_I | 50 | 0.632021 | 2.20764 | 0.000178 | 0.00517 | 0.004001 | 1484 | tags=46%, l | 4192/8555 | C42-NCOR2 |
| KANG_DOX | KANG_DOX | KANG_DOX | 46 | 0.625312 | 2.141882 | 0.000179 | 0.00517 | 0.004001 | 1990 | tags=65%, l | 4173/5587 | C42-NCOR2 |
| REACTOME | REACTOME | REACTOME | 51 | 0.613295 | 2.150371 | 0.000179 | 0.00517 | 0.004001 | 1912 | tags=39%, l | 3012/8335 | C42-NCOR2 |
| TANG_SEN | TANG_SEN | TANG_SEN | 46 | 0.727441 | 2.491705 | 0.000179 | 0.00517 | 0.004001 | 2004 | tags=72%, l | 7544/5587 | C42-NCOR2 |
| REACTOME | REACTOME | REACTOME | 57 | 0.619463 | 2.217559 | 0.000179 | 0.00517 | 0.004001 | 2300 | tags=46%, l | 3012/8335 | C42-NCOR2 |
| REACTOME | REACTOME | REACTOME | 43 | 0.651356 | 2.203652 | 0.000179 | 0.00517 | 0.004001 | 2300 | tags=47%, l | 3012/8335 | C42-NCOR2 |
| WU_APOPT | WU_APOPT | WU_APOPT | 43 | 0.654191 | 2.213242 | 0.000179 | 0.00517 | 0.004001 | 1016 | tags=47%, l | 4173/5444 | C42-NCOR2 |
| GRAHAM_I | GRAHAM_I | GRAHAM_I | 70 | 0.639968 | 2.379168 | 0.000179 | 0.00517 | 0.004001 | 2054 | tags=49%, l | 4173/360/ | C42-NCOR2 |
| REACTOME | REACTOME | REACTOME | 70 | 0.56781 | 2.11091 | 0.000179 | 0.00517 | 0.004001 | 1912 | tags=40%, l | 11073/834 | C42-NCOR2 |
| REACTOME | REACTOME | REACTOME | 58 | 0.601988 | 2.161037 | 0.000179 | 0.00517 | 0.004001 | 2370 | tags=43%, l | 3012/8335 | C42-NCOR2 |
| CROONQUI | CROONQUI | CROONQUI | 67 | 0.594321 | 2.191885 | 0.000179 | 0.00517 | 0.004001 | 3439 | tags=66%, l | 4173/1061 | C42-NCOR2 |
| GAVIN_FOX | GAVIN_FOX | GAVIN_FOX | 67 | 0.610089 | 2.25004 | 0.000179 | 0.00517 | 0.004001 | 2991 | tags=61%, l | 55872/106 | C42-NCOR2 |
| REACTOME | REACTOME | REACTOME | 42 | 0.682524 | 2.297565 | 0.000179 | 0.00517 | 0.004001 | 2300 | tags=52%, l | 3817/3012 | C42-NCOR2 |
| REACTOME | REACTOME | REACTOME | 69 | 0.592538 | 2.196907 | 0.000179 | 0.00517 | 0.004001 | 1912 | tags=41%, l | 11073/834 | C42-NCOR2 |
| REACTOME | REACTOME | REACTOME | 73 | 0.566263 | 2.117251 | 0.000179 | 0.00517 | 0.004001 | 2109 | tags=37%, l | 3012/8335 | C42-NCOR2 |
| ZHOU_CELI | ZHOU_CELI | ZHOU_CELI | 66 | 0.642604 | 2.36296 | 0.000179 | 0.00517 | 0.004001 | 2722 | tags=59%, l | 11073/291 | C42-NCOR2 |
| REACTOME | REACTOME | REACTOME | 41 | 0.680251 | 2.276406 | 0.000179 | 0.00517 | 0.004001 | 2300 | tags=51%, l | 3012/8335 | C42-NCOR2 |
| MOLENAAF | MOLENAAF | MOLENAAF | 40 | 0.696586 | 2.319824 | 0.00018 | 0.00517 | 0.004001 | 929 | tags=45%, l | 360/15731 | C42-NCOR2 |
| REACTOME | REACTOME | REACTOME | 64 | 0.639532 | 2.339878 | 0.00018 | 0.00517 | 0.004001 | 2109 | tags=45%, l | 3817/3012 | C42-NCOR2 |
| WHITFIELD | WHITFIELD | WHITFIELD | 40 | 0.665871 | 2.217533 | 0.00018 | 0.00517 | 0.004001 | 1872 | tags=52%, l | 29127/715 | C42-NCOR2 |

| | | | | | | | | | | | | |
|---|---|---|---|---|---|---|---|---|---|---|---|---|
| KAMMING/ | KAMMING/ | KAMMING/ | 39 | 0.633982 | 2.099127 | 0.00018 | 0.00517 | 0.004001 | 3226 | tags=62%, l | 4173/1107 | C42-NCOR2 |
| ZHAN_MUI | ZHAN_MUI | ZHAN_MUI | 39 | 0.673744 | 2.230782 | 0.00018 | 0.00517 | 0.004001 | 2008 | tags=64%, l | 29127/715 | C42-NCOR2 |
| REACTOME | REACTOME | REACTOME | 63 | 0.65077 | 2.372358 | 0.00018 | 0.00517 | 0.004001 | 2387 | tags=48%, l | 3012/8335 | C42-NCOR2 |
| GRAHAM_( | GRAHAM_( | GRAHAM_( | 62 | 0.597135 | 2.169416 | 0.000181 | 0.00517 | 0.004001 | 1600 | tags=42%, l | 360/29127 | C42-NCOR2 |
| NAKAYAM/ | NAKAYAM/ | NAKAYAM/ | 62 | 0.615059 | 2.234534 | 0.000181 | 0.00517 | 0.004001 | 1913 | tags=56%, l | 4885/7227 | C42-NCOR2 |
| REACTOME | REACTOME | REACTOME | 38 | 0.62515 | 2.058369 | 0.000181 | 0.00517 | 0.004001 | 2055 | tags=45%, l | 3012/8335 | C42-NCOR2 |
| BURTON_A | BURTON_A | BURTON_A | 36 | 0.643243 | 2.093694 | 0.000182 | 0.00517 | 0.004001 | 1484 | tags=44%, l | 54443/291 | C42-NCOR2 |
| PID_ATR_P | PID_ATR_P | PID_ATR_P | 36 | 0.638668 | 2.078803 | 0.000182 | 0.00517 | 0.004001 | 1850 | tags=53%, l | 11073/598 | C42-NCOR2 |
| NADERI_BI | NADERI_BI | NADERI_BI | 37 | 0.637176 | 2.081781 | 0.000183 | 0.00517 | 0.004001 | 1341 | tags=46%, l | 29127/106 | C42-NCOR2 |
| PID_PLK1_I | PID_PLK1_I | PID_PLK1_I | 37 | 0.621746 | 2.031368 | 0.000183 | 0.00517 | 0.004001 | 2759 | tags=65%, l | 8555/2297 | C42-NCOR2 |
| VANTVEER_ | VANTVEER_ | VANTVEER_ | 37 | 0.629771 | 2.057587 | 0.000183 | 0.00517 | 0.004001 | 2316 | tags=51%, l | 9134/5156 | C42-NCOR2 |
| FARMER_B | FARMER_B | FARMER_B | 31 | 0.823048 | 2.592541 | 0.000183 | 0.00517 | 0.004001 | 1257 | tags=77%, l | 4173/2297 | C42-NCOR2 |
| NIKOLSKY_ | NIKOLSKY_ | NIKOLSKY_ | 27 | 0.710061 | 2.163487 | 0.000184 | 0.00517 | 0.004001 | 2360 | tags=67%, l | 7419/8437 | C42-NCOR2 |
| EGUCHI_CE | EGUCHI_CE | EGUCHI_CE | 23 | 0.764018 | 2.247241 | 0.000186 | 0.00517 | 0.004001 | 2664 | tags=87%, l | 4173/9134 | C42-NCOR2 |
| REACTOME | REACTOME | REACTOME | 23 | 0.739357 | 2.174704 | 0.000186 | 0.00517 | 0.004001 | 1819 | tags=70%, l | 993/28227 | C42-NCOR2 |
| KUNINGER_ | KUNINGER_ | KUNINGER_ | 22 | 0.739444 | 2.15021 | 0.000186 | 0.00517 | 0.004001 | 2269 | tags=59%, l | 54443/106 | C42-NCOR2 |
| FINETTI_BI | FINETTI_BI | FINETTI_BI | 15 | 0.800437 | 2.118852 | 0.000188 | 0.00517 | 0.004001 | 1952 | tags=73%, l | 55872/475 | C42-NCOR2 |
| ZHENG_IL2 | ZHENG_IL2 | ZHENG_IL2 | 17 | 0.760714 | 2.07866 | 0.000188 | 0.00517 | 0.004001 | 1205 | tags=47%, l | 29127/568 | C42-NCOR2 |
| PYEON_CA | PYEON_CA | PYEON_CA | 11 | 0.93232 | 2.265116 | 0.000192 | 0.00517 | 0.004001 | 19 | tags=45%, l | 8653/6192 | C42-NCOR2 |
| KEGG_ASC( | KEGG_ASC( | KEGG_ASC( | 10 | -0.820291 | -1.975062 | 0.000208 | 0.00517 | 0.004001 | 988 | tags=70%, l | 217/10720 | C42-NCOR2 |
| GARGALOV | GARGALOV | GARGALOV | 14 | -0.86382 | -2.290386 | 0.000211 | 0.00517 | 0.004001 | 1641 | tags=86%, l | 1051/1054 | C42-NCOR2 |
| LUI_THYRC | LUI_THYRC | LUI_THYRC | 20 | -0.750759 | -2.1943 | 0.000215 | 0.00517 | 0.004001 | 1592 | tags=60%, l | 10430/619 | C42-NCOR2 |
| REACTOME | REACTOME | REACTOME | 22 | -0.759392 | -2.272773 | 0.000216 | 0.00517 | 0.004001 | 803 | tags=50%, l | 7407/4677 | C42-NCOR2 |
| KRIGE_AMI | KRIGE_AMI | KRIGE_AMI | 27 | -0.743022 | -2.338364 | 0.000219 | 0.00517 | 0.004001 | 1004 | tags=63%, l | 1051/440/ | C42-NCOR2 |
| WARTERS_ | WARTERS_ | WARTERS_ | 30 | -0.67191 | -2.161548 | 0.000222 | 0.00517 | 0.004001 | 1952 | tags=53%, l | 282991/55 | C42-NCOR2 |
| SENGUPTA | SENGUPTA | SENGUPTA | 62 | -0.509757 | -1.911442 | 0.000224 | 0.00517 | 0.004001 | 525 | tags=19%, l | 26154/488 | C42-NCOR2 |
| TARTE_PLA | TARTE_PLA | TARTE_PLA | 63 | -0.524106 | -1.968814 | 0.000225 | 0.00517 | 0.004001 | 790 | tags=32%, l | 10960/109 | C42-NCOR2 |
| REACTOME | REACTOME | REACTOME | 40 | -0.619865 | -2.122609 | 0.000226 | 0.00517 | 0.004001 | 1575 | tags=42%, l | 51067/546 | C42-NCOR2 |
| KEGG_AMI | KEGG_AMI | KEGG_AMI | 41 | -0.62252 | -2.142219 | 0.000226 | 0.00517 | 0.004001 | 1575 | tags=41%, l | 123283/23 | C42-NCOR2 |
| MOOTHA_' | MOOTHA_' | MOOTHA_' | 66 | -0.640664 | -2.429536 | 0.000226 | 0.00517 | 0.004001 | 2517 | tags=61%, l | 9167/6835 | C42-NCOR2 |
| REACTOME | REACTOME | REACTOME | 42 | -0.645092 | -2.228925 | 0.000226 | 0.00517 | 0.004001 | 3303 | tags=71%, l | 4702/5596 | C42-NCOR2 |
| NIKOLSKY_ | NIKOLSKY_ | NIKOLSKY_ | 70 | -0.491613 | -1.883403 | 0.000227 | 0.00517 | 0.004001 | 2411 | tags=40%, l | 84219/531 | C42-NCOR2 |
| SCHUHMA( | SCHUHMA( | SCHUHMA( | 71 | -0.486589 | -1.869252 | 0.000227 | 0.00517 | 0.004001 | 3351 | tags=55%, l | 4522/6566 | C42-NCOR2 |
| KEGG_RIBC | KEGG_RIBC | KEGG_RIBC | 74 | -0.554884 | -2.150125 | 0.000227 | 0.00517 | 0.004001 | 1586 | tags=51%, l | 6204/6147 | C42-NCOR2 |
| YAO_TEMP | YAO_TEMP | YAO_TEMP | 51 | -0.554482 | -1.995696 | 0.000227 | 0.00517 | 0.004001 | 2412 | tags=45%, l | 5685/5688 | C42-NCOR2 |
| ANDERSEN | ANDERSEN | ANDERSEN | 50 | -0.623923 | -2.233833 | 0.000228 | 0.00517 | 0.004001 | 1595 | tags=48%, l | 2052/1376 | C42-NCOR2 |
| WARTERS_ | WARTERS_ | WARTERS_ | 48 | -0.589129 | -2.090026 | 0.000228 | 0.00517 | 0.004001 | 1820 | tags=48%, l | 8824/5776 | C42-NCOR2 |
| REACTOME | REACTOME | REACTOME | 76 | -0.674506 | -2.61989 | 0.000229 | 0.00517 | 0.004001 | 2907 | tags=68%, l | 51103/542 | C42-NCOR2 |
| BURTON_A | BURTON_A | BURTON_A | 91 | -0.49115 | -1.971251 | 0.000229 | 0.00517 | 0.004001 | 2760 | tags=51%, l | 10105/122 | C42-NCOR2 |
| LI_DCP2_B | LI_DCP2_B | LI_DCP2_B | 77 | -0.526605 | -2.049836 | 0.000229 | 0.00517 | 0.004001 | 2093 | tags=45%, l | 8214/5115 | C42-NCOR2 |
| REACTOME | REACTOME | REACTOME | 82 | -0.543266 | -2.14094 | 0.000229 | 0.00517 | 0.004001 | 1586 | tags=50%, l | 11224/620 | C42-NCOR2 |
| IVANOVA_I | IVANOVA_I | IVANOVA_I | 95 | -0.478338 | -1.93598 | 0.00023 | 0.00517 | 0.004001 | 2055 | tags=31%, l | 1161/6341 | C42-NCOR2 |
| KEGG_PAR | KEGG_PAR | KEGG_PAR | 81 | -0.573872 | -2.25505 | 0.00023 | 0.00517 | 0.004001 | 2517 | tags=54%, l | 9167/842/ | C42-NCOR2 |
| REACTOME | REACTOME | REACTOME | 99 | -0.547667 | -2.229982 | 0.00023 | 0.00517 | 0.004001 | 1586 | tags=47%, l | 6204/2058 | C42-NCOR2 |
| REACTOME | REACTOME | REACTOME | 93 | -0.541046 | -2.176792 | 0.00023 | 0.00517 | 0.004001 | 2005 | tags=47%, l | 6204/6147 | C42-NCOR2 |
| REACTOME | REACTOME | REACTOME | 79 | -0.65886 | -2.57509 | 0.00023 | 0.00517 | 0.004001 | 2907 | tags=66%, l | 51103/542 | C42-NCOR2 |
| KEGG_OXII | KEGG_OXII | KEGG_OXII | 84 | -0.495444 | -1.960495 | 0.00023 | 0.00517 | 0.004001 | 1616 | tags=40%, l | 4723/8992 | C42-NCOR2 |
| REACTOME | REACTOME | REACTOME | 98 | -0.482218 | -1.9596 | 0.000231 | 0.00517 | 0.004001 | 1586 | tags=38%, l | 6209/6165 | C42-NCOR2 |
| REACTOME | REACTOME | REACTOME | 98 | -0.599831 | -2.437546 | 0.000231 | 0.00517 | 0.004001 | 1586 | tags=47%, l | 6209/6165 | C42-NCOR2 |
| SMID_BRE/ | SMID_BRE/ | SMID_BRE/ | 86 | -0.483084 | -1.917893 | 0.000231 | 0.00517 | 0.004001 | 1473 | tags=31%, l | 367/2867/ | C42-NCOR2 |
| HALLMARK | HALLMARK | HALLMARK | 100 | -0.533489 | -2.174119 | 0.000231 | 0.00517 | 0.004001 | 1947 | tags=40%, l | 4189/5515 | C42-NCOR2 |
| BROWN_M | BROWN_M | BROWN_M | 89 | -0.471789 | -1.88238 | 0.000231 | 0.00517 | 0.004001 | 2134 | tags=39%, l | 2774/328/ | C42-NCOR2 |
| PECE_MAN | PECE_MAN | PECE_MAN | 103 | -0.537129 | -2.199701 | 0.000232 | 0.00517 | 0.004001 | 1830 | tags=42%, l | 6147/6902 | C42-NCOR2 |
| COULOUAF | COULOUAF | COULOUAF | 104 | -0.455683 | -1.865907 | 0.000234 | 0.00517 | 0.004001 | 653 | tags=23%, l | 2040/9380 | C42-NCOR2 |
| YAO_TEMP | YAO_TEMP | YAO_TEMP | 113 | -0.429083 | -1.780486 | 0.000236 | 0.00517 | 0.004001 | 2513 | tags=42%, l | 581/29927 | C42-NCOR2 |
| YAO_TEMP | YAO_TEMP | YAO_TEMP | 123 | -0.438341 | -1.846339 | 0.000236 | 0.00517 | 0.004001 | 2269 | tags=41%, l | 51293/346 | C42-NCOR2 |
| REACTOME | REACTOME | REACTOME | 117 | -0.560553 | -2.340539 | 0.000237 | 0.00517 | 0.004001 | 2517 | tags=54%, l | 9167/5129 | C42-NCOR2 |
| KEGG_ALZI | KEGG_ALZI | KEGG_ALZI | 115 | -0.458027 | -1.906112 | 0.000237 | 0.00517 | 0.004001 | 1783 | tags=36%, l | 8772/6390 | C42-NCOR2 |
| KIM_ALL_D | KIM_ALL_D | KIM_ALL_D | 115 | -0.52283 | -2.175793 | 0.000237 | 0.00517 | 0.004001 | 2015 | tags=38%, l | 7167/4728 | C42-NCOR2 |
| PODAR_RE | PODAR_RE | PODAR_RE | 121 | -0.426055 | -1.786869 | 0.000238 | 0.00517 | 0.004001 | 1498 | tags=26%, l | 1051/1054 | C42-NCOR2 |
| REACTOME | REACTOME | REACTOME | 131 | -0.494532 | -2.10449 | 0.000238 | 0.00517 | 0.004001 | 3145 | tags=46%, l | 5189/2010 | C42-NCOR2 |
| REACTOME | REACTOME | REACTOME | 134 | -0.451818 | -1.928961 | 0.00024 | 0.00517 | 0.004001 | 1649 | tags=34%, l | 6194/6209 | C42-NCOR2 |
| SHEPARD_( | SHEPARD_( | SHEPARD_( | 142 | -0.435554 | -1.878736 | 0.000241 | 0.00517 | 0.004001 | 2583 | tags=41%, l | 8574/5743 | C42-NCOR2 |
| YAO_TEMP | YAO_TEMP | YAO_TEMP | 148 | -0.440692 | -1.910755 | 0.000241 | 0.00517 | 0.004001 | 2968 | tags=46%, l | 223/4706/ | C42-NCOR2 |
| LI_AMPLIFI | LI_AMPLIFI | LI_AMPLIFI | 141 | -0.466195 | -2.006794 | 0.000242 | 0.00517 | 0.004001 | 2005 | tags=40%, l | 10899/620 | C42-NCOR2 |
| REACTOME | REACTOME | REACTOME | 154 | -0.492505 | -2.144358 | 0.000242 | 0.00517 | 0.004001 | 1716 | tags=37%, l | 10885/574 | C42-NCOR2 |

| | | | | | | | | | | | | |
|---|---|---|---|---|---|---|---|---|---|---|---|---|
| HALLMARK | HALLMARK | HALLMARK | 153 | -0.45737 | -1.988404 | 0.000243 | 0.00517 | 0.004001 | 2220 | tags=41%, | 334/1738/ | C42-NCOR2 |
| REACTOME | REACTOME | REACTOME | 159 | -0.489627 | -2.143918 | 0.000244 | 0.00517 | 0.004001 | 1716 | tags=36%, | 10885/574 | C42-NCOR2 |
| HALLMARK | HALLMARK | HALLMARK | 180 | -0.4349 | -1.936147 | 0.000248 | 0.005197 | 0.004022 | 1991 | tags=35%, | 1595/1009 | C42-NCOR2 |
| TIEN_INTES | TIEN_INTES | TIEN_INTES | 179 | -0.486587 | -2.163645 | 0.000249 | 0.005197 | 0.004022 | 1906 | tags=34%, | 55154/647 | C42-NCOR2 |
| HALLMARK | HALLMARK | HALLMARK | 177 | -0.541811 | -2.405607 | 0.000249 | 0.005197 | 0.004022 | 2551 | tags=55%, | 2230/4695, | C42-NCOR2 |
| WONG_MI | WONG_MI | WONG_MI | 191 | -0.43243 | -1.937655 | 0.000253 | 0.005248 | 0.004062 | 2445 | tags=47%, | 9946/6038 | C42-NCOR2 |
| BLUM_RES | BLUM_RES | BLUM_RES | 210 | -0.440213 | -1.996622 | 0.000254 | 0.005257 | 0.004068 | 2889 | tags=46%, | 9616/8535, | C42-NCOR2 |
| ACEVEDO_ | ACEVEDO_ | ACEVEDO_ | 258 | -0.373718 | -1.732691 | 0.000258 | 0.005291 | 0.004094 | 2491 | tags=32%, | 23483/940 | C42-NCOR2 |
| REACTOME | REACTOME | REACTOME | 258 | -0.518003 | -2.40165 | 0.000258 | 0.005291 | 0.004094 | 2260 | tags=43%, | 6730/1975, | C42-NCOR2 |
| REACTOME | REACTOME | REACTOME | 276 | -0.484679 | -2.262005 | 0.000263 | 0.005304 | 0.004105 | 2041 | tags=41%, | 11224/299 | C42-NCOR2 |
| STEIN_ESRI | STEIN_ESRI | STEIN_ESRI | 300 | -0.369571 | -1.743616 | 0.000265 | 0.005304 | 0.004105 | 2395 | tags=35%, | 51295/316 | C42-NCOR2 |
| REACTOME | REACTOME | REACTOME | 319 | -0.32861 | -1.5595 | 0.000268 | 0.005304 | 0.004105 | 2799 | tags=34%, | 5695/1022, | C42-NCOR2 |
| SWEET_LU | SWEET_LU | SWEET_LU | 317 | -0.362337 | -1.717611 | 0.000269 | 0.005304 | 0.004105 | 2103 | tags=31%, | 1512/328/ | C42-NCOR2 |
| HSIAO_HO | HSIAO_HO | HSIAO_HO | 326 | -0.459125 | -2.1825 | 0.000269 | 0.005304 | 0.004105 | 1816 | tags=34%, | 1892/5663, | C42-NCOR2 |
| MOOTHA_ | MOOTHA_ | MOOTHA_ | 356 | -0.44653 | -2.142204 | 0.00027 | 0.005304 | 0.004105 | 3005 | tags=49%, | 10352/706, | C42-NCOR2 |
| MOOTHA_ | MOOTHA_ | MOOTHA_ | 331 | -0.371796 | -1.770747 | 0.000271 | 0.005304 | 0.004105 | 2384 | tags=34%, | 5019/6648, | C42-NCOR2 |
| FULCHER_I | FULCHER_I | FULCHER_I | 368 | -0.306241 | -1.473623 | 0.000272 | 0.005304 | 0.004105 | 2276 | tags=29%, | 2806/1090 | C42-NCOR2 |
| MOOTHA_ | MOOTHA_ | MOOTHA_ | 361 | -0.433824 | -2.082393 | 0.000273 | 0.005304 | 0.004105 | 2617 | tags=42%, | 64981/258 | C42-NCOR2 |
| STARK_PRE | STARK_PRE | STARK_PRE | 365 | -0.360829 | -1.734193 | 0.000273 | 0.005304 | 0.004105 | 1998 | tags=36%, | 9533/5128 | C42-NCOR2 |
| STEIN_ESRI | STEIN_ESRI | STEIN_ESRI | 403 | -0.298277 | -1.447989 | 0.000274 | 0.005304 | 0.004105 | 2198 | tags=28%, | 1738/7417, | C42-NCOR2 |
| ENK_UV_R | ENK_UV_R | ENK_UV_R | 391 | -0.313086 | -1.515462 | 0.000275 | 0.005304 | 0.004105 | 2821 | tags=34%, | 5742/5261, | C42-NCOR2 |
| SPIELMAN_ | SPIELMAN_ | SPIELMAN_ | 398 | -0.37888 | -1.836244 | 0.000275 | 0.005304 | 0.004105 | 2110 | tags=32%, | 476/7922/ | C42-NCOR2 |
| FLECHNER_ | FLECHNER_ | FLECHNER_ | 409 | -0.377555 | -1.833841 | 0.000278 | 0.005324 | 0.00412 | 2363 | tags=37%, | 6391/4043, | C42-NCOR2 |
| REACTOME | REACTOME | REACTOME | 524 | -0.349366 | -1.735651 | 0.000293 | 0.005564 | 0.004306 | 2789 | tags=35%, | 11056/102 | C42-NCOR2 |
| KRIGE_RES | KRIGE_RES | KRIGE_RES | 523 | -0.303698 | -1.508125 | 0.000293 | 0.005564 | 0.004306 | 1955 | tags=21%, | 23339/289 | C42-NCOR2 |
| KIM_BIPOL | KIM_BIPOL | KIM_BIPOL | 525 | -0.343613 | -1.706226 | 0.000294 | 0.005564 | 0.004306 | 2358 | tags=30%, | 4043/1212, | C42-NCOR2 |
| MARTENS_ | MARTENS_ | MARTENS_ | 575 | -0.380004 | -1.904229 | 0.000296 | 0.005564 | 0.004306 | 2094 | tags=30%, | 10376/101 | C42-NCOR2 |
| KIM_ALL_D | KIM_ALL_D | KIM_ALL_D | 581 | -0.36735 | -1.84197 | 0.000297 | 0.005564 | 0.004306 | 2341 | tags=32%, | 51157/512 | C42-NCOR2 |
| WAKABAYA | WAKABAYA | WAKABAYA | 591 | -0.301134 | -1.511923 | 0.000298 | 0.005568 | 0.004309 | 2551 | tags=28%, | 8672/1260 | C42-NCOR2 |
| KRIEG_HYP | KRIEG_HYP | KRIEG_HYP | 553 | 0.310702 | 1.488023 | 0.000302 | 0.00561 | 0.004341 | 2353 | tags=28%, | 5236/7431, | C42-NCOR2 |
| KRIGE_RES | KRIGE_RES | KRIGE_RES | 658 | -0.321972 | -1.630255 | 0.000306 | 0.005648 | 0.004371 | 2951 | tags=36%, | 2193/2840 | C42-NCOR2 |
| KRIGE_RES | KRIGE_RES | KRIGE_RES | 655 | -0.328326 | -1.661774 | 0.000307 | 0.005648 | 0.004371 | 2052 | tags=25%, | 160428/63 | C42-NCOR2 |
| LOPEZ_MB | LOPEZ_MB | LOPEZ_MB | 688 | -0.297187 | -1.511144 | 0.000309 | 0.005672 | 0.00439 | 2572 | tags=31%, | 23223/223 | C42-NCOR2 |
| KRIGE_RES | KRIGE_RES | KRIGE_RES | 737 | -0.342385 | -1.750247 | 0.000316 | 0.005767 | 0.004463 | 3226 | tags=41%, | 10471/706 | C42-NCOR2 |
| SENGUPTA | SENGUPTA | SENGUPTA | 254 | 0.350035 | 1.558096 | 0.000328 | 0.005966 | 0.004617 | 1218 | tags=19%, | 8653/1575 | C42-NCOR2 |
| REACTOME | REACTOME | REACTOME | 187 | 0.397867 | 1.710495 | 0.000334 | 0.006021 | 0.004659 | 2109 | tags=28%, | 2787/3012, | C42-NCOR2 |
| HALLMARK | HALLMARK | HALLMARK | 185 | 0.376657 | 1.617665 | 0.000334 | 0.006021 | 0.004659 | 1905 | tags=26%, | 54443/291 | C42-NCOR2 |
| PYEON_CA | PYEON_CA | PYEON_CA | 138 | 0.414598 | 1.712229 | 0.000343 | 0.006093 | 0.004715 | 1753 | tags=28%, | 11073/307 | C42-NCOR2 |
| PEREZ_TP5 | PEREZ_TP5 | PEREZ_TP5 | 137 | 0.421439 | 1.738623 | 0.000343 | 0.006093 | 0.004715 | 2231 | tags=32%, | 10570/360, | C42-NCOR2 |
| REACTOME | REACTOME | REACTOME | 112 | 0.443217 | 1.775554 | 0.000348 | 0.006093 | 0.004715 | 2109 | tags=29%, | 3012/8335, | C42-NCOR2 |
| CHEMNITZ_ | CHEMNITZ_ | CHEMNITZ_ | 109 | 0.47294 | 1.888597 | 0.000348 | 0.006093 | 0.004715 | 1902 | tags=38%, | 4173/1061 | C42-NCOR2 |
| RUIZ_TNC_ | RUIZ_TNC_ | RUIZ_TNC_ | 107 | 0.455849 | 1.814727 | 0.000349 | 0.006093 | 0.004715 | 1400 | tags=32%, | 4173/5587 | C42-NCOR2 |
| REACTOME | REACTOME | REACTOME | 84 | 0.490207 | 1.87918 | 0.000353 | 0.006093 | 0.004715 | 2334 | tags=38%, | 3012/8335, | C42-NCOR2 |
| REACTOME | REACTOME | REACTOME | 96 | 0.494683 | 1.939944 | 0.000354 | 0.006093 | 0.004715 | 2938 | tags=49%, | 64946/921 | C42-NCOR2 |
| VERNELL_R | VERNELL_R | VERNELL_R | 60 | 0.564043 | 2.041985 | 0.000356 | 0.006093 | 0.004715 | 3146 | tags=60%, | 157570/41 | C42-NCOR2 |
| REACTOME | REACTOME | REACTOME | 61 | 0.559686 | 2.032184 | 0.000357 | 0.006093 | 0.004715 | 2109 | tags=43%, | 3012/8335, | C42-NCOR2 |
| SU_TESTIS | SU_TESTIS | SU_TESTIS | 50 | 0.570115 | 1.991401 | 0.000357 | 0.006093 | 0.004715 | 2820 | tags=56%, | 11073/514 | C42-NCOR2 |
| REACTOME | REACTOME | REACTOME | 45 | 0.602746 | 2.057525 | 0.000357 | 0.006093 | 0.004715 | 2055 | tags=42%, | 3012/8335, | C42-NCOR2 |
| FURUKAWA | FURUKAWA | FURUKAWA | 56 | 0.581989 | 2.074486 | 0.000357 | 0.006093 | 0.004715 | 2629 | tags=59%, | 4173/5587 | C42-NCOR2 |
| REACTOME | REACTOME | REACTOME | 72 | 0.536987 | 2.00356 | 0.000358 | 0.006093 | 0.004715 | 2055 | tags=33%, | 3012/8335, | C42-NCOR2 |
| REACTOME | REACTOME | REACTOME | 42 | 0.592764 | 1.995408 | 0.000358 | 0.006093 | 0.004715 | 2337 | tags=45%, | 7431/3009, | C42-NCOR2 |
| GREENBAU | GREENBAU | GREENBAU | 30 | 0.622485 | 1.949895 | 0.000364 | 0.006142 | 0.004753 | 2257 | tags=50%, | 54443/291 | C42-NCOR2 |
| YU_MYC_T | YU_MYC_T | YU_MYC_T | 35 | 0.600893 | 1.941723 | 0.000366 | 0.006142 | 0.004753 | 1872 | tags=46%, | 54443/291 | C42-NCOR2 |
| REACTOME | REACTOME | REACTOME | 29 | 0.64046 | 1.986083 | 0.000367 | 0.006142 | 0.004753 | 1850 | tags=41%, | 4173/5982, | C42-NCOR2 |
| BIOCARTA_ | BIOCARTA_ | BIOCARTA_ | 33 | 0.610471 | 1.948085 | 0.000367 | 0.006142 | 0.004753 | 1561 | tags=36%, | 5567/5295, | C42-NCOR2 |
| NIKOLSKY_ | NIKOLSKY_ | NIKOLSKY_ | 20 | 0.70551 | 2.002304 | 0.000374 | 0.006167 | 0.004772 | 1216 | tags=40%, | 390927/92 | C42-NCOR2 |
| PETROVA_I | PETROVA_I | PETROVA_I | 21 | 0.711581 | 2.043021 | 0.000375 | 0.006167 | 0.004772 | 1987 | tags=62%, | 23089/715 | C42-NCOR2 |
| SCIAN_CEL | SCIAN_CEL | SCIAN_CEL | 21 | 0.722901 | 2.075521 | 0.000375 | 0.006167 | 0.004772 | 1974 | tags=76%, | 22974/417 | C42-NCOR2 |
| LY_AGING_ | LY_AGING_ | LY_AGING_ | 16 | 0.757462 | 2.038947 | 0.000376 | 0.006167 | 0.004772 | 2189 | tags=81%, | 3832/1106 | C42-NCOR2 |
| REACTOME | REACTOME | REACTOME | 16 | 0.77439 | 2.084514 | 0.000376 | 0.006167 | 0.004772 | 1819 | tags=69%, | 993/4605/ | C42-NCOR2 |
| REACTOME | REACTOME | REACTOME | 38 | -0.609043 | -2.065707 | 0.000448 | 0.00732 | 0.005665 | 4235 | tags=76%, | 84128/259 | C42-NCOR2 |
| ZHAN_MUI | ZHAN_MUI | ZHAN_MUI | 41 | -0.607684 | -2.091164 | 0.000452 | 0.007324 | 0.005668 | 2050 | tags=54%, | 3033/2995 | C42-NCOR2 |
| MACAEVA_ | MACAEVA_ | MACAEVA_ | 73 | -0.485452 | -1.87817 | 0.000452 | 0.007324 | 0.005668 | 1885 | tags=47%, | 196513/16 | C42-NCOR2 |
| TAKAO_RE | TAKAO_RE | TAKAO_RE | 72 | -0.491712 | -1.894634 | 0.000453 | 0.007324 | 0.005668 | 2320 | tags=44%, | 667/10482, | C42-NCOR2 |
| SHAFFER_II | SHAFFER_II | SHAFFER_II | 84 | -0.465772 | -1.843081 | 0.000461 | 0.007419 | 0.005741 | 2022 | tags=30%, | 9777/1595, | C42-NCOR2 |
| MULLIGHA | MULLIGHA | MULLIGHA | 89 | -0.432761 | -1.726663 | 0.000463 | 0.007422 | 0.005743 | 1888 | tags=31%, | 5366/586/ | C42-NCOR2 |

| | | | | | | | | | | | | |
|---|---|---|---|---|---|---|---|---|---|---|---|---|
| HALLMARK | HALLMARK | HALLMARK | 134 | -0.431642 | -1.842823 | 0.000479 | 0.007586 | 0.005871 | 1839 | tags=36%, | 9564/629/ | C42-NCOR2 |
| KEGG_HUN | KEGG_HUN | KEGG_HUN | 134 | -0.404568 | -1.727233 | 0.000479 | 0.007586 | 0.005871 | 2517 | tags=42%, | 9167/581/ | C42-NCOR2 |
| MCBRYAN_ | MCBRYAN_ | MCBRYAN_ | 133 | -0.411696 | -1.754449 | 0.00048 | 0.007586 | 0.005871 | 1492 | tags=31%, | 590/31/10 | C42-NCOR2 |
| GRUETZMA | GRUETZMA | GRUETZMA | 135 | -0.411621 | -1.757458 | 0.000481 | 0.007586 | 0.005871 | 1986 | tags=35%, | 6204/6296 | C42-NCOR2 |
| REACTOME | REACTOME | REACTOME | 145 | -0.422444 | -1.825017 | 0.000483 | 0.007586 | 0.005871 | 2789 | tags=43%, | 5689/6223 | C42-NCOR2 |
| BURTON_A | BURTON_A | BURTON_A | 144 | -0.4245 | -1.832444 | 0.000484 | 0.007586 | 0.005871 | 2422 | tags=39%, | 4257/6391 | C42-NCOR2 |
| KOYAMA_S | KOYAMA_S | KOYAMA_S | 156 | -0.387648 | -1.690714 | 0.000488 | 0.007592 | 0.005876 | 1513 | tags=19%, | 10284/384 | C42-NCOR2 |
| LINDGREN_ | LINDGREN_ | LINDGREN_ | 159 | -0.409259 | -1.792011 | 0.000488 | 0.007592 | 0.005876 | 2463 | tags=33%, | 595/11214 | C42-NCOR2 |
| HALLMARK | HALLMARK | HALLMARK | 164 | -0.368432 | -1.619441 | 0.000492 | 0.0076 | 0.005881 | 2509 | tags=40%, | 3162/5100 | C42-NCOR2 |
| JIANG_HYP | JIANG_HYP | JIANG_HYP | 164 | -0.380443 | -1.672239 | 0.000492 | 0.0076 | 0.005881 | 1792 | tags=28%, | 4907/4598 | C42-NCOR2 |
| HALLMARK | HALLMARK | HALLMARK | 172 | -0.376755 | -1.666763 | 0.000495 | 0.007619 | 0.005896 | 2582 | tags=38%, | 3326/5781 | C42-NCOR2 |
| FORTSCHE | FORTSCHE | FORTSCHE | 191 | -0.376507 | -1.68707 | 0.000505 | 0.007684 | 0.005947 | 1402 | tags=27%, | 7855/5529 | C42-NCOR2 |
| FISCHER_G | FISCHER_G | FISCHER_G | 164 | 0.391964 | 1.659355 | 0.000505 | 0.007684 | 0.005947 | 1990 | tags=27%, | 11073/307 | C42-NCOR2 |
| WAMUNYO | WAMUNYO | WAMUNYO | 200 | -0.381054 | -1.719405 | 0.000505 | 0.007684 | 0.005947 | 1699 | tags=24%, | 9092/9145 | C42-NCOR2 |
| NIKOLSKY_ | NIKOLSKY_ | NIKOLSKY_ | 197 | -0.373712 | -1.681776 | 0.000506 | 0.007684 | 0.005947 | 2977 | tags=39%, | 10513/574 | C42-NCOR2 |
| ACEVEDO_ | ACEVEDO_ | ACEVEDO_ | 105 | 0.432721 | 1.718904 | 0.000523 | 0.007888 | 0.006104 | 2689 | tags=37%, | 3012/5444 | C42-NCOR2 |
| TAKEDA_T | TAKEDA_T | TAKEDA_T | 97 | 0.445483 | 1.749702 | 0.000529 | 0.007888 | 0.006104 | 1479 | tags=27%, | 4173/3899 | C42-NCOR2 |
| REACTOME | REACTOME | REACTOME | 79 | 0.498552 | 1.889783 | 0.00053 | 0.007888 | 0.006104 | 2300 | tags=38%, | 3012/8335 | C42-NCOR2 |
| RIGGI_EWI | RIGGI_EWI | RIGGI_EWI | 95 | 0.485467 | 1.9004 | 0.000531 | 0.007888 | 0.006104 | 2254 | tags=41%, | 54443/558 | C42-NCOR2 |
| MORI_IMM | MORI_IMM | MORI_IMM | 78 | 0.496625 | 1.877449 | 0.000532 | 0.007888 | 0.006104 | 1836 | tags=42%, | 3012/7153 | C42-NCOR2 |
| PETROVA_I | PETROVA_I | PETROVA_I | 92 | 0.492339 | 1.916237 | 0.000532 | 0.007888 | 0.006104 | 2157 | tags=50%, | 4885/5468 | C42-NCOR2 |
| KAUFFMAN | KAUFFMAN | KAUFFMAN | 55 | 0.563729 | 2.007154 | 0.000533 | 0.007888 | 0.006104 | 1782 | tags=35%, | 4173/7019 | C42-NCOR2 |
| ZHAN_MUI | ZHAN_MUI | ZHAN_MUI | 32 | 0.613301 | 1.943449 | 0.000551 | 0.008126 | 0.006288 | 2215 | tags=47%, | 5087/1047 | C42-NCOR2 |
| REICHERT_ | REICHERT_ | REICHERT_ | 25 | 0.678903 | 2.035157 | 0.000554 | 0.008129 | 0.006291 | 2629 | tags=84%, | 157313/71 | C42-NCOR2 |
| REACTOME | REACTOME | REACTOME | 24 | 0.675771 | 2.006524 | 0.000556 | 0.008129 | 0.006291 | 2294 | tags=50%, | 2787/5567 | C42-NCOR2 |
| BIOCARTA_ | BIOCARTA_ | BIOCARTA_ | 19 | 0.697973 | 1.958475 | 0.000561 | 0.008129 | 0.006291 | 1466 | tags=47%, | 5567/5295 | C42-NCOR2 |
| BIOCARTA_ | BIOCARTA_ | BIOCARTA_ | 19 | 0.697294 | 1.95657 | 0.000561 | 0.008129 | 0.006291 | 1561 | tags=47%, | 5295/5530 | C42-NCOR2 |
| REACTOME | REACTOME | REACTOME | 20 | 0.694617 | 1.97139 | 0.000561 | 0.008129 | 0.006291 | 2825 | tags=65%, | 6925/1499 | C42-NCOR2 |
| PARENT_M | PARENT_M | PARENT_M | 421 | -0.29336 | -1.427256 | 0.000563 | 0.008129 | 0.006291 | 2645 | tags=29%, | 79758/542 | C42-NCOR2 |
| MONTERO_ | MONTERO_ | MONTERO_ | 12 | 0.798026 | 1.986975 | 0.000571 | 0.008215 | 0.006358 | 1080 | tags=75%, | 54443/783 | C42-NCOR2 |
| SMID_BRE | SMID_BRE | SMID_BRE | 386 | 0.316026 | 1.467314 | 0.000629 | 0.00902 | 0.006981 | 1724 | tags=24%, | 4885/4173 | C42-NCOR2 |
| GABRIELY_ | GABRIELY_ | GABRIELY_ | 231 | 0.351642 | 1.551124 | 0.000655 | 0.00936 | 0.007243 | 2817 | tags=33%, | 63934/551 | C42-NCOR2 |
| GENTILE_U | GENTILE_U | GENTILE_U | 233 | 0.355495 | 1.567756 | 0.000657 | 0.00936 | 0.007243 | 1994 | tags=25%, | 11073/795 | C42-NCOR2 |
| NATSUME_ | NATSUME_ | NATSUME_ | 36 | -0.606671 | -2.033016 | 0.000664 | 0.009437 | 0.007303 | 774 | tags=39%, | 4792/5052 | C42-NCOR2 |
| REACTOME | REACTOME | REACTOME | 173 | 0.383632 | 1.635752 | 0.00067 | 0.00949 | 0.007344 | 2211 | tags=28%, | 3012/8335 | C42-NCOR2 |
| TIEN_INTE | TIEN_INTE | TIEN_INTE | 40 | -0.566643 | -1.940361 | 0.000677 | 0.009551 | 0.007391 | 1816 | tags=42%, | 6194/8073 | C42-NCOR2 |
| LI_WILMS_ | LI_WILMS_ | LI_WILMS_ | 134 | 0.410144 | 1.687124 | 0.000686 | 0.009648 | 0.007467 | 2094 | tags=37%, | 11073/106 | C42-NCOR2 |
| ALCALAY_A | ALCALAY_A | ALCALAY_A | 128 | 0.42294 | 1.727573 | 0.000691 | 0.009678 | 0.007489 | 2269 | tags=35%, | 9185/4173 | C42-NCOR2 |
| REACTOME | REACTOME | REACTOME | 112 | 0.424506 | 1.700595 | 0.000695 | 0.009708 | 0.007513 | 2711 | tags=37%, | 5567/2912 | C42-NCOR2 |
| HALLMARK | HALLMARK | HALLMARK | 132 | -0.398957 | -1.699884 | 0.000715 | 0.009876 | 0.007643 | 2352 | tags=38%, | 6391/3422 | C42-NCOR2 |
| SCHLOSSER | SCHLOSSER | SCHLOSSER | 132 | -0.402616 | -1.715474 | 0.000715 | 0.009876 | 0.007643 | 1533 | tags=25%, | 7307/1051 | C42-NCOR2 |
| REACTOME | REACTOME | REACTOME | 70 | 0.502847 | 1.869401 | 0.000716 | 0.009876 | 0.007643 | 1405 | tags=30%, | 3012/8335 | C42-NCOR2 |
| REACTOME | REACTOME | REACTOME | 69 | 0.504364 | 1.86999 | 0.000716 | 0.009876 | 0.007643 | 1405 | tags=28%, | 3012/2355 | C42-NCOR2 |
| REACTOME | REACTOME | REACTOME | 27 | 0.645303 | 1.966175 | 0.000737 | 0.010114 | 0.007827 | 2294 | tags=44%, | 2787/5567 | C42-NCOR2 |
| REACTOME | REACTOME | REACTOME | 25 | 0.648506 | 1.944036 | 0.000739 | 0.010114 | 0.007827 | 1193 | tags=40%, | 7153/9134 | C42-NCOR2 |
| PID_EPHRI | PID_EPHRI | PID_EPHRI | 20 | 0.684854 | 1.943679 | 0.000748 | 0.01021 | 0.007901 | 1690 | tags=35%, | 5295/2048 | C42-NCOR2 |
| BIOCARTA_ | BIOCARTA_ | BIOCARTA_ | 12 | 0.778729 | 1.938929 | 0.000761 | 0.010353 | 0.008012 | 1466 | tags=50%, | 5567/2323 | C42-NCOR2 |
| GARY_CD5 | GARY_CD5 | GARY_CD5 | 374 | -0.302142 | -1.455458 | 0.000822 | 0.01115 | 0.008629 | 1874 | tags=22%, | 29780/687 | C42-NCOR2 |
| ZHONG_RE | ZHONG_RE | ZHONG_RE | 121 | 0.413368 | 1.677797 | 0.000862 | 0.011648 | 0.009014 | 2125 | tags=30%, | 9185/1105 | C42-NCOR2 |
| LUI_TARGE | LUI_TARGE | LUI_TARGE | 24 | -0.680244 | -2.0827 | 0.000868 | 0.0117 | 0.009055 | 1592 | tags=50%, | 10430/567 | C42-NCOR2 |
| ELLWOOD_ | ELLWOOD_ | ELLWOOD_ | 28 | -0.627363 | -1.993281 | 0.000877 | 0.011711 | 0.009062 | 1958 | tags=50%, | 9802/5847 | C42-NCOR2 |
| ONGUSAH | ONGUSAH | ONGUSAH | 28 | -0.628798 | -1.99784 | 0.000877 | 0.011711 | 0.009062 | 2220 | tags=57%, | 1512/2052 | C42-NCOR2 |
| TOMLINS_I | TOMLINS_I | TOMLINS_I | 33 | -0.624674 | -2.055846 | 0.000879 | 0.011711 | 0.009062 | 1422 | tags=45%, | 5987/1060 | C42-NCOR2 |
| HALLMARK | HALLMARK | HALLMARK | 93 | 0.444969 | 1.735891 | 0.000884 | 0.011711 | 0.009062 | 1650 | tags=33%, | 3817/8555 | C42-NCOR2 |
| LIU_VAV3_ | LIU_VAV3_ | LIU_VAV3_ | 44 | 0.552264 | 1.878784 | 0.000889 | 0.011711 | 0.009062 | 2060 | tags=43%, | 354/5468/ | C42-NCOR2 |
| BOYLAN_M | BOYLAN_M | BOYLAN_M | 52 | 0.545283 | 1.920788 | 0.000891 | 0.011711 | 0.009062 | 1896 | tags=37%, | 7227/1076 | C42-NCOR2 |
| REN_BOUN | REN_BOUN | REN_BOUN | 57 | 0.526227 | 1.88379 | 0.000893 | 0.011711 | 0.009062 | 3400 | tags=58%, | 7153/4292 | C42-NCOR2 |
| MORI_LAR | MORI_LAR | MORI_LAR | 74 | 0.479454 | 1.797239 | 0.000893 | 0.011711 | 0.009062 | 3226 | tags=55%, | 3012/5885 | C42-NCOR2 |
| YAO_HOXA | YAO_HOXA | YAO_HOXA | 43 | 0.575184 | 1.945949 | 0.000894 | 0.011711 | 0.009062 | 1953 | tags=44%, | 5295/1397 | C42-NCOR2 |
| KEGG_ARG | KEGG_ARG | KEGG_ARG | 41 | -0.564273 | -1.941778 | 0.000903 | 0.01176 | 0.009101 | 1501 | tags=44%, | 6723/2805 | C42-NCOR2 |
| ZHOU_TNF | ZHOU_TNF | ZHOU_TNF | 41 | -0.565168 | -1.944858 | 0.000903 | 0.01176 | 0.009101 | 1630 | tags=46%, | 4691/2593 | C42-NCOR2 |
| CUI_TCF21 | CUI_TCF21 | CUI_TCF21 | 563 | 0.295647 | 1.417583 | 0.000906 | 0.011761 | 0.009102 | 2977 | tags=33%, | 7431/7544 | C42-NCOR2 |
| REACTOME | REACTOME | REACTOME | 48 | -0.540464 | -1.91738 | 0.000911 | 0.011779 | 0.009116 | 2454 | tags=54%, | 617/6901/ | C42-NCOR2 |
| PID_AURO | PID_AURO | PID_AURO | 35 | 0.575968 | 1.861179 | 0.000916 | 0.011779 | 0.009116 | 2303 | tags=57%, | 7431/2912 | C42-NCOR2 |
| REACTOME | REACTOME | REACTOME | 91 | -0.418346 | -1.679051 | 0.000916 | 0.011779 | 0.009116 | 2676 | tags=44%, | 64981/123 | C42-NCOR2 |
| HALLMARK | HALLMARK | HALLMARK | 79 | -0.454737 | -1.777296 | 0.000921 | 0.011811 | 0.00914 | 1168 | tags=27%, | 189/5191/ | C42-NCOR2 |

| | | | | | | | | | | | | |
|---|---|---|---|---|---|---|---|---|---|---|---|---|
| BIOCARTA_ | BIOCARTA_ | BIOCARTA_ | 17 | 0.708527 | 1.936058 | 0.000941 | 0.012027 | 0.009307 | 1466 | tags=47%, l | 5567/5563, | C42-NCOR2 |
| KIM_MYC_ | KIM_MYC_ | KIM_MYC_ | 138 | -0.389651 | -1.672579 | 0.00096 | 0.012232 | 0.009466 | 3327 | tags=50%, l | 51116/600 | C42-NCOR2 |
| KEGG_PATI | KEGG_PATI | KEGG_PATI | 237 | 0.34545 | 1.526421 | 0.000985 | 0.012517 | 0.009687 | 2193 | tags=28%, l | 354/5295/5 | C42-NCOR2 |
| YOSHIMUR | YOSHIMUR | YOSHIMUR | 247 | -0.331815 | -1.532803 | 0.001023 | 0.01294 | 0.010014 | 2045 | tags=27%, l | 10923/109 | C42-NCOR2 |
| FISCHER_D | FISCHER_D | FISCHER_D | 229 | -0.335006 | -1.533635 | 0.001026 | 0.01294 | 0.010014 | 1885 | tags=24%, l | 5366/5531 | C42-NCOR2 |
| STARK_PRE | STARK_PRE | STARK_PRE | 139 | 0.393418 | 1.627576 | 0.001028 | 0.01294 | 0.010014 | 2733 | tags=35%, l | 6925/5295, | C42-NCOR2 |
| KEGG_PEN | KEGG_PEN | KEGG_PEN | 13 | -0.759219 | -1.963524 | 0.001054 | 0.013152 | 0.010178 | 988 | tags=54%, l | 51181/107 | C42-NCOR2 |
| NIKOLSKY_ | NIKOLSKY_ | NIKOLSKY_ | 12 | -0.769438 | -1.943214 | 0.001054 | 0.013152 | 0.010178 | 2590 | tags=83%, l | 104/1292/8 | C42-NCOR2 |
| KEGG_OOC | KEGG_OOC | KEGG_OOC | 87 | 0.440551 | 1.700378 | 0.001054 | 0.013152 | 0.010178 | 1973 | tags=33%, l | 5567/817/9 | C42-NCOR2 |
| LY_AGING_ | LY_AGING_ | LY_AGING_ | 47 | 0.549087 | 1.888746 | 0.00107 | 0.013287 | 0.010283 | 2303 | tags=49%, l | 9928/3832 | C42-NCOR2 |
| REACTOME | REACTOME | REACTOME | 51 | 0.539826 | 1.892772 | 0.001071 | 0.013287 | 0.010283 | 1912 | tags=41%, l | 8342/8343, | C42-NCOR2 |
| HU_GENOT | HU_GENOT | HU_GENOT | 25 | -0.642243 | -1.982147 | 0.00109 | 0.013486 | 0.010437 | 1501 | tags=36%, l | 3149/9709, | C42-NCOR2 |
| BIOCARTA_ | BIOCARTA_ | BIOCARTA_ | 20 | 0.666978 | 1.892948 | 0.001122 | 0.013834 | 0.010705 | 1869 | tags=65%, l | 993/1031/1 | C42-NCOR2 |
| LI_WILMS_ | LI_WILMS_ | LI_WILMS_ | 15 | 0.721131 | 1.908919 | 0.001129 | 0.013834 | 0.010705 | 2189 | tags=67%, l | 5984/1063 | C42-NCOR2 |
| PEDERSEN_ | PEDERSEN_ | PEDERSEN_ | 285 | 0.332049 | 1.498761 | 0.001129 | 0.013834 | 0.010705 | 2243 | tags=31%, l | 4173/5865, | C42-NCOR2 |
| CHNG_MU | CHNG_MU | CHNG_MU | 42 | -0.545442 | -1.884613 | 0.001131 | 0.013834 | 0.010705 | 2005 | tags=48%, l | 9960/6194 | C42-NCOR2 |
| BHATI_G2N | BHATI_G2N | BHATI_G2N | 86 | -0.425941 | -1.691029 | 0.001156 | 0.014087 | 0.010901 | 2557 | tags=35%, l | 4636/6164, | C42-NCOR2 |
| WAKABAYA | WAKABAYA | WAKABAYA | 108 | -0.410382 | -1.692721 | 0.001172 | 0.014193 | 0.010983 | 1198 | tags=22%, l | 10961/551 | C42-NCOR2 |
| REACTOME | REACTOME | REACTOME | 170 | 0.3739 | 1.591538 | 0.001173 | 0.014193 | 0.010983 | 2872 | tags=36%, l | 5885/8461 | C42-NCOR2 |
| BIDUS_MET | BIDUS_MET | BIDUS_MET | 172 | 0.368266 | 1.569201 | 0.001174 | 0.014193 | 0.010983 | 3025 | tags=36%, l | 55872/291 | C42-NCOR2 |
| AFFAR_YY1 | AFFAR_YY1 | AFFAR_YY1 | 169 | 0.376131 | 1.599088 | 0.001178 | 0.0142 | 0.010989 | 2213 | tags=36%, l | 7404/4173, | C42-NCOR2 |
| PLASARI_T | PLASARI_T | PLASARI_T | 157 | 0.382586 | 1.609888 | 0.001187 | 0.014262 | 0.011037 | 2362 | tags=39%, l | 7227/9185, | C42-NCOR2 |
| KOINUMA_ | KOINUMA_ | KOINUMA_ | 624 | 0.286246 | 1.384454 | 0.001195 | 0.014321 | 0.011083 | 2351 | tags=28%, l | 7431/2787, | C42-NCOR2 |
| WONG_AD | WONG_AD | WONG_AD | 447 | 0.302743 | 1.425802 | 0.001234 | 0.014731 | 0.0114 | 2213 | tags=28%, l | 10769/556 | C42-NCOR2 |
| REACTOME | REACTOME | REACTOME | 84 | 0.448435 | 1.719053 | 0.001236 | 0.014731 | 0.0114 | 2300 | tags=33%, l | 3012/8335, | C42-NCOR2 |
| REACTOME | REACTOME | REACTOME | 95 | 0.425694 | 1.666413 | 0.00124 | 0.014734 | 0.011402 | 2179 | tags=31%, l | 3012/8335, | C42-NCOR2 |
| CROONQUI | CROONQUI | CROONQUI | 67 | 0.479739 | 1.769301 | 0.001254 | 0.014855 | 0.011496 | 3438 | tags=61%, l | 6192/2297 | C42-NCOR2 |
| RUTELLA_R | RUTELLA_R | RUTELLA_R | 237 | -0.324669 | -1.495299 | 0.001278 | 0.015079 | 0.01167 | 1687 | tags=22%, l | 10134/115 | C42-NCOR2 |
| REACTOME | REACTOME | REACTOME | 29 | 0.596631 | 1.850169 | 0.001283 | 0.015079 | 0.01167 | 3760 | tags=62%, l | 4173/5982, | C42-NCOR2 |
| KEGG_GLY | KEGG_GLY | KEGG_GLY | 21 | -0.668674 | -1.979351 | 0.001287 | 0.015079 | 0.01167 | 3114 | tags=71%, l | 2731/6470, | C42-NCOR2 |
| REACTOME | REACTOME | REACTOME | 21 | -0.678142 | -2.007377 | 0.001287 | 0.015079 | 0.01167 | 790 | tags=38%, l | 8501/8140, | C42-NCOR2 |
| HORTON_S | HORTON_S | HORTON_S | 23 | -0.662046 | -2.006825 | 0.001296 | 0.015148 | 0.011723 | 2304 | tags=57%, l | 10654/159 | C42-NCOR2 |
| REACTOME | REACTOME | REACTOME | 16 | 0.693828 | 1.867657 | 0.001315 | 0.015306 | 0.011845 | 2058 | tags=62%, l | 2787/784/9 | C42-NCOR2 |
| REACTOME | REACTOME | REACTOME | 18 | 0.671785 | 1.858724 | 0.001321 | 0.015306 | 0.011845 | 2656 | tags=61%, l | 7404/8354, | C42-NCOR2 |
| SCHAEFFER | SCHAEFFER | SCHAEFFER | 197 | 0.351887 | 1.523983 | 0.001322 | 0.015306 | 0.011845 | 3070 | tags=39%, l | 5865/6925, | C42-NCOR2 |
| SCHLOSSER | SCHLOSSER | SCHLOSSER | 31 | -0.613843 | -1.99342 | 0.001325 | 0.015306 | 0.011845 | 3289 | tags=68%, l | 9112/2340 | C42-NCOR2 |
| REACTOME | REACTOME | REACTOME | 14 | 0.733537 | 1.898655 | 0.001329 | 0.015314 | 0.011851 | 1450 | tags=36%, l | 7431/7171, | C42-NCOR2 |
| BENPORAT | BENPORAT | BENPORAT | 188 | 0.352269 | 1.515144 | 0.001332 | 0.015314 | 0.011851 | 1896 | tags=26%, l | 7227/8555, | C42-NCOR2 |
| LIM_MAMI | LIM_MAMI | LIM_MAMI | 325 | -0.310507 | -1.474643 | 0.001349 | 0.015464 | 0.011967 | 1918 | tags=26%, l | 11022/551 | C42-NCOR2 |
| RODRIGUES | RODRIGUES | RODRIGUES | 560 | 0.289474 | 1.386645 | 0.001367 | 0.015625 | 0.012092 | 2252 | tags=26%, l | 10769/417 | C42-NCOR2 |
| REACTOME | REACTOME | REACTOME | 132 | 0.395616 | 1.62264 | 0.001378 | 0.015705 | 0.012154 | 1990 | tags=28%, l | 4173/7153, | C42-NCOR2 |
| SONG_TAR | SONG_TAR | SONG_TAR | 50 | 0.526613 | 1.83945 | 0.001427 | 0.016192 | 0.012531 | 1400 | tags=28%, l | 11073/701 | C42-NCOR2 |
| TURASHVIL | TURASHVIL | TURASHVIL | 51 | 0.505591 | 1.772735 | 0.001428 | 0.016192 | 0.012531 | 1255 | tags=33%, l | 54443/551 | C42-NCOR2 |
| BILD_MYC_ | BILD_MYC_ | BILD_MYC_ | 149 | -0.362808 | -1.575726 | 0.001438 | 0.016259 | 0.012583 | 3109 | tags=41%, l | 617/6832/6 | C42-NCOR2 |
| PID_CDC42 | PID_CDC42 | PID_CDC42 | 62 | 0.493182 | 1.791749 | 0.001445 | 0.016295 | 0.01261 | 2825 | tags=44%, l | 5295/1499, | C42-NCOR2 |
| DEMAGALH | DEMAGALH | DEMAGALH | 10 | -0.775666 | -1.867617 | 0.001459 | 0.016385 | 0.01268 | 1053 | tags=50%, l | 7037/2706 | C42-NCOR2 |
| KEGG_GLY | KEGG_GLY | KEGG_GLY | 11 | -0.764786 | -1.885748 | 0.001462 | 0.016385 | 0.01268 | 815 | tags=91%, l | 50/48/419 | C42-NCOR2 |
| SUNG_MET | SUNG_MET | SUNG_MET | 35 | 0.559585 | 1.808239 | 0.001465 | 0.016385 | 0.01268 | 2851 | tags=51%, l | 3070/2009, | C42-NCOR2 |
| REACTOME | REACTOME | REACTOME | 34 | 0.562184 | 1.804461 | 0.00147 | 0.016385 | 0.01268 | 1782 | tags=35%, l | 11073/598 | C42-NCOR2 |
| HERNANDE | HERNANDE | HERNANDE | 26 | 0.620499 | 1.874629 | 0.001472 | 0.016385 | 0.01268 | 3451 | tags=69%, l | 11065/593 | C42-NCOR2 |
| WILLIAMS_ | WILLIAMS_ | WILLIAMS_ | 23 | 0.642899 | 1.890988 | 0.001489 | 0.016524 | 0.012788 | 1729 | tags=61%, l | 1836/8161 | C42-NCOR2 |
| REACTOME | REACTOME | REACTOME | 158 | 0.379406 | 1.598049 | 0.001523 | 0.016862 | 0.013049 | 2672 | tags=35%, l | 5885/5468, | C42-NCOR2 |
| KEGG_CITR | KEGG_CITR | KEGG_CITR | 28 | -0.617005 | -1.96037 | 0.001534 | 0.01692 | 0.013094 | 2351 | tags=57%, l | 1738/48/6 | C42-NCOR2 |
| HOLLMANN | HOLLMANN | HOLLMANN | 146 | 0.388724 | 1.619355 | 0.001536 | 0.01692 | 0.013094 | 2384 | tags=36%, l | 8653/6192, | C42-NCOR2 |
| REACTOME | REACTOME | REACTOME | 139 | 0.389375 | 1.610847 | 0.001542 | 0.016934 | 0.013105 | 1990 | tags=25%, l | 157570/41 | C42-NCOR2 |
| BILANGES_ | BILANGES_ | BILANGES_ | 37 | -0.564664 | -1.906925 | 0.001547 | 0.016948 | 0.013116 | 1586 | tags=43%, l | 6193/6207, | C42-NCOR2 |
| REACTOME | REACTOME | REACTOME | 117 | 0.403953 | 1.630485 | 0.001559 | 0.017033 | 0.013181 | 2334 | tags=31%, l | 3012/8335, | C42-NCOR2 |
| WHITFIELD | WHITFIELD | WHITFIELD | 114 | 0.403291 | 1.620318 | 0.001564 | 0.017051 | 0.013195 | 1782 | tags=29%, l | 157570/30 | C42-NCOR2 |
| SHAFFER_I | SHAFFER_I | SHAFFER_I | 67 | -0.456525 | -1.736596 | 0.001584 | 0.017221 | 0.013327 | 1751 | tags=34%, l | 4598/7416, | C42-NCOR2 |
| REACTOME | REACTOME | REACTOME | 47 | 0.519218 | 1.786004 | 0.001605 | 0.017402 | 0.013467 | 1912 | tags=36%, l | 8342/8343, | C42-NCOR2 |
| LEE_NEUR | LEE_NEUR | LEE_NEUR | 73 | 0.459813 | 1.719236 | 0.001613 | 0.017441 | 0.013497 | 1693 | tags=33%, l | 10769/110 | C42-NCOR2 |
| DANG_MY | DANG_MY | DANG_MY | 104 | -0.399197 | -1.634611 | 0.001636 | 0.017648 | 0.013657 | 2751 | tags=42%, l | 6470/6223, | C42-NCOR2 |
| PUJANA_CI | PUJANA_CI | PUJANA_CI | 607 | 0.286551 | 1.381755 | 0.001654 | 0.017793 | 0.01377 | 2340 | tags=24%, l | 4173/1107 | C42-NCOR2 |
| REACTOME | REACTOME | REACTOME | 162 | 0.372131 | 1.573185 | 0.001686 | 0.018096 | 0.014004 | 2102 | tags=28%, l | 84617/229 | C42-NCOR2 |
| REACTOME | REACTOME | REACTOME | 151 | 0.377386 | 1.579032 | 0.001703 | 0.018235 | 0.014111 | 1300 | tags=20%, l | 7431/3009, | C42-NCOR2 |
| NAKAMUR | NAKAMUR | NAKAMUR | 432 | -0.281654 | -1.373177 | 0.001709 | 0.018254 | 0.014126 | 2154 | tags=25%, l | 90637/104 | C42-NCOR2 |

| | | | | | | | | | | | | |
|---|---|---|---|---|---|---|---|---|---|---|---|---|
| DAVICIONI_ | DAVICIONI_ | DAVICIONI_ | 111 | 0.405739 | 1.623585 | 0.001736 | 0.018489 | 0.014308 | 1862 | tags=26%, l | 7227/1076 | C42-NCOR2 |
| HALLMARK | HALLMARK | HALLMARK | 38 | -0.561442 | -1.904258 | 0.001791 | 0.019023 | 0.014722 | 2387 | tags=55%, l | 3163/6648, | C42-NCOR2 |
| STEIN_ESR: | STEIN_ESR: | STEIN_ESR1 | 64 | 0.482738 | 1.766211 | 0.001795 | 0.019023 | 0.014722 | 1729 | tags=30%, l | 4173/1109 | C42-NCOR2 |
| PID_E2F_P/ | PID_E2F_P/ | PID_E2F_P/ | 63 | 0.482289 | 1.758167 | 0.001801 | 0.019041 | 0.014735 | 1243 | tags=27%, l | 11073/913 | C42-NCOR2 |
| DEURIG_T_ | DEURIG_T_ | DEURIG_T_ | 199 | 0.349563 | 1.515026 | 0.00182 | 0.019189 | 0.01485 | 2395 | tags=28%, l | 10892/988 | C42-NCOR2 |
| REACTOME | REACTOME | REACTOME | 19 | 0.65752 | 1.844965 | 0.00187 | 0.019665 | 0.015218 | 1231 | tags=42%, l | 51606/364 | C42-NCOR2 |
| FLORIO_NE | FLORIO_NE | FLORIO_NE | 124 | 0.392769 | 1.597449 | 0.001906 | 0.019996 | 0.015475 | 1893 | tags=26%, l | 93664/137 | C42-NCOR2 |
| COLLER_M | COLLER_M | COLLER_M | 18 | -0.690594 | -1.96866 | 0.001914 | 0.020034 | 0.015504 | 1548 | tags=56%, l | 708/191/7 | C42-NCOR2 |
| REACTOME | REACTOME | REACTOME | 78 | 0.447889 | 1.693205 | 0.001949 | 0.020352 | 0.01575 | 2300 | tags=33%, l | 3012/8335, | C42-NCOR2 |
| ACEVEDO_ | ACEVEDO_ | ACEVEDO_ | 68 | 0.464093 | 1.71602 | 0.001969 | 0.020509 | 0.015871 | 777 | tags=24%, l | 3012/1573 | C42-NCOR2 |
| RODRIGUES | RODRIGUES | RODRIGUES | 511 | 0.287772 | 1.369518 | 0.00198 | 0.020566 | 0.015916 | 1699 | tags=23%, l | 4173/1107 | C42-NCOR2 |
| HU_GENOT | HU_GENOT | HU_GENOT | 33 | 0.559735 | 1.78618 | 0.002018 | 0.020917 | 0.016188 | 1965 | tags=39%, l | 54443/475 | C42-NCOR2 |
| BIOCARTA_ | BIOCARTA_ | BIOCARTA_ | 16 | 0.674175 | 1.814755 | 0.002067 | 0.021312 | 0.016493 | 1255 | tags=44%, l | 5295/2323 | C42-NCOR2 |
| REACTOME | REACTOME | REACTOME | 16 | 0.67367 | 1.813394 | 0.002067 | 0.021312 | 0.016493 | 2027 | tags=38%, l | 2787/2323 | C42-NCOR2 |
| REACTOME | REACTOME | REACTOME | 80 | 0.439438 | 1.670074 | 0.002118 | 0.021789 | 0.016862 | 2300 | tags=30%, l | 3012/8335, | C42-NCOR2 |
| LEE_BMP2_ | LEE_BMP2_ | LEE_BMP2_ | 467 | 0.292809 | 1.384744 | 0.002146 | 0.022023 | 0.017043 | 2221 | tags=29%, l | 7227/5560 | C42-NCOR2 |
| JOHNSTON | JOHNSTON | JOHNSTON | 329 | -0.301848 | -1.435784 | 0.002168 | 0.022196 | 0.017177 | 1563 | tags=21%, l | 1509/3728, | C42-NCOR2 |
| CHICAS_RB | CHICAS_RB | CHICAS_RB | 337 | 0.312251 | 1.43346 | 0.002218 | 0.022649 | 0.017528 | 2079 | tags=28%, l | 4192/7431, | C42-NCOR2 |
| ACOSTA_PI | ACOSTA_PI | ACOSTA_PI | 67 | -0.448993 | -1.707945 | 0.002263 | 0.023056 | 0.017842 | 2472 | tags=39%, l | 10560/274 | C42-NCOR2 |
| DAZARD_R | DAZARD_R | DAZARD_R | 238 | 0.334761 | 1.480095 | 0.002301 | 0.023392 | 0.018103 | 2480 | tags=29%, l | 79572/715 | C42-NCOR2 |
| PID_BETA_ | PID_BETA_ | PID_BETA_ | 49 | 0.504173 | 1.753803 | 0.002314 | 0.023467 | 0.01816 | 1913 | tags=33%, l | 6925/1499, | C42-NCOR2 |
| TOOKER_G | TOOKER_G | TOOKER_G | 103 | -0.392115 | -1.605825 | 0.002323 | 0.023502 | 0.018187 | 1528 | tags=28%, l | 1528/8073, | C42-NCOR2 |
| DOANE_RE | DOANE_RE | DOANE_RE | 182 | 0.35237 | 1.510826 | 0.00234 | 0.023608 | 0.01827 | 2691 | tags=32%, l | 4173/8193 | C42-NCOR2 |
| REACTOME | REACTOME | REACTOME | 112 | -0.384827 | -1.596093 | 0.002355 | 0.023648 | 0.0183 | 2564 | tags=39%, l | 2230/3326, | C42-NCOR2 |
| DOUGLAS_ | DOUGLAS_ | DOUGLAS_ | 407 | 0.301056 | 1.404045 | 0.002356 | 0.023648 | 0.0183 | 2164 | tags=29%, l | 10157/795 | C42-NCOR2 |
| REACTOME | REACTOME | REACTOME | 30 | 0.567417 | 1.777396 | 0.002368 | 0.023648 | 0.0183 | 1160 | tags=30%, l | 7431/1499, | C42-NCOR2 |
| REACTOME | REACTOME | REACTOME | 30 | 0.566379 | 1.774144 | 0.002368 | 0.023648 | 0.0183 | 2801 | tags=60%, l | 23191/574 | C42-NCOR2 |
| LASTOWSK | LASTOWSK | LASTOWSK | 130 | -0.366106 | -1.555835 | 0.002372 | 0.023648 | 0.0183 | 3049 | tags=37%, l | 84282/218 | C42-NCOR2 |
| OLSSON_E2 | OLSSON_E2 | OLSSON_E2 | 29 | 0.575027 | 1.783175 | 0.002383 | 0.023648 | 0.0183 | 2322 | tags=45%, l | 7431/2297 | C42-NCOR2 |
| ZHANG_AN | ZHANG_AN | ZHANG_AN | 29 | 0.577671 | 1.791376 | 0.002383 | 0.023648 | 0.0183 | 1522 | tags=34%, l | 10769/154 | C42-NCOR2 |
| LUI_THYRC | LUI_THYRC | LUI_THYRC | 33 | -0.571401 | -1.880522 | 0.002417 | 0.023927 | 0.018517 | 1675 | tags=45%, l | 51614/104 | C42-NCOR2 |
| REACTOME | REACTOME | REACTOME | 19 | 0.645886 | 1.812322 | 0.00243 | 0.024008 | 0.018579 | 726 | tags=32%, l | 7153/7029, | C42-NCOR2 |
| BERNARD_ | BERNARD_ | BERNARD_ | 29 | -0.59625 | -1.911726 | 0.00264 | 0.026014 | 0.020132 | 184 | tags=17%, l | 1349/5689 | C42-NCOR2 |
| REACTOME | REACTOME | REACTOME | 11 | -0.745915 | -1.839216 | 0.002716 | 0.026701 | 0.020663 | 176 | tags=55%, l | 3714/5753 | C42-NCOR2 |
| MULLIGHA | MULLIGHA | MULLIGHA | 53 | -0.475887 | -1.726054 | 0.002733 | 0.026807 | 0.020745 | 2203 | tags=32%, l | 3784/473/ | C42-NCOR2 |
| KEGG_PRO | KEGG_PRO | KEGG_PRO | 13 | -0.722695 | -1.869065 | 0.002739 | 0.026807 | 0.020745 | 2461 | tags=62%, l | 476/483/4 | C42-NCOR2 |
| MAYBURD_ | MAYBURD_ | MAYBURD_ | 21 | -0.638382 | -1.889683 | 0.002789 | 0.027227 | 0.02107 | 174 | tags=29%, l | 9968/2617, | C42-NCOR2 |
| REACTOME | REACTOME | REACTOME | 84 | 0.425057 | 1.629432 | 0.002826 | 0.027528 | 0.021303 | 1405 | tags=21%, l | 3012/8335, | C42-NCOR2 |
| PACHER_T/ | PACHER_T/ | PACHER_T/ | 27 | -0.603962 | -1.900727 | 0.002843 | 0.027626 | 0.021379 | 1910 | tags=48%, l | 4189/7422, | C42-NCOR2 |
| REACTOME | REACTOME | REACTOME | 633 | 0.278271 | 1.34609 | 0.002849 | 0.027626 | 0.021379 | 1831 | tags=21%, l | 6192/3012, | C42-NCOR2 |
| LENAOUR_ | LENAOUR_ | LENAOUR_ | 69 | -0.441735 | -1.688046 | 0.002942 | 0.02846 | 0.022025 | 1333 | tags=35%, l | 3959/1065 | C42-NCOR2 |
| WEST_ADR | WEST_ADR | WEST_ADR | 320 | -0.301748 | -1.431972 | 0.002955 | 0.028526 | 0.022076 | 1521 | tags=20%, l | 79754/936 | C42-NCOR2 |
| REACTOME | REACTOME | REACTOME | 18 | -0.677262 | -1.930653 | 0.002977 | 0.028659 | 0.022178 | 3121 | tags=67%, l | 819/3162/ | C42-NCOR2 |
| HOSHIDA_I | HOSHIDA_I | HOSHIDA_I | 175 | -0.33388 | -1.480031 | 0.002983 | 0.028659 | 0.022178 | 2098 | tags=33%, l | 178/83758, | C42-NCOR2 |
| BENPORAT | BENPORAT | BENPORAT | 185 | -0.332186 | -1.483912 | 0.00299 | 0.028659 | 0.022178 | 1816 | tags=37%, l | 5689/5860, | C42-NCOR2 |
| REACTOME | REACTOME | REACTOME | 57 | 0.480576 | 1.72037 | 0.003036 | 0.02899 | 0.022435 | 1243 | tags=26%, l | 3012/8335, | C42-NCOR2 |
| NELSON_RI | NELSON_RI | NELSON_RI | 74 | 0.438634 | 1.644226 | 0.003038 | 0.02899 | 0.022435 | 1289 | tags=31%, l | 3817/8555, | C42-NCOR2 |
| WELCSH_B | WELCSH_B | WELCSH_B | 121 | -0.371859 | -1.559571 | 0.003096 | 0.029478 | 0.022812 | 3460 | tags=45%, l | 9790/7520, | C42-NCOR2 |
| LEE_LIVER_ | LEE_LIVER_ | LEE_LIVER_ | 27 | 0.592282 | 1.804624 | 0.003131 | 0.029742 | 0.023017 | 1150 | tags=33%, l | 10157/604 | C42-NCOR2 |
| LEE_DIFFEF | LEE_DIFFEF | LEE_DIFFEF | 101 | 0.404289 | 1.596165 | 0.003175 | 0.030096 | 0.023291 | 2790 | tags=43%, l | 360/5150/ | C42-NCOR2 |
| FRASOR_RI | FRASOR_RI | FRASOR_RI | 47 | 0.503239 | 1.731039 | 0.00321 | 0.030355 | 0.023491 | 2054 | tags=45%, l | 5982/3832, | C42-NCOR2 |
| REACTOME | REACTOME | REACTOME | 31 | 0.558036 | 1.757774 | 0.003289 | 0.03104 | 0.024021 | 1782 | tags=35%, l | 11073/598 | C42-NCOR2 |
| BENPORAT | BENPORAT | BENPORAT | 642 | -0.256295 | -1.29491 | 0.003341 | 0.03146 | 0.024346 | 2589 | tags=28%, l | 6945/8574, | C42-NCOR2 |
| REACTOME | REACTOME | REACTOME | 15 | 0.677957 | 1.794633 | 0.003387 | 0.031815 | 0.024621 | 1193 | tags=40%, l | 7029/4605, | C42-NCOR2 |
| BIOCARTA_ | BIOCARTA_ | BIOCARTA_ | 18 | -0.654801 | -1.866624 | 0.003403 | 0.031896 | 0.024683 | 2702 | tags=56%, l | 1647/3337, | C42-NCOR2 |
| REACTOME | REACTOME | REACTOME | 14 | 0.700693 | 1.813644 | 0.003418 | 0.031962 | 0.024735 | 1990 | tags=57%, l | 7029/1869, | C42-NCOR2 |
| REACTOME | REACTOME | REACTOME | 60 | -0.457001 | -1.696705 | 0.003427 | 0.03198 | 0.024749 | 3482 | tags=45%, l | 79796/459 | C42-NCOR2 |
| DUTERTRE_ | DUTERTRE_ | DUTERTRE_ | 377 | 0.299046 | 1.386276 | 0.003455 | 0.032165 | 0.024891 | 2212 | tags=31%, l | 10769/101 | C42-NCOR2 |
| ELVIDGE_H | ELVIDGE_H | ELVIDGE_H | 120 | 0.39077 | 1.582872 | 0.003462 | 0.032165 | 0.024891 | 2151 | tags=34%, l | 5236/1057 | C42-NCOR2 |
| REACTOME | REACTOME | REACTOME | 181 | -0.330671 | -1.473183 | 0.003487 | 0.032325 | 0.025016 | 2005 | tags=30%, l | 6204/5699, | C42-NCOR2 |
| CAIRO_LIVI | CAIRO_LIVI | CAIRO_LIVI | 101 | 0.40258 | 1.589417 | 0.003528 | 0.032585 | 0.025217 | 2246 | tags=31%, l | 4192/7227, | C42-NCOR2 |
| SABATES_C | SABATES_C | SABATES_C | 94 | 0.410352 | 1.603958 | 0.00353 | 0.032585 | 0.025217 | 2597 | tags=39%, l | 5865/3899, | C42-NCOR2 |
| FUKUSHIM | FUKUSHIM | FUKUSHIM | 11 | -0.733686 | -1.809062 | 0.003551 | 0.032706 | 0.02531 | 316 | tags=27%, l | 4851/182 | C42-NCOR2 |
| FEVR_CTNI | FEVR_CTNI | FEVR_CTNI | 417 | 0.293428 | 1.373054 | 0.003575 | 0.032845 | 0.025418 | 1600 | tags=21%, l | 4173/3070, | C42-NCOR2 |
| KEGG_NITF | KEGG_NITF | KEGG_NITF | 13 | -0.713962 | -1.846479 | 0.003582 | 0.032845 | 0.025418 | 1004 | tags=54%, l | 2747/2746, | C42-NCOR2 |
| HALLMARK | HALLMARK | HALLMARK | 65 | 0.458197 | 1.678424 | 0.003598 | 0.032924 | 0.025479 | 2868 | tags=46%, l | 11073/741 | C42-NCOR2 |

| | | | | | | | | | | | | |
|---|---|---|---|---|---|---|---|---|---|---|---|---|
| REACTOME | REACTOME | REACTOME | 404 | 0.296628 | 1.382586 | 0.003611 | 0.03297 | 0.025515 | 2369 | tags=26%, | 3009/3012, | C42-NCOR2 |
| BIOCARTA_ | BIOCARTA_ | BIOCARTA_ | 10 | 0.76492 | 1.810187 | 0.00365 | 0.033183 | 0.02568 | 1456 | tags=60%, | 10810/854 | C42-NCOR2 |
| REACTOME | REACTOME | REACTOME | 10 | 0.761457 | 1.801993 | 0.00365 | 0.033183 | 0.02568 | 1699 | tags=60%, | 9918/6415 | C42-NCOR2 |
| PID_FOXM | PID_FOXM | PID_FOXM | 35 | 0.538413 | 1.739825 | 0.003662 | 0.033221 | 0.025709 | 3936 | tags=63%, | 4751/9212 | C42-NCOR2 |
| MODY_HIP | MODY_HIP | MODY_HIP | 23 | -0.616192 | -1.867832 | 0.003673 | 0.033249 | 0.02573 | 2092 | tags=57%, | 10376/690 | C42-NCOR2 |
| PAL_PRMT | PAL_PRMT | PAL_PRMT | 160 | 0.355398 | 1.499586 | 0.003714 | 0.033548 | 0.025962 | 1438 | tags=22%, | 4173/1107 | C42-NCOR2 |
| WANG_TAI | WANG_TAI | WANG_TAI | 33 | -0.559677 | -1.841937 | 0.003735 | 0.033634 | 0.026028 | 1817 | tags=45%, | 51068/741 | C42-NCOR2 |
| GARGALOV | GARGALOV | GARGALOV | 20 | 0.634928 | 1.801986 | 0.00374 | 0.033634 | 0.026028 | 2294 | tags=45%, | 148418/98 | C42-NCOR2 |
| REACTOME | REACTOME | REACTOME | 67 | 0.450351 | 1.660916 | 0.003761 | 0.033757 | 0.026124 | 2109 | tags=30%, | 3012/8335 | C42-NCOR2 |
| REACTOME | REACTOME | REACTOME | 40 | 0.514933 | 1.714869 | 0.00377 | 0.033758 | 0.026124 | 2192 | tags=40%, | 2787/1499 | C42-NCOR2 |
| RHODES_U | RHODES_U | RHODES_U | 62 | 0.464711 | 1.688312 | 0.003793 | 0.033882 | 0.026221 | 2189 | tags=44%, | 5885/7153 | C42-NCOR2 |
| ACEVEDO_ | ACEVEDO_ | ACEVEDO_ | 340 | -0.291081 | -1.38985 | 0.003803 | 0.033882 | 0.026221 | 2484 | tags=28%, | 3155/595/ | C42-NCOR2 |
| KAUFFMAN | KAUFFMAN | KAUFFMAN | 116 | 0.38891 | 1.567512 | 0.003811 | 0.033882 | 0.026221 | 2897 | tags=30%, | 4173/1107 | C42-NCOR2 |
| VECCHI_GA | VECCHI_GA | VECCHI_GA | 326 | 0.309986 | 1.419338 | 0.003816 | 0.033882 | 0.026221 | 1782 | tags=25%, | 157570/41 | C42-NCOR2 |
| REACTOME | REACTOME | REACTOME | 175 | 0.347508 | 1.484048 | 0.003847 | 0.034054 | 0.026354 | 1684 | tags=21%, | 4192/3012 | C42-NCOR2 |
| YAGI_AML_ | YAGI_AML_ | YAGI_AML_ | 230 | -0.315292 | -1.444628 | 0.003851 | 0.034054 | 0.026354 | 1859 | tags=25%, | 3077/9452 | C42-NCOR2 |
| DAVICIONI_ | DAVICIONI_ | DAVICIONI_ | 168 | 0.35207 | 1.496652 | 0.003868 | 0.034069 | 0.026365 | 2638 | tags=38%, | 360/55872 | C42-NCOR2 |
| LIU_NASOF | LIU_NASOF | LIU_NASOF | 47 | -0.500584 | -1.768563 | 0.003869 | 0.034069 | 0.026365 | 2153 | tags=45%, | 7049/3628 | C42-NCOR2 |
| MCBRYAN_ | MCBRYAN_ | MCBRYAN_ | 83 | -0.409984 | -1.618167 | 0.003922 | 0.034428 | 0.026643 | 1074 | tags=22%, | 3988/2854 | C42-NCOR2 |
| REACTOME | REACTOME | REACTOME | 19 | 0.633883 | 1.778642 | 0.003926 | 0.034428 | 0.026643 | 2027 | tags=47%, | 2787/5567 | C42-NCOR2 |
| RICKMAN_ | RICKMAN_ | RICKMAN_ | 41 | 0.51418 | 1.720662 | 0.003947 | 0.034539 | 0.026729 | 1853 | tags=39%, | 4885/2308 | C42-NCOR2 |
| KEGG_CAR | KEGG_CAR | KEGG_CAR | 36 | -0.534943 | -1.792649 | 0.003987 | 0.034816 | 0.026943 | 1442 | tags=36%, | 7385/1337 | C42-NCOR2 |
| RODWELL_ | RODWELL_ | RODWELL_ | 111 | -0.378497 | -1.567784 | 0.004008 | 0.034877 | 0.026991 | 2363 | tags=37%, | 3032/2806 | C42-NCOR2 |
| AMUNDSO | AMUNDSO | AMUNDSO | 14 | -0.698188 | -1.851218 | 0.004013 | 0.034877 | 0.026991 | 1552 | tags=79%, | 23654/849 | C42-NCOR2 |
| SARRIO_EP | SARRIO_EP | SARRIO_EP | 116 | -0.370046 | -1.543463 | 0.00402 | 0.034877 | 0.026991 | 2337 | tags=34%, | 667/9961/ | C42-NCOR2 |
| BIOCARTA_ | BIOCARTA_ | BIOCARTA_ | 11 | 0.736973 | 1.790512 | 0.004027 | 0.034877 | 0.026991 | 1466 | tags=45%, | 5567/6256 | C42-NCOR2 |
| KEGG_COL | KEGG_COL | KEGG_COL | 54 | 0.474609 | 1.685991 | 0.004081 | 0.035273 | 0.027297 | 2193 | tags=35%, | 5295/1499 | C42-NCOR2 |
| PID_IL2_1P | PID_IL2_1P | PID_IL2_1P | 45 | 0.508411 | 1.735504 | 0.004106 | 0.035415 | 0.027407 | 1561 | tags=33%, | 5295/596/ | C42-NCOR2 |
| REACTOME | REACTOME | REACTOME | 42 | 0.506516 | 1.705073 | 0.00412 | 0.035463 | 0.027444 | 2566 | tags=40%, | 29127/846 | C42-NCOR2 |
| VANTVEER_ | VANTVEER_ | VANTVEER_ | 28 | -0.577966 | -1.836335 | 0.004164 | 0.035771 | 0.027682 | 2428 | tags=50%, | 9522/9703 | C42-NCOR2 |
| REACTOME | REACTOME | REACTOME | 15 | -0.686182 | -1.855471 | 0.004267 | 0.036583 | 0.028311 | 1790 | tags=40%, | 7416/1045 | C42-NCOR2 |
| PEDERSEN_ | PEDERSEN_ | PEDERSEN_ | 53 | 0.473632 | 1.675493 | 0.004277 | 0.036589 | 0.028316 | 2186 | tags=38%, | 5865/1134 | C42-NCOR2 |
| SCHLOSSER | SCHLOSSER | SCHLOSSER | 43 | -0.506724 | -1.756013 | 0.004309 | 0.036795 | 0.028475 | 1720 | tags=49%, | 10969/365 | C42-NCOR2 |
| LE_NEURO | LE_NEURO | LE_NEURO | 17 | 0.645822 | 1.764716 | 0.004328 | 0.036881 | 0.028542 | 2090 | tags=53%, | 11065/949 | C42-NCOR2 |
| TIEN_INTES | TIEN_INTES | TIEN_INTES | 25 | -0.595473 | -1.837802 | 0.004361 | 0.037087 | 0.028701 | 1586 | tags=48%, | 8073/567/ | C42-NCOR2 |
| REACTOME | REACTOME | REACTOME | 10 | -0.747982 | -1.800961 | 0.004377 | 0.037146 | 0.028746 | 2461 | tags=80%, | 2806/5832 | C42-NCOR2 |
| PEREZ_TP6 | PEREZ_TP6 | PEREZ_TP6 | 221 | 0.331712 | 1.456795 | 0.004434 | 0.037557 | 0.029065 | 2166 | tags=26%, | 10570/360 | C42-NCOR2 |
| PID_INTEG | PID_INTEG | PID_INTEG | 24 | 0.599046 | 1.778711 | 0.004449 | 0.037603 | 0.0291 | 1399 | tags=33%, | 4192/5567 | C42-NCOR2 |
| MEISSNER_ | MEISSNER_ | MEISSNER_ | 119 | 0.383098 | 1.550618 | 0.004494 | 0.037915 | 0.029341 | 2670 | tags=34%, | 93664/552 | C42-NCOR2 |
| REACTOME | REACTOME | REACTOME | 21 | -0.620722 | -1.837408 | 0.004505 | 0.037924 | 0.029349 | 1790 | tags=33%, | 7416/1921 | C42-NCOR2 |
| REACTOME | REACTOME | REACTOME | 40 | -0.510828 | -1.749232 | 0.004514 | 0.037925 | 0.02935 | 2337 | tags=38%, | 5908/7297 | C42-NCOR2 |
| REACTOME | REACTOME | REACTOME | 43 | -0.501791 | -1.738918 | 0.004536 | 0.038039 | 0.029437 | 1586 | tags=47%, | 6209/1965 | C42-NCOR2 |
| REACTOME | REACTOME | REACTOME | 10 | 0.746208 | 1.765905 | 0.004611 | 0.038589 | 0.029863 | 551 | tags=40%, | 5567/1134 | C42-NCOR2 |
| PID_FANCC | PID_FANCC | PID_FANCC | 40 | 0.511619 | 1.703834 | 0.004667 | 0.038968 | 0.030156 | 3124 | tags=45%, | 11073/598 | C42-NCOR2 |
| REACTOME | REACTOME | REACTOME | 20 | 0.623163 | 1.768596 | 0.004675 | 0.038968 | 0.030156 | 1231 | tags=40%, | 51606/528 | C42-NCOR2 |
| MARIADAS | MARIADAS | MARIADAS | 20 | -0.628146 | -1.83593 | 0.004727 | 0.039327 | 0.030435 | 2035 | tags=55%, | 3632/35/7 | C42-NCOR2 |
| KEGG_N_G | KEGG_N_G | KEGG_N_G | 42 | -0.502578 | -1.736511 | 0.004752 | 0.039458 | 0.030536 | 882 | tags=29%, | 29880/618 | C42-NCOR2 |
| KEGG_REG | KEGG_REG | KEGG_REG | 141 | 0.367264 | 1.523294 | 0.004777 | 0.039582 | 0.030632 | 2801 | tags=36%, | 5295/3693 | C42-NCOR2 |
| REACTOME | REACTOME | REACTOME | 11 | 0.73378 | 1.782754 | 0.004794 | 0.039603 | 0.030648 | 2565 | tags=64%, | 9334/9331 | C42-NCOR2 |
| REACTOME | REACTOME | REACTOME | 60 | -0.452185 | -1.678822 | 0.004798 | 0.039603 | 0.030648 | 1000 | tags=30%, | 7367/191/ | C42-NCOR2 |
| REACTOME | REACTOME | REACTOME | 43 | 0.49924 | 1.689017 | 0.004827 | 0.03977 | 0.030777 | 2334 | tags=44%, | 8354/9070 | C42-NCOR2 |
| HADDAD_T | HADDAD_T | HADDAD_T | 23 | 0.593811 | 1.746601 | 0.004838 | 0.03978 | 0.030785 | 2213 | tags=57%, | 4885/4173 | C42-NCOR2 |
| RICKMAN_ | RICKMAN_ | RICKMAN_ | 21 | 0.605861 | 1.739488 | 0.004869 | 0.039955 | 0.030921 | 738 | tags=29%, | 8287/9086 | C42-NCOR2 |
| TAKEDA_TA | TAKEDA_TA | TAKEDA_TA | 107 | 0.389022 | 1.548689 | 0.004886 | 0.040015 | 0.030967 | 1534 | tags=27%, | 8821/4940 | C42-NCOR2 |
| VILLANUEV | VILLANUEV | VILLANUEV | 38 | -0.514284 | -1.74431 | 0.004925 | 0.040259 | 0.031156 | 2713 | tags=53%, | 1647/1119 | C42-NCOR2 |
| REACTOME | REACTOME | REACTOME | 95 | 0.396284 | 1.551284 | 0.00496 | 0.040468 | 0.031317 | 2109 | tags=29%, | 3012/8335 | C42-NCOR2 |
| FIGUEROA_ | FIGUEROA_ | FIGUEROA_ | 66 | 0.44189 | 1.6249 | 0.005017 | 0.040853 | 0.031615 | 1334 | tags=24%, | 63934/846 | C42-NCOR2 |
| FLECHNER_ | FLECHNER_ | FLECHNER_ | 423 | -0.273453 | -1.331014 | 0.005073 | 0.041127 | 0.031827 | 2470 | tags=31%, | 2744/9648 | C42-NCOR2 |
| BIOCARTA_ | BIOCARTA_ | BIOCARTA_ | 15 | 0.666297 | 1.763768 | 0.00508 | 0.041127 | 0.031827 | 1939 | tags=60%, | 5295/2323 | C42-NCOR2 |
| SIMBULAN_ | SIMBULAN_ | SIMBULAN_ | 15 | 0.665767 | 1.762363 | 0.00508 | 0.041127 | 0.031827 | 1550 | tags=60%, | 3012/991/ | C42-NCOR2 |
| REACTOME | REACTOME | REACTOME | 46 | 0.484189 | 1.658495 | 0.005177 | 0.0417 | 0.032271 | 2801 | tags=48%, | 23191/574 | C42-NCOR2 |
| REACTOME | REACTOME | REACTOME | 11 | 0.731914 | 1.778219 | 0.005177 | 0.0417 | 0.032271 | 2604 | tags=64%, | 1543/1573 | C42-NCOR2 |
| CROONQUI | CROONQUI | CROONQUI | 10 | 0.742379 | 1.756843 | 0.005187 | 0.0417 | 0.032271 | 2072 | tags=60%, | 10123/231 | C42-NCOR2 |
| AMUNDSO | AMUNDSO | AMUNDSO | 171 | -0.324995 | -1.437084 | 0.00519 | 0.0417 | 0.032271 | 1851 | tags=27%, | 586/3726/ | C42-NCOR2 |
| WANG_TUI | WANG_TUI | WANG_TUI | 290 | -0.299311 | -1.407231 | 0.005245 | 0.042051 | 0.032543 | 2203 | tags=28%, | 7417/1351 | C42-NCOR2 |
| KEGG_POR | KEGG_POR | KEGG_POR | 26 | -0.572479 | -1.783642 | 0.005254 | 0.042051 | 0.032543 | 2362 | tags=50%, | 326625/20 | C42-NCOR2 |

| | | | | | | | | | | | | |
|---|---|---|---|---|---|---|---|---|---|---|---|---|
| REACTOME | REACTOME | REACTOME | 133 | -0.352432 | -1.501896 | 0.005276 | 0.04212 | 0.032596 | 3089 | tags=42%, | 83594/796 | C42-NCOR2 |
| GOBERT_O | GOBERT_O | GOBERT_O | 725 | 0.262842 | 1.284515 | 0.005282 | 0.04212 | 0.032596 | 2842 | tags=30%, | 9185/1134 | C42-NCOR2 |
| MORI_PLA | MORI_PLA | MORI_PLA | 31 | -0.547996 | -1.779586 | 0.005298 | 0.042164 | 0.03263 | 2091 | tags=52%, | 148867/68 | C42-NCOR2 |
| REACTOME | REACTOME | REACTOME | 12 | 0.71192 | 1.772584 | 0.005326 | 0.042309 | 0.032742 | 1990 | tags=58%, | 7029/1869 | C42-NCOR2 |
| KEGG_ARR | KEGG_ARR | KEGG_ARR | 40 | 0.506807 | 1.687807 | 0.005385 | 0.04264 | 0.032998 | 2186 | tags=48%, | 1824/1499 | C42-NCOR2 |
| GROSS_HY | GROSS_HY | GROSS_HY | 166 | 0.346676 | 1.470212 | 0.005388 | 0.04264 | 0.032998 | 2296 | tags=32%, | 11073/558 | C42-NCOR2 |
| MARTINEZ_ | MARTINEZ_ | MARTINEZ_ | 435 | -0.270896 | -1.321681 | 0.005425 | 0.042855 | 0.033165 | 1473 | tags=17%, | 10284/715 | C42-NCOR2 |
| BACOLOD_ | BACOLOD_ | BACOLOD_ | 13 | 0.698577 | 1.77614 | 0.005518 | 0.043408 | 0.033592 | 1724 | tags=46%, | 4192/3479 | C42-NCOR2 |
| MCBRYAN_ | MCBRYAN_ | MCBRYAN_ | 139 | -0.349497 | -1.501508 | 0.005524 | 0.043408 | 0.033592 | 1402 | tags=22%, | 6662/5886 | C42-NCOR2 |
| BASAKI_YB | BASAKI_YB | BASAKI_YB | 284 | -0.298365 | -1.397647 | 0.005526 | 0.043408 | 0.033592 | 1712 | tags=22%, | 163131/74 | C42-NCOR2 |
| KARLSSON_ | KARLSSON_ | KARLSSON_ | 89 | -0.39633 | -1.581309 | 0.005553 | 0.043536 | 0.033692 | 1947 | tags=38%, | 1104/6950 | C42-NCOR2 |
| ODONNELL | ODONNELL | ODONNELL | 38 | 0.519962 | 1.712025 | 0.005601 | 0.043829 | 0.033918 | 2769 | tags=50%, | 3070/7153 | C42-NCOR2 |
| KEGG_PRO | KEGG_PRO | KEGG_PRO | 26 | -0.565615 | -1.762256 | 0.005692 | 0.044372 | 0.034339 | 2999 | tags=54%, | 223/219/84 | C42-NCOR2 |
| CAVARD_LI | CAVARD_LI | CAVARD_LI | 14 | -0.690839 | -1.831731 | 0.005702 | 0.044372 | 0.034339 | 2030 | tags=50%, | 5052/9362 | C42-NCOR2 |
| PID_BCR_5 | PID_BCR_5 | PID_BCR_5 | 51 | 0.470963 | 1.651318 | 0.005712 | 0.044372 | 0.034339 | 2360 | tags=35%, | 10892/529 | C42-NCOR2 |
| TAVAZOIE_ | TAVAZOIE_ | TAVAZOIE_ | 51 | 0.469526 | 1.646281 | 0.005712 | 0.044372 | 0.034339 | 1387 | tags=25%, | 8287/7404 | C42-NCOR2 |
| OSMAN_BL | OSMAN_BL | OSMAN_BL | 292 | -0.295334 | -1.388919 | 0.005774 | 0.044772 | 0.034648 | 2032 | tags=23%, | 11224/262 | C42-NCOR2 |
| MULLIGHA | MULLIGHA | MULLIGHA | 89 | 0.404343 | 1.565862 | 0.00581 | 0.044965 | 0.034797 | 2226 | tags=31%, | 117178/38 | C42-NCOR2 |
| SMIRNOV_ | SMIRNOV_ | SMIRNOV_ | 76 | 0.424289 | 1.598793 | 0.005857 | 0.045249 | 0.035017 | 2189 | tags=38%, | 22974/475 | C42-NCOR2 |
| KEGG_VALI | KEGG_VALI | KEGG_VALI | 42 | -0.498473 | -1.722325 | 0.005884 | 0.04537 | 0.035111 | 3334 | tags=57%, | 3157/6406 | C42-NCOR2 |
| REACTOME | REACTOME | REACTOME | 22 | -0.594037 | -1.777883 | 0.006036 | 0.046458 | 0.035953 | 578 | tags=45%, | 8650/2322 | C42-NCOR2 |
| DAVICIONI_ | DAVICIONI_ | DAVICIONI_ | 11 | 0.72783 | 1.768297 | 0.006328 | 0.048593 | 0.037605 | 1132 | tags=27%, | 10769/692 | C42-NCOR2 |
| CHAUHAN_ | CHAUHAN_ | CHAUHAN_ | 42 | -0.496208 | -1.714499 | 0.006336 | 0.048593 | 0.037605 | 1396 | tags=33%, | 4609/440/ | C42-NCOR2 |
| DIAZ_CHRO | DIAZ_CHRO | DIAZ_CHRO | 61 | -0.437794 | -1.630994 | 0.006371 | 0.04877 | 0.037742 | 2256 | tags=39%, | 7107/2367 | C42-NCOR2 |
| AGUIRRE_I | AGUIRRE_I | AGUIRRE_I | 181 | -0.317781 | -1.415756 | 0.006476 | 0.049483 | 0.038294 | 2015 | tags=27%, | 9841/5565 | C42-NCOR2 |
| LEE_NAIVE | LEE_NAIVE | LEE_NAIVE | 11 | 0.726436 | 1.764912 | 0.00652 | 0.049729 | 0.038484 | 664 | tags=36%, | 6192/8821 | C42-NCOR2 |
| REACTOME | REACTOME | REACTOME | 36 | 0.515492 | 1.677879 | 0.006561 | 0.049953 | 0.038658 | 1969 | tags=33%, | 55193/86/ | C42-NCOR2 |

C42_DHT

| | ID | Descriptior | setSize | enrichmen | NES | pvalue | p.adjust | qvalues | rank | leading_ed | core_enricl | Group |
|---|---|---|---|---|---|---|---|---|---|---|---|---|
| DACOSTA_ | DACOSTA_ | DACOSTA_ | 674 | 0.476554 | 1.73974 | 0.000104 | 0.024717 | 0.021688 | 1523 | tags=26%, | 129642/26 | C42+DHT |
| JOHNSTON | JOHNSTON | JOHNSTON | 647 | 0.422002 | 1.537874 | 0.000104 | 0.024717 | 0.021688 | 2425 | tags=28%, | 2289/2597 | C42+DHT |
| KOINUMA_ | KOINUMA_ | KOINUMA_ | 624 | 0.431719 | 1.571178 | 0.000104 | 0.024717 | 0.021688 | 1038 | tags=17%, | 79098/705 | C42+DHT |
| CUI_TCF21 | CUI_TCF21 | CUI_TCF21 | 563 | 0.440466 | 1.596516 | 0.000105 | 0.024717 | 0.021688 | 1348 | tags=21%, | 7057/2305 | C42+DHT |
| RODRIGUE: | RODRIGUE: | RODRIGUE: | 553 | 0.446143 | 1.614898 | 0.000105 | 0.024717 | 0.021688 | 1097 | tags=16%, | 7057/2601 | C42+DHT |
| CREIGHTOI | CREIGHTOI | CREIGHTOI | 522 | 0.490515 | 1.771491 | 0.000106 | 0.024717 | 0.021688 | 1034 | tags=19%, | 7113/7909 | C42+DHT |
| ZWANG_CI | ZWANG_CI | ZWANG_CI | 337 | 0.48135 | 1.696507 | 0.000112 | 0.024717 | 0.021688 | 1140 | tags=24%, | 7057/1105 | C42+DHT |
| LU_EZH2_T | LU_EZH2_T | LU_EZH2_T | 281 | 0.494678 | 1.721684 | 0.000115 | 0.024717 | 0.021688 | 1063 | tags=16%, | 7057/2601 | C42+DHT |
| SCHAEFFEF | SCHAEFFEF | SCHAEFFEF | 259 | 0.513927 | 1.775475 | 0.000117 | 0.024717 | 0.021688 | 1005 | tags=17%, | 7113/7909 | C42+DHT |
| FARMER_B | FARMER_B | FARMER_B | 247 | 0.549421 | 1.889944 | 0.000117 | 0.024717 | 0.021688 | 1138 | tags=23%, | 7113/7704 | C42+DHT |
| FARMER_B | FARMER_B | FARMER_B | 234 | 0.571462 | 1.955813 | 0.000118 | 0.024717 | 0.021688 | 903 | tags=18%, | 7113/7057 | C42+DHT |
| GENTILE_U | GENTILE_U | GENTILE_U | 233 | 0.51829 | 1.773279 | 0.000118 | 0.024717 | 0.021688 | 1474 | tags=30%, | 7057/2597 | C42+DHT |
| CREIGHTOI | CREIGHTOI | CREIGHTOI | 195 | 0.559655 | 1.884147 | 0.000122 | 0.024717 | 0.021688 | 1576 | tags=24%, | 26018/228 | C42+DHT |
| DURCHDEV | DURCHDEV | DURCHDEV | 188 | 0.527992 | 1.77045 | 0.000123 | 0.024717 | 0.021688 | 886 | tags=19%, | 25976/107 | C42+DHT |
| DUTERTRE_ | DUTERTRE_ | DUTERTRE_ | 184 | 0.576721 | 1.928049 | 0.000123 | 0.024717 | 0.021688 | 1350 | tags=21%, | 23052/260 | C42+DHT |
| MCBRYAN_ | MCBRYAN_ | MCBRYAN_ | 171 | 0.536757 | 1.7803 | 0.000125 | 0.024717 | 0.021688 | 631 | tags=16%, | 7113/7057 | C42+DHT |
| MASSARW | MASSARW | MASSARW | 163 | 0.614072 | 2.026431 | 0.000126 | 0.024717 | 0.021688 | 764 | tags=20%, | 26018/228 | C42+DHT |
| HALLMARK | HALLMARK | HALLMARK | 153 | 0.560134 | 1.836459 | 0.000128 | 0.024717 | 0.021688 | 510 | tags=16%, | 23052/260 | C42+DHT |
| VECCHI_GA | VECCHI_GA | VECCHI_GA | 150 | 0.555361 | 1.816206 | 0.000128 | 0.024717 | 0.021688 | 800 | tags=17%, | 2289/2293 | C42+DHT |
| DOANE_RE | DOANE_RE | DOANE_RE | 135 | 0.739153 | 2.38827 | 0.00013 | 0.024717 | 0.021688 | 804 | tags=27%, | 7113/7909 | C42+DHT |
| YEGNASUB | YEGNASUB | YEGNASUB | 97 | 0.673186 | 2.086583 | 0.000138 | 0.024717 | 0.021688 | 707 | tags=25%, | 7113/3817 | C42+DHT |
| HALLMARK | HALLMARK | HALLMARK | 93 | 0.885599 | 2.733151 | 0.000139 | 0.024717 | 0.021688 | 612 | tags=55%, | 7113/3817 | C42+DHT |
| LIU_PROST | LIU_PROST | LIU_PROST | 74 | 0.674393 | 2.015767 | 0.000144 | 0.024717 | 0.021688 | 719 | tags=26%, | 10645/260 | C42+DHT |
| NELSON_RI | NELSON_RI | NELSON_RI | 74 | 0.91807 | 2.744119 | 0.000144 | 0.024717 | 0.021688 | 612 | tags=65%, | 7113/3817 | C42+DHT |
| CHEN_LVAI | CHEN_LVAI | CHEN_LVAI | 67 | 0.663525 | 1.953779 | 0.000146 | 0.024717 | 0.021688 | 368 | tags=27%, | 7704/2289 | C42+DHT |
| WANG_RES | WANG_RES | WANG_RES | 26 | 0.870081 | 2.165654 | 0.000171 | 0.026162 | 0.022956 | 651 | tags=50%, | 3817/1064 | C42+DHT |
| BURTON_A | BURTON_A | BURTON_A | 25 | 0.819691 | 2.028008 | 0.000172 | 0.026162 | 0.022956 | 413 | tags=52%, | 22936/884 | C42+DHT |
| WANG_RES | WANG_RES | WANG_RES | 21 | 0.85682 | 2.062347 | 0.000174 | 0.026162 | 0.022956 | 630 | tags=43%, | 3817/1064 | C42+DHT |
| BUYTAERT_ | BUYTAERT_ | BUYTAERT_ | 638 | 0.418479 | 1.524065 | 0.000208 | 0.027723 | 0.024325 | 1348 | tags=21%, | 25976/884 | C42+DHT |
| ONKEN_UV | ONKEN_UV | ONKEN_UV | 580 | 0.422274 | 1.532747 | 0.00021 | 0.027723 | 0.024325 | 2072 | tags=25%, | 65986/884 | C42+DHT |
| SMID_BREA | SMID_BREA | SMID_BREA | 421 | 0.461746 | 1.649822 | 0.000218 | 0.027723 | 0.024325 | 927 | tags=18%, | 7704/2601 | C42+DHT |
| ENK_UV_R | ENK_UV_R | ENK_UV_R | 400 | 0.46503 | 1.656885 | 0.000219 | 0.027723 | 0.024325 | 1341 | tags=22%, | 7704/2293 | C42+DHT |
| DACOSTA_ | DACOSTA_ | DACOSTA_ | 367 | 0.456971 | 1.619742 | 0.000222 | 0.027723 | 0.024325 | 1816 | tags=28%, | 25976/218 | C42+DHT |
| NELSON_RI | NELSON_RI | NELSON_RI | 16 | -0.822444 | -2.113978 | 0.000223 | 0.027723 | 0.024325 | 619 | tags=62%, | 7982/1487 | C42+DHT |
| WANG_LM | WANG_LM | WANG_LM | 270 | 0.490373 | 1.701044 | 0.000231 | 0.027831 | 0.02442 | 1351 | tags=20%, | 58480/705 | C42+DHT |
| PASINI_SUZ | PASINI_SUZ | PASINI_SUZ | 226 | 0.503905 | 1.720807 | 0.000238 | 0.027831 | 0.02442 | 582 | tags=19%, | 7113/7057 | C42+DHT |
| REACTOME | REACTOME | REACTOME | 41 | -0.629397 | -2.005125 | 0.000265 | 0.029478 | 0.025865 | 1362 | tags=46%, | 8367/8347 | C42+DHT |
| SHEDDEN_ | SHEDDEN_ | SHEDDEN_ | 122 | 0.56663 | 1.809361 | 0.000266 | 0.029478 | 0.025865 | 915 | tags=19%, | 79098/770 | C42+DHT |
| NABA_ECM | NABA_ECM | NABA_ECM | 56 | -0.638895 | -2.160958 | 0.000292 | 0.030114 | 0.026423 | 2333 | tags=39%, | 309/8416/ | C42+DHT |
| REACTOME | REACTOME | REACTOME | 55 | -0.587776 | -1.976699 | 0.000293 | 0.030114 | 0.026423 | 1362 | tags=35%, | 8367/8347 | C42+DHT |
| REACTOME | REACTOME | REACTOME | 58 | -0.547432 | -1.86786 | 0.000297 | 0.030114 | 0.026423 | 1362 | tags=34%, | 8367/8347 | C42+DHT |
| KEGG_SYST | KEGG_SYST | KEGG_SYST | 61 | -0.639098 | -2.203395 | 0.000303 | 0.030114 | 0.026423 | 1864 | tags=49%, | 8332/8523 | C42+DHT |
| MASSARW | MASSARW | MASSARW | 45 | 0.71861 | 1.973535 | 0.000314 | 0.030114 | 0.026423 | 524 | tags=27%, | 26018/259 | C42+DHT |
| WONG_AD | WONG_AD | WONG_AD | 447 | 0.443335 | 1.591028 | 0.000325 | 0.030114 | 0.026423 | 960 | tags=16%, | 58480/705 | C42+DHT |
| BERENJENC | BERENJENC | BERENJENC | 413 | 0.435433 | 1.555107 | 0.000328 | 0.030114 | 0.026423 | 1833 | tags=20%, | 7057/1076 | C42+DHT |
| YAMASHIT. | YAMASHIT. | YAMASHIT. | 34 | 0.787607 | 2.061802 | 0.000328 | 0.030114 | 0.026423 | 325 | tags=21%, | 7057/2289 | C42+DHT |
| CHARAFE_I | CHARAFE_I | CHARAFE_I | 292 | 0.465963 | 1.626656 | 0.000343 | 0.030738 | 0.026971 | 902 | tags=17%, | 7704/2293 | C42+DHT |
| MISSIAGLIA | MISSIAGLIA | MISSIAGLIA | 92 | -0.526267 | -1.956235 | 0.000356 | 0.03125 | 0.02742 | 1863 | tags=37%, | 2896/9636 | C42+DHT |
| MIKKELSEN | MIKKELSEN | MIKKELSEN | 112 | -0.492642 | -1.894568 | 0.000385 | 0.03294 | 0.028903 | 2641 | tags=41%, | 54716/925 | C42+DHT |
| SHETH_LIV | SHETH_LIV | SHETH_LIV | 115 | -0.449873 | -1.738246 | 0.00039 | 0.03294 | 0.028903 | 2316 | tags=33%, | 8416/1649 | C42+DHT |
| HALLMARK | HALLMARK | HALLMARK | 131 | -0.448545 | -1.765024 | 0.000422 | 0.034665 | 0.030416 | 2058 | tags=38%, | 5743/3665 | C42+DHT |
| HAMAI_AP | HAMAI_AP | HAMAI_AP | 499 | 0.431772 | 1.555982 | 0.000427 | 0.034665 | 0.030416 | 2518 | tags=32%, | 55808/100 | C42+DHT |
| ZHANG_RE | ZHANG_RE | ZHANG_RE | 147 | -0.409826 | -1.644214 | 0.000449 | 0.035581 | 0.03122 | 942 | tags=37%, | 5606/2303 | C42+DHT |
| SMID_BREA | SMID_BREA | SMID_BREA | 55 | 0.679679 | 1.933767 | 0.000456 | 0.035581 | 0.03122 | 760 | tags=27%, | 10769/105 | C42+DHT |
| MARZEC_IL | MARZEC_IL | MARZEC_IL | 28 | -0.654491 | -1.912332 | 0.000486 | 0.037238 | 0.032674 | 952 | tags=36%, | 1027/5140 | C42+DHT |
| MIKKELSEN | MIKKELSEN | MIKKELSEN | 35 | -0.637829 | -1.957206 | 0.00051 | 0.038362 | 0.03366 | 1027 | tags=34%, | 57626/128 | C42+DHT |
| LEE_BMP2_ | LEE_BMP2_ | LEE_BMP2_ | 673 | 0.401471 | 1.465266 | 0.000518 | 0.038362 | 0.03366 | 2341 | tags=27%, | 2289/6675 | C42+DHT |
| REACTOME | REACTOME | REACTOME | 77 | 0.619952 | 1.865448 | 0.00057 | 0.0405 | 0.035536 | 1071 | tags=18%, | 7057/2305 | C42+DHT |
| BOQUEST_ | BOQUEST_ | BOQUEST_ | 256 | 0.481612 | 1.663193 | 0.000583 | 0.0405 | 0.035536 | 1020 | tags=20%, | 7057/7704 | C42+DHT |
| MIKKELSEN | MIKKELSEN | MIKKELSEN | 54 | -0.557389 | -1.868232 | 0.000583 | 0.0405 | 0.035536 | 1825 | tags=28%, | 85445/321 | C42+DHT |

| | | | | | | | | | | | | |
|---|---|---|---|---|---|---|---|---|---|---|---|---|
| KINSEY_TA | KINSEY_TA | KINSEY_TA | 210 | -0.422432 | -1.774012 | 0.000589 | 0.0405 | 0.035536 | 1433 | tags=28%, | 29926/834 | C42+DHT |
| HOLLERN_S | HOLLERN_S | HOLLERN_S | 61 | -0.5078 | -1.750725 | 0.000605 | 0.0405 | 0.035536 | 1005 | tags=28%, | 5074/6599 | C42+DHT |
| OHGUCHI_ | OHGUCHI_ | OHGUCHI_ | 54 | 0.676453 | 1.918234 | 0.000609 | 0.0405 | 0.035536 | 582 | tags=17%, | 85414/378 | C42+DHT |
| REACTOME | REACTOME | REACTOME | 63 | -0.527859 | -1.83202 | 0.000618 | 0.0405 | 0.035536 | 1362 | tags=35%, | 8367/8347 | C42+DHT |
| AMIT_EGF_ | AMIT_EGF_ | AMIT_EGF_ | 46 | 0.688637 | 1.899397 | 0.000625 | 0.0405 | 0.035536 | 995 | tags=24%, | 7057/1076 | C42+DHT |
| REACTOME | REACTOME | REACTOME | 69 | -0.524782 | -1.854731 | 0.000634 | 0.0405 | 0.035536 | 1362 | tags=32%, | 8367/8347 | C42+DHT |
| REACTOME | REACTOME | REACTOME | 430 | 0.43069 | 1.540988 | 0.000653 | 0.041111 | 0.036073 | 1433 | tags=17%, | 85414/850 | C42+DHT |
| FOSTER_TC | FOSTER_TC | FOSTER_TC | 281 | 0.459416 | 1.598959 | 0.00069 | 0.042779 | 0.037536 | 1035 | tags=18%, | 85414/230 | C42+DHT |
| STEARMAN | STEARMAN | STEARMAN | 32 | -0.628777 | -1.888336 | 0.000755 | 0.045464 | 0.039892 | 354 | tags=19%, | 8751/1168 | C42+DHT |
| PID_AR_TF | PID_AR_TF | PID_AR_TF | 44 | 0.701947 | 1.919237 | 0.00079 | 0.045464 | 0.039892 | 443 | tags=23%, | 7113/3817 | C42+DHT |
| MEISSNER_ | MEISSNER_ | MEISSNER_ | 119 | -0.436594 | -1.694363 | 0.000797 | 0.045464 | 0.039892 | 1884 | tags=36%, | 6692/1305 | C42+DHT |
| CHARAFE_I | CHARAFE_I | CHARAFE_I | 288 | 0.459991 | 1.603693 | 0.000801 | 0.045464 | 0.039892 | 850 | tags=17%, | 10257/182 | C42+DHT |
| MCMURRA | MCMURRA | MCMURRA | 43 | -0.593097 | -1.903225 | 0.000809 | 0.045464 | 0.039892 | 917 | tags=23%, | 143098/38 | C42+DHT |
| REACTOME | REACTOME | REACTOME | 43 | -0.582474 | -1.869136 | 0.000809 | 0.045464 | 0.039892 | 1464 | tags=47%, | 8349/8367 | C42+DHT |
| REACTOME | REACTOME | REACTOME | 50 | -0.576595 | -1.903425 | 0.000855 | 0.045464 | 0.039892 | 1362 | tags=40%, | 3720/8367 | C42+DHT |
| KEGG_GLY( | KEGG_GLY( | KEGG_GLY( | 11 | -0.804239 | -1.893763 | 0.000856 | 0.045464 | 0.039892 | 1284 | tags=45%, | 9197/2583 | C42+DHT |
| WANG_TH | WANG_TH | WANG_TH | 11 | -0.812557 | -1.913352 | 0.000856 | 0.045464 | 0.039892 | 700 | tags=55%, | 4851/3081 | C42+DHT |
| BIOCARTA_ | BIOCARTA_ | BIOCARTA_ | 10 | -0.816811 | -1.872352 | 0.000857 | 0.045464 | 0.039892 | 189 | tags=20%, | 330 | C42+DHT |
| BILD_HRAS | BILD_HRAS | BILD_HRAS | 173 | 0.504427 | 1.676272 | 0.000873 | 0.045464 | 0.039892 | 841 | tags=21%, | 129642/79 | C42+DHT |
| GAVIN_FO: | GAVIN_FO: | GAVIN_FO: | 66 | 0.629896 | 1.852005 | 0.000878 | 0.045464 | 0.039892 | 651 | tags=20%, | 23052/260 | C42+DHT |
| HOEGERKC | HOEGERKC | HOEGERKC | 15 | -0.762959 | -1.935714 | 0.000897 | 0.045464 | 0.039892 | 278 | tags=33%, | 4925/1944 | C42+DHT |
| BRACHAT_ | BRACHAT_ | BRACHAT_ | 17 | -0.734703 | -1.91783 | 0.000901 | 0.045464 | 0.039892 | 348 | tags=53%, | 3983/4193 | C42+DHT |
| STARK_PRE | STARK_PRE | STARK_PRE | 139 | 0.537993 | 1.744486 | 0.000906 | 0.045464 | 0.039892 | 1284 | tags=25%, | 58480/230 | C42+DHT |
| SENGUPTA | SENGUPTA | SENGUPTA | 254 | 0.469208 | 1.618801 | 0.000934 | 0.045464 | 0.039892 | 2060 | tags=30%, | 22936/659 | C42+DHT |
| REACTOME | REACTOME | REACTOME | 67 | -0.505958 | -1.778281 | 0.000945 | 0.045464 | 0.039892 | 1464 | tags=36%, | 8349/6442 | C42+DHT |
| CHARAFE_I | CHARAFE_I | CHARAFE_I | 337 | -0.393222 | -1.730267 | 0.000946 | 0.045464 | 0.039892 | 1863 | tags=33%, | 10653/674 | C42+DHT |
| RODWELL_ | RODWELL_ | RODWELL_ | 111 | 0.560245 | 1.767833 | 0.000946 | 0.045464 | 0.039892 | 1279 | tags=23%, | 7113/6675 | C42+DHT |
| GROSS_HYI | GROSS_HYI | GROSS_HYI | 109 | 0.56325 | 1.772796 | 0.000949 | 0.045464 | 0.039892 | 995 | tags=21%, | 10769/742 | C42+DHT |
| KOBAYASH | KOBAYASH | KOBAYASH | 74 | -0.515109 | -1.840794 | 0.000985 | 0.046678 | 0.040957 | 1474 | tags=39%, | 113146/50 | C42+DHT |
| PID_FAS_P. | PID_FAS_P. | PID_FAS_P. | 34 | -0.602814 | -1.833449 | 0.001022 | 0.047921 | 0.042048 | 182 | tags=12%, | 355/8503/3 | C42+DHT |

Table S6

| LNCaP_NCOR2_UP | LNCaP_NCOR2_DOWN | LNCaP_DHT_UP | LNCaP_DHT_DOWN | LNCaP_NCOR2_DHT_COMMON |
|---|---|---|---|---|
| NFKB2 | E2F7 | FOXA1 | EBF1 | CDX1 |
| FOXN2 | TBP | NANOS1 | RELB | NFKB2 |
| FOXO4 | CASP8AP2 | FOXA2 | BRCA1 | RUNX3 |
| TP63 | TBX5 | NR3C1 | SOX4 | PAX2 |
| HIVEP1 | BCLAF1 | CEBPB | CEBPA | FOXA1 |
| GATA3 | DLX1 | | FOXA2 | NFKB1 |
| IRF1 | POU2F2 | | HDAC2 | FOXP3 |
| FOXA1 | NPAT | | EP300 | GLIS1 |
| ERM | TAF7 | | TCF12 | GATA3 |
| | TCF3 | | GTF2B | PAX4 |
| | GATA4 | | | DBP |
| | NFYA | | | NKX3-1 |
| | E2F1 | | | CCNT2 |
| | TGIF1 | | | E2F1 |
| | PML | | | SMC1A |
| | POLR2A | | | E2F2 |
| | ARID3A | | | GATA2 |
| | NONO | | | |
| | ATF2 | | | |
| | MYBL2 | | | |
| | HMX3 | | | |
| | CEBPB | | | |
| | POU2F1 | | | |
| | HNF1B | | | |
| | ONECUT2 | | | |
| | SMAD1 | | | |
| | PROP1 | | | |
| | NFIC | | | |
| | TFAP2C | | | |
| | GATA1 | | | |
| | PIR | | | |
| | SIN3A | | | |
| | NHLH1 | | | |
| | GTF2B | | | |
| | FOXN1 | | | |
| | ZBTB7C | | | |
| | POLE4 | | | |
| | SNAI2 | | | |

| C42_NCOR2_UP | C42_NCOR2_DOWN | C42_DHT_UP | C42_DHT_DOWN | C42_NCOR2_DHT_COMMON |
|---|---|---|---|---|
| GATA5 | ATF4 | FOXA1 | SOX9 | FOXA2 |
| FOXM1 | E2F6 | CEBPB | HDAC2 | ERM |
| E2F4 | FOXD2 | TFAP2A | ETS2 | MYBL2 |
| HSF1 | MYB | HLF | TCF12 | HDAC2 |
| MYCN | TCF12 | CEBPD | EP300 | SMC1A |
| SMARCB1 | CEBPB | NR3C1 | ERM | EBF1 |
| ZNF652 | NFIB | SMC1A | EBF1 | SMC3 |
| | TRIP10 | RAD21 | NFIC | VDR |
| | HNF4A | | NR2F1 | RAD21 |
| | EBF1 | | RELA | TFAP4 |
| | ERM | | GATA3 | FOXK1 |
| | MYF5 | | | |
| | NR2F1 | | | |
| | TP53 | | | |
| | ZBTB7A | | | |
| | HOXB9 | | | |

Table S7

| miRNA | logFC | logPV | Treatment | Cell |
|---|---|---|---|---|
| hsa−let−7e−5p | 1.2087570 | 1.643947 | shNCOR2 | C42 |
| hsa−miR−10a−5p | −0.7527059 | 1.848248 | shNCOR2 | C42 |
| hsa−miR−125a−3p | 1.4515265 | 1.536970 | shNCOR2 | C42 |
| hsa−miR−126−3p | 0.7074766 | 1.180791 | shNCOR2 | C42 |
| hsa−miR−196a−5p | −1.7707560 | 1.519020 | shNCOR2 | C42 |
| hsa−miR−200a−3p | 0.9571056 | 1.500430 | shNCOR2 | C42 |
| hsa−miR−21−5p | 0.7891277 | 1.667443 | shNCOR2 | C42 |
| hsa−miR−221−3p | −0.7604984 | 1.372126 | shNCOR2 | C42 |
| hsa−miR−26b−5p | 0.7058614 | 1.112302 | shNCOR2 | C42 |
| hsa−miR−30d−5p | 0.6042127 | 1.019740 | shNCOR2 | C42 |
| hsa−miR−375 | 0.9336564 | 1.709246 | shNCOR2 | C42 |
| hsa−miR−484 | −0.7245349 | 1.396001 | shNCOR2 | C42 |
| hsa−miR−497−5p | 1.1798807 | 1.199564 | shNCOR2 | C42 |
| hsa−miR−582−3p | −1.2128948 | 1.025480 | shNCOR2 | C42 |
| hsa−miR−615−3p | −0.7109062 | 1.085126 | shNCOR2 | C42 |
| hsa−let−7c−3p | −0.9639780 | 1.320481 | shNCOR2_DHT | C42 |
| hsa−miR−181a−2−3p | −0.7361510 | 1.030782 | shNCOR2_DHT | C42 |
| hsa−miR−30a−3p | −1.0059203 | 1.106527 | shNCOR2_DHT | C42 |
| hsa−miR−30c−2−3p | −1.8125451 | 1.287132 | shNCOR2_DHT | C42 |
| hsa−miR−454−3p | 0.5840674 | 1.067151 | shNCOR2_DHT | C42 |
| hsa−miR−671−3p | −0.5594290 | 1.111110 | shNCOR2_DHT | C42 |
| hsa−miR−10a−5p | −0.4565993 | 1.112900 | shNCOR2 | LNCaP |
| hsa−miR−22−3p | 0.3762707 | 1.181616 | shNCOR2 | LNCaP |

| Seed | miRNA family | p-value | FDR | Odd ratio | Number of interactions |
|---|---|---|---|---|---|
| GAGGUAG | hsa-let-7a-5p/hsa-let-7b-5p/hsa-let-7c-5p/hsa-let-7d-5p/hsa-let-7 | 8.60201E-05 | 0.003957824 | 0.751847056 | 153 |
| AAUACUG | hsa-miR-200b-3p/hsa-miR-200c-3p/hsa-miR-429 | 1.95653E-05 | 0.002103264 | 0.716623357 | 139 |
| ACAUUCA | hsa-miR-181a-5p/hsa-miR-181b-5p/hsa-miR-181c-5p/hsa-miR-18 | 0.000110451 | 0.003957824 | 0.722113983 | 120 |
| UAAGGCA | hsa-miR-124-3p.2/hsa-miR-506-3p | 9.42949E-05 | 0.003957824 | 0.715780005 | 117 |
| CUUUGGU | hsa-miR-9-5p | 0.00060225 | 0.009960294 | 0.749813319 | 117 |
| GUGCAAA | hsa-miR-19a-3p/hsa-miR-19b-3p | 0.000491697 | 0.009960294 | 0.742378305 | 114 |
| UCACAGU | hsa-miR-27a-3p/hsa-miR-27b-3p | 0.003910506 | 0.036554727 | 0.787841353 | 114 |
| GUAAACA | hsa-miR-30a-5p/hsa-miR-30b-5p/hsa-miR-30c-5p/hsa-miR-30d-5 | 0.004551868 | 0.037093819 | 0.788635533 | 111 |
| UUGGCAA | hsa-miR-182-5p | 0.002376035 | 0.023878276 | 0.770633757 | 109 |
| UAUUGCU | hsa-miR-137 | 0.002322791 | 0.023878276 | 0.768990821 | 108 |
| CACAGUG | hsa-miR-128-3p/hsa-miR-216a-3p/hsa-miR-3681-3p | 0.000974158 | 0.013962938 | 0.744478732 | 104 |
| UUGGCAC | hsa-miR-1271-5p/hsa-miR-96-5p | 0.000579415 | 0.009960294 | 0.732070262 | 103 |
| AAGUGCU | hsa-miR-302a-3p/hsa-miR-302b-3p/hsa-miR-302c-3p.1/hsa-miR-3 | 0.000168902 | 0.005187712 | 0.692533492 | 93 |
| AGUGCAA | hsa-miR-130a-3p/hsa-miR-130b-3p/hsa-miR-301a-3p/hsa-miR-30 | 0.000237923 | 0.006394167 | 0.699329699 | 93 |
| UCAAGUA | hsa-miR-1297/hsa-miR-26a-5p/hsa-miR-26b-5p/hsa-miR-4465 | 0.00624917 | 0.043341019 | 0.768010903 | 86 |
| ACAGUAU | hsa-miR-144-3p | 0.00672897 | 0.04384026 | 0.770215719 | 86 |
| UGGCACU | hsa-miR-183-5p.2 | 1.55085E-06 | 0.000333432 | 0.598585925 | 85 |
| AUUGCAC | hsa-miR-25-3p/hsa-miR-32-5p/hsa-miR-363-3p/hsa-miR-367-3p/ | 0.007987703 | 0.049067316 | 0.774071985 | 85 |
| AUAAAGU | hsa-miR-142-5p/hsa-miR-5590-3p | 0.004658294 | 0.037093819 | 0.75055611 | 80 |
| UGCAUAG | hsa-miR-153-3p | 0.002443358 | 0.023878276 | 0.727264758 | 77 |
| UCCAGUU | hsa-miR-145-5p/hsa-miR-5195-3p | 0.006492914 | 0.043624263 | 0.754298495 | 76 |
| CAGUGCA | hsa-miR-148a-3p/hsa-miR-148b-3p/hsa-miR-152-3p | 0.000585016 | 0.009960294 | 0.68500256 | 74 |
| UAAUACA | hsa-miR-374c-5p/hsa-miR-655-3p | 0.004220145 | 0.037093819 | 0.735945428 | 73 |
| AUGGCUU | hsa-miR-135a-5p/hsa-miR-135b-5p | 0.005876577 | 0.043341019 | 0.744411206 | 72 |
| UCCCUUU | hsa-miR-204-5p/hsa-miR-211-5p | 0.002438322 | 0.023878276 | 0.714213393 | 70 |
| GGCAGUG | hsa-miR-34a-5p/hsa-miR-34c-5p/hsa-miR-449a/hsa-miR-449b-5p | 0.006228532 | 0.043341019 | 0.73317481 | 65 |
| UGAAACA | hsa-miR-653-5p | 0.000966629 | 0.013962938 | 0.663113979 | 59 |
| UACAGUU | hsa-miR-582-5p | 0.001903017 | 0.023878276 | 0.682396777 | 59 |
| AUAAUAC | hsa-miR-369-3p | 0.004502986 | 0.037093819 | 0.70202391 | 56 |
| UGCAUUG | hsa-miR-33a-5p/hsa-miR-33b-5p | 0.00110649 | 0.014868465 | 0.64993538 | 53 |
| GAUUGUC | hsa-miR-219a-5p/hsa-miR-4782-3p/hsa-miR-6766-3p | 3.16489E-05 | 0.00226817 | 0.556449442 | 51 |
| UGAAAGG | hsa-miR-488-3p | 0.00771969 | 0.048815684 | 0.676753031 | 41 |
| UCAACAC | hsa-miR-505-3p.2 | 0.002159698 | 0.023878276 | 0.620702812 | 39 |
| UCAUAGA | hsa-miR-376a-3p/hsa-miR-376b-3p | 0.000423298 | 0.009960294 | 0.526026421 | 31 |

DMR-enh_LNCaP Table S8

| | ID | Descriptior | GeneRatio | BgRatio | pvalue | p.adjust | qvalue | geneID | Count | Group |
|---|---|---|---|---|---|---|---|---|---|---|
| CHARAFE_I | CHARAFE_I | CHARAFE_I | 17/196 | 451/20690 | 1.50E-06 | 0.00362 | 0.00311 | 84988/27 | 17 | LNCaP_DMR.enh |
| HALLMARK | HALLMARK | HALLMARK | 8/196 | 97/20690 | 3.95E-06 | 0.00362 | 0.00311 | 841/3430/ | 8 | LNCaP_DMR.enh |
| GRAESSMA | GRAESSMA | GRAESSMA | 19/196 | 596/20690 | 4.21E-06 | 0.00362 | 0.00311 | 79038/15 | 19 | LNCaP_DMR.enh |
| RADAEVA_ | RADAEVA_ | RADAEVA_ | 6/196 | 48/20690 | 5.90E-06 | 0.003802 | 0.003267 | 11068/34 | 6 | LNCaP_DMR.enh |
| DOANE_BR | DOANE_BR | DOANE_BR | 8/196 | 111/20690 | 1.08E-05 | 0.005562 | 0.004778 | 27134/70 | 8 | LNCaP_DMR.enh |
| BROWNE_I | BROWNE_I | BROWNE_I | 16/196 | 482/20690 | 1.52E-05 | 0.006536 | 0.005615 | 115817/2 | 16 | LNCaP_DMR.enh |
| CHARAFE_I | CHARAFE_I | CHARAFE_I | 14/196 | 383/20690 | 1.83E-05 | 0.006758 | 0.005806 | 84988/27 | 14 | LNCaP_DMR.enh |
| MARSON_F | MARSON_F | MARSON_F | 6/196 | 62/20690 | 2.65E-05 | 0.007913 | 0.006798 | 6004/464 | 6 | LNCaP_DMR.enh |
| GRUETZMA | GRUETZMA | GRUETZMA | 13/196 | 346/20690 | 2.76E-05 | 0.007913 | 0.006798 | 7033/600 | 13 | LNCaP_DMR.enh |
| JAEGER_M | JAEGER_M | JAEGER_M | 11/196 | 258/20690 | 3.72E-05 | 0.009097 | 0.007816 | 1241/407 | 11 | LNCaP_DMR.enh |
| SMID_BREA | SMID_BREA | SMID_BREA | 19/196 | 699/20690 | 3.88E-05 | 0.009097 | 0.007816 | 27134/70 | 19 | LNCaP_DMR.enh |
| HOLLMANN | HOLLMANN | HOLLMANN | 11/196 | 266/20690 | 4.90E-05 | 0.010365 | 0.008904 | 9123/299 | 11 | LNCaP_DMR.enh |
| ENK_UV_R | ENK_UV_R | ENK_UV_R | 16/196 | 534/20690 | 5.23E-05 | 0.010365 | 0.008904 | 4833/600 | 16 | LNCaP_DMR.enh |
| MULLIGHA | MULLIGHA | MULLIGHA | 12/196 | 332/20690 | 8.17E-05 | 0.015036 | 0.012917 | 3203/648 | 12 | LNCaP_DMR.enh |
| STAMBOLS | STAMBOLS | STAMBOLS | 5/196 | 51/20690 | 0.00012 | 0.018211 | 0.015645 | 25771/30 | 5 | LNCaP_DMR.enh |
| HALLMARK | HALLMARK | HALLMARK | 9/196 | 200/20690 | 0.000127 | 0.018211 | 0.015645 | 7517/271 | 9 | LNCaP_DMR.enh |
| HALLMARK | HALLMARK | HALLMARK | 9/196 | 200/20690 | 0.000127 | 0.018211 | 0.015645 | 6450/172 | 9 | LNCaP_DMR.enh |
| COLINA_TA | COLINA_TA | COLINA_TA | 12/196 | 348/20690 | 0.000127 | 0.018211 | 0.015645 | 9123/115 | 12 | LNCaP_DMR.enh |
| WANG_ESC | WANG_ESC | WANG_ESC | 7/196 | 119/20690 | 0.000141 | 0.019109 | 0.016417 | 2001/571 | 7 | LNCaP_DMR.enh |
| SANA_TNF_ | SANA_TNF_ | SANA_TNF_ | 6/196 | 85/20690 | 0.000157 | 0.020277 | 0.01742 | 4071/172 | 6 | LNCaP_DMR.enh |
| MULLIGHA | MULLIGHA | MULLIGHA | 10/196 | 262/20690 | 0.000208 | 0.025499 | 0.021906 | 3203/464 | 10 | LNCaP_DMR.enh |
| PID_INTEG | PID_INTEG | PID_INTEG | 4/196 | 33/20690 | 0.000258 | 0.029503 | 0.025345 | 9564/671 | 4 | LNCaP_DMR.enh |
| LEI_MYB_T | LEI_MYB_T | LEI_MYB_T | 11/196 | 322/20690 | 0.000263 | 0.029503 | 0.025345 | 9123/407 | 11 | LNCaP_DMR.enh |
| KIM_BIPOL | KIM_BIPOL | KIM_BIPOL | 17/196 | 681/20690 | 0.000275 | 0.029531 | 0.02537 | 23396/15 | 17 | LNCaP_DMR.enh |
| FEVR_CTNI | FEVR_CTNI | FEVR_CTNI | 17/196 | 687/20690 | 0.000304 | 0.031372 | 0.026952 | 84988/28 | 17 | LNCaP_DMR.enh |
| ZHOU_INFL | ZHOU_INFL | ZHOU_INFL | 13/196 | 443/20690 | 0.000325 | 0.032229 | 0.027688 | 1263/600 | 13 | LNCaP_DMR.enh |
| GRANDVAL | GRANDVAL | GRANDVAL | 3/196 | 15/20690 | 0.00035 | 0.032258 | 0.027713 | 3430/851 | 3 | LNCaP_DMR.enh |
| LI_AMPLIFI | LI_AMPLIFI | LI_AMPLIFI | 8/196 | 182/20690 | 0.000353 | 0.032258 | 0.027713 | 7033/233 | 8 | LNCaP_DMR.enh |
| YANG_BRE. | YANG_BRE. | YANG_BRE. | 4/196 | 36/20690 | 0.000363 | 0.032258 | 0.027713 | 7033/633 | 4 | LNCaP_DMR.enh |
| REACTOME | REACTOME | REACTOME | 8/196 | 185/20690 | 0.000394 | 0.03317 | 0.028496 | 6484/163 | 8 | LNCaP_DMR.enh |
| WANG_BAI | WANG_BAI | WANG_BAI | 4/196 | 37/20690 | 0.000404 | 0.03317 | 0.028496 | 6484/409 | 4 | LNCaP_DMR.enh |
| BROWNE_I | BROWNE_I | BROWNE_I | 6/196 | 102/20690 | 0.000424 | 0.03317 | 0.028496 | 23321/17 | 6 | LNCaP_DMR.enh |
| BIOCARTA_ | BIOCARTA_ | BIOCARTA_ | 3/196 | 16/20690 | 0.000428 | 0.03317 | 0.028496 | 1018/281 | 3 | LNCaP_DMR.enh |
| MARTINEZ_ | MARTINEZ_ | MARTINEZ_ | 15/196 | 580/20690 | 0.000438 | 0.03317 | 0.028496 | 23396/50 | 15 | LNCaP_DMR.enh |
| MIKKELSEN | MIKKELSEN | MIKKELSEN | 17/196 | 711/20690 | 0.00045 | 0.03317 | 0.028496 | 27134/64 | 17 | LNCaP_DMR.enh |
| CREIGHTOI | CREIGHTOI | CREIGHTOI | 17/196 | 721/20690 | 0.000527 | 0.036132 | 0.031041 | 9123/200 | 17 | LNCaP_DMR.enh |
| CASTELLAN | CASTELLAN | CASTELLAN | 5/196 | 70/20690 | 0.000533 | 0.036132 | 0.031041 | 1728/139 | 5 | LNCaP_DMR.enh |
| BOYAULT_I | BOYAULT_I | BOYAULT_I | 4/196 | 40/20690 | 0.000547 | 0.036132 | 0.031041 | 23321/85 | 4 | LNCaP_DMR.enh |
| TRAYNOR_ | TRAYNOR_ | TRAYNOR_ | 4/196 | 40/20690 | 0.000547 | 0.036132 | 0.031041 | 1803/771/ | 4 | LNCaP_DMR.enh |
| ZHU_CMV_ | ZHU_CMV_ | ZHU_CMV_ | 6/196 | 108/20690 | 0.000575 | 0.037067 | 0.031844 | 3612/667 | 6 | LNCaP_DMR.enh |
| HALLMARK | HALLMARK | HALLMARK | 8/196 | 200/20690 | 0.000659 | 0.040665 | 0.034935 | 841/8673/ | 8 | LNCaP_DMR.enh |
| WEST_ADR | WEST_ADR | WEST_ADR | 14/196 | 540/20690 | 0.000662 | 0.040665 | 0.034935 | 2875/115 | 14 | LNCaP_DMR.enh |
| DER_IFN_A | DER_IFN_A | DER_IFN_A | 5/196 | 74/20690 | 0.000688 | 0.040837 | 0.035083 | 841/3430/ | 5 | LNCaP_DMR.enh |
| LIEN_BREA | LIEN_BREA | LIEN_BREA | 6/196 | 112/20690 | 0.000697 | 0.040837 | 0.035083 | 3169/703 | 6 | LNCaP_DMR.enh |
| ONDER_CD | ONDER_CD | ONDER_CD | 9/196 | 256/20690 | 0.000776 | 0.043241 | 0.037148 | 155066/2 | 9 | LNCaP_DMR.enh |
| HALLMARK | HALLMARK | HALLMARK | 7/196 | 158/20690 | 0.000787 | 0.043241 | 0.037148 | 2907/887 | 7 | LNCaP_DMR.enh |
| KEGG_VAS | KEGG_VAS | KEGG_VAS | 4/196 | 44/20690 | 0.000788 | 0.043241 | 0.037148 | 6844/556 | 4 | LNCaP_DMR.enh |
| ROZANOV_ | ROZANOV_ | ROZANOV_ | 9/196 | 261/20690 | 0.000889 | 0.047759 | 0.041029 | 6337/771/ | 9 | LNCaP_DMR.enh |
| HOLLERN_I | HOLLERN_I | HOLLERN_I | 6/196 | 118/20690 | 0.000916 | 0.047759 | 0.041029 | 27134/20 | 6 | LNCaP_DMR.enh |
| PID_P73PA | PID_P73PA | PID_P73PA | 5/196 | 79/20690 | 0.000926 | 0.047759 | 0.041029 | 1263/556 | 5 | LNCaP_DMR.enh |
| REACTOME | REACTOME | REACTOME | 3/196 | 21/20690 | 0.000982 | 0.049631 | 0.042637 | 6484/252 | 3 | LNCaP_DMR.enh |

DMR-enh_LNCaP

| | ID | Descriptior | GeneRatio | BgRatio | pvalue | p.adjust | qvalue | geneID | Count | Group |
|---|---|---|---|---|---|---|---|---|---|---|
| GRAESSMA | GRAESSMA | GRAESSMA | 74/950 | 596/20690 | 5.75E-15 | 2.42E-11 | 1.86E-11 | 10417/139 | 74 | C42_DMR.enh |
| NIKOLSKY_ | NIKOLSKY_ | NIKOLSKY_ | 30/950 | 119/20690 | 1.17E-14 | 2.47E-11 | 1.90E-11 | 10607/899 | 30 | C42_DMR.enh |
| CHARAFE_I | CHARAFE_I | CHARAFE_I | 61/950 | 451/20690 | 3.38E-14 | 4.73E-11 | 3.64E-11 | 7033/8498 | 61 | C42_DMR.enh |
| ENK_UV_R | ENK_UV_R | ENK_UV_R | 65/950 | 534/20690 | 6.91E-13 | 7.25E-10 | 5.59E-10 | 6002/6150 | 65 | C42_DMR.enh |
| DELACROIX | DELACROIX | DELACROIX | 49/950 | 361/20690 | 1.04E-11 | 8.73E-09 | 6.73E-09 | 51299/468 | 49 | C42_DMR.enh |
| LIM_MAMI | LIM_MAMI | LIM_MAMI | 50/950 | 408/20690 | 2.62E-10 | 1.84E-07 | 1.42E-07 | 5590/8470 | 50 | C42_DMR.enh |
| WEST_ADR | WEST_ADR | WEST_ADR | 58/950 | 540/20690 | 1.68E-09 | 1.01E-06 | 7.77E-07 | 2550/975/1 | 58 | C42_DMR.enh |
| DACOSTA_ | DACOSTA_ | DACOSTA_ | 41/950 | 316/20690 | 2.02E-09 | 1.06E-06 | 8.15E-07 | 25855/569 | 41 | C42_DMR.enh |
| HALLMARK | HALLMARK | HALLMARK | 27/950 | 158/20690 | 3.31E-09 | 1.54E-06 | 1.19E-06 | 1543/6150 | 27 | C42_DMR.enh |
| SMID_BREA | SMID_BREA | SMID_BREA | 68/950 | 699/20690 | 4.23E-09 | 1.77E-06 | 1.37E-06 | 7033/2678 | 68 | C42_DMR.enh |
| DELYS_THY | DELYS_THY | DELYS_THY | 50/950 | 445/20690 | 5.17E-09 | 1.97E-06 | 1.52E-06 | 1263/9961 | 50 | C42_DMR.enh |
| FEVR_CTNI | FEVR_CTNI | FEVR_CTNI | 65/950 | 687/20690 | 2.74E-08 | 9.58E-06 | 7.38E-06 | 2550/6293 | 65 | C42_DMR.enh |
| JOHNSTON | JOHNSTON | JOHNSTON | 46/950 | 432/20690 | 1.07E-07 | 3.47E-05 | 2.67E-05 | 4681/8979 | 46 | C42_DMR.enh |
| HUTTMANI | HUTTMANI | HUTTMANI | 35/950 | 285/20690 | 1.21E-07 | 3.47E-05 | 2.67E-05 | 7474/2332 | 35 | C42_DMR.enh |
| MULLIGHA | MULLIGHA | MULLIGHA | 45/950 | 421/20690 | 1.32E-07 | 3.47E-05 | 2.67E-05 | 9123/8030 | 45 | C42_DMR.enh |
| SARRIO_EP | SARRIO_EP | SARRIO_EP | 24/950 | 153/20690 | 1.32E-07 | 3.47E-05 | 2.67E-05 | 288/604/9 | 24 | C42_DMR.enh |
| HALLMARK | HALLMARK | HALLMARK | 28/950 | 200/20690 | 1.47E-07 | 3.63E-05 | 2.80E-05 | 25937/263 | 28 | C42_DMR.enh |
| CREIGHTOI | CREIGHTOI | CREIGHTOI | 65/950 | 721/20690 | 1.65E-07 | 3.76E-05 | 2.89E-05 | 81846/468 | 65 | C42_DMR.enh |
| LI_INDUCEI | LI_INDUCEI | LI_INDUCEI | 36/950 | 302/20690 | 1.70E-07 | 3.76E-05 | 2.89E-05 | 58496/996 | 36 | C42_DMR.enh |
| WONG_AD | WONG_AD | WONG_AD | 64/950 | 710/20690 | 2.06E-07 | 4.34E-05 | 3.34E-05 | 2550/5178 | 64 | C42_DMR.enh |
| ONDER_CD | ONDER_CD | ONDER_CD | 48/950 | 473/20690 | 2.44E-07 | 4.88E-05 | 3.76E-05 | 288/3985/ | 48 | C42_DMR.enh |
| LIAO_META | LIAO_META | LIAO_META | 52/950 | 535/20690 | 3.01E-07 | 5.74E-05 | 4.42E-05 | 2550/1041 | 52 | C42_DMR.enh |
| REACTOME | REACTOME | REACTOME | 53/950 | 554/20690 | 3.81E-07 | 6.95E-05 | 5.35E-05 | 6222/2668 | 53 | C42_DMR.enh |
| RICKMAN_ | RICKMAN_ | RICKMAN_ | 32/950 | 261/20690 | 4.34E-07 | 7.60E-05 | 5.85E-05 | 9123/6891 | 32 | C42_DMR.enh |
| RODRIGUES | RODRIGUES | RODRIGUES | 20/950 | 119/20690 | 4.67E-07 | 7.85E-05 | 6.05E-05 | 23321/604 | 20 | C42_DMR.enh |
| HALLMARK | HALLMARK | HALLMARK | 27/950 | 200/20690 | 5.12E-07 | 8.28E-05 | 6.38E-05 | 10891/704 | 27 | C42_DMR.enh |
| GRAESSMA | GRAESSMA | GRAESSMA | 29/950 | 228/20690 | 6.97E-07 | 0.000108 | 8.35E-05 | 57332/991 | 29 | C42_DMR.enh |
| CHARAFE_I | CHARAFE_I | CHARAFE_I | 40/950 | 383/20690 | 1.19E-06 | 0.000178 | 0.000137 | 7033/8498 | 40 | C42_DMR.enh |
| UROSEVIC_ | UROSEVIC_ | UROSEVIC_ | 8/950 | 20/20690 | 1.48E-06 | 0.000214 | 0.000165 | 8519/4599 | 8 | C42_DMR.enh |
| COLDREN_ | COLDREN_ | COLDREN_ | 28/950 | 225/20690 | 1.67E-06 | 0.000234 | 0.000181 | 64063/991 | 28 | C42_DMR.enh |
| KEGG_AXO | KEGG_AXO | KEGG_AXO | 20/950 | 129/20690 | 1.75E-06 | 0.000237 | 0.000182 | 3985/2051 | 20 | C42_DMR.enh |
| COLINA_TA | COLINA_TA | COLINA_TA | 37/950 | 348/20690 | 1.94E-06 | 0.000242 | 0.000186 | 7474/5129 | 37 | C42_DMR.enh |
| SANSOM_A | SANSOM_A | SANSOM_A | 37/950 | 348/20690 | 1.94E-06 | 0.000242 | 0.000186 | 4681/6484 | 37 | C42_DMR.enh |
| CAIRO_HEF | CAIRO_HEF | CAIRO_HEF | 27/950 | 214/20690 | 1.96E-06 | 0.000242 | 0.000186 | 975/4681/ | 27 | C42_DMR.enh |
| FERNANDE | FERNANDE | FERNANDE | 24/950 | 180/20690 | 2.71E-06 | 0.000325 | 0.000251 | 9910/7043 | 24 | C42_DMR.enh |
| DOANE_BR | DOANE_BR | DOANE_BR | 18/950 | 111/20690 | 2.95E-06 | 0.000344 | 0.000265 | 7033/1396 | 18 | C42_DMR.enh |
| DUTERTRE_ | DUTERTRE_ | DUTERTRE_ | 47/950 | 504/20690 | 3.50E-06 | 0.000397 | 0.000306 | 288/1543/ | 47 | C42_DMR.enh |
| BENNETT_S | BENNETT_S | BENNETT_S | 9/950 | 29/20690 | 3.81E-06 | 0.00041 | 0.000316 | 5359/1041 | 9 | C42_DMR.enh |
| PID_IGF1_F | PID_IGF1_F | PID_IGF1_F | 9/950 | 29/20690 | 3.81E-06 | 0.00041 | 0.000316 | 5590/9564 | 9 | C42_DMR.enh |
| HOSHIDA_I | HOSHIDA_I | HOSHIDA_I | 28/950 | 235/20690 | 3.95E-06 | 0.000414 | 0.000319 | 7033/4681 | 28 | C42_DMR.enh |
| BERENJENC | BERENJENC | BERENJENC | 39/950 | 389/20690 | 4.34E-06 | 0.000436 | 0.000336 | 288/4837/ | 39 | C42_DMR.enh |
| SAGIV_CD2 | SAGIV_CD2 | SAGIV_CD2 | 11/950 | 45/20690 | 4.36E-06 | 0.000436 | 0.000336 | 2051/2635 | 11 | C42_DMR.enh |
| ACEVEDO_ | ACEVEDO_ | ACEVEDO_ | 49/950 | 540/20690 | 4.65E-06 | 0.000454 | 0.00035 | 975/7033/ | 49 | C42_DMR.enh |
| SWEET_LU | SWEET_LU | SWEET_LU | 40/950 | 407/20690 | 5.33E-06 | 0.00049 | 0.000378 | 5178/5699 | 40 | C42_DMR.enh |
| BROWNE_I | BROWNE_I | BROWNE_I | 45/950 | 482/20690 | 5.46E-06 | 0.00049 | 0.000378 | 3985/1158 | 45 | C42_DMR.enh |
| HALLMARK | HALLMARK | HALLMARK | 25/950 | 200/20690 | 5.49E-06 | 0.00049 | 0.000378 | 975/1263/ | 25 | C42_DMR.enh |
| HALLMARK | HALLMARK | HALLMARK | 25/950 | 200/20690 | 5.49E-06 | 0.00049 | 0.000378 | 5699/1543 | 25 | C42_DMR.enh |
| NIKOLSKY_ | NIKOLSKY_ | NIKOLSKY_ | 10/950 | 38/20690 | 5.80E-06 | 0.000499 | 0.000385 | 90780/646 | 10 | C42_DMR.enh |
| LEI_MYB_T | LEI_MYB_T | LEI_MYB_T | 34/950 | 322/20690 | 5.83E-06 | 0.000499 | 0.000385 | 7033/3985 | 34 | C42_DMR.enh |
| PID_LYSOP | PID_LYSOP | PID_LYSOP | 13/950 | 65/20690 | 6.62E-06 | 0.000556 | 0.000429 | 9564/7205 | 13 | C42_DMR.enh |
| PID_RET_P | PID_RET_P | PID_RET_P | 10/950 | 39/20690 | 7.49E-06 | 0.000617 | 0.000475 | 2668/9564 | 10 | C42_DMR.enh |
| ROZANOV_ | ROZANOV_ | ROZANOV_ | 29/950 | 261/20690 | 1.06E-05 | 0.000853 | 0.000657 | 4837/5507 | 29 | C42_DMR.enh |
| KAAB_HEA | KAAB_HEA | KAAB_HEA | 29/950 | 262/20690 | 1.14E-05 | 0.000901 | 0.000694 | 2875/1057 | 29 | C42_DMR.enh |
| REACTOME | REACTOME | REACTOME | 6/950 | 13/20690 | 1.20E-05 | 0.000932 | 0.000718 | 8717/9099 | 6 | C42_DMR.enh |
| HAN_SATB | HAN_SATB | HAN_SATB | 40/950 | 423/20690 | 1.33E-05 | 0.001017 | 0.000784 | 7474/2585 | 40 | C42_DMR.enh |
| RICKMAN_ | RICKMAN_ | RICKMAN_ | 37/950 | 379/20690 | 1.38E-05 | 0.001036 | 0.000798 | 1241/3754 | 37 | C42_DMR.enh |

| | | | | | | | | | | | |
|---|---|---|---|---|---|---|---|---|---|---|---|
| SHIN_B_CE | SHIN_B_CE | SHIN_B_CE | 8/950 | 26/20690 | 1.43E-05 | 0.001054 | 0.000812 | 604/7517/! | | 8 | C42_DMR.enh |
| MIKKELSEN | MIKKELSEN | MIKKELSEN | 58/950 | 711/20690 | 1.67E-05 | 0.001197 | 0.000922 | 288/23321, | | 58 | C42_DMR.enh |
| HALLMARK | HALLMARK | HALLMARK | 24/950 | 200/20690 | 1.68E-05 | 0.001197 | 0.000922 | 7726/5699, | | 24 | C42_DMR.enh |
| MULLIGHA | MULLIGHA | MULLIGHA | 37/950 | 383/20690 | 1.74E-05 | 0.00122 | 0.00094 | 9123/1241, | | 37 | C42_DMR.enh |
| ELVIDGE_H | ELVIDGE_H | ELVIDGE_H | 15/950 | 92/20690 | 1.82E-05 | 0.001252 | 0.000964 | 1543/4016, | | 15 | C42_DMR.enh |
| KAAB_HEA | KAAB_HEA | KAAB_HEA | 28/950 | 256/20690 | 2.00E-05 | 0.001344 | 0.001036 | 975/6910/! | | 28 | C42_DMR.enh |
| ZHANG_AD | ZHANG_AD | ZHANG_AD | 6/950 | 14/20690 | 2.02E-05 | 0.001344 | 0.001036 | 4641/3265, | | 6 | C42_DMR.enh |
| KIM_WT1_ | KIM_WT1_ | KIM_WT1_ | 25/950 | 217/20690 | 2.28E-05 | 0.001465 | 0.001129 | 84159/259 | | 25 | C42_DMR.enh |
| SCHAEFFER | SCHAEFFER | SCHAEFFER | 40/950 | 433/20690 | 2.29E-05 | 0.001465 | 0.001129 | 51299/796, | | 40 | C42_DMR.enh |
| JAEGER_M | JAEGER_M | JAEGER_M | 28/950 | 258/20690 | 2.30E-05 | 0.001465 | 0.001129 | 7474/288/2 | | 28 | C42_DMR.enh |
| MEISSNER_ | MEISSNER_ | MEISSNER_ | 43/950 | 480/20690 | 2.40E-05 | 0.001505 | 0.00116 | 8705/6464 | | 43 | C42_DMR.enh |
| PID_PDGFR | PID_PDGFR | PID_PDGFR | 18/950 | 129/20690 | 2.51E-05 | 0.001551 | 0.001195 | 9564/5155, | | 18 | C42_DMR.enh |
| ZHANG_AN | ZHANG_AN | ZHANG_AN | 8/950 | 28/20690 | 2.62E-05 | 0.001577 | 0.001215 | 975/6662/8 | | 8 | C42_DMR.enh |
| LU_EZH2_T | LU_EZH2_T | LU_EZH2_T | 30/950 | 288/20690 | 2.63E-05 | 0.001577 | 0.001215 | 5699/1136 | | 30 | C42_DMR.enh |
| WANG_TUI | WANG_TUI | WANG_TUI | 35/950 | 361/20690 | 2.71E-05 | 0.001598 | 0.001231 | 2550/4720, | | 35 | C42_DMR.enh |
| HOLLERN_I | HOLLERN_I | HOLLERN_I | 17/950 | 118/20690 | 2.75E-05 | 0.001598 | 0.001231 | 84283/288 | | 17 | C42_DMR.enh |
| TONKS_TAI | TONKS_TAI | TONKS_TAI | 23/950 | 193/20690 | 2.80E-05 | 0.001598 | 0.001231 | 9123/3757, | | 23 | C42_DMR.enh |
| HALLMARK | HALLMARK | HALLMARK | 11/950 | 54/20690 | 2.81E-05 | 0.001598 | 0.001231 | 25937/649 | | 11 | C42_DMR.enh |
| PETROVA_I | PETROVA_I | PETROVA_I | 18/950 | 131/20690 | 3.10E-05 | 0.001736 | 0.001338 | 7033/2290 | | 18 | C42_DMR.enh |
| LIM_MAMI | LIM_MAMI | LIM_MAMI | 42/950 | 470/20690 | 3.15E-05 | 0.001743 | 0.001343 | 5178/4681, | | 42 | C42_DMR.enh |
| GRANDVAL | GRANDVAL | GRANDVAL | 6/950 | 15/20690 | 3.23E-05 | 0.001762 | 0.001357 | 8519/1041 | | 6 | C42_DMR.enh |
| ANDERSEN | ANDERSEN | ANDERSEN | 13/950 | 75/20690 | 3.34E-05 | 0.001798 | 0.001385 | 2875/2714 | | 13 | C42_DMR.enh |
| TSUDA_ALV | TSUDA_ALV | TSUDA_ALV | 5/950 | 10/20690 | 4.20E-05 | 0.002206 | 0.0017 | 26353/559 | | 5 | C42_DMR.enh |
| EBAUER_TA | EBAUER_TA | EBAUER_TA | 23/950 | 198/20690 | 4.20E-05 | 0.002206 | 0.0017 | 22885/259 | | 23 | C42_DMR.enh |
| KEGG_PATI | KEGG_PATI | KEGG_PATI | 32/950 | 325/20690 | 4.38E-05 | 0.002271 | 0.00175 | 7474/7043, | | 32 | C42_DMR.enh |
| LINDGREN_ | LINDGREN_ | LINDGREN_ | 25/950 | 226/20690 | 4.52E-05 | 0.002299 | 0.001771 | 975/6002/2 | | 25 | C42_DMR.enh |
| FISCHER_D | FISCHER_D | FISCHER_D | 31/950 | 311/20690 | 4.54E-05 | 0.002299 | 0.001771 | 1263/604/3 | | 31 | C42_DMR.enh |
| PHONG_TN | PHONG_TN | PHONG_TN | 33/950 | 341/20690 | 4.72E-05 | 0.002359 | 0.001817 | 1543/8415 | | 33 | C42_DMR.enh |
| HALLMARK | HALLMARK | HALLMARK | 16/950 | 112/20690 | 5.22E-05 | 0.002578 | 0.001986 | 9409/5827, | | 16 | C42_DMR.enh |
| RADAEVA_ | RADAEVA_ | RADAEVA_ | 10/950 | 48/20690 | 5.29E-05 | 0.002585 | 0.001991 | 11068/535 | | 10 | C42_DMR.enh |
| ZHANG_INT | ZHANG_INT | ZHANG_INT | 7/950 | 23/20690 | 5.40E-05 | 0.002587 | 0.001993 | 8519/1041 | | 7 | C42_DMR.enh |
| ENK_UV_R | ENK_UV_R | ENK_UV_R | 44/950 | 513/20690 | 5.48E-05 | 0.002587 | 0.001993 | 10417/287 | | 44 | C42_DMR.enh |
| LOCKWOOI | LOCKWOOI | LOCKWOOI | 24/950 | 215/20690 | 5.51E-05 | 0.002587 | 0.001993 | 84988/615 | | 24 | C42_DMR.enh |
| SWEET_LU | SWEET_LU | SWEET_LU | 41/950 | 466/20690 | 5.54E-05 | 0.002587 | 0.001993 | 6222/2583, | | 41 | C42_DMR.enh |
| HUMMERI( | HUMMERI( | HUMMERI( | 15/950 | 101/20690 | 5.62E-05 | 0.002594 | 0.001998 | 2678/5598, | | 15 | C42_DMR.enh |
| CHARAFE_I | CHARAFE_I | CHARAFE_I | 17/950 | 125/20690 | 5.83E-05 | 0.002659 | 0.002049 | 3985/5496 | | 17 | C42_DMR.enh |
| HELLER_HD | HELLER_HD | HELLER_HD | 31/950 | 316/20690 | 6.14E-05 | 0.002773 | 0.002136 | 604/1543/! | | 31 | C42_DMR.enh |
| KOBAYASH | KOBAYASH | KOBAYASH | 15/950 | 102/20690 | 6.31E-05 | 0.002821 | 0.002174 | 5699/2332 | | 15 | C42_DMR.enh |
| MCBRYAN_ | MCBRYAN_ | MCBRYAN_ | 27/950 | 259/20690 | 6.45E-05 | 0.002849 | 0.002195 | 10417/153 | | 27 | C42_DMR.enh |
| KINSEY_TA | KINSEY_TA | KINSEY_TA | 32/950 | 332/20690 | 6.60E-05 | 0.002849 | 0.002195 | 4681/8415 | | 32 | C42_DMR.enh |
| PID_ER_NC | PID_ER_NC | PID_ER_NC | 9/950 | 40/20690 | 6.60E-05 | 0.002849 | 0.002195 | 9564/6464, | | 9 | C42_DMR.enh |
| DUTERTRE_ | DUTERTRE_ | DUTERTRE_ | 14/950 | 91/20690 | 6.65E-05 | 0.002849 | 0.002195 | 84159/704 | | 14 | C42_DMR.enh |
| BOWIE_RES | BOWIE_RES | BOWIE_RES | 6/950 | 17/20690 | 7.38E-05 | 0.00313 | 0.002412 | 5699/8519, | | 6 | C42_DMR.enh |
| REACTOME | REACTOME | REACTOME | 14/950 | 92/20690 | 7.51E-05 | 0.003134 | 0.002415 | 3985/2051, | | 14 | C42_DMR.enh |
| TSAI_RESP( | TSAI_RESP( | TSAI_RESP( | 8/950 | 32/20690 | 7.54E-05 | 0.003134 | 0.002415 | 7042/8519, | | 8 | C42_DMR.enh |
| BOQUEST_ | BOQUEST_ | BOQUEST_ | 27/950 | 262/20690 | 7.85E-05 | 0.003183 | 0.002452 | 7474/1041 | | 27 | C42_DMR.enh |
| REACTOME | REACTOME | REACTOME | 27/950 | 262/20690 | 7.85E-05 | 0.003183 | 0.002452 | 7033/5590, | | 27 | C42_DMR.enh |
| REACTOME | REACTOME | REACTOME | 17/950 | 128/20690 | 7.88E-05 | 0.003183 | 0.002452 | 5320/2819, | | 17 | C42_DMR.enh |
| KIM_WT1_ | KIM_WT1_ | KIM_WT1_ | 20/950 | 167/20690 | 8.51E-05 | 0.003405 | 0.002623 | 5178/4681, | | 20 | C42_DMR.enh |
| WANG_HC | WANG_HC | WANG_HC | 15/950 | 105/20690 | 8.87E-05 | 0.003513 | 0.002706 | 7474/1396, | | 15 | C42_DMR.enh |
| REACTOME | REACTOME | REACTOME | 10/950 | 51/20690 | 9.13E-05 | 0.003533 | 0.002722 | 2051/1945, | | 10 | C42_DMR.enh |
| STAMBOLS | STAMBOLS | STAMBOLS | 10/950 | 51/20690 | 9.13E-05 | 0.003533 | 0.002722 | 6876/3006 | | 10 | C42_DMR.enh |
| JECHLINGE | JECHLINGE | JECHLINGE | 11/950 | 61/20690 | 9.17E-05 | 0.003533 | 0.002722 | 604/5590/7 | | 11 | C42_DMR.enh |
| DURAND_S | DURAND_S | DURAND_S | 29/950 | 294/20690 | 9.51E-05 | 0.003631 | 0.002798 | 2583/1041 | | 29 | C42_DMR.enh |
| PEREZ_TP6 | PEREZ_TP6 | PEREZ_TP6 | 33/950 | 354/20690 | 9.75E-05 | 0.003689 | 0.002842 | 283078/10 | | 33 | C42_DMR.enh |
| KEGG_CHR | KEGG_CHR | KEGG_CHR | 12/950 | 73/20690 | 0.000113 | 0.004252 | 0.003275 | 7043/6464, | | 12 | C42_DMR.enh |
| LINDGREN_ | LINDGREN_ | LINDGREN_ | 35/950 | 388/20690 | 0.000117 | 0.004342 | 0.003345 | 51299/139 | | 35 | C42_DMR.enh |

| | | | | | | | | | | |
|---|---|---|---|---|---|---|---|---|---|---|
| QI_HYPOXI | QI_HYPOXI | QI_HYPOXI | 8/950 | 34/20690 | 0.00012 | 0.004418 | 0.003404 | 9123/4355, | 8 | C42_DMR.enh |
| PID_ENDOT | PID_ENDOT | PID_ENDOT | 11/950 | 63/20690 | 0.000124 | 0.004533 | 0.003493 | 9564/2774, | 11 | C42_DMR.enh |
| RAY_TUMC | RAY_TUMC | RAY_TUMC | 26/950 | 255/20690 | 0.000126 | 0.004533 | 0.003493 | 1397/1158 | 26 | C42_DMR.enh |
| REACTOME | REACTOME | REACTOME | 40/950 | 468/20690 | 0.000127 | 0.004533 | 0.003493 | 5590/1298, | 40 | C42_DMR.enh |
| PID_INSULI | PID_INSULI | PID_INSULI | 7/950 | 26/20690 | 0.000128 | 0.004533 | 0.003493 | 5590/9322, | 7 | C42_DMR.enh |
| REACTOME | REACTOME | REACTOME | 7/950 | 26/20690 | 0.000128 | 0.004533 | 0.003493 | 7249/3164, | 7 | C42_DMR.enh |
| KEGG_REG | KEGG_REG | KEGG_REG | 23/950 | 213/20690 | 0.00013 | 0.004534 | 0.003493 | 3985/5496 | 23 | C42_DMR.enh |
| MARTINEZ_ | MARTINEZ_ | MARTINEZ_ | 52/950 | 665/20690 | 0.000133 | 0.004632 | 0.003569 | 5178/7474, | 52 | C42_DMR.enh |
| BILD_E2F3_ | BILD_E2F3_ | BILD_E2F3_ | 25/950 | 242/20690 | 0.000137 | 0.004717 | 0.003634 | 7474/6450, | 25 | C42_DMR.enh |
| PARENT_M | PARENT_M | PARENT_M | 46/950 | 567/20690 | 0.00014 | 0.004772 | 0.003676 | 7033/5320, | 46 | C42_DMR.enh |
| KAAB_FAIL | KAAB_FAIL | KAAB_FAIL | 9/950 | 44/20690 | 0.000145 | 0.004836 | 0.003726 | 10581/851 | 9 | C42_DMR.enh |
| PID_CERAN | PID_CERAN | PID_CERAN | 9/950 | 44/20690 | 0.000145 | 0.004836 | 0.003726 | 5590/8717, | 9 | C42_DMR.enh |
| PID_INSULI | PID_INSULI | PID_INSULI | 9/950 | 44/20690 | 0.000145 | 0.004836 | 0.003726 | 5590/9322, | 9 | C42_DMR.enh |
| UDAYAKUN | UDAYAKUN | UDAYAKUN | 25/950 | 243/20690 | 0.000146 | 0.004837 | 0.003726 | 6484/8415 | 25 | C42_DMR.enh |
| BANDRES_I | BANDRES_I | BANDRES_I | 19/950 | 161/20690 | 0.000154 | 0.005057 | 0.003896 | 4016/3201, | 19 | C42_DMR.enh |
| HAMAI_AP | HAMAI_AP | HAMAI_AP | 22/950 | 203/20690 | 0.000171 | 0.005557 | 0.004282 | 10417/514 | 22 | C42_DMR.enh |
| TARTE_PLA | TARTE_PLA | TARTE_PLA | 35/950 | 396/20690 | 0.000174 | 0.005612 | 0.004324 | 6222/4681, | 35 | C42_DMR.enh |
| AFFAR_YY1 | AFFAR_YY1 | AFFAR_YY1 | 21/950 | 190/20690 | 0.000181 | 0.005774 | 0.004449 | 64641/18/2 | 21 | C42_DMR.enh |
| CHIARADO | CHIARADO | CHIARADO | 17/950 | 137/20690 | 0.000184 | 0.005774 | 0.004449 | 7106/5507 | 17 | C42_DMR.enh |
| KEGG_INSL | KEGG_INSL | KEGG_INSL | 17/950 | 137/20690 | 0.000184 | 0.005774 | 0.004449 | 10891/559 | 17 | C42_DMR.enh |
| PID_PI3K_F | PID_PI3K_F | PID_PI3K_F | 8/950 | 36/20690 | 0.000184 | 0.005774 | 0.004449 | 6464/3265, | 8 | C42_DMR.enh |
| GAJATE_RE | GAJATE_RE | GAJATE_RE | 11/950 | 66/20690 | 0.000191 | 0.005872 | 0.004524 | 1396/1847, | 11 | C42_DMR.enh |
| FRIDMAN_ | FRIDMAN_ | FRIDMAN_ | 12/950 | 77/20690 | 0.000192 | 0.005872 | 0.004524 | 4692/1410, | 12 | C42_DMR.enh |
| NAKAYAMA | NAKAYAMA | NAKAYAMA | 12/950 | 77/20690 | 0.000192 | 0.005872 | 0.004524 | 5178/4692, | 12 | C42_DMR.enh |
| REACTOME | REACTOME | REACTOME | 23/950 | 219/20690 | 0.000196 | 0.005964 | 0.004595 | 7033/5590, | 23 | C42_DMR.enh |
| MODY_HIP | MODY_HIP | MODY_HIP | 10/950 | 56/20690 | 0.000207 | 0.006199 | 0.004776 | 2026/4720, | 10 | C42_DMR.enh |
| PID_TXA2P | PID_TXA2P | PID_TXA2P | 10/950 | 56/20690 | 0.000207 | 0.006199 | 0.004776 | 5590/7052, | 10 | C42_DMR.enh |
| BIOCARTA_ | BIOCARTA_ | BIOCARTA_ | 7/950 | 28/20690 | 0.000213 | 0.006355 | 0.004896 | 7043/7042, | 7 | C42_DMR.enh |
| KEGG_ADIF | KEGG_ADIF | KEGG_ADIF | 11/950 | 67/20690 | 0.000219 | 0.006486 | 0.004997 | 10891/871 | 11 | C42_DMR.enh |
| PASINI_SUZ | PASINI_SUZ | PASINI_SUZ | 29/950 | 309/20690 | 0.000225 | 0.006496 | 0.005005 | 25937/515 | 29 | C42_DMR.enh |
| LEE_AGING | LEE_AGING | LEE_AGING | 8/950 | 37/20690 | 0.000226 | 0.006496 | 0.005005 | 4692/2671, | 8 | C42_DMR.enh |
| WANG_BAI | WANG_BAI | WANG_BAI | 8/950 | 37/20690 | 0.000226 | 0.006496 | 0.005005 | 6484/5083, | 8 | C42_DMR.enh |
| KYNG_DNA | KYNG_DNA | KYNG_DNA | 22/950 | 207/20690 | 0.000226 | 0.006496 | 0.005005 | 7474/1847, | 22 | C42_DMR.enh |
| CHICAS_RB | CHICAS_RB | CHICAS_RB | 45/950 | 563/20690 | 0.000229 | 0.006553 | 0.005049 | 975/4016/9 | 45 | C42_DMR.enh |
| JACKSON_I | JACKSON_I | JACKSON_I | 12/950 | 79/20690 | 0.000245 | 0.006857 | 0.005283 | 1571/1410, | 12 | C42_DMR.enh |
| PID_MET_F | PID_MET_F | PID_MET_F | 12/950 | 79/20690 | 0.000245 | 0.006857 | 0.005283 | 5590/9564, | 12 | C42_DMR.enh |
| DELACROIX | DELACROIX | DELACROIX | 9/950 | 47/20690 | 0.000246 | 0.006857 | 0.005283 | 7043/6406 | 9 | C42_DMR.enh |
| ZWANG_EC | ZWANG_EC | ZWANG_EC | 24/950 | 237/20690 | 0.00025 | 0.006857 | 0.005283 | 2550/2886, | 24 | C42_DMR.enh |
| BLALOCK_A | BLALOCK_A | BLALOCK_A | 34/950 | 388/20690 | 0.000251 | 0.006857 | 0.005283 | 3985/2593 | 34 | C42_DMR.enh |
| YAO_TEMP | YAO_TEMP | YAO_TEMP | 11/950 | 68/20690 | 0.000251 | 0.006857 | 0.005283 | 2052/5710 | 11 | C42_DMR.enh |
| ENK_UV_R | ENK_UV_R | ENK_UV_R | 28/950 | 296/20690 | 0.000251 | 0.006857 | 0.005283 | 6472/6406 | 28 | C42_DMR.enh |
| CHANDRAN | CHANDRAN | CHANDRAN | 29/950 | 312/20690 | 0.000265 | 0.007168 | 0.005523 | 2550/4681, | 29 | C42_DMR.enh |
| PID_S1P_S: | PID_S1P_S: | PID_S1P_S: | 7/950 | 29/20690 | 0.00027 | 0.007227 | 0.005568 | 5155/2768, | 7 | C42_DMR.enh |
| REACTOME | REACTOME | REACTOME | 7/950 | 29/20690 | 0.00027 | 0.007227 | 0.005568 | 1945/1944, | 7 | C42_DMR.enh |
| KANNAN_T | KANNAN_T | KANNAN_T | 10/950 | 58/20690 | 0.000279 | 0.007362 | 0.005672 | 1263/604/4 | 10 | C42_DMR.enh |
| REACTOME | REACTOME | REACTOME | 10/950 | 58/20690 | 0.000279 | 0.007362 | 0.005672 | 1298/9564, | 10 | C42_DMR.enh |
| FULCHER_I | FULCHER_I | FULCHER_I | 46/950 | 585/20690 | 0.000283 | 0.007422 | 0.005718 | 7474/1263, | 46 | C42_DMR.enh |
| BIOCARTA_ | BIOCARTA_ | BIOCARTA_ | 5/950 | 14/20690 | 0.000286 | 0.007422 | 0.005718 | 5598/6464, | 5 | C42_DMR.enh |
| PID_MTOR | PID_MTOR | PID_MTOR | 11/950 | 69/20690 | 0.000286 | 0.007422 | 0.005718 | 10891/724 | 11 | C42_DMR.enh |
| MILI_PSEU | MILI_PSEU | MILI_PSEU | 36/950 | 423/20690 | 0.000294 | 0.007497 | 0.005776 | 25855/689 | 36 | C42_DMR.enh |
| REACTOME | REACTOME | REACTOME | 22/950 | 211/20690 | 0.000296 | 0.007497 | 0.005776 | 5320/8184 | 22 | C42_DMR.enh |
| IVANOVA_I | IVANOVA_I | IVANOVA_I | 20/950 | 183/20690 | 0.000296 | 0.007497 | 0.005776 | 1571/1847, | 20 | C42_DMR.enh |
| WU_CELL_ | WU_CELL_ | WU_CELL_ | 20/950 | 183/20690 | 0.000296 | 0.007497 | 0.005776 | 7474/8415 | 20 | C42_DMR.enh |
| WANG_ML | WANG_ML | WANG_ML | 27/950 | 285/20690 | 0.000314 | 0.007792 | 0.006003 | 8705/1041 | 27 | C42_DMR.enh |
| ELVIDGE_H | ELVIDGE_H | ELVIDGE_H | 14/950 | 105/20690 | 0.000315 | 0.007792 | 0.006003 | 4016/1057 | 14 | C42_DMR.enh |
| REACTOME | REACTOME | REACTOME | 14/950 | 105/20690 | 0.000315 | 0.007792 | 0.006003 | 5590/9564, | 14 | C42_DMR.enh |
| GRUETZMA | GRUETZMA | GRUETZMA | 31/950 | 346/20690 | 0.000315 | 0.007792 | 0.006003 | 7474/7033, | 31 | C42_DMR.enh |

| | | | | | | | | | | |
|---|---|---|---|---|---|---|---|---|---|---|
| ONDER_CD | ONDER_CD | ONDER_CD | 25/950 | 256/20690 | 0.000327 | 0.008038 | 0.006193 | 7474/1397 | 25 | C42_DMR.enh |
| PID_ECADH | PID_ECADH | PID_ECADH | 8/950 | 39/20690 | 0.000332 | 0.008096 | 0.006237 | 26999/369 | 8 | C42_DMR.enh |
| REACTOME | REACTOME | REACTOME | 28/950 | 302/20690 | 0.000349 | 0.008471 | 0.006526 | 5699/5733 | 28 | C42_DMR.enh |
| ELVIDGE_H | ELVIDGE_H | ELVIDGE_H | 19/950 | 172/20690 | 0.000363 | 0.008758 | 0.006747 | 4016/1057 | 19 | C42_DMR.enh |
| HALLMARK | HALLMARK | HALLMARK | 21/950 | 200/20690 | 0.000366 | 0.008785 | 0.006768 | 7033/4681 | 21 | C42_DMR.enh |
| REACTOME | REACTOME | REACTOME | 15/950 | 120/20690 | 0.000398 | 0.009434 | 0.007268 | 5699/1089 | 15 | C42_DMR.enh |
| PID_EPHB_ | PID_EPHB_ | PID_EPHB_ | 8/950 | 40/20690 | 0.000398 | 0.009434 | 0.007268 | 2886/6464 | 8 | C42_DMR.enh |
| HSIAO_HOI | HSIAO_HOI | HSIAO_HOI | 34/950 | 398/20690 | 0.0004 | 0.009434 | 0.007268 | 6222/975/ | 34 | C42_DMR.enh |
| MASSARWI | MASSARWI | MASSARWI | 45/950 | 578/20690 | 0.000406 | 0.009535 | 0.007346 | 8705/288/ | 45 | C42_DMR.enh |
| REACTOME | REACTOME | REACTOME | 25/950 | 260/20690 | 0.000413 | 0.009627 | 0.007417 | 5590/7043 | 25 | C42_DMR.enh |
| OISHI_CHO | OISHI_CHO | OISHI_CHO | 26/950 | 275/20690 | 0.000415 | 0.009627 | 0.007417 | 80307/667 | 26 | C42_DMR.enh |
| PID_TRKR_ | PID_TRKR_ | PID_TRKR_ | 10/950 | 61/20690 | 0.000425 | 0.009806 | 0.007555 | 5590/6464 | 10 | C42_DMR.enh |
| SENESE_HD | SENESE_HD | SENESE_HD | 43/950 | 546/20690 | 0.000427 | 0.009806 | 0.007555 | 1727/1063 | 43 | C42_DMR.enh |
| BUYTAERT_ | BUYTAERT_ | BUYTAERT_ | 49/950 | 648/20690 | 0.000445 | 0.010166 | 0.007832 | 7474/5699 | 49 | C42_DMR.enh |
| ANDERSEN | ANDERSEN | ANDERSEN | 19/950 | 175/20690 | 0.000451 | 0.010192 | 0.007852 | 9123/1090 | 19 | C42_DMR.enh |
| MOREAUX_ | MOREAUX_ | MOREAUX_ | 32/950 | 369/20690 | 0.000451 | 0.010192 | 0.007852 | 2550/6002 | 32 | C42_DMR.enh |
| DOUGLAS_ | DOUGLAS_ | DOUGLAS_ | 44/950 | 565/20690 | 0.000466 | 0.010395 | 0.008009 | 7474/6910 | 44 | C42_DMR.enh |
| ACEVEDO_ | ACEVEDO_ | ACEVEDO_ | 31/950 | 354/20690 | 0.000467 | 0.010395 | 0.008009 | 7033/1158 | 31 | C42_DMR.enh |
| REACTOME | REACTOME | REACTOME | 8/950 | 41/20690 | 0.000475 | 0.010395 | 0.008009 | 2668/5598 | 8 | C42_DMR.enh |
| PID_PRL_SI | PID_PRL_SI | PID_PRL_SI | 6/950 | 23/20690 | 0.000475 | 0.010395 | 0.008009 | 9564/5595 | 6 | C42_DMR.enh |
| WANG_IMI | WANG_IMI | WANG_IMI | 6/950 | 23/20690 | 0.000475 | 0.010395 | 0.008009 | 51523/369 | 6 | C42_DMR.enh |
| MANALO_H | MANALO_H | MANALO_H | 21/950 | 204/20690 | 0.000477 | 0.010395 | 0.008009 | 8705/604/ | 21 | C42_DMR.enh |
| VART_KSHV | VART_KSHV | VART_KSHV | 18/950 | 162/20690 | 0.000478 | 0.010395 | 0.008009 | 7474/7043 | 18 | C42_DMR.enh |
| MARSON_F | MARSON_F | MARSON_F | 10/950 | 62/20690 | 0.000487 | 0.010536 | 0.008117 | 975/1396/ | 10 | C42_DMR.enh |
| IVANOVA_I | IVANOVA_I | IVANOVA_I | 43/950 | 551/20690 | 0.000516 | 0.011109 | 0.008559 | 89796/320 | 43 | C42_DMR.enh |
| PID_PTP1B | PID_PTP1B | PID_PTP1B | 9/950 | 52/20690 | 0.000542 | 0.011614 | 0.008948 | 9564/5155 | 9 | C42_DMR.enh |
| DANG_REG | DANG_REG | DANG_REG | 24/950 | 250/20690 | 0.000547 | 0.011651 | 0.008976 | 5155/4641 | 24 | C42_DMR.enh |
| AMIT_EGF_ | AMIT_EGF_ | AMIT_EGF_ | 18/950 | 164/20690 | 0.000554 | 0.011651 | 0.008976 | 7052/1534 | 18 | C42_DMR.enh |
| DURAND_S | DURAND_S | DURAND_S | 18/950 | 164/20690 | 0.000554 | 0.011651 | 0.008976 | 9123/9143 | 18 | C42_DMR.enh |
| REACTOME | REACTOME | REACTOME | 10/950 | 63/20690 | 0.000555 | 0.011651 | 0.008976 | 2668/1298 | 10 | C42_DMR.enh |
| CHARAFE_I | CHARAFE_I | CHARAFE_I | 37/950 | 454/20690 | 0.000558 | 0.011651 | 0.008976 | 1727/6891 | 37 | C42_DMR.enh |
| ROPERO_H | ROPERO_H | ROPERO_H | 14/950 | 111/20690 | 0.00056 | 0.011651 | 0.008976 | 3651/2813 | 14 | C42_DMR.enh |
| REACTOME | REACTOME | REACTOME | 8/950 | 42/20690 | 0.000563 | 0.011651 | 0.008976 | 3985/2051 | 8 | C42_DMR.enh |
| BILD_MYC_ | BILD_MYC_ | BILD_MYC_ | 21/950 | 207/20690 | 0.000579 | 0.011871 | 0.009146 | 115817/54 | 21 | C42_DMR.enh |
| HOLLMANN | HOLLMANN | HOLLMANN | 25/950 | 266/20690 | 0.000579 | 0.011871 | 0.009146 | 9123/6891 | 25 | C42_DMR.enh |
| CHIANG_LI | CHIANG_LI | CHIANG_LI | 19/950 | 179/20690 | 0.000598 | 0.012187 | 0.009389 | 2875/1158 | 19 | C42_DMR.enh |
| BIOCARTA_ | BIOCARTA_ | BIOCARTA_ | 6/950 | 24/20690 | 0.000609 | 0.012243 | 0.009432 | 7033/6464 | 6 | C42_DMR.enh |
| DAZARD_U | DAZARD_U | DAZARD_U | 6/950 | 24/20690 | 0.000609 | 0.012243 | 0.009432 | 604/6280/ | 6 | C42_DMR.enh |
| CAIRO_HEF | CAIRO_HEF | CAIRO_HEF | 25/950 | 267/20690 | 0.000612 | 0.012243 | 0.009432 | 10891/115 | 25 | C42_DMR.enh |
| KEGG_MAF | KEGG_MAF | KEGG_MAF | 25/950 | 267/20690 | 0.000612 | 0.012243 | 0.009432 | 5320/7043 | 25 | C42_DMR.enh |
| BORLAK_LI | BORLAK_LI | BORLAK_LI | 9/950 | 53/20690 | 0.000627 | 0.012356 | 0.00952 | 7033/5359 | 9 | C42_DMR.enh |
| REACTOME | REACTOME | REACTOME | 9/950 | 53/20690 | 0.000627 | 0.012356 | 0.00952 | 3164/5916 | 9 | C42_DMR.enh |
| WANG_BAI | WANG_BAI | WANG_BAI | 9/950 | 53/20690 | 0.000627 | 0.012356 | 0.00952 | 64063/308 | 9 | C42_DMR.enh |
| LIU_PROST | LIU_PROST | LIU_PROST | 39/950 | 491/20690 | 0.000659 | 0.012942 | 0.009971 | 4681/7043 | 39 | C42_DMR.enh |
| LEONARD_ | LEONARD_ | LEONARD_ | 8/950 | 43/20690 | 0.000664 | 0.012978 | 0.009998 | 9123/3201 | 8 | C42_DMR.enh |
| FONTAINE_ | FONTAINE_ | FONTAINE_ | 11/950 | 76/20690 | 0.000672 | 0.012998 | 0.010014 | 2051/2365 | 11 | C42_DMR.enh |
| KEGG_VEG | KEGG_VEG | KEGG_VEG | 11/950 | 76/20690 | 0.000672 | 0.012998 | 0.010014 | 5320/3265 | 11 | C42_DMR.enh |
| MARTINEZ_ | MARTINEZ_ | MARTINEZ_ | 44/950 | 576/20690 | 0.000695 | 0.013393 | 0.010318 | 5178/7474 | 44 | C42_DMR.enh |
| BECKER_TA | BECKER_TA | BECKER_TA | 9/950 | 54/20690 | 0.000722 | 0.01378 | 0.010616 | 7033/6876 | 9 | C42_DMR.enh |
| PID_TAP63 | PID_TAP63 | PID_TAP63 | 9/950 | 54/20690 | 0.000722 | 0.01378 | 0.010616 | 6604/9176 | 9 | C42_DMR.enh |
| SMID_BREA | SMID_BREA | SMID_BREA | 18/950 | 168/20690 | 0.000737 | 0.013861 | 0.010679 | 7033/1396 | 18 | C42_DMR.enh |
| RUTELLA_F | RUTELLA_F | RUTELLA_F | 34/950 | 412/20690 | 0.000739 | 0.013861 | 0.010679 | 975/4720/ | 34 | C42_DMR.enh |
| BIOCARTA_ | BIOCARTA_ | BIOCARTA_ | 4/950 | 10/20690 | 0.000743 | 0.013861 | 0.010679 | 8717/8737 | 4 | C42_DMR.enh |
| GOUYER_T | GOUYER_T | GOUYER_T | 4/950 | 10/20690 | 0.000743 | 0.013861 | 0.010679 | 7033/7704 | 4 | C42_DMR.enh |
| REACTOME | REACTOME | REACTOME | 4/950 | 10/20690 | 0.000743 | 0.013861 | 0.010679 | 3164/207/ | 4 | C42_DMR.enh |
| KEGG_GLY | KEGG_GLY | KEGG_GLY | 11/950 | 77/20690 | 0.000752 | 0.013906 | 0.010713 | 5320/2819 | 11 | C42_DMR.enh |
| MUNSHI_M | MUNSHI_M | MUNSHI_M | 11/950 | 77/20690 | 0.000752 | 0.013906 | 0.010713 | 7043/1106 | 11 | C42_DMR.enh |

| | | | | | | | | | | |
|---|---|---|---|---|---|---|---|---|---|---|
| CREIGHTOI | CREIGHTOI | CREIGHTOI | 7/950 | 34/20690 | 0.000762 | 0.014042 | 0.010818 | 25855/153 | 7 | C42_DMR.enh |
| WONG_EN | WONG_EN | WONG_EN | 6/950 | 25/20690 | 0.00077 | 0.01413 | 0.010886 | 7033/8348 | 6 | C42_DMR.enh |
| REACTOME | REACTOME | REACTOME | 8/950 | 44/20690 | 0.00078 | 0.014239 | 0.01097 | 1163/25/1( | 8 | C42_DMR.enh |
| SA_TRKA_F | SA_TRKA_F | SA_TRKA_F | 5/950 | 17/20690 | 0.000787 | 0.014284 | 0.011005 | 6464/3265 | 5 | C42_DMR.enh |
| REACTOME | REACTOME | REACTOME | 13/950 | 102/20690 | 0.000789 | 0.014284 | 0.011005 | 5155/7249 | 13 | C42_DMR.enh |
| ZWANG_CI | ZWANG_CI | ZWANG_CI | 42/950 | 546/20690 | 0.000794 | 0.014312 | 0.011027 | 1263/8415 | 42 | C42_DMR.enh |
| MARTINEZ_ | MARTINEZ_ | MARTINEZ_ | 44/950 | 580/20690 | 0.0008 | 0.014363 | 0.011066 | 7474/288/2 | 44 | C42_DMR.enh |
| GARGALOV | GARGALOV | GARGALOV | 11/950 | 78/20690 | 0.000839 | 0.014954 | 0.011521 | 8470/5155 | 11 | C42_DMR.enh |
| IVANOVA_I | IVANOVA_I | IVANOVA_I | 21/950 | 213/20690 | 0.00084 | 0.014954 | 0.011521 | 85442/320 | 21 | C42_DMR.enh |
| RUTELLA_F | RUTELLA_F | RUTELLA_F | 28/950 | 320/20690 | 0.000869 | 0.015407 | 0.01187 | 1241/4286 | 28 | C42_DMR.enh |
| BENPORAT | BENPORAT | BENPORAT | 22/950 | 229/20690 | 0.000904 | 0.015951 | 0.012289 | 7043/4913 | 22 | C42_DMR.enh |
| FIGUEROA_ | FIGUEROA_ | FIGUEROA_ | 8/950 | 45/20690 | 0.000911 | 0.016004 | 0.01233 | 6293/6222 | 8 | C42_DMR.enh |
| PID_ERBB1 | PID_ERBB1 | PID_ERBB1 | 7/950 | 35/20690 | 0.000916 | 0.016022 | 0.012343 | 6464/8440 | 7 | C42_DMR.enh |
| HALLMARK | HALLMARK | HALLMARK | 20/950 | 200/20690 | 0.000927 | 0.016065 | 0.012377 | 7106/2886 | 20 | C42_DMR.enh |
| KOYAMA_S | KOYAMA_S | KOYAMA_S | 30/950 | 353/20690 | 0.000928 | 0.016065 | 0.012377 | 5178/2668 | 30 | C42_DMR.enh |
| YAGI_AML_ | YAGI_AML_ | YAGI_AML_ | 31/950 | 369/20690 | 0.000929 | 0.016065 | 0.012377 | 4681/3985 | 31 | C42_DMR.enh |
| PID_P73PA | PID_P73PA | PID_P73PA | 11/950 | 79/20690 | 0.000935 | 0.016101 | 0.012405 | 1263/4720 | 11 | C42_DMR.enh |
| BRUINS_U\ | BRUINS_U\ | BRUINS_U\ | 41/950 | 534/20690 | 0.000945 | 0.0162 | 0.012481 | 1263/6483 | 41 | C42_DMR.enh |
| VERHAAK_ | VERHAAK_ | VERHAAK_ | 19/950 | 186/20690 | 0.000954 | 0.016244 | 0.012515 | 2678/604/ | 19 | C42_DMR.enh |
| BIOCARTA_ | BIOCARTA_ | BIOCARTA_ | 6/950 | 26/20690 | 0.000963 | 0.016244 | 0.012515 | 3265/5595 | 6 | C42_DMR.enh |
| PID_RXR_V | PID_RXR_V | PID_RXR_V | 6/950 | 26/20690 | 0.000963 | 0.016244 | 0.012515 | 3164/5916 | 6 | C42_DMR.enh |
| JOHNSTON | JOHNSTON | JOHNSTON | 16/950 | 144/20690 | 0.000963 | 0.016244 | 0.012515 | 396/5045/2 | 16 | C42_DMR.enh |
| HAN_SATB | HAN_SATB | HAN_SATB | 32/950 | 386/20690 | 0.000967 | 0.01625 | 0.01252 | 375449/66 | 32 | C42_DMR.enh |
| ENGELMAN | ENGELMAN | ENGELMAN | 10/950 | 68/20690 | 0.001027 | 0.017187 | 0.013242 | 7033/4681 | 10 | C42_DMR.enh |
| HALLMARK | HALLMARK | HALLMARK | 13/950 | 105/20690 | 0.001038 | 0.017237 | 0.01328 | 7249/396/3 | 13 | C42_DMR.enh |
| WOO_LIVE | WOO_LIVE | WOO_LIVE | 13/950 | 105/20690 | 0.001038 | 0.017237 | 0.01328 | 1298/1396 | 13 | C42_DMR.enh |
| BOWIE_RE | BOWIE_RE | BOWIE_RE | 5/950 | 18/20690 | 0.001049 | 0.017342 | 0.013361 | 8519/4599 | 5 | C42_DMR.enh |
| RASHI_RES | RASHI_RES | RASHI_RES | 8/950 | 46/20690 | 0.001059 | 0.017369 | 0.013382 | 80305/901 | 8 | C42_DMR.enh |
| SIG_BCR_S | SIG_BCR_S | SIG_BCR_S | 8/950 | 46/20690 | 0.001059 | 0.017369 | 0.013382 | 975/6464/ | 8 | C42_DMR.enh |
| WIEDERSCI | WIEDERSCI | WIEDERSCI | 9/950 | 57/20690 | 0.00108 | 0.017651 | 0.013598 | 89796/228 | 9 | C42_DMR.enh |
| KRIEG_HYP | KRIEG_HYP | KRIEG_HYP | 53/950 | 745/20690 | 0.001088 | 0.017718 | 0.013651 | 84988/649 | 53 | C42_DMR.enh |
| MARTENS_ | MARTENS_ | MARTENS_ | 36/950 | 455/20690 | 0.001118 | 0.018027 | 0.013889 | 1241/1847 | 36 | C42_DMR.enh |
| MCBRYAN_ | MCBRYAN_ | MCBRYAN_ | 17/950 | 160/20690 | 0.001122 | 0.018027 | 0.013889 | 7474/7052 | 17 | C42_DMR.enh |
| NIKOLSKY_ | NIKOLSKY_ | NIKOLSKY_ | 4/950 | 11/20690 | 0.001125 | 0.018027 | 0.013889 | 3200/3201 | 4 | C42_DMR.enh |
| WHITESIDE | WHITESIDE | WHITESIDE | 4/950 | 11/20690 | 0.001125 | 0.018027 | 0.013889 | 8519/4599 | 4 | C42_DMR.enh |
| SANSOM_A | SANSOM_A | SANSOM_A | 21/950 | 218/20690 | 0.00113 | 0.018046 | 0.013903 | 6002/8426 | 21 | C42_DMR.enh |
| BIOCARTA_ | BIOCARTA_ | BIOCARTA_ | 11/950 | 81/20690 | 0.001155 | 0.018374 | 0.014155 | 7043/5598 | 11 | C42_DMR.enh |
| PID_GLYPI( | PID_GLYPI( | PID_GLYPI( | 6/950 | 27/20690 | 0.001191 | 0.018663 | 0.014378 | 5320/7043 | 6 | C42_DMR.enh |
| REACTOME | REACTOME | REACTOME | 6/950 | 27/20690 | 0.001191 | 0.018663 | 0.014378 | 9564/6464 | 6 | C42_DMR.enh |
| SHIN_B_CE | SHIN_B_CE | SHIN_B_CE | 6/950 | 27/20690 | 0.001191 | 0.018663 | 0.014378 | 7474/4772 | 6 | C42_DMR.enh |
| BILD_HRAS | BILD_HRAS | BILD_HRAS | 23/950 | 249/20690 | 0.001191 | 0.018663 | 0.014378 | 153572/60 | 23 | C42_DMR.enh |
| HECKER_IF | HECKER_IF | HECKER_IF | 12/950 | 94/20690 | 0.001221 | 0.01906 | 0.014685 | 10417/535 | 12 | C42_DMR.enh |
| BURTON_A | BURTON_A | BURTON_A | 8/950 | 47/20690 | 0.001225 | 0.01906 | 0.014685 | 55076/236 | 8 | C42_DMR.enh |
| LIN_SILENC | LIN_SILENC | LIN_SILENC | 13/950 | 107/20690 | 0.001239 | 0.019196 | 0.014789 | 23321/184 | 13 | C42_DMR.enh |
| BHAT_ESR1 | BHAT_ESR1 | BHAT_ESR1 | 25/950 | 281/20690 | 0.001271 | 0.019633 | 0.015126 | 7517/2635 | 25 | C42_DMR.enh |
| PID_AP1_P | PID_AP1_P | PID_AP1_P | 10/950 | 70/20690 | 0.001291 | 0.019855 | 0.015297 | 4097/7205 | 10 | C42_DMR.enh |
| RIGGINS_T. | RIGGINS_T. | RIGGINS_T. | 21/950 | 221/20690 | 0.001342 | 0.020573 | 0.01585 | 22903/841 | 21 | C42_DMR.enh |
| LIM_MAMI | LIM_MAMI | LIM_MAMI | 13/950 | 108/20690 | 0.00135 | 0.020622 | 0.015888 | 7474/8073 | 13 | C42_DMR.enh |
| BOQUEST_ | BOQUEST_ | BOQUEST_ | 34/950 | 427/20690 | 0.001362 | 0.020623 | 0.015889 | 5178/7474 | 34 | C42_DMR.enh |
| MILI_PSEU | MILI_PSEU | MILI_PSEU | 48/950 | 665/20690 | 0.001364 | 0.020623 | 0.015889 | 2668/1127 | 48 | C42_DMR.enh |
| PID_EPHA2 | PID_EPHA2 | PID_EPHA2 | 5/950 | 19/20690 | 0.00137 | 0.020623 | 0.015889 | 9564/6464 | 5 | C42_DMR.enh |
| REACTOME | REACTOME | REACTOME | 5/950 | 19/20690 | 0.00137 | 0.020623 | 0.015889 | 2051/8440 | 5 | C42_DMR.enh |
| PID_ATF2_ | PID_ATF2_ | PID_ATF2_ | 9/950 | 59/20690 | 0.00139 | 0.020853 | 0.016065 | 10891/184 | 9 | C42_DMR.enh |
| REACTOME | REACTOME | REACTOME | 46/950 | 631/20690 | 0.001396 | 0.020869 | 0.016078 | 2668/5699 | 46 | C42_DMR.enh |
| BROWNE_I | BROWNE_I | BROWNE_I | 22/950 | 237/20690 | 0.001413 | 0.020978 | 0.016162 | 7726/9910 | 22 | C42_DMR.enh |
| SENESE_HI | SENESE_HI | SENESE_HI | 22/950 | 237/20690 | 0.001413 | 0.020978 | 0.016162 | 7474/8415 | 22 | C42_DMR.enh |
| LIN_APC_T. | LIN_APC_T. | LIN_APC_T. | 10/950 | 71/20690 | 0.001442 | 0.021243 | 0.016366 | 1727/1019 | 10 | C42_DMR.enh |

| | | | | | | | | | | |
|---|---|---|---|---|---|---|---|---|---|---|
| RAMALHO_ | RAMALHO_ | RAMALHO_ | 10/950 | 71/20690 | 0.001442 | 0.021243 | 0.016366 | 2583/5119 | 10 | C42_DMR.enh |
| HELLER_SIL | HELLER_SIL | HELLER_SIL | 25/950 | 284/20690 | 0.001474 | 0.021644 | 0.016675 | 604/1298/ | 25 | C42_DMR.enh |
| RUTELLA_R | RUTELLA_R | RUTELLA_R | 22/950 | 238/20690 | 0.001492 | 0.021832 | 0.01682 | 1396/1241 | 22 | C42_DMR.enh |
| DAZARD_R | DAZARD_R | DAZARD_R | 21/950 | 223/20690 | 0.001502 | 0.0219 | 0.016872 | 9961/396/ | 21 | C42_DMR.enh |
| HUANG_FC | HUANG_FC | HUANG_FC | 7/950 | 38/20690 | 0.001524 | 0.022155 | 0.017068 | 1397/1063 | 7 | C42_DMR.enh |
| NEMETH_II | NEMETH_II | NEMETH_II | 11/950 | 84/20690 | 0.001562 | 0.022626 | 0.017431 | 5045/3557 | 11 | C42_DMR.enh |
| LEE_LIVER_ | LEE_LIVER_ | LEE_LIVER_ | 9/950 | 60/20690 | 0.00157 | 0.022661 | 0.017459 | 10417/703 | 9 | C42_DMR.enh |
| HALLMARK | HALLMARK | HALLMARK | 12/950 | 97/20690 | 0.001608 | 0.023132 | 0.017821 | 7726/5359 | 12 | C42_DMR.enh |
| REACTOME | REACTOME | REACTOME | 8/950 | 49/20690 | 0.001621 | 0.023226 | 0.017894 | 10273/288 | 8 | C42_DMR.enh |
| CAIRO_PM | CAIRO_PM | CAIRO_PM | 4/950 | 12/20690 | 0.001626 | 0.023226 | 0.017894 | 7015/595/ | 4 | C42_DMR.enh |
| MIKKELSEN | MIKKELSEN | MIKKELSEN | 17/950 | 166/20690 | 0.001679 | 0.02391 | 0.018421 | 10417/335 | 17 | C42_DMR.enh |
| KEGG_TGF_ | KEGG_TGF_ | KEGG_TGF_ | 11/950 | 85/20690 | 0.001722 | 0.024428 | 0.01882 | 7043/4052 | 11 | C42_DMR.enh |
| GAZDA_DI/ | GAZDA_DI/ | GAZDA_DI/ | 36/950 | 467/20690 | 0.001755 | 0.024488 | 0.018866 | 79862/271 | 36 | C42_DMR.enh |
| QI_PLASM/ | QI_PLASM/ | QI_PLASM/ | 12/950 | 98/20690 | 0.001758 | 0.024488 | 0.018866 | 8996/5034 | 12 | C42_DMR.enh |
| LAMB_CCN | LAMB_CCN | LAMB_CCN | 5/950 | 20/20690 | 0.001758 | 0.024488 | 0.018866 | 2771/4088 | 5 | C42_DMR.enh |
| REACTOME | REACTOME | REACTOME | 5/950 | 20/20690 | 0.001758 | 0.024488 | 0.018866 | 6464/3265 | 5 | C42_DMR.enh |
| KEGG_LINC | KEGG_LINC | KEGG_LINC | 6/950 | 29/20690 | 0.001767 | 0.024488 | 0.018866 | 5320/1571 | 6 | C42_DMR.enh |
| KEGG_THYI | KEGG_THYI | KEGG_THYI | 6/950 | 29/20690 | 0.001767 | 0.024488 | 0.018866 | 3265/5595 | 6 | C42_DMR.enh |
| REACTOME | REACTOME | REACTOME | 6/950 | 29/20690 | 0.001767 | 0.024488 | 0.018866 | 9564/2886 | 6 | C42_DMR.enh |
| BHAT_ESR1 | BHAT_ESR1 | BHAT_ESR1 | 20/950 | 211/20690 | 0.001778 | 0.02452 | 0.018891 | 7517/7052 | 20 | C42_DMR.enh |
| REACTOME | REACTOME | REACTOME | 7/950 | 39/20690 | 0.001785 | 0.02452 | 0.018891 | 5320/2819 | 7 | C42_DMR.enh |
| REACTOME | REACTOME | REACTOME | 10/950 | 73/20690 | 0.001786 | 0.02452 | 0.018891 | 5590/1027 | 10 | C42_DMR.enh |
| DOANE_RE | DOANE_RE | DOANE_RE | 22/950 | 242/20690 | 0.001843 | 0.02522 | 0.01943 | 22885/375 | 22 | C42_DMR.enh |
| LIU_SMAR( | LIU_SMAR( | LIU_SMAR( | 8/950 | 50/20690 | 0.001853 | 0.025272 | 0.019471 | 283078/57 | 8 | C42_DMR.enh |
| SHEDDEN_ | SHEDDEN_ | SHEDDEN_ | 19/950 | 197/20690 | 0.001875 | 0.025441 | 0.0196 | 23321/263 | 19 | C42_DMR.enh |
| ICHIBA_GR | ICHIBA_GR | ICHIBA_GR | 13/950 | 112/20690 | 0.001884 | 0.025441 | 0.0196 | 7033/5699 | 13 | C42_DMR.enh |
| LIEN_BREA | LIEN_BREA | LIEN_BREA | 13/950 | 112/20690 | 0.001884 | 0.025441 | 0.0196 | 7033/18/8 | 13 | C42_DMR.enh |
| SMID_BRE/ | SMID_BRE/ | SMID_BRE/ | 12/950 | 99/20690 | 0.001919 | 0.025827 | 0.019898 | 7033/1396 | 12 | C42_DMR.enh |
| DELACROIX | DELACROIX | DELACROIX | 35/950 | 453/20690 | 0.001943 | 0.026068 | 0.020083 | 2550/1263 | 35 | C42_DMR.enh |
| KEGG_NEU | KEGG_NEU | KEGG_NEU | 14/950 | 126/20690 | 0.001951 | 0.026099 | 0.020107 | 5598/6464 | 14 | C42_DMR.enh |
| LI_WILMS_ | LI_WILMS_ | LI_WILMS_ | 18/950 | 183/20690 | 0.001962 | 0.026163 | 0.020156 | 288/23321 | 18 | C42_DMR.enh |
| KEGG_COL | KEGG_COL | KEGG_COL | 9/950 | 62/20690 | 0.001986 | 0.026394 | 0.020335 | 7043/7042 | 9 | C42_DMR.enh |
| KEGG_ERBI | KEGG_ERBI | KEGG_ERBI | 11/950 | 87/20690 | 0.00208 | 0.02736 | 0.021079 | 6464/8440 | 11 | C42_DMR.enh |
| KIM_BIPOL | KIM_BIPOL | KIM_BIPOL | 11/950 | 87/20690 | 0.00208 | 0.02736 | 0.021079 | 7517/8814 | 11 | C42_DMR.enh |
| HALLMARK | HALLMARK | HALLMARK | 12/950 | 100/20690 | 0.002091 | 0.02736 | 0.021079 | 84159/855 | 12 | C42_DMR.enh |
| KEGG_FOC | KEGG_FOC | KEGG_FOC | 19/950 | 199/20690 | 0.002105 | 0.02736 | 0.021079 | 9564/5155 | 19 | C42_DMR.enh |
| SIG_INSULI | SIG_INSULI | SIG_INSULI | 8/950 | 51/20690 | 0.002111 | 0.02736 | 0.021079 | 6464/8660 | 8 | C42_DMR.enh |
| SENGUPTA | SENGUPTA | SENGUPTA | 29/950 | 356/20690 | 0.002122 | 0.02736 | 0.021079 | 7033/2616 | 29 | C42_DMR.enh |
| KASLER_HD | KASLER_HD | KASLER_HD | 6/950 | 30/20690 | 0.002124 | 0.02736 | 0.021079 | 4355/8309 | 6 | C42_DMR.enh |
| PID_RETIN( | PID_RETIN( | PID_RETIN( | 6/950 | 30/20690 | 0.002124 | 0.02736 | 0.021079 | 5916/5595 | 6 | C42_DMR.enh |
| REACTOME | REACTOME | REACTOME | 6/950 | 30/20690 | 0.002124 | 0.02736 | 0.021079 | 8717/9099 | 6 | C42_DMR.enh |
| YAO_TEMP | YAO_TEMP | YAO_TEMP | 6/950 | 30/20690 | 0.002124 | 0.02736 | 0.021079 | 1571/9099 | 6 | C42_DMR.enh |
| GARY_CD5_ | GARY_CD5_ | GARY_CD5_ | 36/950 | 473/20690 | 0.002177 | 0.027939 | 0.021525 | 9143/2290 | 36 | C42_DMR.enh |
| CHEBOTAE | CHEBOTAE | CHEBOTAE | 10/950 | 75/20690 | 0.002195 | 0.027939 | 0.021525 | 7043/2290 | 10 | C42_DMR.enh |
| REACTOME | REACTOME | REACTOME | 10/950 | 75/20690 | 0.002195 | 0.027939 | 0.021525 | 5590/6464 | 10 | C42_DMR.enh |
| MURATA_\ | MURATA_\ | MURATA_\ | 5/950 | 21/20690 | 0.002221 | 0.027939 | 0.021525 | 604/9961/ | 5 | C42_DMR.enh |
| PID_ECADH | PID_ECADH | PID_ECADH | 5/950 | 21/20690 | 0.002221 | 0.027939 | 0.021525 | 342184/20 | 5 | C42_DMR.enh |
| REACTOME | REACTOME | REACTOME | 5/950 | 21/20690 | 0.002221 | 0.027939 | 0.021525 | 8705/6484 | 5 | C42_DMR.enh |
| PID_MYC_I | PID_MYC_I | PID_MYC_I | 9/950 | 63/20690 | 0.002224 | 0.027939 | 0.021525 | 7474/7249 | 9 | C42_DMR.enh |
| HALLMARK | HALLMARK | HALLMARK | 19/950 | 200/20690 | 0.002228 | 0.027939 | 0.021525 | 2668/5158 | 19 | C42_DMR.enh |
| HALLMARK | HALLMARK | HALLMARK | 19/950 | 200/20690 | 0.002228 | 0.027939 | 0.021525 | 604/9322/ | 19 | C42_DMR.enh |
| MULLIGHA | MULLIGHA | MULLIGHA | 24/950 | 277/20690 | 0.002237 | 0.027958 | 0.02154 | 4681/4016 | 24 | C42_DMR.enh |
| BIOCARTA_ | BIOCARTA_ | BIOCARTA_ | 4/950 | 13/20690 | 0.002264 | 0.028203 | 0.021728 | 8818/3265 | 4 | C42_DMR.enh |
| KANG_IMN | KANG_IMN | KANG_IMN | 12/950 | 101/20690 | 0.002276 | 0.028203 | 0.021728 | 796/6002/ | 12 | C42_DMR.enh |
| KEGG_GNR | KEGG_GNR | KEGG_GNR | 12/950 | 101/20690 | 0.002276 | 0.028203 | 0.021728 | 5320/5598 | 12 | C42_DMR.enh |
| JIANG_HYP | JIANG_HYP | JIANG_HYP | 19/950 | 201/20690 | 0.002358 | 0.02913 | 0.022443 | 5699/2714 | 19 | C42_DMR.enh |
| GAVIN_FO) | GAVIN_FO) | GAVIN_FO) | 16/950 | 157/20690 | 0.002382 | 0.029335 | 0.0226 | 2678/9910 | 16 | C42_DMR.enh |

| | | | | | | | | | | |
|---|---|---|---|---|---|---|---|---|---|---|
| PID_ECADH | PID_ECADH | PID_ECADH | 7/950 | 41/20690 | 0.00241 | 0.029598 | 0.022803 | 3801/2699 | 7 | C42_DMR.enh |
| ZWANG_CL | ZWANG_CL | ZWANG_CL | 21/950 | 232/20690 | 0.002435 | 0.029817 | 0.022972 | 22903/139 | 21 | C42_DMR.enh |
| REACTOME | REACTOME | REACTOME | 51/950 | 738/20690 | 0.002457 | 0.029996 | 0.02311 | 2583/5320 | 51 | C42_DMR.enh |
| BROWNE_I | BROWNE_I | BROWNE_I | 12/950 | 102/20690 | 0.002475 | 0.030126 | 0.02321 | 23321/991 | 12 | C42_DMR.enh |
| BURTON_A | BURTON_A | BURTON_A | 6/950 | 31/20690 | 0.002533 | 0.030742 | 0.023684 | 8660/8848 | 6 | C42_DMR.enh |
| FARMER_B | FARMER_B | FARMER_B | 27/950 | 328/20690 | 0.002563 | 0.031028 | 0.023905 | 22903/267 | 27 | C42_DMR.enh |
| GROSS_HYI | GROSS_HYI | GROSS_HYI | 19/950 | 203/20690 | 0.002637 | 0.031822 | 0.024517 | 975/6484/( | 19 | C42_DMR.enh |
| PID_NFAT_ | PID_NFAT_ | PID_NFAT_ | 8/950 | 53/20690 | 0.002711 | 0.032628 | 0.025137 | 5590/3164 | 8 | C42_DMR.enh |
| BIOCARTA_ | BIOCARTA_ | BIOCARTA_ | 5/950 | 22/20690 | 0.002767 | 0.033043 | 0.025457 | 3265/5595 | 5 | C42_DMR.enh |
| MULLIGAN | MULLIGAN | MULLIGAN | 5/950 | 22/20690 | 0.002767 | 0.033043 | 0.025457 | 6484/2992 | 5 | C42_DMR.enh |
| KEGG_GLIC | KEGG_GLIC | KEGG_GLIC | 9/950 | 65/20690 | 0.002769 | 0.033043 | 0.025457 | 5155/6464 | 9 | C42_DMR.enh |
| MARTIN_IN | MARTIN_IN | MARTIN_IN | 7/950 | 42/20690 | 0.00278 | 0.033071 | 0.025479 | 604/5595/ | 7 | C42_DMR.enh |
| HOSHIDA_I | HOSHIDA_I | HOSHIDA_I | 23/950 | 266/20690 | 0.002802 | 0.033239 | 0.025608 | 975/5320/ | 23 | C42_DMR.enh |
| ALFANO_M | ALFANO_M | ALFANO_M | 21/950 | 235/20690 | 0.002839 | 0.033587 | 0.025877 | 4681/1397 | 21 | C42_DMR.enh |
| REACTOME | REACTOME | REACTOME | 16/950 | 160/20690 | 0.002883 | 0.034018 | 0.026208 | 1263/5497 | 16 | C42_DMR.enh |
| LE_EGR2_T | LE_EGR2_T | LE_EGR2_T | 12/950 | 104/20690 | 0.002914 | 0.034181 | 0.026334 | 2051/5833 | 12 | C42_DMR.enh |
| PLASARI_T | PLASARI_T | PLASARI_T | 12/950 | 104/20690 | 0.002914 | 0.034181 | 0.026334 | 5178/7474 | 12 | C42_DMR.enh |
| REACTOME | REACTOME | REACTOME | 10/950 | 78/20690 | 0.002946 | 0.034466 | 0.026554 | 6464/8660 | 10 | C42_DMR.enh |
| SANSOM_A | SANSOM_A | SANSOM_A | 21/950 | 236/20690 | 0.002985 | 0.03461 | 0.026664 | 1263/6448 | 21 | C42_DMR.enh |
| PID_ALPHA | PID_ALPHA | PID_ALPHA | 6/950 | 32/20690 | 0.002998 | 0.03461 | 0.026664 | 10273/559 | 6 | C42_DMR.enh |
| REACTOME | REACTOME | REACTOME | 6/950 | 32/20690 | 0.002998 | 0.03461 | 0.026664 | 6464/3265 | 6 | C42_DMR.enh |
| PEDERSEN_ | PEDERSEN_ | PEDERSEN_ | 31/950 | 398/20690 | 0.003026 | 0.03461 | 0.026664 | 7033/2593 | 31 | C42_DMR.enh |
| BIOCARTA_ | BIOCARTA_ | BIOCARTA_ | 4/950 | 14/20690 | 0.003055 | 0.03461 | 0.026664 | 6464/8818 | 4 | C42_DMR.enh |
| REACTOME | REACTOME | REACTOME | 4/950 | 14/20690 | 0.003055 | 0.03461 | 0.026664 | 7249/207/ | 4 | C42_DMR.enh |
| REACTOME | REACTOME | REACTOME | 4/950 | 14/20690 | 0.003055 | 0.03461 | 0.026664 | 6464/8660 | 4 | C42_DMR.enh |
| REACTOME | REACTOME | REACTOME | 4/950 | 14/20690 | 0.003055 | 0.03461 | 0.026664 | 54971/207 | 4 | C42_DMR.enh |
| REACTOME | REACTOME | REACTOME | 4/950 | 14/20690 | 0.003055 | 0.03461 | 0.026664 | 604/9540/ | 4 | C42_DMR.enh |
| STARK_BRA | STARK_BRA | STARK_BRA | 4/950 | 14/20690 | 0.003055 | 0.03461 | 0.026664 | 22906/787 | 4 | C42_DMR.enh |
| ZAIDI_OSTE | ZAIDI_OSTE | ZAIDI_OSTE | 4/950 | 14/20690 | 0.003055 | 0.03461 | 0.026664 | 4488/3727 | 4 | C42_DMR.enh |
| REACTOME | REACTOME | REACTOME | 8/950 | 54/20690 | 0.003057 | 0.03461 | 0.026664 | 9564/3265 | 8 | C42_DMR.enh |
| KARLSSON_ | KARLSSON_ | KARLSSON_ | 18/950 | 191/20690 | 0.003129 | 0.035331 | 0.02722 | 1396/2593 | 18 | C42_DMR.enh |
| CHAUHAN_ | CHAUHAN_ | CHAUHAN_ | 12/950 | 105/20690 | 0.003155 | 0.035531 | 0.027374 | 2678/2026 | 12 | C42_DMR.enh |
| DORN_ADE | DORN_ADE | DORN_ADE | 7/950 | 43/20690 | 0.003191 | 0.03576 | 0.027551 | 3164/1969 | 7 | C42_DMR.enh |
| BROWNE_I | BROWNE_I | BROWNE_I | 22/950 | 253/20690 | 0.003193 | 0.03576 | 0.027551 | 3985/9910 | 22 | C42_DMR.enh |
| ELVIDGE_H | ELVIDGE_H | ELVIDGE_H | 14/950 | 133/20690 | 0.003229 | 0.035972 | 0.027714 | 10570/109 | 14 | C42_DMR.enh |
| KEGG_FC_E | KEGG_FC_E | KEGG_FC_E | 10/950 | 79/20690 | 0.003237 | 0.035972 | 0.027714 | 5320/3265 | 10 | C42_DMR.enh |
| SCIBETTA_I | SCIBETTA_I | SCIBETTA_I | 10/950 | 79/20690 | 0.003237 | 0.035972 | 0.027714 | 8996/6662 | 10 | C42_DMR.enh |
| PID_TELON | PID_TELON | PID_TELON | 9/950 | 67/20690 | 0.003415 | 0.037843 | 0.029155 | 23381/588 | 9 | C42_DMR.enh |
| PID_FGF_P | PID_FGF_P | PID_FGF_P | 8/950 | 55/20690 | 0.003437 | 0.037983 | 0.029263 | 6464/632/ | 8 | C42_DMR.enh |
| MCBRYAN_ | MCBRYAN_ | MCBRYAN_ | 19/950 | 208/20690 | 0.003456 | 0.038094 | 0.029349 | 1727/1535 | 19 | C42_DMR.enh |
| SERVITJA_I | SERVITJA_I | SERVITJA_I | 16/950 | 163/20690 | 0.00347 | 0.038148 | 0.02939 | 7043/8996 | 16 | C42_DMR.enh |
| BONOME_( | BONOME_( | BONOME_( | 37/950 | 504/20690 | 0.003492 | 0.038292 | 0.029501 | 5178/7474 | 37 | C42_DMR.enh |
| KUMAR_TA | KUMAR_TA | KUMAR_TA | 31/950 | 402/20690 | 0.003511 | 0.038343 | 0.02954 | 2550/2583 | 31 | C42_DMR.enh |
| PID_SYNDE | PID_SYNDE | PID_SYNDE | 6/950 | 33/20690 | 0.003524 | 0.038343 | 0.02954 | 3265/5595 | 6 | C42_DMR.enh |
| REACTOME | REACTOME | REACTOME | 6/950 | 33/20690 | 0.003524 | 0.038343 | 0.02954 | 8717/9099 | 6 | C42_DMR.enh |
| HUANG_DA | HUANG_DA | HUANG_DA | 10/950 | 80/20690 | 0.003551 | 0.038442 | 0.029617 | 4837/5532 | 10 | C42_DMR.enh |
| YAO_HOXA | YAO_HOXA | YAO_HOXA | 10/950 | 80/20690 | 0.003551 | 0.038442 | 0.029617 | 5178/4681 | 10 | C42_DMR.enh |
| REACTOME | REACTOME | REACTOME | 15/950 | 149/20690 | 0.003605 | 0.038918 | 0.029983 | 5699/5426 | 15 | C42_DMR.enh |
| AMIT_EGF_ | AMIT_EGF_ | AMIT_EGF_ | 9/950 | 68/20690 | 0.003779 | 0.040593 | 0.031274 | 10607/687 | 9 | C42_DMR.enh |
| YAO_TEMP | YAO_TEMP | YAO_TEMP | 9/950 | 68/20690 | 0.003779 | 0.040593 | 0.031274 | 5178/4681 | 9 | C42_DMR.enh |
| MOOTHA_I | MOOTHA_I | MOOTHA_I | 32/950 | 421/20690 | 0.003793 | 0.040644 | 0.031313 | 5699/2635 | 32 | C42_DMR.enh |
| GRAHAM_( | GRAHAM_( | GRAHAM_( | 11/950 | 94/20690 | 0.00384 | 0.040931 | 0.031534 | 9645/3557 | 11 | C42_DMR.enh |
| LIM_MAMI | LIM_MAMI | LIM_MAMI | 11/950 | 94/20690 | 0.00384 | 0.040931 | 0.031534 | 8660/3714 | 11 | C42_DMR.enh |
| BIOCARTA_ | BIOCARTA_ | BIOCARTA_ | 8/950 | 56/20690 | 0.003851 | 0.040952 | 0.03155 | 8717/8837 | 8 | C42_DMR.enh |
| CREIGHTOI | CREIGHTOI | CREIGHTOI | 36/950 | 490/20690 | 0.003881 | 0.041142 | 0.031697 | 6484/288/ | 36 | C42_DMR.enh |
| ROSS_AML | ROSS_AML | ROSS_AML | 10/950 | 81/20690 | 0.003889 | 0.041142 | 0.031697 | 1298/9961 | 10 | C42_DMR.enh |
| AMBROSIN | AMBROSIN | AMBROSIN | 12/950 | 108/20690 | 0.00398 | 0.041438 | 0.031925 | 5441/8309 | 12 | C42_DMR.enh |

| | | | | | | | | | | |
|---|---|---|---|---|---|---|---|---|---|---|
| RICKMAN_ | RICKMAN_ | RICKMAN_ | 12/950 | 108/20690 | 0.00398 | 0.041438 | 0.031925 | 26160/233 | 12 | C42_DMR.enh |
| CHEBOTAE | CHEBOTAE | CHEBOTAE | 13/950 | 122/20690 | 0.004008 | 0.041438 | 0.031925 | 2668/4016 | 13 | C42_DMR.enh |
| REACTOME | REACTOME | REACTOME | 13/950 | 122/20690 | 0.004008 | 0.041438 | 0.031925 | 81846/724 | 13 | C42_DMR.enh |
| BIOCARTA_ | BIOCARTA_ | BIOCARTA_ | 4/950 | 15/20690 | 0.004016 | 0.041438 | 0.031925 | 6464/3265 | 4 | C42_DMR.enh |
| BIOCARTA_ | BIOCARTA_ | BIOCARTA_ | 4/950 | 15/20690 | 0.004016 | 0.041438 | 0.031925 | 8717/8737 | 4 | C42_DMR.enh |
| KYNG_NOR | KYNG_NOR | KYNG_NOR | 4/950 | 15/20690 | 0.004016 | 0.041438 | 0.031925 | 2774/2364 | 4 | C42_DMR.enh |
| MARKS_HD | MARKS_HD | MARKS_HD | 4/950 | 15/20690 | 0.004016 | 0.041438 | 0.031925 | 3091/8837 | 4 | C42_DMR.enh |
| SANCHEZ_I | SANCHEZ_I | SANCHEZ_I | 4/950 | 15/20690 | 0.004016 | 0.041438 | 0.031925 | 3557/3625 | 4 | C42_DMR.enh |
| SCHAVOLT | SCHAVOLT | SCHAVOLT | 4/950 | 15/20690 | 0.004016 | 0.041438 | 0.031925 | 1969/2810 | 4 | C42_DMR.enh |
| ACEVEDO_ | ACEVEDO_ | ACEVEDO_ | 23/950 | 274/20690 | 0.004044 | 0.04161 | 0.032057 | 10891/281 | 23 | C42_DMR.enh |
| SMID_BRE/ | SMID_BRE/ | SMID_BRE/ | 15/950 | 151/20690 | 0.004088 | 0.04161 | 0.032057 | 2678/2886 | 15 | C42_DMR.enh |
| ZHOU_INFL | ZHOU_INFL | ZHOU_INFL | 37/950 | 509/20690 | 0.004101 | 0.04161 | 0.032057 | 7474/6910 | 37 | C42_DMR.enh |
| ABE_VEGF/ | ABE_VEGF/ | ABE_VEGF/ | 6/950 | 34/20690 | 0.004116 | 0.04161 | 0.032057 | 1543/3164 | 6 | C42_DMR.enh |
| PID_MAPK | PID_MAPK | PID_MAPK | 6/950 | 34/20690 | 0.004116 | 0.04161 | 0.032057 | 5598/3265 | 6 | C42_DMR.enh |
| BERTUCCI_ | BERTUCCI_ | BERTUCCI_ | 5/950 | 24/20690 | 0.00414 | 0.04161 | 0.032057 | 10273/288 | 5 | C42_DMR.enh |
| REACTOME | REACTOME | REACTOME | 5/950 | 24/20690 | 0.00414 | 0.04161 | 0.032057 | 25937/632 | 5 | C42_DMR.enh |
| RIZ_ERYTH | RIZ_ERYTH | RIZ_ERYTH | 5/950 | 24/20690 | 0.00414 | 0.04161 | 0.032057 | 3651/3201 | 5 | C42_DMR.enh |
| SA_B_CELL | SA_B_CELL | SA_B_CELL | 5/950 | 24/20690 | 0.00414 | 0.04161 | 0.032057 | 6464/3265 | 5 | C42_DMR.enh |
| ZHAN_MUI | ZHAN_MUI | ZHAN_MUI | 5/950 | 24/20690 | 0.00414 | 0.04161 | 0.032057 | 2261/3959 | 5 | C42_DMR.enh |
| BEGUM_T/ | BEGUM_T/ | BEGUM_T/ | 7/950 | 45/20690 | 0.004151 | 0.04161 | 0.032057 | 6662/3714 | 7 | C42_DMR.enh |
| REACTOME | REACTOME | REACTOME | 7/950 | 45/20690 | 0.004151 | 0.04161 | 0.032057 | 4016/7043 | 7 | C42_DMR.enh |
| WINTER_H | WINTER_H | WINTER_H | 11/950 | 95/20690 | 0.004167 | 0.041631 | 0.032074 | 56901/956 | 11 | C42_DMR.enh |
| KYNG_RESI | KYNG_RESI | KYNG_RESI | 9/950 | 69/20690 | 0.004173 | 0.041631 | 0.032074 | 7106/1410 | 9 | C42_DMR.enh |
| COULOUAF | COULOUAF | COULOUAF | 14/950 | 137/20690 | 0.004223 | 0.042028 | 0.03238 | 2875/8415 | 14 | C42_DMR.enh |
| PID_REG_G | PID_REG_G | PID_REG_G | 10/950 | 82/20690 | 0.004251 | 0.042211 | 0.03252 | 3164/632/ | 10 | C42_DMR.enh |
| KEGG_ACU | KEGG_ACU | KEGG_ACU | 8/950 | 57/20690 | 0.004304 | 0.042556 | 0.032787 | 3265/5595 | 8 | C42_DMR.enh |
| REACTOME | REACTOME | REACTOME | 49/950 | 723/20690 | 0.004306 | 0.042556 | 0.032787 | 6293/7474 | 49 | C42_DMR.enh |
| GRUETZMA | GRUETZMA | GRUETZMA | 18/950 | 197/20690 | 0.004342 | 0.04281 | 0.032982 | 4016/2886 | 18 | C42_DMR.enh |
| BROWNE_I | BROWNE_I | BROWNE_I | 17/950 | 182/20690 | 0.004394 | 0.043123 | 0.033223 | 5699/2051 | 17 | C42_DMR.enh |
| JISON_SICK | JISON_SICK | JISON_SICK | 17/950 | 182/20690 | 0.004394 | 0.043123 | 0.033223 | 22885/106 | 17 | C42_DMR.enh |
| HELLER_HD | HELLER_HD | HELLER_HD | 34/950 | 460/20690 | 0.004487 | 0.043933 | 0.033847 | 604/1543/ | 34 | C42_DMR.enh |
| KEGG_FC_( | KEGG_FC_( | KEGG_FC_( | 11/950 | 96/20690 | 0.004516 | 0.044111 | 0.033984 | 3985/5595 | 11 | C42_DMR.enh |
| KEGG_PAN | KEGG_PAN | KEGG_PAN | 9/950 | 70/20690 | 0.004598 | 0.044602 | 0.034363 | 7043/7042 | 9 | C42_DMR.enh |
| MARTINEZ_ | MARTINEZ_ | MARTINEZ_ | 9/950 | 70/20690 | 0.004598 | 0.044602 | 0.034363 | 3625/1058 | 9 | C42_DMR.enh |
| REACTOME | REACTOME | REACTOME | 9/950 | 70/20690 | 0.004598 | 0.044602 | 0.034363 | 10581/851 | 9 | C42_DMR.enh |
| WOOD_EB' | WOOD_EB' | WOOD_EB' | 12/950 | 110/20690 | 0.004621 | 0.044716 | 0.03445 | 10417/396 | 12 | C42_DMR.enh |
| HUMMERI( | HUMMERI( | HUMMERI( | 10/950 | 83/20690 | 0.00464 | 0.044731 | 0.034462 | 3557/1041 | 10 | C42_DMR.enh |
| TAKEDA_T/ | TAKEDA_T/ | TAKEDA_T/ | 17/950 | 183/20690 | 0.004644 | 0.044731 | 0.034462 | 1543/8415 | 17 | C42_DMR.enh |
| AIGNER_ZE | AIGNER_ZE | AIGNER_ZE | 6/950 | 35/20690 | 0.004778 | 0.04571 | 0.035216 | 11187/834 | 6 | C42_DMR.enh |
| PID_ARF6_ | PID_ARF6_ | PID_ARF6_ | 6/950 | 35/20690 | 0.004778 | 0.04571 | 0.035216 | 1969/3674 | 6 | C42_DMR.enh |
| PID_PI3KCI | PID_PI3KCI | PID_PI3KCI | 6/950 | 35/20690 | 0.004778 | 0.04571 | 0.035216 | 5566/2810 | 6 | C42_DMR.enh |
| HUPER_BR | HUPER_BR | HUPER_BR | 8/950 | 58/20690 | 0.004796 | 0.045781 | 0.035271 | 4016/2026 | 8 | C42_DMR.enh |
| REACTOME | REACTOME | REACTOME | 46/950 | 674/20690 | 0.004905 | 0.046718 | 0.035993 | 5590/9123 | 46 | C42_DMR.enh |
| PID_INTEG | PID_INTEG | PID_INTEG | 5/950 | 25/20690 | 0.004982 | 0.047232 | 0.036388 | 9564/7052 | 5 | C42_DMR.enh |
| REACTOME | REACTOME | REACTOME | 5/950 | 25/20690 | 0.004982 | 0.047232 | 0.036388 | 6464/3265 | 5 | C42_DMR.enh |
| KEGG_SMA | KEGG_SMA | KEGG_SMA | 10/950 | 84/20690 | 0.005056 | 0.047416 | 0.036531 | 1163/3914 | 10 | C42_DMR.enh |
| KEGG_MEL | KEGG_MEL | KEGG_MEL | 9/950 | 71/20690 | 0.005057 | 0.047416 | 0.036531 | 5155/4286 | 9 | C42_DMR.enh |
| MORI_SMA | MORI_SMA | MORI_SMA | 9/950 | 71/20690 | 0.005057 | 0.047416 | 0.036531 | 1397/5147 | 9 | C42_DMR.enh |
| REACTOME | REACTOME | REACTOME | 9/950 | 71/20690 | 0.005057 | 0.047416 | 0.036531 | 4641/2810 | 9 | C42_DMR.enh |
| FOSTER_KD | FOSTER_KD | FOSTER_KD | 18/950 | 200/20690 | 0.00508 | 0.047416 | 0.036531 | 26160/139 | 18 | C42_DMR.enh |
| HALLMARK | HALLMARK | HALLMARK | 18/950 | 200/20690 | 0.00508 | 0.047416 | 0.036531 | 7033/4681 | 18 | C42_DMR.enh |
| HALLMARK | HALLMARK | HALLMARK | 18/950 | 200/20690 | 0.00508 | 0.047416 | 0.036531 | 5178/2290 | 18 | C42_DMR.enh |
| ONDER_CD | ONDER_CD | ONDER_CD | 14/950 | 140/20690 | 0.005121 | 0.047688 | 0.03674 | 604/1397/ | 14 | C42_DMR.enh |
| MAINA_VH | MAINA_VH | MAINA_VH | 4/950 | 16/20690 | 0.005162 | 0.047856 | 0.03687 | 9123/2026 | 4 | C42_DMR.enh |
| REACTOME | REACTOME | REACTOME | 4/950 | 16/20690 | 0.005162 | 0.047856 | 0.03687 | 57332/732 | 4 | C42_DMR.enh |
| VERHAAK_ | VERHAAK_ | VERHAAK_ | 19/950 | 216/20690 | 0.0052 | 0.048102 | 0.037059 | 1263/9123 | 19 | C42_DMR.enh |
| MIYAGAW/ | MIYAGAW/ | MIYAGAW/ | 22/950 | 264/20690 | 0.00529 | 0.048828 | 0.037618 | 6495/7052 | 22 | C42_DMR.enh |

| | | | | | | | | | | |
|---|---|---|---|---|---|---|---|---|---|---|
| KEGG_TYPI | KEGG_TYPI | KEGG_TYPI | 7/950 | 47/20690 | 0.005315 | 0.048849 | 0.037634 | 3651/5590 | 7 | C42_DMR.enh |
| MCDOWEL | MCDOWEL | MCDOWEL | 7/950 | 47/20690 | 0.005315 | 0.048849 | 0.037634 | 4052/3164 | 7 | C42_DMR.enh |
| WEIGEL_O. | WEIGEL_O. | WEIGEL_O. | 8/950 | 59/20690 | 0.00533 | 0.048868 | 0.037649 | 1945/5499 | 8 | C42_DMR.enh |
| RICKMAN_ | RICKMAN_ | RICKMAN_ | 12/950 | 112/20690 | 0.005341 | 0.048868 | 0.037649 | 5742/3091 | 12 | C42_DMR.enh |

### Table S9

| LNCaP.EtO | LNCaP.EtO | LNCaP.DHT | LNCaP.DHT | C42.EtOH_ | C42.EtOH_ | C42.DHT_S | C42.DHT_C | Factor | CellLine | Tissue | GSM | peaks | CistromeDB.ID |
|---|---|---|---|---|---|---|---|---|---|---|---|---|---|
| 86.23856 | 33 | 86.88206 | 37 | 25.31615 | 29 | 35.43384 | 36 | BTAF1 | HeLa | Cervix | GSM44802 | 3367 | 1 |
| 82.91712 | 25 | 90.58088 | 29 | 61.69315 | 31 | 276.1568 | 72 | ZNF645 | HEK293 | Embryonic | GSM28289 | 1717 | 100000 |
| 83.13103 | 25 | 58.69494 | 23 | 102.4947 | 40 | 209.6222 | 62 | ZNF645 | HEK293 | Embryonic | GSM28289 | 1736 | 100001 |
| 651.2529 | 172 | 748.3203 | 204 | 418.3513 | 197 | 697.2918 | 269 | ZNF10 | HEK293 | Embryonic | GSM28250 | 10402 | 100002 |
| 361.6421 | 108 | 382.416 | 123 | 291.2205 | 140 | 477.8731 | 190 | ZNF10 | HEK293 | Embryonic | GSM28250 | 7620 | 100003 |
| 890.4614 | 361 | 916.8486 | 466 | 683.6969 | 404 | 653.5751 | 426 | CTCF | None | gastrocnen | GSM28250 | 23729 | 100004 |
| 809.3333 | 282 | 842.0828 | 378 | 508.4503 | 326 | 534.5905 | 359 | CTCF | None | Esophagus | GSM28250 | 23968 | 100006 |
| 743.5776 | 140 | 1047.525 | 194 | 538.1637 | 161 | 518.2928 | 168 | ZNF579 | MCF-7 | Breast | GSM28250 | 5113 | 100008 |
| 560.5775 | 172 | 723.7605 | 217 | 218.8402 | 163 | 369.2382 | 220 | ZNF24 | GM12878 | Blood | GSM28250 | 12905 | 100010 |
| 329.0214 | 220 | 536.4904 | 305 | 236.6284 | 302 | 276.3605 | 347 | CTCF | A673 | Muscle | GSM34984 | 35771 | 100017 |
| 588.1282 | 348 | 720.2518 | 510 | 492.4445 | 500 | 489.2986 | 543 | CTCF | A673 | Muscle | GSM34984 | 54398 | 100018 |
| 707.1131 | 349 | 751.7823 | 475 | 474.5818 | 441 | 450.566 | 470 | CTCF | None | Muscle | GSM34984 | 44449 | 100026 |
| 451.7686 | 307 | 656.363 | 427 | 319.5843 | 423 | 355.4887 | 478 | CTCF | OCI-Ly3 | None | GSM34984 | 53541 | 100028 |
| 336.7801 | 53 | 601.6225 | 82 | 178.4503 | 50 | 333.7693 | 75 | SREBF1 | A549 | Lung | GSM24241 | 1485 | 100043 |
| 204.1382 | 40 | 301.4936 | 55 | 86.18102 | 35 | 365.6886 | 81 | SREBF1 | A549 | Lung | GSM24241 | 1620 | 100044 |
| 603.185 | 302 | 741.1494 | 429 | 410.395 | 392 | 443.5398 | 438 | CTCF | A549 | Lung | GSM24377 | 38821 | 100048 |
| 446.3426 | 250 | 683.7955 | 338 | 331.2708 | 339 | 348.4978 | 375 | CTCF | A549 | Lung | GSM24377 | 34727 | 100049 |
| 375.2141 | 218 | 605.8386 | 302 | 274.7246 | 295 | 312.8849 | 336 | CTCF | A549 | Lung | GSM24377 | 30837 | 100050 |
| 113.8235 | 76 | 88.08957 | 77 | 117.0243 | 119 | 257.7815 | 178 | JUNB | A549 | Lung | GSM24377 | 11716 | 100052 |
| 148.7499 | 107 | 134.2597 | 116 | 155.0063 | 169 | 274.3445 | 229 | JUNB | A549 | Lung | GSM24377 | 17469 | 100053 |
| 349.5181 | 56 | 267.4812 | 52 | 152.4951 | 48 | 317.5393 | 76 | JUN | A549 | Lung | GSM24377 | 1681 | 100055 |
| 1092.616 | 596 | 1093.004 | 724 | 799.8657 | 587 | 792.911 | 659 | JUN | A549 | Lung | GSM24377 | 25326 | 100056 |
| 507.4532 | 270 | 732.3583 | 396 | 374.6677 | 364 | 384.7667 | 399 | CTCF | A549 | Lung | GSM24377 | 36480 | 100057 |
| 297.7621 | 121 | 465.9998 | 168 | 126.5088 | 125 | 172.3877 | 153 | CTCF | A549 | Lung | GSM24377 | 12100 | 100058 |
| 380.1576 | 188 | 597.7891 | 259 | 261.0402 | 243 | 314.7756 | 283 | CTCF | A549 | Lung | GSM24377 | 22834 | 100059 |
| 1039.751 | 323 | 1051.426 | 405 | 835.452 | 359 | 815.6532 | 390 | JUN | A549 | Lung | GSM24377 | 12586 | 100060 |
| 839.2745 | 133 | 1070.695 | 167 | 540.4453 | 141 | 692.4829 | 171 | JUN | A549 | Lung | GSM24377 | 3912 | 100061 |
| 494.5785 | 262 | 707.1796 | 360 | 338.4975 | 343 | 347.5057 | 376 | CTCF | A549 | Lung | GSM24377 | 35115 | 100062 |
| 430.5531 | 251 | 684.5338 | 349 | 334.2752 | 347 | 327.1069 | 374 | CTCF | A549 | Lung | GSM24377 | 35646 | 100063 |
| 417.6637 | 242 | 701.3036 | 352 | 324.6956 | 335 | 342.6249 | 371 | CTCF | A549 | Lung | GSM24377 | 34457 | 100064 |
| 1049.88 | 404 | 1055.531 | 498 | 841.3496 | 470 | 825.6564 | 510 | ZHX2 | MCF-7 | Mammary | GSM25345 | 17992 | 100071 |
| 1051.356 | 437 | 1053.683 | 534 | 772.8567 | 427 | 749.2832 | 454 | KLF16 | HepG2 | None | GSM25273 | 19743 | 100076 |
| 1191.853 | 164 | 1210.788 | 212 | 799.2946 | 168 | 676.978 | 162 | TARDBP | GM12878 | None | GSM25273 | 3885 | 100079 |
| 1124.939 | 333 | 1117.621 | 397 | 849.1233 | 316 | 809.5608 | 320 | TARDBP | GM12878 | None | GSM25273 | 11196 | 100080 |
| 1039.374 | 372 | 1074.454 | 487 | 790.2936 | 383 | 790.3509 | 433 | SP5 | HepG2 | None | GSM25273 | 16949 | 100085 |
| 1113.078 | 609 | 1086.674 | 699 | 790.607 | 566 | 775.8583 | 625 | SP5 | HepG2 | None | GSM25273 | 27332 | 100086 |
| 863.1993 | 416 | 900.7001 | 551 | 645.4127 | 467 | 642.8017 | 529 | CTCF | None | gastroesop | GSM28255 | 33371 | 100099 |
| 965.6594 | 409 | 1002.408 | 539 | 756.5645 | 465 | 778.7904 | 559 | PKNOX1 | MCF-7 | Breast | GSM24242 | 22700 | 100104 |
| 1091.776 | 297 | 1113.674 | 383 | 875.1141 | 319 | 868.7957 | 359 | PKNOX1 | MCF-7 | Breast | GSM24242 | 9840 | 100105 |
| 1174.128 | 258 | 1212.412 | 338 | 910.8949 | 233 | 941.2998 | 288 | SP1 | K562 | None | GSM24242 | 5751 | 100107 |
| 116.9785 | 91 | 116.5368 | 103 | 79.21212 | 124 | 114.8257 | 154 | ZNF316 | K562 | None | GSM24231 | 15960 | 100128 |
| 78.38829 | 205 | 74.28025 | 235 | 55.00293 | 323 | 62.14149 | 367 | ZNF316 | K562 | None | GSM24231 | 74660 | 100129 |
| 974.8877 | 213 | 995.8859 | 276 | 561.1905 | 217 | 709.5706 | 260 | SMARCA5 | GM12878 | Blood | GSM28284 | 9590 | 100130 |
| 928.3845 | 372 | 943.0327 | 473 | 697.1936 | 397 | 685.7774 | 437 | SMARCA5 | GM12878 | Blood | GSM28284 | 22080 | 100131 |
| 735.8155 | 86 | 487.7265 | 72 | 508.6624 | 91 | 745.2556 | 120 | MTA1 | K562 | None | GSM28284 | 1564 | 100132 |
| 898.9458 | 223 | 824.6692 | 235 | 540.8763 | 246 | 648.7821 | 287 | MTA1 | K562 | None | GSM28284 | 12613 | 100133 |
| 51.23676 | 36 | 63.88886 | 45 | 25.93125 | 44 | 73.08485 | 71 | NFE2L2 | A549 | Lung | GSM24237 | 6230 | 100134 |
| 66.62843 | 43 | 58.5771 | 46 | 26.93692 | 48 | 80.79611 | 79 | NFE2L2 | A549 | Lung | GSM24237 | 7060 | 100135 |
| 769.374 | 371 | 808.1897 | 498 | 590.2589 | 457 | 579.0451 | 505 | CTCF | None | Liver | GSM28284 | 38205 | 100136 |
| 669.1718 | 368 | 716.6953 | 500 | 453.1692 | 479 | 479.4325 | 532 | CTCF | DOHH-2 | None | GSM26986 | 51344 | 100139 |
| 84.73122 | 31 | 97.35336 | 37 | 26.71075 | 27 | 86.47942 | 49 | CTCF | DOHH-2 | None | GSM26986 | 2610 | 100140 |
| 889.116 | 456 | 922.6462 | 591 | 661.3501 | 502 | 657.5693 | 566 | CTCF | None | Lung | GSM28287 | 33321 | 100143 |
| 848.7054 | 291 | 883.1935 | 385 | 580.1467 | 327 | 642.8349 | 371 | CTCF | None | Aorta | GSM28287 | 21625 | 100144 |
| 907.1756 | 550 | 932.7447 | 695 | 669.1469 | 607 | 651.8271 | 665 | CTCF | None | Thyroid | GSM28287 | 42448 | 100147 |
| 555.1834 | 111 | 632.7655 | 131 | 319.3209 | 116 | 553.9348 | 164 | SMARCA5 | MCF-7 | Breast | GSM28275 | 4921 | 100162 |
| 414.1262 | 79 | 461.1611 | 92 | 222.7959 | 79 | 373.1544 | 110 | SMARCA5 | MCF-7 | Breast | GSM28275 | 3274 | 100163 |
| 214.517 | 34 | 253.88 | 41 | 144.9969 | 37 | 255.6647 | 54 | USF2 | A549 | Lung | GSM28275 | 1038 | 100164 |
| 514.1699 | 83 | 472.6169 | 86 | 250.1839 | 76 | 559.2831 | 125 | USF2 | A549 | Lung | GSM28275 | 2580 | 100165 |
| 23.00096 | 19 | 21.54426 | 21 | 7.420098 | 20 | 183.9469 | 81 | ZNF157 | HEK293 | Embryonic | GSM28275 | 3430 | 100166 |
| 41.46375 | 25 | 34.76873 | 26 | 17.15482 | 27 | 206.1751 | 86 | ZNF157 | HEK293 | Embryonic | GSM28275 | 3458 | 100167 |
| 39.64198 | 18 | 28.01802 | 17 | 22.68048 | 21 | 149.1996 | 55 | ZNF473 | HEK293 | Embryonic | GSM28275 | 1934 | 100170 |
| 90.22212 | 25 | 36.02521 | 17 | 30.5529 | 21 | 202.0005 | 58 | ZNF473 | HEK293 | Embryonic | GSM28275 | 1557 | 100171 |
| 97.1179 | 108 | 145.9958 | 144 | 42.92298 | 134 | 86.80592 | 182 | CEBPG | K562 | None | GSM28275 | 23249 | 100172 |
| 106.7188 | 95 | 147.7685 | 123 | 70.36343 | 129 | 115.7213 | 167 | CEBPG | K562 | None | GSM28275 | 17290 | 100173 |
| 745.8602 | 309 | 755.7605 | 387 | 469.0123 | 374 | 566.5102 | 436 | SMC3 | A549 | Lung | GSM31064 | 32902 | 100174 |
| 760.7256 | 341 | 779.7039 | 434 | 591.4907 | 431 | 579.382 | 476 | SMC3 | A549 | Lung | GSM31064 | 36741 | 100175 |
| 771.8116 | 403 | 813.896 | 537 | 606.4824 | 515 | 605.5733 | 589 | SMC3 | A549 | Lung | GSM31064 | 42635 | 100176 |
| 718.2865 | 321 | 735.9849 | 410 | 502.0625 | 413 | 566.8592 | 478 | RAD21 | A549 | Lung | GSM31064 | 37063 | 100177 |
| 735.2227 | 343 | 766.5872 | 452 | 560.9816 | 441 | 567.5045 | 495 | RAD21 | A549 | Lung | GSM31064 | 39706 | 100178 |
| 575.3538 | 271 | 728.1114 | 362 | 427.2746 | 361 | 498.2846 | 415 | RAD21 | A549 | Lung | GSM31064 | 32897 | 100179 |
| 1128.238 | 267 | 1145.07 | 338 | 878.1318 | 251 | 891.2904 | 296 | LARP7 | GM12878 | Blood | GSM28273 | 6663 | 100188 |

| | | | | | | | | | | | | | |
|---|---|---|---|---|---|---|---|---|---|---|---|---|---|
| 1108.563 | 196 | 1099.235 | 234 | 659.4601 | 181 | 656.48 | 192 | LARP7 | GM12878 | Blood | GSM28273 | 4893 | 100189 |
| 128.7895 | 38 | 152.4046 | 46 | 71.23283 | 41 | 245.5154 | 81 | ZNF24 | HepG2 | None | GSM28273 | 2613 | 100190 |
| 177.2011 | 51 | 250.9098 | 68 | 148.2841 | 67 | 294.9118 | 101 | ZNF24 | HepG2 | None | GSM28273 | 3417 | 100191 |
| 72.95129 | 19 | 73.20892 | 21 | 57.87309 | 24 | 195.174 | 49 | ZNF302 | HEK293 | Embryonic | GSM28273 | 1144 | 100194 |
| 108.0772 | 28 | 78.9681 | 26 | 79.62392 | 34 | 269.8098 | 69 | ZNF302 | HEK293 | Embryonic | GSM28273 | 1643 | 100195 |
| 37.71252 | 15 | 16.19892 | 11 | 26.80856 | 19 | 263.0763 | 65 | RBAK | HEK293 | Embryonic | GSM28273 | 1472 | 100197 |
| 76.99033 | 24 | 49.11163 | 21 | 39.15056 | 25 | 275.2255 | 72 | RBAK | HEK293 | Embryonic | GSM28273 | 1744 | 100198 |
| 20.80497 | 12 | 19.09075 | 13 | 24.73476 | 20 | 232.7312 | 65 | ZNF211 | HEK293 | Embryonic | GSM28273 | 1698 | 100201 |
| 27.34422 | 11 | 18.25961 | 10 | 5.412065 | 8 | 288.6593 | 61 | ZNF211 | HEK293 | Embryonic | GSM28273 | 1137 | 100202 |
| 443.33 | 68 | 474.3582 | 77 | 258.9048 | 68 | 265.0031 | 73 | BRD4 | HepG2 | Liver | GSM28273 | 1944 | 100204 |
| 1125.246 | 536 | 1127.82 | 653 | 846.6303 | 536 | 808.7226 | 551 | DMAP1 | HepG2 | None | GSM25342 | 19434 | 100205 |
| 1121.256 | 416 | 1151.183 | 539 | 868.924 | 428 | 837.9976 | 445 | DMAP1 | HepG2 | None | GSM25342 | 14198 | 100206 |
| 690.8734 | 223 | 819.4279 | 284 | 500.3959 | 283 | 665.7239 | 345 | NR2F1 | HepG2 | None | GSM25342 | 19515 | 100211 |
| 261.4021 | 93 | 516.8132 | 147 | 314.4628 | 150 | 355.2482 | 170 | NR2F1 | HepG2 | None | GSM25342 | 8592 | 100212 |
| 854.6263 | 338 | 843.2179 | 401 | 654.8161 | 398 | 655.2749 | 451 | TEAD3 | HepG2 | None | GSM25342 | 27618 | 100215 |
| 888.5363 | 526 | 904.8288 | 658 | 663.1684 | 603 | 653.5384 | 673 | TEAD3 | HepG2 | None | GSM25342 | 44859 | 100216 |
| 1118.596 | 251 | 1151.299 | 336 | 883.8198 | 257 | 817.5148 | 246 | U2AF1 | HepG2 | None | GSM28289 | 7505 | 100219 |
| 782.0392 | 121 | 1145.065 | 176 | 485.1285 | 126 | 522.1358 | 139 | U2AF1 | HepG2 | None | GSM28289 | 3604 | 100220 |
| 1210.308 | 213 | 1239.457 | 277 | 861.4989 | 190 | 828.9641 | 197 | ELF4 | HEK293T | Embryonic | GSM28289 | 4306 | 100221 |
| 1145.408 | 242 | 1186.977 | 323 | 882.181 | 224 | 911.4692 | 272 | ELF4 | HEK293T | Embryonic | GSM28289 | 6094 | 100222 |
| 599.6781 | 140 | 593.0886 | 153 | 271.45 | 133 | 418.2852 | 175 | ARID3A | GM12878 | None | GSM28289 | 7246 | 100226 |
| 811.4886 | 218 | 865.1228 | 264 | 378.2006 | 218 | 429.9202 | 248 | ARID3A | GM12878 | None | GSM28289 | 14064 | 100227 |
| 1146.173 | 424 | 1185.278 | 566 | 855.759 | 397 | 802.9559 | 390 | PCBP1 | HepG2 | None | GSM28289 | 13822 | 100228 |
| 1138.772 | 285 | 1182.478 | 387 | 856.1153 | 255 | 827.6216 | 267 | PCBP1 | HepG2 | None | GSM28289 | 8296 | 100229 |
| 957.6629 | 113 | 971.444 | 123 | 351.9994 | 83 | 459.8902 | 102 | RFX5 | A549 | Lung | GSM28250 | 1973 | 100232 |
| 1170.668 | 152 | 1181.641 | 188 | 632.5069 | 142 | 614.9551 | 148 | RFX5 | A549 | Lung | GSM28250 | 3250 | 100233 |
| 521.5176 | 332 | 681.0964 | 467 | 378.8098 | 455 | 425.3232 | 516 | CTCF | SK-N-SH | Brain | GSM34983 | 53008 | 100236 |
| 654.7775 | 348 | 732.1917 | 490 | 411.1569 | 442 | 533.2918 | 539 | CTCF | SK-N-SH | Brain | GSM34983 | 47074 | 100237 |
| 1021.11 | 400 | 1038.475 | 504 | 774.6957 | 401 | 774.7688 | 457 | ASH2L | HepG2 | Liver | GSM34983 | 16679 | 100238 |
| 1018.367 | 367 | 1048.16 | 477 | 795.4117 | 387 | 770.1104 | 415 | ASH2L | HepG2 | Liver | GSM34983 | 15043 | 100239 |
| 1012.217 | 294 | 1053.279 | 400 | 788.9885 | 305 | 755.5194 | 317 | ASH2L | HepG2 | Liver | GSM34983 | 11259 | 100240 |
| 449.4812 | 345 | 646.1664 | 494 | 327.4803 | 485 | 369.3551 | 551 | CTCF | LOUCY | Peripheral | GSM34983 | 64947 | 100241 |
| 359.9167 | 288 | 645.4113 | 429 | 278.11 | 412 | 332.3498 | 477 | CTCF | LOUCY | Peripheral | GSM34983 | 53258 | 100242 |
| 359.9695 | 211 | 503.9773 | 274 | 246.3292 | 275 | 304.2557 | 323 | CTCF | KMS11 | None | GSM34983 | 27597 | 100243 |
| 349.5128 | 219 | 513.2551 | 290 | 196.5184 | 269 | 216.4151 | 303 | CTCF | KMS11 | None | GSM34983 | 30707 | 100244 |
| 438.949 | 291 | 642.2531 | 387 | 292.8672 | 391 | 378.215 | 465 | CTCF | MCF-7 | Breast | GSM34984 | 49015 | 100249 |
| 514.6394 | 362 | 657.2306 | 504 | 368.9307 | 504 | 424.0201 | 576 | CTCF | MCF-7 | Breast | GSM34984 | 66179 | 100250 |
| 462.7089 | 82 | 577.4181 | 102 | 231.2427 | 78 | 288.0051 | 93 | RFX3 | HepG2 | None | GSM25342 | 3081 | 100269 |
| 428.3732 | 203 | 560.3287 | 257 | 258.1007 | 249 | 208.3778 | 249 | ETV6 | GM12878 | None | GSM25342 | 24353 | 100272 |
| 462.2565 | 158 | 468.3208 | 177 | 286.9662 | 188 | 235.1497 | 186 | ETV6 | GM12878 | None | GSM25342 | 14215 | 100273 |
| 597.0442 | 102 | 843.7135 | 136 | 288.152 | 93 | 383.9392 | 115 | NCOA6 | K562 | None | GSM28255 | 3148 | 100277 |
| 491.354 | 82 | 546.6802 | 95 | 222.6265 | 72 | 341.1196 | 96 | NCOA6 | K562 | None | GSM28255 | 2476 | 100278 |
| 506.3655 | 80 | 507.5207 | 87 | 312.8663 | 82 | 346.9517 | 92 | MYBL2 | K562 | None | GSM28255 | 2182 | 100283 |
| 948.5441 | 158 | 1041.167 | 185 | 522.8906 | 156 | 529.6274 | 167 | MYBL2 | K562 | None | GSM28255 | 5128 | 100284 |
| 1074.393 | 517 | 1100.871 | 652 | 817.9365 | 534 | 772.2396 | 539 | NRF1 | K562 | None | GSM24242 | 20788 | 100311 |
| 1074.668 | 476 | 1091.395 | 595 | 807.6239 | 472 | 782.7698 | 504 | NRF1 | K562 | None | GSM24242 | 18283 | 100312 |
| 665.1925 | 85 | 800.6558 | 103 | 397.0745 | 83 | 432.0245 | 92 | NFRKB | HEK293T | Embryonic | GSM24242 | 1682 | 100313 |
| 581.57 | 73 | 810.653 | 97 | 235.5479 | 57 | 299.2792 | 69 | NFRKB | HEK293T | Embryonic | GSM24242 | 1405 | 100314 |
| 117.2787 | 28 | 153.6339 | 36 | 66.67158 | 30 | 158.2625 | 50 | JUN | A549 | Lung | GSM24242 | 1667 | 100315 |
| 97.82037 | 28 | 128.9393 | 36 | 87.4597 | 38 | 269.8527 | 73 | JUN | MCF-7 | Mammary | GSM24231 | 1874 | 100339 |
| 171.6244 | 43 | 262.1224 | 60 | 98.48617 | 46 | 300.6481 | 87 | JUN | MCF-7 | Mammary | GSM24231 | 2430 | 100340 |
| 459.7438 | 336 | 663.129 | 486 | 323.368 | 460 | 398.0135 | 538 | CTCF | MRC-iPS-9! | None | GSM26985 | 57464 | 100348 |
| 415.517 | 327 | 643.0771 | 472 | 295.8272 | 454 | 337.8901 | 518 | CTCF | MRC-iPS-9! | None | GSM26985 | 59603 | 100349 |
| 88.94005 | 22 | 80.95565 | 23 | 40.78232 | 21 | 72.57597 | 30 | EHMT2 | A549 | Lung | GSM26985 | 1267 | 100351 |
| 65.84156 | 17 | 131.6371 | 28 | 57.8243 | 23 | 215.9567 | 50 | EHMT2 | A549 | Lung | GSM26985 | 1138 | 100352 |
| 1258.843 | 622 | 1314.046 | 814 | 939.3443 | 580 | 889.6492 | 586 | NONO | HepG2 | None | GSM28285 | 16536 | 100355 |
| 1259.864 | 579 | 1321.615 | 770 | 947.0271 | 542 | 904.7221 | 554 | NONO | HepG2 | None | GSM28285 | 14807 | 100356 |
| 1072.874 | 428 | 1081.265 | 525 | 823.4594 | 424 | 787.6762 | 440 | EP400 | K562 | None | GSM28285 | 14156 | 100357 |
| 1051.862 | 369 | 1059.828 | 454 | 802.3369 | 363 | 777.8997 | 386 | EP400 | K562 | None | GSM28285 | 12983 | 100358 |
| 687.0615 | 253 | 801.119 | 340 | 433.9408 | 304 | 446.1843 | 332 | CTCF | None | Sigmoid Cc | GSM28285 | 23216 | 100359 |
| 219.1692 | 50 | 205.7684 | 53 | 87.63051 | 44 | 320.2535 | 91 | ZSCAN18 | HEK293 | Embryonic | GSM28285 | 2462 | 100360 |
| 248.1681 | 51 | 204.2614 | 50 | 123.6838 | 49 | 405.9617 | 98 | ZSCAN18 | HEK293 | Embryonic | GSM28285 | 2166 | 100361 |
| 818.7677 | 346 | 818.4382 | 419 | 514.8622 | 363 | 587.1816 | 414 | RELB | GM12878 | Blood | GSM28273 | 26754 | 100362 |
| 871.113 | 260 | 903.2244 | 348 | 552.6081 | 285 | 534.7652 | 302 | RELB | GM12878 | Blood | GSM28273 | 15978 | 100363 |
| 164.1912 | 31 | 235.8176 | 42 | 85.6777 | 30 | 304.0464 | 64 | IRF5 | GM12878 | None | GSM28288 | 1194 | 100371 |
| 189.3437 | 45 | 383.4915 | 74 | 159.1982 | 58 | 395.1526 | 100 | IRF5 | GM12878 | Blood | GSM28288 | 2397 | 100372 |
| 1047.744 | 535 | 1054.754 | 659 | 765.7978 | 528 | 736.082 | 561 | ELF1 | GM12878 | Blood | GSM28287 | 25398 | 100373 |
| 1061.937 | 547 | 1085.767 | 686 | 763.8547 | 519 | 746.2773 | 567 | ELF1 | GM12878 | Blood | GSM28287 | 25294 | 100374 |
| 164.3007 | 45 | 191.9065 | 54 | 91.60414 | 48 | 258.0317 | 86 | ZNF148 | HEK293 | Embryonic | GSM28287 | 2762 | 100375 |
| 78.35774 | 27 | 50.73976 | 24 | 54.52283 | 33 | 187.638 | 65 | ZNF148 | HEK293 | Embryonic | GSM28287 | 2144 | 100376 |
| 414.1425 | 95 | 501.5368 | 116 | 230.6198 | 99 | 487.4166 | 154 | BMI1 | MCF-7 | Breast | GSM28287 | 4762 | 100377 |
| 354.945 | 95 | 471.3484 | 122 | 242.7563 | 112 | 417.074 | 156 | BMI1 | MCF-7 | Breast | GSM28287 | 5775 | 100378 |

| | | | | | | | | | | | | | |
|---|---|---|---|---|---|---|---|---|---|---|---|---|---|
| 784.205 | 345 | 815.143 | 454 | 601.7783 | 417 | 578.9636 | 447 | CTCF | None | Esophagus | GSM28287 | 32814 | 100379 |
| 558.7672 | 319 | 709.6443 | 450 | 425.8155 | 434 | 422.632 | 471 | CTCF | LNCaP | Prostate | GSM28272 | 43039 | 100386 |
| 411.3556 | 192 | 623.921 | 260 | 245.6619 | 231 | 290.2536 | 267 | CTCF | LNCaP | Prostate | GSM28272 | 20868 | 100387 |
| 278.2357 | 252 | 402.4498 | 330 | 271.7661 | 397 | 390.4232 | 489 | FOS | MCF-7 | Mammary | GSM28275 | 52578 | 100391 |
| 439.7828 | 243 | 522.0736 | 294 | 371.3698 | 344 | 487.7838 | 414 | FOS | MCF-7 | Mammary | GSM28275 | 32485 | 100392 |
| 743.0507 | 360 | 773.7028 | 472 | 523.9049 | 434 | 545.3511 | 478 | CTCF | None | Thyroid | GSM28275 | 38398 | 100393 |
| 940.8833 | 381 | 974.8482 | 505 | 707.5094 | 405 | 675.8316 | 421 | CTCF | None | Ovary | GSM28275 | 21665 | 100394 |
| 1082.95 | 452 | 1096.33 | 565 | 830.9527 | 464 | 796.7181 | 487 | EP300 | None | tibial nerve | GSM28275 | 17791 | 100395 |
| 210.164 | 79 | 230.2056 | 92 | 154.1425 | 101 | 189.5062 | 119 | ARID2 | K562 | None | GSM28275 | 7184 | 100396 |
| 238.7708 | 87 | 297.0273 | 108 | 182.0784 | 113 | 190.9282 | 124 | ARID2 | K562 | None | GSM28275 | 7775 | 100397 |
| 232.9845 | 40 | 254.9553 | 46 | 109.3516 | 36 | 264.6939 | 62 | BCLAF1 | K562 | None | GSM28275 | 1329 | 100398 |
| 100.2122 | 25 | 138.004 | 33 | 87.26552 | 33 | 273.6617 | 65 | BCLAF1 | K562 | None | GSM28275 | 1427 | 100399 |
| 836.9048 | 263 | 861.8373 | 345 | 490.7505 | 291 | 511.7526 | 319 | CTCF | None | Testis | GSM28275 | 19729 | 100400 |
| 1098.023 | 182 | 1113.377 | 231 | 673.8192 | 182 | 685.6114 | 195 | GTF2F1 | HepG2 | None | GSM28273 | 5633 | 100403 |
| 1144.595 | 209 | 1119.893 | 237 | 749.9536 | 194 | 792.6262 | 212 | GTF2F1 | HepG2 | None | GSM28273 | 5721 | 100404 |
| 130.9037 | 41 | 145.0968 | 48 | 58.98211 | 41 | 227.767 | 84 | TOE1 | HepG2 | None | GSM28273 | 3151 | 100405 |
| 121.2679 | 28 | 151.6317 | 35 | 57.54764 | 27 | 314.6301 | 71 | TOE1 | HepG2 | None | GSM28273 | 1509 | 100406 |
| 869.7645 | 554 | 899.7232 | 712 | 652.2908 | 651 | 641.3886 | 726 | CTCF | None | Peyer's pat | GSM28273 | 49861 | 100408 |
| 762.7651 | 135 | 814.439 | 153 | 454.532 | 140 | 585.9298 | 170 | RAD51 | MCF-7 | Breast | GSM28273 | 4773 | 100409 |
| 524.7286 | 93 | 434.2351 | 91 | 343.3507 | 101 | 534.604 | 136 | RAD51 | MCF-7 | Breast | GSM28273 | 3246 | 100410 |
| 727.5941 | 104 | 807.0928 | 120 | 357.6235 | 93 | 295.1998 | 89 | BRD4 | K562 | None | GSM27004 | 2494 | 100415 |
| 787.9942 | 336 | 814.3039 | 440 | 598.5898 | 406 | 542.7242 | 422 | CTCF | None | Esophagus | GSM28289 | 32013 | 100428 |
| 859.9406 | 499 | 908.081 | 667 | 633.0104 | 567 | 630.2392 | 640 | CTCF | None | Adrenal Gl | GSM28289 | 44621 | 100429 |
| 635.2217 | 96 | 936.6939 | 132 | 202.8253 | 70 | 466.5767 | 116 | U2AF2 | HepG2 | None | GSM28288 | 2826 | 100431 |
| 798.5509 | 119 | 1139.16 | 161 | 462.2663 | 119 | 468.1503 | 127 | U2AF2 | HepG2 | None | GSM28288 | 3423 | 100432 |
| 15.4537 | 9 | 18.12693 | 11 | 7.041534 | 10 | 177.2365 | 50 | ATM | HepG2 | None | GSM28288 | 1315 | 100433 |
| 42.19633 | 13 | 34.83281 | 13 | 11.19072 | 10 | 250.3978 | 53 | ATM | HepG2 | None | GSM28288 | 1003 | 100434 |
| 1.051778 | 6 | 1.04882 | 7 | 1.008876 | 11 | 34.49593 | 37 | ZNF555 | HEK293 | Embryonic | GSM28288 | 3395 | 100436 |
| 950.9443 | 643 | 959.5185 | 787 | 673.3862 | 677 | 646.0579 | 726 | BHLHE40 | GM12878 | Blood | GSM28288 | 49009 | 100437 |
| 1023.184 | 485 | 1031.683 | 604 | 736.7271 | 470 | 726.2292 | 519 | BHLHE40 | GM12878 | Blood | GSM28288 | 24554 | 100438 |
| 510.7077 | 139 | 710.1779 | 183 | 459.0065 | 190 | 512.0433 | 214 | RERE | HepG2 | None | GSM25347 | 9526 | 100439 |
| 860.0783 | 202 | 898.0147 | 235 | 664.6308 | 252 | 742.1505 | 284 | RERE | HepG2 | None | GSM25347 | 11578 | 100440 |
| 581.888 | 379 | 679.5607 | 527 | 421.4204 | 521 | 410.4848 | 564 | CTCF | OCI-Ly1 | Bone Marr | GSM34983 | 62596 | 100454 |
| 444.818 | 240 | 647.4272 | 318 | 308.9617 | 318 | 370.0826 | 369 | CTCF | OCI-Ly1 | Bone Marr | GSM34983 | 33466 | 100455 |
| 57.04115 | 16 | 135.0877 | 29 | 46.18593 | 21 | 186.413 | 47 | SUZ12 | GM12878 | Blood | GSM34983 | 1196 | 100456 |
| 451.3506 | 299 | 655.9859 | 409 | 313.027 | 406 | 301.3014 | 437 | CTCF | None | Embryo | GSM34983 | 49616 | 100457 |
| 694.1574 | 452 | 762.6129 | 634 | 531.7687 | 593 | 506.5212 | 636 | CTCF | None | Embryo | GSM34983 | 65982 | 100458 |
| 678.1872 | 411 | 732.6102 | 568 | 499.3883 | 533 | 479.723 | 573 | CTCF | None | Embryo | GSM34983 | 60065 | 100459 |
| 402.3663 | 296 | 643.7543 | 418 | 291.3075 | 412 | 324.5432 | 466 | CTCF | HCT-116 | HCT116 | GSM34983 | 52566 | 100462 |
| 334.4113 | 287 | 560.0184 | 397 | 244.4948 | 407 | 322.0752 | 486 | CTCF | HCT-116 | HCT116 | GSM34983 | 56307 | 100463 |
| 285.795 | 87 | 419.0692 | 118 | 157.9321 | 96 | 150.8984 | 101 | TCF7 | GM12878 | None | GSM25340 | 6732 | 100468 |
| 638.4108 | 196 | 670.4247 | 223 | 312.1264 | 207 | 252.6317 | 204 | TCF7 | GM12878 | None | GSM25340 | 15759 | 100469 |
| 877.382 | 518 | 896.287 | 648 | 670.2737 | 619 | 671.6651 | 710 | TEAD1 | HepG2 | None | GSM25340 | 46054 | 100472 |
| 825.4275 | 345 | 839.2538 | 433 | 646.287 | 424 | 651.7804 | 490 | TEAD1 | HepG2 | None | GSM25340 | 30458 | 100473 |
| 866.2064 | 501 | 873.6704 | 617 | 678.991 | 627 | 677.9486 | 714 | RARA | HepG2 | None | GSM25340 | 45158 | 100474 |
| 893.0315 | 579 | 910.0413 | 723 | 695.9642 | 724 | 687.5563 | 814 | RARA | HepG2 | None | GSM25340 | 52760 | 100475 |
| 1010.23 | 211 | 996.1761 | 251 | 567.4176 | 208 | 506.3887 | 210 | ZFP64 | HepG2 | None | GSM25340 | 9203 | 100479 |
| 420.4997 | 81 | 422.9613 | 89 | 140.8105 | 64 | 223.7419 | 86 | ZFP64 | HepG2 | None | GSM25340 | 3418 | 100480 |
| 235.8116 | 89 | 402.59 | 130 | 179.6056 | 117 | 180.3702 | 126 | ZNF652 | HepG2 | None | GSM25340 | 8682 | 100485 |
| 779.881 | 315 | 811.8669 | 418 | 596.4765 | 396 | 609.7387 | 452 | ZNF652 | HepG2 | None | GSM25340 | 31891 | 100486 |
| 330.5017 | 77 | 376.6496 | 91 | 305.204 | 104 | 485.9585 | 141 | ESRRA | MCF-7 | Mammary | GSM24241 | 4022 | 100495 |
| 252.7388 | 69 | 417.4229 | 100 | 273.4422 | 102 | 406.1143 | 133 | ESRRA | MCF-7 | Mammary | GSM24241 | 4215 | 100496 |
| 760.4225 | 252 | 831.861 | 333 | 626.9987 | 336 | 570.1994 | 347 | REST | K562 | None | GSM24230 | 21206 | 100557 |
| 760.2769 | 441 | 782.0959 | 560 | 589.8801 | 562 | 562.1466 | 596 | REST | K562 | None | GSM24230 | 49606 | 100558 |
| 993.5364 | 249 | 977.4714 | 294 | 680.4355 | 257 | 582.9792 | 255 | RNF2 | K562 | None | GSM24230 | 12094 | 100559 |
| 322.8809 | 54 | 399.9817 | 67 | 117.1988 | 43 | 196.3319 | 60 | RNF2 | K562 | None | GSM24230 | 1918 | 100560 |
| 188.425 | 150 | 170.3193 | 164 | 53.7359 | 159 | 98.22813 | 209 | MAFF | HeLa-S3 | Cervix | GSM24230 | 29752 | 100561 |
| 744.4295 | 340 | 731.0546 | 401 | 423.4925 | 391 | 493.5093 | 449 | MAFF | HeLa-S3 | Cervix | GSM24230 | 40494 | 100562 |
| 906.2879 | 511 | 941.311 | 661 | 678.9511 | 573 | 657.3262 | 624 | CTCF | None | Prostate | GSM28285 | 38277 | 100571 |
| 988.6182 | 218 | 959.5393 | 244 | 658.7967 | 232 | 596.7606 | 236 | MTA3 | K562 | None | GSM28285 | 9151 | 100572 |
| 1060.911 | 396 | 1069.128 | 483 | 820.4458 | 398 | 780.2017 | 408 | MTA3 | K562 | None | GSM28285 | 13799 | 100573 |
| 1181.718 | 250 | 1208.176 | 323 | 931.1033 | 233 | 921.0739 | 259 | MAZ | MCF-7 | Mammary | GSM24232 | 5428 | 100574 |
| 1207.849 | 332 | 1217.604 | 404 | 924.7443 | 294 | 922.9799 | 332 | MAZ | MCF-7 | Mammary | GSM24232 | 7036 | 100575 |
| 31.02158 | 12 | 29.29853 | 13 | 31.71976 | 18 | 120.165 | 38 | PAX8 | MCF-7 | Breast | GSM28286 | 1152 | 100588 |
| 44.03675 | 19 | 56.01663 | 24 | 30.34476 | 24 | 142.6126 | 54 | PAX8 | MCF-7 | Breast | GSM28286 | 2004 | 100589 |
| 1035.252 | 142 | 1150.358 | 173 | 626.3112 | 142 | 741.088 | 165 | MNT | HepG2 | None | GSM28286 | 3239 | 100590 |
| 1084.605 | 211 | 1140.651 | 298 | 789.9969 | 217 | 872.5254 | 259 | MNT | HepG2 | None | GSM28286 | 6642 | 100591 |
| 579.473 | 75 | 863.0724 | 104 | 346.5891 | 74 | 627.1173 | 110 | FOXM1 | HEK293T | Embryonic | GSM28286 | 1621 | 100592 |
| 1124.887 | 152 | 1194.419 | 207 | 601.7927 | 143 | 864.7923 | 185 | FOXM1 | HEK293T | Embryonic | GSM28286 | 3644 | 100593 |
| 633.154 | 82 | 907.2269 | 111 | 344.8485 | 77 | 444.9501 | 94 | SRSF1 | K562 | None | GSM28286 | 1820 | 100596 |
| 733.4901 | 96 | 1040.083 | 129 | 468.9109 | 99 | 554.5163 | 115 | SRSF1 | K562 | None | GSM28286 | 2175 | 100597 |
| 213.1153 | 59 | 250.965 | 71 | 115.8766 | 62 | 311.9784 | 108 | ZNF785 | HEK293 | Embryonic | GSM28286 | 3510 | 100598 |

| | | | | | | | | | | | | | |
|---|---|---|---|---|---|---|---|---|---|---|---|---|---|
| 188.3765 | 48 | 183.0979 | 52 | 94.84054 | 48 | 301.7159 | 92 | ZNF785 | HEK293 | Embryonic | GSM28286 | 2634 | 100599 |
| 614.8671 | 521 | 626.1311 | 648 | 385.7199 | 676 | 437.8151 | 777 | EOMES | H9 | Embryo | GSM64069 | 109214 | 1006 |
| 270.4511 | 63 | 262.1891 | 68 | 122.0671 | 59 | 239.0481 | 88 | LEF1 | K562 | None | GSM28286 | 3124 | 100600 |
| 295.3083 | 52 | 391.1853 | 67 | 166.721 | 52 | 329.7679 | 80 | LEF1 | K562 | None | GSM28286 | 1796 | 100601 |
| 54.42167 | 22 | 155.9186 | 43 | 60.32156 | 35 | 137.7228 | 56 | CEBPB | GM12878 | Blood | GSM24238 | 2446 | 100602 |
| 275.3407 | 48 | 354.8781 | 61 | 140.5626 | 46 | 225.8119 | 63 | CEBPB | GM12878 | Blood | GSM24238 | 1798 | 100603 |
| 509.621 | 229 | 705.4306 | 298 | 280.67 | 273 | 356.3498 | 324 | CTCF | None | Embryo | GSM26991 | 28406 | 100604 |
| 707.3498 | 327 | 760.513 | 455 | 440.6944 | 411 | 462.3228 | 454 | CTCF | None | Embryo | GSM26991 | 42024 | 100605 |
| 688.6235 | 336 | 740.1971 | 462 | 419.9755 | 421 | 490.4938 | 484 | CTCF | None | Embryo | GSM26991 | 44555 | 100606 |
| 236.491 | 45 | 277.7132 | 54 | 114.7928 | 42 | 136.7561 | 49 | CHD4 | A549 | Lung | GSM27004 | 1714 | 100615 |
| 1039.298 | 172 | 1056.7 | 208 | 612.9982 | 175 | 631.6403 | 189 | FOXK2 | GM12878 | Blood | GSM28289 | 5311 | 100620 |
| 1093.667 | 182 | 1100.933 | 223 | 669.5014 | 180 | 661.3204 | 190 | FOXK2 | GM12878 | None | GSM28289 | 5052 | 100621 |
| 631.3045 | 227 | 823.439 | 318 | 369.7119 | 264 | 439.381 | 306 | CTCF | None | gastrocnen | GSM28289 | 20672 | 100622 |
| 36.78376 | 19 | 19.8173 | 16 | 5.901632 | 14 | 206.3263 | 71 | ZNF791 | HEK293 | Embryonic | GSM28289 | 2366 | 100623 |
| 21.88835 | 13 | 7.367273 | 9 | 21.58165 | 20 | 172.3289 | 58 | ZNF791 | HEK293 | Embryonic | GSM28289 | 1884 | 100624 |
| 74.99013 | 22 | 56.99758 | 21 | 42.98296 | 24 | 295.7857 | 70 | ZNF701 | HEK293 | Embryonic | GSM28275 | 1498 | 100625 |
| 89.50301 | 25 | 74.52837 | 25 | 54.97704 | 28 | 295.7694 | 72 | ZNF701 | HEK293 | Embryonic | GSM28275 | 1601 | 100626 |
| 1049.426 | 181 | 1079.161 | 242 | 658.4451 | 192 | 701.1604 | 211 | YY2 | HEK293 | Embryonic | GSM28275 | 6075 | 100627 |
| 883.1775 | 161 | 1058.672 | 222 | 638.4321 | 185 | 639.9118 | 197 | YY2 | HEK293 | Embryonic | GSM28275 | 5789 | 100628 |
| 951.0311 | 402 | 986.7963 | 531 | 734.9177 | 447 | 715.6084 | 484 | CTCF | None | Ovary | GSM28275 | 23016 | 100629 |
| 14.4823 | 16 | 91.55176 | 42 | 17.3809 | 28 | 98.43563 | 62 | ZNF316 | K562 | None | GSM28254 | 3706 | 100633 |
| 141.4381 | 101 | 289.7633 | 156 | 104.7263 | 139 | 167.6732 | 180 | ZNF316 | K562 | None | GSM28254 | 15531 | 100634 |
| 1059.052 | 750 | 1089.485 | 943 | 769.5836 | 816 | 745.7614 | 890 | ARID4B | HepG2 | None | GSM25347 | 48326 | 100639 |
| 1079.615 | 654 | 1119.079 | 843 | 803.8714 | 702 | 784.866 | 765 | ARID4B | HepG2 | None | GSM25347 | 34415 | 100640 |
| 469.0846 | 278 | 678.9824 | 379 | 343.458 | 370 | 325.2954 | 395 | CTCF | 22RV1 | Prostate | GSM28288 | 35597 | 100643 |
| 385.181 | 236 | 630.659 | 327 | 308.43 | 326 | 281.6347 | 343 | CTCF | 22RV1 | Prostate | GSM28288 | 31082 | 100644 |
| 1090.843 | 180 | 1091.583 | 218 | 623.8464 | 174 | 448.9452 | 156 | REST | A549 | Lung | GSM27017 | 5395 | 100656 |
| 361.8557 | 93 | 407.3874 | 109 | 142.5078 | 83 | 63.48964 | 63 | REST | A549 | Lung | GSM27017 | 4962 | 100657 |
| 1070.048 | 686 | 1125.613 | 895 | 791.5455 | 742 | 768.1377 | 803 | GABPB1 | HepG2 | None | GSM25340 | 39675 | 100658 |
| 1102.638 | 664 | 1132.767 | 838 | 811.6657 | 679 | 774.6231 | 709 | GABPB1 | HepG2 | None | GSM25340 | 30134 | 100659 |
| 787.5325 | 141 | 813.5621 | 157 | 356.4589 | 128 | 573.7742 | 174 | ZSCAN29 | GM12878 | Blood | GSM28271 | 5305 | 100660 |
| 404.6811 | 83 | 529.6715 | 106 | 189.6749 | 78 | 358.6184 | 115 | ZSCAN29 | GM12878 | Blood | GSM28271 | 3654 | 100661 |
| 679.1856 | 251 | 791.8932 | 329 | 405.7949 | 296 | 534.4439 | 358 | CTCF | None | vagina | GSM28271 | 23686 | 100664 |
| 62.13449 | 23 | 59.43954 | 25 | 45.33135 | 29 | 195.5588 | 64 | ZSCAN26 | HEK293 | Embryonic | GSM28271 | 1974 | 100667 |
| 97.11826 | 28 | 50.8444 | 22 | 50.40107 | 29 | 218.0241 | 65 | ZSCAN26 | HEK293 | Embryonic | GSM28271 | 1823 | 100668 |
| 546.8279 | 254 | 558.6555 | 288 | 337.8193 | 318 | 396.0316 | 366 | FOS | IMR90 | Lung | GSM28254 | 32475 | 100684 |
| 800.5856 | 421 | 775.8506 | 482 | 547.6053 | 455 | 560.1057 | 517 | FOS | IMR90 | Lung | GSM28254 | 44627 | 100685 |
| 24.72891 | 10 | 16.239 | 9 | 8.26149 | 9 | 203.1846 | 48 | NBN | HepG2 | None | GSM28254 | 1006 | 100687 |
| 877.761 | 649 | 901.7429 | 812 | 615.0203 | 714 | 582.4403 | 760 | EP300 | H9 | Embryo | GSM60229 | 66848 | 1007 |
| 638.1422 | 208 | 708.6702 | 243 | 356.441 | 232 | 452.909 | 277 | CC2D1A | K562 | None | GSM28272 | 16397 | 100700 |
| 592.5446 | 172 | 573.0748 | 187 | 294.1688 | 178 | 394.2842 | 218 | CC2D1A | K562 | None | GSM28272 | 11700 | 100701 |
| 86.93506 | 27 | 48.24684 | 22 | 62.56407 | 33 | 221.411 | 67 | ZNF776 | HEK293 | Embryonic | GSM28257 | 1886 | 100710 |
| 64.60953 | 21 | 48.38645 | 20 | 43.23108 | 25 | 167.8378 | 53 | ZNF776 | HEK293 | Embryonic | GSM28257 | 1583 | 100711 |
| 68.50554 | 21 | 56.58839 | 21 | 38.95363 | 23 | 158.8341 | 50 | ZNF248 | HEK293 | Embryonic | GSM28272 | 1514 | 100753 |
| 26.21726 | 15 | 17.48152 | 14 | 28.8521 | 24 | 179.7687 | 62 | ZNF248 | HEK293 | Embryonic | GSM28272 | 2041 | 100754 |
| 1075.109 | 676 | 1108.635 | 857 | 772.3518 | 690 | 760.1362 | 766 | THAP11 | HepG2 | None | GSM25341 | 37941 | 100755 |
| 1057.223 | 666 | 1086.465 | 842 | 776.8701 | 709 | 754.9381 | 773 | THAP11 | HepG2 | None | GSM25341 | 38394 | 100756 |
| 276.131 | 42 | 177.3258 | 35 | 77.25924 | 28 | 215.7321 | 52 | ZC3H8 | K562 | None | GSM28254 | 1164 | 100779 |
| 353.0299 | 57 | 196.3822 | 44 | 142.9124 | 47 | 253.8109 | 68 | ZC3H8 | K562 | None | GSM28254 | 1723 | 100780 |
| 654.9205 | 170 | 801.3064 | 209 | 361.1249 | 184 | 388.8514 | 204 | ZKSCAN1 | HepG2 | None | GSM24231 | 11315 | 100782 |
| 454.3611 | 103 | 496.4049 | 119 | 274.8213 | 113 | 281.2585 | 122 | ZKSCAN1 | HepG2 | None | GSM24231 | 5533 | 100783 |
| 437.262 | 301 | 660.6301 | 428 | 321.4637 | 417 | 347.8386 | 467 | CTCF | OCI-Ly7 | Blood | GSM26984 | 50158 | 100785 |
| 412.1851 | 298 | 644.5629 | 418 | 277.4948 | 400 | 328.7514 | 462 | CTCF | OCI-Ly7 | Blood | GSM26984 | 49130 | 100786 |
| 521.6533 | 169 | 769.1706 | 228 | 273.7097 | 183 | 331.4924 | 214 | CTCF | None | Sigmoid Cc | GSM28285 | 13197 | 100788 |
| 975.8962 | 622 | 1007.434 | 790 | 698.5598 | 641 | 680.3359 | 701 | CTCF | None | tibial nerve | GSM28285 | 39957 | 100790 |
| 1120.055 | 593 | 1167.198 | 777 | 832.5182 | 588 | 773.127 | 577 | RBFOX2 | K562 | None | GSM28285 | 22128 | 100791 |
| 1076.403 | 129 | 1242.562 | 169 | 726.2176 | 135 | 568.7185 | 124 | RBFOX2 | K562 | None | GSM28285 | 2418 | 100792 |
| 6.097034 | 21 | 18.13777 | 35 | 8.013663 | 40 | 27.00437 | 63 | ZNF34 | HEK293 | Embryonic | GSM28285 | 8520 | 100794 |
| 4.330686 | 15 | 1.745879 | 14 | 4.788171 | 28 | 22.102 | 48 | ZNF34 | HEK293 | Embryonic | GSM28285 | 6396 | 100795 |
| 1061.373 | 178 | 1075.592 | 227 | 621.1121 | 182 | 642.6468 | 197 | RFX5 | MCF-7 | Mammary | GSM28285 | 5919 | 100796 |
| 1075.358 | 194 | 1055.269 | 225 | 647.1073 | 189 | 749.866 | 217 | RFX5 | MCF-7 | Mammary | GSM28285 | 6166 | 100797 |
| 693.2735 | 124 | 682.287 | 134 | 365.5437 | 120 | 333.7187 | 122 | TRIM22 | GM12878 | None | GSM24237 | 4224 | 100798 |
| 1038.187 | 276 | 1036.772 | 337 | 691.8364 | 250 | 691.093 | 267 | TRIM22 | GM12878 | None | GSM24237 | 9941 | 100799 |
| 1231.682 | 344 | 1266.679 | 450 | 959.6059 | 326 | 919.433 | 335 | HCFC1 | HepG2 | None | GSM24236 | 7965 | 100811 |
| 1230.172 | 214 | 1256.348 | 277 | 962.9806 | 199 | 950.6514 | 217 | HCFC1 | HepG2 | None | GSM24236 | 4460 | 100812 |
| 1183.902 | 360 | 1203.401 | 459 | 913.061 | 337 | 890.9738 | 356 | NKRF | GM12878 | Blood | GSM28286 | 8419 | 100813 |
| 1213.36 | 399 | 1198.452 | 472 | 917.5946 | 353 | 903.2172 | 389 | NKRF | GM12878 | Blood | GSM28286 | 8920 | 100814 |
| 168.6477 | 56 | 142.4915 | 57 | 80.95805 | 58 | 277.8319 | 111 | LCORL | HepG2 | None | GSM28286 | 4404 | 100815 |
| 91.76229 | 39 | 121.1068 | 50 | 54.05107 | 46 | 223.8666 | 95 | LCORL | HepG2 | None | GSM28286 | 4000 | 100816 |
| 551.1546 | 161 | 785.7823 | 214 | 379.4609 | 194 | 467.5147 | 229 | CTCF | None | Skin | GSM28255 | 11316 | 100824 |
| 1069.772 | 582 | 1080.885 | 723 | 772.0972 | 571 | 766.6247 | 646 | KAT8 | HepG2 | None | GSM25346 | 28582 | 100827 |
| 1036.746 | 606 | 1063.978 | 761 | 770.0605 | 640 | 736.4062 | 676 | KAT8 | HepG2 | None | GSM25346 | 33638 | 100828 |

| | | | | | | | | | | | | | |
|---|---|---|---|---|---|---|---|---|---|---|---|---|---|
| 1245.322 | 446 | 1277.617 | 578 | 954.2714 | 411 | 906.6863 | 410 | PHF8 | A549 | Lung | GSM27003 | 9684 | 100836 |
| 1236.787 | 289 | 1253.181 | 367 | 965.3161 | 270 | 942.6548 | 290 | PHF8 | A549 | Lung | GSM27003 | 6051 | 100837 |
| 934.9098 | 453 | 994.9202 | 628 | 706.4735 | 504 | 702.9197 | 571 | CTCF | None | Uterus | GSM28289 | 30188 | 100844 |
| 41.71283 | 17 | 22.58156 | 14 | 17.2743 | 17 | 241.137 | 66 | ZNF548 | HEK293 | Embryonic | GSM28289 | 1694 | 100845 |
| 53.64763 | 26 | 49.39495 | 28 | 53.19476 | 39 | 290.7566 | 95 | ZNF548 | HEK293 | Embryonic | GSM28289 | 2987 | 100846 |
| 74.05232 | 19 | 108.2844 | 26 | 22.73567 | 15 | 274.2708 | 59 | NONO | HepG2 | None | GSM28274 | 1137 | 100849 |
| 57.78266 | 22 | 46.56545 | 22 | 24.02169 | 22 | 241.2252 | 72 | NONO | HepG2 | None | GSM28274 | 2138 | 100850 |
| 1145.425 | 152 | 1217.205 | 214 | 733.3807 | 158 | 780.7071 | 173 | U2AF1 | K562 | None | GSM28274 | 3620 | 100851 |
| 1204.277 | 196 | 1215.246 | 250 | 758.357 | 174 | 938.9745 | 208 | U2AF1 | K562 | None | GSM28274 | 4334 | 100852 |
| 801.4096 | 176 | 936.8198 | 211 | 501.7509 | 193 | 520.819 | 210 | GATAD2B | K562 | None | GSM28274 | 8177 | 100853 |
| 109.1202 | 40 | 88.24462 | 40 | 70.24366 | 48 | 331.4284 | 109 | FOXP1 | HepG2 | None | GSM28253 | 3542 | 100862 |
| 110.8205 | 48 | 134.7086 | 59 | 134.0269 | 78 | 342.5531 | 131 | FOXP1 | HepG2 | None | GSM28253 | 4983 | 100863 |
| 11.97799 | 9 | 6.877478 | 8 | 17.96035 | 17 | 152.6715 | 51 | ZNF521 | HEK293 | Embryonic | GSM28273 | 1629 | 100864 |
| 25.31037 | 12 | 10.99208 | 9 | 16.55715 | 15 | 233.7095 | 60 | ZNF521 | HEK293 | Embryonic | GSM28273 | 1410 | 100865 |
| 1167.169 | 144 | 1188.613 | 169 | 655.5205 | 136 | 633.0694 | 141 | KDM5A | A549 | Lung | GSM27017 | 2772 | 100867 |
| 1134.302 | 148 | 1172.352 | 176 | 614.3699 | 139 | 667.4313 | 154 | KDM5A | A549 | Lung | GSM27017 | 3281 | 100868 |
| 982.1156 | 553 | 995.2346 | 687 | 713.783 | 579 | 715.7073 | 663 | MIER3 | HepG2 | None | GSM25341 | 36371 | 100871 |
| 979.5615 | 200 | 1000.731 | 259 | 722.5783 | 238 | 624.6423 | 236 | MIER3 | HepG2 | None | GSM25341 | 9540 | 100872 |
| 964.6082 | 672 | 1016.926 | 873 | 717.2853 | 770 | 701.1969 | 850 | CTCF | None | Pancreas | GSM28271 | 52411 | 100877 |
| 480.8023 | 224 | 754.7019 | 322 | 333.3802 | 284 | 337.2239 | 309 | CTCF | RWPE1 | Prostate | GSM28271 | 23303 | 100878 |
| 478.4894 | 220 | 750.9287 | 310 | 315.8072 | 275 | 347.1316 | 309 | CTCF | RWPE1 | Prostate | GSM28271 | 23702 | 100879 |
| 137.1851 | 59 | 173.0066 | 74 | 65.38883 | 65 | 79.441 | 76 | ZNF7 | HepG2 | None | GSM25341 | 6688 | 100882 |
| 252.9065 | 91 | 292.4107 | 109 | 154.8549 | 108 | 119.0537 | 104 | ZNF7 | HepG2 | None | GSM25341 | 8464 | 100883 |
| 1152.287 | 401 | 1183.324 | 523 | 888.9973 | 404 | 827.5936 | 389 | ZHX2 | HepG2 | None | GSM25340 | 13069 | 100886 |
| 1131.116 | 335 | 1154.752 | 429 | 870.826 | 331 | 852.4946 | 356 | ZHX2 | HepG2 | None | GSM25340 | 10841 | 100887 |
| 963.6144 | 430 | 971.6096 | 534 | 733.3088 | 471 | 713.0344 | 512 | ZNF792 | HepG2 | None | GSM25339 | 26091 | 100892 |
| 1019.556 | 328 | 1018.766 | 401 | 788.092 | 349 | 733.511 | 342 | ZNF792 | HepG2 | None | GSM25339 | 14946 | 100893 |
| 739.5631 | 220 | 726.609 | 242 | 582.7428 | 286 | 661.5283 | 325 | NFIA | HepG2 | None | GSM25276 | 17183 | 100927 |
| 839.4066 | 455 | 840.4594 | 552 | 659.4175 | 570 | 672.9401 | 671 | NFIA | HepG2 | None | GSM25276 | 41536 | 100928 |
| 478.1777 | 62 | 513.7637 | 70 | 163.2689 | 44 | 241.7225 | 58 | ZNF207 | GM12878 | None | GSM24230 | 1203 | 100957 |
| 1133.235 | 185 | 1128.483 | 220 | 660.4218 | 169 | 706.4051 | 186 | ZNF207 | GM12878 | None | GSM24230 | 4273 | 100958 |
| 1015.743 | 258 | 1005.205 | 311 | 548.7362 | 226 | 758.0522 | 284 | TCF12 | GM12878 | Blood | GSM28285 | 10962 | 100959 |
| 1006.38 | 241 | 1018.587 | 300 | 527.7649 | 214 | 725.7132 | 267 | TCF12 | GM12878 | Blood | GSM28285 | 10068 | 100960 |
| 263.1745 | 126 | 298.7452 | 149 | 101.5428 | 126 | 153.2118 | 160 | IKZF2 | GM12878 | None | GSM28285 | 12432 | 100964 |
| 745.5582 | 372 | 742.2016 | 445 | 508.1774 | 433 | 425.0146 | 440 | IKZF2 | GM12878 | Blood | GSM28285 | 37882 | 100965 |
| 32.08837 | 20 | 33.37938 | 23 | 17.00348 | 24 | 91.06599 | 53 | PHB2 | K562 | None | GSM28285 | 2971 | 100967 |
| 456.6543 | 117 | 589.3935 | 148 | 355.3685 | 147 | 620.5794 | 207 | EP300 | None | Sigmoid Cc | GSM28285 | 7063 | 100968 |
| 928.9785 | 121 | 1020.081 | 139 | 437.0203 | 106 | 421.411 | 110 | ZHX1 | K562 | None | GSM24236 | 2905 | 100985 |
| 867.6698 | 124 | 858.2902 | 134 | 411.2912 | 111 | 290.3984 | 98 | ZHX1 | K562 | None | GSM24236 | 3460 | 100986 |
| 759.1658 | 169 | 761.7232 | 186 | 367.9452 | 165 | 419.8357 | 188 | RAD51 | GM12878 | Blood | GSM28275 | 8350 | 100989 |
| 786.9324 | 197 | 704.37 | 205 | 293.0889 | 175 | 340.3681 | 201 | RAD51 | GM12878 | Blood | GSM28275 | 11427 | 100990 |
| 369.8837 | 162 | 315.1676 | 168 | 181.1 | 178 | 282.452 | 230 | ZNF24 | K562 | None | GSM28275 | 16371 | 100991 |
| 288.4616 | 133 | 290.0354 | 149 | 147.7988 | 150 | 222.4207 | 191 | ZNF24 | K562 | None | GSM28275 | 14186 | 100992 |
| 117.9477 | 174 | 178.3188 | 231 | 104.2643 | 279 | 133.999 | 329 | FOXH1 | H9 | Embryo | GSM72758 | 49674 | 1010 |
| 607.0468 | 277 | 701.2335 | 332 | 307.5008 | 315 | 309.8325 | 343 | IKZF2 | GM12878 | Blood | GSM28274 | 31023 | 101001 |
| 124.2086 | 32 | 92.24336 | 30 | 33.29392 | 24 | 235.8202 | 68 | IKZF2 | GM12878 | None | GSM28274 | 1906 | 101002 |
| 950.9266 | 525 | 991.9837 | 684 | 699.0705 | 553 | 680.1755 | 602 | CTCF | None | tibial nerve | GSM28274 | 32904 | 101003 |
| 852.4997 | 173 | 979.2191 | 217 | 612.8763 | 203 | 783.6111 | 245 | EP300 | None | Stomach | GSM28274 | 7786 | 101004 |
| 33.92891 | 12 | 23.43621 | 11 | 24.18752 | 15 | 236.3723 | 53 | ZNF514 | HEK293 | Embryonic | GSM28289 | 1064 | 101007 |
| 28.18986 | 13 | 22.37956 | 13 | 43.14215 | 24 | 245.4242 | 63 | ZNF514 | HEK293 | Embryonic | GSM28289 | 1491 | 101008 |
| 727.5841 | 336 | 763.6823 | 445 | 500.8367 | 417 | 568.8336 | 489 | CTCF | None | Adrenal Gl | GSM28289 | 37145 | 101009 |
| 111.062 | 42 | 50.73168 | 32 | 27.78473 | 34 | 76.25497 | 56 | ZNF184 | K562 | None | GSM28274 | 3807 | 101010 |
| 134.945 | 65 | 131.9919 | 72 | 44.03624 | 62 | 109.2266 | 96 | ZNF184 | K562 | None | GSM28274 | 7329 | 101011 |
| 639.2841 | 100 | 507.9448 | 95 | 299.6612 | 89 | 525.8778 | 128 | KHSRP | K562 | None | GSM28274 | 2870 | 101012 |
| 771.8947 | 109 | 663.4676 | 108 | 438.7463 | 106 | 556.4968 | 128 | KHSRP | K562 | None | GSM28274 | 2686 | 101013 |
| 892.6574 | 392 | 898.6096 | 483 | 701.692 | 473 | 707.8347 | 549 | RXRB | HepG2 | None | GSM25341 | 29105 | 101015 |
| 914.4861 | 600 | 913.4418 | 720 | 673.6583 | 680 | 676.8286 | 782 | RXRB | HepG2 | None | GSM25341 | 50685 | 101016 |
| 496.652 | 75 | 641.3086 | 95 | 273.7777 | 72 | 660.416 | 125 | ZBTB7B | MCF-7 | Breast | GSM28258 | 1978 | 101024 |
| 611.7685 | 107 | 746.4085 | 131 | 362.1623 | 110 | 436.5941 | 129 | ZBTB7B | MCF-7 | Breast | GSM28258 | 3653 | 101025 |
| 963.5426 | 281 | 951.4951 | 333 | 672.3661 | 284 | 652.9649 | 300 | ZNF511 | HepG2 | None | GSM25339 | 14628 | 101031 |
| 1006.038 | 318 | 1028.656 | 414 | 751.9951 | 318 | 748.6158 | 361 | ZNF511 | HepG2 | None | GSM25339 | 15350 | 101032 |
| 449.0056 | 62 | 572.686 | 78 | 250.3548 | 59 | 485.1561 | 91 | NFRKB | HepG2 | None | GSM28254 | 1425 | 101033 |
| 314.0015 | 48 | 343.0454 | 55 | 193.1366 | 49 | 432.7453 | 82 | NFRKB | HepG2 | None | GSM28254 | 1305 | 101034 |
| 571.3759 | 67 | 642.0068 | 78 | 204.4274 | 49 | 347.0171 | 70 | MAZ | A549 | Lung | GSM24237 | 1340 | 101035 |
| 1231.334 | 166 | 1217.874 | 192 | 493.7761 | 121 | 657.7174 | 150 | MAZ | A549 | Lung | GSM24237 | 3379 | 101036 |
| 1057.72 | 502 | 1069.557 | 627 | 776.2447 | 501 | 735.1603 | 512 | CREM | GM12878 | None | GSM25345 | 24635 | 101037 |
| 942.6404 | 585 | 955.7778 | 725 | 682.6399 | 637 | 635.1862 | 645 | CREM | GM12878 | None | GSM25345 | 45577 | 101038 |
| 134.501 | 131 | 116.6637 | 141 | 118.1372 | 196 | 225.3133 | 268 | ARID1B | K562 | None | GSM24240 | 24138 | 101045 |
| 455.9167 | 262 | 503.0387 | 307 | 323.2344 | 347 | 419.4549 | 415 | ARID1B | K562 | None | GSM24240 | 35013 | 101046 |
| 9.623404 | 8 | 17.3035 | 12 | 3.747475 | 9 | 167.178 | 53 | ZNF26 | HEK293 | Embryonic | GSM28251 | 1610 | 101059 |
| 13.41573 | 12 | 9.798477 | 12 | 11.11333 | 18 | 58.61957 | 39 | ZNF26 | HEK293 | Embryonic | GSM28251 | 2442 | 101060 |
| 1002.239 | 520 | 1000.004 | 620 | 744.1088 | 542 | 733.9242 | 602 | ZNF580 | HepG2 | None | GSM25275 | 29248 | 101063 |

| | | | | | | | | | | | | | |
|---|---|---|---|---|---|---|---|---|---|---|---|---|---|
| 1050.759 | 264 | 1049.829 | 324 | 803.8206 | 264 | 650.0436 | 252 | ZNF580 | HepG2 | None | GSM25275 | 10434 | 101064 |
| 23.1116 | 13 | 12.50774 | 11 | 31.95795 | 23 | 147.5087 | 52 | ZNF311 | HEK293 | Embryonic | GSM28251 | 1731 | 101073 |
| 32.95889 | 14 | 13.21375 | 10 | 23.92688 | 18 | 157.2217 | 49 | ZNF311 | HEK293 | Embryonic | GSM28251 | 1440 | 101074 |
| 1065.834 | 371 | 1086.547 | 464 | 818.0914 | 359 | 781.2598 | 371 | ZFX | HCT-116 | HCT116 | GSM28272 | 11582 | 101076 |
| 931.1893 | 455 | 976.2102 | 605 | 695.5539 | 494 | 695.4917 | 561 | CTCF | None | Lung | GSM28251 | 30015 | 101077 |
| 896.7906 | 546 | 878.6582 | 640 | 669.8871 | 629 | 646.2256 | 678 | CREB1 | MCF-7 | Breast | GSM28272 | 45643 | 101078 |
| 1004.658 | 487 | 993.4626 | 572 | 785.3537 | 553 | 771.6822 | 606 | CREB1 | MCF-7 | Breast | GSM28272 | 25553 | 101079 |
| 845.6817 | 208 | 925.5579 | 271 | 491.1357 | 226 | 567.3315 | 259 | FOXM1 | K562 | None | GSM28272 | 11906 | 101081 |
| 1097.419 | 322 | 1112.374 | 403 | 858.0155 | 314 | 837.1046 | 337 | ZFX | C4-2B | Prostate | GSM28251 | 8700 | 101082 |
| 1180.085 | 171 | 1188.304 | 218 | 607.638 | 147 | 682.4206 | 166 | ZFX | C4-2B | Prostate | GSM28251 | 3561 | 101083 |
| 1036.985 | 224 | 1037.563 | 276 | 705.2221 | 224 | 700.1547 | 238 | SKIL | K562 | None | GSM28272 | 7345 | 101084 |
| 285.6079 | 41 | 345.6518 | 50 | 136.8862 | 36 | 176.7801 | 44 | BHLHE40 | HEK293T | Embryonic | GSM24239 | 1010 | 101091 |
| 112.895 | 58 | 62.69882 | 50 | 110.4579 | 87 | 113.8423 | 95 | NCOR1 | K562 | None | GSM24239 | 7115 | 101094 |
| 336.9849 | 149 | 348.2411 | 169 | 299.4548 | 210 | 314.0964 | 231 | NCOR1 | K562 | None | GSM24239 | 15637 | 101095 |
| 403.015 | 172 | 587.0811 | 230 | 259.3754 | 215 | 326.8891 | 255 | FOXH1 | H9 | Embryo | GSM72756 | 19597 | 1011 |
| 925.33 | 260 | 915.4754 | 313 | 735.8689 | 310 | 677.5591 | 313 | MLX | HepG2 | None | GSM25274 | 15589 | 101106 |
| 855.949 | 190 | 940.9079 | 233 | 532.7761 | 212 | 445.2499 | 208 | MLX | HepG2 | None | GSM25274 | 10228 | 101107 |
| 906.8018 | 355 | 909.8763 | 433 | 681.2693 | 387 | 655.3722 | 409 | HOMEZ | HepG2 | None | GSM25274 | 24376 | 101112 |
| 884.088 | 458 | 877.5908 | 553 | 616.3898 | 467 | 642.7912 | 568 | HOMEZ | HepG2 | None | GSM25274 | 37912 | 101113 |
| 814.0107 | 446 | 831.9237 | 563 | 617.9145 | 528 | 599.3551 | 574 | CTCF | None | Pancreas | GSM28271 | 40713 | 101134 |
| 29.34684 | 13 | 35.29642 | 16 | 27.60191 | 19 | 232.0526 | 60 | ZNF416 | HEK293 | Embryonic | GSM28271 | 1432 | 101137 |
| 319.2103 | 77 | 476.5363 | 106 | 240.0211 | 95 | 210.4214 | 95 | ETV6 | K562 | None | GSM24230 | 4502 | 101140 |
| 807.0342 | 220 | 899.5653 | 296 | 718.3746 | 299 | 759.437 | 373 | EP300 | None | Lung | GSM28286 | 15290 | 101151 |
| 452.223 | 61 | 472.2728 | 68 | 162.495 | 46 | 343.9284 | 74 | USF2 | SK-N-SH | Brain | GSM28286 | 1526 | 101154 |
| 713.0785 | 111 | 879.0098 | 137 | 438.211 | 116 | 472.099 | 128 | USF2 | SK-N-SH | Bone Marr | GSM28286 | 3547 | 101155 |
| 955.4283 | 569 | 980.7319 | 721 | 706.2074 | 623 | 699.8715 | 702 | CTCF | None | Prostate | GSM28286 | 39815 | 101156 |
| 394.1085 | 75 | 471.2313 | 91 | 191.3534 | 71 | 183.0981 | 74 | PTBP1 | K562 | None | GSM28286 | 3068 | 101157 |
| 332.9759 | 62 | 451.2214 | 81 | 181.5528 | 62 | 108.767 | 51 | PTBP1 | K562 | None | GSM28286 | 2459 | 101158 |
| 761.5323 | 165 | 969.0832 | 214 | 601.4504 | 204 | 828.0774 | 262 | CTBP1 | HEK293T | Embryonic | GSM24232 | 7894 | 101159 |
| 261.8324 | 64 | 481.6184 | 99 | 358.1513 | 106 | 429.5688 | 124 | CTBP1 | HEK293T | Embryonic | GSM24232 | 3394 | 101160 |
| 1198.143 | 196 | 1189.311 | 235 | 831.1595 | 183 | 928.9321 | 206 | GTF2F1 | MCF-7 | Mammary | GSM24236 | 4086 | 101171 |
| 1174.02 | 193 | 1159.242 | 228 | 772.2773 | 179 | 866.4574 | 202 | GTF2F1 | MCF-7 | Mammary | GSM24236 | 4154 | 101172 |
| 297.9149 | 43 | 333.844 | 50 | 130.8473 | 36 | 123.3723 | 37 | SUZ12 | HEK293T | Embryonic | GSM24229 | 1063 | 101177 |
| 57.62761 | 25 | 80.23005 | 33 | 78.12105 | 43 | 144.3304 | 62 | NFXL1 | K562 | None | GSM24229 | 2604 | 101179 |
| 75.64795 | 25 | 52.7286 | 23 | 47.31827 | 29 | 125.5412 | 50 | NFXL1 | K562 | None | GSM24229 | 1948 | 101180 |
| 137.8895 | 27 | 141.3204 | 30 | 140.7484 | 38 | 355.0553 | 68 | SMARCC2 | HepG2 | None | GSM28289 | 1149 | 101186 |
| 77.85118 | 19 | 59.55688 | 18 | 78.53761 | 27 | 323.2791 | 63 | ZNF624 | HEK293 | Embryonic | GSM28289 | 1048 | 101187 |
| 103.7111 | 22 | 81.45378 | 21 | 60.49962 | 23 | 298.3553 | 59 | ZNF624 | HEK293 | Embryonic | GSM28289 | 1010 | 101188 |
| 1174.193 | 584 | 1233.117 | 778 | 890.0697 | 574 | 830.9885 | 571 | RBFOX2 | HepG2 | None | GSM28289 | 18284 | 101189 |
| 1194.003 | 618 | 1233.36 | 786 | 889.0121 | 588 | 838.5672 | 594 | RBFOX2 | HepG2 | None | GSM28289 | 18807 | 101190 |
| 245.1556 | 64 | 427.3736 | 96 | 159.3606 | 74 | 485.413 | 139 | EP300 | None | Lung | GSM28272 | 4027 | 101191 |
| 984.8063 | 227 | 971.7641 | 268 | 530.6787 | 215 | 744.5601 | 271 | ZBTB48 | HEK293 | Embryonic | GSM28289 | 9711 | 101192 |
| 961.1168 | 296 | 979.1773 | 378 | 743.8326 | 314 | 762.3572 | 377 | ZBTB48 | HEK293 | Embryonic | GSM28289 | 13829 | 101193 |
| 280.9554 | 57 | 278.0716 | 62 | 103.8564 | 47 | 258.8763 | 80 | ZBTB5 | K562 | None | GSM28289 | 2354 | 101194 |
| 264.3501 | 53 | 264.6963 | 58 | 109.3703 | 46 | 384.3518 | 95 | ZBTB5 | K562 | None | GSM28289 | 2129 | 101195 |
| 1101.083 | 373 | 1094.157 | 451 | 814.3572 | 339 | 799.919 | 369 | RB1 | GM12878 | Blood | GSM28289 | 11928 | 101196 |
| 453.762 | 64 | 730.5797 | 93 | 252.4965 | 61 | 489.8094 | 94 | RB1 | GM12878 | None | GSM28289 | 1511 | 101197 |
| 840.3369 | 149 | 1061.844 | 200 | 645.1822 | 177 | 704.975 | 197 | NONO | K562 | None | GSM28289 | 5359 | 101198 |
| 1056.241 | 202 | 1085.058 | 267 | 702.5022 | 208 | 835.149 | 242 | NONO | K562 | None | GSM28289 | 6955 | 101199 |
| 952.3479 | 604 | 973.9581 | 756 | 677.7457 | 637 | 625.2093 | 643 | FOXP1 | H9 | Embryo | GSM76831 | 47374 | 1012 |
| 1142.174 | 266 | 1173.84 | 353 | 914.4053 | 279 | 892.7556 | 296 | NR2C1 | GM12878 | Blood | GSM28289 | 7276 | 101200 |
| 1131.828 | 305 | 1154.066 | 396 | 877.3318 | 297 | 869.8088 | 330 | NR2C1 | GM12878 | Blood | GSM28289 | 8964 | 101201 |
| 1113.146 | 371 | 1112.865 | 447 | 888.8617 | 402 | 863.6589 | 423 | ZNF217 | MCF-7 | Breast | GSM28274 | 11899 | 101202 |
| 1063.741 | 312 | 1051.564 | 367 | 874.1659 | 358 | 833.8846 | 362 | ZNF217 | MCF-7 | Breast | GSM28274 | 10803 | 101203 |
| 872.6673 | 224 | 918.3601 | 298 | 463.3077 | 234 | 657.5214 | 295 | ZNF398 | HEK293 | Embryonic | GSM28274 | 13002 | 101206 |
| 913.954 | 274 | 949.4461 | 365 | 631.9634 | 290 | 718.4034 | 348 | ZNF398 | HEK293 | Embryonic | GSM28274 | 14865 | 101207 |
| 63.16502 | 20 | 57.04877 | 21 | 63.19663 | 29 | 244.7933 | 63 | ZNF23 | HEK293 | Embryonic | GSM28274 | 1502 | 101209 |
| 78.06503 | 27 | 50.42398 | 24 | 34.49449 | 27 | 84.87119 | 44 | ZNF266 | HEK293 | Embryonic | GSM28274 | 2232 | 101213 |
| 21.98753 | 13 | 14.23305 | 12 | 19.1738 | 19 | 96.0348 | 43 | ZNF266 | HEK293 | Embryonic | GSM28274 | 1897 | 101214 |
| 614.7906 | 147 | 832.7175 | 191 | 464.8442 | 182 | 391.8104 | 179 | ZKSCAN8 | HepG2 | None | GSM25344 | 8628 | 101224 |
| 911.9222 | 362 | 888.5614 | 418 | 682.9816 | 390 | 680.9847 | 441 | ZKSCAN8 | HepG2 | None | GSM25344 | 23896 | 101225 |
| 1119.349 | 197 | 1109.049 | 234 | 727.306 | 193 | 586.134 | 183 | RFXANK | HepG2 | None | GSM25344 | 5964 | 101226 |
| 1045.222 | 196 | 1068.916 | 257 | 643.5092 | 200 | 518.8843 | 191 | RFXANK | HepG2 | None | GSM25344 | 7274 | 101227 |
| 862.1408 | 529 | 887.9103 | 673 | 653.0128 | 622 | 653.5774 | 711 | CTCF | None | Pancreas | GSM28256 | 45254 | 101228 |
| 453.7471 | 292 | 663.0555 | 400 | 311.526 | 387 | 303.1134 | 418 | CTCF | VCaP | Prostate | GSM28256 | 42498 | 101229 |
| 528.251 | 295 | 704.0289 | 410 | 343.3613 | 376 | 342.8254 | 409 | CTCF | VCaP | Prostate | GSM28256 | 37689 | 101230 |
| 85.73483 | 21 | 98.42899 | 25 | 48.03084 | 22 | 252.2365 | 57 | THRA | K562 | None | GSM28256 | 1159 | 101231 |
| 95.35866 | 22 | 80.74676 | 22 | 29.69727 | 17 | 198.4307 | 49 | THRA | K562 | None | GSM28256 | 1112 | 101232 |
| 879.9138 | 310 | 890.8725 | 386 | 486.0853 | 291 | 527.6217 | 325 | ZFX | HEK293T | Embryonic | GSM28254 | 17214 | 101235 |
| 1044.361 | 306 | 1072.313 | 393 | 813.9071 | 300 | 801.2649 | 329 | ZFX | HEK293T | Embryonic | GSM28254 | 9145 | 101236 |
| 199.1507 | 59 | 272.0446 | 77 | 147.6786 | 73 | 286.2499 | 108 | GATAD2A | K562 | None | GSM28255 | 3951 | 101238 |
| 486.9865 | 170 | 557.5229 | 202 | 398.7843 | 227 | 341.0672 | 228 | RCOR2 | HepG2 | None | GSM25340 | 14765 | 101239 |

| | | | | | | | | | | | | | |
|---|---|---|---|---|---|---|---|---|---|---|---|---|---|
| 727.2229 | 242 | 686.7043 | 262 | 575.0391 | 319 | 491.906 | 321 | RCOR2 | HepG2 | None | GSM25340 | 21340 | 101240 |
| 523.3851 | 200 | 694.5028 | 255 | 281.4653 | 225 | 347.8398 | 265 | CTCF | None | Sigmoid Cc | GSM28255 | 18524 | 101243 |
| 1047.852 | 260 | 1073.342 | 337 | 830.7166 | 273 | 657.9587 | 248 | MXD4 | HepG2 | None | GSM25340 | 9140 | 101244 |
| 1125.129 | 541 | 1136.607 | 670 | 838.0488 | 540 | 796.3068 | 551 | MXD4 | HepG2 | None | GSM25340 | 21862 | 101245 |
| 1103.119 | 443 | 1117.614 | 550 | 804.0228 | 394 | 800.2367 | 443 | L3MBTL2 | HEK293T | Embryonic | GSM24240 | 14538 | 101246 |
| 1115.382 | 393 | 1142.048 | 505 | 831.9366 | 358 | 822.2464 | 396 | L3MBTL2 | HEK293T | Embryonic | GSM24240 | 11972 | 101247 |
| 870.3701 | 470 | 885.6092 | 589 | 649.6727 | 527 | 639.2069 | 584 | ZFHX2 | HEK293 | Embryonic | GSM28272 | 36686 | 101252 |
| 966.255 | 468 | 958.4403 | 560 | 707.8619 | 478 | 690.3947 | 522 | ZFHX2 | HEK293 | Embryonic | GSM28272 | 27356 | 101253 |
| 456.6092 | 101 | 486.5375 | 115 | 208.6952 | 96 | 359.5504 | 134 | CTCF | None | omental fa | GSM28251 | 5183 | 101260 |
| 1128.81 | 232 | 1112.773 | 275 | 800.7002 | 217 | 790.4148 | 229 | TARDBP | K562 | None | GSM28272 | 6697 | 101262 |
| 1126.942 | 234 | 1143.016 | 296 | 909.9881 | 249 | 832.6114 | 236 | TARDBP | K562 | None | GSM28272 | 6737 | 101263 |
| 283.6379 | 77 | 456.461 | 110 | 254.8031 | 104 | 511.9978 | 158 | ZNF24 | MCF-7 | Breast | GSM28272 | 4796 | 101264 |
| 633.3951 | 166 | 827.9748 | 211 | 506.9029 | 212 | 679.1092 | 261 | ZNF24 | MCF-7 | Breast | GSM28272 | 10352 | 101265 |
| 78.41075 | 21 | 34.83129 | 15 | 38.24731 | 21 | 214.6989 | 55 | FOXA1 | K562 | None | GSM24240 | 1348 | 101271 |
| 277.3417 | 51 | 331.8378 | 62 | 156.4554 | 52 | 165.8051 | 57 | MYB | GM12878 | Blood | GSM24240 | 2055 | 101272 |
| 422.1528 | 83 | 515.5823 | 102 | 308.4479 | 98 | 211.5378 | 86 | MYB | GM12878 | Blood | GSM24240 | 3644 | 101273 |
| 406.8612 | 51 | 606.3452 | 71 | 170.391 | 41 | 327.2735 | 63 | SREBF1 | K562 | None | GSM24240 | 1098 | 101274 |
| 908.3755 | 127 | 1176.711 | 173 | 546.3482 | 129 | 513.5298 | 132 | SREBF1 | K562 | None | GSM24240 | 3414 | 101275 |
| 909.3652 | 684 | 919.3923 | 837 | 625.8733 | 737 | 589.8018 | 780 | FOXP1 | H9 | Embryo | GSM76831 | 68906 | 1013 |
| 681.8342 | 286 | 779.5613 | 396 | 498.8608 | 376 | 547.845 | 422 | SMC3 | A549 | Lung | GSM31064 | 32568 | 101305 |
| 594.8821 | 257 | 767.2593 | 359 | 391.8256 | 324 | 434.2642 | 365 | SMC3 | A549 | Lung | GSM31064 | 29271 | 101307 |
| 854.8445 | 438 | 853.7917 | 532 | 663.201 | 524 | 664.9942 | 604 | ZIC2 | HEK293 | Embryonic | GSM28285 | 35038 | 101318 |
| 878.729 | 503 | 874.62 | 601 | 681.1597 | 603 | 675.5149 | 680 | ZIC2 | HEK293 | Embryonic | GSM28285 | 39775 | 101319 |
| 993.9327 | 296 | 986.7419 | 353 | 772.9441 | 309 | 748.7065 | 328 | GATAD2B | GM12878 | Blood | GSM28285 | 11992 | 101321 |
| 1076.621 | 396 | 1066.178 | 468 | 812.1771 | 377 | 805.766 | 422 | GATAD2B | GM12878 | Blood | GSM28285 | 13598 | 101322 |
| 587.9475 | 78 | 737.333 | 97 | 330.204 | 74 | 298.5119 | 74 | STAT1 | GM12878 | None | GSM24233 | 1650 | 101324 |
| 421.9525 | 213 | 591.6262 | 278 | 286.9796 | 276 | 370.6722 | 330 | RAD21 | A549 | Lung | GSM31064 | 26976 | 101326 |
| 728.858 | 357 | 751.3733 | 460 | 583.9661 | 482 | 578.4122 | 540 | RAD21 | A549 | Lung | GSM31064 | 41618 | 101327 |
| 573.98 | 280 | 725.7467 | 377 | 459.2118 | 387 | 466.8989 | 422 | RAD21 | A549 | Lung | GSM31064 | 36064 | 101328 |
| 1094.453 | 403 | 1121.562 | 518 | 820.0101 | 379 | 806.5121 | 420 | NRF1 | K562 | None | GSM24235 | 13442 | 101329 |
| 1070.979 | 516 | 1094.513 | 648 | 810.0034 | 524 | 761.7875 | 531 | NRF1 | K562 | None | GSM24235 | 20973 | 101330 |
| 305.6379 | 150 | 464.6152 | 204 | 164.2746 | 176 | 268.4763 | 231 | RAD21 | A549 | Lung | GSM31064 | 18036 | 101331 |
| 717.1664 | 341 | 729.2692 | 429 | 486.3023 | 430 | 556.0462 | 504 | RAD21 | A549 | Lung | GSM31064 | 42326 | 101332 |
| 667.1894 | 304 | 737.2977 | 398 | 514.8403 | 410 | 560.5908 | 459 | RAD21 | A549 | Lung | GSM31064 | 36281 | 101333 |
| 1062.184 | 335 | 1052.619 | 393 | 815.5289 | 328 | 790.6194 | 350 | CREB3L1 | K562 | None | GSM24230 | 11175 | 101334 |
| 1155.128 | 217 | 1143.965 | 258 | 825.0245 | 202 | 730.9164 | 201 | CREB3L1 | K562 | None | GSM24230 | 5010 | 101335 |
| 254.4013 | 51 | 263.1328 | 57 | 135.1303 | 51 | 191.3188 | 65 | IRF2 | K562 | None | GSM24233 | 2274 | 101336 |
| 981.3301 | 265 | 1009.114 | 350 | 776.5905 | 299 | 731.6339 | 299 | IRF2 | K562 | None | GSM24233 | 13271 | 101337 |
| 935.6434 | 478 | 949.582 | 591 | 676.1742 | 476 | 653.0151 | 512 | IKZF1 | GM12878 | Blood | GSM28273 | 27396 | 101338 |
| 923.4193 | 536 | 907.4959 | 626 | 663.755 | 548 | 634.126 | 581 | IKZF1 | GM12878 | Blood | GSM28273 | 34690 | 101339 |
| 35.2261 | 20 | 67.59376 | 31 | 18.44724 | 24 | 55.0628 | 41 | KAT2B | HepG2 | None | GSM24237 | 3049 | 101341 |
| 616.1525 | 106 | 654.7941 | 120 | 280.0239 | 96 | 346.5075 | 114 | USF2 | IMR90 | Lung | GSM28273 | 3920 | 101344 |
| 989.7128 | 480 | 1003.018 | 605 | 719.7636 | 494 | 679.2554 | 506 | USF2 | IMR90 | Embryo | GSM28273 | 29764 | 101345 |
| 1183.515 | 311 | 1209.74 | 396 | 921.4256 | 290 | 906.0264 | 315 | SIN3B | K562 | None | GSM28273 | 7226 | 101346 |
| 1238.872 | 234 | 1226.203 | 272 | 954.8838 | 203 | 971.2798 | 238 | SIN3B | K562 | None | GSM28273 | 4412 | 101347 |
| 281.7579 | 171 | 266.0588 | 188 | 342.9281 | 282 | 517.1288 | 360 | PRDM6 | HEK293 | Embryonic | GSM28289 | 24364 | 101351 |
| 392.3314 | 213 | 410.0788 | 244 | 395.6434 | 323 | 601.897 | 418 | PRDM6 | HEK293 | Embryonic | GSM28289 | 28028 | 101352 |
| 775.1234 | 316 | 751.335 | 361 | 476.494 | 361 | 613.5161 | 455 | NFIC | K562 | None | GSM28289 | 29636 | 101353 |
| 742.349 | 328 | 723.1215 | 377 | 471.0404 | 390 | 547.8655 | 448 | NFIC | K562 | None | GSM28289 | 34612 | 101354 |
| 819.6459 | 355 | 801.3942 | 416 | 640.0486 | 430 | 602.0939 | 445 | SOX6 | K562 | None | GSM28289 | 29528 | 101355 |
| 532.0307 | 198 | 571.3167 | 228 | 405.9806 | 257 | 489.6168 | 300 | SOX6 | K562 | None | GSM28289 | 17859 | 101356 |
| 83.26723 | 22 | 48.17753 | 18 | 21.53456 | 16 | 142.7204 | 44 | ZNF354B | K562 | None | GSM28274 | 1254 | 101358 |
| 60.79459 | 17 | 35.43201 | 14 | 14.43296 | 12 | 139.187 | 40 | ZNF354B | K562 | None | GSM28274 | 1049 | 101359 |
| 1170.086 | 286 | 1202.763 | 371 | 882.6282 | 250 | 882.8467 | 282 | SRSF1 | HepG2 | None | GSM28274 | 7310 | 101361 |
| 28.6476 | 12 | 15.94628 | 10 | 35.83951 | 20 | 242.1063 | 58 | ZNF195 | HEK293 | Embryonic | GSM28274 | 1256 | 101366 |
| 54.73138 | 20 | 32.08729 | 17 | 20.77168 | 19 | 205.6257 | 62 | ZNF560 | HEK293 | Embryonic | GSM28274 | 1846 | 101367 |
| 941.9768 | 646 | 968.8801 | 813 | 706.0773 | 758 | 711.2396 | 876 | NR2F6 | HepG2 | None | GSM25344 | 54801 | 101378 |
| 369.879 | 130 | 540.0344 | 175 | 333.8298 | 183 | 554.721 | 248 | NR2F6 | HepG2 | None | GSM25344 | 11899 | 101379 |
| 452.084 | 102 | 605.8298 | 132 | 337.3704 | 124 | 724.3691 | 196 | EP300 | None | Lung | GSM28259 | 5388 | 101384 |
| 169.4182 | 39 | 123.9642 | 36 | 55.27067 | 31 | 322.7755 | 82 | ZNF777 | HEK293 | Embryonic | GSM28259 | 1908 | 101386 |
| 612.5798 | 116 | 832.4699 | 151 | 349.6597 | 118 | 524.3384 | 155 | ZNF777 | HEK293 | Embryonic | GSM28259 | 4284 | 101387 |
| 882.8374 | 239 | 900.5157 | 306 | 528.7927 | 263 | 591.9665 | 297 | ZNF48 | HepG2 | None | GSM25340 | 15653 | 101390 |
| 1030.04 | 652 | 1044.16 | 801 | 753.5079 | 687 | 731.1255 | 747 | ZNF48 | HepG2 | None | GSM25340 | 38215 | 101391 |
| 28.4023 | 14 | 16.16472 | 12 | 47.33923 | 27 | 245.9479 | 67 | ZNF544 | HEK293 | Embryonic | GSM28255 | 1716 | 101392 |
| 58.4192 | 21 | 10.00663 | 10 | 40.68989 | 26 | 260.046 | 71 | ZNF544 | HEK293 | Embryonic | GSM28255 | 1836 | 101393 |
| 0.044107 | 9 | 0.41522 | 14 | 0.003857 | 17 | 16.90316 | 55 | ZNF426 | HEK293 | Embryonic | GSM28255 | 8634 | 101394 |
| 0.162618 | 9 | 0.130347 | 11 | 0.053718 | 17 | 3.548909 | 34 | ZNF426 | HEK293 | Embryonic | GSM28255 | 7783 | 101395 |
| 730.603 | 230 | 808.1257 | 277 | 515.1548 | 285 | 639.1039 | 337 | THRB | HepG2 | None | GSM25340 | 18488 | 101400 |
| 292.6949 | 87 | 408.7445 | 115 | 276.4308 | 123 | 343.5535 | 146 | THRB | HepG2 | None | GSM25340 | 6546 | 101401 |
| 687.5587 | 362 | 713.4057 | 471 | 554.4814 | 514 | 562.4527 | 598 | CEBPA | HepG2 | None | GSM25275 | 53284 | 101421 |
| 690.35 | 510 | 715.4861 | 654 | 534.2549 | 695 | 530.8704 | 788 | CEBPA | HepG2 | None | GSM25275 | 84180 | 101422 |
| 869.8806 | 342 | 862.978 | 412 | 653.0834 | 383 | 647.821 | 429 | PPARG | HepG2 | None | GSM25275 | 26092 | 101423 |

| | | | | | | | | | | | | | |
|---|---|---|---|---|---|---|---|---|---|---|---|---|---|
| 845.2411 | 319 | 871.0053 | 417 | 674.6034 | 401 | 658.2094 | 439 | PPARG | HepG2 | None | GSM25275 | 25788 | 101424 |
| 899.2067 | 366 | 904.7057 | 453 | 660.7182 | 376 | 651.0235 | 419 | NR2F1 | GM12878 | Blood | GSM28273 | 22913 | 101425 |
| 905.3083 | 366 | 904.3081 | 439 | 668.4939 | 378 | 655.5072 | 416 | NR2F1 | GM12878 | Blood | GSM28273 | 22555 | 101426 |
| 789.9793 | 189 | 870.8891 | 219 | 635.6891 | 238 | 686.4622 | 264 | ZBTB21 | HEK293 | Embryonic | GSM28272 | 10095 | 101427 |
| 772.1373 | 176 | 776.3608 | 194 | 612.5049 | 219 | 684.7851 | 247 | ZBTB21 | HEK293 | Embryonic | GSM28272 | 8881 | 101428 |
| 1080.202 | 182 | 1091.757 | 234 | 778.4651 | 201 | 817.3633 | 219 | ZNF524 | HEK293 | Embryonic | GSM28272 | 5579 | 101429 |
| 570.2723 | 107 | 743.2724 | 136 | 351.7917 | 113 | 408.8242 | 130 | ZNF524 | HEK293 | Embryonic | GSM28272 | 3873 | 101430 |
| 1152.848 | 537 | 1183.034 | 682 | 889.6207 | 553 | 865.5915 | 595 | ELF1 | MCF-7 | Breast | GSM28272 | 18237 | 101433 |
| 1190.949 | 256 | 1212.251 | 329 | 968.6031 | 267 | 941.0172 | 282 | ELF1 | MCF-7 | Breast | GSM28272 | 5868 | 101434 |
| 648.554 | 91 | 843.034 | 116 | 199.4994 | 64 | 252.3595 | 77 | SUZ12 | HepG2 | None | GSM24239 | 2420 | 101437 |
| 338.8049 | 50 | 645.2882 | 81 | 176.5027 | 47 | 254.5213 | 61 | SUZ12 | HepG2 | None | GSM24239 | 1445 | 101438 |
| 951.9319 | 257 | 954.2248 | 319 | 728.5621 | 286 | 617.1789 | 283 | ZBTB26 | HepG2 | None | GSM25275 | 13637 | 101439 |
| 989.1674 | 464 | 1000.185 | 572 | 724.6681 | 470 | 709.5531 | 515 | ZBTB26 | HepG2 | None | GSM25275 | 25799 | 101440 |
| 123.2117 | 36 | 189 | 50 | 85.6026 | 42 | 40.68352 | 32 | EZH2 | None | Embryo | GSM34983 | 1991 | 101447 |
| 138.5751 | 41 | 186.5395 | 53 | 48.47545 | 35 | 71.65472 | 45 | EZH2 | None | Embryo | GSM34983 | 2389 | 101448 |
| 932.7585 | 253 | 936.7053 | 308 | 550.6105 | 251 | 646.6441 | 290 | ESRRA | K562 | None | GSM24235 | 12622 | 101470 |
| 940.726 | 461 | 941.4618 | 562 | 703.2576 | 497 | 670.673 | 522 | ESRRA | K562 | None | GSM24235 | 29322 | 101471 |
| 87.52872 | 69 | 150.7221 | 99 | 75.00683 | 104 | 101.0614 | 126 | NFE2L2 | HepG2 | None | GSM24235 | 13169 | 101472 |
| 19.22805 | 24 | 30.11338 | 33 | 17.57357 | 39 | 68.36099 | 70 | NFE2L2 | HepG2 | None | GSM24235 | 6432 | 101473 |
| 304.259 | 145 | 374.303 | 179 | 148.8158 | 165 | 172.3521 | 189 | ZNF384 | HepG2 | None | GSM24230 | 18028 | 101474 |
| 600.1645 | 226 | 702.9749 | 272 | 253.6364 | 235 | 274.057 | 262 | ZNF384 | HepG2 | None | GSM24230 | 23232 | 101475 |
| 141.843 | 75 | 179.7502 | 94 | 90.44612 | 95 | 93.1159 | 104 | KDM1A | hESC | Embryo | GSM26999 | 9658 | 101478 |
| 304.2866 | 109 | 377.7209 | 135 | 179.797 | 126 | 128.6413 | 118 | KDM1A | hESC | Embryo | GSM26999 | 9454 | 101479 |
| 411.4667 | 239 | 572.3881 | 310 | 211.054 | 286 | 214.2816 | 313 | KDM1A | hESC | Embryo | GSM26999 | 34871 | 101480 |
| 537.8875 | 83 | 546.9577 | 91 | 233.0932 | 70 | 292.7322 | 84 | ZZZ3 | K562 | None | GSM24239 | 2195 | 101483 |
| 526.9829 | 79 | 366.0287 | 69 | 215.6691 | 64 | 326.1613 | 85 | ZZZ3 | K562 | None | GSM24239 | 1943 | 101484 |
| 429.9292 | 189 | 629.7722 | 252 | 249.1429 | 222 | 269.9027 | 248 | CTCF | LNCaP | Prostate | GSM28274 | 19009 | 101489 |
| 499.0386 | 286 | 693.1618 | 396 | 356.4327 | 380 | 362.8639 | 416 | CTCF | LNCaP | Prostate | GSM28274 | 38412 | 101490 |
| 1074.974 | 482 | 1087.608 | 595 | 807.2624 | 474 | 779.2426 | 498 | ZBTB20 | HEK293 | Embryonic | GSM28274 | 17342 | 101491 |
| 956.5841 | 316 | 962.1697 | 391 | 747.0573 | 348 | 748.1094 | 398 | ZBTB20 | HEK293 | Embryonic | GSM28274 | 15537 | 101492 |
| 1096.122 | 238 | 1102.423 | 295 | 870.9118 | 246 | 892.966 | 297 | KLF16 | HEK293 | Embryonic | GSM28274 | 6952 | 101493 |
| 1099.78 | 247 | 1094.045 | 298 | 878.0481 | 260 | 874.2097 | 292 | KLF16 | HEK293 | Embryonic | GSM28274 | 7291 | 101494 |
| 259.5201 | 74 | 209.0182 | 73 | 327.9504 | 119 | 450.2772 | 149 | BCL6B | HEK293 | Embryonic | GSM28274 | 4856 | 101495 |
| 283.5041 | 79 | 185.7698 | 70 | 338.3373 | 123 | 441.3196 | 150 | BCL6B | HEK293 | Embryonic | GSM28274 | 4994 | 101496 |
| 569.1688 | 92 | 567.4952 | 100 | 313.3289 | 90 | 297.631 | 93 | ASH2L | GM12878 | Blood | GSM27011 | 2849 | 101511 |
| 26.35977 | 24 | 23.31193 | 26 | 29.37576 | 40 | 118.0117 | 77 | ZNF133 | HEK293 | Embryonic | GSM28259 | 4728 | 101513 |
| 19.07104 | 25 | 10.29566 | 23 | 9.570143 | 33 | 106.3774 | 86 | ZNF133 | HEK293 | Embryonic | GSM28259 | 6194 | 101514 |
| 1.595829 | 11 | 10.50705 | 23 | 1.501371 | 21 | 26.18334 | 50 | ZC3H11A | A549 | Lung | GSM28258 | 6342 | 101523 |
| 1.30494 | 9 | 1.945366 | 12 | 0 | 10 | 44.06209 | 54 | ZC3H11A | A549 | Lung | GSM28258 | 5347 | 101524 |
| 607.6921 | 96 | 683.7197 | 112 | 221.0206 | 75 | 505.6177 | 124 | E2F7 | K562 | None | GSM28255 | 2834 | 101540 |
| 341.4618 | 62 | 406.9408 | 75 | 130.6262 | 51 | 474.1719 | 108 | E2F7 | K562 | None | GSM28255 | 2249 | 101541 |
| 438.2983 | 131 | 588.4048 | 169 | 218.2129 | 137 | 325.0054 | 176 | CTCF | None | omental fa | GSM28255 | 9580 | 101542 |
| 781.3212 | 156 | 908.2307 | 186 | 389.4154 | 152 | 485.9948 | 181 | ZNF592 | GM12878 | Blood | GSM28255 | 6900 | 101547 |
| 981.0157 | 185 | 985.8857 | 212 | 536.3995 | 187 | 567.0688 | 205 | ZNF592 | GM12878 | Blood | GSM28255 | 7515 | 101548 |
| 740.7624 | 386 | 763.6837 | 497 | 581.4278 | 509 | 571.7521 | 569 | CEBPG | HepG2 | None | GSM25342 | 48019 | 101549 |
| 802.485 | 468 | 827.697 | 603 | 609.6408 | 580 | 622.7037 | 685 | CEBPG | HepG2 | None | GSM25342 | 52020 | 101550 |
| 793.1835 | 415 | 834.7706 | 554 | 600.4388 | 500 | 609.5842 | 581 | CTCF | None | Peyer's pat | GSM28272 | 41650 | 101571 |
| 24.28895 | 11 | 10.05304 | 8 | 22.52701 | 16 | 206.7547 | 53 | ZNF571 | HEK293 | Embryonic | GSM28272 | 1236 | 101572 |
| 24.59643 | 13 | 8.632682 | 9 | 27.90795 | 21 | 214.9783 | 62 | ZNF571 | HEK293 | Embryonic | GSM28272 | 1669 | 101573 |
| 1358.537 | 195 | 1397.467 | 268 | 1078.227 | 173 | 1102.237 | 211 | DDX20 | MCF-7 | Breast | GSM28272 | 2628 | 101574 |
| 1516.23 | 155 | 1537.408 | 202 | 945.2062 | 120 | 1254.91 | 163 | DDX20 | MCF-7 | Breast | GSM28272 | 1240 | 101575 |
| 82.48451 | 20 | 51.8289 | 17 | 50.83434 | 22 | 330.3586 | 65 | ZNF300 | HEK293 | Embryonic | GSM28272 | 1108 | 101576 |
| 47.8828 | 18 | 20.07818 | 13 | 28.95081 | 21 | 274.8737 | 70 | ZNF300 | HEK293 | Embryonic | GSM28272 | 1654 | 101577 |
| 857.7254 | 387 | 877.129 | 495 | 665.9936 | 457 | 619.9451 | 463 | CTCF | None | gastrocnen | GSM28272 | 29077 | 101578 |
| 417.0718 | 77 | 458.0522 | 89 | 132.7189 | 59 | 183.6508 | 74 | BMI1 | K562 | None | GSM24239 | 3168 | 101583 |
| 604.0908 | 94 | 680.9582 | 110 | 224.8923 | 75 | 265.7748 | 87 | BMI1 | K562 | None | GSM24239 | 2968 | 101584 |
| 1033.617 | 206 | 1043.467 | 258 | 782.0509 | 231 | 578.2788 | 211 | ZBED1 | K562 | None | GSM25276 | 8214 | 101590 |
| 624.0338 | 119 | 752.3315 | 145 | 377.2185 | 127 | 365.2719 | 133 | ZBED1 | K562 | None | GSM25276 | 5043 | 101591 |
| 409.1528 | 243 | 691.5968 | 348 | 331.8561 | 343 | 328.1894 | 371 | CTCF | A549 | Lung | GSM24378 | 35567 | 101597 |
| 374.314 | 202 | 649.361 | 290 | 252.3403 | 262 | 286.3412 | 298 | CTCF | A549 | Lung | GSM24377 | 26406 | 101598 |
| 432.6689 | 252 | 692.6848 | 357 | 371.1843 | 363 | 360.9113 | 390 | CTCF | A549 | Lung | GSM24377 | 36708 | 101599 |
| 676.8117 | 156 | 863.551 | 196 | 451.6279 | 180 | 412.1128 | 184 | ZFP1 | HepG2 | None | GSM25341 | 8484 | 101650 |
| 926.5325 | 262 | 931.7914 | 329 | 712.7849 | 300 | 704.2275 | 321 | ZFP1 | HepG2 | None | GSM25341 | 15786 | 101651 |
| 780.2411 | 131 | 1030.553 | 168 | 518.3526 | 143 | 698.0652 | 178 | TBX3 | HepG2 | None | GSM24237 | 4347 | 101654 |
| 222.4924 | 48 | 271.8667 | 59 | 108.0837 | 46 | 380.7831 | 95 | TBX3 | HepG2 | None | GSM24237 | 2207 | 101655 |
| 1163.868 | 507 | 1182.264 | 629 | 885.1117 | 480 | 872.6232 | 534 | SIN3A | A549 | Lung | GSM24235 | 14089 | 101660 |
| 1136.849 | 490 | 1148.131 | 603 | 853.6538 | 461 | 829.703 | 494 | SIN3A | A549 | Lung | GSM24235 | 14449 | 101661 |
| 1253.247 | 347 | 1282.542 | 453 | 960.5375 | 311 | 930.5985 | 327 | HCFC1 | GM12878 | None | GSM24235 | 7409 | 101662 |
| 1229.574 | 458 | 1246.777 | 567 | 910.3546 | 398 | 875.9769 | 415 | HCFC1 | GM12878 | None | GSM24235 | 11683 | 101663 |
| 1108.351 | 468 | 1115.637 | 574 | 827.4058 | 455 | 805.0426 | 486 | RB1 | K562 | None | GSM28274 | 17067 | 101666 |
| 1111.051 | 279 | 1130.773 | 359 | 862.8936 | 278 | 853.6786 | 306 | RB1 | K562 | None | GSM28274 | 8586 | 101667 |
| 1085.113 | 320 | 1089.504 | 388 | 840.0685 | 312 | 792.6268 | 311 | MNT | K562 | None | GSM28288 | 9777 | 101668 |

| | | | | | | | | | | | | | |
|---|---|---|---|---|---|---|---|---|---|---|---|---|---|
| 1077.96 | 366 | 1088.49 | 447 | 824.8751 | 353 | 801.3539 | 376 | MNT | K562 | None | GSM28288 | 12033 | 101669 |
| 471.6163 | 281 | 681.3824 | 385 | 349.771 | 377 | 339.533 | 406 | CTCF | 22RV1 | Prostate | GSM28288 | 36838 | 101670 |
| 435.7463 | 258 | 685.7266 | 358 | 315.1263 | 342 | 303.286 | 367 | CTCF | 22RV1 | Prostate | GSM28288 | 33099 | 101671 |
| 444.5943 | 73 | 579.3353 | 93 | 183.3334 | 61 | 381.2369 | 96 | SUZ12 | MCF-7 | Breast | GSM28288 | 2223 | 101672 |
| 540.0453 | 97 | 586.5492 | 111 | 218.1844 | 82 | 378.1575 | 116 | SUZ12 | MCF-7 | Breast | GSM28288 | 3393 | 101673 |
| 213.5479 | 54 | 191.3174 | 56 | 136.8869 | 61 | 288.9386 | 95 | ZNF512B | MCF-7 | Breast | GSM28288 | 3032 | 101676 |
| 120.8195 | 27 | 102.5354 | 27 | 83.39425 | 31 | 195.0974 | 52 | ZNF512B | MCF-7 | Breast | GSM28288 | 1293 | 101677 |
| 952.3897 | 513 | 996.626 | 680 | 699.6703 | 546 | 681.5052 | 592 | CTCF | None | Uterus | GSM28274 | 33704 | 101678 |
| 368.6425 | 70 | 612.2043 | 103 | 355.9822 | 95 | 551.6971 | 128 | ARNT | HEK293T | Embryonic | GSM28288 | 2806 | 101679 |
| 500.6313 | 86 | 738.1007 | 118 | 350.9556 | 97 | 562.6069 | 133 | ARNT | HEK293T | Embryonic | GSM28288 | 2990 | 101680 |
| 752.8631 | 377 | 774.1474 | 483 | 581.2793 | 474 | 576.3035 | 534 | RAD21 | A549 | Lung | GSM31063 | 41866 | 101683 |
| 699.2801 | 343 | 738.5242 | 460 | 558.4066 | 467 | 565.246 | 540 | RAD21 | A549 | Lung | GSM31063 | 43290 | 101684 |
| 629.1618 | 288 | 751.8653 | 396 | 485.3631 | 388 | 530.6337 | 435 | RAD21 | A549 | Lung | GSM31063 | 34089 | 101685 |
| 1086.605 | 598 | 1112.467 | 753 | 799.4721 | 607 | 775.0283 | 655 | DRAP1 | HepG2 | None | GSM25344 | 28680 | 101690 |
| 1139.868 | 461 | 1173.821 | 599 | 868.6284 | 462 | 814.5773 | 457 | DRAP1 | HepG2 | None | GSM25344 | 15863 | 101691 |
| 987.3143 | 590 | 1021.837 | 756 | 720.4004 | 632 | 735.903 | 745 | ELF3 | HepG2 | None | GSM25344 | 39953 | 101696 |
| 944.433 | 562 | 979.9747 | 730 | 703.6704 | 629 | 707.0564 | 727 | ELF3 | HepG2 | None | GSM25344 | 41608 | 101697 |
| 784.3156 | 145 | 1048.696 | 199 | 566.6078 | 166 | 639.0958 | 188 | ZBTB10 | HEK293 | Embryonic | GSM28243 | 5108 | 101702 |
| 1040.695 | 211 | 1064.893 | 276 | 677.6525 | 212 | 820.0807 | 249 | ZBTB10 | HEK293 | Embryonic | GSM28243 | 7041 | 101703 |
| 48.98541 | 22 | 38.93811 | 22 | 39.86923 | 30 | 240.1603 | 77 | SALL1 | HEK293 | Embryonic | GSM28243 | 2363 | 101704 |
| 58.85618 | 19 | 39.09185 | 17 | 60.49835 | 28 | 264.3213 | 65 | SALL1 | HEK293 | Embryonic | GSM28243 | 1467 | 101705 |
| 377.9969 | 49 | 471.4268 | 61 | 166.7813 | 41 | 320.9774 | 63 | HSF1 | GM12878 | None | GSM28243 | 1153 | 101712 |
| 854.0718 | 120 | 937.8949 | 138 | 344.3427 | 98 | 382.6208 | 110 | HSF1 | GM12878 | None | GSM28243 | 3237 | 101713 |
| 113.0246 | 36 | 119.0116 | 41 | 185.5631 | 67 | 479.4088 | 118 | NCOA3 | MCF-7 | Breast | GSM28275 | 2722 | 101716 |
| 354.1039 | 100 | 308.9384 | 103 | 596.7065 | 187 | 870.6721 | 254 | NCOA3 | MCF-7 | Breast | GSM28275 | 6838 | 101717 |
| 915.8792 | 750 | 945.5546 | 940 | 667.6044 | 882 | 650.5573 | 980 | EP300 | None | tibial nerve | GSM28275 | 74227 | 101719 |
| 1224.613 | 196 | 1220.457 | 236 | 869.9946 | 181 | 874.5344 | 192 | CTCF | None | Skin | GSM28275 | 3807 | 101720 |
| 59.23478 | 17 | 39.06739 | 15 | 48.49773 | 22 | 286.1252 | 61 | ZNF747 | HEK293 | Embryonic | GSM28275 | 1125 | 101721 |
| 25.83209 | 12 | 14.10084 | 10 | 17.0977 | 15 | 224.0057 | 58 | ZNF747 | HEK293 | Embryonic | GSM28275 | 1365 | 101722 |
| 991.1976 | 559 | 1009.069 | 698 | 762.9594 | 646 | 760.6725 | 731 | EP300 | None | Lung | GSM28241 | 35549 | 101723 |
| 824.5398 | 255 | 815.4563 | 303 | 466.2683 | 287 | 417.4225 | 295 | MIER2 | HepG2 | None | GSM25274 | 20760 | 101744 |
| 1120.894 | 560 | 1131.305 | 686 | 824.0664 | 542 | 800.8829 | 578 | GATAD1 | HepG2 | None | GSM25274 | 22460 | 101750 |
| 1092.398 | 615 | 1124.717 | 784 | 810.0243 | 631 | 774.6416 | 666 | GATAD1 | HepG2 | None | GSM25274 | 29331 | 101751 |
| 479.3564 | 115 | 448.1619 | 122 | 222.7637 | 111 | 382.2282 | 154 | PHF21A | K562 | None | GSM28254 | 6160 | 101760 |
| 271.0126 | 52 | 288.9655 | 59 | 88.88403 | 40 | 346.8178 | 87 | PHF21A | K562 | None | GSM28254 | 2042 | 101761 |
| 929.418 | 399 | 930.7027 | 483 | 742.7874 | 473 | 732.2042 | 528 | ZNF629 | HEK293 | Embryonic | GSM28273 | 23175 | 101762 |
| 925.78 | 388 | 909.6299 | 456 | 734.4065 | 459 | 730.3031 | 520 | ZNF629 | HEK293 | Embryonic | GSM28273 | 23140 | 101763 |
| 1126.46 | 262 | 1151.504 | 344 | 869.4652 | 253 | 832.36 | 257 | AGO1 | K562 | None | GSM28273 | 7605 | 101764 |
| 1120.882 | 176 | 1140.964 | 228 | 623.7916 | 166 | 642.5316 | 179 | AGO1 | K562 | None | GSM28273 | 4847 | 101765 |
| 574.8828 | 64 | 412.716 | 56 | 302.2851 | 57 | 382.7986 | 69 | CSDE1 | MCF-7 | Breast | GSM28254 | 1041 | 101766 |
| 885.1508 | 427 | 914.7493 | 557 | 674.6169 | 489 | 649.2725 | 524 | CTCF | None | Esophagus | GSM28273 | 31773 | 101770 |
| 929.6969 | 401 | 926.765 | 479 | 685.2816 | 406 | 659.1403 | 430 | DPF2 | GM12878 | Blood | GSM28273 | 22252 | 101771 |
| 970.3702 | 403 | 966.3363 | 483 | 710.9151 | 393 | 687.2582 | 420 | DPF2 | GM12878 | Blood | GSM28273 | 19785 | 101772 |
| 278.479 | 64 | 475.0278 | 95 | 183.1572 | 72 | 349.4289 | 107 | TARDBP | HEK293T | Embryonic | GSM24239 | 3104 | 101774 |
| 438.5493 | 74 | 673.4544 | 104 | 251.5638 | 74 | 490.3531 | 113 | TARDBP | HEK293T | Embryonic | GSM24239 | 2335 | 101775 |
| 177.7055 | 54 | 161.9466 | 57 | 128.3889 | 67 | 125.4453 | 71 | ZNF3 | HepG2 | None | GSM28255 | 4031 | 101782 |
| 481.9083 | 138 | 571.7212 | 167 | 335.661 | 168 | 377.3214 | 190 | ZNF3 | HepG2 | None | GSM28255 | 9973 | 101783 |
| 26.04451 | 17 | 10.62571 | 13 | 20.77942 | 24 | 130.5039 | 59 | ZNF274 | HEK293 | Embryonic | GSM28255 | 2552 | 101784 |
| 13.70792 | 11 | 6.337711 | 9 | 14.79369 | 18 | 192.4502 | 64 | ZNF274 | HEK293 | Embryonic | GSM28255 | 1995 | 101785 |
| 900.249 | 256 | 929.6981 | 343 | 708.3045 | 305 | 565.3157 | 295 | NR2C1 | K562 | None | GSM28255 | 15502 | 101792 |
| 907.471 | 258 | 918.3661 | 328 | 622.4318 | 285 | 669.1963 | 316 | NR2C1 | K562 | None | GSM28255 | 15175 | 101793 |
| 520.8761 | 152 | 881.5732 | 221 | 409.4529 | 193 | 444.9104 | 215 | ZBTB2 | K562 | None | GSM28255 | 10008 | 101794 |
| 651.1176 | 132 | 971.0143 | 181 | 370.452 | 136 | 450.3441 | 160 | ZBTB2 | K562 | None | GSM28255 | 5471 | 101795 |
| 25.1993 | 13 | 6.81652 | 8 | 20.27905 | 18 | 199.4773 | 59 | ZNF670 | HEK293 | Embryonic | GSM28255 | 1646 | 101798 |
| 21.44683 | 12 | 11.24615 | 10 | 25.72697 | 20 | 199.5849 | 59 | ZNF670 | HEK293 | Embryonic | GSM28255 | 1636 | 101799 |
| 199.7413 | 48 | 201.4409 | 53 | 120.2693 | 52 | 223.3917 | 76 | NCOR1 | K562 | None | GSM24232 | 2484 | 101840 |
| 299.4696 | 74 | 276.8293 | 78 | 219.5391 | 89 | 273.5524 | 106 | NCOR1 | K562 | None | GSM24232 | 3995 | 101841 |
| 110.34 | 23 | 148.0536 | 30 | 90.32897 | 29 | 379.5394 | 69 | CTCF | None | Skin | GSM28272 | 1100 | 101850 |
| 1281.431 | 237 | 1296.976 | 302 | 992.0207 | 203 | 1023.956 | 257 | IRF3 | GM12878 | None | GSM24234 | 4062 | 101851 |
| 1355.47 | 207 | 1373.665 | 270 | 1080.294 | 188 | 1098.154 | 221 | IRF3 | GM12878 | None | GSM24234 | 2741 | 101852 |
| 1444.73 | 193 | 1440.338 | 230 | 945.1151 | 144 | 1048.532 | 161 | CEBPZ | K562 | None | GSM24237 | 2065 | 101857 |
| 135.0421 | 91 | 112.1488 | 95 | 143.6133 | 144 | 263.2088 | 199 | JUNB | A549 | Lung | GSM24377 | 13936 | 101867 |
| 371.7892 | 205 | 304.9392 | 212 | 282.8515 | 280 | 474.6721 | 371 | JUNB | A549 | Lung | GSM24377 | 27854 | 101868 |
| 169.4313 | 33 | 201.886 | 40 | 98.3809 | 34 | 272.761 | 63 | NFXL1 | GM12878 | Blood | GSM28288 | 1331 | 101873 |
| 274.6366 | 45 | 211.9104 | 42 | 122.0392 | 39 | 309.0288 | 69 | NFXL1 | GM12878 | Blood | GSM28288 | 1387 | 101874 |
| 311.5787 | 109 | 328.3295 | 124 | 101.1459 | 96 | 223.9569 | 145 | ZNF217 | GM12878 | Blood | GSM28288 | 8330 | 101876 |
| 434.6735 | 110 | 518.9121 | 133 | 179.0078 | 100 | 368.4417 | 151 | ZNF217 | GM12878 | Blood | GSM28288 | 5730 | 101877 |
| 579.4989 | 361 | 684.0258 | 494 | 443.9318 | 490 | 437.3808 | 532 | CTCF | C4-2B | Prostate | GSM28274 | 47813 | 101878 |
| 451.3008 | 259 | 693.1033 | 360 | 335.005 | 347 | 344.3755 | 381 | CTCF | C4-2B | Prostate | GSM28274 | 33400 | 101879 |
| 692.6004 | 223 | 814.9765 | 278 | 373.5799 | 248 | 513.2551 | 306 | CTCF | None | Peyer's pat | GSM28274 | 18956 | 101880 |
| 34.57325 | 15 | 9.388183 | 9 | 24.20238 | 19 | 187.5595 | 56 | ZNF114 | HEK293 | Embryonic | GSM28288 | 1558 | 101883 |
| 20.33058 | 11 | 5.919299 | 7 | 18.09018 | 16 | 173.6657 | 52 | ZNF114 | HEK293 | Embryonic | GSM28288 | 1456 | 101884 |

| | | | | | | | | | | | | | |
|---|---|---|---|---|---|---|---|---|---|---|---|---|---|
| 1037.172 | 280 | 1063.517 | 363 | 787.3236 | 270 | 803.4733 | 323 | ZNF687 | GM12878 | Blood | GSM28288 | 10529 | 101886 |
| 1033.18 | 391 | 1025.418 | 469 | 757.0547 | 365 | 753.0903 | 413 | ZNF687 | GM12878 | Blood | GSM28288 | 16118 | 101887 |
| 66.74067 | 21 | 40.93376 | 18 | 60.93907 | 29 | 231.1264 | 62 | ZNF621 | HEK293 | Embryonic | GSM28288 | 1538 | 101888 |
| 51.82294 | 19 | 33.81601 | 17 | 52.37559 | 28 | 210.8897 | 61 | ZNF621 | HEK293 | Embryonic | GSM28288 | 1646 | 101889 |
| 42.42339 | 17 | 13.88097 | 11 | 20.39216 | 18 | 232.5803 | 64 | ZNF658 | HEK293 | Embryonic | GSM28288 | 1624 | 101890 |
| 1034.262 | 292 | 1035.294 | 360 | 792.7193 | 297 | 794.7839 | 339 | EGR1 | K562 | None | GSM28251 | 11920 | 101892 |
| 1038.688 | 292 | 1034.276 | 354 | 780.0229 | 280 | 775.361 | 317 | EGR1 | K562 | None | GSM28251 | 11691 | 101893 |
| 1101.557 | 430 | 1098.687 | 521 | 804.5236 | 399 | 789.8518 | 439 | KLF6 | HepG2 | None | GSM25343 | 16683 | 101894 |
| 1093.12 | 407 | 1089.167 | 495 | 821.8223 | 400 | 805.0555 | 435 | KLF6 | HepG2 | None | GSM25343 | 15501 | 101895 |
| 830.7116 | 257 | 841.8571 | 323 | 697.8037 | 355 | 714.7488 | 427 | BCL6 | HepG2 | None | GSM25344 | 21026 | 101898 |
| 867.8886 | 437 | 884.6184 | 552 | 675.9523 | 532 | 678.0913 | 610 | BCL6 | HepG2 | None | GSM25344 | 35933 | 101899 |
| 862.3817 | 404 | 862.6202 | 492 | 656.4223 | 473 | 634.1735 | 506 | RCOR1 | SK-N-SH | Bone Marr | GSM28243 | 33953 | 101905 |
| 860.0687 | 555 | 886.1692 | 706 | 645.0651 | 664 | 619.2338 | 715 | RCOR1 | SK-N-SH | None | GSM28243 | 55255 | 101906 |
| 33.74913 | 15 | 38.72966 | 18 | 18.1074 | 17 | 189.1451 | 57 | SALL2 | HEK293 | Embryonic | GSM28243 | 1621 | 101907 |
| 132.1012 | 35 | 134.8776 | 39 | 86.4664 | 40 | 209.064 | 67 | SALL2 | HEK293 | Embryonic | GSM28243 | 2030 | 101908 |
| 826.6873 | 363 | 857.7408 | 477 | 627.7853 | 424 | 621.5515 | 475 | CTCF | None | Adrenal Gl | GSM28243 | 31102 | 101909 |
| 26.8032 | 42 | 52.33195 | 62 | 88.57146 | 106 | 258.4778 | 177 | NFIB | MCF-7 | Breast | GSM28275 | 11556 | 101910 |
| 64.42768 | 60 | 92.10361 | 79 | 130.4924 | 125 | 261.2264 | 180 | NFIB | MCF-7 | Breast | GSM28275 | 11747 | 101911 |
| 1221.52 | 303 | 1233.498 | 377 | 901.8019 | 246 | 887.4148 | 268 | FUS | HepG2 | None | GSM28275 | 6819 | 101912 |
| 1202.302 | 292 | 1220.885 | 371 | 913.236 | 261 | 911.9208 | 291 | FUS | HepG2 | None | GSM28276 | 6981 | 101913 |
| 906.5038 | 130 | 1127.326 | 161 | 332.665 | 101 | 457.2164 | 127 | HNRNPL | K562 | None | GSM28276 | 3490 | 101916 |
| 1131.966 | 231 | 1137.218 | 290 | 720.8217 | 204 | 860.2206 | 241 | HNRNPL | K562 | None | GSM28276 | 6661 | 101917 |
| 878.4058 | 399 | 903.6764 | 510 | 666.0509 | 449 | 639.4278 | 477 | CTCF | None | Esophagus | GSM28243 | 29238 | 101918 |
| 530.1125 | 175 | 656.9687 | 216 | 374.4864 | 216 | 341.0433 | 223 | KDM1A | HepG2 | Liver | GSM28254 | 13732 | 101921 |
| 605.3842 | 209 | 805.231 | 270 | 421.0715 | 258 | 518.5394 | 304 | KDM1A | HepG2 | Liver | GSM28254 | 17343 | 101922 |
| 761.1036 | 398 | 750.6523 | 472 | 579.3496 | 482 | 585.9103 | 560 | KDM1A | HepG2 | Liver | GSM28254 | 39997 | 101923 |
| 891.741 | 492 | 924.5902 | 635 | 682.5984 | 564 | 659.966 | 611 | CTBP1 | K562 | None | GSM28254 | 34952 | 101932 |
| 1002.019 | 202 | 1055.919 | 286 | 663.4248 | 218 | 725.0328 | 243 | CTBP1 | K562 | None | GSM28254 | 8072 | 101933 |
| 109.8143 | 24 | 67.53258 | 20 | 165.7079 | 42 | 298.2625 | 62 | HES1 | MCF-7 | Breast | GSM28254 | 1118 | 101934 |
| 124.553 | 32 | 135.9836 | 37 | 173.8473 | 54 | 436.702 | 95 | HES1 | MCF-7 | Breast | GSM28254 | 1827 | 101935 |
| 1033.128 | 176 | 1036.161 | 208 | 476.3835 | 160 | 441.1659 | 164 | RBM22 | HepG2 | None | GSM28273 | 6129 | 101936 |
| 364.4668 | 60 | 449.2317 | 74 | 132.7121 | 47 | 410.1354 | 92 | SNIP1 | K562 | None | GSM28273 | 1863 | 101939 |
| 475.0857 | 165 | 672.111 | 218 | 262.537 | 186 | 414.674 | 244 | CTCF | None | vagina | GSM28273 | 14486 | 101940 |
| 1034.666 | 399 | 1030.301 | 476 | 783.2232 | 400 | 759.535 | 425 | MNT | K562 | None | GSM28273 | 16249 | 101942 |
| 1017.019 | 356 | 1002.192 | 415 | 779.4354 | 364 | 762.6658 | 398 | MNT | K562 | None | GSM28273 | 14928 | 101943 |
| 637.9296 | 146 | 818.2621 | 184 | 399.9886 | 163 | 689.6093 | 228 | EP300 | None | Esophagus | GSM28273 | 7795 | 101944 |
| 294.6919 | 58 | 393.4202 | 75 | 217.3061 | 68 | 314.5035 | 88 | TBX3 | HepG2 | None | GSM28255 | 2347 | 101959 |
| 454.2049 | 102 | 636.7458 | 135 | 339.7771 | 122 | 438.309 | 148 | TBX3 | HepG2 | None | GSM28255 | 4738 | 101960 |
| 694.6475 | 124 | 966.2887 | 164 | 357.7713 | 120 | 500.9631 | 152 | HNRNPUL1 | HepG2 | None | GSM28255 | 4628 | 101965 |
| 1092.662 | 313 | 1084.492 | 374 | 843.8055 | 318 | 794.9218 | 314 | TARDBP | K562 | None | GSM25339 | 11525 | 101967 |
| 1132.079 | 418 | 1131.465 | 509 | 862.1037 | 419 | 810.4733 | 415 | TARDBP | K562 | None | GSM25339 | 15125 | 101968 |
| 120.5262 | 71 | 156.2019 | 90 | 177.2938 | 129 | 228.8914 | 155 | KDM6A | HepG2 | None | GSM25273 | 10374 | 101969 |
| 224.8916 | 105 | 329.5064 | 141 | 220.4868 | 157 | 314.3852 | 197 | KDM6A | HepG2 | None | GSM25274 | 12346 | 101970 |
| 405.7598 | 120 | 327.9454 | 119 | 205.8751 | 125 | 259.399 | 149 | ARNT | K562 | None | GSM28272 | 8122 | 101975 |
| 386.8248 | 113 | 336.0869 | 116 | 217.5416 | 122 | 296.4395 | 151 | ARNT | K562 | None | GSM28272 | 7114 | 101976 |
| 320.681 | 55 | 251.8568 | 52 | 159.78 | 51 | 350.3211 | 83 | HDAC1 | K562 | None | GSM24236 | 1763 | 101991 |
| 247.6082 | 48 | 153.5794 | 40 | 81.46089 | 37 | 220.728 | 66 | HDAC1 | K562 | None | GSM24236 | 1902 | 101992 |
| 1013.187 | 232 | 991.612 | 268 | 830.179 | 280 | 848.0213 | 332 | EP300 | None | Colon | GSM28272 | 10004 | 101993 |
| 1163.917 | 560 | 1204 | 725 | 866.1655 | 520 | 827.6986 | 541 | ZBTB26 | HEK293 | Embryonic | GSM28255 | 16048 | 102004 |
| 1135.42 | 535 | 1183.069 | 706 | 843.8698 | 500 | 821.8035 | 537 | ZBTB26 | HEK293 | Embryonic | GSM28255 | 16207 | 102005 |
| 453.7828 | 84 | 571.9081 | 105 | 320.2341 | 96 | 511.4559 | 131 | RUNX1 | K562 | None | GSM24234 | 3241 | 102008 |
| 362.3081 | 77 | 594.2848 | 112 | 236.3963 | 86 | 469.9756 | 131 | RUNX1 | K562 | None | GSM24234 | 3558 | 102009 |
| 655.4406 | 75 | 667.3362 | 82 | 284.8824 | 61 | 371.6794 | 75 | GTF2F1 | HepG2 | None | GSM24231 | 1431 | 102010 |
| 926.4642 | 109 | 803.5191 | 108 | 439.4306 | 94 | 460.9562 | 102 | GTF2F1 | HepG2 | None | GSM24231 | 2190 | 102011 |
| 799.1245 | 442 | 850.2459 | 596 | 605.9765 | 534 | 607.4854 | 610 | CTCF | None | Peyer's pat | GSM28289 | 44781 | 102012 |
| 882.2301 | 179 | 833.4014 | 190 | 382.8739 | 162 | 381.5564 | 173 | TOE1 | K562 | None | GSM24231 | 7498 | 102013 |
| 954.9726 | 267 | 941.3631 | 314 | 626.3592 | 266 | 611.8921 | 282 | TOE1 | K562 | None | GSM24231 | 12212 | 102014 |
| 1092.792 | 509 | 1116 | 642 | 842.1098 | 540 | 806.761 | 560 | MNT | MCF-7 | Mammary | GSM24238 | 20295 | 102019 |
| 1087.031 | 478 | 1109.625 | 602 | 848.78 | 511 | 826.2358 | 547 | MNT | MCF-7 | Mammary | GSM24238 | 17909 | 102020 |
| 40.34114 | 15 | 37.05369 | 16 | 32.4017 | 20 | 340.0013 | 73 | NCOA3 | MCF-7 | Breast | GSM28274 | 1398 | 102024 |
| 31.07854 | 14 | 14.98955 | 11 | 19.27259 | 17 | 196.5652 | 57 | NCOA3 | MCF-7 | Breast | GSM28274 | 1572 | 102025 |
| 878.9815 | 593 | 889.0419 | 730 | 655.5062 | 705 | 670.7567 | 830 | NFIL3 | HepG2 | None | GSM25275 | 57973 | 102026 |
| 832.404 | 383 | 833.704 | 470 | 655.6075 | 480 | 664.9554 | 560 | NFIL3 | HepG2 | None | GSM25275 | 34750 | 102027 |
| 534.9242 | 313 | 672.0642 | 412 | 342.0235 | 401 | 322.1073 | 428 | CTCF | PC-9 | Lung | GSM26987 | 42958 | 102028 |
| 366.1274 | 227 | 567.4284 | 308 | 234.1887 | 294 | 282.7718 | 342 | CTCF | PC-9 | Lung | GSM26987 | 33318 | 102029 |
| 1056.83 | 465 | 1066.816 | 572 | 803.1878 | 469 | 776.4606 | 494 | MNT | K562 | None | GSM28273 | 17634 | 102035 |
| 1071.044 | 485 | 1064.761 | 581 | 808.6213 | 489 | 771.4723 | 503 | MNT | K562 | None | GSM28273 | 19161 | 102036 |
| 717.3976 | 330 | 755.8662 | 439 | 455.5574 | 402 | 446.3086 | 433 | CTCF | None | Adrenal Gl | GSM28273 | 36121 | 102037 |
| 755.4181 | 416 | 795.3191 | 554 | 571.9384 | 510 | 579.115 | 592 | CTCF | None | Esophagus | GSM28273 | 46199 | 102038 |
| 1073.847 | 330 | 1100.753 | 423 | 851.9559 | 351 | 842.8133 | 389 | JUN | A549 | Lung | GSM24378 | 11332 | 102039 |
| 1098.005 | 517 | 1106.535 | 632 | 822.0975 | 510 | 813.3093 | 572 | JUN | A549 | Lung | GSM24378 | 19871 | 102040 |
| 131.7285 | 83 | 88.43017 | 79 | 92.14893 | 111 | 193.1455 | 161 | JUNB | A549 | Lung | GSM24378 | 12059 | 102041 |

| | | | | | | | | | | | | | |
|---|---|---|---|---|---|---|---|---|---|---|---|---|---|
| 70.32357 | 42 | 48.01699 | 40 | 68.47364 | 64 | 207.3825 | 112 | JUNB | A549 | Lung | GSM24378 | 5985 | 102042 |
| 109.0103 | 66 | 68.82734 | 61 | 90.12401 | 94 | 229.8719 | 150 | JUNB | A549 | Lung | GSM24378 | 9414 | 102043 |
| 973.6174 | 228 | 977.1854 | 284 | 785.1297 | 266 | 780.8655 | 295 | JUN | A549 | Lung | GSM24378 | 10336 | 102044 |
| 476.0388 | 90 | 542.0167 | 106 | 349.2961 | 105 | 520.9368 | 138 | JUN | A549 | Lung | GSM24378 | 3499 | 102045 |
| 1092.996 | 367 | 1100.188 | 451 | 837.828 | 358 | 819.5291 | 385 | JUN | A549 | Lung | GSM24378 | 11974 | 102046 |
| 1060.921 | 428 | 1045.946 | 500 | 795.091 | 417 | 785.4639 | 461 | JUN | A549 | Lung | GSM24378 | 16947 | 102047 |
| 1036.006 | 221 | 1004.357 | 246 | 720.0446 | 227 | 720.8313 | 242 | JUN | A549 | Lung | GSM24378 | 8070 | 102048 |
| 1145.636 | 481 | 1162.615 | 599 | 868.1935 | 462 | 848.4293 | 503 | JUN | A549 | Lung | GSM24378 | 14882 | 102049 |
| 1016.187 | 223 | 1013.491 | 272 | 765.6208 | 243 | 811.3655 | 277 | JUN | A549 | Lung | GSM24378 | 8683 | 102050 |
| 1073.141 | 624 | 1082.478 | 763 | 787.9519 | 635 | 772.9446 | 701 | JUN | A549 | Lung | GSM24378 | 30120 | 102051 |
| 54.35808 | 28 | 42.40045 | 28 | 34.16499 | 35 | 125.5182 | 67 | JUNB | A549 | Lung | GSM24378 | 3517 | 102052 |
| 150.885 | 75 | 70.52905 | 60 | 91.96291 | 92 | 202.1158 | 138 | JUNB | A549 | Lung | GSM24378 | 8864 | 102053 |
| 81.88751 | 59 | 53.68043 | 56 | 73.66594 | 88 | 154.4252 | 128 | JUNB | A549 | Lung | GSM24378 | 9416 | 102054 |
| 1048.951 | 270 | 1056.351 | 331 | 812.8867 | 275 | 807.1665 | 306 | JUN | A549 | Lung | GSM24378 | 10083 | 102055 |
| 1068.118 | 323 | 1067.605 | 393 | 831.5564 | 329 | 819.6971 | 362 | JUN | A549 | Lung | GSM24378 | 11010 | 102056 |
| 1071.482 | 331 | 1072.31 | 400 | 840.0646 | 347 | 830.8142 | 380 | JUN | A549 | Lung | GSM24378 | 11850 | 102057 |
| 0 | 5 | -0.007892 | 5 | -2.712247 | 3 | 3.82757 | 26 | ZNF507 | K562 | None | GSM28276 | 5232 | 102058 |
| 0.017405 | 6 | -0.071681 | 5 | -2.440395 | 4 | 2.256298 | 26 | ZNF507 | K562 | None | GSM28276 | 6009 | 102059 |
| 1101.259 | 516 | 1119.822 | 645 | 811.783 | 493 | 798.7215 | 545 | KMT2B | HepG2 | None | GSM25274 | 20708 | 102076 |
| 1102.434 | 432 | 1113.979 | 536 | 825.3445 | 410 | 785.0382 | 420 | KMT2B | HepG2 | None | GSM25274 | 15649 | 102077 |
| 283.0819 | 113 | 296.4801 | 129 | 278.4007 | 167 | 313.1571 | 189 | HMG20A | HepG2 | None | GSM25274 | 11548 | 102078 |
| 710.7356 | 327 | 723.698 | 410 | 574.8764 | 449 | 611.8389 | 565 | HMG20A | HepG2 | None | GSM25274 | 40671 | 102079 |
| 1026.307 | 765 | 1088.909 | 994 | 745.0075 | 861 | 716.0092 | 929 | CREM | K562 | None | GSM25274 | 60421 | 102084 |
| 1057.881 | 743 | 1109.464 | 957 | 779.8405 | 831 | 734.7734 | 864 | CREM | K562 | None | GSM25274 | 50792 | 102085 |
| 1061.32 | 197 | 1120.504 | 280 | 708.8991 | 203 | 800.5884 | 230 | BACH1 | GM12878 | None | GSM28272 | 6341 | 102086 |
| 967.6266 | 347 | 1001.587 | 454 | 729.6345 | 358 | 721.4467 | 398 | BACH1 | GM12878 | Blood | GSM28272 | 17929 | 102087 |
| 1224.155 | 227 | 1257.824 | 302 | 970.8764 | 221 | 914.1981 | 213 | GMEB2 | HepG2 | None | GSM25343 | 4777 | 102091 |
| 1203.381 | 172 | 1236.298 | 230 | 850.8277 | 173 | 770.7263 | 173 | GMEB2 | HepG2 | None | GSM25343 | 3778 | 102092 |
| 984.3309 | 338 | 1022.173 | 456 | 733.7169 | 341 | 717.5825 | 371 | CTCF | 11.9 | Blood | GSM27006 | 17021 | 102099 |
| 1013.296 | 460 | 1078.738 | 640 | 760.9755 | 476 | 726.0963 | 499 | CTCF | 11.9 | Blood | GSM27006 | 23119 | 102100 |
| 858.5738 | 246 | 815.4115 | 265 | 557.9647 | 289 | 412.2069 | 272 | CBX5 | K562 | None | GSM25276 | 18891 | 102104 |
| 333.3496 | 115 | 386.5229 | 138 | 218.5889 | 141 | 175.6569 | 138 | CBX5 | K562 | None | GSM25276 | 10759 | 102105 |
| 1170.376 | 163 | 1130.842 | 179 | 662.5385 | 155 | 582.436 | 153 | ZHX1 | HeLa-S3 | Cervix | GSM24241 | 4104 | 102128 |
| 918.7897 | 118 | 1050.814 | 139 | 499.8217 | 112 | 504.355 | 119 | ZHX1 | HeLa-S3 | Cervix | GSM24241 | 2807 | 102129 |
| 168.5065 | 33 | 115.2241 | 29 | 103.8263 | 35 | 242.5189 | 59 | ZNF704 | HEK293 | Embryonic | GSM28255 | 1309 | 102130 |
| 153.6518 | 32 | 110.1518 | 29 | 109.4923 | 37 | 322.3339 | 71 | ZNF704 | HEK293 | Embryonic | GSM28255 | 1386 | 102131 |
| 593.1084 | 365 | 687.1725 | 501 | 382.9156 | 472 | 418.6724 | 531 | CTCF | None | Prostate | GSM28255 | 51972 | 102132 |
| 567.5149 | 339 | 694.5446 | 469 | 387.8208 | 443 | 428.4783 | 500 | CTCF | None | Prostate | GSM28255 | 45423 | 102133 |
| 620.4427 | 86 | 862.3365 | 114 | 497.5261 | 101 | 404.8191 | 95 | LEF1 | HEK293T | Embryonic | GSM28255 | 2028 | 102134 |
| 800.7378 | 399 | 848.7849 | 536 | 568.9609 | 434 | 566.5145 | 480 | RAD21 | hESC | Embryo | GSM28255 | 36283 | 102136 |
| 908.7573 | 517 | 949.4468 | 682 | 642.9325 | 530 | 628.7468 | 584 | RAD21 | hESC | Embryo | GSM28255 | 38029 | 102137 |
| 894.5873 | 551 | 930.2232 | 711 | 634.295 | 571 | 600.2062 | 599 | RAD21 | hESC | Embryo | GSM28255 | 41783 | 102138 |
| 976.7543 | 302 | 1002.704 | 391 | 770.4385 | 341 | 804.6081 | 426 | EP300 | None | Sigmoid Cc | GSM28255 | 15297 | 102139 |
| 210.4247 | 120 | 283.0435 | 154 | 116.983 | 145 | 167.2048 | 180 | ZNF384 | HEK293T | Embryonic | GSM24240 | 15893 | 102140 |
| 205.5033 | 118 | 247.1182 | 144 | 101.9135 | 137 | 202.8063 | 193 | ZNF384 | HEK293T | Embryonic | GSM24240 | 15829 | 102141 |
| 1087.183 | 386 | 1077.65 | 460 | 794.3464 | 354 | 754.6433 | 358 | GABPA | GM12878 | None | GSM25339 | 14668 | 102151 |
| 1023.544 | 451 | 1039.288 | 566 | 729.6175 | 425 | 697.0602 | 443 | GABPA | GM12878 | None | GSM25339 | 23600 | 102152 |
| 401.5145 | 126 | 408.1247 | 141 | 485.2791 | 201 | 792.3852 | 280 | SMARCE1 | MCF-7 | Breast | GSM28272 | 10050 | 102168 |
| 454.8577 | 152 | 516.6276 | 180 | 546.2271 | 242 | 765.3699 | 326 | SMARCE1 | MCF-7 | Breast | GSM28272 | 12909 | 102169 |
| 1242.676 | 677 | 1291.265 | 876 | 913.2351 | 642 | 867.2745 | 660 | SMAD5 | K562 | None | GSM25273 | 21407 | 102176 |
| 1239.356 | 485 | 1282.448 | 636 | 947.2985 | 458 | 907.8494 | 469 | SMAD5 | K562 | None | GSM25273 | 12388 | 102177 |
| 812.3888 | 191 | 940.63 | 252 | 727.7042 | 257 | 495.0692 | 228 | ETV6 | K562 | None | GSM25273 | 11182 | 102178 |
| 166.8948 | 55 | 253.3504 | 76 | 89.19371 | 60 | 124.6716 | 75 | ETV6 | K562 | None | GSM25273 | 4440 | 102179 |
| 262.2596 | 178 | 330.391 | 222 | 190.0242 | 248 | 197.0311 | 273 | NFE2 | K562 | None | GSM25273 | 30844 | 102182 |
| 703.8059 | 343 | 742.4365 | 464 | 559.9259 | 475 | 480.8397 | 484 | NFE2 | K562 | None | GSM25273 | 47791 | 102183 |
| 940.2851 | 337 | 972.7625 | 444 | 689.6987 | 337 | 589.8201 | 334 | PKNOX1 | GM12878 | Blood | GSM24238 | 19004 | 102184 |
| 884.9011 | 415 | 895.7611 | 514 | 638.1606 | 425 | 621.9166 | 466 | PKNOX1 | GM12878 | None | GSM24238 | 29413 | 102185 |
| 236.2797 | 142 | 216.4699 | 154 | 141.0343 | 178 | 253.3517 | 241 | ZSCAN4 | HEK293 | Embryonic | GSM28254 | 19924 | 102204 |
| 297.2234 | 151 | 284.3851 | 166 | 154.2114 | 175 | 254.0417 | 230 | ZSCAN4 | HEK293 | Embryonic | GSM28254 | 18239 | 102205 |
| 1151.727 | 140 | 955.8784 | 135 | 811.1806 | 150 | 947.1015 | 173 | MTA1 | MCF-7 | Mammary | GSM24233 | 2609 | 102206 |
| 1155.54 | 145 | 1165.853 | 161 | 728.9951 | 147 | 918.228 | 177 | MTA1 | MCF-7 | Mammary | GSM24233 | 2933 | 102207 |
| 1034.55 | 384 | 1028.179 | 454 | 780.5057 | 379 | 750.4453 | 397 | EGR1 | K562 | None | GSM28254 | 16013 | 102208 |
| 1012.443 | 386 | 1020.459 | 478 | 777.496 | 403 | 735.7643 | 413 | EGR1 | K562 | None | GSM28254 | 17314 | 102209 |
| 1475.173 | 177 | 1506.497 | 235 | 1061.704 | 143 | 1162.986 | 161 | IRF3 | SK-N-SH | Brain | GSM24234 | 1682 | 102212 |
| 1375.048 | 208 | 1399.753 | 277 | 1010.015 | 169 | 1054.06 | 189 | IRF3 | SK-N-SH | None | GSM24234 | 2806 | 102213 |
| 865.7073 | 379 | 906.1332 | 507 | 664.1055 | 438 | 653.4435 | 484 | PKNOX1 | HEK293T | Embryonic | GSM24232 | 28846 | 102218 |
| 832.3004 | 397 | 874.1097 | 536 | 640.7359 | 478 | 607.3254 | 498 | PKNOX1 | HEK293T | Embryonic | GSM24232 | 34724 | 102219 |
| 87.73588 | 41 | 65.5025 | 40 | 64.48341 | 54 | 250.5379 | 108 | ZBTB33 | MCF-7 | Breast | GSM24232 | 4669 | 102220 |
| 53.67672 | 21 | 21.57318 | 15 | 33.07094 | 25 | 174.2006 | 60 | ZBTB33 | MCF-7 | Breast | GSM24232 | 2061 | 102221 |
| 909.9307 | 264 | 902.8846 | 317 | 709.75 | 307 | 744.2605 | 379 | EP300 | None | Sigmoid Cc | GSM28284 | 16056 | 102222 |
| 778.7022 | 115 | 780.3703 | 125 | 483.7925 | 118 | 617.8066 | 143 | TARDBP | MCF-7 | Breast | GSM28284 | 2979 | 102223 |
| 1044.22 | 184 | 1060.784 | 238 | 747.4529 | 208 | 819.4046 | 232 | TARDBP | MCF-7 | Breast | GSM28284 | 6317 | 102224 |

| | | | | | | | | | | | | | |
|---|---|---|---|---|---|---|---|---|---|---|---|---|---|
| 1210.833 | 129 | 1305.065 | 165 | 591.4039 | 110 | 1104.454 | 168 | SP1 | HEK293T | Embryonic | GSM28284 | 1970 | 102225 |
| 1205.294 | 127 | 1295.653 | 154 | 579.0866 | 107 | 1095.991 | 164 | SP1 | HEK293T | Embryonic | GSM28284 | 1894 | 102226 |
| 508.3497 | 100 | 772.338 | 139 | 233.7861 | 92 | 304.0786 | 112 | RNF2 | A549 | Lung | GSM27006 | 4003 | 102228 |
| 26.79545 | 13 | 14.90485 | 11 | 40.59625 | 24 | 85.44868 | 37 | ZNF112 | HEK293 | Embryonic | GSM28288 | 1589 | 102238 |
| 42.86838 | 17 | 14.05817 | 11 | 26.001 | 20 | 152.9349 | 51 | ZNF112 | HEK293 | Embryonic | GSM28288 | 1642 | 102239 |
| 74.04368 | 25 | 43.02035 | 21 | 77.16875 | 37 | 248.2317 | 72 | ZNF169 | HEK293 | Embryonic | GSM28273 | 1941 | 102241 |
| 38.01371 | 17 | 42.00135 | 20 | 24.78681 | 21 | 229.9105 | 67 | ZNF169 | HEK293 | Embryonic | GSM28273 | 1833 | 102242 |
| 157.8597 | 46 | 158.6212 | 51 | 311.9531 | 94 | 510.6415 | 130 | GLI2 | HEK293 | Embryonic | GSM28288 | 3160 | 102243 |
| 272.2354 | 42 | 327.1472 | 51 | 212.7301 | 49 | 308.221 | 64 | ZNF350 | HEK293 | Embryonic | GSM28288 | 1141 | 102248 |
| 830.3271 | 359 | 870.2035 | 480 | 622.8974 | 408 | 615.1226 | 457 | CTCF | None | gastroesop | GSM28273 | 30387 | 102250 |
| 754.2537 | 149 | 812.06 | 170 | 609.4138 | 184 | 743.4599 | 217 | JUN | A549 | Lung | GSM24378 | 6195 | 102257 |
| 1035.209 | 342 | 1017.819 | 397 | 800.0843 | 352 | 791.2837 | 393 | JUN | A549 | Lung | GSM24378 | 13793 | 102258 |
| 1085.94 | 578 | 1092.238 | 707 | 802.2002 | 573 | 774.0862 | 613 | JUN | A549 | Lung | GSM24378 | 24595 | 102259 |
| 1077.023 | 276 | 1096.996 | 351 | 858.3509 | 292 | 834.451 | 311 | JUN | A549 | Lung | GSM24378 | 8962 | 102260 |
| 1081.909 | 295 | 1087.536 | 364 | 851.2286 | 302 | 846.3593 | 337 | JUN | A549 | Lung | GSM24378 | 9561 | 102261 |
| 500.7741 | 278 | 702.014 | 386 | 369.5005 | 378 | 385.5931 | 417 | CTCF | A549 | Lung | GSM24378 | 39931 | 102262 |
| 542.3786 | 285 | 709.5535 | 388 | 373.6092 | 374 | 406.6723 | 419 | CTCF | A549 | Lung | GSM24378 | 38240 | 102263 |
| 697.0361 | 341 | 731.4886 | 455 | 436.6129 | 430 | 433.5935 | 466 | CTCF | A549 | Lung | GSM24378 | 45842 | 102264 |
| 60.44847 | 40 | 66.05171 | 47 | 67.52041 | 65 | 144.7919 | 97 | JUNB | A549 | Lung | GSM24378 | 6172 | 102265 |
| 75.92594 | 48 | 54.77377 | 47 | 72.80412 | 73 | 154.362 | 108 | JUNB | A549 | Lung | GSM24378 | 7147 | 102266 |
| 65.51533 | 46 | 74.4249 | 55 | 73.27902 | 75 | 165.8463 | 114 | JUNB | A549 | Lung | GSM24378 | 7397 | 102267 |
| 493.417 | 272 | 702.2037 | 379 | 378.5231 | 374 | 373.1906 | 404 | CTCF | A549 | Lung | GSM24378 | 38228 | 102268 |
| 454.1024 | 249 | 697.9567 | 345 | 334.518 | 336 | 322.8014 | 360 | CTCF | A549 | Lung | GSM24378 | 34045 | 102269 |
| 470.415 | 237 | 716.4523 | 324 | 329.1409 | 312 | 331.2499 | 339 | CTCF | A549 | Lung | GSM24378 | 31001 | 102270 |
| 1085.987 | 430 | 1109.563 | 545 | 835.3786 | 443 | 772.5019 | 425 | NFYC | HepG2 | None | GSM25341 | 16751 | 102273 |
| 1111.72 | 364 | 1131.713 | 471 | 852.7562 | 362 | 792.2823 | 348 | NFYC | HepG2 | None | GSM25341 | 12574 | 102274 |
| 436.867 | 227 | 470.6983 | 264 | 326.8518 | 310 | 409.4196 | 366 | ZNF644 | HepG2 | None | GSM25341 | 31832 | 102279 |
| 418.4006 | 199 | 508.1994 | 244 | 341.2074 | 278 | 372.283 | 311 | ZNF644 | HepG2 | None | GSM25341 | 25493 | 102280 |
| 882.6616 | 254 | 837.6223 | 272 | 492.5862 | 260 | 413.7374 | 259 | ZNF592 | K562 | None | GSM28255 | 14496 | 102283 |
| 742.4301 | 190 | 811.5516 | 219 | 402.5374 | 198 | 340.5517 | 197 | ZNF592 | K562 | None | GSM28255 | 10190 | 102284 |
| 915.2793 | 584 | 913.0669 | 707 | 681.4454 | 671 | 691.9054 | 785 | FOXA3 | HepG2 | None | GSM25274 | 49014 | 102285 |
| 886.3519 | 564 | 894.6757 | 697 | 682.8186 | 694 | 689.3922 | 804 | FOXA3 | HepG2 | None | GSM25274 | 51743 | 102286 |
| 332.7313 | 127 | 563.155 | 184 | 410.6809 | 208 | 464.9531 | 236 | KDM1A | HepG2 | None | GSM25274 | 12769 | 102293 |
| 846.7112 | 246 | 855.3276 | 312 | 642.7079 | 315 | 674.8905 | 351 | KDM1A | HepG2 | None | GSM25274 | 19372 | 102294 |
| 987.6512 | 589 | 1014.95 | 744 | 715.811 | 608 | 687.5425 | 650 | CTCF | None | tibial nerve | GSM28272 | 35464 | 102295 |
| 1081.183 | 299 | 1094.285 | 370 | 819.5754 | 279 | 814.7848 | 312 | ELF4 | K562 | None | GSM28272 | 9775 | 102296 |
| 1049.943 | 334 | 1060.245 | 410 | 776.8461 | 306 | 777.5706 | 352 | ELF4 | K562 | None | GSM28272 | 12576 | 102297 |
| 286.2238 | 43 | 269.1012 | 45 | 116.5484 | 35 | 425.0257 | 77 | BCLAF1 | GM12878 | Blood | GSM28272 | 1180 | 102300 |
| 402.0496 | 57 | 469.5167 | 68 | 197.4811 | 51 | 405.316 | 81 | BCLAF1 | GM12878 | Blood | GSM28272 | 1427 | 102301 |
| 136.4477 | 44 | 177.6153 | 56 | 75.36842 | 48 | 176.2785 | 77 | NFATC3 | K562 | None | GSM28274 | 3258 | 102304 |
| 238.1303 | 58 | 226.1893 | 62 | 112.3153 | 56 | 244.3275 | 88 | NFATC3 | K562 | None | GSM28274 | 3074 | 102305 |
| 833.9341 | 403 | 819.5946 | 476 | 646.0849 | 494 | 640.1033 | 555 | SOX13 | HepG2 | None | GSM25343 | 36495 | 102308 |
| 912.448 | 713 | 943.9538 | 896 | 697.8887 | 893 | 694.2651 | 1014 | SOX13 | HepG2 | None | GSM25343 | 72112 | 102309 |
| 231.8839 | 60 | 181.6817 | 58 | 242.1439 | 87 | 525.34 | 139 | GATAD2B | MCF-7 | Breast | GSM28288 | 3505 | 102314 |
| 308.4804 | 80 | 340.5331 | 93 | 372.5937 | 125 | 658.0949 | 179 | GATAD2B | MCF-7 | Breast | GSM28288 | 4723 | 102315 |
| 820.0702 | 351 | 846.1784 | 457 | 593.9375 | 397 | 607.2858 | 454 | CTCF | None | Testis | GSM28288 | 31697 | 102316 |
| 505.5903 | 65 | 610.9088 | 79 | 265.9775 | 59 | 350.6644 | 73 | FOXK2 | HEK293T | Embryonic | GSM24240 | 1296 | 102343 |
| 383.9315 | 112 | 386.4094 | 124 | 295.8385 | 140 | 463.8159 | 186 | ZSCAN21 | HEK293 | Embryonic | GSM28255 | 7033 | 102351 |
| 1137.467 | 302 | 1148.914 | 379 | 866.1878 | 280 | 824.6775 | 281 | MXD3 | HepG2 | None | GSM25346 | 8427 | 102380 |
| 1127.993 | 258 | 1174.6 | 349 | 898.0091 | 265 | 854.1325 | 263 | MXD3 | HepG2 | None | GSM25346 | 7369 | 102381 |
| 424.8067 | 101 | 540.0767 | 127 | 396.7064 | 139 | 272.3022 | 123 | MITF | K562 | None | GSM25273 | 5879 | 102442 |
| 960.9061 | 209 | 918.0568 | 230 | 594.1654 | 230 | 480.2748 | 222 | MITF | K562 | None | GSM25273 | 11087 | 102443 |
| 38.48996 | 13 | 27.09043 | 12 | 7.643978 | 9 | 305.8749 | 62 | ZNF530 | HEK293 | Embryonic | GSM28283 | 1085 | 102446 |
| 40.17347 | 13 | 54.12056 | 17 | 18.46457 | 13 | 298.9695 | 60 | ZNF530 | HEK293 | Embryonic | GSM28283 | 1041 | 102447 |
| 34.99197 | 14 | 11.41709 | 9 | 25.68174 | 18 | 260.6867 | 63 | ZNF155 | HEK293 | Embryonic | GSM28283 | 1381 | 102448 |
| 20.98478 | 11 | 16.42927 | 11 | 30.4035 | 20 | 258.4036 | 64 | ZNF155 | HEK293 | Embryonic | GSM28283 | 1450 | 102449 |
| 180.1523 | 38 | 290.4978 | 55 | 126.778 | 44 | 437.8486 | 92 | EP300 | None | gastroesop | GSM28283 | 1743 | 102450 |
| 153.4037 | 36 | 163.4923 | 41 | 102.4684 | 41 | 306.5293 | 78 | ZNF223 | HEK293 | Embryonic | GSM28283 | 1842 | 102451 |
| 133.2178 | 35 | 197.428 | 48 | 95.80572 | 42 | 369.5692 | 91 | ZNF223 | HEK293 | Embryonic | GSM28283 | 2054 | 102452 |
| 943.0012 | 312 | 943.7934 | 386 | 788.167 | 404 | 809.8264 | 490 | NFIB | MCF-7 | Breast | GSM28283 | 17211 | 102453 |
| 958.3381 | 254 | 939.3204 | 295 | 804.042 | 329 | 830.7533 | 401 | NFIB | MCF-7 | Breast | GSM28283 | 12732 | 102454 |
| 348.9612 | 56 | 761.4215 | 98 | 311.6285 | 71 | 372.3315 | 83 | SNIP1 | MCF-7 | Breast | GSM28243 | 1652 | 102457 |
| 964.1755 | 445 | 976.0208 | 556 | 716.7449 | 463 | 702.4453 | 510 | CTCF | None | Lung | GSM28287 | 25008 | 102459 |
| 52.87197 | 18 | 79.71835 | 25 | 46.28851 | 25 | 293.6543 | 70 | SMARCE1 | HepG2 | None | GSM28287 | 1616 | 102460 |
| 967.9674 | 577 | 991.0646 | 722 | 708.7811 | 613 | 683.8662 | 658 | CTCF | None | Lung | GSM28288 | 37313 | 102462 |
| 902.684 | 181 | 994.0659 | 231 | 481.4851 | 181 | 526.6951 | 202 | NR2C1 | K562 | None | GSM28288 | 7506 | 102463 |
| 938.1139 | 205 | 972.7331 | 266 | 520.4376 | 212 | 535.8602 | 230 | NR2C1 | K562 | None | GSM28288 | 9470 | 102464 |
| 493.0503 | 96 | 596.9044 | 117 | 192.9542 | 81 | 289.8368 | 106 | ZNF407 | K562 | None | GSM24228 | 3690 | 102477 |
| 369.4001 | 61 | 484.1343 | 78 | 142.079 | 49 | 267.4339 | 73 | ZNF407 | K562 | None | GSM24228 | 1842 | 102478 |
| 369.6365 | 201 | 615.3568 | 283 | 251.4838 | 261 | 290.7133 | 299 | CTCF | A549 | Lung | GSM24377 | 25913 | 102486 |
| 514.1075 | 272 | 723.3994 | 386 | 375.7932 | 365 | 401.4829 | 406 | CTCF | A549 | Lung | GSM24377 | 36530 | 102487 |
| 452.2925 | 247 | 712.6389 | 353 | 352.0591 | 340 | 373.2098 | 377 | CTCF | A549 | Lung | GSM24377 | 33578 | 102488 |

| | | | | | | | | | | | | | |
|---|---|---|---|---|---|---|---|---|---|---|---|---|---|
| 503.9783 | 284 | 699.7147 | 395 | 372.2842 | 386 | 363.7005 | 416 | CTCF | A549 | Lung | GSM24377 | 40699 | 102489 |
| 689.8824 | 345 | 741.4894 | 475 | 467.0726 | 445 | 416.2035 | 464 | CTCF | A549 | Lung | GSM24377 | 44736 | 102490 |
| 427.3382 | 236 | 704.0736 | 333 | 316.3826 | 319 | 330.9145 | 352 | CTCF | A549 | Lung | GSM24377 | 32202 | 102491 |
| 572.8451 | 222 | 512.434 | 235 | 332.2232 | 259 | 433.7089 | 312 | JUNB | A549 | Lung | GSM24377 | 21241 | 102492 |
| 118.7643 | 80 | 105.8585 | 86 | 124.6399 | 126 | 243.4432 | 179 | JUNB | A549 | Lung | GSM24377 | 12226 | 102493 |
| 42.66679 | 27 | 38.18489 | 29 | 62.02522 | 49 | 124.4608 | 72 | JUNB | A549 | Lung | GSM24377 | 4017 | 102494 |
| 115.1302 | 74 | 98.4098 | 78 | 125.508 | 118 | 233.8509 | 165 | JUNB | A549 | Lung | GSM24377 | 10911 | 102495 |
| 414.0976 | 247 | 699.8752 | 362 | 347.9486 | 354 | 327.3696 | 376 | CTCF | A549 | Lung | GSM24377 | 36705 | 102496 |
| 409.0412 | 249 | 692.6959 | 364 | 332.3838 | 353 | 338.3315 | 386 | CTCF | A549 | Lung | GSM24377 | 37490 | 102497 |
| 532.7696 | 285 | 721.6805 | 407 | 394.8245 | 387 | 425.8556 | 432 | CTCF | A549 | Lung | GSM24377 | 40049 | 102498 |
| 267.8483 | 73 | 355.8617 | 94 | 239.1012 | 99 | 620.6351 | 172 | EP300 | None | Sigmoid Cc | GSM28271 | 4784 | 102500 |
| 1069.522 | 447 | 1084.851 | 558 | 816.1504 | 462 | 812.3837 | 522 | HNRNPL | HepG2 | None | GSM28271 | 18890 | 102501 |
| 1057.369 | 353 | 1065.863 | 438 | 811.3557 | 363 | 818.7262 | 423 | HNRNPL | HepG2 | None | GSM28271 | 14230 | 102502 |
| 1025.192 | 244 | 1015.729 | 290 | 817.5088 | 270 | 812.2452 | 300 | JUN | A549 | Lung | GSM24378 | 9571 | 102503 |
| 793.5718 | 157 | 887.0287 | 183 | 470.6616 | 166 | 530.934 | 188 | JUN | A549 | Lung | GSM24378 | 6678 | 102505 |
| 954.3929 | 266 | 947.8331 | 317 | 594.5937 | 257 | 677.9802 | 293 | RNF2 | K562 | None | GSM28271 | 11604 | 102506 |
| 462.7688 | 227 | 505.3458 | 264 | 344.0042 | 298 | 447.457 | 358 | RNF2 | K562 | None | GSM28271 | 24963 | 102507 |
| 1219.737 | 301 | 1261.822 | 402 | 980.0549 | 302 | 918.0445 | 287 | NONO | K562 | None | GSM28271 | 6630 | 102508 |
| 1205.365 | 209 | 1242.091 | 279 | 820.3687 | 185 | 821.2654 | 196 | NONO | K562 | None | GSM28271 | 4502 | 102509 |
| 30.40726 | 13 | 20.71557 | 12 | 22.62173 | 17 | 210.4786 | 56 | ZNF662 | HEK293 | Embryonic | GSM28275 | 1391 | 102511 |
| 24.52864 | 14 | 11.16481 | 11 | 18.39638 | 19 | 190.8114 | 62 | ZNF493 | HEK293 | Embryonic | GSM28275 | 1911 | 102512 |
| 815.4478 | 355 | 850.4749 | 470 | 619.1513 | 412 | 627.2716 | 480 | CTCF | None | Breast | GSM28275 | 30202 | 102514 |
| 1114.933 | 239 | 1107.507 | 285 | 835.9412 | 228 | 883.1257 | 276 | EP300 | None | Ovary | GSM28275 | 6978 | 102515 |
| 114.0656 | 146 | 155.7446 | 187 | 136.956 | 251 | 199.1775 | 310 | CBFA2T3 | K562 | None | GSM28275 | 35177 | 102517 |
| 218.4562 | 207 | 255.5695 | 250 | 195.354 | 316 | 269.8942 | 384 | CBFA2T3 | K562 | None | GSM28275 | 42233 | 102518 |
| 942.3506 | 383 | 937.5879 | 462 | 717.6916 | 420 | 708.5128 | 469 | TFE3 | HepG2 | None | GSM25341 | 23752 | 102527 |
| 984.8909 | 394 | 990.1787 | 484 | 746.9687 | 418 | 736.8785 | 464 | TFE3 | HepG2 | None | GSM25341 | 21166 | 102528 |
| 752.0912 | 372 | 773.9646 | 476 | 590.4412 | 480 | 588.946 | 545 | SMC3 | A549 | Lung | GSM31063 | 40856 | 102529 |
| 664.8533 | 297 | 743.5923 | 391 | 457.6083 | 383 | 505.1307 | 431 | SMC3 | A549 | Lung | GSM31063 | 35212 | 102531 |
| 789.0791 | 407 | 808.3899 | 513 | 607.8056 | 494 | 607.9815 | 563 | RAD21 | A549 | Lung | GSM31063 | 38911 | 102532 |
| 656.1795 | 306 | 722.8076 | 394 | 473.1239 | 406 | 528.6127 | 459 | RAD21 | A549 | Lung | GSM31063 | 39053 | 102533 |
| 530.4664 | 262 | 718.5528 | 353 | 385.9542 | 348 | 470.8081 | 407 | RAD21 | A549 | Lung | GSM31063 | 33020 | 102534 |
| 770.9565 | 421 | 807.0101 | 552 | 605.0082 | 536 | 613.4918 | 625 | RAD21 | A549 | Lung | GSM31063 | 44507 | 102535 |
| 752.322 | 357 | 780.8263 | 466 | 579.9749 | 449 | 584.544 | 518 | RAD21 | A549 | Lung | GSM31063 | 39441 | 102537 |
| 764.6638 | 384 | 766.9341 | 473 | 577.7181 | 465 | 580.3092 | 534 | RAD21 | A549 | Lung | GSM31063 | 40845 | 102538 |
| 743.5144 | 376 | 776.8287 | 498 | 586.1894 | 491 | 594.6924 | 572 | RAD21 | A549 | Lung | GSM31063 | 42913 | 102539 |
| 818.0847 | 501 | 828.4321 | 621 | 604.9373 | 590 | 605.7941 | 673 | RAD21 | A549 | Lung | GSM31063 | 52061 | 102540 |
| 764.6438 | 371 | 798.3674 | 490 | 590.3197 | 464 | 582.2926 | 518 | SMC3 | A549 | Lung | GSM31063 | 39849 | 102541 |
| 765.7395 | 396 | 789.4062 | 511 | 601.8292 | 510 | 596.3166 | 573 | SMC3 | A549 | Lung | GSM31063 | 42415 | 102543 |
| 725.6209 | 320 | 751.496 | 418 | 498.8987 | 409 | 568.0909 | 474 | SMC3 | A549 | Lung | GSM31063 | 38278 | 102545 |
| 733.8869 | 314 | 764.9589 | 414 | 498.574 | 395 | 568.6082 | 451 | SMC3 | A549 | Lung | GSM31063 | 34387 | 102546 |
| 228.281 | 82 | 313.2059 | 107 | 67.88781 | 71 | 103.1896 | 91 | SMC3 | A549 | Lung | GSM31063 | 7096 | 102547 |
| 742.4994 | 352 | 760.0798 | 449 | 577.4272 | 452 | 580.9542 | 521 | SMC3 | A549 | Lung | GSM31063 | 40105 | 102548 |
| 657.7732 | 282 | 728.2883 | 350 | 440.9942 | 359 | 496.6579 | 407 | SMC3 | A549 | Lung | GSM31063 | 32693 | 102549 |
| 634.4275 | 265 | 732.1173 | 327 | 411.4507 | 329 | 463.186 | 373 | SMC3 | A549 | Lung | GSM31063 | 28374 | 102550 |
| 572.4825 | 270 | 735.6977 | 368 | 431.0088 | 363 | 474.5285 | 408 | SMC3 | A549 | Lung | GSM31063 | 33602 | 102551 |
| 681.3853 | 298 | 745.1782 | 389 | 476.4123 | 388 | 527.546 | 437 | SMC3 | A549 | Lung | GSM31063 | 36032 | 102552 |
| 672.9985 | 81 | 899.2948 | 105 | 284.318 | 65 | 474.6062 | 92 | FUS | K562 | None | GSM28243 | 1587 | 102553 |
| 748.312 | 81 | 926.186 | 100 | 260.8968 | 57 | 365.2002 | 73 | FUS | K562 | None | GSM28243 | 1315 | 102554 |
| 231.3729 | 62 | 229.3718 | 68 | 74.21413 | 51 | 74.52355 | 55 | REST | hESC | Embryo | GSM27005 | 3649 | 102566 |
| 992.8609 | 489 | 1010.288 | 620 | 731.165 | 511 | 717.9129 | 564 | ZSCAN9 | HepG2 | None | GSM25343 | 29041 | 102567 |
| 988.466 | 260 | 984.9827 | 313 | 771.8665 | 280 | 738.7721 | 292 | ZSCAN9 | HepG2 | None | GSM25343 | 12350 | 102568 |
| 1019.59 | 532 | 1009.673 | 624 | 764.4571 | 552 | 720.5111 | 566 | ZGPAT | HepG2 | None | GSM25343 | 26362 | 102571 |
| 957.4708 | 357 | 950.1287 | 426 | 715.742 | 366 | 705.6963 | 404 | ZGPAT | HepG2 | None | GSM25343 | 18899 | 102572 |
| 41.89383 | 16 | 15.87802 | 11 | 36.11919 | 22 | 160.9888 | 50 | ZIK1 | HEK293 | Embryonic | GSM28250 | 1474 | 102577 |
| 30.28281 | 14 | 9.384439 | 9 | 33.11467 | 22 | 218.6603 | 61 | ZIK1 | HEK293 | Embryonic | GSM28250 | 1592 | 102578 |
| 45.08796 | 17 | 7.508219 | 8 | 19.46038 | 17 | 217.3133 | 60 | ZNF280C | HEK293 | Embryonic | GSM28251 | 1543 | 102579 |
| 26.33811 | 12 | 11.56864 | 9 | 29.36444 | 19 | 256.3309 | 62 | ZNF280C | HEK293 | Embryonic | GSM28251 | 1341 | 102580 |
| 1040.557 | 411 | 1061.68 | 522 | 772.8182 | 395 | 730.655 | 404 | ASH2L | hESC | Embryo | GSM34984 | 16876 | 102586 |
| 987.4376 | 389 | 994.6815 | 476 | 730.6974 | 374 | 686.9821 | 380 | ASH2L | hESC | Embryo | GSM34984 | 17266 | 102587 |
| 333.5067 | 48 | 291.7278 | 48 | 118.9498 | 36 | 154.0956 | 44 | HDAC6 | GM12878 | Blood | GSM34984 | 1212 | 102588 |
| 442.6147 | 163 | 354.3238 | 163 | 285.8656 | 196 | 301.347 | 216 | NCOR1 | K562 | None | GSM24241 | 14190 | 102607 |
| 367.0479 | 164 | 303.1037 | 168 | 272.9427 | 215 | 275.2012 | 233 | NCOR1 | K562 | None | GSM24241 | 17525 | 102608 |
| 862.0793 | 413 | 868.2224 | 513 | 648.5988 | 478 | 652.6974 | 551 | MIXL1 | HepG2 | None | GSM25346 | 35233 | 102615 |
| 907.7303 | 386 | 880.6922 | 440 | 664.2228 | 406 | 637.9535 | 431 | MIXL1 | HepG2 | None | GSM25346 | 27342 | 102616 |
| 1040.54 | 734 | 1097.083 | 953 | 758.8249 | 806 | 753.7304 | 910 | SAP130 | HepG2 | None | GSM25346 | 48978 | 102617 |
| 1080.571 | 866 | 1126.918 | 1088 | 761.8532 | 951 | 766.7241 | 1101 | SAP130 | HepG2 | None | GSM25346 | 65054 | 102618 |
| 812.3087 | 268 | 795.2866 | 313 | 544.7812 | 325 | 480.3498 | 332 | SOX5 | HepG2 | None | GSM25346 | 23243 | 102619 |
| 898.6512 | 602 | 902.0946 | 731 | 672.2265 | 707 | 671.5274 | 809 | SOX5 | HepG2 | None | GSM25346 | 54168 | 102620 |
| 85.14406 | 54 | 46.25016 | 47 | 65.41785 | 75 | 182.9904 | 124 | JUNB | A549 | Lung | GSM24377 | 7926 | 102621 |
| 214.8924 | 109 | 163.0506 | 108 | 130.522 | 134 | 267.4493 | 194 | JUNB | A549 | Lung | GSM24377 | 13107 | 102622 |
| 48.30181 | 38 | 45.3775 | 42 | 79.11763 | 73 | 169.2573 | 109 | JUNB | A549 | Lung | GSM24377 | 6724 | 102623 |

| | | | | | | | | | | | | | |
|---|---|---|---|---|---|---|---|---|---|---|---|---|---|
| 114.4271 | 64 | 88.03372 | 64 | 96.95439 | 91 | 237.9275 | 144 | JUNB | A549 | Lung | GSM24377 | 8381 | 102624 |
| 438.1434 | 252 | 697.8282 | 360 | 314.009 | 338 | 342.4614 | 379 | CTCF | A549 | Lung | GSM24377 | 35550 | 102626 |
| 646.5728 | 323 | 735.2735 | 448 | 466.6018 | 430 | 445.2512 | 459 | CTCF | A549 | Lung | GSM24377 | 42495 | 102628 |
| 511.6326 | 291 | 706.0909 | 413 | 402.2744 | 405 | 396.9159 | 438 | CTCF | A549 | Lung | GSM24377 | 42042 | 102629 |
| 1058.437 | 223 | 1070.01 | 280 | 718.6481 | 222 | 692.9085 | 232 | KLF9 | HepG2 | None | GSM25346 | 8153 | 102643 |
| 784.6298 | 127 | 1103.812 | 171 | 414.8674 | 123 | 346.3599 | 119 | KLF9 | HepG2 | None | GSM25346 | 4226 | 102644 |
| 843.684 | 540 | 867.8238 | 683 | 653.7603 | 673 | 666.2504 | 792 | GATAD2A | HepG2 | None | GSM25346 | 53230 | 102645 |
| 855.4488 | 579 | 867.9162 | 713 | 654.7189 | 715 | 674.0574 | 853 | GATAD2A | HepG2 | None | GSM25346 | 57896 | 102646 |
| 698.0753 | 170 | 878.6088 | 212 | 566.187 | 218 | 517.4951 | 223 | HMG20B | HepG2 | None | GSM25346 | 10129 | 102653 |
| 809.6593 | 201 | 900.1734 | 249 | 546.669 | 237 | 552.7967 | 255 | HMG20B | HepG2 | None | GSM25346 | 12445 | 102654 |
| 520.7438 | 280 | 717.7247 | 395 | 402.9287 | 378 | 442.104 | 425 | CTCF | PANC-1 | Pancreas | GSM28254 | 34097 | 102718 |
| 398.0888 | 297 | 633.3146 | 410 | 326.412 | 422 | 308.1243 | 452 | CTCF | PANC-1 | Pancreas | GSM28254 | 46171 | 102719 |
| 899.6175 | 283 | 906.8321 | 351 | 500.2003 | 270 | 562.2918 | 306 | ZFX | MCF-7 | Breast | GSM28273 | 14687 | 102720 |
| 1136.023 | 249 | 1160.011 | 322 | 902.8579 | 242 | 887.5204 | 264 | ZFX | MCF-7 | Breast | GSM28273 | 5581 | 102721 |
| 1163.642 | 303 | 1171.62 | 376 | 911.9261 | 298 | 878.5564 | 308 | SIN3B | HepG2 | None | GSM24235 | 8484 | 102726 |
| 1248.105 | 190 | 1255.644 | 235 | 896.9399 | 176 | 880.9489 | 184 | SIN3B | HepG2 | None | GSM24235 | 3651 | 102727 |
| 268.9354 | 43 | 392.5034 | 59 | 82.86316 | 31 | 214.868 | 55 | ESRRA | A549 | Lung | GSM24235 | 1425 | 102730 |
| 204.2569 | 47 | 373.1034 | 73 | 69.64202 | 39 | 250.0225 | 79 | ESRRA | A549 | Lung | GSM24235 | 2600 | 102731 |
| 277.6212 | 55 | 292.3734 | 62 | 124.9392 | 50 | 260.3531 | 78 | BMI1 | GM12878 | Blood | GSM24235 | 2266 | 102733 |
| 973.2036 | 344 | 994.2371 | 439 | 720.0747 | 338 | 729.4217 | 398 | HDAC1 | K562 | None | GSM28285 | 16665 | 102736 |
| 1012.802 | 180 | 1032.327 | 221 | 541.1998 | 178 | 680.0671 | 213 | TOE1 | MCF-7 | Breast | GSM28284 | 6718 | 102738 |
| 964.9476 | 385 | 954.9066 | 452 | 711.5629 | 379 | 705.5203 | 423 | MGA | K562 | None | GSM28284 | 18928 | 102740 |
| 975.8536 | 317 | 971.5401 | 383 | 731.7563 | 314 | 715.2557 | 339 | MGA | K562 | None | GSM28284 | 13957 | 102741 |
| 788.6581 | 290 | 833.7635 | 398 | 544.5466 | 354 | 594.2006 | 396 | CTCF | None | Thyroid | GSM28288 | 26844 | 102747 |
| 995.8592 | 439 | 1012.573 | 548 | 756.2851 | 460 | 725.3024 | 481 | ATF7 | K562 | None | GSM28288 | 21184 | 102748 |
| 63.01676 | 17 | 185.5059 | 35 | 38.81812 | 19 | 141.9946 | 40 | KDM1A | SET-2 | None | GSM34362 | 1082 | 103010 |
| 117.5861 | 43 | 218.2042 | 66 | 58.6142 | 46 | 126.5337 | 70 | KDM1A | SET-2 | None | GSM34362 | 3758 | 103014 |
| 836.0295 | 469 | 861.0115 | 600 | 625.7494 | 548 | 614.9319 | 608 | RAD21 | HeLa | Cervix | GSM29904 | 44825 | 103198 |
| 907.6855 | 678 | 961.5732 | 890 | 672.4772 | 805 | 658.8768 | 897 | RAD21 | HeLa | Cervix | GSM29904 | 66749 | 103199 |
| 1.330533 | 29 | 2.241925 | 38 | 0.840844 | 53 | 10.86396 | 91 | ZFP57 | H1 | Embryo | GSM31771 | 20152 | 103253 |
| 1067.851 | 144 | 1171.764 | 180 | 619.2958 | 141 | 686.6092 | 158 | KMT2A | HEK293T | Kidney | GSM34449 | 3331 | 103306 |
| 1028.926 | 128 | 1097.519 | 144 | 618.0777 | 126 | 521.0802 | 121 | KMT2A | HEK293T | Kidney | GSM34449 | 2558 | 103307 |
| 348.3405 | 47 | 514.5482 | 65 | 151.2253 | 39 | 117.5095 | 36 | ELL2 | HEK293T | Kidney | GSM34449 | 1124 | 103309 |
| 642.6484 | 96 | 694.8262 | 109 | 240.1552 | 74 | 218.8698 | 75 | KMT2A | MCF-7 | Breast | GSM34449 | 2297 | 103310 |
| 621.1173 | 91 | 812.7136 | 116 | 317.4989 | 83 | 317.195 | 88 | KMT2A | MCF-7 | Breast | GSM34449 | 2177 | 103311 |
| 927.5298 | 138 | 1121.342 | 174 | 430.9507 | 120 | 390.1429 | 121 | KMT2A | HEK293T | Kidney | GSM34449 | 3314 | 103317 |
| 855.3467 | 123 | 1052.032 | 151 | 370.5848 | 103 | 420.4666 | 117 | KMT2A | HEK293T | Kidney | GSM34449 | 2981 | 103320 |
| 1024.702 | 169 | 1043.858 | 192 | 475.6882 | 149 | 371.0738 | 140 | KMT2A | HEK293T | Kidney | GSM34449 | 4472 | 103323 |
| 1122.478 | 191 | 1098.523 | 217 | 646.298 | 173 | 513.3056 | 163 | KMT2A | HEK293T | Kidney | GSM34449 | 4586 | 103324 |
| 1138.501 | 221 | 1178.423 | 297 | 916.085 | 225 | 816.991 | 219 | KMT2D | HEK293T | Kidney | GSM34449 | 5375 | 103325 |
| 181.9954 | 65 | 221.7902 | 80 | 457.5756 | 151 | 685.3061 | 198 | SRC | MCF-7 | Breast | GSM33810 | 5885 | 103654 |
| 184.0136 | 72 | 290.4127 | 101 | 465.0429 | 167 | 731.9315 | 224 | SRC | MCF-7 | Breast | GSM33810 | 7111 | 103655 |
| 695.569 | 290 | 672.8628 | 320 | 679.633 | 497 | 673.9543 | 560 | SRC | MCF-7 | Breast | GSM33810 | 32990 | 103656 |
| 178.8111 | 61 | 174.5612 | 67 | 431.5404 | 139 | 527.4079 | 164 | SRC | MCF-7 | Breast | GSM33810 | 5269 | 103657 |
| 343.4605 | 135 | 359.4166 | 154 | 722.4664 | 283 | 797.8561 | 375 | ESR1 | MCF-7 | Breast | GSM33810 | 13650 | 103658 |
| 314.1542 | 138 | 361.6121 | 165 | 606.7482 | 278 | 753.769 | 373 | ESR1 | MCF-7 | Breast | GSM33810 | 15578 | 103659 |
| 388.9265 | 153 | 419.0735 | 177 | 742.1897 | 314 | 793.548 | 417 | ESR1 | MCF-7 | Breast | GSM33810 | 15561 | 103660 |
| 157.5931 | 72 | 171.6627 | 84 | 414.1027 | 170 | 573.9448 | 213 | ESR1 | MCF-7 | Breast | GSM33810 | 8346 | 103661 |
| 452.3788 | 77 | 659.188 | 105 | 527.9766 | 115 | 546.9862 | 124 | TRPS1 | T47D | Mammary | GSM31380 | 2647 | 103662 |
| 300.2559 | 46 | 519.2935 | 70 | 340.7137 | 67 | 343.3332 | 71 | TRPS1 | T47D | Mammary | GSM31380 | 1322 | 103663 |
| 322.4686 | 71 | 330.4454 | 79 | 153.4349 | 68 | 301.6722 | 102 | RUNX1 | None | Blood | GSM30402 | 3420 | 103694 |
| 1253.541 | 462 | 1288.709 | 594 | 959.2557 | 424 | 915.5889 | 429 | RUNX1 | None | Bone Marr | GSM30402 | 10219 | 103695 |
| 1213.714 | 519 | 1255.522 | 679 | 916.7096 | 490 | 861.7078 | 488 | RUNX1 | None | Blood | GSM30402 | 13988 | 103696 |
| 1250.428 | 492 | 1261.722 | 611 | 945.8881 | 452 | 884.6182 | 437 | RUNX1 | None | Blood | GSM30402 | 11491 | 103697 |
| 937.369 | 251 | 941.1164 | 314 | 649.3683 | 274 | 681.158 | 300 | REST | None | Liver | ENCSR893( | 13964 | 103701 |
| 904.7133 | 230 | 902.4489 | 277 | 352.5311 | 205 | 488.8852 | 255 | ATF7 | GM12878 | Blood | ENCSR014Y | 13263 | 103704 |
| 793.4529 | 320 | 783.2674 | 380 | 422.4025 | 336 | 465.9569 | 378 | ATF7 | GM12878 | Blood | ENCSR014Y | 28135 | 103705 |
| 917.4704 | 373 | 939.6515 | 483 | 753.2596 | 482 | 718.4646 | 502 | MAX | None | Liver | ENCSR521I | 24584 | 103706 |
| 54.09958 | 173 | 40.72007 | 188 | 14.79975 | 231 | 33.7247 | 300 | ATF2 | HepG2 | Liver | ENCSR047E | 67827 | 103708 |
| 64.1931 | 93 | 67.75542 | 109 | 15.70606 | 105 | 77.03285 | 178 | ATF2 | HepG2 | Liver | ENCSR047E | 24852 | 103709 |
| 1106.314 | 332 | 1154.115 | 452 | 860.0328 | 333 | 846.7758 | 366 | TAF1 | None | Liver | ENCSR016E | 10926 | 103710 |
| 383.7798 | 135 | 352.1048 | 144 | 337.6018 | 187 | 504.2787 | 241 | FOXA2 | None | Liver | ENCSR080X | 12024 | 103711 |
| 831.4725 | 413 | 792.45 | 460 | 668.6074 | 545 | 667.2104 | 619 | FOXA2 | None | Liver | ENCSR080X | 39044 | 103712 |
| 1228.763 | 276 | 1238.024 | 342 | 950.2391 | 249 | 908.6666 | 247 | GABPA | None | Liver | ENCSR350( | 5728 | 103713 |
| 1181.481 | 337 | 1202.39 | 426 | 891.7832 | 305 | 862.8135 | 317 | GABPA | None | Liver | ENCSR350( | 9097 | 103714 |
| 850.3802 | 323 | 857.7426 | 402 | 688.5021 | 415 | 690.2124 | 473 | JUND | None | Liver | ENCSR196H | 25347 | 103715 |
| 430.5996 | 130 | 541.1984 | 162 | 221.4704 | 138 | 324.4969 | 176 | JUND | None | Liver | ENCSR196H | 9539 | 103716 |
| 451.8966 | 305 | 647.462 | 413 | 290.0342 | 406 | 391.8763 | 490 | CTCF | PC-3 | Prostate | ENCSR359L | 52604 | 103717 |
| 377.4963 | 313 | 631.6328 | 456 | 272.403 | 438 | 335.5585 | 512 | CTCF | PC-3 | Prostate | ENCSR359L | 58304 | 103718 |
| 471.421 | 183 | 400.2838 | 189 | 187.8905 | 184 | 250.5028 | 223 | ATF2 | K562 | Bone Marr | ENCSR869I | 17824 | 103719 |
| 400.3633 | 193 | 406.4538 | 218 | 171.1977 | 209 | 257.2942 | 264 | ATF2 | K562 | Bone Marr | ENCSR869I | 23809 | 103720 |
| 497.5946 | 407 | 623.069 | 559 | 329.3498 | 572 | 368.3877 | 648 | CTCF | GM23338 | None | ENCSR932 | 92019 | 103721 |

| | | | | | | | | | | | | | |
|---|---|---|---|---|---|---|---|---|---|---|---|---|---|
| 632.9911 | 508 | 679.595 | 677 | 470.4349 | 690 | 456.6043 | 762 | CTCF | GM23338 | None | ENCSR932 | 104292 | 103722 |
| 277.8264 | 166 | 383.212 | 215 | 213.3129 | 229 | 266.1594 | 270 | TCF12 | K562 | Bone Marr | ENCSR744 | 23588 | 103723 |
| 171.0476 | 105 | 163.3564 | 116 | 147.1567 | 152 | 206.563 | 188 | TCF12 | K562 | Bone Marr | ENCSR744 | 15202 | 103724 |
| 273.0166 | 214 | 288.8635 | 248 | 178.4307 | 288 | 254.3948 | 354 | ATF3 | K562 | Bone Marr | ENCSR028 | 37819 | 103725 |
| 505.8618 | 315 | 555.4459 | 369 | 340.8197 | 417 | 461.5725 | 505 | ATF3 | K562 | Bone Marr | ENCSR028 | 47473 | 103726 |
| 1148.396 | 338 | 1130.571 | 390 | 868.8169 | 305 | 852.0292 | 332 | E2F1 | K562 | Bone Marr | ENCSR720 | 9186 | 103727 |
| 1117.452 | 320 | 1100.11 | 373 | 848.9152 | 301 | 840.8126 | 333 | E2F1 | K562 | Bone Marr | ENCSR720 | 9728 | 103728 |
| 381.0351 | 285 | 440.5143 | 342 | 283.0746 | 402 | 524.3003 | 552 | FOXA2 | HepG2 | Liver | ENCSR066 | 52008 | 103729 |
| 497.7808 | 325 | 562.9865 | 386 | 382.4401 | 457 | 543.4377 | 603 | FOXA2 | HepG2 | Liver | ENCSR066 | 54056 | 103730 |
| 46.22457 | 40 | 45.14744 | 45 | 71.90891 | 75 | 93.08193 | 90 | TCF12 | K562 | Bone Marr | ENCSR189 | 7322 | 103731 |
| 216.1418 | 104 | 218.4161 | 117 | 206.8845 | 153 | 219.2345 | 169 | TCF12 | K562 | Bone Marr | ENCSR189 | 11903 | 103732 |
| 1056.915 | 429 | 1073.784 | 537 | 835.4098 | 482 | 792.5329 | 490 | MAX | None | Liver | ENCSR847 | 18702 | 103733 |
| 1065.061 | 606 | 1080.508 | 751 | 814.9032 | 672 | 783.801 | 713 | MAX | None | Liver | ENCSR847 | 31151 | 103734 |
| 1148.654 | 174 | 1156.456 | 216 | 727.9927 | 173 | 661.1958 | 174 | EGR1 | None | Liver | ENCSR290 | 4564 | 103735 |
| 1113.923 | 305 | 1107.694 | 367 | 868.0759 | 311 | 827.8217 | 316 | EGR1 | None | Liver | ENCSR290 | 10159 | 103736 |
| 551.506 | 185 | 576.575 | 210 | 341.5334 | 217 | 541.6616 | 286 | ATF7 | MCF-7 | Breast | ENCSR866 | 15233 | 103737 |
| 337.0642 | 103 | 384.0146 | 122 | 166.1023 | 107 | 369.4568 | 166 | ATF7 | MCF-7 | Breast | ENCSR866 | 7500 | 103738 |
| 823.8699 | 633 | 843.4312 | 788 | 657.9277 | 847 | 658.976 | 973 | HNF4A | None | Liver | ENCSR601 | 74064 | 103739 |
| 852.6749 | 668 | 877.2424 | 838 | 678.2817 | 892 | 664.7775 | 994 | HNF4A | None | Liver | ENCSR601 | 75588 | 103740 |
| 862.7287 | 421 | 842.4666 | 486 | 682.9896 | 523 | 693.332 | 614 | JUND | None | Liver | ENCSR837 | 34452 | 103741 |
| 844.5672 | 328 | 819.8672 | 380 | 668.6154 | 412 | 703.5484 | 512 | JUND | None | Liver | ENCSR837 | 27251 | 103742 |
| 830.3774 | 626 | 832.95 | 761 | 647.6986 | 812 | 629.3459 | 891 | HNF4A | None | Liver | ENCSR445 | 69963 | 103743 |
| 823.674 | 641 | 847.793 | 805 | 658.5374 | 870 | 643.7629 | 969 | HNF4A | None | Liver | ENCSR445 | 78536 | 103744 |
| 1008.481 | 568 | 1013.613 | 693 | 754.6101 | 612 | 751.4657 | 693 | REST | None | Liver | ENCSR867 | 33384 | 103745 |
| 954.5499 | 398 | 977.7016 | 505 | 741.2986 | 448 | 741.1335 | 513 | REST | None | Liver | ENCSR867 | 23551 | 103746 |
| 888.8028 | 380 | 881.5017 | 452 | 725.3335 | 491 | 716.8839 | 551 | FOXA2 | None | Liver | ENCSR310 | 28564 | 103747 |
| 567.0685 | 197 | 608.0847 | 227 | 462.6969 | 265 | 662.7644 | 334 | FOXA2 | None | Liver | ENCSR310 | 18215 | 103748 |
| 1099.861 | 544 | 1129.598 | 696 | 855.3373 | 601 | 820.9214 | 628 | GABPA | None | Liver | ENCSR038 | 23259 | 103749 |
| 1126.371 | 398 | 1172.119 | 539 | 888.8184 | 424 | 856.9679 | 441 | GABPA | None | Liver | ENCSR038 | 13393 | 103750 |
| 601.9803 | 224 | 466.6544 | 222 | 502.1406 | 306 | 646.102 | 371 | FOXA1 | None | Liver | ENCSR324 | 22111 | 103751 |
| 477.2646 | 134 | 546.1234 | 159 | 444.2804 | 186 | 558.3365 | 222 | FOXA1 | None | Liver | ENCSR324 | 9206 | 103752 |
| 907.388 | 231 | 887.7241 | 268 | 733.4153 | 292 | 733.9699 | 325 | FOXA1 | None | Liver | ENCSR735 | 14540 | 103753 |
| 949.9204 | 320 | 944.536 | 382 | 765.9682 | 391 | 773.851 | 452 | FOXA1 | None | Liver | ENCSR735 | 18804 | 103754 |
| 221.5262 | 142 | 237.1625 | 165 | 78.41569 | 151 | 138.8627 | 200 | ATF7 | HepG2 | Liver | ENCSR545 | 21793 | 103755 |
| 873.9127 | 461 | 845.2533 | 525 | 648.436 | 515 | 646.6563 | 585 | CREB1 | MCF-7 | Breast | ENCSR897J | 37559 | 103756 |
| 984.8073 | 531 | 966.3897 | 615 | 743.9898 | 581 | 732.5887 | 647 | CREB1 | MCF-7 | Breast | ENCSR897J | 31762 | 103757 |
| 1196.55 | 184 | 1226.63 | 244 | 767.8087 | 170 | 904.2841 | 197 | E2F1 | K562 | Bone Marr | ENCSR563 | 3814 | 103758 |
| 1161.739 | 178 | 1184.806 | 233 | 658.0711 | 160 | 826.14 | 192 | E2F1 | K562 | Bone Marr | ENCSR563 | 3992 | 103759 |
| 748.0111 | 272 | 738.5033 | 303 | 680.1892 | 418 | 719.8081 | 530 | FOXA1 | MCF-7 | Breast | ENCSR126 | 25885 | 103760 |
| 591.4242 | 223 | 598.1325 | 250 | 627.6405 | 337 | 711.7531 | 446 | FOXA1 | MCF-7 | Breast | ENCSR126 | 21603 | 103761 |
| 248.7732 | 263 | 450.2585 | 373 | 192.6605 | 388 | 235.9866 | 452 | CTCF | GM23338 | None | ENCSR987 | 58930 | 103762 |
| 283.7469 | 282 | 471.3617 | 387 | 212.5931 | 407 | 228.8552 | 456 | CTCF | GM23338 | None | ENCSR987 | 59045 | 103763 |
| 798.0739 | 121 | 931.5667 | 144 | 458.1204 | 119 | 410.1265 | 119 | BRD4 | HAP-1 | None | GSM35732 | 3174 | 103819 |
| 878.5137 | 129 | 1120.262 | 162 | 481.7166 | 124 | 512.7562 | 136 | BRD4 | HAP-1 | None | GSM35732 | 3403 | 103823 |
| 171.0945 | 54 | 288.2013 | 79 | 111.7565 | 64 | 112.9443 | 69 | VDR | None | Prostate | GSM35370 | 4108 | 103922 |
| 208.2963 | 39 | 254.6819 | 48 | 55.90047 | 27 | 106.6822 | 40 | VDR | None | Prostate | GSM35370 | 1532 | 103923 |
| 1057.64 | 455 | 1019.488 | 507 | 793.356 | 455 | 780.7277 | 501 | POU2F1 | 22RV1 | Prostate | GSM35072 | 19230 | 103924 |
| 1026.229 | 370 | 998.4205 | 420 | 752.2832 | 344 | 766.6048 | 411 | POU2F1 | 22RV1 | Prostate | GSM35072 | 15656 | 103925 |
| 922.1116 | 811 | 822.7687 | 840 | 693.7019 | 1005 | 707.3245 | 1178 | FOXA1 | 22RV1 | Prostate | GSM35072 | 83619 | 103928 |
| 867.1051 | 761 | 805.6326 | 828 | 641.8753 | 928 | 667.5704 | 1117 | FOXA1 | 22RV1 | Prostate | GSM35072 | 84882 | 103929 |
| 879.8936 | 139 | 1085.031 | 171 | 455.9951 | 131 | 382.2957 | 127 | BRD4 | 402-91 | None | GSM34787 | 4033 | 103932 |
| 500.7288 | 257 | 546.5551 | 300 | 245.3633 | 297 | 281.2077 | 339 | DDIT3 | 402-91 | None | GSM30272 | 33929 | 103939 |
| 169.866 | 50 | 262.1925 | 70 | 148.5494 | 68 | 162.9058 | 76 | RUNX2 | None | None | GSM30272 | 3686 | 103944 |
| 117.9152 | 140 | 105.0026 | 154 | 80.45522 | 203 | 141.9444 | 265 | FOSL2 | None | None | GSM30272 | 34442 | 103945 |
| 1144.274 | 290 | 1178.406 | 379 | 902.8015 | 286 | 833.4668 | 266 | BRD4 | None | None | GSM30272 | 7373 | 103946 |
| 1130.365 | 240 | 1171.582 | 332 | 891.9129 | 244 | 846.6822 | 243 | BRD3 | None | None | GSM30272 | 6759 | 103947 |
| 1213.698 | 418 | 1243.407 | 545 | 918.4835 | 380 | 872.4661 | 381 | BRD2 | None | None | GSM30272 | 10341 | 103948 |
| 1209.565 | 180 | 1179.933 | 203 | 530.6755 | 136 | 599.8831 | 154 | SUPT5H | U2OS | Bone | GSM31766 | 4022 | 104036 |
| 1256.369 | 498 | 1295.012 | 647 | 951.384 | 464 | 923.4093 | 489 | SUPT5H | U2OS | Bone | GSM31766 | 12612 | 104037 |
| 296.2475 | 78 | 322.5074 | 90 | 213.077 | 94 | 141.899 | 83 | MYH11 | ME-1 | None | GSM32720 | 4651 | 104083 |
| 795.4986 | 500 | 793.2616 | 603 | 560.62 | 551 | 515.6073 | 561 | GATA2 | ME-1 | None | GSM32720 | 52875 | 104084 |
| 667.835 | 365 | 674.1328 | 450 | 303.0212 | 416 | 301.0471 | 453 | GATA2 | ME-1 | None | GSM32720 | 52023 | 104085 |
| 45.94263 | 19 | 53.75031 | 23 | 26.21424 | 22 | 158.5818 | 56 | CBFB | ME-1 | None | GSM32720 | 1955 | 104091 |
| 898.3343 | 314 | 918.8682 | 406 | 626.1314 | 327 | 567.7815 | 337 | RUNX1 | ME-1 | None | GSM32720 | 20051 | 104092 |
| 906.7838 | 345 | 919.0452 | 436 | 670.6007 | 359 | 652.8628 | 390 | RUNX1 | ME-1 | None | GSM32720 | 21853 | 104093 |
| 222.3432 | 67 | 198.9712 | 70 | 406.0224 | 131 | 517.2663 | 158 | ESR1 | MCF-7 | Breast | GSM36670 | 4808 | 104111 |
| 251.4871 | 74 | 252.2916 | 82 | 374.0933 | 130 | 602.676 | 177 | ESR1 | MCF-7 | Breast | GSM36670 | 5176 | 104112 |
| 242.6795 | 70 | 211.3912 | 72 | 414.3354 | 132 | 539.1427 | 161 | ESR1 | MCF-7 | Breast | GSM36670 | 4786 | 104113 |
| 206.0946 | 74 | 214.661 | 84 | 423.1087 | 154 | 565.565 | 190 | ESR1 | MCF-7 | Breast | GSM36670 | 6504 | 104114 |
| 213.199 | 68 | 227.7973 | 78 | 400.626 | 135 | 548.8979 | 169 | ESR1 | MCF-7 | Breast | GSM36670 | 5187 | 104115 |
| 28.32492 | 11 | 31.48477 | 13 | 91.87012 | 30 | 223.8728 | 52 | ESR1 | MCF-7 | Breast | GSM36670 | 1106 | 104116 |
| 152.8882 | 29 | 172.2152 | 34 | 64.05573 | 25 | 300.7432 | 62 | POLR3A | HEK293 | Embryonic | GSM33462 | 1120 | 104118 |

| | | | | | | | | | | | | | |
|---|---|---|---|---|---|---|---|---|---|---|---|---|---|
| 1063.825 | 691 | 1072.755 | 847 | 771.0543 | 721 | 741.2418 | 767 | JUN | MDA-MB-2 | Breast | GSM30702 | 38191 | 104129 |
| 1000.381 | 728 | 1021.807 | 900 | 717.5953 | 783 | 691.7918 | 847 | JUN | MDA-MB-2 | Breast | GSM30702 | 52490 | 104130 |
| 1084.865 | 399 | 1100.876 | 499 | 824.4245 | 398 | 789.7361 | 411 | JUN | MDA-MB-2 | Breast | GSM30702 | 15274 | 104131 |
| 961.714 | 726 | 988.0261 | 904 | 684.6721 | 794 | 652.6864 | 849 | JUN | MDA-MB-2 | Breast | GSM30702 | 59881 | 104132 |
| 915.3435 | 720 | 937.5335 | 893 | 652.4152 | 811 | 634.3118 | 893 | JUN | MDA-MB-2 | Breast | GSM30702 | 67384 | 104133 |
| 911.9015 | 742 | 937.4798 | 923 | 650.8504 | 847 | 621.7288 | 914 | JUN | MDA-MB-2 | Breast | GSM30702 | 73059 | 104134 |
| 1007.776 | 702 | 1013.631 | 850 | 720.8657 | 747 | 708.8092 | 831 | JUN | MDA-MB-2 | Breast | GSM30702 | 47488 | 104135 |
| 951.9721 | 738 | 965.3584 | 905 | 679.2761 | 819 | 645.056 | 877 | JUN | MDA-MB-2 | Breast | GSM30702 | 64013 | 104136 |
| 959.0254 | 679 | 1004.969 | 875 | 729.4089 | 801 | 686.5066 | 841 | MYOD1 | IMR90 | Lung | GSM25891 | 51968 | 104262 |
| 1068.828 | 645 | 1116.827 | 842 | 784.6977 | 670 | 752.037 | 707 | MYOD1 | IMR90 | Lung | GSM25891 | 33429 | 104263 |
| 130.6199 | 244 | 162.3327 | 301 | 108.9063 | 385 | 125.3888 | 440 | GATA6 | hESC | None | GSM33819 | 72893 | 104308 |
| 634.1454 | 331 | 674.4625 | 404 | 287.8132 | 374 | 258.0058 | 393 | NANOG | hESC | None | GSM33819 | 44040 | 104312 |
| 43.95451 | 115 | 66.27623 | 151 | 35.05517 | 184 | 54.74086 | 227 | GATA6 | hESC | None | GSM33819 | 38169 | 104318 |
| 91.30256 | 132 | 124.8285 | 169 | 86.51924 | 212 | 99.1823 | 242 | GATA6 | hESC | None | GSM33819 | 33539 | 104320 |
| 314.3654 | 184 | 362.2818 | 220 | 188.5668 | 230 | 240.1307 | 273 | JUN | hESC | None | GSM33819 | 25076 | 104321 |
| 984.331 | 328 | 977.235 | 392 | 737.1622 | 328 | 711.8178 | 346 | JUN | hESC | None | GSM29458 | 15553 | 104360 |
| 999.8398 | 192 | 1054.787 | 271 | 630.243 | 207 | 626.5421 | 220 | SMAD2 | hESC | None | GSM29458 | 7590 | 104361 |
| 13.76238 | 85 | 20.47108 | 110 | 17.13017 | 159 | 29.44654 | 198 | GATA6 | hESC | None | GSM29458 | 39447 | 104362 |
| 10.29546 | 45 | 20.85143 | 64 | 14.5847 | 86 | 21.77813 | 105 | SMAD2 | hESC | None | GSM29458 | 18762 | 104363 |
| 28.2908 | 85 | 36.99176 | 107 | 33.97235 | 153 | 57.55674 | 194 | GATA6 | hESC | None | GSM29458 | 30450 | 104364 |
| 183.2368 | 138 | 191.5081 | 159 | 107.0192 | 177 | 152.6018 | 218 | JUN | hESC | None | GSM29458 | 22992 | 104365 |
| 6.400326 | 33 | 13.8246 | 48 | 16.05316 | 74 | 23.14135 | 90 | SMAD2 | hESC | None | GSM29458 | 14848 | 104366 |
| 183.4485 | 53 | 179.3357 | 58 | 77.14311 | 51 | 164.9157 | 78 | CTCF | A375 | None | GSM36716 | 3859 | 104418 |
| 395.8661 | 122 | 641.0971 | 174 | 310.9713 | 159 | 313.4623 | 171 | RAD21 | None | Peripheral | GSM36122 | 9750 | 104439 |
| 1179.238 | 532 | 1217.063 | 692 | 902.0527 | 527 | 830.5033 | 500 | MAX | P493-6 | None | GSM35836 | 15862 | 104444 |
| 1135.348 | 197 | 1118.476 | 232 | 630.866 | 173 | 600.2692 | 179 | MAX | P493-6 | None | GSM35836 | 5324 | 104445 |
| 0.126552 | 7 | -0.064201 | 5 | -0.098632 | 9 | 0.998646 | 22 | HMGA1 | IMR90 | Lung | GSM30408 | 6374 | 104449 |
| 2.44942 | 4 | 43.62731 | 18 | 15.14146 | 15 | 158.5145 | 50 | HMGA1 | IMR90 | Lung | GSM30408 | 1694 | 104450 |
| 508.7307 | 138 | 734.1027 | 185 | 301.2752 | 153 | 342.3785 | 174 | CTCF | None | Peripheral | GSM36122 | 8748 | 104501 |
| 864.1241 | 414 | 901.8307 | 543 | 655.623 | 468 | 634.8632 | 504 | CTCF | None | Peripheral | GSM36122 | 30405 | 104502 |
| 814.219 | 240 | 877.348 | 323 | 519.1861 | 279 | 563.2525 | 311 | CTCF | None | Peripheral | GSM36122 | 16677 | 104503 |
| 454.7577 | 150 | 739.9631 | 213 | 309.5407 | 182 | 344.0438 | 205 | CTCF | None | Peripheral | GSM36122 | 11724 | 104504 |
| 867.2035 | 344 | 916.4575 | 473 | 679.3232 | 408 | 650.3705 | 429 | CTCF | None | Peripheral | GSM36122 | 24922 | 104505 |
| 878.1583 | 504 | 919.4896 | 665 | 666.9141 | 585 | 639.7466 | 622 | RAD21 | None | Peripheral | GSM36122 | 38758 | 104506 |
| 898.4926 | 355 | 945.4929 | 483 | 671.0569 | 373 | 657.5225 | 409 | RAD21 | None | Peripheral | GSM36122 | 22285 | 104507 |
| 958.3056 | 300 | 993.1681 | 399 | 726.8715 | 312 | 730.0637 | 360 | SMC3 | None | Peripheral | GSM36122 | 15521 | 104508 |
| 998.3592 | 504 | 1042.471 | 666 | 739.7588 | 515 | 707.8018 | 543 | SMC3 | None | Peripheral | GSM36122 | 25998 | 104509 |
| 982.4704 | 395 | 1062.898 | 570 | 762.7205 | 429 | 730.2949 | 449 | SMC3 | None | Peripheral | GSM36122 | 19688 | 104510 |
| 747.2339 | 479 | 791.4601 | 640 | 550.4705 | 583 | 499.9462 | 585 | IKZF1 | MM1S | MM1S mye | GSM36605 | 60263 | 104512 |
| 624.8 | 297 | 761.1803 | 430 | 459.0225 | 396 | 395.1838 | 406 | IKZF1 | ARP-1 | ARP1 myel | GSM36605 | 37427 | 104513 |
| 667.1696 | 130 | 1047.45 | 188 | 415.4192 | 140 | 530.5848 | 169 | SMARCA4 | BIN-67 | None | GSM34941 | 5286 | 104548 |
| 869.4424 | 251 | 892.8847 | 326 | 723.1659 | 332 | 706.1593 | 360 | SMARCA4 | BIN-67 | None | GSM34941 | 17274 | 104549 |
| 151.5344 | 64 | 176.7367 | 77 | 196.3422 | 107 | 274.0653 | 134 | SMARCA4 | BIN-67 | None | GSM34941 | 6478 | 104551 |
| 309.2193 | 140 | 368.3595 | 170 | 267.1942 | 197 | 297.0126 | 222 | SMARCA4 | BIN-67 | None | GSM34941 | 15760 | 104552 |
| 1021.428 | 685 | 1046.729 | 855 | 728.4402 | 693 | 703.8699 | 752 | SMARCA4 | BIN-67 | None | GSM33079 | 38489 | 104553 |
| 1163.868 | 631 | 1187.52 | 777 | 834.0372 | 552 | 809.3469 | 592 | SMARCA4 | BIN-67 | None | GSM33079 | 18294 | 104555 |
| 965.7871 | 270 | 980.8079 | 339 | 760.0759 | 297 | 738.7229 | 317 | SMARCA4 | BIN-67 | None | GSM33079 | 13066 | 104562 |
| 662.5505 | 167 | 703.7475 | 190 | 405.1439 | 186 | 421.8518 | 203 | SMARCA4 | BIN-67 | None | GSM33079 | 9695 | 104563 |
| 1103.86 | 421 | 1126.672 | 537 | 819.7257 | 388 | 781.3981 | 398 | SMARCA4 | BIN-67 | None | GSM33079 | 13803 | 104564 |
| 1057.464 | 392 | 1055.697 | 472 | 791.6723 | 376 | 740.5735 | 373 | SMARCA4 | BIN-67 | None | GSM33079 | 15273 | 104565 |
| 1008.102 | 413 | 1024.403 | 512 | 774.0128 | 433 | 717.6285 | 423 | SMARCA4 | BIN-67 | None | GSM33079 | 18513 | 104566 |
| 712.9506 | 187 | 926.3713 | 265 | 456.5099 | 215 | 472.0274 | 234 | SMARCA4 | BIN-67 | None | GSM33079 | 11710 | 104568 |
| 1032.403 | 491 | 1033.001 | 589 | 778.6597 | 507 | 730.5222 | 511 | SMARCA4 | BIN-67 | None | GSM33078 | 22951 | 104570 |
| 1133.933 | 422 | 1181.35 | 561 | 856.6836 | 400 | 814.2633 | 401 | SMARCA4 | BIN-67 | None | GSM33078 | 12892 | 104571 |
| 1122.691 | 336 | 1138.572 | 428 | 846.7621 | 314 | 799.8453 | 312 | ARID2 | BIN-67 | None | GSM33078 | 10241 | 104573 |
| 1087.007 | 292 | 1085.581 | 356 | 802.8395 | 262 | 693.9455 | 260 | DPF2 | SCCOHT-1 | None | GSM33078 | 10058 | 104575 |
| 1139.828 | 532 | 1149.552 | 653 | 832.3891 | 493 | 779.3551 | 486 | SMARCC1 | SCCOHT-1 | None | GSM33078 | 18907 | 104576 |
| 1151.432 | 430 | 1177.468 | 555 | 853.0377 | 389 | 821.9517 | 406 | SMARCC1 | BIN-67 | None | GSM33078 | 13225 | 104578 |
| 749.2311 | 311 | 735.546 | 367 | 615.2091 | 431 | 604.3875 | 477 | SMARCA4 | BIN-67 | None | GSM33078 | 33261 | 104580 |
| 814.2131 | 432 | 796.8326 | 501 | 626.8178 | 521 | 596.4421 | 553 | SMARCA4 | BIN-67 | None | GSM33078 | 39572 | 104583 |
| 723.6625 | 480 | 677.1356 | 524 | 596.5913 | 677 | 599.5212 | 781 | FOXA1 | ZR75-1 | Breast | GSM30925 | 61097 | 104593 |
| 740.5876 | 456 | 706.1753 | 510 | 620.8824 | 660 | 613.7258 | 742 | FOXA1 | ZR75-1 | Breast | GSM30925 | 57381 | 104594 |
| 745.3798 | 430 | 700.7866 | 471 | 613.0655 | 602 | 621.8073 | 701 | FOXA1 | ZR75-1 | Breast | GSM30925 | 50828 | 104595 |
| 720.9797 | 449 | 679.5509 | 497 | 596.0513 | 643 | 601.4366 | 745 | FOXA1 | ZR75-1 | Breast | GSM30925 | 58691 | 104596 |
| 750.3 | 530 | 693.7483 | 569 | 592.0737 | 708 | 597.05 | 816 | FOXA1 | ZR75-1 | Breast | GSM30925 | 67630 | 104597 |
| 748.2976 | 477 | 695.8275 | 516 | 607.7854 | 660 | 607.2642 | 754 | FOXA1 | ZR75-1 | Breast | GSM30925 | 59255 | 104598 |
| 761.0938 | 308 | 689.669 | 320 | 634.3331 | 428 | 655.337 | 516 | FOXA1 | ZR75-1 | Breast | GSM30925 | 30531 | 104599 |
| 749.9071 | 206 | 602.6562 | 204 | 580.4755 | 261 | 738.5483 | 331 | FOXA1 | ZR75-1 | Breast | GSM30925 | 13864 | 104600 |
| 757.6298 | 336 | 700.5988 | 353 | 625.1357 | 462 | 643.8891 | 554 | FOXA1 | ZR75-1 | Breast | GSM30925 | 34131 | 104601 |
| 767.3472 | 307 | 700.9072 | 319 | 629.6689 | 415 | 658.7484 | 514 | FOXA1 | ZR75-1 | Breast | GSM30925 | 29675 | 104602 |
| 763.6891 | 309 | 659.8052 | 314 | 628.2226 | 421 | 641.199 | 497 | FOXA1 | ZR75-1 | Breast | GSM30925 | 29435 | 104603 |
| 777.6789 | 336 | 707.1765 | 340 | 635.5894 | 449 | 651.1683 | 533 | FOXA1 | ZR75-1 | Breast | GSM30925 | 32017 | 104604 |

| | | | | | | | | | | | | | |
|---|---|---|---|---|---|---|---|---|---|---|---|---|---|
| 763.4886 | 443 | 694.415 | 456 | 653.687 | 651 | 665.2023 | 765 | FOXA1 | MCF-7 | Breast | GSM30925 | 49337 | 104605 |
| 756.4543 | 416 | 689.5921 | 430 | 652.9339 | 615 | 666.1041 | 728 | FOXA1 | MCF-7 | Breast | GSM30925 | 44896 | 104606 |
| 758.2897 | 417 | 703.1846 | 444 | 651.0487 | 615 | 655.3717 | 708 | FOXA1 | MCF-7 | Breast | GSM30925 | 45424 | 104607 |
| 798.8055 | 390 | 741.8027 | 413 | 696.3784 | 582 | 704.8386 | 679 | FOXA1 | MCF-7 | Breast | GSM30925 | 37348 | 104608 |
| 817.8927 | 367 | 749.2073 | 375 | 705.2147 | 534 | 718.4055 | 633 | FOXA1 | MCF-7 | Breast | GSM30925 | 32287 | 104613 |
| 806.096 | 355 | 728.6964 | 353 | 679.8177 | 497 | 693.1149 | 589 | FOXA1 | MCF-7 | Breast | GSM30924 | 31672 | 104614 |
| 828.4039 | 113 | 1191.77 | 156 | 700.0223 | 136 | 570.396 | 128 | MLLT3 | None | None | GSM30320 | 2513 | 104636 |
| 444.6338 | 61 | 571.752 | 77 | 283.6881 | 62 | 294.725 | 67 | MLLT3 | None | None | GSM30320 | 1280 | 104638 |
| 316.9193 | 53 | 264.4693 | 52 | 103.9363 | 40 | 78.2866 | 37 | NR3C1 | JHU-011 | Peripheral | GSM29429 | 1871 | 104675 |
| 788.5464 | 344 | 820.8474 | 454 | 617.1428 | 433 | 592.9861 | 462 | TP63 | None | Tongue | GSM27905 | 32842 | 104722 |
| 916.8865 | 289 | 924.7401 | 359 | 688.9871 | 314 | 606.4228 | 318 | SOX2 | None | Tongue | GSM27905 | 17295 | 104723 |
| 273.1174 | 63 | 236.6931 | 64 | 157.5082 | 67 | 141.8325 | 68 | PITX1 | None | Tongue | GSM27905 | 3273 | 104724 |
| 890.0423 | 444 | 908.6417 | 559 | 650.4376 | 479 | 641.5879 | 534 | SMAD3 | PC-3 | Prostate | GSM31455 | 34753 | 104796 |
| 5.241224 | 60 | 1.653778 | 59 | 0.331047 | 82 | 6.554113 | 131 | CBX8 | PC-3 | Prostate | GSM31455 | 36464 | 104797 |
| 6.342982 | 20 | 1.752814 | 17 | 1.803791 | 27 | 3.271043 | 34 | CBX8 | PC-3 | Prostate | GSM31455 | 8433 | 104798 |
| 977.4781 | 561 | 975.2674 | 671 | 705.0975 | 576 | 683.6534 | 625 | SMAD3 | PC-3 | Prostate | GSM31455 | 36367 | 104799 |
| 554.6724 | 125 | 697.6139 | 156 | 371.3956 | 144 | 373.0693 | 154 | HIF1A | PC-3 | Prostate | GSM31455 | 6615 | 104800 |
| 991.398 | 539 | 1002.536 | 666 | 717.5218 | 555 | 707.8166 | 616 | HIF1A | PC-3 | Prostate | GSM31455 | 33488 | 104801 |
| 133.3715 | 66 | 121.7024 | 71 | 114.391 | 94 | 207.73 | 131 | EPAS1 | PC-3 | Prostate | GSM31455 | 8212 | 104802 |
| 898.3961 | 424 | 925.9219 | 546 | 663.0693 | 444 | 653.2938 | 495 | WDR5 | SMMC-772 | None | GSM36193 | 27177 | 104843 |
| 9.266043 | 106 | 11.3359 | 131 | 4.021457 | 173 | 4.361714 | 196 | PAX3 | None | differentia | GSM35651 | 59121 | 104895 |
| 186.0017 | 30 | 257.9064 | 40 | 124.0663 | 33 | 234.0225 | 50 | LMNB1 | None | Adipose | GSM33947 | 1059 | 104912 |
| 161.5275 | 84 | 195.6475 | 103 | 128.9612 | 116 | 186.9537 | 146 | STAT1 | HNSCC cell | None | GSM20696 | 10541 | 105143 |
| 142.5468 | 70 | 164.9134 | 84 | 156.8603 | 110 | 197.9048 | 131 | STAT1 | HNSCC cell | None | GSM20696 | 8266 | 105144 |
| 1139.229 | 176 | 1139.211 | 213 | 617.9522 | 160 | 612.6385 | 169 | STAT3 | HNSCC cell | None | GSM20696 | 4441 | 105145 |
| 462.6485 | 148 | 736.1207 | 208 | 338.3372 | 185 | 378.7968 | 209 | STAT3 | HNSCC cell | None | GSM20696 | 11302 | 105146 |
| -0.045768 | 125 | -3.030475 | 114 | -0.045514 | 267 | 0.049507 | 339 | CDX2 | None | Endoderm | GSM31755 | 143477 | 105170 |
| 89.63837 | 23 | 81.0504 | 24 | 29.78318 | 19 | 126.7469 | 42 | KDM6A | None | None | GSM28902 | 1406 | 105185 |
| 111.8056 | 47 | 127.4227 | 56 | 41.50056 | 46 | 71.06196 | 62 | KDM6A | None | None | GSM28902 | 4904 | 105189 |
| 1031.074 | 221 | 1033.913 | 273 | 614.4083 | 213 | 483.154 | 202 | TP53BP1 | None | None | GSM28902 | 8534 | 105195 |
| 999.7342 | 572 | 1019.281 | 719 | 717.581 | 563 | 660.1775 | 559 | TP53BP1 | None | None | GSM28902 | 29827 | 105196 |
| 1230.493 | 284 | 1262.334 | 372 | 973.2359 | 270 | 930.5882 | 268 | TP53BP1 | None | None | GSM28902 | 5679 | 105197 |
| 1156.245 | 518 | 1190.257 | 669 | 857.0906 | 489 | 815.8442 | 503 | TP53BP1 | None | None | GSM28902 | 17623 | 105198 |
| 143.7062 | 37 | 236.3505 | 54 | 44.13484 | 30 | 139.9172 | 56 | TP53 | None | Foreskin | GSM33785 | 2383 | 105304 |
| 37.20371 | 50 | 55.86846 | 67 | 42.69292 | 87 | 69.51282 | 112 | TP63 | MCF-10A | Breast | GSM33785 | 13284 | 105310 |
| 432.2588 | 137 | 619.8241 | 183 | 224.9701 | 149 | 211.2644 | 156 | TP53 | MCF-10A | Breast | GSM33785 | 11321 | 105311 |
| 497.4567 | 68 | 616.3877 | 84 | 155.7548 | 47 | 180.2106 | 54 | SMARCA4 | parental T( | None | GSM29913 | 1531 | 105317 |
| 288.0934 | 102 | 365.9482 | 128 | 146.2305 | 111 | 250.2206 | 151 | EP300 | IMR90 | Lung | GSM28304 | 9136 | 105544 |
| 111.9324 | 40 | 139.9053 | 50 | 61.11321 | 45 | 189.8914 | 82 | EP300 | IMR90 | Lung | GSM28304 | 3726 | 105545 |
| 109.076 | 80 | 70.92787 | 76 | 69.8778 | 106 | 57.76823 | 108 | EP300 | IMR90 | Lung | GSM28304 | 13369 | 105546 |
| 144.9173 | 37 | 246.0693 | 55 | 113.6572 | 47 | 200.3876 | 67 | EP300 | IMR90 | Lung | GSM28304 | 2348 | 105547 |
| 387.1063 | 94 | 424.8319 | 109 | 271.2312 | 112 | 301.9029 | 126 | CREBBP | IMR90 | Lung | GSM28304 | 5512 | 105548 |
| 424.4708 | 160 | 360.4348 | 165 | 276.5961 | 196 | 224.9903 | 194 | CREBBP | IMR90 | Lung | GSM28304 | 15367 | 105550 |
| 279.4787 | 61 | 332.7097 | 74 | 231.1255 | 78 | 258.9817 | 88 | CREBBP | IMR90 | Lung | GSM28304 | 3128 | 105551 |
| 14.8138 | 26 | 19.07588 | 33 | 50.89763 | 68 | 51.25765 | 74 | SMAD3 | H9 | Embryo | GSM72758 | 8320 | 1084 |
| 971.4725 | 211 | 986.942 | 272 | 582.5896 | 223 | 541.932 | 230 | SMAD3 | H9 | Embryo | GSM72755 | 10474 | 1085 |
| 58.19427 | 58 | 51.06026 | 63 | 76.65656 | 104 | 57.46463 | 102 | SMAD4 | H9 | Embryo | GSM72758 | 12803 | 1086 |
| 997.2083 | 288 | 1004.834 | 357 | 763.9124 | 302 | 665.1618 | 293 | SMAD4 | H9 | Embryo | GSM72756 | 14067 | 1087 |
| 152.469 | 61 | 161.3348 | 70 | 275.1289 | 120 | 415.9356 | 157 | TCF4 | LS174T | Colon | GSM46012 | 6134 | 11 |
| 95.85547 | 32 | 82.76294 | 33 | 27.83282 | 27 | 81.89897 | 47 | AFF4 | HCT-116 | Colon | GSM74982 | 2807 | 1140 |
| 305.3586 | 49 | 301.3704 | 53 | 77.57859 | 32 | 186.3764 | 54 | ELL2 | HCT-116 | Colon | GSM74982 | 1606 | 1143 |
| 738.7423 | 91 | 1069.271 | 124 | 354.2553 | 79 | 639.5363 | 117 | ELK4 | HEK293 | Embryonic | GSM70036 | 1879 | 1173 |
| 743.1801 | 125 | 853.8888 | 148 | 417.4923 | 126 | 321.1652 | 117 | TRIM28 | HEK293 | Embryonic | GSM70035 | 4456 | 1174 |
| 803.4277 | 304 | 809.0586 | 379 | 544.5928 | 359 | 606.6304 | 408 | TRIM28 | HEK293 | Embryonic | GSM70035 | 28287 | 1175 |
| 222.9355 | 42 | 214.827 | 45 | 115.3205 | 41 | 155.3075 | 51 | TRIM28 | HEK293 | Embryonic | GSM70035 | 1742 | 1177 |
| 550.2493 | 213 | 644.2609 | 256 | 395.4245 | 273 | 450.9558 | 311 | TRIM28 | HEK293 | Embryonic | GSM70035 | 21114 | 1178 |
| 519.393 | 204 | 633.4245 | 250 | 287.6477 | 235 | 238.7291 | 236 | TRIM28 | HEK293 | Embryonic | GSM70035 | 20337 | 1194 |
| 252.072 | 81 | 315.6161 | 101 | 129.4896 | 87 | 217.0094 | 118 | TRIM28 | HEK293 | Embryonic | GSM70035 | 6530 | 1195 |
| 358.5572 | 129 | 428.3554 | 157 | 210.3491 | 151 | 256.7294 | 177 | CTCF | HeLa | Cervix | GSM32589 | 12286 | 1203 |
| 1105.315 | 204 | 1128.339 | 268 | 805.5827 | 211 | 738.4707 | 214 | DDX5 | HeLa | Cervix | GSM59399 | 6351 | 1208 |
| 1218.753 | 371 | 1255.992 | 485 | 928.3103 | 339 | 903.3411 | 357 | E2F1 | HeLa | Cervix | GSM55846 | 9086 | 1212 |
| 855.7071 | 118 | 1184.302 | 161 | 476.9437 | 114 | 508.435 | 125 | ELK1 | HeLa-S3 | Cervix | GSM60816 | 2975 | 1213 |
| 1164.062 | 197 | 1181.072 | 253 | 789.1931 | 188 | 765.6693 | 196 | ELK4 | HeLa-S3 | Cervix | GSM60816 | 4970 | 1214 |
| 1123.724 | 145 | 1221.665 | 199 | 642.506 | 141 | 753.8981 | 163 | ELK4 | HeLa-S3 | Cervix | GSM60816 | 3327 | 1215 |
| 1082.249 | 240 | 1055.148 | 272 | 829.508 | 242 | 783.1305 | 250 | GATAD1 | HeLa | Cervix | GSM56616 | 8420 | 1218 |
| 1167.258 | 491 | 1190.908 | 622 | 891.1397 | 484 | 859.3835 | 509 | GATAD1 | HeLa | Cervix | GSM56615 | 15834 | 1219 |
| 47.5598 | 14 | 44.5382 | 15 | 28.84972 | 16 | 134.4088 | 38 | GLYR1 | HeLa | Cervix | GSM56616 | 1107 | 1221 |
| 523.7276 | 74 | 653.6641 | 92 | 212.1029 | 60 | 273.0645 | 73 | TRRAP | HeLa | Cervix | GSM56616 | 1927 | 1222 |
| 1042.879 | 171 | 1049.853 | 198 | 586.7265 | 172 | 587.852 | 183 | C17orf49 | HeLa | Cervix | GSM56616 | 5851 | 1223 |
| 267.7747 | 52 | 327.1082 | 64 | 257.1553 | 71 | 189.2931 | 64 | GLYR1 | HeLa | Cervix | GSM56616 | 2255 | 1225 |
| 1088.127 | 259 | 1082.906 | 314 | 864.1403 | 275 | 803.2163 | 263 | TRRAP | HeLa | Cervix | GSM56615 | 8977 | 1226 |
| 1015.976 | 195 | 1041.192 | 255 | 634.2483 | 207 | 586.4721 | 212 | C17orf49 | HeLa | Cervix | GSM56615 | 8116 | 1227 |

| | | | | | | | | | | | | | |
|---|---|---|---|---|---|---|---|---|---|---|---|---|---|
| 970.9239 | 139 | 1180.959 | 200 | 650.2801 | 150 | 712.9137 | 167 | NR2C2 | HeLa-S3 | Cervix | GSM60816 | 3798 | 1298 |
| 840.2491 | 116 | 1178.79 | 158 | 521.3191 | 119 | 470.5183 | 119 | NR2C2 | HeLa-S3 | Cervix | GSM60815 | 2938 | 1299 |
| 430.2723 | 158 | 429.6644 | 176 | 388.0437 | 223 | 545.7645 | 279 | NR3C1 | HeLa | Cervix | GSM60466 | 15115 | 1300 |
| 283.3728 | 44 | 218.4461 | 41 | 107.0706 | 35 | 185.1483 | 50 | NR3C1 | HeLa | Cervix | GSM60466 | 1367 | 1301 |
| 393.3157 | 65 | 262.6251 | 56 | 134.5336 | 50 | 227.3583 | 70 | NR3C1 | HeLa | Cervix | GSM60465 | 2217 | 1302 |
| 181.7791 | 68 | 159.3037 | 71 | 182.5068 | 101 | 292.4364 | 135 | NR3C1 | HeLa | Cervix | GSM60465 | 6463 | 1304 |
| 279.726 | 91 | 188.6289 | 83 | 266.4896 | 130 | 471.4371 | 183 | NR3C1 | HeLa | Cervix | GSM60464 | 7453 | 1306 |
| 1152.491 | 370 | 1163.538 | 469 | 869.5599 | 349 | 830.5821 | 352 | PHF8 | HeLa | Cervix | GSM56616 | 11095 | 1308 |
| 1226.254 | 521 | 1249.659 | 657 | 923.1996 | 483 | 886.1387 | 501 | PHF8 | HeLa | Cervix | GSM56615 | 14166 | 1309 |
| 1313.341 | 509 | 1360.811 | 669 | 995.2829 | 463 | 970.6412 | 492 | PHF8 | HeLa | Cervix | GSM55847 | 10843 | 1312 |
| 924.1122 | 109 | 1054.368 | 128 | 541.8057 | 106 | 603.9785 | 119 | PHF8 | HeLa | Cervix | GSM53196 | 2172 | 1313 |
| 1337.372 | 685 | 1398.804 | 900 | 990.7681 | 629 | 955.674 | 655 | PHF8 | HeLa | Cervix | GSM52038 | 16007 | 1314 |
| 507.3134 | 73 | 579.1344 | 86 | 207.9281 | 60 | 235.0362 | 68 | RELA | HeLa | Cervix | GSM60466 | 2008 | 1333 |
| 330.8262 | 129 | 360.169 | 150 | 305.2552 | 185 | 440.1162 | 234 | RELA | HeLa | Cervix | GSM60465 | 12988 | 1335 |
| 326.9237 | 140 | 346.6737 | 161 | 289.2119 | 199 | 464.1652 | 263 | RELA | HeLa | Cervix | GSM60465 | 15506 | 1336 |
| 241.1696 | 36 | 270.2717 | 42 | 67.00019 | 24 | 120.492 | 35 | RELA | HeLa | Cervix | GSM60465 | 1051 | 1338 |
| 187.8345 | 52 | 357.9184 | 82 | 131.5223 | 63 | 146.6589 | 71 | RELA | HeLa | Cervix | GSM60465 | 3574 | 1339 |
| 262.2336 | 58 | 249.4084 | 62 | 143.2251 | 60 | 228.2483 | 81 | SMARCA4 | HeLa | Cervix | GSM60139 | 2989 | 1344 |
| 1132.524 | 492 | 1153.203 | 620 | 844.3568 | 474 | 824.805 | 510 | SMARCB1 | HeLa | Cervix | GSM60139 | 17387 | 1345 |
| 639.2508 | 188 | 770.2008 | 229 | 358.3287 | 208 | 456.8453 | 249 | SMARCC1 | HeLa | Cervix | GSM60140 | 13884 | 1346 |
| 352.9367 | 90 | 577.2256 | 130 | 161.934 | 88 | 283.6604 | 123 | SMARCC2 | HeLa | Cervix | GSM60140 | 5524 | 1347 |
| 260.9248 | 83 | 254.6904 | 91 | 229.038 | 114 | 245.184 | 126 | STAT1 | HeLa-S3 | Cervix | GSM38550 | 6691 | 1356 |
| 241.9906 | 81 | 236.5519 | 89 | 181.439 | 104 | 240.6875 | 127 | STAT1 | HeLa-S3 | Cervix | GSM38550 | 6901 | 1357 |
| 62.05686 | 32 | 62.0516 | 36 | 76.79305 | 54 | 127.5007 | 73 | STAT1 | HeLa-S3 | Cervix | GSM38550 | 4253 | 1358 |
| 202.6752 | 84 | 198.7188 | 93 | 149.183 | 110 | 222.9614 | 141 | STAT1 | HeLa-S3 | Cervix | GSM38550 | 8958 | 1359 |
| 12.3353 | 7 | 6.904639 | 6 | 17.21646 | 13 | 32.88691 | 19 | STAT1 | HeLa-S3 | Cervix | GSM38550 | 1184 | 1360 |
| 18.73584 | 9 | 6.351604 | 6 | 35.46903 | 19 | 41.52255 | 22 | STAT1 | HeLa-S3 | Cervix | GSM38550 | 1257 | 1361 |
| 966.4152 | 566 | 982.8367 | 709 | 727.4803 | 638 | 721.0208 | 720 | STAT1 | HeLa-S3 | Cervix | GSM32073 | 39031 | 1362 |
| 282.0951 | 267 | 381.5039 | 341 | 240.0535 | 408 | 291.3607 | 474 | CEBPA | HepG2 | Liver | GSM61463 | 61146 | 1374 |
| 250.1214 | 244 | 430.8326 | 342 | 260.5961 | 399 | 296.3189 | 454 | CEBPA | HepG2 | Liver | GSM61463 | 56502 | 1375 |
| 666.509 | 516 | 716.1421 | 693 | 523.7047 | 727 | 512.5299 | 808 | CTCF | HepG2 | Liver | GSM61463 | 92752 | 1380 |
| 432.4662 | 243 | 693.1505 | 339 | 321.6581 | 334 | 306.0522 | 356 | CTCF | HepG2 | Liver | GSM61463 | 36429 | 1381 |
| 689.5433 | 412 | 732.5879 | 556 | 486.2291 | 524 | 504.2521 | 586 | CTCF | HepG2 | Liver | GSM61463 | 62092 | 1382 |
| 689.4993 | 405 | 726.3503 | 537 | 503.5689 | 522 | 517.7622 | 593 | CTCF | HepG2 | Liver | GSM61463 | 60432 | 1383 |
| 759.5559 | 390 | 806.2537 | 530 | 612.7326 | 533 | 620.9657 | 620 | HNF4A | HepG2 | Liver | GSM61463 | 45556 | 1447 |
| 56.25302 | 44 | 116.7872 | 69 | 58.06001 | 71 | 123.8107 | 104 | HNF4A | HepG2 | Liver | GSM46986 | 8239 | 1448 |
| 89.33787 | 66 | 154.2945 | 95 | 119.6271 | 117 | 197.3208 | 155 | HNF4A | HepG2 | Liver | GSM46986 | 11732 | 1449 |
| 804.5386 | 107 | 1144.211 | 144 | 498.7607 | 109 | 528.388 | 119 | NR2C2 | HepG2 | Liver | GSM60815 | 2537 | 1463 |
| 884.5518 | 118 | 1204.218 | 162 | 504.8434 | 115 | 568.9914 | 130 | NR2C2 | HepG2 | Liver | GSM60815 | 2839 | 1464 |
| 777.5538 | 395 | 801.7739 | 511 | 580.6989 | 476 | 565.2934 | 521 | RAD21 | HepG2 | Liver | GSM61462 | 43961 | 1470 |
| 446.767 | 238 | 646.3992 | 315 | 299.5362 | 313 | 337.8802 | 355 | RAD21 | HepG2 | Liver | GSM61462 | 34268 | 1471 |
| 828.8189 | 584 | 851.2308 | 737 | 614.9562 | 707 | 593.5862 | 769 | STAG1 | HepG2 | Liver | GSM61462 | 66534 | 1492 |
| 794.0486 | 513 | 819.6609 | 656 | 596.4752 | 635 | 578.2359 | 696 | STAG1 | HepG2 | Liver | GSM61462 | 61020 | 1493 |
| 1244.672 | 326 | 1265.009 | 416 | 958.1287 | 298 | 953.2446 | 335 | PHF8 | Hs68 | Foreskin | GSM52038 | 7187 | 1628 |
| 1113.865 | 139 | 1209.977 | 176 | 596.845 | 130 | 791.3559 | 161 | PHF8 | Hs68 | Foreskin | GSM52038 | 3092 | 1629 |
| 142.2091 | 200 | 142.2544 | 230 | 104.7453 | 301 | 198.2132 | 398 | FOS | None | Umbilical V | GSM73070 | 54088 | 1740 |
| 771.8073 | 284 | 795.4238 | 373 | 447.2103 | 335 | 606.2246 | 414 | FOS | None | Umbilical V | GSM73070 | 28990 | 1741 |
| 332.6051 | 210 | 315.038 | 232 | 269.6339 | 300 | 310.5895 | 343 | GATA2 | None | Umbilical V | GSM73070 | 33171 | 1742 |
| 103.2187 | 113 | 74.86258 | 115 | 114.0025 | 189 | 117.3908 | 208 | GATA2 | None | Umbilical V | GSM73070 | 25632 | 1743 |
| 565.3918 | 230 | 647.9613 | 274 | 410.4587 | 300 | 373.9209 | 312 | RBPJ | IB4 | Blood | GSM72985 | 25094 | 1792 |
| 43.60134 | 25 | 20.08276 | 20 | 33.82377 | 35 | 84.53758 | 56 | RBPJ | IB4 | Blood | GSM72985 | 3706 | 1793 |
| 572.5838 | 170 | 727.4206 | 213 | 246.0822 | 168 | 249.7809 | 182 | SND1 | IB4 | Blood | GSM72985 | 12894 | 1794 |
| 89.26191 | 33 | 44.07798 | 26 | 13.38649 | 22 | 54.10914 | 42 | SND1 | IB4 | Blood | GSM72985 | 3165 | 1795 |
| 1053.299 | 158 | 1130.385 | 206 | 628.6965 | 162 | 589.5183 | 166 | RB1 | IMR90 | Lung | GSM49749 | 4806 | 1870 |
| 79.31606 | 28 | 88.38085 | 33 | 42.20411 | 31 | 88.91622 | 47 | RB1 | IMR90 | Lung | GSM49749 | 2580 | 1871 |
| 626.4702 | 102 | 922.7261 | 140 | 364.7996 | 104 | 467.3091 | 126 | RB1 | IMR90 | Lung | GSM49749 | 3430 | 1872 |
| 391.784 | 74 | 591.4345 | 103 | 227.2745 | 77 | 384.5047 | 108 | RB1 | IMR90 | Lung | GSM49748 | 3082 | 1873 |
| 1189.973 | 274 | 1196.956 | 346 | 911.6217 | 252 | 909.0047 | 284 | RBL2 | IMR90 | Lung | GSM49749 | 6918 | 1874 |
| 791.5618 | 119 | 934.4919 | 143 | 322.9605 | 99 | 495.206 | 132 | RBL2 | IMR90 | Lung | GSM49749 | 3526 | 1876 |
| 1170.508 | 275 | 1187.921 | 351 | 914.169 | 265 | 876.7317 | 271 | RBL2 | IMR90 | Lung | GSM49749 | 7296 | 1878 |
| 684.4824 | 103 | 1013.615 | 142 | 293.9831 | 88 | 461.562 | 119 | RBL2 | IMR90 | Lung | GSM49749 | 3069 | 1879 |
| 1161.132 | 205 | 1201.702 | 283 | 845.7714 | 200 | 900.2515 | 220 | RBL2 | IMR90 | Lung | GSM49749 | 5336 | 1880 |
| 67.76953 | 20 | 55.86252 | 20 | 27.81235 | 19 | 70.88526 | 32 | SNAPC2 | IMR90 | Lung | GSM45459 | 1550 | 1881 |
| 934.9122 | 278 | 986.1174 | 387 | 745.1519 | 327 | 689.7405 | 321 | CREBBP | Jurkat E6-1 | Blood | GSM44952 | 15712 | 1935 |
| 427.014 | 151 | 569.2668 | 194 | 213.6178 | 165 | 272.6887 | 197 | CTCF | Jurkat | Blood | GSM32589 | 14207 | 1936 |
| 1103.972 | 435 | 1140.588 | 573 | 843.9336 | 440 | 801.4677 | 450 | ETS1 | Jurkat E6-1 | Blood | GSM44952 | 16359 | 1938 |
| 7.688088 | 6 | 20.16742 | 11 | 7.878833 | 10 | 81.3823 | 32 | RUNX1 | Jurkat E6-1 | Blood | GSM44952 | 1326 | 1955 |
| 8.021406 | 9 | 21.45882 | 16 | 6.371939 | 14 | 67.20021 | 40 | TAL1 | Jurkat | Blood | GSM61400 | 2446 | 1957 |
| 1088.035 | 678 | 1087.18 | 818 | 772.4741 | 676 | 727.5729 | 696 | BCL6 | JURL_MK1 | Bone Marr | GSM64042 | 37164 | 1958 |
| 1138.756 | 578 | 1184.091 | 764 | 854.1556 | 586 | 806.3529 | 596 | E2F6 | K562 | Bone Marr | GSM70186 | 23001 | 1978 |
| 141.7186 | 29 | 218.0126 | 41 | 69.6871 | 28 | 100.0505 | 36 | EGR1 | K562 | Bone Marr | GSM54072 | 1382 | 1982 |
| 658.4518 | 412 | 642.4928 | 484 | 476.2493 | 550 | 496.0686 | 622 | FOS | K562 | Bone Marr | GSM61033 | 68724 | 1990 |

| | | | | | | | | | | | | | |
|---|---|---|---|---|---|---|---|---|---|---|---|---|---|
| 583.8196 | 299 | 667.0931 | 358 | 400.9765 | 396 | 443.4174 | 446 | FOS | K562 | Bone Marr | GSM61033 | 42682 | 1991 |
| 960.6909 | 368 | 973.0517 | 462 | 708.3106 | 376 | 723.4515 | 447 | FOS | K562 | Bone Marr | GSM48742 | 21739 | 1992 |
| 0 | 0 | 0.804889 | 2 | 0.101847 | 2 | -0.011416 | 1 | GAPDH | HeLa | Cervix | GSM44802 | 1027 | 2 |
| 26.61561 | 48 | 51.00496 | 70 | 54.97599 | 103 | 63.74996 | 118 | GATA1 | K562 | Bone Marr | GSM61033 | 15114 | 2000 |
| 219.9558 | 84 | 325.3673 | 114 | 243.8092 | 131 | 269.4026 | 147 | GATA1 | K562 | Bone Marr | GSM46764 | 8276 | 2001 |
| 216.441 | 132 | 277.3244 | 166 | 260.5566 | 220 | 304.487 | 253 | GATA2 | K562 | Bone Marr | GSM46764 | 20178 | 2002 |
| 940.4466 | 532 | 964.837 | 672 | 722.1098 | 622 | 688.5065 | 655 | JUN | K562 | Bone Marr | GSM48742 | 39481 | 2078 |
| 1097.336 | 560 | 1129.189 | 716 | 818.8304 | 567 | 784.0545 | 594 | MYC | K562 | Bone Marr | GSM48742 | 25049 | 2090 |
| 1107.405 | 533 | 1129.518 | 677 | 821.4078 | 528 | 782.4403 | 544 | NFYA | K562 | Bone Marr | GSM64858 | 22541 | 2094 |
| 1061.524 | 352 | 1094.285 | 463 | 776.7986 | 327 | 769.5175 | 365 | NFYB | K562 | Bone Marr | GSM64858 | 14322 | 2095 |
| 1095.885 | 498 | 1104.171 | 611 | 825.2645 | 505 | 783.535 | 516 | PRAME | K562 | Bone Marr | GSM64858 | 21168 | 2110 |
| 528.9646 | 203 | 738.4401 | 266 | 385.5091 | 264 | 410.2611 | 292 | RUNX1 | K562 | Bone Marr | GSM61033 | 21196 | 2121 |
| 168.3807 | 58 | 291.9093 | 86 | 86.0973 | 63 | 185.9544 | 96 | RUNX1 | K562 | Bone Marr | GSM61033 | 5131 | 2122 |
| 271.5178 | 67 | 350.2672 | 85 | 145.3835 | 70 | 180.099 | 83 | SETDB1 | K562 | Bone Marr | GSM60749 | 3970 | 2123 |
| 715.3025 | 134 | 926.1108 | 170 | 399.7245 | 137 | 322.1388 | 131 | SETDB1 | K562 | Bone Marr | GSM60748 | 5527 | 2124 |
| 4.441653 | 7 | 4.194743 | 8 | 0.550988 | 7 | 35.91832 | 30 | TRIM28 | K562 | Bone Marr | GSM60749 | 2389 | 2145 |
| 4.78596 | 8 | 7.291571 | 11 | -0.002368 | 4 | 22.86583 | 27 | TRIM28 | K562 | Bone Marr | GSM60749 | 2684 | 2146 |
| 181.4083 | 51 | 197.2715 | 59 | 111.5589 | 58 | 111.0874 | 62 | ZNF263 | K562 | Bone Marr | GSM47665 | 3501 | 2157 |
| 283.1032 | 64 | 287.6791 | 71 | 126.1279 | 60 | 137.938 | 67 | ZNF263 | K562 | Bone Marr | GSM47665 | 3334 | 2158 |
| 684.21 | 215 | 818.0033 | 265 | 771.2991 | 384 | 854.8918 | 547 | AR | LNCaP | Prostate | GSM75966 | 17982 | 2171 |
| 760.4334 | 226 | 866.7878 | 303 | 816.0293 | 431 | 869.2271 | 564 | AR | LNCaP | Prostate | GSM75965 | 17844 | 2172 |
| 1055.305 | 243 | 1043.557 | 285 | 906.4767 | 319 | 1004.402 | 471 | AR | LNCaP | Prostate | GSM75965 | 9294 | 2173 |
| 1133.543 | 404 | 1100.214 | 450 | 948.8685 | 498 | 1022.979 | 676 | AR | LNCaP | Prostate | GSM75965 | 13832 | 2174 |
| 853.646 | 430 | 811.8976 | 475 | 675.6735 | 545 | 753.9177 | 753 | AR | LNCaP | Prostate | GSM35364 | 38671 | 2181 |
| 1242.264 | 249 | 1132.943 | 226 | 1069.27 | 308 | 1151.183 | 431 | FOXA1 | LNCaP | Prostate | GSM75966 | 5224 | 2189 |
| 1201.323 | 261 | 1135.592 | 268 | 1061.359 | 355 | 1134.524 | 483 | FOXA1 | LNCaP | Prostate | GSM75966 | 6347 | 2190 |
| 1088.409 | 578 | 982.7916 | 566 | 911.4414 | 752 | 957.5 | 949 | FOXA1 | LNCaP | Prostate | GSM75966 | 28312 | 2191 |
| 1103.193 | 522 | 1001.539 | 503 | 909.3843 | 643 | 972.3803 | 848 | FOXA1 | LNCaP | Prostate | GSM75966 | 22517 | 2192 |
| 1014.249 | 566 | 931.4676 | 581 | 825.3073 | 719 | 877.3457 | 915 | FOXA1 | LNCaP | Prostate | GSM35363 | 35088 | 2195 |
| 251.206 | 110 | 343.3309 | 143 | 346.6641 | 191 | 726.9796 | 289 | NR3C1 | LNCaP | Prostate | GSM75967 | 12480 | 2242 |
| 1035.805 | 212 | 1002.88 | 236 | 886.9999 | 279 | 978.0682 | 410 | NR3C1 | LNCaP | Prostate | GSM75966 | 8427 | 2243 |
| 761.3808 | 472 | 806.8971 | 629 | 559.9747 | 571 | 550.5054 | 636 | CTCF | MCF-7 | Breast | GSM61461 | 59488 | 2254 |
| 730.1186 | 365 | 765.1657 | 486 | 542.7959 | 463 | 524.8779 | 496 | CTCF | MCF-7 | Breast | GSM61461 | 46337 | 2255 |
| 1218.506 | 790 | 1247.592 | 976 | 945.0249 | 894 | 918.349 | 974 | E2F1 | MCF-7 | Breast | GSM69998 | 31805 | 2278 |
| 1176.819 | 817 | 1192.984 | 994 | 901.6666 | 932 | 873.5087 | 1014 | E2F1 | MCF-7 | Breast | GSM69998 | 38292 | 2279 |
| 1254.069 | 694 | 1285.411 | 875 | 935.9674 | 685 | 901.4219 | 721 | E2F1 | MCF-7 | Breast | GSM69998 | 21534 | 2280 |
| 106.202 | 27 | 127.2767 | 33 | 106.941 | 39 | 130.2168 | 46 | E2F1 | MCF-7 | Breast | GSM69998 | 1717 | 2281 |
| 1115.365 | 451 | 1130.84 | 567 | 862.0747 | 470 | 837.8233 | 501 | E2F1 | MCF-7 | Breast | GSM69998 | 16248 | 2282 |
| 1175.905 | 660 | 1190.962 | 810 | 888.789 | 677 | 855.5289 | 721 | E2F1 | MCF-7 | Breast | GSM69998 | 23841 | 2283 |
| 106.0261 | 24 | 110.1318 | 27 | 190.2154 | 47 | 427.9181 | 79 | EP300 | MCF-7 | Breast | GSM72042 | 1302 | 2285 |
| 831.5897 | 270 | 813.9679 | 317 | 769.4795 | 459 | 808.5772 | 576 | ESR1 | MCF-7 | Breast | GSM61461 | 22508 | 2294 |
| 668.0716 | 383 | 666.0821 | 464 | 639.4474 | 694 | 699.6215 | 899 | ESR1 | MCF-7 | Breast | GSM61461 | 59589 | 2295 |
| 158.862 | 190 | 290.2778 | 272 | 121.9701 | 289 | 173.7232 | 352 | ESR1 | MCF-7 | Breast | GSM59460 | 50301 | 2296 |
| 421.675 | 252 | 584.9496 | 327 | 472.6061 | 409 | 567.1713 | 487 | ESR1 | MCF-7 | Breast | GSM59460 | 41423 | 2297 |
| 322.7335 | 140 | 427.5223 | 179 | 651.2001 | 288 | 769.5446 | 390 | ESR1 | MCF-7 | Breast | GSM58923 | 15734 | 2298 |
| 70.33299 | 17 | 41.68089 | 14 | 35.10586 | 17 | 298.7114 | 58 | ESR1 | MCF-7 | Breast | GSM58923 | 1029 | 2299 |
| 160.0748 | 61 | 204.7845 | 77 | 245.1846 | 111 | 525.1306 | 173 | ESR1 | MCF-7 | Breast | GSM47041 | 5958 | 2301 |
| 387.2297 | 105 | 355.4896 | 111 | 589.9855 | 187 | 857.7963 | 247 | ESR1 | MCF-7 | Breast | GSM36592 | 7103 | 2303 |
| 873.0314 | 225 | 754.4975 | 231 | 789.8738 | 349 | 812.8605 | 420 | ESR1 | MCF-7 | Breast | GSM36592 | 15081 | 2304 |
| 459.608 | 178 | 480.279 | 203 | 705.3465 | 325 | 688.7707 | 350 | ESR1 | MCF-7 | Breast | GSM36592 | 18382 | 2305 |
| 292.403 | 88 | 300.7844 | 99 | 260.1681 | 121 | 483.1909 | 175 | FOS | MCF-7 | Breast | GSM65978 | 6700 | 2318 |
| 754.1224 | 126 | 612.07 | 122 | 792.2637 | 178 | 986.7811 | 238 | FOXA1 | MCF-7 | Breast | GSM65978 | 4455 | 2319 |
| 694.181 | 168 | 643.1512 | 178 | 856.3909 | 298 | 877.7184 | 361 | FOXA1 | MCF-7 | Breast | GSM65978 | 10064 | 2320 |
| 818.3979 | 489 | 759.6638 | 520 | 712.7148 | 739 | 736.6344 | 887 | FOXA1 | MCF-7 | Breast | GSM58893 | 51807 | 2321 |
| 822.9583 | 525 | 754.13 | 554 | 697.0587 | 765 | 717.3207 | 913 | FOXA1 | MCF-7 | Breast | GSM58892 | 57535 | 2322 |
| 366.964 | 141 | 336.5469 | 151 | 625.8305 | 268 | 756.1325 | 342 | GATA3 | MCF-7 | Breast | GSM72042 | 14248 | 2324 |
| 558.9106 | 256 | 526.6854 | 280 | 672.5584 | 473 | 722.7402 | 611 | GATA3 | MCF-7 | Breast | GSM72042 | 32199 | 2325 |
| 707.1315 | 525 | 718.0876 | 653 | 566.2698 | 743 | 574.1053 | 863 | RAD21 | MCF-7 | Breast | GSM61462 | 83830 | 2387 |
| 801.8001 | 340 | 823.5237 | 436 | 641.5692 | 444 | 642.5667 | 507 | RAD21 | MCF-7 | Breast | GSM61462 | 32937 | 2388 |
| 777.3817 | 538 | 766.8814 | 640 | 600.1863 | 704 | 591.1343 | 788 | RAD21 | MCF-7 | Breast | GSM61461 | 69542 | 2389 |
| 721.7515 | 451 | 732.007 | 562 | 589.4422 | 646 | 566.5169 | 697 | RAD21 | MCF-7 | Breast | GSM61461 | 63803 | 2390 |
| 713.7929 | 369 | 740.5228 | 479 | 567.8205 | 506 | 559.5112 | 563 | RAD21 | MCF-7 | Breast | GSM61461 | 49713 | 2391 |
| 733.7318 | 406 | 739.4979 | 501 | 580.8178 | 546 | 575.4349 | 614 | RAD21 | MCF-7 | Breast | GSM61461 | 52717 | 2392 |
| 779.5528 | 624 | 786.709 | 768 | 610.6357 | 843 | 607.947 | 962 | STAG1 | MCF-7 | Breast | GSM61462 | 88472 | 2393 |
| 955.684 | 676 | 970.0962 | 831 | 732.8088 | 822 | 745.1683 | 964 | STAG1 | MCF-7 | Breast | GSM61462 | 57093 | 2394 |
| 835.4742 | 640 | 875.6958 | 825 | 654.1215 | 840 | 644.77 | 943 | STAG1 | MCF-7 | Breast | GSM61461 | 76576 | 2395 |
| 835.8348 | 611 | 835.8169 | 737 | 645.4804 | 783 | 628.4428 | 866 | STAG1 | MCF-7 | Breast | GSM61461 | 70737 | 2396 |
| 851.7222 | 619 | 873.3777 | 777 | 654.1261 | 785 | 652.5775 | 894 | STAG1 | MCF-7 | Breast | GSM61460 | 69111 | 2397 |
| 841.4156 | 570 | 839.998 | 684 | 642.773 | 710 | 633.0054 | 792 | STAG1 | MCF-7 | Breast | GSM61460 | 61450 | 2398 |
| 836.4979 | 105 | 1063.957 | 132 | 489.7026 | 103 | 507.5237 | 111 | TDRD3 | MCF-7 | Breast | GSM56088 | 2327 | 2400 |
| 990.3958 | 599 | 1007.087 | 745 | 817.0944 | 786 | 813.1299 | 896 | TFAP2A | MCF-7 | Breast | GSM58892 | 40579 | 2401 |
| 991.2392 | 567 | 1029.471 | 739 | 809.5468 | 732 | 834.4816 | 883 | TFAP2A | MCF-7 | Breast | GSM58892 | 37230 | 2402 |

| | | | | | | | | | | | | | |
|---|---|---|---|---|---|---|---|---|---|---|---|---|---|
| 831.8371 | 592 | 881.3552 | 779 | 720.4006 | 884 | 721.9118 | 1016 | TFAP2C | MCF-7 | Breast | GSM53091 | 67442 | 2403 |
| 113.5497 | 91 | 198.3011 | 131 | 134.3136 | 155 | 100.1688 | 151 | TRIM24 | MCF-7 | Breast | GSM59460 | 17816 | 2404 |
| 265.0677 | 217 | 348.0507 | 275 | 210.9554 | 321 | 176.1308 | 330 | TRIM24 | MCF-7 | Breast | GSM59460 | 45172 | 2405 |
| 549.348 | 144 | 700.2737 | 181 | 492.841 | 196 | 452.692 | 201 | PML | MR4 | Bone Marr | GSM46820 | 9484 | 2447 |
| 944.02 | 231 | 921.0753 | 265 | 770.9171 | 287 | 591.2595 | 265 | PML | MR4 | Bone Marr | GSM46820 | 12798 | 2448 |
| 173.7187 | 39 | 167.3445 | 42 | 107.6101 | 43 | 187.1691 | 61 | PML | NB4 | Blood | GSM46818 | 2049 | 2460 |
| 172.738 | 57 | 308.0201 | 86 | 117.0977 | 70 | 106.5684 | 72 | RARA | NB4 | Blood | GSM46820 | 4901 | 2464 |
| 968.5646 | 200 | 983.7273 | 256 | 686.9595 | 236 | 417.7571 | 198 | RARA | NB4 | Blood | GSM46818 | 9917 | 2465 |
| 203.1552 | 48 | 211.0697 | 54 | 123.9056 | 53 | 153.7506 | 63 | RXRG | NB4 | Blood | GSM46818 | 2697 | 2467 |
| 0.842059 | 3 | 6.34363 | 8 | 8.232582 | 13 | 51.85697 | 31 | PAX3 | Rh4 | Muscle | GSM47181 | 1927 | 2654 |
| 2.53983 | 11 | 7.02202 | 18 | 33.74 | 46 | 87.63055 | 73 | PAX3 | Rh4 | Muscle | GSM47181 | 5720 | 2655 |
| 855.2136 | 147 | 1062.666 | 192 | 503.9944 | 153 | 555.1791 | 171 | ERG | RWPE1 | Prostate | GSM35364 | 5514 | 2660 |
| 1272.655 | 558 | 1303.559 | 707 | 944.8037 | 500 | 919.1722 | 532 | PHF8 | SH-SY5Y | Bone Marr | GSM51855 | 13688 | 2684 |
| 519.2112 | 70 | 745.7021 | 95 | 334.5064 | 73 | 318.2408 | 75 | ZNF711 | SH-SY5Y | Bone Marr | GSM51856 | 1714 | 2685 |
| 116.879 | 41 | 173.1332 | 56 | 206.1606 | 80 | 459.022 | 129 | ESR1 | T47D | Mammary | GSM58923 | 3658 | 2725 |
| 960.721 | 504 | 920.1657 | 561 | 780.2905 | 646 | 790.5328 | 752 | FOXA1 | T47D | Mammary | GSM65979 | 34232 | 2735 |
| 954.6114 | 171 | 665.1907 | 153 | 816.9219 | 218 | 910.4384 | 280 | FOXA1 | T47D | Mammary | GSM65979 | 6849 | 2736 |
| 890.7964 | 298 | 847.3076 | 326 | 692.1815 | 345 | 669.6998 | 367 | CREBBP | T98G | Brain | GSM52526 | 20353 | 2748 |
| 312.7387 | 68 | 243.2344 | 65 | 231.7801 | 82 | 313.521 | 102 | CREBBP | T98G | Brain | GSM52526 | 3435 | 2749 |
| 174.38 | 31 | 118.3312 | 27 | 90.55982 | 30 | 206.3272 | 50 | CREBBP | T98G | Brain | GSM52526 | 1202 | 2750 |
| 969.9426 | 313 | 957.3132 | 372 | 761.4272 | 354 | 747.9836 | 390 | EP300 | T98G | Brain | GSM52527 | 16900 | 2751 |
| 1025.236 | 574 | 1010.847 | 681 | 779.3801 | 641 | 751.0513 | 682 | EP300 | T98G | Brain | GSM52526 | 33450 | 2752 |
| 983.728 | 279 | 1003.53 | 357 | 778.9315 | 315 | 762.7367 | 345 | EP300 | T98G | Brain | GSM52526 | 13977 | 2753 |
| 1098.51 | 782 | 1127.735 | 976 | 757.5254 | 783 | 704.9159 | 796 | SMAD3 | BG03 | Embryo | GSM53954 | 48748 | 276 |
| 172.0386 | 31 | 296.1737 | 47 | 95.12505 | 31 | 196.9146 | 49 | VDR | THP-1 | Blood | GSM67826 | 1199 | 2774 |
| 103.3232 | 91 | 157.0942 | 123 | 103.6681 | 146 | 153.5267 | 183 | ESR1 | U2OS | Bone | GSM54254 | 18458 | 2778 |
| 272.3112 | 102 | 268.6795 | 113 | 148.3802 | 116 | 168.9196 | 132 | ESR1 | U2OS | Bone | GSM54254 | 9937 | 2779 |
| 1048.216 | 245 | 1035.813 | 295 | 676.4876 | 233 | 787.7315 | 268 | SMAD3 | BG03 | Embryo | GSM53954 | 9804 | 278 |
| 499.6553 | 124 | 637.5199 | 156 | 344.0234 | 147 | 435.0347 | 176 | TRIM28 | U2OS | Bone | GSM70036 | 7484 | 2780 |
| 190.4716 | 82 | 334.5796 | 121 | 186.7963 | 122 | 340.9925 | 172 | TRIM28 | U2OS | Bone | GSM70036 | 8979 | 2781 |
| 12.02248 | 7 | 9.239364 | 7 | 5.502984 | 8 | 79.80825 | 30 | TP53 | U2OS | Bone | GSM54581 | 1207 | 2793 |
| 12.70824 | 7 | 12.87114 | 8 | 2.939047 | 6 | 78.33718 | 29 | TP53 | U2OS | Bone | GSM54580 | 1152 | 2794 |
| 44.66342 | 22 | 28.45174 | 20 | 12.45092 | 20 | 40.76202 | 35 | TP53 | U2OS | Bone | GSM54580 | 2864 | 2795 |
| 46.42619 | 18 | 50.39678 | 21 | 18.84449 | 18 | 15.58432 | 18 | TP53 | U2OS | Bone | GSM54580 | 1883 | 2796 |
| 654.6743 | 235 | 774.3076 | 290 | 431.6862 | 289 | 584.1919 | 354 | TRIM28 | U2OS | Bone | GSM70036 | 22266 | 2797 |
| 358.3655 | 99 | 412.3674 | 118 | 244.9518 | 119 | 308.3119 | 142 | BRD7 | BJ | Skin | GSM50879 | 6915 | 280 |
| 488.8621 | 152 | 630.3614 | 192 | 278.1635 | 171 | 336.9609 | 200 | PML | UPR9 | Bone Marr | GSM46821 | 12098 | 2809 |
| 867.7793 | 266 | 867.6781 | 329 | 596.8705 | 307 | 630.8605 | 338 | RARA | UPR9 | Bone Marr | GSM46821 | 19227 | 2811 |
| 876.5195 | 296 | 875.4427 | 361 | 604.3897 | 325 | 503.4873 | 323 | RXRG | UPR9 | Bone Marr | GSM46821 | 21803 | 2813 |
| 120.4204 | 27 | 129.6052 | 31 | 80.22851 | 31 | 169.3314 | 49 | RXRG | UPR9 | Bone Marr | GSM46821 | 1468 | 2814 |
| 26.01955 | 14 | 50.51225 | 22 | 21.13441 | 20 | 29.60059 | 25 | SMC1A | None | None | GSM56035 | 2080 | 2894 |
| 161.9066 | 32 | 123.307 | 30 | 71.19495 | 29 | 71.43254 | 31 | SMC1A | None | None | GSM56035 | 1452 | 2895 |
| 106.3573 | 41 | 191.6762 | 62 | 116.5437 | 64 | 114.3788 | 68 | SMC1A | V6.5 | None | GSM56034 | 4135 | 2896 |
| 118.3834 | 35 | 97.40958 | 35 | 18.37257 | 22 | 42.89019 | 34 | SMC3 | V6.5 | None | GSM56034 | 2654 | 2899 |
| 677.4328 | 359 | 676.48 | 438 | 600.4308 | 580 | 655.6231 | 756 | AR | VCaP | Prostate | GSM69684 | 53112 | 2910 |
| 366.4943 | 69 | 387.359 | 78 | 345.8672 | 93 | 827.3414 | 160 | AR | VCaP | Prostate | GSM69684 | 2866 | 2911 |
| 305.9831 | 86 | 323.4069 | 98 | 194.7972 | 100 | 520.1217 | 173 | AR | VCaP | Prostate | GSM35364 | 6058 | 2912 |
| 1243.871 | 863 | 1319.018 | 1116 | 975.7749 | 1031 | 976.1394 | 1182 | ERG | VCaP | Prostate | GSM35364 | 39062 | 2916 |
| 784.0785 | 114 | 1075.233 | 150 | 649.0366 | 139 | 1009.514 | 191 | ERG | VCaP | Prostate | GSM35363 | 3245 | 2918 |
| 1089.973 | 200 | 1026.873 | 202 | 973.7499 | 286 | 1004.27 | 358 | FOXA1 | VCaP | Prostate | GSM35363 | 6679 | 2919 |
| 433.6061 | 88 | 453.0955 | 99 | 150.4064 | 72 | 236.5171 | 96 | ATF7IP | WM262 | Skin | GSM69762 | 4109 | 2972 |
| 915.0968 | 143 | 866.1986 | 151 | 469.9472 | 136 | 479.7421 | 146 | SETDB1 | WM262 | Skin | GSM69762 | 4599 | 2974 |
| 243.2881 | 55 | 246.8015 | 61 | 118.4479 | 54 | 91.44435 | 51 | SETDB1 | WM451-lu | Skin | GSM69762 | 2940 | 2976 |
| 613.4142 | 100 | 803.6771 | 128 | 661.0313 | 143 | 1034.72 | 211 | AR | None | Prostate | GSM35365 | 3379 | 2986 |
| 396.5626 | 260 | 516.394 | 328 | 239.6633 | 340 | 298.4074 | 399 | CEBPA | None | Adipose | GSM59983 | 45319 | 3006 |
| 1211.814 | 401 | 1265.227 | 541 | 918.2746 | 368 | 891.6262 | 386 | CREBBP | None | Blood | GSM39394 | 10420 | 3038 |
| 791.5111 | 340 | 831.4315 | 459 | 601.2835 | 413 | 579.6232 | 442 | CTCF | None | Bone Marr | GSM65154 | 34376 | 3049 |
| 756.674 | 342 | 788.6864 | 453 | 544.1463 | 423 | 556.7168 | 462 | CTCF | None | Bone Marr | GSM65154 | 38895 | 3050 |
| 412.8115 | 172 | 605.9725 | 231 | 257.9944 | 212 | 290.1459 | 240 | CTCF | None | Blood | GSM65154 | 19298 | 3051 |
| 429.9309 | 203 | 603.8195 | 266 | 249.983 | 248 | 319.1593 | 295 | CTCF | None | Pancreatic | GSM58688 | 25788 | 3053 |
| 772.4333 | 307 | 786.6301 | 390 | 488.2495 | 365 | 588.9999 | 426 | CTCF | None | Pancreatic | GSM58688 | 32050 | 3054 |
| 715.3621 | 351 | 762.3211 | 480 | 472.1115 | 437 | 539.3113 | 502 | CTCF | None | Adipose | GSM53449 | 44854 | 3056 |
| 138.908 | 133 | 175.4305 | 166 | 95.52293 | 188 | 98.04562 | 207 | CTCF | None | Adipose | GSM53448 | 28039 | 3057 |
| 353.577 | 248 | 650.9296 | 361 | 247.1401 | 342 | 317.7287 | 406 | CTCF | None | Adipose | GSM53447 | 44198 | 3058 |
| 441.2833 | 281 | 664.0414 | 388 | 308.439 | 383 | 364.9652 | 442 | CTCF | None | Adipose | GSM53447 | 46680 | 3059 |
| 776.141 | 256 | 831.1093 | 340 | 465.5043 | 301 | 454.0051 | 321 | CTCF | None | Blood | GSM32589 | 23239 | 3060 |
| 1246.639 | 290 | 1272.108 | 377 | 971.1401 | 267 | 951.541 | 288 | E2F4 | None | Adipose | GSM59983 | 6148 | 3225 |
| 912.9341 | 415 | 928.0633 | 523 | 663.001 | 434 | 633.5868 | 454 | EP300 | None | None | GSM60229 | 29933 | 3232 |
| 977.2405 | 129 | 1178.367 | 161 | 518.8113 | 121 | 665.5835 | 147 | EP300 | None | Blood | GSM39394 | 3129 | 3238 |
| 744.9015 | 141 | 1025.862 | 187 | 383.2708 | 139 | 408.8649 | 153 | FLI1 | None | Cord blood | GSM60795 | 6045 | 3252 |
| 155.5033 | 30 | 117.8497 | 28 | 135.9659 | 39 | 339.2699 | 69 | FOXA2 | None | Liver | GSM63461 | 1318 | 3262 |
| 828.3367 | 367 | 843.6705 | 463 | 655.1002 | 464 | 617.8774 | 479 | GATA1 | None | Bone Marr | GSM65154 | 33347 | 3272 |

| | | | | | | | | | | | | | |
|---|---|---|---|---|---|---|---|---|---|---|---|---|---|
| 995.1379 | 514 | 1028.469 | 671 | 759.3469 | 576 | 723.8419 | 603 | GATA1 | None | Bone Marr | GSM65154 | 31324 | 3273 |
| 153.5253 | 54 | 229.393 | 74 | 108.4083 | 68 | 166.9876 | 89 | GATA1 | None | Cord blood | GSM60794 | 4904 | 3274 |
| 102.3875 | 31 | 69.20289 | 28 | 31.36844 | 26 | 33.85636 | 29 | BMI1 | None | None | GSM83867 | 2386 | 32776 |
| 4.038595 | 17 | 4.441469 | 21 | 2.665378 | 29 | 27.55877 | 60 | BMI1 | None | None | GSM83866 | 8541 | 32777 |
| 23.74478 | 30 | 30.44371 | 38 | 17.11487 | 45 | 37.5737 | 64 | BMI1 | None | None | GSM83867 | 7968 | 32781 |
| 156.1687 | 161 | 153.8292 | 183 | 102.1018 | 227 | 248.5254 | 332 | ATF3 | None | None | GSM83867 | 36638 | 32784 |
| 1123.597 | 372 | 1171.834 | 501 | 869.9198 | 370 | 814.6608 | 360 | BRD4 | P493-6 | None | GSM10364 | 11994 | 32790 |
| 1177.99 | 400 | 1225.08 | 539 | 878.689 | 361 | 852.3648 | 378 | MYC | P493-6 | None | GSM10364 | 11255 | 32791 |
| 846.2996 | 106 | 984.7796 | 126 | 482.9336 | 102 | 430.1598 | 101 | MED1 | P493-6 | None | GSM10364 | 2260 | 32803 |
| 1038.877 | 373 | 1096.941 | 518 | 784.6686 | 373 | 749.0711 | 382 | BRD4 | P493-6 | None | GSM10364 | 15658 | 32808 |
| 1114.848 | 536 | 1128.554 | 663 | 824.3707 | 522 | 755.3762 | 496 | BRD4 | P493-6 | None | GSM10364 | 20368 | 32812 |
| 1209.74 | 491 | 1261.924 | 657 | 911.8586 | 465 | 874.4993 | 480 | MAX | P493-6 | None | GSM10364 | 14175 | 32814 |
| 1044.082 | 131 | 1203.308 | 164 | 556.1394 | 122 | 564.1001 | 130 | T | P493-6 | None | GSM10364 | 2908 | 32815 |
| 402.2017 | 52 | 398.0254 | 56 | 285.2468 | 57 | 361.7164 | 69 | T | P493-6 | None | GSM10364 | 1202 | 32816 |
| 10.77896 | 130 | 7.88252 | 147 | 8.104045 | 235 | 20.6957 | 304 | HOTAIR | MDA-231 | Mammary | GSM82043 | 80658 | 32847 |
| 251.853 | 446 | 318.4045 | 552 | 314.9606 | 788 | 260.083 | 832 | TERC | HeLa | Cervix | GSM82043 | 155675 | 32848 |
| 19.31374 | 35 | 11.22131 | 34 | 7.341218 | 46 | 17.7882 | 64 | HOTAIR | MDA-231 | Mammary | GSM82043 | 11094 | 32849 |
| 357.1105 | 507 | 485.4842 | 640 | 413.2841 | 866 | 317.1838 | 897 | TERC | HeLa | Cervix | GSM82043 | 162603 | 32850 |
| 3.392892 | 64 | 22.66369 | 114 | 12.67718 | 154 | 13.06432 | 172 | HOTAIR | MDA-231 | Mammary | GSM82043 | 44013 | 32852 |
| 402.5948 | 454 | 555.821 | 608 | 320.9745 | 692 | 314.4725 | 758 | TERC | HeLa | Cervix | GSM82043 | 123934 | 32853 |
| 3.459399 | 78 | 16.61481 | 126 | 16.1822 | 194 | 15.28358 | 213 | HOTAIR | MDA-231 | Mammary | GSM82043 | 55274 | 32854 |
| 556.9795 | 652 | 642.4193 | 903 | 456.2122 | 1015 | 430.1208 | 1109 | TERC | HeLa | Cervix | GSM82043 | 186569 | 32856 |
| 81.36771 | 28 | 118.2237 | 38 | 31.76559 | 27 | 71.87204 | 42 | FOXM1 | U2OS | Bone | GSM93620 | 2563 | 32890 |
| 966.4384 | 345 | 993.3432 | 447 | 735.0668 | 369 | 721.4003 | 404 | RELA | None | Umbilical V | GSM85030 | 19075 | 32899 |
| 552.9541 | 147 | 794.7431 | 197 | 291.9835 | 156 | 429.6991 | 200 | ERG | RWPE1 | Prostate | GSM92707 | 9868 | 32939 |
| 106.1789 | 295 | 142.6966 | 373 | 148.688 | 555 | 168.1318 | 631 | ARNT | MCF-7 | Breast | GSM10248 | 122503 | 33032 |
| 307.9056 | 344 | 324.5533 | 402 | 383.4555 | 599 | 399.0497 | 664 | AHR | MCF-7 | Breast | GSM10247 | 91348 | 33035 |
| 237.2916 | 141 | 303.0577 | 177 | 129.5593 | 173 | 109.1428 | 177 | TP53 | U2OS | Bone | GSM11334 | 21066 | 33075 |
| 23.6863 | 15 | 20.74371 | 16 | 10.33746 | 17 | 61.90302 | 39 | TP53 | U2OS | Bone | GSM11334 | 2480 | 33077 |
| 112.1044 | 46 | 119.31 | 53 | 56.49638 | 51 | 144.5109 | 83 | TP53 | U2OS | Bone | GSM11334 | 4800 | 33080 |
| 355.3121 | 116 | 334.6106 | 125 | 517.9408 | 203 | 804.2964 | 278 | ESR1 | MCF-7 | Breast | GSM11159 | 9555 | 33100 |
| 227.1448 | 161 | 257.7983 | 192 | 437.7834 | 324 | 638.6539 | 437 | ESR1 | MCF-7 | Breast | GSM11159 | 27144 | 33101 |
| 1125.447 | 626 | 1154.98 | 793 | 805.3078 | 595 | 769.5257 | 620 | CDK9 | MM1.S | Blood | GSM10857 | 27245 | 33123 |
| 1092.68 | 655 | 1109.654 | 808 | 769.9643 | 625 | 734.5118 | 659 | MED1 | MM1.S | Blood | GSM10857 | 32334 | 33124 |
| 1046.399 | 624 | 1064.703 | 772 | 735.4144 | 597 | 708.5671 | 641 | CDK9 | MM1.S | Blood | GSM10857 | 34023 | 33125 |
| 891.4949 | 487 | 855.7253 | 546 | 765.3785 | 700 | 762.0816 | 797 | EP300 | MCF-7 | Breast | GSM98608 | 40201 | 33126 |
| 783.5528 | 637 | 768.9895 | 751 | 698.2509 | 1000 | 728.341 | 1207 | ESR1 | MCF-7 | Breast | GSM98606 | 88542 | 33127 |
| 769.0056 | 242 | 751.3728 | 266 | 750.074 | 405 | 800.3663 | 525 | EP300 | MCF-7 | Breast | GSM98608 | 20343 | 33128 |
| 444.774 | 204 | 436.4726 | 227 | 665.924 | 380 | 698.5169 | 473 | GATA3 | MCF-7 | Breast | GSM98607 | 25296 | 33129 |
| 724.7432 | 585 | 700.2966 | 674 | 668.2885 | 972 | 699.0833 | 1173 | ESR1 | MCF-7 | Breast | GSM98606 | 92200 | 33135 |
| 108.0601 | 25 | 132.8938 | 31 | 293.2156 | 62 | 451.9311 | 84 | GATA3 | MCF-7 | Breast | GSM98607 | 1409 | 33136 |
| 819.3874 | 729 | 764.7297 | 806 | 685.5938 | 1064 | 714.0373 | 1275 | FOXA1 | MCF-7 | Breast | GSM98606 | 104340 | 33137 |
| 351.9156 | 93 | 267.8096 | 89 | 453.5658 | 152 | 806.1004 | 218 | GATA3 | MCF-7 | Breast | GSM98607 | 6100 | 33138 |
| 741.6656 | 297 | 690.4132 | 322 | 678.141 | 493 | 733.3629 | 647 | ESR1 | MCF-7 | Breast | GSM98606 | 33312 | 33139 |
| 795.1137 | 718 | 747.3552 | 801 | 656.914 | 1041 | 681.1789 | 1240 | FOXA1 | MCF-7 | Breast | GSM98606 | 109470 | 33143 |
| 521.397 | 130 | 330.4233 | 113 | 721.89 | 219 | 852.3652 | 273 | GATA3 | MCF-7 | Breast | GSM98607 | 7969 | 33144 |
| 577.6214 | 259 | 531.1286 | 280 | 673.2793 | 468 | 720.1976 | 604 | ESR1 | MCF-7 | Breast | GSM98605 | 31687 | 33145 |
| 778.4862 | 316 | 735.1528 | 346 | 712.3148 | 526 | 741.7108 | 644 | ESR1 | MCF-7 | Breast | GSM98606 | 32357 | 33146 |
| 70.24894 | 24 | 141.7766 | 39 | 80.66307 | 38 | 233.8117 | 70 | ESR1 | ZR-75-1 | Mammary | GSM98608 | 2135 | 33147 |
| 543.4858 | 304 | 443.4442 | 316 | 606.8403 | 536 | 641.0331 | 663 | GATA3 | MCF-7 | Breast | GSM98606 | 47011 | 33148 |
| 260.0068 | 68 | 175.5695 | 61 | 357.0448 | 115 | 637.5093 | 166 | GATA3 | MCF-7 | Breast | GSM98607 | 4389 | 33149 |
| 637.7225 | 427 | 586.5269 | 461 | 603.2142 | 760 | 641.8608 | 938 | GATA3 | MCF-7 | Breast | GSM98606 | 76230 | 33150 |
| 856.7224 | 361 | 810.8862 | 392 | 752.7366 | 544 | 793.3483 | 681 | EP300 | MCF-7 | Breast | GSM98608 | 29360 | 33154 |
| 909.2238 | 551 | 869.827 | 617 | 786.2487 | 799 | 800.1075 | 941 | EP300 | MCF-7 | Breast | GSM98608 | 45686 | 33155 |
| 816.8097 | 310 | 774.018 | 339 | 740.9348 | 502 | 770.6063 | 619 | ESR1 | MCF-7 | Breast | GSM98606 | 27763 | 33156 |
| 235.3757 | 45 | 135.6126 | 36 | 301.6498 | 72 | 613.9135 | 114 | GATA3 | MCF-7 | Breast | GSM98607 | 1912 | 33157 |
| 391.9047 | 85 | 262.6612 | 75 | 511.4004 | 138 | 757.8067 | 181 | GATA3 | MCF-7 | Breast | GSM98607 | 4293 | 33158 |
| 445.622 | 118 | 304.4334 | 107 | 507.7007 | 181 | 842.6248 | 259 | GATA3 | MCF-7 | Breast | GSM98606 | 7811 | 33159 |
| 1087.304 | 243 | 1089.654 | 303 | 820.3624 | 239 | 799.5437 | 251 | MED1 | P493-6 | None | GSM89407 | 8012 | 33160 |
| 1087.819 | 655 | 1108.137 | 814 | 786.9914 | 661 | 729.2104 | 658 | MAX | P493-6 | None | GSM89407 | 32671 | 33162 |
| 1100.03 | 143 | 1193.522 | 183 | 874.4312 | 168 | 719.9526 | 159 | MED1 | P493-6 | None | GSM89408 | 3371 | 33164 |
| 1163.075 | 219 | 1201.127 | 294 | 926.9392 | 220 | 839.3826 | 217 | MYC | P493-6 | None | GSM89405 | 5610 | 33168 |
| 979.2782 | 527 | 1004.717 | 670 | 728.4257 | 569 | 704.4256 | 615 | MED1 | H2171 | Lung | GSM89408 | 33226 | 33171 |
| 1122.927 | 524 | 1149.115 | 669 | 829.9245 | 504 | 794.6172 | 524 | MYC | U87 | Brain | GSM89406 | 19784 | 33174 |
| 1036.695 | 674 | 1050.055 | 833 | 740.7747 | 694 | 685.8794 | 699 | MAX | P493-6 | None | GSM89407 | 40885 | 33175 |
| 1050.507 | 781 | 1057.524 | 941 | 747.032 | 827 | 716.3791 | 893 | MAX | H128 | Lung | GSM89409 | 51072 | 33177 |
| 1185.35 | 424 | 1205.783 | 534 | 892.4228 | 394 | 880.6333 | 433 | MYC | H128 | Lung | GSM89409 | 12252 | 33180 |
| 1194.451 | 472 | 1226.864 | 616 | 894.1409 | 441 | 846.8347 | 442 | MYC | P493-6 | None | GSM89406 | 13573 | 33189 |
| 1107.321 | 300 | 1179.14 | 437 | 832.8762 | 280 | 832.8675 | 315 | MAX | MM1.S | Blood | GSM89408 | 9864 | 33191 |
| 1054.595 | 653 | 1078.517 | 822 | 762.4474 | 669 | 696.6592 | 658 | MAX | P493-6 | None | GSM89407 | 36465 | 33194 |
| 1155.853 | 579 | 1180.809 | 730 | 860.1619 | 567 | 789.8462 | 539 | MYC | P493-6 | None | GSM89405 | 21083 | 33195 |
| 887.4161 | 748 | 897.4017 | 910 | 620.6563 | 850 | 606.1567 | 943 | MAX | U87 | Brain | GSM89407 | 80965 | 33197 |

| | | | | | | | | | | | | | |
|---|---|---|---|---|---|---|---|---|---|---|---|---|---|
| 845.0141 | 474 | 841.3012 | 571 | 595.2749 | 496 | 585.7281 | 554 | MED1 | U87 | Brain | GSM89408 | 41950 | 33198 |
| 880.1029 | 141 | 1113.697 | 193 | 606.2405 | 157 | 602.7497 | 166 | MED1 | P493-6 | None | GSM89407 | 4596 | 33199 |
| 700.2561 | 404 | 673.2609 | 460 | 659.9584 | 705 | 678.5251 | 840 | ESR1 | MCF-7 | Breast | GSM11987 | 56013 | 33216 |
| 19.85475 | 33 | 27.63782 | 43 | 7.125459 | 42 | 12.74508 | 54 | SOX2 | ReN-VM | None | GSM11989 | 10037 | 33217 |
| 128.7051 | 116 | 143.9174 | 138 | 76.39936 | 155 | 59.99197 | 157 | SOX2 | ReN-VM | None | GSM11989 | 23338 | 33219 |
| 179.3742 | 76 | 211.1486 | 92 | 200.4393 | 120 | 313.5896 | 158 | NR5A2 | MCF-7 | Breast | GSM11987 | 8300 | 33220 |
| 746.5995 | 335 | 702.9339 | 366 | 693.4214 | 574 | 714.752 | 689 | ESR1 | MCF-7 | Breast | GSM11987 | 38057 | 33221 |
| 748.6022 | 392 | 703.1408 | 427 | 676.1988 | 639 | 719.8995 | 805 | ESR1 | MCF-7 | Breast | GSM11987 | 46947 | 33223 |
| 742.4015 | 398 | 705.6689 | 446 | 668.7829 | 648 | 704.0387 | 800 | ESR1 | MCF-7 | Breast | GSM11987 | 48857 | 33224 |
| 16.41152 | 9 | 3.51398 | 5 | 1.864687 | 6 | 13.10596 | 14 | NFE2 | None | Bone Marr | GSM90805 | 1451 | 33278 |
| 1.802897 | 4 | 19.02452 | 13 | 4.23877 | 10 | 2.331765 | 9 | TAL1 | None | Fetal Liver | GSM90805 | 1868 | 33284 |
| 7.680784 | 11 | 15.6548 | 17 | 10.49281 | 21 | 9.263469 | 22 | TAL1 | None | Bone Marr | GSM90805 | 3338 | 33293 |
| 6.544161 | 5 | 4.851627 | 5 | 5.875071 | 8 | 12.43203 | 12 | CTCF | None | Bone Marr | GSM90806 | 1183 | 33297 |
| 9.351311 | 9 | 10.73459 | 11 | 5.104261 | 12 | 22.05878 | 23 | EP300 | None | Blood | GSM97133 | 2210 | 33429 |
| 1069.977 | 147 | 1188.764 | 202 | 697.5131 | 156 | 689.5998 | 164 | SMARCA4 | None | Blood | GSM97133 | 3878 | 33430 |
| 1075.043 | 392 | 1104.366 | 509 | 806.7798 | 385 | 782.8294 | 410 | SMARCA4 | None | Blood | GSM97134 | 16091 | 33434 |
| 90.22555 | 60 | 172.8698 | 91 | 151.3082 | 116 | 115.785 | 112 | EP300 | None | Blood | GSM97133 | 9671 | 33436 |
| 1123.442 | 449 | 1149.281 | 577 | 843.1728 | 437 | 824.3529 | 469 | SMARCA4 | None | Blood | GSM97133 | 16582 | 33437 |
| 1016.574 | 427 | 1029.214 | 536 | 745.0087 | 425 | 696.0957 | 421 | SMARCA4 | None | Blood | GSM97133 | 22355 | 33440 |
| 356.0826 | 53 | 523.3489 | 73 | 313.5907 | 67 | 262.338 | 64 | CDK9 | P493-6 | None | GSM93513 | 1543 | 33445 |
| 740.5076 | 139 | 1041.6 | 189 | 378.6697 | 136 | 318.3154 | 133 | CDK9 | P493-6 | None | GSM93513 | 5774 | 33446 |
| 365.0973 | 56 | 469.4585 | 71 | 158.9113 | 48 | 291.8529 | 71 | CDK9 | P493-6 | None | GSM93513 | 1691 | 33447 |
| 1022.038 | 802 | 1075.591 | 1027 | 724.0269 | 889 | 682.903 | 939 | ERG | None | None | GSM58560 | 65813 | 33458 |
| 1045.445 | 632 | 1104.468 | 842 | 757.9196 | 654 | 704.5873 | 659 | ERG | None | None | GSM58560 | 36469 | 33459 |
| 270.971 | 49 | 486.7459 | 76 | 256.8537 | 66 | 396.8096 | 89 | ELF5 | T47D | Mammary | GSM77391 | 1931 | 33467 |
| 892.1186 | 519 | 872.0954 | 600 | 821.4767 | 833 | 828.1543 | 968 | ESR1 | BT-474 | Mammary | GSM79843 | 43658 | 33484 |
| 807.4065 | 612 | 775.1265 | 697 | 676.7778 | 883 | 710.5342 | 1079 | ESR1 | None | Mammary | GSM79842 | 76380 | 33487 |
| 853.0029 | 693 | 797.6403 | 763 | 716.8739 | 999 | 719.878 | 1151 | FOXA1 | MCF-7 | Breast | GSM79843 | 83483 | 33489 |
| 229.3822 | 92 | 273.6708 | 112 | 415.4517 | 181 | 637.3616 | 238 | ESR1 | None | Mammary | GSM79840 | 9508 | 33490 |
| 775.4899 | 497 | 751.2928 | 573 | 694.5474 | 788 | 725.5051 | 963 | ESR1 | MCF-7 | Breast | GSM79843 | 60346 | 33491 |
| 751.6065 | 520 | 695.6183 | 562 | 635.4231 | 767 | 662.5182 | 927 | ESR1 | None | Mammary | GSM79842 | 68419 | 33494 |
| 754.2277 | 451 | 721.3869 | 511 | 691.8311 | 750 | 713.4493 | 897 | ESR1 | TAM_R | Mammary | GSM79843 | 55916 | 33495 |
| 907.8758 | 715 | 864.5476 | 804 | 753.7595 | 999 | 764.8573 | 1169 | FOXA1 | MCF-7 | Breast | GSM79843 | 74679 | 33496 |
| 766.2746 | 429 | 715.3278 | 464 | 696.8083 | 702 | 725.24 | 855 | ESR1 | MCF-7 | Breast | GSM79842 | 50385 | 33504 |
| 1196.397 | 994 | 1170.8 | 1154 | 857.1596 | 1148 | 850.7966 | 1305 | ESR1 | None | Mammary | GSM79840 | 69702 | 33506 |
| 717.6018 | 171 | 664.5438 | 181 | 791.3439 | 255 | 833.3429 | 315 | ESR1 | None | Mammary | GSM79840 | 9944 | 33507 |
| 711.6087 | 218 | 592.2274 | 221 | 739.2634 | 342 | 760.5607 | 413 | ESR1 | T47D | Mammary | GSM79842 | 16900 | 33508 |
| 777.6894 | 392 | 740.948 | 436 | 716.6153 | 651 | 732.0969 | 774 | ESR1 | TAM_R | Mammary | GSM79843 | 42053 | 33509 |
| 798.281 | 549 | 774.5722 | 635 | 693.3927 | 831 | 706.4595 | 975 | FOXA1 | MCF-7 | Breast | GSM79843 | 66250 | 33510 |
| 893.4771 | 523 | 884.0056 | 619 | 811.2783 | 825 | 833.9885 | 984 | ESR1 | BT-474 | Mammary | GSM79843 | 44127 | 33512 |
| 889.0503 | 362 | 837.1166 | 390 | 774.3541 | 531 | 797.0309 | 640 | ESR1 | T47D | Mammary | GSM79842 | 26432 | 33513 |
| 806.3284 | 737 | 786.7494 | 864 | 684.4624 | 1100 | 686.9982 | 1265 | FOXA1 | MCF-7 | Breast | GSM79843 | 111891 | 33518 |
| 275.7329 | 64 | 272.5305 | 70 | 356.5531 | 104 | 512.1581 | 134 | ESR1 | None | Mammary | GSM79840 | 3521 | 33519 |
| 749.0365 | 524 | 733.9571 | 616 | 682.6184 | 852 | 718.7012 | 1046 | ESR1 | MCF-7 | Breast | GSM79843 | 70602 | 33520 |
| 845.0891 | 629 | 842.5905 | 757 | 749.4315 | 970 | 762.2689 | 1138 | ESR1 | MCF-7 | Breast | GSM79842 | 71520 | 33525 |
| 773.108 | 530 | 740.2454 | 604 | 696.1299 | 853 | 718.9387 | 1019 | ESR1 | MCF-7 | Breast | GSM79842 | 67352 | 33526 |
| 94.54387 | 20 | 94.67185 | 22 | 59.40895 | 22 | 188.6533 | 44 | IRF2 | None | Bone Marr | GSM97026 | 1000 | 33542 |
| 895.6756 | 345 | 955.9824 | 484 | 718.0986 | 431 | 644.8811 | 400 | GATA1 | None | Fetal Liver | GSM97025 | 24682 | 33543 |
| 678.0546 | 627 | 707.8382 | 797 | 529.5607 | 884 | 499.8593 | 952 | GATA1 | None | Bone Marr | GSM97025 | 120643 | 33545 |
| 356.6334 | 124 | 460.8068 | 157 | 195.0039 | 140 | 272.1011 | 174 | CTCF | None | Blood | GSM10565 | 11373 | 33556 |
| 119.9309 | 84 | 188.9856 | 116 | 101.3838 | 124 | 121.7688 | 144 | CTCF | None | None | GSM10565 | 14438 | 33561 |
| 896.3954 | 388 | 929.2483 | 508 | 655.3971 | 413 | 604.4887 | 408 | TBX21 | None | None | GSM77655 | 28973 | 33678 |
| 966.4429 | 404 | 988.307 | 518 | 719.6682 | 426 | 693.6887 | 450 | GATA3 | None | None | GSM77655 | 23856 | 33679 |
| 855.8988 | 246 | 850.8549 | 298 | 475.7956 | 269 | 448.945 | 282 | GATA3 | None | None | GSM77655 | 18303 | 33680 |
| 0.421743 | 50 | 0.100925 | 55 | 0.324928 | 100 | 0.47559 | 116 | SOX2 | BJ | Skin | GSM89698 | 44619 | 33707 |
| 26.35289 | 71 | 15.21495 | 71 | 8.809141 | 96 | 19.86823 | 127 | POU5F1 | BJ | Skin | GSM89698 | 26461 | 33713 |
| 234.0943 | 191 | 327.0699 | 248 | 102.2217 | 230 | 149.0237 | 283 | KLF4 | BJ | Skin | GSM89698 | 37175 | 33718 |
| 850.9881 | 397 | 832.3719 | 464 | 608.6088 | 422 | 612.4048 | 485 | MYC | BJ | Skin | GSM89698 | 34483 | 33720 |
| 358.9904 | 59 | 413.3922 | 70 | 269.837 | 69 | 541.6798 | 108 | EP300 | MCF-7 | Breast | GSM10593 | 1990 | 33736 |
| 57.75722 | 16 | 53.56888 | 17 | 20.79813 | 14 | 54.12802 | 24 | SF1 | H295R | Adrenal Gl | GSM10809 | 1137 | 33852 |
| 124.9719 | 89 | 260.3508 | 139 | 85.2462 | 120 | 114.0256 | 145 | POU5F1 | NCCIT | Embryo | GSM88171 | 14429 | 33864 |
| 190.7353 | 37 | 148.1315 | 35 | 64.6124 | 29 | 207.7209 | 57 | SMARCB1 | NCCIT | Embryo | GSM88171 | 1526 | 33865 |
| 10.76013 | 11 | 21.22891 | 17 | 6.264978 | 15 | 59.2531 | 40 | POU5F1 | NCCIT | Embryo | GSM88171 | 2662 | 33866 |
| 845.7702 | 102 | 780.3642 | 105 | 611.2399 | 112 | 498.0598 | 105 | MED1 | None | None | GSM10337 | 2067 | 33874 |
| 978.0587 | 594 | 994.254 | 736 | 681.9049 | 581 | 668.5886 | 642 | MED1 | None | None | GSM10337 | 38770 | 33875 |
| 1095.332 | 558 | 1117.598 | 707 | 792.3488 | 534 | 750.0506 | 543 | CDK9 | None | None | GSM10337 | 23998 | 33888 |
| 1195.851 | 157 | 1222.798 | 205 | 750.659 | 156 | 501.1772 | 132 | MYC | None | None | GSM10337 | 3495 | 33895 |
| 1149.321 | 478 | 1181.916 | 618 | 848.9455 | 443 | 799.2439 | 444 | BRD4 | None | None | GSM10337 | 15708 | 33904 |
| 66.52189 | 57 | 97.37117 | 76 | 55.76942 | 85 | 45.25314 | 86 | GRHL3 | NHEK | Skin | GSM10345 | 10866 | 33916 |
| 429.414 | 248 | 527.6842 | 305 | 232.5845 | 303 | 218.933 | 323 | WDR5 | NHEK | Skin | GSM10345 | 36525 | 33917 |
| 995.5094 | 676 | 899.0789 | 688 | 829.6428 | 920 | 884.8649 | 1163 | FOXA1 | LNCaP | Prostate | GSM91652 | 51289 | 33918 |
| 974.6551 | 96 | 574.9643 | 74 | 704.2689 | 103 | 871.7405 | 123 | FOXA1 | LNCaP | Prostate | GSM91652 | 1390 | 33919 |

| | | | | | | | | | | | | | |
|---|---|---|---|---|---|---|---|---|---|---|---|---|---|
| 823.486 | 147 | 842.7456 | 163 | 561.2717 | 166 | 968.6521 | 284 | AR | LNCaP | Prostate | GSM91652 | 5814 | 33920 |
| 758.6584 | 171 | 863.4838 | 202 | 866.9227 | 289 | 957.8514 | 422 | AR | LNCaP | Prostate | GSM91652 | 9453 | 33921 |
| 1019.55 | 669 | 1035.031 | 828 | 702.2145 | 655 | 663.5575 | 682 | BRD4 | MM1.S | Blood | GSM10701 | 42686 | 33932 |
| 1025.397 | 764 | 1061.778 | 961 | 723.409 | 813 | 673.8659 | 841 | CTCF | MM1.S | Blood | GSM10701 | 56150 | 33934 |
| 1111.981 | 233 | 1162.681 | 326 | 878.0309 | 235 | 866.7322 | 259 | TET3 | HEK293T | Embryonic | GSM89757 | 7198 | 33988 |
| 1114.405 | 249 | 1111.168 | 297 | 901.7913 | 270 | 855.2833 | 266 | TET2 | HEK293T | Embryonic | GSM89757 | 7646 | 33992 |
| 1168.212 | 224 | 1197.19 | 294 | 805.8934 | 201 | 901.3788 | 236 | OGT | HEK293T | Embryonic | GSM89757 | 5732 | 34001 |
| 309.778 | 142 | 421.8616 | 184 | 251.4211 | 197 | 276.6565 | 221 | SMC1A | BG03 | Embryo | GSM89712 | 17257 | 34008 |
| 325.4696 | 183 | 502.9936 | 250 | 190.5699 | 230 | 240.853 | 272 | SMC3 | BG03 | Embryo | GSM89711 | 27076 | 34011 |
| 102.953 | 49 | 141.0809 | 64 | 69.49793 | 63 | 82.46203 | 73 | SMAD3 | BG03 | Embryo | GSM89711 | 5977 | 34012 |
| 231.7335 | 49 | 240.736 | 55 | 155.8496 | 56 | 123.4198 | 53 | SETDB1 | WM853.2 | None | GSM89712 | 2404 | 34015 |
| 346.5044 | 101 | 481.1546 | 133 | 236.7151 | 122 | 278.6951 | 141 | SMAD3 | BG03 | Embryo | GSM89711 | 7396 | 34016 |
| 1009.942 | 177 | 1031.454 | 214 | 503.8031 | 170 | 595.1841 | 197 | SMAD3 | BG03 | Embryo | GSM89711 | 6888 | 34017 |
| 1036.924 | 283 | 1036.159 | 343 | 818.6052 | 302 | 787.3378 | 316 | CDK8 | None | Colon | GSM93756 | 10688 | 34074 |
| 1110.168 | 426 | 1124.025 | 534 | 858.36 | 432 | 814.2928 | 437 | CDK8 | None | Colon | GSM93756 | 14277 | 34078 |
| 1107.689 | 368 | 1142.356 | 482 | 863.1604 | 374 | 812.5729 | 371 | CDK8 | None | Colon | GSM93756 | 12038 | 34081 |
| 20.44534 | 60 | 48.71208 | 92 | 10.11955 | 89 | 20.8092 | 116 | NANOG | HUES64 | Embryo | GSM11240 | 22853 | 34089 |
| 10.94365 | 11 | 4.68479 | 9 | 1.663958 | 10 | 47.1587 | 36 | SOX2 | HUES64 | Embryo | GSM11240 | 2689 | 34092 |
| 162.7897 | 106 | 189.8549 | 128 | 35.50995 | 97 | 73.61135 | 134 | POU5F1 | HUES64 | Embryo | GSM11240 | 16833 | 34100 |
| 53.12495 | 74 | 68.00138 | 93 | 13.13871 | 83 | 49.75001 | 131 | NANOG | HUES64 | Embryo | GSM11240 | 19376 | 34102 |
| 7.265923 | 6 | 12.75754 | 9 | 2.953583 | 7 | 201.2088 | 53 | SOX2 | HUES64 | Embryo | GSM11240 | 1386 | 34118 |
| 585.414 | 381 | 627.4253 | 470 | 363.2167 | 505 | 491.2842 | 619 | STAT3 | MCF-10A | Breast | GSM93555 | 68718 | 34236 |
| 1238.196 | 557 | 1270.67 | 715 | 919.8099 | 507 | 877.9192 | 518 | MYC | HEK293T | Embryonic | GSM10540 | 14424 | 34318 |
| 1065.531 | 196 | 1072.55 | 243 | 824.0396 | 221 | 852.6583 | 239 | EZH2 | LNCaP-abl | Prostate | GSM96956 | 6780 | 34326 |
| 100.8221 | 23 | 127.507 | 29 | 38.61719 | 20 | 61.51415 | 27 | EZH2 | LNCaP | Prostate | GSM96957 | 1286 | 34329 |
| 1069.143 | 433 | 1012.729 | 467 | 919.0435 | 594 | 979.7885 | 777 | AR | LNCaP-abl | Prostate | GSM96956 | 19545 | 34330 |
| 76.05486 | 31 | 86.65529 | 37 | 40.12319 | 35 | 60.19931 | 45 | SUZ12 | LNCaP | Prostate | GSM96957 | 3267 | 34333 |
| 162.1122 | 62 | 190.5706 | 75 | 356.6126 | 135 | 561.5654 | 181 | FOXO3 | DL23 | Colon | GSM86930 | 6140 | 34350 |
| 172.8438 | 66 | 302.975 | 98 | 93.41491 | 75 | 111.1475 | 87 | SUMO1 | HeLa | Cervix | GSM10127 | 6605 | 34353 |
| 449.5424 | 148 | 591.6526 | 189 | 285.8074 | 177 | 228.8938 | 173 | SUMO1 | HeLa | Cervix | GSM10127 | 12811 | 34354 |
| 388.3924 | 170 | 533.3013 | 221 | 262.8554 | 218 | 218.741 | 219 | SUMO1 | HeLa | Cervix | GSM10127 | 20036 | 34355 |
| 222.0093 | 72 | 284.6339 | 91 | 105.1596 | 75 | 124.9721 | 87 | SUMO1 | HeLa | Cervix | GSM10127 | 6013 | 34356 |
| 483.3208 | 195 | 699.8473 | 260 | 297.1321 | 238 | 288.5894 | 254 | SUMO1 | HeLa | Cervix | GSM10127 | 21402 | 34357 |
| 222.1651 | 90 | 252.2649 | 107 | 102.7551 | 97 | 93.6223 | 101 | SUMO1 | HeLa | Cervix | GSM10127 | 9551 | 34361 |
| 294.0898 | 110 | 428.6083 | 148 | 153.3909 | 123 | 143.4146 | 129 | SUMO1 | HeLa | Cervix | GSM10127 | 10793 | 34364 |
| 152.0944 | 53 | 167.5524 | 62 | 47.89241 | 47 | 63.35822 | 57 | SUMO1 | HeLa | Cervix | GSM10127 | 4799 | 34365 |
| 0 | 6 | 0.017542 | 8 | -0.000988 | 12 | -0.775933 | 8 | SUMO1 | HeLa | Cervix | GSM10127 | 7256 | 34366 |
| 346.5542 | 135 | 475.9953 | 176 | 206.3178 | 161 | 174.0491 | 162 | SUMO1 | HeLa | Cervix | GSM10127 | 13950 | 34368 |
| 338.0321 | 136 | 463.003 | 177 | 183.5635 | 157 | 158.8194 | 160 | SUMO1 | HeLa | Cervix | GSM10127 | 14543 | 34369 |
| 307.2693 | 135 | 388.7592 | 169 | 153.494 | 153 | 214.7255 | 189 | SUMO1 | HeLa | Cervix | GSM10127 | 15747 | 34370 |
| 381.1383 | 176 | 472.6738 | 218 | 240.2197 | 221 | 240.1709 | 239 | SUMO1 | HeLa | Cervix | GSM10127 | 21822 | 34371 |
| 413.6898 | 175 | 499.3188 | 214 | 199.3445 | 195 | 204.3064 | 213 | SUMO1 | HeLa | Cervix | GSM10127 | 19971 | 34373 |
| 0.147593 | 7 | 0.187403 | 9 | -2.410018 | 4 | -0.008858 | 12 | SUMO1 | HeLa | Cervix | GSM10127 | 6687 | 34374 |
| 9.212672 | 144 | 10.06376 | 175 | 13.56423 | 289 | 31.52605 | 371 | SMARCA4 | None | None | GSM65155 | 92917 | 34389 |
| 40.05686 | 172 | 66.61004 | 230 | 39.30005 | 304 | 48.4573 | 352 | SMARCA4 | None | None | GSM65155 | 77383 | 34390 |
| 1115.892 | 663 | 1150.543 | 846 | 816.8687 | 676 | 764.7889 | 688 | MECP2 | IMR90 | Lung | GSM11545 | 32611 | 34392 |
| 1132.493 | 246 | 1163.752 | 328 | 882.1865 | 244 | 866.5102 | 266 | MECP2 | IMR90 | Lung | GSM11545 | 7488 | 34394 |
| 1146.691 | 619 | 1157.17 | 761 | 820.3511 | 582 | 792.838 | 617 | MECP2 | IMR90 | Lung | GSM11545 | 25989 | 34396 |
| 1035.472 | 472 | 1050.679 | 593 | 746.6745 | 455 | 709.1281 | 471 | MECP2 | HCT-116 | Colon | GSM11545 | 24044 | 34399 |
| 510.1241 | 131 | 681.062 | 169 | 219.5188 | 126 | 264.1571 | 147 | MECP2 | HCT-116 | Colon | GSM11545 | 8571 | 34400 |
| 71.60197 | 32 | 80.48683 | 38 | 43.22815 | 39 | 64.69975 | 50 | LMNA | HGFDFN16 | Forearm Sk | GSM10236 | 3768 | 34444 |
| 371.8846 | 146 | 572.7107 | 201 | 234.2669 | 177 | 234.6247 | 191 | PAX5 | Namalwa | None | GSM10862 | 14443 | 34466 |
| 185.7104 | 85 | 380.0338 | 135 | 105.7755 | 101 | 117.8668 | 114 | PAX5 | L428-PAX5 | None | GSM10862 | 9728 | 34467 |
| 285.5462 | 128 | 460.8255 | 180 | 131.8932 | 140 | 151.7062 | 160 | PAX5 | L428-PAX5 | None | GSM10862 | 14528 | 34472 |
| 326.5534 | 126 | 662.5842 | 200 | 188.4632 | 147 | 268.3288 | 184 | PAX5 | Raji | Blood | GSM10862 | 12343 | 34473 |
| 356.8673 | 111 | 535.3465 | 152 | 220.0697 | 129 | 266.6662 | 151 | PAX5 | Raji | Blood | GSM10862 | 8688 | 34475 |
| 201.1789 | 69 | 304.3212 | 95 | 85.72262 | 69 | 199.1467 | 108 | PAX5 | Namalwa | None | GSM10862 | 5931 | 34476 |
| 15.80766 | 13 | 27.25224 | 19 | 6.210235 | 15 | 46.595 | 36 | TP63 | None | Foreskin | GSM79486 | 2746 | 34559 |
| 29.93112 | 32 | 49.45626 | 45 | 22.90546 | 48 | 29.38937 | 57 | TP63 | None | Foreskin | GSM79486 | 7612 | 34560 |
| 546.132 | 103 | 653.0003 | 125 | 274.7513 | 100 | 282.3563 | 108 | EGR2 | None | None | GSM78550 | 4356 | 34563 |
| 83.13634 | 127 | 119.3479 | 166 | 57.34505 | 191 | 79.63804 | 230 | CEBPB | None | None | GSM78550 | 37056 | 34564 |
| 225.8171 | 283 | 309.4387 | 362 | 180.8786 | 436 | 185.4977 | 482 | SPI1 | None | None | GSM78550 | 77532 | 34565 |
| 794.049 | 121 | 1042.562 | 155 | 439.5676 | 119 | 425.1789 | 124 | EGR2 | None | None | GSM78550 | 3709 | 34567 |
| 787.5329 | 552 | 822.1895 | 712 | 563.2857 | 642 | 539.5423 | 693 | RUNX1 | SEM | None | GSM10320 | 65771 | 34568 |
| 332.3116 | 89 | 423.0793 | 112 | 155.6445 | 89 | 385.0606 | 147 | CHRM2 | C4-12 | None | GSM11684 | 5930 | 34675 |
| 743.6257 | 477 | 775.7604 | 617 | 540.982 | 575 | 525.1589 | 633 | SPI1 | None | None | GSM78549 | 64236 | 34929 |
| 1072.934 | 180 | 1105.305 | 242 | 539.6028 | 167 | 571.7287 | 183 | KMT2A | RS4 | None | GSM94154 | 6038 | 34962 |
| 316.4093 | 55 | 344.524 | 63 | 232.5114 | 63 | 161.1669 | 55 | AFF1 | RS4 | None | GSM94154 | 1792 | 34966 |
| 1082.121 | 213 | 1056.928 | 243 | 952.1076 | 297 | 960.7478 | 347 | FOXM1 | MCF-7 | Breast | GSM10010 | 7262 | 34995 |
| 1010.817 | 508 | 1057.56 | 679 | 756.0514 | 540 | 731.7305 | 580 | FOXM1 | MDA-MB-2 | Breast | GSM10010 | 28305 | 34997 |
| 1017.208 | 276 | 1024.881 | 348 | 776.2439 | 283 | 754.2828 | 301 | FOXM1 | MDA-MB-2 | Breast | GSM10010 | 12321 | 34999 |
| 1086.317 | 211 | 1088.078 | 259 | 936.8408 | 276 | 938.9214 | 316 | FOXM1 | MCF-7 | Breast | GSM10010 | 7110 | 35002 |

| | | | | | | | | | | | | | |
|---|---|---|---|---|---|---|---|---|---|---|---|---|---|
| 1061.075 | 228 | 1059.211 | 277 | 737.6151 | 228 | 825.5651 | 265 | FOXM1 | MDA-MB-2 | Breast | GSM10010 | 8400 | 35011 |
| 596.7067 | 101 | 768.8232 | 128 | 469.8788 | 122 | 421.4234 | 122 | SPDEF | MCF-7 | Breast | GSM11871 | 3628 | 35041 |
| 785.3221 | 287 | 758.6059 | 328 | 715.6867 | 475 | 739.6455 | 579 | ESR1 | MCF-7 | Breast | GSM11871 | 27933 | 35044 |
| 1052.871 | 625 | 1034.347 | 728 | 850.0214 | 776 | 832.8163 | 858 | ESR1 | MCF-7 | Breast | GSM11871 | 35833 | 35046 |
| 894.7948 | 414 | 869.3822 | 473 | 767.6246 | 587 | 797.9472 | 721 | ESR1 | MCF-7 | Breast | GSM11871 | 31060 | 35051 |
| 284.3019 | 58 | 413.4605 | 79 | 196.9195 | 67 | 159.3924 | 64 | SPDEF | MCF-7 | Breast | GSM11871 | 2711 | 35052 |
| 117.928 | 26 | 51.31614 | 18 | 47.77867 | 23 | 129.0466 | 41 | HSF1 | None | None | GSM95190 | 1345 | 35064 |
| 929.8905 | 302 | 951.859 | 387 | 714.2821 | 325 | 595.1614 | 315 | BCL6 | None | None | GSM89806 | 16585 | 35085 |
| 1045.04 | 435 | 1067.917 | 555 | 783.1561 | 437 | 734.1811 | 434 | BCOR | None | None | GSM89806 | 19360 | 35087 |
| 975.6686 | 384 | 976.0585 | 467 | 728.9555 | 399 | 690.4002 | 405 | EP300 | OCI-Ly1 | Bone Marr | GSM89805 | 21336 | 35089 |
| 206.5554 | 85 | 337.1599 | 121 | 161.6649 | 114 | 127.3907 | 111 | NCOR1 | OCI-Ly1 | Bone Marr | GSM89805 | 8906 | 35091 |
| 383.2837 | 106 | 515.4478 | 137 | 280.619 | 130 | 174.8803 | 112 | NCOR2 | None | None | GSM89806 | 6808 | 35093 |
| 342.1607 | 57 | 416.9756 | 70 | 155.6572 | 51 | 320.5863 | 80 | HSF1 | M0-91 | None | GSM11166 | 1991 | 35219 |
| 1104.834 | 203 | 1066.303 | 225 | 573.1921 | 177 | 519.1587 | 179 | HSF1 | M0-91 | None | GSM11166 | 6528 | 35220 |
| 298.0925 | 42 | 426.5999 | 57 | 169.039 | 41 | 281.7848 | 58 | HSF1 | M0-91 | None | GSM11166 | 1123 | 35224 |
| 986.1649 | 300 | 1026.238 | 401 | 751.59 | 311 | 703.0916 | 312 | HSF1 | M0-91 | None | GSM11166 | 15045 | 35228 |
| 66.95834 | 31 | 52.54559 | 31 | 40.17182 | 38 | 114.5425 | 65 | HSF1 | MCF-7 | Breast | GSM11166 | 3840 | 35230 |
| 563.6585 | 84 | 687.064 | 103 | 332.9531 | 85 | 388.7423 | 98 | HSF1 | M0-91 | None | GSM11166 | 2460 | 35231 |
| 898.3302 | 106 | 1127.535 | 132 | 444.4956 | 93 | 542.4729 | 110 | SNAPC1 | MCF-10A | Breast | GSM91781 | 2021 | 35268 |
| 797.3458 | 99 | 1141.354 | 134 | 418.3523 | 91 | 480.8601 | 104 | ORC1 | HeLa-S3 | Cervix | GSM92279 | 2142 | 35296 |
| 1054.984 | 815 | 1101.238 | 1035 | 732.2633 | 869 | 713.3739 | 955 | ERG | SKNO-1 | Bone Marr | GSM58559 | 61602 | 35298 |
| 52.12245 | 22 | 86.75507 | 32 | 38.3316 | 29 | 105.6556 | 50 | RUNX1T1 | Kasumi-1 | Blood | GSM58558 | 2490 | 35300 |
| 1117.341 | 303 | 1150.934 | 403 | 876.64 | 314 | 803.6914 | 290 | ERG | None | None | GSM58559 | 10112 | 35302 |
| 833.3208 | 388 | 850.8041 | 491 | 626.8512 | 452 | 594.0489 | 472 | TCF12 | Kasumi-1 | Blood | GSM58558 | 35453 | 35308 |
| 1026.311 | 710 | 1057.654 | 895 | 732.514 | 746 | 716.1354 | 822 | FLI1 | SKNO-1 | Bone Marr | GSM58559 | 46911 | 35309 |
| 539.0034 | 341 | 578.3061 | 397 | 533.2757 | 533 | 538.2569 | 615 | GRHL2 | None | Bronchia | GSM11259 | 58405 | 35315 |
| 393.1779 | 245 | 417.844 | 284 | 452.7079 | 400 | 496.5113 | 449 | GRHL2 | None | Bronchia | GSM11259 | 39712 | 35317 |
| 458.1718 | 249 | 471.3612 | 284 | 467.5674 | 384 | 578.7841 | 460 | GRHL2 | None | Bronchia | GSM11259 | 35989 | 35323 |
| 709.9733 | 335 | 756.3253 | 460 | 514.2392 | 441 | 490.6759 | 470 | CTCF | NHLF | Lung | GSM64647 | 43417 | 35380 |
| 741.1386 | 309 | 792.2051 | 428 | 505.4364 | 391 | 511.1451 | 425 | CTCF | None | Umbilical V | GSM64641 | 34736 | 35387 |
| 493.776 | 300 | 662.2949 | 397 | 321.1004 | 398 | 384.118 | 461 | CTCF | NHEK | Skin | GSM64645 | 49220 | 35392 |
| 431.7796 | 251 | 687.5975 | 350 | 274.7897 | 327 | 353.4604 | 389 | CTCF | None | Umbilical V | GSM64641 | 38502 | 35405 |
| 587.6118 | 313 | 692.3568 | 410 | 375.2734 | 404 | 353.2393 | 430 | CTCF | NHEK | Skin | GSM64645 | 45069 | 35406 |
| 523.2978 | 237 | 734.6433 | 317 | 344.9099 | 302 | 367.5698 | 335 | CTCF | NHLF | Lung | GSM64647 | 28906 | 35412 |
| 549.7175 | 307 | 689.3297 | 412 | 375.2082 | 409 | 430.4674 | 467 | CTCF | K562 | Bone Marr | GSM64643 | 46458 | 35416 |
| 451.1249 | 292 | 684.0697 | 424 | 362.6965 | 417 | 415.5045 | 476 | CTCF | K562 | Bone Marr | GSM64643 | 48044 | 35447 |
| 370.9538 | 123 | 446.4133 | 150 | 386.6879 | 183 | 485.2186 | 218 | NKX2-1 | NCI-H1819 | Lung | GSM98310 | 10165 | 35471 |
| 750.6185 | 383 | 708.4956 | 422 | 587.9876 | 501 | 605.8758 | 598 | FOXA1 | NCI-H3122 | Lung | GSM98310 | 45834 | 35473 |
| 958.3447 | 663 | 933.814 | 763 | 699.9569 | 742 | 693.8495 | 836 | NKX2-1 | NCI-H3122 | Lung | GSM98310 | 53803 | 35474 |
| 789.837 | 167 | 1002.226 | 235 | 527.5917 | 189 | 605.8515 | 216 | NKX2-1 | NCI-H2087 | Lung | GSM98310 | 7669 | 35475 |
| 10.45695 | 13 | 18.5898 | 19 | 30.74604 | 33 | 62.24762 | 48 | TAL1 | CCRF-CEM | Blood | GSM83799 | 3566 | 35476 |
| 14.3322 | 11 | 21.07241 | 15 | 10.78005 | 16 | 63.28443 | 37 | GATA3 | CCRF-CEM | Blood | GSM83798 | 2266 | 35477 |
| 9.553465 | 8 | 14.15003 | 11 | 5.910729 | 11 | 34.93726 | 25 | TCF12 | CCRF-CEM | Blood | GSM83798 | 1845 | 35478 |
| 39.87928 | 15 | 36.37069 | 16 | 6.156912 | 10 | 18.13134 | 17 | TAL1 | None | Lymph Noc | GSM83799 | 1581 | 35479 |
| 70.06357 | 21 | 52.60597 | 20 | 15.07951 | 15 | 103.2681 | 40 | TCF12 | CCRF-CEM | Blood | GSM83798 | 1647 | 35481 |
| 46.74665 | 19 | 54.38983 | 23 | 34.31303 | 25 | 26.38048 | 24 | TAL1 | None | Lymph Noc | GSM83799 | 2129 | 35482 |
| 13.72192 | 10 | 24.48401 | 15 | 4.985104 | 11 | 18.54772 | 20 | RUNX1 | CCRF-CEM | Blood | GSM83799 | 2026 | 35483 |
| 93.42068 | 35 | 133.8035 | 47 | 40.64723 | 36 | 89.65229 | 55 | LMO2 | CCRF-CEM | Blood | GSM83799 | 3455 | 35484 |
| 36.79434 | 14 | 107.1609 | 28 | 10.50929 | 12 | 74.22419 | 32 | LMO2 | CCRF-CEM | Blood | GSM83799 | 1488 | 35486 |
| 10.37957 | 11 | 9.082715 | 12 | 4.728829 | 14 | 50.5165 | 38 | TAL1 | CCRF-CEM | Blood | GSM83799 | 2834 | 35489 |
| 707.1279 | 211 | 864.1745 | 282 | 347.6049 | 222 | 441.6718 | 265 | CTCF | None | Blood | GSM11389 | 16268 | 35498 |
| 1017.657 | 792 | 1035.721 | 969 | 724.589 | 875 | 669.6775 | 899 | BCL6 | OCI-Ly1 | Bone Marr | GSM76340 | 63913 | 35517 |
| 572.9964 | 190 | 754.0617 | 242 | 395.8683 | 235 | 288.553 | 221 | BCL6 | OCI-Ly1 | Bone Marr | GSM76340 | 16164 | 35524 |
| 250.4745 | 91 | 339.0007 | 118 | 141.1362 | 104 | 138.1907 | 111 | NCOR2 | OCI-Ly1 | Bone Marr | GSM76340 | 8388 | 35525 |
| 1033.159 | 197 | 1033.651 | 238 | 582.2094 | 189 | 512.2342 | 189 | BCOR | OCI-Ly1 | Bone Marr | GSM76341 | 6238 | 35528 |
| 336.6267 | 49 | 524.1551 | 70 | 265.7997 | 58 | 335.7555 | 70 | SFMBT1 | HeLa-S3 | Cervix | GSM11043 | 1371 | 35569 |
| 235.6651 | 42 | 290.7856 | 52 | 113.2857 | 39 | 275.1808 | 67 | RCOR1 | HeLa-S3 | Cervix | GSM11043 | 1595 | 35580 |
| 1040.561 | 494 | 1067.49 | 632 | 853.6884 | 618 | 822.1087 | 655 | CDX2 | Caco-2 | Colon | GSM57522 | 25419 | 356 |
| 26.36047 | 19 | 39.82412 | 26 | 13.24301 | 23 | 17.84513 | 28 | REST | H295R | Adrenal Gl | GSM11921 | 3265 | 35605 |
| 32.01353 | 21 | 49.22403 | 29 | 11.30778 | 22 | 18.96085 | 29 | REST | H295R | Adrenal Gl | GSM11921 | 3338 | 35607 |
| 210.2281 | 60 | 338.9696 | 86 | 160.1162 | 76 | 193.3697 | 89 | NR2F2 | None | Cranial | GSM71481 | 4260 | 35621 |
| 263.799 | 72 | 461.6793 | 108 | 162.8912 | 82 | 275.1492 | 113 | NR2F1 | None | Cranial | GSM71481 | 4857 | 35628 |
| 1272.002 | 188 | 1295.534 | 247 | 998.5911 | 181 | 998.9098 | 191 | PBX1 | MCF7-LTEC | Mammary | GSM98935 | 3261 | 35658 |
| 904.1989 | 170 | 994.0368 | 203 | 516.714 | 177 | 449.7234 | 176 | SUMO1 | WI-38 | Lung | GSM10354 | 7304 | 35694 |
| 674.196 | 140 | 938.5427 | 185 | 664.3766 | 195 | 706.489 | 214 | CDX2 | Caco-2 | Colon | GSM57522 | 6792 | 357 |
| 928.4374 | 202 | 922.2241 | 228 | 450.7441 | 200 | 406.7915 | 204 | SUMO1 | WI-38 | Lung | GSM10354 | 10806 | 35700 |
| 772.6756 | 284 | 799.0699 | 371 | 414.3733 | 323 | 438.3914 | 357 | SUMO2 | WI-38 | Lung | GSM10354 | 28851 | 35704 |
| 89.2474 | 25 | 91.08249 | 28 | 41.73089 | 25 | 157.69 | 52 | PIAS4 | WI-38 | Lung | GSM10354 | 1794 | 35708 |
| 574.2649 | 117 | 635.8192 | 136 | 238.2722 | 105 | 259.959 | 117 | UBE2I | WI-38 | Lung | GSM10354 | 5554 | 35717 |
| 268.1999 | 43 | 288.6424 | 49 | 63.37665 | 27 | 108.4374 | 38 | SUMO1 | WI-38 | Lung | GSM10354 | 1412 | 35719 |
| 936.5697 | 292 | 923.7307 | 347 | 566.1772 | 282 | 550.5688 | 299 | SUMO1 | WI-38 | Lung | GSM10354 | 16700 | 35724 |
| 227.2125 | 102 | 297.9166 | 130 | 130.374 | 122 | 140.9892 | 136 | SUMO2 | WI-38 | Lung | GSM10354 | 11871 | 35725 |

| | | | | | | | | | | | | | |
|---|---|---|---|---|---|---|---|---|---|---|---|---|---|
| 187.6192 | 40 | 227.1713 | 49 | 139.751 | 48 | 239.7832 | 68 | AR | C4-2B | Prostate | GSM98438 | 1918 | 35764 |
| 229.9035 | 35 | 290.9468 | 44 | 56.85309 | 22 | 339.3797 | 63 | AR | 22RV1 | Prostate | GSM98439 | 1038 | 35765 |
| 818.5559 | 173 | 898.5234 | 200 | 739.7949 | 230 | 915.1402 | 351 | AR | C4-2B | Prostate | GSM98438 | 8443 | 35766 |
| 1044.131 | 530 | 966.1846 | 545 | 893.9835 | 725 | 949.5141 | 927 | FOXA1 | C4-2B | Prostate | GSM98439 | 27450 | 35768 |
| 485.7698 | 95 | 605.0758 | 118 | 319.5101 | 106 | 857.5055 | 191 | AR | LNCaP | Prostate | GSM98438 | 4104 | 35775 |
| 1108.478 | 401 | 1005.123 | 379 | 981.2088 | 568 | 1032.805 | 733 | FOXA1 | C4-2B | Prostate | GSM98439 | 14601 | 35776 |
| 67.43564 | 38 | 78.69049 | 46 | 104.5069 | 71 | 109.735 | 78 | AR | C4-2B | Prostate | GSM98438 | 5394 | 35777 |
| 305.2579 | 53 | 297.3585 | 57 | 169.6789 | 53 | 633.5623 | 117 | AR | C4-2B | Prostate | GSM98438 | 1936 | 35778 |
| 657.9458 | 149 | 820.3769 | 185 | 594.0056 | 200 | 922.9972 | 339 | AR | LNCaP | Prostate | GSM98438 | 8088 | 35779 |
| 204.834 | 33 | 186.7598 | 34 | 105.4632 | 31 | 255.8464 | 54 | AR | C4-2B | Prostate | GSM98438 | 1044 | 35780 |
| 1061.832 | 240 | 1072.659 | 298 | 894.0837 | 299 | 928.6432 | 377 | REPIN1 | LNCaP | Prostate | GSM71461 | 8989 | 35828 |
| 886.6887 | 645 | 955.9849 | 868 | 649.5419 | 754 | 621.3171 | 808 | CTCF | GM19240 | Blood | GSM82227 | 65050 | 35832 |
| 153.1976 | 34 | 203.2946 | 44 | 82.60954 | 35 | 163.0052 | 53 | MYC | H1 | Embryo | GSM82227 | 1814 | 35837 |
| 852.5296 | 687 | 939.2112 | 940 | 653.9117 | 881 | 615.6505 | 936 | CTCF | NHEK | Skin | GSM82227 | 82596 | 35840 |
| 892.8953 | 653 | 944.7768 | 857 | 645.8466 | 753 | 617.6644 | 813 | CTCF | GM19239 | Blood | GSM82227 | 66447 | 35845 |
| 713.5918 | 463 | 764.2887 | 626 | 548.1334 | 615 | 533.2418 | 675 | CTCF | H1 | Embryo | GSM82229 | 68304 | 35846 |
| 897.9438 | 690 | 938.1943 | 884 | 647.1232 | 796 | 609.3784 | 841 | CTCF | GM12892 | Blood | GSM82229 | 71095 | 35851 |
| 914.5461 | 681 | 972.7149 | 898 | 670.2288 | 798 | 636.5703 | 850 | CTCF | GM12891 | Blood | GSM82229 | 66600 | 35854 |
| 888.9197 | 663 | 957.3084 | 888 | 651.9014 | 778 | 621.985 | 833 | CTCF | GM19238 | Blood | GSM82227 | 67862 | 35855 |
| 274.0346 | 75 | 339.1947 | 93 | 166.965 | 85 | 364.2145 | 133 | BRD3 | HEK293 | Embryonic | GSM97194 | 5105 | 35874 |
| 1051.273 | 689 | 1094.257 | 884 | 783.978 | 760 | 773.1751 | 854 | BRD2 | HEK293 | Embryonic | GSM97194 | 42551 | 35878 |
| 392.8302 | 176 | 634.6608 | 247 | 253.7962 | 222 | 284.3885 | 251 | CTCF | WI-38 | Lung | GSM10079 | 21086 | 35924 |
| 457.496 | 156 | 752.9968 | 223 | 249.0543 | 175 | 335.0318 | 214 | CTCF | WI-38 | Lung | GSM10079 | 13834 | 35929 |
| 351.5252 | 123 | 282.2212 | 123 | 300.5428 | 169 | 523.1461 | 234 | FOXA1 | Huh7 | Liver | GSM95875 | 11240 | 35942 |
| 334.6952 | 121 | 272.4389 | 122 | 302.1459 | 171 | 499.5189 | 231 | FOXA1 | Huh7 | Liver | GSM95875 | 11444 | 35945 |
| 281.6481 | 88 | 250.5666 | 92 | 239.4366 | 119 | 431.5124 | 169 | FOXA1 | Huh7 | Liver | GSM95875 | 7029 | 35946 |
| 290.1192 | 81 | 250.8643 | 83 | 220.0158 | 102 | 236.9232 | 113 | HIRA | None | Cervix | GSM10959 | 5564 | 36026 |
| 870.3524 | 207 | 899.5818 | 245 | 471.6903 | 218 | 555.3731 | 252 | RB1 | None | Cervix | GSM10959 | 11907 | 36028 |
| 1037.276 | 333 | 1050.884 | 419 | 812.9972 | 359 | 780.2237 | 373 | SPI1 | None | Cervix | GSM10959 | 14343 | 36032 |
| 733.2499 | 341 | 775.0863 | 459 | 569.7332 | 446 | 527.0647 | 469 | MYOD1 | None | None | GSM12188 | 41622 | 36075 |
| 654.617 | 342 | 733.6163 | 483 | 447.4822 | 455 | 490.351 | 511 | MYOD1 | None | Muscle | GSM12188 | 51053 | 36077 |
| 496.4256 | 114 | 821.8408 | 166 | 294.0122 | 124 | 374.1535 | 149 | YY1 | None | Embryo | GSM95612 | 6201 | 36082 |
| 1060.181 | 486 | 1040.308 | 562 | 878.2862 | 611 | 866.0315 | 676 | CTBP1 | MCF-7 | Breast | GSM89674 | 23000 | 36085 |
| 225.3635 | 54 | 176.1085 | 52 | 236.8332 | 79 | 428.8309 | 115 | ESR1 | None | Mammary | GSM79838 | 3121 | 36092 |
| 930.8449 | 418 | 928.319 | 508 | 836.417 | 645 | 856.3432 | 769 | ESR1 | None | Mammary | GSM79839 | 28392 | 36094 |
| 15.12924 | 13 | 20.46351 | 17 | 2.910851 | 12 | 7.533666 | 18 | ESR1 | None | Mammary | GSM79839 | 2839 | 36096 |
| 10.93633 | 11 | 21.48704 | 17 | 21.6944 | 24 | 49.89033 | 37 | ESR1 | None | Mammary | GSM79839 | 2725 | 36097 |
| 428.3014 | 107 | 555.8583 | 136 | 575.9099 | 178 | 867.9807 | 236 | ESR1 | None | Mammary | GSM79839 | 6579 | 36098 |
| 439.7684 | 74 | 608.8815 | 98 | 433.4484 | 101 | 542.0615 | 121 | ESR1 | None | Mammary | GSM79838 | 2610 | 36100 |
| 581.9406 | 126 | 562.8968 | 136 | 493.8015 | 163 | 735.3012 | 213 | ESR1 | None | Mammary | GSM79838 | 6400 | 36101 |
| 133.2152 | 41 | 153.3238 | 49 | 187.3724 | 71 | 274.6344 | 92 | ESR1 | None | Mammary | GSM79838 | 3184 | 36103 |
| 1210.73 | 906 | 1240.93 | 1110 | 927.5757 | 1075 | 898.9068 | 1176 | ESR1 | None | Mammary | GSM79839 | 48751 | 36104 |
| 397.2433 | 79 | 408.4981 | 88 | 414.9536 | 113 | 560.7486 | 141 | ESR1 | None | Mammary | GSM79839 | 3531 | 36105 |
| 936.9333 | 411 | 939.2864 | 501 | 831.5992 | 613 | 851.616 | 733 | ESR1 | None | Mammary | GSM79838 | 27055 | 36106 |
| 97.33585 | 26 | 124.7152 | 33 | 62.29126 | 30 | 143.6701 | 49 | ESR1 | None | Mammary | GSM79839 | 1752 | 36107 |
| 338.2541 | 65 | 477.4164 | 87 | 465.2521 | 108 | 669.3777 | 140 | ESR1 | None | Mammary | GSM79839 | 2777 | 36109 |
| 713.3611 | 152 | 751.8602 | 172 | 877.7648 | 248 | 887.3033 | 292 | ESR1 | None | Mammary | GSM79839 | 7694 | 36110 |
| 17.08496 | 39 | 18.91488 | 47 | 11.21931 | 61 | 25.48895 | 84 | FANCD2 | HEK293T | Embryonic | GSM11403 | 14381 | 36114 |
| 680.9755 | 197 | 907.0078 | 290 | 501.1823 | 248 | 420.9416 | 246 | TP63 | None | None | GSM83012 | 15105 | 36140 |
| 990.1537 | 397 | 1024.918 | 523 | 724.6165 | 394 | 710.7002 | 432 | TP63 | None | None | GSM83012 | 21731 | 36141 |
| 27.5831 | 42 | 19.47081 | 43 | 109.2592 | 115 | 194.956 | 157 | GATA6 | Caco-2 | Colon | GSM57522 | 12031 | 362 |
| 1246.55 | 426 | 1301.105 | 583 | 947.1937 | 389 | 938.8996 | 435 | DCP1A | HEK293 | Embryonic | GSM89124 | 10321 | 36226 |
| 1238.295 | 412 | 1269.997 | 536 | 925.7144 | 360 | 916.4482 | 400 | XRN2 | HEK293 | Embryonic | GSM89124 | 10002 | 36228 |
| 1305.962 | 647 | 1360.822 | 850 | 942.7456 | 561 | 917.5001 | 604 | TTF2 | HEK293 | Embryonic | GSM89123 | 16377 | 36230 |
| 1229.723 | 267 | 1253.169 | 345 | 936.3007 | 238 | 922.0348 | 255 | DCP1A | HeLa | Cervix | GSM89123 | 5947 | 36231 |
| 130.3465 | 34 | 133.1148 | 38 | 186.2776 | 59 | 260.8921 | 75 | TTF2 | HeLa | Cervix | GSM89123 | 2199 | 36232 |
| 1292.321 | 588 | 1369.739 | 807 | 954.1201 | 522 | 903.3571 | 523 | XRN2 | HEK293 | Embryonic | GSM89124 | 14363 | 36234 |
| 862.3379 | 240 | 915.2114 | 338 | 613.8899 | 294 | 578.0671 | 307 | RUNX1 | None | Blood | GSM10978 | 17454 | 36244 |
| 839.6403 | 240 | 871.1802 | 312 | 515.4839 | 278 | 433.3596 | 277 | FLI1 | None | Blood | GSM10978 | 18196 | 36245 |
| 72.99911 | 25 | 112.2571 | 35 | 68.77923 | 36 | 146.5539 | 56 | ERG | None | Blood | GSM10978 | 2250 | 36246 |
| 12.73032 | 11 | 38.42924 | 21 | 9.309856 | 16 | 64.23721 | 39 | LYL1 | None | Blood | GSM10978 | 2424 | 36247 |
| 57.23583 | 16 | 143.6713 | 30 | 17.39369 | 13 | 103.9582 | 34 | LMO2 | None | Blood | GSM10978 | 1171 | 36248 |
| 51.34314 | 52 | 121.5647 | 85 | 68.96309 | 94 | 104.011 | 119 | GATA2 | None | Blood | GSM10978 | 11550 | 36250 |
| 1117.25 | 910 | 1050.839 | 1009 | 906.5644 | 1211 | 978.4346 | 1523 | NKX3-1 | LNCaP | Prostate | GSM98964 | 69449 | 36252 |
| 1215.998 | 668 | 1273.762 | 877 | 882.9707 | 628 | 819.2114 | 616 | BRD4 | None | Embryo | GSM82338 | 23272 | 36299 |
| 1142.024 | 517 | 1160.269 | 644 | 849.7109 | 501 | 807.5421 | 511 | BRD4 | None | Blood | GSM82337 | 18743 | 36300 |
| 1201.003 | 372 | 1204.012 | 454 | 916.0252 | 347 | 878.321 | 355 | BRD4 | None | Blood | GSM82337 | 9800 | 36301 |
| 342.8176 | 131 | 618.9804 | 196 | 225.3393 | 162 | 185.5628 | 161 | RELA | GM19099 | Blood | GSM48628 | 13063 | 36303 |
| 433.3871 | 98 | 910.9543 | 164 | 342.6357 | 123 | 435.7031 | 148 | RELA | GM12892 | Blood | GSM48629 | 5278 | 36310 |
| 851.0694 | 176 | 977.9582 | 225 | 443.5188 | 178 | 472.4189 | 196 | RELA | GM15510 | Blood | GSM48628 | 8645 | 36311 |
| 1018.112 | 201 | 1023.696 | 250 | 601.6656 | 205 | 487.7743 | 197 | RELA | GM12892 | Blood | GSM48629 | 8481 | 36312 |
| 885.0355 | 418 | 896.6262 | 526 | 656.5958 | 466 | 592.4261 | 447 | RELA | GM19099 | Blood | GSM48628 | 33165 | 36314 |

| | | | | | | | | | | | | | |
|---|---|---|---|---|---|---|---|---|---|---|---|---|---|
| 803.1289 | 142 | 1044.911 | 184 | 435.0191 | 142 | 463.0944 | 156 | RELA | GM10847 | Blood | GSM48629 | 5522 | 36316 |
| 930.6728 | 265 | 943.9387 | 340 | 628.5888 | 283 | 552.6915 | 286 | RELA | GM12892 | Blood | GSM48629 | 15828 | 36317 |
| 346.7595 | 69 | 542.2689 | 98 | 172.729 | 67 | 260.0792 | 88 | RELA | GM15510 | Blood | GSM48628 | 3101 | 36318 |
| 1055.404 | 319 | 1076.006 | 410 | 807.1135 | 324 | 777.1901 | 339 | RELA | GM15510 | Blood | GSM48628 | 13032 | 36320 |
| 1075.777 | 350 | 1110.838 | 468 | 826.4551 | 359 | 750.7933 | 328 | RELA | GM15510 | Blood | GSM48629 | 13831 | 36321 |
| 295.751 | 50 | 614.5082 | 85 | 210.2993 | 57 | 268.0495 | 69 | RELA | GM15510 | Blood | GSM48628 | 1781 | 36322 |
| 936.3357 | 137 | 1131.331 | 176 | 454.9207 | 125 | 463.9042 | 134 | RELA | GM10847 | Blood | GSM48629 | 3987 | 36323 |
| 953.615 | 275 | 992.5891 | 368 | 738.5363 | 299 | 637.4722 | 298 | RELA | GM19099 | Blood | GSM48628 | 15002 | 36324 |
| 150.3921 | 28 | 207.0813 | 37 | 147.1687 | 38 | 31.08382 | 18 | CHD1 | A549 | Lung | GSM87464 | 1044 | 36348 |
| 750.1032 | 124 | 967.0111 | 157 | 561.8916 | 145 | 477.4876 | 141 | PTPN11 | A549 | Lung | GSM87463 | 4224 | 36349 |
| 1197.594 | 586 | 1243.998 | 763 | 883.4309 | 553 | 862.7663 | 594 | POLR2B | IMR90 | Lung | GSM93840 | 18653 | 36356 |
| 1185.893 | 248 | 1194.104 | 311 | 924.5326 | 242 | 900.051 | 253 | GTF2B | IMR90 | Lung | GSM93841 | 6282 | 36362 |
| 189.0637 | 53 | 156.0317 | 53 | 60.68404 | 45 | 74.47282 | 53 | TAL1 | None | Lymph Noc | GSM83800 | 3778 | 36366 |
| 142.5858 | 41 | 151.9068 | 47 | 61.86559 | 40 | 68.48419 | 45 | TAL1 | None | Lymph Noc | GSM83800 | 3022 | 36368 |
| 294.7699 | 184 | 482.1997 | 257 | 202.2796 | 248 | 264.6629 | 297 | CTCF | None | Mammary | GSM64637 | 29318 | 36573 |
| 394.8924 | 199 | 634.0745 | 277 | 304.1884 | 273 | 335.8994 | 307 | CTCF | GM12878 | Blood | GSM64631 | 26519 | 36575 |
| 271.6771 | 250 | 448.36 | 345 | 201.2464 | 364 | 246.4133 | 424 | CTCF | H1 | Embryo | GSM64633 | 55396 | 36578 |
| 421.3399 | 303 | 636.0534 | 415 | 305.6161 | 426 | 378.6615 | 499 | CTCF | HepG2 | Liver | GSM64635 | 56816 | 36602 |
| 477.9142 | 278 | 673.8576 | 371 | 292.7845 | 359 | 348.2285 | 415 | CTCF | None | Mammary | GSM64637 | 43876 | 36610 |
| 376.7591 | 226 | 615.7012 | 315 | 269.8005 | 309 | 325.0335 | 359 | CTCF | HepG2 | Liver | GSM64635 | 35832 | 36612 |
| 470.9323 | 257 | 710.5476 | 365 | 354.7172 | 353 | 409.6149 | 404 | CTCF | GM12878 | Blood | GSM64631 | 37249 | 36616 |
| 554.7971 | 403 | 650.5335 | 555 | 400.7705 | 570 | 446.6626 | 645 | CTCF | H1 | Embryo | GSM64633 | 80533 | 36632 |
| 465.5852 | 274 | 667.6065 | 362 | 341.5494 | 376 | 391.0976 | 429 | CTCF | None | Skeletal Mu | GSM64639 | 42424 | 36636 |
| 609.3414 | 351 | 689.9873 | 474 | 437.8408 | 481 | 501.7293 | 549 | CTCF | None | Skeletal Mu | GSM64639 | 56940 | 36650 |
| 0.404921 | 12 | 0.287751 | 14 | 1.784186 | 31 | 17.68132 | 59 | CDX2 | COLO-320 | Colon | GSM74303 | 10082 | 36712 |
| 357.1472 | 162 | 449.225 | 202 | 162.0333 | 178 | 178.4196 | 200 | ZNF76 | FLP76 | Embryonic | GSM95905 | 19437 | 36764 |
| 956.9973 | 609 | 977.9413 | 766 | 704.9409 | 669 | 651.6351 | 672 | NOTCH1 | HPB-ALL | Blood | GSM95905 | 42227 | 36765 |
| 542.7946 | 84 | 806.7469 | 116 | 200.4945 | 66 | 257.2207 | 80 | ZNF143 | HEK293T | Embryonic | GSM95904 | 2469 | 36766 |
| 694.0484 | 100 | 1104.773 | 144 | 247.7194 | 76 | 356.1744 | 98 | ZNF143 | HPB-ALL | Blood | GSM95904 | 2602 | 36768 |
| 481.5632 | 171 | 651.0705 | 221 | 253.3653 | 191 | 274.9023 | 213 | ZNF143 | FLP143HA | Embryonic | GSM95905 | 16112 | 36770 |
| 527.099 | 73 | 884.371 | 109 | 201.6268 | 57 | 320.9928 | 78 | ZNF143 | HeLa-S3 | Cervix | GSM95904 | 1809 | 36771 |
| 789.8513 | 150 | 1047.411 | 210 | 365.742 | 140 | 315.6416 | 139 | ZNF143 | K562 | Bone Marr | GSM95904 | 6330 | 36774 |
| 1114.014 | 167 | 1120.437 | 208 | 458.8979 | 139 | 448.3215 | 146 | ZNF143 | HeLa-S3 | Cervix | GSM95904 | 4802 | 36775 |
| 148.1809 | 96 | 223.247 | 130 | 76.46073 | 117 | 100.0492 | 140 | ZNF143 | FLP143HA | Embryonic | GSM95905 | 15472 | 36777 |
| 946.6771 | 276 | 944.1389 | 333 | 844.1548 | 418 | 870.9114 | 509 | GREB1 | MCF-7 | Breast | GSM10191 | 15507 | 36802 |
| 1049.287 | 805 | 1110.776 | 1033 | 833.2879 | 1029 | 812.614 | 1137 | NR2F2 | MCF-7 | Breast | GSM10191 | 60658 | 36804 |
| 958.4058 | 149 | 1015.174 | 168 | 988.3136 | 224 | 981.9434 | 250 | GREB1 | MCF-7 | Breast | GSM10191 | 4742 | 36806 |
| 1217.673 | 275 | 1245.095 | 362 | 954.9183 | 266 | 929.1854 | 283 | E2F4 | MCF-7 | Breast | GSM10191 | 6378 | 36807 |
| 133.0003 | 38 | 176.1462 | 49 | 241.8718 | 75 | 448.1315 | 111 | GREB1 | MCF-7 | Breast | GSM10191 | 2701 | 36810 |
| 918.6142 | 203 | 924.6457 | 238 | 864.188 | 329 | 871.6414 | 381 | GREB1 | MCF-7 | Breast | GSM10191 | 11012 | 36811 |
| 652.0105 | 79 | 919.1993 | 106 | 205.1383 | 54 | 407.2277 | 84 | E2F4 | MCF-7 | Breast | GSM10191 | 1606 | 36812 |
| 472.7518 | 130 | 487.0358 | 146 | 847.4742 | 260 | 860.5329 | 309 | TLE3 | MCF-7 | Breast | GSM10191 | 8807 | 36814 |
| 956.9836 | 579 | 991.345 | 745 | 800.1008 | 787 | 798.0723 | 895 | ESR1 | MCF-7 | Breast | GSM10191 | 43001 | 36819 |
| 332.623 | 77 | 496.1688 | 106 | 331.6984 | 109 | 469.9174 | 139 | RXRA | MCF-7 | Breast | GSM10191 | 4217 | 36820 |
| 604.5411 | 115 | 936.5014 | 162 | 549.8381 | 152 | 689.6066 | 182 | KLF4 | MCF-7 | Breast | GSM10191 | 4864 | 36821 |
| 1010.917 | 165 | 1028.968 | 182 | 948.8493 | 234 | 967.2065 | 280 | GREB1 | MCF-7 | Breast | GSM10191 | 5700 | 36822 |
| 845.8332 | 633 | 846.9417 | 768 | 755.0528 | 986 | 767.9452 | 1153 | ESR1 | MCF-7 | Breast | GSM10191 | 72324 | 36825 |
| 770.9027 | 437 | 721.2175 | 475 | 700.6968 | 712 | 727.5797 | 865 | ESR1 | MCF-7 | Breast | GSM10191 | 50824 | 36834 |
| 1110.767 | 699 | 1138.048 | 872 | 871.2186 | 812 | 838.6095 | 873 | TCF7L2 | MDA-MB-4 | Mammary | GSM10990 | 34142 | 36841 |
| 1232.226 | 1066 | 1237.99 | 1267 | 883.9853 | 1288 | 878.2393 | 1472 | FOXA1 | MDA-MB-4 | Mammary | GSM10990 | 82968 | 36842 |
| 1160.255 | 311 | 1180.247 | 399 | 888.0615 | 291 | 868.129 | 314 | TCF7L2 | MDA-MB-4 | Mammary | GSM10990 | 8685 | 36843 |
| 1148.789 | 705 | 1191.504 | 897 | 842.0015 | 703 | 805.75 | 745 | MYC | MDA-MB-4 | Mammary | GSM10990 | 29149 | 36844 |
| 1154.558 | 799 | 1178.042 | 986 | 874.1445 | 902 | 853.5462 | 991 | AR | MDA-MB-4 | Mammary | GSM10990 | 39247 | 36845 |
| 916.7302 | 653 | 909.894 | 778 | 656.1935 | 728 | 638.8441 | 799 | EP300 | IMR90 | Lung | GSM92909 | 60044 | 36878 |
| 141.2952 | 65 | 199.2658 | 86 | 79.07802 | 77 | 117.8853 | 98 | EP300 | SKNO-1 | Bone Marr | GSM83586 | 7655 | 36908 |
| 948.3436 | 461 | 987.3556 | 613 | 709.841 | 509 | 688.1464 | 549 | TFAP2A | None | Cranial | GSM90277 | 31256 | 36958 |
| 143.9641 | 92 | 199.3989 | 120 | 123.4608 | 135 | 158.5181 | 161 | FLI1 | NB4 | Blood | GSM86315 | 14580 | 37055 |
| 1112.084 | 638 | 1140.854 | 807 | 806.7106 | 634 | 737.9589 | 612 | FLI1 | U937 | Pleura | GSM86315 | 29323 | 37056 |
| 1151.773 | 637 | 1195.652 | 817 | 851.6101 | 632 | 782.9528 | 610 | FLI1 | U937 | Pleura | GSM86315 | 25448 | 37057 |
| 469.2651 | 55 | 662.2254 | 74 | 319.0602 | 58 | 938.6554 | 117 | AR | LNCaP | Prostate | GSM98064 | 1069 | 37089 |
| 1038.728 | 476 | 1001.909 | 534 | 938.2618 | 722 | 956.8348 | 853 | NR3C1 | VCaP | Prostate | GSM98066 | 24071 | 37091 |
| 736.833 | 98 | 837.4697 | 115 | 496.4157 | 105 | 1089.135 | 181 | AR | LNCaP | Prostate | GSM98064 | 2392 | 37092 |
| 897.7608 | 335 | 880.9681 | 391 | 821.2348 | 537 | 861.6305 | 676 | NR3C1 | LNCaP | Prostate | GSM98066 | 23328 | 37093 |
| 803.2544 | 755 | 739.6915 | 821 | 673.1784 | 1106 | 723.2762 | 1365 | AR | VCaP | Prostate | GSM98065 | 115772 | 37094 |
| 954.4754 | 361 | 905.838 | 389 | 796.049 | 475 | 897.7943 | 690 | AR | LNCaP | Prostate | GSM98066 | 21467 | 37096 |
| 742.7765 | 466 | 721.1772 | 538 | 665.251 | 748 | 731.0015 | 979 | AR | VCaP | Prostate | GSM98065 | 61769 | 37099 |
| 802.6222 | 390 | 807.4916 | 478 | 645.866 | 518 | 650.0464 | 597 | HNF4A | Caco-2 | Colon | GSM57522 | 39336 | 371 |
| 892.2582 | 415 | 865.2089 | 475 | 781.4776 | 620 | 851.1262 | 826 | NR3C1 | LNCaP | Prostate | GSM98066 | 32139 | 37100 |
| 1017.249 | 325 | 943.219 | 323 | 879.7071 | 443 | 934.2423 | 583 | NR3C1 | LNCaP | Prostate | GSM98065 | 15091 | 37101 |
| 712.0426 | 101 | 917.2912 | 128 | 600.5729 | 124 | 1063.128 | 189 | AR | LNCaP | Prostate | GSM98064 | 2743 | 37104 |
| 1010.277 | 281 | 972.7713 | 310 | 865.5032 | 375 | 960.6793 | 549 | AR | LNCaP | Prostate | GSM98066 | 12851 | 37106 |
| 47.07997 | 15 | 54.09297 | 18 | 24.08023 | 16 | 123.4272 | 39 | HSF1 | MCF-10A | Breast | GSM95186 | 1286 | 37108 |

| | | | | | | | | | | | | | |
|---|---|---|---|---|---|---|---|---|---|---|---|---|---|
| 1001.438 | 157 | 999.6028 | 171 | 467.2246 | 142 | 455.6488 | 149 | HSF1 | NCI-H838 | Lung | GSM95189 | 5037 | 37110 |
| 54.55337 | 17 | 49.8555 | 18 | 72.98522 | 29 | 229.3687 | 57 | HSF1 | SKBR-3 | Mammary | GSM95188 | 1393 | 37111 |
| 113.1789 | 30 | 218.4151 | 48 | 122.1003 | 45 | 176.2594 | 58 | HSF1 | BT-20 | Mammary | GSM95189 | 1983 | 37116 |
| 1019.5 | 171 | 1038.846 | 197 | 511.5611 | 163 | 568.0535 | 183 | HSF1 | NCI-H838 | Lung | GSM95189 | 6136 | 37123 |
| 203.574 | 63 | 269.175 | 81 | 124.4538 | 73 | 192.9382 | 96 | HSF1 | BT-20 | Mammary | GSM95189 | 5005 | 37126 |
| 10.93173 | 10 | 36.44324 | 20 | 13.54005 | 18 | 78.76022 | 42 | HSF1 | None | Mammary | GSM95189 | 2336 | 37130 |
| 46.8944 | 22 | 51.99866 | 26 | 23.58601 | 25 | 115.3404 | 55 | HSF1 | None | Mammary | GSM95189 | 2736 | 37134 |
| 148.2897 | 32 | 154.3517 | 36 | 67.28921 | 30 | 203.3081 | 57 | HSF1 | HT29 | Colon | GSM95188 | 1639 | 37144 |
| 58.02955 | 26 | 70.03629 | 32 | 33.44463 | 31 | 113.9777 | 58 | HSF1 | MCF-10A | Breast | GSM95186 | 3048 | 37153 |
| 18.65891 | 8 | 8.406715 | 6 | 5.325118 | 7 | 117.8091 | 34 | HSF1 | None | Mammary | GSM95185 | 1021 | 37156 |
| 1069.73 | 684 | 1088.861 | 849 | 752.7393 | 674 | 712.5426 | 704 | MYC | MM1.S | Blood | GSM89410 | 37620 | 37169 |
| 926.4274 | 774 | 959.0318 | 965 | 666.3222 | 897 | 647.6767 | 989 | MAX | H2171 | Lung | GSM89410 | 76811 | 37173 |
| 1018.74 | 136 | 1161.565 | 163 | 585.6844 | 133 | 565.1513 | 138 | MED1 | MM1.S | Blood | GSM89410 | 3237 | 37174 |
| 943.4087 | 745 | 975.8972 | 933 | 688.262 | 862 | 656.1019 | 922 | MYC | H2171 | Lung | GSM89410 | 66871 | 37175 |
| 153.9883 | 509 | 176.4291 | 616 | 125.8861 | 862 | 111.2798 | 941 | STAT4 | None | Blood | GSM94561 | 250235 | 37177 |
| 67.87844 | 423 | 90.41407 | 530 | 90.09017 | 806 | 101.7729 | 917 | IRF5 | None | Blood | GSM94560 | 246590 | 37185 |
| 589.3551 | 185 | 657.1451 | 217 | 443.3539 | 237 | 473.4478 | 262 | HNF4A | Caco-2 | Colon | GSM57522 | 15247 | 372 |
| 1289.774 | 363 | 1315.111 | 465 | 1005.201 | 334 | 984.0781 | 358 | CREB1 | A549 | Lung | GSM10107 | 7041 | 37305 |
| 946.5712 | 242 | 969.3679 | 316 | 608.6265 | 256 | 473.8456 | 244 | REST | GM12878 | Blood | GSM10107 | 13109 | 37311 |
| 1211.257 | 463 | 1221.662 | 574 | 936.3867 | 462 | 907.3429 | 489 | CREB1 | A549 | Lung | GSM10107 | 13072 | 37334 |
| 1190.197 | 734 | 1210.011 | 901 | 871.9356 | 743 | 848.6974 | 801 | CREB1 | A549 | Lung | GSM10107 | 30189 | 37349 |
| 1040.474 | 576 | 1049.402 | 708 | 775.4821 | 616 | 769.5785 | 692 | JUN | A549 | Lung | GSM10107 | 31560 | 37354 |
| 930.7652 | 548 | 903.9886 | 628 | 735.1035 | 680 | 745.3302 | 798 | FOXA2 | A549 | Lung | GSM10107 | 42927 | 37366 |
| 102.8032 | 52 | 187.1309 | 78 | 104.0035 | 80 | 143.478 | 99 | RUNX1 | Jurkat E6-1 | Blood | GSM72217 | 6756 | 37384 |
| 23.88568 | 12 | 41.92721 | 18 | 27.75915 | 20 | 64.42693 | 32 | RUNX1 | Jurkat E6-1 | Blood | GSM72216 | 1696 | 37388 |
| 15.58071 | 11 | 16.86664 | 13 | 4.518871 | 11 | 34.77028 | 27 | TCF12 | Jurkat E6-1 | Blood | GSM72216 | 2075 | 37389 |
| 6.502573 | 13 | 13.72184 | 20 | 12.24355 | 28 | 46.6618 | 50 | TAL1 | Jurkat E6-1 | Blood | GSM72216 | 4695 | 37390 |
| 19.07545 | 24 | 16.20757 | 26 | 11.49002 | 34 | 52.25962 | 63 | GATA3 | Jurkat E6-1 | Blood | GSM72216 | 6386 | 37391 |
| 523.682 | 186 | 820.4455 | 264 | 628.4539 | 298 | 731.9318 | 382 | REPIN1 | DLD-1 | Colon | GSM11415 | 17499 | 37565 |
| 904.2672 | 550 | 895.3637 | 653 | 618.9218 | 547 | 599.8961 | 593 | EP300 | ME-1 | None | GSM11223 | 43625 | 37576 |
| 1209.265 | 602 | 1253.77 | 782 | 887.2508 | 566 | 836.054 | 564 | ELF1 | ME-1 | None | GSM11223 | 19241 | 37578 |
| 1178.499 | 250 | 1174.785 | 304 | 911.3902 | 235 | 819.5926 | 226 | FLI1 | ME-1 | None | GSM11223 | 6341 | 37579 |
| 1093.708 | 791 | 1130.52 | 992 | 760.1629 | 805 | 725.3052 | 858 | ERG | ME-1 | None | GSM11223 | 48905 | 37583 |
| 1248.564 | 459 | 1287.607 | 598 | 941.8778 | 412 | 904.8556 | 424 | TBP | ME-1 | None | GSM11223 | 10704 | 37584 |
| 1121.611 | 324 | 1148.082 | 416 | 839.7541 | 297 | 813.5824 | 311 | GATA2 | ME-1 | None | GSM11223 | 10427 | 37587 |
| 1081.603 | 324 | 1096.311 | 405 | 793.486 | 293 | 741.4253 | 288 | SPI1 | ME-1 | None | GSM11223 | 11943 | 37588 |
| 1181.024 | 308 | 1215.015 | 405 | 946.4802 | 322 | 893.9591 | 314 | MYH11 | None | None | GSM11223 | 8100 | 37589 |
| 1054.191 | 453 | 1094.553 | 595 | 780.0099 | 445 | 745.4984 | 461 | TAL1 | ME-1 | None | GSM11223 | 20662 | 37596 |
| 1024.391 | 551 | 1053.914 | 702 | 730.2226 | 538 | 717.4021 | 596 | HDAC1 | ME-1 | None | GSM11223 | 30605 | 37597 |
| 1096.44 | 573 | 1121.192 | 716 | 785.7576 | 539 | 746.7998 | 555 | TCF12 | ME-1 | None | GSM11223 | 25213 | 37602 |
| 959.0223 | 199 | 989.8241 | 261 | 542.2835 | 210 | 511.2724 | 218 | CBFB | U937 | Pleura | GSM11223 | 9905 | 37603 |
| 1054.362 | 687 | 1096.11 | 878 | 771.3162 | 736 | 727.9503 | 761 | CBFB | ME-1 | None | GSM11223 | 41875 | 37604 |
| 1197.479 | 266 | 1247.133 | 372 | 932.2616 | 252 | 901.3034 | 264 | CBFB | None | None | GSM11223 | 6525 | 37606 |
| 1209.764 | 664 | 1234.337 | 830 | 880.9573 | 635 | 823.5617 | 628 | CBFB | ME-1 | None | GSM11223 | 23225 | 37607 |
| 1110.846 | 471 | 1119.443 | 584 | 811.981 | 444 | 781.5699 | 464 | MYH11 | ME-1 | None | GSM11223 | 18476 | 37608 |
| 1050.843 | 587 | 1102.355 | 776 | 760.9715 | 590 | 716.2911 | 602 | RUNX1 | ME-1 | None | GSM11223 | 31162 | 37619 |
| 1184.725 | 590 | 1197.748 | 732 | 880.4491 | 573 | 822.204 | 564 | CBFB | U937 | Pleura | GSM11223 | 20356 | 37621 |
| 930.9427 | 270 | 966.5572 | 362 | 655.6682 | 290 | 542.3858 | 285 | MYH11 | ME-1 | None | GSM11223 | 16022 | 37623 |
| 772.7598 | 284 | 735.0473 | 311 | 264.3078 | 269 | 225.218 | 275 | MYH11 | U937 | Pleura | GSM11223 | 27562 | 37625 |
| 1194.463 | 759 | 1275.079 | 1003 | 864.5233 | 754 | 823.996 | 789 | ERG | MCF-7 | Breast | GSM72698 | 31742 | 37633 |
| 1112.369 | 393 | 1139.261 | 506 | 816.3607 | 358 | 781.546 | 369 | ERG | K562 | Bone Marr | GSM72698 | 14153 | 37635 |
| 1046.43 | 749 | 1072.925 | 930 | 719.0452 | 752 | 668.0057 | 768 | RUNX1 | SKNO-1 | Bone Marr | GSM72697 | 51152 | 37638 |
| 1160.27 | 561 | 1207.503 | 742 | 862.1768 | 545 | 816.0517 | 552 | ERG | K562 | Bone Marr | GSM72698 | 20278 | 37639 |
| 1200.247 | 503 | 1240.544 | 663 | 899.3199 | 479 | 869.2377 | 502 | CBFB | SKNO-1 | Bone Marr | GSM72697 | 15203 | 37640 |
| 228.6593 | 70 | 417.2508 | 107 | 152.0614 | 84 | 194.7087 | 101 | EP300 | None | Umbilical V | GSM10096 | 5405 | 37921 |
| 105.0092 | 28 | 180.9261 | 42 | 70.22941 | 33 | 124.3581 | 47 | EP300 | None | Umbilical V | GSM10096 | 1873 | 37924 |
| 750.1063 | 198 | 900.5741 | 260 | 462.3836 | 226 | 452.8481 | 240 | EP300 | None | Umbilical V | GSM10096 | 13312 | 37926 |
| 1054.122 | 252 | 1079.669 | 329 | 758.4488 | 247 | 661.5413 | 246 | ETS1 | None | Umbilical V | GSM10096 | 9735 | 37927 |
| 634.5617 | 178 | 717.7383 | 210 | 319.8315 | 187 | 338.5862 | 206 | EP300 | None | Umbilical V | GSM10096 | 12768 | 37930 |
| 451.5354 | 118 | 469.0725 | 133 | 201.4564 | 115 | 203.6994 | 124 | EP300 | None | Umbilical V | GSM10096 | 7598 | 37931 |
| 136.6255 | 32 | 128.1067 | 34 | 72.97681 | 33 | 161.9704 | 53 | EP300 | None | Umbilical V | GSM10096 | 1803 | 37932 |
| 1077.972 | 240 | 1093.194 | 306 | 740.5622 | 230 | 670.925 | 233 | ETS1 | None | Umbilical V | GSM10096 | 8560 | 37934 |
| 977.3537 | 364 | 1011.333 | 481 | 726.7769 | 370 | 708.364 | 401 | ETS1 | None | Umbilical V | GSM10096 | 19813 | 37936 |
| 151.6244 | 49 | 218.5379 | 66 | 92.70019 | 57 | 156.2831 | 78 | EP300 | None | Umbilical V | GSM10096 | 4039 | 37941 |
| 410.5204 | 108 | 597.2327 | 146 | 268.5424 | 126 | 295.34 | 141 | EP300 | None | Umbilical V | GSM10096 | 7023 | 37942 |
| 990.4906 | 453 | 1000.494 | 564 | 725.6846 | 462 | 699.1579 | 489 | ETS1 | None | Umbilical V | GSM10096 | 26034 | 37945 |
| 517.0034 | 175 | 710.1795 | 228 | 384.4545 | 225 | 345.7654 | 231 | EP300 | NB4 | Blood | GSM74952 | 15567 | 37976 |
| 12.98286 | 15 | 16.56256 | 19 | 4.270487 | 17 | 55.16453 | 47 | SSRP1 | HT1080 | Connective | GSM11035 | 3491 | 38073 |
| 310.3561 | 59 | 314.4807 | 65 | 159.0067 | 57 | 155.3259 | 60 | SSRP1 | HT1080 | Connective | GSM11035 | 2319 | 38077 |
| 441.5218 | 89 | 461.6979 | 100 | 175.5728 | 77 | 225.3459 | 93 | SSRP1 | HT1080 | Connective | GSM11035 | 3869 | 38078 |
| 103.4942 | 39 | 115.8313 | 46 | 40.48824 | 38 | 79.90986 | 55 | SSRP1 | HT1080 | Connective | GSM11035 | 3629 | 38081 |
| 13.49412 | 16 | 5.345641 | 13 | 14.82467 | 27 | 49.9406 | 47 | SSRP1 | HT1080 | Connective | GSM11035 | 3698 | 38083 |

| | | | | | | | | | | | | | |
|---|---|---|---|---|---|---|---|---|---|---|---|---|---|
| 29.47222 | 20 | 33.59323 | 24 | 3.834522 | 15 | 67.82384 | 48 | SSRP1 | HT1080 | Connective | GSM11035 | 3109 | 38084 |
| 1183.663 | 708 | 1229.732 | 910 | 872.1549 | 721 | 852.4818 | 792 | MYC | HeLa | Cervix | GSM10589 | 29721 | 38115 |
| 1093.941 | 807 | 1143.455 | 1027 | 810.6199 | 914 | 805.8371 | 1040 | MAX | HeLa | Cervix | GSM10589 | 53042 | 38119 |
| 137.2805 | 45 | 369.8463 | 85 | 92.05534 | 53 | 103.1245 | 60 | SUZ12 | None | Bone Marr | GSM10312 | 3054 | 38163 |
| 271.162 | 82 | 417.0664 | 114 | 110.0337 | 78 | 101.2125 | 81 | REST | None | Bone Marr | GSM10312 | 5959 | 38164 |
| 464.2123 | 185 | 490.3663 | 212 | 247.7006 | 211 | 296.2816 | 245 | SND1 | Mutu III | None | GSM11537 | 19079 | 38173 |
| 678.0626 | 141 | 1028.507 | 212 | 496.698 | 169 | 411.9469 | 164 | MYC | FB0167P | None | GSM80876 | 6972 | 38182 |
| 1103.512 | 806 | 1141.205 | 1007 | 828.6758 | 926 | 821.8263 | 1051 | MYC | MCF-7 | Breast | GSM80875 | 51251 | 38183 |
| 730.5742 | 461 | 760.1667 | 598 | 539.9945 | 573 | 527.5422 | 633 | CTCF | H54 | Brain | GSM80877 | 63367 | 38185 |
| 830.3381 | 461 | 877.1483 | 620 | 618.2246 | 543 | 597.7603 | 588 | CTCF | FB8470 | None | GSM80875 | 46179 | 38188 |
| 473.0836 | 71 | 630.7395 | 92 | 323.5436 | 78 | 312.6809 | 81 | MYC | FB8470 | None | GSM80876 | 2111 | 38189 |
| 190.6939 | 54 | 168.4745 | 56 | 58.29235 | 45 | 77.83307 | 55 | MYC | FB0167P | None | GSM80876 | 3896 | 38190 |
| 714.0529 | 429 | 766.6743 | 587 | 539.6367 | 554 | 532.4652 | 619 | CTCF | H54 | Brain | GSM80877 | 60717 | 38192 |
| 1161.873 | 907 | 1183.091 | 1107 | 832.355 | 1007 | 795.4476 | 1081 | MYC | MCF-7 | Breast | GSM80875 | 58842 | 38193 |
| 817.2897 | 582 | 872.7618 | 779 | 622.3207 | 739 | 590.0566 | 786 | CTCF | MCF-7 | Breast | GSM80875 | 69561 | 38194 |
| 371.4518 | 58 | 525.7925 | 78 | 218.9561 | 59 | 276.1386 | 71 | MYC | FB8470 | None | GSM80876 | 1836 | 38195 |
| 747.6736 | 379 | 797.2515 | 520 | 565.6213 | 472 | 560.9342 | 531 | CTCF | FB0167P | None | GSM80876 | 45967 | 38196 |
| 723.3774 | 350 | 771.6009 | 484 | 493.0861 | 440 | 540.8586 | 497 | CTCF | FB0167P | None | GSM80876 | 45075 | 38197 |
| 667.8404 | 131 | 853.1933 | 165 | 511.4551 | 159 | 431.062 | 155 | MYC | H54 | Brain | GSM80877 | 5866 | 38200 |
| 782.0128 | 574 | 816.7292 | 744 | 597.9866 | 743 | 563.9179 | 790 | CTCF | MCF-7 | Breast | GSM80875 | 76228 | 38203 |
| 791.8052 | 402 | 840.4378 | 547 | 606.1861 | 498 | 578.4641 | 530 | CTCF | FB8470 | None | GSM80876 | 43135 | 38204 |
| 569.9785 | 103 | 828.1872 | 140 | 357.2085 | 111 | 430.0637 | 130 | MYC | H54 | Brain | GSM80877 | 4038 | 38207 |
| 879.9895 | 706 | 918.8751 | 897 | 649.1906 | 840 | 624.3634 | 914 | MAX | H2171 | Lung | GSM10086 | 71172 | 38287 |
| 1057.228 | 562 | 1096.592 | 729 | 782.2665 | 580 | 731.2211 | 584 | MYC | H2171 | Lung | GSM10086 | 29182 | 38288 |
| 1061.818 | 702 | 1076.076 | 862 | 778.2644 | 743 | 751.5001 | 800 | MAX | H128 | Lung | GSM10086 | 38668 | 38289 |
| 1232.656 | 256 | 1237.83 | 312 | 949.6869 | 229 | 944.617 | 255 | MYC | H128 | Lung | GSM10086 | 5443 | 38299 |
| 1046.758 | 312 | 1040.549 | 372 | 776.9834 | 292 | 747.0354 | 305 | BRD4 | None | Blood | GSM10229 | 12158 | 38340 |
| 1193.539 | 359 | 1191.909 | 438 | 905.0694 | 324 | 874.6481 | 338 | BRD4 | None | Blood | GSM10229 | 8979 | 38389 |
| 114.7481 | 59 | 183.9166 | 83 | 80.44993 | 78 | 59.63109 | 75 | RUNX1 | CMK | Blood | GSM61033 | 7789 | 384 |
| 259.84 | 103 | 219.3675 | 106 | 490.5531 | 206 | 781.3635 | 280 | ESR1 | MCF-7 | Breast | GSM85469 | 10426 | 38474 |
| 769.3603 | 128 | 1144.341 | 203 | 526.2908 | 143 | 521.5076 | 151 | RBL2 | IMR90 | Lung | GSM80240 | 4482 | 38488 |
| 950.6002 | 274 | 941.0901 | 325 | 556.0823 | 259 | 452.1675 | 253 | RBL1 | IMR90 | Lung | GSM80240 | 13808 | 38496 |
| 49.77219 | 15 | 57.29268 | 18 | 16.80691 | 13 | 112.937 | 36 | HSF1 | K562 | Bone Marr | GSM10657 | 1192 | 38582 |
| 1277.519 | 634 | 1335.93 | 828 | 944.6602 | 577 | 913.2802 | 608 | TAF3 | None | Colon | GSM10650 | 15834 | 38585 |
| 568.0632 | 152 | 600.3815 | 173 | 229.5627 | 143 | 355.5897 | 187 | E2F7 | IMR90 | Lung | GSM99166 | 10215 | 38586 |
| 176.315 | 30 | 206.2431 | 36 | 144.7635 | 37 | 239.1108 | 52 | E2F7 | IMR90 | Lung | GSM99165 | 1076 | 38588 |
| 465.383 | 148 | 550.9383 | 179 | 222.4729 | 155 | 268.9444 | 181 | E2F7 | IMR90 | Lung | GSM99166 | 12134 | 38589 |
| 670.6572 | 158 | 735.6173 | 183 | 276.0191 | 146 | 446.7939 | 196 | E2F7 | IMR90 | Lung | GSM99166 | 9070 | 38590 |
| 832.8393 | 295 | 839.3737 | 365 | 483.2442 | 317 | 576.6791 | 368 | E2F7 | IMR90 | Lung | GSM99166 | 24605 | 38591 |
| 1159.067 | 180 | 1133.808 | 202 | 708.6902 | 169 | 875.8673 | 201 | FOXD2 | LoVo | Colon | GSM12087 | 4385 | 38635 |
| 1174.712 | 274 | 1170.293 | 330 | 958.3536 | 300 | 978.5119 | 362 | ETV7 | LoVo | Colon | GSM12087 | 7196 | 38647 |
| 1325.114 | 183 | 1342.598 | 230 | 793.1509 | 145 | 914.6414 | 166 | NFYA | LoVo | Colon | GSM12087 | 2594 | 38650 |
| 1121.936 | 580 | 1161.223 | 744 | 877.1774 | 636 | 869.6443 | 712 | MED12 | LoVo | Colon | GSM12086 | 23501 | 38653 |
| 1245.602 | 511 | 1274.935 | 652 | 919.6612 | 444 | 894.3652 | 476 | NFATC1 | None | Umbilical V | GSM12083 | 12373 | 38654 |
| 738.002 | 117 | 875.3852 | 141 | 568.6236 | 138 | 717.2837 | 166 | EHF | LoVo | Colon | GSM12086 | 3701 | 38657 |
| 1172.105 | 447 | 1201.46 | 569 | 923.9637 | 461 | 924.9299 | 531 | SOX2 | LoVo | Colon | GSM12087 | 13030 | 38658 |
| 679.2437 | 136 | 967.9309 | 182 | 700.4768 | 192 | 893.3789 | 244 | CAMTA2 | LoVo | Colon | GSM12085 | 5947 | 38661 |
| 1056.497 | 559 | 1092.019 | 714 | 841.5152 | 654 | 856.4183 | 772 | MYC | LoVo | Colon | GSM12086 | 26922 | 38663 |
| 1034.136 | 153 | 1143.582 | 204 | 587.2892 | 151 | 634.9236 | 167 | ZBTB7B | LoVo | Colon | GSM12087 | 4252 | 38666 |
| 391.8591 | 86 | 521.6475 | 111 | 280.3515 | 102 | 428.3154 | 135 | GLIS1 | LoVo | Colon | GSM12087 | 4396 | 38673 |
| 322.2998 | 52 | 185.8237 | 41 | 245.8013 | 61 | 607.0682 | 108 | HOXA13 | LoVo | Colon | GSM12087 | 1676 | 38678 |
| 1049.625 | 216 | 1047.679 | 266 | 897.2302 | 280 | 913.2312 | 333 | LHX2 | LoVo | Colon | GSM12086 | 7878 | 38679 |
| 302.8299 | 41 | 234.4398 | 38 | 189.5431 | 42 | 375.6006 | 66 | PHF20 | LoVo | Colon | GSM12087 | 1015 | 38681 |
| 554.5602 | 76 | 594.3778 | 86 | 438.7579 | 89 | 651.9232 | 118 | EGR1 | LoVo | Colon | GSM12086 | 1853 | 38684 |
| 890.9255 | 693 | 920.3996 | 871 | 696.2554 | 891 | 687.2416 | 1003 | RAD21 | LoVo | Colon | GSM12086 | 71954 | 38691 |
| 1384.93 | 224 | 1385.48 | 279 | 1073.074 | 186 | 1094.559 | 221 | RBCK1 | LoVo | Colon | GSM12087 | 2791 | 38695 |
| 1230.781 | 262 | 1266.509 | 355 | 972.2961 | 253 | 953.4208 | 272 | E2F7 | LoVo | Colon | GSM12087 | 5530 | 38698 |
| 671.4939 | 99 | 706.1628 | 111 | 832.5114 | 152 | 1071.323 | 201 | HES1 | LoVo | Colon | GSM12086 | 2841 | 38702 |
| 1154.443 | 272 | 1152.267 | 333 | 915.3256 | 280 | 894.2959 | 302 | HINFP | LoVo | Colon | GSM12086 | 7791 | 38704 |
| 1225.236 | 158 | 1237.197 | 202 | 740.4336 | 151 | 962.2865 | 185 | VEZF1 | LoVo | Colon | GSM12088 | 3253 | 38706 |
| 305.6916 | 55 | 184.7946 | 45 | 140.6311 | 50 | 182.7582 | 61 | HOXA2 | LoVo | Colon | GSM12087 | 2083 | 38708 |
| 1060.785 | 419 | 1066.464 | 514 | 843.0402 | 470 | 842.9644 | 538 | ETS2 | LoVo | Colon | GSM12087 | 17272 | 38712 |
| 19.92948 | 13 | 73.07779 | 28 | 56.6499 | 33 | 199.2508 | 66 | HNF4A | LoVo | Colon | GSM12086 | 2274 | 38713 |
| 732.5265 | 90 | 1065.629 | 123 | 486.9204 | 94 | 564.8138 | 108 | YY1 | LoVo | Colon | GSM12086 | 1827 | 38716 |
| 381.2258 | 67 | 415.9213 | 77 | 182.6849 | 62 | 488.7211 | 112 | CREM | LoVo | Colon | GSM12087 | 2441 | 38717 |
| 1473.448 | 141 | 1491.386 | 184 | 1132.478 | 136 | 1225.919 | 157 | GLI2 | LoVo | Colon | GSM12086 | 1298 | 38718 |
| 1157.251 | 366 | 1150.425 | 433 | 920.1601 | 379 | 950.5592 | 467 | DLX1 | LoVo | Colon | GSM12087 | 10500 | 38719 |
| 1119.076 | 121 | 1236.292 | 139 | 562.9089 | 106 | 932.4393 | 150 | CREB3L4 | LoVo | Colon | GSM12087 | 2030 | 38721 |
| 1288.472 | 409 | 1306.997 | 512 | 1004.175 | 381 | 988.6859 | 416 | GMEB2 | LoVo | Colon | GSM12086 | 7978 | 38724 |
| 826.3194 | 154 | 1013.109 | 190 | 894.7939 | 223 | 933.9889 | 282 | HOXB13 | LoVo | Colon | GSM12086 | 6223 | 38729 |
| 1218.949 | 268 | 1266.216 | 366 | 978.5218 | 269 | 960.0054 | 291 | TFDP1 | LoVo | Colon | GSM12086 | 5709 | 38731 |
| 356.6954 | 54 | 400.1966 | 63 | 154.5222 | 46 | 454.0788 | 89 | NKX2-2 | LoVo | Colon | GSM12087 | 1590 | 38736 |

| | | | | | | | | | | | | | |
|---|---|---|---|---|---|---|---|---|---|---|---|---|---|
| 1130.107 | 232 | 1187.052 | 326 | 903.0712 | 241 | 881.4199 | 256 | MED1 | LoVo | Colon | GSM12086 | 6573 | 38737 |
| 1094.909 | 120 | 1281.016 | 152 | 570.9116 | 107 | 616.95 | 118 | ELF2 | LoVo | Colon | GSM12086 | 1971 | 38739 |
| 1249.914 | 116 | 1346.624 | 135 | 1060.245 | 136 | 1231.095 | 172 | GATA4 | LoVo | Colon | GSM12086 | 1472 | 38741 |
| 1242.946 | 467 | 1273.875 | 601 | 927.6364 | 410 | 914.7767 | 454 | NFATC1 | None | Umbilical V | GSM12083 | 10984 | 38742 |
| 329.3838 | 65 | 335.8403 | 72 | 274.2354 | 82 | 285.746 | 89 | JUN | LoVo | Colon | GSM12086 | 2804 | 38743 |
| 869.9201 | 84 | 1164.899 | 109 | 537.4113 | 81 | 568.2964 | 88 | RFX5 | LoVo | Colon | GSM12086 | 1087 | 38746 |
| 365.5635 | 49 | 506.3068 | 65 | 192.2631 | 45 | 288.8059 | 60 | ZNF407 | LoVo | Colon | GSM12087 | 1108 | 38750 |
| 996.5609 | 539 | 1017.824 | 680 | 797.3991 | 654 | 788.0205 | 732 | JUND | LoVo | Colon | GSM12087 | 30959 | 38758 |
| 352.8143 | 72 | 385.4242 | 83 | 336.8685 | 98 | 568.8441 | 138 | ETV5 | LoVo | Colon | GSM12087 | 3236 | 38759 |
| 998.4061 | 145 | 1121.612 | 177 | 427.1878 | 122 | 670.4871 | 165 | RFX2 | LoVo | Colon | GSM12086 | 3800 | 38761 |
| 105.7656 | 27 | 120.0339 | 32 | 244.4943 | 61 | 502.415 | 97 | EP300 | LoVo | Colon | GSM12086 | 1711 | 38762 |
| 481.6624 | 113 | 812.0324 | 166 | 444.2562 | 153 | 781.4513 | 218 | CEBPB | LoVo | Colon | GSM12085 | 6104 | 38763 |
| 1536.855 | 142 | 1571.414 | 188 | 1039.704 | 119 | 1170.444 | 134 | SP1 | LoVo | Colon | GSM12086 | 1031 | 38771 |
| 1087.478 | 304 | 1074.31 | 362 | 883.8578 | 348 | 903.1898 | 417 | CLOCK | LoVo | Colon | GSM12087 | 10609 | 38772 |
| 1271.738 | 301 | 1268.965 | 362 | 1002.591 | 288 | 998.505 | 318 | NFAT5 | LoVo | Colon | GSM12086 | 5861 | 38773 |
| 877.0504 | 422 | 902.4841 | 541 | 651.9855 | 465 | 638.0514 | 509 | CTCF | LoVo | Colon | GSM12086 | 32407 | 38774 |
| 953.0494 | 647 | 963.9335 | 792 | 738.0601 | 787 | 731.9433 | 889 | SMC1A | LoVo | Colon | GSM12086 | 51272 | 38778 |
| 1225.085 | 394 | 1247.85 | 507 | 933.8336 | 355 | 912.2595 | 381 | RFX1 | LoVo | Colon | GSM12086 | 8920 | 38784 |
| 1001.762 | 516 | 1029.424 | 659 | 822.3198 | 647 | 838.0671 | 771 | NIPBL | LoVo | Colon | GSM12086 | 28628 | 38785 |
| 727.2252 | 103 | 931.4955 | 130 | 570.5582 | 121 | 687.3743 | 142 | KLF3 | LoVo | Colon | GSM12086 | 2728 | 38790 |
| 837.2266 | 80 | 1240.663 | 111 | 460.0107 | 72 | 765.7688 | 103 | SMAD2 | LoVo | Colon | GSM12086 | 1048 | 38793 |
| 447.7674 | 79 | 375.8179 | 78 | 335.0666 | 93 | 659.7817 | 143 | ZFAT | LoVo | Colon | GSM12086 | 2945 | 38796 |
| 1397.892 | 183 | 1418.166 | 234 | 1129.003 | 169 | 1099.388 | 173 | GMEB1 | LoVo | Colon | GSM12086 | 2138 | 38799 |
| 1036.667 | 302 | 1062.877 | 392 | 820.0024 | 332 | 822.4699 | 381 | E2F8 | LoVo | Colon | GSM12087 | 12360 | 38804 |
| 885.5302 | 602 | 933.7107 | 788 | 691.6765 | 754 | 689.3038 | 859 | SMC3 | LoVo | Colon | GSM12086 | 55576 | 38811 |
| 850.9687 | 114 | 1153.897 | 149 | 596.4892 | 125 | 901.6182 | 167 | ASCL2 | LoVo | Colon | GSM12085 | 2800 | 38812 |
| 350.794 | 79 | 345.234 | 86 | 116.3816 | 64 | 254.3236 | 100 | REST | LoVo | Colon | GSM12087 | 3924 | 38814 |
| 1209.874 | 172 | 1234.075 | 223 | 954.5715 | 183 | 1015.909 | 219 | HOXA1 | LoVo | Colon | GSM12086 | 3586 | 38817 |
| 1349.661 | 170 | 1402.611 | 238 | 1005.068 | 156 | 1096.429 | 181 | BARX2 | LoVo | Colon | GSM12085 | 2153 | 38820 |
| 1228.677 | 406 | 1228.884 | 490 | 961.0689 | 393 | 944.8932 | 429 | MNT | LoVo | Colon | GSM12087 | 9542 | 38824 |
| 326.9802 | 49 | 237.6706 | 44 | 305.6527 | 64 | 550.1562 | 95 | ZIC2 | LoVo | Colon | GSM12088 | 1417 | 38826 |
| 480.3491 | 70 | 633.0964 | 90 | 233.2059 | 63 | 416.0057 | 92 | FOXG1 | LoVo | Colon | GSM12087 | 1913 | 38827 |
| 1462.566 | 151 | 1431.078 | 167 | 1096.643 | 138 | 1126.297 | 147 | LYL1 | LoVo | Colon | GSM12087 | 1433 | 38829 |
| 527.2078 | 71 | 778.0921 | 98 | 300.0641 | 69 | 495.5496 | 97 | FEV | LoVo | Colon | GSM12087 | 1733 | 38830 |
| 1091.057 | 375 | 1149.568 | 521 | 886.6987 | 439 | 885.3447 | 498 | KLF5 | LoVo | Colon | GSM12086 | 14349 | 38831 |
| 344.0195 | 63 | 531.8127 | 89 | 426.2337 | 98 | 726.1464 | 140 | TP73 | LoVo | Colon | GSM12088 | 2427 | 38832 |
| 232.574 | 37 | 421.2552 | 58 | 175.7786 | 43 | 508.6129 | 84 | CREB3 | LoVo | Colon | GSM12087 | 1154 | 38841 |
| 1106.372 | 122 | 1288.375 | 160 | 581.8851 | 110 | 756.4877 | 135 | NFE2L2 | LoVo | Colon | GSM12086 | 2113 | 38850 |
| 1347.647 | 117 | 1373.475 | 127 | 790.1611 | 108 | 1091.776 | 138 | ATF1 | LoVo | Colon | GSM12087 | 1254 | 38852 |
| 216.6055 | 61 | 297.4361 | 80 | 386.1175 | 118 | 837.7444 | 190 | DLX2 | LoVo | Colon | GSM12087 | 4119 | 38854 |
| 817.0655 | 100 | 1124.35 | 132 | 518.464 | 102 | 488.1235 | 104 | ATOH1 | LoVo | Colon | GSM12085 | 2072 | 38856 |
| 677.0119 | 147 | 891.7231 | 188 | 735.2764 | 215 | 858.5843 | 263 | MECOM | LoVo | Colon | GSM12087 | 7274 | 38858 |
| 1161.046 | 118 | 1336.254 | 155 | 841.0616 | 127 | 1142.683 | 162 | CASP8AP2 | LoVo | Colon | GSM12085 | 1768 | 38860 |
| 1038.688 | 222 | 1058.085 | 287 | 891.7332 | 290 | 889.159 | 329 | RXRA | LoVo | Colon | GSM12086 | 8585 | 38864 |
| 1005.069 | 201 | 1035.798 | 268 | 839.6692 | 248 | 849.8721 | 292 | E2F2 | LoVo | Colon | GSM12086 | 8344 | 38865 |
| 1076.214 | 316 | 1078.678 | 387 | 808.2263 | 298 | 772.1481 | 303 | BRD4 | MM1.S | Blood | GSM10382 | 11151 | 39001 |
| 1088.347 | 667 | 1117.212 | 838 | 781.2167 | 657 | 741.3756 | 680 | BRD4 | MM1.S | Blood | GSM10382 | 31502 | 39002 |
| 1041.431 | 420 | 1052.988 | 520 | 772.5175 | 408 | 725.8104 | 409 | BRD4 | MM1.S | Blood | GSM10382 | 18625 | 39014 |
| 1068.508 | 688 | 1061.702 | 822 | 738.8083 | 663 | 695.3475 | 683 | MED1 | MM1.S | Blood | GSM10382 | 38422 | 39017 |
| 900.438 | 566 | 909.462 | 694 | 604.5483 | 548 | 584.7397 | 594 | BRD4 | U87 | Brain | GSM10382 | 44758 | 39025 |
| 931.3196 | 653 | 948.4293 | 811 | 660.9771 | 704 | 635.6474 | 761 | BRD4 | H2171 | Lung | GSM10382 | 54301 | 39035 |
| 986.7602 | 554 | 1003.429 | 690 | 706.8809 | 547 | 663.0423 | 560 | BRD4 | MM1.S | Blood | GSM10382 | 32069 | 39047 |
| 1006.953 | 508 | 1025.29 | 639 | 727.0047 | 498 | 680.8948 | 504 | BRD4 | MM1.S | Blood | GSM10382 | 26859 | 39058 |
| 1164.322 | 551 | 1177.278 | 681 | 871.7364 | 541 | 817.0853 | 532 | NOTCH1 | CUTLL1 | Blood | GSM73290 | 19517 | 392 |
| 1080.801 | 254 | 1099.363 | 321 | 839.5534 | 254 | 770.7751 | 254 | NOTCH1 | CUTLL1 | Blood | GSM73290 | 8879 | 393 |
| 1008.911 | 874 | 1066.24 | 1109 | 728.2815 | 1033 | 702.6329 | 1134 | KLF1 | None | None | GSM10672 | 84894 | 39350 |
| 996.3681 | 774 | 1033.685 | 977 | 714.0333 | 872 | 677.6227 | 929 | GATA1 | None | None | GSM10672 | 66409 | 39352 |
| 1203.351 | 655 | 1237.822 | 835 | 890.2553 | 649 | 842.7464 | 658 | EP300 | None | None | GSM10672 | 24147 | 39356 |
| 996.1056 | 835 | 1031.418 | 1041 | 715.2021 | 969 | 671.0536 | 1021 | NFE2 | None | None | GSM10672 | 79109 | 39357 |
| 711.6035 | 260 | 790.7263 | 337 | 388.104 | 297 | 608.872 | 386 | SMAD3 | LX2 | None | GSM93461 | 25360 | 39359 |
| 48.97916 | 18 | 74.00624 | 25 | 17.93723 | 17 | 100.1123 | 41 | VDR | LX2 | None | GSM93461 | 1771 | 39360 |
| 506.1307 | 222 | 774.9264 | 323 | 387.3287 | 298 | 555.7458 | 373 | VDR | LX2 | None | GSM93461 | 25700 | 39364 |
| 85.76368 | 106 | 98.89081 | 128 | 102.7816 | 184 | 186.2529 | 246 | SMAD3 | LX2 | None | GSM93461 | 26098 | 39365 |
| 188.4924 | 52 | 284.534 | 72 | 99.40351 | 55 | 253.5886 | 93 | VDR | LX2 | None | GSM93461 | 3564 | 39366 |
| 113.7423 | 37 | 140.7761 | 46 | 59.30225 | 40 | 152.3298 | 67 | VDR | LX2 | None | GSM93461 | 3072 | 39367 |
| 1163.367 | 577 | 1182.664 | 722 | 952.5503 | 698 | 951.063 | 796 | ETV1 | LNCaP | Prostate | GSM11453 | 21845 | 39372 |
| 337.3416 | 53 | 389.7221 | 63 | 166.9787 | 49 | 732.9067 | 120 | AR | LNCaP | Prostate | GSM11453 | 1685 | 39378 |
| 1180.359 | 463 | 1219.457 | 607 | 903.6702 | 459 | 828.2905 | 427 | RBPJ | CUTLL1 | Blood | GSM73290 | 14273 | 394 |
| 947.6517 | 457 | 922.5089 | 526 | 795.9652 | 613 | 812.0807 | 731 | FOXM1 | MCF-7 | Breast | GSM10009 | 30027 | 39443 |
| 1136.987 | 349 | 1133.314 | 418 | 842.1545 | 308 | 841.3326 | 349 | FOXM1 | MDA-MB-2 | Breast | GSM10009 | 10878 | 39460 |
| 414.864 | 63 | 322.6723 | 59 | 344.1462 | 77 | 503.6039 | 101 | FOXM1 | MCF-7 | Breast | GSM10009 | 1863 | 39484 |
| 925.4787 | 387 | 907.2105 | 451 | 783.9477 | 530 | 818.8578 | 655 | FOXM1 | MCF-7 | Breast | GSM10009 | 25434 | 39495 |

| | | | | | | | | | | | | | |
|---|---|---|---|---|---|---|---|---|---|---|---|---|---|
| 1170.759 | 190 | 1163.74 | 227 | 787.6174 | 183 | 590.3492 | 166 | RBPJ | CUTLL1 | Blood | GSM73290 | 4760 | 395 |
| 234.1133 | 39 | 247.4596 | 44 | 52.21288 | 24 | 200.2067 | 52 | POU5F1 | None | None | GSM10005 | 1358 | 39513 |
| 984.2179 | 197 | 985.0846 | 245 | 547.8161 | 205 | 628.2438 | 234 | MYC | BJ | Skin | GSM10005 | 9326 | 39560 |
| 1080.609 | 295 | 1107.33 | 382 | 829.6674 | 293 | 805.3829 | 309 | BCL6 | OCI-Ly7 | Blood | GSM10848 | 10743 | 39572 |
| 817.9769 | 206 | 863.6469 | 234 | 357.8447 | 199 | 454.7413 | 238 | BACH2 | OCI-Ly7 | Blood | GSM10848 | 12862 | 39577 |
| 995.5882 | 208 | 1040.763 | 286 | 509.4849 | 198 | 522.1538 | 214 | ZNF143 | CUTLL1 | Blood | GSM73290 | 9154 | 396 |
| 83.2928 | 24 | 150.7931 | 37 | 92.59888 | 37 | 187.6681 | 57 | SRC | LS180 | Colon | GSM95962 | 1804 | 39652 |
| 480.2685 | 90 | 554.0566 | 107 | 390.6262 | 112 | 558.8045 | 144 | CREBBP | LS180 | Colon | GSM95962 | 3744 | 39654 |
| 441.5365 | 96 | 483.3076 | 111 | 343.3629 | 119 | 552.7191 | 162 | NCOR1 | LS180 | Colon | GSM95963 | 4940 | 39655 |
| 469.29 | 106 | 578.5914 | 131 | 492.1472 | 154 | 756.4396 | 205 | NCOR1 | LS180 | Colon | GSM95963 | 5764 | 39656 |
| 118.2444 | 34 | 144.4538 | 42 | 65.00742 | 37 | 127.9851 | 55 | MED1 | LS180 | Colon | GSM95963 | 2516 | 39658 |
| 363.4787 | 75 | 639.6268 | 114 | 322.8915 | 99 | 414.6361 | 120 | CREBBP | LS180 | Colon | GSM95962 | 3587 | 39659 |
| 49.41325 | 35 | 72.08275 | 47 | 62.12965 | 61 | 136.8442 | 92 | SRC | LS180 | Colon | GSM95962 | 6243 | 39660 |
| 175.5262 | 53 | 133.4613 | 51 | 56.78943 | 46 | 130.9767 | 72 | MED1 | LS180 | Colon | GSM95963 | 4131 | 39661 |
| 61.38931 | 22 | 63.97687 | 25 | 56.45705 | 31 | 59.13653 | 34 | RUNX1 | Jurkat | Blood | GSM10453 | 1895 | 39674 |
| 1113.899 | 281 | 1128.204 | 360 | 845.6946 | 270 | 827.0514 | 289 | TET3 | HEK293T | Embryonic | GSM10189 | 9323 | 39719 |
| 432.4883 | 253 | 656.025 | 341 | 296.3895 | 340 | 347.2474 | 391 | CTCF | PrEC | Prostate | GSM94752 | 39851 | 39736 |
| 655.3718 | 302 | 752.1975 | 419 | 470.2975 | 401 | 500.8099 | 445 | CTCF | LNCaP | Prostate | GSM94752 | 39077 | 39737 |
| 112.3309 | 89 | 218.2635 | 134 | 98.03212 | 133 | 125.2185 | 158 | RUNX1 | Kasumi-1 | Blood | GSM11134 | 15747 | 39775 |
| 372.2818 | 106 | 485.1428 | 135 | 175.6352 | 108 | 278.3756 | 143 | REPIN1 | Kasumi-1 | Blood | GSM11134 | 7716 | 39776 |
| 710.9317 | 409 | 728.1263 | 516 | 494.2485 | 501 | 509.681 | 554 | REPIN1 | Kasumi-1 | Blood | GSM11134 | 55869 | 39777 |
| 311.0614 | 326 | 352.7493 | 390 | 157.5918 | 433 | 123.8819 | 447 | REPIN1 | Kasumi-1 | Blood | GSM11134 | 80678 | 39779 |
| 197.3332 | 123 | 324.7768 | 173 | 106.761 | 151 | 127.755 | 175 | RUNX1 | Kasumi-1 | Blood | GSM11134 | 18781 | 39780 |
| 208.7761 | 56 | 333.0815 | 80 | 105.9417 | 58 | 213.0548 | 87 | REPIN1 | Kasumi-1 | Blood | GSM11134 | 3750 | 39781 |
| 764.373 | 341 | 810.5846 | 468 | 561.5134 | 419 | 531.9005 | 444 | RUNX1 | Kasumi-1 | Blood | GSM11134 | 36589 | 39782 |
| 51.48118 | 66 | 145.9555 | 114 | 68.70189 | 119 | 85.90953 | 140 | RUNX1 | Kasumi-1 | Blood | GSM11134 | 16137 | 39783 |
| 253.8962 | 185 | 415.3015 | 257 | 142.1008 | 236 | 220.9221 | 299 | RUNX1T1 | Kasumi-1 | Blood | GSM11134 | 32379 | 39784 |
| 57.48452 | 16 | 47.65414 | 16 | 14.64673 | 12 | 98.43397 | 33 | RUNX1 | Kasumi-1 | Blood | GSM11134 | 1180 | 39785 |
| 63.95044 | 69 | 130.6504 | 105 | 51.61379 | 104 | 99.26409 | 142 | RUNX1T1 | Kasumi-1 | Blood | GSM11134 | 15456 | 39786 |
| 178.9459 | 49 | 177.7641 | 54 | 111.2479 | 56 | 206.0255 | 81 | ESR1 | DL23 | Colon | GSM10375 | 3343 | 39873 |
| 807.7926 | 250 | 823.3937 | 306 | 326.7913 | 247 | 362.4362 | 278 | HCFC1 | HEK293T | Embryonic | GSM10375 | 21437 | 39886 |
| 857.0309 | 131 | 1183.36 | 208 | 539.2842 | 138 | 688.4928 | 167 | OGT | HEK293T | Embryonic | GSM10375 | 3990 | 39911 |
| 1007.218 | 514 | 1009.952 | 626 | 736.2596 | 528 | 721.1194 | 577 | ICE1 | HCT-116 | Colon | GSM11627 | 29511 | 39914 |
| 876.9517 | 144 | 762.0716 | 145 | 424.1458 | 134 | 387.1899 | 136 | ZC3H8 | HCT-116 | Colon | GSM11627 | 5016 | 39918 |
| 1013.68 | 558 | 1021.865 | 691 | 747.5384 | 590 | 727.3425 | 643 | ICE1 | HCT-116 | Colon | GSM11627 | 33354 | 39920 |
| 294.0278 | 55 | 227.9737 | 52 | 107.6849 | 45 | 167.0033 | 60 | ICE2 | HCT-116 | Colon | GSM11627 | 2253 | 39924 |
| 1035.313 | 295 | 1061.603 | 379 | 788.6082 | 290 | 602.7922 | 262 | EP300 | None | None | GSM10527 | 11332 | 39938 |
| 718.7104 | 350 | 775.4854 | 494 | 544.0025 | 459 | 529.9022 | 493 | CTCF | GM12868 | Blood | GSM84930 | 44883 | 39970 |
| 729.2303 | 340 | 788.4858 | 482 | 545.9778 | 441 | 550.1303 | 480 | CTCF | GM12867 | Blood | GSM84930 | 41612 | 39971 |
| 754.8516 | 426 | 812.2789 | 592 | 567.8607 | 529 | 555.3029 | 584 | CTCF | GM12871 | Blood | GSM84930 | 51832 | 39972 |
| 712.1851 | 328 | 785.5549 | 478 | 502.3602 | 428 | 482.2573 | 457 | CTCF | GM12870 | Blood | GSM84930 | 41714 | 39973 |
| 728.5066 | 370 | 794.3706 | 525 | 572.2809 | 492 | 544.6523 | 520 | CTCF | GM12869 | Blood | GSM84930 | 46651 | 39974 |
| 732.6427 | 367 | 794.2106 | 518 | 565.3341 | 472 | 552.7318 | 520 | CTCF | GM12866 | Blood | GSM84930 | 45290 | 39975 |
| 1008.109 | 782 | 1035.014 | 969 | 709.9596 | 852 | 707.6466 | 974 | TFAP2C | A375 | Skin | GSM10115 | 63934 | 39978 |
| 50.31343 | 20 | 99.55292 | 32 | 17.82082 | 19 | 138.413 | 53 | EGR1 | K562 | Bone Marr | GSM54070 | 2136 | 4 |
| 1050.413 | 542 | 1067.352 | 677 | 755.7639 | 526 | 720.0049 | 545 | MED1 | MM1.S | Blood | GSM11210 | 27278 | 40016 |
| 1080.095 | 590 | 1101.51 | 743 | 770.545 | 562 | 727.1036 | 577 | BRD4 | None | Blood | GSM11210 | 27512 | 40022 |
| 549.2945 | 119 | 772.5999 | 158 | 274.0785 | 117 | 273.924 | 125 | BRD4 | SKMEL5 | None | GSM11210 | 5699 | 40032 |
| 1066.966 | 520 | 1086.005 | 650 | 775.4685 | 506 | 736.5318 | 521 | CDK7 | None | Blood | GSM11210 | 24537 | 40053 |
| 1033.277 | 454 | 1052.113 | 574 | 743.8895 | 425 | 733.1648 | 469 | CDK7 | None | Blood | GSM11210 | 21376 | 40055 |
| 481.0825 | 60 | 579.3992 | 73 | 228.7178 | 52 | 358.0731 | 71 | BRD4 | None | Blood | GSM11210 | 1307 | 40056 |
| 24.45077 | 29 | 43.62432 | 42 | 32.32259 | 53 | 41.32529 | 63 | LMO2 | Kasumi-1 | Blood | GSM10823 | 7292 | 40156 |
| 863.9803 | 262 | 884.189 | 336 | 597.982 | 304 | 608.1258 | 329 | TCF12 | Kasumi-1 | Blood | GSM10823 | 18637 | 40162 |
| 1075.353 | 541 | 1105.87 | 692 | 787.8261 | 529 | 757.1375 | 560 | RUNX1T1 | Kasumi-1 | Blood | GSM10823 | 24911 | 40164 |
| 808.7767 | 571 | 854.4009 | 746 | 599.7216 | 687 | 592.0138 | 769 | TCF7L1 | Kasumi-1 | Blood | GSM10823 | 64087 | 40166 |
| 886.0474 | 601 | 909.5863 | 753 | 625.6689 | 651 | 581.3454 | 666 | IRF4 | None | Blood | GSM11955 | 54689 | 40215 |
| 1163.736 | 639 | 1167.161 | 773 | 830.5588 | 595 | 781.9353 | 600 | MED1 | None | Blood | GSM11955 | 24948 | 40216 |
| 879.4894 | 716 | 889.9732 | 871 | 601.4972 | 784 | 578.7635 | 855 | BRD4 | U87 | Brain | GSM11955 | 78681 | 40217 |
| 984.7237 | 397 | 980.1256 | 477 | 731.0106 | 403 | 684.9356 | 406 | BRD4 | H2171 | Lung | GSM11955 | 21052 | 40220 |
| 902.8675 | 729 | 889.2347 | 857 | 612.522 | 780 | 583.7808 | 839 | BRD4 | U87 | Brain | GSM11955 | 72954 | 40221 |
| 705.8742 | 137 | 764.1498 | 157 | 405.9751 | 143 | 372.6654 | 146 | BRD4 | H2171 | Lung | GSM11955 | 5943 | 40225 |
| 1113.466 | 533 | 1141.119 | 679 | 819.353 | 509 | 791.0674 | 542 | CDK9 | None | Blood | GSM11955 | 21232 | 40226 |
| 1175.942 | 629 | 1200.259 | 788 | 855.3449 | 596 | 817.5748 | 617 | CDK9 | None | Blood | GSM11955 | 22948 | 40227 |
| 985.4795 | 372 | 1001.061 | 467 | 738.5402 | 376 | 711.5653 | 398 | BRD4 | None | None | GSM11955 | 18623 | 40230 |
| 1057.316 | 679 | 1096.571 | 866 | 751.869 | 683 | 695.4625 | 684 | MED1 | None | Blood | GSM11955 | 39064 | 40233 |
| 1035.829 | 612 | 1052.87 | 757 | 737.2126 | 602 | 684.7237 | 607 | MED1 | None | Blood | GSM11955 | 34181 | 40234 |
| 4.802171 | 6 | 0.348703 | 3 | 0.122638 | 4 | 33.48498 | 25 | TP53 | IMR90 | Lung | GSM10488 | 1887 | 40287 |
| 2.011699 | 3 | 1.352416 | 3 | 0 | 2 | 41.77038 | 22 | TP53 | IMR90 | Lung | GSM10488 | 1254 | 40289 |
| 259.0511 | 108 | 336.7871 | 137 | 180.6589 | 138 | 158.6677 | 141 | MBD2 | HeLa | Cervix | GSM10067 | 11438 | 40334 |
| 361.4931 | 53 | 496.4076 | 70 | 263.4747 | 60 | 267.7169 | 64 | MBD3 | HeLa | Cervix | GSM10067 | 1499 | 40335 |
| 396.1363 | 55 | 547.9748 | 73 | 265.712 | 59 | 770.6474 | 116 | AR | LNCaP | Prostate | GSM11744 | 1434 | 40346 |
| 15.52565 | 9 | 47.47771 | 18 | 17.70194 | 15 | 49.13212 | 26 | CHD5 | SH-SY5Y | Bone Marr | GSM11747 | 1415 | 40348 |

| | | | | | | | | | | | | | |
|---|---|---|---|---|---|---|---|---|---|---|---|---|---|
| 244.2421 | 73 | 299.0393 | 90 | 436.4039 | 142 | 684.9836 | 191 | ESR1 | ZR-75-1 | Mammary | GSM63148 | 5468 | 40371 |
| 263.8298 | 87 | 281.927 | 100 | 383.9518 | 153 | 489.2897 | 184 | ESR1 | ZR-75-1 | Mammary | GSM63148 | 7331 | 40372 |
| 251.823 | 89 | 309.7899 | 110 | 332.9857 | 150 | 443.2964 | 184 | ESR1 | ZR-75-1 | Mammary | GSM63148 | 8111 | 40373 |
| 295.464 | 95 | 320.6234 | 110 | 468.6919 | 174 | 762.3264 | 237 | ESR1 | ZR-75-1 | Mammary | GSM63148 | 7791 | 40375 |
| 811.0458 | 546 | 782.8428 | 626 | 655.8259 | 749 | 697.0334 | 932 | FOXA1 | ZR-75-1 | Mammary | GSM63147 | 63661 | 40376 |
| 8.028269 | 50 | 10.36846 | 63 | 13.67268 | 104 | 40.31614 | 150 | CTCF | BCBL1 | None | GSM97082 | 26634 | 40379 |
| 241.5788 | 49 | 267.7878 | 57 | 299.3907 | 77 | 440.7494 | 101 | ESR1 | None | Mammary | GSM10037 | 2209 | 41121 |
| 966.665 | 316 | 942.5967 | 359 | 833.4872 | 436 | 872.8753 | 548 | ESR1 | None | Mammary | GSM10037 | 16833 | 41124 |
| 290.6309 | 65 | 428.8321 | 89 | 369.8688 | 104 | 473.5169 | 126 | ESR1 | None | Mammary | GSM10037 | 3299 | 41127 |
| 982.1666 | 226 | 966.366 | 268 | 863.4898 | 325 | 870.4849 | 376 | ESR1 | None | Mammary | GSM10037 | 10813 | 41130 |
| 433.1595 | 105 | 421.3178 | 114 | 538.7586 | 167 | 750.3051 | 211 | ESR1 | None | Mammary | GSM10037 | 6093 | 41133 |
| 118.4017 | 29 | 168.2349 | 39 | 85.6542 | 35 | 287.7962 | 71 | ESR1 | None | Mammary | GSM10037 | 1704 | 41136 |
| 572.9237 | 114 | 691.334 | 139 | 751.764 | 184 | 935.9131 | 235 | ESR1 | None | Mammary | GSM10037 | 5086 | 41141 |
| 993.4141 | 270 | 1011.892 | 346 | 869.0597 | 383 | 893.0797 | 466 | ESR1 | None | Mammary | GSM10037 | 12710 | 41144 |
| 38.77588 | 13 | 103.3905 | 25 | 74.57004 | 27 | 132.1384 | 39 | ESR1 | None | Mammary | GSM10037 | 1175 | 41147 |
| 21.37645 | 32 | 28.35024 | 41 | 57.17586 | 76 | 83.10547 | 95 | PPARG | SGBS | Adipose | GSM10203 | 9231 | 41166 |
| 6.09384 | 28 | 9.399275 | 37 | 9.047126 | 56 | 23.85502 | 80 | CEBPA | SGBS | Adipose | GSM10203 | 13560 | 41167 |
| 1205.169 | 276 | 1271.4 | 393 | 940.8564 | 265 | 932.3366 | 292 | HDAC1 | None | Blood | GSM39395 | 6649 | 4118 |
| 153.5456 | 212 | 199.4597 | 266 | 87.84982 | 295 | 166.4937 | 388 | EED | DU145 | Prostate | GSM10454 | 54875 | 41187 |
| 83.93009 | 107 | 82.64477 | 122 | 39.20678 | 139 | 59.58347 | 172 | EED | DU145 | Prostate | GSM10454 | 26254 | 41188 |
| 294.6068 | 42 | 409.8872 | 56 | 243.9267 | 51 | 411.892 | 73 | HDAC3 | None | Blood | GSM39395 | 1146 | 4125 |
| 1058.626 | 222 | 1086.558 | 292 | 649.536 | 211 | 651.0746 | 225 | HDAC6 | None | Blood | GSM39395 | 8251 | 4126 |
| 506.1001 | 93 | 662.595 | 119 | 261.4154 | 91 | 407.8297 | 122 | RELA | IMR90 | Lung | GSM10558 | 3781 | 41268 |
| 916.7302 | 653 | 909.894 | 778 | 656.1935 | 728 | 638.8441 | 799 | EP300 | IMR90 | Lung | GSM10558 | 60044 | 41270 |
| 834.3676 | 391 | 848.6029 | 490 | 629.8479 | 456 | 613.8962 | 495 | EP300 | IMR90 | Lung | GSM10558 | 34864 | 41271 |
| 753.53 | 371 | 793.0787 | 498 | 570.8435 | 461 | 544.8321 | 490 | CTCF | IMR90 | Lung | GSM10558 | 43300 | 41283 |
| 14.55331 | 25 | 36.63603 | 41 | 11.68794 | 40 | 20.27781 | 52 | TP63 | EP156T | Prostate | GSM10568 | 7876 | 41284 |
| 1101.041 | 201 | 1071.592 | 226 | 669.8491 | 189 | 518.1137 | 176 | STAT1 | None | Blood | GSM10570 | 5959 | 41286 |
| 980.5761 | 608 | 986.4602 | 744 | 715.507 | 651 | 655.9126 | 644 | STAT1 | None | Blood | GSM10570 | 40387 | 41287 |
| 949.1305 | 357 | 936.0414 | 423 | 720.1191 | 382 | 641.7239 | 355 | STAT1 | None | Blood | GSM10570 | 19893 | 41288 |
| 947.0197 | 639 | 966.715 | 799 | 694.1552 | 714 | 632.2844 | 709 | STAT1 | None | Blood | GSM10570 | 49857 | 41289 |
| 1013.256 | 388 | 1005.886 | 464 | 761.1926 | 390 | 710.4005 | 385 | STAT1 | None | Blood | GSM10570 | 17577 | 41290 |
| 1008.123 | 276 | 992.7354 | 318 | 772.6905 | 276 | 593.8657 | 260 | STAT1 | None | Blood | GSM10570 | 11186 | 41291 |
| 232.1944 | 68 | 371.9421 | 97 | 108.4898 | 69 | 166.0864 | 90 | IRF1 | None | Blood | GSM10570 | 5049 | 41300 |
| 832.657 | 391 | 869.2843 | 517 | 613.6515 | 439 | 593.7285 | 475 | IRF1 | None | Blood | GSM10570 | 35327 | 41301 |
| 495.7225 | 177 | 720.7619 | 237 | 326.2212 | 217 | 294.9476 | 224 | IRF1 | None | Blood | GSM10570 | 16507 | 41302 |
| 681.457 | 280 | 736.8332 | 342 | 351.8788 | 322 | 434.0463 | 378 | IRF1 | None | Blood | GSM10570 | 32362 | 41303 |
| 1155.509 | 163 | 1148.04 | 191 | 490.7779 | 133 | 550.2623 | 150 | ZFX | NOMO-1 | None | GSM10573 | 4082 | 41307 |
| 532.3868 | 249 | 713.4955 | 322 | 302.3996 | 303 | 321.3998 | 336 | SMC3 | HEK293T | Embryonic | GSM10815 | 32497 | 41382 |
| 85.11797 | 45 | 90.14026 | 52 | 40.21943 | 51 | 131.885 | 90 | TEAD4 | MKN28 | None | GSM10847 | 6057 | 41389 |
| 134.4235 | 132 | 128.1099 | 148 | 159.743 | 226 | 258.5904 | 292 | TEAD4 | SNU216 | None | GSM10847 | 28988 | 41391 |
| 80.79422 | 28 | 56.7303 | 26 | 43.87128 | 31 | 155.7048 | 61 | MBD3 | MDA-231 | Mammary | GSM10898 | 2443 | 41402 |
| 432.6407 | 182 | 481.9115 | 214 | 138.2813 | 172 | 305.2455 | 253 | ERCC3 | HT1080 | Connective | GSM10925 | 19749 | 41411 |
| 301.9268 | 129 | 385.6611 | 162 | 123.4573 | 133 | 234.5227 | 186 | ERCC2 | HT1080 | Connective | GSM10925 | 13768 | 41412 |
| 275.3432 | 41 | 398.6771 | 56 | 182.5641 | 44 | 293.9361 | 61 | SFPQ | LNCaP | Prostate | GSM10974 | 1179 | 41423 |
| 1050.77 | 473 | 1069.771 | 600 | 763.1734 | 455 | 742.2943 | 490 | BRD4 | OCI-Ly1 | Bone Marr | GSM11336 | 22157 | 41475 |
| 1061.378 | 434 | 1066.578 | 539 | 779.2166 | 421 | 759.9463 | 455 | BRD4 | OCI-Ly1 | Bone Marr | GSM11336 | 19342 | 41476 |
| 1259.855 | 414 | 1292.481 | 545 | 981.3183 | 403 | 942.0457 | 411 | E2F1 | OCI-Ly1 | Bone Marr | GSM11336 | 9300 | 41481 |
| 536.4942 | 146 | 611.9162 | 173 | 314.4039 | 163 | 377.244 | 190 | BCL6 | OCI-Ly1 | Bone Marr | GSM11336 | 10003 | 41482 |
| 42.52552 | 107 | 24.81883 | 107 | 18.93127 | 151 | 19.90685 | 169 | RING1 | DU145 | Prostate | GSM11385 | 37200 | 41489 |
| 19.91626 | 119 | 24.98295 | 148 | 13.0986 | 199 | 37.63604 | 272 | RNF2 | DU145 | Prostate | GSM11385 | 58391 | 41490 |
| 12.9005 | 16 | 15.81169 | 20 | 6.121022 | 21 | 32.38981 | 41 | RNF2 | DU145 | Prostate | GSM11385 | 4288 | 41491 |
| 82.57791 | 236 | 106.7999 | 295 | 33.26027 | 335 | 67.99747 | 430 | BMI1 | DU145 | Prostate | GSM11385 | 90248 | 41492 |
| 17.0506 | 84 | 6.430131 | 80 | 14.35357 | 145 | 20.93485 | 174 | BMI1 | DU145 | Prostate | GSM11385 | 38211 | 41493 |
| 979.8374 | 356 | 1005.601 | 468 | 754.0299 | 393 | 694.623 | 379 | TP53 | GM06993, | Blood | GSM11426 | 19737 | 41496 |
| 17.55805 | 10 | 4.362943 | 6 | 0.66714 | 5 | 40.8081 | 25 | TP53 | GM12878 | Blood | GSM11427 | 1604 | 41500 |
| 297.8097 | 59 | 358.9523 | 72 | 527.8998 | 113 | 882.6966 | 160 | NR5A2 | MCF-7 | Breast | GSM11431 | 2582 | 41515 |
| 571.6373 | 116 | 902.4118 | 165 | 343.9476 | 125 | 337.7742 | 132 | SPI1 | None | Blood | GSM11464 | 5444 | 41568 |
| 693.1343 | 113 | 944.5275 | 148 | 468.0407 | 125 | 539.0334 | 143 | SPI1 | None | Blood | GSM11464 | 3844 | 41569 |
| 1161.638 | 410 | 1187.902 | 526 | 871.332 | 369 | 825.4019 | 370 | ING2 | RKO | Colon | GSM11464 | 10632 | 41572 |
| 1115.955 | 204 | 1109.151 | 239 | 648.0991 | 179 | 515.0926 | 169 | ING5 | RKO | Colon | GSM11464 | 4853 | 41573 |
| 1215.245 | 477 | 1259.554 | 636 | 931.8422 | 452 | 878.8434 | 450 | BRD1 | RKO | Colon | GSM11464 | 11627 | 41574 |
| 652.9612 | 166 | 911.614 | 222 | 531.3317 | 214 | 403.8845 | 201 | RELA | None | Umbilical V | GSM11540 | 10328 | 41619 |
| 593.078 | 175 | 819.847 | 229 | 359.6189 | 201 | 323.3953 | 206 | MAFK | OCI-Ly7 | Blood | GSM11596 | 13146 | 41625 |
| 262.8585 | 77 | 278.9519 | 88 | 83.8849 | 66 | 143.7694 | 90 | POLR3D | IMR90 | Lung | GSM11606 | 5665 | 41628 |
| 148.0024 | 115 | 152.4916 | 132 | 183.8129 | 196 | 297.3722 | 256 | NR3C1 | HEK293 | Embryonic | GSM11766 | 20428 | 41735 |
| 144.5383 | 142 | 156.7562 | 167 | 132.9076 | 221 | 278.475 | 312 | NR3C1 | HEK293 | Embryonic | GSM11767 | 30133 | 41737 |
| 105.4692 | 33 | 142.602 | 43 | 208.7005 | 68 | 241.1801 | 78 | NR3C1 | HEK293 | Embryonic | GSM11767 | 2522 | 41738 |
| 346.8806 | 137 | 484.7607 | 180 | 151.9195 | 144 | 229.8536 | 184 | SUMO2 | HEK293 | Embryonic | GSM11767 | 14093 | 41740 |
| 645.8077 | 251 | 778.4001 | 330 | 291.5018 | 268 | 354.2788 | 313 | SUMO2 | HEK293 | Embryonic | GSM11767 | 25631 | 41741 |
| 647.301 | 219 | 769.2825 | 265 | 254.9574 | 215 | 327.3774 | 257 | SUMO2 | HEK293 | Embryonic | GSM11767 | 19244 | 41742 |
| 834.6958 | 287 | 827.6079 | 343 | 352.2743 | 270 | 439.0361 | 319 | SUMO2 | HEK293 | Embryonic | GSM11767 | 22601 | 41743 |

| | | | | | | | | | | | | | |
|---|---|---|---|---|---|---|---|---|---|---|---|---|---|
| 1106.585 | 578 | 1144.041 | 744 | 817.1951 | 570 | 758.5654 | 564 | EZH2 | None | None | GSM11801 | 24405 | 41788 |
| 558.8822 | 149 | 664.9063 | 180 | 297.3889 | 156 | 192.2883 | 138 | JARID2 | None | None | GSM11801 | 8921 | 41790 |
| 1009.817 | 428 | 1015.678 | 525 | 735.0092 | 412 | 695.8533 | 420 | JARID2 | None | None | GSM11801 | 20600 | 41793 |
| 927.2994 | 229 | 937.0584 | 286 | 496.132 | 229 | 343.8956 | 208 | EZH2 | None | None | GSM11801 | 12050 | 41794 |
| 690.9521 | 167 | 832.2951 | 203 | 351.6425 | 169 | 381.0977 | 188 | JARID2 | None | None | GSM11801 | 9042 | 41797 |
| 21.41 | 9 | 17.14104 | 9 | 19.33656 | 13 | 88.73739 | 30 | EZH2 | None | None | GSM11801 | 1025 | 41798 |
| 20.187 | 9 | 23.69801 | 11 | 24.09195 | 15 | 94.33109 | 32 | JARID2 | Edom-iPS-2 | None | GSM11801 | 1114 | 41807 |
| 23.00333 | 13 | 61.4599 | 24 | 43.86745 | 27 | 113.574 | 46 | JARID2 | None | None | GSM11801 | 1867 | 41808 |
| 555.2841 | 150 | 714.9269 | 189 | 276.7106 | 153 | 260.8674 | 160 | EZH2 | None | None | GSM11801 | 9229 | 41809 |
| 14.24563 | 8 | 48.28002 | 17 | 18.41448 | 14 | 119.0586 | 38 | JARID2 | None | None | GSM11801 | 1180 | 41812 |
| 32.77641 | 12 | 30.99983 | 13 | 29.74544 | 17 | 70.28127 | 28 | EZH2 | None | None | GSM11801 | 1133 | 41813 |
| 16.92307 | 9 | 56.36945 | 19 | 19.57572 | 15 | 95.71833 | 35 | JARID2 | None | None | GSM11801 | 1316 | 41816 |
| 171.4519 | 56 | 240.0474 | 74 | 126.4059 | 70 | 129.7298 | 76 | EZH2 | None | None | GSM11801 | 4201 | 41817 |
| 28.39215 | 14 | 70.30225 | 25 | 40.63728 | 25 | 47.80019 | 29 | JARID2 | None | None | GSM11801 | 1665 | 41820 |
| 34.05943 | 19 | 87.04941 | 34 | 36.68879 | 30 | 99.59355 | 51 | EZH2 | None | None | GSM11801 | 2509 | 41821 |
| 202.6366 | 65 | 376.0457 | 100 | 127.9555 | 76 | 158.6142 | 90 | JARID2 | None | None | GSM11801 | 5006 | 41824 |
| 127.1751 | 44 | 238.1013 | 68 | 87.9701 | 54 | 132.0917 | 70 | EZH2 | None | None | GSM11801 | 3610 | 41825 |
| 115.4678 | 37 | 219.8143 | 58 | 49.28466 | 36 | 101.6894 | 54 | JARID2 | None | None | GSM11801 | 2803 | 41828 |
| 1159.114 | 553 | 1192.703 | 715 | 856.6988 | 535 | 801.2724 | 533 | ZBTB17 | MDA-MD-2 | Mammary | GSM11819 | 20495 | 41831 |
| 172.8253 | 210 | 281.038 | 287 | 178.7857 | 349 | 208.237 | 400 | ASCL1 | None | None | GSM11872 | 55566 | 41850 |
| 49.78409 | 64 | 48.67389 | 73 | 37.27831 | 97 | 38.28531 | 107 | ASCL1 | NHEK | Skin | GSM11872 | 16898 | 41852 |
| 99.09212 | 38 | 88.03967 | 40 | 58.94824 | 45 | 60.46952 | 49 | EZH2 | HEK293T | Embryonic | GSM12151 | 3858 | 41974 |
| 461.8207 | 269 | 686.5371 | 372 | 306.7999 | 357 | 343.577 | 404 | CTCF | HCT-116 | Colon | GSM12246 | 42098 | 42148 |
| 543.1984 | 320 | 687.5534 | 438 | 365.1047 | 428 | 397.9199 | 480 | CTCF | HCT-116 | Colon | GSM12246 | 51489 | 42149 |
| 409.5747 | 265 | 675.2363 | 384 | 321.9912 | 378 | 335.2531 | 417 | CTCF | HCT-116 | Colon | GSM12246 | 45067 | 42150 |
| 446.1321 | 244 | 715.0454 | 353 | 340.803 | 338 | 372.7998 | 379 | CTCF | HCT-116 | Colon | GSM12246 | 35968 | 42151 |
| 518.2338 | 312 | 701.2648 | 450 | 351.473 | 420 | 369.0675 | 465 | CTCF | HCT-116 | Colon | GSM12246 | 51223 | 42152 |
| 482.0551 | 294 | 676.9415 | 406 | 332.7169 | 399 | 364.1121 | 448 | CTCF | HCT-116 | Colon | GSM12246 | 48771 | 42153 |
| 447.574 | 274 | 688.7874 | 396 | 329.0504 | 380 | 356.4243 | 425 | CTCF | HCT-116 | Colon | GSM12246 | 45164 | 42154 |
| 604.193 | 325 | 728.8128 | 467 | 434.3196 | 440 | 505.3233 | 505 | CTCF | K562 | Bone Marr | GSM12246 | 48358 | 42155 |
| 267.764 | 122 | 422.9594 | 170 | 187.0818 | 159 | 267.9634 | 199 | CTCF | K562 | Bone Marr | GSM12246 | 14692 | 42156 |
| 398.907 | 250 | 638.169 | 344 | 291.379 | 345 | 309.0071 | 383 | CTCF | K562 | Bone Marr | GSM12246 | 40514 | 42157 |
| 529.5341 | 308 | 693.8767 | 429 | 431.5257 | 441 | 477.5318 | 497 | CTCF | K562 | Bone Marr | GSM12246 | 49303 | 42158 |
| 346.1944 | 219 | 536.3938 | 298 | 267.9971 | 310 | 337.4187 | 366 | CTCF | K562 | Bone Marr | GSM12246 | 36247 | 42159 |
| 929.7652 | 464 | 949.2563 | 585 | 680.7055 | 493 | 665.4408 | 540 | MITF | None | None | GSM12262 | 33270 | 42176 |
| 539.641 | 169 | 770.5811 | 225 | 283.3492 | 185 | 313.9912 | 208 | MITF | None | None | GSM12262 | 14098 | 42177 |
| 79.14492 | 66 | 151.303 | 99 | 85.71864 | 109 | 112.7187 | 131 | MITF | COLO-829 | Skin | GSM12262 | 13080 | 42178 |
| 876.4765 | 465 | 909.2607 | 602 | 637.5408 | 505 | 634.7874 | 572 | MITF | COLO-829 | Skin | GSM12262 | 39324 | 42179 |
| 1118.308 | 444 | 1140.44 | 565 | 821.3717 | 410 | 783.5602 | 423 | STAT3 | OCI-Ly10 | Bone Marr | GSM12272 | 16063 | 42212 |
| 1136.02 | 369 | 1123.408 | 437 | 848.0916 | 336 | 793.9317 | 330 | STAT3 | OCI-Ly19 | Bone Marr | GSM12272 | 11677 | 42213 |
| 1011.727 | 280 | 1015.288 | 344 | 658.0375 | 261 | 566.087 | 260 | STAT3 | OCI-Ly3 | Bone Marr | GSM12272 | 12525 | 42214 |
| 1040.488 | 373 | 1053.577 | 469 | 765.4735 | 353 | 740.0966 | 376 | STAT3 | OCI-Ly7 | Blood | GSM12272 | 16287 | 42215 |
| 897.4406 | 133 | 798.646 | 135 | 452.2464 | 123 | 326.3712 | 110 | STAT3 | SU-DHL10 | None | GSM12272 | 3711 | 42216 |
| 1009.603 | 417 | 1039.969 | 543 | 743.5583 | 417 | 702.1209 | 425 | STAT3 | SU-DHL2 | None | GSM12272 | 21740 | 42217 |
| 1044.443 | 342 | 1052.145 | 425 | 753.3253 | 309 | 732.4003 | 332 | STAT3 | SU-DHL4 | Peritoneal | GSM12272 | 14564 | 42218 |
| 1059.356 | 415 | 1080.891 | 526 | 795.2217 | 410 | 755.3229 | 420 | STAT3 | SU-DHL6 | Lymph Noc | GSM12272 | 16847 | 42219 |
| 964.237 | 440 | 969.2778 | 539 | 682.1587 | 425 | 671.3209 | 471 | STAT3 | U-2932 | None | GSM12272 | 26910 | 42220 |
| 656.0217 | 324 | 756.0427 | 467 | 461.2887 | 424 | 399.4487 | 437 | CTCF | GM10847 | Blood | GSM12338 | 40560 | 42255 |
| 429.8856 | 269 | 676.8901 | 380 | 331.4647 | 373 | 280.5938 | 383 | CTCF | GM10847 | Blood | GSM12338 | 40046 | 42256 |
| 312.4974 | 88 | 292.8892 | 94 | 176.7364 | 96 | 200.0918 | 109 | CTCF | GM10847 | Blood | GSM12338 | 5872 | 42257 |
| 628.9675 | 273 | 703.4929 | 321 | 397.842 | 336 | 353.7135 | 348 | STAG1 | GM10847 | Blood | GSM12338 | 29516 | 42269 |
| 783.0224 | 348 | 779.1315 | 420 | 534.8317 | 398 | 551.3219 | 436 | STAG1 | GM10847 | Blood | GSM12338 | 32314 | 42270 |
| 709.1518 | 470 | 751.89 | 624 | 536.1193 | 595 | 506.8723 | 632 | CTCF | GM12878 | Blood | GSM12338 | 61858 | 42271 |
| 743.5644 | 524 | 766.9675 | 666 | 552.7688 | 644 | 509.4441 | 662 | CTCF | GM12878 | Blood | GSM12338 | 64799 | 42272 |
| 807.9498 | 571 | 825.7117 | 714 | 592.5868 | 667 | 549.9808 | 691 | STAG1 | GM12878 | Blood | GSM12339 | 58866 | 42293 |
| 613.2767 | 298 | 682.8948 | 357 | 398.3245 | 377 | 354.0084 | 392 | STAG1 | GM12878 | Blood | GSM12339 | 36239 | 42294 |
| 833.999 | 610 | 868.5115 | 783 | 604.5875 | 706 | 566.0668 | 737 | STAG1 | GM12878 | Blood | GSM12339 | 62107 | 42295 |
| 727.7259 | 480 | 769.9682 | 633 | 534.0046 | 579 | 504.5144 | 615 | CTCF | GM12890 | Blood | GSM12339 | 60537 | 42296 |
| 717.8266 | 484 | 760.0667 | 636 | 534.3457 | 594 | 516.3841 | 650 | CTCF | GM12890 | Blood | GSM12339 | 61004 | 42297 |
| 762.056 | 444 | 795.5093 | 580 | 566.0548 | 525 | 532.7801 | 551 | CTCF | GM12890 | Blood | GSM12339 | 47128 | 42298 |
| 786.0061 | 456 | 797.9436 | 570 | 581.8164 | 534 | 565.2288 | 583 | STAG1 | GM12890 | Blood | GSM12339 | 45980 | 42311 |
| 863.5324 | 564 | 881.4356 | 705 | 623.8843 | 626 | 592.8077 | 661 | STAG1 | GM12890 | Blood | GSM12339 | 49840 | 42312 |
| 667.6745 | 438 | 718.3422 | 594 | 515.4898 | 582 | 487.4728 | 620 | CTCF | GM12891 | Blood | GSM12339 | 63828 | 42313 |
| 694.5239 | 515 | 738.3669 | 680 | 517.3015 | 654 | 498.4785 | 714 | CTCF | GM12891 | Blood | GSM12339 | 73667 | 42314 |
| 774.138 | 560 | 785.2153 | 692 | 559.6774 | 654 | 531.2678 | 698 | STAG1 | GM12891 | Blood | GSM12339 | 63592 | 42331 |
| 821.7928 | 608 | 828.1728 | 742 | 588.8083 | 693 | 558.8859 | 741 | STAG1 | GM12891 | Blood | GSM12339 | 63777 | 42332 |
| 713.6746 | 472 | 744.4442 | 610 | 530.5981 | 582 | 510.1443 | 631 | CTCF | GM12892 | Blood | GSM12339 | 60973 | 42333 |
| 712.6494 | 500 | 747.187 | 650 | 534.0954 | 627 | 489.3251 | 644 | CTCF | GM12892 | Blood | GSM12339 | 66161 | 42334 |
| 796.3046 | 560 | 804.1891 | 684 | 571.4687 | 637 | 544.7477 | 685 | STAG1 | GM12892 | Blood | GSM12339 | 59289 | 42351 |
| 828.5701 | 602 | 848.1679 | 753 | 589.7947 | 678 | 552.8296 | 712 | STAG1 | GM12892 | Blood | GSM12339 | 61722 | 42352 |
| 751.2736 | 513 | 795.7258 | 680 | 556.6852 | 627 | 528.9126 | 670 | CTCF | GM18486 | Blood | GSM12339 | 62968 | 42353 |
| 573.8515 | 250 | 734.1226 | 321 | 396.966 | 318 | 381.9722 | 339 | CTCF | GM18486 | Blood | GSM12339 | 26618 | 42354 |

| | | | | | | | | | | | | | |
|---|---|---|---|---|---|---|---|---|---|---|---|---|---|
| 763.163 | 444 | 797.0909 | 582 | 573.2525 | 535 | 550.1322 | 577 | CTCF | GM18486 | Blood | GSM12339 | 48344 | 42355 |
| 843.7054 | 620 | 862.4641 | 772 | 604.5464 | 700 | 566.6091 | 734 | STAG1 | GM18486 | Blood | GSM12339 | 60992 | 42367 |
| 857.9415 | 596 | 886.3954 | 753 | 621.4387 | 670 | 582.0909 | 698 | STAG1 | GM18486 | Blood | GSM12339 | 55116 | 42368 |
| 642.8039 | 325 | 742.8048 | 459 | 427.4618 | 418 | 381.1839 | 436 | CTCF | GM18505 | Blood | GSM12339 | 41648 | 42369 |
| 412.7687 | 263 | 680.9472 | 379 | 304.9685 | 360 | 312.0204 | 395 | CTCF | GM18505 | Blood | GSM12339 | 39759 | 42370 |
| 369.8685 | 131 | 278.9603 | 127 | 217.2603 | 151 | 221.6147 | 164 | STAG1 | GM18505 | Blood | GSM12340 | 11292 | 42382 |
| 739.7399 | 330 | 747.9995 | 407 | 511.6786 | 405 | 425.0742 | 409 | STAG1 | GM18505 | Blood | GSM12340 | 34354 | 42383 |
| 693.6936 | 369 | 745.0512 | 507 | 488.9885 | 472 | 436.2354 | 493 | CTCF | GM18526 | Blood | GSM12340 | 47337 | 42384 |
| 405.6421 | 258 | 670.4248 | 364 | 296.2724 | 351 | 249.3245 | 360 | CTCF | GM18526 | Blood | GSM12340 | 38479 | 42385 |
| 622.7455 | 283 | 708.5792 | 347 | 426.9447 | 362 | 352.5435 | 365 | STAG1 | GM18526 | Blood | GSM12340 | 32177 | 42397 |
| 730.4924 | 328 | 735.6751 | 404 | 507.7073 | 411 | 391.2318 | 404 | STAG1 | GM18526 | Blood | GSM12340 | 35802 | 42398 |
| 471.4807 | 279 | 681.5701 | 384 | 353.9078 | 380 | 313.4904 | 396 | CTCF | GM18951 | Blood | GSM12340 | 39231 | 42399 |
| 386.2665 | 261 | 654.7948 | 366 | 301.2002 | 366 | 274.2575 | 386 | CTCF | GM18951 | Blood | GSM12340 | 40968 | 42400 |
| 617.1197 | 291 | 692.1679 | 348 | 378.1449 | 357 | 374.1451 | 386 | STAG1 | GM18951 | Blood | GSM12340 | 33390 | 42412 |
| 801.9663 | 406 | 805.8621 | 495 | 600.8363 | 468 | 555.732 | 476 | STAG1 | GM18951 | Blood | GSM12340 | 37311 | 42413 |
| 637.7757 | 338 | 718.0204 | 462 | 449.3376 | 447 | 349.2025 | 445 | CTCF | GM19099 | Blood | GSM12340 | 45428 | 42414 |
| 408.399 | 266 | 656.2822 | 365 | 300.0248 | 364 | 275.589 | 385 | CTCF | GM19099 | Blood | GSM12340 | 40742 | 42415 |
| 461.3662 | 203 | 457.4424 | 226 | 318.4061 | 258 | 307.0838 | 275 | STAG1 | GM19099 | Blood | GSM12340 | 21616 | 42427 |
| 749.4567 | 371 | 750.9728 | 455 | 574.3931 | 459 | 481.1489 | 462 | STAG1 | GM19099 | Blood | GSM12340 | 39436 | 42428 |
| 591.5196 | 317 | 712.4414 | 434 | 418.9962 | 420 | 351.3512 | 429 | CTCF | GM19193 | Blood | GSM12340 | 42534 | 42429 |
| 479.1177 | 305 | 669.4716 | 420 | 347.376 | 415 | 280.9027 | 421 | CTCF | GM19193 | Blood | GSM12340 | 46558 | 42430 |
| 733.7804 | 330 | 718.0422 | 385 | 427.1261 | 384 | 377.3896 | 398 | STAG1 | GM19193 | Blood | GSM12340 | 35517 | 42442 |
| 744.9855 | 357 | 770.826 | 462 | 563.9 | 443 | 500.3247 | 459 | STAG1 | GM19193 | Blood | GSM12340 | 38072 | 42443 |
| 681.5674 | 429 | 746.763 | 600 | 493.1328 | 537 | 492.479 | 587 | CTCF | GM19238 | Blood | GSM12340 | 58041 | 42444 |
| 720.1521 | 532 | 771.0957 | 710 | 538.569 | 668 | 510.7835 | 717 | CTCF | GM19238 | Blood | GSM12340 | 71350 | 42445 |
| 773.2281 | 497 | 776.0899 | 605 | 568.4987 | 583 | 522.3442 | 595 | STAG1 | GM19238 | Blood | GSM12340 | 53755 | 42461 |
| 826.4475 | 593 | 848.8531 | 743 | 607.0284 | 691 | 575.9388 | 737 | STAG1 | GM19238 | Blood | GSM12340 | 60764 | 42462 |
| 687.4398 | 452 | 739.626 | 611 | 507.2159 | 561 | 496.8664 | 622 | CTCF | GM19239 | Blood | GSM12340 | 62649 | 42463 |
| 735.147 | 493 | 784.1071 | 659 | 548.3817 | 607 | 525.9037 | 655 | CTCF | GM19239 | Blood | GSM12341 | 61289 | 42464 |
| 804.1091 | 534 | 832.0946 | 678 | 577.3321 | 602 | 550.4467 | 643 | STAG1 | GM19239 | Blood | GSM12341 | 54648 | 42481 |
| 820.4394 | 602 | 842.0724 | 753 | 597.24 | 699 | 553.4661 | 726 | STAG1 | GM19239 | Blood | GSM12341 | 62439 | 42482 |
| 727.0643 | 491 | 771.6758 | 652 | 527.5712 | 583 | 507.0021 | 633 | CTCF | GM19240 | Blood | GSM12341 | 61383 | 42483 |
| 720.3239 | 505 | 767.9876 | 672 | 530.3481 | 617 | 516.4115 | 682 | CTCF | GM19240 | Blood | GSM12341 | 65593 | 42484 |
| 802.0196 | 509 | 813.9407 | 629 | 591.4475 | 591 | 560.8837 | 626 | STAG1 | GM19240 | Blood | GSM12341 | 50536 | 42502 |
| 828.4078 | 547 | 845.7219 | 686 | 604.7889 | 625 | 570.2878 | 657 | STAG1 | GM19240 | Blood | GSM12341 | 52857 | 42503 |
| 742.5976 | 479 | 788.5581 | 641 | 545.8495 | 578 | 526.6581 | 629 | CTCF | GM2255 | Blood | GSM12341 | 58324 | 42504 |
| 743.8445 | 526 | 802.1086 | 710 | 554.8467 | 648 | 539.7721 | 713 | CTCF | GM2255 | Blood | GSM12341 | 65342 | 42505 |
| 803.4382 | 494 | 829.0587 | 630 | 583.1769 | 562 | 545.255 | 583 | CTCF | GM2255 | Blood | GSM12341 | 49346 | 42506 |
| 812.8567 | 529 | 836.535 | 671 | 590.6708 | 599 | 559.3177 | 636 | STAG1 | GM2255 | Blood | GSM12341 | 51860 | 42518 |
| 857.9559 | 602 | 876.4715 | 754 | 628.0254 | 690 | 588.2349 | 718 | STAG1 | GM2255 | Blood | GSM12341 | 55116 | 42519 |
| 713.9414 | 473 | 758.8703 | 631 | 527.7838 | 582 | 500.2261 | 620 | CTCF | GM2588 | Blood | GSM12341 | 62683 | 42520 |
| 700.6051 | 485 | 751.0735 | 651 | 517.2918 | 598 | 487.5761 | 635 | CTCF | GM2588 | Blood | GSM12341 | 65514 | 42521 |
| 764.8007 | 462 | 793.7555 | 591 | 552.1521 | 523 | 515.3706 | 544 | CTCF | GM2588 | Blood | GSM12341 | 48879 | 42522 |
| 777.357 | 523 | 780.8602 | 634 | 541.2994 | 574 | 513.9689 | 610 | STAG1 | GM2588 | Blood | GSM12341 | 56838 | 42534 |
| 817.9425 | 603 | 835.5561 | 750 | 586.099 | 687 | 555.1899 | 732 | STAG1 | GM2588 | Blood | GSM12341 | 63139 | 42535 |
| 688.2238 | 443 | 723.6143 | 581 | 521.047 | 570 | 495.8232 | 610 | CTCF | GM2610 | Blood | GSM12341 | 61791 | 42536 |
| 660.6347 | 449 | 713.5392 | 612 | 484.6449 | 577 | 486.4228 | 643 | CTCF | GM2610 | Blood | GSM12341 | 67001 | 42537 |
| 756.5471 | 467 | 796.0363 | 616 | 572.8937 | 574 | 524.9804 | 580 | CTCF | GM2610 | Blood | GSM12341 | 51695 | 42538 |
| 759.2402 | 503 | 774.9653 | 631 | 556.7125 | 595 | 539.5893 | 653 | STAG1 | GM2610 | Blood | GSM12341 | 57177 | 42550 |
| 796.1307 | 563 | 805.1871 | 691 | 571.4184 | 643 | 549.468 | 697 | STAG1 | GM2610 | Blood | GSM12341 | 60792 | 42551 |
| 693.8838 | 471 | 733.8573 | 620 | 516.0927 | 591 | 501.0523 | 652 | CTCF | GM2630 | Blood | GSM12341 | 66194 | 42552 |
| 684.971 | 479 | 729.113 | 636 | 505.9668 | 600 | 483.3917 | 649 | CTCF | GM2630 | Blood | GSM12341 | 68513 | 42553 |
| 748.7478 | 471 | 785.8873 | 617 | 547.104 | 552 | 511.8551 | 578 | CTCF | GM2630 | Blood | GSM12342 | 53584 | 42554 |
| 775.6123 | 533 | 806.5576 | 681 | 563.003 | 621 | 527.99 | 649 | STAG1 | GM2630 | Blood | GSM12342 | 59206 | 42566 |
| 813.2643 | 602 | 828.9407 | 748 | 596.2484 | 710 | 555.4492 | 742 | STAG1 | GM2630 | Blood | GSM12342 | 63818 | 42567 |
| 773.9383 | 528 | 814.5883 | 689 | 573.7537 | 636 | 544.0934 | 677 | CTCF | MS1 | Blood | GSM12342 | 60798 | 42568 |
| 727.0022 | 516 | 775.7791 | 684 | 535.3891 | 630 | 513.6414 | 684 | CTCF | MS1 | Blood | GSM12342 | 67080 | 42569 |
| 688.5796 | 502 | 741.8419 | 679 | 517.5832 | 645 | 489.5899 | 690 | CTCF | MS1 | Blood | GSM12342 | 72922 | 42570 |
| 774.7208 | 472 | 816.8248 | 621 | 578.4473 | 562 | 532.2257 | 569 | CTCF | MS1 | Blood | GSM12342 | 49972 | 42571 |
| 819.63 | 589 | 828.5899 | 721 | 575.4807 | 648 | 547.5761 | 693 | STAG1 | MS1 | Blood | GSM12342 | 59950 | 42587 |
| 874.6819 | 590 | 892.8917 | 734 | 632.5757 | 652 | 605.0599 | 699 | STAG1 | MS1 | Blood | GSM12342 | 50899 | 42588 |
| 125.9019 | 34 | 99.02852 | 33 | 80.20299 | 39 | 210.8838 | 68 | ESR1 | MDA-MB-1 | Mammary | GSM12350 | 2221 | 42615 |
| 1088.696 | 377 | 1121.77 | 492 | 809.0526 | 353 | 772.5291 | 363 | EP300 | HEK293 | Embryonic | GSM12390 | 13689 | 42633 |
| 1075.374 | 406 | 1101.943 | 521 | 804.3261 | 387 | 753.4317 | 384 | EP300 | HEK293 | Embryonic | GSM12390 | 15313 | 42634 |
| 1076.296 | 487 | 1112.387 | 631 | 776.245 | 450 | 750.2333 | 480 | CREBBP | HEK293 | Embryonic | GSM12390 | 20139 | 42637 |
| 1015.301 | 531 | 1063.764 | 703 | 741.5815 | 533 | 688.0517 | 529 | CREBBP | HEK293 | Embryonic | GSM12390 | 28178 | 42638 |
| 199.6706 | 101 | 332.2134 | 144 | 113.0304 | 122 | 246.8173 | 181 | CTCF | LoVo | Colon | GSM12393 | 12876 | 42650 |
| 313.1148 | 71 | 298.3703 | 76 | 440.55 | 120 | 734.2413 | 168 | FOXA2 | LoVo | Colon | GSM12393 | 3685 | 42655 |
| 53.15886 | 18 | 34.4382 | 16 | 20.24281 | 17 | 98.91679 | 39 | RAD21 | LoVo | Colon | GSM12394 | 1632 | 42686 |
| 22.58681 | 9 | 31.22048 | 12 | 20.33103 | 13 | 103.8502 | 32 | MAFG | LoVo | Colon | GSM12394 | 1016 | 42692 |
| 260.4424 | 117 | 376.2473 | 156 | 156.8313 | 142 | 231.9241 | 180 | RAD21 | LoVo | Colon | GSM12394 | 13214 | 42706 |
| 930.0715 | 124 | 1208.829 | 172 | 570.17 | 126 | 934.0262 | 176 | ARNT2 | LoVo | Colon | GSM12394 | 3018 | 42707 |

| | | | | | | | | | | | | | |
|---|---|---|---|---|---|---|---|---|---|---|---|---|---|
| 1186.737 | 473 | 1210.178 | 594 | 921.6985 | 469 | 930.8119 | 547 | ATF2 | LoVo | Colon | GSM12394 | 13009 | 42708 |
| 1215.903 | 377 | 1226.551 | 474 | 937.7785 | 358 | 936.1415 | 405 | CREB1 | LoVo | Colon | GSM12394 | 9170 | 42710 |
| 549.3404 | 63 | 722.8309 | 81 | 289.7437 | 57 | 423.2887 | 75 | DR1 | LoVo | Colon | GSM12394 | 1165 | 42711 |
| 1171.15 | 590 | 1180.242 | 719 | 880.5956 | 587 | 882.5531 | 668 | E2F3 | LoVo | Colon | GSM12394 | 20218 | 42712 |
| 1098.681 | 409 | 1155.559 | 558 | 881.859 | 461 | 892.3032 | 538 | ETV5 | LoVo | Colon | GSM12394 | 14985 | 42714 |
| 1157.399 | 512 | 1173.761 | 638 | 893.4175 | 530 | 884.7015 | 593 | GLI2 | LoVo | Colon | GSM12394 | 17360 | 42715 |
| 658.0147 | 70 | 763.4917 | 83 | 313.5684 | 59 | 393.3666 | 71 | HEYL | LoVo | Colon | GSM12394 | 1139 | 42717 |
| 1138.262 | 505 | 1142.765 | 615 | 863.1963 | 500 | 854.9007 | 556 | HOXA4 | LoVo | Colon | GSM12394 | 17119 | 42720 |
| 1185.474 | 542 | 1191.229 | 668 | 885.8478 | 520 | 898.0844 | 606 | HOXA6 | LoVo | Colon | GSM12394 | 17030 | 42722 |
| 1137.15 | 637 | 1157.924 | 788 | 865.2823 | 666 | 855.6453 | 747 | IRF1 | LoVo | Colon | GSM12394 | 25059 | 42725 |
| 1105.326 | 550 | 1125.031 | 688 | 860.5927 | 603 | 862.4328 | 688 | JUN | LoVo | Colon | GSM12394 | 22535 | 42727 |
| 1250.023 | 257 | 1274.792 | 333 | 959.5698 | 220 | 964.2366 | 256 | NFYA | LoVo | Colon | GSM12394 | 4828 | 42738 |
| 1174.311 | 466 | 1191.76 | 581 | 895.5579 | 448 | 885.5613 | 498 | RELA | LoVo | Colon | GSM12394 | 13525 | 42744 |
| 800.0899 | 96 | 981.7694 | 118 | 406.0548 | 86 | 550.2021 | 108 | REST | LoVo | Colon | GSM12394 | 1948 | 42746 |
| 1150.913 | 237 | 1172.693 | 302 | 930.597 | 250 | 930.5059 | 287 | SOX9 | LoVo | Colon | GSM12394 | 6411 | 42747 |
| 1176.394 | 132 | 1288.04 | 177 | 544.0926 | 111 | 780.0035 | 144 | ARNTL | LoVo | Colon | GSM12394 | 2368 | 42749 |
| 1195.385 | 266 | 1215.925 | 345 | 951.876 | 271 | 971.9409 | 324 | BARX1 | LoVo | Colon | GSM12394 | 6451 | 42752 |
| 1097.879 | 393 | 1126.553 | 510 | 905.5872 | 469 | 872.0943 | 491 | FOXP1 | LoVo | Colon | GSM12395 | 14445 | 42760 |
| 1253.195 | 195 | 1298.011 | 266 | 1015.835 | 196 | 1002.253 | 210 | GMEB2 | LoVo | Colon | GSM12395 | 3558 | 42762 |
| 1231.731 | 257 | 1276.86 | 347 | 983.224 | 253 | 996.3563 | 297 | JUNB | LoVo | Colon | GSM12395 | 5414 | 42770 |
| 711.9492 | 95 | 1090.55 | 134 | 551.2818 | 110 | 840.7481 | 148 | MXD3 | LoVo | Colon | GSM12395 | 2291 | 42772 |
| 1297.376 | 366 | 1284.548 | 437 | 1003.7 | 329 | 982.597 | 353 | NFYC | LoVo | Colon | GSM12395 | 6802 | 42774 |
| 1200.138 | 441 | 1214.499 | 545 | 928.7751 | 432 | 930.0552 | 492 | NPAT | LoVo | Colon | GSM12395 | 11801 | 42775 |
| 354.9648 | 46 | 439.6416 | 57 | 223.3402 | 47 | 432.6445 | 73 | OVOL2 | LoVo | Colon | GSM12395 | 1062 | 42778 |
| 1242.808 | 143 | 1262.201 | 183 | 729.4567 | 138 | 777.7317 | 151 | TAL1 | LoVo | Colon | GSM12395 | 2706 | 42788 |
| 963.5323 | 655 | 981.7386 | 813 | 724.5993 | 752 | 719.0908 | 854 | RAD21 | LoVo | Colon | GSM12395 | 48737 | 42798 |
| 1180.163 | 280 | 1191.475 | 351 | 950.4785 | 294 | 934.725 | 321 | STAT2 | LoVo | Colon | GSM12395 | 7152 | 42800 |
| 1130.296 | 150 | 1187.307 | 197 | 753.5977 | 160 | 901.7996 | 187 | ZBTB10 | LoVo | Colon | GSM12395 | 3665 | 42804 |
| 148.0511 | 56 | 175.3777 | 68 | 97.69825 | 69 | 201.3614 | 103 | CTCFL | LoVo | Colon | GSM12395 | 5432 | 42819 |
| 172.9393 | 104 | 257.4006 | 140 | 107.2009 | 133 | 191.006 | 180 | RAD21 | LoVo | Colon | GSM12395 | 15125 | 42845 |
| 295.8317 | 87 | 282.2253 | 94 | 174.7866 | 98 | 275.7986 | 130 | STAG1 | LoVo | Colon | GSM12395 | 6299 | 42846 |
| 484.2941 | 196 | 706.9501 | 262 | 290.0295 | 235 | 369.304 | 280 | CTCF | LoVo | Colon | GSM12395 | 20606 | 42848 |
| 454.2406 | 72 | 592.8368 | 92 | 359.0627 | 86 | 685.1798 | 131 | GRHL1 | LoVo | Colon | GSM12395 | 2255 | 42850 |
| 425.6365 | 92 | 471.5708 | 107 | 334.4963 | 114 | 510.2546 | 151 | MAFK | LoVo | Colon | GSM12395 | 4535 | 42854 |
| 621.2793 | 73 | 819.964 | 94 | 422.2641 | 77 | 572.7001 | 97 | ZNF12 | LoVo | Colon | GSM12396 | 1415 | 42882 |
| 1142.639 | 249 | 1159.67 | 315 | 872.5538 | 229 | 881.067 | 263 | ZNF143 | LoVo | Colon | GSM12396 | 6676 | 42883 |
| 1040.502 | 352 | 1075.235 | 462 | 843.195 | 412 | 846.1245 | 471 | ZNF83 | LoVo | Colon | GSM12396 | 15238 | 42890 |
| 1206.575 | 440 | 1229.507 | 565 | 958.9908 | 468 | 935.2678 | 499 | XBP1 | T47D | Mammary | GSM12396 | 12086 | 42895 |
| 24.54944 | 13 | 54.81835 | 22 | 35.63032 | 24 | 122.9206 | 47 | RELA | AC16 | None | GSM12399 | 1892 | 42900 |
| 431.5206 | 69 | 560.8376 | 88 | 207.0811 | 63 | 269.4142 | 77 | RELA | AC16 | None | GSM12399 | 2236 | 42901 |
| 663.6349 | 216 | 812.0756 | 270 | 424.8193 | 259 | 447.4938 | 285 | KMT2B | HCT-116 | Colon | GSM12401 | 18591 | 42906 |
| 531.1262 | 191 | 720.6524 | 247 | 356.738 | 237 | 432.3576 | 277 | EP300 | HCT-116 | Colon | GSM12401 | 18212 | 42907 |
| 314.3516 | 245 | 530.843 | 342 | 235.9169 | 345 | 295.5729 | 406 | CTCF | GP5d | Colon | GSM12408 | 43216 | 42918 |
| 170.1299 | 78 | 353.9633 | 125 | 277.3826 | 146 | 617.5572 | 229 | HNF4A | GP5d | Colon | GSM12408 | 8811 | 42920 |
| 263.9419 | 53 | 320.6469 | 65 | 244.2035 | 71 | 418.6406 | 101 | JUND | GP5d | Colon | GSM12408 | 2368 | 42924 |
| 769.1415 | 152 | 1020.761 | 210 | 679.7162 | 198 | 864.2468 | 245 | KLF5 | GP5d | Colon | GSM12408 | 6656 | 42925 |
| 1107.106 | 356 | 1158.377 | 485 | 886.4597 | 389 | 896.1954 | 454 | MYC | GP5d | Colon | GSM12408 | 12121 | 42926 |
| 1082.674 | 283 | 1125.184 | 385 | 889.0624 | 330 | 900.7115 | 384 | NIPBL | GP5d | Colon | GSM12408 | 9827 | 42927 |
| 735.7509 | 352 | 781.3604 | 476 | 582.8979 | 461 | 579.7854 | 523 | RAD21 | GP5d | Colon | GSM12408 | 39962 | 42928 |
| 17.08862 | 15 | 30.27116 | 22 | 10.61937 | 20 | 49.25246 | 40 | REST | GP5d | Colon | GSM12408 | 2857 | 42929 |
| 756.7886 | 365 | 786.1568 | 476 | 587.9806 | 457 | 566.3774 | 492 | SMC3 | GP5d | Colon | GSM12408 | 38656 | 42931 |
| 448.5063 | 201 | 728.4895 | 282 | 318.3382 | 262 | 358.7568 | 297 | CTCF | GP5d | Colon | GSM12408 | 23436 | 42932 |
| 97.14879 | 27 | 157.0837 | 39 | 117.4312 | 43 | 228.1083 | 65 | HNF4A | GP5d | Colon | GSM12408 | 1850 | 42934 |
| 1137.819 | 200 | 1155.852 | 261 | 895.6714 | 211 | 916.8244 | 241 | JUND | GP5d | Colon | GSM12408 | 5531 | 42938 |
| 772.9095 | 105 | 1070.662 | 139 | 493.4547 | 109 | 611.7144 | 130 | MYC | GP5d | Colon | GSM12408 | 2562 | 42940 |
| 1048.602 | 116 | 1276.569 | 145 | 679.3112 | 118 | 753.2148 | 132 | NIPBL | GP5d | Colon | GSM12408 | 2002 | 42941 |
| 403.8529 | 156 | 560.492 | 204 | 254.0144 | 189 | 308.3717 | 221 | RAD21 | GP5d | Colon | GSM12408 | 15398 | 42942 |
| 15.47226 | 13 | 26.94409 | 19 | 9.173343 | 17 | 50.78183 | 37 | REST | GP5d | Colon | GSM12408 | 2465 | 42943 |
| 117.9462 | 57 | 216.9953 | 86 | 116.0513 | 86 | 175.7274 | 111 | SMC3 | GP5d | Colon | GSM12408 | 6979 | 42945 |
| 286.893 | 90 | 395.9194 | 118 | 261.3807 | 125 | 440.4311 | 172 | HNF4A | LoVo | Colon | GSM12422 | 6924 | 42947 |
| 205.8191 | 34 | 304.0832 | 47 | 110.7479 | 33 | 307.2614 | 62 | KLF5 | LoVo | Colon | GSM12422 | 1149 | 42949 |
| 1133.613 | 159 | 1169.814 | 212 | 657.6334 | 158 | 821.35 | 189 | MYC | LoVo | Colon | GSM12422 | 4176 | 42950 |
| 211.1016 | 118 | 293.1528 | 154 | 147.6852 | 157 | 178.7794 | 183 | RAD21 | LoVo | Colon | GSM12422 | 16450 | 42951 |
| 264.5031 | 145 | 350.4111 | 185 | 137.236 | 172 | 217.9711 | 222 | RAD21 | LoVo | Colon | GSM12422 | 19962 | 42957 |
| 1226.029 | 387 | 1225.441 | 475 | 923.0402 | 344 | 888.4979 | 355 | BRD2 | MM1.S | Blood | GSM12478 | 9387 | 43045 |
| 1171.971 | 331 | 1198.806 | 434 | 912.007 | 326 | 882.8449 | 340 | BRD2 | MM1.S | Blood | GSM12478 | 9136 | 43046 |
| 1146.775 | 176 | 1175.85 | 233 | 736.2736 | 176 | 884.2036 | 206 | BRD2 | MM1.S | Blood | GSM12478 | 4723 | 43047 |
| 1117.56 | 576 | 1153.816 | 738 | 805.0898 | 542 | 767.0114 | 561 | BRD3 | MM1.S | Blood | GSM12478 | 24273 | 43050 |
| 1128.305 | 288 | 1158.462 | 381 | 864.7033 | 276 | 839.7202 | 294 | BRD3 | MM1.S | Blood | GSM12478 | 8790 | 43051 |
| 1131.753 | 298 | 1164.069 | 398 | 849.8954 | 273 | 836.0581 | 297 | BRD3 | MM1.S | Blood | GSM12478 | 9162 | 43052 |
| 1117.371 | 260 | 1161.023 | 353 | 855.0728 | 246 | 838.7456 | 266 | BRD3 | MM1.S | Blood | GSM12478 | 8030 | 43053 |
| 559.6897 | 203 | 834.5247 | 300 | 471.4644 | 277 | 493.5737 | 304 | TCF7L2 | LNCaP | Prostate | GSM12494 | 19097 | 43061 |

| | | | | | | | | | | | | | |
|---|---|---|---|---|---|---|---|---|---|---|---|---|---|
| 213.1743 | 54 | 292.5269 | 71 | 159.9063 | 67 | 180.8752 | 76 | JMJD6 | HEK293T | Embryonic | GSM12498 | 3374 | 43063 |
| 339.7184 | 113 | 444.2332 | 144 | 174.7229 | 123 | 212.3923 | 144 | JMJD6 | HEK293T | Embryonic | GSM12498 | 9755 | 43064 |
| 1139.679 | 178 | 1159.756 | 229 | 792.5797 | 185 | 601.0215 | 169 | BRD4 | HEK293T | Embryonic | GSM12498 | 4597 | 43065 |
| 1235.12 | 457 | 1282.878 | 608 | 950.4864 | 429 | 910.6118 | 441 | BRD4 | HEK293T | Embryonic | GSM12498 | 10833 | 43066 |
| 734.054 | 115 | 1157.071 | 175 | 367.4421 | 107 | 400.9891 | 119 | EP300 | HEK293T | Embryonic | GSM12498 | 3519 | 43074 |
| 1295.049 | 564 | 1319.7 | 713 | 946.2746 | 486 | 926.1552 | 522 | CDK9 | HEK293T | Embryonic | GSM12498 | 13103 | 43081 |
| 627.8296 | 77 | 660.2288 | 86 | 358.8053 | 74 | 382.4061 | 81 | CDK9 | HEK293T | Embryonic | GSM12498 | 1621 | 43082 |
| 1147.76 | 329 | 1130.282 | 385 | 839.8431 | 286 | 825.653 | 313 | CDK9 | HEK293T | Embryonic | GSM12498 | 10247 | 43083 |
| 391.1873 | 96 | 522.1318 | 124 | 232.7612 | 106 | 244.4667 | 116 | JMJD6 | HeLa | Cervix | GSM12499 | 5784 | 43089 |
| 306.6957 | 45 | 427.0895 | 60 | 186.4515 | 46 | 344.82 | 69 | BRD4 | HeLa | Cervix | GSM12499 | 1283 | 43090 |
| 570.8297 | 158 | 630.705 | 184 | 252.0188 | 155 | 232.9491 | 161 | CDK8 | MM1.S | Blood | GSM12520 | 10732 | 43095 |
| 1004.473 | 648 | 1020.117 | 801 | 725.9322 | 680 | 690.2927 | 718 | ETS1 | CUTLL1 | Blood | GSM12529 | 41591 | 43097 |
| 376.5951 | 225 | 501.2918 | 287 | 144.6034 | 248 | 132.5504 | 263 | GABPA | CUTLL1 | Blood | GSM12529 | 35865 | 43098 |
| 1003.989 | 539 | 1029.667 | 687 | 760.5898 | 593 | 683.5275 | 562 | RUNX1 | CUTLL1 | Blood | GSM12529 | 32040 | 43099 |
| 1263.687 | 354 | 1283.574 | 452 | 977.0556 | 326 | 946.3808 | 336 | NOTCH1 | CUTLL1 | Blood | GSM12529 | 7480 | 43101 |
| 1180.81 | 411 | 1207.842 | 533 | 903.6951 | 397 | 852.4578 | 390 | RBPJ | CUTLL1 | Blood | GSM12529 | 11914 | 43102 |
| 1164.014 | 492 | 1193.677 | 632 | 899.6736 | 498 | 846.6867 | 496 | NOTCH1 | CUTLL1 | Blood | GSM12529 | 16128 | 43104 |
| 1145.287 | 482 | 1176.362 | 627 | 862.9919 | 475 | 808.9836 | 468 | RBPJ | CUTLL1 | Blood | GSM12529 | 17073 | 43105 |
| 1161.024 | 652 | 1181.022 | 812 | 847.6 | 632 | 782.4534 | 618 | BRD4 | CUTLL1 | Blood | GSM12529 | 24930 | 43109 |
| 1134.826 | 559 | 1158.734 | 705 | 851.5223 | 558 | 781.3257 | 530 | MED1 | CUTLL1 | Blood | GSM12529 | 21090 | 43110 |
| 1039.309 | 516 | 1066.578 | 656 | 765.3646 | 520 | 720.4305 | 528 | EP300 | CUTLL1 | Blood | GSM12529 | 26535 | 43111 |
| 24.31808 | 12 | 15.8895 | 11 | 57.60416 | 28 | 74.43613 | 34 | GATA6 | AGS | Stomach | GSM12555 | 1647 | 43161 |
| 88.57023 | 36 | 141.1461 | 51 | 338.9687 | 105 | 389.5452 | 120 | GATA6 | HUG1N | Stomach | GSM12555 | 3818 | 43163 |
| 763.1209 | 364 | 758.2921 | 439 | 572.7421 | 444 | 570.3905 | 499 | NR2F2 | None | None | GSM12573 | 40414 | 43165 |
| 1208.966 | 146 | 1210.583 | 174 | 570.0864 | 126 | 679.8606 | 147 | FOXM1 | HeLa | Cervix | GSM12593 | 2992 | 43194 |
| 644.7766 | 69 | 708.9313 | 79 | 353.545 | 63 | 435.2727 | 75 | FOXM1 | HeLa | Cervix | GSM12593 | 1101 | 43195 |
| 979.8416 | 494 | 958.961 | 572 | 723.4024 | 514 | 704.8355 | 559 | CHRM2 | HeLa | Cervix | GSM12641 | 28422 | 43234 |
| 45.41208 | 17 | 54.98334 | 21 | 8.126419 | 12 | 75.31185 | 35 | TEAD4 | WA09 | Embryo | GSM12668 | 1734 | 43300 |
| 54.42388 | 18 | 59.3735 | 21 | 20.91037 | 17 | 111.5506 | 41 | TEAD4 | WA09 | Embryo | GSM12668 | 1601 | 43301 |
| 3.760112 | 26 | 14.93999 | 45 | 5.336067 | 53 | 13.38199 | 73 | SMAD2 | WA09 | Embryo | GSM12668 | 14973 | 43302 |
| 1168.481 | 471 | 1193.927 | 606 | 888.3083 | 463 | 848.9584 | 479 | CDK6 | KB Cell | HeLa conta | GSM12674 | 15332 | 43305 |
| 1005.251 | 279 | 1051.017 | 384 | 775.054 | 294 | 757.0105 | 318 | RELA | KB Cell | HeLa conta | GSM12674 | 12959 | 43313 |
| 935.6344 | 634 | 866.4259 | 670 | 714.817 | 760 | 771.5464 | 980 | AR | LNCaP | Prostate | GSM12748 | 53592 | 43395 |
| 963.0282 | 235 | 921.7628 | 259 | 658.2528 | 256 | 847.2269 | 392 | AR | LNCaP | Prostate | GSM12748 | 12235 | 43396 |
| 970.191 | 812 | 865.0779 | 836 | 768.6531 | 1070 | 812.347 | 1313 | FOXA1 | LNCaP | Prostate | GSM12748 | 82135 | 43397 |
| 450.6046 | 69 | 723.6151 | 100 | 431.8813 | 91 | 346.4837 | 85 | XBP1 | MDA-MB-2 | Breast | GSM12761 | 1959 | 43398 |
| 326.9207 | 50 | 384.6141 | 60 | 164.4026 | 46 | 135.5122 | 44 | XBP1 | MDA-MB-2 | Breast | GSM12761 | 1436 | 43399 |
| 1237.16 | 525 | 1252.225 | 649 | 976.3601 | 549 | 944.7192 | 571 | XBP1 | T47D | Mammary | GSM12761 | 13783 | 43400 |
| 661.4117 | 96 | 1048.83 | 138 | 504.8442 | 111 | 336.7458 | 94 | XBP1 | HS578T | Mammary | GSM12761 | 2575 | 43401 |
| 678.0546 | 627 | 707.8382 | 797 | 529.5607 | 884 | 499.8593 | 952 | GATA1 | None | Bone Marr | GSM12782 | 120643 | 43446 |
| 388.7561 | 182 | 562.3337 | 242 | 377.5226 | 272 | 301.3056 | 268 | TAL1 | None | Bone Marr | GSM12782 | 22558 | 43447 |
| 60.97429 | 29 | 132.2683 | 48 | 60.1942 | 44 | 80.89878 | 54 | GFI1B | None | Bone Marr | GSM12782 | 3587 | 43448 |
| 966.0451 | 147 | 1132.233 | 197 | 481.3341 | 135 | 466.1429 | 141 | SUPT5H | HeLa | Cervix | GSM12802 | 4063 | 43450 |
| 1145.066 | 312 | 1173.189 | 407 | 884.5132 | 296 | 842.8998 | 297 | NELFA | HeLa | Cervix | GSM12802 | 8342 | 43451 |
| 46.52404 | 14 | 54.65744 | 17 | 78.35595 | 27 | 113.3899 | 35 | C17orf96 | HEK293T | Embryonic | GSM12949 | 1122 | 43550 |
| 319.4903 | 55 | 516.1467 | 80 | 216.0606 | 61 | 355.6056 | 85 | CBX4 | HEK293T | Embryonic | GSM12949 | 1970 | 43555 |
| 506.5516 | 72 | 548.2169 | 82 | 203.5693 | 58 | 188.0409 | 59 | CBX6 | None | Mammary | GSM12950 | 1847 | 43556 |
| 21.62985 | 11 | 31.55186 | 15 | 30.63303 | 20 | 45.82419 | 26 | CBX8 | None | Mammary | GSM12950 | 1520 | 43558 |
| 98.64991 | 25 | 82.56224 | 25 | 27.33111 | 19 | 118.8506 | 42 | RING1 | None | Mammary | GSM12950 | 1531 | 43560 |
| 276.2008 | 51 | 475.7296 | 77 | 98.77842 | 41 | 256.9415 | 72 | RNF2 | None | Mammary | GSM12950 | 2028 | 43561 |
| 117.604 | 47 | 179.9838 | 65 | 42.69375 | 45 | 73.69842 | 61 | RING1 | None | Foreskin | GSM12950 | 4607 | 43571 |
| 685.1097 | 172 | 918.0821 | 228 | 305.0112 | 166 | 459.1411 | 215 | RNF2 | None | Foreskin | GSM12950 | 10260 | 43572 |
| 1169.565 | 439 | 1159.24 | 517 | 865.538 | 387 | 818.4131 | 385 | CDK7 | Jurkat | Blood | GSM12963 | 11992 | 43592 |
| 1092.164 | 490 | 1103.695 | 608 | 802.4788 | 467 | 772.0326 | 490 | BRD4 | OCI-Ly1 | Bone Marr | GSM12966 | 19377 | 43631 |
| 1048.12 | 540 | 1060.125 | 665 | 764.6223 | 532 | 739.6686 | 567 | BRD4 | OCI-Ly1 | Bone Marr | GSM12966 | 25986 | 43633 |
| 1092.046 | 491 | 1118.242 | 620 | 808.684 | 474 | 778.836 | 498 | MED1 | OCI-Ly1 | Bone Marr | GSM12966 | 19662 | 43634 |
| 980.5865 | 294 | 962.0039 | 344 | 738.854 | 303 | 722.8751 | 331 | CDK8 | SW480 | Colon | GSM12966 | 15061 | 43642 |
| 62.68364 | 91 | 97.5106 | 123 | 37.17243 | 131 | 31.86959 | 138 | DPPA2 | HEK293T | Embryonic | GSM13011 | 26255 | 43668 |
| 78.97777 | 77 | 191.7114 | 127 | 53.83241 | 110 | 78.41071 | 136 | TET3 | HEK293T | Embryonic | GSM13011 | 17289 | 43669 |
| 15.38931 | 100 | 50.409 | 160 | 16.58121 | 188 | 34.32555 | 242 | CDK2 | T47D-MTVI | Mammary | GSM13021 | 54318 | 43676 |
| 47.37192 | 48 | 45.32268 | 54 | 49.85135 | 80 | 75.41019 | 101 | CDK2 | T47D-MTVI | Mammary | GSM13021 | 11050 | 43677 |
| 1168.473 | 603 | 1236.159 | 815 | 853.4098 | 567 | 837.8479 | 621 | KDM4C | KYSE-150 | Esophagus | GSM13037 | 21031 | 43684 |
| 1166.065 | 343 | 1174.624 | 426 | 878.5868 | 316 | 848.8614 | 331 | CDK9 | MM1.S | Blood | GSM13139 | 9890 | 43741 |
| 1123.963 | 520 | 1143.18 | 652 | 825.2731 | 486 | 777.328 | 492 | BRD4 | KOPT-K1 | Blood | GSM13141 | 19018 | 43742 |
| 1137.35 | 476 | 1143.723 | 586 | 835.4357 | 441 | 779.8324 | 431 | BRD4 | KOPT-K1 | Blood | GSM13141 | 16282 | 43749 |
| 1133.598 | 325 | 1168.256 | 430 | 866.6161 | 309 | 827.8261 | 311 | BRD4 | DND-41 | Blood | GSM13141 | 9737 | 43756 |
| 1148.869 | 446 | 1201.113 | 602 | 858.2957 | 414 | 822.656 | 430 | BRD4 | DND-41 | Blood | GSM13141 | 13961 | 43763 |
| 236.8125 | 96 | 249.6065 | 110 | 148.3213 | 117 | 142.3585 | 124 | TERF1 | LCL | Blood | GSM13288 | 10016 | 43825 |
| 248.2139 | 76 | 252.2324 | 85 | 141.306 | 85 | 90.15455 | 75 | TERF2 | LCL | Blood | GSM13288 | 5903 | 43826 |
| 560.9159 | 273 | 734.6152 | 378 | 394.0777 | 361 | 451.676 | 412 | CTCF | THP-1 | Blood | GSM13355 | 36336 | 43831 |
| 296.1203 | 133 | 376.9948 | 167 | 149.6514 | 152 | 212.9759 | 189 | RAD21 | THP-1 | Blood | GSM13355 | 15848 | 43832 |
| 802.9182 | 596 | 743.7565 | 644 | 729.0243 | 954 | 792.2561 | 1220 | AR | VCaP | Prostate | GSM81734 | 74342 | 43853 |

| | | | | | | | | | | | | | |
|---|---|---|---|---|---|---|---|---|---|---|---|---|---|
| 977.4512 | 481 | 912.6133 | 505 | 873.5773 | 722 | 920.2625 | 913 | AR | VCaP | Prostate | GSM81734 | 29675 | 43854 |
| 827.3373 | 592 | 747.3655 | 611 | 741.5569 | 926 | 796.0395 | 1172 | AR | VCaP | Prostate | GSM81734 | 67834 | 43855 |
| 912.4832 | 547 | 825.4831 | 553 | 804.0098 | 823 | 858.431 | 1046 | AR | VCaP | Prostate | GSM81735 | 45562 | 43856 |
| 984.7341 | 445 | 937.3284 | 486 | 878.6871 | 667 | 939.3793 | 865 | AR | VCaP | Prostate | GSM81735 | 25955 | 43857 |
| 885.5198 | 538 | 801.4807 | 546 | 777.8816 | 805 | 839.3221 | 1039 | AR | VCaP | Prostate | GSM81735 | 47923 | 43858 |
| 942.0661 | 495 | 877.0385 | 524 | 839.3788 | 752 | 891.1768 | 957 | AR | VCaP | Prostate | GSM81735 | 35169 | 43859 |
| 973.925 | 575 | 894.169 | 592 | 861.5558 | 856 | 900.5783 | 1059 | AR | VCaP | Prostate | GSM81735 | 40396 | 43860 |
| 705.3877 | 344 | 670.5598 | 386 | 491.9114 | 447 | 511.1818 | 492 | HOXC9 | BE2-C | Brain | GSM84878 | 46643 | 43893 |
| 630.6721 | 289 | 651.0827 | 329 | 413.808 | 369 | 444.7761 | 411 | HOXC9 | BE2-C | Brain | GSM84878 | 36212 | 43894 |
| 100.4802 | 30 | 233.3032 | 53 | 52.82702 | 32 | 129.6597 | 53 | T | None | Artery | GSM91697 | 2257 | 43972 |
| 348.6044 | 92 | 533.9013 | 128 | 173.3532 | 94 | 251.8062 | 120 | T | None | Artery | GSM91697 | 5883 | 43975 |
| 494.9468 | 120 | 686.3175 | 158 | 295.5252 | 132 | 312.7097 | 145 | T | None | Artery | GSM91697 | 6923 | 43978 |
| 83.55011 | 21 | 179.2828 | 36 | 115.7737 | 36 | 77.49485 | 31 | TP53 | H9 | Embryo | GSM98123 | 1337 | 43998 |
| 395.763 | 120 | 518.4445 | 153 | 225.0701 | 134 | 208.067 | 139 | TP53 | H9 | Embryo | GSM98123 | 9151 | 43999 |
| 55.11129 | 21 | 44.28084 | 21 | 20.00494 | 20 | 45.57317 | 31 | MEIS1 | None | Cord blood | GSM63831 | 2156 | 4400 |
| 398.2146 | 88 | 736.2844 | 137 | 283.1732 | 104 | 254.0482 | 105 | TP53 | H9 | Embryo | GSM98123 | 4494 | 44000 |
| 45.19936 | 18 | 28.51594 | 16 | 6.576602 | 12 | 38.05937 | 27 | MEIS1 | None | Cord blood | GSM63831 | 1956 | 4402 |
| 990.8694 | 284 | 999.7255 | 353 | 765.7447 | 303 | 746.7599 | 329 | AGO1 | PC-3 | Prostate | GSM99594 | 13977 | 44026 |
| 1215.554 | 364 | 1248.999 | 477 | 941.3745 | 341 | 896.6171 | 339 | BRCA1 | MCF-10A | Breast | GSM99754 | 8357 | 44028 |
| 1168.881 | 223 | 1219.57 | 311 | 922.11 | 223 | 727.4349 | 203 | KAT8 | None | Blood | GSM39394 | 5842 | 4403 |
| 1072.974 | 191 | 1100.675 | 250 | 715.6033 | 200 | 758.3261 | 219 | ETS1 | None | Blood | GSM10569 | 6601 | 44090 |
| 908.8832 | 128 | 1152.239 | 163 | 471.6822 | 120 | 548.4452 | 138 | ETS1 | None | Blood | GSM10569 | 3473 | 44091 |
| 1083.334 | 376 | 1113.797 | 492 | 831.7925 | 387 | 793.8038 | 393 | ETS1 | None | Blood | GSM10569 | 14700 | 44092 |
| 1112.271 | 268 | 1144.004 | 357 | 858.0272 | 265 | 819.4264 | 270 | ETS1 | None | Blood | GSM10569 | 8828 | 44093 |
| 1087.885 | 369 | 1132.181 | 495 | 840.0252 | 377 | 792.4633 | 375 | ETS1 | None | Blood | GSM10569 | 13949 | 44094 |
| 346.6027 | 48 | 423.6305 | 59 | 191.3265 | 46 | 253.2736 | 57 | RUNX1 | None | Blood | GSM10569 | 1236 | 44095 |
| 708.1842 | 97 | 949.512 | 126 | 403.7067 | 95 | 482.7654 | 111 | RUNX1 | None | Blood | GSM10569 | 2481 | 44096 |
| 1065.749 | 198 | 1122.921 | 283 | 705.5377 | 204 | 638.466 | 206 | RUNX1 | None | Blood | GSM10569 | 6976 | 44097 |
| 1110.266 | 176 | 1109.386 | 218 | 681.1058 | 179 | 674.9149 | 189 | FOXP3 | None | Blood | GSM10569 | 5465 | 44098 |
| 613.3722 | 89 | 712.9564 | 106 | 335.5352 | 86 | 419.7795 | 103 | FOXP3 | None | Blood | GSM10569 | 2503 | 44099 |
| 711.4406 | 267 | 781.0325 | 343 | 686.4652 | 435 | 699.7333 | 515 | TFAP2A | MCF-7 | Breast | GSM10813 | 27988 | 44122 |
| 153.8269 | 79 | 299.6523 | 122 | 98.46686 | 101 | 143.5351 | 127 | INO80 | None | Blood | GSM10829 | 10590 | 44130 |
| 46.07534 | 69 | 70.87828 | 93 | 18.7362 | 90 | 44.7683 | 126 | SRCAP | None | Blood | GSM10829 | 20090 | 44131 |
| 100.4002 | 38 | 154.7688 | 53 | 124.7639 | 63 | 155.9478 | 75 | TAL1 | None | Cord blood | GSM10854 | 3800 | 44136 |
| 768.3589 | 96 | 1086.885 | 129 | 315.5972 | 77 | 408.0794 | 94 | PHF6 | Jurkat | Blood | GSM11167 | 2105 | 44144 |
| 636.2844 | 80 | 932.396 | 110 | 369.6356 | 78 | 423.6254 | 89 | KAT2B | None | Blood | GSM39394 | 1769 | 4418 |
| 468.2335 | 132 | 688.0102 | 179 | 322.9685 | 160 | 266.4589 | 157 | PML | None | Bone Marr | GSM46820 | 9512 | 4420 |
| 354.6297 | 84 | 423.8503 | 102 | 172.5032 | 83 | 303.6808 | 117 | SOX2 | KYSE-70 | None | GSM11390 | 4619 | 44224 |
| 56.7127 | 26 | 106.2009 | 40 | 45.87077 | 36 | 88.19665 | 52 | TP63 | KYSE-70 | None | GSM11390 | 3116 | 44225 |
| 329.5197 | 93 | 433.2937 | 119 | 284.8851 | 125 | 342.4135 | 146 | SOX2 | TT | Thyroid | GSM11390 | 6500 | 44227 |
| 41.37853 | 34 | 81.22244 | 52 | 55.18065 | 61 | 105.6918 | 86 | TP63 | TT | Thyroid | GSM11390 | 6648 | 44228 |
| 36.29221 | 21 | 67.63105 | 32 | 51.13543 | 38 | 90.75425 | 53 | SOX2 | HCC95 | Lung | GSM11390 | 3143 | 44230 |
| 42.29202 | 33 | 38.33856 | 36 | 53.97668 | 58 | 63.6269 | 67 | TP63 | HCC95 | Lung | GSM11390 | 6202 | 44231 |
| 27.91471 | 16 | 21.44278 | 16 | 18.33925 | 21 | 50.22561 | 35 | SOX2 | H9 | Embryo | GSM11390 | 2448 | 44233 |
| 967.5999 | 334 | 923.9667 | 367 | 799.162 | 428 | 887.6402 | 609 | AR | LNCaP | Prostate | GSM12078 | 18855 | 44310 |
| 962.4098 | 220 | 951.0142 | 261 | 799.393 | 278 | 892.7532 | 412 | AR | LNCaP | Prostate | GSM12078 | 11390 | 44311 |
| 1359.062 | 811 | 1404.952 | 1028 | 987.682 | 776 | 964.4645 | 843 | E2F1 | LNCaP | Prostate | GSM12078 | 22268 | 44312 |
| 1344.063 | 662 | 1376.832 | 843 | 996.8483 | 608 | 958.4004 | 638 | E2F1 | LNCaP | Prostate | GSM12078 | 15380 | 44313 |
| 871.1736 | 127 | 911.6082 | 142 | 524.6601 | 130 | 1059.881 | 245 | AR | LNCaP | Prostate | GSM12078 | 3663 | 44314 |
| 858.5668 | 123 | 915.1452 | 139 | 621.6277 | 139 | 1063.711 | 237 | AR | LNCaP | Prostate | GSM12078 | 3443 | 44315 |
| 1343.689 | 696 | 1389.653 | 897 | 995.1338 | 650 | 952.2692 | 667 | E2F1 | LNCaP | Prostate | GSM12078 | 16724 | 44316 |
| 1321.199 | 661 | 1392.195 | 885 | 994.6829 | 632 | 956.9931 | 653 | E2F1 | LNCaP | Prostate | GSM12078 | 16149 | 44317 |
| 982.1517 | 463 | 904.5624 | 472 | 788.72 | 569 | 872.1785 | 785 | AR | LNCaP | Prostate | GSM12078 | 28047 | 44318 |
| 814.6226 | 172 | 918.5294 | 202 | 514.4247 | 192 | 892.2157 | 327 | AR | LNCaP | Prostate | GSM12078 | 8741 | 44319 |
| 1330.759 | 751 | 1389.678 | 965 | 989.9552 | 726 | 949.0352 | 754 | E2F1 | LNCaP | Prostate | GSM12078 | 20087 | 44320 |
| 1314.434 | 605 | 1352.191 | 774 | 984.0497 | 557 | 951.5406 | 585 | E2F1 | LNCaP | Prostate | GSM12078 | 14154 | 44321 |
| 249.9587 | 37 | 237.3092 | 39 | 132.0633 | 35 | 511.8809 | 81 | AR | LNCaP | Prostate | GSM12079 | 1077 | 44323 |
| 1349.45 | 702 | 1407.358 | 907 | 993.0706 | 642 | 958.5735 | 671 | E2F1 | LNCaP | Prostate | GSM12079 | 16567 | 44324 |
| 1327.229 | 637 | 1368.992 | 822 | 991.2161 | 588 | 958.2593 | 622 | E2F1 | LNCaP | Prostate | GSM12079 | 14892 | 44325 |
| 957.0384 | 417 | 887.4679 | 431 | 787.8081 | 535 | 861.2627 | 731 | AR | LNCaP | Prostate | GSM12079 | 26201 | 44326 |
| 944.7652 | 256 | 917.3948 | 290 | 769.7069 | 314 | 880.8075 | 485 | AR | LNCaP | Prostate | GSM12079 | 14409 | 44327 |
| 1286.976 | 426 | 1339.943 | 569 | 1000.869 | 400 | 963.8812 | 414 | E2F1 | LNCaP | Prostate | GSM12079 | 8904 | 44328 |
| 1281.649 | 407 | 1302.27 | 516 | 983.0527 | 367 | 965.9581 | 399 | E2F1 | LNCaP | Prostate | GSM12079 | 8556 | 44329 |
| 1100.015 | 271 | 1032.412 | 279 | 894.3803 | 312 | 990.8278 | 460 | AR | LNCaP | Prostate | GSM12079 | 9373 | 44330 |
| 771.7141 | 118 | 890.9306 | 140 | 608.8516 | 141 | 1039.588 | 230 | AR | LNCaP | Prostate | GSM12079 | 3652 | 44331 |
| 1288.76 | 384 | 1297.754 | 468 | 989.3537 | 343 | 974.7566 | 371 | E2F1 | LNCaP | Prostate | GSM12079 | 7676 | 44332 |
| 1301.316 | 493 | 1342.153 | 652 | 990.1453 | 458 | 963.8106 | 484 | E2F1 | LNCaP | Prostate | GSM12079 | 10727 | 44333 |
| 989.2015 | 342 | 925.3487 | 354 | 794.6049 | 408 | 879.2279 | 580 | AR | LNCaP | Prostate | GSM12079 | 17974 | 44334 |
| 977.1915 | 320 | 931.381 | 344 | 787.7887 | 385 | 871.7066 | 545 | AR | LNCaP | Prostate | GSM12079 | 17118 | 44335 |
| 1311.305 | 539 | 1342.8 | 694 | 999.1443 | 506 | 956.2277 | 520 | E2F1 | LNCaP | Prostate | GSM12079 | 11965 | 44336 |
| 1338.09 | 656 | 1387.969 | 857 | 993.9233 | 600 | 951.7971 | 623 | E2F1 | LNCaP | Prostate | GSM12079 | 15232 | 44337 |
| 924.745 | 130 | 954.0689 | 144 | 532.3756 | 129 | 1040.699 | 224 | AR | LNCaP | Prostate | GSM12079 | 3521 | 44338 |

| | | | | | | | | | | | | | |
|---|---|---|---|---|---|---|---|---|---|---|---|---|---|
| 1273.183 | 470 | 1307.799 | 618 | 974.1041 | 444 | 945.4661 | 467 | E2F1 | LNCaP | Prostate | GSM12079 | 10973 | 44339 |
| 1283.385 | 282 | 1304.695 | 365 | 993.282 | 248 | 981.4595 | 275 | E2F1 | LNCaP | Prostate | GSM12079 | 5291 | 44340 |
| 727.3165 | 104 | 694.8334 | 110 | 432.5133 | 105 | 489.6417 | 119 | PPARD | None | Umbilical V | GSM12146 | 2845 | 44350 |
| 637.4099 | 73 | 551.3002 | 72 | 416.868 | 75 | 491.5149 | 87 | PPARD | None | Umbilical V | GSM12146 | 1352 | 44351 |
| 18.40513 | 20 | 14.63354 | 21 | 15.18038 | 31 | 11.63022 | 31 | ERCC6 | None | None | GSM12150 | 4946 | 44357 |
| 844.7112 | 388 | 852.72 | 483 | 646.2762 | 464 | 650.6085 | 533 | TP53 | Saos-2 | Bone | GSM12414 | 34625 | 44394 |
| 434.7497 | 221 | 457.2824 | 254 | 639.8719 | 406 | 675.3908 | 511 | GATA3 | MCF-7 | Breast | GSM12417 | 29345 | 44395 |
| 621.1483 | 232 | 673.2938 | 269 | 610.6718 | 341 | 686.3326 | 432 | GATA3 | T47D | Mammary | GSM12417 | 23337 | 44397 |
| 724.1219 | 288 | 725.1636 | 334 | 662.1326 | 456 | 704.0971 | 582 | GATA3 | MCF-7 | Breast | GSM12417 | 31483 | 44399 |
| 614.86 | 183 | 527.7135 | 188 | 532.7793 | 248 | 749.1256 | 332 | GATA3 | T47D | Mammary | GSM12417 | 13993 | 44401 |
| 650.321 | 383 | 649.5221 | 467 | 532.4389 | 565 | 524.263 | 631 | TAL1 | None | Cord blood | GSM12911 | 63023 | 44433 |
| 528.3067 | 394 | 590.407 | 478 | 428.1976 | 578 | 493.0731 | 694 | TAL1 | None | Cord blood | GSM12911 | 78751 | 44435 |
| 94.56784 | 51 | 123.5993 | 65 | 263.3873 | 124 | 394.0047 | 161 | ESR1 | MCF-7 | Breast | GSM12955 | 6824 | 44437 |
| 50.21811 | 19 | 74.0095 | 26 | 148.2096 | 49 | 292.0463 | 75 | ESR1 | MCF-7 | Breast | GSM12955 | 1890 | 44438 |
| 87.85479 | 20 | 101.9321 | 24 | 36.31252 | 18 | 160.1874 | 42 | HIF1A | None | Kidney | GSM13129 | 1112 | 44489 |
| 1042.943 | 677 | 1105.39 | 897 | 751.6743 | 715 | 714.0086 | 750 | EMX1 | 293FT | Embryonic | GSM13271 | 43312 | 44562 |
| 1063.635 | 303 | 1109.879 | 410 | 818.2531 | 308 | 828.8455 | 358 | EMX1 | 293FT | Embryonic | GSM13271 | 11875 | 44563 |
| 981.617 | 190 | 1012.178 | 244 | 611.3885 | 208 | 553.4256 | 211 | EMX1 | 293FT | Embryonic | GSM13271 | 8599 | 44564 |
| 995.8728 | 422 | 969.088 | 480 | 894.4155 | 634 | 931.0638 | 786 | AR | VCaP | Prostate | GSM13289 | 22868 | 44574 |
| 1067.669 | 553 | 1060.023 | 658 | 905.0092 | 741 | 948.8456 | 923 | AR | VCaP | Prostate | GSM13289 | 27161 | 44575 |
| 801.9016 | 442 | 778.0584 | 511 | 682.7207 | 645 | 744.9893 | 845 | AR | VCaP | Prostate | GSM13289 | 45871 | 44576 |
| 863.0993 | 551 | 874.539 | 680 | 715.8646 | 754 | 762.9618 | 953 | AR | VCaP | Prostate | GSM13289 | 50843 | 44577 |
| 562.8303 | 240 | 705.7425 | 298 | 523.5031 | 348 | 681.3429 | 487 | AR | VCaP | Prostate | GSM13289 | 27204 | 44578 |
| 547.1085 | 240 | 696.5514 | 300 | 588.2479 | 370 | 694.3939 | 515 | AR | VCaP | Prostate | GSM13289 | 27672 | 44579 |
| 854.376 | 299 | 832.0067 | 348 | 731.2416 | 429 | 797.0312 | 576 | AR | VCaP | Prostate | GSM13289 | 22883 | 44580 |
| 853.1647 | 443 | 869.3793 | 560 | 725.9033 | 633 | 760.231 | 784 | AR | VCaP | Prostate | GSM13289 | 38632 | 44581 |
| 922.2604 | 564 | 893.9952 | 644 | 794.5483 | 807 | 831.6877 | 994 | AR | VCaP | Prostate | GSM13289 | 44342 | 44582 |
| 1000.758 | 686 | 974.7649 | 789 | 829.9496 | 915 | 858.4482 | 1105 | AR | VCaP | Prostate | GSM13289 | 48617 | 44583 |
| 1173.503 | 349 | 1194.386 | 440 | 960.6743 | 380 | 921.9872 | 389 | BRD4 | VCaP | Prostate | GSM13289 | 8827 | 44584 |
| 1154.962 | 320 | 1185.645 | 421 | 931.2922 | 343 | 911.3666 | 368 | BRD4 | VCaP | Prostate | GSM13289 | 8528 | 44585 |
| 372.016 | 51 | 569.6649 | 72 | 371.9611 | 68 | 357.1454 | 70 | BRD4 | VCaP | Prostate | GSM13289 | 1182 | 44587 |
| 1138.011 | 230 | 1166.783 | 304 | 970.4955 | 282 | 927.1376 | 281 | BRD4 | VCaP | Prostate | GSM13289 | 6051 | 44588 |
| 980.6887 | 295 | 1044.76 | 424 | 771.6585 | 329 | 760.618 | 365 | BRD2 | VCaP | Prostate | GSM13289 | 14458 | 44594 |
| 568.1741 | 151 | 643.5524 | 178 | 338.2036 | 168 | 362.8978 | 186 | BRD2 | VCaP | Prostate | GSM13289 | 9597 | 44595 |
| 1112.681 | 334 | 1138.544 | 430 | 877.8083 | 347 | 840.7093 | 356 | BRD2 | VCaP | Prostate | GSM13289 | 10687 | 44596 |
| 1002.298 | 360 | 1047.258 | 485 | 790.2909 | 400 | 755.864 | 415 | BRD3 | VCaP | Prostate | GSM13289 | 17022 | 44597 |
| 423.3335 | 140 | 688.618 | 199 | 392.3336 | 196 | 313.397 | 190 | BRD3 | VCaP | Prostate | GSM13289 | 11158 | 44598 |
| 993.6761 | 176 | 1022.158 | 211 | 616.7981 | 188 | 586.7861 | 195 | BRD3 | VCaP | Prostate | GSM13289 | 6573 | 44599 |
| 1034.403 | 309 | 1085.358 | 428 | 824.7154 | 347 | 788.4129 | 354 | BRD4 | VCaP | Prostate | GSM13289 | 12743 | 44600 |
| 775.8383 | 181 | 964.0322 | 254 | 512.4371 | 208 | 559.8195 | 232 | BRD4 | VCaP | Prostate | GSM13289 | 9923 | 44601 |
| 1035.523 | 297 | 1055.861 | 386 | 820.4209 | 326 | 783.7067 | 334 | BRD4 | VCaP | Prostate | GSM13289 | 12204 | 44602 |
| 1107.381 | 547 | 1102.566 | 661 | 926.8438 | 698 | 909.4365 | 762 | ERG | VCaP | Prostate | GSM13289 | 22680 | 44603 |
| 1094.971 | 611 | 1101.588 | 750 | 911.8417 | 781 | 880.995 | 833 | ERG | VCaP | Prostate | GSM13289 | 28024 | 44604 |
| 1071.164 | 479 | 1074.736 | 581 | 895.1774 | 605 | 883.804 | 673 | ERG | VCaP | Prostate | GSM13289 | 20282 | 44605 |
| 1058.047 | 443 | 1073.858 | 554 | 890.6313 | 565 | 863.9123 | 605 | ERG | VCaP | Prostate | GSM13289 | 18861 | 44606 |
| 255.2614 | 56 | 412.8876 | 81 | 160.4576 | 62 | 145.7211 | 63 | BRCA1 | MCF-10A | Breast | GSM13405 | 2826 | 44621 |
| 26.09176 | 82 | 57.98176 | 122 | 74.86609 | 192 | 60.7538 | 198 | TP63 | None | Foreskin | GSM13666 | 32749 | 44759 |
| 7.352336 | 13 | 15.28377 | 20 | 8.791739 | 24 | 27.27484 | 39 | TP63 | None | Foreskin | GSM13666 | 4448 | 44760 |
| 15.48437 | 48 | 30.19854 | 69 | 29.46918 | 101 | 42.19869 | 123 | TP63 | None | Foreskin | GSM13666 | 19467 | 44761 |
| 37.37286 | 20 | 35.48017 | 22 | 11.1668 | 19 | 59.54541 | 41 | TP53 | None | Foreskin | GSM13666 | 2807 | 44762 |
| 14.57859 | 20 | 19.84725 | 26 | 5.358205 | 25 | 33.01921 | 49 | TP53 | None | Foreskin | GSM13666 | 5599 | 44763 |
| 18.16878 | 31 | 35.77493 | 46 | 10.65616 | 45 | 27.87267 | 66 | TP53 | None | Foreskin | GSM13666 | 9476 | 44764 |
| 7.542672 | 38 | 25.99595 | 64 | 22.4729 | 91 | 23.17416 | 101 | TP63 | None | Foreskin | GSM13666 | 18742 | 44765 |
| 6.206429 | 7 | 20.92294 | 14 | 6.246361 | 12 | 36.75254 | 27 | TP63 | None | Foreskin | GSM13666 | 2006 | 44766 |
| 40.70476 | 79 | 66.62528 | 108 | 52.71822 | 145 | 68.74604 | 172 | TP63 | None | Foreskin | GSM13666 | 25428 | 44767 |
| 14.78358 | 14 | 5.72462 | 11 | 1.9566 | 12 | 42.2273 | 38 | TP53 | None | Foreskin | GSM13666 | 3195 | 44768 |
| 38.53963 | 41 | 76.22385 | 62 | 26.68032 | 59 | 53.88506 | 82 | TP53 | None | Foreskin | GSM13666 | 9366 | 44769 |
| 16.88849 | 24 | 28.77274 | 34 | 7.910521 | 32 | 24.42795 | 50 | TP53 | None | Foreskin | GSM13666 | 6801 | 44770 |
| 226.4286 | 51 | 367.8294 | 74 | 147.7404 | 58 | 215.7639 | 75 | POLR3A | None | Blood | GSM50171 | 2741 | 4487 |
| 30.56705 | 26 | 31.42036 | 30 | 47.73158 | 50 | 104.3496 | 75 | PPARG | SGBS | Adipose | GSM63463 | 5289 | 4492 |
| 35.46555 | 32 | 31.92478 | 35 | 68.48545 | 67 | 115.7906 | 90 | PPARG | SGBS | Adipose | GSM63462 | 6753 | 4493 |
| 122.502 | 99 | 57.25784 | 84 | 112.007 | 152 | 197.9229 | 204 | PPARG | None | Adipose | GSM53449 | 18650 | 4494 |
| 283.445 | 185 | 244.7043 | 197 | 213.6198 | 259 | 301.3269 | 319 | PPARG | None | Adipose | GSM53449 | 30314 | 4495 |
| 992.5171 | 437 | 1006.787 | 545 | 709.7126 | 420 | 679.9525 | 440 | RUNX1 | None | Cord blood | GSM60795 | 24833 | 4514 |
| 453.9907 | 316 | 662.6291 | 451 | 350.958 | 447 | 387.3792 | 504 | CTCF | A549 | Lung | GSM10035 | 54103 | 45144 |
| 538.5706 | 363 | 660.4349 | 497 | 337.9651 | 478 | 398.4445 | 551 | CTCF | A549 | Lung | GSM10035 | 61530 | 45146 |
| 190.825 | 67 | 235.2716 | 83 | 129.6812 | 83 | 125.8805 | 88 | RXRG | None | Bone Marr | GSM46820 | 6103 | 4515 |
| 747.0867 | 380 | 796.2321 | 520 | 562.311 | 462 | 544.4277 | 502 | CTCF | None | Blood | GSM10034 | 41478 | 45178 |
| 704.2557 | 436 | 775.9899 | 618 | 554.0413 | 588 | 509.2529 | 599 | CTCF | DND-41 | Blood | GSM10034 | 61204 | 45189 |
| 259.1331 | 41 | 355.7874 | 54 | 75.62915 | 28 | 108.0226 | 36 | EZH2 | DND-41 | Blood | GSM10035 | 1141 | 45192 |
| 586.4998 | 315 | 714.5621 | 435 | 452.8336 | 436 | 397.4911 | 452 | CTCF | GM12878 | Blood | GSM73375 | 45660 | 45218 |
| 1083.975 | 547 | 1114.27 | 699 | 798.9458 | 547 | 756.6233 | 557 | HDAC2 | H1 | Embryo | GSM10034 | 24975 | 45222 |

| | | | | | | | | | | | | | |
|---|---|---|---|---|---|---|---|---|---|---|---|---|---|
| 1204.735 | 748 | 1282.074 | 992 | 840.6965 | 683 | 803.4308 | 717 | KDM4A | H1 | Embryo | GSM10034 | 28179 | 45228 |
| 209.7004 | 62 | 363.0741 | 92 | 162.194 | 78 | 168.3165 | 85 | EZH2 | H1 | Embryo | GSM10035 | 4046 | 45229 |
| 546.439 | 409 | 640.9832 | 556 | 406.973 | 586 | 446.604 | 660 | CTCF | H1 | Embryo | GSM73367 | 83652 | 45230 |
| 220.7742 | 93 | 342.9602 | 129 | 110.5818 | 104 | 163.9531 | 132 | CHD7 | H1 | Embryo | GSM10034 | 10102 | 45231 |
| 1102.182 | 357 | 1101.252 | 432 | 812.6425 | 324 | 795.7645 | 352 | RBBP5 | H1 | Embryo | GSM10035 | 12537 | 45232 |
| 791.0347 | 143 | 1057.817 | 197 | 378.7232 | 134 | 379.8379 | 143 | SUZ12 | H1 | Embryo | GSM10035 | 5506 | 45233 |
| 1141.296 | 495 | 1196.138 | 667 | 865.9607 | 493 | 832.6675 | 516 | SAP30 | H1 | Embryo | GSM10035 | 17660 | 45234 |
| 1068.049 | 227 | 1084.356 | 292 | 757.1441 | 226 | 686.9297 | 229 | CHD1 | H1 | Embryo | GSM10034 | 7486 | 45237 |
| 1307.608 | 716 | 1385.597 | 955 | 941.8912 | 638 | 915.1109 | 686 | PHF8 | H1 | Embryo | GSM10035 | 19149 | 45239 |
| 397.8128 | 96 | 618.6638 | 135 | 216.2674 | 101 | 216.2023 | 108 | EP300 | H1 | Embryo | GSM10035 | 5591 | 45242 |
| 1217.858 | 146 | 1218.265 | 178 | 671.8667 | 137 | 682.3551 | 146 | KDM5A | H1 | Embryo | GSM10034 | 2869 | 45244 |
| 434.5244 | 309 | 645.0287 | 428 | 281.9637 | 415 | 319.5609 | 472 | CTCF | HeLa-S3 | Cervix | GSM73378 | 55030 | 45279 |
| 376.7591 | 226 | 615.7012 | 315 | 269.8005 | 309 | 325.0335 | 359 | CTCF | HepG2 | Liver | GSM73364 | 35832 | 45303 |
| 30.70536 | 16 | 143.7688 | 40 | 29.73479 | 24 | 68.95351 | 38 | EZH2 | None | Mammary | GSM10035 | 2007 | 45312 |
| 452.2277 | 275 | 685.5402 | 391 | 291.6456 | 363 | 347.7317 | 420 | CTCF | None | Mammary | GSM73372 | 44468 | 45323 |
| 465.5852 | 274 | 667.6065 | 362 | 341.5494 | 376 | 391.0976 | 429 | CTCF | None | Skeletal Mu | GSM73376 | 42424 | 45334 |
| 574.699 | 338 | 681.5282 | 455 | 369.9477 | 444 | 414.6951 | 503 | CTCF | None | Skeletal Mu | GSM73378 | 53489 | 45347 |
| 748.0261 | 317 | 792.6649 | 437 | 505.0222 | 394 | 522.0882 | 432 | CTCF | None | Umbilical V | GSM73371 | 35274 | 45359 |
| 223.4233 | 72 | 440.3737 | 114 | 172.6162 | 91 | 137.4618 | 88 | EZH2 | None | Umbilical V | GSM10035 | 4986 | 45370 |
| 628.976 | 83 | 832.9604 | 107 | 287.1533 | 71 | 336.0705 | 82 | HDAC6 | K562 | Bone Marr | GSM10035 | 1936 | 45379 |
| 118.2287 | 56 | 195.6672 | 80 | 48.85712 | 58 | 81.39692 | 77 | REST | K562 | Bone Marr | GSM10035 | 6125 | 45385 |
| 895.3049 | 141 | 1108.89 | 185 | 596.5006 | 154 | 635.9093 | 169 | SETDB1 | K562 | Bone Marr | GSM10034 | 4516 | 45386 |
| 1201.708 | 579 | 1255.27 | 767 | 885.61 | 537 | 857.3518 | 569 | KDM5B | K562 | Bone Marr | GSM10035 | 17261 | 45390 |
| 143.0667 | 34 | 182.6658 | 43 | 37.64567 | 25 | 179.3852 | 58 | RNF2 | K562 | Bone Marr | GSM10035 | 1934 | 45391 |
| 386.718 | 59 | 404.8139 | 66 | 184.1526 | 53 | 176.6162 | 55 | CHD7 | K562 | Bone Marr | GSM10034 | 1752 | 45392 |
| 536.3722 | 344 | 683.0572 | 488 | 399.3127 | 479 | 475.0883 | 554 | CTCF | K562 | Bone Marr | GSM73371 | 57670 | 45395 |
| 185.6831 | 112 | 238.4168 | 141 | 183.9777 | 172 | 199.4433 | 192 | KDM1A | K562 | Bone Marr | GSM10035 | 16503 | 45400 |
| 81.31087 | 19 | 88.59226 | 22 | 36.89077 | 18 | 128.8959 | 37 | NCOR1 | K562 | Bone Marr | GSM10035 | 1093 | 45405 |
| 114.436 | 38 | 118.6209 | 43 | 47.36363 | 37 | 37.18541 | 36 | EP300 | K562 | Bone Marr | GSM10035 | 3120 | 45408 |
| 1264.888 | 788 | 1352.949 | 1044 | 898.7058 | 731 | 855.6183 | 757 | PHF8 | K562 | Bone Marr | GSM10034 | 25247 | 45411 |
| 1035.563 | 192 | 1075.164 | 264 | 527.3283 | 182 | 646.9337 | 215 | HDAC2 | K562 | Bone Marr | GSM10034 | 7402 | 45414 |
| 483.7087 | 64 | 734.3611 | 90 | 222.4339 | 55 | 382.3684 | 79 | KAT2B | K562 | Bone Marr | GSM10035 | 1525 | 45418 |
| 1235.519 | 476 | 1277.298 | 619 | 940.6776 | 440 | 902.025 | 457 | SAP30 | K562 | Bone Marr | GSM10034 | 11580 | 45419 |
| 644.0062 | 170 | 788.1689 | 209 | 263.265 | 161 | 329.2167 | 191 | SMARCA4 | None | Bone Marr | GSM65155 | 11465 | 4542 |
| 1101.822 | 504 | 1147.95 | 666 | 820.5873 | 484 | 810.4337 | 537 | RBBP5 | K562 | Bone Marr | GSM10034 | 18429 | 45421 |
| 29.27328 | 27 | 34.41997 | 33 | 12.92562 | 33 | 24.84504 | 45 | CREBBP | K562 | Bone Marr | GSM10035 | 5809 | 45422 |
| 124.2138 | 24 | 191.2404 | 34 | 57.23063 | 22 | 50.44841 | 22 | EZH2 | K562 | Bone Marr | GSM10035 | 1024 | 45424 |
| 1146.583 | 268 | 1202.841 | 377 | 901.6412 | 263 | 860.1184 | 267 | CHD1 | K562 | Bone Marr | GSM10035 | 7066 | 45426 |
| 1150.346 | 471 | 1172.004 | 592 | 869.5234 | 451 | 835.2141 | 470 | HDAC1 | K562 | Bone Marr | GSM10034 | 14962 | 45428 |
| 80.21107 | 26 | 139.6952 | 39 | 38.40976 | 27 | 54.20425 | 34 | CBX3 | K562 | Bone Marr | GSM10035 | 2182 | 45429 |
| 1043.176 | 372 | 1039.412 | 451 | 772.4617 | 361 | 718.5411 | 354 | SMARCA4 | None | Blood | GSM65155 | 16879 | 4543 |
| 749.2536 | 338 | 801.4135 | 470 | 569.5152 | 425 | 568.767 | 466 | CTCF | None | Blood | GSM10035 | 35366 | 45444 |
| 433.8205 | 262 | 676.8062 | 365 | 272.6895 | 341 | 319.0719 | 392 | CTCF | None | Brain | GSM73376 | 41087 | 45455 |
| 68.67383 | 26 | 158.7441 | 45 | 69.49515 | 38 | 81.58038 | 44 | EZH2 | None | Brain | GSM10035 | 2152 | 45459 |
| 83.08577 | 30 | 159.591 | 47 | 81.16713 | 43 | 126.0161 | 57 | EZH2 | NHDF-Ad | Skin | GSM10035 | 2392 | 45466 |
| 625.2468 | 351 | 700.8047 | 479 | 427.2397 | 469 | 455.3624 | 522 | CTCF | NHDF-Ad | Skin | GSM73374 | 54233 | 45472 |
| 54.44655 | 28 | 113.493 | 45 | 43.71386 | 38 | 59.9127 | 47 | EZH2 | NHEK | Skin | GSM10034 | 3108 | 45490 |
| 617.1312 | 320 | 701.4229 | 418 | 373.5233 | 404 | 363.2924 | 435 | CTCF | NHEK | Skin | GSM73363 | 45263 | 45494 |
| 84.75842 | 29 | 146.5673 | 43 | 33.12698 | 27 | 78.08291 | 43 | EZH2 | NHLF | Lung | GSM10035 | 2144 | 45498 |
| 33.14652 | 45 | 35.2234 | 53 | 39.92037 | 80 | 29.21867 | 79 | SPI1 | None | Cord blood | GSM63831 | 12268 | 4550 |
| 715.8675 | 365 | 767.6902 | 503 | 532.3021 | 468 | 536.7549 | 517 | CTCF | NHLF | Lung | GSM73369 | 46932 | 45504 |
| 38.37103 | 51 | 63.45748 | 71 | 56.77795 | 97 | 62.25503 | 109 | SPI1 | None | Cord blood | GSM63831 | 13676 | 4551 |
| 269.3173 | 71 | 321.5845 | 86 | 159.2801 | 77 | 204.6514 | 93 | EP300 | None | Bone | GSM10035 | 3937 | 45517 |
| 627.7615 | 375 | 699.0655 | 523 | 428.1318 | 503 | 492.1099 | 575 | CTCF | None | Bone | GSM73378 | 59962 | 45526 |
| 455.9778 | 238 | 712.2971 | 333 | 318.7675 | 317 | 374.2667 | 365 | CTCF | A549 | Lung | GSM10226 | 33603 | 45550 |
| 561.1999 | 346 | 688.5178 | 483 | 389.71 | 471 | 457.2856 | 542 | CTCF | A549 | Lung | GSM10226 | 58204 | 45551 |
| 1143.092 | 432 | 1149.003 | 528 | 858.1234 | 405 | 806.0657 | 400 | MAX | A549 | Lung | GSM93529 | 13917 | 45553 |
| 495.4227 | 225 | 608.3437 | 277 | 260.3016 | 263 | 300.7844 | 301 | RAD21 | A549 | Lung | GSM93530 | 27642 | 45554 |
| 696.5316 | 358 | 737.3756 | 481 | 433.6386 | 444 | 436.9854 | 484 | CTCF | A549 | Lung | GSM10036 | 49204 | 45556 |
| 735.9823 | 121 | 1020.742 | 160 | 405.854 | 120 | 517.942 | 145 | BHLHE40 | A549 | Lung | GSM93544 | 4094 | 45557 |
| 733.5333 | 550 | 791.706 | 742 | 551.543 | 710 | 528.5813 | 771 | CTCF | A549 | Lung | GSM82228 | 81289 | 45558 |
| 307.6294 | 348 | 375.0727 | 427 | 182.0625 | 489 | 246.6467 | 584 | CEBPB | A549 | Lung | GSM93563 | 92264 | 45559 |
| 1116.932 | 343 | 1138.26 | 438 | 849.2052 | 326 | 824.7948 | 347 | MYC | A549 | Lung | GSM10036 | 11008 | 45560 |
| 734.568 | 380 | 779.1039 | 514 | 546.0205 | 472 | 552.0917 | 536 | CTCF | AG04449 | Skin | GSM74967 | 47524 | 45561 |
| 836.2297 | 517 | 890.2721 | 699 | 619.5967 | 613 | 598.6903 | 663 | CTCF | AG04449 | Skin | GSM74969 | 53908 | 45562 |
| 678.0689 | 335 | 746.5184 | 470 | 452.5813 | 437 | 474.8902 | 483 | CTCF | AG04450 | Fetal Lung | GSM74976 | 46495 | 45563 |
| 700.8227 | 334 | 737.1883 | 449 | 443.887 | 423 | 453.7109 | 463 | CTCF | AG04450 | Fetal Lung | GSM10226 | 43901 | 45564 |
| 677.346 | 334 | 743.1606 | 465 | 466.0506 | 440 | 491.0431 | 487 | CTCF | AG09309 | Skin | GSM74975 | 45961 | 45565 |
| 503.2483 | 213 | 736.9896 | 285 | 294.7879 | 257 | 357.8681 | 300 | CTCF | AG09309 | Skin | GSM74968 | 24592 | 45566 |
| 748.5256 | 419 | 798.0147 | 567 | 560.1123 | 514 | 558.7604 | 583 | CTCF | AG09319 | Gingiva | GSM74972 | 50860 | 45567 |
| 716.8231 | 363 | 762.7693 | 498 | 513.4863 | 462 | 537.4559 | 516 | CTCF | AG09319 | Gingiva | GSM74972 | 46997 | 45568 |
| 707.9562 | 336 | 770.5091 | 480 | 503.4563 | 440 | 519.6832 | 483 | CTCF | AG10803 | Skin | GSM74971 | 43526 | 45569 |

| | | | | | | | | | | | | | |
|---|---|---|---|---|---|---|---|---|---|---|---|---|---|
| 660.3669 | 343 | 722.2 | 471 | 418.9554 | 442 | 502.8625 | 513 | CTCF | AG10803 | Skin | GSM74975 | 49617 | 45570 |
| 654.0666 | 262 | 790.2616 | 364 | 432.0177 | 330 | 461.187 | 366 | CTCF | None | Umbilical \ | GSM74973 | 29208 | 45571 |
| 733.1176 | 348 | 790.8707 | 488 | 534.8679 | 442 | 519.8746 | 474 | CTCF | None | Umbilical \ | GSM74966 | 42614 | 45572 |
| 569.9263 | 337 | 700.884 | 475 | 420.2008 | 466 | 471.4804 | 528 | CTCF | BE2-C | Brain | GSM10226 | 54701 | 45573 |
| 482.0102 | 302 | 686.1104 | 431 | 362.0754 | 421 | 396.5486 | 473 | CTCF | BE2-C | Brain | GSM10226 | 49493 | 45574 |
| 726.8675 | 351 | 794.0449 | 500 | 519.9251 | 443 | 546.6427 | 494 | CTCF | BJ | Skin | GSM74975 | 43327 | 45575 |
| 759.2157 | 443 | 804.14 | 596 | 563.9268 | 538 | 550.9033 | 593 | CTCF | BJ | Skin | GSM74967 | 53271 | 45576 |
| 702.1796 | 333 | 759.7137 | 468 | 459.6126 | 428 | 505.3217 | 481 | CTCF | Caco-2 | Colon | GSM74968 | 44964 | 45577 |
| 697.6119 | 361 | 768.3402 | 522 | 502.3555 | 474 | 537.3046 | 544 | CTCF | Caco-2 | Colon | GSM74974 | 50898 | 45578 |
| 716.1394 | 504 | 758.8716 | 669 | 531.9524 | 641 | 517.0911 | 706 | CTCF | None | Brain | GSM82230 | 74611 | 45579 |
| 696.1297 | 342 | 744.2779 | 473 | 491.0232 | 452 | 534.618 | 510 | CTCF | GM06990 | Blood | GSM74970 | 46936 | 45581 |
| 652.1559 | 342 | 736.0975 | 488 | 460.1403 | 459 | 486.5869 | 509 | CTCF | GM06990 | Blood | GSM74970 | 50256 | 45582 |
| 11.06033 | 8 | 10.7509 | 9 | 1.461176 | 6 | 22.86469 | 19 | ZNF274 | GM08714 | Blood | GSM93532 | 1486 | 45583 |
| 753.2771 | 364 | 811.7842 | 509 | 588.2602 | 471 | 566.7206 | 508 | CTCF | GM10248 | Blood | GSM10068 | 42454 | 45584 |
| 758.4961 | 398 | 814.959 | 550 | 570.7826 | 489 | 560.2587 | 544 | CTCF | GM10266 | Blood | GSM10068 | 46962 | 45585 |
| 935.2044 | 426 | 942.7239 | 528 | 672.2608 | 428 | 640.0723 | 446 | RELA | GM10847 | Blood | GSM93552 | 27717 | 45587 |
| 358.1298 | 171 | 539.9818 | 232 | 237.3674 | 220 | 232.3449 | 236 | CTCF | GM12801 | Blood | GSM74971 | 21833 | 45588 |
| 458.9577 | 276 | 687.8556 | 392 | 327.5403 | 376 | 365.3281 | 425 | CTCF | GM12864 | Blood | GSM10226 | 43519 | 45589 |
| 695.8612 | 317 | 778.5324 | 459 | 491.8308 | 418 | 483.0053 | 450 | CTCF | GM12864 | Blood | GSM74976 | 40739 | 45590 |
| 692.7576 | 338 | 765.2914 | 495 | 490.2608 | 450 | 535.157 | 506 | CTCF | GM12864 | Blood | GSM74967 | 46524 | 45591 |
| 726.3253 | 352 | 785.4834 | 493 | 519.4633 | 445 | 549.2387 | 501 | CTCF | GM12865 | Blood | GSM74972 | 43912 | 45592 |
| 436.8198 | 242 | 709.9297 | 349 | 342.0033 | 338 | 395.4631 | 387 | CTCF | GM12865 | Blood | GSM10226 | 35530 | 45593 |
| 766.1316 | 352 | 803.4333 | 471 | 560.5991 | 427 | 567.7962 | 472 | CTCF | GM12865 | Blood | GSM74974 | 38605 | 45594 |
| 539.2857 | 313 | 693.5887 | 435 | 404.3328 | 433 | 414.2796 | 475 | CTCF | GM12872 | Blood | GSM10226 | 48803 | 45595 |
| 553.2393 | 281 | 730.4285 | 397 | 414.0693 | 383 | 429.361 | 421 | CTCF | GM12872 | Blood | GSM74969 | 39162 | 45596 |
| 705.8746 | 333 | 763.851 | 471 | 496.6238 | 437 | 544.9862 | 493 | CTCF | GM12872 | Blood | GSM74969 | 43765 | 45597 |
| 595.4857 | 344 | 716.4834 | 497 | 422.2162 | 467 | 480.9409 | 532 | CTCF | GM12873 | Blood | GSM10226 | 54190 | 45598 |
| 666.6625 | 318 | 768.1488 | 465 | 448.0061 | 414 | 514.8378 | 473 | CTCF | GM12873 | Blood | GSM74968 | 42729 | 45599 |
| 700.8769 | 351 | 764.7561 | 500 | 517.9549 | 466 | 535.2411 | 518 | CTCF | GM12873 | Blood | GSM74973 | 47696 | 45600 |
| 609.3475 | 318 | 732.7678 | 451 | 449.4414 | 433 | 472.8524 | 479 | CTCF | GM12874 | Blood | GSM74974 | 46037 | 45601 |
| 773.7918 | 378 | 833.6798 | 532 | 592.1574 | 471 | 571.9235 | 509 | CTCF | GM12874 | Blood | GSM74975 | 41599 | 45602 |
| 735.4629 | 280 | 796.9834 | 381 | 467.4119 | 345 | 516.6269 | 388 | CTCF | GM12875 | Blood | GSM74967 | 29815 | 45603 |
| 668.6698 | 339 | 763.4719 | 505 | 520.8401 | 470 | 534.2006 | 521 | CTCF | GM12875 | Blood | GSM74976 | 48434 | 45604 |
| 360.7543 | 96 | 379.7365 | 109 | 181.6092 | 99 | 170.0609 | 103 | YY1 | GM12878 | Blood | GSM93548 | 6272 | 45605 |
| 1274.264 | 246 | 1297.122 | 315 | 980.1387 | 211 | 975.4557 | 236 | FOS | GM12878 | Blood | GSM93540 | 4559 | 45606 |
| 1036.785 | 128 | 1211.736 | 160 | 385.6156 | 97 | 436.83 | 110 | SREBF2 | GM12878 | Blood | GSM10036 | 2699 | 45607 |
| 608.5593 | 85 | 562.2027 | 88 | 266.0936 | 72 | 251.1809 | 74 | NFE2 | GM12878 | Blood | GSM93565 | 2222 | 45608 |
| 1374.421 | 204 | 1416.593 | 273 | 1023.388 | 167 | 1025.074 | 176 | IRF3 | GM12878 | Blood | GSM93565 | 2627 | 45609 |
| 734.1494 | 191 | 873.7738 | 232 | 357.0955 | 195 | 311.4011 | 197 | JUND | GM12878 | Blood | GSM10036 | 12568 | 45610 |
| 991.2596 | 313 | 970.966 | 367 | 729.6388 | 306 | 582.7037 | 294 | USF2 | GM12878 | Blood | GSM93555 | 15789 | 45611 |
| 440.5228 | 149 | 514.5149 | 179 | 193.4153 | 152 | 199.409 | 166 | STAT3 | GM12878 | Blood | GSM93555 | 12919 | 45612 |
| 434.9803 | 261 | 624.4008 | 343 | 289.7214 | 347 | 311.2238 | 387 | RAD21 | GM12878 | Blood | GSM93533 | 41106 | 45614 |
| 169.2935 | 80 | 269.4363 | 112 | 85.73196 | 91 | 96.10363 | 103 | CUX1 | GM12878 | Blood | GSM10036 | 9347 | 45615 |
| 1219.889 | 278 | 1231.548 | 350 | 922.2117 | 238 | 890.2149 | 248 | E2F4 | GM12878 | Blood | GSM93533 | 6193 | 45617 |
| 1042.232 | 491 | 1082.866 | 648 | 768.4353 | 488 | 728.3282 | 505 | TBP | GM12878 | Blood | GSM93527 | 23635 | 45618 |
| 1147.877 | 208 | 1162.99 | 262 | 699.2713 | 184 | 598.8356 | 180 | SREBF1 | GM12878 | Blood | GSM10036 | 5487 | 45620 |
| 213.7123 | 95 | 294.1561 | 124 | 91.91059 | 101 | 95.55176 | 111 | MAFK | GM12878 | Blood | GSM10036 | 10877 | 45621 |
| 643.7431 | 322 | 757.9215 | 474 | 500.1561 | 445 | 537.7484 | 496 | CTCF | GM12878 | Blood | GSM74970 | 45078 | 45622 |
| 486.0394 | 250 | 726.0927 | 358 | 344.3007 | 334 | 416.5009 | 389 | CTCF | GM12878 | Blood | GSM74970 | 34887 | 45623 |
| 994.6137 | 127 | 1221.645 | 172 | 418.5949 | 104 | 629.445 | 138 | NRF1 | GM12878 | Blood | GSM93530 | 2892 | 45624 |
| 947.7686 | 342 | 989.2669 | 463 | 689.8299 | 341 | 692.1215 | 389 | NFYB | GM12878 | Blood | GSM93550 | 20109 | 45625 |
| 1315.775 | 219 | 1350.621 | 288 | 844.712 | 167 | 954.6766 | 189 | NFYA | GM12878 | Blood | GSM93550 | 3437 | 45626 |
| 1038.966 | 213 | 1067.69 | 281 | 527.8561 | 191 | 561.1791 | 210 | ESRRA | GM12878 | Blood | GSM10036 | 8192 | 45627 |
| 1220.281 | 298 | 1258.151 | 395 | 949.8598 | 278 | 937.0886 | 304 | MYC | GM12878 | Blood | GSM82229 | 6755 | 45628 |
| 968.0491 | 670 | 1014.448 | 870 | 688.1495 | 720 | 652.45 | 758 | ZNF143 | GM12878 | Blood | GSM93561 | 53337 | 45629 |
| 397.9537 | 55 | 562.383 | 74 | 164.515 | 45 | 192.5867 | 52 | STAT1 | None | Blood | GSM67139 | 1437 | 4563 |
| 787.9998 | 192 | 899.7372 | 232 | 372.9823 | 191 | 315.5684 | 190 | WRNIP1 | GM12878 | Blood | GSM93549 | 11552 | 45630 |
| 815.1265 | 599 | 862.5893 | 789 | 581.1095 | 696 | 553.7824 | 745 | CTCF | GM12878 | Blood | GSM93561 | 72056 | 45631 |
| 953.2458 | 612 | 973.78 | 763 | 676.6368 | 638 | 640.6242 | 672 | BHLHE40 | GM12878 | Blood | GSM93543 | 46265 | 45632 |
| 1095.656 | 548 | 1111.804 | 683 | 793.1366 | 516 | 746.7684 | 521 | MXI1 | GM12878 | Blood | GSM93543 | 23072 | 45633 |
| 330.6516 | 162 | 472.1495 | 214 | 153.5668 | 181 | 116.7623 | 178 | ZNF384 | GM12878 | Blood | GSM10036 | 20169 | 45634 |
| 1171.966 | 321 | 1177.713 | 397 | 884.0535 | 287 | 857.2684 | 304 | ELK1 | GM12878 | Blood | GSM93534 | 8620 | 45636 |
| 1064.901 | 544 | 1078.027 | 678 | 749.8175 | 504 | 721.7991 | 535 | CHD2 | GM12878 | Blood | GSM93537 | 25970 | 45638 |
| 1204.088 | 426 | 1219.748 | 531 | 883.6285 | 360 | 870.2469 | 391 | SIN3A | GM12878 | Blood | GSM93533 | 10672 | 45639 |
| 508.3029 | 263 | 699.1242 | 349 | 351.4275 | 347 | 355.2323 | 378 | SMC3 | GM12878 | Blood | GSM93537 | 36049 | 45640 |
| 787.3015 | 311 | 836.1418 | 431 | 459.3275 | 347 | 390.2958 | 352 | RELA | GM12878 | Blood | GSM93547 | 29662 | 45641 |
| 1107.501 | 317 | 1155.481 | 430 | 855.9976 | 312 | 833.6949 | 336 | STAT1 | GM12878 | Blood | GSM93561 | 10511 | 45643 |
| 620.5424 | 247 | 751.8204 | 311 | 274.4926 | 263 | 282.4104 | 288 | EP300 | GM12878 | Blood | GSM93529 | 25958 | 45644 |
| 702.2733 | 428 | 763.4804 | 595 | 535.9984 | 559 | 532.7425 | 633 | CTCF | GM12878 | Blood | GSM82231 | 62441 | 45645 |
| 1111.467 | 174 | 1141.665 | 233 | 594.98 | 163 | 568.6268 | 169 | CHD1 | GM12878 | Blood | GSM93530 | 5023 | 45646 |
| 78.20008 | 73 | 98.32091 | 91 | 59.39814 | 105 | 48.37995 | 107 | IKZF1 | GM12878 | Blood | GSM93544 | 14009 | 45647 |
| 55.42956 | 17 | 225.7687 | 42 | 39.26006 | 21 | 56.8784 | 27 | ZZZ3 | GM12878 | Blood | GSM93548 | 1356 | 45648 |

| | | | | | | | | | | | | | |
|---|---|---|---|---|---|---|---|---|---|---|---|---|---|
| 1158.616 | 343 | 1162.345 | 422 | 874.6185 | 316 | 829.1922 | 317 | RFX5 | GM12878 | Blood | GSM93555 | 10057 | 45649 |
| 1153.903 | 548 | 1140.142 | 644 | 825.2687 | 488 | 788.6439 | 503 | MAX | GM12878 | Blood | GSM93551 | 19414 | 45650 |
| 974.9659 | 369 | 966.1031 | 442 | 684.7644 | 337 | 671.6361 | 372 | EP300 | GM12878 | Blood | GSM93555 | 20265 | 45651 |
| 930.9092 | 294 | 958.4721 | 384 | 697.9425 | 314 | 645.4768 | 325 | RCOR1 | GM12878 | Blood | GSM93558 | 17323 | 45652 |
| 266.0256 | 82 | 363.0869 | 107 | 135.3818 | 87 | 182.8891 | 107 | NR2C2 | GM12878 | Blood | GSM93548 | 6280 | 45653 |
| 1092.653 | 126 | 1266.21 | 166 | 679.3384 | 126 | 645.1241 | 129 | BRCA1 | GM12878 | Blood | GSM93537 | 2351 | 45654 |
| 6.076997 | 21 | 19.40109 | 36 | 7.229053 | 40 | 24.20177 | 62 | JUND | GM12878 | Blood | GSM93554 | 9385 | 45655 |
| 741.1557 | 461 | 760.6619 | 583 | 502.9209 | 522 | 514.5667 | 590 | EBF1 | GM12878 | Blood | GSM93537 | 58636 | 45656 |
| 668.2456 | 294 | 710.0427 | 354 | 352.4338 | 343 | 300.4566 | 351 | EP300 | GM12878 | Blood | GSM93556 | 35154 | 45658 |
| 943.0445 | 384 | 938.729 | 463 | 667.7731 | 367 | 609.4725 | 371 | TBL1XR1 | GM12878 | Blood | GSM93565 | 23445 | 45660 |
| 1107.096 | 597 | 1142.858 | 762 | 808.4141 | 581 | 764.5775 | 596 | MAZ | GM12878 | Blood | GSM93528 | 25810 | 45661 |
| 811.8863 | 540 | 853.2095 | 708 | 572.0728 | 601 | 552.3087 | 657 | RELA | GM12891 | Blood | GSM93552 | 59301 | 45662 |
| 975.6138 | 444 | 1009.33 | 583 | 703.4744 | 439 | 694.0897 | 487 | RELA | GM12892 | Blood | GSM93528 | 26228 | 45664 |
| 761.7415 | 316 | 803.1438 | 430 | 541.4688 | 395 | 503.4653 | 415 | CTCF | GM13976 | Blood | GSM10068 | 34555 | 45666 |
| 750.7153 | 410 | 789.1165 | 545 | 567.8111 | 514 | 540.6173 | 546 | CTCF | GM13977 | Blood | GSM10068 | 50283 | 45667 |
| 1010.769 | 464 | 1031.201 | 584 | 732.0838 | 450 | 686.5394 | 454 | RELA | GM15510 | Blood | GSM93552 | 24093 | 45668 |
| 875.7498 | 362 | 886.6793 | 453 | 624.115 | 376 | 564.8419 | 389 | RELA | GM18505 | Blood | GSM93528 | 27088 | 45670 |
| 843.9612 | 204 | 940.055 | 274 | 423.269 | 209 | 390.5838 | 216 | RELA | GM18526 | Blood | GSM93528 | 12453 | 45672 |
| 920.5307 | 496 | 927.5786 | 613 | 665.3535 | 523 | 635.0486 | 553 | RELA | GM18951 | Blood | GSM93553 | 36778 | 45674 |
| 866.2056 | 447 | 910.892 | 597 | 644.4361 | 506 | 599.7868 | 512 | RELA | GM19099 | Blood | GSM93527 | 37529 | 45676 |
| 909.084 | 436 | 919.1243 | 542 | 654.7354 | 450 | 613.9041 | 458 | RELA | GM19193 | Blood | GSM93527 | 31597 | 45679 |
| 656.2824 | 282 | 774.0376 | 394 | 450.7134 | 365 | 430.3759 | 388 | CTCF | GM20000 | Blood | GSM10068 | 34184 | 45680 |
| 364.5214 | 252 | 520.0546 | 330 | 147.4963 | 291 | 206.8268 | 354 | CEBPB | H1 | Embryo | GSM93529 | 45379 | 45681 |
| 890.2786 | 357 | 912.3963 | 462 | 654.182 | 382 | 623.4557 | 398 | MAX | H1 | Embryo | GSM93534 | 26529 | 45682 |
| 1129.704 | 322 | 1114.604 | 378 | 857.6887 | 303 | 813.3763 | 300 | GTF2F1 | H1 | Embryo | GSM93558 | 10054 | 45683 |
| 1139.428 | 658 | 1169.359 | 836 | 802.8318 | 613 | 781.7867 | 662 | TBP | H1 | Embryo | GSM93530 | 28240 | 45684 |
| 408.6418 | 101 | 490.4785 | 123 | 173.4402 | 95 | 268.6145 | 125 | JUN | H1 | Embryo | GSM93561 | 6071 | 45685 |
| 477.6093 | 77 | 865.8994 | 120 | 273.0765 | 77 | 361.005 | 95 | CHD1 | H1 | Embryo | GSM93529 | 2429 | 45686 |
| 172.009 | 125 | 197.3283 | 150 | 75.57329 | 147 | 80.92691 | 164 | MAFK | H1 | Embryo | GSM93529 | 21934 | 45687 |
| 759.8712 | 493 | 777.7838 | 622 | 543.2808 | 579 | 520.8166 | 624 | ZNF143 | H1 | Embryo | GSM93551 | 63391 | 45688 |
| 546.756 | 400 | 634.5849 | 531 | 359.9925 | 549 | 335.3855 | 588 | RAD21 | H1 | Embryo | GSM93537 | 79460 | 45689 |
| 912.07 | 317 | 947.6234 | 426 | 644.3847 | 329 | 657.4316 | 359 | BACH1 | H1 | Embryo | GSM93558 | 20494 | 45690 |
| 1030.837 | 297 | 1045.171 | 376 | 790.0129 | 308 | 740.7852 | 304 | MYC | H1 | Embryo | GSM93550 | 12850 | 45691 |
| 410.7957 | 87 | 510.9688 | 108 | 222.88 | 89 | 190.7639 | 88 | RFX5 | H1 | Embryo | GSM93538 | 4159 | 45692 |
| 914.0798 | 347 | 932.4525 | 441 | 687.3547 | 375 | 666.6546 | 403 | CTBP2 | H1 | Embryo | GSM93546 | 22912 | 45693 |
| 930.5405 | 221 | 900.5156 | 248 | 418.8513 | 210 | 366.4029 | 212 | USF2 | H1 | Embryo | GSM93538 | 12577 | 45694 |
| 1059.374 | 351 | 1072.944 | 441 | 801.5316 | 351 | 752.5213 | 348 | MXI1 | H1 | Embryo | GSM93529 | 14302 | 45695 |
| 806.9911 | 358 | 814.4925 | 446 | 603.2227 | 416 | 591.1826 | 458 | JUND | H1 | Embryo | GSM93543 | 34316 | 45696 |
| 1130.925 | 192 | 1182.981 | 275 | 523.9891 | 157 | 718.9554 | 197 | NRF1 | H1 | Embryo | GSM93530 | 5445 | 45697 |
| 1002.74 | 212 | 978.3644 | 241 | 460.1652 | 190 | 518.3023 | 215 | BRCA1 | H1 | Embryo | GSM93551 | 9485 | 45698 |
| 1020.237 | 585 | 1045.607 | 740 | 736.8144 | 593 | 709.9122 | 635 | SIN3A | H1 | Embryo | GSM93528 | 33006 | 45699 |
| 14.79032 | 13 | 22.79144 | 18 | 12.71606 | 20 | 53.01057 | 39 | SUZ12 | H1 | Embryo | GSM93535 | 2791 | 45700 |
| 1065.153 | 490 | 1072.135 | 598 | 764.7158 | 456 | 740.1251 | 486 | CHD2 | H1 | Embryo | GSM93529 | 22372 | 45701 |
| 6.106115 | 6 | 6.286893 | 7 | 3.472794 | 8 | 57.86145 | 29 | ZNF274 | H1 | Embryo | GSM10036 | 1419 | 45702 |
| 541.1908 | 281 | 723.8371 | 395 | 396.3931 | 381 | 411.1875 | 419 | CTCF | HA-sp | Spinal Corc | GSM10226 | 39985 | 45703 |
| 690.6536 | 342 | 738.1747 | 470 | 455.6293 | 445 | 497.3306 | 499 | CTCF | HA-sp | Spinal Corc | GSM74969 | 48008 | 45704 |
| 690.471 | 368 | 751.7646 | 518 | 452.2719 | 465 | 494.5095 | 522 | CTCF | None | Cerebellun | GSM10226 | 51711 | 45705 |
| 495.417 | 248 | 731.3628 | 352 | 316.1127 | 316 | 344.8082 | 354 | CTCF | None | Cerebellun | GSM10226 | 32899 | 45706 |
| 647.0331 | 331 | 732.136 | 462 | 420.8343 | 430 | 466.7424 | 485 | CTCF | None | Umbilical \ | GSM74971 | 47559 | 45707 |
| 702.4283 | 471 | 769.3899 | 658 | 537.4713 | 624 | 525.2792 | 692 | CTCF | None | Umbilical \ | GSM74974 | 70995 | 45708 |
| 658.347 | 354 | 736.9053 | 511 | 469.9987 | 479 | 522.3543 | 549 | CTCF | None | Heart | GSM74973 | 53535 | 45709 |
| 581.7492 | 293 | 728.4745 | 408 | 354.6742 | 368 | 444.0265 | 434 | CTCF | None | Heart | GSM10226 | 39505 | 45710 |
| 686.8757 | 369 | 759.124 | 533 | 517.1005 | 491 | 524.7752 | 544 | CTCF | None | Heart | GSM10226 | 51821 | 45711 |
| 605.3481 | 352 | 699.9499 | 488 | 390.1664 | 464 | 438.2943 | 526 | CTCF | None | None | GSM74974 | 56756 | 45712 |
| 764.5524 | 515 | 821.4071 | 700 | 566.6003 | 637 | 552.7353 | 701 | CTCF | None | None | GSM74973 | 66031 | 45713 |
| 1055.277 | 684 | 1094.885 | 869 | 757.6388 | 703 | 738.1475 | 770 | TCF7L2 | HCT-116 | Colon | GSM78212 | 40831 | 45714 |
| 543.386 | 320 | 684.5213 | 438 | 365.2781 | 428 | 398.1113 | 480 | CTCF | HCT-116 | Colon | GSM10226 | 51474 | 45716 |
| 462.1363 | 269 | 685.951 | 371 | 307.0899 | 357 | 346.296 | 405 | CTCF | HCT-116 | Colon | GSM10226 | 42073 | 45717 |
| 444.5036 | 260 | 676.5364 | 352 | 284.1007 | 339 | 333.0459 | 390 | CTCF | None | Esophagus | GSM74972 | 39752 | 45718 |
| 702.8238 | 367 | 758.1134 | 510 | 504.0021 | 472 | 540.1997 | 543 | CTCF | None | Esophagus | GSM74971 | 49548 | 45719 |
| 669.1505 | 437 | 716.6425 | 591 | 501.6451 | 574 | 499.0559 | 648 | CTCF | HEK293 | Embryonic | GSM74966 | 71313 | 45720 |
| 606.0202 | 317 | 714.608 | 434 | 365.9316 | 402 | 402.309 | 452 | CTCF | HEK293 | Embryonic | GSM74968 | 46290 | 45721 |
| 338.8638 | 131 | 512.8631 | 179 | 458.3708 | 222 | 546.0609 | 258 | TCF7L2 | HEK293 | Embryonic | GSM78212 | 12616 | 45723 |
| 826.6313 | 97 | 1151.993 | 129 | 385.7984 | 82 | 429.0295 | 92 | ELK4 | HEK293 | Embryonic | GSM93559 | 1800 | 45724 |
| 495.7031 | 209 | 663.0341 | 267 | 335.4174 | 261 | 305.9943 | 272 | TRIM28 | HEK293 | Embryonic | GSM93559 | 20806 | 45725 |
| 812.6598 | 488 | 817.0701 | 598 | 619.8381 | 594 | 588.8827 | 632 | ZNF263 | HEK293T | Embryonic | GSM93557 | 48830 | 45726 |
| 390.3794 | 250 | 326.0488 | 263 | 250.8458 | 330 | 432.6241 | 436 | EP300 | HeLa-S3 | Cervix | GSM93550 | 40740 | 45728 |
| 1128.023 | 306 | 1161.346 | 402 | 892.807 | 316 | 846.1686 | 314 | ELK1 | HeLa-S3 | Cervix | GSM93532 | 9304 | 45729 |
| 956.8924 | 607 | 962.776 | 743 | 689.6283 | 654 | 690.9967 | 747 | RFX5 | HeLa-S3 | Cervix | GSM93556 | 45926 | 45730 |
| 639.2508 | 188 | 770.2008 | 229 | 358.3287 | 208 | 456.8453 | 249 | SMARCC1 | HeLa-S3 | Cervix | GSM93563 | 13884 | 45732 |
| 631.2232 | 292 | 761.5628 | 415 | 414.4341 | 375 | 457.7959 | 422 | CTCF | HeLa-S3 | Cervix | GSM74972 | 37893 | 45733 |
| 628.1191 | 345 | 710.8476 | 477 | 409.9799 | 454 | 444.059 | 508 | CTCF | HeLa-S3 | Cervix | GSM74973 | 53458 | 45734 |

| | | | | | | | | | | | | | |
|---|---|---|---|---|---|---|---|---|---|---|---|---|---|
| 1017.417 | 484 | 1018.949 | 590 | 777.1497 | 527 | 740.1962 | 545 | BRCA1 | HeLa-S3 | Cervix | GSM93555 | 25790 | 45736 |
| 996.3444 | 514 | 997.3478 | 619 | 737.0035 | 537 | 734.7562 | 609 | GTF2F1 | HeLa-S3 | Cervix | GSM93555 | 29972 | 45737 |
| 1020.339 | 389 | 1032.377 | 483 | 753.5996 | 380 | 716.6234 | 393 | ZNF143 | HeLa-S3 | Cervix | GSM93543 | 18791 | 45738 |
| 890.699 | 356 | 878.0288 | 421 | 672.3705 | 394 | 656.4344 | 431 | USF2 | HeLa-S3 | Cervix | GSM93556 | 25597 | 45741 |
| 713.5921 | 438 | 721.9268 | 541 | 517.3418 | 535 | 551.3152 | 661 | JUND | HeLa-S3 | Cervix | GSM93532 | 61396 | 45743 |
| 344.0434 | 210 | 303.0077 | 225 | 194.5238 | 263 | 348.4152 | 353 | JUN | HeLa-S3 | Cervix | GSM93534 | 32681 | 45744 |
| 1128.237 | 779 | 1178.948 | 999 | 817.6074 | 831 | 822.2624 | 962 | MYC | HeLa-S3 | Cervix | GSM93532 | 43783 | 45745 |
| 745.3027 | 309 | 743.6891 | 374 | 401.4121 | 355 | 377.9598 | 376 | MAFK | HeLa-S3 | Cervix | GSM93529 | 34623 | 45746 |
| 343.6296 | 109 | 259.1921 | 105 | 190.2392 | 121 | 196.8199 | 132 | PRDM1 | HeLa-S3 | Cervix | GSM93555 | 8693 | 45747 |
| 1262.302 | 214 | 1274.483 | 271 | 843.4566 | 178 | 857.9287 | 190 | IRF3 | HeLa-S3 | Cervix | GSM93557 | 4053 | 45748 |
| 1093.941 | 807 | 1143.455 | 1027 | 810.6199 | 914 | 805.8371 | 1040 | MAX | HeLa-S3 | Cervix | GSM93531 | 53042 | 45749 |
| 1218.96 | 517 | 1270.654 | 690 | 917.2591 | 490 | 881.2132 | 505 | E2F1 | HeLa-S3 | Cervix | GSM93536 | 14313 | 45750 |
| 885.8664 | 324 | 893.7659 | 399 | 695.7707 | 380 | 695.2049 | 432 | TCF7L2 | HeLa-S3 | Cervix | GSM93562 | 21430 | 45751 |
| 1227.203 | 351 | 1270.208 | 479 | 947.7182 | 329 | 913.5479 | 340 | E2F4 | HeLa-S3 | Cervix | GSM93536 | 8155 | 45752 |
| 982.2414 | 479 | 996.7047 | 594 | 777.622 | 563 | 756.8834 | 610 | TFAP2A | HeLa-S3 | Cervix | GSM93536 | 27804 | 45753 |
| 1202.185 | 444 | 1244.767 | 594 | 916.8852 | 420 | 878.8323 | 431 | E2F1 | HeLa-S3 | Cervix | GSM93548 | 12073 | 45754 |
| 1148.379 | 236 | 1191.432 | 326 | 911.8959 | 242 | 881.257 | 250 | ELK4 | HeLa-S3 | Cervix | GSM93535 | 6422 | 45755 |
| 483.2552 | 111 | 519.0292 | 127 | 182.0229 | 97 | 278.4793 | 127 | GTF3C2 | HeLa-S3 | Cervix | GSM93534 | 5954 | 45756 |
| 733.7224 | 538 | 754.6436 | 678 | 562.3974 | 709 | 559.3534 | 805 | RAD21 | HeLa-S3 | Cervix | GSM93557 | 77923 | 45757 |
| 1051.158 | 306 | 1066.636 | 385 | 773.1317 | 281 | 788.3747 | 333 | NFYB | HeLa-S3 | Cervix | GSM93540 | 12190 | 45758 |
| 828.8271 | 543 | 850.3016 | 686 | 618.2478 | 650 | 620.0547 | 749 | SMC3 | HeLa-S3 | Cervix | GSM93538 | 57115 | 45759 |
| 714.0821 | 510 | 752.8287 | 671 | 536.4481 | 661 | 518.2803 | 725 | CTCF | HeLa-S3 | Cervix | GSM82228 | 78170 | 45760 |
| 1137.545 | 611 | 1158.835 | 761 | 831.7872 | 587 | 819.8669 | 653 | MAZ | HeLa-S3 | Cervix | GSM93527 | 24320 | 45761 |
| 751.097 | 734 | 763.4142 | 902 | 541.6524 | 945 | 544.4407 | 1087 | CEBPB | HeLa-S3 | Cervix | GSM93555 | 130057 | 45762 |
| 172.2359 | 60 | 112.7156 | 54 | 47.79509 | 50 | 104.3694 | 75 | POLR3A | HeLa-S3 | Cervix | GSM93548 | 5072 | 45763 |
| 1087.018 | 669 | 1120.045 | 846 | 789.5144 | 682 | 777.5626 | 762 | CHD2 | HeLa-S3 | Cervix | GSM93543 | 34906 | 45764 |
| 923.4787 | 132 | 1181.218 | 188 | 372.1558 | 108 | 583.8362 | 146 | NRF1 | HeLa-S3 | Cervix | GSM93563 | 3593 | 45765 |
| 1154.039 | 519 | 1196.593 | 682 | 864.5951 | 504 | 843.0402 | 541 | HCFC1 | HeLa-S3 | Cervix | GSM10036 | 17745 | 45767 |
| 262.2336 | 58 | 249.4084 | 62 | 143.2251 | 60 | 228.2483 | 81 | SMARCA4 | HeLa-S3 | Cervix | GSM93551 | 2989 | 45768 |
| 1144.442 | 362 | 1174.809 | 472 | 876.4842 | 352 | 855.6595 | 373 | NFYA | HeLa-S3 | Cervix | GSM93550 | 11228 | 45769 |
| 994.4272 | 560 | 1005.782 | 687 | 766.8997 | 640 | 750.4944 | 705 | TFAP2C | HeLa-S3 | Cervix | GSM93537 | 33886 | 45770 |
| 740.623 | 299 | 669.9237 | 320 | 427.9167 | 357 | 549.0566 | 426 | STAT3 | HeLa-S3 | Cervix | GSM93527 | 33251 | 45771 |
| 858.0697 | 377 | 842.2285 | 441 | 658.592 | 446 | 688.5808 | 546 | FOS | HeLa-S3 | Cervix | GSM93531 | 31398 | 45772 |
| 1097.756 | 515 | 1133.211 | 669 | 826.9614 | 529 | 819.2354 | 589 | MXI1 | HeLa-S3 | Cervix | GSM93549 | 21513 | 45774 |
| 986.0334 | 592 | 997.6973 | 735 | 729.6668 | 646 | 729.8103 | 742 | STAT1 | HeLa-S3 | Cervix | GSM93536 | 39431 | 45775 |
| 1165.485 | 406 | 1212.282 | 541 | 900.6 | 396 | 861.5559 | 409 | E2F6 | HeLa-S3 | Cervix | GSM93547 | 11912 | 45776 |
| 953.0862 | 506 | 954.639 | 615 | 713.0643 | 553 | 704.7206 | 617 | RCOR1 | HeLa-S3 | Cervix | GSM93557 | 33536 | 45777 |
| 1064.963 | 718 | 1095.323 | 905 | 757.5079 | 739 | 758.1686 | 845 | MAX | HeLa-S3 | Cervix | GSM93540 | 43204 | 45778 |
| 1132.524 | 492 | 1153.203 | 620 | 844.3568 | 474 | 824.805 | 510 | SMARCB1 | HeLa-S3 | Cervix | GSM93563 | 17387 | 45779 |
| 1021.305 | 271 | 1046.06 | 356 | 824.2364 | 316 | 832.2941 | 365 | MYC | HeLa-S3 | Cervix | GSM82228 | 12003 | 45780 |
| 1072.784 | 621 | 1095.034 | 779 | 789.2177 | 640 | 779.3642 | 714 | TBP | HeLa-S3 | Cervix | GSM93560 | 31572 | 45781 |
| 202.8961 | 55 | 405.2432 | 89 | 214.5679 | 81 | 303.395 | 103 | TCF7L2 | HeLa-S3 | Cervix | GSM81643 | 3517 | 45782 |
| 1100.654 | 177 | 1151.58 | 247 | 801.6588 | 196 | 711.5564 | 195 | NR2C2 | HeLa-S3 | Cervix | GSM93536 | 5377 | 45783 |
| 991.4462 | 275 | 949.2509 | 301 | 743.1335 | 283 | 607.2308 | 275 | ZKSCAN1 | HeLa-S3 | Cervix | GSM93536 | 13185 | 45786 |
| 352.9367 | 90 | 577.2256 | 130 | 161.934 | 88 | 283.6604 | 123 | SMARCC2 | HeLa-S3 | Cervix | GSM93563 | 5524 | 45787 |
| 1149.245 | 517 | 1201.929 | 684 | 869.224 | 510 | 840.329 | 537 | MAZ | HepG2 | Liver | GSM93533 | 17969 | 45788 |
| 862.1693 | 552 | 878.712 | 688 | 658.4469 | 677 | 641.3165 | 742 | CEBPB | HepG2 | Liver | GSM93562 | 54783 | 45789 |
| 891.2153 | 468 | 913.6594 | 602 | 680.9534 | 554 | 678.952 | 627 | HNF4A | HepG2 | Liver | GSM93561 | 38377 | 45790 |
| 1136.7 | 528 | 1207.028 | 728 | 852.422 | 519 | 846.4302 | 586 | TBP | HepG2 | Liver | GSM93528 | 19258 | 45791 |
| 1203.293 | 411 | 1247.798 | 547 | 928.8815 | 395 | 885.0969 | 398 | SREBF1 | HepG2 | Liver | GSM93562 | 10965 | 45793 |
| 653.226 | 402 | 711.5568 | 562 | 450.984 | 535 | 505.3043 | 629 | CTCF | HepG2 | Liver | GSM74971 | 67209 | 45794 |
| 603.6349 | 311 | 721.9289 | 430 | 420.455 | 415 | 458.3577 | 465 | CTCF | HepG2 | Liver | GSM74968 | 45122 | 45795 |
| 923.7267 | 171 | 1028.73 | 222 | 473.2981 | 168 | 619.2577 | 205 | ESRRA | HepG2 | Liver | GSM93562 | 7172 | 45796 |
| 1139.521 | 518 | 1173.123 | 665 | 869.5965 | 525 | 823.0995 | 531 | MAX | HepG2 | Liver | GSM93540 | 18801 | 45797 |
| 1047.939 | 313 | 1049.716 | 382 | 785.4785 | 301 | 723.8521 | 293 | RCOR1 | HepG2 | Liver | GSM93557 | 12555 | 45798 |
| 275.5983 | 56 | 389.2884 | 75 | 102.0143 | 47 | 255.1865 | 80 | PPARGC1A | HepG2 | Liver | GSM93562 | 2596 | 45799 |
| 1147.262 | 170 | 1197.34 | 244 | 772.955 | 178 | 666.7853 | 174 | NR2C2 | HepG2 | Liver | GSM93559 | 4508 | 45800 |
| 227.8901 | 316 | 239.5085 | 370 | 113.7268 | 435 | 103.2146 | 469 | MAFF | HepG2 | Liver | GSM93530 | 92759 | 45801 |
| 1213.175 | 399 | 1244.892 | 521 | 927.6588 | 372 | 895.6299 | 390 | CHD2 | HepG2 | Liver | GSM93530 | 10045 | 45802 |
| 630.4995 | 162 | 749.0467 | 196 | 408.2142 | 188 | 352.2634 | 188 | USF2 | HepG2 | Liver | GSM93564 | 10406 | 45804 |
| 809.0366 | 379 | 812.2706 | 467 | 618.108 | 458 | 612.7095 | 515 | ARID3A | HepG2 | Liver | GSM93527 | 36396 | 45806 |
| 984.0855 | 472 | 1029.746 | 623 | 749.817 | 517 | 725.6157 | 552 | BHLHE40 | HepG2 | Liver | GSM93556 | 27514 | 45807 |
| 517.3371 | 346 | 648.6511 | 462 | 344.6794 | 471 | 425.6333 | 551 | CTCF | HepG2 | Liver | GSM82228 | 63465 | 45808 |
| 1200.118 | 151 | 1190.406 | 182 | 699.9594 | 148 | 514.1803 | 132 | BRCA1 | HepG2 | Liver | GSM93560 | 3335 | 45809 |
| 1122.134 | 638 | 1165.277 | 819 | 825.4058 | 639 | 799.8016 | 682 | MXI1 | HepG2 | Liver | GSM93543 | 27472 | 45810 |
| 679.5832 | 96 | 1139.133 | 143 | 293.6493 | 81 | 417.3949 | 104 | NRF1 | HepG2 | Liver | GSM93564 | 2541 | 45813 |
| 207.77 | 347 | 370.9426 | 480 | 157.796 | 543 | 182.7077 | 621 | CEBPB | HepG2 | Liver | GSM93549 | 115065 | 45814 |
| 117.5918 | 299 | 86.4907 | 322 | 27.89286 | 387 | 37.81651 | 454 | MAFK | HepG2 | Liver | GSM93530 | 117620 | 45815 |
| 438.2333 | 262 | 602.3356 | 338 | 348.4275 | 371 | 382.9187 | 417 | SMC3 | HepG2 | Liver | GSM93554 | 41106 | 45816 |
| 128.6542 | 243 | 115.8075 | 273 | 42.16631 | 316 | 40.86337 | 348 | MAFK | HepG2 | Liver | GSM93561 | 81735 | 45817 |
| 538.4129 | 314 | 678.3672 | 419 | 382.018 | 427 | 473.1021 | 501 | RAD21 | HepG2 | Liver | GSM93564 | 49721 | 45818 |
| 969.3429 | 175 | 999.2704 | 199 | 422.9999 | 158 | 499.4821 | 183 | HSF1 | HepG2 | Liver | GSM93562 | 7109 | 45820 |

| | | | | | | | | | | | | | |
|---|---|---|---|---|---|---|---|---|---|---|---|---|---|
| 899.1883 | 228 | 923.5147 | 298 | 634.1893 | 272 | 709.6951 | 307 | EP300 | HepG2 | Liver | GSM93554 | 14167 | 45822 |
| 1127.846 | 366 | 1129.364 | 450 | 853.0046 | 350 | 811.9551 | 353 | RFX5 | HepG2 | Liver | GSM93530 | 11987 | 45823 |
| 76.79881 | 36 | 75.50231 | 40 | 95.87511 | 60 | 182.6766 | 87 | TCF7L2 | HepG2 | Liver | GSM78212 | 4138 | 45825 |
| 1084.686 | 516 | 1135.186 | 686 | 837.9896 | 555 | 816.5091 | 600 | MYC | HepG2 | Liver | GSM82229 | 22753 | 45827 |
| 439.3316 | 379 | 436.9664 | 432 | 243.7201 | 506 | 336.7765 | 610 | JUND | HepG2 | Liver | GSM93564 | 85617 | 45828 |
| 203.5111 | 163 | 208.2234 | 187 | 108.9597 | 209 | 173.4767 | 266 | JUN | HepG2 | Liver | GSM93536 | 31580 | 45830 |
| 665.7286 | 337 | 737.6814 | 471 | 433.7638 | 437 | 475.8775 | 491 | CTCF | None | Foreskin | GSM10226 | 47688 | 45832 |
| 547.4582 | 288 | 722.9407 | 407 | 387.7633 | 385 | 408.4167 | 426 | CTCF | None | Foreskin | GSM10226 | 40673 | 45833 |
| 468.1392 | 266 | 685.0253 | 361 | 302.6013 | 347 | 349.2241 | 397 | CTCF | None | Foreskin | GSM10226 | 39793 | 45834 |
| 483.9382 | 255 | 675.6387 | 332 | 347.8909 | 344 | 365.5127 | 380 | CTCF | HL-60 | Blood | GSM74968 | 36625 | 45835 |
| 656.7074 | 343 | 727.1126 | 477 | 468.443 | 461 | 452.5969 | 495 | CTCF | None | Mammary | GSM10226 | 49610 | 45836 |
| 776.672 | 500 | 822.6751 | 668 | 565.6608 | 596 | 563.1459 | 676 | CTCF | None | Mammary | GSM74975 | 61367 | 45837 |
| 404.9847 | 275 | 659.8527 | 392 | 278.241 | 376 | 363.8707 | 449 | CTCF | None | Mammary | GSM74966 | 48340 | 45838 |
| 651.9761 | 349 | 727.1024 | 491 | 422.1324 | 456 | 454.1363 | 509 | CTCF | None | Mammary | GSM74967 | 52642 | 45839 |
| 616.9302 | 346 | 729.831 | 503 | 417.2435 | 461 | 452.0966 | 516 | CTCF | None | Pulmonary | GSM74975 | 53671 | 45840 |
| 675.0261 | 393 | 732.038 | 550 | 471.6269 | 512 | 510.6657 | 585 | CTCF | None | Pulmonary | GSM74968 | 59572 | 45841 |
| 657.1221 | 333 | 752.1502 | 483 | 450.5872 | 440 | 469.0889 | 485 | CTCF | None | Lung | GSM74969 | 47167 | 45842 |
| 611.9346 | 305 | 751.4207 | 442 | 399.0663 | 394 | 433.7741 | 441 | CTCF | None | Lung | GSM74971 | 41957 | 45843 |
| 679.2688 | 353 | 741.0516 | 503 | 459.9698 | 466 | 530.4116 | 545 | CTCF | HRE | None | GSM74973 | 51899 | 45844 |
| 688.0928 | 368 | 748.1728 | 518 | 423.0702 | 457 | 507.0598 | 530 | CTCF | HRE | None | GSM74972 | 52793 | 45845 |
| 574.5662 | 340 | 706.8058 | 485 | 389.823 | 457 | 435.4041 | 517 | CTCF | None | None | GSM74967 | 55403 | 45846 |
| 651.5133 | 331 | 741.6019 | 471 | 438.0564 | 435 | 486.9163 | 491 | CTCF | None | None | GSM10226 | 47390 | 45847 |
| 406.4667 | 273 | 379.206 | 301 | 232.9915 | 351 | 311.9019 | 422 | JUN | None | Umbilical V | GSM93527 | 47826 | 45848 |
| 1079.147 | 602 | 1118.747 | 774 | 793.9949 | 616 | 764.9901 | 655 | MAX | None | Umbilical V | GSM93557 | 30627 | 45849 |
| 188.6848 | 68 | 317.7663 | 99 | 170.3284 | 96 | 170.4994 | 103 | STAT4 | None | Blood | GSM67139 | 6296 | 4585 |
| 732.68 | 348 | 785.4977 | 482 | 494.6485 | 428 | 550.9087 | 487 | CTCF | None | Umbilical V | GSM74974 | 42414 | 45850 |
| 800.4959 | 457 | 860.3535 | 635 | 619.0231 | 582 | 584.5427 | 609 | CTCF | None | Umbilical V | GSM74967 | 49898 | 45851 |
| 213.191 | 182 | 210.8871 | 206 | 194.0931 | 278 | 215.8589 | 314 | GATA2 | None | Umbilical V | GSM93534 | 35008 | 45852 |
| 392.4241 | 303 | 448.1093 | 362 | 218.3868 | 392 | 334.5606 | 491 | FOS | None | Umbilical V | GSM93558 | 57624 | 45853 |
| 453.8263 | 299 | 650.5847 | 406 | 314.285 | 410 | 374.2364 | 474 | CTCF | None | Umbilical V | GSM82227 | 52681 | 45855 |
| 1099.519 | 595 | 1109.846 | 733 | 795.0956 | 578 | 773.8567 | 629 | MYC | None | Umbilical V | GSM82229 | 28038 | 45857 |
| 617.3462 | 311 | 730.037 | 431 | 395.7486 | 399 | 452.3291 | 455 | CTCF | None | Connective | GSM10226 | 42681 | 45858 |
| 610.1276 | 321 | 738.4944 | 466 | 432.3205 | 431 | 473.7404 | 484 | CTCF | None | Connective | GSM10226 | 46730 | 45859 |
| 423.129 | 536 | 488.894 | 645 | 277.1553 | 793 | 300.704 | 894 | CEBPB | IMR90 | Lung | GSM93551 | 167234 | 45861 |
| 763.2741 | 458 | 770.893 | 567 | 573.3476 | 567 | 547.8199 | 606 | RAD21 | IMR90 | Lung | GSM93562 | 55182 | 45862 |
| 855.1648 | 165 | 977.7953 | 195 | 380.7927 | 152 | 309.9436 | 147 | RFX5 | IMR90 | Lung | GSM10036 | 7235 | 45863 |
| 303.6481 | 331 | 302.3436 | 379 | 116.1616 | 411 | 126.8006 | 463 | MAFK | IMR90 | Lung | GSM93540 | 84225 | 45864 |
| 772.1662 | 442 | 823.2215 | 598 | 580.7131 | 540 | 556.2388 | 581 | CTCF | IMR90 | Lung | GSM93540 | 51216 | 45865 |
| 1115.86 | 619 | 1137.676 | 777 | 800.1551 | 594 | 772.7055 | 634 | MXI1 | IMR90 | Lung | GSM10036 | 27697 | 45866 |
| 1080.159 | 684 | 1129.804 | 885 | 781.7905 | 701 | 754.146 | 751 | MAZ | IMR90 | Lung | GSM10036 | 37038 | 45867 |
| 950.4277 | 235 | 964.3438 | 303 | 469.1813 | 222 | 554.3181 | 257 | CHD1 | IMR90 | Lung | GSM10036 | 12484 | 45868 |
| 771.319 | 262 | 780.1409 | 309 | 455.871 | 306 | 452.3828 | 329 | RCOR1 | IMR90 | Lung | GSM10036 | 23760 | 45869 |
| 1097.336 | 560 | 1129.189 | 716 | 818.8304 | 567 | 784.0545 | 594 | MYC | K562 | Bone Marr | GSM93541 | 25049 | 45870 |
| 943.0482 | 477 | 956.5939 | 599 | 711.3842 | 533 | 660.9414 | 534 | ATF1 | K562 | Bone Marr | GSM93534 | 32865 | 45871 |
| 343.9306 | 227 | 524.2742 | 306 | 307.004 | 338 | 306.5489 | 367 | TAL1 | K562 | Bone Marr | GSM93549 | 37723 | 45872 |
| 113.1648 | 29 | 82.87269 | 27 | 40.47408 | 25 | 51.25191 | 30 | SETDB1 | K562 | Bone Marr | GSM93557 | 1812 | 45873 |
| 940.4466 | 532 | 964.837 | 672 | 722.1098 | 622 | 688.5065 | 655 | JUN | K562 | Bone Marr | GSM93541 | 39481 | 45874 |
| 879.8963 | 302 | 866.7378 | 356 | 655.998 | 338 | 617.9558 | 354 | TBL1XR1 | K562 | Bone Marr | GSM93557 | 21230 | 45876 |
| 1201.754 | 247 | 1194.07 | 292 | 923.5613 | 219 | 904.5007 | 235 | GTF2F1 | K562 | Bone Marr | GSM93550 | 5569 | 45878 |
| 1093.198 | 296 | 1111.374 | 379 | 831.7991 | 284 | 795.5169 | 288 | MXI1 | K562 | Bone Marr | GSM93549 | 10067 | 45879 |
| 846.2525 | 616 | 866.168 | 769 | 632.7365 | 748 | 611.7839 | 816 | JUND | K562 | Bone Marr | GSM93556 | 67102 | 45880 |
| 1115.179 | 588 | 1152.855 | 757 | 827.3434 | 588 | 777.1553 | 596 | TBP | K562 | Bone Marr | GSM93549 | 24589 | 45881 |
| 1111.558 | 336 | 1125.098 | 422 | 852.4735 | 330 | 804.4224 | 328 | UBTF | K562 | Bone Marr | GSM93533 | 11158 | 45882 |
| 1016.818 | 632 | 1054.248 | 809 | 744.3844 | 675 | 710.3502 | 712 | MYC | K562 | Bone Marr | GSM93551 | 39039 | 45883 |
| 261.1886 | 156 | 334.5089 | 196 | 328.4879 | 264 | 275.0041 | 266 | GATA1 | K562 | Bone Marr | GSM10036 | 23646 | 45884 |
| 996.1807 | 588 | 1035.67 | 762 | 747.4761 | 653 | 720.2843 | 699 | MYC | K562 | Bone Marr | GSM93559 | 37958 | 45885 |
| 1132.179 | 169 | 1200.654 | 250 | 568.7329 | 153 | 807.5686 | 196 | NRF1 | K562 | Bone Marr | GSM93536 | 4702 | 45886 |
| 931.4329 | 192 | 970.7542 | 236 | 523.3951 | 201 | 396.6773 | 188 | SETDB1 | K562 | Bone Marr | GSM93559 | 9234 | 45887 |
| 923.4619 | 385 | 943.8193 | 493 | 677.0479 | 402 | 679.8252 | 459 | JUN | K562 | Bone Marr | GSM93552 | 25970 | 45888 |
| 149.0626 | 46 | 165.8637 | 54 | 98.04593 | 55 | 71.15619 | 51 | STAT5A | None | Blood | GSM67140 | 3618 | 4589 |
| 1115.307 | 526 | 1125.15 | 655 | 806.7614 | 491 | 772.2165 | 513 | HCFC1 | K562 | Bone Marr | GSM10036 | 20882 | 45890 |
| 881.2029 | 420 | 882.2641 | 511 | 657.9897 | 471 | 633.2116 | 504 | JUN | K562 | Bone Marr | GSM10036 | 33326 | 45892 |
| 1081.192 | 151 | 1118.139 | 174 | 495.2908 | 132 | 522.0118 | 144 | RFX5 | K562 | Bone Marr | GSM93556 | 3981 | 45893 |
| 972.6302 | 687 | 1024.615 | 893 | 720.5648 | 782 | 698.8527 | 856 | MYC | K562 | Bone Marr | GSM93554 | 53830 | 45894 |
| 1121.776 | 624 | 1158.45 | 802 | 844.2497 | 656 | 783.6108 | 648 | CCNT2 | K562 | Bone Marr | GSM93554 | 27695 | 45895 |
| 818.8053 | 249 | 810.8411 | 291 | 448.9416 | 277 | 441.0568 | 296 | ARID3A | K562 | Bone Marr | GSM93533 | 20143 | 45897 |
| 1133.815 | 678 | 1200.069 | 899 | 826.9095 | 683 | 794.9672 | 722 | E2F6 | K562 | Bone Marr | GSM93559 | 30671 | 45898 |
| 643.3492 | 428 | 711.387 | 604 | 510.4833 | 609 | 494.7371 | 668 | CTCF | K562 | Bone Marr | GSM82231 | 76017 | 45899 |
| 139.7165 | 55 | 202.6463 | 74 | 111.8245 | 74 | 85.95661 | 71 | BACH1 | K562 | Bone Marr | GSM93557 | 5457 | 45900 |
| 818.4072 | 174 | 970.6279 | 223 | 468.7359 | 185 | 505.8589 | 205 | GTF3C2 | K562 | Bone Marr | GSM93534 | 8818 | 45901 |
| 1128.224 | 131 | 1056.061 | 136 | 606.8808 | 121 | 758.0321 | 145 | ATF3 | K562 | Bone Marr | GSM93539 | 2529 | 45902 |
| 824.1274 | 623 | 852.6741 | 791 | 618.5158 | 773 | 588.2877 | 826 | RCOR1 | K562 | Bone Marr | GSM93543 | 73729 | 45903 |

| | | | | | | | | | | | | | |
|---|---|---|---|---|---|---|---|---|---|---|---|---|---|
| 629.612 | 157 | 733.6656 | 188 | 405.476 | 181 | 376.3646 | 187 | STAT1 | K562 | Bone Marr | GSM93547 | 9746 | 45904 |
| 1151.452 | 394 | 1186.451 | 523 | 870.6157 | 371 | 823.7061 | 370 | CHD2 | K562 | Bone Marr | GSM93550 | 12129 | 45905 |
| 262.8677 | 83 | 256.5511 | 91 | 185.4161 | 103 | 191.2262 | 112 | NFE2 | K562 | Bone Marr | GSM93541 | 6705 | 45906 |
| 1031.908 | 744 | 1078.487 | 946 | 746.6778 | 809 | 714.9376 | 869 | MAX | K562 | Bone Marr | GSM93534 | 52028 | 45907 |
| 688.8805 | 420 | 741.5255 | 576 | 535.3978 | 567 | 502.4816 | 598 | CTCF | K562 | Bone Marr | GSM93540 | 63165 | 45908 |
| 993.4112 | 296 | 965.6802 | 336 | 755.3943 | 307 | 703.3251 | 307 | STAT1 | K562 | Bone Marr | GSM93548 | 14556 | 45910 |
| 370.9738 | 133 | 484.4442 | 169 | 286.1574 | 174 | 213.5786 | 165 | RCOR1 | K562 | Bone Marr | GSM93538 | 12045 | 45911 |
| 560.7097 | 318 | 720.7894 | 465 | 425.3251 | 442 | 449.3107 | 490 | CTCF | K562 | Bone Marr | GSM74969 | 49636 | 45912 |
| 543.6547 | 280 | 730.4209 | 400 | 364.6389 | 367 | 448.5689 | 430 | CTCF | K562 | Bone Marr | GSM74973 | 39524 | 45913 |
| 558.7946 | 249 | 726.3224 | 319 | 374.1722 | 319 | 405.0474 | 356 | SMC3 | K562 | Bone Marr | GSM93531 | 30186 | 45914 |
| 922.9706 | 484 | 954.1159 | 624 | 693.6785 | 543 | 663.0551 | 571 | BHLHE40 | K562 | Bone Marr | GSM93561 | 34827 | 45916 |
| 919.1021 | 261 | 916.5611 | 321 | 726.4278 | 305 | 603.5391 | 300 | ZMIZ1 | K562 | Bone Marr | GSM10036 | 15714 | 45919 |
| 678.9409 | 205 | 857.2477 | 275 | 441.1832 | 245 | 353.112 | 239 | STAT5B | None | Blood | GSM67140 | 16339 | 4592 |
| 1170.2 | 612 | 1229.17 | 811 | 861.1734 | 592 | 815.5802 | 605 | HMGN3 | K562 | Bone Marr | GSM93554 | 22483 | 45920 |
| 593.0109 | 271 | 700.5603 | 329 | 301.9793 | 314 | 269.6289 | 327 | ZNF384 | K562 | Bone Marr | GSM10036 | 33683 | 45922 |
| 286.4952 | 336 | 439.1449 | 446 | 217.5873 | 507 | 227.6991 | 564 | CEBPB | K562 | Bone Marr | GSM93549 | 90229 | 45923 |
| 1012.789 | 686 | 1052.34 | 877 | 747.46 | 764 | 719.3308 | 819 | MAZ | K562 | Bone Marr | GSM93533 | 46864 | 45924 |
| 953.2003 | 381 | 975.3494 | 485 | 725.4129 | 411 | 698.9928 | 435 | IRF1 | K562 | Bone Marr | GSM93550 | 22258 | 45925 |
| 1057.883 | 362 | 1057.996 | 445 | 817.9591 | 385 | 773.8823 | 386 | MAX | K562 | Bone Marr | GSM93553 | 15529 | 45926 |
| 111.8505 | 31 | 128.6719 | 37 | 68.56148 | 35 | 118.292 | 49 | POLR3A | K562 | Bone Marr | GSM93537 | 2082 | 45927 |
| 1087.469 | 569 | 1126.819 | 735 | 809.1407 | 585 | 768.8021 | 599 | MYC | K562 | Bone Marr | GSM82231 | 26476 | 45928 |
| 304.7405 | 96 | 268.9971 | 100 | 163.0685 | 105 | 160.3141 | 112 | SIRT6 | K562 | Bone Marr | GSM93537 | 7692 | 45929 |
| 585.6695 | 78 | 842.7324 | 106 | 349.4356 | 78 | 449.2472 | 95 | STAT5B | None | Blood | GSM67140 | 1916 | 4593 |
| 271.8626 | 123 | 285.5448 | 141 | 197.0361 | 162 | 275.9343 | 201 | JUN | K562 | Bone Marr | GSM93546 | 14408 | 45930 |
| 1177.208 | 205 | 1163.83 | 243 | 701.4621 | 178 | 648.234 | 181 | GTF2B | K562 | Bone Marr | GSM93539 | 5161 | 45932 |
| 16.45859 | 20 | 14.68766 | 22 | 3.002145 | 20 | 17.71612 | 37 | TRIM28 | K562 | Bone Marr | GSM93546 | 4952 | 45933 |
| 169.6444 | 74 | 255.179 | 101 | 124.8689 | 97 | 143.8769 | 111 | CUX1 | K562 | Bone Marr | GSM10036 | 8024 | 45934 |
| 685.0666 | 141 | 943.0978 | 185 | 382.0149 | 146 | 348.7396 | 149 | TBL1XR1 | K562 | Bone Marr | GSM93557 | 6522 | 45935 |
| 916.4675 | 310 | 946.6963 | 409 | 631.5228 | 319 | 649.6309 | 347 | NFYB | K562 | Bone Marr | GSM93542 | 19733 | 45936 |
| 590.0192 | 146 | 674.194 | 173 | 361.6077 | 164 | 325.3447 | 167 | STAT1 | K562 | Bone Marr | GSM93548 | 8914 | 45937 |
| 422.8774 | 142 | 467.3047 | 166 | 230.7206 | 159 | 353.0902 | 206 | JUN | K562 | Bone Marr | GSM93546 | 12395 | 45938 |
| 143.2947 | 65 | 165.5399 | 78 | 111.7886 | 88 | 106.6453 | 93 | SMARCA4 | K562 | Bone Marr | GSM93563 | 7432 | 45939 |
| 139.2405 | 55 | 196.5891 | 73 | 111.0125 | 74 | 96.94021 | 75 | SMARCB1 | K562 | Bone Marr | GSM93563 | 5520 | 45940 |
| 559.5135 | 105 | 542.9197 | 113 | 346.1313 | 113 | 225.8483 | 97 | USF2 | K562 | Bone Marr | GSM93535 | 4330 | 45941 |
| 78.24568 | 33 | 70.40687 | 35 | 33.41423 | 34 | 48.63534 | 43 | ZNF274 | K562 | Bone Marr | GSM93550 | 3410 | 45942 |
| 216.441 | 132 | 277.3244 | 166 | 260.5566 | 220 | 304.487 | 253 | GATA2 | K562 | Bone Marr | GSM93537 | 20178 | 45944 |
| 1110.823 | 262 | 1137.154 | 342 | 834.9562 | 243 | 836.9306 | 274 | NFYA | K562 | Bone Marr | GSM93543 | 8372 | 45945 |
| 817.5535 | 429 | 849.1007 | 559 | 604.8564 | 497 | 569.4977 | 515 | ZNF143 | K562 | Bone Marr | GSM93556 | 42550 | 45946 |
| 784.1298 | 93 | 998.842 | 117 | 484.7801 | 93 | 572.9319 | 108 | NELFE | K562 | Bone Marr | GSM93539 | 1828 | 45947 |
| 1100.214 | 575 | 1135.014 | 742 | 833.6847 | 609 | 794.8065 | 631 | IRF1 | K562 | Bone Marr | GSM93554 | 25920 | 45949 |
| 101.1715 | 148 | 113.008 | 177 | 60.22741 | 212 | 76.84352 | 249 | MAFF | K562 | Bone Marr | GSM93552 | 41661 | 45950 |
| 1237.045 | 573 | 1258.056 | 720 | 926.9611 | 544 | 879.6761 | 554 | E2F4 | K562 | Bone Marr | GSM93560 | 16454 | 45951 |
| 484.1345 | 191 | 416.5835 | 199 | 305.1394 | 234 | 359.3141 | 270 | JUN | K562 | Bone Marr | GSM93560 | 20118 | 45952 |
| 1177.074 | 186 | 1207.149 | 244 | 788.9013 | 178 | 659.7715 | 171 | ELK1 | K562 | Bone Marr | GSM10036 | 4385 | 45953 |
| 190.7108 | 166 | 188.1489 | 188 | 96.5071 | 211 | 131.0744 | 254 | MAFK | K562 | Bone Marr | GSM93531 | 33556 | 45954 |
| 567.7559 | 140 | 840.926 | 191 | 390.0013 | 166 | 338.8438 | 166 | STAT1 | K562 | Bone Marr | GSM93547 | 8505 | 45955 |
| 1114.16 | 253 | 1131.619 | 328 | 867.3009 | 255 | 769.5769 | 243 | YY1 | K562 | Bone Marr | GSM93536 | 8001 | 45956 |
| 689.0643 | 164 | 918.1631 | 211 | 426.7303 | 184 | 405.1245 | 192 | ZNF263 | K562 | Bone Marr | GSM93553 | 9505 | 45958 |
| 960.6909 | 368 | 973.0517 | 462 | 708.3106 | 376 | 723.4515 | 447 | FOS | K562 | Bone Marr | GSM93535 | 21739 | 45959 |
| 412.2415 | 116 | 486.2279 | 140 | 266.7169 | 136 | 248.283 | 141 | STAT2 | K562 | Bone Marr | GSM93546 | 8179 | 45960 |
| 1085.028 | 561 | 1111.557 | 715 | 811.9762 | 581 | 749.4254 | 567 | UBTF | K562 | Bone Marr | GSM93564 | 25845 | 45961 |
| 786.0391 | 359 | 783.6174 | 437 | 577.9529 | 423 | 598.8195 | 496 | EP300 | K562 | Bone Marr | GSM93540 | 36759 | 45963 |
| 1054.437 | 491 | 1064.284 | 608 | 787.8467 | 505 | 752.0906 | 527 | MYC | K562 | Bone Marr | GSM93546 | 23645 | 45965 |
| 304.7805 | 178 | 416.3371 | 230 | 228.3722 | 246 | 240.7378 | 272 | RAD21 | K562 | Bone Marr | GSM93531 | 26937 | 45966 |
| 945.7777 | 250 | 955.17 | 317 | 726.8011 | 285 | 688.047 | 297 | STAT2 | K562 | Bone Marr | GSM93547 | 13864 | 45967 |
| 1002.031 | 426 | 1018.276 | 540 | 749.1657 | 442 | 711.9315 | 455 | IRF1 | K562 | Bone Marr | GSM93550 | 22304 | 45969 |
| 219.9558 | 84 | 325.3673 | 114 | 243.8092 | 131 | 269.4026 | 147 | GATA1 | K562 | Bone Marr | GSM93554 | 8276 | 45970 |
| 357.5732 | 93 | 498.8168 | 123 | 293.9277 | 121 | 234.8762 | 116 | IRF1 | K562 | Bone Marr | GSM93554 | 5954 | 45971 |
| 167.9021 | 53 | 215.1005 | 67 | 136.2371 | 70 | 117.553 | 70 | EP300 | K562 | Bone Marr | GSM93549 | 4197 | 45972 |
| 1046.991 | 428 | 1081.417 | 561 | 798.3455 | 452 | 772.5193 | 477 | MYC | K562 | Bone Marr | GSM93542 | 19752 | 45973 |
| 867.2493 | 489 | 918.678 | 655 | 656.1641 | 575 | 623.9644 | 606 | CTCF | None | Kidney | GSM10068 | 43853 | 45974 |
| 673.0114 | 339 | 738.9866 | 476 | 510.5703 | 466 | 499.649 | 502 | CTCF | LNCaP | Prostate | GSM10068 | 48742 | 45975 |
| 743.0268 | 420 | 802.9897 | 586 | 582.9325 | 560 | 564.228 | 611 | CTCF | LNCaP | Prostate | GSM10068 | 53863 | 45976 |
| 872.0543 | 422 | 894.6645 | 544 | 660.9438 | 495 | 633.8138 | 522 | CTCF | None | Lung | GSM10068 | 35473 | 45977 |
| 448.7766 | 156 | 462.1984 | 176 | 755.6178 | 296 | 781.4419 | 362 | ZNF217 | MCF-7 | Breast | GSM93556 | 13561 | 45978 |
| 656.8673 | 195 | 862.1937 | 260 | 788.4703 | 344 | 805.8519 | 413 | TCF7L2 | MCF-7 | Breast | GSM81643 | 14485 | 45979 |
| 924.9166 | 816 | 946.0203 | 1003 | 670.7901 | 984 | 635.928 | 1060 | MYC | MCF-7 | Breast | GSM10068 | 93048 | 45980 |
| 391.0983 | 103 | 244.464 | 89 | 545.6128 | 174 | 801.8855 | 226 | GATA3 | MCF-7 | Breast | GSM82571 | 6427 | 45981 |
| 382.8097 | 147 | 340.7558 | 155 | 667.5589 | 280 | 758.6385 | 354 | GATA3 | MCF-7 | Breast | GSM93544 | 14099 | 45982 |
| 798.6528 | 675 | 848.2284 | 881 | 611.2868 | 888 | 567.0286 | 929 | CTCF | MCF-7 | Breast | GSM82230 | 95400 | 45983 |
| 865.1291 | 797 | 911.7041 | 1013 | 639.7601 | 1012 | 613.274 | 1107 | CTCF | MCF-7 | Breast | GSM10068 | 108931 | 45984 |
| 428.5879 | 295 | 665.1546 | 427 | 313.4667 | 414 | 332.0057 | 460 | CTCF | MCF-7 | Breast | GSM10226 | 53556 | 45987 |

| | | | | | | | | | | | | | |
|---|---|---|---|---|---|---|---|---|---|---|---|---|---|
| 542.9906 | 355 | 669.355 | 490 | 373.3099 | 486 | 435.4326 | 558 | CTCF | MCF-7 | Breast | GSM10226 | 63347 | 45988 |
| 1090.553 | 898 | 1126.655 | 1110 | 787.5449 | 1030 | 780.7331 | 1168 | MYC | MCF-7 | Breast | GSM82230 | 70048 | 45989 |
| 890.7404 | 451 | 883.8888 | 538 | 695.5762 | 544 | 663.9103 | 575 | MYC | MCF-7 | Breast | GSM82230 | 35069 | 45990 |
| 1123.56 | 629 | 1141.584 | 777 | 859.7317 | 666 | 819.9047 | 698 | E2F1 | MCF-7 | Breast | GSM93547 | 25437 | 45991 |
| 769.9933 | 713 | 773.4085 | 864 | 570.0132 | 928 | 549.3058 | 1020 | CTCF | MCF-7 | Breast | GSM10068 | 115846 | 45992 |
| 714.0856 | 586 | 749.3922 | 755 | 543.9389 | 783 | 513.894 | 839 | CTCF | MCF-7 | Breast | GSM82230 | 95843 | 45994 |
| 645.5626 | 453 | 681.2161 | 598 | 493.1177 | 619 | 478.3743 | 682 | CTCF | MCF-7 | Breast | GSM82230 | 81046 | 45995 |
| 662.961 | 645 | 659.3611 | 775 | 506.8703 | 903 | 466.2165 | 955 | MYC | MCF-7 | Breast | GSM10068 | 134833 | 45996 |
| 648.2463 | 433 | 648.919 | 530 | 369.5728 | 536 | 470.726 | 633 | STAT3 | MCF-10A | Breast | GSM93545 | 74730 | 45997 |
| 388.9873 | 452 | 426.911 | 535 | 248.326 | 652 | 347.3249 | 784 | FOS | MCF-10A | Breast | GSM93539 | 125466 | 45999 |
| 1117.603 | 648 | 1141.125 | 817 | 789.8665 | 621 | 757.6325 | 657 | E2F4 | MCF-10A | Breast | GSM93540 | 31113 | 46000 |
| 793.2926 | 469 | 791.6963 | 567 | 584.0326 | 553 | 578.7399 | 623 | MYC | MCF-10A | Breast | GSM93549 | 51993 | 46001 |
| 560.7362 | 370 | 613.917 | 448 | 343.8014 | 489 | 480.8728 | 596 | STAT3 | MCF-10A | Breast | GSM93539 | 67064 | 46002 |
| 584.8837 | 483 | 586.4914 | 589 | 330.0803 | 641 | 433.9537 | 761 | FOS | MCF-10A | Breast | GSM93539 | 108412 | 46004 |
| 361.9025 | 430 | 399.5335 | 510 | 263.4219 | 642 | 389.7195 | 785 | FOS | MCF-10A | Breast | GSM93539 | 119111 | 46005 |
| 683.8585 | 259 | 759.9501 | 318 | 417.1607 | 313 | 504.1923 | 365 | STAT3 | MCF-10A | Breast | GSM93559 | 27095 | 46006 |
| 615.2681 | 489 | 599.7053 | 576 | 411.9194 | 654 | 465.9409 | 775 | FOS | MCF-10A | Breast | GSM93543 | 100855 | 46007 |
| 963.5324 | 740 | 1014.851 | 957 | 690.5829 | 834 | 684.212 | 944 | MYC | MCF-10A | Breast | GSM93544 | 66731 | 46008 |
| 690.6663 | 372 | 727.0939 | 496 | 458.2939 | 472 | 472.6957 | 519 | CTCF | None | Brain | GSM10068 | 53191 | 46009 |
| 967.3958 | 657 | 1001.872 | 839 | 708.1181 | 732 | 653.0413 | 738 | MYC | NB4 | Blood | GSM93564 | 50249 | 46011 |
| 914.2793 | 717 | 951.5994 | 908 | 659.8147 | 829 | 614.2816 | 862 | MAX | NB4 | Blood | GSM93564 | 71695 | 46012 |
| 499.9524 | 284 | 678.8113 | 379 | 334.5279 | 377 | 380.4242 | 429 | CTCF | NB4 | Blood | GSM10226 | 43831 | 46013 |
| 712.4362 | 365 | 755.1475 | 494 | 481.8304 | 455 | 530.3559 | 514 | CTCF | NHDF-neo | Skin | GSM10226 | 47928 | 46014 |
| 654.7614 | 309 | 759.0668 | 439 | 446.7844 | 402 | 483.2213 | 449 | CTCF | NHDF-neo | Skin | GSM10226 | 39825 | 46015 |
| 522.5807 | 309 | 668.7164 | 406 | 294.4649 | 388 | 381.6352 | 462 | CTCF | NHEK | Skin | GSM74974 | 49047 | 46016 |
| 754.168 | 453 | 812.7652 | 622 | 564.9822 | 559 | 541.574 | 601 | CTCF | NHEK | Skin | GSM74970 | 56630 | 46017 |
| 660.0456 | 304 | 751.2906 | 419 | 434.3818 | 389 | 443.2387 | 425 | CTCF | NHLF | Lung | GSM10226 | 38385 | 46018 |
| 106.4601 | 25 | 146.3257 | 33 | 82.84285 | 31 | 97.72203 | 36 | SUZ12 | NT2-D1 | Embryo | GSM93542 | 1352 | 46019 |
| 13.81091 | 8 | 13.63815 | 9 | 0.545498 | 4 | 14.93784 | 14 | ZNF274 | NT2-D1 | Embryo | GSM93558 | 1205 | 46020 |
| 1040.926 | 292 | 1081.44 | 399 | 812.8552 | 310 | 721.7605 | 286 | YY1 | NT2-D1 | Embryo | GSM93542 | 12161 | 46021 |
| 1030.567 | 345 | 1025.935 | 410 | 867.0415 | 435 | 847.096 | 468 | TCF7L2 | PANC-1 | Pancreas | GSM81643 | 14640 | 46022 |
| 848.8592 | 485 | 899.1416 | 655 | 646.2173 | 586 | 624.1038 | 634 | CTCF | None | Pancreas | GSM10068 | 46552 | 46023 |
| 844.955 | 374 | 859.0385 | 469 | 651.5942 | 442 | 618.1214 | 459 | GATA1 | PBDE | Blood | GSM93546 | 29890 | 46025 |
| 49.15557 | 23 | 121.8813 | 41 | 45.99801 | 34 | 65.87273 | 43 | GATA1 | PBDE | Blood | GSM93533 | 2791 | 46026 |
| 472.8758 | 308 | 665.4762 | 428 | 304.2872 | 409 | 357.9916 | 471 | CTCF | RPTEC | Kidney | GSM10226 | 52073 | 46028 |
| 663.5681 | 381 | 712.6251 | 521 | 417.6378 | 490 | 477.6718 | 559 | CTCF | RPTEC | Kidney | GSM10226 | 59674 | 46029 |
| 770.8456 | 448 | 814.9165 | 600 | 577.0695 | 549 | 561.161 | 603 | CTCF | SAEC | Lung | GSM74977 | 52875 | 46030 |
| 795.1486 | 438 | 845.5861 | 594 | 594.1836 | 524 | 591.849 | 597 | CTCF | SAEC | Lung | GSM74968 | 47422 | 46031 |
| 432.4496 | 295 | 544.4044 | 366 | 358.1654 | 428 | 461.2309 | 509 | GATA2 | SH-SY5Y | Bone Marr | GSM93558 | 50625 | 46032 |
| 32.52545 | 60 | 39.47559 | 74 | 86.40433 | 138 | 107.2446 | 162 | GATA3 | SH-SY5Y | Bone Marr | GSM93556 | 17812 | 46033 |
| 1134.063 | 474 | 1191.099 | 651 | 845.4502 | 455 | 807.7646 | 468 | NRF1 | SK-N-SH | Brain | GSM10036 | 17290 | 46034 |
| 786.1419 | 496 | 842.6378 | 676 | 597.0961 | 621 | 579.0068 | 678 | CTCF | SK-N-SH | Brain | GSM10036 | 58234 | 46035 |
| 728.0465 | 403 | 725.0177 | 487 | 452.9882 | 466 | 529.9218 | 550 | JUND | SK-N-SH | Brain | GSM10036 | 52270 | 46036 |
| 1025.995 | 489 | 1019.129 | 585 | 729.4658 | 465 | 704.3499 | 494 | RFX5 | SK-N-SH | Brain | GSM10036 | 25565 | 46037 |
| 1080.851 | 750 | 1125.974 | 959 | 792.8471 | 816 | 749.9179 | 857 | MXI1 | SK-N-SH | Brain | GSM10036 | 45316 | 46038 |
| 745.8346 | 477 | 736.3907 | 570 | 578.4762 | 622 | 545.0253 | 657 | EP300 | SK-N-SH | Brain | GSM10036 | 60644 | 46039 |
| 750.1726 | 388 | 774.3134 | 497 | 569.2845 | 481 | 561.5375 | 538 | SMC3 | SK-N-SH | Brain | GSM10036 | 45541 | 46040 |
| 715.0081 | 507 | 744.5303 | 656 | 546.2453 | 668 | 544.8869 | 761 | RAD21 | SK-N-SH_R | Brain | GSM10036 | 74598 | 46041 |
| 802.8778 | 497 | 860.5912 | 678 | 604.2805 | 606 | 585.9878 | 663 | CTCF | SK-N-SH_R | Brain | GSM74969 | 55570 | 46042 |
| 811.6576 | 498 | 862.4126 | 667 | 602.0329 | 593 | 585.0468 | 648 | CTCF | SK-N-SH_R | Brain | GSM74966 | 54111 | 46043 |
| 976.4882 | 642 | 1036.813 | 854 | 687.5276 | 665 | 670.0491 | 730 | CTCF | None | Spleen | GSM10068 | 47937 | 46044 |
| 154.6401 | 186 | 185.1915 | 227 | 36.03034 | 201 | 55.80223 | 247 | SETDB1 | U2OS | Bone | GSM93545 | 44333 | 46045 |
| 674.6202 | 230 | 726.7788 | 265 | 371.9962 | 258 | 499.2912 | 315 | TRIM28 | U2OS | Bone | GSM93558 | 19460 | 46046 |
| 678.5523 | 443 | 735.0212 | 609 | 517.3059 | 590 | 513.3143 | 666 | CTCF | WERI-Rb-1 | Eye | GSM74976 | 70965 | 46047 |
| 633.8266 | 345 | 725.3956 | 490 | 397.2833 | 447 | 465.0899 | 514 | CTCF | WERI-Rb-1 | Eye | GSM74967 | 53016 | 46048 |
| 536.0277 | 218 | 774.7391 | 300 | 320.8357 | 263 | 370.4384 | 301 | CTCF | WI-38 | Lung | GSM10226 | 23819 | 46049 |
| 471.5835 | 235 | 732.74 | 337 | 345.1299 | 316 | 382.5868 | 356 | CTCF | WI-38 | Lung | GSM10226 | 31431 | 46050 |
| 390.6627 | 74 | 442.1687 | 87 | 293.0391 | 88 | 439.019 | 116 | BCL3 | A549 | Lung | GSM10107 | 3047 | 46051 |
| 1092.52 | 694 | 1127.647 | 877 | 827.1201 | 768 | 812.4279 | 854 | NR3C1 | A549 | Lung | GSM80337 | 37115 | 46052 |
| 634.0774 | 326 | 700.9689 | 424 | 409.1342 | 423 | 450.959 | 476 | CTCF | A549 | Lung | GSM80345 | 47319 | 46053 |
| 1172.462 | 259 | 1177.858 | 314 | 902.3534 | 234 | 883.8971 | 251 | SIX5 | A549 | Lung | GSM10108 | 6152 | 46054 |
| 965.6304 | 224 | 926.6044 | 249 | 634.3129 | 242 | 615.9663 | 255 | ATF3 | A549 | Lung | GSM10107 | 10858 | 46055 |
| 859.77 | 294 | 845.7029 | 346 | 684.5149 | 362 | 713.8356 | 446 | EP300 | A549 | Lung | GSM10108 | 20965 | 46056 |
| 1069.369 | 215 | 1061.453 | 256 | 735.3064 | 215 | 598.2455 | 206 | SIN3A | A549 | Lung | GSM10108 | 6850 | 46057 |
| 894.05 | 345 | 861.3261 | 388 | 738.324 | 450 | 748.7946 | 526 | GATA3 | A549 | Lung | GSM10107 | 23403 | 46058 |
| 1134.798 | 371 | 1141.005 | 458 | 863.9811 | 352 | 819.1329 | 351 | ELF1 | A549 | Lung | GSM10107 | 11340 | 46059 |
| 1070.943 | 697 | 1056.787 | 823 | 794.6274 | 754 | 771.6787 | 821 | SP1 | A549 | Lung | GSM10108 | 38052 | 46060 |
| 1057.705 | 477 | 1051.518 | 575 | 791.5992 | 490 | 773.7273 | 532 | USF1 | A549 | Lung | GSM80342 | 22760 | 46062 |
| 524.1398 | 286 | 710.2521 | 400 | 401.9773 | 397 | 422.7821 | 439 | CTCF | A549 | Lung | GSM80345 | 42677 | 46063 |
| 1007.762 | 271 | 946.0246 | 282 | 724.9403 | 269 | 686.3602 | 280 | USF1 | A549 | Lung | GSM10108 | 11920 | 46064 |
| 1074.244 | 466 | 1090.083 | 583 | 788.0665 | 440 | 752.2765 | 458 | YY1 | A549 | Lung | GSM10107 | 19081 | 46065 |
| 1010.714 | 561 | 963.5247 | 620 | 762.2965 | 612 | 760.7584 | 694 | TCF12 | A549 | Lung | GSM10107 | 31495 | 46066 |

| | | | | | | | | | | | | | |
|---|---|---|---|---|---|---|---|---|---|---|---|---|---|
| 801.7206 | 487 | 767.3146 | 547 | 638.8675 | 648 | 659.4933 | 775 | FOXA1 | A549 | Lung | GSM10108 | 54218 | 46067 |
| 1135.14 | 277 | 1141.147 | 338 | 857.997 | 246 | 846.4105 | 270 | ETS1 | A549 | Lung | GSM10108 | 7569 | 46068 |
| 1040.774 | 412 | 1032.699 | 491 | 798.1299 | 426 | 789.0363 | 478 | REST | A549 | Lung | GSM10107 | 17365 | 46069 |
| 1121.517 | 750 | 1113.791 | 892 | 794.6162 | 749 | 775.9015 | 822 | MAX | A549 | Lung | GSM10107 | 37512 | 46070 |
| 1134.225 | 461 | 1162.045 | 589 | 856.9563 | 447 | 805.654 | 440 | E2F6 | A549 | Lung | GSM10107 | 15236 | 46071 |
| 726.3899 | 318 | 672.6256 | 343 | 437.6086 | 387 | 581.6893 | 489 | FOSL2 | A549 | Lung | GSM10107 | 36994 | 46072 |
| 1129.688 | 309 | 1107.672 | 360 | 859.4672 | 292 | 841.9442 | 316 | ZBTB33 | A549 | Lung | GSM10108 | 9224 | 46073 |
| 646.7976 | 314 | 669.6167 | 363 | 432.7949 | 408 | 488.2923 | 463 | CEBPB | A549 | Lung | GSM10108 | 42163 | 46074 |
| 678.4411 | 218 | 803.9093 | 265 | 523.8129 | 283 | 600.3062 | 323 | TEAD4 | A549 | Lung | GSM10108 | 18362 | 46075 |
| 547.4341 | 110 | 464.0336 | 110 | 335.4289 | 119 | 425.1889 | 143 | NR3C1 | A549 | Lung | GSM80335 | 4968 | 46076 |
| 1249.762 | 456 | 1276.641 | 586 | 937.0845 | 399 | 915.0949 | 428 | TAF1 | A549 | Lung | GSM10108 | 9980 | 46077 |
| 1056.705 | 544 | 1023.287 | 617 | 769.6778 | 545 | 758.781 | 608 | USF1 | A549 | Lung | GSM80345 | 27979 | 46078 |
| 824.5162 | 403 | 809.0523 | 472 | 665.982 | 532 | 683.7 | 632 | NR3C1 | A549 | Lung | GSM80335 | 37619 | 46079 |
| 1109.222 | 415 | 1096.816 | 490 | 835.2368 | 397 | 813.6352 | 430 | GABPA | A549 | Lung | GSM10108 | 14271 | 46080 |
| 438.1725 | 169 | 430.4561 | 187 | 364.0876 | 231 | 543.1649 | 296 | NR3C1 | A549 | Lung | GSM80335 | 16898 | 46081 |
| 1188.213 | 341 | 1184.333 | 412 | 920.6841 | 326 | 891.5176 | 339 | PBX3 | A549 | Lung | GSM10107 | 8717 | 46082 |
| 506.2026 | 315 | 664.4669 | 422 | 403.4057 | 449 | 399.3115 | 487 | RAD21 | A549 | Lung | GSM10107 | 52197 | 46084 |
| 1001.984 | 546 | 1013.694 | 677 | 754.3342 | 593 | 731.2416 | 641 | TCF12 | ECC-1 | Endometri | GSM10108 | 31468 | 46085 |
| 1114.788 | 363 | 1132.17 | 466 | 856.1999 | 364 | 815.9884 | 370 | EGR1 | ECC-1 | Endometri | GSM10108 | 12410 | 46086 |
| 715.1484 | 248 | 812.8278 | 327 | 613.5843 | 341 | 609.8747 | 366 | ESR1 | ECC-1 | Endometri | GSM80354 | 23186 | 46087 |
| 680.4214 | 372 | 715.2737 | 493 | 517.9192 | 503 | 511.25 | 546 | RAD21 | ECC-1 | Endometri | GSM10108 | 54682 | 46088 |
| 505.4129 | 236 | 590.5725 | 284 | 318.8975 | 297 | 360.1743 | 337 | CEBPB | ECC-1 | Endometri | GSM10108 | 29653 | 46089 |
| 2.312029 | 3 | 9.885526 | 7 | 2.960572 | 6 | 20.71647 | 15 | TAL1 | None | Blood | GSM71061 | 1150 | 4609 |
| 557.336 | 145 | 460.3182 | 145 | 405.3195 | 178 | 478.3656 | 206 | FOXA1 | ECC-1 | Endometri | GSM80344 | 9407 | 46090 |
| 359.8172 | 187 | 510.4869 | 246 | 218.1523 | 235 | 259.766 | 272 | CTCF | ECC-1 | Endometri | GSM10107 | 25622 | 46091 |
| 1037.985 | 409 | 1033.718 | 488 | 781.9081 | 415 | 731.0212 | 410 | SRF | ECC-1 | Endometri | GSM10107 | 18570 | 46092 |
| 928.3555 | 247 | 954.6578 | 323 | 709.682 | 286 | 632.3524 | 290 | FOXM1 | ECC-1 | Endometri | GSM10108 | 13888 | 46093 |
| 1086.019 | 732 | 1135.979 | 938 | 788.0375 | 766 | 759.1412 | 823 | MAX | ECC-1 | Endometri | GSM10108 | 39975 | 46094 |
| 1242.834 | 388 | 1258.741 | 490 | 965.7603 | 370 | 918.9741 | 367 | CREB1 | ECC-1 | Endometri | GSM10108 | 8782 | 46095 |
| 338.2639 | 133 | 416.4108 | 164 | 213.6638 | 161 | 169.6423 | 158 | REST | ECC-1 | Endometri | GSM10108 | 12955 | 46096 |
| 866.1533 | 378 | 870.9725 | 466 | 672.1889 | 450 | 648.8808 | 481 | EP300 | ECC-1 | Endometri | GSM10107 | 29192 | 46097 |
| 938.6155 | 469 | 965.5407 | 604 | 695.7615 | 508 | 667.0497 | 537 | USF1 | ECC-1 | Endometri | GSM10108 | 31946 | 46098 |
| 897.0538 | 397 | 939.5304 | 531 | 715.9159 | 489 | 710.8686 | 551 | NR3C1 | ECC-1 | Endometri | GSM80334 | 28913 | 46099 |
| 813.1505 | 339 | 831.167 | 432 | 628.172 | 413 | 607.6234 | 442 | TAL1 | None | Bone Marr | GSM65154 | 31359 | 4610 |
| 523.0833 | 161 | 827.461 | 226 | 397.5704 | 206 | 480.9816 | 241 | ESR1 | ECC-1 | Endometri | GSM10107 | 12657 | 46100 |
| 797.3607 | 422 | 793.2336 | 504 | 627.9088 | 536 | 645.2748 | 640 | NFIC | ECC-1 | Endometri | GSM10108 | 42180 | 46101 |
| 1085.988 | 462 | 1107.835 | 588 | 812.8961 | 449 | 775.4467 | 463 | ZBTB7A | ECC-1 | Endometri | GSM10107 | 17763 | 46102 |
| 600.295 | 246 | 751.9844 | 317 | 486.2236 | 336 | 616.4106 | 406 | ESR1 | ECC-1 | Endometri | GSM80342 | 27141 | 46103 |
| 1070.041 | 627 | 1094.345 | 784 | 800.0312 | 665 | 763.4943 | 700 | YY1 | ECC-1 | Endometri | GSM10107 | 32277 | 46104 |
| 1254.367 | 569 | 1299.263 | 740 | 937.493 | 516 | 882.9353 | 509 | TAF1 | ECC-1 | Endometri | GSM10107 | 13471 | 46106 |
| 838.2888 | 478 | 848.5773 | 594 | 646.3053 | 588 | 632.2723 | 646 | TEAD4 | ECC-1 | Endometri | GSM10108 | 45170 | 46107 |
| 1186.238 | 550 | 1219.944 | 705 | 875.3095 | 510 | 826.703 | 509 | TAF1 | GM12878 | Blood | GSM80353 | 17233 | 46108 |
| 314.1961 | 277 | 359.7685 | 332 | 169.9286 | 363 | 143.9762 | 379 | BATF | GM12878 | Blood | GSM80353 | 58789 | 46109 |
| 806.6733 | 232 | 862.3162 | 298 | 504.6995 | 271 | 446.6813 | 276 | TAL1 | None | Bone Marr | GSM65154 | 17612 | 4611 |
| 189.0828 | 109 | 233.8733 | 135 | 89.80666 | 126 | 103.3514 | 144 | NFATC1 | GM12878 | Blood | GSM10107 | 15603 | 46110 |
| 650.87 | 314 | 682.7623 | 372 | 254.0431 | 332 | 341.5607 | 401 | NFIC | GM12878 | Blood | GSM10108 | 39246 | 46111 |
| 738.4583 | 329 | 733.8158 | 395 | 360.92 | 358 | 398.1141 | 403 | ATF2 | GM12878 | Blood | GSM10107 | 36118 | 46112 |
| 1034.459 | 346 | 1045.249 | 428 | 802.4157 | 364 | 757.7863 | 368 | REST | GM12878 | Blood | GSM80334 | 15000 | 46113 |
| 1139.482 | 244 | 1154.756 | 310 | 882.6665 | 236 | 846.3146 | 241 | SRF | GM12878 | Blood | GSM80335 | 7149 | 46114 |
| 909.7642 | 483 | 916.6459 | 595 | 643.7021 | 490 | 617.0675 | 522 | POU2F2 | GM12878 | Blood | GSM80353 | 35482 | 46115 |
| 858.8489 | 453 | 905.1614 | 605 | 629.1427 | 500 | 596.3685 | 525 | PAX5 | GM12878 | Blood | GSM80336 | 39165 | 46116 |
| 1065.223 | 471 | 1086.679 | 596 | 788.2218 | 464 | 754.7636 | 485 | ETS1 | GM12878 | Blood | GSM80351 | 21113 | 46117 |
| 644.6275 | 404 | 679.9535 | 536 | 458.9357 | 549 | 324.5289 | 536 | SPI1 | GM12878 | Blood | GSM80353 | 69452 | 46119 |
| 976.6434 | 591 | 1014.067 | 765 | 684.5324 | 596 | 669.4925 | 657 | TCF12 | GM12878 | Blood | GSM80338 | 40919 | 46120 |
| 946.0636 | 418 | 982.1956 | 548 | 677.1304 | 411 | 638.9538 | 421 | PBX3 | GM12878 | Blood | GSM80353 | 26547 | 46121 |
| 353.1679 | 108 | 557.2902 | 152 | 217.2889 | 125 | 298.2453 | 155 | EP300 | GM12878 | Blood | GSM80338 | 8272 | 46122 |
| 827.1328 | 450 | 836.521 | 561 | 575.9561 | 474 | 550.0949 | 506 | MEF2A | GM12878 | Blood | GSM80351 | 43966 | 46123 |
| 1038.239 | 418 | 1062.229 | 537 | 774.3945 | 422 | 735.907 | 434 | USF1 | GM12878 | Blood | GSM80334 | 20012 | 46124 |
| 1021.158 | 251 | 1020.85 | 305 | 636.5337 | 238 | 595.0269 | 246 | RXRA | GM12878 | Blood | GSM80334 | 10954 | 46125 |
| 1122.7 | 605 | 1160.223 | 782 | 826.1486 | 605 | 774.9186 | 609 | EGR1 | GM12878 | Blood | GSM80343 | 25977 | 46126 |
| 1062.182 | 691 | 1081.893 | 857 | 758.7168 | 706 | 717.6749 | 731 | ELF1 | GM12878 | Blood | GSM80349 | 39973 | 46127 |
| 781.9547 | 394 | 779.2275 | 476 | 524.1023 | 439 | 550.2362 | 490 | IRF4 | GM12878 | Blood | GSM80339 | 42175 | 46128 |
| 817.402 | 538 | 830.9028 | 673 | 571.236 | 593 | 566.7488 | 671 | EBF1 | GM12878 | Blood | GSM80338 | 59170 | 46129 |
| 770.915 | 391 | 786.691 | 496 | 508.3146 | 441 | 487.4237 | 471 | BCL3 | GM12878 | Blood | GSM80334 | 43434 | 46130 |
| 750.8159 | 604 | 766.9915 | 750 | 530.5754 | 716 | 508.4843 | 780 | RUNX3 | GM12878 | Blood | GSM10108 | 84048 | 46131 |
| 1182.61 | 331 | 1206.822 | 430 | 904.358 | 313 | 880.5893 | 330 | EGR1 | GM12878 | Blood | GSM10107 | 8962 | 46132 |
| 1105.348 | 287 | 1069.884 | 317 | 820.3655 | 256 | 749.7394 | 259 | ATF3 | GM12878 | Blood | GSM80350 | 9577 | 46133 |
| 825.343 | 407 | 816.718 | 483 | 556.4347 | 422 | 575.5133 | 475 | FOXM1 | GM12878 | Blood | GSM10107 | 36453 | 46134 |
| 917.4659 | 563 | 944.0087 | 716 | 671.2644 | 621 | 625.18 | 631 | SP1 | GM12878 | Blood | GSM80336 | 44941 | 46135 |
| 1243.757 | 506 | 1238.525 | 606 | 923.2786 | 453 | 881.749 | 465 | GABPA | GM12878 | Blood | GSM80335 | 12979 | 46136 |
| 946.8668 | 575 | 979.7174 | 739 | 685.9708 | 615 | 636.8284 | 623 | PAX5 | GM12878 | Blood | GSM80339 | 43156 | 46138 |
| 1084.466 | 488 | 1103.246 | 612 | 824.1508 | 503 | 767.7156 | 495 | ZEB1 | GM12878 | Blood | GSM80341 | 20980 | 46139 |

| | | | | | | | | | | | | | |
|---|---|---|---|---|---|---|---|---|---|---|---|---|---|
| 1000.117 | 551 | 1021.124 | 693 | 712.1774 | 545 | 697.379 | 602 | TCF7L1 | GM12878 | Blood | GSM10107 | 33448 | 46140 |
| 965.9853 | 635 | 961.1782 | 759 | 678.4196 | 656 | 641.8076 | 685 | CREB1 | GM12878 | Blood | GSM10107 | 46678 | 46141 |
| 1156.335 | 336 | 1180.696 | 434 | 843.1008 | 287 | 843.8207 | 328 | SIX5 | GM12878 | Blood | GSM80333 | 9930 | 46142 |
| 1024.517 | 481 | 1026.571 | 586 | 726.6067 | 441 | 708.7909 | 480 | PML | GM12878 | Blood | GSM10107 | 22898 | 46143 |
| 886.2249 | 317 | 893.9975 | 392 | 587.963 | 329 | 587.9335 | 354 | MEF2C | GM12878 | Blood | GSM80342 | 22401 | 46144 |
| 1125.615 | 172 | 1108.251 | 200 | 456.0723 | 138 | 476.1533 | 150 | ZBTB33 | GM12878 | Blood | GSM80339 | 4841 | 46145 |
| 477.7683 | 181 | 622.6147 | 229 | 195.0506 | 182 | 293.551 | 232 | STAT5A | GM12878 | Blood | GSM10107 | 16741 | 46146 |
| 446.1396 | 186 | 457.9291 | 210 | 161.0777 | 181 | 190.0715 | 209 | MTA3 | GM12878 | Blood | GSM10107 | 18353 | 46147 |
| 1005.044 | 495 | 1018.129 | 619 | 723.9832 | 490 | 698.6853 | 522 | SRF | GM12878 | Blood | GSM80347 | 28260 | 46148 |
| 249.4772 | 111 | 406.5402 | 157 | 184.6022 | 146 | 145.3842 | 143 | CEBPB | GM12878 | Blood | GSM10108 | 12195 | 46149 |
| 30.43194 | 28 | 73.86892 | 47 | 37.26805 | 49 | 53.27727 | 61 | TAL1 | None | Bone Marr | GSM61400 | 5948 | 4615 |
| 682.7746 | 406 | 716.79 | 533 | 487.0025 | 523 | 504.0026 | 583 | RAD21 | GM12878 | Blood | GSM80341 | 61667 | 46150 |
| 307.1396 | 208 | 409.144 | 265 | 205.5264 | 280 | 186.7732 | 295 | BCL11A | GM12878 | Blood | GSM80338 | 34943 | 46151 |
| 1076.377 | 625 | 1091.289 | 778 | 769.2941 | 619 | 731.9953 | 647 | YY1 | GM12891 | Blood | GSM80353 | 32999 | 46153 |
| 1197.503 | 491 | 1231.886 | 634 | 888.6346 | 443 | 851.848 | 459 | TAF1 | GM12891 | Blood | GSM80339 | 13895 | 46154 |
| 666.1305 | 479 | 687.8074 | 610 | 493.6644 | 625 | 395.5433 | 630 | SPI1 | GM12891 | Blood | GSM80339 | 82800 | 46156 |
| 863.5757 | 407 | 883.1395 | 516 | 628.7651 | 437 | 588.7983 | 445 | POU2F2 | GM12891 | Blood | GSM80339 | 32565 | 46157 |
| 1012.265 | 440 | 1050.588 | 580 | 738.6714 | 438 | 701.2576 | 447 | PAX5 | GM12891 | Blood | GSM80333 | 23390 | 46158 |
| 17.25528 | 13 | 84.55615 | 32 | 23.92267 | 24 | 78.77514 | 44 | TAL1 | None | Cord blood | GSM60795 | 2540 | 4616 |
| 1158.865 | 653 | 1193.802 | 829 | 825.8514 | 612 | 790.2371 | 642 | PAX5 | GM12892 | Blood | GSM80333 | 26781 | 46160 |
| 1213.868 | 483 | 1246.998 | 626 | 895.2248 | 426 | 879.3148 | 462 | TAF1 | GM12892 | Blood | GSM80348 | 12943 | 46162 |
| 1023.993 | 607 | 1040.632 | 755 | 725.9251 | 599 | 704.373 | 649 | YY1 | GM12892 | Blood | GSM80351 | 35937 | 46163 |
| 688.2474 | 645 | 731.7366 | 835 | 500.3757 | 844 | 477.9288 | 920 | RAD21 | H1 | Embryo | GSM80346 | 122741 | 46164 |
| 961.1919 | 521 | 967.5212 | 638 | 682.8767 | 529 | 672.5797 | 585 | EP300 | H1 | Embryo | GSM80354 | 35718 | 46165 |
| 870.5086 | 477 | 898.5964 | 618 | 627.0815 | 521 | 627.328 | 592 | TCF12 | H1 | Embryo | GSM80342 | 42220 | 46166 |
| 997.9765 | 261 | 977.0945 | 305 | 708.1413 | 266 | 661.1992 | 275 | SRF | H1 | Embryo | GSM80342 | 12366 | 46167 |
| 984.8664 | 391 | 1002.39 | 495 | 740.9171 | 412 | 699.371 | 415 | EGR1 | H1 | Embryo | GSM80343 | 21427 | 46169 |
| 847.4233 | 664 | 870.6882 | 834 | 591.502 | 752 | 571.911 | 824 | YY1 | H1 | Embryo | GSM80351 | 77604 | 46170 |
| 768.9923 | 506 | 796.7035 | 647 | 531.2881 | 566 | 517.871 | 624 | USF1 | H1 | Embryo | GSM80342 | 64634 | 46171 |
| 793.4847 | 321 | 810.3869 | 408 | 486.1384 | 360 | 463.9226 | 382 | REST | H1 | Embryo | GSM80336 | 30926 | 46172 |
| 872.3101 | 468 | 887.7516 | 587 | 629.289 | 506 | 595.4143 | 530 | NANOG | H1 | Embryo | GSM80343 | 40673 | 46173 |
| 1019.001 | 257 | 993.1528 | 292 | 761.9105 | 263 | 704.2169 | 270 | ATF3 | H1 | Embryo | GSM80351 | 11210 | 46174 |
| 685.3384 | 106 | 1107.737 | 154 | 363.7639 | 102 | 441.1657 | 120 | FOSL1 | H1 | Embryo | GSM80338 | 3302 | 46175 |
| 1312.17 | 213 | 1357.348 | 286 | 908.9373 | 171 | 1001.746 | 194 | SP2 | H1 | Embryo | GSM10107 | 3257 | 46176 |
| 952.4255 | 575 | 966.2735 | 716 | 675.0577 | 593 | 634.0633 | 611 | E2F6 | H1 | Embryo | GSM10108 | 40687 | 46177 |
| 79.28756 | 43 | 57.91424 | 42 | 31.65621 | 46 | 58.02545 | 63 | BCL11A | H1 | Embryo | GSM80339 | 5974 | 46178 |
| 1078.76 | 460 | 1090.004 | 569 | 793.4495 | 439 | 753.2404 | 450 | SP4 | H1 | Embryo | GSM10107 | 18910 | 46179 |
| 961.5875 | 762 | 984.4994 | 941 | 678.0657 | 841 | 636.4275 | 884 | MAX | H1 | Embryo | GSM10108 | 68273 | 46180 |
| 849.7451 | 363 | 887.8502 | 487 | 628.7546 | 406 | 514.1383 | 394 | HDAC2 | H1 | Embryo | GSM80334 | 30208 | 46181 |
| 1053.036 | 641 | 1066.687 | 793 | 755.1173 | 650 | 708.3605 | 666 | CREB1 | H1 | Embryo | GSM10108 | 36373 | 46182 |
| 1040.07 | 508 | 1079.776 | 666 | 773.7752 | 521 | 732.9111 | 533 | SIN3A | H1 | Embryo | GSM80342 | 25283 | 46183 |
| 547.5905 | 363 | 662.3404 | 494 | 356.6743 | 489 | 375.7062 | 543 | CTCF | H1 | Embryo | GSM80341 | 65533 | 46184 |
| 954.8404 | 448 | 964.8713 | 552 | 707.3391 | 472 | 663.4272 | 478 | GABPA | H1 | Embryo | GSM80342 | 28132 | 46185 |
| 144.7824 | 45 | 118.5095 | 45 | 38.54839 | 36 | 92.63167 | 57 | BCL11A | H1 | Embryo | GSM80347 | 3561 | 46186 |
| 954.4945 | 150 | 1122.7 | 203 | 454.9719 | 137 | 597.5021 | 168 | SIX5 | H1 | Embryo | GSM80340 | 4776 | 46188 |
| 751.0046 | 507 | 758.9911 | 625 | 518.1526 | 573 | 508.2417 | 640 | TEAD4 | H1 | Embryo | GSM10108 | 68023 | 46189 |
| 926.8813 | 589 | 942.4768 | 734 | 658.0027 | 627 | 633.9681 | 678 | SP1 | H1 | Embryo | GSM80337 | 48091 | 46190 |
| 983.0296 | 556 | 998.6047 | 692 | 702.9315 | 562 | 668.4974 | 587 | JUND | H1 | Embryo | GSM80352 | 35636 | 46191 |
| 610.0695 | 155 | 794.0783 | 197 | 303.5666 | 159 | 350.2984 | 182 | RXRA | H1 | Embryo | GSM80350 | 9922 | 46192 |
| 845.019 | 294 | 805.1466 | 326 | 478.0975 | 307 | 447.4037 | 322 | ATF2 | H1 | Embryo | GSM10107 | 22539 | 46193 |
| 233.7329 | 103 | 211.2668 | 110 | 75.07115 | 98 | 122.3572 | 128 | POU5F1 | H1 | Embryo | GSM80343 | 11797 | 46194 |
| 1122.318 | 514 | 1165.536 | 675 | 827.8153 | 485 | 773.9414 | 482 | TAF7 | H1 | Embryo | GSM80350 | 19061 | 46195 |
| 1157.338 | 731 | 1214.84 | 946 | 825.7179 | 710 | 782.6033 | 734 | TAF1 | H1 | Embryo | GSM80345 | 31694 | 46196 |
| 1181.597 | 448 | 1209.738 | 573 | 905.9384 | 429 | 863.4396 | 433 | TAF1 | H1 | Embryo | GSM10108 | 12031 | 46197 |
| 949.4359 | 554 | 964.392 | 691 | 652.6982 | 535 | 602.8952 | 536 | REST | H1 | Embryo | GSM10108 | 37604 | 46198 |
| 1143.07 | 332 | 1149.668 | 413 | 864.9226 | 314 | 861.7462 | 351 | ZBTB33 | HCT-116 | Colon | GSM80345 | 10298 | 46200 |
| 1085.669 | 539 | 1112.012 | 685 | 811.8367 | 544 | 787.5368 | 583 | ELF1 | HCT-116 | Colon | GSM10107 | 22766 | 46202 |
| 466.82 | 126 | 607.2041 | 160 | 346.57 | 156 | 351.4262 | 168 | REST | HCT-116 | Colon | GSM10108 | 8191 | 46203 |
| 1105.939 | 732 | 1113.747 | 889 | 775.5453 | 724 | 765.7345 | 809 | YY1 | HCT-116 | Colon | GSM80335 | 40648 | 46204 |
| 982.9031 | 576 | 985.1522 | 699 | 698.7497 | 584 | 685.9057 | 646 | USF1 | HCT-116 | Colon | GSM10108 | 37830 | 46205 |
| 895.2119 | 524 | 887.5274 | 619 | 654.7956 | 572 | 645.8494 | 640 | CEBPB | HCT-116 | Colon | GSM10108 | 42061 | 46206 |
| 799.525 | 598 | 837.9997 | 775 | 596.497 | 742 | 580.8159 | 819 | RAD21 | HCT-116 | Colon | GSM10108 | 73058 | 46207 |
| 1148.79 | 645 | 1188.026 | 826 | 848.376 | 649 | 809.6696 | 679 | SIN3A | HCT-116 | Colon | GSM10109 | 26734 | 46208 |
| 928.919 | 709 | 937.5449 | 866 | 635.0331 | 750 | 633.5673 | 854 | CBX3 | HCT-116 | Colon | GSM10107 | 65057 | 46209 |
| 945.2898 | 464 | 937.6696 | 552 | 706.907 | 504 | 698.3723 | 563 | TEAD4 | HCT-116 | Colon | GSM10107 | 31031 | 46210 |
| 1003.776 | 595 | 1029.041 | 748 | 719.1665 | 602 | 710.1881 | 676 | SRF | HCT-116 | Colon | GSM10108 | 36325 | 46211 |
| 857.1742 | 535 | 869.5987 | 665 | 620.1789 | 595 | 620.9725 | 684 | FOSL1 | HCT-116 | Colon | GSM10107 | 49611 | 46212 |
| 895.4843 | 554 | 890.4862 | 663 | 645.6716 | 606 | 640.571 | 681 | JUND | HCT-116 | Colon | GSM10108 | 46236 | 46213 |
| 1081.03 | 645 | 1108.229 | 819 | 778.7774 | 647 | 753.6314 | 696 | EGR1 | HCT-116 | Colon | GSM10108 | 32713 | 46214 |
| 998.4116 | 699 | 1030.832 | 878 | 721.2843 | 753 | 706.6891 | 835 | SP1 | HCT-116 | Colon | GSM10109 | 49266 | 46215 |
| 1013.933 | 723 | 1057.85 | 921 | 735.8147 | 784 | 726.2256 | 876 | MAX | HCT-116 | Colon | GSM10109 | 49162 | 46216 |
| 937.7781 | 431 | 931.7212 | 518 | 710.9667 | 476 | 679.4832 | 499 | ATF3 | HCT-116 | Colon | GSM10107 | 27633 | 46217 |

| | | | | | | | | | | | | |
|---|---|---|---|---|---|---|---|---|---|---|---|---|
| 732.7175 | 399 | 769.7477 | 530 | 556.6368 | 505 | 535.3619 | 543 | CTCF | HCT-116 | Colon | GSM10109 | 49965 | 46218 |
| 1174.814 | 396 | 1181.364 | 487 | 885.2158 | 360 | 849.5158 | 372 | GABPA | HeLa-S3 | Cervix | GSM80345 | 11198 | 46219 |
| 1201.258 | 618 | 1267.012 | 824 | 873.6472 | 568 | 866.2023 | 636 | TAF1 | HeLa-S3 | Cervix | GSM80345 | 20095 | 46221 |
| 771.1439 | 244 | 820.6946 | 305 | 403.266 | 268 | 400.6199 | 288 | REST | HeLa-S3 | Cervix | GSM80347 | 20751 | 46222 |
| 1034.175 | 325 | 1033.659 | 395 | 762.7068 | 311 | 745.7766 | 337 | SRF | HepG2 | Liver | GSM80350 | 14314 | 46223 |
| 999.7783 | 415 | 1039.637 | 552 | 769.927 | 453 | 744.0232 | 479 | ZBTB7A | HepG2 | Liver | GSM10108 | 21639 | 46224 |
| 827.4202 | 468 | 854.2277 | 598 | 650.3282 | 594 | 669.2501 | 707 | EP300 | HepG2 | Liver | GSM80349 | 45523 | 46225 |
| 1132.088 | 691 | 1159.795 | 870 | 832.5252 | 713 | 796.7083 | 753 | YY1 | HepG2 | Liver | GSM80338 | 32187 | 46226 |
| 733.2866 | 533 | 740.8907 | 658 | 569.1021 | 715 | 603.2365 | 879 | FOXA2 | HepG2 | Liver | GSM80340 | 77574 | 46227 |
| 899.5248 | 267 | 908.211 | 338 | 729.4559 | 334 | 643.479 | 324 | TCF12 | HepG2 | Liver | GSM80333 | 17380 | 46228 |
| 882.5987 | 159 | 1065.555 | 220 | 568.2398 | 174 | 538.1876 | 180 | ZBTB33 | HepG2 | Liver | GSM80344 | 6226 | 46229 |
| 1114.086 | 555 | 1158.511 | 727 | 829.5065 | 552 | 797.7822 | 583 | CEBPD | HepG2 | Liver | GSM10107 | 23186 | 46230 |
| 1044.5 | 683 | 1073.207 | 855 | 758.6554 | 715 | 728.3308 | 763 | CREB1 | HepG2 | Liver | GSM10108 | 40374 | 46231 |
| 942.7584 | 580 | 975.5615 | 749 | 709.9832 | 671 | 702.7252 | 753 | JUND | HepG2 | Liver | GSM80350 | 44403 | 46232 |
| 652.9968 | 280 | 752.9599 | 369 | 477.2866 | 371 | 558.8069 | 427 | CEBPB | HepG2 | Liver | GSM10107 | 33684 | 46233 |
| 951.2516 | 585 | 997.9453 | 770 | 723.7148 | 677 | 726.7253 | 781 | SP1 | HepG2 | Liver | GSM80350 | 43012 | 46234 |
| 929.8556 | 551 | 947.1359 | 686 | 705.7589 | 633 | 690.2222 | 700 | HDAC2 | HepG2 | Liver | GSM80349 | 41191 | 46235 |
| 1146.072 | 623 | 1189.565 | 803 | 852.4398 | 617 | 831.5228 | 673 | SIN3A | HepG2 | Liver | GSM80353 | 23770 | 46236 |
| 1192.774 | 586 | 1234.879 | 762 | 879.0723 | 559 | 836.5409 | 570 | GABPA | HepG2 | Liver | GSM80334 | 19675 | 46237 |
| 590.4416 | 253 | 756.2959 | 342 | 483.6589 | 348 | 499.881 | 381 | TEAD4 | HepG2 | Liver | GSM10108 | 28723 | 46238 |
| 777.9409 | 324 | 801.7951 | 418 | 618.3335 | 418 | 623.2204 | 483 | HNF4G | HepG2 | Liver | GSM80340 | 32478 | 46239 |
| 895.1184 | 384 | 882.5827 | 452 | 701.0702 | 451 | 680.3212 | 489 | NFIC | HepG2 | Liver | GSM10107 | 26236 | 46240 |
| 906.6746 | 408 | 905.7137 | 497 | 676.545 | 440 | 671.8353 | 497 | MYBL2 | HepG2 | Liver | GSM10108 | 27313 | 46241 |
| 849.57 | 487 | 863.6774 | 610 | 650.7932 | 591 | 670.2264 | 708 | RXRA | HepG2 | Liver | GSM80345 | 45653 | 46242 |
| 906.1747 | 586 | 906.3948 | 711 | 658.3345 | 654 | 676.8176 | 778 | FOSL2 | HepG2 | Liver | GSM80345 | 51000 | 46243 |
| 1027.709 | 744 | 1056.233 | 927 | 740.3854 | 799 | 712.5882 | 861 | MAX | HepG2 | Liver | GSM10108 | 49688 | 46244 |
| 939.8137 | 434 | 936.2631 | 523 | 726.8817 | 493 | 708.0902 | 535 | ZEB1 | HepG2 | Liver | GSM10108 | 26894 | 46245 |
| 1531.816 | 171 | 1527.687 | 210 | 965.7627 | 125 | 1088.854 | 141 | SP2 | HepG2 | Liver | GSM10107 | 1345 | 46246 |
| 883.4711 | 500 | 896.0981 | 622 | 665.7682 | 581 | 613.6672 | 581 | USF1 | HepG2 | Liver | GSM80352 | 42853 | 46248 |
| 900.3108 | 491 | 906.7969 | 603 | 677.8322 | 555 | 680.5722 | 638 | NR2F2 | HepG2 | Liver | GSM10108 | 37380 | 46249 |
| 659.2175 | 411 | 717.7672 | 570 | 476.6568 | 546 | 487.1926 | 602 | CTCF | HepG2 | Liver | GSM80348 | 67232 | 46250 |
| 621.4332 | 179 | 596.6953 | 194 | 312.9568 | 187 | 341.8341 | 209 | MBD4 | HepG2 | Liver | GSM10107 | 12408 | 46251 |
| 724.9353 | 533 | 744.8881 | 673 | 538.3791 | 680 | 535.2661 | 770 | RAD21 | HepG2 | Liver | GSM80351 | 79478 | 46252 |
| 972.5413 | 382 | 1019.089 | 516 | 754.9648 | 427 | 744.5261 | 474 | REST | HepG2 | Liver | GSM10107 | 20811 | 46253 |
| 1078.668 | 617 | 1113.77 | 790 | 786.3821 | 628 | 750.5308 | 656 | ELF1 | HepG2 | Liver | GSM80341 | 31417 | 46254 |
| 1034.431 | 137 | 1184.772 | 178 | 472.0391 | 118 | 444.3478 | 121 | ZBTB33 | HepG2 | Liver | GSM80341 | 3269 | 46255 |
| 1041.717 | 527 | 1063.752 | 665 | 771.638 | 544 | 724.0792 | 547 | REST | HepG2 | Liver | GSM80334 | 27001 | 46257 |
| 1122.228 | 699 | 1147.257 | 872 | 803.8133 | 690 | 761.7006 | 710 | TAF1 | HepG2 | Liver | GSM80336 | 32991 | 46258 |
| 602.9161 | 480 | 628.8895 | 615 | 383.6142 | 643 | 337.2022 | 680 | SPI1 | HL-60 | Blood | GSM10108 | 101377 | 46259 |
| 933.6708 | 295 | 956.2603 | 384 | 661.0346 | 304 | 500.4557 | 288 | REST | HL-60 | Blood | GSM10107 | 16521 | 46260 |
| 1037.209 | 327 | 1043.821 | 410 | 763.6436 | 310 | 717.1258 | 308 | GABPA | HL-60 | Blood | GSM10108 | 13937 | 46261 |
| 492.9763 | 227 | 634.2688 | 285 | 344.4446 | 294 | 357.9905 | 323 | NR2F2 | K562 | Bone Marr | GSM10107 | 26608 | 46265 |
| 1010.496 | 681 | 1071.991 | 901 | 747.8231 | 762 | 691.3653 | 771 | ELF1 | K562 | Bone Marr | GSM80349 | 47978 | 46266 |
| 1080.514 | 562 | 1128.297 | 738 | 816.6469 | 592 | 756.8113 | 583 | GABPA | K562 | Bone Marr | GSM80352 | 26569 | 46268 |
| 1125.566 | 699 | 1158.865 | 876 | 816.624 | 704 | 782.1866 | 743 | YY1 | K562 | Bone Marr | GSM80344 | 33857 | 46269 |
| 656.7917 | 124 | 846.9807 | 157 | 276.7909 | 110 | 256.1001 | 113 | ZBTB33 | K562 | Bone Marr | GSM80350 | 5155 | 46270 |
| 716.6436 | 412 | 788.4753 | 591 | 554.8709 | 543 | 534.3714 | 587 | CTCF | K562 | Bone Marr | GSM10108 | 56404 | 46271 |
| 711.8319 | 398 | 729.3319 | 510 | 539.124 | 514 | 497.0578 | 539 | SPI1 | K562 | Bone Marr | GSM80338 | 55856 | 46272 |
| 968.6559 | 129 | 1216.217 | 185 | 562.0663 | 127 | 597.7386 | 139 | SIX5 | K562 | Bone Marr | GSM80338 | 3112 | 46273 |
| 1087.453 | 469 | 1111.238 | 604 | 829.7055 | 485 | 784.6781 | 493 | CREB1 | K562 | Bone Marr | GSM10108 | 19021 | 46274 |
| 791.9971 | 531 | 840.8764 | 710 | 600.0327 | 667 | 580.3604 | 728 | RAD21 | K562 | Bone Marr | GSM80344 | 64501 | 46275 |
| 927.0841 | 418 | 910.7673 | 486 | 693.2095 | 454 | 650.8784 | 461 | ATF3 | K562 | Bone Marr | GSM80338 | 28242 | 46276 |
| 1164.751 | 458 | 1181.162 | 570 | 864.3553 | 413 | 840.6086 | 442 | SIN3A | K562 | Bone Marr | GSM80352 | 13484 | 46277 |
| 1126.086 | 491 | 1163.677 | 641 | 847.4436 | 480 | 805.3268 | 493 | ETS1 | K562 | Bone Marr | GSM80344 | 17526 | 46278 |
| 861.3705 | 493 | 859.9844 | 597 | 616.3404 | 537 | 595.1976 | 580 | USF1 | K562 | Bone Marr | GSM80344 | 45845 | 46279 |
| 250.9489 | 125 | 278.3461 | 147 | 177.9502 | 164 | 157.2659 | 169 | STAT5A | K562 | Bone Marr | GSM10108 | 15333 | 46280 |
| 1177.627 | 621 | 1247.215 | 844 | 872.8662 | 612 | 828.0566 | 622 | TAF1 | K562 | Bone Marr | GSM80343 | 22224 | 46282 |
| 428.6758 | 134 | 402.1957 | 144 | 260.0096 | 155 | 254.7367 | 165 | MEF2A | K562 | Bone Marr | GSM80337 | 10653 | 46283 |
| 595.8635 | 333 | 660.1258 | 409 | 489.6944 | 472 | 447.5707 | 497 | TEAD4 | K562 | Bone Marr | GSM10108 | 49107 | 46284 |
| 485.9012 | 284 | 666.7591 | 372 | 358.2934 | 391 | 346.3678 | 420 | CEBPB | K562 | Bone Marr | GSM10108 | 44386 | 46285 |
| 920.622 | 502 | 935.3401 | 625 | 678.0913 | 546 | 651.939 | 581 | REST | K562 | Bone Marr | GSM80344 | 37240 | 46286 |
| 1229.073 | 274 | 1270.254 | 371 | 945.8899 | 249 | 913.9991 | 259 | SP2 | K562 | Bone Marr | GSM80340 | 6073 | 46287 |
| 838.2178 | 164 | 908.1555 | 188 | 403.8292 | 157 | 415.8215 | 170 | BCLAF1 | K562 | Bone Marr | GSM80351 | 7178 | 46288 |
| 1097.376 | 224 | 1094.815 | 271 | 726.1093 | 210 | 775.3115 | 231 | TAF7 | K562 | Bone Marr | GSM80340 | 7053 | 46289 |
| 140.559 | 134 | 196.0327 | 174 | 189.8427 | 239 | 172.314 | 251 | GATA2 | K562 | Bone Marr | GSM80354 | 28403 | 46290 |
| 749.3837 | 368 | 739.5419 | 436 | 519.0087 | 437 | 437.832 | 445 | CBX3 | K562 | Bone Marr | GSM10107 | 40834 | 46291 |
| 932.2928 | 800 | 999.9995 | 1038 | 682.7918 | 961 | 645.4974 | 1031 | MAX | K562 | Bone Marr | GSM80352 | 84727 | 46292 |
| 957.1344 | 384 | 1025.864 | 546 | 745.3624 | 442 | 690.8915 | 434 | CTCFL | K562 | Bone Marr | GSM80340 | 23302 | 46293 |
| 831.6968 | 301 | 832.468 | 368 | 559.2975 | 342 | 511.1093 | 355 | CEBPD | K562 | Bone Marr | GSM10109 | 25043 | 46294 |
| 1083.839 | 663 | 1125.661 | 847 | 779.7082 | 663 | 738.7745 | 685 | ZBTB7A | K562 | Bone Marr | GSM80347 | 33466 | 46295 |
| 457.0895 | 187 | 534.0581 | 225 | 324.9837 | 242 | 346.4477 | 268 | FOSL1 | K562 | Bone Marr | GSM80343 | 20416 | 46296 |
| 1087.562 | 725 | 1177.932 | 980 | 808.3234 | 789 | 739.701 | 780 | E2F6 | K562 | Bone Marr | GSM80346 | 40726 | 46297 |

| | | | | | | | | | | | | | |
|---|---|---|---|---|---|---|---|---|---|---|---|---|---|
| 1013.636 | 713 | 1093.704 | 957 | 739.2958 | 787 | 693.4501 | 815 | EGR1 | K562 | Bone Marr | GSM80341 | 51429 | 46298 |
| 1047.239 | 270 | 1033.73 | 316 | 807.5925 | 277 | 771.2716 | 282 | SRF | K562 | Bone Marr | GSM80352 | 10854 | 46299 |
| 1157.014 | 372 | 1179.406 | 475 | 878.6484 | 351 | 854.8925 | 374 | SP1 | K562 | Bone Marr | GSM80350 | 11120 | 46300 |
| 968.3852 | 386 | 976.8354 | 478 | 720.525 | 393 | 703.2438 | 429 | PML | K562 | Bone Marr | GSM10107 | 20871 | 46301 |
| 1188.507 | 199 | 1178.312 | 235 | 880.3633 | 194 | 808.4646 | 196 | SIX5 | K562 | Bone Marr | GSM80337 | 4672 | 46302 |
| 999.9555 | 682 | 1057.78 | 895 | 726.5359 | 747 | 674.4214 | 761 | YY1 | K562 | Bone Marr | GSM80347 | 49257 | 46303 |
| 764.8989 | 227 | 835.2155 | 279 | 563.3044 | 286 | 423.3647 | 271 | HDAC2 | K562 | Bone Marr | GSM80347 | 17647 | 46304 |
| 580.7161 | 232 | 685.1717 | 280 | 331.0086 | 273 | 359.6235 | 305 | TRIM28 | K562 | Bone Marr | GSM10108 | 24492 | 46305 |
| 1198.033 | 422 | 1213.247 | 529 | 926.5283 | 409 | 855.5942 | 385 | THAP1 | K562 | Bone Marr | GSM80340 | 11317 | 46306 |
| 1212.775 | 490 | 1245.325 | 625 | 951.5387 | 497 | 901.7667 | 503 | PML | MCF-7 | Breast | GSM10108 | 13052 | 46307 |
| 881.5799 | 289 | 881.0655 | 354 | 772.2961 | 425 | 786.2745 | 505 | JUND | MCF-7 | Breast | GSM10108 | 20147 | 46308 |
| 887.1599 | 579 | 889.1101 | 703 | 765.2476 | 841 | 781.9122 | 992 | NR2F2 | MCF-7 | Breast | GSM10108 | 52370 | 46309 |
| 1232.651 | 386 | 1270.485 | 505 | 955.5898 | 365 | 921.0807 | 375 | KAT5 | None | Blood | GSM39395 | 9222 | 4631 |
| 875.2155 | 708 | 836.4992 | 803 | 718.5142 | 986 | 745.4391 | 1179 | GATA3 | MCF-7 | Breast | GSM10107 | 79994 | 46310 |
| 953.4658 | 316 | 952.5451 | 384 | 829.4515 | 447 | 816.771 | 493 | FOXM1 | MCF-7 | Breast | GSM10107 | 17645 | 46311 |
| 1122.681 | 465 | 1137.91 | 576 | 873.5351 | 484 | 848.2647 | 515 | SRF | MCF-7 | Breast | GSM10108 | 16593 | 46312 |
| 863.2784 | 238 | 855.7208 | 287 | 730.3495 | 329 | 695.9268 | 338 | REST | MCF-7 | Breast | GSM10108 | 16650 | 46313 |
| 1167.987 | 564 | 1187.972 | 705 | 922.926 | 615 | 896.5749 | 662 | GABPA | MCF-7 | Breast | GSM10108 | 19167 | 46314 |
| 831.9701 | 229 | 858.6713 | 280 | 774.7464 | 359 | 758.8976 | 392 | TEAD4 | MCF-7 | Breast | GSM10108 | 16286 | 46315 |
| 727.4324 | 484 | 723.8689 | 584 | 615.4486 | 722 | 617.8381 | 833 | CEBPB | MCF-7 | Breast | GSM10108 | 68717 | 46316 |
| 735.1057 | 284 | 726.2809 | 322 | 635.2189 | 410 | 658.7154 | 497 | FOSL2 | MCF-7 | Breast | GSM10107 | 30409 | 46317 |
| 467.1956 | 326 | 641.1382 | 442 | 312.5026 | 445 | 341.5588 | 500 | CTCF | MCF-7 | Breast | GSM10107 | 60273 | 46318 |
| 1078.258 | 333 | 1118.146 | 445 | 858.5842 | 368 | 823.821 | 380 | EGR1 | MCF-7 | Breast | GSM10108 | 12424 | 46319 |
| 732.5818 | 108 | 1109.806 | 151 | 387.7179 | 103 | 526.8848 | 129 | KAT5 | None | Blood | GSM39394 | 3141 | 4632 |
| 1228.565 | 397 | 1257.364 | 508 | 934.3034 | 357 | 897.9324 | 367 | TAF1 | MCF-7 | Breast | GSM10108 | 9011 | 46320 |
| 952.2415 | 299 | 951.1654 | 366 | 833.6253 | 433 | 859.4548 | 525 | TCF12 | MCF-7 | Breast | GSM10108 | 16684 | 46321 |
| 850.525 | 249 | 847.0271 | 299 | 778.551 | 404 | 802.1637 | 488 | EP300 | MCF-7 | Breast | GSM10108 | 18379 | 46322 |
| 1114.356 | 655 | 1123.619 | 807 | 870.0931 | 753 | 847.6131 | 817 | ELF1 | MCF-7 | Breast | GSM10107 | 30859 | 46323 |
| 1143.631 | 866 | 1154.969 | 1047 | 848.2591 | 983 | 821.1341 | 1072 | MAX | MCF-7 | Breast | GSM10108 | 51336 | 46324 |
| 630.374 | 348 | 668.5857 | 433 | 551.9048 | 521 | 532.8976 | 568 | RAD21 | MCF-7 | Breast | GSM10107 | 54305 | 46325 |
| 948.3999 | 568 | 950.4024 | 693 | 778.8825 | 750 | 787.7485 | 872 | HDAC2 | MCF-7 | Breast | GSM10108 | 41407 | 46326 |
| 1061.433 | 723 | 1082.813 | 893 | 838.3831 | 877 | 838.7143 | 1005 | SIN3A | MCF-7 | Breast | GSM10108 | 43273 | 46327 |
| 1093.406 | 579 | 1105.968 | 713 | 808.1307 | 586 | 748.1979 | 577 | REST | PANC-1 | Pancreas | GSM80337 | 26776 | 46328 |
| 1246.535 | 509 | 1286.55 | 662 | 952.7908 | 478 | 913.8623 | 494 | SIN3A | PANC-1 | Pancreas | GSM10107 | 12298 | 46329 |
| 1065.755 | 269 | 1105.122 | 365 | 867.8226 | 310 | 818.7309 | 307 | REST | PANC-1 | Pancreas | GSM10107 | 10097 | 46330 |
| 415.9398 | 117 | 530.607 | 147 | 344.4933 | 153 | 306.8418 | 155 | REST | PANC-1 | Pancreas | GSM10108 | 7832 | 46331 |
| 1112.999 | 609 | 1150.854 | 777 | 816.4474 | 606 | 795.421 | 660 | FOXP2 | PFSK-1 | Brain | GSM80348 | 27999 | 46333 |
| 1196.612 | 582 | 1210.45 | 725 | 859.773 | 518 | 841.5079 | 565 | TAF1 | PFSK-1 | Brain | GSM10107 | 18583 | 46334 |
| 1037.855 | 508 | 1087.769 | 672 | 789.8789 | 542 | 756.0637 | 571 | SIN3A | PFSK-1 | Brain | GSM10107 | 25480 | 46335 |
| 897.4473 | 496 | 931.6835 | 643 | 670.4343 | 561 | 639.7909 | 591 | REST | PFSK-1 | Brain | GSM10107 | 39518 | 46337 |
| 1017.543 | 604 | 1067.159 | 791 | 761.9101 | 665 | 744.117 | 725 | REST | PFSK-1 | Brain | GSM80336 | 37097 | 46338 |
| 1008.782 | 545 | 1034.991 | 697 | 740.8294 | 572 | 692.7712 | 578 | FOXP2 | SK-N-MC | Brain | GSM80335 | 33008 | 46340 |
| 1037.147 | 709 | 1059.877 | 879 | 752.0867 | 757 | 717.8822 | 803 | REST | SK-N-SH | Brain | GSM80333 | 46197 | 46341 |
| 1016.419 | 545 | 1027.724 | 679 | 739.3489 | 554 | 712.6085 | 594 | YY1 | SK-N-SH | Brain | GSM10108 | 30596 | 46342 |
| 756.4876 | 466 | 737.2922 | 543 | 596.022 | 619 | 584.5752 | 687 | NFIC | SK-N-SH | Brain | GSM10108 | 56843 | 46343 |
| 1116.908 | 468 | 1128.739 | 578 | 837.7645 | 456 | 802.5494 | 474 | GABPA | SK-N-SH | Brain | GSM10107 | 16920 | 46344 |
| 1187.126 | 613 | 1217.317 | 775 | 865.7768 | 572 | 838.4302 | 606 | TAF1 | SK-N-SH | Brain | GSM10107 | 20644 | 46345 |
| 529.6769 | 340 | 522.3018 | 383 | 350.5133 | 457 | 377.7316 | 511 | FOSL2 | SK-N-SH | Brain | GSM10108 | 58538 | 46346 |
| 989.6034 | 429 | 997.1015 | 531 | 730.7985 | 435 | 693.3301 | 450 | USF1 | SK-N-SH | Brain | GSM10108 | 23707 | 46347 |
| 686.4492 | 305 | 726.2173 | 384 | 523.8986 | 411 | 487.7296 | 433 | TEAD4 | SK-N-SH | Brain | GSM10108 | 37058 | 46348 |
| 1033.873 | 770 | 1056.071 | 954 | 742.6288 | 839 | 713.9271 | 908 | MAX | SK-N-SH | Brain | GSM10108 | 54858 | 46349 |
| 884.4383 | 251 | 881.7824 | 307 | 476.5568 | 260 | 414.8709 | 263 | FOXM1 | SK-N-SH | Brain | GSM10107 | 16797 | 46350 |
| 324.6679 | 252 | 351.3805 | 295 | 263.8342 | 368 | 310.9014 | 424 | JUND | SK-N-SH | Brain | GSM10109 | 47066 | 46351 |
| 455.8743 | 174 | 490.4659 | 201 | 310.4124 | 218 | 288.6822 | 228 | MEF2A | SK-N-SH | Brain | GSM10108 | 17158 | 46352 |
| 829.8504 | 487 | 809.7451 | 567 | 619.1078 | 571 | 587.3341 | 604 | EP300 | SK-N-SH | Brain | GSM10107 | 47613 | 46354 |
| 1090.495 | 467 | 1120.253 | 599 | 827.4433 | 472 | 786.4895 | 483 | ELF1 | SK-N-SH | Brain | GSM10107 | 18760 | 46355 |
| 657.2216 | 255 | 739.709 | 304 | 455.7212 | 324 | 499.0291 | 363 | RXRA | SK-N-SH | Brain | GSM10107 | 26388 | 46356 |
| 1010.853 | 405 | 1026.428 | 505 | 745.197 | 398 | 732.0965 | 440 | ZBTB33 | SK-N-SH | Brain | GSM10107 | 20095 | 46357 |
| 811.1024 | 626 | 815.8577 | 758 | 603.8831 | 770 | 557.4077 | 795 | PBX3 | SK-N-SH | Brain | GSM10108 | 75617 | 46358 |
| 924.4629 | 458 | 936.6335 | 573 | 705.9601 | 523 | 679.2027 | 557 | REST | SK-N-SH | Brain | GSM10107 | 32153 | 46359 |
| 1043.586 | 567 | 1084.386 | 732 | 765.9015 | 577 | 754.4472 | 640 | SIN3A | SK-N-SH | Brain | GSM10108 | 29488 | 46360 |
| 603.4929 | 478 | 588.2082 | 560 | 486.5791 | 702 | 488.9771 | 807 | GATA3 | SK-N-SH | Brain | GSM10107 | 95559 | 46361 |
| 746.7221 | 602 | 751.0275 | 735 | 564.9311 | 783 | 547.7941 | 863 | TCF12 | SK-N-SH | Brain | GSM10109 | 86717 | 46362 |
| 819.0096 | 736 | 836.3561 | 905 | 610.8957 | 942 | 605.6631 | 1066 | EP300 | SK-N-SH_R | Brain | GSM80349 | 103335 | 46363 |
| 1002.724 | 677 | 1015.312 | 832 | 721.0793 | 723 | 697.7241 | 782 | YY1 | SK-N-SH_R | Brain | GSM80349 | 47258 | 46364 |
| 1010.162 | 672 | 1028.954 | 834 | 711.9397 | 700 | 701.1655 | 780 | USF1 | SK-N-SH_R | Brain | GSM80347 | 48071 | 46365 |
| 706.0885 | 534 | 750.4417 | 706 | 559.1435 | 746 | 557.2093 | 848 | RAD21 | SK-N-SH_R | Brain | GSM80349 | 85174 | 46366 |
| 746.0504 | 415 | 803.409 | 573 | 574.6603 | 536 | 553.4879 | 578 | CTCF | SK-N-SH_R | Brain | GSM80333 | 52010 | 46367 |
| 1111.555 | 1002 | 1095.502 | 1177 | 811.776 | 1243 | 799.7332 | 1404 | FOXA1 | T47D | Mammary | GSM80340 | 98475 | 46368 |
| 439.9887 | 121 | 422.6454 | 131 | 660.4659 | 213 | 832.8523 | 264 | ESR1 | T47D | Mammary | GSM10108 | 8165 | 46369 |
| 1013.57 | 464 | 1003.76 | 548 | 853.8323 | 614 | 849.5528 | 693 | EP300 | T47D | Mammary | GSM80352 | 24569 | 46370 |
| 884.3351 | 290 | 816.7937 | 297 | 775.8177 | 433 | 801.5348 | 525 | GATA3 | T47D | Mammary | GSM80351 | 20530 | 46371 |

| | | | | | | | | | | | | | |
|---|---|---|---|---|---|---|---|---|---|---|---|---|---|
| 402.9321 | 78 | 441.0478 | 90 | 613.6939 | 137 | 766.5749 | 164 | JUND | T47D | Mammary | GSM10107 | 3358 | 46372 |
| 529.651 | 256 | 714.7881 | 337 | 326.9362 | 322 | 376.273 | 368 | CTCF | T47D | Mammary | GSM80334 | 33799 | 46373 |
| 846.6203 | 290 | 811.986 | 322 | 742.8875 | 438 | 744.5952 | 501 | ESR1 | T47D | Mammary | GSM80353 | 22824 | 46374 |
| 895.1646 | 278 | 837.1977 | 292 | 770.0557 | 397 | 759.3347 | 437 | ESR1 | T47D | Mammary | GSM80337 | 18477 | 46375 |
| 1043.11 | 686 | 1054.095 | 840 | 741.6563 | 703 | 709.2828 | 747 | REST | U87 | Brain | GSM80337 | 42766 | 46377 |
| 438.5255 | 90 | 464.2212 | 102 | 198.474 | 84 | 173.3607 | 84 | TAL1 | None | None | GSM10386 | 4238 | 47077 |
| 78.87388 | 51 | 125.0517 | 71 | 87.32661 | 83 | 65.21867 | 80 | TAL1 | None | None | GSM10386 | 8082 | 47078 |
| 905.3621 | 181 | 1000.296 | 237 | 540.5905 | 195 | 494.1571 | 199 | TAL1 | None | None | GSM10386 | 8582 | 47079 |
| 89.69688 | 46 | 102.2778 | 55 | 82.65683 | 68 | 59.22822 | 64 | TAL1 | None | None | GSM10386 | 5996 | 47080 |
| 161.6181 | 31 | 308.9086 | 50 | 127.4973 | 38 | 224.332 | 55 | HIF1A | None | None | GSM10568 | 1332 | 47084 |
| 79.59022 | 19 | 60.75283 | 18 | 24.6843 | 15 | 145.0413 | 40 | EPAS1 | None | None | GSM10568 | 1125 | 47088 |
| 85.16041 | 64 | 172.6887 | 99 | 148.1201 | 126 | 184.9274 | 149 | TCF4 | CAL-1 | None | GSM10727 | 11363 | 47101 |
| 960.002 | 322 | 975.1245 | 408 | 758.5992 | 373 | 745.831 | 406 | MYC | HeLa | Cervix | GSM10886 | 17896 | 47102 |
| 413.5469 | 60 | 531.2518 | 76 | 194.1641 | 53 | 327.2642 | 75 | ZBTB17 | HeLa | Cervix | GSM10886 | 1649 | 47103 |
| 114.5109 | 113 | 157.084 | 146 | 154.2197 | 203 | 167.2579 | 227 | TP63 | None | Skin | GSM43977 | 25027 | 4718 |
| 32.22674 | 70 | 24.73267 | 75 | 46.76999 | 134 | 58.61673 | 157 | TP63 | None | Skin | GSM43977 | 24025 | 4719 |
| 1023.748 | 651 | 1041.957 | 804 | 774.4506 | 736 | 716.6284 | 736 | KDM5B | MCF-7 | Breast | GSM11226 | 41047 | 47265 |
| 1076.776 | 700 | 1094.451 | 862 | 806.768 | 769 | 755.3558 | 785 | KDM5B | MCF-7 | Breast | GSM11226 | 39188 | 47266 |
| 1132.035 | 793 | 1143.025 | 965 | 857.6703 | 901 | 831.1511 | 979 | KDM5B | MCF-7 | Breast | GSM11226 | 42343 | 47267 |
| 1079.563 | 885 | 1123.675 | 1102 | 792.7109 | 1033 | 768.8489 | 1137 | KDM5B | T47D | Mammary | GSM11226 | 67730 | 47273 |
| 962.1984 | 477 | 950.1833 | 562 | 717.4036 | 515 | 691.16 | 549 | KDM5B | SUM185PE | Mammary | GSM11226 | 31215 | 47275 |
| 1121.209 | 856 | 1122.005 | 1021 | 832.1893 | 972 | 788.3938 | 1032 | KDM5B | SUM185PE | Mammary | GSM11226 | 52622 | 47276 |
| 877.9247 | 476 | 902.2183 | 609 | 618.1014 | 495 | 617.4859 | 566 | KDM5B | SUM159PT | Mammary | GSM11226 | 41248 | 47278 |
| 870.1125 | 489 | 893.4946 | 619 | 636.2001 | 535 | 604.9488 | 563 | CTCF | SUM159PT | Mammary | GSM11226 | 40372 | 47281 |
| 1044.222 | 737 | 1081.373 | 930 | 748.0513 | 779 | 706.1347 | 815 | KDM5B | MDA-MB-2 | Breast | GSM11226 | 48404 | 47283 |
| 1053.987 | 860 | 1102.197 | 1081 | 776.0762 | 1007 | 744.9284 | 1093 | KDM5B | HCC2157 | Mammary | GSM11226 | 69154 | 47285 |
| 643.5953 | 253 | 758.4572 | 319 | 375.7805 | 300 | 432.4522 | 343 | FOSL1 | BT-549 | Mammary | GSM11252 | 26322 | 47287 |
| 810.6739 | 276 | 818.6244 | 346 | 495.1317 | 318 | 582.2652 | 367 | JUN | BT-549 | Mammary | GSM11252 | 23698 | 47288 |
| 781.6738 | 287 | 811.5714 | 377 | 572.611 | 365 | 639.0445 | 442 | AR | DU145 | Prostate | GSM11641 | 28306 | 47323 |
| 904.8743 | 513 | 904.8403 | 626 | 637.1012 | 533 | 616.313 | 578 | FOXA1 | DU145 | Prostate | GSM11641 | 42556 | 47324 |
| 664.4133 | 151 | 963.0854 | 207 | 287.8449 | 142 | 505.9439 | 199 | FOXA1 | DU145 | Prostate | GSM11641 | 8356 | 47325 |
| 977.3574 | 222 | 1019.166 | 303 | 794.7813 | 266 | 809.1822 | 314 | FOXA1 | DU145 | Prostate | GSM11641 | 10844 | 47326 |
| 128.5143 | 270 | 108.5771 | 299 | 31.3937 | 338 | 42.33746 | 397 | FOXA1 | DU145 | Prostate | GSM11641 | 95690 | 47327 |
| 48.71298 | 241 | 25.90586 | 249 | 5.431808 | 311 | 13.76536 | 391 | FOXA1 | DU145 | Prostate | GSM11641 | 117878 | 47328 |
| 128.24 | 323 | 116.4362 | 367 | 44.5641 | 446 | 39.33584 | 485 | FOXA1 | DU145 | Prostate | GSM11641 | 125267 | 47329 |
| 80.01418 | 236 | 62.62698 | 259 | 32.77088 | 341 | 37.19783 | 388 | FOXA1 | DU145 | Prostate | GSM11641 | 95838 | 47330 |
| 365.5746 | 57 | 708.1938 | 93 | 258.3403 | 64 | 371.7182 | 83 | FOXA1 | DU145 | Prostate | GSM11641 | 1783 | 47332 |
| 696.75 | 354 | 709.0066 | 448 | 455.3326 | 451 | 533.8688 | 528 | AR | DU145 | Prostate | GSM11641 | 49314 | 47333 |
| 318.8747 | 232 | 330.2731 | 267 | 157.5241 | 287 | 197.4572 | 337 | FOXA1 | DU145 | Prostate | GSM11641 | 42809 | 47334 |
| 86.60268 | 21 | 79.04554 | 22 | 61.03123 | 25 | 191.124 | 49 | ERG | VCaP | Prostate | GSM11936 | 1242 | 47407 |
| 1033.021 | 258 | 999.8201 | 291 | 948.5689 | 410 | 989.5279 | 517 | ERG | VCaP | Prostate | GSM11936 | 10653 | 47408 |
| 936.043 | 280 | 919.4788 | 328 | 844.4746 | 436 | 884.5227 | 553 | ERG | VCaP | Prostate | GSM11936 | 16279 | 47409 |
| 1175.522 | 300 | 1205.797 | 390 | 934.6254 | 308 | 899.0955 | 315 | GABPA | VCaP | Prostate | GSM11936 | 7870 | 47410 |
| 1186.745 | 606 | 1218.647 | 770 | 952.0121 | 692 | 919.629 | 736 | GABPA | VCaP | Prostate | GSM11936 | 21062 | 47411 |
| 1166.048 | 571 | 1208.995 | 738 | 910.8514 | 611 | 898.5146 | 675 | GABPA | VCaP | Prostate | GSM11936 | 20049 | 47412 |
| 966.5591 | 551 | 1015.345 | 727 | 707.3377 | 586 | 658.5291 | 590 | GABPA | Jurkat | Blood | GSM11936 | 36809 | 47413 |
| 954.9946 | 499 | 990.9412 | 650 | 694.4673 | 523 | 641.3295 | 518 | GABPA | Jurkat | Blood | GSM11936 | 33462 | 47414 |
| 369.1295 | 84 | 380.7909 | 94 | 126.9772 | 70 | 161.7034 | 84 | ERG | Jurkat | Blood | GSM11936 | 4456 | 47415 |
| 1069.977 | 397 | 1066.148 | 478 | 767.4666 | 353 | 721.6519 | 354 | ERG | Jurkat | Blood | GSM11936 | 16058 | 47416 |
| 396.2403 | 130 | 382.8632 | 142 | 632.4501 | 238 | 786.7703 | 297 | GATA6 | LS174T | Colon | GSM11973 | 10944 | 47431 |
| 208.6774 | 81 | 230.9837 | 95 | 517.3652 | 186 | 768.4849 | 242 | GATA6 | LS174T | Colon | GSM11973 | 8028 | 47432 |
| 212.1854 | 76 | 189.5805 | 80 | 497.2669 | 170 | 806.6788 | 232 | GATA6 | LS174T | Colon | GSM11973 | 6938 | 47433 |
| 1027.896 | 211 | 1049.147 | 274 | 612.9647 | 209 | 642.0057 | 228 | REST | None | Blood | GSM12019 | 8459 | 47435 |
| 336.6624 | 179 | 493.2454 | 239 | 255.4948 | 246 | 307.4095 | 286 | CTCF | K562 | Bone Marr | GSM12246 | 25100 | 47474 |
| 560.887 | 315 | 730.006 | 465 | 427.3584 | 438 | 468.5394 | 492 | CTCF | K562 | Bone Marr | GSM12246 | 48705 | 47475 |
| 501.5611 | 257 | 720.8648 | 359 | 349.6121 | 341 | 422.52 | 397 | CTCF | K562 | Bone Marr | GSM12246 | 35709 | 47476 |
| 543.6547 | 280 | 730.4209 | 400 | 364.6389 | 367 | 448.5689 | 430 | CTCF | K562 | Bone Marr | GSM12246 | 39524 | 47477 |
| 1073.184 | 414 | 1075.462 | 512 | 798.5528 | 409 | 765.5463 | 425 | MYC | U2OS | Bone | GSM12315 | 17710 | 47487 |
| 1107.896 | 170 | 1108.591 | 210 | 718.9805 | 180 | 651.0149 | 181 | MYC | U2OS | Bone | GSM12315 | 5163 | 47488 |
| 117.9164 | 74 | 146.3297 | 92 | 79.61936 | 99 | 113.8431 | 123 | ZBTB17 | U2OS | Bone | GSM12315 | 11599 | 47489 |
| 234.0418 | 113 | 284.9837 | 139 | 88.81283 | 117 | 160.721 | 159 | ZBTB17 | U2OS | Bone | GSM12316 | 14235 | 47490 |
| 331.2804 | 44 | 477.6346 | 60 | 169.0374 | 40 | 242.982 | 52 | MYC | P493-6 | None | GSM12344 | 1055 | 47549 |
| 728.4805 | 137 | 1061.915 | 200 | 417.3124 | 142 | 432.6561 | 154 | MYC | P493-6 | None | GSM12345 | 5693 | 47550 |
| 1042.195 | 534 | 1051.662 | 657 | 759.8091 | 527 | 712.298 | 535 | MYC | P493-6 | None | GSM12345 | 26793 | 47551 |
| 847.4497 | 397 | 856.9591 | 493 | 673.1351 | 495 | 657.3765 | 545 | SMAD3 | NCI-H441 | Lung | GSM12467 | 32728 | 47576 |
| 783.4731 | 437 | 805.8349 | 558 | 630.6776 | 583 | 614.8646 | 642 | SMAD3 | NCI-H441 | Lung | GSM12467 | 46348 | 47577 |
| 766.5372 | 548 | 789.4715 | 693 | 603.3586 | 729 | 627.3442 | 876 | TTF1 | NCI-H441 | Lung | GSM12467 | 68599 | 47578 |
| 708.0929 | 400 | 720.4281 | 503 | 565.9173 | 546 | 569.0637 | 629 | TTF1 | NCI-H441 | Lung | GSM12467 | 52097 | 47579 |
| 23.27647 | 13 | 21.02341 | 14 | 19.93453 | 19 | 138.2058 | 51 | SMAD4 | NCI-H441 | Lung | GSM12467 | 1953 | 47581 |
| 104.2214 | 65 | 116.301 | 77 | 105.0777 | 101 | 198.3564 | 142 | SMAD4 | NCI-H441 | Lung | GSM12467 | 9765 | 47582 |
| 612.5966 | 171 | 716.6666 | 205 | 553.8686 | 233 | 690.7785 | 277 | SMAD3 | A549 | Lung | GSM12467 | 11462 | 47584 |
| 420.1365 | 90 | 557.7162 | 116 | 270.5479 | 101 | 335.6217 | 120 | KLF5 | YCC3 | None | GSM12508 | 4501 | 47593 |

| | | | | | | | | | | | | | |
|---|---|---|---|---|---|---|---|---|---|---|---|---|---|
| 85.23018 | 52 | 133.201 | 72 | 300.0493 | 141 | 479.6426 | 189 | GATA4 | YCC3 | None | GSM12508 | 7854 | 47594 |
| 803.3527 | 332 | 810.1613 | 412 | 682.4151 | 480 | 713.8176 | 590 | GATA6 | YCC3 | None | GSM12508 | 31448 | 47595 |
| 197.0088 | 42 | 250.825 | 53 | 44.38367 | 28 | 197.7958 | 63 | KLF5 | AGS | Stomach | GSM12508 | 2079 | 47597 |
| 935.5 | 817 | 918.6123 | 958 | 693.8459 | 1011 | 689.2424 | 1147 | GATA4 | AGS | Stomach | GSM12508 | 90048 | 47598 |
| 200.7818 | 153 | 234.7355 | 185 | 296.1082 | 279 | 360.7882 | 326 | GATA6 | AGS | Stomach | GSM12508 | 27968 | 47599 |
| 1115.298 | 564 | 1146.376 | 723 | 835.2029 | 571 | 818.4764 | 629 | KLF5 | KATO III | Stomach | GSM12508 | 23722 | 47601 |
| 273.829 | 81 | 299.3138 | 94 | 301.1404 | 123 | 267.4416 | 124 | GATA4 | KATO III | Stomach | GSM12509 | 5949 | 47602 |
| 113.8243 | 57 | 154.4506 | 74 | 275.8009 | 130 | 403.9893 | 167 | GATA6 | KATO III | Stomach | GSM12509 | 7276 | 47603 |
| 0.472386 | 13 | 0.987872 | 18 | 0.17769 | 24 | 11.43252 | 55 | SOX11 | GRANTA-5 | None | GSM12603 | 10949 | 47710 |
| 8.080777 | 11 | 32.41909 | 23 | 2.345896 | 13 | 35.48531 | 36 | SOX11 | GRANTA-5 | None | GSM12603 | 3364 | 47711 |
| 80.76241 | 93 | 202.1777 | 153 | 154.9285 | 191 | 134.2653 | 197 | LDB1 | None | Blood | GSM12731 | 22360 | 47774 |
| 874.2028 | 707 | 907.6437 | 898 | 617.2021 | 815 | 575.1514 | 854 | CTCF | None | Blood | GSM12731 | 81344 | 47776 |
| 64.06417 | 37 | 128.4569 | 58 | 44.2593 | 49 | 57.72917 | 59 | REST | GM12878 | Blood | GSM12901 | 5207 | 47847 |
| 79.5416 | 43 | 112.8174 | 57 | 56.45393 | 57 | 82.49192 | 72 | REST | GM12878 | Blood | GSM12901 | 5662 | 47848 |
| 59.52086 | 36 | 101.3213 | 52 | 55.68369 | 54 | 62.00617 | 61 | REST | GM12878 | Blood | GSM12901 | 5163 | 47849 |
| 33.00574 | 21 | 78.96553 | 36 | 11.84931 | 22 | 39.03096 | 38 | REST | GM12878 | Blood | GSM12901 | 3264 | 47850 |
| 29.62988 | 22 | 94.80482 | 43 | 25.86108 | 33 | 32.16511 | 39 | REST | GM12878 | Blood | GSM12901 | 3872 | 47851 |
| 33.55846 | 23 | 97.02651 | 43 | 17.56685 | 28 | 37.94586 | 41 | REST | GM12878 | Blood | GSM12901 | 3735 | 47852 |
| 8.429008 | 8 | 24.39007 | 15 | 9.561428 | 14 | 65.19731 | 35 | REST | GM12878 | Blood | GSM12901 | 1946 | 47853 |
| 18.61901 | 12 | 37.13704 | 19 | 10.36518 | 15 | 57.57481 | 34 | REST | GM12878 | Blood | GSM12901 | 2057 | 47854 |
| 9.251845 | 9 | 18.22771 | 14 | 3.005571 | 10 | 47.02657 | 32 | REST | GM12878 | Blood | GSM12901 | 2164 | 47855 |
| 26.64433 | 21 | 65.63158 | 36 | 27.71947 | 34 | 55.89787 | 49 | REST | GM12878 | Blood | GSM12901 | 3905 | 47856 |
| 29.09805 | 22 | 57.36975 | 34 | 27.10906 | 34 | 54.83337 | 49 | REST | GM12878 | Blood | GSM12901 | 3979 | 47857 |
| 10.98238 | 9 | 31.7526 | 17 | 5.123081 | 11 | 69.55883 | 36 | REST | GM12878 | Blood | GSM12901 | 1940 | 47858 |
| 28.68825 | 17 | 38.12157 | 22 | 9.303631 | 17 | 45.30978 | 35 | REST | Jurkat | Blood | GSM12901 | 2621 | 47859 |
| 28.94559 | 20 | 60.55744 | 32 | 25.33412 | 30 | 50.07094 | 43 | REST | Jurkat | Blood | GSM12901 | 3446 | 47860 |
| 5.206506 | 6 | 23.92219 | 14 | 6.176243 | 11 | 73.0153 | 35 | REST | Jurkat | Blood | GSM12901 | 1765 | 47861 |
| 25.88655 | 17 | 45.04772 | 25 | 21.76988 | 25 | 40.38559 | 35 | REST | Jurkat | Blood | GSM12901 | 2844 | 47862 |
| 31.65735 | 27 | 67.92218 | 43 | 27.54161 | 41 | 52.37985 | 57 | REST | Jurkat | Blood | GSM12901 | 5248 | 47863 |
| 10.57131 | 15 | 46.02451 | 32 | 25.86422 | 35 | 49.29828 | 49 | REST | Jurkat | Blood | GSM12901 | 4287 | 47864 |
| 26.38849 | 10 | 82.50405 | 21 | 13.45897 | 11 | 36.32529 | 19 | REST | Jurkat | Blood | GSM12901 | 1078 | 47865 |
| 21.71459 | 19 | 48.62704 | 31 | 17.06024 | 28 | 51.62406 | 47 | REST | Jurkat | Blood | GSM12901 | 3871 | 47866 |
| 17.80777 | 18 | 66.72662 | 37 | 24.16972 | 33 | 45.56113 | 46 | REST | Jurkat | Blood | GSM12901 | 4059 | 47867 |
| 15.50489 | 10 | 31.32124 | 16 | 7.738168 | 12 | 39.21326 | 26 | REST | Jurkat | Blood | GSM12901 | 1759 | 47868 |
| 42.60972 | 28 | 63.47028 | 38 | 27.92198 | 37 | 51.17372 | 51 | REST | Jurkat | Blood | GSM12901 | 4440 | 47869 |
| 44.57383 | 30 | 90.18465 | 47 | 35.65685 | 43 | 61.5932 | 58 | REST | Jurkat | Blood | GSM12901 | 4863 | 47870 |
| 38.02461 | 22 | 68.81254 | 33 | 27.89291 | 30 | 57.4079 | 44 | REST | Jurkat | Blood | GSM12901 | 3215 | 47871 |
| 19.59224 | 14 | 25.61553 | 18 | 14.87721 | 20 | 38.17108 | 32 | REST | Jurkat | Blood | GSM12901 | 2541 | 47872 |
| 21.09972 | 13 | 32.05292 | 18 | 8.181182 | 14 | 30.54724 | 26 | REST | Jurkat | Blood | GSM12901 | 2123 | 47873 |
| 56.03675 | 49 | 51.66834 | 54 | 49.37578 | 75 | 66.78888 | 91 | EP300 | GM12878 | Blood | GSM12901 | 9768 | 47875 |
| 25.66077 | 29 | 21.32597 | 31 | 29.11775 | 50 | 45.57106 | 64 | EP300 | GM12878 | Blood | GSM12901 | 7052 | 47876 |
| 6.31095 | 16 | 12.95806 | 24 | 12.61071 | 35 | 46.06183 | 60 | EP300 | GM12878 | Blood | GSM12901 | 6317 | 47884 |
| 5.554258 | 10 | 3.585179 | 10 | 5.022602 | 17 | 36.52603 | 38 | EP300 | GM12878 | Blood | GSM12901 | 3531 | 47885 |
| 785.2023 | 395 | 803.021 | 505 | 598.8463 | 487 | 534.9064 | 478 | EP300 | GM12878 | Blood | GSM12901 | 42090 | 47886 |
| 794.3326 | 445 | 825.1763 | 580 | 593.8771 | 534 | 515.3457 | 508 | EP300 | GM12878 | Blood | GSM12901 | 48123 | 47887 |
| 513.0862 | 284 | 707.4851 | 400 | 356.5293 | 380 | 368.9849 | 418 | CTCF | T47D-MTV | Mammary | GSM12940 | 42283 | 47896 |
| 544.5984 | 297 | 725.6358 | 429 | 382.5705 | 398 | 383.8713 | 433 | CTCF | T47D-MTV | Mammary | GSM12940 | 43583 | 47897 |
| 974.8237 | 247 | 996.4271 | 317 | 752.7393 | 273 | 712.5353 | 284 | BRD4 | None | Umbilical V | GSM13052 | 11910 | 48060 |
| 986.0357 | 292 | 995.4341 | 370 | 718.048 | 290 | 721.3864 | 315 | BRD4 | None | Umbilical V | GSM13052 | 13953 | 48061 |
| 462.9398 | 119 | 440.923 | 128 | 262.3514 | 129 | 261.8215 | 138 | BRD4 | None | Umbilical V | GSM13052 | 7425 | 48062 |
| 438.1774 | 54 | 319.155 | 48 | 260.7271 | 53 | 329.1825 | 64 | RELA | None | Umbilical V | GSM13052 | 1136 | 48069 |
| 515.3811 | 190 | 725.478 | 250 | 423.0373 | 257 | 406.8338 | 272 | RELA | None | Umbilical V | GSM13052 | 18210 | 48070 |
| 610.4322 | 227 | 780.3215 | 294 | 435.2477 | 290 | 359.1805 | 289 | RELA | None | Umbilical V | GSM13052 | 22364 | 48071 |
| 161.4984 | 278 | 248.8775 | 368 | 272.4716 | 538 | 400.3466 | 661 | AR | HEK293 | Embryonic | GSM13082 | 88395 | 48085 |
| 163.4529 | 258 | 206.9002 | 321 | 226.651 | 471 | 362.1918 | 594 | AR | HEK293 | Embryonic | GSM13082 | 77745 | 48086 |
| 184.4748 | 229 | 184.6954 | 263 | 241.3964 | 407 | 356.1965 | 504 | AR | HEK293 | Embryonic | GSM13082 | 59528 | 48088 |
| 172.5498 | 305 | 228.1095 | 384 | 254.8952 | 567 | 413.9943 | 715 | AR | HEK293 | Embryonic | GSM13082 | 98916 | 48089 |
| 99.11993 | 22 | 90.67778 | 23 | 60.57152 | 24 | 160.2621 | 43 | AR | PC-3 | Prostate | GSM13082 | 1156 | 48091 |
| 280.643 | 188 | 269.3122 | 209 | 306.1722 | 301 | 477.1604 | 387 | AR | PC-3 | Prostate | GSM13082 | 30483 | 48092 |
| 315.9296 | 261 | 319.271 | 298 | 359.1545 | 430 | 524.8895 | 543 | AR | PC-3 | Prostate | GSM13082 | 51324 | 48093 |
| 361.4144 | 253 | 346.884 | 282 | 367.7493 | 396 | 519.4866 | 488 | AR | PC-3 | Prostate | GSM13082 | 43794 | 48096 |
| 297.6051 | 207 | 322.065 | 242 | 351.7932 | 343 | 512.2585 | 429 | AR | PC-3 | Prostate | GSM13082 | 35256 | 48097 |
| 27.13441 | 25 | 62.48626 | 41 | 47.08777 | 50 | 55.98334 | 58 | MBD2 | MCF-7 | Breast | GSM13222 | 5196 | 48127 |
| 509.5299 | 62 | 337.6503 | 52 | 246.6907 | 54 | 335.3662 | 68 | ESR1 | MCF-7 | Breast | GSM13252 | 1283 | 48167 |
| 233.8421 | 60 | 245.5007 | 68 | 526.3754 | 132 | 858.1948 | 183 | ESR1 | MCF-7 | Breast | GSM13252 | 3743 | 48168 |
| 303.5056 | 83 | 336.8126 | 97 | 595.8455 | 169 | 867.0097 | 219 | ESR1 | MCF-7 | Breast | GSM13252 | 5601 | 48169 |
| 616.0958 | 189 | 536.971 | 196 | 771.1568 | 331 | 817.4537 | 431 | ESR1 | MCF-7 | Breast | GSM13252 | 15159 | 48170 |
| 333.4634 | 116 | 478.9911 | 155 | 515.2667 | 210 | 654.9798 | 252 | ESR1 | MCF-7 | Breast | GSM13252 | 10436 | 48171 |
| 96.37853 | 25 | 80.34748 | 25 | 101.1786 | 37 | 196.5679 | 56 | ESR1 | MCF-7 | Breast | GSM13252 | 1637 | 48172 |
| 106.272 | 34 | 91.78733 | 35 | 49.77648 | 35 | 148.452 | 63 | EBF1 | None | Adipose | GSM13262 | 2826 | 48179 |
| 1028.201 | 246 | 1023.962 | 300 | 877.7499 | 327 | 950.2689 | 457 | AR | LNCaP | Prostate | GSM13278 | 10561 | 48182 |
| 952.0953 | 311 | 956.7127 | 384 | 855.7398 | 479 | 923.1183 | 646 | AR | LNCaP | Prostate | GSM13278 | 17934 | 48183 |

| | | | | | | | | | | | | |
|---|---|---|---|---|---|---|---|---|---|---|---|---|
| 193.0064 | 145 | 212.5608 | 171 | 134.5355 | 200 | 138.7568 | 220 | TEAD1 | RD | Muscle | GSM13312 | 25768 | 48240 |
| 1266.392 | 160 | 1257.643 | 190 | 668.2123 | 135 | 757.1456 | 153 | WDR5 | LNCaP | Prostate | GSM13333 | 2880 | 48250 |
| 1196.723 | 343 | 1224.144 | 446 | 923.7923 | 322 | 884.7846 | 330 | CEBPA | GM12878 | Blood | GSM13339 | 8607 | 48263 |
| 52.12972 | 33 | 98.41579 | 50 | 36.464 | 44 | 34.98267 | 47 | REST | GM12878 | Blood | GSM13339 | 4749 | 48267 |
| 84.18753 | 79 | 91.39455 | 93 | 54.42434 | 109 | 57.33696 | 121 | EP300 | GM12878 | Blood | GSM13339 | 16236 | 48268 |
| 25.45802 | 49 | 34.56922 | 63 | 24.84966 | 83 | 37.6701 | 103 | EP300 | GM12878 | Blood | GSM13339 | 15725 | 48269 |
| 867.9631 | 374 | 878.87 | 467 | 643.2597 | 409 | 602.837 | 418 | EP300 | GM12878 | Blood | GSM13339 | 29060 | 48270 |
| 900.951 | 383 | 895.0451 | 459 | 662.2926 | 407 | 621.5966 | 412 | EP300 | GM12878 | Blood | GSM13339 | 26964 | 48271 |
| 763.2601 | 481 | 785.8508 | 610 | 577.1536 | 598 | 505.8488 | 572 | EP300 | GM12878 | Blood | GSM13339 | 57997 | 48272 |
| 192.9335 | 160 | 276.1461 | 210 | 175.6224 | 245 | 126.7027 | 239 | EP300 | GM12878 | Blood | GSM13339 | 30788 | 48273 |
| 202.3655 | 161 | 394.8177 | 241 | 180.299 | 244 | 154.9998 | 252 | EP300 | GM12878 | Blood | GSM13339 | 30254 | 48274 |
| 12.60445 | 25 | 23.03298 | 36 | 34.40511 | 60 | 32.36409 | 64 | SPI1 | GM12878 | Blood | GSM13339 | 8544 | 48275 |
| 1115.577 | 446 | 1122.084 | 549 | 828.9587 | 423 | 777.3192 | 421 | CEBPA | GM12878 | Blood | GSM13339 | 16085 | 48277 |
| 1166.132 | 313 | 1169.135 | 383 | 887.8875 | 286 | 843.2542 | 287 | CEBPA | GM12878 | Blood | GSM13339 | 8333 | 48278 |
| 1166.73 | 410 | 1173.887 | 500 | 856.4455 | 357 | 819.2654 | 365 | CEBPA | GM12878 | Blood | GSM13339 | 11844 | 48279 |
| 1110.988 | 340 | 1122.715 | 424 | 844.4486 | 322 | 796.2958 | 320 | CEBPA | GM12878 | Blood | GSM13339 | 10943 | 48280 |
| 964.4335 | 205 | 962.9256 | 250 | 532.1319 | 211 | 357.2958 | 187 | REST | GM12878 | Blood | GSM13339 | 9778 | 48293 |
| 17.97127 | 19 | 44.38412 | 32 | 10.92691 | 26 | 47.8497 | 49 | REST | GM12878 | Blood | GSM13339 | 4253 | 48294 |
| 940.6266 | 212 | 939.1198 | 257 | 483.6424 | 212 | 396.5478 | 207 | REST | GM12878 | Blood | GSM13339 | 10751 | 48295 |
| 44.39074 | 35 | 45.20032 | 40 | 21.19694 | 42 | 42.95193 | 59 | REST | GM12878 | Blood | GSM13339 | 6076 | 48296 |
| 375.8795 | 113 | 435.1666 | 135 | 227.5673 | 129 | 190.7097 | 128 | REST | GM12878 | Blood | GSM13339 | 8279 | 48297 |
| 35.02264 | 28 | 95.35158 | 50 | 40.7517 | 47 | 40.83963 | 51 | REST | GM12878 | Blood | GSM13339 | 5027 | 48298 |
| 91.18322 | 56 | 104.4572 | 67 | 44.82693 | 65 | 47.14597 | 72 | REST | GM12878 | Blood | GSM13339 | 7755 | 48299 |
| 40.93138 | 36 | 65.54475 | 50 | 38.65146 | 56 | 71.70188 | 77 | REST | GM12878 | Blood | GSM13339 | 6817 | 48300 |
| 20.40819 | 21 | 62.51485 | 39 | 27.58599 | 38 | 47.51738 | 51 | REST | GM12878 | Blood | GSM13339 | 4506 | 48301 |
| 100.5324 | 59 | 116.7144 | 71 | 59.37814 | 73 | 65.0684 | 82 | REST | GM12878 | Blood | GSM13339 | 7943 | 48302 |
| 6.965915 | 10 | 52.56113 | 28 | 14.14895 | 22 | 47.5492 | 39 | REST | GM12878 | Blood | GSM13339 | 2961 | 48303 |
| 75.20861 | 53 | 128.6344 | 76 | 55.37258 | 73 | 54.81831 | 79 | REST | GM12878 | Blood | GSM13339 | 8257 | 48304 |
| 120.9263 | 65 | 176.3652 | 87 | 84.68911 | 85 | 87.09228 | 93 | REST | GM12878 | Blood | GSM13339 | 8128 | 48305 |
| 36.06057 | 24 | 90.92283 | 42 | 18.64536 | 29 | 48.80459 | 46 | REST | GM12878 | Blood | GSM13339 | 3837 | 48306 |
| 17.79428 | 13 | 37.61913 | 21 | 9.130119 | 16 | 20.78049 | 24 | REST | GM12878 | Blood | GSM13339 | 2391 | 48307 |
| 90.63206 | 38 | 156.5982 | 56 | 56.1791 | 46 | 96.98813 | 63 | REST | GM12878 | Blood | GSM13340 | 3991 | 48308 |
| 461.4703 | 73 | 456.3213 | 79 | 200.3511 | 63 | 206.3829 | 68 | EP300 | GM12878 | Blood | GSM13340 | 2305 | 48309 |
| 509.9067 | 89 | 581.8484 | 105 | 202.5867 | 75 | 218.353 | 83 | EP300 | GM12878 | Blood | GSM13340 | 3261 | 48311 |
| 478.4134 | 296 | 624.2249 | 373 | 341.3279 | 405 | 257.6146 | 401 | SPI1 | GM12878 | Blood | GSM13340 | 48205 | 48313 |
| 215.6033 | 228 | 260.426 | 279 | 168.1441 | 339 | 118.496 | 335 | SPI1 | GM12878 | Blood | GSM13340 | 52257 | 48314 |
| 357.978 | 254 | 536.8488 | 339 | 282.9618 | 365 | 175.5956 | 340 | SPI1 | GM12878 | Blood | GSM13340 | 45096 | 48315 |
| 28.81239 | 55 | 51.76405 | 78 | 60.56488 | 118 | 39.96425 | 113 | SPI1 | GM12878 | Blood | GSM13340 | 17431 | 48316 |
| 1134.653 | 281 | 1135.256 | 342 | 863.4433 | 260 | 815.5602 | 257 | ZBTB33 | GM12878 | Blood | GSM13340 | 8301 | 48317 |
| 785.62 | 115 | 871.5858 | 133 | 285.5445 | 89 | 237.3949 | 86 | ZBTB33 | GM12878 | Blood | GSM13340 | 3146 | 48318 |
| 1159.374 | 193 | 1157.542 | 235 | 651.1408 | 169 | 493.7098 | 155 | ZBTB33 | GM12878 | Blood | GSM13340 | 4920 | 48319 |
| 637.2607 | 97 | 779.1944 | 119 | 246.3019 | 78 | 224.3035 | 79 | ZBTB33 | GM12878 | Blood | GSM13340 | 2806 | 48320 |
| 627.8734 | 255 | 750.2237 | 323 | 272.7084 | 273 | 265.7362 | 293 | SNAI2 | None | None | GSM13361 | 28714 | 48331 |
| 559.8716 | 235 | 746.581 | 306 | 303.6179 | 275 | 380.574 | 325 | SNAI2 | None | None | GSM13361 | 27042 | 48332 |
| 102.3461 | 270 | 149.3197 | 350 | 73.04759 | 436 | 76.39488 | 487 | SNAI2 | None | None | GSM13361 | 110190 | 48333 |
| 107.1653 | 346 | 191.2853 | 470 | 90.18462 | 588 | 126.2382 | 698 | SNAI2 | None | None | GSM13361 | 157474 | 48334 |
| 666.0444 | 251 | 794.7875 | 340 | 411.72 | 304 | 555.0806 | 371 | SNAI2 | None | None | GSM13361 | 25732 | 48337 |
| 741.7972 | 221 | 855.2851 | 288 | 436.5641 | 252 | 455.508 | 276 | SNAI2 | None | None | GSM13361 | 17212 | 48338 |
| 685.0272 | 551 | 708.2383 | 696 | 483.7636 | 680 | 471.9943 | 758 | SNAI2 | None | None | GSM13361 | 94328 | 48339 |
| 705.1557 | 564 | 715.3782 | 699 | 493.2879 | 683 | 493.4084 | 780 | SNAI2 | None | None | GSM13361 | 91783 | 48340 |
| 346.9346 | 93 | 506.437 | 126 | 155.7132 | 91 | 369.7734 | 147 | SNAI2 | None | None | GSM13361 | 6102 | 48341 |
| 336.7108 | 75 | 466.0205 | 99 | 139.8962 | 68 | 357.037 | 116 | SNAI2 | None | None | GSM13361 | 3882 | 48342 |
| 770.2464 | 506 | 782.2945 | 630 | 538.711 | 573 | 520.8035 | 625 | SNAI2 | None | None | GSM13361 | 63046 | 48343 |
| 738.7734 | 519 | 742.045 | 632 | 514.4342 | 599 | 495.4229 | 655 | SNAI2 | None | None | GSM13361 | 72160 | 48344 |
| 1293.189 | 167 | 1292.147 | 208 | 616.9054 | 127 | 665.4517 | 140 | PRKDC | None | None | GSM13400 | 2746 | 48353 |
| 1049.226 | 195 | 1053.716 | 242 | 490.9849 | 173 | 462.3659 | 179 | PRKDC | None | None | GSM13400 | 7229 | 48354 |
| 306.7508 | 97 | 422.7951 | 127 | 166.2281 | 106 | 159.8364 | 112 | PCGF2 | None | None | GSM13400 | 7464 | 48355 |
| 109.7636 | 46 | 135.3229 | 57 | 50.26914 | 49 | 62.68832 | 58 | PCGF2 | None | None | GSM13400 | 4790 | 48356 |
| 620.2464 | 144 | 630.252 | 160 | 365.2744 | 157 | 308.7479 | 155 | KDM6B | BJ | Skin | GSM13483 | 8080 | 48442 |
| 836.3424 | 316 | 808.65 | 361 | 439.1846 | 317 | 615.1568 | 394 | KDM6B | BJ | Skin | GSM13483 | 26054 | 48443 |
| 327.3603 | 82 | 317.9287 | 89 | 116.8058 | 71 | 148.352 | 85 | TP53 | BJ | Skin | GSM13483 | 4898 | 48445 |
| 1090.645 | 154 | 1132.054 | 186 | 692.7806 | 161 | 609.2357 | 159 | BRD4 | MCF-7 | Breast | GSM13485 | 4090 | 48448 |
| 1151.564 | 165 | 1201.469 | 228 | 788.9466 | 174 | 756.9701 | 180 | BRD4 | MCF-7 | Breast | GSM13485 | 4123 | 48450 |
| 981.8645 | 135 | 1183.335 | 184 | 558.9683 | 132 | 690.4382 | 157 | BRD4 | MCF-7 | Breast | GSM13485 | 3388 | 48451 |
| 63.64256 | 19 | 63.14897 | 21 | 121.7156 | 39 | 308.4021 | 69 | ESR1 | MCF-7 | Breast | GSM13485 | 1479 | 48453 |
| 82.28202 | 23 | 30.08224 | 15 | 122.1077 | 41 | 285.1788 | 69 | ESR1 | MCF-7 | Breast | GSM13485 | 1632 | 48454 |
| 108.8701 | 37 | 93.29431 | 38 | 207.4038 | 75 | 409.7213 | 114 | ESR1 | MCF-7 | Breast | GSM13485 | 3163 | 48455 |
| 57.16045 | 20 | 37.73009 | 18 | 140.9651 | 47 | 299.6332 | 75 | ESR1 | MCF-7 | Breast | GSM13485 | 1849 | 48457 |
| 35.17274 | 12 | 24.32224 | 11 | 48.64823 | 21 | 186.7787 | 46 | ESR1 | MCF-7 | Breast | GSM13485 | 1097 | 48458 |
| 530.5931 | 178 | 766.6706 | 238 | 337.3791 | 214 | 382.4409 | 243 | CTCF | BEAS-2B | Lung | GSM13544 | 15954 | 48481 |
| 711.944 | 254 | 813.0817 | 343 | 457.4355 | 311 | 466.023 | 338 | CTCF | BEAS-2B | Lung | GSM13544 | 25061 | 48482 |
| 300.321 | 138 | 363.0973 | 169 | 154.3666 | 159 | 80.85421 | 136 | POU5F1 | None | None | GSM13640 | 16563 | 48550 |

| | | | | | | | | | | | | | |
|---|---|---|---|---|---|---|---|---|---|---|---|---|---|
| 611.9918 | 300 | 693.52 | 373 | 202.3132 | 304 | 222.4506 | 342 | SOX2 | None | None | GSM13640 | 40791 | 48551 |
| 292.6221 | 82 | 351.3507 | 100 | 169.1754 | 91 | 167.22 | 97 | KLF4 | None | None | GSM13640 | 5725 | 48552 |
| 1057.334 | 633 | 1095.223 | 811 | 770.6237 | 655 | 720.646 | 662 | KDM6B | CUTLL1 | Blood | GSM13669 | 35321 | 48576 |
| 145.5272 | 53 | 186.251 | 67 | 51.85432 | 50 | 129.2782 | 80 | AGO1 | MCF-7 | Breast | GSM13699 | 5046 | 48594 |
| 3.621727 | 13 | 12.23513 | 23 | 9.064251 | 31 | 27.80694 | 49 | CBX5 | MCF-7 | Breast | GSM13699 | 6266 | 48598 |
| 145.5272 | 53 | 186.251 | 67 | 51.85432 | 50 | 129.2782 | 80 | AGO1 | None | None | GSM13699 | 5046 | 48601 |
| 15.60187 | 14 | 23.01407 | 19 | 12.30783 | 21 | 75.03335 | 48 | BATF | OCI-Ly3 | Bone Marr | GSM13702 | 3118 | 48621 |
| 148.5851 | 320 | 242.828 | 429 | 147.8914 | 548 | 121.1653 | 578 | IRF4 | OCI-Ly3 | Bone Marr | GSM13702 | 121300 | 48623 |
| 24.43127 | 157 | 16.69699 | 172 | 20.95664 | 281 | 27.84205 | 329 | SPI1 | OCI-Ly3 | Bone Marr | GSM13702 | 85241 | 48624 |
| 345.2366 | 383 | 328.5931 | 433 | 217.4143 | 549 | 139.8741 | 540 | SPIB | OCI-Ly3 | Bone Marr | GSM13702 | 103759 | 48625 |
| 99.8599 | 62 | 123.5079 | 77 | 86.19026 | 91 | 76.38019 | 94 | BATF | OCI-Ly10 | Bone Marr | GSM13702 | 9566 | 48626 |
| 481.1968 | 163 | 585.6532 | 200 | 331.6669 | 203 | 270.3708 | 200 | IRF4 | OCI-Ly10 | Bone Marr | GSM13702 | 14635 | 48628 |
| 45.20197 | 103 | 50.51307 | 124 | 47.02586 | 180 | 36.08331 | 185 | SPI1 | OCI-Ly10 | Bone Marr | GSM13702 | 37044 | 48629 |
| 78.08576 | 170 | 65.84044 | 188 | 58.98347 | 272 | 66.68639 | 309 | SPIB | OCI-Ly10 | Bone Marr | GSM13702 | 60741 | 48630 |
| 1017.08 | 413 | 1018.552 | 509 | 745.4073 | 407 | 710.9071 | 421 | IRF4 | H929 | Bone Marr | GSM13702 | 21258 | 48632 |
| 863.9279 | 358 | 886.1916 | 465 | 642.1906 | 401 | 604.4846 | 411 | SPI1 | H929 | Bone Marr | GSM13702 | 29243 | 48633 |
| 794.0465 | 444 | 800.4921 | 546 | 453.3576 | 448 | 462.0313 | 490 | SPIB | H929 | Bone Marr | GSM13702 | 49882 | 48634 |
| 394.7413 | 128 | 399.044 | 143 | 228.1533 | 146 | 186.5901 | 144 | IRF4 | OCI-Ly3 | Bone Marr | GSM13702 | 10716 | 48636 |
| 36.49614 | 29 | 53.49984 | 39 | 71.30777 | 61 | 45.65442 | 55 | SPI1 | OCI-Ly3 | Bone Marr | GSM13702 | 5607 | 48637 |
| 307.6671 | 214 | 407.1549 | 272 | 294.9585 | 330 | 220.8954 | 324 | SPIB | OCI-Ly3 | Bone Marr | GSM13702 | 37792 | 48638 |
| 220.5326 | 87 | 309.9856 | 115 | 132.6577 | 104 | 141.0889 | 115 | IRF4 | OCI-Ly3 | Bone Marr | GSM13702 | 8940 | 48640 |
| 30.31787 | 24 | 20.06598 | 23 | 35.39163 | 41 | 40.84945 | 47 | SPI1 | OCI-Ly3 | Bone Marr | GSM13702 | 4676 | 48641 |
| 8.018192 | 54 | 8.01492 | 64 | 6.731852 | 97 | 12.23991 | 121 | SPIB | OCI-Ly3 | Bone Marr | GSM13702 | 29254 | 48642 |
| 15.86442 | 14 | 26.21626 | 20 | 6.979652 | 17 | 8.585168 | 20 | H3F3B | None | None | GSM13752 | 3076 | 48692 |
| 65.3104 | 19 | 34.62642 | 15 | 63.06249 | 27 | 82.18811 | 33 | SPI1 | None | Liver | GSM13775 | 1349 | 48704 |
| 40.19087 | 20 | 42.20111 | 23 | 80.60842 | 42 | 84.39729 | 46 | SPI1 | None | Liver | GSM13775 | 2467 | 48706 |
| 1202.329 | 541 | 1228.216 | 678 | 901.4412 | 497 | 859.3961 | 512 | CBX8 | PC-3 | Prostate | GSM13838 | 13916 | 48755 |
| 633.3753 | 349 | 717.2291 | 486 | 442.6426 | 467 | 482.8439 | 524 | CTCF | PC-3 | Prostate | GSM13838 | 51719 | 48757 |
| 914.0067 | 232 | 881.0924 | 260 | 600.161 | 260 | 521.2528 | 261 | EZH2 | NTKO | None | GSM13857 | 12996 | 48763 |
| 656.9398 | 193 | 677.5765 | 217 | 379.1014 | 215 | 331.6452 | 218 | EZH2 | None | None | GSM13857 | 13673 | 48765 |
| 1070.049 | 291 | 1125.574 | 406 | 835.1976 | 303 | 789.374 | 301 | MYC | P493-6 | None | GSM13863 | 10826 | 48778 |
| 1065.002 | 301 | 1063.846 | 367 | 795.0952 | 285 | 757.0247 | 291 | MYC | P493-6 | None | GSM13863 | 11572 | 48779 |
| 759.8468 | 347 | 762.677 | 427 | 596.0009 | 447 | 583.9996 | 496 | ZNF750 | None | Foreskin | GSM13870 | 38361 | 48799 |
| 904.7919 | 392 | 929.0124 | 508 | 688.4708 | 448 | 690.3189 | 512 | KLF4 | None | Foreskin | GSM13870 | 28601 | 48800 |
| 443.0759 | 263 | 603.0967 | 338 | 324.8867 | 362 | 321.5819 | 392 | RCOR1 | None | Foreskin | GSM13870 | 41460 | 48801 |
| 154.3722 | 95 | 177.3566 | 114 | 132.722 | 139 | 141.0177 | 154 | KDM1A | None | Foreskin | GSM13870 | 14614 | 48802 |
| 34.06082 | 20 | 49.60087 | 27 | 26.42293 | 28 | 48.15543 | 39 | MBD2 | MCF-7 | Breast | GSM13881 | 2990 | 48822 |
| 1329.116 | 677 | 1366.975 | 863 | 1007.492 | 663 | 983.7676 | 713 | MBD2 | MCF-7 | Breast | GSM13881 | 16248 | 48824 |
| 1320.305 | 691 | 1352.02 | 873 | 946.8587 | 598 | 921.4801 | 636 | MBD2 | MCF-7 | Breast | GSM13881 | 16121 | 48825 |
| 1245.956 | 294 | 1266.972 | 377 | 941.8292 | 250 | 956.4966 | 293 | MBD2 | MCF-7 | Breast | GSM13881 | 6033 | 48826 |
| 1298.545 | 661 | 1335.618 | 842 | 954.1598 | 597 | 920.7228 | 628 | MBD2 | MCF-7 | Breast | GSM13881 | 15592 | 48827 |
| 114.4972 | 209 | 145.7872 | 261 | 63.1033 | 303 | 77.19521 | 351 | TRIM28 | H1 | Embryo | GSM13992 | 67991 | 48923 |
| 0.153822 | 22 | -0.180125 | 17 | 0.118625 | 44 | 0.003018 | 44 | TRIM28 | H1 | Embryo | GSM13992 | 19044 | 48930 |
| 1056.483 | 324 | 1071.642 | 411 | 807.7698 | 329 | 785.3273 | 348 | AFF4 | HeLa | Cervix | GSM14050 | 13086 | 48993 |
| 1153.374 | 626 | 1190.174 | 801 | 848.9852 | 614 | 827.5409 | 668 | AFF4 | HeLa | Cervix | GSM14050 | 25036 | 48994 |
| 1037.808 | 389 | 1053.252 | 490 | 783.3585 | 396 | 783.4227 | 452 | AFF4 | HeLa | Cervix | GSM14050 | 17742 | 48995 |
| 1047.142 | 434 | 1075.819 | 562 | 812.2364 | 475 | 797.4382 | 518 | AFF4 | HeLa | Cervix | GSM14050 | 19760 | 48996 |
| 1153.62 | 175 | 1191.864 | 236 | 723.4395 | 171 | 650.2617 | 171 | AFF4 | HeLa | Cervix | GSM14050 | 4467 | 48997 |
| 1171.402 | 193 | 1223.835 | 271 | 956.6929 | 206 | 717.9792 | 185 | AFF4 | HeLa | Cervix | GSM14050 | 4650 | 48998 |
| 1169.988 | 256 | 1235.915 | 365 | 936.8321 | 261 | 879.1098 | 249 | AFF4 | HeLa | Cervix | GSM14050 | 6467 | 48999 |
| 1177.157 | 161 | 1192.15 | 205 | 695.278 | 156 | 649.1911 | 159 | AFF4 | HeLa | Cervix | GSM14050 | 3759 | 49000 |
| 1137.149 | 363 | 1169.912 | 469 | 884.3576 | 367 | 858.0547 | 384 | ELL2 | HeLa | Cervix | GSM14050 | 11529 | 49001 |
| 1197.428 | 397 | 1213.415 | 496 | 916.6282 | 378 | 887.2607 | 395 | ELL2 | HeLa | Cervix | GSM14050 | 10833 | 49002 |
| 1130.386 | 416 | 1122.76 | 495 | 875.5292 | 423 | 844.1923 | 443 | ELL2 | HeLa | Cervix | GSM14050 | 14286 | 49003 |
| 1153.972 | 321 | 1165.149 | 399 | 891.6691 | 313 | 875.7349 | 339 | ELL2 | HeLa | Cervix | GSM14050 | 9437 | 49004 |
| 1304.95 | 184 | 1297.691 | 219 | 833.4884 | 154 | 856.6413 | 165 | ELL2 | HeLa | Cervix | GSM14050 | 2901 | 49005 |
| 1299.564 | 138 | 1298.902 | 168 | 645.5111 | 118 | 633.0028 | 123 | ELL2 | HeLa | Cervix | GSM14050 | 2157 | 49006 |
| 1328.429 | 155 | 1348.658 | 200 | 874.7403 | 142 | 688.2298 | 130 | ELL2 | HeLa | Cervix | GSM14050 | 2203 | 49007 |
| 1263.639 | 248 | 1271.481 | 307 | 989.8229 | 229 | 938.3045 | 224 | ELL2 | HeLa | Cervix | GSM14050 | 4734 | 49008 |
| 506.9862 | 62 | 446.5155 | 62 | 130.5958 | 38 | 259.7102 | 59 | ERCC6 | None | Skin | GSM14051 | 1301 | 49027 |
| 267.549 | 39 | 372.2654 | 52 | 116.4457 | 33 | 228.6124 | 51 | ERCC6 | None | Skin | GSM14051 | 1095 | 49033 |
| 1130.898 | 294 | 1173.805 | 393 | 865.0132 | 275 | 840.2024 | 293 | ERCC6 | None | Skin | GSM14051 | 8812 | 49036 |
| 8.334277 | 11 | 8.797472 | 13 | 13.27358 | 22 | 28.13945 | 32 | SFPQ | MCF-7 | Breast | GSM14112 | 3094 | 49154 |
| 449.9058 | 69 | 460.0498 | 76 | 292.2385 | 73 | 530.7773 | 108 | AR | 22RV1 | Prostate | GSM14123 | 1935 | 49158 |
| 180.7066 | 60 | 312.6791 | 89 | 83.24011 | 62 | 162.4452 | 90 | EMX1 | HeLa | Cervix | GSM14145 | 5172 | 49183 |
| 995.1012 | 133 | 1231.921 | 196 | 607.6531 | 134 | 556.1975 | 135 | CSNK2A1 | LNCaP | Prostate | GSM14151 | 3072 | 49193 |
| 0.531194 | 65 | 1.147991 | 86 | 0.246855 | 126 | 1.757657 | 166 | DDX5 | NTERA-2 | Testis | GSM14159 | 57890 | 49196 |
| 130.3864 | 79 | 191.4377 | 106 | 120.9053 | 119 | 188.7067 | 154 | PDX1 | None | Embryo | GSM14169 | 11994 | 49200 |
| 1036.258 | 326 | 1072.34 | 432 | 877.6103 | 424 | 851.2816 | 449 | ERG | VCaP | Prostate | GSM14224 | 13904 | 49225 |
| 1071.972 | 210 | 1084.509 | 266 | 912.9397 | 266 | 887.3695 | 282 | ERG | VCaP | Prostate | GSM14224 | 7166 | 49226 |
| 1199.807 | 332 | 1212.416 | 409 | 933.7547 | 309 | 869.8937 | 293 | ETS1 | DU145 | Prostate | GSM14245 | 7308 | 49238 |
| 1207.87 | 151 | 1246.724 | 208 | 769.213 | 154 | 641.6173 | 147 | GABPA | DU145 | Prostate | GSM14245 | 3181 | 49241 |

| | | | | | | | | | | | | | |
|---|---|---|---|---|---|---|---|---|---|---|---|---|---|
| 107.2656 | 25 | 131.9765 | 31 | 77.41629 | 30 | 148.2695 | 45 | JUND | DU145 | Prostate | GSM14245 | 1407 | 49242 |
| 1214.912 | 167 | 1206.006 | 203 | 693.2221 | 151 | 680.024 | 158 | ETS1 | PANC-1 | Pancreas | GSM14245 | 3559 | 49243 |
| 196.4761 | 37 | 389.6426 | 61 | 80.28255 | 32 | 275.895 | 66 | SCML2 | 293T-Rex | None | GSM14245 | 1561 | 49258 |
| 126.3612 | 68 | 227.4107 | 101 | 68.24049 | 82 | 148.2545 | 121 | BMI1 | 293T-Rex | None | GSM14245 | 9563 | 49259 |
| 519.3884 | 85 | 531.5807 | 94 | 254.4574 | 79 | 296.6676 | 91 | BMI1 | 293T-Rex | None | GSM14245 | 2872 | 49263 |
| 483.1228 | 167 | 510.3675 | 191 | 246.3335 | 183 | 289.4064 | 211 | SCML2 | 293T-Rex | None | GSM14245 | 15184 | 49266 |
| 93.51244 | 27 | 94.00341 | 30 | 83.10181 | 37 | 148.3234 | 53 | BMI1 | 293T-Rex | None | GSM14245 | 1974 | 49267 |
| 674.0757 | 206 | 855.7381 | 277 | 560.5253 | 276 | 669.7515 | 321 | ZBTB17 | LS174T | Colon | GSM14309 | 16825 | 49276 |
| 570.5804 | 88 | 795.2122 | 117 | 375.4133 | 95 | 556.2496 | 125 | MYC | LS174T | Colon | GSM14309 | 2718 | 49277 |
| 298.41 | 58 | 326.9936 | 67 | 105.0378 | 47 | 243.3763 | 77 | ZBTB17 | LS174T | Colon | GSM14309 | 2521 | 49279 |
| 2.922275 | 7 | 4.922527 | 10 | 0.376677 | 8 | 11.34135 | 22 | EP300 | H1 | Embryo | GSM44758 | 3071 | 4928 |
| 12.15561 | 27 | 4.19356 | 23 | 0.172492 | 22 | 16.79422 | 57 | SOX2 | H1 | Embryo | GSM45657 | 9847 | 4931 |
| 3.492388 | 32 | 1.628538 | 33 | 2.349719 | 57 | 4.478763 | 71 | NANOG | H1 | Embryo | GSM45657 | 19332 | 4932 |
| 7.887373 | 44 | 1.913979 | 39 | 4.983272 | 74 | 5.567044 | 84 | NANOG | H1 | Embryo | GSM45657 | 22508 | 4933 |
| 299.5002 | 93 | 548.692 | 142 | 230.9998 | 120 | 202.2517 | 121 | RELA | GM19193 | Blood | GSM48626 | 7360 | 4938 |
| 974.8817 | 417 | 1001.653 | 542 | 720.0112 | 431 | 665.6075 | 426 | RELA | GM19193 | Blood | GSM48627 | 24540 | 4939 |
| 124.3041 | 81 | 231.4895 | 121 | 86.48539 | 110 | 89.79443 | 121 | RELA | GM19193 | Blood | GSM48627 | 13008 | 4940 |
| 304.7813 | 71 | 369.4253 | 87 | 208.0873 | 83 | 210.3149 | 89 | RELA | GM18505 | Blood | GSM48627 | 3953 | 4941 |
| 908.4081 | 565 | 906.2939 | 679 | 662.8729 | 624 | 649.2085 | 690 | EP300 | IMR90 | Lung | GSM14427 | 46494 | 49414 |
| 884.5159 | 404 | 885.849 | 498 | 672.6403 | 465 | 631.3094 | 476 | EP300 | IMR90 | Lung | GSM14427 | 31269 | 49415 |
| 1111.712 | 288 | 1157.16 | 390 | 858.4936 | 279 | 817.0702 | 284 | RB1 | IMR90 | Lung | GSM14428 | 9149 | 49416 |
| 1016.809 | 288 | 1049.94 | 382 | 769.1341 | 288 | 774.5188 | 329 | RB1 | IMR90 | Lung | GSM14428 | 12641 | 49417 |
| 407.6311 | 74 | 516.9618 | 93 | 204.6522 | 71 | 234.2277 | 81 | RELA | GM18505 | Blood | GSM48627 | 2930 | 4942 |
| 267.1926 | 136 | 395.584 | 183 | 199.256 | 184 | 179.3919 | 191 | RELA | GM18505 | Blood | GSM48627 | 17844 | 4943 |
| 843.2537 | 136 | 1073.815 | 171 | 477.812 | 137 | 431.9397 | 138 | RELA | GM18505 | Blood | GSM48627 | 4597 | 4944 |
| 1131.516 | 961 | 1139.497 | 1150 | 807.221 | 1106 | 787.3409 | 1235 | HIF1A | T47D | Mammary | GSM14624 | 73122 | 49446 |
| 1162.807 | 872 | 1184.793 | 1067 | 856.9927 | 978 | 823.3157 | 1054 | HIF1A | T47D | Mammary | GSM14624 | 49223 | 49447 |
| 1121.707 | 561 | 1124.674 | 676 | 880.9135 | 613 | 859.8247 | 666 | EGLN2 | T47D | Mammary | GSM14624 | 21948 | 49448 |
| 1128.144 | 236 | 1202.699 | 348 | 879.0016 | 228 | 895.7112 | 272 | NRF1 | T47D | Mammary | GSM14624 | 6740 | 49449 |
| 630.4589 | 168 | 854.7424 | 218 | 393.8992 | 193 | 260.9799 | 172 | RELA | GM18505 | Blood | GSM48627 | 11359 | 4945 |
| 2.895287 | 3 | 3.795556 | 4 | 0 | 1 | 113.9374 | 33 | HES1 | SW1353 | Bone | GSM14634 | 1004 | 49475 |
| 538.4284 | 282 | 718.3805 | 395 | 384.8735 | 379 | 383.9692 | 411 | CTCF | None | None | GSM14639 | 40244 | 49479 |
| 754.3838 | 90 | 1152.736 | 127 | 300.267 | 70 | 424.7074 | 90 | E2F4 | None | None | GSM14639 | 1780 | 49480 |
| 1291.134 | 640 | 1330.729 | 822 | 943.6976 | 573 | 898.6823 | 583 | RPE | None | None | GSM14639 | 16072 | 49481 |
| 134.4253 | 125 | 161.1399 | 153 | 86.27707 | 173 | 99.94033 | 198 | ZNF92 | None | None | GSM14639 | 26467 | 49482 |
| 830.1668 | 403 | 819.7853 | 481 | 623.2357 | 471 | 605.02 | 509 | MED1 | None | None | GSM14639 | 36716 | 49484 |
| 1281.964 | 402 | 1320.059 | 527 | 983.394 | 365 | 953.6394 | 376 | CDK7 | Jurkat | Blood | GSM14639 | 8186 | 49493 |
| 1191.87 | 474 | 1201.823 | 594 | 877.415 | 428 | 844.8516 | 447 | CDK7 | Jurkat | Blood | GSM14639 | 14253 | 49494 |
| 1152.879 | 762 | 1168.217 | 930 | 814.5797 | 749 | 773.111 | 785 | RUNX1 | None | None | GSM14660 | 37079 | 49536 |
| 17.79277 | 8 | 14.07481 | 8 | 2.139038 | 5 | 58.57077 | 24 | EP300 | Kasumi-1 | Blood | GSM14660 | 1075 | 49537 |
| 310.8449 | 57 | 566.2598 | 89 | 248.4377 | 70 | 361.2121 | 91 | BRD2 | H9 | Embryo | GSM14668 | 2282 | 49543 |
| 125.2968 | 34 | 153.2304 | 42 | 82.92457 | 40 | 96.2404 | 46 | RELA | GM10847 | Blood | GSM48630 | 2319 | 4959 |
| 229.673 | 77 | 391.6069 | 113 | 179.8955 | 101 | 150.5743 | 100 | RELA | GM10847 | Blood | GSM48630 | 6615 | 4960 |
| 0.264076 | 10 | -0.001414 | 8 | 0 | 15 | -0.030803 | 15 | ZNF486 | None | Embryo | GSM14677 | 8163 | 49603 |
| -0.084025 | 13 | -2.247388 | 9 | -4.365526 | 15 | -0.928239 | 26 | ZNF486 | None | Embryo | GSM14677 | 18566 | 49604 |
| 870.0163 | 434 | 904.673 | 567 | 665.0155 | 498 | 633.6541 | 522 | TRIM28 | None | Embryo | GSM14677 | 30919 | 49606 |
| 72.78119 | 87 | 97.06808 | 111 | 50.33437 | 126 | 76.09085 | 157 | TRIM28 | None | Embryo | GSM14677 | 20701 | 49608 |
| 30.46005 | 13 | 45.70829 | 18 | 14.21194 | 14 | 89.96276 | 36 | TRIM28 | None | Embryo | GSM14677 | 1504 | 49609 |
| 881.6837 | 274 | 884.3532 | 336 | 477.3599 | 276 | 496.7356 | 302 | RELA | GM10847 | Blood | GSM48630 | 19124 | 4961 |
| 268.5649 | 78 | 290.4278 | 90 | 611.7891 | 172 | 906.792 | 234 | ESR1 | MCF-7 | Breast | GSM14699 | 5641 | 49611 |
| 227.1448 | 161 | 257.7983 | 192 | 437.7834 | 324 | 638.6539 | 437 | ESR1 | MCF-7 | Breast | GSM14699 | 27144 | 49612 |
| 1005.389 | 196 | 1033.66 | 260 | 912.739 | 303 | 928.7476 | 361 | RARA | MCF-7 | Breast | GSM14699 | 8452 | 49614 |
| 817.4753 | 147 | 1046.072 | 195 | 955.5691 | 243 | 960.409 | 280 | RARG | MCF-7 | Breast | GSM14699 | 5751 | 49617 |
| 179.3311 | 37 | 247.9536 | 49 | 380.8647 | 79 | 693.8488 | 118 | RARG | MCF-7 | Breast | GSM14699 | 1721 | 49618 |
| 124.409 | 27 | 173.6395 | 36 | 314.344 | 64 | 554.6486 | 94 | RARG | MCF-7 | Breast | GSM14699 | 1358 | 49619 |
| 288.1122 | 74 | 447.1789 | 104 | 188.6744 | 86 | 202.059 | 95 | RELA | GM18951 | Blood | GSM48630 | 4670 | 4962 |
| 1033.156 | 284 | 1053.58 | 361 | 915.4762 | 407 | 923.7227 | 470 | RARA | MCF-7 | Breast | GSM14699 | 11804 | 49620 |
| 391.6482 | 65 | 562.0876 | 88 | 794.4846 | 134 | 995.1672 | 161 | RARA | MCF-7 | Breast | GSM14699 | 2216 | 49621 |
| 845.5376 | 125 | 1156.623 | 174 | 957.4188 | 182 | 989.628 | 199 | RARG | MCF-7 | Breast | GSM14699 | 3613 | 49622 |
| 1089.111 | 196 | 1129.998 | 271 | 966.183 | 278 | 971.442 | 318 | RARG | MCF-7 | Breast | GSM14699 | 6377 | 49623 |
| 258.1878 | 89 | 222.833 | 92 | 480.4072 | 177 | 670.4736 | 223 | GATA3 | MCF-7 | Breast | GSM14699 | 7872 | 49625 |
| 339.2779 | 120 | 301.1143 | 126 | 533.4808 | 219 | 716.9931 | 270 | GATA3 | MCF-7 | Breast | GSM14699 | 10954 | 49626 |
| 366.8413 | 94 | 342.5151 | 100 | 573.0967 | 169 | 751.3722 | 207 | GATA3 | MCF-7 | Breast | GSM14699 | 5828 | 49627 |
| 284.3073 | 69 | 263.848 | 73 | 554.699 | 140 | 712.4735 | 170 | GATA3 | MCF-7 | Breast | GSM14699 | 3996 | 49628 |
| 655.0658 | 127 | 892.908 | 166 | 614.34 | 171 | 873.9391 | 219 | TFAP2C | MCF-7 | Breast | GSM14699 | 5542 | 49629 |
| 961.8641 | 491 | 992.7778 | 637 | 716.3031 | 535 | 663.5172 | 532 | RELA | GM18951 | Blood | GSM48630 | 32551 | 4963 |
| 980.1352 | 406 | 1003.354 | 518 | 838.2244 | 552 | 841.0382 | 633 | TFAP2C | MCF-7 | Breast | GSM14699 | 22772 | 49630 |
| 375.2017 | 63 | 463.8247 | 78 | 228.8324 | 66 | 286.2187 | 79 | PRKDC | MCF-7 | Breast | GSM14700 | 2209 | 49634 |
| 131.6722 | 25 | 217.6077 | 37 | 118.92 | 33 | 179.6715 | 44 | PRKDC | MCF-7 | Breast | GSM14700 | 1071 | 49636 |
| 10.06173 | 9 | 19.63611 | 14 | 7.052924 | 13 | 23.83983 | 23 | RELA | GM18951 | Blood | GSM48630 | 2093 | 4964 |
| 114.1615 | 24 | 96.95772 | 24 | 183.0563 | 44 | 382.6045 | 71 | ESR1 | MCF-7 | Breast | GSM14700 | 1180 | 49640 |
| 36.13763 | 13 | 48.49492 | 17 | 163.8441 | 43 | 260.3096 | 59 | ESR1 | MCF-7 | Breast | GSM14700 | 1288 | 49643 |

| | | | | | | | | | | | | | |
|---|---|---|---|---|---|---|---|---|---|---|---|---|---|
| 33.68116 | 12 | 46.76187 | 16 | 102.5453 | 32 | 260.9331 | 57 | ESR1 | MCF-7 | Breast | GSM14700 | 1186 | 49644 |
| 1183.176 | 663 | 1195.737 | 810 | 921.1799 | 724 | 890.6581 | 769 | EP300 | MCF-7 | Breast | GSM14700 | 24076 | 49645 |
| 1086.056 | 307 | 1067.405 | 354 | 877.9308 | 344 | 879.2288 | 392 | EP300 | MCF-7 | Breast | GSM14700 | 10776 | 49646 |
| 768.9649 | 118 | 920.3848 | 143 | 679.9094 | 150 | 745.8524 | 167 | EP300 | MCF-7 | Breast | GSM14700 | 3619 | 49647 |
| 1049.012 | 432 | 1066.993 | 542 | 850.5467 | 508 | 815.8117 | 531 | EP300 | MCF-7 | Breast | GSM14700 | 18419 | 49648 |
| 1206.007 | 436 | 1205.502 | 530 | 935.2534 | 433 | 889.8102 | 434 | MED1 | MCF-7 | Breast | GSM14700 | 11658 | 49649 |
| 907.9794 | 247 | 946.1453 | 339 | 580.3892 | 272 | 622.0956 | 301 | RELA | GM18951 | Blood | GSM48630 | 15682 | 4965 |
| 1174.058 | 250 | 1154.064 | 283 | 914.1466 | 235 | 897.4861 | 252 | MED1 | MCF-7 | Breast | GSM14700 | 6179 | 49650 |
| 1085.699 | 450 | 1094.169 | 556 | 850.1085 | 482 | 826.0601 | 519 | MED1 | MCF-7 | Breast | GSM14700 | 17784 | 49651 |
| 1199.273 | 606 | 1205.273 | 739 | 914.8656 | 614 | 854.2697 | 610 | MED1 | MCF-7 | Breast | GSM14700 | 19596 | 49652 |
| 100.1115 | 29 | 99.38661 | 32 | 99.4976 | 42 | 261.2198 | 74 | ESR1 | MCF-7 | Breast | GSM14700 | 2111 | 49653 |
| 152.3401 | 42 | 143.6742 | 45 | 149.4775 | 60 | 340.7253 | 98 | ESR1 | MCF-7 | Breast | GSM14700 | 2843 | 49654 |
| 66.69115 | 19 | 132.7412 | 31 | 93.66805 | 33 | 222.5526 | 56 | ESR1 | MCF-7 | Breast | GSM14700 | 1390 | 49656 |
| 889.7409 | 233 | 858.2155 | 259 | 783.1924 | 347 | 797.8972 | 409 | FOXA1 | MCF-7 | Breast | GSM14700 | 15297 | 49657 |
| 809.6142 | 415 | 787.4951 | 482 | 683.5511 | 597 | 691.9582 | 692 | FOXA1 | MCF-7 | Breast | GSM14700 | 41588 | 49658 |
| 139.8871 | 51 | 198.4845 | 68 | 83.97751 | 60 | 68.04903 | 59 | RELA | GM18526 | Blood | GSM48631 | 4801 | 4966 |
| 897.0691 | 552 | 920.6509 | 700 | 697.0084 | 681 | 671.7124 | 735 | PROX1 | SW480 | Colon | GSM14775 | 48145 | 49669 |
| 575.899 | 110 | 684.3393 | 133 | 316.5845 | 112 | 347.3046 | 125 | RELA | GM18526 | Blood | GSM48631 | 4728 | 4968 |
| 14.41364 | 22 | 16.9066 | 27 | 6.087967 | 29 | 38.08386 | 57 | RELA | GM12878 | Blood | GSM48631 | 6540 | 4969 |
| 111.5457 | 55 | 80.67089 | 53 | 50.03003 | 59 | 115.9883 | 90 | DEK | U937 | Pleura | GSM14859 | 6090 | 49697 |
| 534.4611 | 208 | 782.124 | 284 | 298.7003 | 242 | 277.7987 | 254 | RELA | GM12878 | Blood | GSM48631 | 21536 | 4970 |
| 395.6738 | 122 | 573.3014 | 164 | 291.6557 | 154 | 218.0276 | 145 | RELA | GM12878 | Blood | GSM48631 | 9596 | 4971 |
| 422.932 | 132 | 664.561 | 185 | 268.565 | 156 | 177.3586 | 140 | RELA | GM12878 | Blood | GSM48631 | 10551 | 4972 |
| 216.2281 | 92 | 337.9172 | 128 | 100.1988 | 100 | 172.7586 | 135 | ASCL1 | NCI-H1184 | Lung | GSM14994 | 10160 | 49723 |
| 99.18384 | 51 | 178.3696 | 76 | 101.692 | 79 | 171.9235 | 107 | ASCL1 | NCI-H128 | Lung | GSM14994 | 6708 | 49724 |
| 195.4447 | 57 | 206.4475 | 65 | 130.2466 | 68 | 150.7076 | 78 | ASCL1 | NCI-H1755 | Lung | GSM14994 | 4181 | 49725 |
| 223.5964 | 118 | 377.4261 | 169 | 184.3436 | 167 | 238.1256 | 200 | ASCL1 | NCI-H2107 | Lung | GSM14994 | 15997 | 49726 |
| 46.36377 | 26 | 52.17921 | 31 | 26.39759 | 32 | 92.91232 | 59 | ASCL1 | NCI-H2107 | Lung | GSM14994 | 3760 | 49727 |
| 189.8721 | 56 | 201.2306 | 64 | 63.65345 | 49 | 189.7179 | 87 | ASCL1 | HCC4018 | Lung | GSM14994 | 4190 | 49728 |
| 1094.384 | 409 | 1148.826 | 559 | 830.3924 | 411 | 778.2287 | 405 | RELA | GM12891 | Blood | GSM48632 | 15887 | 4973 |
| 58.12257 | 22 | 69.68855 | 27 | 37.65805 | 27 | 123.1557 | 51 | ASCL1 | HCC4018 | Lung | GSM14994 | 2243 | 49730 |
| 43.8105 | 38 | 93.12774 | 60 | 31.18432 | 54 | 67.20911 | 78 | CEBPA | Kasumi-1 | Blood | GSM15011 | 7650 | 49737 |
| 429.8917 | 176 | 549.6191 | 221 | 258.5392 | 212 | 247.509 | 225 | LMO2 | Kasumi-1 | Blood | GSM15011 | 18868 | 49738 |
| 349.4154 | 137 | 627.3497 | 204 | 263.3626 | 180 | 273.7449 | 197 | LMO2 | Kasumi-1 | Blood | GSM15011 | 13901 | 49739 |
| 860.1868 | 570 | 883.6855 | 720 | 620.9003 | 647 | 579.2264 | 670 | RELA | GM12891 | Blood | GSM48632 | 57553 | 4974 |
| 268.4615 | 209 | 327.8628 | 257 | 141.8882 | 266 | 137.976 | 288 | SPI1 | Kasumi-1 | Blood | GSM15011 | 39812 | 49740 |
| 539.475 | 274 | 662.5164 | 337 | 243.1461 | 310 | 200.9592 | 316 | RUNX1 | Kasumi-1 | Blood | GSM15011 | 37936 | 49741 |
| 648.903 | 246 | 773.9305 | 314 | 266.3071 | 253 | 241.5014 | 264 | SSU72 | H1 | Embryo | GSM15038 | 25177 | 49742 |
| 119.6135 | 37 | 148.6426 | 46 | 72.53318 | 42 | 83.00261 | 48 | BRD4 | None | None | GSM15038 | 2687 | 49744 |
| 787.6055 | 377 | 816.6303 | 495 | 580.2372 | 439 | 497.1536 | 448 | RELA | GM12891 | Blood | GSM48632 | 39456 | 4975 |
| 1130.007 | 461 | 1151.182 | 580 | 835.1459 | 412 | 769.5354 | 390 | CHD8 | None | None | GSM15060 | 12523 | 49778 |
| 167.3976 | 72 | 201.5799 | 88 | 97.62225 | 85 | 131.7835 | 104 | CHD7 | None | None | GSM15060 | 7498 | 49781 |
| 16.98568 | 9 | 10.39679 | 8 | 7.961076 | 10 | 0 | 2 | EZH2 | None | None | GSM15089 | 1250 | 49798 |
| 9.555193 | 6 | 24.47659 | 11 | 6.426145 | 8 | 11.02053 | 11 | EZH2 | SF7761 | None | GSM15089 | 1013 | 49801 |
| 30.38193 | 20 | 59.27387 | 31 | 20.77097 | 27 | 50.19774 | 42 | FLI1 | SK-N-MC | Brain | GSM15175 | 3285 | 49847 |
| 1160.567 | 450 | 1193.926 | 583 | 861.5053 | 411 | 837.6654 | 439 | WDR5 | SK-N-MC | Brain | GSM15175 | 14017 | 49852 |
| 202.4539 | 59 | 268.7008 | 76 | 89.37871 | 58 | 158.0389 | 81 | FLI1 | SK-N-MC | Brain | GSM15175 | 4253 | 49854 |
| 12.0405 | 14 | 36.78164 | 26 | 6.589442 | 19 | 37.06162 | 39 | FLI1 | SK-N-MC | Brain | GSM15175 | 3594 | 49856 |
| 1097.358 | 331 | 1103.863 | 411 | 810.4582 | 301 | 782.6399 | 318 | GABPA | SK-N-MC | Brain | GSM15175 | 11549 | 49860 |
| 1118.558 | 179 | 1144.08 | 238 | 642.9493 | 172 | 728.0808 | 195 | ELF1 | SK-N-MC | Brain | GSM15175 | 5304 | 49861 |
| 963.8426 | 146 | 1157.845 | 208 | 552.3272 | 146 | 587.0711 | 160 | EP300 | SK-N-MC | Brain | GSM15175 | 4368 | 49862 |
| 1070.423 | 335 | 1102.965 | 441 | 822.2389 | 335 | 779.102 | 338 | GABPA | SK-N-MC | Brain | GSM15175 | 12681 | 49869 |
| 1005.106 | 479 | 1036.355 | 616 | 777.3265 | 537 | 719.0693 | 527 | ELF1 | SK-N-MC | Brain | GSM15175 | 26292 | 49870 |
| 390.7392 | 104 | 564.3726 | 140 | 267.7888 | 124 | 355.8357 | 152 | EP300 | SK-N-MC | Brain | GSM15175 | 6762 | 49871 |
| 13.61687 | 16 | 46.0469 | 31 | 11.2637 | 25 | 53.08052 | 49 | FLI1 | A673 | Muscle | GSM15175 | 4144 | 49872 |
| 1158.547 | 474 | 1204.151 | 627 | 858.9953 | 438 | 824.8887 | 455 | WDR5 | A673 | Muscle | GSM15175 | 15526 | 49877 |
| 0.18758 | 4 | 2.611997 | 9 | -0.018625 | 5 | 15.01809 | 27 | FLI1 | A673 | Muscle | GSM15175 | 3524 | 49879 |
| 537.4627 | 147 | 667.8395 | 182 | 278.2315 | 154 | 269.6971 | 163 | OTX2 | None | None | GSM15217 | 9642 | 49947 |
| 1104.795 | 822 | 1198.343 | 1094 | 794.383 | 897 | 756.2583 | 952 | KLF9 | GBM1A | None | GSM15225 | 52866 | 49949 |
| 771.91 | 118 | 1128.017 | 162 | 434.781 | 117 | 421.1548 | 122 | KLF9 | GBM1B | None | GSM15225 | 3592 | 49951 |
| 152.6034 | 33 | 320.479 | 56 | 88.26251 | 35 | 204.3391 | 58 | KDM3A | Jurkat | Blood | GSM15251 | 1688 | 49953 |
| 451.5721 | 78 | 612.4794 | 102 | 237.3617 | 76 | 310.2177 | 93 | KDM3A | Jurkat | Blood | GSM15251 | 2873 | 49954 |
| 574.6241 | 106 | 636.0502 | 123 | 373.5868 | 117 | 431.1551 | 134 | KDM3A | Jurkat | Blood | GSM15251 | 4326 | 49955 |
| 833.7622 | 497 | 820.5367 | 591 | 593.873 | 551 | 583.1772 | 609 | BRD4 | 90-8TL | Brain | GSM15279 | 49609 | 49977 |
| 1040.675 | 551 | 1052.711 | 682 | 753.5028 | 548 | 723.3684 | 582 | BRD4 | 90-8TL | Brain | GSM15279 | 29593 | 49980 |
| 377.5615 | 61 | 612.5773 | 89 | 313.7218 | 75 | 325.4615 | 81 | BRD4 | GM12878 | Blood | GSM15361 | 1951 | 49989 |
| 19.29491 | 9 | 15.1948 | 9 | 9.129231 | 10 | 59.63923 | 26 | POU5F1 | NT2-D1 | Embryo | GSM15376 | 1234 | 49996 |
| 197.6718 | 34 | 344.5518 | 52 | 141.858 | 39 | 214.5779 | 52 | SCML2 | 293T-Rex | None | GSM93786 | 1244 | 50013 |
| 915.269 | 191 | 932.7215 | 213 | 507.8673 | 200 | 529.7724 | 218 | SCML2 | K562 | Bone Marr | GSM93787 | 9629 | 50020 |
| 409.6372 | 99 | 494.6191 | 121 | 243.2342 | 109 | 295.6642 | 128 | BMI1 | K562 | Bone Marr | GSM93787 | 5870 | 50021 |
| -0.028311 | 5 | -0.598869 | 4 | -0.227072 | 10 | 0.454449 | 23 | MYC | A549 | Lung | GSM99104 | 8096 | 50033 |
| -0.107626 | 3 | -1.062877 | 2 | -1.040519 | 5 | 0.357599 | 17 | MYC | A549 | Lung | GSM99104 | 6128 | 50034 |

| | | | | | | | | | | | | | |
|---|---|---|---|---|---|---|---|---|---|---|---|---|---|
| 0 | 1 | -0.015062 | 1 | 0.243514 | 5 | 1.670136 | 9 | NME2 | A549 | Lung | GSM99104 | 2078 | 50035 |
| 0.395842 | 3 | -0.027799 | 1 | 0 | 4 | 0 | 4 | NME2 | A549 | Lung | GSM99104 | 2227 | 50036 |
| 904.5969 | 980 | 811.2133 | 1067 | 682.4187 | 1320 | 731.6391 | 1603 | FOXA1 | LNCaP | Prostate | GSM10681 | 156594 | 50039 |
| 880.4609 | 895 | 784.8655 | 951 | 684.0759 | 1212 | 735.8242 | 1491 | FOXA1 | LNCaP | Prostate | GSM10681 | 131115 | 50040 |
| 1340.534 | 173 | 1256.734 | 162 | 1030.686 | 162 | 1140.979 | 216 | AR | LNCaP | Prostate | GSM10696 | 2397 | 50041 |
| 1330.689 | 171 | 1249.88 | 162 | 982.0876 | 159 | 1157.499 | 233 | AR | LNCaP | Prostate | GSM10696 | 2456 | 50042 |
| 1003.885 | 318 | 949.2011 | 337 | 795.9665 | 358 | 886.7313 | 518 | AR | LNCaP | Prostate | GSM10696 | 14575 | 50043 |
| 1021.483 | 312 | 972.8008 | 332 | 798.3488 | 333 | 906.8715 | 508 | AR | LNCaP | Prostate | GSM10696 | 13501 | 50044 |
| 1075.302 | 511 | 972.998 | 495 | 895.1197 | 648 | 967.4018 | 870 | AR | LNCaP | Prostate | GSM10712 | 23441 | 50045 |
| 863.9989 | 761 | 773.8765 | 793 | 674.5738 | 1005 | 743.8721 | 1286 | AR | LNCaP | Prostate | GSM10712 | 93100 | 50047 |
| 930.1674 | 486 | 849.9889 | 491 | 739.1963 | 599 | 822.7966 | 829 | AR | LNCaP | Prostate | GSM10712 | 35235 | 50048 |
| 395.0129 | 96 | 348.7817 | 99 | 159.2258 | 88 | 508.5917 | 166 | AR | None | Prostate | GSM10712 | 5727 | 50049 |
| 129.2384 | 51 | 129.5199 | 57 | 75.59655 | 60 | 203.8915 | 101 | AR | None | Prostate | GSM10712 | 5174 | 50051 |
| 105.7173 | 45 | 140.6698 | 58 | 54.99363 | 51 | 180.4469 | 93 | AR | None | Prostate | GSM10712 | 4937 | 50052 |
| 413.214 | 71 | 603.9239 | 97 | 310.1161 | 83 | 688.1443 | 137 | AR | None | Prostate | GSM10712 | 2453 | 50054 |
| 668.3391 | 106 | 654.8032 | 114 | 411.7986 | 110 | 851.2923 | 174 | AR | None | Prostate | GSM10712 | 3204 | 50056 |
| 868.2081 | 237 | 839.7897 | 270 | 584.8515 | 284 | 776.3644 | 422 | AR | None | Prostate | GSM10712 | 16898 | 50057 |
| 859.9802 | 235 | 842.2187 | 271 | 675.2994 | 302 | 780.7405 | 423 | AR | None | Prostate | GSM10712 | 16698 | 50058 |
| 195.321 | 68 | 256.4238 | 87 | 122.1724 | 81 | 324.4963 | 137 | AR | None | Prostate | GSM10712 | 6107 | 50059 |
| 254.0259 | 87 | 234.5533 | 93 | 167.5983 | 106 | 379.3582 | 166 | AR | None | Prostate | GSM10712 | 7709 | 50060 |
| 0.897988 | 6 | 1.310712 | 8 | 0.379342 | 10 | 25.88138 | 35 | ESR1 | VCaP | Prostate | GSM10761 | 3859 | 50062 |
| 131.2087 | 30 | 117.4522 | 31 | 91.0174 | 35 | 125.1859 | 44 | ESR1 | VCaP | Prostate | GSM10761 | 1583 | 50063 |
| 207.0395 | 64 | 207.0339 | 71 | 95.07643 | 65 | 101.7221 | 72 | ESR1 | NCI-H660 | Prostate | GSM10761 | 5017 | 50064 |
| 1219.491 | 262 | 1217.316 | 324 | 933.0467 | 234 | 906.9922 | 247 | ZFX | Daoy | Cerebellum | GSM11035 | 5814 | 50077 |
| 1207.012 | 351 | 1094.527 | 323 | 1016.059 | 425 | 1068.236 | 548 | AR | LNCaP | Prostate | GSM12369 | 8990 | 50100 |
| 1114.124 | 477 | 990.9682 | 435 | 924.4674 | 576 | 955.6754 | 711 | AR | LNCaP | Prostate | GSM12369 | 17378 | 50102 |
| 1148.128 | 313 | 1137.96 | 377 | 874.6545 | 298 | 833.9259 | 300 | CHRM2 | U937 | Pleura | GSM12759 | 9543 | 50107 |
| 1138.546 | 251 | 1119.354 | 292 | 790.7413 | 222 | 817.2834 | 240 | CHRM2 | U937 | Pleura | GSM12759 | 7455 | 50108 |
| 118.5506 | 34 | 114.7119 | 37 | 73.28423 | 39 | 102.1305 | 49 | MAF | None | Foreskin | GSM12785 | 2466 | 50111 |
| 1213.218 | 179 | 1223.754 | 225 | 754.4615 | 163 | 898.7065 | 190 | MAFB | None | Foreskin | GSM12785 | 3838 | 50112 |
| 1139.245 | 258 | 1066.41 | 254 | 946.0037 | 299 | 1035.78 | 438 | KDM1A | LNCaP | Prostate | GSM12797 | 7648 | 50114 |
| 220.9115 | 49 | 282.8145 | 62 | 90.42723 | 44 | 200.1419 | 70 | VDR | THP-1 | Blood | GSM12808 | 2564 | 50116 |
| 424.1591 | 246 | 401.0507 | 271 | 144.5637 | 257 | 114.7599 | 261 | LMNB1 | IMR90 | Lung | GSM12894 | 36325 | 50122 |
| 21.46234 | 11 | 44.53355 | 18 | 44.33106 | 24 | 97.30746 | 38 | TCF7 | DLD-1 | Colon | GSM12956 | 1549 | 50135 |
| 32.00416 | 11 | 35.55892 | 13 | 42.61608 | 19 | 101.8185 | 32 | TCF7 | DLD-1 | Colon | GSM12956 | 1050 | 50136 |
| 30.32908 | 15 | 60.81287 | 24 | 73.4947 | 35 | 136.2948 | 51 | TCF7 | DLD-1 | Colon | GSM12956 | 1960 | 50139 |
| 1289.619 | 858 | 1269.807 | 1007 | 1114.959 | 1185 | 1174.328 | 1473 | ARRB1 | C4-2 | None | GSM13402 | 33587 | 50152 |
| 1288.281 | 781 | 1264.114 | 906 | 1125.984 | 1071 | 1157.135 | 1293 | ARRB1 | C4-2 | None | GSM13402 | 26095 | 50162 |
| 1068.553 | 508 | 1084.579 | 637 | 787.0378 | 505 | 763.9479 | 545 | TBP | K562 | Bone Marr | GSM13418 | 23961 | 50163 |
| 337.65 | 83 | 320.9481 | 89 | 186.7066 | 88 | 142.6733 | 83 | HIRA | IMR90 | Lung | GSM13588 | 4854 | 50172 |
| 386.2638 | 106 | 410.1995 | 121 | 217.79 | 116 | 208.9545 | 122 | HIRA | IMR90 | Lung | GSM13588 | 7226 | 50174 |
| 3.511258 | 6 | 3.465824 | 7 | 5.007213 | 12 | 67.21501 | 38 | HIRA | IMR90 | Lung | GSM13588 | 2163 | 50176 |
| 100.5685 | 92 | 158.3117 | 126 | 73.05983 | 132 | 91.11593 | 155 | HIRA | IMR90 | Lung | GSM13588 | 18896 | 50178 |
| 1206.151 | 304 | 1231.154 | 391 | 932.5791 | 279 | 895.3148 | 281 | DDX21 | HEK293 | Embryonic | GSM13693 | 6828 | 50210 |
| 16.28053 | 26 | 51.41752 | 47 | 34.72647 | 57 | 75.8688 | 83 | NR3C1 | None | None | GSM14054 | 7794 | 50246 |
| 783.8078 | 414 | 761.4754 | 478 | 660.1447 | 588 | 721.8928 | 775 | AR | VCaP | Prostate | GSM14107 | 41204 | 50254 |
| 1018.992 | 285 | 971.2017 | 306 | 892.2288 | 390 | 902.7297 | 456 | FOXP1 | VCaP | Prostate | GSM14107 | 11030 | 50256 |
| 317.345 | 76 | 327.0865 | 85 | 150.9652 | 74 | 384.9464 | 126 | RUNX1 | VCaP | Prostate | GSM14107 | 4090 | 50257 |
| 769.0193 | 286 | 754.2038 | 337 | 696.1009 | 459 | 751.8829 | 605 | AR | VCaP | Prostate | GSM14107 | 27222 | 50258 |
| 186.7573 | 35 | 129.5721 | 31 | 70.79783 | 29 | 282.8574 | 65 | AR | VCaP | Prostate | GSM14107 | 1442 | 50259 |
| 795.6608 | 274 | 781.057 | 323 | 688.1981 | 400 | 762.0941 | 556 | AR | VCaP | Prostate | GSM14107 | 23100 | 50260 |
| 944.0139 | 184 | 963.7882 | 207 | 631.1919 | 206 | 876.1763 | 295 | FOXA1 | VCaP | Prostate | GSM14107 | 7557 | 50261 |
| 1127.595 | 183 | 845.501 | 157 | 981.9818 | 235 | 1035.135 | 308 | FOXA1 | VCaP | Prostate | GSM14107 | 4665 | 50262 |
| 937.6814 | 218 | 925.0806 | 258 | 675.025 | 256 | 865.6775 | 401 | AR | LNCaP | Prostate | GSM14107 | 11249 | 50268 |
| 204.5469 | 43 | 258.7164 | 54 | 209.5336 | 61 | 429.57 | 96 | AR | VCaP | Prostate | GSM14107 | 2011 | 50270 |
| 1051.732 | 290 | 980.9183 | 292 | 848.8079 | 328 | 939.285 | 473 | AR | LNCaP | Prostate | GSM14107 | 11132 | 50273 |
| 1193.184 | 188 | 1117.499 | 187 | 1049.323 | 251 | 1130.793 | 356 | FOXA1 | LNCaP | Prostate | GSM14107 | 4291 | 50274 |
| 907.2007 | 594 | 821.094 | 603 | 751.3032 | 804 | 794.0962 | 1002 | FOXA1 | LNCaP | Prostate | GSM14107 | 50368 | 50275 |
| 20.27528 | 16 | 30.86918 | 22 | 5.511689 | 16 | 26.58597 | 31 | TP53 | HCT-116 | Colon | GSM14127 | 3063 | 50345 |
| 131.0134 | 51 | 100.9459 | 50 | 78.8575 | 60 | 68.72506 | 61 | TP53 | IMR90 | Lung | GSM14189 | 4802 | 50361 |
| 172.8371 | 307 | 291.3464 | 417 | 165.9718 | 508 | 132.809 | 530 | SPI1 | None | None | GSM14210 | 99159 | 50365 |
| 177.9604 | 291 | 248.165 | 373 | 157.3333 | 469 | 112.477 | 475 | SPI1 | None | None | GSM14210 | 90172 | 50367 |
| 1231.4 | 417 | 1251.357 | 530 | 943.1073 | 391 | 878.2446 | 369 | SUZ12 | None | Fetal Liver | GSM14270 | 10147 | 50372 |
| 1186.213 | 163 | 1167.368 | 187 | 643.9494 | 147 | 708.4986 | 164 | EZH1 | None | Fetal Liver | GSM14270 | 3586 | 50373 |
| 1042.391 | 147 | 1152.672 | 188 | 629.5473 | 150 | 491.3986 | 139 | EZH1 | None | Fetal Liver | GSM14270 | 4017 | 50374 |
| 181.3052 | 35 | 247.1185 | 46 | 125.9609 | 40 | 196.3948 | 54 | EZH2 | None | Fetal Liver | GSM14270 | 1488 | 50375 |
| 166.5387 | 97 | 186.1636 | 115 | 75.04081 | 111 | 124.0428 | 145 | EZH2 | None | Fetal Liver | GSM14270 | 14589 | 50376 |
| 258.4412 | 40 | 389.7418 | 56 | 99.46736 | 32 | 153.5199 | 43 | EZH1 | None | Fetal Liver | GSM14270 | 1217 | 50378 |
| 125.1736 | 29 | 147.1437 | 35 | 102.1502 | 37 | 172.8744 | 52 | EZH2 | None | Fetal Liver | GSM14270 | 1593 | 50379 |
| 1037.731 | 713 | 1083.38 | 918 | 760.6854 | 788 | 702.5679 | 792 | NFE2 | None | Fetal Liver | GSM14270 | 48280 | 50381 |
| 173.4623 | 153 | 384.735 | 241 | 255.4177 | 281 | 199.0854 | 279 | TAL1 | None | Fetal Liver | GSM14270 | 31409 | 50382 |
| 618.0492 | 318 | 734.9712 | 451 | 465.1316 | 437 | 427.1892 | 460 | CTCF | GM12878 | Blood | GSM48929 | 46434 | 5039 |

| | | | | | | | | | | | | | |
|---|---|---|---|---|---|---|---|---|---|---|---|---|---|
| 1180.665 | 172 | 1175.104 | 206 | 659.764 | 154 | 607.6143 | 156 | MED1 | Jurkat | Blood | GSM14420 | 3723 | 50396 |
| 860.394 | 482 | 887.3192 | 617 | 633.2203 | 539 | 556.0768 | 503 | MYB | Jurkat | Blood | GSM14420 | 41400 | 50397 |
| 865.979 | 602 | 926.0019 | 806 | 619.2516 | 673 | 566.7394 | 679 | MYB | Jurkat | Blood | GSM14420 | 58783 | 50398 |
| 553.6118 | 286 | 704.807 | 382 | 381.6093 | 380 | 408.9598 | 423 | CTCF | GM12878 | Blood | GSM48929 | 41258 | 5040 |
| 7.643557 | 12 | 9.241432 | 15 | 17.88615 | 28 | 64.85687 | 51 | CREBBP | hMADS-3 | Adipose | GSM14438 | 3890 | 50402 |
| 17.22203 | 12 | 4.953676 | 8 | 21.06239 | 21 | 15.36501 | 20 | CREBBP | hMADS-3 | Adipose | GSM14438 | 2255 | 50403 |
| 178.8999 | 264 | 321.8255 | 370 | 162.2942 | 426 | 248.9126 | 528 | PPARG | hMADS-3 | Adipose | GSM14438 | 77617 | 50408 |
| 39.27411 | 153 | 68.16757 | 208 | 63.81276 | 304 | 112.2801 | 386 | PPARG | hMADS-3 | Adipose | GSM14438 | 67738 | 50409 |
| 690.789 | 358 | 732.3396 | 485 | 469.6404 | 465 | 489.7288 | 513 | CTCF | GM12878 | Blood | GSM48929 | 51459 | 5041 |
| 742.2473 | 510 | 784.6248 | 677 | 598.38 | 710 | 554.6199 | 737 | PPARG | hMADS-3 | Adipose | GSM14438 | 69909 | 50411 |
| 1043.333 | 671 | 1079.416 | 857 | 760.7378 | 714 | 730.977 | 760 | KLF11 | hMADS-3 | Adipose | GSM14438 | 40720 | 50412 |
| 1025.152 | 618 | 1076.461 | 810 | 746.5384 | 642 | 721.7723 | 694 | KLF11 | hMADS-3 | Adipose | GSM14438 | 37028 | 50413 |
| 1058.37 | 564 | 1108.39 | 743 | 795.9358 | 594 | 760.3273 | 622 | KLF11 | hMADS-3 | Adipose | GSM14438 | 28367 | 50414 |
| 1056.571 | 555 | 1118.486 | 746 | 801.4052 | 594 | 745.1031 | 589 | KLF11 | hMADS-3 | Adipose | GSM14438 | 27961 | 50415 |
| 901.803 | 539 | 956.6726 | 725 | 658.2396 | 597 | 642.9368 | 656 | CTCF | GM12891 | Blood | GSM48929 | 45803 | 5042 |
| 6.540298 | 30 | 6.213521 | 35 | 15.60667 | 68 | 6.757851 | 61 | DMC1 | None | Testis | GSM14473 | 14487 | 50422 |
| 892.6624 | 563 | 933.9334 | 737 | 667.5917 | 659 | 633.0337 | 695 | CTCF | GM12891 | Blood | GSM48929 | 51085 | 5043 |
| 1175.992 | 371 | 1207.112 | 479 | 893.2975 | 342 | 854.1142 | 351 | ELK3 | None | Umbilical V | GSM14666 | 10494 | 50434 |
| 888.3156 | 634 | 934.0292 | 825 | 653.9008 | 741 | 623.6411 | 794 | CTCF | GM12892 | Blood | GSM48929 | 62952 | 5044 |
| 911.5918 | 610 | 956.3852 | 794 | 664.6986 | 689 | 634.1672 | 733 | CTCF | GM12892 | Blood | GSM48929 | 54654 | 5045 |
| 334.9761 | 53 | 526.476 | 76 | 276.2188 | 65 | 580.2224 | 105 | AUTS2 | 293T-Rex | None | GSM14792 | 1697 | 50456 |
| 931.2472 | 602 | 985.9185 | 799 | 690.9637 | 691 | 655.441 | 727 | CTCF | GM19240 | Blood | GSM48929 | 50239 | 5046 |
| 797.6513 | 453 | 862.4826 | 630 | 603.7414 | 552 | 583.8647 | 601 | CTCF | GM19240 | Blood | GSM48929 | 49543 | 5047 |
| 872.8508 | 576 | 934.9204 | 781 | 639.7984 | 665 | 606.446 | 701 | CTCF | GM19238 | Blood | GSM48929 | 56467 | 5048 |
| 849.8735 | 532 | 901.6495 | 714 | 642.5984 | 646 | 598.2504 | 665 | CTCF | GM19238 | Blood | GSM48930 | 53742 | 5049 |
| 897.9049 | 620 | 946.7747 | 816 | 655.2115 | 714 | 631.1813 | 772 | CTCF | GM19239 | Blood | GSM48930 | 59303 | 5050 |
| 797.1512 | 341 | 842.8734 | 466 | 608.2727 | 415 | 588.6013 | 449 | CTCF | GM19239 | Blood | GSM48930 | 34176 | 5051 |
| 952.68 | 436 | 956.529 | 533 | 679.1633 | 424 | 646.1362 | 443 | NIPBL | Ramos | None | GSM15191 | 26883 | 50516 |
| 879.1598 | 362 | 900.9148 | 464 | 579.2119 | 361 | 612.6613 | 400 | NIPBL | Ramos | None | GSM15191 | 26967 | 50517 |
| 932.2731 | 555 | 973.885 | 723 | 676.8939 | 593 | 619.9762 | 587 | MYB | Jurkat | Blood | GSM15196 | 41184 | 50565 |
| 942.9859 | 528 | 968.9397 | 674 | 691.6078 | 562 | 624.2159 | 539 | MYB | Jurkat | Blood | GSM15196 | 35296 | 50569 |
| 964.8639 | 519 | 993.1852 | 665 | 713.4014 | 557 | 635.3245 | 520 | MYB | Jurkat | Blood | GSM15196 | 33288 | 50571 |
| 1001.766 | 635 | 1035.811 | 810 | 735.9076 | 690 | 665.6501 | 669 | MYB | MOLT-3 | Peripheral | GSM15196 | 42270 | 50573 |
| 720.934 | 333 | 717.1416 | 404 | 432.0164 | 403 | 394.3224 | 423 | TAL1 | MOLT-3 | Peripheral | GSM15196 | 40746 | 50576 |
| 228.9909 | 138 | 318.7627 | 180 | 220.4397 | 210 | 222.3498 | 228 | MED1 | hMADS-3 | Adipose | GSM15376 | 20812 | 50690 |
| 244.4154 | 201 | 365.2208 | 268 | 169.5138 | 281 | 237.5299 | 343 | MED1 | hMADS-3 | Adipose | GSM15376 | 39383 | 50691 |
| 519.7517 | 257 | 708.9096 | 338 | 350.5341 | 334 | 432.2826 | 392 | MED1 | hMADS-3 | Adipose | GSM15376 | 33837 | 50692 |
| 302.3072 | 215 | 361.6827 | 262 | 238.629 | 309 | 263.5047 | 348 | MED1 | hMADS-3 | Adipose | GSM15376 | 37949 | 50693 |
| 66.59637 | 22 | 74.56722 | 26 | 45.2128 | 27 | 83.92967 | 39 | GTF2I | None | None | GSM15390 | 1923 | 50694 |
| 43.04971 | 47 | 43.36336 | 54 | 41.35887 | 76 | 44.64871 | 85 | GTF2I | None | None | GSM15390 | 11107 | 50695 |
| 41.97952 | 19 | 57.65194 | 25 | 42.40291 | 29 | 68.79038 | 39 | GTF2I | None | None | GSM15390 | 2244 | 50697 |
| 655.8827 | 205 | 817.005 | 254 | 349.617 | 225 | 342.1174 | 240 | GTF2I | None | None | GSM15390 | 16584 | 50698 |
| 1045.584 | 159 | 1097.485 | 193 | 413.8771 | 131 | 605.6651 | 170 | GTF2I | None | None | GSM15390 | 4815 | 50699 |
| 378.1921 | 119 | 466.3542 | 147 | 195.372 | 128 | 213.2656 | 143 | GTF2I | None | None | GSM15390 | 9411 | 50700 |
| 61.99642 | 34 | 95.54108 | 47 | 30.19251 | 39 | 39.57143 | 47 | GTF2I | None | None | GSM15390 | 4630 | 50708 |
| 83.51081 | 55 | 116.2953 | 72 | 55.71088 | 73 | 41.05809 | 71 | GTF2I | None | None | GSM15390 | 8517 | 50709 |
| 51.78963 | 47 | 68.26934 | 60 | 51.12837 | 75 | 45.11857 | 78 | GTF2I | None | None | GSM15390 | 9455 | 50712 |
| 33.22028 | 25 | 66.95911 | 39 | 47.56691 | 46 | 45.86688 | 49 | GTF2I | None | None | GSM15390 | 4556 | 50713 |
| 79.97782 | 39 | 73.31488 | 42 | 39.27849 | 44 | 53.66715 | 54 | GTF2I | None | None | GSM15390 | 4769 | 50714 |
| 64.63336 | 29 | 88.55769 | 38 | 64.9475 | 44 | 55.02801 | 44 | GTF2I | None | None | GSM15390 | 3346 | 50715 |
| 362.6083 | 71 | 264.097 | 65 | 143.8621 | 61 | 138.7849 | 64 | UBTF | None | None | GSM15445 | 3018 | 50729 |
| 97.90068 | 400 | 194.6237 | 552 | 109.2603 | 727 | 101.8322 | 798 | SPI1 | None | None | GSM15445 | 201889 | 50730 |
| 86.25542 | 23 | 191.1652 | 40 | 127.9726 | 41 | 165.624 | 50 | EHF | Calu-3 | Lung | GSM15480 | 1569 | 50820 |
| 2.787192 | 5 | 5.903553 | 8 | 3.672034 | 10 | 67.17213 | 36 | MITF | 501-Mel | None | GSM15650 | 2047 | 50853 |
| 150.7674 | 32 | 322.5802 | 55 | 108.6955 | 38 | 174.7476 | 52 | RELA | KB Cell | HeLa conta | GSM15665 | 1612 | 50875 |
| 1265.41 | 616 | 1306.508 | 794 | 926.6518 | 551 | 883.3069 | 562 | SPI1 | KB Cell | HeLa conta | GSM15665 | 15625 | 50876 |
| 1252.842 | 600 | 1291.592 | 770 | 917.7391 | 534 | 888.599 | 564 | SPI1 | KB Cell | HeLa conta | GSM15665 | 15141 | 50877 |
| 1249.192 | 646 | 1307.874 | 845 | 915.3141 | 586 | 885.2155 | 623 | SPI1 | KB Cell | HeLa conta | GSM15665 | 17608 | 50878 |
| 1236.301 | 657 | 1293.368 | 852 | 907.6245 | 606 | 868.698 | 629 | SPI1 | KB Cell | HeLa conta | GSM15665 | 18505 | 50879 |
| 1163.595 | 409 | 1200.97 | 538 | 882.195 | 391 | 838.874 | 395 | RELA | KB Cell | HeLa conta | GSM15665 | 12641 | 50892 |
| 1082.874 | 335 | 1129.913 | 455 | 824.8485 | 329 | 804.9339 | 353 | RELA | KB Cell | HeLa conta | GSM15665 | 12465 | 50893 |
| 1270.101 | 403 | 1339.16 | 570 | 978.6311 | 376 | 952.9393 | 401 | SPI1 | KB Cell | HeLa conta | GSM15665 | 8821 | 50894 |
| 49.23602 | 44 | 41.8568 | 47 | 19.92143 | 52 | 81.34493 | 93 | OTX2 | 6540 cells | None | GSM15670 | 9064 | 50896 |
| 136.5764 | 43 | 160.598 | 52 | 127.8141 | 61 | 203.8871 | 82 | MITF | 6540 cells | None | GSM15670 | 3463 | 50897 |
| 1077.869 | 277 | 1103.347 | 363 | 816.027 | 269 | 794.0717 | 288 | PAX6 | 6540 cells | None | GSM15670 | 10244 | 50898 |
| 48.04951 | 59 | 36.0244 | 61 | 36.79487 | 89 | 53.89859 | 110 | LHX2 | 6540 cells | None | GSM15670 | 14642 | 50899 |
| 235.5247 | 75 | 380.017 | 107 | 148.0577 | 88 | 218.7274 | 113 | NR2F2 | WI-38 | Lung | GSM11268 | 5959 | 50972 |
| 584.1203 | 90 | 1092.237 | 143 | 379.5972 | 96 | 334.5608 | 95 | NR2C2 | WI-38 | Lung | GSM11268 | 2670 | 50973 |
| 460.1123 | 63 | 692.5749 | 88 | 203.0473 | 53 | 333.1492 | 74 | NR2C2 | HeLa | Cervix | GSM11268 | 1531 | 50977 |
| 1008.715 | 224 | 1035.242 | 295 | 881.6437 | 317 | 855.8044 | 330 | GATA6 | PaTu8988S | Pancreatic | GSM11516 | 9910 | 50979 |
| 1218.595 | 156 | 1227.899 | 198 | 701.6053 | 146 | 707.5908 | 155 | TET2 | HCT-116 | Colon | GSM11528 | 3264 | 50981 |
| 619.3895 | 505 | 682.37 | 697 | 470.7628 | 705 | 457.5404 | 781 | CTCF | H1 | Embryo | GSM12672 | 104467 | 50987 |

| | | | | | | | | | | | | | |
|---|---|---|---|---|---|---|---|---|---|---|---|---|---|
| 651.0761 | 501 | 733.6033 | 714 | 497.8212 | 687 | 484.4884 | 759 | CTCF | H1 | Embryo | GSM12672 | 92520 | 50988 |
| 711.3772 | 348 | 786.8098 | 510 | 519.2809 | 455 | 508.7459 | 490 | CTCF | H1 | Embryo | GSM12672 | 45621 | 50989 |
| 724.4942 | 410 | 778.9094 | 564 | 540.6909 | 513 | 531.3959 | 571 | CTCF | H1 | Embryo | GSM12672 | 54943 | 50990 |
| 667.7047 | 485 | 728.2515 | 668 | 509.3897 | 653 | 496.5362 | 723 | CTCF | H1 | Embryo | GSM12672 | 82467 | 50991 |
| 687.9069 | 536 | 635.0376 | 582 | 601.2906 | 843 | 658.6734 | 1080 | AR | VCaP | Prostate | GSM13548 | 90391 | 51048 |
| 688.2215 | 372 | 671.7888 | 438 | 604.6321 | 594 | 674.3244 | 796 | AR | VCaP | Prostate | GSM13548 | 54128 | 51049 |
| 1112.66 | 299 | 1062.188 | 319 | 1022.944 | 459 | 1053.159 | 564 | PIAS1 | VCaP | Prostate | GSM13548 | 9898 | 51050 |
| 1147.849 | 201 | 1077.863 | 199 | 1073.562 | 327 | 1110.593 | 408 | PIAS1 | VCaP | Prostate | GSM13548 | 5505 | 51051 |
| 906.5722 | 226 | 852.0081 | 240 | 764.9624 | 305 | 841.8461 | 435 | PIAS1 | VCaP | Prostate | GSM13548 | 14091 | 51052 |
| 994.3593 | 952 | 931.5046 | 1065 | 823.7836 | 1357 | 854.3552 | 1610 | FOXA1 | VCaP | Prostate | GSM13548 | 118320 | 51054 |
| 1014.532 | 975 | 933.5596 | 1075 | 834.9793 | 1379 | 861.9382 | 1631 | FOXA1 | VCaP | Prostate | GSM13548 | 119444 | 51055 |
| 879.3433 | 879 | 848.1974 | 1016 | 759.042 | 1317 | 787.5987 | 1564 | FOXA1 | VCaP | Prostate | GSM13548 | 132978 | 51056 |
| 890.316 | 859 | 829.6957 | 953 | 747.3106 | 1252 | 783.7298 | 1503 | FOXA1 | VCaP | Prostate | GSM13548 | 121313 | 51057 |
| 1139.268 | 454 | 1189.564 | 601 | 853.7969 | 405 | 814.4219 | 419 | CHD8 | None | None | GSM13812 | 11664 | 51103 |
| 613.09 | 90 | 691.1584 | 105 | 416.604 | 97 | 250.6524 | 78 | CHD8 | None | None | GSM13812 | 2333 | 51104 |
| 230.1564 | 132 | 366.0641 | 183 | 191.6171 | 188 | 192.836 | 204 | BCL6 | None | None | GSM14389 | 18568 | 51245 |
| 323.4894 | 198 | 485.7848 | 266 | 245.536 | 275 | 230.8681 | 292 | TCF7L1 | None | None | GSM14389 | 30324 | 51246 |
| 616.5148 | 209 | 774.6081 | 260 | 278.5055 | 217 | 317.1574 | 247 | PBX1 | None | None | GSM14389 | 18429 | 51248 |
| 210.3212 | 156 | 386.3761 | 228 | 183.5363 | 231 | 125.0237 | 220 | EP300 | None | None | GSM14389 | 26379 | 51249 |
| 0 | 2 | 0 | 3 | 0.071235 | 7 | 23.06317 | 30 | KDM1B | HepG2 | Liver | GSM14429 | 3319 | 51251 |
| 654.1209 | 437 | 611.1384 | 483 | 581.5872 | 714 | 660.6533 | 962 | AR | VCaP | Prostate | GSM14634 | 74696 | 51267 |
| 658.3266 | 447 | 620.2968 | 498 | 578.4384 | 716 | 656.2445 | 964 | AR | VCaP | Prostate | GSM14634 | 75718 | 51268 |
| 770.5428 | 717 | 686.2208 | 753 | 666.7774 | 1097 | 714.67 | 1351 | FOXA1 | VCaP | Prostate | GSM14634 | 116327 | 51269 |
| 772.4164 | 772 | 687.9941 | 817 | 670.0025 | 1183 | 701.4854 | 1416 | FOXA1 | VCaP | Prostate | GSM14634 | 134352 | 51270 |
| 769.409 | 746 | 696.8627 | 805 | 646.6408 | 1113 | 704.344 | 1388 | FOXA1 | VCaP | Prostate | GSM14634 | 127316 | 51271 |
| 784.7328 | 804 | 709.2943 | 870 | 676.3364 | 1226 | 701.9147 | 1454 | FOXA1 | VCaP | Prostate | GSM14634 | 142675 | 51272 |
| 1059.643 | 392 | 1004.386 | 419 | 902.5795 | 516 | 916.8899 | 606 | ASH2L | VCaP | Prostate | GSM14899 | 16600 | 51332 |
| 1022.587 | 459 | 1031.876 | 568 | 864.8481 | 606 | 894.1411 | 743 | ASH2L | VCaP | Prostate | GSM14899 | 23375 | 51333 |
| 13.51079 | 44 | 16.78188 | 55 | 20.70878 | 88 | 32.23016 | 109 | PSIP1 | None | None | GSM14954 | 18551 | 51360 |
| 7.239547 | 11 | 17.11124 | 18 | 3.632256 | 15 | 43.30271 | 40 | PRDM1 | HUES64 | Embryo | GSM15056 | 3251 | 51434 |
| 225.8607 | 34 | 440.0506 | 56 | 111.3541 | 31 | 499.4012 | 78 | PRDM1 | HUES64 | Embryo | GSM15056 | 1012 | 51436 |
| 613.4203 | 398 | 676.7758 | 545 | 407.9514 | 535 | 467.4477 | 611 | CTCF | HUES64 | Embryo | GSM15056 | 68278 | 51438 |
| 121.9057 | 63 | 230.7624 | 96 | 41.98558 | 63 | 57.75573 | 77 | CTCF | HUES64 | Embryo | GSM15056 | 8096 | 51439 |
| 484.5451 | 176 | 824.6753 | 258 | 340.286 | 222 | 371.0437 | 248 | CTCF | HUES64 | Embryo | GSM15056 | 16517 | 51441 |
| 274.7116 | 173 | 476.421 | 248 | 182.9091 | 230 | 210.903 | 263 | CTCF | HUES64 | Embryo | GSM15056 | 27130 | 51442 |
| 466.9765 | 187 | 748.6899 | 262 | 262.5687 | 219 | 427.5434 | 289 | CTCF | HUES64 | Embryo | GSM15056 | 19908 | 51443 |
| 786.5422 | 254 | 874.5704 | 370 | 446.479 | 290 | 533.9426 | 337 | CTCF | HUES64 | Embryo | GSM15056 | 22143 | 51444 |
| 60.96414 | 31 | 76.82627 | 39 | 32.88846 | 37 | 164.8335 | 81 | EOMES | HUES64 | Embryo | GSM15056 | 4228 | 51445 |
| 22.31835 | 26 | 43.01712 | 39 | 23.48322 | 44 | 26.47155 | 50 | EOMES | HUES64 | Embryo | GSM15056 | 6620 | 51447 |
| 36.51728 | 46 | 67.23548 | 67 | 39.05774 | 78 | 87.59874 | 113 | EOMES | HUES64 | Embryo | GSM15056 | 11859 | 51448 |
| 6.021707 | 5 | 11.90973 | 8 | 1.590811 | 5 | 1.978856 | 6 | EOMES | HUES64 | Embryo | GSM15056 | 1218 | 51449 |
| 268.3241 | 324 | 240.2439 | 359 | 191.3677 | 480 | 256.1099 | 572 | FOXA1 | HUES64 | Embryo | GSM15056 | 86453 | 51450 |
| 49.93086 | 24 | 42.5895 | 25 | 53.48265 | 38 | 135.5793 | 63 | FOXA1 | HUES64 | Embryo | GSM15056 | 3086 | 51453 |
| 164.8068 | 153 | 142.2614 | 165 | 131.6459 | 227 | 218.5189 | 294 | FOXA2 | HUES64 | Embryo | GSM15056 | 32154 | 51455 |
| 27.28288 | 14 | 31.65637 | 17 | 15.30368 | 17 | 16.4611 | 19 | FOXA2 | HUES64 | Embryo | GSM15056 | 2005 | 51457 |
| 139.4703 | 104 | 125.3872 | 113 | 144.8938 | 167 | 238.0031 | 219 | FOXA2 | HUES64 | Embryo | GSM15056 | 18955 | 51459 |
| 58.842 | 48 | 168.1332 | 87 | 55.13194 | 75 | 185.0735 | 132 | GATA4 | HUES64 | Embryo | GSM15056 | 9322 | 51463 |
| 17.23217 | 23 | 53.45497 | 42 | 19.32346 | 40 | 90.18165 | 78 | GATA4 | HUES64 | Embryo | GSM15056 | 6204 | 51464 |
| 32.62732 | 34 | 49.25457 | 46 | 18.7781 | 46 | 88.66692 | 88 | GATA4 | HUES64 | Embryo | GSM15056 | 7743 | 51465 |
| 337.0312 | 150 | 457.0354 | 194 | 200.5914 | 183 | 258.9408 | 219 | GATA4 | HUES64 | Embryo | GSM15056 | 17899 | 51466 |
| 374.6203 | 145 | 450.3345 | 177 | 196.4225 | 164 | 221.9018 | 186 | GATA4 | HUES64 | Embryo | GSM15056 | 14930 | 51467 |
| 218.142 | 108 | 228.7236 | 124 | 151.874 | 142 | 179.4433 | 164 | GATA4 | HUES64 | Embryo | GSM15056 | 13970 | 51468 |
| 119.1433 | 60 | 207.6612 | 88 | 186.9863 | 112 | 273.7227 | 143 | GATA4 | HUES64 | Embryo | GSM15056 | 7797 | 51469 |
| 72.12145 | 19 | 127.0886 | 29 | 64.75147 | 26 | 44.49192 | 23 | GATA4 | HUES64 | Embryo | GSM15056 | 1281 | 51470 |
| 224.334 | 61 | 314.1702 | 81 | 157.7141 | 74 | 307.2139 | 110 | GATA4 | HUES64 | Embryo | GSM15056 | 4143 | 51471 |
| 333.0121 | 115 | 392.5088 | 139 | 254.3565 | 150 | 329.8589 | 181 | GATA4 | HUES64 | Embryo | GSM15056 | 10374 | 51472 |
| 2.123151 | 24 | 9.142658 | 41 | 16.11415 | 72 | 26.67845 | 91 | GATA6 | HUES64 | Embryo | GSM15056 | 15259 | 51476 |
| 1.255773 | 7 | 6.9398 | 14 | 4.212668 | 18 | 5.094271 | 21 | GATA6 | HUES64 | Embryo | GSM15056 | 4150 | 51478 |
| 22.64211 | 27 | 24.53538 | 32 | 21.69196 | 44 | 56.82107 | 68 | GATA6 | HUES64 | Embryo | GSM15056 | 6737 | 51480 |
| 14.76826 | 24 | 31.46709 | 37 | 44.1 | 60 | 141.8688 | 104 | GATA6 | HUES64 | Embryo | GSM15056 | 7517 | 51481 |
| 4.564426 | 13 | 4.288371 | 15 | 6.66072 | 26 | 4.666934 | 26 | GATA6 | HUES64 | Embryo | GSM15056 | 5435 | 51482 |
| 14.6427 | 10 | 29.74565 | 16 | 7.128949 | 12 | 82.17252 | 38 | HNF1B | HUES64 | Embryo | GSM15056 | 1838 | 51498 |
| 144.9203 | 35 | 213.3216 | 48 | 72.01563 | 35 | 297.8335 | 78 | KLF5 | HUES64 | Embryo | GSM15056 | 2033 | 51508 |
| 238.868 | 42 | 374.4465 | 60 | 146.1949 | 44 | 260.1057 | 64 | NR5A2 | HUES64 | Embryo | GSM15056 | 1498 | 51512 |
| 65.55639 | 82 | 81.70817 | 102 | 17.70186 | 92 | 54.03847 | 138 | NANOG | HUES64 | Embryo | GSM15056 | 20141 | 51513 |
| 752.0051 | 365 | 788.6269 | 485 | 364.8888 | 382 | 407.8991 | 432 | NANOG | HUES64 | Embryo | GSM15056 | 42310 | 51514 |
| 23.57614 | 64 | 58.76199 | 100 | 12.8863 | 96 | 18.90093 | 116 | NANOG | HUES64 | Embryo | GSM15056 | 23529 | 51516 |
| 135.0369 | 82 | 157.3426 | 99 | 57.75092 | 92 | 23.56854 | 75 | NANOG | HUES64 | Embryo | GSM15057 | 12271 | 51518 |
| 16.70409 | 38 | 30.26451 | 54 | 10.08666 | 58 | 40.0172 | 94 | NANOG | HUES64 | Embryo | GSM15057 | 13506 | 51520 |
| 698.0223 | 492 | 708.29 | 611 | 499.8006 | 605 | 479.6263 | 656 | OTX2 | HUES64 | Embryo | GSM15057 | 76163 | 51522 |
| 383.5471 | 175 | 476.9222 | 217 | 224.8949 | 213 | 291.1514 | 255 | OTX2 | HUES64 | Embryo | GSM15057 | 21334 | 51524 |
| 380.8283 | 95 | 329.5719 | 97 | 151.942 | 87 | 246.8631 | 117 | OTX2 | HUES64 | Embryo | GSM15057 | 5795 | 51525 |

| | | | | | | | | | | | | | |
|---|---|---|---|---|---|---|---|---|---|---|---|---|---|
| 55.50147 | 25 | 81.60239 | 34 | 22.96557 | 26 | 10.51655 | 21 | OTX2 | HUES64 | Embryo | GSM15057 | 2953 | 51529 |
| 39.91344 | 24 | 42.73696 | 28 | 12.64326 | 24 | 12.35489 | 26 | OTX2 | HUES64 | Embryo | GSM15057 | 3709 | 51530 |
| 5.83196 | 5 | 30.16739 | 13 | 1.50756 | 5 | 81.98996 | 31 | PAX6 | HUES64 | Embryo | GSM15057 | 1245 | 51532 |
| 94.65371 | 21 | 210.6827 | 37 | 49.27439 | 21 | 70.62027 | 27 | POU5F1 | HUES64 | Embryo | GSM15057 | 1077 | 51542 |
| 144.1405 | 28 | 171.8634 | 34 | 112.7729 | 34 | 95.17414 | 33 | POU5F1 | HUES64 | Embryo | GSM15057 | 1172 | 51543 |
| 161.4705 | 106 | 209.1181 | 134 | 39.90528 | 101 | 74.33812 | 135 | POU5F1 | HUES64 | Embryo | GSM15057 | 16926 | 51546 |
| 40.72977 | 15 | 46.68541 | 18 | 10.21359 | 12 | 0.998308 | 6 | SALL4 | HUES64 | Embryo | GSM15057 | 1481 | 51551 |
| 813.436 | 138 | 1078.729 | 185 | 454.3721 | 139 | 473.8287 | 151 | SMAD1 | HUES64 | Embryo | GSM15057 | 4993 | 51554 |
| 55.50204 | 15 | 64.01427 | 18 | 11.25062 | 10 | 152.7687 | 40 | SMAD1 | HUES64 | Embryo | GSM15057 | 1055 | 51556 |
| 33.86903 | 14 | 48.89823 | 19 | 4.453281 | 9 | 138.3818 | 46 | SMAD1 | HUES64 | Embryo | GSM15057 | 1598 | 51557 |
| 12.54909 | 10 | 25.8302 | 16 | 25.39454 | 22 | 166.0926 | 58 | SMAD1 | HUES64 | Embryo | GSM15057 | 2133 | 51559 |
| 39.24631 | 19 | 76.98593 | 30 | 61.88368 | 36 | 223.9562 | 73 | SMAD1 | HUES64 | Embryo | GSM15057 | 2462 | 51560 |
| 37.85873 | 16 | 42.38359 | 19 | 22.03452 | 19 | 5.711394 | 12 | SMAD1 | HUES64 | Embryo | GSM15057 | 1830 | 51563 |
| 59.67561 | 24 | 40.01426 | 22 | 13.54538 | 19 | 75.35961 | 43 | SMAD1 | HUES64 | Embryo | GSM15057 | 2470 | 51564 |
| 799.1422 | 87 | 925.6867 | 103 | 393.2705 | 75 | 286.4838 | 66 | SMAD4 | HUES64 | Embryo | GSM15057 | 1452 | 51574 |
| 9.145692 | 7 | 21.13653 | 12 | 3.627751 | 8 | 5.125548 | 10 | SOX17 | HUES64 | Embryo | GSM15057 | 1518 | 51581 |
| 6.715408 | 10 | 10.72885 | 14 | 7.538481 | 18 | 42.47766 | 38 | SOX2 | HUES64 | Embryo | GSM15057 | 3204 | 51582 |
| 18.56627 | 10 | 46.18944 | 18 | 3.581932 | 8 | 156.553 | 48 | SOX2 | HUES64 | Embryo | GSM15057 | 1515 | 51585 |
| 11.92478 | 12 | 5.33272 | 10 | 2.413057 | 12 | 45.28826 | 37 | SOX2 | HUES64 | Embryo | GSM15057 | 2903 | 51586 |
| 1309.419 | 224 | 1334.564 | 289 | 1000.057 | 187 | 1025.315 | 224 | SP1 | HUES64 | Embryo | GSM15057 | 3636 | 51587 |
| 667.0341 | 389 | 712.7325 | 527 | 442.0183 | 501 | 502.7992 | 575 | SRF | HUES64 | Embryo | GSM15057 | 59136 | 51590 |
| 1054.331 | 516 | 1104.416 | 686 | 761.8852 | 497 | 729.8269 | 518 | SRF | HUES64 | Embryo | GSM15057 | 24428 | 51591 |
| 478.9099 | 141 | 591.6236 | 174 | 187.8063 | 132 | 187.3179 | 142 | SRF | HUES64 | Embryo | GSM15057 | 9892 | 51592 |
| 216.8639 | 74 | 347.9736 | 105 | 167.7228 | 96 | 80.45403 | 75 | SRF | HUES64 | Embryo | GSM15057 | 6185 | 51594 |
| 429.3318 | 117 | 596.1256 | 154 | 207.5032 | 119 | 316.2093 | 155 | SRF | HUES64 | Embryo | GSM15057 | 7834 | 51595 |
| 684.3987 | 202 | 906.6617 | 302 | 430.7893 | 237 | 520.749 | 277 | SRF | HUES64 | Embryo | GSM15057 | 15599 | 51596 |
| 26.20148 | 10 | 95.98005 | 23 | 32.86087 | 17 | 228.4886 | 51 | T | HUES64 | Embryo | GSM15057 | 1110 | 51600 |
| 96.49167 | 22 | 62.64631 | 19 | 19.40043 | 14 | 233.0059 | 54 | T | HUES64 | Embryo | GSM15057 | 1218 | 51602 |
| 324.881 | 46 | 546.8344 | 69 | 171.9519 | 43 | 450.0256 | 79 | TAL1 | HUES64 | Embryo | GSM15057 | 1185 | 51603 |
| 176.1536 | 35 | 334.1874 | 56 | 74.88095 | 31 | 119.1019 | 42 | TAL1 | HUES64 | Embryo | GSM15057 | 1520 | 51604 |
| 39.85556 | 33 | 101.8147 | 57 | 20.99209 | 42 | 115.0792 | 88 | TCF4 | HUES64 | Embryo | GSM15057 | 6490 | 51609 |
| 251.6884 | 53 | 555.5217 | 92 | 102.2581 | 46 | 309.9126 | 87 | THAP11 | HUES64 | Embryo | GSM15057 | 2259 | 51611 |
| 2.821623 | 5 | 7.901629 | 9 | 0 | 3 | 0.828427 | 7 | TRIM28 | HUES64 | Embryo | GSM15057 | 1932 | 51616 |
| 0 | 1 | 17.68596 | 10 | 0.570796 | 4 | 0.073775 | 3 | TRIM28 | HUES64 | Embryo | GSM15057 | 1248 | 51617 |
| 1144.883 | 349 | 1199.649 | 484 | 886.0726 | 338 | 858.0826 | 354 | MYC | HUES64 | Embryo | GSM15058 | 9915 | 51627 |
| 291.9431 | 46 | 310.6087 | 52 | 142.4088 | 42 | 163.2609 | 48 | MYC | HUES64 | Embryo | GSM15058 | 1428 | 51628 |
| 2.694502 | 3 | 18.21767 | 9 | 2.266666 | 5 | 1.737593 | 5 | HAND1 | HUES64 | Embryo | GSM15058 | 1038 | 51631 |
| 840.5889 | 280 | 821.7282 | 328 | 602.7711 | 333 | 775.7105 | 532 | AR | LNCaP | Prostate | GSM15278 | 22667 | 51737 |
| 806.1694 | 414 | 785.2247 | 481 | 659.6207 | 561 | 730.3551 | 760 | AR | LNCaP | Prostate | GSM15278 | 40506 | 51748 |
| 836.626 | 377 | 808.6163 | 432 | 664.4471 | 480 | 753.5021 | 681 | AR | LNCaP | Prostate | GSM15278 | 32612 | 51749 |
| 117.8838 | 82 | 130.572 | 97 | 75.63067 | 109 | 97.8114 | 130 | FOXP1 | LNCaP | Prostate | GSM15278 | 13861 | 51750 |
| 865.217 | 285 | 850.0538 | 333 | 690.6762 | 351 | 780.7366 | 511 | AR | LNCaP | Prostate | GSM15278 | 20703 | 51757 |
| 779.1142 | 191 | 891.041 | 227 | 533.9546 | 226 | 815.9127 | 343 | AR | LNCaP | Prostate | GSM15278 | 11498 | 51758 |
| 507.0153 | 118 | 643.1744 | 148 | 251.3529 | 118 | 786.2823 | 223 | AR | LNCaP | Prostate | GSM15437 | 6510 | 51837 |
| 1239.533 | 397 | 1233.208 | 472 | 986.0113 | 405 | 991.3583 | 473 | MRE11A | LNCaP | Prostate | GSM15437 | 9609 | 51838 |
| 1237.771 | 525 | 1209.819 | 598 | 990.9877 | 570 | 976.093 | 626 | MRE11A | LNCaP | Prostate | GSM15437 | 14470 | 51839 |
| 862.3056 | 391 | 803.132 | 411 | 673.4166 | 470 | 723.24 | 607 | TOP1 | LNCaP | Prostate | GSM15437 | 30854 | 51840 |
| 871.4106 | 444 | 839.2071 | 504 | 679.0344 | 541 | 758.6409 | 749 | TOP1 | LNCaP | Prostate | GSM15437 | 36862 | 51841 |
| 823.817 | 578 | 831.1562 | 708 | 597.8369 | 676 | 558.7982 | 704 | HSF1 | 451Lu | None | GSM15532 | 64770 | 51863 |
| 1078.415 | 463 | 1098.486 | 587 | 785.4009 | 438 | 751.2359 | 457 | HSF1 | 451Lu | None | GSM15532 | 19938 | 51864 |
| 998.8915 | 258 | 950.7219 | 279 | 857.7806 | 349 | 864.8885 | 408 | SUMO2 | VCaP | Prostate | GSM15555 | 12173 | 51866 |
| 1159.867 | 847 | 1126.294 | 972 | 933.0868 | 1075 | 938.8424 | 1239 | SUMO2 | VCaP | Prostate | GSM15555 | 47991 | 51867 |
| 202.222 | 61 | 198.6841 | 67 | 78.02318 | 57 | 352.5453 | 125 | SUMO2 | VCaP | Prostate | GSM15555 | 4630 | 51868 |
| 953.0019 | 617 | 951.4511 | 738 | 778.5504 | 813 | 814.9384 | 1001 | SUMO2 | VCaP | Prostate | GSM15555 | 48649 | 51869 |
| 1194.94 | 371 | 1211.445 | 469 | 918.6227 | 353 | 880.3522 | 362 | GABPA | None | Blood | GSM61296 | 10012 | 5188 |
| 1211.234 | 383 | 1242.822 | 500 | 922.0397 | 353 | 917.2808 | 397 | HMGN1 | None | Blood | GSM63080 | 9971 | 5194 |
| 1212.379 | 429 | 1269.952 | 589 | 923.1416 | 404 | 864.0762 | 392 | YY1 | None | Blood | GSM63081 | 11515 | 5195 |
| 324.8472 | 53 | 355.6114 | 61 | 108.5039 | 40 | 227.2499 | 63 | HJURP | SW480 | Colon | GSM15983 | 1760 | 52054 |
| 21.29058 | 9 | 16.97933 | 9 | 13.14287 | 11 | 52.18621 | 23 | HJURP | HeLa | Cervix | GSM15983 | 1089 | 52055 |
| 958.3207 | 648 | 871.2661 | 666 | 785.5384 | 866 | 820.7063 | 1060 | GATA2 | LNCaP | Prostate | GSM16005 | 52418 | 52057 |
| 64.04472 | 141 | 87.04631 | 180 | 105.5394 | 277 | 138.8245 | 329 | TP63 | None | Skin | GSM53893 | 48664 | 5206 |
| 784.6409 | 840 | 825.0915 | 1052 | 568.1977 | 1096 | 504.1658 | 1132 | TRIM28 | None | None | GSM12887 | 153261 | 52087 |
| 1181.921 | 157 | 1202.768 | 202 | 602.3415 | 141 | 629.9425 | 153 | TP53 | IMR90 | Lung | GSM12948 | 3558 | 52090 |
| 1240.08 | 416 | 1310.782 | 585 | 953.2496 | 392 | 924.4368 | 408 | TP53 | IMR90 | Lung | GSM12948 | 9577 | 52091 |
| 1163.406 | 430 | 1222.522 | 591 | 888.366 | 409 | 853.4233 | 426 | TP53 | IMR90 | Lung | GSM12948 | 12588 | 52092 |
| 1.977335 | 3 | 11.5251 | 8 | 0.233445 | 3 | 53.30091 | 25 | TP53 | IMR90 | Lung | GSM12948 | 1280 | 52103 |
| 685.7077 | 286 | 713.5868 | 334 | 398.4541 | 345 | 285.7616 | 330 | PDX1 | CyT49 | Embryo | GSM13163 | 33483 | 52147 |
| 46.01611 | 37 | 43.33345 | 41 | 20.58651 | 44 | 74.75675 | 77 | TP53 | GM06170 | Fetal Skin | GSM13424 | 6839 | 52165 |
| 72.99519 | 43 | 81.62488 | 51 | 43.64088 | 54 | 78.87993 | 74 | TP53 | GM00011 | Fetal Skin | GSM13424 | 6168 | 52167 |
| 1112.429 | 218 | 1180.287 | 316 | 828.5892 | 218 | 860.8576 | 237 | PRDM11 | U-2932 | None | GSM13545 | 6512 | 52170 |
| 269.4599 | 117 | 413.222 | 161 | 123.4384 | 128 | 154.0188 | 151 | PRDM11 | U-2932 | None | GSM13545 | 13614 | 52171 |
| 3.96354 | 5 | 8.14722 | 8 | 2.15869 | 7 | 33.6903 | 23 | LMNA | HeLa | Cervix | GSM13761 | 1585 | 52222 |

| | | | | | | | | | | | | | |
|---|---|---|---|---|---|---|---|---|---|---|---|---|---|
| 44.74278 | 29 | 48.34494 | 34 | 34.28532 | 41 | 102.309 | 70 | AR | None | Ovary | GSM14156 | 4751 | 52249 |
| 687.8052 | 380 | 689.3186 | 466 | 345.9572 | 439 | 382.6276 | 495 | SMARCA4 | H9 | Embryo | GSM60229 | 55553 | 5227 |
| 84.66568 | 156 | 179.6667 | 232 | 156.6842 | 315 | 221.5441 | 384 | TP63 | None | None | GSM14469 | 49067 | 52299 |
| 104.9253 | 164 | 185.8641 | 230 | 155.4489 | 307 | 222.6073 | 376 | TP63 | None | None | GSM14469 | 47076 | 52300 |
| 79.57835 | 138 | 118.4024 | 182 | 138.8012 | 274 | 144.0969 | 303 | TP63 | None | None | GSM14469 | 41968 | 52301 |
| 59.64919 | 124 | 105.6409 | 173 | 134.462 | 268 | 140.5466 | 297 | TP63 | None | None | GSM14469 | 41179 | 52302 |
| 1078.09 | 303 | 1094.98 | 381 | 823.06 | 292 | 767.2492 | 282 | BCL6 | None | Tonsil | GSM14624 | 10282 | 52303 |
| 1145.141 | 407 | 1173.847 | 529 | 875.8986 | 399 | 837.5025 | 411 | MYC | P493-6 | None | GSM14683 | 13253 | 52358 |
| 1122.165 | 307 | 1195.332 | 447 | 898.2162 | 331 | 851.5905 | 325 | MYC | P493-6 | None | GSM14683 | 9773 | 52359 |
| 519.2652 | 62 | 594.5324 | 73 | 245.3303 | 53 | 326.9731 | 66 | MYC | P493-6 | None | GSM14683 | 1230 | 52360 |
| 284.2878 | 41 | 354.1526 | 51 | 154.0527 | 39 | 126.0881 | 37 | MYC | HEK293 | Embryonic | GSM14930 | 1104 | 52424 |
| 936.8385 | 169 | 1053.628 | 227 | 525.9722 | 171 | 542.4568 | 185 | MYC | HEK293 | Embryonic | GSM14930 | 6233 | 52427 |
| 1061.701 | 196 | 1042.326 | 230 | 546.5488 | 179 | 577.1072 | 196 | MYC | HEK293 | Embryonic | GSM14930 | 6641 | 52428 |
| 354.3462 | 56 | 394.0877 | 65 | 238.066 | 61 | 131.1555 | 47 | MYC | HEK293 | Embryonic | GSM14930 | 1735 | 52429 |
| 1073.467 | 684 | 1118.852 | 878 | 780.8291 | 700 | 734.6751 | 720 | WDR5 | HEK293 | Embryonic | GSM14930 | 35813 | 52432 |
| 1230.834 | 670 | 1286.53 | 873 | 903.8773 | 623 | 855.8097 | 638 | WDR5 | HEK293 | Embryonic | GSM14930 | 19968 | 52436 |
| 286.4652 | 172 | 448.3639 | 236 | 257.3434 | 255 | 388.4462 | 324 | TCF21 | None | Coronary a | GSM15032 | 26347 | 52495 |
| 525.4454 | 254 | 661.3274 | 316 | 273.9814 | 300 | 408.923 | 378 | TCF21 | None | Coronary a | GSM15032 | 33533 | 52496 |
| 15.4535 | 18 | 52.91188 | 35 | 27.38111 | 37 | 108.7844 | 71 | TCF21 | None | Coronary a | GSM15032 | 4603 | 52497 |
| 58.45208 | 34 | 53.97693 | 37 | 41.74931 | 46 | 135.1545 | 82 | TCF21 | None | Coronary a | GSM15032 | 5012 | 52498 |
| 130.1323 | 211 | 143.3185 | 251 | 87.974 | 315 | 117.1803 | 374 | JUN | None | Coronary a | GSM15032 | 61996 | 52500 |
| 134.8601 | 304 | 150.3913 | 364 | 95.01058 | 475 | 120.9603 | 555 | JUND | None | Coronary a | GSM15032 | 108585 | 52501 |
| 707.9256 | 204 | 841.9829 | 251 | 644.5246 | 283 | 802.0521 | 417 | AR | R1-AD1 | Prostate | GSM15155 | 15505 | 52540 |
| 628.8902 | 163 | 675.5372 | 187 | 485.8774 | 206 | 835.6128 | 335 | AR | R1-AD1 | Prostate | GSM15155 | 10582 | 52541 |
| 433.0035 | 101 | 535.5661 | 125 | 331.5441 | 125 | 786.7244 | 208 | AR | R1-AD1 | Prostate | GSM15155 | 5595 | 52542 |
| 150.7445 | 72 | 264.3678 | 106 | 138.6875 | 106 | 259.1849 | 150 | AR | R1-D567 | Prostate | GSM15155 | 9048 | 52543 |
| 105.2318 | 27 | 216.8165 | 45 | 105.8465 | 39 | 421.6054 | 88 | AR | R1-D567 | Prostate | GSM15155 | 1733 | 52545 |
| 199.2924 | 286 | 229.5959 | 345 | 96.48984 | 391 | 108.7081 | 444 | MITF | 501-Mel | None | GSM15177 | 83873 | 52548 |
| 15.01538 | 21 | 21.87102 | 28 | 14.63535 | 35 | 37.19624 | 53 | SOX10 | 501-Mel | None | GSM15177 | 5846 | 52549 |
| 858.006 | 321 | 874.3262 | 407 | 455.0548 | 312 | 476.2952 | 343 | SMARCA4 | 501-Mel | None | GSM15177 | 24739 | 52550 |
| 91.7054 | 64 | 81.59085 | 69 | 72.29927 | 92 | 68.81661 | 98 | SMARCA4 | 501-Mel | None | GSM15177 | 10934 | 52551 |
| 1069.613 | 471 | 1098.487 | 604 | 782.0709 | 450 | 763.0962 | 487 | SMARCA4 | 501-Mel | None | GSM15189 | 21022 | 52555 |
| 59.94857 | 39 | 39.95477 | 37 | 48.65333 | 56 | 69.84105 | 70 | CHRM2 | None | Endometri | GSM15275 | 6143 | 52573 |
| 50.85467 | 54 | 72.39289 | 71 | 30.6577 | 74 | 65.05475 | 104 | CHRM2 | None | Endometri | GSM15275 | 12076 | 52574 |
| 559.8791 | 194 | 413.7976 | 187 | 594.1924 | 293 | 735.809 | 393 | ESR1 | MCF-7 | Breast | GSM15347 | 17709 | 52608 |
| 475.1233 | 188 | 474.9031 | 210 | 634.5179 | 318 | 727.1483 | 419 | ESR1 | MCF-7 | Breast | GSM15347 | 19689 | 52609 |
| 573.6517 | 66 | 356.3814 | 53 | 367.4631 | 67 | 484.1707 | 83 | ESR1 | MCF-7 | Breast | GSM15347 | 1232 | 52611 |
| 377.9747 | 143 | 356.1469 | 155 | 505.7306 | 242 | 741.3235 | 324 | ESR1 | MCF-7 | Breast | GSM15347 | 14039 | 52612 |
| 419.9623 | 165 | 433.2848 | 187 | 555.6415 | 278 | 737.8081 | 382 | ESR1 | MCF-7 | Breast | GSM15347 | 16946 | 52613 |
| 319.6507 | 43 | 595.7689 | 69 | 199.8587 | 44 | 259.5984 | 54 | RELA | MCF-7 | Breast | GSM15347 | 1055 | 52614 |
| 344.6947 | 47 | 559.1291 | 69 | 229.3737 | 50 | 303.4616 | 62 | RELA | MCF-7 | Breast | GSM15347 | 1185 | 52616 |
| 402.577 | 55 | 683.4049 | 83 | 287.9068 | 61 | 269.1511 | 62 | RELA | MCF-7 | Breast | GSM15347 | 1385 | 52617 |
| 498.5652 | 70 | 815.7984 | 103 | 372.7474 | 80 | 323.1285 | 78 | RELA | MCF-7 | Breast | GSM15347 | 1848 | 52619 |
| 383.7541 | 62 | 627.9913 | 91 | 271.3415 | 70 | 392.9904 | 91 | RELA | MCF-7 | Breast | GSM15347 | 2049 | 52620 |
| 549.024 | 84 | 802.6054 | 115 | 401.6424 | 96 | 456.2132 | 109 | RELA | MCF-7 | Breast | GSM15347 | 2554 | 52621 |
| 909.7032 | 292 | 872.5678 | 322 | 800.4514 | 437 | 808.8681 | 507 | FOXA1 | MCF-7 | Breast | GSM15347 | 19304 | 52622 |
| 919.2722 | 341 | 889.9146 | 389 | 799.6944 | 496 | 815.5766 | 586 | FOXA1 | MCF-7 | Breast | GSM15347 | 22767 | 52623 |
| 926.0931 | 333 | 872.4335 | 354 | 775.3044 | 450 | 808.3428 | 553 | FOXA1 | MCF-7 | Breast | GSM15347 | 21709 | 52624 |
| 901.7147 | 388 | 861.1471 | 431 | 757.0409 | 535 | 796.9545 | 671 | FOXA1 | MCF-7 | Breast | GSM15347 | 28768 | 52625 |
| 912.4515 | 340 | 876.3649 | 377 | 784.1501 | 482 | 789.9017 | 556 | FOXA1 | MCF-7 | Breast | GSM15347 | 23116 | 52626 |
| 912.1235 | 325 | 866.8649 | 353 | 788.8666 | 467 | 804.9819 | 555 | FOXA1 | MCF-7 | Breast | GSM15347 | 21812 | 52627 |
| 913.4707 | 268 | 851.3129 | 274 | 778.1235 | 371 | 802.9988 | 449 | FOXA1 | MCF-7 | Breast | GSM15347 | 17019 | 52628 |
| 899.2991 | 371 | 846.9159 | 401 | 773.3684 | 535 | 795.5339 | 644 | FOXA1 | MCF-7 | Breast | GSM15347 | 27259 | 52629 |
| 894.9561 | 313 | 918.8424 | 407 | 731.9546 | 405 | 730.8994 | 461 | ESR1 | MCF-7 | Breast | GSM15347 | 21900 | 52632 |
| 382.7759 | 153 | 471.7761 | 189 | 435.5133 | 242 | 548.2599 | 288 | ESR1 | MCF-7 | Breast | GSM15347 | 16236 | 52633 |
| 198.1009 | 115 | 228.2487 | 138 | 350.8255 | 225 | 538.8014 | 292 | ESR1 | MCF-7 | Breast | GSM15347 | 16759 | 52636 |
| 145.6827 | 67 | 188.6146 | 85 | 260.7777 | 132 | 335.1686 | 159 | ESR1 | MCF-7 | Breast | GSM15347 | 7880 | 52637 |
| 1013.653 | 216 | 1050.309 | 291 | 621.0765 | 219 | 565.2061 | 223 | NOTCH1 | None | None | GSM15379 | 9494 | 52638 |
| 101.1834 | 65 | 65.3785 | 61 | 56.65049 | 81 | 163.72 | 133 | FOXA1 | CyT49 | Embryo | GSM15514 | 10151 | 52677 |
| 437.1061 | 201 | 326.3418 | 198 | 352.1114 | 277 | 461.7356 | 334 | FOXA1 | CyT49 | Embryo | GSM15514 | 24239 | 52678 |
| 354.892 | 152 | 258.3258 | 147 | 243.1832 | 194 | 358.4887 | 246 | FOXA1 | CyT49 | Embryo | GSM15514 | 16821 | 52679 |
| 486.0017 | 199 | 432.8012 | 211 | 383.7028 | 269 | 514.5546 | 328 | FOXA1 | CyT49 | Embryo | GSM15514 | 21769 | 52680 |
| 723.2961 | 487 | 725.4547 | 597 | 527.515 | 600 | 483.6667 | 616 | FOXA2 | CyT49 | Embryo | GSM15514 | 70058 | 52681 |
| 752.547 | 485 | 734.1977 | 566 | 563.2879 | 605 | 537.8409 | 648 | FOXA2 | CyT49 | Embryo | GSM15514 | 62432 | 52682 |
| 816.1097 | 489 | 801.0447 | 578 | 609.1459 | 585 | 574.3149 | 612 | FOXA2 | CyT49 | Embryo | GSM15514 | 51670 | 52683 |
| 767.5777 | 434 | 769.7252 | 532 | 571.7753 | 525 | 554.9165 | 575 | FOXA2 | CyT49 | Embryo | GSM15514 | 50830 | 52684 |
| 255.0995 | 41 | 185.9133 | 37 | 92.39556 | 32 | 216.6978 | 54 | DYRK1A | T98G | Brain | GSM15560 | 1313 | 52715 |
| 1161.132 | 406 | 1184.485 | 516 | 874.0425 | 375 | 842.2053 | 390 | BCL6 | None | Tonsil | GSM15897 | 12332 | 52774 |
| 178.7088 | 35 | 252.5474 | 47 | 123.472 | 40 | 219.5626 | 58 | CIITA | Raji | Blood | GSM16022 | 1528 | 52784 |
| 481.9655 | 59 | 597.1903 | 73 | 333.7523 | 63 | 347.6008 | 68 | CIITA | Raji | Blood | GSM16022 | 1192 | 52788 |
| 86.25765 | 91 | 149.2186 | 129 | 79.36985 | 143 | 94.01447 | 165 | NR3C1 | Nalm6 | Blood | GSM16373 | 20211 | 52959 |
| 190.992 | 148 | 274.9743 | 195 | 135.5618 | 206 | 138.6002 | 226 | NR3C1 | Nalm6 | Blood | GSM16373 | 26788 | 52960 |

| | | | | | | | | | | | | | |
|---|---|---|---|---|---|---|---|---|---|---|---|---|---|
| 35.93185 | 33 | 73.07145 | 51 | 75.0408 | 71 | 107.5994 | 89 | NR3C1 | Nalm6 | Blood | GSM16373 | 6869 | 52962 |
| 7.163428 | 5 | 21.5374 | 10 | 3.387528 | 6 | 101.065 | 32 | GATA3 | MCF-7 | Breast | GSM16425 | 1055 | 52964 |
| 112.717 | 47 | 210.6513 | 72 | 82.47465 | 61 | 150.8296 | 86 | EHMT2 | MCF-7 | Breast | GSM16425 | 4842 | 52965 |
| 425.242 | 81 | 445.1605 | 91 | 293.1949 | 92 | 500.7485 | 130 | MTA3 | MCF-7 | Breast | GSM16425 | 3308 | 52966 |
| 1266.174 | 726 | 1267.949 | 876 | 958.7479 | 740 | 919.842 | 780 | E2F1 | LNCaP-abl | Prostate | GSM16564 | 21320 | 53013 |
| 730.487 | 84 | 1005.38 | 111 | 532.616 | 92 | 524.528 | 96 | RBP2 | U937 | Pleura | GSM70053 | 1552 | 53075 |
| 1209.815 | 227 | 1258.977 | 316 | 970.4552 | 228 | 942.8627 | 236 | RBP2 | U937 | Pleura | GSM70053 | 4952 | 53076 |
| 613.1692 | 68 | 801.4093 | 87 | 368.7297 | 66 | 410.2154 | 74 | RBP2 | U937 | Pleura | GSM70053 | 1167 | 53077 |
| 654.9144 | 70 | 648.7384 | 75 | 368.9093 | 65 | 490.4954 | 81 | RBP2 | U937 | Pleura | GSM70053 | 1124 | 53078 |
| 391.1456 | 87 | 393.5234 | 96 | 207.1002 | 89 | 220.528 | 98 | MYC | None | None | GSM11457 | 4547 | 53087 |
| 1132.063 | 226 | 1180.586 | 310 | 747.6464 | 204 | 890.1026 | 252 | NRF1 | Namalwa | None | GSM12828 | 6438 | 53105 |
| 1047.22 | 539 | 1037.476 | 643 | 779.1653 | 566 | 755.1473 | 607 | MYC | MCF-10A | Breast | GSM14237 | 28602 | 53192 |
| 1227.249 | 148 | 1204.753 | 170 | 766.6044 | 149 | 747.0821 | 155 | FOXM1 | HEK293 | Embryonic | GSM14640 | 3018 | 53251 |
| 583.085 | 74 | 791.9606 | 97 | 468.9687 | 87 | 361.5977 | 79 | FOXM1 | HEK293 | Embryonic | GSM14640 | 1636 | 53252 |
| 1137.547 | 375 | 1152.984 | 468 | 898.3198 | 393 | 864.1069 | 405 | FOXM1 | HEK293 | Embryonic | GSM14640 | 12095 | 53253 |
| 1156.415 | 386 | 1190.31 | 509 | 918.2447 | 407 | 872.9939 | 412 | FOXM1 | HEK293 | Embryonic | GSM14640 | 11748 | 53254 |
| 1183.764 | 415 | 1220.927 | 540 | 929.2291 | 425 | 892.562 | 435 | FOXM1 | HEK293 | Embryonic | GSM14640 | 11888 | 53256 |
| 1064.306 | 139 | 1195.196 | 181 | 659.2837 | 142 | 603.9757 | 143 | FOXM1 | HEK293 | Embryonic | GSM14640 | 3326 | 53257 |
| 1199.731 | 386 | 1217.334 | 490 | 927.0153 | 370 | 893.6493 | 386 | FOXM1 | HEK293 | Embryonic | GSM14640 | 10263 | 53259 |
| 1204.442 | 335 | 1235.176 | 437 | 949.0496 | 331 | 912.9942 | 340 | FOXM1 | HEK293 | Embryonic | GSM14640 | 8313 | 53260 |
| 915.1262 | 115 | 1080.355 | 138 | 512.2829 | 110 | 441.4908 | 107 | FOXM1 | HEK293 | Embryonic | GSM14640 | 2544 | 53261 |
| 1188.481 | 164 | 1194.909 | 204 | 792.4275 | 167 | 647.9834 | 158 | FOXM1 | HEK293 | Embryonic | GSM14640 | 3808 | 53262 |
| 948.4325 | 114 | 1136.652 | 138 | 556.0653 | 111 | 481.3695 | 108 | FOXM1 | HEK293 | Embryonic | GSM14640 | 2311 | 53263 |
| 869.4352 | 104 | 1178.792 | 136 | 509.4979 | 101 | 463.3801 | 101 | FOXM1 | HEK293 | Embryonic | GSM14640 | 2072 | 53264 |
| 1108.949 | 761 | 1138.438 | 950 | 780.8716 | 764 | 719.8716 | 765 | ERG | TSU-1621-I | None | GSM14807 | 42436 | 53296 |
| 797.4816 | 561 | 822.0315 | 712 | 570.1665 | 656 | 543.8875 | 704 | ERG | TSU-1621-I | None | GSM14807 | 68543 | 53297 |
| 1142.84 | 707 | 1189.451 | 913 | 806.2654 | 675 | 763.5355 | 701 | SPI1 | TSU-1621-I | None | GSM14807 | 31949 | 53298 |
| 1052.117 | 542 | 1086.954 | 698 | 773.4695 | 542 | 723.7202 | 547 | SPI1 | TSU-1621-I | None | GSM14807 | 26392 | 53299 |
| 1190.331 | 650 | 1242.766 | 851 | 875.6133 | 631 | 820.0962 | 628 | RARA | TSU-1621-I | None | GSM14807 | 23367 | 53300 |
| 1035.943 | 465 | 1067.716 | 602 | 787.3844 | 492 | 749.4872 | 508 | FLI1 | TSU-1621-I | None | GSM14807 | 23327 | 53301 |
| 965.163 | 462 | 996.7111 | 599 | 729.0429 | 510 | 674.844 | 501 | T | TSU-1621-I | None | GSM14807 | 29512 | 53302 |
| 677.6177 | 223 | 813.0411 | 283 | 325.0665 | 237 | 320.0608 | 254 | FUS | TSU-1621-I | None | GSM14807 | 19440 | 53303 |
| 442.0597 | 150 | 647.016 | 202 | 172.624 | 146 | 170.9998 | 157 | FUS | TSU-1621-I | None | GSM14807 | 13126 | 53304 |
| 758.3388 | 526 | 790.2306 | 676 | 557.5336 | 640 | 536.1199 | 696 | RUNX1 | TSU-1621-I | None | GSM14807 | 68622 | 53305 |
| 41.60711 | 86 | 50.30248 | 106 | 67.47339 | 169 | 66.25518 | 184 | LMO2 | TSU-1621-I | None | GSM14807 | 28414 | 53306 |
| 10.84111 | 20 | 25.64276 | 32 | 21.23848 | 43 | 44.42122 | 61 | GATA2 | TSU-1621-I | None | GSM14807 | 6659 | 53307 |
| 1.369687 | 15 | 16.06271 | 35 | 14.339 | 50 | 19.3558 | 60 | LYL1 | TSU-1621-I | None | GSM14807 | 9795 | 53308 |
| 170.3427 | 133 | 288.2856 | 188 | 174.2858 | 212 | 178.1968 | 232 | TAL1 | TSU-1621-I | None | GSM14807 | 24683 | 53309 |
| 1125.594 | 452 | 1128.88 | 549 | 844.1835 | 429 | 786.2139 | 419 | INTS3 | HeLa | Cervix | GSM14828 | 15367 | 53323 |
| 1253.209 | 640 | 1301.581 | 837 | 896.6536 | 559 | 864.8676 | 589 | NELFE | HeLa | Cervix | GSM14828 | 18442 | 53324 |
| 1203.33 | 702 | 1238.058 | 891 | 868.2741 | 669 | 848.6621 | 732 | SUPT5H | HeLa | Cervix | GSM14828 | 25586 | 53325 |
| 313.5654 | 183 | 552.5543 | 265 | 176.1122 | 228 | 276.4555 | 292 | NELFE | H9 | Embryo | GSM14831 | 28063 | 53337 |
| 199.8479 | 131 | 278.669 | 171 | 105.9262 | 162 | 198.4468 | 221 | NELFE | H9 | Embryo | GSM14831 | 21582 | 53338 |
| 6.121243 | 19 | 16.49093 | 31 | 7.915371 | 37 | 15.29728 | 49 | T | H9 | Embryo | GSM14831 | 8356 | 53340 |
| 1225.411 | 550 | 1288.933 | 734 | 918.8385 | 505 | 882.2069 | 524 | CDK9 | HeLa | Cervix | GSM14944 | 13777 | 53346 |
| 1296.718 | 691 | 1352.291 | 900 | 932.1389 | 605 | 923.8707 | 677 | CDK9 | HeLa | Cervix | GSM14944 | 17087 | 53347 |
| 1204.49 | 339 | 1218.549 | 427 | 949.7301 | 338 | 890.3308 | 326 | CHD8 | T47D | Mammary | GSM15268 | 8637 | 53396 |
| 24.17053 | 17 | 23.26886 | 19 | 15.92447 | 23 | 93.97904 | 53 | CHD8 | T47D | Mammary | GSM15268 | 3076 | 53397 |
| 229.7931 | 162 | 324.0681 | 212 | 135.9548 | 211 | 109.7278 | 214 | CHD8 | T47D | Mammary | GSM15268 | 28656 | 53398 |
| 293.1235 | 98 | 315.0081 | 113 | 175.8145 | 114 | 338.877 | 165 | CHD8 | T47D | Mammary | GSM15268 | 8438 | 53399 |
| 1012.601 | 425 | 1057.945 | 572 | 760.9938 | 440 | 727.5404 | 460 | ETV1 | GIST48 | Stomach | GSM55867 | 21961 | 534 |
| 208.408 | 146 | 197.8889 | 162 | 87.26842 | 169 | 95.73864 | 190 | CHD8 | T47D | Mammary | GSM15268 | 25630 | 53400 |
| 875.8356 | 354 | 896.5339 | 452 | 655.9226 | 391 | 605.6726 | 393 | RUNX1 | Kasumi-1 | Blood | GSM15344 | 26692 | 53417 |
| 851.8473 | 373 | 881.0746 | 486 | 624.33 | 409 | 617.6784 | 455 | RUNX1 | Kasumi-1 | Blood | GSM15344 | 31007 | 53418 |
| 806.4229 | 252 | 835.8327 | 324 | 376.3773 | 263 | 420.6278 | 297 | NCOR1 | Kasumi-1 | Blood | GSM15344 | 20971 | 53419 |
| 1106.025 | 551 | 1120.568 | 684 | 792.2618 | 508 | 756.0188 | 530 | EP300 | Kasumi-1 | Blood | GSM15344 | 23169 | 53420 |
| 1112.412 | 160 | 1170.352 | 223 | 673.7791 | 164 | 646.924 | 170 | KDM1A | SH-SY5Y | Bone Marr | GSM15422 | 4494 | 53467 |
| 1166.589 | 256 | 1198.053 | 336 | 921.963 | 254 | 895.229 | 266 | KDM1A | SH-SY5Y | Bone Marr | GSM15422 | 6740 | 53468 |
| 1119.279 | 277 | 1149.78 | 367 | 886.9254 | 290 | 858.3648 | 305 | KDM1A | SH-SY5Y | Bone Marr | GSM15422 | 8769 | 53469 |
| 679.9475 | 254 | 798.6215 | 342 | 505.6322 | 331 | 495.3479 | 354 | KDM1A | SH-SY5Y | Bone Marr | GSM15422 | 25482 | 53470 |
| 100.5508 | 23 | 134.897 | 30 | 90.78102 | 31 | 85.63487 | 32 | SVIL | SH-SY5Y | Bone Marr | GSM15422 | 1274 | 53472 |
| 17.7216 | 9 | 10.85731 | 8 | 26.52899 | 17 | 109.2615 | 37 | MBD2 | HMEC-hTEI | None | GSM15441 | 1322 | 53477 |
| 4.867951 | 31 | 5.972982 | 39 | 3.992695 | 56 | 18.48762 | 88 | MBD2 | HMLER | None | GSM15441 | 17145 | 53479 |
| 1183.071 | 700 | 1170.368 | 821 | 887.5184 | 734 | 869.6748 | 807 | SOX9 | HT29 | Colon | GSM15542 | 28780 | 53493 |
| 654.0509 | 534 | 668.3781 | 664 | 433.7127 | 665 | 423.1681 | 724 | SPI1 | TSU-1621-I | None | GSM15576 | 100844 | 53549 |
| 717.7489 | 397 | 728.9509 | 497 | 542.9852 | 505 | 532.7911 | 559 | MED1 | None | None | GSM15667 | 52399 | 53592 |
| 995.9762 | 607 | 1048.152 | 801 | 746.8085 | 678 | 703.4368 | 700 | MED1 | None | None | GSM15667 | 40071 | 53593 |
| 1048.25 | 625 | 1113.162 | 838 | 782.2162 | 678 | 737.5537 | 696 | MED1 | None | None | GSM15667 | 35540 | 53594 |
| 145.2786 | 115 | 283.2345 | 173 | 183.2797 | 198 | 195.8126 | 220 | MED1 | None | None | GSM15667 | 21084 | 53595 |
| 322.9605 | 206 | 470.299 | 273 | 268.642 | 299 | 307.5671 | 341 | RELA | None | None | GSM15667 | 33810 | 53596 |
| 604.6404 | 299 | 700.7778 | 380 | 447.2046 | 405 | 449.2753 | 440 | RELA | None | None | GSM15667 | 41179 | 53597 |
| 196.711 | 291 | 270.6591 | 372 | 179.2516 | 471 | 205.0396 | 537 | RELA | None | None | GSM15667 | 87359 | 53598 |

| | | | | | | | | | | | | |
|---|---|---|---|---|---|---|---|---|---|---|---|---|
| 832.1115 | 499 | 856.539 | 636 | 641.0814 | 618 | 609.3612 | 650 | RELA | None | None | GSM15667 | 49707 | 53599 |
| 557.0764 | 163 | 751.6123 | 211 | 283.4363 | 174 | 214.6472 | 166 | STAT5B | None | Blood | GSM15777 | 12652 | 53628 |
| 855.8761 | 424 | 881.798 | 549 | 628.8766 | 472 | 604.8938 | 505 | STAT5B | None | Blood | GSM15777 | 36973 | 53629 |
| 929.7468 | 496 | 972.9763 | 657 | 680.998 | 532 | 628.1471 | 528 | STAT5B | None | Blood | GSM15777 | 36321 | 53630 |
| 276.9586 | 129 | 395.2928 | 171 | 159.6123 | 156 | 172.7464 | 174 | STAT5B | None | Blood | GSM15777 | 15812 | 53631 |
| 873.8505 | 401 | 885.2431 | 499 | 589.0311 | 398 | 609.8384 | 461 | STAT5B | None | Blood | GSM15777 | 33341 | 53632 |
| 220.6001 | 51 | 293.7853 | 66 | 167.8492 | 63 | 172.2248 | 68 | STAT5B | None | Blood | GSM15777 | 2848 | 53633 |
| 394.7576 | 144 | 615.148 | 200 | 270.074 | 180 | 223.8303 | 179 | STAT5B | None | Blood | GSM15777 | 13694 | 53634 |
| 279.0987 | 94 | 389.3261 | 124 | 144.8826 | 103 | 154.7313 | 114 | STAT5B | None | Blood | GSM15777 | 8288 | 53636 |
| 69.45977 | 33 | 142.4509 | 53 | 77.6254 | 53 | 122.2973 | 70 | STAT5B | None | Blood | GSM15777 | 4123 | 53637 |
| 17.0706 | 8 | 10.29589 | 7 | 6.328646 | 8 | 61.00792 | 25 | SMAD2 | H9 | Embryo | GSM76175 | 1113 | 5365 |
| 59.9078 | 17 | 79.06747 | 22 | 51.57037 | 23 | 137.5932 | 41 | LEF1 | None | None | GSM15793 | 1269 | 53650 |
| 142.5382 | 75 | 232.1919 | 106 | 94.80923 | 97 | 163.8788 | 131 | LEF1 | None | None | GSM15793 | 9879 | 53651 |
| 218.5466 | 205 | 365.4221 | 285 | 82.84739 | 243 | 121.7974 | 298 | CTNNB1 | None | None | GSM15793 | 44304 | 53653 |
| 14.1919 | 14 | 12.00344 | 15 | 10.89885 | 21 | 20.63254 | 29 | SMAD2 | H9 | Embryo | GSM76175 | 3311 | 5366 |
| 884.4166 | 503 | 913.5352 | 645 | 636.2865 | 539 | 589.329 | 545 | NIPBL | None | None | GSM15793 | 41376 | 53670 |
| 945.0626 | 410 | 956.0813 | 513 | 661.5945 | 389 | 650.1666 | 428 | NIPBL | None | None | GSM15793 | 25899 | 53671 |
| 945.8099 | 497 | 946.3733 | 604 | 674.2366 | 501 | 637.941 | 519 | NIPBL | None | None | GSM15793 | 34015 | 53672 |
| 927.1086 | 460 | 978.605 | 618 | 682.1361 | 491 | 645.4838 | 508 | NIPBL | None | None | GSM15793 | 32155 | 53673 |
| 167.789 | 37 | 200.1849 | 45 | 31.92739 | 23 | 97.17551 | 42 | TAT | Jurkat | Blood | GSM16032 | 1922 | 53704 |
| 745.2199 | 415 | 698.4942 | 453 | 674.5591 | 673 | 718.3663 | 851 | RUNX2 | C4-2B | Prostate | GSM83840 | 49856 | 5373 |
| 188.4772 | 116 | 182.9351 | 129 | 307.3871 | 217 | 238.6892 | 211 | MEF2B | HEK293A | None | GSM16471 | 16552 | 53873 |
| 290.4435 | 214 | 284.0301 | 240 | 360.4356 | 358 | 277.6894 | 354 | MEF2B | HEK293A | None | GSM16471 | 35489 | 53874 |
| 260.0523 | 143 | 257.8213 | 160 | 262.478 | 217 | 215.41 | 217 | MEF2B | HEK293A | None | GSM16471 | 18391 | 53877 |
| 207.7498 | 78 | 167.1678 | 78 | 243.357 | 123 | 183.7131 | 116 | MEF2B | HEK293A | None | GSM16471 | 6898 | 53878 |
| 1074.019 | 754 | 1112.577 | 956 | 784.6882 | 831 | 763.2489 | 907 | RELA | A549 | Lung | GSM84787 | 48567 | 5388 |
| 484.9786 | 128 | 825.2386 | 188 | 366.0395 | 159 | 352.3733 | 167 | CTCF | K562 | Bone Marr | GSM16473 | 8003 | 53885 |
| 1070.794 | 766 | 1128.585 | 993 | 792.4389 | 864 | 760.6873 | 931 | RELA | A549 | Lung | GSM84787 | 51080 | 5389 |
| 993.3535 | 745 | 1039.687 | 954 | 703.6599 | 807 | 673.96 | 872 | SP1 | HCT-116 | Colon | GSM85544 | 59941 | 5395 |
| 270.6322 | 74 | 323.5797 | 90 | 488.4699 | 144 | 644.1137 | 177 | ESR1 | T47D | Mammary | GSM16690 | 4904 | 53955 |
| 235.5306 | 60 | 281.7375 | 73 | 399.7151 | 113 | 505.7558 | 136 | ESR1 | T47D | Mammary | GSM16690 | 3642 | 53956 |
| 129.0129 | 38 | 204.0802 | 54 | 265.3713 | 80 | 472.8916 | 116 | ESR1 | T47D | Mammary | GSM16690 | 2769 | 53957 |
| 551.9301 | 191 | 597.8343 | 221 | 609.669 | 293 | 728.131 | 377 | ESR1 | T47D | Mammary | GSM16690 | 17093 | 53958 |
| 492.8718 | 182 | 588.3061 | 221 | 523.1614 | 276 | 708.6297 | 360 | ESR1 | T47D | Mammary | GSM16690 | 17517 | 53959 |
| 234.4093 | 89 | 354.1541 | 122 | 313.6611 | 151 | 662.5274 | 232 | ESR1 | T47D | Mammary | GSM16690 | 8563 | 53960 |
| 363.3461 | 130 | 422.9768 | 156 | 404.6822 | 201 | 686.3143 | 276 | ESR1 | T47D | Mammary | GSM16690 | 11804 | 53961 |
| 210.0286 | 77 | 318.7944 | 106 | 278.2773 | 130 | 562.2383 | 196 | ESR1 | T47D | Mammary | GSM16690 | 7166 | 53962 |
| 326.4815 | 130 | 468.4317 | 173 | 405.5038 | 213 | 714.6659 | 297 | ESR1 | T47D | Mammary | GSM16690 | 13213 | 53963 |
| 468.6411 | 94 | 594.9009 | 118 | 741.2549 | 168 | 892.6041 | 197 | EP300 | T47D | Mammary | GSM16690 | 4147 | 53967 |
| 360.3028 | 60 | 384.3404 | 68 | 498.8283 | 99 | 672.0789 | 124 | EP300 | T47D | Mammary | GSM16690 | 2037 | 53968 |
| 193.266 | 40 | 259.103 | 52 | 391.33 | 83 | 581.9339 | 110 | EP300 | T47D | Mammary | GSM16690 | 1860 | 53969 |
| 612.2464 | 202 | 677.3627 | 236 | 617.0764 | 296 | 719.8363 | 372 | EP300 | T47D | Mammary | GSM16690 | 17250 | 53970 |
| 855.468 | 282 | 860.2252 | 349 | 726.9296 | 397 | 748.8689 | 481 | EP300 | T47D | Mammary | GSM16690 | 21343 | 53971 |
| 502.7657 | 156 | 635.9142 | 195 | 578.598 | 242 | 771.4747 | 316 | EP300 | T47D | Mammary | GSM16690 | 12188 | 53972 |
| 424.8205 | 150 | 505.9847 | 182 | 547.7578 | 248 | 742.9759 | 319 | EP300 | T47D | Mammary | GSM16690 | 13576 | 53973 |
| 911.0835 | 228 | 927.7534 | 295 | 787.0417 | 325 | 794.9266 | 377 | EP300 | T47D | Mammary | GSM16690 | 13625 | 53974 |
| 904.4612 | 237 | 941.3437 | 321 | 782.9911 | 336 | 803.2341 | 401 | EP300 | T47D | Mammary | GSM16690 | 14450 | 53975 |
| 764.2565 | 287 | 722.1661 | 313 | 699.0066 | 476 | 744.6301 | 607 | ESR1 | MCF-7 | Breast | GSM16690 | 29107 | 53986 |
| 389.189 | 139 | 355.1019 | 148 | 675.1866 | 265 | 796.1676 | 353 | ESR1 | MCF-7 | Breast | GSM16690 | 12711 | 53987 |
| 543.5794 | 201 | 517.6353 | 219 | 703.1607 | 343 | 764.77 | 464 | ESR1 | MCF-7 | Breast | GSM16690 | 19464 | 53988 |
| 795.9837 | 298 | 767.9352 | 339 | 711.0892 | 470 | 764.9045 | 613 | ESR1 | MCF-7 | Breast | GSM16690 | 27690 | 53989 |
| 764.4239 | 394 | 720.7721 | 435 | 688.177 | 637 | 725.6505 | 792 | ESR1 | MCF-7 | Breast | GSM16690 | 44281 | 53990 |
| 755.1998 | 333 | 707.528 | 361 | 683.5161 | 546 | 726.773 | 693 | ESR1 | MCF-7 | Breast | GSM16690 | 36621 | 53991 |
| 701.7448 | 254 | 663.672 | 276 | 694.1283 | 409 | 749.3387 | 540 | ESR1 | MCF-7 | Breast | GSM16690 | 24608 | 53992 |
| 741.6798 | 340 | 706.1356 | 377 | 669.739 | 553 | 718.0981 | 711 | ESR1 | MCF-7 | Breast | GSM16690 | 38890 | 53993 |
| 767.772 | 373 | 735.9224 | 421 | 695.9056 | 606 | 730.1844 | 750 | ESR1 | MCF-7 | Breast | GSM16690 | 40544 | 53994 |
| 609.8587 | 203 | 606.0962 | 225 | 736.4675 | 344 | 766.2993 | 426 | EP300 | MCF-7 | Breast | GSM16690 | 17303 | 53999 |
| 112.8319 | 29 | 100.2619 | 30 | 326.2298 | 74 | 485.5648 | 98 | EP300 | MCF-7 | Breast | GSM16690 | 1814 | 54000 |
| 439.063 | 106 | 426.8743 | 115 | 839.1029 | 211 | 922.0802 | 263 | EP300 | MCF-7 | Breast | GSM16690 | 6023 | 54001 |
| 600.5118 | 170 | 562.4176 | 182 | 788.5675 | 292 | 822.8883 | 365 | EP300 | MCF-7 | Breast | GSM16690 | 12043 | 54003 |
| 838.8725 | 269 | 810.9142 | 305 | 757.6301 | 431 | 798.5133 | 548 | EP300 | MCF-7 | Breast | GSM16690 | 21375 | 54004 |
| 891.4779 | 441 | 864.6956 | 505 | 758.5447 | 620 | 789.5252 | 760 | EP300 | MCF-7 | Breast | GSM16690 | 33816 | 54005 |
| 42.02822 | 13 | 39.55699 | 14 | 22.03549 | 14 | 92.69436 | 31 | EP300 | MCF-7 | Breast | GSM16690 | 1113 | 54007 |
| 938.586 | 425 | 932.5122 | 510 | 803.7189 | 593 | 828.6044 | 716 | EP300 | MCF-7 | Breast | GSM16690 | 27305 | 54008 |
| 867.0375 | 439 | 843.6769 | 502 | 750.088 | 639 | 768.766 | 762 | EP300 | MCF-7 | Breast | GSM16690 | 36298 | 54009 |
| 915.4233 | 405 | 904.4529 | 482 | 800.2982 | 593 | 817.4333 | 704 | EP300 | MCF-7 | Breast | GSM16690 | 27829 | 54010 |
| 349.62 | 66 | 322.9993 | 69 | 560.7701 | 119 | 721.733 | 145 | ESR1 | MCF-7 | Breast | GSM16690 | 2675 | 54022 |
| 237.4628 | 80 | 270.629 | 95 | 374.511 | 146 | 610.598 | 199 | ESR1 | MCF-7 | Breast | GSM16690 | 6692 | 54023 |
| 235.7358 | 49 | 253.2708 | 56 | 356.2927 | 86 | 460.7907 | 105 | ESR1 | MCF-7 | Breast | GSM16690 | 2271 | 54024 |
| 355.5749 | 121 | 306.7903 | 125 | 577.7682 | 223 | 785.8858 | 284 | ESR1 | MCF-7 | Breast | GSM16690 | 10265 | 54025 |
| 145.9956 | 55 | 164.1135 | 65 | 292.7544 | 114 | 432.8537 | 148 | ESR1 | MCF-7 | Breast | GSM16690 | 5204 | 54027 |
| 392.4099 | 124 | 334.8082 | 127 | 561.0134 | 214 | 769.8835 | 267 | ESR1 | MCF-7 | Breast | GSM16690 | 9719 | 54029 |

| | | | | | | | | | | | | | |
|---|---|---|---|---|---|---|---|---|---|---|---|---|---|
| 239.7198 | 79 | 274.3916 | 94 | 554.9434 | 175 | 746.273 | 217 | ESR1 | MCF-7 | Breast | GSM16690 | 6442 | 54031 |
| 118.7653 | 39 | 122.3954 | 44 | 278.7297 | 88 | 466.6025 | 123 | ESR1 | MCF-7 | Breast | GSM16690 | 3200 | 54033 |
| 85.019 | 19 | 175.1386 | 32 | 68.55754 | 24 | 116.7013 | 34 | ESR1 | T47D | Mammary | GSM16691 | 1012 | 54047 |
| 568.6025 | 112 | 798.9793 | 149 | 689.4127 | 173 | 750.6012 | 192 | ESR1 | MCF-7 | Breast | GSM16691 | 4905 | 54078 |
| 145.0485 | 37 | 254.5226 | 56 | 161.5165 | 56 | 315.5677 | 85 | ESR1 | MCF-7 | Breast | GSM16691 | 2273 | 54079 |
| 1205.236 | 165 | 1229.213 | 221 | 832.0071 | 169 | 990.3442 | 201 | ESR1 | MCF-7 | Breast | GSM16691 | 3655 | 54080 |
| 243.3692 | 43 | 288.7412 | 52 | 226.0488 | 57 | 441.3972 | 88 | ESR1 | MCF-7 | Breast | GSM16691 | 1611 | 54081 |
| 145.7936 | 43 | 197.0193 | 56 | 291.3041 | 89 | 542.1857 | 132 | ESR1 | MCF-7 | Breast | GSM16691 | 3130 | 54082 |
| 198.6152 | 41 | 214.338 | 47 | 208.2636 | 59 | 426.7868 | 93 | ESR1 | MCF-7 | Breast | GSM16691 | 1915 | 54083 |
| 268.9942 | 67 | 297.1402 | 78 | 305.506 | 102 | 572.4921 | 151 | ESR1 | MCF-7 | Breast | GSM16691 | 3980 | 54084 |
| 165.4963 | 38 | 211.034 | 48 | 149.0322 | 51 | 389.047 | 91 | ESR1 | MCF-7 | Breast | GSM16691 | 2058 | 54085 |
| 869.3756 | 205 | 910.4006 | 246 | 804.1424 | 300 | 839.4751 | 375 | ESR1 | MCF-7 | Breast | GSM16691 | 11801 | 54086 |
| 227.128 | 58 | 332.7609 | 79 | 260.8652 | 89 | 463.3663 | 128 | ESR1 | MCF-7 | Breast | GSM16691 | 3542 | 54087 |
| 266.3652 | 79 | 389.317 | 107 | 344.8796 | 130 | 577.4394 | 180 | ESR1 | MCF-7 | Breast | GSM16691 | 5773 | 54088 |
| 198.8344 | 60 | 288.2019 | 81 | 375.0041 | 120 | 577.3441 | 160 | ESR1 | MCF-7 | Breast | GSM16691 | 4457 | 54089 |
| 819.775 | 594 | 835.826 | 743 | 636.5415 | 771 | 638.0549 | 883 | DAXX | PC-3 | Prostate | GSM16780 | 70252 | 54104 |
| 821.8949 | 481 | 802.6986 | 560 | 648.5734 | 624 | 649.743 | 714 | DAXX | PC-3 | Prostate | GSM16780 | 50068 | 54105 |
| 67.18021 | 25 | 87.51294 | 32 | 16.25766 | 20 | 124.9575 | 54 | DNMT1 | PC-3 | Prostate | GSM16780 | 2445 | 54106 |
| 2.176549 | 4 | 1.403519 | 4 | 3.77977 | 9 | 5.071863 | 11 | AGO2 | K562 | Bone Marr | GSM16822 | 1767 | 54115 |
| 0.568489 | 2 | 3.275768 | 5 | 0.786865 | 5 | 27.74209 | 20 | AGO2 | K562 | Bone Marr | GSM16822 | 1511 | 54118 |
| 236.7722 | 246 | 295.8323 | 305 | 156.6562 | 350 | 163.0591 | 388 | CEBPB | None | None | GSM16846 | 58283 | 54125 |
| 408.4684 | 400 | 510.2331 | 494 | 286.0362 | 579 | 314.2453 | 653 | CEBPB | None | None | GSM16846 | 97685 | 54126 |
| 366.6047 | 343 | 427.7825 | 414 | 273.3795 | 502 | 271.537 | 549 | CEBPB | None | None | GSM16846 | 79182 | 54127 |
| 21.63619 | 25 | 54.23129 | 42 | 30.89936 | 47 | 108.6295 | 84 | NR3C1 | None | None | GSM16846 | 6198 | 54128 |
| 43.58916 | 21 | 57.22439 | 27 | 7.661001 | 16 | 38.855 | 33 | ATF4 | None | None | GSM16846 | 2687 | 54129 |
| -1.053017 | 62 | -1.55208 | 75 | -0.708246 | 151 | 0 | 211 | CEBPB | None | None | GSM16846 | 102120 | 54131 |
| -3.274178 | 137 | -9.134347 | 149 | -1.469333 | 350 | -0.976057 | 411 | CEBPB | None | None | GSM16846 | 231737 | 54132 |
| -10.26854 | 223 | -20.99697 | 254 | -9.255154 | 555 | -3.729785 | 687 | CEBPB | None | None | GSM16846 | 395736 | 54133 |
| -13.72056 | 326 | -37.02129 | 357 | -13.74236 | 808 | -4.751864 | 997 | CEBPB | None | None | GSM16846 | 558492 | 54134 |
| -2.638933 | 68 | -2.534275 | 88 | -0.29631 | 198 | -0.106085 | 236 | CEBPB | None | None | GSM16846 | 124133 | 54136 |
| -0.198821 | 60 | -0.680607 | 68 | -0.287142 | 133 | -0.004224 | 172 | CEBPB | None | None | GSM16846 | 85402 | 54137 |
| -1.373252 | 49 | -1.068329 | 65 | -3.137867 | 105 | -0.178678 | 157 | CEBPB | None | None | GSM16846 | 85766 | 54138 |
| 2.043667 | 5 | 4.478788 | 8 | 1.631604 | 9 | 40.18424 | 31 | ATF4 | None | None | GSM16846 | 2374 | 54139 |
| 0.028448 | 12 | 1.400213 | 23 | 0.014922 | 25 | 0.347915 | 35 | ATF4 | None | None | GSM16846 | 13122 | 54140 |
| 756.3249 | 478 | 809.1808 | 647 | 568.218 | 594 | 531.1969 | 618 | CTCF | Jurkat | Blood | GSM16891 | 58997 | 54151 |
| 1.18308 | 2 | 5.682888 | 5 | 1.271599 | 4 | 26.88022 | 16 | FOXP1 | OCI-Ly1 | Bone Marr | GSM16900 | 1048 | 54152 |
| 7.738616 | 7 | 3.902498 | 6 | 1.873868 | 7 | 8.043452 | 13 | FOXP1 | OCI-Ly3 | Bone Marr | GSM16900 | 1736 | 54153 |
| 1.170739 | 3 | 4.10827 | 6 | 5.416598 | 10 | 4.220817 | 10 | FOXP1 | OCI-Ly7 | Blood | GSM16900 | 1685 | 54154 |
| 162.445 | 81 | 159.2655 | 90 | 48.34046 | 76 | 55.7411 | 87 | FOXP1 | OCI-Ly10 | Bone Marr | GSM16900 | 9781 | 54155 |
| 513.1097 | 65 | 827.4209 | 95 | 196.8507 | 50 | 417.3272 | 81 | E2F4 | GM06990 | Blood | GSM53706 | 1429 | 542 |
| 921.7051 | 506 | 958.0371 | 660 | 677.633 | 551 | 601.0618 | 515 | RUNX1 | Jurkat | Blood | GSM16978 | 38407 | 54234 |
| 339.9529 | 201 | 366.5099 | 234 | 229.3493 | 266 | 164.2778 | 257 | GATA3 | Jurkat | Blood | GSM16978 | 29794 | 54236 |
| 125.7796 | 86 | 150.0069 | 105 | 103.3952 | 125 | 164.7164 | 162 | PGR | None | Endometri | GSM17035 | 14325 | 54240 |
| 417.5158 | 272 | 498.5871 | 331 | 309.3722 | 381 | 359.9193 | 437 | FOSL2 | None | Endometri | GSM17035 | 47084 | 54241 |
| 769.5665 | 490 | 744.1922 | 564 | 554.4237 | 577 | 550.9523 | 652 | FOXO1 | None | Endometri | GSM17036 | 60281 | 54243 |
| 78.74001 | 62 | 100.428 | 78 | 53.13871 | 85 | 132.1714 | 130 | BRD4 | None | None | GSM17099 | 11313 | 54332 |
| 112.9687 | 61 | 103.6611 | 66 | 81.89486 | 82 | 135.725 | 109 | BRD4 | None | None | GSM17099 | 8261 | 54333 |
| 103.7675 | 78 | 213.2229 | 121 | 73.89628 | 108 | 82.26722 | 122 | FLI1 | EWS502 | None | GSM79020 | 13487 | 5437 |
| 437.7546 | 73 | 300.864 | 64 | 272.3518 | 77 | 312.7643 | 88 | LMTK3 | MCF-7 | Breast | GSM17263 | 2470 | 54380 |
| 425.8563 | 67 | 415.9834 | 72 | 305.0843 | 76 | 388.7374 | 92 | LMTK3 | MDA-MB-2 | Breast | GSM17263 | 2140 | 54381 |
| 873.3132 | 527 | 897.4242 | 669 | 632.1157 | 583 | 600.835 | 617 | FLI1 | EWS502 | None | GSM79020 | 46681 | 5439 |
| 1094.414 | 212 | 956.8792 | 189 | 973.1807 | 300 | 999.1559 | 366 | GATA2 | LNCaP | Prostate | GSM94119 | 7036 | 54396 |
| 979.8627 | 617 | 896.33 | 634 | 822.2922 | 845 | 843.8299 | 1005 | GATA2 | LNCaP | Prostate | GSM94119 | 44978 | 54397 |
| 164.4323 | 138 | 250.5511 | 186 | 143.5576 | 208 | 184.1592 | 247 | FLI1 | None | Umbilical V | GSM79020 | 26252 | 5441 |
| 832.9022 | 515 | 847.4636 | 641 | 605.0586 | 585 | 600.0144 | 657 | FLI1 | None | Umbilical V | GSM79021 | 50643 | 5443 |
| 275.3382 | 75 | 403.9661 | 102 | 350.9806 | 122 | 603.887 | 172 | VDR | LS180 | Colon | GSM79140 | 5006 | 5451 |
| 1036.715 | 327 | 1044.274 | 405 | 794.1666 | 335 | 788.962 | 375 | MYC | MCF-10A | Breast | GSM14294 | 14307 | 54511 |
| 499.3744 | 84 | 873.7241 | 128 | 253.1726 | 80 | 312.7027 | 95 | MYC | MCF-10A | Breast | GSM14294 | 2986 | 54512 |
| 1058.861 | 227 | 1058.128 | 277 | 803.5982 | 238 | 676.1762 | 232 | MYC | MCF-10A | Breast | GSM14294 | 8527 | 54513 |
| 470.4298 | 71 | 477.7091 | 78 | 237.5465 | 66 | 414.2199 | 95 | ZBTB17 | MCF-10A | Breast | GSM14294 | 2118 | 54514 |
| 740.2154 | 107 | 969.9694 | 137 | 459.505 | 111 | 506.8587 | 124 | MYC | MCF-10A | Breast | GSM14294 | 3014 | 54515 |
| 508.9269 | 77 | 551.8017 | 88 | 265.7926 | 73 | 372.254 | 93 | MYC | MCF-10A | Breast | GSM14294 | 2312 | 54516 |
| 901.6429 | 147 | 1098.069 | 199 | 500.8385 | 147 | 582.3125 | 169 | MYC | MCF-10A | Breast | GSM14294 | 5087 | 54517 |
| 701.7612 | 104 | 648.5854 | 108 | 329.4924 | 93 | 390.051 | 108 | MYC | MCF-10A | Breast | GSM14294 | 3043 | 54518 |
| 276.9983 | 124 | 407.63 | 167 | 313.8721 | 198 | 506.0732 | 263 | RXRA | LS180 | Colon | GSM79140 | 14719 | 5452 |
| 559.5428 | 159 | 457.2605 | 159 | 513.0304 | 221 | 774.337 | 303 | RXRA | LS180 | Colon | GSM79140 | 11735 | 5453 |
| 922.301 | 188 | 956.3949 | 218 | 698.6938 | 229 | 818.9156 | 277 | BCAT1 | LS180 | Colon | GSM79140 | 9179 | 5454 |
| 686.821 | 127 | 1034.29 | 176 | 669.7069 | 174 | 807.3994 | 204 | BCAT1 | LS180 | Colon | GSM79140 | 5234 | 5455 |
| 131.9745 | 56 | 184.7948 | 74 | 215.9574 | 106 | 416.8254 | 156 | TCF4 | LS180 | Colon | GSM79140 | 6202 | 5456 |
| 257.8974 | 88 | 347.4152 | 114 | 254.7297 | 129 | 419.3458 | 175 | TCF4 | LS180 | Colon | GSM79141 | 7792 | 5457 |
| 830.8975 | 319 | 846.1985 | 405 | 555.4013 | 356 | 611.9116 | 402 | HSF1 | U2OS | Bone | GSM14951 | 27322 | 54573 |
| 665.2706 | 265 | 763.0708 | 343 | 390.3991 | 317 | 405.5448 | 348 | HSF1 | U2OS | Bone | GSM14951 | 28753 | 54574 |

| | | | | | | | | | | | | | |
|---|---|---|---|---|---|---|---|---|---|---|---|---|---|
| 664.768 | 513 | 672.309 | 635 | 552.5917 | 773 | 548.5239 | 877 | CEBPB | LS180 | Colon | GSM79141 | 93547 | 5458 |
| 613.2239 | 527 | 624.1177 | 656 | 503.3553 | 802 | 507.1361 | 925 | CEBPB | LS180 | Colon | GSM79141 | 114797 | 5459 |
| 773.2393 | 295 | 776.9265 | 366 | 663.5284 | 440 | 725.6464 | 588 | CDX2 | LS180 | Colon | GSM79141 | 30415 | 5460 |
| 461.9004 | 224 | 519.9179 | 265 | 297.2568 | 284 | 244.1412 | 286 | YAP1 | SF268 | None | GSM15157 | 28190 | 54608 |
| 844.1813 | 259 | 835.6668 | 310 | 480.355 | 284 | 373.847 | 275 | YAP1 | SF268 | None | GSM15157 | 19640 | 54609 |
| 825.0412 | 440 | 814.0426 | 522 | 694.0322 | 625 | 701.5385 | 725 | CDX2 | LS180 | Colon | GSM79141 | 43136 | 5461 |
| 37.09657 | 89 | 30.99764 | 98 | 44.79267 | 160 | 44.89697 | 176 | TEAD1 | SF268 | None | GSM15157 | 30240 | 54611 |
| 5.614374 | 43 | 4.650882 | 49 | 9.210974 | 88 | 12.95692 | 105 | TEAD1 | SF268 | None | GSM15157 | 22937 | 54612 |
| 104.6641 | 27 | 153.6982 | 37 | 194.5182 | 54 | 380.0659 | 83 | ESR1 | MCF-7 | Breast | GSM15230 | 1714 | 54620 |
| 355.3121 | 116 | 334.6106 | 125 | 517.9408 | 203 | 804.2964 | 278 | ESR1 | MCF-7 | Breast | GSM15230 | 9555 | 54621 |
| 169.6321 | 51 | 155.5232 | 54 | 255.0201 | 91 | 436.2759 | 128 | NCAPG | MCF-7 | Breast | GSM15230 | 3817 | 54623 |
| 108.441 | 37 | 107.706 | 41 | 160.3617 | 66 | 330.9872 | 102 | NCAPG | MCF-7 | Breast | GSM15230 | 3188 | 54625 |
| 221.6045 | 74 | 225.8268 | 83 | 385.1977 | 142 | 566.8557 | 184 | NCAPG2 | MCF-7 | Breast | GSM15230 | 6175 | 54631 |
| 1173.98 | 185 | 1192.058 | 236 | 1009.148 | 229 | 983.8106 | 241 | NCAPG | MCF-7 | Breast | GSM15230 | 4561 | 54632 |
| 1160.874 | 200 | 1173.952 | 256 | 998.7666 | 255 | 957.9658 | 255 | NCAPG | MCF-7 | Breast | GSM15230 | 5282 | 54633 |
| 317.1962 | 43 | 389.4108 | 53 | 106.2198 | 31 | 274.191 | 56 | HECTD1 | MCF-7 | Breast | GSM15230 | 1062 | 54634 |
| 300.5995 | 48 | 409.029 | 63 | 134.7921 | 42 | 239.7632 | 61 | EP300 | MCF-7 | Breast | GSM15230 | 1521 | 54636 |
| 104.4379 | 27 | 199.308 | 43 | 95.34305 | 37 | 294.8268 | 72 | EP300 | MCF-7 | Breast | GSM15230 | 1712 | 54637 |
| 245.4908 | 56 | 331.8094 | 73 | 268.2324 | 83 | 448.0594 | 116 | EP300 | MCF-7 | Breast | GSM15230 | 2974 | 54639 |
| 169.5961 | 34 | 347.4329 | 57 | 111.8988 | 38 | 185.899 | 53 | EP300 | MCF-7 | Breast | GSM15230 | 1522 | 54640 |
| 263.8646 | 136 | 296.0713 | 161 | 437.6302 | 257 | 690.0592 | 349 | ESR1 | MCF-7 | Breast | GSM15230 | 17910 | 54645 |
| 174.6733 | 82 | 222.135 | 103 | 321.6228 | 163 | 591.5105 | 233 | ESR1 | MCF-7 | Breast | GSM15230 | 9710 | 54646 |
| 51.85949 | 20 | 79.83979 | 28 | 127.9494 | 47 | 235.8538 | 69 | ESR1 | MCF-7 | Breast | GSM15230 | 2054 | 54647 |
| 50.77879 | 23 | 103.1539 | 37 | 150.2854 | 59 | 269.8376 | 85 | ESR1 | MCF-7 | Breast | GSM15230 | 2751 | 54648 |
| 629.2464 | 201 | 778.7148 | 248 | 264.8677 | 199 | 180.1545 | 184 | TLX1 | ALL-SIL | Blood | GSM15242 | 15971 | 54657 |
| 102.7577 | 30 | 150.8667 | 41 | 58.01463 | 33 | 185.7818 | 63 | HNF4A | HCT-116 | Colon | GSM15356 | 2213 | 54660 |
| 28.00483 | 18 | 58.71711 | 29 | 61.34184 | 40 | 125.1532 | 60 | TCF4 | HCT-116 | Colon | GSM15356 | 2978 | 54661 |
| 29.50786 | 14 | 61.75815 | 23 | 67.87961 | 32 | 112.193 | 44 | TCF4 | HCT-116 | Colon | GSM15356 | 1807 | 54663 |
| 14.29258 | 10 | 22.11269 | 14 | 45.68324 | 27 | 89.12393 | 40 | TCF4 | HCT-116 | Colon | GSM15357 | 1884 | 54664 |
| 323.5981 | 76 | 445.8801 | 100 | 183.7828 | 81 | 303.6374 | 111 | HNF4A | HCT-116 | Colon | GSM15357 | 4203 | 54665 |
| 209.9396 | 55 | 211.7425 | 61 | 78.72519 | 49 | 167.0456 | 75 | HNF4A | HCT-116 | Colon | GSM15357 | 3473 | 54666 |
| 10.76425 | 9 | 35.04922 | 18 | 43.27936 | 27 | 122.1788 | 48 | TCF4 | HCT-116 | Colon | GSM15357 | 1967 | 54667 |
| 22.58664 | 11 | 37.23248 | 16 | 31.89841 | 20 | 128.5868 | 43 | TCF4 | HCT-116 | Colon | GSM15357 | 1494 | 54669 |
| 7.394105 | 5 | 18.08136 | 9 | 20.17541 | 13 | 134.6574 | 37 | TCF4 | HCT-116 | Colon | GSM15357 | 1033 | 54670 |
| 341.6522 | 171 | 434.41 | 214 | 304.017 | 247 | 383.1234 | 293 | HNF4A | HCT-116 | Colon | GSM15357 | 21805 | 54673 |
| 259.5493 | 122 | 372.1962 | 162 | 211.1564 | 169 | 344.489 | 224 | HNF4A | HCT-116 | Colon | GSM15357 | 14641 | 54674 |
| 238.4572 | 299 | 224.2541 | 336 | 131.7804 | 409 | 141.6592 | 459 | YAP1 | NCI-H2052 | Lung | GSM15425 | 75038 | 54675 |
| 116.9305 | 220 | 124.3408 | 259 | 118.6591 | 368 | 104.5262 | 392 | YAP1 | NCI-H2052 | Lung | GSM15425 | 67509 | 54676 |
| 210.9123 | 66 | 226.3467 | 76 | 134.2238 | 78 | 172.8994 | 94 | PPARG | None | Adipose | GSM15717 | 5289 | 55122 |
| 143.8157 | 48 | 203.4509 | 64 | 145.0816 | 71 | 201.5654 | 89 | PPARG | None | Adipose | GSM15717 | 4122 | 55123 |
| 331.9231 | 100 | 443.4363 | 129 | 236.8976 | 124 | 265.3192 | 140 | PPARG | None | Adipose | GSM15717 | 7666 | 55124 |
| 82.17239 | 29 | 140.9975 | 43 | 43.13604 | 32 | 218.1958 | 75 | PPARG | None | Adipose | GSM15717 | 2709 | 55125 |
| 115.486 | 26 | 72.23634 | 22 | 77.82194 | 30 | 251.1094 | 60 | PPARG | None | Adipose | GSM15717 | 1409 | 55126 |
| 237.1883 | 55 | 269.2831 | 65 | 123.1786 | 56 | 306.6176 | 95 | PPARG | None | Adipose | GSM15717 | 3037 | 55127 |
| 696.7203 | 119 | 733.9607 | 134 | 318.3483 | 108 | 485.04 | 143 | PPARG | None | Adipose | GSM15717 | 4348 | 55128 |
| 967.0443 | 139 | 798.3937 | 135 | 461.0829 | 125 | 425.0781 | 127 | PPARG | None | Adipose | GSM15717 | 3898 | 55129 |
| 1064.994 | 188 | 1024.888 | 208 | 547.6854 | 176 | 592.5986 | 195 | PPARG | None | Adipose | GSM15717 | 6729 | 55130 |
| 234.5288 | 35 | 285.7178 | 43 | 116.4127 | 32 | 345.3254 | 63 | PPARG | None | Adipose | GSM15717 | 1030 | 55131 |
| 232.1089 | 41 | 279.0914 | 50 | 219.582 | 55 | 253.1145 | 63 | PPARG | None | Adipose | GSM15717 | 1555 | 55132 |
| 146.6685 | 32 | 207.6332 | 43 | 105.4511 | 38 | 201.294 | 57 | PPARG | None | Adipose | GSM15717 | 1646 | 55133 |
| 226.6904 | 74 | 324.1399 | 99 | 186.8753 | 99 | 269.5783 | 126 | PPARG | None | Adipose | GSM15717 | 6142 | 55135 |
| 555.81 | 93 | 656.8599 | 112 | 249.5077 | 83 | 446.2556 | 120 | PPARG | None | Adipose | GSM15717 | 3260 | 55136 |
| 219.1484 | 41 | 254.4478 | 49 | 147.4762 | 46 | 362.2894 | 80 | PPARG | None | Adipose | GSM15717 | 1696 | 55137 |
| 32.59005 | 16 | 19.22495 | 14 | 64.8664 | 34 | 109.3606 | 47 | EPAS1 | 786-O | Kidney | GSM85679 | 2121 | 5516 |
| 901.4904 | 471 | 834.0793 | 491 | 734.0394 | 607 | 809.881 | 824 | AR | LNCaP | Prostate | GSM15764 | 35379 | 55162 |
| 1143.463 | 702 | 1131.501 | 833 | 938.5704 | 890 | 966.6564 | 1067 | VDR | LNCaP | Prostate | GSM15764 | 33063 | 55164 |
| 729.1366 | 631 | 610.5983 | 607 | 591.1157 | 899 | 645.5228 | 1137 | FOXA1 | LNCaP | Prostate | GSM15764 | 100956 | 55166 |
| 828.4198 | 157 | 1019.087 | 204 | 565.8781 | 179 | 414.846 | 163 | ARNT | 786-O | Kidney | GSM85679 | 6757 | 5517 |
| 815.3013 | 518 | 818.6595 | 630 | 610.2443 | 626 | 577.942 | 661 | RUNX1 | None | None | GSM15817 | 56309 | 55172 |
| 810.3557 | 174 | 949.1257 | 210 | 634.7033 | 215 | 892.051 | 335 | AR | BicR | Prostate | GSM16133 | 8524 | 55273 |
| 216.0478 | 92 | 244.0809 | 109 | 99.5173 | 99 | 116.5136 | 114 | YAP1 | MDA-MB-2 | Breast | GSM16140 | 9727 | 55291 |
| 87.26186 | 52 | 91.93254 | 60 | 39.7137 | 59 | 50.38932 | 70 | YAP1 | MDA-MB-2 | Breast | GSM16140 | 7471 | 55292 |
| 98.11037 | 65 | 158.7623 | 91 | 46.23465 | 76 | 84.40718 | 103 | WWTR1 | MDA-MB-2 | Breast | GSM16140 | 10055 | 55293 |
| 105.2628 | 60 | 124.9732 | 73 | 34.79975 | 60 | 80.35436 | 89 | WWTR1 | MDA-MB-2 | Breast | GSM16140 | 8145 | 55294 |
| 182.4026 | 97 | 121.1526 | 91 | 99.9385 | 116 | 76.50403 | 114 | TEAD4 | MDA-MB-2 | Breast | GSM16140 | 12560 | 55297 |
| 127.3725 | 57 | 75.60538 | 50 | 41.99062 | 54 | 44.67547 | 60 | TEAD4 | MDA-MB-2 | Breast | GSM16140 | 6247 | 55298 |
| 361.1094 | 273 | 412.6111 | 326 | 249.559 | 375 | 347.3893 | 457 | JUN | MDA-MB-2 | Breast | GSM16140 | 49031 | 55299 |
| 329.3181 | 206 | 434.4662 | 261 | 267.7921 | 291 | 313.742 | 335 | JUN | MDA-MB-2 | Breast | GSM16140 | 30126 | 55300 |
| 571.9978 | 314 | 491.0762 | 333 | 567.1585 | 493 | 581.6665 | 583 | FOXA2 | Caco-2 | Colon | GSM16174 | 47808 | 55321 |
| 265.4044 | 146 | 252.9165 | 161 | 452.6289 | 280 | 658.1672 | 356 | FOXA2 | Caco-2 | Colon | GSM16174 | 20675 | 55322 |
| 398.5257 | 138 | 364.6405 | 147 | 214.1787 | 154 | 187.7196 | 157 | KDM1A | NCI-H526 | Lung | GSM16190 | 12304 | 55323 |
| 838.206 | 346 | 807.5928 | 390 | 585.633 | 377 | 488.5051 | 378 | KDM1A | NCI-H526 | Lung | GSM16190 | 29172 | 55324 |

| | | | | | | | | | | | | | |
|---|---|---|---|---|---|---|---|---|---|---|---|---|---|
| 669.8836 | 169 | 618.4866 | 179 | 228.5114 | 146 | 327.037 | 184 | SUMO2 | K562 | Bone Marr | GSM16226 | 10672 | 55336 |
| 719.4864 | 445 | 720.2151 | 539 | 504.8321 | 529 | 430.8978 | 546 | SUMO2 | K562 | Bone Marr | GSM16226 | 60180 | 55337 |
| 981.9253 | 548 | 1012.23 | 706 | 714.55 | 573 | 673.3445 | 588 | SUMO2 | K562 | Bone Marr | GSM16226 | 34387 | 55338 |
| 970.5092 | 506 | 991.102 | 640 | 707.4033 | 522 | 675.1523 | 548 | SUMO2 | K562 | Bone Marr | GSM16226 | 31467 | 55339 |
| 250.1612 | 132 | 337.0648 | 170 | 102.0593 | 143 | 157.3052 | 182 | SUMO2 | K562 | Bone Marr | GSM16226 | 17995 | 55342 |
| 196.2941 | 61 | 181.5539 | 65 | 49.6717 | 48 | 100.6545 | 70 | SUMO2 | K562 | Bone Marr | GSM16226 | 4816 | 55343 |
| 1112.766 | 338 | 1134.42 | 434 | 851.6199 | 330 | 814.8347 | 339 | SUMO2 | K562 | Bone Marr | GSM16226 | 11410 | 55344 |
| 1088.037 | 136 | 1216.58 | 177 | 719.336 | 143 | 694.2889 | 148 | SUMO2 | K562 | Bone Marr | GSM16226 | 2987 | 55345 |
| 92.77765 | 41 | 110.3374 | 50 | 28.61365 | 38 | 106.7729 | 71 | SUMO2 | K562 | Bone Marr | GSM16226 | 4708 | 55346 |
| 331.9555 | 113 | 377.0072 | 134 | 108.356 | 102 | 133.7865 | 120 | SUMO2 | K562 | Bone Marr | GSM16226 | 9942 | 55347 |
| 1126.73 | 243 | 1124.856 | 295 | 809.8743 | 223 | 847.4301 | 245 | SUMO2 | K562 | Bone Marr | GSM16226 | 7212 | 55348 |
| 358.8276 | 48 | 438.6744 | 59 | 231.3911 | 50 | 306.066 | 62 | SUMO2 | K562 | Bone Marr | GSM16226 | 1170 | 55349 |
| 1160.5 | 610 | 1123.998 | 688 | 978.3234 | 780 | 991.4031 | 919 | SUMO2 | VCaP | Prostate | GSM16226 | 23685 | 55350 |
| 1222.63 | 912 | 1227.334 | 1086 | 987.1944 | 1167 | 991.5942 | 1343 | SUMO2 | VCaP | Prostate | GSM16226 | 47893 | 55351 |
| 1208.655 | 853 | 1153.349 | 956 | 943.8611 | 1023 | 969.5711 | 1220 | SUMO2 | VCaP | Prostate | GSM16226 | 41940 | 55352 |
| 1176.553 | 706 | 1153.164 | 819 | 957.1763 | 869 | 963.3866 | 1004 | SUMO2 | VCaP | Prostate | GSM16226 | 29874 | 55353 |
| 1135.426 | 478 | 1093.302 | 526 | 1009.74 | 685 | 1034.282 | 823 | PIAS1 | VCaP | Prostate | GSM16226 | 17488 | 55354 |
| 1127.073 | 547 | 1080.003 | 603 | 1000.096 | 780 | 1005.639 | 906 | PIAS1 | VCaP | Prostate | GSM16226 | 22011 | 55355 |
| 586.2404 | 73 | 481.3825 | 70 | 590.3717 | 98 | 1056.763 | 146 | PIAS1 | VCaP | Prostate | GSM16226 | 1591 | 55356 |
| 1116.486 | 272 | 1057.066 | 284 | 1028.805 | 425 | 1060.61 | 527 | PIAS1 | VCaP | Prostate | GSM16226 | 8772 | 55357 |
| 18.18969 | 12 | 19.24209 | 14 | 15.729 | 18 | 23.34898 | 23 | TP53 | SW480 | Colon | GSM16389 | 2112 | 55387 |
| 581.8103 | 198 | 817.3883 | 261 | 393.6248 | 238 | 546.6087 | 296 | SPI1 | 786-O | Kidney | GSM16427 | 14794 | 55389 |
| 39.4498 | 20 | 84.85449 | 33 | 98.09783 | 47 | 264.6989 | 83 | ESR1 | H3396 | Mammary | GSM81120 | 2645 | 5539 |
| 34.44866 | 27 | 69.84067 | 42 | 65.03049 | 55 | 51.60042 | 54 | SPI1 | 786-O | Kidney | GSM16427 | 4653 | 55390 |
| 956.4532 | 220 | 972.8522 | 283 | 588.9934 | 231 | 520.6412 | 233 | SPI1 | 786-O | Kidney | GSM16427 | 9930 | 55392 |
| 1074.395 | 317 | 1131.413 | 440 | 847.9079 | 328 | 807.8737 | 331 | SPI1 | 786-O | Kidney | GSM16427 | 10209 | 55393 |
| 40.7148 | 34 | 73.87298 | 50 | 51.96477 | 59 | 42.01789 | 59 | SPI1 | 786-O | Kidney | GSM16427 | 6123 | 55394 |
| 186.9743 | 57 | 348.2486 | 88 | 223.333 | 89 | 195.3469 | 89 | SPI1 | 786-O | Kidney | GSM16427 | 3842 | 55396 |
| 422.5258 | 182 | 556.396 | 231 | 371.9587 | 254 | 285.6438 | 246 | SPI1 | 786-O | Kidney | GSM16427 | 18382 | 55397 |
| 1150.992 | 500 | 1185.005 | 644 | 860.2828 | 482 | 817.4042 | 491 | KMT2D | OCI-Ly7 | Blood | GSM16446 | 17036 | 55405 |
| 23.16365 | 10 | 56.05426 | 18 | 28.74241 | 17 | 223.1125 | 53 | ESR1 | H3396 | Mammary | GSM81120 | 1227 | 5541 |
| 6.990267 | 5 | 39.63137 | 14 | 19.02984 | 13 | 148.9881 | 40 | ESR1 | H3396 | Mammary | GSM81121 | 1089 | 5543 |
| 369.6126 | 170 | 424.1845 | 203 | 621.9551 | 321 | 717.4146 | 421 | ESR1 | H3396 | Mammary | GSM81121 | 20353 | 5546 |
| 1281.053 | 486 | 1309.939 | 617 | 972.27 | 438 | 934.4799 | 450 | NELFA | HeLa | Cervix | GSM16708 | 10589 | 55465 |
| 1298.444 | 561 | 1341.376 | 731 | 955.1047 | 491 | 909.8694 | 493 | NELFA | HeLa | Cervix | GSM16708 | 12814 | 55466 |
| 1187.199 | 302 | 1197.892 | 373 | 904.0144 | 275 | 865.1244 | 277 | NELFA | HeLa | Cervix | GSM16708 | 7800 | 55467 |
| 1233.944 | 493 | 1269.651 | 638 | 929.6653 | 452 | 911.2067 | 485 | NELFA | HeLa | Cervix | GSM16708 | 12494 | 55468 |
| 212.2437 | 87 | 288.7604 | 113 | 403.6339 | 175 | 656.6346 | 237 | ESR1 | H3396 | Mammary | GSM81121 | 9034 | 5547 |
| 136.8519 | 29 | 116.0615 | 29 | 143.4701 | 42 | 268.957 | 63 | CREB1 | H3396 | Mammary | GSM80104 | 1442 | 5548 |
| 5.702806 | 12 | 1.815894 | 10 | 0.283349 | 11 | 19.59002 | 35 | FOS | MV4-11 | Blood | GSM16905 | 4541 | 55489 |
| 1079.403 | 206 | 1102.371 | 268 | 818.505 | 221 | 807.0888 | 233 | CREB1 | H3396 | Mammary | GSM80105 | 7079 | 5549 |
| 35.81637 | 15 | 40.8738 | 18 | 8.130355 | 12 | 113.166 | 43 | FOS | MV4-11 | Blood | GSM16905 | 1729 | 55490 |
| 948.2548 | 505 | 964.708 | 636 | 700.5752 | 548 | 686.3247 | 600 | RUNX1 | None | None | GSM16933 | 34538 | 55492 |
| 1152.879 | 762 | 1168.217 | 930 | 814.5797 | 749 | 773.111 | 785 | RUNX1 | None | Blood | GSM16933 | 37079 | 55493 |
| 91.00864 | 50 | 194.7308 | 81 | 116.9211 | 86 | 146.3141 | 102 | CREB1 | H3396 | Mammary | GSM80105 | 7064 | 5551 |
| 81.81887 | 94 | 59.72538 | 96 | 50.94542 | 131 | 43.59792 | 137 | YAP1 | IMR90 | Lung | GSM16962 | 21681 | 55512 |
| 127.1075 | 133 | 99.96712 | 139 | 92.47221 | 192 | 60.76533 | 185 | YAP1 | IMR90 | Lung | GSM16962 | 28630 | 55513 |
| 20.311 | 57 | 45.59388 | 86 | 8.351052 | 81 | 27.48457 | 119 | ATF4 | HAP-1 | None | GSM16973 | 21874 | 55516 |
| 49.4389 | 117 | 73.92656 | 154 | 19.02691 | 160 | 28.13396 | 193 | ATF4 | HAP-1 | None | GSM16973 | 41155 | 55517 |
| 44.41794 | 86 | 73.61956 | 118 | 16.99843 | 114 | 46.94497 | 163 | ATF4 | HAP-1 | None | GSM16973 | 27651 | 55518 |
| 33.18958 | 25 | 32.02094 | 28 | 4.120175 | 20 | 18.55184 | 35 | CREB1 | H3396 | Mammary | GSM80105 | 4692 | 5552 |
| 105.8684 | 23 | 202.105 | 37 | 85.05127 | 29 | 96.5648 | 33 | CREB1 | H3396 | Mammary | GSM80105 | 1203 | 5553 |
| 237.6684 | 41 | 179.7203 | 38 | 144.3821 | 43 | 231.1633 | 59 | CREB1 | H3396 | Mammary | GSM80105 | 1509 | 5554 |
| 1105.387 | 664 | 1137.214 | 837 | 809.927 | 664 | 751.4626 | 662 | RAG2 | None | None | GSM17018 | 29112 | 55543 |
| 25.65497 | 15 | 36.15128 | 20 | 6.974512 | 14 | 32.2373 | 28 | MEF2B | None | None | GSM17038 | 2312 | 55547 |
| 31.52418 | 22 | 43.91891 | 29 | 18.06263 | 28 | 20.58973 | 32 | MEF2B | None | None | GSM17038 | 3782 | 55548 |
| 82.20913 | 38 | 57.60409 | 36 | 71.26497 | 54 | 141.2854 | 79 | BACH2 | None | None | GSM17038 | 4439 | 55549 |
| 232.436 | 49 | 234.4045 | 54 | 142.8798 | 53 | 338.3185 | 89 | ESR1 | H3396 | Mammary | GSM80105 | 2264 | 5555 |
| 5.737911 | 5 | 66.25717 | 20 | 15.61214 | 13 | 85.89271 | 32 | PAX5 | None | None | GSM17038 | 1273 | 55554 |
| 668.163 | 449 | 682.4339 | 565 | 378.9983 | 537 | 436.4771 | 614 | SPI1 | None | None | GSM17039 | 74265 | 55555 |
| 0.016825 | 15 | 0 | 17 | 0.283056 | 38 | 2.400047 | 56 | MEF2B | None | None | GSM17039 | 15975 | 55563 |
| 21.91627 | 19 | 22.65231 | 22 | 9.809601 | 23 | 30.6196 | 38 | MEF2B | KARPAS-42 | None | GSM17039 | 3926 | 55566 |
| 84.1837 | 32 | 55.40829 | 29 | 57.63025 | 40 | 118.4849 | 60 | BACH2 | None | None | GSM17039 | 3105 | 55571 |
| 4.116444 | 6 | 17.14278 | 13 | 7.370837 | 13 | 58.69048 | 34 | EP300 | None | None | GSM17039 | 2012 | 55572 |
| 169.586 | 73 | 349.539 | 117 | 141.1561 | 101 | 197.9479 | 126 | PAX5 | None | None | GSM17039 | 7939 | 55576 |
| 463.894 | 241 | 629.4982 | 310 | 254.8724 | 293 | 238.9764 | 311 | SPI1 | None | None | GSM17039 | 33630 | 55577 |
| 10.46773 | 11 | 18.05225 | 16 | 7.192219 | 16 | 51.38226 | 38 | MEF2B | None | None | GSM17039 | 2747 | 55584 |
| 26.41983 | 30 | 27.64353 | 35 | 19.10116 | 44 | 17.73873 | 47 | MEF2B | None | None | GSM17039 | 7061 | 55585 |
| 364.4595 | 62 | 453.477 | 77 | 326.9496 | 80 | 488.9947 | 106 | RAC3 | H3396 | Mammary | GSM80106 | 2139 | 5562 |
| 976.0029 | 123 | 1226.748 | 165 | 446.9693 | 105 | 483.2016 | 116 | ELL2 | HCT-116 | Colon | GSM17270 | 2690 | 55645 |
| 685.449 | 75 | 740.8891 | 85 | 271.398 | 57 | 285.9092 | 62 | ELL2 | HCT-116 | Colon | GSM17270 | 1267 | 55646 |
| 649.2573 | 77 | 828.4534 | 97 | 245.9753 | 58 | 265.2662 | 64 | ELL2 | HCT-116 | Colon | GSM17270 | 1491 | 55647 |

| | | | | | | | | | | | | | |
|---|---|---|---|---|---|---|---|---|---|---|---|---|---|
| 396.2688 | 49 | 444.5641 | 57 | 156.6472 | 38 | 155.1393 | 40 | ELL2 | HCT-116 | Colon | GSM17270 | 1030 | 55648 |
| 1298.305 | 565 | 1339.946 | 736 | 973.4004 | 518 | 927.221 | 521 | PAF1 | HCT-116 | Colon | GSM17270 | 13093 | 55664 |
| 1236.565 | 289 | 1263.917 | 379 | 952.3738 | 262 | 942.5124 | 292 | PAF1 | HCT-116 | Colon | GSM17270 | 6299 | 55665 |
| 1268.167 | 285 | 1284.835 | 368 | 1008.384 | 277 | 980.594 | 290 | PAF1 | HCT-116 | Colon | GSM17270 | 5685 | 55666 |
| 1189.409 | 597 | 1245.332 | 787 | 878.0184 | 551 | 843.7309 | 575 | PAF1 | HCT-116 | Colon | GSM17270 | 17855 | 55667 |
| 1189.612 | 370 | 1222.298 | 482 | 911.2192 | 345 | 879.4714 | 355 | PAF1 | HCT-116 | Colon | GSM17270 | 9380 | 55668 |
| 1218.755 | 346 | 1256.746 | 458 | 938.4784 | 324 | 913.9225 | 344 | PAF1 | HCT-116 | Colon | GSM17270 | 8226 | 55669 |
| 1242.051 | 190 | 1261.197 | 243 | 1013.477 | 195 | 920.8673 | 191 | PAF1 | MCF-7 | Breast | GSM17271 | 3677 | 55682 |
| 1131.107 | 246 | 1126.191 | 298 | 900.777 | 259 | 884.5095 | 282 | POLR2M | HeLa | Cervix | GSM80274 | 7515 | 5569 |
| 1138.493 | 582 | 1159.791 | 731 | 849.6167 | 590 | 837.2447 | 649 | POLR2M | HeLa | Cervix | GSM80274 | 23795 | 5570 |
| 1016.773 | 645 | 976.8541 | 724 | 817.6735 | 816 | 857.8648 | 1009 | SUMO2 | VCaP | Prostate | GSM17775 | 42625 | 55715 |
| 1037.293 | 641 | 1007.005 | 729 | 880.7755 | 877 | 901.0776 | 1044 | SUMO2 | VCaP | Prostate | GSM17775 | 39637 | 55716 |
| 1209.893 | 965 | 1158.609 | 1096 | 957.6714 | 1240 | 981.0252 | 1469 | SUMO2 | VCaP | Prostate | GSM17775 | 61222 | 55717 |
| 1379.61 | 1157 | 1282.753 | 1304 | 994.665 | 1460 | 1005.526 | 1698 | SUMO2 | VCaP | Prostate | GSM17776 | 86103 | 55718 |
| 1299.073 | 1063 | 1241.762 | 1209 | 977.0557 | 1321 | 999.9777 | 1560 | SUMO2 | VCaP | Prostate | GSM17776 | 66423 | 55721 |
| 1220.588 | 926 | 1175.868 | 1047 | 974.7123 | 1172 | 963.506 | 1321 | SUMO2 | VCaP | Prostate | GSM17776 | 49938 | 55722 |
| 971.0529 | 508 | 933.2569 | 565 | 844.9619 | 729 | 865.4794 | 870 | SUMO2 | VCaP | Prostate | GSM17776 | 32827 | 55723 |
| 956.0834 | 540 | 921.7783 | 609 | 836.2326 | 789 | 864.3789 | 952 | SUMO2 | VCaP | Prostate | GSM17776 | 38178 | 55724 |
| 1014.32 | 670 | 967.8147 | 740 | 857.8516 | 922 | 883.3329 | 1100 | SUMO2 | VCaP | Prostate | GSM17776 | 45888 | 55725 |
| 1052.599 | 789 | 987.2391 | 862 | 862.3181 | 1052 | 884.9293 | 1248 | SUMO2 | VCaP | Prostate | GSM17776 | 57847 | 55726 |
| 639.7675 | 314 | 733.9265 | 429 | 440.0883 | 413 | 472.1453 | 460 | CTCF | K562 | Bone Marr | GSM17827 | 42703 | 55727 |
| 445.9746 | 257 | 696.3759 | 366 | 328.161 | 353 | 353.94 | 394 | CTCF | K562 | Bone Marr | GSM17827 | 39520 | 55728 |
| 137.3448 | 147 | 230.8473 | 205 | 177.0315 | 260 | 197.8156 | 294 | GATA1 | K562 | Bone Marr | GSM17827 | 34354 | 55729 |
| 807.5671 | 290 | 810.1096 | 357 | 642.7644 | 368 | 570.0502 | 377 | GATA1 | None | Blood | GSM80401 | 24990 | 5573 |
| 162.6712 | 214 | 204.1057 | 266 | 156.8352 | 349 | 167.8179 | 390 | GATA1 | K562 | Bone Marr | GSM17827 | 58399 | 55730 |
| 500.126 | 163 | 538.7422 | 188 | 263.4786 | 179 | 242.2707 | 186 | SPI1 | K562 | Bone Marr | GSM17827 | 13783 | 55738 |
| 552.2585 | 314 | 665.2811 | 399 | 348.4079 | 410 | 345.7419 | 445 | SPI1 | K562 | Bone Marr | GSM17827 | 49832 | 55739 |
| 554.9112 | 186 | 800.6824 | 248 | 412.128 | 237 | 383.5608 | 247 | GATA1 | None | Blood | GSM80401 | 15637 | 5574 |
| 492.3147 | 98 | 590.127 | 119 | 261.744 | 98 | 215.9166 | 95 | REST | K562 | Bone Marr | GSM17827 | 4233 | 55740 |
| 420.4854 | 82 | 561.1881 | 106 | 225.5087 | 82 | 255.0585 | 93 | REST | K562 | Bone Marr | GSM17827 | 3446 | 55741 |
| 780.9216 | 331 | 769.8499 | 393 | 684.2512 | 509 | 728.7606 | 646 | AR | DUCaP | Prostate | GSM18163 | 33906 | 55752 |
| 236.2242 | 40 | 305.8482 | 51 | 170.8004 | 46 | 248.5599 | 60 | EP300 | None | Heart | GSM80773 | 1429 | 5577 |
| 733.7577 | 508 | 796.6373 | 700 | 566.3681 | 672 | 548.0025 | 737 | CTCF | K562 | Bone Marr | GSM18176 | 72049 | 55770 |
| 906.0883 | 266 | 977.9628 | 396 | 727.352 | 327 | 710.1404 | 353 | CTCFL | K562 | Bone Marr | GSM18176 | 17152 | 55771 |
| 779.5013 | 397 | 824.3328 | 536 | 599.5034 | 496 | 569.7517 | 523 | CTCF | Delta47 | None | GSM18176 | 42494 | 55774 |
| 1045.655 | 562 | 1076.305 | 722 | 765.3335 | 574 | 724.3008 | 591 | CTCFL | Delta47 | None | GSM18176 | 30248 | 55775 |
| 437.1047 | 236 | 676.903 | 322 | 289.0326 | 310 | 295.2384 | 339 | CTCF | Ovcar8 | None | GSM18176 | 34800 | 55778 |
| 976.828 | 620 | 1002.962 | 780 | 698.7028 | 651 | 646.9289 | 656 | CTCFL | Ovcar8 | None | GSM18176 | 43980 | 55779 |
| 1050.789 | 260 | 1074.832 | 340 | 804.3182 | 261 | 594.0897 | 239 | EP300 | None | Heart | GSM80773 | 10197 | 5578 |
| 570.0848 | 357 | 685.3549 | 500 | 401.315 | 491 | 382.7495 | 527 | CTCF | MCF-7 | Breast | GSM18176 | 62287 | 55781 |
| 629.7502 | 402 | 683.886 | 553 | 437.1609 | 552 | 467.4809 | 616 | CTCF | MCF-7 | Breast | GSM18176 | 72691 | 55782 |
| 665.2313 | 450 | 721.6298 | 621 | 519.8784 | 623 | 502.901 | 685 | CTCF | MCF-7 | Breast | GSM18176 | 75993 | 55783 |
| 949.9029 | 286 | 1010.949 | 409 | 724.6148 | 309 | 706.707 | 334 | CTCFL | MCF-7 | Breast | GSM18176 | 16304 | 55784 |
| 932.2617 | 348 | 987.5659 | 486 | 738.3971 | 413 | 681.3815 | 403 | CTCFL | MCF-7 | Breast | GSM18176 | 22046 | 55785 |
| 499.7093 | 101 | 684.2649 | 132 | 321.7779 | 110 | 258.4442 | 105 | BRD4 | TC-797 | None | GSM18209 | 3987 | 55812 |
| 551.4743 | 73 | 633.4877 | 86 | 308.0031 | 69 | 183.9845 | 55 | BRD4 | 293T | None | GSM18209 | 1541 | 55816 |
| 431.4962 | 66 | 505.5476 | 79 | 271.3476 | 69 | 250.666 | 70 | BRD4 | TC-797 | None | GSM18209 | 1940 | 55821 |
| 594.1548 | 100 | 780.1626 | 128 | 321.7766 | 98 | 332.8205 | 106 | BRD4 | TC-797 | None | GSM18209 | 3371 | 55822 |
| 101.1633 | 24 | 126.3949 | 30 | 100.8587 | 34 | 184.1659 | 50 | BRD4 | TC-797 | None | GSM18209 | 1330 | 55825 |
| 152.726 | 34 | 136.1112 | 35 | 119.5307 | 42 | 195.6147 | 58 | BRD4 | TC-797 | None | GSM18209 | 1702 | 55826 |
| 219.1814 | 63 | 285.723 | 80 | 122.2633 | 67 | 91.29158 | 63 | BRD4 | TC-797 | None | GSM18209 | 3739 | 55844 |
| 711.2475 | 100 | 729.5377 | 110 | 342.4749 | 88 | 648.5953 | 133 | BRD4 | 293T-Rex | None | GSM18209 | 2306 | 55868 |
| 833.9895 | 339 | 847.5844 | 431 | 626.2564 | 387 | 599.4344 | 410 | T | None | None | GSM18257 | 28368 | 55895 |
| 800.1208 | 239 | 847.9377 | 305 | 381.547 | 250 | 425.6351 | 282 | VDR | GM10855 | Blood | GSM55863 | 19158 | 559 |
| 268.678 | 47 | 257.1735 | 50 | 85.60482 | 35 | 79.48276 | 36 | T | None | None | GSM18257 | 1683 | 55903 |
| 871.8822 | 294 | 911.7058 | 399 | 574.6724 | 320 | 535.7725 | 334 | VDR | GM10855 | Blood | GSM55863 | 21888 | 560 |
| 1040.791 | 215 | 1045.181 | 266 | 676.0382 | 218 | 541.8757 | 208 | CDK12 | Jurkat | Blood | GSM18502 | 8394 | 56010 |
| 126.5356 | 65 | 269.1087 | 105 | 91.92867 | 87 | 131.9523 | 109 | RNF2 | OS25 | Embryo | GSM85047 | 8475 | 5603 |
| 789.6651 | 553 | 832.4598 | 728 | 588.4094 | 685 | 575.3005 | 760 | RAD21 | HepG2 | Liver | GSM18619 | 69655 | 56037 |
| 640.3499 | 380 | 660.1638 | 480 | 468.836 | 526 | 503.9305 | 600 | RAD21 | HepG2 | Liver | GSM18619 | 63527 | 56038 |
| 741.9576 | 460 | 752.5433 | 573 | 584.2385 | 618 | 583.7916 | 703 | RAD21 | HepG2 | Liver | GSM18619 | 60876 | 56039 |
| 786.5186 | 583 | 782.0208 | 703 | 607.3663 | 765 | 606.5609 | 873 | RAD21 | HepG2 | Liver | GSM18619 | 75862 | 56040 |
| 786.3915 | 580 | 784.6251 | 702 | 609.3581 | 761 | 601.3909 | 854 | RAD21 | HepG2 | Liver | GSM18619 | 75382 | 56041 |
| 1177.265 | 367 | 1207.719 | 479 | 905.3607 | 347 | 881.1555 | 371 | GABPA | HepG2 | Liver | GSM18619 | 9906 | 56042 |
| 1194.441 | 322 | 1223.563 | 417 | 915.1821 | 294 | 891.2178 | 312 | GABPA | HepG2 | Liver | GSM18619 | 7950 | 56043 |
| 1126.357 | 438 | 1165.47 | 574 | 867.4053 | 447 | 834.3554 | 465 | GABPA | HepG2 | Liver | GSM18619 | 15361 | 56044 |
| 1021.202 | 606 | 1061.355 | 787 | 768.7644 | 668 | 734.4905 | 706 | GABPA | HepG2 | Liver | GSM18619 | 36611 | 56045 |
| 1060.528 | 592 | 1085.828 | 746 | 804.4868 | 644 | 760.1331 | 664 | GABPA | HepG2 | Liver | GSM18619 | 31030 | 56046 |
| 1053.042 | 732 | 1066.874 | 896 | 764.2476 | 789 | 732.0339 | 839 | CREB1 | HepG2 | Liver | GSM18619 | 46641 | 56047 |
| 1117.165 | 562 | 1138.636 | 707 | 840.3833 | 575 | 796.7128 | 592 | CREB1 | HepG2 | Liver | GSM18619 | 23371 | 56048 |
| 1179.128 | 405 | 1216.594 | 527 | 912.4075 | 398 | 859.0359 | 393 | CREB1 | HepG2 | Liver | GSM18619 | 11727 | 56049 |
| 1102.226 | 611 | 1126.567 | 767 | 817.5671 | 629 | 791.9793 | 677 | CREB1 | HepG2 | Liver | GSM18619 | 29130 | 56050 |

| | | | | | | | | | | | | | |
|---|---|---|---|---|---|---|---|---|---|---|---|---|---|
| 1143.92 | 654 | 1150.087 | 800 | 841.8489 | 660 | 800.8266 | 685 | CREB1 | HepG2 | Liver | GSM18619 | 28662 | 56051 |
| 1126.195 | 583 | 1162.188 | 745 | 841.0654 | 586 | 809.6101 | 618 | CREB1 | HepG2 | Liver | GSM18619 | 24102 | 56052 |
| 1009.131 | 634 | 1066.477 | 836 | 753.7088 | 701 | 755.3015 | 806 | ATF1 | HepG2 | Liver | GSM18619 | 41427 | 56053 |
| 1067.823 | 607 | 1103.787 | 780 | 798.0914 | 646 | 777.8638 | 701 | ATF1 | HepG2 | Liver | GSM18619 | 31805 | 56054 |
| 690.9775 | 387 | 690.0922 | 473 | 589.5387 | 592 | 571.4287 | 649 | RAD21 | MCF-7 | Breast | GSM18619 | 56515 | 56055 |
| 539.1446 | 320 | 660.6561 | 412 | 544.9295 | 498 | 522.568 | 535 | RAD21 | MCF-7 | Breast | GSM18619 | 52390 | 56056 |
| 498.7491 | 283 | 637.0144 | 354 | 385.2046 | 396 | 425.8551 | 446 | RAD21 | MCF-7 | Breast | GSM18619 | 43882 | 56057 |
| 676.2248 | 370 | 701.3452 | 480 | 591.8304 | 589 | 590.8256 | 668 | RAD21 | MCF-7 | Breast | GSM18619 | 55293 | 56058 |
| 684.4905 | 375 | 689.5831 | 463 | 574.367 | 561 | 570.0351 | 634 | RAD21 | MCF-7 | Breast | GSM18619 | 54722 | 56059 |
| 705.8328 | 429 | 713.1352 | 533 | 569.069 | 604 | 567.6025 | 687 | RAD21 | MCF-7 | Breast | GSM18619 | 61517 | 56060 |
| 1105.565 | 300 | 1096.979 | 360 | 802.3824 | 262 | 806.762 | 299 | VDR | GM10855 | Blood | GSM55863 | 10384 | 561 |
| 57.60671 | 23 | 38.49193 | 21 | 64.47526 | 36 | 84.85426 | 44 | NR3C1 | MDA-MB-2 | Breast | GSM13505 | 2236 | 56103 |
| 479.4964 | 126 | 579.695 | 154 | 295.3296 | 143 | 586.0725 | 213 | AR | None | Prostate | GSM13583 | 8229 | 56106 |
| 973.5621 | 393 | 902.064 | 405 | 832.9937 | 543 | 904.5131 | 733 | AR | None | Prostate | GSM13583 | 22747 | 56107 |
| 372.3173 | 111 | 613.4139 | 160 | 334.0648 | 153 | 586.9944 | 215 | AR | None | Prostate | GSM13583 | 8375 | 56108 |
| 966.9916 | 294 | 909.7535 | 312 | 801.256 | 374 | 879.9076 | 525 | AR | None | Prostate | GSM13583 | 15744 | 56109 |
| 980.0615 | 180 | 993.4315 | 206 | 810.0744 | 226 | 948.2727 | 340 | AR | None | Prostate | GSM13583 | 7493 | 56110 |
| 948.53 | 378 | 900.4321 | 408 | 796.1089 | 503 | 845.5187 | 649 | AR | None | Prostate | GSM13584 | 23031 | 56111 |
| 1086.047 | 269 | 1108.753 | 347 | 899.2118 | 319 | 917.0539 | 386 | AR | None | Prostate | GSM13584 | 9612 | 56112 |
| 1161.61 | 263 | 1202.868 | 357 | 944.8441 | 288 | 977.8694 | 360 | AR | None | Prostate | GSM13584 | 7225 | 56113 |
| 987.4893 | 169 | 943.5559 | 180 | 730.9733 | 198 | 952.927 | 292 | AR | None | Prostate | GSM13584 | 6235 | 56114 |
| 1085.088 | 616 | 1082.135 | 734 | 866.6031 | 733 | 908.077 | 915 | AR | None | Prostate | GSM13584 | 30073 | 56115 |
| 547.6327 | 150 | 802.8181 | 203 | 461.7488 | 199 | 783.1411 | 282 | AR | None | Prostate | GSM13584 | 10394 | 56116 |
| 977.3693 | 250 | 928.6488 | 266 | 819.4125 | 325 | 900.0919 | 461 | AR | None | Prostate | GSM13584 | 12457 | 56118 |
| 963.6861 | 327 | 903.4614 | 341 | 811.2712 | 428 | 860.1 | 555 | AR | None | Prostate | GSM13584 | 17926 | 56119 |
| 137.7796 | 38 | 189.1157 | 50 | 158.4416 | 59 | 375.6593 | 99 | AR | None | Prostate | GSM13584 | 2612 | 56120 |
| 964.2493 | 274 | 924.9652 | 301 | 809.888 | 357 | 880.7067 | 493 | AR | None | Prostate | GSM13584 | 14465 | 56121 |
| 1125.776 | 830 | 1109.404 | 966 | 900.6211 | 1045 | 924.8498 | 1242 | AR | None | Prostate | GSM13584 | 49882 | 56122 |
| 1007.185 | 371 | 987.211 | 433 | 849.3949 | 489 | 900.5518 | 630 | AR | None | Prostate | GSM13584 | 18771 | 56123 |
| 965.2335 | 376 | 906.7043 | 394 | 817.4789 | 501 | 873.0208 | 650 | AR | None | Prostate | GSM13584 | 20671 | 56124 |
| 888.0881 | 314 | 854.2443 | 351 | 771.0662 | 451 | 827.538 | 597 | AR | None | Prostate | GSM13584 | 21431 | 56125 |
| 48.75275 | 17 | 53.96941 | 20 | 125.5575 | 41 | 413.0222 | 84 | APOBEC3B | MCF-7 | Breast | GSM13725 | 1576 | 56126 |
| 651.5118 | 166 | 656.883 | 184 | 795.1699 | 261 | 780.18 | 282 | APOBEC3B | MCF-7 | Breast | GSM13725 | 10236 | 56127 |
| 44.7585 | 18 | 35.80461 | 18 | 93.76504 | 39 | 253.7241 | 70 | H2AFX | MCF-7 | Breast | GSM13824 | 1946 | 56129 |
| 265.872 | 105 | 265.0248 | 117 | 468.2151 | 203 | 780.3237 | 285 | H2AFX | MCF-7 | Breast | GSM13824 | 10576 | 56131 |
| 132.5378 | 50 | 153.3653 | 60 | 302.7482 | 111 | 525.808 | 157 | H2AFX | MCF-7 | Breast | GSM13824 | 4799 | 56132 |
| 820.455 | 162 | 1002.042 | 211 | 369.0182 | 151 | 416.4104 | 171 | VDR | GM10855 | Blood | GSM55863 | 7458 | 562 |
| 490.9921 | 108 | 561.9118 | 128 | 322.1138 | 123 | 906.861 | 233 | AR | LNCaP | Prostate | GSM15011 | 5633 | 56205 |
| 485.0786 | 97 | 531.0644 | 112 | 307.9189 | 107 | 914.3715 | 203 | AR | LNCaP | Prostate | GSM15011 | 4429 | 56206 |
| 977.4619 | 274 | 969.2815 | 326 | 795.2728 | 331 | 903.3656 | 504 | AR | LNCaP | Prostate | GSM15011 | 14063 | 56207 |
| 1102.798 | 135 | 1189.217 | 154 | 870.4643 | 157 | 1106.2 | 225 | AR | C4-2B | Prostate | GSM15011 | 2902 | 56208 |
| 701.3163 | 112 | 709.7712 | 123 | 484.4213 | 125 | 1015.968 | 220 | AR | C4-2B | Prostate | GSM15011 | 3680 | 56209 |
| 712.6632 | 108 | 636.484 | 110 | 556.1471 | 128 | 1061.048 | 222 | AR | C4-2B | Prostate | GSM15011 | 3271 | 56211 |
| 836.8605 | 401 | 819.4357 | 468 | 666.7327 | 512 | 665.405 | 581 | EP300 | HeLa | Cervix | GSM15170 | 35294 | 56231 |
| 757.5066 | 274 | 761.5264 | 322 | 516.1282 | 343 | 647.4008 | 439 | EP300 | HeLa | Cervix | GSM15170 | 26715 | 56232 |
| 767.9306 | 359 | 762.3588 | 432 | 615.2192 | 472 | 611.1313 | 536 | EP300 | HeLa | Cervix | GSM15170 | 37898 | 56233 |
| 863.4863 | 472 | 870.4965 | 577 | 677.7628 | 585 | 668.3373 | 649 | EP300 | HeLa | Cervix | GSM15170 | 40456 | 56234 |
| 25.37934 | 15 | 22.33224 | 16 | 11.34166 | 17 | 48.94107 | 34 | YAP1 | CClp1 | None | GSM15243 | 2387 | 56248 |
| 1229.158 | 496 | 1262.683 | 641 | 924.8477 | 453 | 898.7742 | 485 | E2F1 | None | Mammary | GSM15268 | 12709 | 56252 |
| 1172.759 | 233 | 1115.774 | 243 | 960.6814 | 257 | 1059.907 | 380 | AR | LNCaP | Prostate | GSM15270 | 6046 | 56254 |
| 899.5803 | 550 | 815.5162 | 558 | 714.0512 | 698 | 799.2976 | 954 | AR | LNCaP | Prostate | GSM15270 | 47429 | 56255 |
| 975.6255 | 567 | 869.5016 | 554 | 783.0049 | 717 | 866.2047 | 975 | AR | LNCaP | Prostate | GSM15270 | 39533 | 56256 |
| 1141.894 | 322 | 1089.889 | 343 | 957.2734 | 387 | 1028.498 | 531 | AR | LNCaP | Prostate | GSM15270 | 9946 | 56257 |
| 1323.573 | 1040 | 1267.05 | 1182 | 964.9937 | 1203 | 1002.406 | 1452 | AR | CWR22Pc | Prostate | GSM15270 | 54097 | 56264 |
| 1415.307 | 1187 | 1291.985 | 1329 | 931.3019 | 1412 | 962.3074 | 1676 | AR | CWR22Pc | Prostate | GSM15270 | 91442 | 56265 |
| 1207.502 | 847 | 1161.223 | 952 | 940.2364 | 991 | 977.6933 | 1209 | AR | CWR22Pc | Prostate | GSM15270 | 38678 | 56267 |
| 14.26136 | 8 | 20.93444 | 11 | 12.59473 | 12 | 32.55451 | 20 | FOXP1 | OF4155 | None | GSM15322 | 1302 | 56275 |
| 260.2393 | 42 | 321.2513 | 52 | 135.554 | 40 | 224.6952 | 56 | BRD1 | HEK293T | Embryonic | GSM15337 | 1381 | 56279 |
| 1065.413 | 313 | 1089.038 | 408 | 812.5214 | 312 | 798.0246 | 339 | CDK8 | MOLM-14 | None | GSM15878 | 11969 | 56351 |
| 1149.285 | 418 | 1193.009 | 560 | 896.7161 | 430 | 852.9642 | 433 | CDK8 | MOLM-14 | None | GSM15879 | 13044 | 56353 |
| 1042.911 | 197 | 1047.656 | 244 | 679.7875 | 206 | 642.1908 | 213 | MED1 | MOLM-14 | None | GSM15879 | 7043 | 56355 |
| 1165.21 | 443 | 1181.003 | 552 | 877.6981 | 404 | 820.3752 | 392 | BRD4 | MOLM-14 | None | GSM15879 | 11839 | 56357 |
| 1137.633 | 187 | 1156.796 | 242 | 762.8099 | 186 | 775.8238 | 199 | BRD4 | MOLM-14 | None | GSM15879 | 4980 | 56359 |
| 1076.404 | 292 | 1102.949 | 376 | 847.638 | 304 | 795.7269 | 296 | CDK8 | SET2 | Blood | GSM15879 | 10251 | 56369 |
| 1125.154 | 298 | 1125.624 | 366 | 871.0831 | 289 | 818.3174 | 281 | CDK8 | MV4-11 | Blood | GSM15879 | 8910 | 56375 |
| 1158.766 | 294 | 1186.064 | 380 | 884.7098 | 272 | 874.4457 | 300 | CDK8 | K562 | Bone Marr | GSM15879 | 7861 | 56379 |
| 1045.9 | 372 | 1014.843 | 420 | 959.6058 | 584 | 1002.582 | 729 | AR | None | Prostate | GSM15982 | 16297 | 56390 |
| 484.0027 | 71 | 534.6039 | 82 | 455.5288 | 93 | 861.9254 | 142 | AR | None | Prostate | GSM15982 | 2006 | 56391 |
| 966.7829 | 169 | 1026.154 | 201 | 921.0094 | 245 | 997.4846 | 344 | AR | None | Prostate | GSM15982 | 6460 | 56392 |
| 839.6977 | 300 | 858.0227 | 383 | 474.3813 | 315 | 475.5302 | 340 | VDR | GM10861 | Blood | GSM55863 | 24872 | 564 |
| 74.07159 | 88 | 124.5492 | 123 | 104.358 | 162 | 125.589 | 188 | GATA3 | KELLY | None | GSM16026 | 21516 | 56401 |
| 29.473 | 47 | 50.53205 | 66 | 86.31478 | 114 | 141.3501 | 149 | GATA3 | SH-SY5Y | Bone Marr | GSM16026 | 13666 | 56402 |

| | | | | | | | | | | | | | |
|---|---|---|---|---|---|---|---|---|---|---|---|---|---|
| 10.44571 | 7 | 23.78201 | 12 | 22.49723 | 16 | 111.8944 | 38 | GATA3 | BE2-C | Brain | GSM16026 | 1360 | 56403 |
| 1048.146 | 294 | 1056.99 | 363 | 789.9674 | 288 | 780.0316 | 316 | MYC | None | Breast | GSM16180 | 11978 | 56427 |
| 28.8803 | 12 | 23.01101 | 12 | 16.03255 | 14 | 87.5518 | 34 | TOP2B | MCF-7 | Breast | GSM16314 | 1425 | 56433 |
| 55.96311 | 25 | 118.4469 | 41 | 64.20634 | 40 | 64.25308 | 43 | EHMT2 | HEK293T | Embryonic | GSM16446 | 2746 | 56446 |
| 34.01287 | 14 | 54.25343 | 20 | 6.077723 | 10 | 35.1112 | 23 | EHMT2 | HEK293T | Embryonic | GSM16446 | 1507 | 56447 |
| 0.307355 | 14 | 0.4572 | 18 | 1.900646 | 37 | 2.127544 | 42 | EHMT2 | HEK293T | Embryonic | GSM16446 | 12104 | 56448 |
| 17.17761 | 17 | 116.3442 | 47 | 15.78138 | 27 | 47.31005 | 45 | EHMT2 | HEK293T | Embryonic | GSM16446 | 3863 | 56449 |
| 129.9551 | 37 | 130.6123 | 41 | 80.76099 | 42 | 96.49603 | 49 | EHMT2 | HEK293T | Embryonic | GSM16446 | 2460 | 56454 |
| 23.40417 | 15 | 36.59054 | 21 | 11.83222 | 18 | 15.95922 | 22 | EHMT2 | HEK293T | Embryonic | GSM16446 | 2490 | 56455 |
| 2.382377 | 10 | 9.922466 | 19 | 3.653921 | 21 | 13.45414 | 34 | EHMT2 | HEK293T | Embryonic | GSM16446 | 5200 | 56456 |
| 24.17453 | 17 | 53.56002 | 28 | 26.63183 | 28 | 32.72868 | 33 | EHMT2 | HEK293T | Embryonic | GSM16446 | 2943 | 56457 |
| 50.33848 | 24 | 80.65497 | 34 | 25.38354 | 27 | 26.99043 | 30 | EHMT2 | HEK293T | Embryonic | GSM16446 | 2792 | 56462 |
| 241.2156 | 84 | 326.8854 | 109 | 111.2665 | 87 | 59.68593 | 73 | EHMT2 | HEK293T | Embryonic | GSM16446 | 7199 | 56463 |
| 78.13888 | 28 | 162.383 | 46 | 33.28586 | 28 | 59.24433 | 39 | EHMT2 | HEK293T | Embryonic | GSM16446 | 2529 | 56466 |
| 365.4634 | 93 | 628.7394 | 138 | 216.8119 | 102 | 191.6352 | 103 | EHMT2 | HEK293T | Embryonic | GSM16446 | 5451 | 56467 |
| 1097.29 | 231 | 1127.492 | 308 | 827.3548 | 231 | 644.0208 | 216 | KMT2D | None | Tonsil | GSM16480 | 7488 | 56478 |
| 1020.394 | 141 | 1162.011 | 183 | 665.6834 | 149 | 585.8992 | 147 | KMT2D | None | Tonsil | GSM16480 | 3600 | 56481 |
| 902.0011 | 370 | 917.2295 | 464 | 614.2241 | 362 | 636.5818 | 412 | VDR | GM10861 | Blood | GSM55863 | 26295 | 565 |
| 12.43358 | 7 | 27.7845 | 12 | 14.65634 | 12 | 115.8164 | 36 | YAP1 | MSTO | None | GSM16649 | 1165 | 56534 |
| 210.0285 | 164 | 232.8413 | 194 | 181.1838 | 245 | 255.1147 | 301 | TEAD1 | MSTO | None | GSM16649 | 30230 | 56535 |
| 6.186612 | 6 | 27.26205 | 14 | 3.299702 | 8 | 68.82759 | 32 | YAP1 | HuCCT1 | None | GSM16671 | 1582 | 56538 |
| 196.2767 | 202 | 238.1287 | 248 | 201.085 | 329 | 266.8398 | 394 | TEAD1 | HuCCT1 | None | GSM16671 | 46788 | 56542 |
| 125.6757 | 92 | 178.1516 | 121 | 126.1359 | 145 | 182.3845 | 181 | TEAD4 | HuCCT1 | None | GSM16671 | 16081 | 56543 |
| 902.7767 | 277 | 908.6285 | 341 | 726.9194 | 333 | 684.4609 | 336 | MED1 | HuCCT1 | None | GSM16707 | 16591 | 56555 |
| 963.2907 | 214 | 938.9297 | 246 | 698.4809 | 247 | 770.1986 | 279 | MED1 | HuCCT1 | None | GSM16707 | 10258 | 56556 |
| 806.7653 | 659 | 835.1231 | 831 | 610.1029 | 842 | 597.8158 | 939 | SMC1A | HuCCT1 | None | GSM16707 | 86222 | 56557 |
| 855.3449 | 762 | 879.3791 | 945 | 636.7823 | 960 | 635.1464 | 1096 | SMC1A | HuCCT1 | None | GSM16707 | 97515 | 56558 |
| 268.5043 | 43 | 299.0254 | 50 | 187.4143 | 48 | 267.5404 | 62 | CDK9 | HuCCT1 | None | GSM16707 | 1384 | 56565 |
| 64.555 | 103 | 113.8602 | 145 | 114.1924 | 206 | 157.3742 | 250 | GATA3 | BE2-C | Brain | GSM16801 | 29669 | 56598 |
| 1077.921 | 358 | 1066.541 | 430 | 799.8208 | 342 | 780.6016 | 368 | VDR | GM10861 | Blood | GSM55863 | 14332 | 566 |
| 9.639314 | 25 | 26.01616 | 41 | 33.87559 | 65 | 72.68115 | 93 | GATA3 | NGP | None | GSM16801 | 9680 | 56601 |
| 205.9034 | 51 | 336.0111 | 74 | 108.5508 | 53 | 153.0641 | 67 | POLR2M | IMR90 | Lung | GSM82053 | 3098 | 5662 |
| 107.2398 | 89 | 168.5937 | 122 | 105.9984 | 140 | 191.6825 | 190 | JUN | MDA-MB-2 | Breast | GSM17007 | 16428 | 56632 |
| 135.9835 | 98 | 170.3917 | 122 | 78.63681 | 125 | 210.3425 | 196 | JUN | MDA-MB-2 | Breast | GSM17007 | 16195 | 56633 |
| 146.4232 | 103 | 163.316 | 122 | 97.66311 | 138 | 226.7179 | 206 | JUN | MDA-MB-2 | Breast | GSM17007 | 16858 | 56634 |
| 939.6644 | 613 | 965.5144 | 773 | 799.3466 | 863 | 826.9404 | 1041 | AR | VCaP | Prostate | GSM80101 | 49684 | 5666 |
| 1144.561 | 227 | 1064.077 | 222 | 946.0472 | 259 | 997.5396 | 338 | FOXA1 | None | Prostate | GSM17167 | 6144 | 56662 |
| 1010.438 | 337 | 933.9943 | 338 | 877.4027 | 474 | 938.9161 | 624 | HOXB13 | None | Prostate | GSM17167 | 16394 | 56663 |
| 990.5247 | 530 | 901.228 | 528 | 831.1069 | 710 | 878.8892 | 899 | HOXB13 | LNCaP | Prostate | GSM17167 | 32910 | 56664 |
| 428.0063 | 438 | 311.9309 | 454 | 270.8563 | 621 | 443.5686 | 800 | FOXA1 | LHSAR | Prostate | GSM17167 | 112132 | 56665 |
| 236.3141 | 376 | 177.5371 | 400 | 211.7262 | 606 | 378.8024 | 778 | HOXB13 | LHSAR | Prostate | GSM17167 | 118089 | 56666 |
| 361.7096 | 483 | 283.5605 | 519 | 321.8193 | 766 | 417.0704 | 917 | HOXB13 | LHSAR | Prostate | GSM17167 | 144211 | 56667 |
| 339.6925 | 551 | 185.3205 | 547 | 212.387 | 820 | 358.5038 | 1020 | HOXB13 | LHSAR | Prostate | GSM17167 | 186049 | 56668 |
| 581.8541 | 477 | 578.6431 | 575 | 474.9695 | 719 | 502.8431 | 873 | AR | LHSAR | Prostate | GSM17167 | 102104 | 56669 |
| 922.6557 | 566 | 935.0151 | 703 | 758.7518 | 760 | 789.6195 | 923 | AR | VCS2 | Prostate | GSM80101 | 46101 | 5667 |
| 247.4499 | 346 | 295.0566 | 421 | 336.7663 | 618 | 458.6637 | 764 | AR | LHSAR | Prostate | GSM17167 | 98096 | 56670 |
| 509.9763 | 115 | 644.2467 | 144 | 201.8888 | 103 | 304.2951 | 134 | VDR | GM10861 | Blood | GSM55863 | 6216 | 567 |
| 1382.052 | 150 | 1303.453 | 149 | 1194.633 | 174 | 1206.559 | 204 | NKX3-1 | LNCaP | Prostate | GSM69963 | 1874 | 5677 |
| 1204.234 | 294 | 1141.678 | 303 | 1008.593 | 342 | 1058.522 | 445 | NKX3-1 | LNCaP | Prostate | GSM69963 | 7160 | 5678 |
| 885.8157 | 684 | 788.2064 | 693 | 725.9143 | 942 | 765.06 | 1155 | FOXA1 | LNCaP | Prostate | GSM69963 | 73478 | 5679 |
| 850.4206 | 596 | 736.9131 | 571 | 675.8934 | 784 | 726.2526 | 992 | FOXA1 | LNCaP | Prostate | GSM69963 | 64243 | 5680 |
| 1032.855 | 581 | 1075.982 | 755 | 746.7076 | 587 | 710.0238 | 608 | EP300 | None | Cranial | GSM18171 | 32751 | 56814 |
| 1074.036 | 617 | 1115.902 | 797 | 773.045 | 607 | 746.6718 | 649 | EP300 | None | Cranial | GSM18171 | 30411 | 56815 |
| 912.7027 | 489 | 922.6056 | 607 | 666.3117 | 523 | 643.5929 | 564 | EP300 | None | Cranial | GSM18171 | 36071 | 56816 |
| 779.0507 | 668 | 824.3405 | 866 | 563.0019 | 824 | 550.7886 | 919 | NR2F1 | None | Cranial | GSM18171 | 95781 | 56817 |
| 827.3531 | 462 | 847.8201 | 587 | 602.9898 | 513 | 599.8529 | 581 | NR2F1 | None | Cranial | GSM18171 | 41349 | 56818 |
| 943.062 | 770 | 1001.645 | 998 | 664.6948 | 873 | 639.9282 | 951 | TFAP2A | None | Cranial | GSM18171 | 75942 | 56819 |
| 895.5384 | 768 | 949.0408 | 990 | 636.1246 | 897 | 606.9933 | 973 | TFAP2A | None | Cranial | GSM18171 | 87066 | 56820 |
| 892.9069 | 760 | 940.4124 | 971 | 645.8999 | 906 | 603.1111 | 955 | TFAP2A | None | Cranial | GSM18172 | 85008 | 56821 |
| 153.8161 | 126 | 119.0417 | 129 | 130.1549 | 184 | 85.63976 | 174 | PBX3 | SMMC-772 | Liver | GSM18316 | 21138 | 56901 |
| 713.4431 | 97 | 1006.884 | 130 | 416.8766 | 96 | 512.9762 | 114 | ESR1 | MCF-7 | Breast | GSM18317 | 2411 | 56903 |
| 438.0056 | 161 | 774.7433 | 238 | 644.754 | 284 | 756.0529 | 368 | ESR1 | MCF-7 | Breast | GSM18317 | 15413 | 56904 |
| 617.4937 | 215 | 795.7457 | 274 | 700.7943 | 344 | 746.4298 | 446 | ESR1 | MCF-7 | Breast | GSM18317 | 19887 | 56905 |
| 104.867 | 129 | 150.5498 | 169 | 97.47511 | 207 | 64.40709 | 200 | SPI1 | RS4 | None | GSM18412 | 32739 | 56943 |
| 10.81682 | 20 | 49.64406 | 42 | 40.06703 | 54 | 65.96297 | 71 | NR3C1 | RS4 | None | GSM18413 | 6599 | 56951 |
| 180.979 | 207 | 274.7502 | 276 | 149.5335 | 318 | 82.38502 | 294 | SPI1 | RS4 | None | GSM18413 | 51505 | 56953 |
| 1221.463 | 201 | 1222.304 | 249 | 841.0519 | 182 | 789.9364 | 186 | GTF2B | HeLa | Cervix | GSM18467 | 4317 | 56958 |
| 1284.153 | 563 | 1314.364 | 708 | 933.8817 | 471 | 918.4697 | 520 | GTF2B | HeLa | Cervix | GSM18467 | 13001 | 56961 |
| 1056.528 | 223 | 1080.093 | 297 | 694.967 | 221 | 646.7726 | 227 | GTF2B | HeLa | Cervix | GSM18467 | 8446 | 56962 |
| 1246.467 | 340 | 1279.737 | 449 | 955.9123 | 308 | 930.5981 | 328 | GTF2B | HeLa | Cervix | GSM18468 | 7481 | 56978 |
| 1240.915 | 265 | 1266.444 | 341 | 956.8456 | 239 | 934.1032 | 253 | GTF2B | HeLa | Cervix | GSM18468 | 5542 | 56982 |
| 1018.562 | 410 | 1083 | 569 | 753.1582 | 405 | 717.3389 | 420 | TRIM28 | HCT-116 | Colon | GSM18666 | 20151 | 57090 |

| | | | | | | | | | | | | | |
|---|---|---|---|---|---|---|---|---|---|---|---|---|---|
| 853.018 | 372 | 871.7443 | 479 | 618.7538 | 402 | 597.8181 | 433 | LARP7 | HCT-116 | Colon | GSM18666 | 31507 | 57091 |
| 1015.83 | 662 | 1061.594 | 863 | 729.4375 | 699 | 697.4849 | 742 | HEXIM1 | HCT-116 | Colon | GSM18666 | 44697 | 57092 |
| 1045.449 | 281 | 1064.527 | 361 | 793.1135 | 278 | 733.3753 | 279 | CDK9 | HCT-116 | Colon | GSM18666 | 11498 | 57093 |
| 538.5646 | 62 | 841.7172 | 89 | 249.0657 | 52 | 416.4669 | 74 | EPO | RL95-2 | Uterus | GSM18751 | 1154 | 57110 |
| 428.8837 | 52 | 703.3877 | 77 | 206.3518 | 45 | 451.1872 | 75 | EPO | RL95-2 | Uterus | GSM18751 | 1063 | 57111 |
| 994.5995 | 117 | 1282.271 | 165 | 491.3431 | 103 | 754.6458 | 139 | LEF1 | RL95-2 | Uterus | GSM18751 | 2294 | 57114 |
| 1272.813 | 187 | 1328.877 | 269 | 800.728 | 159 | 959.3686 | 186 | LEF1 | RL95-2 | Uterus | GSM18751 | 3324 | 57115 |
| 981.2184 | 225 | 957.7438 | 256 | 849.8551 | 312 | 868.2546 | 372 | FOXM1 | MCF-7 | Breast | GSM18761 | 10808 | 57117 |
| 627.7455 | 109 | 503.9248 | 105 | 580.0371 | 144 | 816.9776 | 184 | FOXM1 | MCF-7 | Breast | GSM18761 | 4055 | 57118 |
| 988.3141 | 275 | 929.9655 | 289 | 840.6703 | 366 | 867.6911 | 451 | FOXM1 | MCF-7 | Breast | GSM18761 | 13667 | 57119 |
| 802.5405 | 593 | 816.262 | 733 | 576.699 | 687 | 549.7476 | 737 | CDK8 | MOLM-14 | None | GSM18939 | 67035 | 57165 |
| 802.5798 | 580 | 811.9333 | 713 | 579.7398 | 677 | 561.8488 | 742 | CDK8 | MOLM-14 | None | GSM18939 | 64893 | 57167 |
| 888.3713 | 509 | 913.657 | 643 | 654.1782 | 555 | 616.5577 | 579 | CDK8 | MOLM-14 | None | GSM18939 | 38850 | 57169 |
| 952.8538 | 427 | 989.9321 | 561 | 695.4624 | 426 | 671.0881 | 454 | CDK8 | MOLM-14 | None | GSM18939 | 23900 | 57171 |
| 1064.221 | 506 | 1060.321 | 607 | 776.2985 | 478 | 739.1716 | 492 | BRD4 | MOLM-14 | None | GSM18939 | 20198 | 57173 |
| 1142.663 | 408 | 1149.519 | 497 | 857.1417 | 368 | 794.1962 | 353 | BRD4 | MOLM-14 | None | GSM18939 | 11429 | 57175 |
| 1084.123 | 471 | 1088.898 | 575 | 776.9281 | 426 | 765.4657 | 473 | BRD4 | MOLM-14 | None | GSM18939 | 18947 | 57177 |
| 31.59816 | 58 | 37.62193 | 71 | 9.653661 | 71 | 24.6003 | 99 | NANOG | None | None | GSM19014 | 16664 | 57188 |
| 746.2703 | 225 | 811.3177 | 262 | 659.1461 | 309 | 765.3443 | 429 | AR | MDA-MB-4 | Mammary | GSM19090 | 18023 | 57277 |
| 796.8976 | 299 | 750.0087 | 320 | 658.5221 | 409 | 730.4474 | 562 | AR | MDA-MB-4 | Mammary | GSM19090 | 28118 | 57278 |
| 485.765 | 117 | 587.4301 | 143 | 505.8963 | 170 | 833.2435 | 234 | AR | MDA-MB-4 | Mammary | GSM19090 | 6841 | 57279 |
| 744.7021 | 206 | 573.2613 | 200 | 703.5187 | 289 | 785.7323 | 385 | AR | MDA-MB-4 | Mammary | GSM19090 | 14726 | 57280 |
| 208.3084 | 173 | 285.9243 | 223 | 133.9439 | 235 | 180.3888 | 283 | CTCF | None | Embryonic | GSM19109 | 33119 | 57283 |
| 307.1876 | 193 | 464.2899 | 260 | 185.8666 | 248 | 246.0871 | 298 | CTCF | HEK293 | Embryonic | GSM19109 | 30337 | 57284 |
| 1.903332 | 5 | 0.45998 | 4 | -0.272189 | 2 | 7.183 | 16 | EP300 | PC-3 | Prostate | GSM19177 | 2474 | 57301 |
| 0.292461 | 1 | -0.019165 | 0 | -0.508651 | 0 | 42.9865 | 20 | EP300 | PC-3 | Prostate | GSM19177 | 1032 | 57302 |
| 0.689029 | 10 | 0.089222 | 9 | 0 | 13 | 0.330068 | 20 | EP300 | PC-3 | Prostate | GSM19177 | 7373 | 57303 |
| 0.264792 | 1 | -0.031038 | 0 | 0.088469 | 2 | 0.87605 | 4 | EP300 | PC-3 | Prostate | GSM19177 | 1077 | 57304 |
| 0.021267 | 21 | -0.042748 | 21 | -0.14052 | 35 | 0 | 48 | EP300 | PC-3 | Prostate | GSM19177 | 23366 | 57305 |
| 459.1306 | 105 | 698.8071 | 146 | 265.1805 | 113 | 275.7986 | 123 | POU5F1 | H9 | Embryo | GSM54520 | 5781 | 5753 |
| 160.3643 | 60 | 190.6846 | 73 | 115.8809 | 77 | 110.3135 | 81 | POU5F1 | H9 | Embryo | GSM54520 | 5816 | 5754 |
| 249.4925 | 82 | 352.6985 | 109 | 75.89801 | 71 | 152.5178 | 103 | POU5F1 | H9 | Embryo | GSM54520 | 6926 | 5755 |
| 894.6226 | 275 | 902.561 | 345 | 585.013 | 297 | 524.6645 | 304 | IKZF1 | Nalm6 | Blood | GSM10809 | 18388 | 57640 |
| 438.6294 | 276 | 568.2597 | 347 | 307.6557 | 376 | 285.6475 | 399 | HDAC1 | Nalm6 | Blood | GSM10809 | 45313 | 57641 |
| 23.2009 | 46 | 28.19815 | 57 | 5.003635 | 54 | 5.844939 | 62 | BAHD1 | HEK293 | Embryonic | GSM12902 | 15310 | 57659 |
| 9.695932 | 7 | 9.706183 | 8 | 5.552279 | 9 | 13.17856 | 14 | BAHD1 | HEK293 | Embryonic | GSM12902 | 1364 | 57661 |
| 7.642435 | 6 | 5.689814 | 6 | 6.233209 | 9 | 12.24084 | 13 | BAHD1 | HEK293 | Embryonic | GSM12902 | 1257 | 57662 |
| 71.28939 | 17 | 224.9705 | 37 | 63.86944 | 23 | 137.2137 | 37 | BMI1 | K562 | Bone Marr | GSM13193 | 1021 | 57669 |
| 733.1347 | 109 | 1119.361 | 153 | 460.8644 | 114 | 441.7181 | 118 | PCGF1 | K562 | Bone Marr | GSM13193 | 3135 | 57670 |
| 86.08173 | 29 | 106.016 | 36 | 17.41628 | 21 | 103.2823 | 50 | CBX2 | K562 | Bone Marr | GSM13193 | 2517 | 57671 |
| 55.46682 | 31 | 62.67288 | 37 | 27.2778 | 36 | 10.55984 | 28 | RNF2 | None | Embryo | GSM58522 | 4431 | 5768 |
| 10.12622 | 10 | 18.50199 | 15 | 5.078329 | 13 | 4.779731 | 14 | RNF2 | None | Embryo | GSM58523 | 2482 | 5769 |
| 12.75531 | 12 | 20.63572 | 17 | 9.826783 | 18 | 8.921825 | 19 | RNF2 | None | Embryo | GSM58523 | 2815 | 5770 |
| 10.1938 | 9 | 9.446214 | 10 | 7.133008 | 13 | 4.416062 | 12 | RNF2 | None | Embryo | GSM58523 | 2097 | 5771 |
| 1040.379 | 275 | 1033.764 | 331 | 785.4372 | 272 | 728.7006 | 279 | BRD2 | LX2 | None | GSM14168 | 11597 | 57804 |
| 982.6011 | 164 | 1072.278 | 209 | 443.0489 | 148 | 574.699 | 180 | BRD2 | LX2 | None | GSM14168 | 5908 | 57805 |
| 981.0213 | 252 | 986.9628 | 317 | 531.4281 | 233 | 624.1101 | 269 | BRD3 | LX2 | None | GSM14168 | 12386 | 57806 |
| 410.8428 | 85 | 620.05 | 118 | 265.5287 | 95 | 346.9905 | 116 | BRD3 | LX2 | None | GSM14168 | 4041 | 57807 |
| 876.0117 | 475 | 885.6862 | 588 | 648.5073 | 534 | 620.9208 | 571 | BRD4 | LX2 | None | GSM14168 | 39993 | 57808 |
| 909.1114 | 377 | 879.9632 | 430 | 670.2737 | 403 | 654.0319 | 438 | BRD4 | LX2 | None | GSM14168 | 26248 | 57809 |
| 1.184606 | 65 | 2.208329 | 85 | 1.360922 | 128 | 1.366495 | 144 | NR3C1 | None | None | GSM15161 | 47226 | 57915 |
| 925.3711 | 385 | 933.1472 | 477 | 698.6111 | 421 | 664.6135 | 438 | JMJD1C | Kasumi-1 | Blood | GSM15506 | 25300 | 57960 |
| 450.2553 | 58 | 702.4449 | 83 | 308.2554 | 62 | 473.913 | 84 | CHD1 | LNCaP | Prostate | GSM15736 | 1286 | 58018 |
| 649.0627 | 84 | 920.3853 | 113 | 369.1493 | 81 | 468.8654 | 98 | CHD1 | LNCaP | Prostate | GSM15736 | 1892 | 58019 |
| 881.7972 | 360 | 908.9741 | 474 | 673.3614 | 415 | 634.0474 | 425 | RUNX1 | Kasumi-1 | Blood | GSM15959 | 27418 | 58030 |
| 442.8066 | 192 | 773.0245 | 281 | 356.7412 | 263 | 366.6552 | 287 | RUNX1 | Kasumi-1 | Blood | GSM15959 | 21933 | 58031 |
| 404.4086 | 160 | 604.7909 | 217 | 273.7238 | 201 | 244.7794 | 207 | RUNX1T1 | Kasumi-1 | Blood | GSM15959 | 16384 | 58032 |
| 30.65721 | 72 | 40.06441 | 91 | 44.36843 | 137 | 88.18845 | 185 | NR3C1 | U2OS | Bone | GSM16075 | 24689 | 58054 |
| 31.89075 | 95 | 70.48775 | 141 | 38.48837 | 173 | 84.57576 | 236 | NR3C1 | U2OS | Bone | GSM16075 | 35988 | 58055 |
| 44.92298 | 32 | 46.94438 | 37 | 29.45482 | 43 | 84.96003 | 71 | NR3C1 | U2OS | Bone | GSM16075 | 5554 | 58056 |
| 33.70749 | 56 | 40.61671 | 69 | 30.03857 | 91 | 60.57418 | 124 | NR3C1 | U2OS | Bone | GSM16075 | 16452 | 58057 |
| 1113.86 | 423 | 1109.885 | 512 | 807.3609 | 374 | 773.4316 | 391 | KDM2B | None | None | GSM16120 | 15080 | 58062 |
| 1134.167 | 184 | 1133.798 | 227 | 631.1471 | 169 | 538.5971 | 165 | KDM2B | None | None | GSM16120 | 5266 | 58068 |
| 1191.439 | 396 | 1204.431 | 492 | 873.2574 | 340 | 826.6965 | 338 | KDM2B | None | None | GSM16120 | 10895 | 58074 |
| 1178.559 | 464 | 1194.704 | 580 | 855.3309 | 405 | 822.3456 | 422 | KDM2B | None | None | GSM16120 | 14197 | 58079 |
| 1159.309 | 370 | 1192.38 | 484 | 873.4099 | 342 | 818.837 | 333 | KDM2B | None | None | GSM16120 | 11092 | 58081 |
| 1111.656 | 521 | 1150.438 | 676 | 815.5611 | 495 | 768.8295 | 500 | KDM2B | None | Blood | GSM16120 | 20554 | 58083 |
| 1109.255 | 367 | 1111.35 | 447 | 819.9694 | 334 | 782.9865 | 338 | KDM2B | None | None | GSM16120 | 12597 | 58086 |
| 155.709 | 99 | 258.4257 | 140 | 113.6928 | 136 | 204.4963 | 185 | RUNX1 | U937 | Pleura | GSM16326 | 15305 | 58139 |
| 944.7179 | 411 | 965.8196 | 526 | 764.7106 | 511 | 749.3832 | 564 | HIF1A | HeLa | Cervix | GSM16349 | 25744 | 58172 |
| 30.04429 | 14 | 17.18401 | 12 | 22.84255 | 19 | 72.30657 | 35 | SOX2 | None | None | GSM16437 | 1781 | 58175 |
| 77.60928 | 36 | 50.62726 | 33 | 26.72824 | 35 | 43.49919 | 46 | SOX2 | None | None | GSM16437 | 4239 | 58176 |

| | | | | | | | | | | | | | |
|---|---|---|---|---|---|---|---|---|---|---|---|---|---|
| 70.94499 | 22 | 31.35858 | 16 | 35.20064 | 23 | 64.80791 | 33 | SOX2 | None | None | GSM16437 | 1764 | 58177 |
| 47.22651 | 24 | 27.49631 | 21 | 22.47273 | 27 | 45.18509 | 39 | SOX2 | None | None | GSM16437 | 3165 | 58178 |
| 54.46934 | 29 | 42.16971 | 29 | 37.13408 | 38 | 56.74525 | 49 | SOX2 | None | None | GSM16437 | 3942 | 58179 |
| 319.0006 | 173 | 335.0997 | 199 | 159.2676 | 203 | 142.5031 | 212 | BANF1 | None | None | GSM16457 | 24173 | 58186 |
| 101.7028 | 29 | 68.36776 | 26 | 43.82217 | 28 | 92.87481 | 43 | BANF1 | None | None | GSM16457 | 2098 | 58187 |
| 12.56395 | 11 | 18.56566 | 15 | 20.42827 | 22 | 119.9926 | 53 | BANF1 | None | None | GSM16457 | 2464 | 58188 |
| 29.5689 | 59 | 57.58683 | 86 | 41.87933 | 112 | 40.04224 | 121 | TP63 | None | None | GSM16457 | 19467 | 58189 |
| 26.51692 | 71 | 57.78767 | 106 | 69.53946 | 163 | 70.47888 | 179 | TP63 | None | None | GSM16457 | 26739 | 58190 |
| 54.63607 | 35 | 52.79496 | 39 | 59.8726 | 57 | 145.0905 | 90 | EP300 | None | None | GSM16457 | 5632 | 58193 |
| 68.15983 | 58 | 51.44037 | 59 | 74.35384 | 96 | 135.1322 | 131 | EP300 | None | None | GSM16457 | 11491 | 58194 |
| 0.893095 | 131 | 0.016266 | 133 | 0.238192 | 248 | 0.042384 | 265 | DUX4 | None | Muscle | GSM83761 | 119534 | 5822 |
| 386.3904 | 169 | 477.3233 | 209 | 256.3254 | 215 | 332.5529 | 258 | CTCF | BJAB | None | GSM78215 | 19931 | 5828 |
| 393.237 | 181 | 539.6416 | 235 | 272.3207 | 236 | 332.0567 | 276 | CTCF | BL-41 | Blood | GSM78215 | 22478 | 5830 |
| 134.7663 | 62 | 128.9844 | 68 | 80.41032 | 75 | 55.37442 | 70 | FOXO1 | None | Tonsil | GSM16689 | 7095 | 58300 |
| 112.3326 | 38 | 116.204 | 43 | 40.1542 | 35 | 86.58281 | 53 | FOXO1 | None | Tonsil | GSM16689 | 3259 | 58301 |
| 502.3185 | 177 | 481.5826 | 193 | 235.8488 | 187 | 134.3994 | 162 | BCL6 | None | Tonsil | GSM16689 | 15838 | 58302 |
| 468.6358 | 156 | 581.1322 | 193 | 333.31 | 195 | 195.1478 | 167 | BCL6 | None | Tonsil | GSM16689 | 13070 | 58303 |
| 1256.237 | 359 | 1305.077 | 479 | 986.9409 | 342 | 953.9782 | 352 | MED1 | LNCaP-abl | Prostate | GSM16790 | 7402 | 58315 |
| 1265.48 | 371 | 1298.281 | 488 | 990.8746 | 353 | 938.2328 | 345 | MED1 | LNCaP-abl | Prostate | GSM16790 | 7527 | 58316 |
| 1158.998 | 344 | 1066.105 | 330 | 963.1414 | 399 | 969.1254 | 463 | MED1 | LNCaP-abl | Prostate | GSM16790 | 9885 | 58319 |
| 1292.974 | 166 | 1267.955 | 193 | 1015.521 | 169 | 1046.947 | 182 | MED1 | None | Prostate | GSM16790 | 2720 | 58321 |
| 511.9143 | 59 | 567.3564 | 68 | 250.2902 | 51 | 276.4859 | 57 | MED1 | None | Liver | GSM16790 | 1084 | 58322 |
| 1254.734 | 170 | 1278.135 | 222 | 878.2368 | 164 | 781.313 | 162 | MED1 | LNCaP-abl | Prostate | GSM16791 | 3031 | 58333 |
| 956.768 | 96 | 966.9928 | 104 | 427.8711 | 77 | 365.1173 | 74 | MED1 | LNCaP-abl | Prostate | GSM16791 | 1354 | 58334 |
| 1268.898 | 546 | 1305.239 | 702 | 954.1533 | 489 | 916.5634 | 510 | SUPT5H | LNCaP-abl | Prostate | GSM16791 | 12154 | 58338 |
| 1187.895 | 153 | 1231.686 | 207 | 682.9217 | 146 | 719.7272 | 159 | MED1 | LNCaP-abl | Prostate | GSM16791 | 3147 | 58339 |
| 1176.244 | 293 | 1209.116 | 385 | 964.0513 | 314 | 907.9537 | 302 | MED1 | LNCaP-abl | Prostate | GSM16791 | 6757 | 58341 |
| 1210.454 | 261 | 1214.498 | 317 | 962.7613 | 251 | 910.7945 | 246 | MED1 | LNCaP-abl | Prostate | GSM16791 | 5345 | 58342 |
| 1225.761 | 159 | 1253.005 | 205 | 958.9677 | 173 | 769.222 | 161 | MED1 | LNCaP-abl | Prostate | GSM16791 | 3161 | 58343 |
| 981.6688 | 124 | 1221.908 | 166 | 733.7921 | 139 | 608.9725 | 132 | MED1 | LNCaP-abl | Prostate | GSM16791 | 2606 | 58347 |
| 898.9015 | 119 | 1213.672 | 169 | 646.6117 | 131 | 515.9389 | 122 | MED1 | LNCaP-abl | Prostate | GSM16791 | 2658 | 58348 |
| 620.4674 | 81 | 948.1254 | 114 | 406.0755 | 84 | 272.3943 | 71 | MED1 | LNCaP-abl | Prostate | GSM16791 | 1728 | 58349 |
| 353.1827 | 253 | 629.7751 | 361 | 288.2403 | 357 | 256.5461 | 374 | CTCF | PANC-1 | Pancreas | GSM16845 | 37472 | 58360 |
| 353.5612 | 278 | 621.1368 | 392 | 315.1251 | 408 | 265.8095 | 421 | CTCF | PANC-1 | Pancreas | GSM16845 | 44460 | 58361 |
| 354.4207 | 272 | 636.4623 | 389 | 299.7412 | 391 | 261.8412 | 408 | CTCF | PANC-1 | Pancreas | GSM16845 | 42487 | 58362 |
| 340.1278 | 247 | 611.8935 | 354 | 276.4601 | 350 | 260.8226 | 374 | CTCF | PANC-1 | Pancreas | GSM16845 | 37973 | 58363 |
| 437.412 | 53 | 520.7461 | 64 | 195.5544 | 44 | 368.0931 | 67 | ARID1A | HepG2 | Liver | GSM17040 | 1087 | 58462 |
| 330.2915 | 236 | 570.0033 | 334 | 228.0221 | 322 | 266.5018 | 370 | CTCF | None | Embryo | GSM17052 | 40528 | 58480 |
| 378.4572 | 253 | 621.9727 | 351 | 249.394 | 337 | 280.0834 | 382 | CTCF | None | Embryo | GSM17052 | 41389 | 58481 |
| 431.2396 | 74 | 397.8456 | 77 | 189.7468 | 65 | 239.0014 | 78 | MED1 | None | Embryo | GSM17052 | 2534 | 58483 |
| 95.23238 | 83 | 96.88444 | 95 | 39.92086 | 98 | 39.63112 | 107 | POU5F1 | None | Embryo | GSM17052 | 15815 | 58485 |
| 57.27576 | 74 | 73.3951 | 93 | 28.65542 | 98 | 24.20623 | 103 | POU5F1 | None | Embryo | GSM17052 | 18221 | 58486 |
| 328.3067 | 180 | 578.3643 | 261 | 192.5332 | 224 | 268.0846 | 275 | CTCF | None | Embryo | GSM17052 | 25089 | 58489 |
| 435.9946 | 213 | 624.1106 | 281 | 209.011 | 242 | 309.2695 | 304 | CTCF | None | Embryo | GSM17052 | 27018 | 58490 |
| 941.3017 | 117 | 747.5323 | 110 | 272.1628 | 77 | 454.2077 | 108 | MED1 | None | Embryo | GSM17052 | 2418 | 58492 |
| 1007.113 | 129 | 1094.137 | 147 | 400.2955 | 102 | 568.1722 | 131 | MED1 | None | Embryo | GSM17052 | 2853 | 58493 |
| 172.4038 | 106 | 184.2585 | 123 | 45.75211 | 101 | 100.4282 | 144 | POU5F1 | None | Embryo | GSM17052 | 15493 | 58494 |
| 133.7744 | 94 | 167.4267 | 117 | 48.59166 | 102 | 80.658 | 132 | POU5F1 | None | Embryo | GSM17052 | 15300 | 58495 |
| 462.2463 | 59 | 417.3186 | 60 | 202.6406 | 49 | 327.2471 | 68 | WDR5 | BE2-C | Brain | GSM17170 | 1327 | 58502 |
| 1267.862 | 673 | 1335.252 | 890 | 923.8105 | 609 | 887.8519 | 640 | WDR5 | BE2-C | Brain | GSM17170 | 18197 | 58506 |
| 1212.705 | 242 | 1267.398 | 342 | 980.4861 | 245 | 943.9745 | 254 | MED1 | LNCaP-abl | Prostate | GSM18473 | 5096 | 58636 |
| 1026.425 | 117 | 1255.436 | 144 | 686.1256 | 121 | 582.033 | 116 | MED1 | LNCaP-abl | Prostate | GSM18473 | 2022 | 58638 |
| 1240.026 | 152 | 1272.821 | 202 | 834.3505 | 153 | 750.825 | 152 | MED1 | LNCaP-abl | Prostate | GSM18473 | 2745 | 58639 |
| 1242.088 | 194 | 1283.753 | 263 | 1022.238 | 201 | 928.9346 | 193 | MED1 | LNCaP-abl | Prostate | GSM18473 | 3611 | 58640 |
| 732.6039 | 177 | 934.3829 | 236 | 347.9263 | 176 | 349.3797 | 189 | MYB | None | None | GSM18590 | 10559 | 58804 |
| 932.3106 | 427 | 974.0053 | 573 | 705.2435 | 476 | 668.5952 | 493 | MYB | None | None | GSM18590 | 28473 | 58805 |
| 438.9316 | 115 | 681.8866 | 161 | 249.0351 | 125 | 261.681 | 137 | MYB | None | None | GSM18590 | 7303 | 58806 |
| 1171.08 | 318 | 1205.278 | 413 | 900.4989 | 299 | 884.5338 | 326 | MAF | None | None | GSM18590 | 8641 | 58807 |
| 1115.528 | 505 | 1161.076 | 669 | 812.4732 | 468 | 807.5794 | 529 | MAF | None | None | GSM18590 | 19611 | 58808 |
| 1159.165 | 578 | 1189.841 | 733 | 845.4027 | 540 | 827.6802 | 585 | MAF | None | None | GSM18590 | 20229 | 58809 |
| 24.34093 | 15 | 18.54292 | 15 | 27.72026 | 25 | 25.29303 | 26 | GATA3 | None | None | GSM18590 | 2439 | 58811 |
| 35.65937 | 21 | 66.52027 | 32 | 72.70352 | 45 | 132.4174 | 64 | PPARG | THP-1 | Blood | GSM62414 | 3213 | 5882 |
| 25.17959 | 17 | 67.87691 | 31 | 1.751334 | 11 | 7.751716 | 19 | CD59 | Raji | Blood | GSM18685 | 3051 | 58829 |
| 7.244373 | 5 | 21.71861 | 10 | 16.61841 | 12 | 101.4151 | 32 | RXRA | THP-1 | Blood | GSM62414 | 1064 | 5883 |
| 722.2084 | 343 | 748.8443 | 449 | 513.6164 | 441 | 409.4294 | 440 | SPI1 | THP-1 | Blood | GSM62414 | 43946 | 5884 |
| 1226.585 | 599 | 1279.367 | 791 | 903.2104 | 557 | 883.5249 | 602 | XRN2 | HEK293 | Embryonic | GSM18722 | 17818 | 58840 |
| 1221.27 | 535 | 1270.298 | 713 | 906.3814 | 494 | 882.9992 | 529 | XRN2 | HEK293 | Embryonic | GSM18722 | 15067 | 58841 |
| 1242.326 | 499 | 1308.192 | 687 | 940.209 | 470 | 917.6296 | 507 | XRN2 | HEK293 | Embryonic | GSM18722 | 13055 | 58842 |
| 1205.88 | 375 | 1261.986 | 515 | 905.0993 | 335 | 918.869 | 390 | XRN2 | HEK293 | Embryonic | GSM18722 | 9541 | 58843 |
| 1251.829 | 549 | 1290.82 | 710 | 932.5826 | 500 | 913.6335 | 537 | XRN2 | HEK293 | Embryonic | GSM18722 | 14097 | 58844 |
| 1231.141 | 456 | 1263.268 | 592 | 911.5346 | 399 | 920.1897 | 468 | XRN2 | HEK293 | Embryonic | GSM18722 | 11504 | 58845 |
| 1180.315 | 160 | 1191.62 | 202 | 958.6977 | 186 | 1001.484 | 216 | XRN2 | HEK293 | Embryonic | GSM18723 | 3818 | 58846 |

| | | | | | | | | | | | | | |
|---|---|---|---|---|---|---|---|---|---|---|---|---|---|
| 95.15965 | 131 | 125.7982 | 166 | 111.8817 | 226 | 91.10128 | 232 | GATA2 | TF-1 | Bone Marr | GSM18885 | 33746 | 58871 |
| 88.44033 | 130 | 132.0785 | 172 | 134.0381 | 244 | 127.1057 | 262 | GATA2 | TF-1 | Bone Marr | GSM18885 | 34642 | 58872 |
| 429.1495 | 221 | 573.1186 | 282 | 255.6184 | 273 | 296.3913 | 313 | GATA2 | TF-1 | Bone Marr | GSM18885 | 28445 | 58873 |
| 374.0422 | 188 | 554.3421 | 252 | 240.6694 | 238 | 266.2492 | 268 | GATA2 | TF-1 | Bone Marr | GSM18885 | 23496 | 58874 |
| 335.2336 | 53 | 357.4739 | 60 | 199.0173 | 54 | 309.1751 | 73 | RUNX1 | TF-1 | Bone Marr | GSM18885 | 1663 | 58875 |
| 448.2094 | 66 | 649.516 | 90 | 305.9566 | 72 | 513.0383 | 102 | RUNX1 | TF-1 | Bone Marr | GSM18885 | 1861 | 58876 |
| 171.6492 | 59 | 216.8917 | 74 | 121.9241 | 74 | 127.2894 | 81 | RUNX1 | TF-1 | Bone Marr | GSM18885 | 5111 | 58877 |
| 181.7073 | 68 | 307.2356 | 99 | 146.1161 | 91 | 160.7248 | 102 | RAD21 | TF-1 | Bone Marr | GSM18885 | 6351 | 58878 |
| 327.1582 | 133 | 543.6098 | 190 | 196.7287 | 158 | 206.6238 | 174 | RAD21 | TF-1 | Bone Marr | GSM18885 | 13136 | 58879 |
| 345.9319 | 84 | 335.2177 | 91 | 272.9727 | 106 | 272.6985 | 113 | LIN9 | HeLa | Cervix | GSM66590 | 4871 | 5888 |
| 38.26792 | 62 | 64.18559 | 86 | 83.87757 | 134 | 68.05777 | 136 | RAD21 | TF-1 | Bone Marr | GSM18885 | 17595 | 58881 |
| 388.5617 | 54 | 460.2873 | 65 | 189.5626 | 48 | 247.6258 | 59 | KDM5C | caki-1 | Kidney | GSM18899 | 1322 | 58885 |
| 273.2648 | 43 | 378.6375 | 57 | 82.24152 | 30 | 119.9966 | 39 | KDM5C | caki-1 | Kidney | GSM18899 | 1239 | 58886 |
| 374.4482 | 62 | 497.4527 | 80 | 189.7652 | 58 | 197.7885 | 63 | KDM5C | caki-1 | Kidney | GSM18899 | 1961 | 58887 |
| 310.2292 | 50 | 334.2472 | 57 | 139.2712 | 44 | 188.6727 | 55 | MYBL2 | HeLa | Cervix | GSM66590 | 1640 | 5890 |
| 1175.026 | 559 | 1222.566 | 735 | 866.2091 | 529 | 836.3384 | 560 | NRF1 | None | None | GSM18916 | 19425 | 58936 |
| 1181.589 | 602 | 1242.902 | 806 | 873.1104 | 582 | 827.8306 | 596 | NRF1 | None | None | GSM18916 | 21598 | 58937 |
| 1223.024 | 663 | 1264.752 | 845 | 913.1352 | 651 | 866.0147 | 670 | NRF1 | HCC1954 | Mammary | GSM18916 | 22154 | 58939 |
| 1214.418 | 626 | 1238.865 | 792 | 898.0871 | 605 | 867.221 | 641 | NRF1 | HCC1954 | Mammary | GSM18916 | 20886 | 58940 |
| 13.8514 | 9 | 47.72683 | 19 | 10.72859 | 13 | 8.701336 | 13 | SMN1 | HEK293 | Embryonic | GSM18922 | 1657 | 58948 |
| 11.60017 | 7 | 8.866264 | 7 | 6.88869 | 9 | 53.94161 | 25 | SMN1 | HEK293 | Embryonic | GSM18922 | 1263 | 58949 |
| 53.09335 | 48 | 105.0745 | 73 | 47.55773 | 74 | 68.73605 | 92 | LYL1 | Kasumi-1 | Blood | GSM19015 | 9694 | 58999 |
| 804.1749 | 514 | 812.4872 | 635 | 564.9394 | 575 | 572.8558 | 668 | JMJD1C | HL-60 | Blood | GSM19015 | 57942 | 59001 |
| 8.032951 | 25 | 18.01387 | 38 | 12.19933 | 50 | 42.92246 | 81 | LYL1 | HL-60 | Blood | GSM19015 | 10449 | 59002 |
| 818.1661 | 458 | 848.3591 | 591 | 621.9607 | 555 | 584.7592 | 578 | JMJD1C | NB4 | Blood | GSM19015 | 46162 | 59004 |
| 29.28981 | 31 | 52.17858 | 45 | 14.40194 | 40 | 40.38677 | 62 | LYL1 | NB4 | Blood | GSM19015 | 7180 | 59005 |
| 202.2736 | 140 | 266.3314 | 178 | 135.5089 | 189 | 154.071 | 215 | JMJD1C | THP-1 | Blood | GSM19015 | 23712 | 59006 |
| 10.48225 | 15 | 45.62872 | 32 | 19.87813 | 32 | 10.49969 | 28 | LYL1 | THP-1 | Blood | GSM19015 | 4449 | 59007 |
| 1006.468 | 365 | 1043.437 | 486 | 761.0384 | 367 | 687.2457 | 340 | KMT2A | SEM | None | GSM19340 | 15791 | 59165 |
| 291.6585 | 55 | 524.7823 | 85 | 195.5857 | 61 | 323.962 | 85 | MLLT1 | SEM | None | GSM19340 | 2111 | 59168 |
| 1185.152 | 541 | 1223.368 | 698 | 891.4722 | 508 | 837.911 | 505 | SETD1A | SEM | None | GSM19341 | 14469 | 59170 |
| 182.4381 | 47 | 274.2358 | 65 | 176.2881 | 66 | 333.9678 | 98 | ETV4 | PC-3 | Prostate | GSM73881 | 2880 | 5924 |
| 86.43781 | 53 | 141.4792 | 75 | 95.57929 | 86 | 243.8919 | 139 | JUND | PC-3 | Prostate | GSM73881 | 8125 | 5925 |
| 729.0684 | 373 | 776.8989 | 507 | 451.2754 | 437 | 544.796 | 522 | EBF1 | MutuI | None | GSM19580 | 45933 | 59344 |
| 363.3044 | 144 | 413.5136 | 171 | 146.452 | 146 | 241.3848 | 193 | EBF1 | LCL | Blood | GSM19580 | 14507 | 59345 |
| 1087.582 | 356 | 1110.405 | 451 | 802.5673 | 325 | 773.7719 | 342 | RBPJ | MutuI | None | GSM19580 | 13155 | 59346 |
| 49.3639 | 58 | 89.72619 | 84 | 39.04088 | 88 | 63.6826 | 113 | RBPJ | LCL | Blood | GSM19580 | 14006 | 59347 |
| 718.9645 | 324 | 720.8813 | 397 | 666.4303 | 557 | 705.5678 | 697 | ESR1 | MCF-7 | Breast | GSM19675 | 39749 | 59373 |
| 646.393 | 278 | 708.9101 | 327 | 666.4921 | 475 | 710.1172 | 606 | ESR1 | MCF-7 | Breast | GSM19675 | 32633 | 59374 |
| 722.3027 | 328 | 703.2539 | 381 | 658.6087 | 546 | 709.8871 | 704 | ESR1 | MCF-7 | Breast | GSM19675 | 39994 | 59375 |
| 700.7153 | 159 | 738.5844 | 180 | 846.5237 | 254 | 862.3984 | 299 | ESR1 | T47D | Mammary | GSM19675 | 8458 | 59376 |
| 622.759 | 139 | 755.9926 | 170 | 787.4427 | 221 | 853.9058 | 253 | ESR1 | T47D | Mammary | GSM19675 | 7217 | 59377 |
| 769.7131 | 168 | 779.5578 | 186 | 814.2603 | 243 | 844.7196 | 290 | ESR1 | T47D | Mammary | GSM19675 | 8576 | 59378 |
| 11.47462 | 17 | 7.640685 | 17 | 21.47607 | 36 | 8.600693 | 29 | DUX4 | None | None | GSM19678 | 5098 | 59381 |
| 386.6193 | 129 | 542.5811 | 170 | 219.3569 | 145 | 303.7493 | 180 | MBD3 | MDA-MB-2 | Breast | GSM19744 | 10389 | 59412 |
| 475.4843 | 148 | 578.3688 | 181 | 316.8248 | 176 | 359.1644 | 200 | MBD3 | MDA-MB-2 | Breast | GSM19744 | 10663 | 59413 |
| 921.7051 | 506 | 958.0371 | 660 | 677.633 | 551 | 601.0618 | 515 | RUNX1 | Jurkat | Blood | GSM19759 | 38407 | 59416 |
| 339.9529 | 201 | 366.5099 | 234 | 229.3493 | 266 | 164.2778 | 257 | GATA3 | Jurkat | Blood | GSM19759 | 29794 | 59417 |
| 847.4265 | 202 | 951.367 | 276 | 374.8056 | 195 | 466.0266 | 231 | RNF2 | None | None | GSM12554 | 12204 | 59423 |
| 679.5901 | 85 | 764.0669 | 99 | 308.6349 | 72 | 375.6336 | 85 | RNF2 | None | None | GSM12554 | 1856 | 59424 |
| 398.4084 | 67 | 523.0548 | 86 | 191.8787 | 62 | 302.8195 | 84 | FOXP1 | None | None | GSM15445 | 2380 | 59437 |
| 161.9282 | 33 | 183.9916 | 39 | 41.56517 | 23 | 179.5455 | 52 | FOXP1 | None | None | GSM15445 | 1543 | 59439 |
| 428.3211 | 88 | 415.6901 | 95 | 183.1487 | 80 | 222.5857 | 94 | FOXP1 | OCI-Ly10 | Bone Marr | GSM15445 | 4196 | 59443 |
| 220.2301 | 82 | 352.5384 | 116 | 216.9855 | 121 | 236.7139 | 135 | FOXP1 | None | None | GSM15445 | 7921 | 59447 |
| 1048.146 | 294 | 1056.99 | 363 | 789.9674 | 288 | 780.0316 | 316 | MYC | IMEC | Breast | GSM17155 | 11978 | 59542 |
| 373.5808 | 58 | 401.8085 | 66 | 285.6172 | 68 | 156.7924 | 52 | MYC | IMEC | Breast | GSM17155 | 1799 | 59543 |
| 616.3357 | 83 | 903.719 | 114 | 353.0063 | 81 | 340.7852 | 84 | MYC | Raji | Blood | GSM76270 | 2023 | 5965 |
| 890.0362 | 128 | 1148.262 | 168 | 524.1118 | 129 | 484.8445 | 131 | MYC | CA46 | None | GSM76270 | 3582 | 5966 |
| 1145.078 | 184 | 1139.802 | 223 | 646.9381 | 169 | 735.6034 | 192 | MYC | None | Blood | GSM76270 | 5094 | 5967 |
| 1223.123 | 579 | 1244.167 | 724 | 912.6208 | 551 | 851.5615 | 539 | MYC | BL-41 | Blood | GSM76271 | 17703 | 5968 |
| 1176.977 | 292 | 1217.873 | 398 | 929.4488 | 294 | 881.7889 | 289 | MYC | Ramos | None | GSM76271 | 7787 | 5969 |
| 1421.092 | 178 | 1442.755 | 226 | 898.0472 | 137 | 1144.714 | 178 | FOS | GM12878 | Blood | GSM75433 | 1947 | 621 |
| 987.2263 | 426 | 1030.129 | 568 | 774.3718 | 482 | 726.3733 | 486 | YY1 | None | Liver | ENCSR994\ | 23009 | 62485 |
| 1021.62 | 530 | 1056.668 | 686 | 800.3686 | 613 | 749.8491 | 622 | YY1 | None | Liver | ENCSR994\ | 29094 | 62486 |
| 20.6518 | 13 | 15.82391 | 13 | 15.71837 | 18 | 20.94445 | 22 | ZNF202 | HEK293 | Embryonic | ENCSR996I | 1952 | 62487 |
| 11.94594 | 12 | 17.01393 | 16 | 26.78986 | 27 | 52.6367 | 39 | ZNF202 | HEK293 | Embryonic | ENCSR996I | 2682 | 62488 |
| 715.6146 | 332 | 727.7235 | 417 | 472.8792 | 420 | 485.4267 | 460 | SMC3 | IMR90 | Lung | ENCSR000I | 42101 | 62504 |
| 521.1554 | 234 | 734.7161 | 310 | 339.155 | 297 | 386.1336 | 338 | SMC3 | IMR90 | Lung | ENCSR000I | 28684 | 62505 |
| 655.1134 | 226 | 795.286 | 282 | 386.7797 | 260 | 436.4379 | 295 | ZEB2 | K562 | Bone Marr | ENCSR004( | 18689 | 62508 |
| 490.8006 | 175 | 575.1033 | 210 | 330.8256 | 213 | 378.1796 | 243 | ZEB2 | K562 | Bone Marr | ENCSR004( | 14544 | 62509 |
| 391.0872 | 102 | 544.3334 | 134 | 301.5429 | 125 | 203.6786 | 111 | ID3 | K562 | Bone Marr | ENCSR005I | 5333 | 62513 |
| 616.717 | 142 | 796.4928 | 179 | 370.4945 | 152 | 282.8542 | 143 | ID3 | K562 | Bone Marr | ENCSR005I | 6497 | 62514 |

| | | | | | | | | | | | | | |
|---|---|---|---|---|---|---|---|---|---|---|---|---|---|
| 724.0877 | 190 | 877.4054 | 233 | 501.1237 | 224 | 418.7307 | 221 | ZNF2 | HEK293 | Embryonic | ENCSR011( | 10956 | 62525 |
| 967.7651 | 241 | 946.3655 | 276 | 614.4956 | 243 | 580.0961 | 253 | ZNF2 | HEK293 | Embryonic | ENCSR011( | 10475 | 62526 |
| 894.5077 | 222 | 896.353 | 271 | 544.8295 | 249 | 459.9683 | 247 | ZNF175 | K562 | Bone Marr | ENCSR011I | 13577 | 62529 |
| 838.7296 | 218 | 870.1752 | 259 | 404.0847 | 221 | 332.7015 | 218 | ZNF175 | K562 | Bone Marr | ENCSR011I | 13977 | 62530 |
| 19.8714 | 24 | 10.8381 | 22 | 6.879649 | 28 | 22.88228 | 45 | ZNF585B | HEK293 | Embryonic | ENCSR011X | 5666 | 62531 |
| 2.889687 | 12 | 6.225825 | 18 | 6.357112 | 27 | 2.681822 | 24 | ZNF585B | HEK293 | Embryonic | ENCSR011X | 5547 | 62532 |
| 1230.272 | 505 | 1224.663 | 605 | 935.7505 | 480 | 901.951 | 499 | BCL3 | A549 | Lung | ENCSR013I | 12997 | 62533 |
| 1190.443 | 249 | 1213.174 | 323 | 963.6435 | 264 | 941.6898 | 283 | BCL3 | A549 | Lung | ENCSR013I | 6085 | 62534 |
| 1188.907 | 550 | 1202.647 | 679 | 891.7204 | 529 | 879.0469 | 579 | BCL3 | A549 | Lung | ENCSR013I | 17274 | 62535 |
| 158.3683 | 62 | 140.337 | 65 | 133.615 | 84 | 163.9757 | 99 | CUX1 | MCF-7 | Breast | ENCSR017( | 5525 | 62547 |
| 495.5656 | 135 | 457.6348 | 143 | 398.4836 | 173 | 398.1256 | 185 | CUX1 | MCF-7 | Breast | ENCSR017( | 8579 | 62548 |
| 1135.746 | 269 | 1148.456 | 341 | 863.1332 | 246 | 843.6341 | 263 | TFDP1 | K562 | Bone Marr | ENCSR017( | 7489 | 62550 |
| 1099.663 | 347 | 1114.988 | 436 | 822.3735 | 315 | 776.2895 | 313 | TFDP1 | K562 | Bone Marr | ENCSR017( | 10600 | 62551 |
| 903.6293 | 327 | 886.2614 | 376 | 656.833 | 334 | 685.8171 | 395 | ZNF600 | HEK293 | Embryonic | ENCSR017( | 18608 | 62554 |
| 878.3089 | 332 | 849.1276 | 372 | 669.2861 | 363 | 639.5385 | 379 | ZNF600 | HEK293 | Embryonic | ENCSR017( | 20981 | 62555 |
| 25.36928 | 10 | 13.23798 | 8 | 22.22481 | 14 | 2.521254 | 6 | PLRG1 | HepG2 | Liver | ENCSR019I | 1005 | 62557 |
| 15.54603 | 8 | 22.91932 | 11 | 2.843217 | 6 | 4.640202 | 8 | PLRG1 | HepG2 | Liver | ENCSR019I | 1079 | 62558 |
| 459.0756 | 113 | 470.5018 | 126 | 436.1849 | 155 | 408.5285 | 160 | BCL11A | HEK293 | Embryonic | ENCSR021I | 6170 | 62560 |
| 258.534 | 63 | 342.6668 | 81 | 277.1655 | 92 | 214.549 | 86 | BCL11A | HEK293 | Embryonic | ENCSR021I | 3386 | 62561 |
| 53.63393 | 49 | 59.14597 | 58 | 23.91143 | 59 | 39.33524 | 76 | ZNF623 | HEK293 | Embryonic | ENCSR022I | 9430 | 62565 |
| 60.21886 | 54 | 54.50663 | 59 | 30.97234 | 68 | 25.61655 | 70 | ZNF623 | HEK293 | Embryonic | ENCSR022I | 10293 | 62566 |
| 927.1585 | 441 | 937.7396 | 548 | 702.4635 | 487 | 659.7651 | 496 | CTCF | None | Spleen | ENCSR028Y | 28756 | 62569 |
| 941.1163 | 177 | 1003.975 | 212 | 461.8377 | 168 | 324.2312 | 151 | TARDBP | K562 | Bone Marr | ENCSR033V | 6753 | 62579 |
| 613.836 | 107 | 655.4343 | 121 | 230.7596 | 86 | 226.9875 | 91 | TARDBP | K562 | Bone Marr | ENCSR033V | 3354 | 62580 |
| 412.1932 | 109 | 518.2047 | 136 | 199.7713 | 110 | 230.6272 | 126 | CTCF | None | Skin | ENCSR038I | 6903 | 62586 |
| 19.12509 | 10 | 14.96587 | 10 | 8.80861 | 11 | 8.831565 | 12 | MCM7 | K562 | Bone Marr | ENCSR038I | 1353 | 62587 |
| 17.46894 | 9 | 8.117553 | 7 | 2.287924 | 6 | 3.797263 | 8 | MCM7 | K562 | Bone Marr | ENCSR038I | 1214 | 62588 |
| 1246.715 | 152 | 1253.876 | 188 | 563.1411 | 122 | 486.8707 | 119 | GABPA | MCF-7 | Breast | ENCSR038I | 2758 | 62589 |
| 526.6729 | 71 | 1045.627 | 118 | 416.1402 | 83 | 436.3946 | 90 | GABPA | MCF-7 | Breast | ENCSR038I | 1700 | 62590 |
| 647.5678 | 179 | 734.1563 | 211 | 328.7405 | 186 | 310.9914 | 195 | RFX1 | K562 | Bone Marr | ENCSR041/ | 11691 | 62593 |
| 584.6401 | 194 | 692.1009 | 234 | 283.0431 | 204 | 252.7131 | 210 | RFX1 | K562 | Bone Marr | ENCSR041/ | 15533 | 62594 |
| 155.4281 | 47 | 97.84735 | 41 | 67.51377 | 46 | 264.5051 | 95 | EP300 | A549 | Lung | ENCSR047I | 3484 | 62601 |
| 201.7361 | 94 | 169.0165 | 97 | 278.5342 | 163 | 496.7074 | 228 | EP300 | A549 | Lung | ENCSR047I | 10838 | 62602 |
| 786.3915 | 580 | 784.6251 | 702 | 609.3581 | 761 | 601.3909 | 854 | RAD21 | HepG2 | Liver | ENCSR054I | 75382 | 62607 |
| 786.5186 | 583 | 782.0208 | 703 | 607.3663 | 765 | 606.5609 | 873 | RAD21 | HepG2 | Liver | ENCSR054I | 75862 | 62608 |
| 984.0521 | 224 | 978.009 | 268 | 737.1521 | 249 | 557.7818 | 232 | KLF17 | HEK293 | Embryonic | ENCSR065V | 9255 | 62617 |
| 1001.262 | 250 | 997.2587 | 298 | 802.6721 | 274 | 783.2403 | 295 | KLF17 | HEK293 | Embryonic | ENCSR065V | 10000 | 62618 |
| 459.7926 | 118 | 558.1671 | 144 | 268.3406 | 127 | 227.4754 | 126 | CHAMP1 | K562 | Bone Marr | ENCSR065X | 6360 | 62619 |
| 452.465 | 122 | 381.8203 | 123 | 217.1771 | 121 | 185.2041 | 121 | CHAMP1 | K562 | Bone Marr | ENCSR065X | 7037 | 62620 |
| 782.652 | 245 | 825.3409 | 305 | 385.0022 | 258 | 428.5289 | 291 | RFX1 | MCF-7 | Breast | ENCSR066T | 18744 | 62623 |
| 1004.568 | 341 | 991.9969 | 401 | 762.2622 | 346 | 743.1371 | 375 | RFX1 | MCF-7 | Breast | ENCSR066T | 15424 | 62624 |
| 63.14523 | 37 | 57.4854 | 40 | 45.13333 | 50 | 160.9285 | 93 | FOSL2 | A549 | Lung | ENCSR067X | 5411 | 62626 |
| 42.31482 | 27 | 40.6152 | 30 | 30.17583 | 37 | 143.1986 | 78 | FOSL2 | A549 | Lung | ENCSR067X | 4350 | 62627 |
| 79.22637 | 48 | 57.34129 | 47 | 47.89941 | 61 | 209.0969 | 122 | FOSL2 | A549 | Lung | ENCSR067X | 7166 | 62628 |
| 477.3954 | 56 | 739.9095 | 80 | 381.4735 | 65 | 425.85 | 73 | STAT1 | None | None | ENCSR071I | 1089 | 62629 |
| 1130.247 | 375 | 1135.486 | 459 | 853.8688 | 345 | 815.1809 | 351 | ZNF76 | HEK293 | Embryonic | ENCSR072I | 10601 | 62632 |
| 1098.93 | 213 | 1115.22 | 267 | 625.4198 | 187 | 759.1786 | 220 | ZNF76 | HEK293 | Embryonic | ENCSR072I | 6121 | 62633 |
| 813.9333 | 269 | 831.2133 | 344 | 533.1552 | 316 | 520.0613 | 337 | HDAC2 | K562 | Bone Marr | ENCSR075I | 20436 | 62634 |
| 613.5541 | 170 | 616.3814 | 188 | 390.2376 | 194 | 361.3609 | 201 | HDAC2 | K562 | Bone Marr | ENCSR075I | 10432 | 62635 |
| 1070.565 | 306 | 1112.811 | 411 | 844.1657 | 321 | 817.2533 | 335 | KLF9 | HEK293 | Embryonic | ENCSR076I | 10414 | 62636 |
| 1088.087 | 398 | 1128.431 | 527 | 839.4946 | 400 | 802.1387 | 410 | KLF9 | HEK293 | Embryonic | ENCSR076I | 13451 | 62637 |
| 1044.335 | 381 | 1049.47 | 472 | 793.8803 | 394 | 756.5663 | 403 | RNF2 | K562 | Bone Marr | ENCSR076Y | 16801 | 62638 |
| 1017.917 | 477 | 1036.602 | 601 | 772.0058 | 514 | 721.9608 | 512 | RNF2 | K562 | Bone Marr | ENCSR076Y | 25104 | 62639 |
| 82.34279 | 37 | 125.0858 | 51 | 57.92125 | 48 | 209.6846 | 93 | HES2 | A549 | Lung | ENCSR078I | 4279 | 62645 |
| 104.6848 | 48 | 101.51 | 53 | 75.44302 | 63 | 239.47 | 114 | HES2 | A549 | Lung | ENCSR078I | 5595 | 62646 |
| 609.1307 | 127 | 616.2099 | 140 | 485.581 | 156 | 380.8732 | 147 | ZNF639 | HEK293 | Embryonic | ENCSR080( | 5475 | 62649 |
| 563.2215 | 100 | 653.8935 | 119 | 470.3698 | 124 | 446.93 | 128 | ZNF639 | HEK293 | Embryonic | ENCSR080( | 3534 | 62650 |
| 179.6476 | 67 | 206.8881 | 80 | 92.16443 | 73 | 92.9527 | 79 | RAD51 | HepG2 | Liver | ENCSR081V | 5959 | 62652 |
| 216.3044 | 48 | 226.1245 | 54 | 113.3546 | 48 | 90.74614 | 46 | RAD51 | HepG2 | Liver | ENCSR081V | 2282 | 62653 |
| 856.8629 | 188 | 685.1297 | 183 | 364.6893 | 171 | 394.0806 | 190 | NBN | K562 | Bone Marr | ENCSR085( | 8627 | 62658 |
| 924.8848 | 282 | 886.8267 | 309 | 598.0271 | 283 | 531.0674 | 288 | NBN | K562 | Bone Marr | ENCSR085( | 14986 | 62659 |
| 289.5442 | 52 | 241.318 | 51 | 140.7169 | 48 | 114.5918 | 46 | KAT8 | K562 | Bone Marr | ENCSR086I | 1793 | 62660 |
| 210.3415 | 34 | 148.0581 | 30 | 122.688 | 34 | 71.26041 | 27 | KAT8 | K562 | Bone Marr | ENCSR086I | 1026 | 62661 |
| 423.9674 | 99 | 356.7344 | 99 | 186.7018 | 92 | 286.3819 | 121 | HLTF | K562 | Bone Marr | ENCSR090J | 4917 | 62665 |
| 204.9803 | 56 | 116.8192 | 46 | 84.94859 | 52 | 99.31125 | 60 | HLTF | K562 | Bone Marr | ENCSR090J | 3411 | 62666 |
| 1119.83 | 339 | 1133.638 | 427 | 897.6272 | 355 | 861.8023 | 366 | ATF1 | K562 | Bone Marr | ENCSR091( | 9694 | 62670 |
| 1045.95 | 438 | 1056.039 | 535 | 796.214 | 438 | 754.5682 | 444 | ATF1 | K562 | Bone Marr | ENCSR091( | 16590 | 62671 |
| 339.8497 | 100 | 397.7232 | 120 | 190.974 | 109 | 220.8389 | 125 | HES1 | K562 | Bone Marr | ENCSR091J | 6768 | 62672 |
| 27.4984 | 13 | 50.78403 | 20 | 65.10722 | 30 | 154.9744 | 50 | HES1 | K562 | Bone Marr | ENCSR091J | 1612 | 62673 |
| 394.1369 | 100 | 427.7449 | 115 | 191.8773 | 99 | 161.2427 | 98 | CREB3 | K562 | Bone Marr | ENCSR093I | 5515 | 62679 |
| 491.7464 | 124 | 594.3252 | 151 | 321.5361 | 141 | 222.2156 | 127 | CREB3 | K562 | Bone Marr | ENCSR093I | 6682 | 62680 |
| 54.08882 | 24 | 32.46647 | 21 | 30.68018 | 28 | 106.2629 | 53 | ZNF16 | HEK293 | Embryonic | ENCSR093I | 2593 | 62681 |

| | | | | | | | | | | | | | |
|---|---|---|---|---|---|---|---|---|---|---|---|---|---|
| 97.83442 | 25 | 82.02046 | 25 | 57.08197 | 27 | 35.38791 | 23 | ZNF16 | HEK293 | Embryonic | ENCSR093I | 1468 | 62682 |
| 71.71653 | 52 | 127.4226 | 76 | 47.21474 | 70 | 83.14572 | 94 | CEBPG | MCF-7 | Breast | ENCSR094Z | 8989 | 62690 |
| 79.95502 | 53 | 102.7848 | 67 | 55.64703 | 72 | 97.45402 | 97 | CEBPG | MCF-7 | Breast | ENCSR094Z | 8543 | 62691 |
| 165.1672 | 36 | 184.2145 | 42 | 92.83659 | 37 | 56.72519 | 31 | ZNF207 | MCF-7 | Breast | ENCSR096I | 1637 | 62697 |
| 268.407 | 182 | 235.8645 | 195 | 167.5431 | 236 | 128.431 | 235 | PRDM1 | HEK293 | Embryonic | ENCSR098Y | 28320 | 62701 |
| 331.8034 | 209 | 343.0767 | 239 | 200.7146 | 266 | 159.9566 | 268 | PRDM1 | HEK293 | Embryonic | ENCSR098Y | 31103 | 62702 |
| 205.2285 | 98 | 236.7582 | 117 | 133.4494 | 121 | 144.1901 | 135 | ZNF24 | K562 | Bone Marr | ENCSR099I | 10208 | 62704 |
| 367.4808 | 142 | 438.5046 | 172 | 272.0834 | 181 | 184.5656 | 166 | ZNF24 | K562 | Bone Marr | ENCSR099I | 12409 | 62705 |
| 842.3792 | 471 | 862.4649 | 599 | 632.5975 | 543 | 601.8578 | 572 | CTCF | None | Colon | ENCSR102( | 39203 | 62707 |
| 100.6905 | 124 | 120.8601 | 152 | 70.31416 | 182 | 78.57851 | 206 | ZMYM3 | K562 | Bone Marr | ENCSR102I | 31278 | 62708 |
| 123.1494 | 170 | 155.7984 | 212 | 98.18916 | 262 | 87.31157 | 278 | ZMYM3 | K562 | Bone Marr | ENCSR102I | 45822 | 62709 |
| 14.90651 | 10 | 5.034364 | 7 | 11.35888 | 14 | 90.77146 | 39 | ZNF175 | HEK293 | Embryonic | ENCSR103( | 1620 | 62713 |
| 15.83646 | 9 | 12.30066 | 9 | 21.47919 | 16 | 18.08019 | 16 | ZNF175 | HEK293 | Embryonic | ENCSR103( | 1291 | 62714 |
| 996.2429 | 615 | 1003.619 | 754 | 755.5062 | 705 | 728.1509 | 754 | TFAP4 | HepG2 | Liver | ENCSR103S | 40916 | 62715 |
| 535.6199 | 146 | 644.853 | 178 | 429.8482 | 189 | 451.8763 | 207 | TFAP4 | HepG2 | Liver | ENCSR103S | 10028 | 62716 |
| 53.51169 | 27 | 26.52977 | 22 | 22.65883 | 29 | 143.328 | 70 | FOSL2 | A549 | Lung | ENCSR103U | 3535 | 62717 |
| 238.3642 | 134 | 156.364 | 126 | 172.9046 | 181 | 358.3125 | 262 | FOSL2 | A549 | Lung | ENCSR103U | 18847 | 62718 |
| 210.4591 | 85 | 274.9989 | 108 | 195.2231 | 121 | 234.0207 | 141 | ZNF366 | HEK293 | Embryonic | ENCSR106E | 8039 | 62731 |
| 333.6944 | 200 | 355.7991 | 231 | 225.8623 | 260 | 268.3844 | 301 | ZNF366 | HEK293 | Embryonic | ENCSR106E | 26421 | 62732 |
| 95.02955 | 85 | 84.0209 | 92 | 117.3506 | 145 | 106.524 | 152 | TAL1 | K562 | Bone Marr | ENCSR106I | 15908 | 62733 |
| 109.9787 | 116 | 147.3924 | 148 | 126.853 | 195 | 184.2631 | 242 | TAL1 | K562 | Bone Marr | ENCSR106I | 24705 | 62734 |
| 452.021 | 139 | 701.1365 | 193 | 300.5755 | 165 | 346.826 | 189 | MLLT1 | K562 | Bone Marr | ENCSR107( | 9941 | 62738 |
| 522.4265 | 140 | 585.791 | 164 | 258.1708 | 142 | 299.2433 | 163 | MLLT1 | K562 | Bone Marr | ENCSR107( | 8540 | 62739 |
| 31.65697 | 13 | 21.64303 | 12 | 40.47687 | 22 | 70.51087 | 31 | YBX1 | K562 | Bone Marr | ENCSR107I | 1389 | 62743 |
| 62.82168 | 48 | 64.26078 | 55 | 62.89162 | 76 | 238.478 | 143 | FOSL2 | A549 | Lung | ENCSR111Z | 8600 | 62753 |
| 127.63 | 82 | 132.0711 | 94 | 104.9796 | 118 | 299.9695 | 196 | FOSL2 | A549 | Lung | ENCSR111Z | 12736 | 62754 |
| 140.05 | 83 | 134.8254 | 92 | 100.0196 | 112 | 228.0798 | 169 | FOSL2 | A549 | Lung | ENCSR111Z | 12042 | 62755 |
| 212.7061 | 104 | 195.035 | 112 | 174.4995 | 144 | 195.9095 | 163 | MTA2 | K562 | Bone Marr | ENCSR113L | 12264 | 62756 |
| 121.9354 | 78 | 152.1994 | 97 | 136.87 | 126 | 124.0127 | 131 | MTA2 | K562 | Bone Marr | ENCSR113L | 11582 | 62757 |
| 815.864 | 397 | 852.8037 | 527 | 606.5749 | 450 | 596.9767 | 502 | PKNOX1 | K562 | Bone Marr | ENCSR115S | 35163 | 62761 |
| 809.9259 | 436 | 829.805 | 552 | 601.191 | 500 | 583.0557 | 542 | PKNOX1 | K562 | Bone Marr | ENCSR115S | 40410 | 62762 |
| 171.2941 | 75 | 185.6373 | 87 | 99.3948 | 88 | 86.72786 | 90 | ZNF24 | K562 | Bone Marr | ENCSR117V | 7619 | 62763 |
| 140.2265 | 65 | 175.8795 | 81 | 78.82381 | 76 | 88.1991 | 86 | ZNF24 | K562 | Bone Marr | ENCSR117V | 7025 | 62764 |
| 733.3606 | 115 | 1007.964 | 151 | 335.4444 | 101 | 610.6014 | 148 | ZNF71 | HEK293 | Embryonic | ENCSR121E | 3263 | 62768 |
| 376.5106 | 97 | 398.135 | 110 | 191.0915 | 98 | 106.9502 | 81 | ZNF71 | HEK293 | Embryonic | ENCSR121E | 5382 | 62769 |
| 252.0143 | 55 | 200.355 | 53 | 73.95936 | 41 | 88.95287 | 48 | CDC5L | K562 | Bone Marr | ENCSR121I | 2407 | 62771 |
| 268.4079 | 62 | 261.1788 | 67 | 97.44174 | 52 | 70.24393 | 48 | CDC5L | K562 | Bone Marr | ENCSR121I | 2960 | 62772 |
| 125.4388 | 54 | 132.5607 | 62 | 133.0573 | 83 | 322.9632 | 135 | EP300 | A549 | Lung | ENCSR124V | 5759 | 62786 |
| 147.9361 | 53 | 93.47758 | 47 | 106.0239 | 67 | 306.5656 | 119 | EP300 | A549 | Lung | ENCSR124V | 4752 | 62787 |
| 115.7678 | 41 | 76.21738 | 37 | 83.97034 | 52 | 281.0224 | 100 | EP300 | A549 | Lung | ENCSR124V | 3629 | 62788 |
| 887.9789 | 280 | 867.9323 | 322 | 584.0348 | 296 | 502.9953 | 298 | ZNF394 | HEK293 | Embryonic | ENCSR125I | 16256 | 62792 |
| 268.7672 | 118 | 288.0654 | 136 | 141.7832 | 133 | 70.93727 | 111 | ZNF394 | HEK293 | Embryonic | ENCSR125I | 11752 | 62793 |
| 930.8227 | 146 | 1137.724 | 209 | 429.1177 | 131 | 640.7603 | 172 | KLF9 | MCF-7 | Breast | ENCSR125Z | 4648 | 62799 |
| 1117.267 | 378 | 1156.836 | 503 | 846.634 | 366 | 864.1499 | 433 | KLF9 | MCF-7 | Breast | ENCSR125Z | 12727 | 62800 |
| 34.76054 | 15 | 20.7133 | 13 | 13.90062 | 15 | 17.65048 | 18 | PTRF | K562 | Bone Marr | ENCSR126I | 1633 | 62801 |
| 80.30771 | 24 | 94.43858 | 29 | 52.35988 | 28 | 16.65237 | 18 | PTRF | K562 | Bone Marr | ENCSR126I | 1684 | 62802 |
| 498.3989 | 133 | 819.9746 | 191 | 267.2121 | 138 | 217.7831 | 135 | NR4A1 | K562 | Bone Marr | ENCSR130I | 7317 | 62814 |
| 613.4306 | 155 | 673.0231 | 179 | 266.1815 | 145 | 199.3793 | 137 | NR4A1 | K562 | Bone Marr | ENCSR130I | 8128 | 62815 |
| 1110.751 | 156 | 1189.672 | 220 | 452.9385 | 127 | 519.1042 | 145 | NRF1 | MCF-7 | Breast | ENCSR135/ | 3831 | 62819 |
| 634.0962 | 91 | 1020.33 | 132 | 298.0312 | 80 | 525.98 | 116 | NRF1 | MCF-7 | Breast | ENCSR135/ | 2337 | 62820 |
| 1163.102 | 525 | 1187.703 | 667 | 879.2841 | 505 | 839.4597 | 522 | SP3 | HEK293 | Embryonic | ENCSR141I | 15665 | 62840 |
| 1170.003 | 490 | 1185.898 | 612 | 879.0345 | 457 | 833.489 | 462 | SP3 | HEK293 | Embryonic | ENCSR141I | 13831 | 62841 |
| 554.0578 | 119 | 783.3902 | 158 | 328.8913 | 125 | 306.8371 | 129 | HDGF | GM12878 | Blood | ENCSR145X | 4916 | 62848 |
| 758.8493 | 137 | 1057.929 | 184 | 381.4252 | 129 | 362.4975 | 134 | HDGF | GM12878 | Blood | ENCSR145X | 4541 | 62849 |
| 142.6079 | 55 | 141.1129 | 61 | 125.5801 | 77 | 347.819 | 134 | EP300 | A549 | Lung | ENCSR145Y | 5310 | 62850 |
| 107.1285 | 35 | 77.94243 | 33 | 81.56043 | 45 | 279.9782 | 89 | EP300 | A549 | Lung | ENCSR145Y | 2874 | 62851 |
| 906.708 | 526 | 950.644 | 693 | 690.7237 | 605 | 639.5326 | 608 | CTCF | None | gastroesop | ENCSR146I | 38524 | 62852 |
| 920.0064 | 239 | 918.7645 | 290 | 620.0195 | 267 | 588.0732 | 279 | ZNF584 | K562 | Bone Marr | ENCSR149Z | 13865 | 62861 |
| 669.425 | 152 | 854.3909 | 191 | 479.5202 | 181 | 404.946 | 178 | ZNF584 | K562 | Bone Marr | ENCSR149Z | 8035 | 62862 |
| 316.371 | 73 | 209.3909 | 64 | 113.5535 | 61 | 83.93747 | 57 | ARNT | K562 | Bone Marr | ENCSR155I | 3481 | 62868 |
| 475.089 | 111 | 366.8333 | 106 | 131.5425 | 83 | 241.6643 | 118 | ARNT | K562 | Bone Marr | ENCSR155I | 5376 | 62869 |
| 924.083 | 123 | 1191.001 | 164 | 437.7661 | 108 | 354.0113 | 102 | ZBTB11 | MCF-7 | Breast | ENCSR155V | 2929 | 62870 |
| 501.6125 | 68 | 806.7572 | 99 | 293.6296 | 67 | 258.4877 | 66 | ZBTB11 | MCF-7 | Breast | ENCSR155V | 1640 | 62871 |
| 1027.91 | 208 | 996.0731 | 232 | 585.4608 | 202 | 514.3634 | 202 | DNMT3B | HepG2 | Liver | ENCSR156( | 8362 | 62874 |
| 897.4991 | 174 | 1014.471 | 228 | 492.359 | 178 | 482.8208 | 188 | DNMT3B | HepG2 | Liver | ENCSR156( | 7677 | 62875 |
| 326.8884 | 194 | 362.8205 | 228 | 259.6817 | 268 | 267.2206 | 294 | SMARCE1 | K562 | Bone Marr | ENCSR157T | 25770 | 62879 |
| 201.0833 | 123 | 253.3386 | 153 | 200.1342 | 186 | 169.789 | 189 | SMARCE1 | K562 | Bone Marr | ENCSR157T | 16455 | 62880 |
| 933.6573 | 242 | 932.0758 | 294 | 612.7417 | 258 | 455.5487 | 241 | ZNF518A | HEK293 | Embryonic | ENCSR159( | 12408 | 62888 |
| 800.4609 | 227 | 845.3372 | 274 | 409.9249 | 239 | 370.066 | 246 | ZNF518A | HEK293 | Embryonic | ENCSR159( | 15573 | 62889 |
| 1257.643 | 223 | 1258.047 | 267 | 1029.585 | 228 | 948.0575 | 206 | ATF1 | K562 | Bone Marr | ENCSR159( | 4226 | 62891 |
| 1271.937 | 246 | 1277.992 | 311 | 1024.7 | 246 | 955.0509 | 228 | ATF1 | K562 | Bone Marr | ENCSR159( | 4684 | 62892 |
| 714.4305 | 138 | 715.8225 | 151 | 520.201 | 160 | 394.5018 | 148 | ZNF346 | HEK293 | Embryonic | ENCSR162I | 5297 | 62899 |

| | | | | | | | | | | | | | |
|---|---|---|---|---|---|---|---|---|---|---|---|---|---|
| 668.6736 | 131 | 650.3743 | 141 | 337.1543 | 126 | 319.8357 | 131 | ZNF346 | HEK293 | Embryonic | ENCSR162I | 5094 | 62900 |
| 954.7947 | 150 | 993.1023 | 167 | 458.4663 | 136 | 442.8038 | 142 | DIDO1 | K562 | Bone Marr | ENCSR167J | 4384 | 62904 |
| 1064.632 | 186 | 1061.55 | 226 | 693.3042 | 195 | 509.5847 | 177 | DIDO1 | K562 | Bone Marr | ENCSR167J | 6014 | 62905 |
| 767.7797 | 324 | 772.2737 | 400 | 627.4549 | 439 | 608.6943 | 478 | NR2F2 | None | Liver | ENCSR168S | 33333 | 62908 |
| 803.7414 | 384 | 837.0341 | 504 | 663.3958 | 524 | 628.9815 | 549 | NR2F2 | None | Liver | ENCSR168S | 37016 | 62909 |
| 525.587 | 98 | 687.0017 | 125 | 334.825 | 106 | 342.0757 | 114 | FOXK2 | HepG2 | Liver | ENCSR171F | 3749 | 62912 |
| 762.9274 | 141 | 888.1715 | 168 | 429.3032 | 143 | 444.393 | 155 | FOXK2 | HepG2 | Liver | ENCSR171F | 5315 | 62913 |
| 807.7001 | 282 | 807.2692 | 346 | 502.2709 | 324 | 423.9267 | 326 | CTCF | None | Stomach | ENCSR173A | 23674 | 62917 |
| 529.4575 | 115 | 664.1136 | 143 | 297.7812 | 119 | 269.1549 | 121 | ZSCAN29 | K562 | Bone Marr | ENCSR175S | 5216 | 62919 |
| 864.201 | 214 | 887.8182 | 251 | 380.8166 | 204 | 487.2728 | 245 | ZSCAN29 | K562 | Bone Marr | ENCSR175S | 11912 | 62920 |
| 368.1625 | 87 | 420.2867 | 103 | 174.635 | 85 | 162.7367 | 88 | CUX1 | K562 | Bone Marr | ENCSR178I | 4801 | 62923 |
| 314.1578 | 69 | 442.4017 | 92 | 147.4853 | 66 | 136.7095 | 68 | CUX1 | K562 | Bone Marr | ENCSR178I | 3450 | 62924 |
| 120.7666 | 38 | 166.2637 | 50 | 70.73249 | 43 | 178.7357 | 72 | CEBPB | A549 | Lung | ENCSR182C | 2999 | 62928 |
| 139.2017 | 60 | 130.237 | 65 | 82.79082 | 72 | 247.9503 | 126 | CEBPB | A549 | Lung | ENCSR182C | 6578 | 62929 |
| 973.2778 | 392 | 1011.178 | 524 | 762.4368 | 439 | 732.6684 | 465 | ZNF341 | HEK293 | Embryonic | ENCSR185F | 19929 | 62930 |
| 992.5134 | 423 | 1038.798 | 568 | 775.0085 | 464 | 729.5351 | 469 | ZNF341 | HEK293 | Embryonic | ENCSR185F | 20447 | 62931 |
| 1192.661 | 633 | 1206.208 | 783 | 879.0517 | 615 | 851.1441 | 657 | BCL3 | A549 | Lung | ENCSR186I | 21801 | 62932 |
| 1184.836 | 482 | 1200.157 | 601 | 895.2753 | 460 | 893.8795 | 518 | BCL3 | A549 | Lung | ENCSR186I | 14162 | 62933 |
| 1167.425 | 574 | 1165.988 | 696 | 866.6028 | 563 | 853.0328 | 621 | BCL3 | A549 | Lung | ENCSR186I | 20644 | 62934 |
| 496.622 | 70 | 679.3084 | 92 | 232.3622 | 61 | 221.4849 | 63 | ZBTB40 | GM12878 | Blood | ENCSR189Y | 1735 | 62936 |
| 905.1445 | 441 | 896.5112 | 523 | 642.0297 | 427 | 609.0234 | 445 | ZBTB40 | GM12878 | Blood | ENCSR189Y | 26365 | 62937 |
| 151.1433 | 83 | 237.184 | 115 | 139.2458 | 123 | 358.3168 | 199 | CEBPB | A549 | Lung | ENCSR190V | 11332 | 62938 |
| 199.7657 | 106 | 210.5141 | 122 | 122.7919 | 133 | 258.238 | 194 | CEBPB | A549 | Lung | ENCSR190V | 14003 | 62939 |
| 403.2418 | 185 | 440.9723 | 216 | 240.8332 | 226 | 392.4038 | 297 | CEBPB | A549 | Lung | ENCSR190V | 22090 | 62940 |
| 871.4394 | 399 | 884.368 | 499 | 669.3383 | 466 | 620.2085 | 464 | ZNF766 | K562 | Bone Marr | ENCSR194I | 30624 | 62946 |
| 882.8974 | 406 | 871.698 | 485 | 658.4905 | 449 | 613.577 | 455 | ZNF766 | K562 | Bone Marr | ENCSR194I | 30584 | 62947 |
| 223.484 | 46 | 236.7745 | 52 | 143.3834 | 50 | 147.2212 | 54 | HDGF | K562 | Bone Marr | ENCSR197A | 1867 | 62954 |
| 184.3822 | 45 | 216.0358 | 54 | 121.838 | 51 | 199.8609 | 70 | HDGF | K562 | Bone Marr | ENCSR197A | 2318 | 62955 |
| 460.2204 | 82 | 748.0551 | 119 | 322.1099 | 92 | 214.1164 | 79 | SREBF1 | MCF-7 | Breast | ENCSR197D | 2800 | 62956 |
| 397.1114 | 79 | 530.1276 | 102 | 231.241 | 82 | 189.2405 | 79 | SREBF1 | MCF-7 | Breast | ENCSR197D | 3220 | 62957 |
| 566.6604 | 226 | 647.7672 | 269 | 301.6771 | 260 | 320.8199 | 288 | NFE2L2 | IMR90 | Lung | ENCSR197V | 24554 | 62958 |
| 137.1566 | 86 | 127.0855 | 94 | 94.75085 | 116 | 95.18225 | 126 | NFE2L2 | IMR90 | Lung | ENCSR197V | 13449 | 62959 |
| 747.133 | 304 | 799.0887 | 423 | 559.8521 | 397 | 476.9463 | 403 | EED | GM12878 | Blood | ENCSR199V | 32758 | 62964 |
| 787.1448 | 361 | 808.1601 | 464 | 595.689 | 432 | 487.7587 | 428 | EED | GM12878 | Blood | ENCSR199V | 35806 | 62965 |
| 929.1959 | 316 | 908.3237 | 364 | 725.4825 | 359 | 702.5176 | 383 | ATF3 | None | Liver | ENCSR205F | 19193 | 62975 |
| 803.2699 | 200 | 884.5221 | 237 | 510.1257 | 229 | 554.7606 | 255 | ATF3 | None | Liver | ENCSR205F | 12356 | 62976 |
| 859.994 | 472 | 883.4181 | 600 | 646.7857 | 538 | 615.5354 | 569 | CTCF | None | gastroesop | ENCSR206E | 37944 | 62979 |
| 696.5827 | 205 | 770.5224 | 239 | 390.8711 | 227 | 320.2964 | 224 | ZBED1 | GM12878 | Blood | ENCSR207F | 15136 | 62980 |
| 302.3737 | 67 | 456.7013 | 93 | 141.0769 | 64 | 130.9437 | 66 | ZBED1 | GM12878 | Blood | ENCSR207F | 3394 | 62981 |
| 750.9895 | 152 | 856.6608 | 179 | 479.7581 | 166 | 512.6634 | 183 | ZNF391 | HEK293 | Embryonic | ENCSR210I | 6255 | 62985 |
| 244.3553 | 50 | 355.3455 | 68 | 174.7248 | 58 | 187.3571 | 64 | ZNF391 | HEK293 | Embryonic | ENCSR210I | 2148 | 62986 |
| 209.0888 | 51 | 265.3649 | 64 | 129.0436 | 56 | 104.2952 | 54 | GLI4 | HEK293 | Embryonic | ENCSR211F | 2663 | 62990 |
| 313.0004 | 67 | 372.5291 | 81 | 151.9044 | 64 | 154.7463 | 69 | GLI4 | HEK293 | Embryonic | ENCSR211F | 2965 | 62991 |
| 57.22972 | 16 | 59.0847 | 18 | 27.97655 | 16 | 50.52457 | 23 | TRIM25 | K562 | Bone Marr | ENCSR213V | 1047 | 62994 |
| 143.9675 | 33 | 78.95504 | 26 | 62.38774 | 30 | 61.86669 | 32 | TRIM25 | K562 | Bone Marr | ENCSR213V | 1601 | 62995 |
| 514.5867 | 65 | 682.9504 | 84 | 297.6314 | 63 | 419.3078 | 81 | BCL3 | A549 | Lung | ENCSR215U | 1414 | 63000 |
| 1143.931 | 377 | 1162.49 | 481 | 887.5913 | 378 | 889.6899 | 430 | BCL3 | A549 | Lung | ENCSR215U | 11801 | 63001 |
| 1169.968 | 634 | 1185.268 | 787 | 881.7166 | 652 | 865.167 | 717 | BCL3 | A549 | Lung | ENCSR215U | 23898 | 63002 |
| 450.3809 | 209 | 510.2046 | 248 | 315.925 | 273 | 408.8291 | 327 | CEBPB | A549 | Lung | ENCSR216C | 25557 | 63003 |
| 262.6425 | 108 | 264.2892 | 121 | 120.8911 | 116 | 324.0908 | 190 | CEBPB | A549 | Lung | ENCSR216C | 11282 | 63004 |
| 91.12928 | 21 | 112.0772 | 26 | 47.45872 | 21 | 58.61984 | 25 | ZNF8 | HEK293 | Embryonic | ENCSR217E | 1080 | 63006 |
| 64.02947 | 32 | 102.1401 | 45 | 46.83788 | 42 | 48.34687 | 46 | ZNF8 | HEK293 | Embryonic | ENCSR217E | 3666 | 63007 |
| 583.4247 | 180 | 585.8973 | 200 | 417.4324 | 223 | 392.4814 | 233 | ATF2 | HEK293 | Embryonic | ENCSR217H | 13735 | 63008 |
| 585.0519 | 185 | 538.5103 | 197 | 460.661 | 240 | 407.274 | 244 | ATF2 | HEK293 | Embryonic | ENCSR217H | 14484 | 63009 |
| 191.9525 | 47 | 236.1243 | 58 | 128.9167 | 54 | 153.911 | 63 | ZNF697 | HEK293 | Embryonic | ENCSR217C | 2517 | 63010 |
| 0 | 6 | 0.045333 | 9 | 0.619449 | 20 | 6.408261 | 36 | ZNF697 | HEK293 | Embryonic | ENCSR217C | 7082 | 63011 |
| 259.8024 | 63 | 193.1187 | 59 | 249.8672 | 87 | 225.6319 | 88 | TSHZ1 | HEK293 | Embryonic | ENCSR217V | 3385 | 63014 |
| 524.9891 | 130 | 476.7582 | 136 | 434.8334 | 167 | 526.8939 | 196 | TSHZ1 | HEK293 | Embryonic | ENCSR217V | 7300 | 63015 |
| 916.8292 | 259 | 878.3385 | 290 | 668.5857 | 295 | 609.6404 | 303 | DEK | HeLa-S3 | Cervix | ENCSR219I | 16349 | 63018 |
| 548.5522 | 159 | 728.2669 | 204 | 342.8242 | 185 | 364.3513 | 204 | DEK | HeLa-S3 | Cervix | ENCSR219I | 11867 | 63019 |
| 1099.977 | 472 | 1125.538 | 607 | 852.4216 | 500 | 802.6615 | 500 | MBD2 | K562 | Bone Marr | ENCSR221C | 18522 | 63028 |
| 1168.439 | 374 | 1191.108 | 475 | 900.6155 | 362 | 868.2755 | 375 | MBD2 | K562 | Bone Marr | ENCSR221C | 10718 | 63029 |
| 873.7444 | 524 | 909.4116 | 678 | 682.487 | 636 | 659.8596 | 687 | PRDM10 | HEK293 | Embryonic | ENCSR223T | 42012 | 63034 |
| 896.948 | 525 | 920.8534 | 662 | 681.4936 | 604 | 676.6321 | 683 | PRDM10 | HEK293 | Embryonic | ENCSR223T | 39013 | 63035 |
| 149.9284 | 59 | 233.1552 | 82 | 137.6296 | 83 | 117.0369 | 83 | ZNF121 | HEK293 | Embryonic | ENCSR224C | 5242 | 63037 |
| 139.19 | 73 | 255.245 | 109 | 181.6438 | 123 | 132.6523 | 116 | ZNF121 | HEK293 | Embryonic | ENCSR224C | 8648 | 63038 |
| 47.00765 | 17 | 26.05258 | 14 | 32.55486 | 21 | 30.68529 | 22 | ZNF292 | HEK293 | Embryonic | ENCSR228F | 1505 | 63049 |
| 849.2976 | 717 | 894.0009 | 916 | 662.0043 | 941 | 640.8031 | 1034 | RAD21 | None | Liver | ENCSR230Z | 86408 | 63055 |
| 850.8368 | 741 | 886.2599 | 936 | 648.7285 | 954 | 626.4882 | 1049 | RAD21 | None | Liver | ENCSR230Z | 91053 | 63056 |
| 766.7569 | 225 | 856.8571 | 289 | 504.6766 | 267 | 367.5314 | 250 | CTCF | None | Spleen | ENCSR234H | 16365 | 63059 |
| 103.9582 | 23 | 102.0007 | 25 | 159.6803 | 41 | 425.4562 | 76 | DPF2 | MCF-7 | Breast | ENCSR234V | 1155 | 63060 |
| 483.4042 | 172 | 337.0596 | 161 | 608.2342 | 278 | 707.0578 | 325 | DPF2 | MCF-7 | Breast | ENCSR234V | 14667 | 63061 |

| | | | | | | | | | | | | | |
|---|---|---|---|---|---|---|---|---|---|---|---|---|---|
| 912.9261 | 395 | 894.6447 | 459 | 642.347 | 374 | 623.8852 | 406 | ZBTB40 | K562 | Bone Marr | ENCSR237\ | 23732 | 63068 |
| 947.319 | 390 | 926.4198 | 452 | 669.2903 | 361 | 639.88 | 377 | ZBTB40 | K562 | Bone Marr | ENCSR237\ | 20743 | 63069 |
| 702.905 | 334 | 700.2873 | 404 | 445.9542 | 417 | 465.3615 | 460 | FOSL1 | K562 | Bone Marr | ENCSR2392 | 40918 | 63076 |
| 614.633 | 289 | 664.0407 | 335 | 379.7151 | 357 | 421.9234 | 403 | FOSL1 | K562 | Bone Marr | ENCSR2392 | 34002 | 63077 |
| 93.71169 | 52 | 57.12319 | 47 | 58.62355 | 66 | 208.9169 | 122 | FOSL2 | A549 | Lung | ENCSR242E | 7125 | 63079 |
| 110.4282 | 72 | 91.26678 | 75 | 112.3224 | 113 | 233.482 | 165 | FOSL2 | A549 | Lung | ENCSR242E | 11419 | 63080 |
| 131.6497 | 98 | 134.2865 | 112 | 113.984 | 146 | 275.9091 | 223 | FOSL2 | A549 | Lung | ENCSR242E | 17150 | 63081 |
| 205.7798 | 36 | 220.6108 | 41 | 114.3562 | 36 | 161.034 | 46 | PRPF4 | HepG2 | Liver | ENCSR243I | 1344 | 63089 |
| 193.8115 | 33 | 193.4811 | 36 | 114.7796 | 34 | 185.6906 | 47 | PRPF4 | HepG2 | Liver | ENCSR243I | 1188 | 63090 |
| 1229.222 | 392 | 1258.449 | 503 | 940.4319 | 357 | 916.5293 | 379 | SMAD5 | GM12878 | Blood | ENCSR251( | 9200 | 63097 |
| 1233.053 | 245 | 1228.673 | 296 | 973.1355 | 230 | 938.1121 | 238 | SMAD5 | GM12878 | Blood | ENCSR251( | 5037 | 63098 |
| 1121.482 | 430 | 1119.237 | 523 | 842.3357 | 418 | 809.8662 | 435 | ATF1 | HepG2 | Liver | ENCSR253( | 15165 | 63104 |
| 1169.085 | 378 | 1186.391 | 474 | 914.0792 | 377 | 850.2825 | 359 | ATF1 | HepG2 | Liver | ENCSR253( | 10623 | 63105 |
| 118.8385 | 42 | 62.56429 | 34 | 78.56383 | 51 | 286.68 | 102 | NR3C1 | A549 | Lung | ENCSR255( | 3688 | 63108 |
| 116.8743 | 26 | 92.19871 | 25 | 60.23636 | 26 | 248.0448 | 59 | NR3C1 | A549 | Lung | ENCSR255( | 1357 | 63109 |
| 378.597 | 85 | 325.5593 | 86 | 225.7021 | 92 | 348.1173 | 122 | NR3C1 | A549 | Lung | ENCSR255( | 4385 | 63110 |
| 340.8111 | 48 | 511.1365 | 67 | 139.6176 | 39 | 296.449 | 63 | HNRNPLL | HepG2 | Liver | ENCSR255I | 1252 | 63112 |
| 811.8602 | 168 | 982.8475 | 218 | 574.5611 | 197 | 464.0665 | 189 | ZNF83 | K562 | Bone Marr | ENCSR257X | 8034 | 63117 |
| 965.0937 | 256 | 957.0355 | 306 | 625.1655 | 259 | 553.9892 | 262 | ZNF83 | K562 | Bone Marr | ENCSR257X | 12652 | 63118 |
| 103.4097 | 30 | 113.8837 | 35 | 62.6836 | 34 | 205.3579 | 66 | EP300 | A549 | Lung | ENCSR260\ | 2156 | 63124 |
| 186.8631 | 68 | 120.2653 | 61 | 177.9662 | 98 | 298.6492 | 134 | EP300 | A549 | Lung | ENCSR260\ | 6158 | 63125 |
| 118.6998 | 39 | 82.78413 | 36 | 98.20164 | 52 | 300.0983 | 97 | EP300 | A549 | Lung | ENCSR260\ | 3142 | 63126 |
| 711.1756 | 246 | 829.3643 | 340 | 508.1393 | 310 | 479.6322 | 326 | MNT | HepG2 | Liver | ENCSR261E | 21742 | 63127 |
| 791.2133 | 242 | 860.0097 | 322 | 442.1133 | 269 | 418.5719 | 283 | MNT | HepG2 | Liver | ENCSR261E | 18476 | 63128 |
| 759.0218 | 297 | 781.6873 | 382 | 458.4834 | 351 | 465.436 | 382 | PBX2 | K562 | Bone Marr | ENCSR263I | 29206 | 63132 |
| 728.4612 | 295 | 746.741 | 363 | 457.3297 | 361 | 419.7998 | 378 | PBX2 | K562 | Bone Marr | ENCSR263I | 31182 | 63133 |
| 587.3964 | 185 | 666.79 | 219 | 269.0342 | 191 | 221.7881 | 190 | POU5F1 | None | None | ENCSR264F | 15394 | 63136 |
| 243.2914 | 83 | 177.8486 | 79 | 77.88032 | 74 | 80.45004 | 81 | POU5F1 | None | None | ENCSR264F | 7268 | 63137 |
| 147.0144 | 39 | 274.401 | 61 | 86.99389 | 43 | 224.1854 | 74 | KLF4 | MCF-7 | Breast | ENCSR265\ | 2468 | 63139 |
| 218.7266 | 54 | 323.8173 | 74 | 173.3216 | 68 | 242.5917 | 86 | KLF4 | MCF-7 | Breast | ENCSR265\ | 3067 | 63140 |
| 144.8211 | 46 | 118.7911 | 46 | 108.7188 | 58 | 131.4938 | 68 | FOXA1 | HepG2 | Liver | ENCSR267I | 3437 | 63146 |
| 145.5267 | 49 | 128.3756 | 51 | 115.7233 | 64 | 132.1311 | 73 | FOXA1 | HepG2 | Liver | ENCSR267I | 3925 | 63147 |
| 566.5187 | 75 | 608.6128 | 85 | 414.9251 | 84 | 401.4155 | 87 | SRSF3 | K562 | Bone Marr | ENCSR268( | 1786 | 63150 |
| 182.2144 | 81 | 196.5667 | 94 | 101.1276 | 95 | 258.6235 | 153 | CEBPB | A549 | Lung | ENCSR269I | 9136 | 63152 |
| 127.8366 | 71 | 124.5769 | 79 | 83.79004 | 92 | 247.7065 | 156 | CEBPB | A549 | Lung | ENCSR269I | 9824 | 63153 |
| 53.04398 | 32 | 80.97352 | 44 | 24.93562 | 37 | 162.749 | 88 | CEBPB | A549 | Lung | ENCSR269I | 4859 | 63154 |
| 40.37494 | 16 | 21.28342 | 13 | 16.27912 | 16 | 108.1115 | 42 | FOSL2 | A549 | Lung | ENCSR274\ | 1717 | 63169 |
| 106.0243 | 59 | 94.24941 | 63 | 75.34636 | 79 | 253.669 | 143 | FOSL2 | A549 | Lung | ENCSR274\ | 8212 | 63170 |
| 49.98476 | 24 | 32.52467 | 22 | 28.88457 | 29 | 178.5626 | 72 | FOSL2 | A549 | Lung | ENCSR274\ | 3033 | 63171 |
| 979.1076 | 322 | 967.8404 | 381 | 733.523 | 322 | 692.2676 | 327 | ETV1 | K562 | Bone Marr | ENCSR277I | 15806 | 63176 |
| 975.8102 | 320 | 972.6317 | 387 | 735.8241 | 326 | 715.234 | 352 | ETV1 | K562 | Bone Marr | ENCSR277I | 15877 | 63177 |
| 19.71096 | 9 | 12.41066 | 8 | 7.896999 | 9 | 15.08987 | 13 | IKZF1 | HepG2 | Liver | ENCSR278J | 1037 | 63180 |
| 1027.268 | 225 | 1038.14 | 284 | 543.743 | 202 | 625.1203 | 231 | NBN | GM12878 | Blood | ENCSR278S | 8501 | 63182 |
| 632.7236 | 101 | 785.7149 | 125 | 418.7049 | 109 | 498.2537 | 127 | NBN | GM12878 | Blood | ENCSR278S | 3074 | 63183 |
| 157.0922 | 54 | 83.95309 | 44 | 95.98734 | 63 | 347.8617 | 125 | NR3C1 | A549 | Lung | ENCSR287F | 4569 | 63195 |
| 122.6842 | 51 | 97.59946 | 51 | 118.5305 | 75 | 327.3339 | 130 | NR3C1 | A549 | Lung | ENCSR287F | 5289 | 63196 |
| 1153.824 | 324 | 1189.583 | 427 | 911.1956 | 331 | 899.3319 | 363 | BCL3 | A549 | Lung | ENCSR288E | 9341 | 63199 |
| 1164.556 | 393 | 1160.544 | 475 | 884.0178 | 376 | 872.7546 | 412 | BCL3 | A549 | Lung | ENCSR288E | 11737 | 63200 |
| 236.3573 | 56 | 161.7674 | 50 | 109.4716 | 53 | 94.73572 | 53 | LARP7 | K562 | Bone Marr | ENCSR288I | 2755 | 63201 |
| 168.5829 | 41 | 135.614 | 40 | 77.58682 | 39 | 140.6781 | 56 | LARP7 | K562 | Bone Marr | ENCSR288I | 2173 | 63202 |
| 25.20932 | 12 | 10.93361 | 9 | 1.537502 | 6 | 32.4209 | 22 | ZNF354C | HEK293 | Embryonic | ENCSR289I | 1421 | 63203 |
| 1025.033 | 485 | 1047.246 | 618 | 771.8857 | 507 | 722.8905 | 506 | GABPA | K562 | Bone Marr | ENCSR290I | 23401 | 63207 |
| 1045.382 | 503 | 1063.727 | 635 | 775.5904 | 506 | 742.449 | 531 | GABPA | K562 | Bone Marr | ENCSR290I | 23609 | 63208 |
| 1111.753 | 591 | 1137.871 | 738 | 842.4523 | 598 | 792.4192 | 608 | MAZ | HEK293 | Embryonic | ENCSR290S | 21801 | 63209 |
| 1090.661 | 400 | 1110.858 | 508 | 852.7494 | 417 | 835.4889 | 458 | MAZ | HEK293 | Embryonic | ENCSR290S | 13872 | 63210 |
| 296.9354 | 189 | 292.5116 | 211 | 112.8216 | 200 | 113.4887 | 219 | MTA2 | GM12878 | Blood | ENCSR293( | 24224 | 63212 |
| 407.6327 | 224 | 513.8238 | 278 | 221.2 | 267 | 226.8404 | 293 | MTA2 | GM12878 | Blood | ENCSR293( | 28500 | 63213 |
| 582.9749 | 207 | 723.0803 | 256 | 419.3578 | 264 | 540.5084 | 317 | HNF4G | None | Liver | ENCSR297( | 19469 | 63224 |
| 510.4163 | 197 | 645.8568 | 246 | 375.5869 | 256 | 461.7915 | 301 | HNF4G | None | Liver | ENCSR297( | 19969 | 63225 |
| 29.00376 | 15 | 83.67691 | 29 | 34.132 | 25 | 145.5255 | 54 | CEBPB | A549 | Lung | ENCSR299I | 2086 | 63230 |
| 331.6694 | 131 | 321.1851 | 144 | 180.4011 | 150 | 352.5956 | 215 | CEBPB | A549 | Lung | ENCSR299I | 13304 | 63231 |
| 434.9351 | 180 | 541.0696 | 223 | 318.1384 | 235 | 505.2173 | 308 | CEBPB | A549 | Lung | ENCSR299I | 19322 | 63232 |
| 1005.246 | 297 | 949.4746 | 314 | 762.012 | 300 | 727.2859 | 308 | FOXK2 | K562 | Bone Marr | ENCSR302/ | 13037 | 63240 |
| 979.5046 | 323 | 974.1936 | 387 | 759.3497 | 345 | 736.0311 | 370 | FOXK2 | K562 | Bone Marr | ENCSR302/ | 15334 | 63241 |
| 678.5363 | 209 | 845.0349 | 272 | 403.4347 | 238 | 350.33 | 241 | CTCF | None | Breast | ENCSR304X | 15733 | 63245 |
| 25.49466 | 27 | 23.73965 | 30 | 21.41793 | 41 | 23.18293 | 46 | ZNF680 | HEK293 | Embryonic | ENCSR307( | 5809 | 63246 |
| 20.95632 | 24 | 43.22384 | 37 | 20.8239 | 39 | 29.27016 | 48 | ZNF680 | HEK293 | Embryonic | ENCSR307( | 5481 | 63247 |
| 23.4492 | 13 | 31.76411 | 17 | 13.82443 | 16 | 13.07025 | 17 | PTTG1 | K562 | Bone Marr | ENCSR314E | 1806 | 63264 |
| 54.88774 | 21 | 57.63713 | 24 | 31.45122 | 24 | 17.10392 | 20 | PTTG1 | K562 | Bone Marr | ENCSR314E | 1958 | 63265 |
| 377.8584 | 68 | 552.3963 | 93 | 235.8244 | 73 | 305.7708 | 89 | DROSHA | HepG2 | Liver | ENCSR316J | 2658 | 63270 |
| 234.8838 | 56 | 372.7902 | 80 | 142.7666 | 62 | 262.7604 | 90 | DROSHA | HepG2 | Liver | ENCSR316J | 3230 | 63271 |
| 1006.222 | 199 | 995.9482 | 234 | 417.0604 | 170 | 318.8146 | 160 | ZBTB40 | MCF-7 | Breast | ENCSR318I | 7705 | 63277 |

| | | | | | | | | | | | | | |
|---|---|---|---|---|---|---|---|---|---|---|---|---|---|
| 999.3632 | 256 | 976.4943 | 292 | 611.4908 | 240 | 440.3734 | 220 | ZBTB40 | MCF-7 | Breast | ENCSR318I | 10562 | 63278 |
| 525.7603 | 190 | 583.3464 | 222 | 391.5474 | 242 | 392.8082 | 261 | ZEB2 | K562 | Bone Marr | ENCSR322( | 16070 | 63282 |
| 439.7599 | 139 | 429.3421 | 152 | 281.123 | 162 | 299.225 | 179 | ZEB2 | K562 | Bone Marr | ENCSR322( | 9946 | 63283 |
| 453.1914 | 132 | 676.0561 | 180 | 240.6087 | 141 | 253.4574 | 155 | CTCF | None | Sigmoid Cc | ENCSR323I | 9183 | 63284 |
| 540.698 | 211 | 641.975 | 255 | 651.8698 | 336 | 670.1713 | 382 | OSR2 | HEK293 | Embryonic | ENCSR324I | 20105 | 63285 |
| 606.6911 | 200 | 698.6151 | 238 | 664.1126 | 302 | 706.2944 | 348 | OSR2 | HEK293 | Embryonic | ENCSR324I | 16143 | 63286 |
| 1023.079 | 630 | 1058.789 | 803 | 767.1917 | 671 | 738.317 | 718 | ZNF335 | HEK293 | Embryonic | ENCSR328! | 31876 | 63294 |
| 1059.751 | 627 | 1094.093 | 798 | 801.1617 | 664 | 742.7884 | 667 | ZNF335 | HEK293 | Embryonic | ENCSR328! | 28088 | 63295 |
| 182.9491 | 105 | 158.2827 | 111 | 179.3803 | 160 | 363.7991 | 232 | FOSL2 | A549 | Lung | ENCSR330I | 14866 | 63298 |
| 76.04644 | 43 | 67.82166 | 46 | 54.72695 | 58 | 218.4323 | 114 | FOSL2 | A549 | Lung | ENCSR330I | 6089 | 63299 |
| 980.7217 | 143 | 968.0523 | 154 | 461.8415 | 126 | 523.3008 | 143 | NKRF | K562 | Bone Marr | ENCSR331I | 3600 | 63301 |
| 1074.264 | 272 | 1057.635 | 311 | 823.5575 | 256 | 802.1956 | 274 | NKRF | K562 | Bone Marr | ENCSR331I | 8166 | 63302 |
| 1056.23 | 179 | 1094.087 | 246 | 484.3703 | 162 | 427.1787 | 162 | ZBTB11 | K562 | Bone Marr | ENCSR331( | 6329 | 63303 |
| 1118.644 | 283 | 1118.944 | 347 | 842.0963 | 260 | 748.3818 | 252 | ZBTB11 | K562 | Bone Marr | ENCSR331( | 8793 | 63304 |
| 1034.504 | 576 | 1065.719 | 738 | 787.4586 | 636 | 748.9967 | 663 | CREB1 | HepG2 | Liver | ENCSR331( | 31564 | 63305 |
| 1048.419 | 520 | 1069.938 | 656 | 798.0833 | 551 | 756.3109 | 564 | CREB1 | HepG2 | Liver | ENCSR331( | 24897 | 63306 |
| 193.8041 | 42 | 137.7857 | 38 | 52.49259 | 30 | 78.3358 | 39 | ZNF318 | K562 | Bone Marr | ENCSR334I | 1843 | 63309 |
| 404.3753 | 54 | 578.2093 | 73 | 180.2025 | 45 | 182.4362 | 48 | SP1 | HepG2 | Liver | ENCSR334I | 1176 | 63312 |
| 95.37748 | 26 | 136.4438 | 35 | 21.13841 | 18 | 88.1878 | 38 | HDAC2 | HepG2 | Liver | ENCSR337I | 1606 | 63314 |
| 177.7789 | 43 | 248.1103 | 57 | 113.2779 | 48 | 149.8956 | 59 | HDAC2 | HepG2 | Liver | ENCSR337I | 2263 | 63315 |
| 517.9023 | 288 | 648.7317 | 356 | 327.6072 | 366 | 331.7677 | 400 | SCRT2 | HEK293 | Embryonic | ENCSR338I | 38488 | 63316 |
| 384.9315 | 231 | 515.2217 | 294 | 267.9641 | 305 | 266.4844 | 331 | SCRT2 | HEK293 | Embryonic | ENCSR338I | 31601 | 63317 |
| 770.8203 | 271 | 809.8359 | 356 | 649.7 | 375 | 644.3773 | 423 | NR2F2 | None | Liver | ENCSR338I | 25902 | 63318 |
| 856.039 | 366 | 883.0184 | 478 | 692.0263 | 478 | 687.235 | 537 | NR2F2 | None | Liver | ENCSR338I | 30250 | 63319 |
| 1082.72 | 396 | 1113.482 | 511 | 822.9482 | 390 | 798.2637 | 419 | ELK1 | K562 | Bone Marr | ENCSR338( | 14929 | 63321 |
| 1073.082 | 377 | 1100.74 | 489 | 830.9437 | 389 | 798.2592 | 402 | ELK1 | K562 | Bone Marr | ENCSR338( | 14350 | 63322 |
| 603.4619 | 226 | 800.293 | 310 | 357.5998 | 265 | 323.9526 | 275 | CTCF | None | Sigmoid Cc | ENCSR341\ | 21068 | 63328 |
| 452.7492 | 142 | 516.6889 | 168 | 308.475 | 170 | 234.9827 | 162 | LEF1 | K562 | Bone Marr | ENCSR343I | 10007 | 63332 |
| 650.4264 | 201 | 601.0729 | 214 | 330.5088 | 212 | 267.7316 | 209 | LEF1 | K562 | Bone Marr | ENCSR343I | 14339 | 63333 |
| 1170.134 | 234 | 1191.456 | 302 | 854.3042 | 212 | 812.7512 | 219 | ZBTB33 | None | Liver | ENCSR345Y | 5935 | 63346 |
| 869.1184 | 107 | 996.4511 | 126 | 392.8945 | 90 | 384.0509 | 94 | ZBTB33 | None | Liver | ENCSR345Y | 2215 | 63347 |
| 1517.817 | 162 | 1558.941 | 214 | 1059.596 | 130 | 1220.341 | 151 | CEBPZ | GM12878 | Blood | ENCSR347I | 1291 | 63354 |
| 1522.882 | 179 | 1524.648 | 215 | 981.5021 | 128 | 1052.037 | 140 | CEBPZ | GM12878 | Blood | ENCSR347I | 1403 | 63355 |
| 77.91198 | 21 | 70.90499 | 22 | 59.22286 | 26 | 83.06183 | 33 | ZFP41 | HEK293 | Embryonic | ENCSR348( | 1291 | 63356 |
| 64.00937 | 17 | 48.02491 | 16 | 58.06541 | 23 | 31.07952 | 18 | ZFP41 | HEK293 | Embryonic | ENCSR348( | 1030 | 63357 |
| 316.7504 | 92 | 266.9815 | 93 | 156.3989 | 94 | 196.5538 | 112 | ZNF318 | K562 | Bone Marr | ENCSR352I | 6013 | 63364 |
| 264.869 | 78 | 263.8731 | 86 | 225.4507 | 103 | 138.6797 | 88 | ZNF318 | K562 | Bone Marr | ENCSR352I | 5165 | 63365 |
| 67.44798 | 25 | 63.29241 | 27 | 27.95475 | 25 | 130.2968 | 55 | HES2 | A549 | Lung | ENCSR356Y | 2428 | 63374 |
| 467.7584 | 90 | 499.0509 | 102 | 310.1944 | 99 | 207.8541 | 86 | ZSCAN5A | HEK293 | Embryonic | ENCSR357( | 3374 | 63376 |
| 318.3733 | 64 | 473.2183 | 88 | 223.1765 | 73 | 172.0144 | 68 | ZSCAN5A | HEK293 | Embryonic | ENCSR357( | 2591 | 63377 |
| 49.53533 | 17 | 35.86714 | 16 | 56.9192 | 27 | 266.7389 | 65 | EP300 | A549 | Lung | ENCSR358I | 1537 | 63378 |
| 149.024 | 57 | 115.2503 | 56 | 132.3398 | 80 | 318.7983 | 130 | EP300 | A549 | Lung | ENCSR358I | 5443 | 63379 |
| 943.703 | 261 | 925.018 | 304 | 642.4453 | 277 | 531.873 | 272 | USF2 | K562 | Bone Marr | ENCSR359I | 14613 | 63382 |
| 916.5538 | 280 | 869.9388 | 305 | 558.2662 | 283 | 482.8434 | 285 | USF2 | K562 | Bone Marr | ENCSR359I | 17292 | 63383 |
| 239.6732 | 142 | 244.4585 | 161 | 174.4975 | 190 | 169.7197 | 204 | KDM1A | K562 | Bone Marr | ENCSR360I | 18861 | 63384 |
| 165.8945 | 84 | 129.8269 | 84 | 100.2689 | 101 | 92.4238 | 106 | KDM1A | K562 | Bone Marr | ENCSR360I | 8947 | 63385 |
| 813.8792 | 114 | 1190.195 | 162 | 504.9032 | 115 | 405.4373 | 108 | CHD1 | MCF-7 | Breast | ENCSR360J | 2527 | 63386 |
| 1110.472 | 183 | 1134.365 | 238 | 813.3667 | 194 | 679.9335 | 187 | CHD1 | MCF-7 | Breast | ENCSR360J | 4597 | 63387 |
| 793.4475 | 287 | 775.3002 | 334 | 422.557 | 316 | 336.3535 | 312 | POU5F1 | None | None | ENCSR362\ | 27213 | 63393 |
| 104.674 | 35 | 100.4368 | 38 | 171.389 | 65 | 132.4445 | 61 | ZNF488 | HEK293 | Embryonic | ENCSR363) | 2741 | 63394 |
| 61.68434 | 26 | 57.37285 | 28 | 173.5459 | 64 | 172.4036 | 68 | ZNF488 | HEK293 | Embryonic | ENCSR363) | 2606 | 63395 |
| 438.3972 | 94 | 486.9655 | 109 | 232.9068 | 93 | 114.5653 | 71 | ZFP37 | HEK293 | Embryonic | ENCSR365( | 3791 | 63398 |
| 504.4147 | 103 | 548.8596 | 118 | 282.1951 | 104 | 210.1907 | 96 | ZFP37 | HEK293 | Embryonic | ENCSR365( | 3946 | 63399 |
| 941.2742 | 487 | 931.4523 | 578 | 661.2238 | 480 | 616.3192 | 484 | CBX5 | GM12878 | Blood | ENCSR372( | 33699 | 63410 |
| 884.4152 | 386 | 892.8042 | 479 | 655.0724 | 421 | 615.8587 | 432 | CBX5 | GM12878 | Blood | ENCSR372( | 29223 | 63411 |
| 231.801 | 96 | 302.303 | 122 | 125.5119 | 110 | 333.552 | 181 | CEBPB | A549 | Lung | ENCSR375I | 9976 | 63414 |
| 669.0338 | 319 | 669.3042 | 364 | 467.5715 | 420 | 550.2194 | 488 | CEBPB | A549 | Lung | ENCSR375I | 41965 | 63415 |
| 937.1506 | 238 | 946.0061 | 296 | 688.3851 | 269 | 629.9113 | 276 | ZFP69B | HEK293 | Embryonic | ENCSR381\ | 11830 | 63423 |
| 887.9581 | 198 | 966.6907 | 261 | 594.7001 | 226 | 649.1012 | 252 | ZFP69B | HEK293 | Embryonic | ENCSR381\ | 9703 | 63424 |
| 85.23458 | 34 | 81.2595 | 37 | 80.77203 | 49 | 24.85392 | 32 | DDX20 | K562 | Bone Marr | ENCSR382/ | 3219 | 63425 |
| 74.20374 | 28 | 87.85815 | 34 | 73.23687 | 41 | 12.5258 | 21 | DDX20 | K562 | Bone Marr | ENCSR382/ | 2539 | 63426 |
| 875.4191 | 174 | 688.5584 | 167 | 523.6044 | 184 | 560.1933 | 203 | INSM2 | HEK293 | Embryonic | ENCSR382( | 7153 | 63427 |
| 535.4295 | 133 | 505.5671 | 142 | 473.1399 | 176 | 547.9093 | 202 | INSM2 | HEK293 | Embryonic | ENCSR382( | 7350 | 63428 |
| 982.5084 | 718 | 1012.967 | 903 | 751.2655 | 865 | 723.5959 | 937 | YY1 | None | Liver | ENCSR382I | 56955 | 63430 |
| 1012.594 | 585 | 1046.762 | 747 | 796.2126 | 691 | 741.6778 | 693 | YY1 | None | Liver | ENCSR382I | 33981 | 63431 |
| 767.6733 | 102 | 1064.355 | 135 | 395.4427 | 93 | 385.0036 | 97 | ELK1 | MCF-7 | Breast | ENCSR382\ | 2290 | 63432 |
| 1208.742 | 275 | 1228.699 | 350 | 946.3619 | 254 | 891.6566 | 244 | ELK1 | MCF-7 | Breast | ENCSR382\ | 5631 | 63433 |
| 811.9715 | 214 | 892.2954 | 272 | 535.4365 | 249 | 438.864 | 244 | ZNF24 | K562 | Bone Marr | ENCSR385/ | 13217 | 63440 |
| 352.3166 | 107 | 479.9647 | 139 | 228.8191 | 125 | 145.9297 | 110 | ZNF24 | K562 | Bone Marr | ENCSR385/ | 7395 | 63441 |
| 161.9942 | 77 | 152.904 | 84 | 154.7023 | 114 | 335.3859 | 173 | NR3C1 | A549 | Lung | ENCSR385I | 9100 | 63442 |
| 107.3865 | 26 | 103.0393 | 28 | 58.00241 | 27 | 227.3286 | 59 | NR3C1 | A549 | Lung | ENCSR385I | 1504 | 63443 |
| 149.7345 | 63 | 131.1312 | 66 | 153.6248 | 95 | 327.6892 | 145 | NR3C1 | A549 | Lung | ENCSR385I | 6508 | 63444 |

| | | | | | | | | | | | | | |
|---|---|---|---|---|---|---|---|---|---|---|---|---|---|
| 1384.041 | 268 | 1396.095 | 330 | 1078.6 | 225 | 1044.73 | 231 | DEAF1 | K562 | Bone Marr | ENCSR387S | 3453 | 63449 |
| 917.8215 | 526 | 913.6247 | 632 | 684.967 | 582 | 656.9421 | 620 | HDAC1 | K562 | Bone Marr | ENCSR387U | 36613 | 63450 |
| 928.3785 | 484 | 930.0874 | 586 | 682.6604 | 509 | 649.9595 | 534 | HDAC1 | K562 | Bone Marr | ENCSR387U | 31312 | 63451 |
| 538.282 | 100 | 553.4665 | 111 | 587.7058 | 144 | 473.9839 | 136 | RCOR1 | MCF-7 | Breast | ENCSR391J | 3750 | 63457 |
| 666.4655 | 127 | 826.7597 | 157 | 625.1615 | 169 | 628.7423 | 180 | RCOR1 | MCF-7 | Breast | ENCSR391J | 5008 | 63458 |
| 204.7666 | 40 | 160.3967 | 38 | 135.0601 | 44 | 90.05033 | 38 | MTA3 | MCF-7 | Breast | ENCSR391H | 1576 | 63459 |
| 771.0553 | 95 | 913.2949 | 114 | 379.348 | 83 | 388.9958 | 89 | MTA3 | MCF-7 | Breast | ENCSR391H | 1830 | 63460 |
| 484.2818 | 249 | 688.7854 | 326 | 294.1737 | 308 | 466.3876 | 398 | IKZF1 | K562 | Bone Marr | ENCSR395H | 30843 | 63462 |
| 582.6642 | 325 | 675.7623 | 417 | 377.1017 | 418 | 521.6757 | 511 | IKZF1 | K562 | Bone Marr | ENCSR395H | 44496 | 63463 |
| 1033.424 | 331 | 1047.279 | 414 | 777.553 | 312 | 752.5032 | 331 | POLR2H | K562 | Bone Marr | ENCSR400F | 11451 | 63477 |
| 1064.672 | 293 | 1074.801 | 363 | 798.5503 | 265 | 762.3023 | 275 | POLR2H | K562 | Bone Marr | ENCSR400F | 8597 | 63478 |
| 565.6079 | 300 | 709.8629 | 408 | 371.0522 | 389 | 359.4218 | 418 | CTCF | MM.1S | Blood | ENCSR402I | 41832 | 63479 |
| 602.7235 | 313 | 723.4275 | 433 | 392.5607 | 403 | 428.2876 | 452 | CTCF | MM.1S | Blood | ENCSR402I | 42436 | 63480 |
| 1179.399 | 167 | 1036.686 | 159 | 512.7107 | 131 | 512.5892 | 139 | CSDE1 | K562 | Bone Marr | ENCSR402X | 3265 | 63484 |
| 1002.644 | 149 | 822.7514 | 144 | 386.0416 | 118 | 371.1349 | 123 | CSDE1 | K562 | Bone Marr | ENCSR402X | 3742 | 63485 |
| 156.9466 | 45 | 130.0145 | 45 | 67.49311 | 43 | 37.98327 | 36 | PYGO2 | K562 | Bone Marr | ENCSR410D | 2979 | 63514 |
| 176.9363 | 50 | 130.3938 | 47 | 95.80785 | 53 | 56.93107 | 45 | PYGO2 | K562 | Bone Marr | ENCSR410D | 3221 | 63515 |
| 681.9861 | 162 | 604.3601 | 167 | 409.5094 | 176 | 389.8063 | 184 | MTA2 | K562 | Bone Marr | ENCSR411U | 8301 | 63517 |
| 609.9281 | 165 | 552.2475 | 173 | 354.6845 | 181 | 326.3645 | 187 | MTA2 | K562 | Bone Marr | ENCSR411U | 10246 | 63518 |
| 503.807 | 89 | 651.2541 | 113 | 231.1183 | 81 | 229.8828 | 86 | SUZ12 | K562 | Bone Marr | ENCSR412( | 3334 | 63519 |
| 572.833 | 88 | 855.6597 | 122 | 226.4213 | 72 | 264.2851 | 83 | SUZ12 | K562 | Bone Marr | ENCSR412( | 2669 | 63520 |
| 911.0087 | 228 | 891.4481 | 263 | 380.367 | 207 | 355.9888 | 216 | TARDBP | GM12878 | Blood | ENCSR412( | 12112 | 63522 |
| 908.9358 | 217 | 867.5722 | 233 | 338.1626 | 190 | 311.7074 | 197 | TARDBP | GM12878 | Blood | ENCSR412( | 11436 | 63523 |
| 1010.444 | 435 | 999.2093 | 511 | 772.1961 | 456 | 735.068 | 470 | ZFP2 | HEK293 | Embryonic | ENCSR413A | 19475 | 63524 |
| 1038.273 | 396 | 1070.213 | 513 | 832.5983 | 445 | 781.2747 | 443 | ZFP2 | HEK293 | Embryonic | ENCSR413A | 16336 | 63525 |
| 184.3032 | 196 | 291.513 | 265 | 133.0421 | 280 | 136.444 | 309 | CEBPB | K562 | Bone Marr | ENCSR416( | 40818 | 63533 |
| 197.3311 | 187 | 291.5432 | 247 | 127.6826 | 255 | 130.6828 | 281 | CEBPB | K562 | Bone Marr | ENCSR416( | 36240 | 63534 |
| 966.7172 | 355 | 965.8313 | 432 | 764.7499 | 400 | 746.4333 | 435 | ZEB2 | HEK293 | Embryonic | ENCSR417V | 17293 | 63535 |
| 990.4811 | 403 | 1003.188 | 497 | 783.6316 | 449 | 776.9307 | 504 | ZEB2 | HEK293 | Embryonic | ENCSR417V | 18664 | 63536 |
| 1006.575 | 408 | 989.4467 | 475 | 755.0303 | 413 | 746.694 | 460 | ZNF692 | HEK293 | Embryonic | ENCSR418I | 18474 | 63537 |
| 992.7044 | 352 | 977.2289 | 407 | 776.6561 | 381 | 754.5116 | 409 | ZNF692 | HEK293 | Embryonic | ENCSR418I | 15722 | 63538 |
| 82.1111 | 29 | 80.64259 | 32 | 56.39407 | 36 | 225.2044 | 76 | HES2 | A549 | Lung | ENCSR419T | 2643 | 63542 |
| 134.222 | 57 | 176.862 | 73 | 118.8166 | 81 | 293.6717 | 132 | HES2 | A549 | Lung | ENCSR419T | 6146 | 63543 |
| 810.4032 | 475 | 799.2123 | 561 | 680.7794 | 662 | 689.0709 | 771 | GATA3 | MCF-7 | Breast | ENCSR423F | 44336 | 63545 |
| 746.3331 | 437 | 733.5663 | 515 | 635.1518 | 636 | 640.8148 | 737 | GATA3 | MCF-7 | Breast | ENCSR423F | 49376 | 63546 |
| 138.2826 | 58 | 105.9515 | 57 | 112.9404 | 79 | 290.3848 | 131 | NR3C1 | A549 | Lung | ENCSR424J | 5978 | 63549 |
| 172.4593 | 61 | 108.6344 | 54 | 127.3898 | 78 | 313.5075 | 128 | NR3C1 | A549 | Lung | ENCSR424J | 5369 | 63550 |
| 68.30273 | 29 | 54.29325 | 29 | 53.17016 | 39 | 159.4157 | 70 | HES2 | A549 | Lung | ENCSR425I | 3169 | 63551 |
| 238.2863 | 112 | 240.2022 | 126 | 170.5503 | 147 | 376.7613 | 222 | HES2 | A549 | Lung | ENCSR425I | 13329 | 63552 |
| 90.09751 | 30 | 59.66848 | 27 | 45.311 | 32 | 161.16 | 63 | HES2 | A549 | Lung | ENCSR425I | 2564 | 63553 |
| 77.57613 | 34 | 73.82821 | 37 | 38.81117 | 37 | 38.68864 | 40 | MIER1 | K562 | Bone Marr | ENCSR426I | 3268 | 63554 |
| 195.2277 | 65 | 351.3849 | 98 | 125.2585 | 76 | 103.3372 | 75 | MIER1 | K562 | Bone Marr | ENCSR426I | 4797 | 63555 |
| 212.5073 | 41 | 274.2231 | 52 | 164.7635 | 49 | 119.6231 | 44 | MLLT1 | MCF-7 | Breast | ENCSR427E | 1562 | 63557 |
| 115.1305 | 36 | 105.1782 | 38 | 67.04575 | 40 | 63.77794 | 42 | ZNF239 | HEK293 | Embryonic | ENCSR440( | 2579 | 63579 |
| 123.379 | 46 | 163.2823 | 59 | 107.0627 | 63 | 130.1625 | 74 | ZNF239 | HEK293 | Embryonic | ENCSR440( | 3980 | 63580 |
| 527.9977 | 151 | 521.7979 | 166 | 295.4894 | 165 | 227.7601 | 158 | ADNP | K562 | Bone Marr | ENCSR440V | 10370 | 63581 |
| 445.7892 | 152 | 457.5623 | 171 | 254.1822 | 172 | 234.4064 | 179 | ADNP | K562 | Bone Marr | ENCSR440V | 12568 | 63582 |
| 174.4608 | 100 | 248.0513 | 132 | 220.0983 | 169 | 354.1151 | 223 | TCF7 | HepG2 | Liver | ENCSR444L | 14093 | 63591 |
| 535.5788 | 211 | 659.4709 | 260 | 487.1199 | 301 | 610.4111 | 357 | TCF7 | HepG2 | Liver | ENCSR444L | 22029 | 63592 |
| 490.8684 | 135 | 403.7665 | 135 | 331.5635 | 161 | 317.4627 | 169 | SOX13 | HepG2 | Liver | ENCSR445A | 9268 | 63593 |
| 914.4342 | 252 | 912.043 | 309 | 732.8799 | 304 | 706.2602 | 319 | SOX13 | HepG2 | Liver | ENCSR445A | 15367 | 63594 |
| 251.3581 | 68 | 232.7533 | 72 | 236.9663 | 94 | 216.5992 | 96 | GFI1B | HEK293 | Embryonic | ENCSR445F | 4197 | 63597 |
| 220.6744 | 56 | 154.8815 | 51 | 202.6481 | 76 | 182.9753 | 77 | GFI1B | HEK293 | Embryonic | ENCSR445F | 3221 | 63598 |
| 1078.015 | 233 | 1084.201 | 290 | 718.0578 | 218 | 599.8109 | 212 | DDX20 | K562 | Bone Marr | ENCSR446L | 7199 | 63599 |
| 1095.465 | 250 | 1078.027 | 294 | 806.7856 | 236 | 780.5945 | 247 | DDX20 | K562 | Bone Marr | ENCSR446L | 7705 | 63600 |
| 1084.332 | 256 | 1100.556 | 321 | 847.8481 | 262 | 863.6858 | 309 | BCL3 | A549 | Lung | ENCSR446S | 8776 | 63602 |
| 231.0808 | 118 | 272.7232 | 143 | 125.7732 | 141 | 270.4655 | 207 | CEBPB | A549 | Lung | ENCSR447Z | 15381 | 63605 |
| 369.1388 | 141 | 412.6724 | 166 | 235.9476 | 172 | 348.6943 | 219 | CEBPB | A549 | Lung | ENCSR447Z | 13908 | 63606 |
| 119.1085 | 43 | 155.9368 | 55 | 55.47659 | 45 | 171.4545 | 81 | CEBPB | A549 | Lung | ENCSR447Z | 3930 | 63607 |
| 35.93563 | 28 | 39.28449 | 33 | 42.4 | 47 | 23.76339 | 41 | ZNF558 | HEK293 | Embryonic | ENCSR447Z | 4827 | 63608 |
| 354.136 | 109 | 274.4348 | 106 | 211.7702 | 123 | 225.9218 | 136 | ZNF558 | HEK293 | Embryonic | ENCSR447Z | 7705 | 63609 |
| 378.7777 | 90 | 393.2384 | 101 | 153.4104 | 81 | 146.504 | 85 | ZKSCAN8 | K562 | Bone Marr | ENCSR448U | 4729 | 63610 |
| 496.1016 | 138 | 623.9572 | 172 | 261.0877 | 146 | 247.4644 | 153 | ZKSCAN8 | K562 | Bone Marr | ENCSR448U | 9220 | 63611 |
| 56.86481 | 25 | 34.65676 | 22 | 70.03565 | 41 | 50.52901 | 38 | ZKSCAN1 | MCF-7 | Breast | ENCSR449U | 2583 | 63614 |
| 44.24499 | 18 | 52.80831 | 22 | 45.18156 | 27 | 55.66237 | 32 | ZKSCAN1 | MCF-7 | Breast | ENCSR449U | 1772 | 63615 |
| 161.1059 | 52 | 214.6767 | 67 | 99.63036 | 60 | 84.95065 | 60 | ZNF280D | HEK293 | Embryonic | ENCSR451( | 3936 | 63618 |
| 132.1187 | 48 | 135.2715 | 54 | 80.50201 | 56 | 79.66316 | 60 | ZNF280D | HEK293 | Embryonic | ENCSR451( | 4132 | 63619 |
| 1019.979 | 212 | 1057.058 | 282 | 780.4749 | 233 | 659.4136 | 228 | ZNF501 | HEK293 | Embryonic | ENCSR461Z | 7082 | 63631 |
| 658.8563 | 117 | 748.0207 | 137 | 397.6094 | 120 | 292.0098 | 109 | ZNF501 | HEK293 | Embryonic | ENCSR461Z | 3610 | 63632 |
| 1045.277 | 186 | 1031.713 | 220 | 678.9222 | 198 | 566.1311 | 192 | ZNF101 | HEK293 | Embryonic | ENCSR462F | 6120 | 63633 |
| 158.2787 | 65 | 213.5126 | 84 | 102.0808 | 79 | 75.0167 | 75 | ZNF101 | HEK293 | Embryonic | ENCSR462F | 6222 | 63634 |
| 20.55674 | 14 | 13.14396 | 13 | 4.091643 | 12 | 27.75527 | 27 | ZNF140 | HEK293 | Embryonic | ENCSR464H | 2298 | 63640 |

| | | | | | | | | | | | | | |
|---|---|---|---|---|---|---|---|---|---|---|---|---|---|
| 6.428175 | 12 | 14.47483 | 19 | 3.785335 | 18 | 16.25132 | 31 | ZNF140 | HEK293 | Embryonic | ENCSR464I | 3991 | 63641 |
| 93.96535 | 22 | 128.9359 | 29 | 83.25992 | 29 | 78.68191 | 30 | FOXK2 | MCF-7 | Breast | ENCSR465\ | 1133 | 63642 |
| 163.3503 | 59 | 101.7064 | 52 | 89.13197 | 66 | 338.1836 | 132 | EP300 | A549 | Lung | ENCSR467\ | 5257 | 63649 |
| 194.3656 | 95 | 180.8001 | 103 | 247.3344 | 159 | 460.722 | 226 | EP300 | A549 | Lung | ENCSR467\ | 11321 | 63650 |
| 114.6576 | 39 | 62.637 | 32 | 58.81983 | 42 | 266.319 | 93 | EP300 | A549 | Lung | ENCSR467\ | 3303 | 63651 |
| 916.1718 | 403 | 888.282 | 459 | 729.1063 | 480 | 698.5066 | 503 | SP7 | HEK293 | Embryonic | ENCSR468I | 24319 | 63653 |
| 907.6632 | 407 | 907.5577 | 492 | 727.2309 | 492 | 687.414 | 506 | SP7 | HEK293 | Embryonic | ENCSR468I | 25109 | 63654 |
| 1162.042 | 471 | 1183.749 | 590 | 891.7192 | 454 | 839.7815 | 449 | SIN3A | MCF-7 | Breast | ENCSR468l | 12581 | 63655 |
| 1151.472 | 518 | 1178.625 | 653 | 901.5328 | 533 | 856.9763 | 546 | SIN3A | MCF-7 | Breast | ENCSR468l | 15695 | 63656 |
| 1166.074 | 353 | 1185.734 | 450 | 919.3354 | 358 | 894.8541 | 382 | BCL3 | A549 | Lung | ENCSR473( | 9947 | 63663 |
| 1201.025 | 550 | 1208.835 | 680 | 902.7922 | 536 | 891.491 | 594 | BCL3 | A549 | Lung | ENCSR473( | 17162 | 63664 |
| 414.6623 | 219 | 489.6914 | 265 | 285.2801 | 289 | 462.1389 | 377 | CEBPB | A549 | Lung | ENCSR474I | 30311 | 63665 |
| 77.4668 | 42 | 110.9833 | 56 | 59.91731 | 58 | 165.2752 | 97 | CEBPB | A549 | Lung | ENCSR474I | 5718 | 63666 |
| 354.6719 | 187 | 380.7346 | 217 | 223.3942 | 238 | 399.5494 | 323 | CEBPB | A549 | Lung | ENCSR474I | 25393 | 63667 |
| 452.1541 | 73 | 706.4533 | 104 | 361.6211 | 88 | 312.3066 | 86 | ELF1 | MCF-7 | Breast | ENCSR475S | 2359 | 63674 |
| 287.0949 | 46 | 397.121 | 61 | 170.9023 | 47 | 239.511 | 60 | ELF1 | MCF-7 | Breast | ENCSR475S | 1473 | 63675 |
| 546.8565 | 108 | 792.5031 | 146 | 429.6678 | 131 | 392.6858 | 133 | ZNF423 | HEK293 | Embryonic | ENCSR477( | 4370 | 63678 |
| 435.4302 | 96 | 677.1349 | 135 | 357.9644 | 121 | 444.5816 | 144 | ZNF423 | HEK293 | Embryonic | ENCSR477( | 4568 | 63679 |
| 855.5526 | 313 | 864.1204 | 394 | 675.1968 | 386 | 691.2308 | 459 | ATF3 | None | Liver | ENCSR480l | 24692 | 63683 |
| 877.1184 | 334 | 846.1608 | 380 | 690.9411 | 407 | 699.7741 | 475 | ATF3 | None | Liver | ENCSR480l | 24765 | 63684 |
| 1046.477 | 462 | 1038.731 | 552 | 805.5953 | 486 | 768.0462 | 500 | ZBTB8A | HEK293 | Embryonic | ENCSR481I | 19029 | 63685 |
| 1033.028 | 462 | 1037.479 | 564 | 787.9147 | 480 | 740.0537 | 483 | ZBTB8A | HEK293 | Embryonic | ENCSR481I | 20166 | 63686 |
| 422.5786 | 202 | 639.7611 | 274 | 285.6134 | 262 | 385.9096 | 319 | SMC3 | A549 | Lung | ENCSR481Y | 25926 | 63687 |
| 239.7016 | 93 | 310.5753 | 118 | 99.10489 | 95 | 181.9155 | 132 | SMC3 | A549 | Lung | ENCSR481Y | 9451 | 63688 |
| 932.0487 | 614 | 957.2281 | 777 | 731.8649 | 758 | 738.4313 | 877 | GLIS1 | HEK293 | Embryonic | ENCSR482I | 46623 | 63689 |
| 891.0901 | 457 | 912.6908 | 577 | 698.4332 | 549 | 716.3748 | 653 | GLIS1 | HEK293 | Embryonic | ENCSR482I | 33258 | 63690 |
| 894.306 | 475 | 900.4731 | 585 | 660.3817 | 518 | 636.2643 | 558 | CTCF | None | Spleen | ENCSR482I | 35200 | 63691 |
| 637.4297 | 155 | 585.8263 | 163 | 306.2406 | 152 | 282.3247 | 157 | ZC3H8 | K562 | Bone Marr | ENCSR494I | 8166 | 63718 |
| 432.338 | 88 | 236.3035 | 69 | 95.4422 | 57 | 155.9881 | 77 | ZC3H8 | K562 | Bone Marr | ENCSR494I | 3595 | 63719 |
| 1058.2 | 284 | 1048.227 | 338 | 805.2032 | 274 | 715.7173 | 268 | ELF1 | K562 | Bone Marr | ENCSR502( | 9772 | 63733 |
| 1057.872 | 228 | 1060.585 | 277 | 742.9554 | 226 | 610.2518 | 218 | ELF1 | K562 | Bone Marr | ENCSR502( | 7541 | 63734 |
| 257.2215 | 169 | 341.0279 | 215 | 177.1408 | 227 | 161.6973 | 239 | ZNF654 | HEK293 | Embryonic | ENCSR504\ | 26288 | 63737 |
| 267.4724 | 175 | 375.1415 | 228 | 193.648 | 239 | 200.5076 | 263 | ZNF654 | HEK293 | Embryonic | ENCSR504\ | 26998 | 63738 |
| 970.6018 | 327 | 954.5766 | 380 | 730.5764 | 336 | 733.8724 | 386 | FOXK2 | K562 | Bone Marr | ENCSR508I | 16854 | 63743 |
| 986.8692 | 338 | 984.0337 | 408 | 772.5952 | 368 | 728.0932 | 369 | FOXK2 | K562 | Bone Marr | ENCSR508I | 15443 | 63744 |
| 271.5652 | 78 | 378.5713 | 103 | 155.3681 | 86 | 140.1418 | 88 | CTCF | None | Skin | ENCSR515l | 5415 | 63753 |
| 982.471 | 376 | 992.6515 | 471 | 775.4363 | 432 | 760.0509 | 475 | ZBTB33 | None | Liver | ENCSR516I | 20261 | 63756 |
| 1037.022 | 432 | 1050.241 | 535 | 804.931 | 470 | 781.8285 | 502 | ZBTB33 | None | Liver | ENCSR516I | 20235 | 63757 |
| 328.8603 | 100 | 462.3079 | 132 | 205.2578 | 114 | 135.8357 | 102 | ZNF414 | HEK293 | Embryonic | ENCSR517( | 6633 | 63765 |
| 245.7511 | 75 | 257.9988 | 85 | 101.6177 | 71 | 122.5191 | 83 | ZNF414 | HEK293 | Embryonic | ENCSR517( | 5033 | 63766 |
| 723.555 | 284 | 803.9291 | 398 | 635.9565 | 413 | 621.1656 | 452 | NR2F6 | HepG2 | Liver | ENCSR518\ | 29885 | 63771 |
| 596.0667 | 257 | 738.9401 | 332 | 484.5906 | 353 | 607.234 | 427 | NR2F6 | HepG2 | Liver | ENCSR518\ | 29560 | 63772 |
| 60.13497 | 21 | 79.0988 | 27 | 35.45152 | 24 | 207.3848 | 62 | ZNF264 | HEK293 | Embryonic | ENCSR519I | 1798 | 63773 |
| 315.8659 | 61 | 239.6192 | 57 | 140.6628 | 55 | 227.1028 | 75 | ZNF264 | HEK293 | Embryonic | ENCSR519I | 2446 | 63774 |
| 904.2992 | 143 | 1132.805 | 203 | 375.7177 | 119 | 455.5256 | 140 | RNF2 | HepG2 | Liver | ENCSR523\ | 3873 | 63791 |
| 1088.563 | 184 | 1132.63 | 252 | 595.481 | 172 | 596.2673 | 183 | RNF2 | HepG2 | Liver | ENCSR523\ | 5412 | 63792 |
| 847.7221 | 272 | 808.9645 | 300 | 523.8955 | 300 | 458.1739 | 305 | RAD51 | K562 | Bone Marr | ENCSR524I | 19551 | 63793 |
| 524.0278 | 141 | 421.2778 | 139 | 276.7194 | 148 | 254.8491 | 153 | RAD51 | K562 | Bone Marr | ENCSR524I | 8825 | 63794 |
| 389.0774 | 79 | 450.2636 | 94 | 177.9078 | 73 | 114.3051 | 63 | ZBTB40 | HepG2 | Liver | ENCSR525Y | 3378 | 63799 |
| 466.256 | 104 | 680.3277 | 141 | 194.408 | 94 | 179.7574 | 97 | ZBTB40 | HepG2 | Liver | ENCSR525Y | 5111 | 63800 |
| 149.6966 | 243 | 239.6452 | 327 | 138.8075 | 398 | 141.7943 | 440 | HLF | HepG2 | Liver | ENCSR528I | 75068 | 63808 |
| 157.4405 | 244 | 243.9044 | 325 | 159.6509 | 408 | 139.6224 | 432 | HLF | HepG2 | Liver | ENCSR528I | 73331 | 63809 |
| 951.0115 | 443 | 979.2668 | 573 | 705.0325 | 456 | 660.4644 | 461 | L3MBTL2 | K562 | Bone Marr | ENCSR530\ | 24160 | 63813 |
| 925.479 | 548 | 956.2041 | 699 | 697.4216 | 610 | 631.6083 | 596 | L3MBTL2 | K562 | Bone Marr | ENCSR530\ | 36274 | 63814 |
| 1148.006 | 178 | 1137.964 | 215 | 678.2564 | 168 | 705.7714 | 182 | GTF2E2 | K562 | Bone Marr | ENCSR532I | 4462 | 63822 |
| 1139.533 | 333 | 1163.65 | 432 | 861.4997 | 313 | 841.9421 | 334 | ETS1 | None | None | ENCSR534\ | 10256 | 63823 |
| 1080.001 | 379 | 1096.496 | 480 | 824.9074 | 380 | 771.4239 | 374 | ETS1 | None | None | ENCSR534\ | 14713 | 63824 |
| 1110.42 | 496 | 1107.153 | 595 | 836.4623 | 481 | 785.736 | 481 | GLIS2 | HEK293 | Embryonic | ENCSR535I | 16786 | 63825 |
| 1091.448 | 390 | 1120.98 | 502 | 834.9504 | 382 | 799.3396 | 396 | GLIS2 | HEK293 | Embryonic | ENCSR535I | 13023 | 63826 |
| 47.54932 | 26 | 25.24142 | 22 | 29.64672 | 33 | 133.8544 | 69 | FOSL2 | A549 | Lung | ENCSR535I | 3644 | 63827 |
| 284.0792 | 114 | 188.7157 | 105 | 130.964 | 122 | 276.2454 | 180 | FOSL2 | A549 | Lung | ENCSR535I | 11617 | 63828 |
| 154.5852 | 95 | 163.984 | 110 | 144.0811 | 143 | 308.0103 | 211 | FOSL2 | A549 | Lung | ENCSR535I | 14335 | 63829 |
| 97.88468 | 53 | 146.8343 | 72 | 133.4321 | 92 | 155.9821 | 106 | ZBTB33 | HEK293 | Embryonic | ENCSR541I | 6732 | 63843 |
| 80.99791 | 53 | 122.4601 | 72 | 110.4699 | 93 | 123.1569 | 105 | ZBTB33 | HEK293 | Embryonic | ENCSR541I | 7934 | 63844 |
| 1083.984 | 245 | 1097.875 | 311 | 881.1862 | 279 | 865.6617 | 304 | BCL3 | A549 | Lung | ENCSR544( | 8542 | 63854 |
| 1172.004 | 285 | 1173.088 | 350 | 926.8904 | 288 | 916.7683 | 315 | BCL3 | A549 | Lung | ENCSR544( | 7537 | 63855 |
| 90.39956 | 52 | 153.6455 | 75 | 119.2128 | 90 | 212.7715 | 125 | FOSL2 | MCF-7 | Breast | ENCSR546I | 7312 | 63858 |
| 110.2231 | 67 | 146.8556 | 86 | 144.491 | 116 | 253.0166 | 159 | FOSL2 | MCF-7 | Breast | ENCSR546I | 9967 | 63859 |
| 624.4521 | 216 | 805.9219 | 283 | 359.7539 | 248 | 371.4692 | 271 | CTCF | None | Stomach | ENCSR549\ | 18911 | 63868 |
| 441.4062 | 69 | 374.9276 | 68 | 195.9156 | 59 | 180.1562 | 60 | KLF1 | K562 | Bone Marr | ENCSR550I | 1864 | 63869 |
| 629.2995 | 101 | 794.6312 | 126 | 348.846 | 98 | 225.8344 | 83 | KLF1 | K562 | Bone Marr | ENCSR550I | 2829 | 63870 |
| 794.1164 | 230 | 835.4446 | 274 | 362.9045 | 228 | 370.9691 | 248 | MLLT1 | GM12878 | Blood | ENCSR552\ | 14608 | 63875 |

| | | | | | | | | | | | | | |
|---|---|---|---|---|---|---|---|---|---|---|---|---|---|
| 840.7745 | 255 | 848.5312 | 318 | 370.6418 | 248 | 402.517 | 277 | MLLT1 | GM12878 | Blood | ENCSR552X | 16750 | 63876 |
| 5.366262 | 12 | 6.484178 | 15 | 15.71116 | 30 | 55.82578 | 53 | NFE2 | K562 | Bone Marr | ENCSR552Y | 4461 | 63877 |
| 50.31776 | 115 | 81.14728 | 155 | 70.20451 | 214 | 47.49154 | 212 | NFE2 | K562 | Bone Marr | ENCSR552Y | 38109 | 63878 |
| 1188.221 | 455 | 1220.36 | 581 | 916.419 | 443 | 903.1328 | 490 | BCL3 | A549 | Lung | ENCSR555A | 12970 | 63883 |
| 1196.287 | 605 | 1206.084 | 733 | 881.7795 | 568 | 866.4019 | 622 | BCL3 | A549 | Lung | ENCSR555A | 19556 | 63884 |
| 1170.866 | 510 | 1191.746 | 644 | 882.8458 | 495 | 875.275 | 551 | BCL3 | A549 | Lung | ENCSR555A | 16165 | 63885 |
| 23.948 | 16 | 9.296964 | 12 | 12.3367 | 19 | 41.90462 | 34 | NFE2L1 | K562 | Bone Marr | ENCSR557I | 2403 | 63894 |
| 29.4255 | 22 | 7.01096 | 14 | 12.88781 | 25 | 18.33091 | 31 | NFE2L1 | K562 | Bone Marr | ENCSR557I | 3470 | 63895 |
| 306.3802 | 78 | 472.3022 | 109 | 188.7291 | 87 | 155.7964 | 85 | CTCF | None | Colon | ENCSR558I | 4551 | 63899 |
| 1068.754 | 423 | 1074.215 | 520 | 837.5184 | 451 | 803.0569 | 465 | ZXDB | HEK293 | Embryonic | ENCSR559I | 15491 | 63902 |
| 1053.416 | 411 | 1063.395 | 504 | 827.0579 | 439 | 789.6703 | 454 | ZXDB | HEK293 | Embryonic | ENCSR559I | 15668 | 63903 |
| 0.265631 | 9 | -0.004562 | 7 | 0.057654 | 16 | 0.01446 | 17 | ZNF438 | HEK293 | Embryonic | ENCSR559S | 7268 | 63904 |
| 1.454906 | 13 | 1.237174 | 15 | 1.230576 | 24 | -0.034271 | 14 | ZNF438 | HEK293 | Embryonic | ENCSR559S | 7498 | 63905 |
| 120.4775 | 49 | 100.1953 | 50 | 99.42505 | 67 | 229.7153 | 106 | EP300 | A549 | Lung | ENCSR561Z | 4982 | 63909 |
| 181.8692 | 84 | 173.4945 | 92 | 211.3989 | 135 | 392.8311 | 192 | EP300 | A549 | Lung | ENCSR561Z | 9618 | 63910 |
| 90.18118 | 34 | 85.99837 | 37 | 73.98183 | 46 | 260.9341 | 91 | EP300 | A549 | Lung | ENCSR561Z | 3224 | 63911 |
| 90.79094 | 67 | 82.62853 | 73 | 79.02764 | 100 | 210.8235 | 160 | FOSL2 | A549 | Lung | ENCSR563C | 11597 | 63917 |
| 73.36217 | 46 | 44.79161 | 42 | 51.03286 | 62 | 189.9827 | 116 | FOSL2 | A549 | Lung | ENCSR563C | 7075 | 63918 |
| 146.1621 | 103 | 159.5774 | 121 | 112.2412 | 146 | 287.5702 | 228 | FOSL2 | A549 | Lung | ENCSR563C | 17333 | 63919 |
| 171.5036 | 44 | 120.3143 | 40 | 52.87042 | 35 | 85.44998 | 47 | HDGF | K562 | Bone Marr | ENCSR563Y | 2454 | 63920 |
| 142.4055 | 41 | 107.5054 | 39 | 58.32459 | 38 | 40.96335 | 35 | HDGF | K562 | Bone Marr | ENCSR563Y | 2590 | 63921 |
| 1095.08 | 100 | 982.9608 | 100 | 686.1528 | 96 | 446.7649 | 78 | YBX3 | K562 | Bone Marr | ENCSR567J | 1101 | 63935 |
| 1050.706 | 111 | 1044.494 | 119 | 599.788 | 103 | 642.9624 | 113 | YBX3 | K562 | Bone Marr | ENCSR567J | 1608 | 63936 |
| 438.3387 | 128 | 476.7843 | 148 | 188.2967 | 125 | 193.3317 | 136 | SSRP1 | HepG2 | Liver | ENCSR571I | 9188 | 63942 |
| 657.6426 | 208 | 728.6593 | 243 | 370.8209 | 234 | 388.6624 | 257 | SSRP1 | HepG2 | Liver | ENCSR571I | 16872 | 63943 |
| 153.4609 | 33 | 123.6038 | 32 | 124.0286 | 41 | 94.94101 | 38 | ZNF641 | HEK293 | Embryonic | ENCSR578I | 1502 | 63951 |
| 127.9591 | 29 | 174.007 | 38 | 90.83423 | 34 | 75.47743 | 33 | ZNF641 | HEK293 | Embryonic | ENCSR578I | 1455 | 63952 |
| 1019.772 | 301 | 1005.952 | 357 | 773.9073 | 307 | 728.8763 | 305 | ZNF197 | K562 | Bone Marr | ENCSR580I | 13517 | 63956 |
| 1006.374 | 286 | 998.7695 | 343 | 749.877 | 283 | 706.3671 | 294 | ZNF197 | K562 | Bone Marr | ENCSR580I | 13067 | 63957 |
| 310.6221 | 216 | 353.7211 | 257 | 233.6226 | 298 | 289.5547 | 350 | SMARCA4 | K562 | Bone Marr | ENCSR587C | 33002 | 63970 |
| 186.1333 | 126 | 153.0926 | 131 | 133.3411 | 170 | 218.1022 | 222 | SMARCA4 | K562 | Bone Marr | ENCSR587C | 18462 | 63971 |
| 734.6596 | 157 | 761.6487 | 176 | 548.4758 | 190 | 461.5861 | 186 | RUNX1 | K562 | Bone Marr | ENCSR588A | 7857 | 63972 |
| 749.0766 | 143 | 865.236 | 170 | 580.8375 | 174 | 538.883 | 178 | RUNX1 | K562 | Bone Marr | ENCSR588A | 6145 | 63973 |
| 1093.027 | 191 | 1109.688 | 246 | 800.488 | 207 | 768.4874 | 215 | GATA4 | HepG2 | Liver | ENCSR590C | 6174 | 63974 |
| 1051.29 | 383 | 1097.04 | 519 | 842.1662 | 437 | 818.8255 | 470 | GATA4 | HepG2 | Liver | ENCSR590C | 16410 | 63975 |
| 6.378527 | 16 | 2.993836 | 15 | 0.007905 | 11 | 2.143325 | 23 | ZNF512 | K562 | Bone Marr | ENCSR591C | 4964 | 63976 |
| 6.60236 | 16 | 4.96515 | 17 | 1.08283 | 18 | 6.299795 | 30 | ZNF512 | K562 | Bone Marr | ENCSR591C | 5269 | 63977 |
| 370.4747 | 59 | 234.4798 | 49 | 244.066 | 63 | 194.0776 | 59 | NBN | MCF-7 | Breast | ENCSR591E | 1707 | 63978 |
| 241.6906 | 49 | 198.9572 | 48 | 205.2042 | 62 | 188.4679 | 63 | NBN | MCF-7 | Breast | ENCSR591E | 2031 | 63979 |
| 2.70109 | 10 | 0.454667 | 8 | 0.009517 | 9 | 0.379401 | 14 | ZFP64 | HEK293 | Embryonic | ENCSR592I | 4725 | 63981 |
| 3.812747 | 7 | 1.587902 | 6 | 1.27772 | 9 | 8.73276 | 18 | ZFP64 | HEK293 | Embryonic | ENCSR592I | 2520 | 63982 |
| 88.51325 | 66 | 83.3475 | 73 | 87.96646 | 104 | 196.1801 | 155 | FOSL2 | A549 | Lung | ENCSR593I | 11700 | 63983 |
| 71.16103 | 56 | 86.30322 | 69 | 96.4006 | 100 | 258.3695 | 163 | FOSL2 | A549 | Lung | ENCSR593I | 10317 | 63984 |
| 864.1578 | 226 | 901.6944 | 293 | 519.0792 | 253 | 517.1948 | 271 | ZNF589 | K562 | Bone Marr | ENCSR603X | 14369 | 64008 |
| 958.1816 | 247 | 930.8356 | 280 | 572.4969 | 247 | 529.3656 | 255 | ZNF589 | K562 | Bone Marr | ENCSR603X | 12675 | 64009 |
| 997.0357 | 182 | 1018.926 | 220 | 744.7837 | 213 | 504.8269 | 186 | KLF7 | HEK293 | Embryonic | ENCSR604V | 6601 | 64010 |
| 235.2529 | 54 | 292.8659 | 67 | 283.0675 | 83 | 156.4031 | 65 | KLF7 | HEK293 | Embryonic | ENCSR604V | 2576 | 64011 |
| 298.5545 | 129 | 321.0429 | 149 | 221.6543 | 168 | 257.7021 | 193 | SCRT1 | HEK293 | Embryonic | ENCSR605I | 13149 | 64012 |
| 245.2102 | 80 | 220.4563 | 84 | 179.6337 | 100 | 193.492 | 111 | SCRT1 | HEK293 | Embryonic | ENCSR605I | 6103 | 64013 |
| 173.5221 | 82 | 190.2892 | 96 | 81.60685 | 91 | 251.6032 | 157 | CEBPB | A549 | Lung | ENCSR606Z | 9811 | 64016 |
| 237.0363 | 103 | 281.0202 | 125 | 146.2833 | 126 | 347.7353 | 197 | CEBPB | A549 | Lung | ENCSR606Z | 11470 | 64017 |
| 1000.522 | 144 | 1073.506 | 163 | 436.8745 | 122 | 389.721 | 122 | KLF13 | K562 | Bone Marr | ENCSR608I | 3659 | 64021 |
| 1039.088 | 167 | 1071.363 | 199 | 555.9345 | 161 | 435.7425 | 151 | KLF13 | K562 | Bone Marr | ENCSR608I | 5032 | 64022 |
| 120.2869 | 57 | 116.8533 | 63 | 126.5847 | 88 | 321.7193 | 145 | EP300 | A549 | Lung | ENCSR610I | 6665 | 64028 |
| 140.8686 | 63 | 107.2588 | 62 | 128.8455 | 91 | 310.815 | 146 | EP300 | A549 | Lung | ENCSR610I | 7003 | 64029 |
| 198.4603 | 81 | 145.6291 | 78 | 215.446 | 125 | 420.3289 | 183 | EP300 | A549 | Lung | ENCSR610I | 8188 | 64030 |
| 1110.918 | 484 | 1111.089 | 583 | 838.2615 | 484 | 807.1462 | 508 | HBP1 | HepG2 | Liver | ENCSR616V | 18928 | 64038 |
| 1100.613 | 374 | 1136.38 | 498 | 867.0959 | 398 | 804.2346 | 384 | HBP1 | HepG2 | Liver | ENCSR616V | 13572 | 64039 |
| 859.8819 | 239 | 898.2024 | 320 | 533.8001 | 272 | 480.9348 | 279 | CTCF | HEK293 | Embryonic | ENCSR617I | 15755 | 64040 |
| 882.1151 | 362 | 910.5883 | 473 | 672.6636 | 409 | 644.8827 | 429 | CTCF | HEK293 | Embryonic | ENCSR617I | 24823 | 64041 |
| 248.4888 | 41 | 133.4659 | 31 | 60.48157 | 26 | 128.1622 | 41 | CTCF | None | Stomach | ENCSR618C | 1294 | 64044 |
| 305.0001 | 72 | 369.0502 | 88 | 215.5448 | 85 | 262.0831 | 100 | HINFP | K562 | Bone Marr | ENCSR619C | 3825 | 64045 |
| 357.8234 | 82 | 553.05 | 115 | 277.0084 | 101 | 337.6643 | 119 | HINFP | K562 | Bone Marr | ENCSR619C | 4195 | 64046 |
| 700.8222 | 399 | 747.4834 | 539 | 499.0948 | 503 | 500.9704 | 548 | CTCF | None | None | ENCSR619I | 56089 | 64047 |
| 685.7447 | 436 | 745.7551 | 608 | 535.5869 | 595 | 495.9594 | 617 | CTCF | None | None | ENCSR619I | 67614 | 64048 |
| 257.5987 | 186 | 329.8383 | 233 | 180.5457 | 255 | 175.5543 | 275 | CEBPG | K562 | Bone Marr | ENCSR620V | 31684 | 64052 |
| 147.951 | 151 | 267.8213 | 217 | 120.2178 | 227 | 128.189 | 253 | CEBPG | K562 | Bone Marr | ENCSR620V | 33424 | 64053 |
| 864.5807 | 454 | 879.4607 | 568 | 674.286 | 540 | 652.2597 | 583 | ZBTB17 | HEK293 | Embryonic | ENCSR631V | 33034 | 64073 |
| 884.1454 | 423 | 883.8973 | 508 | 682.0632 | 480 | 670.1983 | 531 | ZBTB17 | HEK293 | Embryonic | ENCSR631V | 27447 | 64074 |
| 405.0984 | 227 | 384.3979 | 249 | 244.4298 | 279 | 202.7847 | 284 | ATF3 | K562 | Bone Marr | ENCSR632I | 27382 | 64076 |
| 378.9745 | 215 | 396.0151 | 246 | 269.2991 | 283 | 231.69 | 291 | ATF3 | K562 | Bone Marr | ENCSR632I | 27338 | 64077 |
| 17.57548 | 15 | 7.397775 | 12 | 19.31159 | 25 | 15.49491 | 25 | NFE2L1 | K562 | Bone Marr | ENCSR632S | 2961 | 64080 |

| | | | | | | | | | | | | | |
|---|---|---|---|---|---|---|---|---|---|---|---|---|---|
| 19.46477 | 30 | 25.17525 | 38 | 38.40631 | 63 | 35.8596 | 67 | NFE2L1 | K562 | Bone Marr | ENCSR632S | 8237 | 64081 |
| 1266.431 | 195 | 1258.659 | 231 | 772.3058 | 161 | 714.8783 | 163 | PBX2 | K562 | Bone Marr | ENCSR633E | 3627 | 64082 |
| 1227.983 | 205 | 1218.668 | 247 | 839.7022 | 182 | 705.3753 | 175 | PBX2 | K562 | Bone Marr | ENCSR633E | 4341 | 64083 |
| 473.6129 | 67 | 795.1703 | 100 | 254.4767 | 63 | 256.401 | 67 | UBTF | HeLa-S3 | Cervix | ENCSR634Z | 1658 | 64087 |
| 243.1124 | 58 | 279.7658 | 69 | 117.7744 | 57 | 67.01861 | 47 | TEAD2 | K562 | Bone Marr | ENCSR635C | 3173 | 64089 |
| 144.644 | 39 | 189.935 | 50 | 89.80487 | 44 | 25.83855 | 27 | TEAD2 | K562 | Bone Marr | ENCSR635C | 2469 | 64090 |
| 1051.992 | 375 | 1077.564 | 481 | 822.9059 | 401 | 797.6972 | 427 | KLF8 | HEK293 | Embryonic | ENCSR635M | 15143 | 64091 |
| 1111.853 | 488 | 1111.948 | 588 | 847.7298 | 486 | 827.2097 | 525 | KLF8 | HEK293 | Embryonic | ENCSR635M | 16491 | 64092 |
| 937.9656 | 281 | 922.9853 | 333 | 803.2532 | 382 | 768.3763 | 396 | CTBP1 | MCF-7 | Breast | ENCSR636E | 15182 | 64093 |
| 980.8608 | 364 | 999.7319 | 459 | 822.1877 | 466 | 813.6655 | 520 | CTBP1 | MCF-7 | Breast | ENCSR636E | 17926 | 64094 |
| 1105.003 | 422 | 1123.272 | 531 | 847.3255 | 428 | 846.4589 | 486 | BCL3 | A549 | Lung | ENCSR639H | 15640 | 64101 |
| 1196.097 | 549 | 1197.965 | 672 | 903.9209 | 538 | 878.8124 | 577 | BCL3 | A549 | Lung | ENCSR639H | 17290 | 64102 |
| 1163.979 | 413 | 1144.911 | 486 | 881.7049 | 395 | 885.6064 | 449 | BCL3 | A549 | Lung | ENCSR639H | 12436 | 64103 |
| 1184.188 | 208 | 1156.162 | 236 | 809.1219 | 191 | 590.3287 | 171 | KDM4B | K562 | Bone Marr | ENCSR642V | 5073 | 64106 |
| 1121.557 | 418 | 1114.736 | 500 | 841.7605 | 400 | 815.6832 | 421 | KDM4B | K562 | Bone Marr | ENCSR642V | 14531 | 64107 |
| 377.4334 | 184 | 310.8428 | 189 | 248.0352 | 231 | 298.6219 | 269 | SMARCA4 | K562 | Bone Marr | ENCSR643V | 20698 | 64111 |
| 956.3146 | 291 | 942.4869 | 342 | 665.2978 | 288 | 666.1622 | 309 | SMARCA4 | K562 | Bone Marr | ENCSR643V | 13792 | 64112 |
| 52.64572 | 26 | 37.82178 | 25 | 41.85383 | 36 | 152.5551 | 70 | HES2 | A549 | Lung | ENCSR646E | 3313 | 64118 |
| 97.60175 | 54 | 98.32366 | 61 | 55.1882 | 66 | 199.6611 | 122 | HES2 | A549 | Lung | ENCSR646E | 7463 | 64119 |
| 107.5318 | 67 | 146.996 | 87 | 126.2671 | 111 | 271.7821 | 166 | HES2 | A549 | Lung | ENCSR646E | 10185 | 64120 |
| 239.7681 | 88 | 207.1812 | 91 | 131.214 | 98 | 104.2984 | 96 | MEF2D | K562 | Bone Marr | ENCSR647Z | 7480 | 64127 |
| 217.4294 | 78 | 300.1072 | 102 | 161.7098 | 99 | 104.2055 | 88 | MEF2D | K562 | Bone Marr | ENCSR647Z | 6432 | 64128 |
| 116.8867 | 28 | 111.339 | 30 | 72.46988 | 31 | 19.64248 | 18 | ILK | K562 | Bone Marr | ENCSR648I | 1554 | 64129 |
| 49.52451 | 17 | 45.09558 | 18 | 31.16955 | 20 | 3.729819 | 9 | ILK | K562 | Bone Marr | ENCSR648I | 1457 | 64130 |
| 285.9105 | 94 | 436.9276 | 130 | 342.4649 | 150 | 345.3106 | 161 | HHEX | HepG2 | Liver | ENCSR656J | 7822 | 64139 |
| 301.6347 | 92 | 385.9275 | 116 | 326.2965 | 139 | 301.4232 | 143 | HHEX | HepG2 | Liver | ENCSR656J | 7056 | 64140 |
| 1023.596 | 207 | 1037.809 | 265 | 748.3279 | 227 | 689.5026 | 232 | NFRKB | K562 | Bone Marr | ENCSR657E | 7368 | 64143 |
| 1019.389 | 264 | 1018.909 | 322 | 766.4432 | 262 | 659.1178 | 260 | NFRKB | K562 | Bone Marr | ENCSR657E | 9560 | 64144 |
| 120.2576 | 43 | 92.72849 | 42 | 110.406 | 61 | 286.7639 | 104 | NR3C1 | A549 | Lung | ENCSR660F | 3878 | 64155 |
| 144.7496 | 54 | 107.8508 | 52 | 112.4724 | 71 | 299.6376 | 121 | NR3C1 | A549 | Lung | ENCSR660F | 4972 | 64156 |
| 821.911 | 374 | 857.1374 | 491 | 631.6604 | 438 | 607.2418 | 470 | CTCF | None | Breast | ENCSR661M | 30658 | 64159 |
| 957.8749 | 250 | 967.0882 | 312 | 767.5777 | 289 | 636.223 | 278 | ZNF580 | HEK293 | Embryonic | ENCSR668H | 12204 | 64173 |
| 937.1711 | 240 | 930.0562 | 287 | 763.408 | 288 | 698.7149 | 292 | ZNF580 | HEK293 | Embryonic | ENCSR668H | 12380 | 64174 |
| 614.9953 | 151 | 748.5908 | 185 | 327.0659 | 158 | 266.2986 | 154 | ATF4 | HepG2 | Liver | ENCSR669L | 8972 | 64175 |
| 477.3128 | 122 | 541.001 | 144 | 281.9516 | 135 | 279.8851 | 144 | ATF4 | HepG2 | Liver | ENCSR669L | 7648 | 64176 |
| 1037.895 | 177 | 1094.613 | 243 | 633.6339 | 185 | 538.4525 | 181 | TAF7 | K562 | Bone Marr | ENCSR671C | 5905 | 64177 |
| 1089.123 | 194 | 1099.465 | 243 | 718.9132 | 195 | 602.4284 | 189 | TAF7 | K562 | Bone Marr | ENCSR671C | 5684 | 64178 |
| 1092.654 | 280 | 1099.069 | 354 | 848.5459 | 285 | 863.7481 | 339 | BCL3 | A549 | Lung | ENCSR673C | 9806 | 64183 |
| 1128.678 | 338 | 1152.816 | 433 | 878.8039 | 341 | 887.9615 | 397 | BCL3 | A549 | Lung | ENCSR673C | 10754 | 64184 |
| 1186.137 | 239 | 1130.402 | 247 | 923.0037 | 221 | 869.8295 | 222 | MLLT1 | K562 | Bone Marr | ENCSR675L | 5423 | 64186 |
| 1104.994 | 174 | 1035.797 | 178 | 620.1334 | 167 | 521.7458 | 162 | MLLT1 | K562 | Bone Marr | ENCSR675L | 4830 | 64187 |
| 202.0223 | 48 | 190.322 | 51 | 142.1866 | 56 | 98.83889 | 50 | ZNF792 | HEK293 | Embryonic | ENCSR679S | 2401 | 64200 |
| 173.688 | 42 | 193.2961 | 49 | 141.2662 | 53 | 148.661 | 58 | ZNF792 | HEK293 | Embryonic | ENCSR679S | 2191 | 64201 |
| 27.6619 | 11 | 30.80295 | 13 | 11.78928 | 11 | 7.77927 | 10 | ZNF207 | HepG2 | Liver | ENCSR683C | 1080 | 64210 |
| 177.2792 | 86 | 148.7403 | 89 | 157.5483 | 124 | 301.8399 | 177 | HES2 | A549 | Lung | ENCSR686E | 10519 | 64216 |
| 63.60149 | 31 | 72.17092 | 37 | 53.8892 | 44 | 174.5878 | 81 | HES2 | A549 | Lung | ENCSR686E | 3871 | 64217 |
| 107.2328 | 41 | 62.03617 | 35 | 46.22736 | 42 | 177.448 | 83 | HES2 | A549 | Lung | ENCSR686E | 4001 | 64218 |
| 115.0569 | 64 | 133.2362 | 77 | 209.0092 | 127 | 352.1442 | 173 | EP300 | A549 | Lung | ENCSR686E | 8697 | 64220 |
| 89.18783 | 29 | 58.91427 | 26 | 109.2859 | 47 | 215.9472 | 71 | EP300 | A549 | Lung | ENCSR686E | 2370 | 64221 |
| 95.91829 | 62 | 95.70418 | 70 | 94.29936 | 96 | 246.203 | 155 | FOSL2 | A549 | Lung | ENCSR686H | 9769 | 64222 |
| 66.52858 | 36 | 51.3243 | 36 | 54.44133 | 51 | 183.1353 | 94 | FOSL2 | A549 | Lung | ENCSR686H | 4964 | 64223 |
| 155.531 | 57 | 91.63392 | 49 | 93.94016 | 67 | 353.4211 | 134 | NR3C1 | A549 | Lung | ENCSR686F | 5243 | 64224 |
| 210.714 | 71 | 180.9058 | 73 | 137.162 | 85 | 336.4152 | 139 | NR3C1 | A549 | Lung | ENCSR686F | 5856 | 64225 |
| 22.47816 | 64 | 17.94934 | 70 | 8.581285 | 89 | 13.10798 | 108 | ZNF146 | HEK293 | Embryonic | ENCSR689Y | 23840 | 64240 |
| 24.78268 | 73 | 28.83049 | 89 | 15.75973 | 114 | 21.39757 | 135 | ZNF146 | HEK293 | Embryonic | ENCSR689Y | 27378 | 64241 |
| 114.1876 | 59 | 169.9181 | 80 | 110.9087 | 89 | 316.5441 | 153 | HES2 | A549 | Lung | ENCSR690H | 7643 | 64242 |
| 75.38707 | 24 | 56.91324 | 23 | 54.28409 | 30 | 257.4274 | 71 | NR3C1 | A549 | Lung | ENCSR691C | 1954 | 64249 |
| 109.5362 | 36 | 89.37234 | 36 | 82.20461 | 46 | 265.7627 | 88 | NR3C1 | A549 | Lung | ENCSR691C | 2971 | 64250 |
| 118.6304 | 39 | 82.68457 | 36 | 68.37991 | 44 | 275.9328 | 93 | NR3C1 | A549 | Lung | ENCSR691C | 3175 | 64251 |
| 690.3102 | 123 | 828.1476 | 149 | 351.2728 | 117 | 509.0839 | 151 | ZNF610 | HEK293 | Embryonic | ENCSR691T | 4228 | 64255 |
| 1023.824 | 203 | 1009.081 | 239 | 557.1073 | 193 | 563.0788 | 207 | ZNF610 | HEK293 | Embryonic | ENCSR691T | 7271 | 64256 |
| 848.3766 | 273 | 863.0558 | 343 | 564.1183 | 309 | 490.3561 | 313 | CTCF | None | Spleen | ENCSR692I | 19472 | 64259 |
| 1196.83 | 276 | 1190.852 | 330 | 912.0969 | 237 | 869.9457 | 237 | HCFC1 | MCF-7 | Breast | ENCSR697C | 5818 | 64265 |
| 1196.4 | 386 | 1196.511 | 466 | 932.7828 | 373 | 870.6431 | 360 | HCFC1 | MCF-7 | Breast | ENCSR697C | 9284 | 64266 |
| 578.5037 | 98 | 696.3321 | 119 | 234.7123 | 82 | 243.4675 | 89 | ZNF592 | MCF-7 | Breast | ENCSR701A | 3137 | 64278 |
| 419.0968 | 64 | 327.4374 | 60 | 151.2775 | 49 | 167.5225 | 55 | ZNF592 | MCF-7 | Breast | ENCSR701A | 1740 | 64279 |
| 187.1855 | 71 | 196.0852 | 81 | 125.3466 | 88 | 274.7496 | 135 | CEBPB | A549 | Lung | ENCSR701T | 6848 | 64280 |
| 197.6008 | 130 | 251.5475 | 163 | 125.5123 | 171 | 231.3559 | 233 | CEBPB | A549 | Lung | ENCSR701T | 21121 | 64281 |
| 702.027 | 462 | 726.8022 | 592 | 609.9869 | 713 | 598.8627 | 795 | RAD21 | MCF-7 | Breast | ENCSR703T | 67154 | 64284 |
| 705.8328 | 429 | 713.1352 | 533 | 569.069 | 604 | 567.6025 | 687 | RAD21 | MCF-7 | Breast | ENCSR703T | 61517 | 64285 |
| 207.6651 | 49 | 237.3943 | 58 | 192.5102 | 66 | 165.1628 | 65 | ZSCAN23 | HEK293 | Embryonic | ENCSR705A | 2480 | 64288 |
| 253.8735 | 67 | 274.69 | 77 | 229.6326 | 90 | 178.7444 | 85 | ZSCAN23 | HEK293 | Embryonic | ENCSR705A | 3860 | 64289 |

| | | | | | | | | | | | | | |
|---|---|---|---|---|---|---|---|---|---|---|---|---|---|
| 1013.407 | 388 | 1037.429 | 492 | 758.3837 | 382 | 711.5957 | 385 | ZBTB11 | K562 | Bone Marr | ENCSR706I | 17563 | 64290 |
| 1009.445 | 426 | 1016.875 | 524 | 738.1781 | 413 | 714.4929 | 439 | ZBTB11 | K562 | Bone Marr | ENCSR706I | 20545 | 64291 |
| 273.5012 | 68 | 261.5958 | 73 | 142.0774 | 69 | 138.5204 | 73 | MYNN | HEK293 | Embryonic | ENCSR707I | 3655 | 64294 |
| 101.5421 | 34 | 92.8246 | 36 | 72.34107 | 42 | 90.27375 | 50 | MYNN | HEK293 | Embryonic | ENCSR707I | 2660 | 64295 |
| 1020.275 | 303 | 995.6477 | 350 | 759.2487 | 295 | 762.0779 | 341 | NFE2L2 | HeLa-S3 | Cervix | ENCSR707I | 13748 | 64298 |
| 372.684 | 98 | 465.5339 | 122 | 251.5293 | 116 | 222.0873 | 117 | NFE2L2 | HeLa-S3 | Cervix | ENCSR707I | 6357 | 64299 |
| 415.6056 | 224 | 609.3514 | 298 | 356.0155 | 321 | 389.7837 | 360 | NR2F6 | K562 | Bone Marr | ENCSR707( | 30909 | 64300 |
| 280.7055 | 116 | 359.4189 | 146 | 151.4043 | 133 | 224.6217 | 169 | NR2F6 | K562 | Bone Marr | ENCSR707( | 12161 | 64301 |
| 976.1 | 392 | 994.4691 | 497 | 738.8682 | 415 | 717.9482 | 444 | E2F5 | K562 | Bone Marr | ENCSR709I | 21156 | 64302 |
| 961.4058 | 427 | 976.9189 | 537 | 715.5338 | 449 | 675.3334 | 458 | E2F5 | K562 | Bone Marr | ENCSR709I | 24934 | 64303 |
| 111.5096 | 32 | 98.18067 | 33 | 81.30997 | 39 | 74.33734 | 40 | ZNF837 | HEK293 | Embryonic | ENCSR711I | 2048 | 64307 |
| 161.3443 | 89 | 133.734 | 92 | 103.2152 | 113 | 89.44368 | 116 | ZNF664 | HEK293 | Embryonic | ENCSR714I | 11553 | 64313 |
| 112.1126 | 62 | 124.1295 | 73 | 84.26147 | 84 | 58.5304 | 79 | ZNF664 | HEK293 | Embryonic | ENCSR714I | 8113 | 64314 |
| 1057.152 | 517 | 1077.046 | 651 | 785.3386 | 533 | 765.5907 | 575 | ETV4 | HepG2 | Liver | ENCSR714Y | 25305 | 64317 |
| 1002.492 | 312 | 1019.231 | 398 | 772.5428 | 332 | 762.8313 | 365 | ETV4 | HepG2 | Liver | ENCSR714Y | 14654 | 64318 |
| 273.0182 | 195 | 257.3778 | 215 | 238.2598 | 284 | 258.3763 | 318 | DPF2 | K562 | Bone Marr | ENCSR715( | 29947 | 64319 |
| 452.8144 | 219 | 392.1783 | 230 | 300.6176 | 275 | 305.6886 | 300 | DPF2 | K562 | Bone Marr | ENCSR715( | 24010 | 64320 |
| 189.072 | 105 | 239.2389 | 131 | 97.83513 | 122 | 60.81265 | 112 | ZNF362 | HEK293 | Embryonic | ENCSR715( | 12989 | 64321 |
| 195.8602 | 101 | 222.6924 | 120 | 119.8002 | 124 | 103.4356 | 127 | ZNF362 | HEK293 | Embryonic | ENCSR715( | 11866 | 64322 |
| 205.6553 | 82 | 155.6864 | 80 | 168.1326 | 111 | 450.8959 | 188 | NR3C1 | A549 | Lung | ENCSR720I | 8014 | 64331 |
| 153.996 | 55 | 74.79839 | 43 | 123.8274 | 73 | 339.8179 | 127 | NR3C1 | A549 | Lung | ENCSR720I | 4809 | 64332 |
| 191.77 | 101 | 147.5527 | 101 | 226.6313 | 165 | 430.5345 | 235 | NR3C1 | A549 | Lung | ENCSR720I | 13163 | 64333 |
| 977.2958 | 255 | 980.339 | 313 | 649.5806 | 259 | 634.7773 | 274 | ZNF644 | K562 | Bone Marr | ENCSR729I | 12397 | 64348 |
| 699.7151 | 180 | 907.5398 | 239 | 502.4199 | 218 | 336.0315 | 194 | ZNF644 | K562 | Bone Marr | ENCSR729I | 11069 | 64349 |
| 596.1162 | 67 | 567.7731 | 70 | 223.5029 | 49 | 272.5295 | 58 | SP1 | MCF-7 | Breast | ENCSR729I | 1080 | 64350 |
| 7.955264 | 23 | 8.036412 | 27 | 5.509764 | 37 | 3.477296 | 37 | ZNF788 | HEK293 | Embryonic | ENCSR732I | 8803 | 64359 |
| 1.151653 | 13 | 0.649345 | 14 | 0.409693 | 21 | 5.515149 | 38 | ZNF788 | HEK293 | Embryonic | ENCSR732I | 7962 | 64360 |
| 889.4716 | 359 | 879.1226 | 429 | 688.4215 | 420 | 640.3608 | 422 | MYNN | K562 | Bone Marr | ENCSR737I | 26302 | 64369 |
| 820.7727 | 231 | 868.6693 | 293 | 493.2733 | 264 | 498.9221 | 285 | MYNN | K562 | Bone Marr | ENCSR737I | 17037 | 64370 |
| 1059.573 | 192 | 1109.498 | 268 | 662.8813 | 196 | 703.8419 | 215 | ZNF740 | K562 | Bone Marr | ENCSR737I | 6695 | 64373 |
| 762.8528 | 121 | 1128.441 | 172 | 338.9056 | 106 | 477.7068 | 135 | ZNF740 | K562 | Bone Marr | ENCSR737I | 3777 | 64374 |
| 713.5106 | 341 | 688.4969 | 392 | 291.3777 | 356 | 274.8887 | 380 | TBX21 | GM12878 | Blood | ENCSR739I | 39720 | 64375 |
| 441.4384 | 202 | 432.9735 | 224 | 160.6912 | 202 | 182.756 | 230 | TBX21 | GM12878 | Blood | ENCSR739I | 22269 | 64376 |
| 922.2609 | 217 | 897.6264 | 250 | 547.1397 | 237 | 529.8126 | 250 | BACH1 | K562 | Bone Marr | ENCSR740I | 11907 | 64377 |
| 398.851 | 113 | 351.2294 | 117 | 210.3602 | 120 | 134.762 | 106 | BACH1 | K562 | Bone Marr | ENCSR740I | 7748 | 64378 |
| 579.8486 | 86 | 640.6041 | 99 | 214.0177 | 66 | 163.3064 | 61 | SUZ12 | GM12878 | Blood | ENCSR744X | 2121 | 64381 |
| 1034.973 | 157 | 1107.619 | 194 | 477.188 | 138 | 389.2787 | 132 | SUZ12 | GM12878 | Blood | ENCSR744X | 4225 | 64382 |
| 958.9324 | 364 | 980.2982 | 465 | 728.2931 | 381 | 692.4456 | 392 | NR2C2 | K562 | Bone Marr | ENCSR750I | 18630 | 64394 |
| 978.9007 | 453 | 987.0686 | 556 | 746.6766 | 485 | 693.5807 | 480 | NR2C2 | K562 | Bone Marr | ENCSR750I | 23322 | 64395 |
| 290.8329 | 128 | 311.8575 | 148 | 185.9133 | 159 | 365.4752 | 228 | CEBPB | A549 | Lung | ENCSR752A | 14330 | 64398 |
| 578.7034 | 240 | 585.862 | 270 | 347.0893 | 290 | 476.8189 | 356 | CEBPB | A549 | Lung | ENCSR752A | 26356 | 64399 |
| 590.3912 | 235 | 525.5706 | 249 | 331.655 | 275 | 469.8659 | 342 | CEBPB | A549 | Lung | ENCSR752A | 24842 | 64400 |
| 446.3536 | 146 | 636.176 | 194 | 247.3172 | 163 | 214.1822 | 165 | CTCF | None | Skin | ENCSR756I | 11978 | 64413 |
| 548.9923 | 270 | 594.2969 | 314 | 355.9346 | 345 | 317.7526 | 359 | HMBOX1 | K562 | Bone Marr | ENCSR757I | 35119 | 64414 |
| 1.545368 | 5 | 0.73661 | 5 | 3.450445 | 12 | 7.346446 | 17 | PRDM12 | HEK293 | Embryonic | ENCSR760A | 2527 | 64421 |
| 576.7202 | 221 | 686.4234 | 268 | 508.4554 | 311 | 425.6658 | 311 | KLF16 | K562 | Bone Marr | ENCSR760I | 22877 | 64424 |
| 635.2438 | 244 | 773.4675 | 317 | 500.3726 | 328 | 451.1919 | 339 | KLF16 | K562 | Bone Marr | ENCSR760I | 25732 | 64425 |
| 398.1931 | 118 | 488.1488 | 145 | 264.386 | 139 | 382.2387 | 177 | ZSCAN20 | HEK293 | Embryonic | ENCSR767I | 8006 | 64441 |
| 425.6938 | 125 | 517.932 | 153 | 241.3096 | 137 | 482.0316 | 203 | ZSCAN20 | HEK293 | Embryonic | ENCSR767I | 8488 | 64442 |
| 957.7196 | 179 | 999.2226 | 210 | 590.684 | 190 | 526.0535 | 191 | ZNF324 | HEK293 | Embryonic | ENCSR768I | 6617 | 64443 |
| 920.4166 | 164 | 1027.492 | 197 | 575.6088 | 174 | 515.4096 | 175 | ZNF324 | HEK293 | Embryonic | ENCSR768I | 5660 | 64444 |
| 455.6119 | 95 | 535.656 | 114 | 538.5311 | 144 | 401.3578 | 131 | OVOL3 | HEK293 | Embryonic | ENCSR768I | 4154 | 64445 |
| 398.9656 | 82 | 542.9537 | 107 | 443.1613 | 120 | 402.332 | 121 | OVOL3 | HEK293 | Embryonic | ENCSR768I | 3478 | 64446 |
| 79.094 | 24 | 93.06111 | 29 | 59.46699 | 30 | 79.31304 | 37 | AEBP2 | HEK293 | Embryonic | ENCSR769J | 1686 | 64454 |
| 78.04974 | 25 | 54.64892 | 23 | 49.42177 | 29 | 96.54334 | 43 | AEBP2 | HEK293 | Embryonic | ENCSR769J | 1869 | 64455 |
| 833.2652 | 392 | 874.484 | 525 | 631.5916 | 454 | 605.7368 | 484 | CTCF | None | Colon | ENCSR769V | 32764 | 64457 |
| 234.7856 | 76 | 171.6305 | 72 | 173.1188 | 96 | 418.4033 | 157 | EP300 | A549 | Lung | ENCSR770( | 6075 | 64458 |
| 174.1483 | 58 | 182.603 | 66 | 141.9201 | 77 | 298.8765 | 118 | EP300 | A549 | Lung | ENCSR770( | 4773 | 64459 |
| 108.6681 | 31 | 84.49891 | 30 | 77.67265 | 38 | 264.2833 | 76 | EP300 | A549 | Lung | ENCSR770( | 2197 | 64460 |
| 684.0299 | 115 | 681.8229 | 125 | 578.4514 | 142 | 389.6837 | 122 | BCL11B | HEK293 | Embryonic | ENCSR770I | 3609 | 64463 |
| 428.6924 | 82 | 527.587 | 101 | 409.421 | 110 | 507.2359 | 131 | BCL11B | HEK293 | Embryonic | ENCSR770I | 3171 | 64464 |
| 81.26248 | 21 | 68.36434 | 21 | 26.9263 | 17 | 32.47724 | 20 | RELA | K562 | Bone Marr | ENCSR772I | 1150 | 64465 |
| 70.50574 | 19 | 64.90918 | 20 | 28.34165 | 17 | 24.21612 | 17 | RELA | K562 | Bone Marr | ENCSR772I | 1098 | 64466 |
| 176.7372 | 64 | 120.5998 | 59 | 144.8946 | 86 | 359.9901 | 142 | NR3C1 | A549 | Lung | ENCSR773I | 5745 | 64471 |
| 207.2057 | 107 | 184.3073 | 114 | 233.674 | 171 | 500.5609 | 257 | NR3C1 | A549 | Lung | ENCSR773I | 13688 | 64472 |
| 199.3584 | 105 | 199.5442 | 118 | 268.2064 | 181 | 498.3816 | 256 | NR3C1 | A549 | Lung | ENCSR773I | 13569 | 64473 |
| 272.8877 | 66 | 255.9347 | 70 | 211.4573 | 81 | 180.8617 | 80 | ZBTB7A | HEK293 | Embryonic | ENCSR773I | 3350 | 64474 |
| 513.2998 | 97 | 790.1615 | 136 | 410.4932 | 118 | 465.6607 | 134 | ZBTB7A | HEK293 | Embryonic | ENCSR773I | 3658 | 64475 |
| 109.5649 | 65 | 83.26718 | 65 | 63.18417 | 81 | 204.3848 | 141 | FOSL2 | A549 | Lung | ENCSR775I | 9560 | 64484 |
| 128.9349 | 72 | 105.0999 | 74 | 74.83803 | 89 | 205.0487 | 145 | FOSL2 | A549 | Lung | ENCSR775I | 9965 | 64485 |
| 51.91511 | 24 | 34.00434 | 22 | 25.72699 | 27 | 128.4944 | 60 | FOSL2 | A549 | Lung | ENCSR775I | 2908 | 64486 |
| 4.979992 | 17 | 1.672665 | 15 | 0.180136 | 17 | 1.814934 | 27 | ZNF781 | HEK293 | Embryonic | ENCSR777Y | 7123 | 64495 |

| | | | | | | | | | | | | | |
|---|---|---|---|---|---|---|---|---|---|---|---|---|---|
| 2.148581 | 12 | 0.055907 | 8 | 0.311775 | 16 | 29.69681 | 52 | ZNF781 | HEK293 | Embryonic | ENCSR777 | 6242 | 64496 |
| 34.29278 | 14 | 13.89777 | 10 | 9.979862 | 12 | 21.19647 | 18 | TSC22D4 | K562 | Bone Marr | ENCSR787 | 1395 | 64519 |
| 16.67116 | 9 | 5.473829 | 6 | 25.72993 | 17 | 24.77765 | 18 | TSC22D4 | K562 | Bone Marr | ENCSR787 | 1259 | 64520 |
| 477.6464 | 140 | 585.2946 | 172 | 282.8309 | 157 | 262.3451 | 163 | RFX1 | MCF-7 | Breast | ENCSR788 | 9675 | 64525 |
| 414.1636 | 133 | 510.6996 | 164 | 212.2974 | 142 | 239.4298 | 161 | RFX1 | MCF-7 | Breast | ENCSR788 | 10217 | 64526 |
| 130.772 | 63 | 124.4831 | 69 | 167.0115 | 106 | 373.2906 | 165 | EP300 | A549 | Lung | ENCSR792 | 7459 | 64534 |
| 130.0001 | 50 | 95.89821 | 48 | 125.7201 | 73 | 328.1643 | 124 | EP300 | A549 | Lung | ENCSR792 | 4788 | 64535 |
| 241.4684 | 91 | 239.2354 | 101 | 207.1177 | 126 | 192.3626 | 131 | JUNB | K562 | Bone Marr | ENCSR795 | 8816 | 64537 |
| 238.4217 | 57 | 275.2864 | 68 | 215.8898 | 77 | 152.7919 | 69 | JUNB | K562 | Bone Marr | ENCSR795 | 3236 | 64538 |
| 217.3603 | 82 | 215.4924 | 91 | 192.7783 | 115 | 338.6899 | 160 | HES2 | A549 | Lung | ENCSR796 | 7820 | 64542 |
| 5.544906 | 9 | 2.661741 | 8 | 2.390758 | 12 | 2.866721 | 14 | YBX1 | HepG2 | Liver | ENCSR799 | 2623 | 64543 |
| 9.801843 | 9 | 7.171295 | 9 | 3.567718 | 10 | 3.529855 | 11 | YBX1 | HepG2 | Liver | ENCSR799 | 1840 | 64544 |
| 924.2789 | 285 | 899.1433 | 324 | 682.8221 | 309 | 632.3133 | 320 | HNF1A | HepG2 | Liver | ENCSR800 | 17208 | 64546 |
| 459.9199 | 138 | 403.1559 | 143 | 290.9211 | 161 | 269.4766 | 167 | HNF1A | HepG2 | Liver | ENCSR800 | 10193 | 64547 |
| 944.6341 | 195 | 937.5291 | 215 | 536.6341 | 201 | 502.2982 | 208 | HIC1 | HEK293 | Embryonic | ENCSR803 | 8013 | 64552 |
| 776.5429 | 154 | 918.7538 | 185 | 465.3246 | 162 | 434.5823 | 167 | HIC1 | HEK293 | Embryonic | ENCSR803 | 6063 | 64553 |
| 926.7884 | 621 | 951.5744 | 777 | 688.3792 | 704 | 650.9578 | 740 | IKZF2 | HEK293 | Embryonic | ENCSR803J | 48647 | 64555 |
| 920.4645 | 699 | 931.0929 | 853 | 674.0426 | 804 | 637.4845 | 849 | IKZF2 | HEK293 | Embryonic | ENCSR803J | 62245 | 64556 |
| 1151.754 | 570 | 1200.613 | 754 | 860.9108 | 555 | 840.8894 | 605 | SP2 | HEK293 | Embryonic | ENCSR807 | 19456 | 64559 |
| 1162.509 | 574 | 1199.928 | 741 | 862.4281 | 551 | 843.4688 | 594 | SP2 | HEK293 | Embryonic | ENCSR807 | 18890 | 64560 |
| 359.6778 | 190 | 448.1628 | 235 | 314.945 | 271 | 271.4313 | 277 | BCOR | K562 | Bone Marr | ENCSR808 | 23771 | 64563 |
| 295.0376 | 154 | 307.6943 | 176 | 215.0483 | 204 | 222.9821 | 224 | BCOR | K562 | Bone Marr | ENCSR808 | 18910 | 64564 |
| 69.46283 | 128 | 66.89663 | 146 | 67.8838 | 212 | 54.74134 | 220 | MAFG | K562 | Bone Marr | ENCSR818 | 37724 | 64575 |
| 71.01107 | 119 | 54.68884 | 126 | 59.68887 | 187 | 51.23514 | 197 | MAFG | K562 | Bone Marr | ENCSR818 | 32874 | 64576 |
| 1078.907 | 505 | 1101.148 | 645 | 806.4979 | 516 | 754.331 | 514 | CBX1 | HepG2 | Liver | ENCSR819 | 22823 | 64581 |
| 983.4801 | 573 | 996.3242 | 710 | 769.1536 | 686 | 752.3559 | 753 | CBX1 | HepG2 | Liver | ENCSR819 | 38577 | 64582 |
| 120.5674 | 30 | 147.9097 | 37 | 35.4406 | 23 | 82.11666 | 37 | RNF2 | K562 | Bone Marr | ENCSR820 | 1606 | 64583 |
| 140.4339 | 48 | 155.0313 | 56 | 75.1786 | 52 | 55.15984 | 49 | RNF2 | K562 | Bone Marr | ENCSR820 | 3714 | 64584 |
| 694.4653 | 253 | 690.4857 | 281 | 696.8503 | 405 | 665.15 | 423 | FEZF1 | HEK293 | Embryonic | ENCSR827 | 23095 | 64604 |
| 555.2299 | 190 | 616.1332 | 222 | 689.1071 | 304 | 701.4131 | 338 | FEZF1 | HEK293 | Embryonic | ENCSR827 | 15562 | 64605 |
| 799.4945 | 447 | 804.6695 | 547 | 585.7987 | 502 | 568.2875 | 550 | CTCF | None | Colon | ENCSR833 | 41795 | 64614 |
| 462.0797 | 94 | 572.5195 | 116 | 268.7667 | 97 | 163.2117 | 81 | HDAC8 | K562 | Bone Marr | ENCSR835 | 3671 | 64617 |
| 215.4504 | 51 | 370.0396 | 76 | 197.777 | 68 | 81.96438 | 47 | HDAC8 | K562 | Bone Marr | ENCSR835 | 2490 | 64618 |
| 646.8729 | 249 | 710.1203 | 290 | 296.1091 | 265 | 275.2563 | 279 | TRIM22 | GM12878 | Blood | ENCSR835 | 24124 | 64619 |
| 561.2505 | 236 | 625.0258 | 277 | 224.3401 | 242 | 263.8524 | 279 | TRIM22 | GM12878 | Blood | ENCSR835 | 24777 | 64620 |
| 1152.725 | 591 | 1187.755 | 765 | 865.2374 | 602 | 806.724 | 590 | NRF1 | K562 | Bone Marr | ENCSR837 | 22827 | 64621 |
| 1150.012 | 523 | 1182 | 668 | 865.3362 | 517 | 817.7951 | 520 | NRF1 | K562 | Bone Marr | ENCSR837 | 18664 | 64622 |
| 628.031 | 537 | 643.1754 | 671 | 458.6114 | 712 | 378.2868 | 737 | SMARCA4 | None | None | ENCSR839 | 107329 | 64633 |
| 428.3354 | 300 | 536.7665 | 372 | 260.5061 | 397 | 190.6817 | 393 | SMARCA4 | None | None | ENCSR839 | 55388 | 64634 |
| 266.9682 | 113 | 227.0443 | 117 | 325.2332 | 184 | 549.3132 | 251 | EP300 | A549 | Lung | ENCSR841 | 11961 | 64638 |
| 148.1077 | 51 | 97.64146 | 46 | 85.48831 | 58 | 279.6179 | 109 | EP300 | A549 | Lung | ENCSR841 | 4324 | 64639 |
| 234.0347 | 92 | 186.6334 | 92 | 169.1909 | 118 | 321.5359 | 169 | EP300 | A549 | Lung | ENCSR841 | 8968 | 64640 |
| 1052.346 | 424 | 1051.234 | 507 | 772.9372 | 400 | 731.0786 | 405 | ZNF639 | K562 | Bone Marr | ENCSR845 | 17823 | 64648 |
| 1041.709 | 436 | 1028.226 | 514 | 767.3736 | 425 | 710.1749 | 413 | ZNF639 | K562 | Bone Marr | ENCSR845 | 19472 | 64649 |
| 1004.341 | 279 | 991.2752 | 328 | 763.0101 | 282 | 703.3249 | 289 | FOXJ2 | K562 | Bone Marr | ENCSR847 | 12553 | 64653 |
| 962.2947 | 240 | 964.8541 | 299 | 742.3304 | 274 | 733.0564 | 291 | FOXJ2 | K562 | Bone Marr | ENCSR847 | 12212 | 64654 |
| 530.2728 | 144 | 621.2553 | 173 | 411.5549 | 183 | 351.8549 | 182 | ZMYM3 | HepG2 | Liver | ENCSR848 | 9709 | 64656 |
| 210.5301 | 88 | 245.6542 | 106 | 204.0002 | 130 | 166.4215 | 128 | ZMYM3 | HepG2 | Liver | ENCSR848 | 9439 | 64657 |
| 1164.653 | 200 | 1195.159 | 267 | 745.3006 | 183 | 788.7725 | 200 | PBX2 | HepG2 | Liver | ENCSR849 | 5056 | 64660 |
| 1145.651 | 207 | 1153.897 | 264 | 758.9708 | 195 | 691.8257 | 197 | PBX2 | HepG2 | Liver | ENCSR849 | 5749 | 64661 |
| 1163.137 | 538 | 1190.177 | 688 | 880.1465 | 533 | 821.1397 | 522 | NRF1 | HepG2 | Liver | ENCSR853 | 18394 | 64667 |
| 1078.184 | 309 | 1131.295 | 429 | 822.3538 | 303 | 816.685 | 340 | NRF1 | HepG2 | Liver | ENCSR853 | 11143 | 64668 |
| 374.6368 | 163 | 363.9634 | 180 | 177.7602 | 180 | 358.0711 | 258 | CEBPB | A549 | Lung | ENCSR853 | 18297 | 64669 |
| 203.05 | 95 | 235.0598 | 114 | 125.3025 | 117 | 265.4219 | 173 | CEBPB | A549 | Lung | ENCSR853 | 11086 | 64670 |
| 274.0708 | 118 | 283.1372 | 134 | 131.538 | 130 | 301.6099 | 198 | CEBPB | A549 | Lung | ENCSR853 | 12879 | 64671 |
| 40.22424 | 23 | 22.71639 | 20 | 19.17544 | 26 | 20.46944 | 29 | ZNF707 | HEK293 | Embryonic | ENCSR854 | 3080 | 64673 |
| 76.28437 | 31 | 49.62058 | 28 | 59.94152 | 41 | 68.99801 | 47 | ZNF707 | HEK293 | Embryonic | ENCSR854 | 2948 | 64674 |
| 42.88495 | 35 | 36.47651 | 37 | 38.82778 | 52 | 34.2227 | 54 | IRF1 | K562 | Bone Marr | ENCSR854 | 5486 | 64675 |
| 54.27679 | 29 | 45.70411 | 30 | 61.76032 | 46 | 41.61641 | 42 | IRF1 | K562 | Bone Marr | ENCSR854 | 3354 | 64676 |
| 937.9959 | 327 | 935.9987 | 398 | 717.9462 | 353 | 701.8985 | 385 | KLF1 | HEK293 | Embryonic | ENCSR859 | 17687 | 64685 |
| 940.8155 | 332 | 953.2807 | 420 | 732.4251 | 371 | 705.3131 | 393 | KLF1 | HEK293 | Embryonic | ENCSR859 | 18061 | 64686 |
| 972.9271 | 318 | 1010.186 | 429 | 725.2737 | 323 | 704.6928 | 349 | CBFB | GM12878 | Blood | ENCSR860 | 16958 | 64691 |
| 997.7064 | 392 | 1014.256 | 490 | 740.9513 | 395 | 699.2781 | 399 | CBFB | GM12878 | Blood | ENCSR860 | 19925 | 64692 |
| 865.2587 | 241 | 862.8094 | 290 | 676.4129 | 299 | 574.9833 | 298 | ZNF382 | HEK293 | Embryonic | ENCSR862 | 14333 | 64693 |
| 840.5757 | 238 | 832.4006 | 275 | 579.1516 | 283 | 537.3228 | 294 | ZNF382 | HEK293 | Embryonic | ENCSR862 | 15026 | 64694 |
| 645.3887 | 128 | 1032.625 | 185 | 504.273 | 157 | 360.8232 | 141 | TCF7 | K562 | Bone Marr | ENCSR863 | 5765 | 64696 |
| 777.6769 | 176 | 910.6802 | 211 | 576.8902 | 214 | 366.3293 | 184 | TCF7 | K562 | Bone Marr | ENCSR863 | 9537 | 64697 |
| 19.43859 | 10 | 7.195652 | 7 | 5.724596 | 9 | 13.49524 | 14 | YBX1 | MCF-7 | Breast | ENCSR864 | 1235 | 64704 |
| 10.96744 | 12 | 0.930953 | 6 | 5.266545 | 15 | 9.285057 | 20 | YBX1 | MCF-7 | Breast | ENCSR864 | 2588 | 64705 |
| 171.8695 | 51 | 224.4523 | 65 | 176.5611 | 74 | 167.3004 | 77 | ZSCAN16 | HEK293 | Embryonic | ENCSR864 | 3420 | 64706 |
| 166.5132 | 46 | 206.3075 | 57 | 117.4228 | 55 | 157.8442 | 68 | ZSCAN16 | HEK293 | Embryonic | ENCSR864 | 2836 | 64707 |
| 167.5941 | 55 | 122.4723 | 52 | 233.9059 | 94 | 204.6025 | 94 | ZNF800 | HEK293 | Embryonic | ENCSR866 | 4261 | 64713 |

| | | | | | | | | | | | | | |
|---|---|---|---|---|---|---|---|---|---|---|---|---|---|
| 100.5775 | 35 | 72.81689 | 33 | 194.4343 | 71 | 196.1259 | 76 | ZNF800 | HEK293 | Embryonic | ENCSR866V | 2910 | 64714 |
| 688.2814 | 256 | 756.9376 | 306 | 386.0437 | 289 | 336.8055 | 296 | IKZF1 | GM12878 | Blood | ENCSR874A | 21371 | 64734 |
| 673.4364 | 400 | 691.0281 | 505 | 444.3929 | 496 | 401.8529 | 523 | IKZF1 | GM12878 | Blood | ENCSR874A | 51478 | 64735 |
| 81.67593 | 142 | 80.81148 | 163 | 51.8866 | 207 | 85.98055 | 261 | ZBTB33 | K562 | Bone Marr | ENCSR876C | 39510 | 64738 |
| 107.4576 | 131 | 111.9071 | 152 | 66.63907 | 182 | 76.67629 | 208 | ZBTB33 | K562 | Bone Marr | ENCSR876C | 29937 | 64739 |
| 898.2387 | 274 | 860.9946 | 303 | 648.6103 | 307 | 679.8752 | 338 | ZKSCAN1 | K562 | Bone Marr | ENCSR882E | 17985 | 64745 |
| 938.3364 | 238 | 916.1899 | 275 | 590.2286 | 253 | 483.6794 | 247 | ZKSCAN1 | K562 | Bone Marr | ENCSR882E | 13264 | 64746 |
| 994.3105 | 318 | 1021.366 | 409 | 732.238 | 302 | 705.1392 | 317 | ZBTB11 | HEK293 | Embryonic | ENCSR882Z | 14325 | 64750 |
| 1051.012 | 365 | 1075.516 | 470 | 785.2283 | 348 | 739.01 | 347 | ZBTB11 | HEK293 | Embryonic | ENCSR882Z | 13887 | 64751 |
| 235.4321 | 46 | 411.6663 | 70 | 142.6243 | 49 | 255.1965 | 71 | TCF7L2 | K562 | Bone Marr | ENCSR888X | 1977 | 64758 |
| 331.0926 | 57 | 473.5942 | 77 | 173.1776 | 55 | 269.1687 | 74 | TCF7L2 | K562 | Bone Marr | ENCSR888X | 2023 | 64759 |
| 497.6346 | 75 | 501.3616 | 82 | 259.8452 | 71 | 346.108 | 88 | SRSF3 | HepG2 | Liver | ENCSR893L | 2230 | 64770 |
| 490.5172 | 72 | 476.9659 | 77 | 329.5845 | 78 | 333.0819 | 83 | SRSF3 | HepG2 | Liver | ENCSR893L | 2052 | 64771 |
| 1139.577 | 397 | 1154.798 | 500 | 871.7606 | 375 | 858.464 | 413 | SUPT5H | K562 | Bone Marr | ENCSR894C | 11106 | 64772 |
| 1125.497 | 406 | 1139.885 | 505 | 858.6777 | 381 | 838.1039 | 409 | SUPT5H | K562 | Bone Marr | ENCSR894C | 11623 | 64773 |
| 152.391 | 66 | 154.2154 | 74 | 127.9201 | 90 | 85.80117 | 82 | SMARCA5 | K562 | Bone Marr | ENCSR895H | 6408 | 64776 |
| 91.97754 | 34 | 93.03161 | 38 | 97.49845 | 51 | 43.28924 | 38 | SMARCA5 | K562 | Bone Marr | ENCSR895H | 2866 | 64777 |
| 259.8385 | 118 | 385.0376 | 159 | 202.442 | 158 | 154.8133 | 153 | REST | HEK293 | Embryonic | ENCSR896U | 12728 | 64778 |
| 418.4489 | 139 | 485.289 | 166 | 314.6512 | 176 | 186.7892 | 151 | REST | HEK293 | Embryonic | ENCSR896U | 10726 | 64779 |
| 37.04759 | 88 | 69.42551 | 125 | 21.66785 | 132 | 20.64389 | 144 | JUNB | GM12878 | Blood | ENCSR897M | 30402 | 64780 |
| 141.7949 | 148 | 201.2045 | 193 | 53.117 | 176 | 71.24111 | 210 | JUNB | GM12878 | Blood | ENCSR897M | 32723 | 64781 |
| 1057.039 | 262 | 1065.86 | 324 | 850.9847 | 286 | 805.7008 | 286 | ZNF687 | MCF-7 | Breast | ENCSR899E | 8949 | 64786 |
| 1055.333 | 249 | 1045.342 | 290 | 772.1744 | 242 | 796.42 | 262 | ZNF687 | MCF-7 | Breast | ENCSR899E | 8324 | 64787 |
| 70.28358 | 46 | 55.62745 | 47 | 48.18788 | 62 | 182.0473 | 116 | FOSL2 | A549 | Lung | ENCSR900F | 7326 | 64788 |
| 160.947 | 97 | 124.8222 | 98 | 106.2662 | 127 | 288.3374 | 205 | FOSL2 | A549 | Lung | ENCSR900F | 14093 | 64789 |
| 111.3932 | 69 | 70.38885 | 64 | 84.76071 | 96 | 199.6558 | 147 | FOSL2 | A549 | Lung | ENCSR900F | 10413 | 64790 |
| 1089.573 | 429 | 1129.332 | 562 | 848.964 | 456 | 809.4797 | 466 | CREM | HepG2 | Liver | ENCSR903E | 16516 | 64797 |
| 1002.102 | 480 | 1018.164 | 605 | 766.4898 | 527 | 742.0138 | 566 | CREM | HepG2 | Liver | ENCSR903E | 25812 | 64798 |
| 1207.34 | 457 | 1207.75 | 550 | 918.3688 | 431 | 905.2249 | 474 | BCL3 | A549 | Lung | ENCSR904C | 12195 | 64800 |
| 1029.149 | 163 | 1078.898 | 200 | 689.7155 | 179 | 869.5066 | 215 | BCL3 | A549 | Lung | ENCSR904C | 5311 | 64801 |
| 1199.383 | 572 | 1202.9 | 686 | 903.1396 | 556 | 888.3905 | 608 | BCL3 | A549 | Lung | ENCSR904C | 18010 | 64802 |
| 304.6618 | 58 | 386.9188 | 73 | 169.1702 | 59 | 214.3449 | 71 | ZMIZ1 | K562 | Bone Marr | ENCSR907J | 2444 | 64817 |
| 511.9752 | 220 | 512.8681 | 246 | 298.8131 | 261 | 283.9679 | 277 | TRIM24 | K562 | Bone Marr | ENCSR907M | 23094 | 64819 |
| 429.4838 | 179 | 412.6254 | 196 | 255.6743 | 212 | 175.5246 | 198 | TRIM24 | K562 | Bone Marr | ENCSR907M | 17734 | 64820 |
| 299.29 | 151 | 250.2726 | 156 | 159.3025 | 174 | 161.9831 | 190 | KDM1A | K562 | Bone Marr | ENCSR908C | 16590 | 64822 |
| 193.0747 | 127 | 257.6376 | 162 | 180.5659 | 189 | 178.4171 | 204 | KDM1A | K562 | Bone Marr | ENCSR908C | 18286 | 64823 |
| 434.4642 | 94 | 537.6392 | 116 | 256.2345 | 99 | 330.2446 | 120 | ZEB1 | HEK293 | Embryonic | ENCSR909H | 4104 | 64828 |
| 449.8548 | 91 | 745.2657 | 133 | 263.9129 | 95 | 367.8252 | 120 | ZEB1 | HEK293 | Embryonic | ENCSR909H | 3756 | 64829 |
| 86.81701 | 31 | 61.81297 | 29 | 66.81787 | 40 | 163.1725 | 66 | ZNF547 | HEK293 | Embryonic | ENCSR909T | 2590 | 64830 |
| 108.0249 | 32 | 111.5037 | 36 | 98.68524 | 44 | 81.97738 | 43 | ZNF547 | HEK293 | Embryonic | ENCSR909T | 2207 | 64831 |
| 102.7531 | 39 | 109.9102 | 45 | 84.49418 | 53 | 161.4462 | 77 | HES2 | A549 | Lung | ENCSR911H | 3779 | 64835 |
| 44.97663 | 15 | 41.471 | 16 | 9.810338 | 11 | 118.2841 | 39 | HES2 | A549 | Lung | ENCSR911H | 1335 | 64836 |
| 84.05896 | 39 | 77.27013 | 42 | 41.90377 | 44 | 155.3133 | 84 | HES2 | A549 | Lung | ENCSR911H | 4622 | 64837 |
| 96.32723 | 26 | 74.87175 | 25 | 63.85003 | 30 | 48.01578 | 28 | NONO | MCF-7 | Breast | ENCSR912M | 1567 | 64839 |
| 163.6299 | 32 | 288.6202 | 49 | 142.6517 | 41 | 99.74813 | 36 | NONO | MCF-7 | Breast | ENCSR912M | 1289 | 64840 |
| 370.9931 | 59 | 500.5404 | 77 | 281.4784 | 69 | 220.9446 | 64 | JUNB | None | None | ENCSR917M | 1880 | 64855 |
| 809.3167 | 657 | 851.54 | 847 | 625.4542 | 855 | 620.4766 | 970 | RAD21 | None | Liver | ENCSR917C | 81934 | 64856 |
| 833.5182 | 614 | 867.1338 | 787 | 632.5858 | 760 | 617.5644 | 845 | RAD21 | None | Liver | ENCSR917C | 66829 | 64857 |
| 865.04 | 500 | 880.1601 | 629 | 631.6217 | 558 | 610.9162 | 605 | EGR2 | HEK293 | Embryonic | ENCSR919C | 43352 | 64860 |
| 902.9528 | 434 | 921.2469 | 548 | 666.3721 | 465 | 663.5886 | 526 | EGR2 | HEK293 | Embryonic | ENCSR919C | 30322 | 64861 |
| 202.6486 | 96 | 165.3821 | 98 | 177.693 | 137 | 385.7684 | 207 | NR3C1 | A549 | Lung | ENCSR919C | 11368 | 64862 |
| 232.7215 | 101 | 153.14 | 93 | 233.5788 | 151 | 414.9282 | 210 | NR3C1 | A549 | Lung | ENCSR919C | 10808 | 64863 |
| 185.903 | 74 | 152.5172 | 75 | 168.252 | 105 | 372.9712 | 163 | NR3C1 | A549 | Lung | ENCSR919C | 7299 | 64864 |
| 1206.874 | 551 | 1250.448 | 722 | 889.3053 | 503 | 859.0486 | 529 | ZFX | K562 | Bone Marr | ENCSR920A | 16017 | 64865 |
| 1182.061 | 589 | 1230.66 | 763 | 875.0687 | 556 | 835.751 | 573 | ZFX | K562 | Bone Marr | ENCSR920A | 19054 | 64866 |
| 1178.66 | 345 | 1217.222 | 455 | 921.8334 | 341 | 896.9428 | 361 | SIN3A | K562 | Bone Marr | ENCSR920E | 9096 | 64867 |
| 1227.407 | 235 | 1251.88 | 306 | 969.9142 | 225 | 950.3178 | 238 | SIN3A | K562 | Bone Marr | ENCSR920E | 4945 | 64868 |
| 110.1659 | 43 | 84.35971 | 42 | 59.03781 | 48 | 44.37413 | 46 | IRF9 | K562 | Bone Marr | ENCSR926H | 4018 | 64874 |
| 72.10797 | 33 | 93.73958 | 42 | 73.85034 | 50 | 46.22109 | 44 | IRF9 | K562 | Bone Marr | ENCSR926H | 3616 | 64875 |
| 322.5285 | 59 | 260.9985 | 57 | 228.0975 | 67 | 159.3475 | 59 | ZBTB1 | HEK293 | Embryonic | ENCSR927U | 2155 | 64878 |
| 262.4807 | 59 | 208.5167 | 57 | 208.2259 | 73 | 276.3574 | 90 | ZBTB1 | HEK293 | Embryonic | ENCSR927U | 2831 | 64879 |
| 819.998 | 193 | 907.0772 | 228 | 400.3308 | 191 | 359.9733 | 195 | NCOA1 | K562 | Bone Marr | ENCSR931H | 10149 | 64887 |
| 655.1356 | 154 | 652.6765 | 169 | 338.2587 | 156 | 312.6046 | 161 | NCOA1 | K562 | Bone Marr | ENCSR931H | 8031 | 64888 |
| 775.0307 | 109 | 1109.09 | 147 | 395.3078 | 100 | 368.3212 | 102 | ZNF143 | GM12878 | Blood | ENCSR936X | 2714 | 64894 |
| 1034.657 | 524 | 1055.859 | 656 | 758.1752 | 518 | 715.6323 | 527 | ZNF143 | GM12878 | Blood | ENCSR936X | 25243 | 64895 |
| 1241.808 | 233 | 1227.919 | 273 | 1049.514 | 274 | 1029.736 | 289 | MBD2 | MCF-7 | Breast | ENCSR940M | 4803 | 64900 |
| 1151.103 | 525 | 1166.849 | 651 | 928.4658 | 603 | 918.0521 | 670 | MBD2 | MCF-7 | Breast | ENCSR940M | 18935 | 64901 |
| 1074.595 | 214 | 1070.411 | 260 | 823.0505 | 229 | 869.7478 | 273 | BCL3 | A549 | Lung | ENCSR948J | 7462 | 64912 |
| 1197.49 | 671 | 1202.562 | 817 | 880.5022 | 655 | 862.5128 | 719 | BCL3 | A549 | Lung | ENCSR948J | 23790 | 64913 |
| 1165.077 | 581 | 1181.04 | 727 | 855.3728 | 554 | 853.9494 | 625 | BCL3 | A549 | Lung | ENCSR948J | 20445 | 64914 |
| 974.0636 | 406 | 966.0949 | 485 | 725.4922 | 424 | 680.1402 | 428 | CBX1 | K562 | Bone Marr | ENCSR948C | 23523 | 64916 |
| 1109.747 | 383 | 1134.087 | 487 | 837.3443 | 364 | 808.4147 | 385 | CBX1 | K562 | Bone Marr | ENCSR948C | 13397 | 64917 |

| | | | | | | | | | | | | | |
|---|---|---|---|---|---|---|---|---|---|---|---|---|---|
| 197.9026 | 76 | 281.6909 | 101 | 140.6897 | 96 | 176.1125 | 114 | IKZF1 | K562 | Bone Marr | ENCSR948\ | 7020 | 64918 |
| 770.736 | 315 | 771.9709 | 388 | 448.7015 | 354 | 587.4537 | 428 | IKZF1 | K562 | Bone Marr | ENCSR948\ | 29283 | 64919 |
| 84.13601 | 42 | 91.17071 | 49 | 63.94691 | 57 | 196.3624 | 101 | HES2 | A549 | Lung | ENCSR951S | 5309 | 64927 |
| 66.99662 | 34 | 73.72278 | 40 | 55.4089 | 48 | 176.5074 | 87 | HES2 | A549 | Lung | ENCSR951S | 4414 | 64928 |
| 38.82819 | 16 | 43.51094 | 19 | 31.59881 | 22 | 162.9155 | 53 | HES2 | A549 | Lung | ENCSR951S | 1777 | 64929 |
| 114.44 | 43 | 100.9151 | 45 | 89.74958 | 57 | 315.8304 | 112 | NR3C1 | A549 | Lung | ENCSR952( | 4070 | 64930 |
| 109.1654 | 54 | 81.68388 | 53 | 85.94824 | 74 | 316.5726 | 143 | NR3C1 | A549 | Lung | ENCSR952( | 6610 | 64931 |
| 109.3666 | 37 | 89.06028 | 37 | 63.40791 | 42 | 287.2605 | 94 | NR3C1 | A549 | Lung | ENCSR952( | 3092 | 64932 |
| 837.7169 | 358 | 839.9851 | 438 | 618.717 | 399 | 607.3305 | 441 | BHLHE40 | IMR90 | Lung | ENCSR957I | 31212 | 64945 |
| 909.1433 | 518 | 959.8784 | 690 | 686.3137 | 598 | 661.2041 | 643 | BHLHE40 | IMR90 | Lung | ENCSR957I | 41751 | 64946 |
| 73.90118 | 63 | 55.94741 | 64 | 69.33517 | 97 | 121.2619 | 130 | TRIM24 | K562 | Bone Marr | ENCSR957l | 11443 | 64947 |
| 118.7709 | 80 | 91.82537 | 81 | 113.0643 | 121 | 112.8643 | 131 | TRIM24 | K562 | Bone Marr | ENCSR957l | 12069 | 64948 |
| 111.4758 | 77 | 94.21419 | 81 | 32.80195 | 77 | 24.44928 | 77 | ATF2 | GM12878 | Blood | ENCSR961I | 11894 | 64957 |
| 430.3112 | 181 | 477.2702 | 212 | 110.5761 | 156 | 126.4697 | 178 | ATF2 | GM12878 | Blood | ENCSR961I | 18273 | 64958 |
| 120.58 | 52 | 119.9057 | 58 | 55.45636 | 56 | 213.8205 | 109 | HES2 | A549 | Lung | ENCSR964I | 5660 | 64961 |
| 119.6016 | 51 | 87.10696 | 49 | 71.16305 | 61 | 164.7095 | 95 | HES2 | A549 | Lung | ENCSR964I | 5496 | 64962 |
| 105.1169 | 41 | 101.6727 | 45 | 35.23132 | 38 | 178.0184 | 84 | HES2 | A549 | Lung | ENCSR964I | 4078 | 64963 |
| 871.1037 | 251 | 854.8528 | 295 | 735.5666 | 338 | 694.0888 | 342 | WT1 | HEK293 | Embryonic | ENCSR966I | 16280 | 64977 |
| 483.5036 | 167 | 592.9455 | 205 | 450.7583 | 234 | 477.8999 | 258 | WT1 | HEK293 | Embryonic | ENCSR966I | 13473 | 64978 |
| 1061.092 | 468 | 1064.861 | 571 | 813.2044 | 484 | 782.7314 | 511 | PATZ1 | HEK293 | Embryonic | ENCSR966l | 18532 | 64981 |
| 1064.553 | 435 | 1067.322 | 527 | 819.8927 | 443 | 781.3603 | 459 | PATZ1 | HEK293 | Embryonic | ENCSR966l | 16247 | 64982 |
| 699.702 | 197 | 727.4298 | 222 | 307.1622 | 192 | 291.2904 | 202 | RFX1 | K562 | Bone Marr | ENCSR968( | 12804 | 64987 |
| 814.6444 | 256 | 820.14 | 315 | 362.8736 | 259 | 327.155 | 268 | RFX1 | K562 | Bone Marr | ENCSR968( | 19722 | 64988 |
| 688.4758 | 263 | 758.9711 | 323 | 421.3681 | 313 | 471.3838 | 354 | NR2F1 | K562 | Bone Marr | ENCSR970I | 24709 | 64996 |
| 709.1871 | 298 | 734.5503 | 367 | 479.7873 | 375 | 588.8254 | 450 | NR2F1 | K562 | Bone Marr | ENCSR970I | 31242 | 64997 |
| 1069.615 | 390 | 1080.921 | 483 | 816.6929 | 393 | 770.4998 | 394 | ELF1 | K562 | Bone Marr | ENCSR975S | 15156 | 65013 |
| 1076.1 | 414 | 1078.002 | 505 | 810.1727 | 409 | 763.01 | 408 | ELF1 | K562 | Bone Marr | ENCSR975S | 15812 | 65014 |
| 295.9705 | 109 | 424.2047 | 145 | 268.7512 | 152 | 455.0985 | 208 | ZNF18 | HEK293 | Embryonic | ENCSR977I | 9322 | 65018 |
| 371.9864 | 129 | 493.8448 | 165 | 322.5647 | 175 | 334.4928 | 191 | ZNF18 | HEK293 | Embryonic | ENCSR977I | 10369 | 65019 |
| 25.39151 | 13 | 34.18987 | 17 | 17.1479 | 17 | 111.5018 | 44 | HES2 | A549 | Lung | ENCSR979I | 1827 | 65022 |
| 43.90182 | 24 | 58.38559 | 31 | 46.20676 | 38 | 120.0755 | 63 | HES2 | A549 | Lung | ENCSR979I | 3399 | 65023 |
| 792.6659 | 158 | 776.8862 | 171 | 593.3538 | 187 | 546.261 | 191 | ZNF24 | HEK293 | Embryonic | ENCSR984I | 6443 | 65031 |
| 944.6071 | 204 | 919.308 | 227 | 711.1417 | 245 | 632.6581 | 247 | ZNF24 | HEK293 | Embryonic | ENCSR984I | 9388 | 65032 |
| 108.8317 | 26 | 91.88282 | 26 | 65.12877 | 28 | 69.83405 | 31 | DNMT1 | K562 | Bone Marr | ENCSR987I | 1355 | 65040 |
| 173.1472 | 34 | 179.8005 | 38 | 90.10818 | 33 | 65.82097 | 30 | DNMT1 | K562 | Bone Marr | ENCSR987I | 1350 | 65041 |
| 113.2607 | 28 | 134.7788 | 34 | 84.03605 | 34 | 45.87675 | 27 | MCM3 | K562 | Bone Marr | ENCSR990/ | 1578 | 65048 |
| 78.53605 | 23 | 70.47246 | 24 | 37.31605 | 23 | 60.04995 | 31 | MCM3 | K562 | Bone Marr | ENCSR990/ | 1586 | 65049 |
| 1029.871 | 432 | 1049.183 | 543 | 782.5382 | 446 | 756.9665 | 479 | TGIF2 | HepG2 | Liver | ENCSR993l | 20275 | 65056 |
| 1034.993 | 324 | 1062.715 | 428 | 813.0191 | 355 | 782.8365 | 371 | TGIF2 | HepG2 | Liver | ENCSR993l | 13904 | 65057 |
| 84.13088 | 22 | 122.1494 | 30 | 85.45947 | 32 | 62.01713 | 29 | ZNF329 | HEK293 | Embryonic | GSM20268 | 1446 | 65060 |
| 821.7375 | 276 | 825.9444 | 343 | 667.322 | 365 | 662.2496 | 411 | ZNF554 | HEK293 | Embryonic | GSM20268 | 24194 | 65061 |
| 919.7952 | 384 | 973.6986 | 532 | 707.1838 | 442 | 685.1577 | 473 | ZNF467 | HEK293 | Embryonic | GSM20268 | 26234 | 65062 |
| 1095.476 | 396 | 1107.655 | 498 | 850.1114 | 418 | 822.5202 | 439 | ZNF574 | HEK293 | Embryonic | GSM20268 | 15165 | 65063 |
| 15.99986 | 9 | 12.33736 | 9 | 13.00304 | 13 | 75.55155 | 32 | ZNF547 | HEK293 | Embryonic | GSM20268 | 1450 | 65067 |
| 0.178452 | 14 | 0.064371 | 16 | 0.206696 | 30 | 1.100959 | 41 | ZNF490 | HEK293 | Embryonic | GSM20268 | 13236 | 65069 |
| 1174.62 | 253 | 1196.826 | 331 | 903.0126 | 235 | 884.078 | 251 | KLF12 | HEK293 | Embryonic | GSM20269 | 6595 | 65071 |
| 1110.753 | 230 | 1155.409 | 317 | 869.7913 | 232 | 847.2508 | 243 | KLF15 | HEK293 | Embryonic | GSM20268 | 7218 | 65072 |
| 108.8012 | 66 | 210.8077 | 101 | 46.30445 | 74 | 55.76622 | 86 | ZNF280A | HEK293 | Embryonic | GSM20267 | 10010 | 65073 |
| 21.7567 | 27 | 27.44148 | 34 | 4.874526 | 29 | 22.87539 | 50 | ZNF382 | HEK293 | Embryonic | GSM20268 | 7086 | 65074 |
| 532.5519 | 207 | 527.6803 | 230 | 263.7173 | 228 | 200.962 | 222 | ZNF382 | HEK293 | Embryonic | GSM20268 | 20658 | 65075 |
| 316.3383 | 112 | 392.8854 | 139 | 141.1239 | 116 | 258.4947 | 162 | ZNF382 | HEK293 | Embryonic | GSM20268 | 10304 | 65076 |
| 80.2587 | 76 | 98.30661 | 94 | 54.10613 | 107 | 69.72529 | 127 | ZNF382 | HEK293 | Embryonic | GSM20268 | 16130 | 65077 |
| 15.35969 | 32 | 20.42622 | 41 | 8.93664 | 48 | 15.37598 | 61 | ZNF317 | HEK293 | Embryonic | GSM20268 | 11143 | 65078 |
| 28.19329 | 78 | 25.02253 | 88 | 28.45441 | 137 | 46.09287 | 171 | ZNF317 | HEK293 | Embryonic | GSM20268 | 30100 | 65079 |
| 4.485729 | 29 | 2.781489 | 31 | 12.01527 | 68 | 23.22219 | 89 | ZIM3 | HEK293 | Embryonic | GSM20268 | 15898 | 65081 |
| 434.0499 | 97 | 584.9843 | 126 | 380.3934 | 128 | 507.4127 | 158 | ZNF778 | HEK293 | Embryonic | GSM20268 | 5103 | 65082 |
| 1248.395 | 781 | 1338.04 | 1041 | 877.2226 | 727 | 857.1173 | 791 | SP2 | HEK293 | Embryonic | GSM20267 | 28642 | 65084 |
| 101.9974 | 40 | 127.4858 | 50 | 71.57924 | 51 | 146.3714 | 76 | ZNF134 | HEK293 | Embryonic | GSM20267 | 4092 | 65085 |
| 25.44625 | 135 | 16.65479 | 145 | 13.55837 | 217 | 32.36991 | 284 | ZSCAN16 | HEK293 | Embryonic | GSM20268 | 66115 | 65086 |
| 46.97851 | 20 | 82.75103 | 30 | 18.06439 | 20 | 101.2872 | 47 | ZNF264 | HEK293 | Embryonic | GSM20267 | 2311 | 65087 |
| 808.267 | 268 | 862.6847 | 383 | 477.6549 | 309 | 484.0033 | 335 | CTCF | HEK293 | Embryonic | GSM20267 | 23022 | 65090 |
| 12.49315 | 8 | 29.72261 | 14 | 20.51267 | 16 | 126.5774 | 42 | ZNF41 | HEK293 | Embryonic | GSM20267 | 1452 | 65091 |
| 707.5204 | 244 | 846.2383 | 352 | 507.5659 | 309 | 385.8038 | 297 | CTCF | HEK293 | Embryonic | GSM20267 | 22017 | 65093 |
| 898.5738 | 447 | 934.9924 | 585 | 679.2077 | 511 | 684.9108 | 595 | CTCF | HEK293 | Embryonic | GSM20267 | 35334 | 65094 |
| 206.7041 | 79 | 355.8312 | 116 | 211.4037 | 119 | 173.9259 | 117 | CTCF | HEK293 | Embryonic | GSM20267 | 7883 | 65095 |
| 347.6465 | 78 | 401.536 | 93 | 184.1131 | 80 | 300.4726 | 109 | CTCF | HEK293 | Embryonic | GSM20267 | 4165 | 65096 |
| 361.7859 | 117 | 601.0654 | 169 | 277.836 | 152 | 308.7654 | 171 | CTCF | HEK293 | Embryonic | GSM20267 | 9940 | 65097 |
| 111.7949 | 24 | 265.0481 | 44 | 100.069 | 32 | 129.1249 | 39 | ZFP42 | HEK293 | Embryonic | GSM20268 | 1229 | 65098 |
| 111.5638 | 22 | 143.3829 | 28 | 64.92806 | 23 | 189.4499 | 44 | ZSCAN31 | HEK293 | Embryonic | GSM20268 | 1001 | 65102 |
| 0.305688 | 151 | 0.074635 | 175 | 1.079479 | 338 | 2.24926 | 401 | ZNF384 | HEK293 | Embryonic | GSM20268 | 153703 | 65106 |
| 650.6053 | 230 | 784.153 | 286 | 693.8723 | 369 | 679.4739 | 401 | OSR2 | HEK293 | Embryonic | GSM20268 | 22215 | 65108 |
| 1195.374 | 554 | 1238.171 | 724 | 883.9571 | 522 | 851.0131 | 549 | SP4 | HEK293 | Embryonic | GSM20267 | 17869 | 65110 |

| | | | | | | | | | | | | |
|---|---|---|---|---|---|---|---|---|---|---|---|---|
| 1196.548 | 584 | 1234.933 | 751 | 885.6798 | 555 | 847.3654 | 569 | SP4 | HEK293 | Embryonic GSM20267 | 19060 | 65111 |
| 314.9779 | 74 | 332.7013 | 84 | 263.6348 | 96 | 360.7058 | 120 | SP4 | HEK293 | Embryonic GSM20267 | 4165 | 65112 |
| 41.36509 | 25 | 90.18245 | 41 | 16.14156 | 27 | 79.93798 | 56 | ZNF140 | HEK293 | Embryonic GSM20267 | 3869 | 65113 |
| 112.6362 | 120 | 181.4589 | 165 | 143.1965 | 212 | 121.0408 | 218 | ZNF121 | HEK293 | Embryonic GSM20267 | 28036 | 65114 |
| 1143.405 | 675 | 1233.511 | 923 | 846.0763 | 697 | 827.8876 | 763 | MAZ | HEK293 | Embryonic GSM20267 | 30371 | 65115 |
| 821.9474 | 294 | 865.8304 | 403 | 729.9409 | 460 | 693.3308 | 472 | FEZF1 | HEK293 | Embryonic GSM20267 | 25679 | 65116 |
| 102.3106 | 30 | 124.5604 | 37 | 108.1878 | 45 | 52.60518 | 34 | ZNF768 | HEK293 | Embryonic GSM20268 | 2254 | 65118 |
| 646.006 | 91 | 839.8037 | 116 | 293.7179 | 79 | 308.5188 | 86 | PATZ1 | HEK293 | Embryonic GSM20268 | 2430 | 65119 |
| 1166.509 | 425 | 1224.182 | 584 | 898.87 | 420 | 874.4299 | 443 | ZBTB26 | HEK293 | Embryonic GSM20268 | 12917 | 65120 |
| 210.0135 | 43 | 208.5576 | 47 | 147.5217 | 50 | 267.4522 | 73 | ZBTB26 | HEK293 | Embryonic GSM20268 | 2015 | 65121 |
| 1059.067 | 405 | 1102.991 | 541 | 820.3624 | 432 | 790.6735 | 454 | ZFP64 | HEK293 | Embryonic GSM20268 | 17369 | 65122 |
| 1054.308 | 575 | 1109.908 | 768 | 780.1041 | 596 | 766.772 | 658 | ZFP64 | HEK293 | Embryonic GSM20268 | 29264 | 65123 |
| 94.35717 | 28 | 184.9607 | 45 | 46.49834 | 29 | 149.2331 | 55 | MYNN | HEK293 | Embryonic GSM20268 | 2125 | 65124 |
| 201.6318 | 130 | 180.1656 | 140 | 217.0358 | 207 | 284.1036 | 249 | PRDM6 | HEK293 | Embryonic GSM20268 | 20157 | 65125 |
| 341.3093 | 62 | 500.9476 | 85 | 192.1889 | 63 | 383.8358 | 97 | ZNF281 | HEK293 | Embryonic GSM20268 | 2429 | 65127 |
| 18.11177 | 18 | 28.56086 | 25 | 28.35979 | 35 | 48.39285 | 47 | ZNF436 | HEK293 | Embryonic GSM20268 | 4100 | 65128 |
| 963.0623 | 331 | 1024.347 | 471 | 759.2171 | 380 | 746.0882 | 419 | ZNF341 | HEK293 | Embryonic GSM20268 | 18519 | 65129 |
| 516.5297 | 79 | 764.4236 | 109 | 212.7736 | 66 | 266.4416 | 79 | ZNF350 | HEK293 | Embryonic GSM20268 | 2396 | 65131 |
| 195.0621 | 84 | 250.0015 | 106 | 132.3703 | 107 | 142.6534 | 119 | PRDM1 | HEK293 | Embryonic GSM20267 | 9402 | 65133 |
| 5.077589 | 7 | 6.44429 | 9 | 10.46572 | 16 | 25.74428 | 25 | SCRT1 | HEK293 | Embryonic GSM20268 | 2315 | 65135 |
| 269.8928 | 41 | 304.1323 | 48 | 142.9288 | 39 | 187.326 | 48 | ZBTB18 | HEK293 | Embryonic GSM20268 | 1233 | 65137 |
| 173.3683 | 160 | 190.018 | 189 | 110.2939 | 221 | 120.1503 | 248 | ZNF382 | HEK293 | Embryonic GSM20268 | 34128 | 65138 |
| 851.8295 | 260 | 851.3344 | 318 | 699.5902 | 341 | 721.2121 | 413 | ZBTB42 | HEK293 | Embryonic GSM20267 | 19846 | 65139 |
| 67.56094 | 40 | 99.488 | 54 | 75.23434 | 65 | 99.84773 | 79 | ZNF320 | HEK293 | Embryonic GSM20267 | 5955 | 65141 |
| 50.45458 | 27 | 107.3489 | 44 | 54.07378 | 43 | 119.0689 | 66 | ZNF320 | HEK293 | Embryonic GSM20267 | 3753 | 65142 |
| 449.9641 | 111 | 653.1777 | 150 | 321.9887 | 134 | 303.0603 | 139 | ZNF324 | HEK293 | Embryonic GSM20267 | 6660 | 65144 |
| 1167.255 | 431 | 1227.827 | 597 | 876.7514 | 406 | 853.4835 | 431 | ZBTB14 | HEK293 | Embryonic GSM20267 | 13160 | 65146 |
| 1031.644 | 732 | 1097.801 | 962 | 735.7287 | 779 | 706.4616 | 836 | EGR2 | HEK293 | Embryonic GSM20267 | 52497 | 65147 |
| 1135.712 | 558 | 1174.407 | 716 | 832.6332 | 529 | 809.1567 | 571 | SP1 | HEK293 | Embryonic GSM20267 | 21728 | 65151 |
| 1200.797 | 417 | 1198.174 | 503 | 893.3724 | 371 | 888.825 | 418 | ZSCAN22 | HEK293 | Embryonic GSM20267 | 11462 | 65153 |
| 1183.498 | 271 | 1187.53 | 332 | 909.8473 | 252 | 870.7256 | 255 | ZSCAN22 | HEK293 | Embryonic GSM20267 | 6924 | 65154 |
| 861.9413 | 592 | 874.8912 | 734 | 667.5194 | 750 | 627.0561 | 783 | ZBTB48 | HEK293 | Embryonic GSM20267 | 60928 | 65155 |
| 1146.817 | 480 | 1176.527 | 621 | 871.2759 | 481 | 840.5383 | 503 | ZSCAN22 | HEK293 | Embryonic GSM20267 | 16750 | 65156 |
| 62.21954 | 31 | 62.75543 | 35 | 29.87098 | 35 | 13.05649 | 28 | ZNF260 | HEK293 | Embryonic GSM20268 | 4035 | 65157 |
| 199.2664 | 54 | 274.6673 | 71 | 132.2474 | 63 | 110.8124 | 62 | ZNF528 | HEK293 | Embryonic GSM20268 | 3416 | 65160 |
| 117.5195 | 49 | 141.2127 | 60 | 91.92242 | 66 | 145.1047 | 87 | ZNF549 | HEK293 | Embryonic GSM20268 | 5286 | 65162 |
| 376.8705 | 275 | 658.1433 | 407 | 319.035 | 409 | 399.1933 | 481 | ZNF770 | HEK293 | Embryonic GSM20268 | 52505 | 65164 |
| 41.60029 | 17 | 70.15071 | 25 | 45.918 | 27 | 123.1445 | 47 | ZNF449 | HEK293 | Embryonic GSM20268 | 1900 | 65165 |
| 68.23102 | 21 | 142.1259 | 35 | 87.40909 | 35 | 147.923 | 49 | ZNF322 | HEK293 | Embryonic GSM20268 | 1703 | 65166 |
| 57.79106 | 49 | 95.30033 | 69 | 50.79069 | 75 | 85.68129 | 99 | ZNF121 | HEK293 | Embryonic GSM20267 | 9754 | 65170 |
| 261.5219 | 80 | 447.8343 | 118 | 240.6893 | 112 | 245.8757 | 121 | ZNF18 | HEK293 | Embryonic GSM20267 | 6197 | 65171 |
| 11.89558 | 16 | 41.34843 | 31 | 9.333834 | 25 | 37.65912 | 45 | ZNF8 | HEK293 | Embryonic GSM20267 | 4575 | 65174 |
| 21.92616 | 9 | 17.51802 | 9 | 3.333335 | 6 | 118.3085 | 35 | ZNF22 | HEK293 | Embryonic GSM20267 | 1080 | 65177 |
| 2.388794 | 63 | 0.288086 | 60 | 0.332889 | 102 | 6.584445 | 159 | ZNF22 | HEK293 | Embryonic GSM20267 | 46278 | 65178 |
| 949.3938 | 648 | 981.3244 | 820 | 673.7653 | 698 | 655.5888 | 763 | EGR2 | HEK293 | Embryonic GSM20267 | 54242 | 65179 |
| 834.2477 | 364 | 875.2889 | 491 | 673.4439 | 475 | 663.5316 | 526 | ZNF35 | HEK293 | Embryonic GSM20267 | 32300 | 65180 |
| 82.07461 | 21 | 136.6156 | 31 | 48.11166 | 23 | 62.32935 | 28 | ZNF85 | HEK293 | Embryonic GSM20268 | 1344 | 65184 |
| 36.15013 | 16 | 73.91595 | 26 | 38.04517 | 25 | 115.7616 | 46 | ZNF816 | HEK293 | Embryonic GSM20268 | 1922 | 65185 |
| 1013.342 | 935 | 1121.675 | 1217 | 717.835 | 1120 | 682.017 | 1214 | EGR3 | HEK293 | Embryonic GSM20268 | 103729 | 65186 |
| 2.772155 | 27 | 8.43233 | 42 | 2.003199 | 49 | 9.705537 | 74 | ZNF146 | HEK293 | Embryonic GSM20268 | 16748 | 65187 |
| 987.0568 | 336 | 1024.385 | 447 | 744.4552 | 349 | 757.545 | 411 | KLF1 | HEK293 | Embryonic GSM20268 | 17494 | 65188 |
| 474.3009 | 120 | 612.6834 | 152 | 347.513 | 147 | 409.2874 | 170 | ZBTB6 | HEK293 | Embryonic GSM20268 | 7416 | 65189 |
| 417.3962 | 103 | 588.292 | 137 | 268.4777 | 118 | 347.9162 | 143 | ZBTB6 | HEK293 | Embryonic GSM20268 | 6170 | 65190 |
| 12.23669 | 7 | 31.85024 | 13 | 9.499633 | 10 | 86.0321 | 31 | ZNF528 | HEK293 | Embryonic GSM20268 | 1198 | 65191 |
| 282.0268 | 105 | 504.815 | 157 | 255.4301 | 149 | 352.3632 | 185 | ZBTB6 | HEK293 | Embryonic GSM20268 | 10167 | 65192 |
| 0.819737 | 3 | 1.942335 | 5 | 2.097379 | 8 | 35.18353 | 26 | ZNF331 | HEK293 | Embryonic GSM20268 | 1907 | 65193 |
| 4.147932 | 7 | 5.283231 | 9 | 1.930119 | 10 | 24.30288 | 26 | ZNF331 | HEK293 | Embryonic GSM20268 | 2469 | 65194 |
| 0 | 3 | 0.640702 | 7 | 1.350114 | 13 | 13.54105 | 28 | ZNF586 | HEK293 | Embryonic GSM20268 | 3897 | 65196 |
| 1138.104 | 551 | 1189.341 | 733 | 865.1337 | 572 | 818.5028 | 583 | KLF15 | HEK293 | Embryonic GSM20268 | 21822 | 65197 |
| 503.3411 | 74 | 695.8371 | 98 | 212.0984 | 62 | 354.4335 | 87 | KLF15 | HEK293 | Embryonic GSM20268 | 2101 | 65198 |
| 113.8451 | 25 | 215.9543 | 40 | 94.15912 | 32 | 127.5345 | 40 | ZNF30 | HEK293 | Embryonic GSM20267 | 1308 | 65200 |
| 1122.987 | 415 | 1137.405 | 522 | 847.3565 | 403 | 813.7751 | 420 | YY1 | HEK293 | Embryonic GSM20267 | 14865 | 65203 |
| 1050.883 | 766 | 1114.153 | 993 | 747.6492 | 817 | 700.9901 | 846 | YY1 | HEK293 | Embryonic GSM20267 | 53068 | 65204 |
| 949.4565 | 156 | 1133.552 | 233 | 627.1809 | 171 | 663.5073 | 187 | YY1 | HEK293 | Embryonic GSM20267 | 5403 | 65205 |
| 1152.721 | 173 | 1184.001 | 236 | 575.4822 | 152 | 635.439 | 170 | ZNF76 | HEK293 | Embryonic GSM20267 | 4594 | 65206 |
| 64.23611 | 17 | 121.0179 | 27 | 61.24439 | 24 | 99.46221 | 33 | MZF1 | HEK293 | Embryonic GSM20267 | 1151 | 65207 |
| 11.28571 | 18 | 36.14641 | 33 | 11.65838 | 31 | 45.7381 | 55 | ZNF680 | HEK293 | Embryonic GSM20268 | 5475 | 65208 |
| 16.26016 | 17 | 47.25039 | 31 | 14.57938 | 27 | 51.97632 | 48 | ZNF680 | HEK293 | Embryonic GSM20268 | 4078 | 65209 |
| 8.215189 | 8 | 2.899462 | 6 | 3.722611 | 10 | 39.50471 | 28 | ZFP28 | HEK293 | Embryonic GSM20268 | 1992 | 65210 |
| 3.679801 | 6 | 2.428824 | 6 | 5.170666 | 12 | 38.07033 | 29 | ZFP28 | HEK293 | Embryonic GSM20268 | 2185 | 65211 |
| 1012.028 | 831 | 1086.726 | 1083 | 775.8916 | 1038 | 740.5777 | 1123 | GLIS1 | HEK293 | Embryonic GSM20268 | 74459 | 65214 |
| 32.70019 | 20 | 63.25025 | 31 | 34.16201 | 32 | 63.01482 | 45 | ZNF563 | HEK293 | Embryonic GSM20268 | 3181 | 65216 |

| | | | | | | | | | | | | |
|---|---|---|---|---|---|---|---|---|---|---|---|---|
| 76.06161 | 107 | 102.8396 | 137 | 71.81508 | 174 | 113.9446 | 220 | TP73 | Saos-2 | Bone | GSM50168 | 28759 | 6543 |
| 11.96277 | 33 | 16.86833 | 43 | 26.29746 | 73 | 30.91487 | 84 | TP73 | Saos-2 | Bone | GSM50168 | 12924 | 6544 |
| 84.88024 | 117 | 117.4221 | 151 | 63.51778 | 177 | 76.64715 | 205 | TP73 | Saos-2 | Bone | GSM50168 | 31126 | 6545 |
| 15.38252 | 67 | 36.22025 | 100 | 24.36298 | 134 | 28.13385 | 153 | TP73 | Saos-2 | Bone | GSM50169 | 30469 | 6546 |
| 673.5788 | 159 | 891.5723 | 204 | 384.9228 | 172 | 367.451 | 180 | TP53 | Saos-2 | Bone | GSM50169 | 9331 | 6547 |
| 268.449 | 208 | 338.2937 | 259 | 169.1224 | 280 | 177 | 310 | TP53 | Saos-2 | Bone | GSM50169 | 39374 | 6548 |
| 157.0482 | 69 | 182.3944 | 83 | 437.4682 | 168 | 831.2745 | 250 | ESR1 | MCF-7 | Breast | GSM63145 | 7725 | 6549 |
| 194.213 | 72 | 217.9936 | 85 | 494.3361 | 168 | 833.2886 | 234 | ESR1 | MCF-7 | Breast | GSM63145 | 6800 | 6550 |
| 283.8752 | 132 | 391.4123 | 172 | 662.0857 | 291 | 743.0777 | 362 | ESR1 | MCF-7 | Breast | GSM63145 | 15853 | 6551 |
| 310.2703 | 145 | 414.4328 | 186 | 628.9999 | 299 | 726.318 | 377 | ESR1 | MCF-7 | Breast | GSM63145 | 17560 | 6552 |
| 488.1551 | 248 | 616.7206 | 309 | 665.395 | 484 | 725.5266 | 645 | ESR1 | MCF-7 | Breast | GSM63145 | 33847 | 6553 |
| 157.4856 | 48 | 173.3739 | 56 | 440.3736 | 119 | 793.2526 | 174 | ESR1 | MCF-7 | Breast | GSM63145 | 3686 | 6555 |
| 463.0382 | 151 | 452.9096 | 166 | 705.5877 | 270 | 796.5602 | 349 | ESR1 | MCF-7 | Breast | GSM63145 | 12803 | 6556 |
| 531.0763 | 162 | 497.6755 | 174 | 777.0624 | 290 | 809.5263 | 362 | ESR1 | MCF-7 | Breast | GSM63146 | 12791 | 6557 |
| 518.4318 | 181 | 443.4625 | 187 | 735.9606 | 325 | 785.6922 | 427 | ESR1 | MCF-7 | Breast | GSM63146 | 16740 | 6558 |
| 319.9109 | 66 | 332.2098 | 74 | 491.3113 | 117 | 621.4482 | 141 | ESR1 | MCF-7 | Breast | GSM63146 | 3110 | 6559 |
| 82.04741 | 30 | 70.63296 | 31 | 175.2823 | 65 | 408.616 | 108 | ESR1 | MCF-7 | Breast | GSM63146 | 2865 | 6560 |
| 75.13364 | 30 | 101.1045 | 39 | 127.4269 | 58 | 346.1063 | 103 | ESR1 | MCF-7 | Breast | GSM63146 | 3141 | 6561 |
| 632.4464 | 333 | 655.5324 | 391 | 634.8643 | 598 | 696.4314 | 786 | ESR1 | MCF-7 | Breast | GSM63146 | 49242 | 6562 |
| 50.66369 | 33 | 62.27173 | 41 | 168.6775 | 88 | 360.4273 | 136 | ESR1 | MCF-7 | Breast | GSM63146 | 5410 | 6563 |
| 136.2152 | 111 | 166.5686 | 137 | 297.5267 | 239 | 553.267 | 334 | ESR1 | MCF-7 | Breast | GSM63146 | 21527 | 6564 |
| 331.2611 | 164 | 294.9882 | 175 | 564.4514 | 313 | 703.0076 | 416 | ESR1 | MCF-7 | Breast | GSM63146 | 21323 | 6565 |
| 854.7303 | 748 | 835.9842 | 871 | 721.2537 | 1087 | 742.3936 | 1290 | FOXA1 | MCF-7 | Breast | GSM63146 | 98546 | 6566 |
| 871.4366 | 749 | 847.1792 | 867 | 734.9008 | 1085 | 745.4223 | 1268 | FOXA1 | MCF-7 | Breast | GSM63147 | 93919 | 6567 |
| 776.8506 | 652 | 699.7785 | 682 | 668.4878 | 985 | 699.1675 | 1187 | FOXA1 | MCF-7 | Breast | GSM63147 | 95052 | 6568 |
| 824.3809 | 544 | 761.3848 | 578 | 669.8814 | 743 | 704.0061 | 914 | FOXA1 | T47D | Mammary | GSM63147 | 60163 | 6570 |
| 748.9833 | 453 | 698.9539 | 492 | 669.486 | 728 | 705.5043 | 896 | FOXA1 | MCF-7 | Breast | GSM63147 | 57963 | 6571 |
| 713.6924 | 343 | 741.5641 | 448 | 488.7823 | 438 | 463.4685 | 466 | CTCF | MCF-7 | Breast | GSM63147 | 45003 | 6572 |
| 717.8339 | 399 | 768.6191 | 544 | 550.6825 | 518 | 529.595 | 558 | CTCF | MCF-7 | Breast | GSM63147 | 54055 | 6573 |
| 730.1186 | 365 | 765.1657 | 486 | 542.7959 | 463 | 524.8779 | 496 | CTCF | MCF-7 | Breast | GSM63147 | 46337 | 6574 |
| 761.3808 | 472 | 806.8971 | 629 | 559.9747 | 571 | 550.5054 | 636 | CTCF | MCF-7 | Breast | GSM63147 | 59488 | 6575 |
| 708.9587 | 356 | 764.9969 | 498 | 489.3666 | 456 | 511.0332 | 503 | CTCF | MCF-7 | Breast | GSM63147 | 48378 | 6576 |
| 401.5273 | 141 | 375.8634 | 152 | 749.4483 | 279 | 793.7229 | 352 | ESR1 | MCF-7 | Breast | GSM63148 | 12888 | 6577 |
| 457.332 | 167 | 585.3688 | 210 | 764.5252 | 342 | 816.1324 | 449 | ESR1 | MCF-7 | Breast | GSM63148 | 16043 | 6578 |
| 572.9185 | 218 | 632.0617 | 255 | 746.9161 | 431 | 773.8853 | 525 | ESR1 | MCF-7 | Breast | GSM63148 | 22358 | 6579 |
| 244.6076 | 86 | 251.2653 | 97 | 555.7769 | 189 | 830.32 | 247 | ESR1 | MCF-7 | Breast | GSM63148 | 7706 | 6580 |
| 221.3842 | 73 | 189.736 | 75 | 432.9396 | 149 | 779.8739 | 215 | ESR1 | MCF-7 | Breast | GSM63148 | 6101 | 6581 |
| 50.472 | 14 | 65.693 | 18 | 48.55303 | 20 | 240.4348 | 51 | ESR1 | T47D | Mammary | GSM63148 | 1021 | 6586 |
| 70.34268 | 18 | 37.10604 | 14 | 70.55271 | 26 | 288.8653 | 60 | ESR1 | T47D | Mammary | GSM63149 | 1167 | 6587 |
| 71.01311 | 18 | 59.40234 | 18 | 61.26824 | 24 | 325.8382 | 64 | ESR1 | T47D | Mammary | GSM63149 | 1152 | 6588 |
| 69.06158 | 17 | 51.96145 | 16 | 72.07432 | 25 | 329.0961 | 62 | ESR1 | T47D | Mammary | GSM63149 | 1055 | 6589 |
| 6.076997 | 21 | 19.40109 | 36 | 7.229053 | 40 | 24.20177 | 62 | JUND | GM12878 | Blood | GSM75433 | 9385 | 660 |
| 989.0734 | 424 | 1014.517 | 547 | 711.2654 | 416 | 673.4929 | 425 | MAX | GM12878 | Blood | GSM75433 | 23952 | 661 |
| 1059.434 | 487 | 1118.756 | 661 | 790.7451 | 492 | 757.9159 | 514 | MYC | GM12878 | Blood | GSM75433 | 22491 | 666 |
| 260.4232 | 59 | 402.2868 | 83 | 185.8902 | 70 | 274.0225 | 91 | NR2C2 | GM12878 | Blood | GSM60815 | 3097 | 669 |
| 175.6958 | 32 | 318.0909 | 50 | 75.2241 | 28 | 175.9477 | 47 | NR2C2 | GM12878 | Blood | GSM60815 | 1255 | 670 |
| 1205.21 | 455 | 1184.542 | 526 | 981.607 | 512 | 939.9573 | 524 | SRC | OCI-Ly2 | Mammary | GSM71797 | 12950 | 6709 |
| 1231.912 | 747 | 1262.952 | 939 | 944.716 | 815 | 909.2681 | 872 | SRC | OCI-Ly2 | Mammary | GSM71797 | 28674 | 6710 |
| 612.3407 | 381 | 666.0935 | 501 | 265.3686 | 444 | 258.7195 | 482 | SOX2 | None | None | GSM17018 | 63587 | 67317 |
| 277.7751 | 183 | 332.289 | 223 | 115.0813 | 208 | 141.6712 | 243 | SOX2 | None | None | GSM17018 | 29805 | 67318 |
| 889.5034 | 284 | 834.9746 | 299 | 785.5404 | 418 | 796.0907 | 490 | ESRRA | BT-474 | Breast | GSM19692 | 16909 | 67367 |
| 933.4942 | 348 | 963.5921 | 454 | 693.7881 | 359 | 611.0642 | 350 | KDM6A | Jurkat | None | GSM15813 | 20573 | 67370 |
| 32.37511 | 16 | 32.26791 | 18 | 19.82273 | 20 | 112.6797 | 48 | SIX2 | None | Embryonic | GSM19045 | 2176 | 67424 |
| 648.0281 | 410 | 664.2086 | 519 | 381.7726 | 515 | 318.0567 | 531 | SIX2 | None | Embryonic | GSM19045 | 67568 | 67425 |
| 414.0635 | 58 | 404.0833 | 62 | 255.2791 | 59 | 404.1287 | 81 | RARA | SK-N-SH | Brain | GSM16931 | 1456 | 67426 |
| 477.6425 | 63 | 497.8661 | 70 | 291.0423 | 63 | 454.87 | 86 | RARA | SK-N-SH | Brain | GSM16931 | 1414 | 67427 |
| 415.5734 | 65 | 499.2484 | 79 | 310.6613 | 75 | 358.9237 | 86 | RARA | SK-N-SH | Brain | GSM16931 | 1970 | 67429 |
| 1245.402 | 219 | 1260.44 | 282 | 936.8209 | 195 | 954.4866 | 214 | E2F1 | None | Umbilical C | GSM20468 | 4383 | 67454 |
| 1234.971 | 170 | 1281.511 | 240 | 797.3049 | 160 | 1009.347 | 198 | E2F1 | None | Umbilical C | GSM20468 | 3388 | 67455 |
| 1246.843 | 351 | 1264.239 | 449 | 954.7237 | 320 | 925.1364 | 333 | E2F1 | None | Umbilical C | GSM20468 | 7860 | 67456 |
| 1240.273 | 218 | 1261.252 | 286 | 949.152 | 198 | 968.4591 | 228 | E2F1 | None | Umbilical C | GSM20468 | 4470 | 67457 |
| 1190.074 | 214 | 1240.521 | 301 | 868.0528 | 199 | 944.6342 | 241 | E2F1 | None | Umbilical C | GSM20468 | 5105 | 67458 |
| 1028.094 | 126 | 1267.177 | 183 | 533.1605 | 115 | 685.3206 | 140 | E2F1 | None | Umbilical C | GSM20468 | 2670 | 67459 |
| 124.4481 | 159 | 146.3459 | 193 | 159.2846 | 280 | 185.1372 | 321 | GATA3 | MDA-MB-2 | Mammary | GSM18559 | 39306 | 67494 |
| 125.9747 | 109 | 156.0607 | 135 | 118.989 | 169 | 192.8845 | 219 | GATA3 | MDA-MB-2 | Mammary | GSM18559 | 20709 | 67495 |
| 913.6417 | 455 | 918.0425 | 553 | 659.6077 | 457 | 631.7519 | 485 | SMARCA4 | MDA-MB-2 | Mammary | GSM18560 | 27963 | 67496 |
| 792.9819 | 298 | 758.7292 | 334 | 380.8601 | 306 | 389.1862 | 334 | SMARCA4 | MDA-MB-2 | Mammary | GSM18560 | 25111 | 67497 |
| 828.2445 | 294 | 835.9291 | 363 | 430.6599 | 296 | 550.1704 | 354 | SMARCA4 | MDA-MB-2 | Mammary | GSM18560 | 20822 | 67498 |
| 789.3311 | 375 | 805.8986 | 471 | 541.6567 | 410 | 556.3343 | 449 | GATA3 | MDA-MB-2 | Mammary | GSM18560 | 31387 | 67499 |
| 673.6437 | 357 | 668.4017 | 426 | 319.3833 | 399 | 367.5104 | 457 | GATA3 | MDA-MB-2 | Mammary | GSM18560 | 39985 | 67500 |
| 534.9672 | 222 | 670.7974 | 275 | 240.1244 | 235 | 301.1861 | 278 | GATA3 | MDA-MB-2 | Mammary | GSM18560 | 21732 | 67501 |
| 164.2931 | 179 | 187.1428 | 214 | 165.8406 | 285 | 160.2441 | 308 | GATA3 | MDA-MB-2 | Mammary | GSM18560 | 37574 | 67502 |

| | | | | | | | | | | | | | |
|---|---|---|---|---|---|---|---|---|---|---|---|---|---|
| 113.3502 | 138 | 160.2044 | 179 | 146.4607 | 241 | 134.8418 | 256 | GATA3 | MDA-MB-2 | Mammary | GSM18560 | 31048 | 67503 |
| 277.8318 | 246 | 350.378 | 305 | 247.8804 | 370 | 331.3293 | 445 | GATA3 | MDA-MB-2 | Mammary | GSM18559 | 46286 | 67516 |
| 282.6757 | 246 | 344.5256 | 301 | 239.7909 | 363 | 269.6458 | 412 | GATA3 | MDA-MB-2 | Mammary | GSM18559 | 45508 | 67517 |
| 171.8697 | 237 | 219.7928 | 295 | 145.6329 | 364 | 189.0574 | 431 | GATA3 | MDA-MB-2 | Mammary | GSM18559 | 58181 | 67518 |
| 182.3029 | 255 | 187.0155 | 295 | 155.3775 | 394 | 192.4883 | 460 | GATA3 | MDA-MB-2 | Mammary | GSM18559 | 64583 | 67519 |
| 1072.383 | 486 | 1106.438 | 634 | 799.0353 | 478 | 741.6482 | 470 | BRD4 | COLO-741 | None | GSM18907 | 19596 | 67532 |
| 970.41 | 125 | 1160.485 | 151 | 454.8198 | 108 | 368.6668 | 102 | BRD4 | LS180 | Colon | GSM18907 | 2724 | 67533 |
| 523.915 | 60 | 404.2492 | 55 | 286.5888 | 55 | 245.0349 | 53 | BRD4 | COLO-320 | None | GSM18907 | 1039 | 67534 |
| 1115.808 | 158 | 1139.132 | 196 | 773.8096 | 172 | 527.3241 | 148 | BRD4 | SW480 | Colon | GSM18907 | 3930 | 67535 |
| 1118.596 | 229 | 1144.027 | 300 | 834.1364 | 220 | 684.6048 | 211 | BRD4 | HCT-15 | Colon | GSM18907 | 5966 | 67536 |
| 941.9199 | 128 | 990.8518 | 143 | 447.4299 | 112 | 442.4679 | 118 | BRD4 | COLO 205 | None | GSM18907 | 2931 | 67537 |
| 612.1897 | 83 | 789.2038 | 105 | 358.2249 | 81 | 281.3673 | 75 | BRD4 | HCT-116 | Colon | GSM18907 | 1840 | 67539 |
| 85.12957 | 47 | 103.7368 | 58 | 47.21091 | 57 | 98.5792 | 83 | JUN | BT-549 | Breast | GSM18488 | 6509 | 67573 |
| 29.72806 | 17 | 39.44996 | 22 | 13.0782 | 19 | 69.98502 | 42 | JUN | BT-549 | Breast | GSM18488 | 2582 | 67574 |
| 240.9759 | 52 | 255.1626 | 59 | 170.2947 | 61 | 196.9975 | 70 | MYC | None | Foreskin | GSM15999 | 2575 | 67584 |
| 290.7568 | 58 | 217.8839 | 54 | 164.4388 | 60 | 170.0149 | 65 | MYC | None | Foreskin | GSM15999 | 2598 | 67585 |
| 365.8125 | 237 | 407.6178 | 279 | 415.3143 | 376 | 482.7591 | 432 | FOXA1 | CFPAC-1 | Pancreatic | GSM19199 | 33348 | 67596 |
| 748.9406 | 345 | 737.5562 | 408 | 511.5247 | 408 | 483.5547 | 433 | ELF3 | CFPAC-1 | Pancreatic | GSM19199 | 33192 | 67597 |
| 481.3572 | 266 | 408.3752 | 279 | 402.8977 | 372 | 539.1497 | 451 | FOXA1 | CFPAC-1 | Pancreatic | GSM19199 | 33728 | 67598 |
| 724.6802 | 288 | 747.4776 | 352 | 549.9548 | 374 | 455.3346 | 375 | ELF3 | CFPAC-1 | Pancreatic | GSM19199 | 27043 | 67599 |
| 1062.683 | 186 | 1079.68 | 237 | 729.1381 | 199 | 580.2072 | 188 | KLF6 | CFPAC-1 | Pancreatic | GSM19199 | 5486 | 67600 |
| 924.0342 | 339 | 895.5995 | 383 | 727.04 | 385 | 705.6138 | 410 | KLF4 | CFPAC-1 | Pancreatic | GSM19199 | 18203 | 67601 |
| 269.3737 | 47 | 199.9413 | 43 | 93.41103 | 36 | 204.4934 | 58 | ZEB1 | PANC-1 | Pancreatic | GSM15742 | 1510 | 67610 |
| 936.4721 | 313 | 925.6913 | 371 | 738.9828 | 347 | 715.0526 | 372 | KLF5 | CFPAC-1 | Pancreatic | GSM15742 | 15006 | 67611 |
| 877.461 | 252 | 893.9515 | 320 | 667.2285 | 300 | 691.144 | 328 | IRF1 | CFPAC-1 | Pancreatic | GSM15742 | 14584 | 67612 |
| 20.88218 | 20 | 28.46718 | 26 | 41.42374 | 42 | 68.41685 | 56 | HNF1B | CFPAC-1 | Pancreatic | GSM15742 | 4064 | 67613 |
| 560.8278 | 303 | 567.1738 | 342 | 502.2287 | 435 | 560.7515 | 511 | FOXA1 | CFPAC-1 | Pancreatic | GSM15742 | 38791 | 67614 |
| 796.045 | 484 | 800.2899 | 593 | 613.399 | 595 | 590.8101 | 646 | ELF3 | CFPAC-1 | Pancreatic | GSM15742 | 47325 | 67615 |
| 1110.813 | 242 | 1159.453 | 337 | 858.2121 | 235 | 843.8292 | 254 | ZBTB7A | None | None | GSM19395 | 7290 | 67683 |
| 1026.543 | 252 | 1056.176 | 334 | 797.1172 | 262 | 666.8414 | 256 | ZBTB7A | None | None | GSM19395 | 9899 | 67685 |
| 1043.328 | 302 | 1064.906 | 391 | 801.6745 | 306 | 699.0355 | 283 | ZBTB7A | None | None | GSM19395 | 11577 | 67686 |
| 382.7319 | 156 | 448.3683 | 188 | 251.9215 | 195 | 287.1739 | 222 | RAD21 | SK-N-SH | Brain | GSM20383 | 16710 | 67698 |
| 668.088 | 294 | 768.4332 | 409 | 449.8393 | 378 | 508.1041 | 429 | CTCF | SK-N-SH | Brain | GSM20383 | 36018 | 67699 |
| 1248.469 | 748 | 1237.815 | 882 | 969.8935 | 817 | 959.908 | 916 | CREB1 | LNCaP | Prostate | GSM18730 | 24358 | 67710 |
| 1263.406 | 595 | 1282.79 | 737 | 970.3987 | 588 | 956.7946 | 650 | CREB1 | LNCaP | Prostate | GSM18730 | 15182 | 67711 |
| 1012.359 | 932 | 889.7451 | 967 | 764.4407 | 1180 | 808.9451 | 1442 | FOXA1 | LNCaP | Prostate | GSM18730 | 95156 | 67712 |
| 982.6019 | 875 | 863.8816 | 900 | 766.8877 | 1142 | 802.5331 | 1381 | FOXA1 | LNCaP | Prostate | GSM18730 | 88689 | 67713 |
| 1323.709 | 769 | 1304.266 | 893 | 998.6596 | 775 | 994.4427 | 874 | CREB1 | LNCaP | Prostate | GSM18730 | 20531 | 67714 |
| 1311.406 | 654 | 1336.381 | 815 | 1024.989 | 671 | 1008.52 | 741 | CREB1 | LNCaP | Prostate | GSM18730 | 15390 | 67715 |
| 967.3765 | 843 | 852.5388 | 860 | 743.4155 | 1072 | 793.6748 | 1324 | FOXA1 | LNCaP | Prostate | GSM18730 | 83052 | 67716 |
| 977.3489 | 816 | 853.7501 | 820 | 764.8576 | 1048 | 797.6707 | 1270 | FOXA1 | LNCaP | Prostate | GSM18730 | 75230 | 67717 |
| 1333.929 | 806 | 1325.109 | 954 | 986.4341 | 802 | 967.2914 | 877 | CREB1 | LNCaP | Prostate | GSM15384 | 23906 | 67718 |
| 1016.144 | 943 | 909.7838 | 1003 | 791.4188 | 1254 | 829.0789 | 1509 | FOXA1 | LNCaP | Prostate | GSM15384 | 107096 | 67719 |
| 1358.877 | 901 | 1366.814 | 1085 | 1026.769 | 988 | 1020.789 | 1119 | CREB1 | LNCaP | Prostate | GSM15384 | 30260 | 67720 |
| 985.1691 | 701 | 899.199 | 727 | 810.7518 | 943 | 871.0475 | 1202 | FOXA1 | LNCaP | Prostate | GSM16911 | 57877 | 67730 |
| 1062.136 | 966 | 952.4164 | 1030 | 850.2693 | 1315 | 893.7114 | 1590 | FOXA1 | LNCaP | Prostate | GSM16911 | 102409 | 67731 |
| 1062.661 | 293 | 970.7134 | 281 | 970.7098 | 457 | 1007.155 | 572 | GATA2 | LNCaP | Prostate | GSM16911 | 11915 | 67734 |
| 1173.809 | 1019 | 1054.833 | 1088 | 887.8218 | 1300 | 930.942 | 1570 | FOXA1 | LNCaP | Prostate | GSM16911 | 88848 | 67735 |
| 998.6778 | 426 | 901.5206 | 411 | 893.9946 | 643 | 935.596 | 806 | GATA2 | LNCaP | Prostate | GSM16911 | 24086 | 67737 |
| 1334.605 | 1150 | 1248.353 | 1301 | 979.9176 | 1490 | 993.1983 | 1734 | FOXA1 | LNCaP | Prostate | GSM16911 | 100518 | 67738 |
| 980.1929 | 279 | 918.7345 | 289 | 902.9484 | 454 | 968.751 | 610 | GATA2 | LNCaP | Prostate | GSM16911 | 14651 | 67740 |
| 991.1917 | 518 | 910.8477 | 531 | 805.2412 | 658 | 862.8373 | 851 | FOXA1 | LNCaP | Prostate | GSM16911 | 33114 | 67741 |
| 929.7331 | 473 | 900.226 | 541 | 754.4553 | 613 | 848.1577 | 862 | AR | LNCaP | Prostate | GSM16911 | 35358 | 67742 |
| 917.5552 | 443 | 852.076 | 466 | 824.9422 | 695 | 876.712 | 886 | GATA2 | LNCaP | Prostate | GSM16911 | 33457 | 67743 |
| 1178.362 | 1075 | 1102.901 | 1211 | 892.2985 | 1416 | 933.9557 | 1695 | FOXA1 | LNCaP | Prostate | GSM16911 | 111714 | 67744 |
| 766.9746 | 214 | 849.2087 | 257 | 583.8966 | 273 | 813.0064 | 439 | AR | LNCaP | Prostate | GSM16911 | 16009 | 67745 |
| 880.8117 | 305 | 794.7859 | 297 | 856.5324 | 565 | 907.917 | 723 | GATA2 | LNCaP | Prostate | GSM16911 | 22341 | 67746 |
| 942.6565 | 387 | 853.5799 | 375 | 859.3994 | 613 | 911.3684 | 785 | GATA2 | LNCaP | Prostate | GSM16911 | 24577 | 67747 |
| 769.3728 | 455 | 703.6467 | 476 | 738.1939 | 811 | 786.6964 | 1021 | GATA2 | LNCaP | Prostate | GSM16911 | 55700 | 67748 |
| 935.9616 | 422 | 817.3686 | 387 | 835.7856 | 642 | 871.2467 | 799 | GATA2 | LNCaP | Prostate | GSM16911 | 28764 | 67749 |
| 1144.602 | 949 | 1038.313 | 1013 | 876.3551 | 1192 | 938.9011 | 1482 | FOXA1 | LNCaP | Prostate | GSM16911 | 73457 | 67750 |
| 768.1836 | 173 | 848.1627 | 201 | 860.884 | 286 | 957.6093 | 425 | AR | LNCaP | Prostate | GSM16911 | 9524 | 67753 |
| 841.274 | 149 | 850.1303 | 164 | 572.9472 | 168 | 969.089 | 286 | AR | LNCaP | Prostate | GSM16911 | 5825 | 67754 |
| 227.2552 | 63 | 424.0309 | 98 | 153.2694 | 75 | 227.016 | 97 | RUNX1 | None | Fetal Liver | GSM18160 | 4334 | 67763 |
| 400.0544 | 376 | 497.0964 | 464 | 258.2169 | 530 | 243.2519 | 572 | SPI1 | None | Bone Marr | GSM18160 | 89742 | 67764 |
| 278.4514 | 245 | 321.8635 | 295 | 175.3603 | 337 | 187.2271 | 376 | SPI1 | None | Fetal Liver | GSM18160 | 52416 | 67765 |
| 22.33206 | 10 | 54.28015 | 18 | 31.01488 | 18 | 112.4065 | 37 | IRF2 | None | Bone Marr | GSM18160 | 1273 | 67766 |
| 224.877 | 50 | 232.3789 | 56 | 119.4487 | 51 | 224.3806 | 75 | NFE2 | None | Bone Marr | GSM18160 | 2592 | 67767 |
| 1110.696 | 827 | 1201.936 | 1093 | 781.0333 | 877 | 728.4424 | 901 | NFE2 | F5 | Fetal Liver | GSM18160 | 55156 | 67768 |
| 448.5326 | 82 | 635.9219 | 110 | 290.0612 | 90 | 328.0374 | 102 | NFE2 | None | Fetal Liver | GSM18160 | 3285 | 67769 |
| 204.3691 | 175 | 424.8054 | 268 | 260.552 | 305 | 225.6614 | 315 | TAL1 | F5 | Fetal Liver | GSM18160 | 36128 | 67771 |
| 591.5744 | 83 | 736.8503 | 103 | 523.3732 | 104 | 352.8314 | 88 | BRD4 | T47D | None | GSM18427 | 2115 | 67793 |

| | | | | | | | | | | | | | |
|---|---|---|---|---|---|---|---|---|---|---|---|---|---|
| 1072.108 | 313 | 1077.669 | 384 | 889.201 | 367 | 850.2256 | 377 | BRD4 | T47D | None | GSM18427 | 10532 | 67794 |
| 366.4615 | 46 | 453.3876 | 57 | 174.4262 | 40 | 357.2558 | 64 | BRD4 | SUM185PE | None | GSM18427 | 1042 | 67795 |
| 1086.296 | 191 | 1073.778 | 226 | 822.7218 | 211 | 721.4354 | 209 | BRD4 | SUM185PE | None | GSM18427 | 6071 | 67796 |
| 1005.998 | 158 | 1085.042 | 193 | 651.114 | 169 | 537.1237 | 162 | BRD4 | SUM149 | None | GSM18427 | 4840 | 67798 |
| 422.8411 | 55 | 485.2212 | 65 | 278.1825 | 58 | 422.9348 | 78 | BRD4 | SUM149R | None | GSM18427 | 1322 | 67799 |
| 382.4116 | 68 | 453.5221 | 82 | 270.2559 | 77 | 184.8723 | 67 | BRD4 | SUM149R | None | GSM18427 | 2402 | 67800 |
| 1118.914 | 263 | 1100.426 | 306 | 828.679 | 236 | 764.4851 | 241 | BRD4 | SUM1315M | None | GSM18427 | 7201 | 67802 |
| 448.2053 | 62 | 490.5093 | 71 | 353.025 | 72 | 297.4482 | 69 | BRD4 | MDA-MB-4 | None | GSM18427 | 1454 | 67804 |
| 332.4038 | 146 | 340.5809 | 165 | 198.6811 | 175 | 233.9642 | 202 | BRD4 | HCC1395 | None | GSM18426 | 15516 | 67805 |
| 861.4308 | 494 | 872.7101 | 611 | 622.7505 | 532 | 605.7724 | 581 | BRD4 | HCC1395 | None | GSM18426 | 39510 | 67806 |
| 1087.156 | 230 | 1080.97 | 276 | 780.288 | 224 | 685.2458 | 223 | BRD4 | SUM159R | None | GSM15531 | 7057 | 67808 |
| 1124.755 | 370 | 1132.727 | 456 | 857.6595 | 348 | 825.1949 | 361 | BRD4 | SUM159PT | Breast | GSM15531 | 10539 | 67828 |
| 1091.653 | 379 | 1104.168 | 474 | 821.4681 | 371 | 802.3897 | 401 | BRD4 | SUM159R | None | GSM15531 | 14343 | 67829 |
| 194.6965 | 43 | 164.0529 | 43 | 80.61205 | 39 | 217.8387 | 69 | BRD4 | LP-1 | None | GSM19611 | 2333 | 67836 |
| 67.54174 | 20 | 71.97694 | 23 | 48.64587 | 25 | 88.33743 | 36 | EP300 | LP-1 | None | GSM18473 | 1603 | 67841 |
| 755.5986 | 182 | 726.9105 | 196 | 354.6105 | 176 | 487.5954 | 219 | TET2 | HEK293T | Embryonic | GSM19375 | 9276 | 67858 |
| 624.5815 | 548 | 659.0146 | 710 | 433.0413 | 719 | 433.8279 | 799 | SPI1 | None | Blood | GSM16814 | 114545 | 67926 |
| 657.633 | 733 | 708.5313 | 949 | 480.5695 | 1005 | 472.4702 | 1135 | SPI1 | None | Blood | GSM16814 | 171581 | 67927 |
| 666.194 | 606 | 701.0331 | 778 | 476.7179 | 787 | 461.0853 | 869 | SPI1 | None | Blood | GSM16814 | 121294 | 67928 |
| 666.5699 | 663 | 708.8574 | 858 | 470.0555 | 859 | 452.6771 | 949 | SPI1 | None | Blood | GSM16814 | 138829 | 67929 |
| 821.928 | 262 | 836.5244 | 336 | 690.9074 | 368 | 681.3651 | 410 | GRHL2 | PEO1 | None | GSM18256 | 22657 | 68001 |
| 798.6526 | 327 | 822.7885 | 425 | 660.5628 | 454 | 667.028 | 528 | GRHL2 | OVCA429 | None | GSM18256 | 32011 | 68002 |
| 350.9298 | 103 | 503.1266 | 138 | 417.6723 | 163 | 633.7679 | 214 | GRHL2 | NIH:OVCAF | None | GSM18256 | 7731 | 68003 |
| 121.8632 | 25 | 203.3796 | 37 | 77.55554 | 27 | 184.4107 | 46 | KDM2B | DND-41 | None | GSM18120 | 1025 | 68017 |
| 863.5496 | 386 | 948.6039 | 574 | 660.0502 | 461 | 605.1307 | 451 | NFKB1 | L1236 | None | GSM15563 | 33528 | 68048 |
| 44.85301 | 16 | 137.042 | 33 | 84.12631 | 33 | 202.9283 | 56 | RELA | L1236 | None | GSM15563 | 1612 | 68050 |
| 1020.608 | 669 | 1068.212 | 868 | 735.9618 | 710 | 675.0389 | 704 | NFKB1 | L1236 | None | GSM15563 | 45801 | 68051 |
| 356.6325 | 232 | 591.3253 | 325 | 183.6242 | 289 | 205.7674 | 327 | RELB | L1236 | None | GSM15563 | 41629 | 68052 |
| 246.2491 | 201 | 385.8405 | 274 | 144.1326 | 270 | 215.8444 | 336 | RELB | L1236 | None | GSM15563 | 42656 | 68053 |
| 108.8421 | 28 | 109.4388 | 31 | 177.485 | 52 | 341.2557 | 79 | ESR1 | LTED | Breast | GSM20616 | 1739 | 68054 |
| 835.5712 | 213 | 738.8745 | 221 | 776.9741 | 314 | 835.9326 | 420 | ESR1 | LTED | Breast | GSM20616 | 13869 | 68055 |
| 922.1117 | 325 | 897.7292 | 370 | 789.6227 | 452 | 809.2516 | 540 | ESR1 | MCF-7 | Breast | GSM20616 | 20698 | 68056 |
| 846.6313 | 461 | 811.8528 | 519 | 715.201 | 657 | 723.2617 | 764 | ESR1 | MCF-7 | Breast | GSM20616 | 43198 | 68057 |
| 80.62957 | 25 | 27.53149 | 16 | 15.64406 | 17 | 58.21397 | 33 | POLR3D | IMR90hTer | Fetal Lung | GSM19063 | 1815 | 68058 |
| 66.99517 | 24 | 34.09451 | 19 | 21.56373 | 21 | 53.7868 | 34 | POLR3D | IMR90hTer | Fetal Lung | GSM19063 | 2043 | 68059 |
| 90.33445 | 23 | 43.29591 | 17 | 30.57605 | 19 | 63.08232 | 29 | POLR3D | IMR90hTer | Fetal Lung | GSM19063 | 1334 | 68060 |
| 65.92702 | 23 | 19.56384 | 14 | 10.74296 | 15 | 22.31618 | 22 | POLR3D | IMR90hTer | Fetal Lung | GSM19063 | 1849 | 68061 |
| 80.15205 | 21 | 31.44973 | 14 | 22.76802 | 16 | 38.23651 | 22 | POLR3D | IMR90hTer | Fetal Lung | GSM19063 | 1235 | 68062 |
| 63.33562 | 30 | 11.76762 | 16 | 9.180171 | 21 | 35.53705 | 38 | POLR3D | IMR90hTer | Fetal Lung | GSM19063 | 3357 | 68063 |
| 76.01498 | 25 | 19.2018 | 14 | 18.5668 | 19 | 40.87331 | 29 | POLR3D | IMR90hTer | Fetal Lung | GSM19063 | 1907 | 68064 |
| 108.8424 | 28 | 27.54158 | 15 | 26.7683 | 20 | 50.83454 | 29 | POLR3D | IMR90hTer | Fetal Lung | GSM19063 | 1603 | 68065 |
| 39.15037 | 29 | 16.40066 | 23 | 6.97064 | 25 | 28.73396 | 44 | POLR3D | IMR90hTer | Fetal Lung | GSM19063 | 4938 | 68067 |
| 58.01751 | 21 | 14.88575 | 12 | 15.70293 | 17 | 41.03502 | 28 | POLR3D | IMR90hTer | Fetal Lung | GSM19063 | 1766 | 68068 |
| 778.9429 | 565 | 714.9715 | 599 | 598.4755 | 728 | 659.6307 | 946 | AR | LNCaP | Prostate | GSM15866 | 71514 | 68073 |
| 746.4199 | 568 | 691.2101 | 616 | 579.0948 | 740 | 627.827 | 939 | AR | LNCaP | Prostate | GSM15866 | 71539 | 68074 |
| 1120.682 | 557 | 1060.651 | 603 | 894.8063 | 643 | 931.8338 | 800 | AR | LNCaP | Prostate | GSM15866 | 22975 | 68075 |
| 984.4926 | 241 | 953.9317 | 269 | 662.744 | 248 | 804.6112 | 324 | AR | LNCaP | Prostate | GSM15866 | 10577 | 68076 |
| 833.0422 | 631 | 770.3313 | 679 | 717.6532 | 937 | 766.5306 | 1167 | AR | C4-2 | Prostate | GSM15866 | 73153 | 68077 |
| 871.1465 | 611 | 814.4981 | 665 | 708.1494 | 821 | 769.3227 | 1060 | AR | C4-2 | Prostate | GSM15866 | 59965 | 68078 |
| 1130.854 | 507 | 1077.757 | 545 | 941.9767 | 631 | 985.1559 | 785 | AR | C4-2 | Prostate | GSM15866 | 18810 | 68079 |
| 1055.196 | 581 | 998.0348 | 625 | 865.1158 | 715 | 900.0576 | 880 | AR | C4-2 | Prostate | GSM15866 | 27880 | 68080 |
| 959.5473 | 189 | 956.3384 | 210 | 746.0784 | 230 | 921.3936 | 350 | AR | C4-2 | Prostate | GSM18434 | 8183 | 68089 |
| 442.9269 | 78 | 363.9699 | 76 | 259.0259 | 80 | 821.2304 | 160 | AR | C4-2 | Prostate | GSM18434 | 2783 | 68090 |
| 176.9424 | 31 | 233.8615 | 40 | 137.809 | 37 | 717.0587 | 102 | AR | C4-2 | Prostate | GSM18434 | 1102 | 68091 |
| 503.5824 | 103 | 473.581 | 109 | 367.7899 | 121 | 769.0295 | 190 | AR | C4-2 | Prostate | GSM18434 | 4429 | 68092 |
| 5.435038 | 17 | 13.58065 | 27 | 5.8173 | 31 | 48.77199 | 67 | CRY1 | U2OS | Bone | GSM10811 | 7104 | 68102 |
| 111.0908 | 72 | 136.2648 | 89 | 43.24227 | 79 | 110.4488 | 121 | CLOCK | U2OS | Bone | GSM10811 | 11324 | 68103 |
| 162.0431 | 40 | 312.3686 | 64 | 63.65912 | 36 | 132.1973 | 55 | ARNTL | U2OS | Bone | GSM10811 | 2422 | 68104 |
| 110.3571 | 86 | 143.7631 | 109 | 59.70171 | 109 | 67.77779 | 124 | KDM1A | Kasumi-1 | Blood | GSM18444 | 15522 | 68116 |
| 114.2186 | 95 | 127.8477 | 113 | 50.73502 | 114 | 77.53373 | 143 | KDM1A | Kasumi-1 | Blood | GSM18444 | 17955 | 68117 |
| 154.7848 | 107 | 136.8554 | 115 | 55.55915 | 116 | 83.60023 | 145 | KDM1A | Kasumi-1 | Blood | GSM18444 | 17649 | 68118 |
| 151.1915 | 83 | 164.4908 | 97 | 94.00431 | 105 | 119.9686 | 125 | KDM1A | SKNO-1 | Bone Marr | GSM18444 | 11244 | 68119 |
| 149.61 | 82 | 241.0395 | 115 | 87.27534 | 101 | 113.2529 | 121 | KDM1A | SKNO-1 | Bone Marr | GSM18444 | 10981 | 68120 |
| 168.5082 | 68 | 207.0931 | 84 | 87.66305 | 76 | 84.98514 | 81 | KDM1A | SKNO-1 | Bone Marr | GSM18444 | 6791 | 68121 |
| 100.2727 | 56 | 141.5308 | 74 | 110.695 | 90 | 110.5984 | 97 | GATA3 | None | Thymus | GSM18446 | 7870 | 68160 |
| 71.50089 | 46 | 105.4812 | 62 | 66.87445 | 70 | 102.524 | 90 | GATA3 | None | Thymus | GSM18446 | 7303 | 68161 |
| 34.52859 | 18 | 75.40625 | 30 | 31.431 | 27 | 124.6725 | 55 | TP63 | None | None | GSM20747 | 2633 | 68211 |
| 827.5461 | 135 | 797.8539 | 144 | 364.1902 | 118 | 795.654 | 190 | AR | LNCaP | Prostate | GSM20588 | 4272 | 68217 |
| 1039.147 | 290 | 962.9986 | 288 | 842.9837 | 331 | 915.4748 | 457 | AR | LNCaP | Prostate | GSM20588 | 11082 | 68218 |
| 896.6463 | 500 | 815.4269 | 504 | 716.6034 | 628 | 759.812 | 791 | FOXA1 | LNCaP | Prostate | GSM20588 | 38060 | 68219 |
| 917.7675 | 341 | 922.4174 | 420 | 717.2789 | 391 | 687.0231 | 412 | NR3C1 | BEAS-2B | Lung | GSM21030 | 20923 | 68222 |
| 914.5945 | 399 | 933.9912 | 511 | 692.1812 | 441 | 674.932 | 478 | NR3C1 | BEAS-2B | Lung | GSM21030 | 25818 | 68223 |

| | | | | | | | | | | | | | |
|---|---|---|---|---|---|---|---|---|---|---|---|---|---|
| 185.0921 | 86 | 273.5534 | 116 | 189.0759 | 130 | 195.6 | 142 | RELA | BEAS-2B | Lung | GSM21030 | 9602 | 68224 |
| 167.7883 | 68 | 242.4828 | 91 | 128.3135 | 89 | 165.0761 | 107 | RELA | BEAS-2B | Lung | GSM21030 | 6492 | 68225 |
| 128.7695 | 54 | 118.8861 | 58 | 91.09129 | 69 | 144.4781 | 91 | RELA | BEAS-2B | Lung | GSM21030 | 5532 | 68226 |
| 812.3941 | 360 | 862.9951 | 496 | 633.3822 | 439 | 635.8569 | 502 | NR3C1 | BEAS-2B | Lung | GSM21030 | 30795 | 68227 |
| 798.4796 | 262 | 830.5355 | 342 | 525.1344 | 316 | 583.149 | 356 | NR3C1 | BEAS-2B | Lung | GSM21030 | 21697 | 68228 |
| 798.0223 | 387 | 839.3341 | 516 | 624.3618 | 480 | 618.7185 | 539 | NR3C1 | BEAS-2B | Lung | GSM21030 | 35693 | 68229 |
| 813.3959 | 386 | 845.4996 | 506 | 638.597 | 479 | 631.8089 | 537 | NR3C1 | BEAS-2B | Lung | GSM21030 | 34682 | 68230 |
| 866.0505 | 167 | 887.3633 | 185 | 578.8477 | 185 | 888.1855 | 270 | FOXA1 | VCaP | Prostate | GSM20588 | 6315 | 68234 |
| 1137.098 | 170 | 780.3732 | 143 | 999.3297 | 217 | 1028 | 268 | FOXA1 | VCaP | Prostate | GSM20588 | 3981 | 68235 |
| 77.74832 | 30 | 119.9913 | 42 | 23.5283 | 27 | 165.4354 | 70 | POU2F1 | LNCaP | Prostate | GSM20588 | 3270 | 68236 |
| 769.0422 | 355 | 747.3159 | 409 | 654.0093 | 508 | 694.3082 | 640 | AR | VCaP | Prostate | GSM20588 | 33469 | 68239 |
| 750.8383 | 378 | 731.9501 | 436 | 650.1826 | 559 | 702.296 | 725 | AR | VCaP | Prostate | GSM20588 | 38321 | 68240 |
| 777.9848 | 399 | 787.3111 | 500 | 553.5917 | 458 | 546.1684 | 500 | NR4A1 | Kasumi-1 | None | GSM20959 | 44783 | 68309 |
| 678.0586 | 309 | 719.4119 | 388 | 363.364 | 368 | 369.8452 | 402 | NR4A1 | Kasumi-1 | None | GSM20959 | 40224 | 68310 |
| 937.0866 | 721 | 902.0163 | 822 | 777.5128 | 997 | 804.2405 | 1196 | ESR1 | LY2 | Breast | GSM15569 | 69398 | 68323 |
| 1000.437 | 585 | 1009.415 | 718 | 849.6384 | 809 | 858.9229 | 939 | ESR1 | LY2 | Breast | GSM15569 | 38385 | 68324 |
| 6.810499 | 29 | 1.702523 | 25 | 1.004383 | 36 | 2.557457 | 47 | DUX4 | LHCN-M2 | Skeletal Mu | GSM20683 | 12084 | 68379 |
| 978.1725 | 291 | 924.3956 | 309 | 804.7593 | 360 | 897.2237 | 528 | AR | C4-2B | Prostate | GSM18688 | 14820 | 68399 |
| 780.5491 | 166 | 849.4619 | 191 | 587.0296 | 201 | 894.4874 | 319 | AR | C4-2B | Prostate | GSM18688 | 8107 | 68401 |
| 1238.626 | 640 | 1295.987 | 840 | 916.2172 | 612 | 866.7321 | 622 | USF2 | None | None | GSM16983 | 21078 | 68455 |
| 1251.094 | 784 | 1290.501 | 990 | 902.0725 | 773 | 852.1405 | 796 | USF1 | None | None | GSM16983 | 30840 | 68456 |
| 882.9766 | 331 | 881.7652 | 401 | 680.7569 | 387 | 690.8702 | 451 | ATF3 | HCT-116 | Colon | GSM19177 | 25135 | 68460 |
| 12.43137 | 10 | 9.233831 | 10 | 4.650124 | 11 | 41.42651 | 29 | TP53 | HCT-116 | Colon | GSM19177 | 1892 | 68462 |
| 855.5069 | 553 | 868.1605 | 689 | 649.4396 | 679 | 654.0608 | 781 | ATF3 | HCT-116 | Colon | GSM19177 | 57573 | 68464 |
| 421.8643 | 58 | 696.5823 | 86 | 213.2835 | 53 | 391.0443 | 79 | RBPJ | None | Brain | GSM21017 | 1495 | 68486 |
| 875.5557 | 119 | 1225.257 | 182 | 391.8697 | 102 | 474.8831 | 120 | RBPJ | None | Brain | GSM21017 | 3010 | 68487 |
| 1192.426 | 166 | 1211.5 | 213 | 753.7678 | 161 | 635.7849 | 155 | KAT5 | K562 | None | GSM17052 | 3680 | 68490 |
| 743.0909 | 121 | 1119.257 | 174 | 315.1792 | 105 | 441.1255 | 133 | POU3F2 | None | None | GSM19344 | 4206 | 68569 |
| 666.4096 | 115 | 932.5389 | 153 | 333.8593 | 110 | 433.7887 | 134 | POU3F2 | None | None | GSM19344 | 4363 | 68570 |
| 92.80273 | 21 | 161.5153 | 32 | 23.44156 | 15 | 187.1568 | 47 | POU3F2 | None | None | GSM19344 | 1229 | 68572 |
| 18.43106 | 9 | 49.21275 | 17 | 10.70405 | 11 | 90.98496 | 33 | SMAD3 | BJ | Skin | GSM16684 | 1275 | 68573 |
| 1059.745 | 244 | 1075.2 | 310 | 842.5176 | 256 | 807.8835 | 265 | KMT2D | None | None | GSM18469 | 8231 | 68627 |
| 1102.071 | 331 | 1135.551 | 442 | 835.8739 | 315 | 835.5198 | 355 | KMT2D | None | None | GSM18468 | 10504 | 68632 |
| 1193.611 | 298 | 1238.718 | 400 | 939.096 | 280 | 901.1929 | 287 | KMT2D | None | None | GSM18468 | 6343 | 68635 |
| 1002.271 | 279 | 1027.026 | 361 | 768.1174 | 285 | 769.1901 | 326 | KMT2D | None | None | GSM18468 | 12104 | 68638 |
| 1004.933 | 591 | 1029.313 | 749 | 722.9069 | 605 | 713.9486 | 678 | BRD4 | IMR90 | None | GSM20981 | 36711 | 68711 |
| 1029.268 | 412 | 1074.74 | 554 | 805.3698 | 459 | 765.7171 | 467 | BRD4 | IMR90 | None | GSM20981 | 19583 | 68712 |
| 442.654 | 107 | 355.157 | 105 | 241.3183 | 114 | 308.2023 | 137 | BRD4 | IMR90 | None | GSM19151 | 6676 | 68713 |
| 1035.382 | 197 | 1016.906 | 233 | 608.0891 | 201 | 659.7102 | 223 | BRD4 | IMR90 | None | GSM19151 | 8276 | 68714 |
| 901.0815 | 231 | 941.0056 | 316 | 545.8678 | 255 | 611.7464 | 288 | BRD4 | IMR90 | None | GSM19151 | 14583 | 68715 |
| 444.6442 | 131 | 497.5751 | 154 | 150.6075 | 117 | 280.3571 | 165 | TOP1 | HCT-116 | Colon | GSM20586 | 10160 | 68774 |
| 211.344 | 57 | 205.0946 | 62 | 127.2437 | 64 | 158.1904 | 76 | TOP1 | HCT-116 | Colon | GSM20586 | 3805 | 68775 |
| 1073.217 | 189 | 1083.093 | 241 | 692.2709 | 197 | 652.3487 | 203 | BRD4 | HCT-116 | Colon | GSM20586 | 6616 | 68777 |
| 352.8989 | 71 | 445.2835 | 89 | 174.6639 | 69 | 186.3835 | 76 | TOP1 | HCT-116 | Colon | GSM20586 | 3280 | 68780 |
| 341.4162 | 94 | 411.9933 | 115 | 198.5958 | 105 | 224.3164 | 119 | TOP1 | HCT-116 | Colon | GSM20586 | 6675 | 68781 |
| 596.3643 | 98 | 973.0471 | 143 | 451.0184 | 115 | 328.9837 | 103 | BRD4 | HCT-116 | Colon | GSM20586 | 3289 | 68782 |
| 1077.605 | 264 | 1055.756 | 302 | 837.2818 | 269 | 821.3204 | 291 | BRD4 | HCT-116 | Colon | GSM20586 | 9455 | 68783 |
| 1.905056 | 24 | 4.105215 | 34 | 0.642198 | 40 | 3.501724 | 58 | SATB1 | SH-SY5Y | None | GSM18981 | 16406 | 68797 |
| 1.54503 | 20 | 8.100153 | 36 | 1.851425 | 41 | 15.64192 | 72 | SATB1 | SH-SY5Y | None | GSM18981 | 14204 | 68798 |
| 82.01986 | 75 | 154.3752 | 111 | 138.0169 | 146 | 133.262 | 156 | GATA2 | None | Bone Marr | GSM14887 | 15252 | 68819 |
| 151.8939 | 54 | 238.9579 | 76 | 114.3344 | 70 | 124.1664 | 78 | GATA2 | None | Bone Marr | GSM14887 | 4914 | 68820 |
| 614.2072 | 121 | 932.5852 | 168 | 373.7112 | 129 | 324.2136 | 128 | HDAC2 | None | Bone Marr | GSM14887 | 5008 | 68821 |
| 1141.726 | 232 | 1164.074 | 295 | 826.1886 | 214 | 804.1896 | 224 | HDAC1 | None | Bone Marr | GSM14887 | 6187 | 68822 |
| 373.7536 | 218 | 504.4338 | 280 | 323.3644 | 320 | 250.2249 | 316 | SPI1 | K562 | None | GSM19402 | 34523 | 68823 |
| 288.6851 | 118 | 341.0573 | 143 | 225.1412 | 159 | 185.7776 | 158 | SPI1 | K562 | None | GSM19401 | 12867 | 68824 |
| 331.932 | 129 | 360.7115 | 150 | 226.21 | 163 | 172.5067 | 157 | SPI1 | K562 | None | GSM19401 | 13384 | 68825 |
| 251.7595 | 118 | 283.7937 | 140 | 194.6296 | 161 | 171.1884 | 165 | SPI1 | K562 | None | GSM19401 | 14643 | 68826 |
| 1152.348 | 232 | 1166.156 | 297 | 887.5987 | 222 | 807.5624 | 224 | HEXIM1 | A375 | None | GSM13824 | 6510 | 68827 |
| 1052.578 | 574 | 1070.302 | 714 | 758.2912 | 569 | 733.8014 | 611 | HEXIM1 | A375 | None | GSM13824 | 31168 | 68828 |
| 177.8619 | 32 | 186.9844 | 36 | 130.0345 | 37 | 116.4246 | 37 | CDK9 | A375 | None | GSM13824 | 1179 | 68830 |
| 463.025 | 75 | 484.5244 | 84 | 249.2515 | 73 | 316.0449 | 88 | FOXM1 | VAL-3 | None | GSM21011 | 2469 | 68837 |
| 888.8163 | 520 | 821.9249 | 544 | 718.869 | 678 | 725.1031 | 783 | FOXA1 | T47D | Breast | GSM18586 | 43987 | 68838 |
| 912.0162 | 469 | 850.9532 | 491 | 738.932 | 593 | 738.7427 | 677 | FOXA1 | T47D | Breast | GSM18586 | 33213 | 68839 |
| 912.6619 | 579 | 833.7198 | 600 | 721.1162 | 726 | 736.03 | 861 | FOXA1 | T47D | Breast | GSM18586 | 48149 | 68840 |
| 918.3054 | 470 | 848.4666 | 487 | 731.0701 | 578 | 731.9387 | 661 | FOXA1 | T47D | Breast | GSM18586 | 32932 | 68841 |
| 940.6399 | 685 | 889.7905 | 757 | 734.336 | 857 | 727.3433 | 967 | FOXA1 | T47D | Breast | GSM18586 | 59352 | 68842 |
| 903.7512 | 490 | 822.3354 | 494 | 702.161 | 584 | 710.8128 | 680 | FOXA1 | T47D | Breast | GSM18586 | 36495 | 68843 |
| 685.6806 | 195 | 578.0156 | 198 | 758.4097 | 302 | 774.7754 | 360 | ESR1 | T47D | Breast | GSM18586 | 13570 | 68844 |
| 872.8919 | 273 | 821.9273 | 292 | 758.9686 | 394 | 769.2728 | 462 | ESR1 | T47D | Breast | GSM18586 | 18128 | 68845 |
| 1043.201 | 342 | 1046.285 | 418 | 873.6449 | 425 | 872.4244 | 484 | ESR1 | T47D | Breast | GSM18586 | 13908 | 68846 |
| 931.1746 | 261 | 875.6606 | 274 | 807.7988 | 365 | 811.9341 | 423 | ESR1 | T47D | Breast | GSM18586 | 14331 | 68847 |
| 221.6477 | 47 | 209.9483 | 50 | 324.8623 | 81 | 405.9017 | 97 | NR3C1 | T47D | Breast | GSM18586 | 2198 | 68848 |

| | | | | | | | | | | | | | |
|---|---|---|---|---|---|---|---|---|---|---|---|---|---|
| 236.8256 | 50 | 180.0143 | 47 | 332.3286 | 84 | 511.5413 | 113 | NR3C1 | T47D | Breast | GSM18586 | 2309 | 68849 |
| 814.7033 | 638 | 736.6088 | 670 | 631.0968 | 828 | 647.5441 | 979 | FOXA1 | ZR-75-1 | Breast | GSM18586 | 76176 | 68852 |
| 788.4973 | 531 | 727.4703 | 563 | 631.1097 | 703 | 645.1699 | 833 | FOXA1 | ZR-75-1 | Breast | GSM18586 | 58741 | 68853 |
| 794.5363 | 618 | 729.197 | 658 | 616.4496 | 800 | 629.0605 | 943 | FOXA1 | ZR-75-1 | Breast | GSM18586 | 75168 | 68854 |
| 822.9139 | 658 | 761.0431 | 708 | 633.6364 | 844 | 648.4166 | 993 | FOXA1 | ZR-75-1 | Breast | GSM18586 | 77725 | 68855 |
| 789.5904 | 582 | 722.5703 | 619 | 612.683 | 755 | 632.375 | 902 | FOXA1 | ZR-75-1 | Breast | GSM18586 | 70174 | 68856 |
| 843.2436 | 712 | 755.1197 | 739 | 630.9362 | 886 | 634.0728 | 1019 | FOXA1 | ZR-75-1 | Breast | GSM18586 | 84896 | 68857 |
| 794.712 | 350 | 740.6733 | 373 | 684.647 | 518 | 719.7249 | 645 | ESR1 | ZR-75-1 | Breast | GSM18586 | 33767 | 68858 |
| 786.3646 | 404 | 716.6204 | 415 | 661.819 | 580 | 687.2317 | 704 | ESR1 | ZR-75-1 | Breast | GSM18586 | 41922 | 68859 |
| 843.3787 | 350 | 791.3802 | 377 | 719.8324 | 500 | 737.7836 | 594 | ESR1 | ZR-75-1 | Breast | GSM18586 | 28350 | 68860 |
| 860.7515 | 301 | 793.2826 | 309 | 722.2866 | 411 | 757.0386 | 514 | ESR1 | ZR-75-1 | Breast | GSM18586 | 22166 | 68861 |
| 817.8495 | 334 | 748.4772 | 340 | 670.5034 | 446 | 700.4821 | 549 | NR3C1 | ZR-75-1 | Breast | GSM18586 | 28694 | 68862 |
| 787.0152 | 294 | 712.7236 | 304 | 663.0498 | 415 | 693.7523 | 513 | NR3C1 | ZR-75-1 | Breast | GSM18586 | 27045 | 68863 |
| 702.2118 | 166 | 592.1319 | 167 | 636.2986 | 223 | 803.9527 | 276 | NR3C1 | ZR-75-1 | Breast | GSM18586 | 9187 | 68864 |
| 504.6524 | 122 | 481.5223 | 131 | 536.8659 | 178 | 693.4035 | 216 | NR3C1 | ZR-75-1 | Breast | GSM18586 | 6828 | 68865 |
| 865.7265 | 620 | 795.3443 | 658 | 713.7325 | 854 | 717.9272 | 988 | FOXA1 | MCF-7 | Breast | GSM18586 | 63206 | 68866 |
| 899.2028 | 635 | 851.3139 | 700 | 749.5821 | 877 | 741.3312 | 985 | FOXA1 | MCF-7 | Breast | GSM18586 | 58205 | 68867 |
| 839.5462 | 550 | 784.8613 | 600 | 709.693 | 789 | 714.9966 | 915 | FOXA1 | MCF-7 | Breast | GSM18586 | 56783 | 68868 |
| 988.1839 | 726 | 944.352 | 816 | 792.41 | 939 | 789.6517 | 1071 | FOXA1 | MCF-7 | Breast | GSM18586 | 57157 | 68869 |
| 842.2706 | 564 | 805.347 | 637 | 698.3783 | 785 | 713.7218 | 928 | FOXA1 | MCF-7 | Breast | GSM18586 | 58406 | 68870 |
| 1025.107 | 727 | 997.8236 | 832 | 806.0416 | 891 | 805.8214 | 1019 | FOXA1 | MCF-7 | Breast | GSM18586 | 49493 | 68871 |
| 136.7416 | 55 | 85.81568 | 49 | 88.82128 | 67 | 146.5457 | 90 | ESR1 | MCF-7 | Breast | GSM18586 | 5242 | 68872 |
| 426.2201 | 148 | 372.4135 | 154 | 728.7266 | 278 | 774.6084 | 337 | ESR1 | MCF-7 | Breast | GSM18586 | 12607 | 68873 |
| 385.4655 | 80 | 284.4354 | 74 | 530.6885 | 132 | 610.6694 | 151 | ESR1 | MCF-7 | Breast | GSM18586 | 3511 | 68874 |
| 563.9865 | 142 | 469.1432 | 142 | 796.6321 | 239 | 847.9539 | 290 | ESR1 | MCF-7 | Breast | GSM18586 | 8147 | 68875 |
| 686.4199 | 229 | 607.8365 | 240 | 717.0683 | 363 | 768.7074 | 474 | NR3C1 | MCF-7 | Breast | GSM18586 | 18994 | 68876 |
| 667.263 | 205 | 557.4444 | 208 | 762.8967 | 343 | 794.6798 | 428 | NR3C1 | MCF-7 | Breast | GSM18586 | 15494 | 68877 |
| 423.8614 | 95 | 414.1978 | 103 | 232.338 | 98 | 238.3249 | 106 | NR3C1 | MCF-7 | Breast | GSM18586 | 4669 | 68878 |
| 699.1923 | 119 | 886.1862 | 149 | 356.2189 | 113 | 489.9367 | 142 | KDM5C | ZR-75-30 | None | GSM20295 | 4004 | 68913 |
| 27.11458 | 35 | 21.58924 | 37 | 33.25573 | 62 | 101.4133 | 102 | EP300 | ZR-75-30 | None | GSM20295 | 8881 | 68917 |
| 761.155 | 452 | 785.4604 | 575 | 555.0926 | 528 | 557.3392 | 606 | EP300 | ZR-75-30 | None | GSM20295 | 51320 | 68918 |
| 996.5789 | 235 | 982.5491 | 277 | 521.1953 | 213 | 627.0882 | 249 | KDM5C | ZR-75-30 | None | GSM18326 | 9892 | 68932 |
| 14.14654 | 7 | 11.08147 | 7 | 6.974494 | 8 | 20.14376 | 14 | SMARCA4 | MCF-10A | None | GSM19307 | 1043 | 68958 |
| 81.04707 | 19 | 150.2416 | 30 | 15.86244 | 12 | 76.33306 | 28 | SMARCA4 | MCF-10A | None | GSM19307 | 1101 | 68959 |
| 38.63063 | 15 | 64.66728 | 22 | 33.70534 | 21 | 69.54953 | 32 | PRDM14 | NCCIT | None | GSM18427 | 1503 | 68962 |
| 103.8167 | 31 | 96.36329 | 33 | 35.42099 | 27 | 146.2787 | 57 | CBFA2T2 | NCCIT | None | GSM18427 | 2261 | 68963 |
| 1039.767 | 260 | 1070.614 | 346 | 822.2453 | 280 | 797.5037 | 299 | ESR1 | T47D | Breast | GSM21128 | 10075 | 68970 |
| 325.0915 | 103 | 549.0446 | 150 | 200.7065 | 119 | 201.3563 | 128 | ESR1 | T47D | Breast | GSM21128 | 7767 | 68972 |
| 812.5668 | 329 | 811.4094 | 400 | 539.7239 | 368 | 599.8931 | 418 | ESR1 | T47D | Breast | GSM21127 | 28726 | 68974 |
| 868.9084 | 288 | 864.6458 | 351 | 465.5017 | 288 | 423.069 | 298 | ESR1 | T47D | Breast | GSM21127 | 20594 | 68976 |
| 1186.204 | 708 | 1250.702 | 934 | 914.551 | 773 | 873.6935 | 812 | TRIM24 | LNCaP | Prostate | GSM16979 | 26800 | 69201 |
| 1142.313 | 441 | 1164.047 | 560 | 864.2281 | 426 | 830.6635 | 441 | TRIM24 | LNCaP | Prostate | GSM16979 | 14126 | 69202 |
| 1129.894 | 428 | 1158.117 | 550 | 846.7398 | 399 | 817.0152 | 421 | TRIM24 | LNCaP | Prostate | GSM16979 | 13795 | 69203 |
| 858.2119 | 201 | 911.5317 | 242 | 372.7569 | 193 | 435.073 | 222 | POU2F2 | HBL1 | None | GSM20958 | 12383 | 69219 |
| 1038.915 | 384 | 1063.897 | 495 | 741.7714 | 348 | 739.7597 | 393 | POU2F2 | HBL1 | None | GSM20958 | 18131 | 69220 |
| 15.14859 | 9 | 11.65403 | 9 | 0.780396 | 5 | 8.381035 | 12 | CENPT | HuRef | None | GSM17098 | 1431 | 69236 |
| 454.6506 | 59 | 871.6182 | 96 | 306.7342 | 62 | 570.4561 | 94 | CENPT | HuRef | None | GSM17098 | 1223 | 69238 |
| 648.0281 | 410 | 664.2086 | 519 | 381.7726 | 515 | 318.0567 | 531 | SIX2 | None | Kidney | GSM19705 | 67568 | 69246 |
| 366.5747 | 292 | 418.1832 | 349 | 207.36 | 382 | 204.2305 | 416 | SIX2 | None | Kidney | GSM19705 | 57333 | 69247 |
| 32.37511 | 16 | 32.26791 | 18 | 19.82273 | 20 | 112.6797 | 48 | SIX2 | None | Kidney | GSM19705 | 2176 | 69248 |
| 123.4451 | 117 | 129.7529 | 136 | 90.56772 | 169 | 74.1517 | 173 | SIX2 | None | Kidney | GSM19705 | 24639 | 69249 |
| 1165.583 | 643 | 1165.396 | 770 | 822.2266 | 561 | 775.3278 | 572 | EP300 | None | Kidney | GSM19705 | 20988 | 69250 |
| 923.5681 | 148 | 1116.022 | 205 | 745.547 | 179 | 936.2462 | 222 | ERG | VCaP | Prostate | GSM20863 | 4698 | 69266 |
| 1077.305 | 181 | 1099.039 | 235 | 761.5508 | 199 | 903.1759 | 245 | ERG | VCaP | Prostate | GSM20863 | 5813 | 69267 |
| 1079.719 | 299 | 1098.571 | 385 | 879.0403 | 342 | 862.6355 | 371 | ERG | VCaP | Prostate | GSM20863 | 10252 | 69268 |
| 1066.181 | 181 | 1108.436 | 244 | 838.3827 | 212 | 883.8639 | 237 | ERG | VCaP | Prostate | GSM20863 | 5960 | 69269 |
| 550.5328 | 99 | 867.215 | 141 | 492.3408 | 128 | 570.7537 | 147 | ERG | VCaP | Prostate | GSM20863 | 3656 | 69270 |
| 812.0361 | 130 | 1052.86 | 165 | 503.0197 | 136 | 689.5685 | 171 | ERG | VCaP | Prostate | GSM20863 | 4078 | 69271 |
| 1093.17 | 197 | 1094.99 | 243 | 742.0346 | 201 | 819.7596 | 225 | ERG | VCaP | Prostate | GSM20863 | 6115 | 69272 |
| 274.2783 | 94 | 372.5332 | 122 | 183.645 | 114 | 507.7176 | 197 | AR | VCaP | Prostate | GSM20863 | 7762 | 69273 |
| 229.759 | 71 | 248.9846 | 82 | 124.5622 | 77 | 409.6806 | 146 | AR | VCaP | Prostate | GSM20863 | 5243 | 69274 |
| 510.9044 | 190 | 629.9467 | 234 | 346.0275 | 235 | 702.0716 | 365 | AR | VCaP | Prostate | GSM20863 | 17440 | 69275 |
| 137.1678 | 62 | 157.14 | 74 | 134.8195 | 92 | 329.7837 | 149 | AR | VCaP | Prostate | GSM20863 | 6719 | 69276 |
| 183.0601 | 81 | 160.2953 | 85 | 201.8692 | 126 | 446.28 | 195 | AR | VCaP | Prostate | GSM20863 | 8555 | 69277 |
| 196.9892 | 113 | 293.3332 | 152 | 180.6658 | 166 | 394.208 | 248 | AR | VCaP | Prostate | GSM20863 | 15132 | 69278 |
| 262.7056 | 145 | 370.4194 | 190 | 266.5293 | 221 | 597.5974 | 335 | AR | VCaP | Prostate | GSM20863 | 18959 | 69279 |
| 199.288 | 96 | 265.4296 | 123 | 173.8525 | 136 | 400.5424 | 211 | AR | VCaP | Prostate | GSM20863 | 11068 | 69280 |
| 169.2956 | 72 | 171.6635 | 81 | 115.1867 | 91 | 357.4676 | 163 | AR | VCaP | Prostate | GSM20863 | 7637 | 69281 |
| 241.3788 | 122 | 329.4548 | 158 | 203.0323 | 172 | 486.0236 | 269 | AR | VCaP | Prostate | GSM20862 | 15263 | 69282 |
| 170.2476 | 70 | 187.8541 | 82 | 109.7284 | 86 | 307.6367 | 147 | AR | VCaP | Prostate | GSM20862 | 7163 | 69283 |
| 349.2923 | 177 | 451.3799 | 223 | 271.763 | 242 | 627.4173 | 369 | AR | VCaP | Prostate | GSM20862 | 22501 | 69284 |
| 288.4489 | 128 | 345.8734 | 156 | 239.103 | 177 | 500.773 | 263 | AR | VCaP | Prostate | GSM20862 | 14293 | 69285 |

| | | | | | | | | | | | | | |
|---|---|---|---|---|---|---|---|---|---|---|---|---|---|
| 236.1823 | 74 | 306.4061 | 94 | 146.9242 | 86 | 478.8996 | 163 | AR | VCaP | Prostate | GSM20862 | 5692 | 69287 |
| 275.9427 | 97 | 319.169 | 116 | 190.6605 | 120 | 569.69 | 215 | AR | VCaP | Prostate | GSM20862 | 8351 | 69288 |
| 369.7002 | 132 | 496.8599 | 170 | 279.6227 | 170 | 722.521 | 283 | AR | VCaP | Prostate | GSM20862 | 11412 | 69289 |
| 118.1439 | 34 | 132.5809 | 40 | 87.20385 | 42 | 156.125 | 60 | AR | VCaP | Prostate | GSM20862 | 2288 | 69290 |
| 783.9011 | 640 | 678.6554 | 632 | 637.3274 | 882 | 662.2858 | 1059 | FOXA1 | LNCaP | Prostate | GSM18630 | 78907 | 69365 |
| 155.145 | 48 | 148.6397 | 52 | 77.18657 | 50 | 145.0965 | 72 | TCF7L1 | Ramos | None | GSM18732 | 3571 | 69437 |
| 236.9845 | 101 | 232.5412 | 112 | 141.8127 | 122 | 170.5106 | 142 | TP53 | None | Foreskin | GSM20462 | 11232 | 69456 |
| 148.9716 | 54 | 119.964 | 54 | 67.03389 | 56 | 107.4079 | 74 | TP53 | None | Foreskin | GSM20462 | 5044 | 69457 |
| 277.1629 | 47 | 499.4752 | 73 | 178.9844 | 50 | 425.5217 | 86 | INO80 | SKmel147 | None | GSM21905 | 1485 | 69516 |
| 72.00712 | 28 | 138.9939 | 44 | 59.21525 | 38 | 80.37257 | 47 | MED1 | A375 | None | GSM21905 | 2719 | 69517 |
| 91.12695 | 27 | 75.28167 | 27 | 10.92228 | 15 | 233.9578 | 68 | MED1 | A375 | None | GSM21905 | 1940 | 69518 |
| 385.9145 | 141 | 633.9884 | 201 | 232.6787 | 166 | 323.6179 | 206 | RUNX1 | Kasumi-1 | Blood | GSM21503 | 12957 | 69541 |
| 71.78869 | 25 | 104.8664 | 34 | 68.18112 | 36 | 111.5588 | 49 | RUNX1 | Kasumi-1 | Blood | GSM21503 | 2213 | 69542 |
| 366.43 | 54 | 432.6865 | 65 | 166.1299 | 47 | 422.4842 | 84 | ASH1L | MV4-11 | None | GSM18973 | 1547 | 69557 |
| 649.0395 | 147 | 776.6962 | 178 | 265.5123 | 132 | 284.1366 | 146 | PSIP1 | MV4-11 | None | GSM18973 | 7157 | 69558 |
| 865.8982 | 447 | 853.3515 | 526 | 608.9005 | 459 | 591.7725 | 499 | BCLAF1 | GM12878 | Blood | GSM80350 | 36680 | 6958 |
| 1085.985 | 455 | 1108.169 | 569 | 799.8748 | 423 | 765.3862 | 442 | CDK7 | Jurkat | Blood | GSM22187 | 17205 | 69615 |
| 1116.56 | 474 | 1153.09 | 617 | 835.8828 | 459 | 782.0018 | 449 | MYC | Jurkat | Blood | GSM22187 | 17247 | 69616 |
| 1004.731 | 531 | 1030.671 | 671 | 726.8151 | 518 | 654.6429 | 491 | BRD4 | Jurkat | Blood | GSM22187 | 26136 | 69617 |
| 852.2665 | 195 | 787.9967 | 206 | 505.2875 | 213 | 429.0215 | 211 | STAG1 | None | None | GSM21406 | 10723 | 69618 |
| 795.4534 | 407 | 845.8731 | 554 | 593.6763 | 482 | 589.952 | 543 | CTCF | None | None | GSM21406 | 41950 | 69619 |
| 477.5015 | 277 | 484.4785 | 314 | 228.191 | 323 | 241.6686 | 359 | STAG1 | None | None | GSM21406 | 39839 | 69620 |
| 654.6136 | 317 | 714.6728 | 409 | 310.5092 | 360 | 360.6651 | 413 | CTCF | None | None | GSM21406 | 40219 | 69621 |
| 870.6461 | 457 | 911.6534 | 606 | 642.8375 | 509 | 625.2902 | 555 | STAG1 | None | None | GSM16557 | 37784 | 69623 |
| 831.313 | 275 | 856.5514 | 359 | 479.7239 | 299 | 518.3439 | 333 | CTCF | None | None | GSM16557 | 20777 | 69624 |
| 476.8536 | 283 | 532.7938 | 334 | 185.9405 | 310 | 193.6028 | 343 | STAG1 | None | None | GSM16557 | 41770 | 69626 |
| 580.4494 | 283 | 717.6327 | 376 | 330.2508 | 346 | 343.3647 | 381 | CTCF | None | None | GSM16557 | 37803 | 69627 |
| 1217.126 | 165 | 1230.579 | 203 | 788.0287 | 157 | 657.5505 | 150 | KDM5A | T47D | Breast | GSM21311 | 2967 | 69731 |
| 1235.523 | 148 | 1249.919 | 185 | 702.1465 | 137 | 637.2207 | 137 | KDM5A | T47D | Breast | GSM21311 | 2526 | 69732 |
| 1090.588 | 446 | 1101.237 | 551 | 800.5136 | 414 | 755.0517 | 418 | BRD4 | MV4;11 | Blood | GSM18451 | 17022 | 69755 |
| 992.5081 | 572 | 1004.062 | 705 | 709.8385 | 574 | 682.0891 | 612 | BRD4 | MV4;11 | Blood | GSM18451 | 34726 | 69758 |
| 1047.37 | 583 | 1061.828 | 728 | 742.0432 | 563 | 712.2703 | 595 | BRD4 | MV4;11 | Blood | GSM18451 | 30283 | 69761 |
| 1111.168 | 168 | 1138.208 | 222 | 547.5362 | 153 | 570.2817 | 166 | BRD4 | MV4;11 | Blood | GSM18451 | 4818 | 69764 |
| 843.1797 | 588 | 869.8428 | 750 | 579.9354 | 632 | 562.8409 | 693 | BRD4 | MV4;11 | Blood | GSM18451 | 60795 | 69767 |
| 1091.548 | 554 | 1107.273 | 685 | 789.3952 | 523 | 768.6917 | 567 | BRD4 | MV4;11 | Blood | GSM18451 | 23896 | 69770 |
| 951.82 | 577 | 979.8222 | 728 | 676.6928 | 589 | 647.023 | 624 | BRD4 | MV4;11 | Blood | GSM18451 | 39313 | 69773 |
| 789.522 | 304 | 778.7133 | 360 | 631.8383 | 393 | 654.7103 | 478 | JUND | HT29 | None | GSM20429 | 28523 | 69774 |
| 840.0609 | 301 | 814.4993 | 344 | 606.2324 | 346 | 686.3622 | 443 | JUND | HT29 | None | GSM20429 | 23152 | 69775 |
| 290.9225 | 135 | 294.1672 | 152 | 343.4918 | 218 | 474.2677 | 270 | PPARG | HT29 | None | GSM20428 | 15741 | 69795 |
| 622.6082 | 251 | 739.0686 | 305 | 642.5247 | 380 | 641.6439 | 433 | PPARG | HT29 | None | GSM20428 | 26058 | 69796 |
| 868.7418 | 457 | 881.1242 | 566 | 720.8167 | 618 | 744.0417 | 745 | PPARG | HT29 | None | GSM20428 | 38698 | 69797 |
| 832.4208 | 467 | 829.911 | 565 | 693.2448 | 653 | 689.3363 | 740 | PPARG | HT29 | None | GSM20428 | 45391 | 69798 |
| 919.3271 | 649 | 934.0702 | 804 | 736.1905 | 846 | 745.9632 | 988 | NR1H3 | HT29 | None | GSM20428 | 57870 | 69799 |
| 880.5 | 477 | 911.3248 | 615 | 737.5191 | 653 | 747.4476 | 766 | NR1H3 | HT29 | None | GSM20428 | 39137 | 69800 |
| 885.3445 | 471 | 881.435 | 566 | 767.7414 | 682 | 773.4268 | 788 | NR1H3 | HT29 | None | GSM20428 | 37565 | 69801 |
| 961.921 | 684 | 934.9444 | 790 | 783.1505 | 904 | 796.3666 | 1062 | NR1H3 | HT29 | None | GSM20428 | 55399 | 69802 |
| 955.113 | 518 | 934.2297 | 606 | 760.1667 | 638 | 745.4935 | 707 | NR1H2 | HT29 | None | GSM20428 | 35294 | 69803 |
| 876.9849 | 386 | 869.5629 | 462 | 729.5488 | 518 | 729.1081 | 588 | NR1H2 | HT29 | None | GSM20428 | 29510 | 69804 |
| 748.7829 | 196 | 862.0985 | 233 | 738.9317 | 278 | 755.1526 | 316 | NR1H2 | HT29 | None | GSM20428 | 12640 | 69805 |
| 929.2234 | 308 | 906.8207 | 357 | 770.3683 | 398 | 789.4486 | 481 | NR1H2 | HT29 | None | GSM20428 | 18557 | 69806 |
| 1253.029 | 386 | 1276.932 | 490 | 963.9965 | 347 | 924.7613 | 355 | NELFE | MCF-7 | Breast | GSM21516 | 7905 | 69807 |
| 1203.744 | 186 | 1225.257 | 241 | 875.1461 | 182 | 955.953 | 203 | NELFE | MCF-7 | Breast | GSM21516 | 3847 | 69808 |
| 1261.841 | 376 | 1278.941 | 467 | 980.713 | 344 | 945.0569 | 353 | NELFE | MCF-7 | Breast | GSM21516 | 7546 | 69809 |
| 1244.517 | 267 | 1275.915 | 356 | 968.0691 | 246 | 967.0649 | 278 | NELFE | MCF-7 | Breast | GSM21516 | 5346 | 69810 |
| 1234.797 | 250 | 1271.115 | 326 | 972.0411 | 232 | 969.1266 | 259 | NELFE | MCF-7 | Breast | GSM21516 | 4878 | 69811 |
| 1245.032 | 279 | 1270.627 | 358 | 954.8294 | 243 | 924.3307 | 253 | NELFE | MCF-7 | Breast | GSM21516 | 5443 | 69812 |
| 1259.961 | 383 | 1296.544 | 497 | 992.1283 | 361 | 962.0117 | 378 | NELFE | MCF-7 | Breast | GSM21516 | 7497 | 69813 |
| 1248.558 | 291 | 1288.604 | 388 | 998.3217 | 283 | 969.5419 | 298 | NELFE | MCF-7 | Breast | GSM21516 | 5614 | 69814 |
| 1270.076 | 392 | 1306.269 | 509 | 977.9044 | 352 | 963.3264 | 385 | NELFE | MCF-7 | Breast | GSM21516 | 7658 | 69815 |
| 1262.75 | 388 | 1289.771 | 504 | 965.6197 | 347 | 924.4851 | 347 | NELFE | MCF-7 | Breast | GSM21516 | 7602 | 69816 |
| 535.373 | 77 | 1030.808 | 125 | 470.4252 | 96 | 500.442 | 105 | NELFE | MCF-7 | Breast | GSM21516 | 1980 | 69817 |
| 1238.577 | 351 | 1260.289 | 450 | 964.6675 | 326 | 940.5085 | 345 | NELFE | MCF-7 | Breast | GSM21516 | 7346 | 69818 |
| 1254.865 | 294 | 1264.044 | 363 | 990.4892 | 276 | 953.305 | 283 | NELFE | MCF-7 | Breast | GSM21516 | 5640 | 69819 |
| 1205.335 | 388 | 1217.785 | 483 | 907.8918 | 349 | 894.2908 | 379 | NELFE | MCF-7 | Breast | GSM21516 | 9538 | 69820 |
| 827.835 | 332 | 756.0796 | 338 | 720.0375 | 491 | 735.9508 | 584 | FOXA1 | MCF-7 | None | GSM21377 | 28034 | 69865 |
| 1214.869 | 131 | 1304.682 | 175 | 635.1379 | 118 | 741.3436 | 136 | CDK9 | None | None | GSM19615 | 2285 | 69866 |
| 1315.137 | 457 | 1354.752 | 605 | 999.988 | 411 | 974.253 | 433 | MED1 | MHH-MED- | None | GSM19615 | 9201 | 69868 |
| 1290.024 | 414 | 1309.646 | 525 | 967.6944 | 357 | 934.206 | 371 | MED1 | MHH-MED- | None | GSM19615 | 8755 | 69869 |
| 1267.572 | 618 | 1308.261 | 791 | 927.6043 | 559 | 898.6016 | 591 | BRD4 | None | None | GSM19615 | 16971 | 69870 |
| 1192.798 | 563 | 1226.704 | 728 | 887.7547 | 544 | 842.1981 | 551 | BRD4 | None | None | GSM19615 | 18816 | 69871 |
| 1274.521 | 510 | 1324.988 | 687 | 954.8479 | 467 | 919.5796 | 483 | AFF4 | None | None | GSM19615 | 12496 | 69872 |
| 1244.939 | 542 | 1281.915 | 704 | 913.8352 | 485 | 879.6587 | 501 | AFF4 | None | None | GSM19615 | 15038 | 69873 |

| | | | | | | | | | | | | | |
|---|---|---|---|---|---|---|---|---|---|---|---|---|---|
| 1244.936 | 596 | 1298.798 | 783 | 941.092 | 576 | 895.2856 | 585 | RELA | None | None | GSM19615 | 17099 | 69874 |
| 1078.726 | 467 | 1108.252 | 607 | 808.325 | 473 | 732.1675 | 442 | RELA | None | None | GSM19615 | 20902 | 69875 |
| 1110.466 | 464 | 1181.251 | 653 | 857.5979 | 494 | 815.801 | 501 | CDK9 | None | None | GSM15277 | 18533 | 69878 |
| 1139.171 | 530 | 1179.485 | 686 | 844.6812 | 515 | 812.6583 | 537 | CDK9 | None | None | GSM15277 | 20426 | 69879 |
| 1064.961 | 211 | 1090.352 | 280 | 784.7471 | 226 | 654.4233 | 219 | CDK9 | None | None | GSM15276 | 7816 | 69880 |
| 1089.123 | 360 | 1124.5 | 481 | 830.7397 | 362 | 788.8269 | 369 | CDK9 | None | None | GSM15276 | 13835 | 69881 |
| 1059.719 | 524 | 1104.031 | 693 | 795.8613 | 546 | 729.8385 | 528 | T | None | None | GSM15276 | 26147 | 69884 |
| 45.63714 | 18 | 97.13604 | 30 | 64.72909 | 32 | 88.80251 | 40 | ESR1 | None | endometri | GSM21447 | 1876 | 69891 |
| 316.9296 | 76 | 221.4527 | 69 | 231.127 | 92 | 408.5514 | 131 | FOXA1 | None | endometri | GSM21447 | 4325 | 69893 |
| 817.0791 | 242 | 814.5507 | 279 | 707.8413 | 334 | 689.8571 | 364 | ESR1 | None | endometri | GSM21447 | 18384 | 69894 |
| 859.0609 | 255 | 862.2077 | 312 | 631.8274 | 309 | 654.5047 | 337 | ESR1 | None | endometri | GSM21447 | 18524 | 69896 |
| 109.5292 | 49 | 159.6716 | 66 | 185.4232 | 94 | 282.3416 | 123 | ESR1 | None | endometri | GSM21447 | 5507 | 69899 |
| 191.4102 | 60 | 228.5397 | 73 | 151.4883 | 78 | 216.4667 | 99 | ESR1 | None | endometri | GSM21447 | 4619 | 69902 |
| 42.89094 | 15 | 44.20891 | 17 | 37.80775 | 21 | 212.9402 | 55 | ESR1 | None | endometri | GSM21447 | 1405 | 69905 |
| 7.501727 | 7 | 24.36896 | 14 | 15.76325 | 16 | 61.29813 | 32 | GATA2 | Kasumi-3 | None | GSM22502 | 1796 | 69925 |
| 421.7275 | 73 | 761.4334 | 113 | 340.7868 | 88 | 603.4275 | 128 | INO80 | A549 | Lung | GSM22579 | 2346 | 69945 |
| 660.2789 | 67 | 865.4052 | 86 | 463.7467 | 71 | 691.4365 | 95 | CHD4 | LNCaP | Prostate | GSM18685 | 1041 | 69995 |
| 258.5819 | 49 | 221.2118 | 49 | 119.1896 | 45 | 139.5889 | 52 | SMARCA5 | LNCaP | Prostate | GSM18684 | 2000 | 69998 |
| 299.1966 | 177 | 438.6256 | 236 | 175.1007 | 224 | 252.4654 | 278 | CTCF | KhES-1 | None | GSM19392 | 27271 | 70062 |
| 1361.291 | 887 | 1457.494 | 1164 | 944.4414 | 809 | 904.9078 | 852 | TET2 | 293T | None | GSM18630 | 25376 | 70093 |
| 1313.169 | 846 | 1404.615 | 1112 | 912.4836 | 763 | 886.659 | 827 | TET2 | 293T | None | GSM18630 | 25989 | 70095 |
| 1.926811 | 46 | 1.449315 | 53 | 4.233173 | 99 | 3.380194 | 107 | TET2 | 293T | None | GSM18630 | 30607 | 70097 |
| -0.060763 | 32 | 0.092937 | 52 | 0.039844 | 84 | 3.249291 | 133 | TET2 | 293T | None | GSM18630 | 39813 | 70098 |
| 0.671371 | 35 | 0.167066 | 37 | 1.683304 | 77 | 4.033768 | 98 | TET2 | 293T | None | GSM18630 | 26970 | 70099 |
| -0.104406 | 8 | -0.044083 | 11 | -0.786841 | 14 | 0.866317 | 38 | TET2 | 293T | None | GSM18630 | 11874 | 70100 |
| 1026.837 | 672 | 930.9541 | 683 | 832.8928 | 869 | 880.7535 | 1085 | FOXA1 | LNCaP | Prostate | GSM18918 | 46379 | 70106 |
| 1008.95 | 764 | 931.0506 | 815 | 825.4739 | 1029 | 867.0411 | 1257 | FOXA1 | LNCaP | Prostate | GSM18918 | 63869 | 70108 |
| 115.2775 | 84 | 151.2829 | 107 | 67.02927 | 109 | 117.3313 | 145 | NEUROD1 | NCI-H524 | None | GSM17006 | 15063 | 70118 |
| 116.6247 | 94 | 166.6884 | 124 | 58.79178 | 118 | 92.72709 | 150 | ASCL1 | NCI-H82 | None | GSM17006 | 18171 | 70119 |
| 418.1685 | 204 | 629.3998 | 276 | 417.2176 | 311 | 411.6599 | 334 | ASCL1 | NCI-H889 | None | GSM17006 | 27341 | 70120 |
| 295.8141 | 136 | 425.5611 | 181 | 224.7202 | 184 | 291.7064 | 221 | ASCL1 | NCI-H2107 | None | GSM17006 | 16732 | 70121 |
| 145.147 | 62 | 198.8266 | 81 | 113.5876 | 84 | 195.9696 | 115 | ASCL1 | H128 | Lung | GSM17006 | 6956 | 70122 |
| 1026.013 | 266 | 1024.796 | 323 | 736.3722 | 255 | 674.6361 | 261 | KDM2B | OCI-Ly1 | Bone Marr | GSM21716 | 9390 | 70133 |
| 643.634 | 101 | 546.1551 | 100 | 505.4458 | 121 | 823.2317 | 168 | PIAS1 | MDA-MB-4 | Breast | GSM17178 | 3301 | 70139 |
| 840.7295 | 256 | 779.5188 | 270 | 670.1639 | 332 | 748.6233 | 459 | AR | MDA-MB-4 | Breast | GSM17178 | 20916 | 70140 |
| 803.2411 | 454 | 738.151 | 478 | 645.2982 | 611 | 714.6896 | 815 | AR | MDA-MB-4 | Breast | GSM17178 | 49798 | 70141 |
| 773.4703 | 418 | 744.3566 | 476 | 658.1496 | 619 | 731.0938 | 834 | AR | MDA-MB-4 | Breast | GSM17178 | 48458 | 70142 |
| 842.411 | 288 | 786.7559 | 304 | 700.1832 | 394 | 766.1809 | 536 | AR | MDA-MB-4 | Breast | GSM17178 | 24076 | 70143 |
| 104.0615 | 168 | 111.8264 | 198 | 90.49854 | 266 | 81.52996 | 284 | TEAD4 | BE2-C | Bone Marr | GSM22329 | 46436 | 70162 |
| 765.8822 | 100 | 891.3138 | 119 | 420.3266 | 95 | 365.6333 | 93 | DNMT3B | None | Foreskin | GSM16073 | 2337 | 70174 |
| 675.0642 | 130 | 613.0786 | 135 | 400.736 | 138 | 318.1669 | 131 | DNMT3A | None | Foreskin | GSM16073 | 5667 | 70175 |
| 1099.463 | 178 | 1121.402 | 229 | 533.2109 | 158 | 545.8897 | 170 | DNMT3B | None | Foreskin | GSM16073 | 5401 | 70178 |
| 567.0891 | 88 | 598.8656 | 99 | 410.0626 | 100 | 309.9382 | 91 | DNMT3A | None | Foreskin | GSM16073 | 2710 | 70179 |
| 15.34919 | 62 | 20.66587 | 79 | 8.263442 | 97 | 15.99038 | 124 | TRIM28 | WIBR3 | Embryo | GSM22328 | 27206 | 70221 |
| 2.083905 | 13 | 1.774236 | 15 | 0.417813 | 19 | 1.837998 | 27 | TRIM28 | WIBR3 | Embryo | GSM22328 | 7483 | 70222 |
| 120.1951 | 25 | 316.3114 | 49 | 148.9074 | 40 | 318.3163 | 65 | DNMT3A | None | Foreskin | GSM22289 | 1292 | 70224 |
| 360.8041 | 63 | 506.3612 | 84 | 215.1114 | 65 | 238.8582 | 73 | DNMT3A | None | Foreskin | GSM22289 | 2188 | 70226 |
| 783.427 | 158 | 986.4079 | 202 | 432.8785 | 163 | 383.8675 | 164 | DNMT3A | None | Foreskin | GSM22289 | 7295 | 70228 |
| 589.7278 | 81 | 668.1741 | 95 | 417.8169 | 90 | 489.8475 | 104 | MYC | U2OS | Bone | GSM21312 | 2145 | 70560 |
| 1244.872 | 877 | 1295.424 | 1105 | 828.2668 | 814 | 797.8637 | 873 | MYC | U2OS | Bone | GSM21312 | 41868 | 70561 |
| 1091.95 | 340 | 1114.439 | 438 | 862.747 | 372 | 816.5909 | 371 | MYC | U2OS | Bone | GSM20506 | 13178 | 70568 |
| 1056.627 | 573 | 1076.772 | 716 | 785.2791 | 606 | 733.4569 | 607 | MYC | U2OS | Bone | GSM20506 | 31638 | 70569 |
| 1000.328 | 694 | 1018.343 | 862 | 708.5153 | 736 | 673.4738 | 778 | YY1 | GM12878 | Blood | GSM80340 | 51089 | 7060 |
| 1231.086 | 552 | 1256.406 | 699 | 950.2723 | 565 | 941.3237 | 631 | FOXK1 | HEK293 | None | GSM12503 | 16319 | 70731 |
| 263.3411 | 191 | 438.0578 | 266 | 221.4028 | 273 | 177.0436 | 275 | MYC | LNCaP | LNCaP cells | GSM23052 | 28392 | 70752 |
| 216.5146 | 170 | 373.1616 | 240 | 215.0396 | 259 | 174.653 | 262 | MYC | LNCaP | LNCaP cells | GSM23052 | 26087 | 70753 |
| 206.2157 | 168 | 370.3925 | 241 | 184.8149 | 247 | 163.4889 | 258 | MYC | LNCaP | LNCaP cells | GSM23052 | 26160 | 70754 |
| 333.9809 | 205 | 501.3649 | 274 | 210.8074 | 259 | 211.0726 | 282 | MYC | LNCaP | LNCaP cells | GSM23052 | 26545 | 70755 |
| 263.678 | 186 | 384.325 | 245 | 190.0534 | 250 | 199.5643 | 277 | MYC | LNCaP | LNCaP cells | GSM23052 | 26438 | 70756 |
| 1210.349 | 730 | 1291.421 | 977 | 894.3065 | 746 | 844.1341 | 763 | OGT | HEK293 | None | GSM12503 | 29794 | 70759 |
| 888.6122 | 380 | 864.0679 | 437 | 598.6366 | 373 | 554.8259 | 390 | HDAC2 | MM1.S | Blood | GSM23028 | 27407 | 70760 |
| 702.9446 | 160 | 986.8641 | 233 | 313.8569 | 152 | 310.6098 | 162 | HDAC3 | MM1.S | Blood | GSM23028 | 8591 | 70761 |
| 1018.076 | 200 | 1003.165 | 239 | 463.9618 | 180 | 438.4715 | 187 | HDAC4 | MM1.S | Blood | GSM23028 | 8281 | 70762 |
| 897.9694 | 509 | 927.4692 | 651 | 642.9072 | 529 | 606.779 | 551 | HDAC1 | MM1.S | Blood | GSM23028 | 37532 | 70763 |
| 380.4787 | 107 | 508.875 | 138 | 361.4999 | 150 | 418.2371 | 172 | TCF7L1 | HCT-116 | Colon | GSM20658 | 7299 | 70802 |
| 567.4258 | 159 | 904.9507 | 231 | 389.7935 | 191 | 442.2763 | 217 | TCF7L1 | HCT-116 | Colon | GSM20658 | 10915 | 70803 |
| 1045.741 | 679 | 1089.435 | 877 | 765.9932 | 724 | 732.6409 | 772 | TCF7L1 | HCT-116 | Colon | GSM20658 | 40721 | 70804 |
| 1195.913 | 319 | 1226.175 | 414 | 888.7403 | 273 | 847.082 | 274 | MYC | HCT-116 | Colon | GSM20658 | 7911 | 70805 |
| 1189.026 | 549 | 1256.619 | 746 | 880.6364 | 512 | 836.705 | 523 | MYC | HCT-116 | Colon | GSM20658 | 16821 | 70806 |
| 1193.087 | 486 | 1257.034 | 667 | 907.5492 | 472 | 861.228 | 478 | MYC | HCT-116 | Colon | GSM20658 | 14205 | 70807 |
| 1014.509 | 586 | 1014.492 | 703 | 695.3402 | 542 | 661.2392 | 565 | MYC | HCT-116 | Colon | GSM20658 | 33831 | 70808 |
| 1172.391 | 361 | 1200.089 | 469 | 910.4252 | 352 | 848.919 | 339 | MYC | HCT-116 | Colon | GSM20658 | 9950 | 70809 |

| | | | | | | | | | | | | | |
|---|---|---|---|---|---|---|---|---|---|---|---|---|---|
| 1125.123 | 355 | 1184.987 | 497 | 832.4348 | 320 | 817.3791 | 346 | MYC | HCT-116 | Colon | GSM20658 | 11284 | 70810 |
| 1160.269 | 626 | 1211.586 | 821 | 851.0604 | 602 | 816.2141 | 632 | SP1 | HCT-116 | Colon | GSM20658 | 22632 | 70811 |
| 1187.57 | 595 | 1239.419 | 782 | 884.1034 | 566 | 835.4568 | 570 | SP1 | HCT-116 | Colon | GSM20658 | 18579 | 70812 |
| 1194.288 | 580 | 1219.703 | 735 | 867.7655 | 523 | 845.3647 | 564 | SP1 | HCT-116 | Colon | GSM20658 | 17951 | 70813 |
| 878.4866 | 655 | 918.5845 | 843 | 623.272 | 739 | 602.8454 | 808 | SP1 | HCT-116 | Colon | GSM20658 | 68016 | 70814 |
| 1059.479 | 568 | 1137.275 | 785 | 792.5274 | 594 | 748.7495 | 610 | SP1 | HCT-116 | Colon | GSM20658 | 28492 | 70815 |
| 1045.647 | 575 | 1082.484 | 739 | 749.9383 | 561 | 723.7359 | 601 | SP1 | HCT-116 | Colon | GSM20658 | 30294 | 70816 |
| 790.6112 | 92 | 1121.581 | 124 | 374.3223 | 79 | 514.0597 | 100 | E2F1 | U2OS | None | GSM16939 | 1814 | 70857 |
| 1274.9 | 156 | 1299.428 | 203 | 588.1632 | 123 | 740.5875 | 148 | E2F1 | hTERT-RPE | None | GSM16939 | 2797 | 70860 |
| 807.9228 | 307 | 787.6729 | 357 | 401.4261 | 311 | 394.1761 | 334 | TRIM28 | BJAB | None | GSM22959 | 25026 | 70926 |
| 1165.132 | 495 | 1176.063 | 611 | 870.1511 | 449 | 823.5415 | 454 | SUPT5H | BJAB | None | GSM22959 | 13447 | 70927 |
| 145.3913 | 84 | 108.2401 | 83 | 166.5325 | 135 | 288.5384 | 184 | NR3C1 | A549 | Lung | GSM20952 | 11071 | 70941 |
| 161.3609 | 100 | 143.2239 | 107 | 194.2328 | 165 | 312.2763 | 217 | NR3C1 | A549 | Lung | GSM20952 | 14207 | 70942 |
| 154.1847 | 66 | 130.222 | 68 | 189.4463 | 108 | 320.6305 | 148 | NR3C1 | A549 | Lung | GSM20952 | 6874 | 70945 |
| 73.1775 | 29 | 62.89792 | 30 | 55.57688 | 38 | 230.3036 | 81 | NR3C1 | A549 | Lung | GSM20952 | 2850 | 70946 |
| 164.9611 | 114 | 150.4525 | 124 | 263.7162 | 213 | 516.7404 | 305 | NR3C1 | A549 | Lung | GSM20952 | 18220 | 70947 |
| 1221.537 | 388 | 1260.841 | 514 | 948.7113 | 362 | 906.2037 | 366 | E2F1 | Raji | Blood | GSM19763 | 8254 | 70953 |
| 1091.882 | 420 | 1123.913 | 548 | 819.4726 | 395 | 768.8083 | 393 | MYC | Raji | Blood | GSM19763 | 13959 | 70955 |
| 1230.437 | 264 | 1281.921 | 369 | 979.0557 | 254 | 959.8278 | 273 | E2F1 | Raji | Blood | GSM19762 | 5166 | 70962 |
| 1115.402 | 331 | 1155.884 | 444 | 853.0969 | 311 | 809.0521 | 312 | MYC | Raji | Blood | GSM19762 | 9491 | 70964 |
| 928.2219 | 354 | 926.0124 | 426 | 628.8397 | 339 | 652.6359 | 379 | MED12 | None | None | GSM19701 | 22404 | 70966 |
| 225.5695 | 89 | 292.9609 | 113 | 109.5967 | 97 | 131.94 | 113 | MED12 | None | None | GSM19701 | 8986 | 70967 |
| 515.1479 | 90 | 723.6546 | 120 | 593.5945 | 134 | 788.1063 | 166 | RUNX1 | MCF-7 | Breast | GSM19421 | 3287 | 70984 |
| 359.821 | 64 | 520.766 | 87 | 473.3881 | 103 | 522.0662 | 115 | RUNX1 | MCF-7 | Breast | GSM19421 | 2405 | 70985 |
| 1028.104 | 227 | 1048.977 | 293 | 662.3903 | 227 | 618.8496 | 234 | MYC | None | Bone Marr | GSM22676 | 9648 | 71013 |
| 1013.764 | 218 | 1010.256 | 269 | 632.924 | 223 | 591.3062 | 230 | MYC | None | Bone Marr | GSM22676 | 9757 | 71014 |
| 1043.628 | 238 | 1038.705 | 286 | 720.3453 | 237 | 591.0357 | 229 | MYC | None | Bone Marr | GSM22676 | 9671 | 71015 |
| 1041.632 | 231 | 1062.278 | 297 | 622.6505 | 218 | 625.719 | 233 | MYC | None | Bone Marr | GSM22676 | 9457 | 71016 |
| 223.7875 | 38 | 285.8545 | 48 | 114.983 | 36 | 260.9562 | 60 | CLOCK | U2OS | None | GSM19206 | 1291 | 71021 |
| 955.368 | 121 | 955.3783 | 131 | 541.3494 | 116 | 490.0129 | 116 | ARNTL | U2OS | None | GSM19206 | 2581 | 71023 |
| 329.9916 | 47 | 476.141 | 64 | 167.7885 | 43 | 440.6008 | 79 | ARNTL | U2OS | None | GSM19206 | 1220 | 71024 |
| 52.68472 | 33 | 75.45966 | 44 | 61.34194 | 55 | 99.80872 | 73 | TP63 | HaCaT | None | GSM14892 | 5196 | 71026 |
| 30.55556 | 43 | 75.02806 | 70 | 71.68906 | 96 | 80.95195 | 109 | TP63 | HaCaT | None | GSM14892 | 11397 | 71027 |
| 33.15944 | 44 | 92.99056 | 76 | 106.8586 | 111 | 106.5345 | 120 | TP63 | HaCaT | None | GSM14892 | 11253 | 71028 |
| 50.19243 | 53 | 99.58031 | 80 | 86.86146 | 105 | 71.45689 | 106 | TP63 | HaCaT | None | GSM14892 | 11788 | 71029 |
| 139.2343 | 99 | 186.5553 | 127 | 146.1083 | 158 | 150.299 | 173 | TP63 | MCFDCIS | Breast | GSM18496 | 16774 | 71034 |
| 722.3862 | 289 | 736.3613 | 344 | 516.8487 | 375 | 438.5532 | 380 | TP63 | MCFDCIS | Breast | GSM18496 | 31794 | 71035 |
| 1144.457 | 114 | 1330.285 | 139 | 552.3575 | 95 | 532.9446 | 98 | EHMT2 | None | Colon | GSM21839 | 1492 | 71037 |
| 1106.694 | 527 | 1131.765 | 670 | 830.1731 | 534 | 812.0723 | 580 | ZNF143 | MCF-7 | None | GSM20248 | 21381 | 71039 |
| 1119.278 | 770 | 1167.353 | 983 | 833.2065 | 848 | 810.0305 | 927 | ZNF143 | MCF-7 | None | GSM20248 | 42586 | 71040 |
| 1078.32 | 479 | 1105.08 | 608 | 825.0443 | 498 | 817.5387 | 556 | ZNF143 | MCF-7 | None | GSM20248 | 20070 | 71041 |
| 1103.184 | 787 | 1138.422 | 987 | 811.5658 | 869 | 793.1168 | 962 | ZNF143 | MCF-7 | None | GSM20248 | 47264 | 71042 |
| 0 | 3 | 0.469509 | 6 | -0.005751 | 5 | 2.44925 | 15 | SETDB1 | HeLa | Cervix | GSM23089 | 2945 | 71056 |
| 0.471851 | 6 | 3.699799 | 12 | 0.991106 | 13 | 2.488129 | 18 | ATF7IP | HeLa | Cervix | GSM23089 | 3668 | 71057 |
| 234.2374 | 53 | 332.5774 | 71 | 197.1872 | 68 | 370.8292 | 101 | ESR1 | MCF-7 | Breast | GSM21354 | 2656 | 71063 |
| 330.9916 | 83 | 411.8967 | 103 | 414.5048 | 132 | 591.3074 | 169 | ESR1 | MCF-7 | Breast | GSM21354 | 4752 | 71065 |
| 1019.838 | 504 | 1029.897 | 622 | 773.1992 | 545 | 731.1281 | 556 | GATAD2B | HeLa | Cervix | GSM21044 | 26882 | 71067 |
| 1359.876 | 1037 | 1510.474 | 1339 | 909.786 | 1037 | 846.8928 | 1064 | MBD3 | HeLa | Cervix | GSM21044 | 45690 | 71069 |
| 440.303 | 234 | 699.4 | 323 | 256.0208 | 292 | 269.5809 | 323 | CTCF | VCaP | Prostate | GSM22356 | 33681 | 71103 |
| 477.9453 | 253 | 699.5412 | 344 | 285.7603 | 319 | 325.178 | 363 | CTCF | VCaP | Prostate | GSM22356 | 36456 | 71104 |
| 435.0876 | 226 | 632.2605 | 300 | 263.781 | 285 | 272.707 | 313 | CTCF | VCaP | Prostate | GSM22356 | 31608 | 71105 |
| 504.6974 | 251 | 699.2394 | 326 | 294.7848 | 311 | 333.2165 | 353 | CTCF | VCaP | Prostate | GSM22356 | 34294 | 71106 |
| 691.0645 | 397 | 676.653 | 469 | 620.4933 | 653 | 688.265 | 870 | AR | VCaP | Prostate | GSM22356 | 58936 | 71107 |
| 679.341 | 425 | 658.7518 | 495 | 625.2928 | 723 | 687.2397 | 942 | AR | VCaP | Prostate | GSM22356 | 66905 | 71108 |
| 741.5661 | 468 | 716.9354 | 539 | 688.1284 | 794 | 736.1319 | 1002 | AR | VCaP | Prostate | GSM22356 | 63378 | 71109 |
| 754.3175 | 456 | 729.1323 | 525 | 700.9406 | 776 | 752.8979 | 988 | AR | VCaP | Prostate | GSM22356 | 58639 | 71110 |
| 1169.812 | 334 | 1207.23 | 446 | 905.1798 | 317 | 859.0759 | 315 | BRD4 | MCF-10A | None | GSM18749 | 8625 | 71136 |
| 1147.836 | 445 | 1205.659 | 606 | 910.7756 | 468 | 834.4712 | 437 | BRD4 | MCF-10A | None | GSM18749 | 13646 | 71137 |
| 1157.972 | 371 | 1217.656 | 511 | 922.5452 | 386 | 862.6461 | 372 | BRD4 | MCF-10A | None | GSM18749 | 10275 | 71138 |
| 954.4004 | 682 | 830.7204 | 660 | 819.9862 | 984 | 900.4832 | 1284 | AR | LNCaP | Prostate | GSM21864 | 59185 | 71166 |
| 1009.58 | 552 | 905.7633 | 536 | 858.2913 | 753 | 929.7539 | 996 | AR | LNCaP | Prostate | GSM21864 | 33714 | 71167 |
| 918.9621 | 637 | 807.3763 | 622 | 794.438 | 928 | 864.347 | 1202 | AR | LNCaP | Prostate | GSM21864 | 58095 | 71168 |
| 937.7503 | 606 | 843.0378 | 608 | 791.8583 | 848 | 860.6334 | 1103 | AR | LNCaP | Prostate | GSM21864 | 50045 | 71170 |
| 113.5713 | 27 | 158.4589 | 36 | 115.4096 | 39 | 189.2407 | 54 | RELA | LNCaP | Prostate | GSM22198 | 1596 | 71196 |
| 1055.992 | 224 | 1026.084 | 252 | 866.9893 | 264 | 894.045 | 325 | RELA | LNCaP | Prostate | GSM22198 | 8481 | 71202 |
| 767.6082 | 150 | 746.779 | 162 | 625.9911 | 188 | 860.8865 | 236 | RELA | LNCaP | Prostate | GSM22198 | 6687 | 71203 |
| 1089.883 | 838 | 986.2719 | 871 | 876.3775 | 1092 | 925.3326 | 1350 | PIAS1 | LNCaP | Prostate | GSM22198 | 60423 | 71204 |
| 1068.001 | 780 | 971.9981 | 814 | 856.043 | 1003 | 912.5236 | 1259 | PIAS1 | LNCaP | Prostate | GSM22198 | 54459 | 71205 |
| 1076.877 | 584 | 949.2538 | 544 | 905.4815 | 771 | 949.0427 | 965 | PIAS1 | LNCaP | Prostate | GSM22198 | 29912 | 71206 |
| 1104.703 | 850 | 1003.756 | 887 | 881.5954 | 1093 | 929.4031 | 1346 | PIAS1 | LNCaP | Prostate | GSM22198 | 59714 | 71207 |
| 1046.409 | 660 | 950.1166 | 669 | 846.3789 | 839 | 906.291 | 1074 | PIAS1 | LNCaP | Prostate | GSM22198 | 41174 | 71208 |
| 1038.653 | 621 | 933.8715 | 617 | 857.3768 | 811 | 909.7672 | 1029 | PIAS1 | LNCaP | Prostate | GSM22198 | 37756 | 71209 |

| | | | | | | | | | | | | | |
|---|---|---|---|---|---|---|---|---|---|---|---|---|---|
| 1075.773 | 490 | 984.9574 | 487 | 887.1509 | 609 | 946.3771 | 791 | PIAS1 | LNCaP | Prostate | GSM22198 | 22217 | 71210 |
| 1055.707 | 748 | 958.8443 | 773 | 848.0878 | 959 | 918.8482 | 1234 | PIAS1 | LNCaP | Prostate | GSM22198 | 51390 | 71211 |
| 817.834 | 825 | 710.6525 | 849 | 676.0315 | 1196 | 706.6515 | 1431 | FOXA1 | LNCaP | Prostate | GSM22198 | 135368 | 71212 |
| 833.9967 | 800 | 724.7468 | 822 | 698.2405 | 1170 | 726.5344 | 1400 | FOXA1 | LNCaP | Prostate | GSM22198 | 121471 | 71213 |
| 843.2155 | 837 | 734.8538 | 865 | 685.5689 | 1191 | 733.1834 | 1458 | FOXA1 | LNCaP | Prostate | GSM22198 | 131177 | 71214 |
| 863.5013 | 886 | 756.2245 | 926 | 691.6942 | 1249 | 746.6089 | 1532 | FOXA1 | LNCaP | Prostate | GSM22198 | 142632 | 71215 |
| 828.5798 | 874 | 740.9792 | 940 | 671.3763 | 1254 | 718.9569 | 1528 | FOXA1 | LNCaP | Prostate | GSM22198 | 151649 | 71216 |
| 878.9676 | 601 | 769.3016 | 588 | 709.7337 | 805 | 775.2838 | 1048 | FOXA1 | LNCaP | Prostate | GSM22198 | 59770 | 71217 |
| 825.4488 | 849 | 721.3835 | 883 | 656.5396 | 1189 | 701.188 | 1451 | FOXA1 | LNCaP | Prostate | GSM22198 | 141263 | 71218 |
| 833.3255 | 862 | 731.3266 | 906 | 672.2911 | 1228 | 702.0422 | 1468 | FOXA1 | LNCaP | Prostate | GSM22198 | 144448 | 71219 |
| 405.5973 | 50 | 403.8335 | 54 | 190.8489 | 43 | 545.554 | 84 | AR | LNCaP | Prostate | GSM22198 | 1083 | 71223 |
| 948.5002 | 306 | 896.5725 | 325 | 727.0126 | 333 | 819.1388 | 488 | AR | LNCaP | Prostate | GSM22198 | 17703 | 71224 |
| 951.3654 | 332 | 874.2465 | 336 | 735.514 | 373 | 821.3184 | 535 | AR | LNCaP | Prostate | GSM22198 | 19654 | 71225 |
| 948.715 | 276 | 910.4802 | 302 | 746.6634 | 316 | 836.6007 | 460 | AR | LNCaP | Prostate | GSM22198 | 15414 | 71226 |
| 957.4493 | 290 | 907.3539 | 309 | 746.6878 | 323 | 826.9975 | 459 | AR | LNCaP | Prostate | GSM22198 | 15995 | 71227 |
| 1148.873 | 265 | 1078.525 | 269 | 989.7954 | 344 | 961.8318 | 358 | RELA | LNCaP | Prostate | GSM22198 | 7639 | 71228 |
| 1130.611 | 303 | 1065.871 | 309 | 985.8377 | 406 | 984.5837 | 458 | RELA | LNCaP | Prostate | GSM22198 | 9427 | 71229 |
| 549.5513 | 151 | 723.0243 | 193 | 848.8084 | 305 | 879.9588 | 376 | RELA | LNCaP | Prostate | GSM22198 | 10686 | 71230 |
| 482.5977 | 126 | 554.9327 | 150 | 779.3634 | 231 | 845.5458 | 273 | RELA | LNCaP | Prostate | GSM22198 | 8330 | 71231 |
| 10.63439 | 25 | 15.49604 | 33 | 0.353115 | 22 | 10.09666 | 47 | BRD4 | L3.6 | Pancreas | GSM21885 | 8661 | 71232 |
| 148.4877 | 51 | 193.8775 | 65 | 93.73752 | 60 | 88.90932 | 63 | BRD4 | L3.6 | Pancreas | GSM21885 | 4152 | 71233 |
| 56.28234 | 29 | 40.65958 | 28 | 15.02658 | 26 | 37.06479 | 40 | STAT3 | None | None | GSM21885 | 3643 | 71234 |
| 7.271593 | 7 | 11.75281 | 10 | 9.634848 | 13 | 39.7177 | 26 | TEAD1 | None | None | GSM21885 | 1681 | 71236 |
| 153.9581 | 162 | 197.3027 | 203 | 104.3451 | 230 | 172.3709 | 295 | CEBPB | None | None | GSM21885 | 36096 | 71242 |
| 182.572 | 166 | 228.0724 | 206 | 102.2288 | 218 | 170.6273 | 281 | CEBPB | None | None | GSM21885 | 33621 | 71243 |
| 447.7224 | 58 | 533.1879 | 70 | 265.8695 | 57 | 279.8765 | 62 | BRD4 | None | None | GSM21884 | 1269 | 71252 |
| 577.0351 | 94 | 661.6697 | 111 | 418.5931 | 107 | 436.3783 | 116 | BRD4 | None | None | GSM21884 | 2963 | 71253 |
| 388.407 | 70 | 523.195 | 91 | 246.5431 | 75 | 376.8054 | 100 | BRD4 | None | None | GSM21884 | 2526 | 71254 |
| 458.2123 | 79 | 610.4168 | 102 | 336.5479 | 91 | 457.1943 | 114 | BRD4 | None | None | GSM21884 | 2668 | 71255 |
| 52.86364 | 28 | 34.78922 | 26 | 11.89606 | 24 | 28.20651 | 36 | SMAD3 | None | None | GSM19490 | 3793 | 71282 |
| 44.49641 | 25 | 37.34543 | 26 | 18.23915 | 27 | 17.59947 | 29 | SMAD3 | None | None | GSM19490 | 3564 | 71283 |
| 48.79735 | 20 | 42.97915 | 21 | 29.52965 | 24 | 57.65295 | 35 | SMAD3 | None | None | GSM19490 | 2169 | 71284 |
| 48.77928 | 20 | 24.59591 | 16 | 9.505715 | 15 | 57.74301 | 35 | SMAD3 | None | None | GSM19490 | 2157 | 71285 |
| 823.2386 | 500 | 858.7222 | 650 | 606.1403 | 571 | 594.6667 | 637 | RAD21 | HCT-116 | Colon | GSM22973 | 47204 | 71319 |
| 884.4736 | 561 | 892.0389 | 691 | 640.7732 | 618 | 628.5643 | 684 | RAD21 | HCT-116 | Colon | GSM22973 | 46661 | 71320 |
| 828.4729 | 548 | 853.3863 | 692 | 608.8437 | 630 | 596.3969 | 698 | RAD21 | HCT-116 | Colon | GSM22973 | 53328 | 71321 |
| 811.1199 | 544 | 843.4548 | 698 | 604.8923 | 649 | 570.0975 | 683 | RAD21 | HCT-116 | Colon | GSM22973 | 56146 | 71322 |
| 790.539 | 492 | 845.9675 | 666 | 596.625 | 597 | 555.3821 | 615 | CTCF | 786-O | Kidney | GSM20675 | 52251 | 71669 |
| 823.0069 | 518 | 859.3063 | 674 | 609.2442 | 601 | 592.2788 | 659 | CTCF | 786-O | Kidney | GSM20675 | 50258 | 71670 |
| 756.1928 | 461 | 818.6558 | 637 | 573.5453 | 577 | 547.7814 | 619 | CTCF | MCF-7 | Breast | GSM20675 | 55806 | 71677 |
| 744.1342 | 428 | 792.6368 | 581 | 573.6096 | 550 | 539.4955 | 575 | CTCF | MCF-7 | Breast | GSM20675 | 52479 | 71678 |
| 768.3 | 439 | 808.6976 | 584 | 580.3872 | 540 | 552.6616 | 573 | CTCF | MCF-7 | Breast | GSM20675 | 49589 | 71679 |
| 801.1482 | 526 | 862.0603 | 716 | 592.2251 | 628 | 569.9465 | 679 | CTCF | MCF-7 | Breast | GSM20675 | 58829 | 71680 |
| 708.4165 | 447 | 684.2313 | 513 | 385.0459 | 496 | 324.1667 | 512 | SOX4 | None | Fetal Lung | GSM19701 | 63951 | 71746 |
| 697.0532 | 538 | 702.8271 | 660 | 491.9395 | 656 | 394.8657 | 657 | SOX4 | None | Fetal Lung | GSM19701 | 87699 | 71747 |
| 722.1075 | 497 | 731.8874 | 613 | 514.3391 | 594 | 377.4532 | 574 | SOX4 | None | Fetal Lung | GSM19701 | 71737 | 71748 |
| 1127.524 | 473 | 1074.21 | 510 | 1006.199 | 690 | 1026.267 | 823 | NANOG | LNCaP | Prostate | GSM19338 | 18079 | 71756 |
| 25.01514 | 13 | 19.48544 | 13 | 12.57742 | 15 | 58.07701 | 32 | NANOG | LNCaP | Prostate | GSM19338 | 1819 | 71757 |
| 1158.932 | 515 | 1086.225 | 535 | 996.8476 | 684 | 1052.892 | 873 | NANOG | LNCaP | Prostate | GSM19338 | 18328 | 71758 |
| 59.49071 | 19 | 58.94469 | 21 | 23.88843 | 18 | 48.63722 | 27 | NANOG | LNCaP | Prostate | GSM19338 | 1533 | 71759 |
| 9.399496 | 6 | 20.04302 | 10 | 3.066093 | 6 | 38.57351 | 20 | NANOG | LNCaP | Prostate | GSM19338 | 1065 | 71760 |
| 144.21 | 49 | 131.4471 | 52 | 76.40356 | 54 | 59.03346 | 52 | PAX6 | None | None | GSM23334 | 4327 | 71776 |
| 152.1781 | 55 | 219.754 | 74 | 85.98721 | 63 | 89.53077 | 69 | PAX6 | None | None | GSM23334 | 5161 | 71777 |
| 795.706 | 566 | 766.5941 | 648 | 650.5376 | 799 | 665.2403 | 940 | FOXA1 | None | Breast | GSM19450 | 72488 | 71807 |
| 924.3919 | 560 | 873.5056 | 613 | 792.1435 | 801 | 817.4824 | 969 | FOXA1 | MCF-7 | Breast | GSM19450 | 46477 | 71808 |
| 352.5521 | 59 | 682.2591 | 96 | 295.7862 | 74 | 466.6785 | 101 | ERG | VCaP | Prostate | GSM18995 | 2149 | 71839 |
| 849.4993 | 278 | 849.8608 | 340 | 755.9205 | 429 | 758.314 | 493 | ESR1 | MCF-7 | None | GSM20712 | 21269 | 71855 |
| 728.0158 | 178 | 856.6043 | 214 | 814.9304 | 280 | 844.1251 | 345 | ESR1 | MCF-7 | None | GSM20712 | 10535 | 71856 |
| 616.4661 | 224 | 633.7586 | 253 | 722.8122 | 393 | 751.2165 | 485 | ESR1 | MCF-7 | None | GSM20712 | 21333 | 71857 |
| 308.5134 | 71 | 315.0963 | 79 | 495.9144 | 129 | 787.1085 | 176 | ESR1 | MCF-7 | None | GSM20712 | 3747 | 71858 |
| 915.5588 | 530 | 907.8878 | 630 | 773.2574 | 739 | 785.9336 | 865 | ESR1 | MCF-7 | Breast | GSM20400 | 42027 | 71859 |
| 700.3484 | 384 | 723.9301 | 494 | 597.0816 | 584 | 590.3347 | 658 | SMC1A | MCF-7 | Breast | GSM20400 | 54622 | 71860 |
| 662.5194 | 120 | 807.3561 | 147 | 594.0256 | 156 | 738.4476 | 186 | MED1 | MCF-7 | Breast | GSM20400 | 4607 | 71861 |
| 1065.851 | 238 | 1058.484 | 286 | 915.097 | 311 | 941.8466 | 383 | NIPBL | MCF-7 | Breast | GSM20400 | 8694 | 71862 |
| 818.22 | 564 | 815.9602 | 682 | 691.1888 | 817 | 719.0762 | 990 | ESR1 | MCF-7 | Breast | GSM20400 | 63773 | 71863 |
| 484.9428 | 279 | 561.8972 | 335 | 426.6241 | 410 | 494.8422 | 470 | SMC1A | MCF-7 | Breast | GSM20400 | 43627 | 71864 |
| 178.7607 | 35 | 96.6816 | 27 | 168.2458 | 47 | 221.8433 | 58 | MED1 | MCF-7 | Breast | GSM20400 | 1464 | 71865 |
| 155.5753 | 48 | 226.412 | 65 | 397.3699 | 113 | 523.596 | 139 | NIPBL | MCF-7 | Breast | GSM20400 | 3661 | 71866 |
| 728.7613 | 370 | 767.5803 | 494 | 579.1596 | 501 | 562.227 | 545 | SMC1A | A549 | Lung | GSM20400 | 47183 | 71867 |
| 659.988 | 118 | 723.7564 | 136 | 397.403 | 124 | 452.2109 | 141 | MED1 | A549 | Lung | GSM20400 | 4455 | 71868 |
| 440.4644 | 81 | 551.892 | 101 | 363.249 | 101 | 541.6329 | 133 | NIPBL | A549 | Lung | GSM20400 | 3204 | 71869 |
| 697.8603 | 340 | 717.8694 | 438 | 539.2541 | 464 | 542.4533 | 512 | SMC1A | A549 | Lung | GSM20400 | 46039 | 71870 |

| | | | | | | | | | | | | | |
|---|---|---|---|---|---|---|---|---|---|---|---|---|---|
| 938.3655 | 193 | 915.9279 | 209 | 550.4279 | 205 | 603.6381 | 229 | MED1 | A549 | Lung | GSM20400 | 8843 | 71871 |
| 1074.104 | 224 | 1065.739 | 264 | 779.8198 | 226 | 860.859 | 273 | NIPBL | A549 | Lung | GSM20400 | 7704 | 71872 |
| 151.023 | 90 | 192.1772 | 113 | 27.7435 | 76 | 42.33214 | 95 | SMC1A | HepG2 | None | GSM20400 | 13237 | 71873 |
| 911.4034 | 169 | 982.2997 | 193 | 625.834 | 192 | 531.2623 | 188 | MED1 | HepG2 | None | GSM20400 | 6885 | 71874 |
| 356.1534 | 66 | 424.6369 | 80 | 222.0709 | 71 | 278.3908 | 85 | NIPBL | HepG2 | None | GSM20400 | 2656 | 71875 |
| 410.3525 | 270 | 551.973 | 345 | 328.5664 | 389 | 376.898 | 444 | SMC1A | HepG2 | None | GSM20400 | 47056 | 71876 |
| 394.128 | 131 | 435.6788 | 153 | 338.1252 | 178 | 378.5373 | 201 | MED1 | HepG2 | None | GSM20400 | 10844 | 71877 |
| 448.9178 | 109 | 400.5128 | 113 | 296.0827 | 126 | 390.093 | 154 | NIPBL | HepG2 | None | GSM20400 | 6358 | 71878 |
| 772.1843 | 111 | 1177.485 | 160 | 493.5746 | 117 | 632.2686 | 142 | GTF2B | K562 | Haematop | GSM20414 | 3110 | 71922 |
| 885.5494 | 120 | 1176.887 | 156 | 554.2939 | 125 | 695.7422 | 150 | GTF2B | K562 | Haematop | GSM20414 | 3257 | 71925 |
| 1093.743 | 452 | 1088.981 | 548 | 826.9885 | 465 | 792.7122 | 481 | MYC | U2OS | Bone | GSM20493 | 19440 | 71933 |
| 1161.02 | 365 | 1178.754 | 471 | 895.29 | 364 | 873.3018 | 389 | MYC | U2OS | Bone | GSM20493 | 11358 | 71934 |
| 1086.596 | 404 | 1117.005 | 528 | 839.8013 | 429 | 803.006 | 443 | MYC | U2OS | Bone | GSM20493 | 16761 | 71935 |
| 1126.026 | 315 | 1150.969 | 412 | 876.9784 | 324 | 851.2515 | 344 | MYC | U2OS | Bone | GSM20493 | 10616 | 71936 |
| 49.34699 | 32 | 82.53223 | 46 | 24.57723 | 39 | 68.34278 | 63 | BANP | HCT-116 | None | GSM17165 | 5655 | 71950 |
| 13.85133 | 15 | 17.36625 | 19 | 0.681489 | 11 | 26.0786 | 35 | BANP | HCT-116 | None | GSM17164 | 4096 | 71951 |
| 329.843 | 116 | 319.0998 | 127 | 471.5172 | 202 | 731.2101 | 267 | ESR1 | T47D | Breast | GSM19085 | 10424 | 71952 |
| 61.50264 | 17 | 87.80089 | 23 | 25.59756 | 16 | 135.2064 | 40 | ESR1 | U2OS | Bone | GSM64142 | 1227 | 71968 |
| 168.8723 | 241 | 267.4592 | 324 | 205.225 | 422 | 327.6837 | 533 | ESR1 | U2OS | Bone | GSM64142 | 68690 | 71969 |
| 1130.895 | 517 | 1164.887 | 672 | 822.1686 | 487 | 783.6344 | 498 | MYC | Gen2.2 | None | GSM17227 | 20677 | 72049 |
| 1092.76 | 232 | 1093.135 | 285 | 838.3669 | 233 | 782.2101 | 239 | AFF1 | None | Bone Marr | GSM22274 | 7436 | 72089 |
| 898.3531 | 508 | 930.4029 | 654 | 655.905 | 547 | 632.5588 | 589 | RUNX1 | NB4 | None | GSM21797 | 38284 | 72094 |
| 1.189932 | 2 | 18.40807 | 9 | 31.34282 | 16 | 129.7323 | 36 | SMARCC1 | HCT-116 | None | GSM18359 | 1024 | 72099 |
| 16.74657 | 20 | 24.96797 | 27 | 37.86006 | 45 | 94.55296 | 71 | SMARCC1 | HCT-116 | None | GSM18359 | 5104 | 72101 |
| 13.14446 | 20 | 16.1691 | 25 | 31.38155 | 46 | 130.9263 | 89 | SMARCA4 | HCT-116 | None | GSM18359 | 5935 | 72102 |
| 86.93251 | 43 | 29.74996 | 30 | 76.08494 | 62 | 135.7492 | 86 | SMARCA4 | TTC-549 | Liver | GSM18358 | 5447 | 72109 |
| 189.5266 | 118 | 160.1465 | 124 | 190.1642 | 183 | 185.2574 | 196 | SMARCC1 | TTC-549 | Liver | GSM18358 | 18112 | 72110 |
| 159.1705 | 55 | 173.7467 | 64 | 79.15016 | 59 | 83.48229 | 65 | SMARCA4 | TTC-549 | Liver | GSM18358 | 4868 | 72113 |
| 391.7412 | 148 | 352.8722 | 157 | 214.5361 | 169 | 287.6353 | 206 | SMARCC1 | TTC-549 | Liver | GSM18358 | 14479 | 72114 |
| 222.9417 | 147 | 276.5467 | 182 | 196.1524 | 217 | 276.4938 | 268 | SMARCA4 | BT-16 | Brain | GSM18358 | 23335 | 72119 |
| 123.6569 | 71 | 168.2195 | 92 | 126.0002 | 110 | 154.1303 | 129 | SMARCC1 | BT-16 | Brain | GSM18358 | 9968 | 72120 |
| 91.93616 | 37 | 177.8916 | 58 | 60.98382 | 46 | 123.2008 | 68 | SMARCA4 | BT-16 | Brain | GSM18358 | 3803 | 72121 |
| 16.52063 | 9 | 30.95238 | 14 | 38.24172 | 21 | 102.999 | 37 | SMARCC1 | G-401 | Kidney | GSM18358 | 1407 | 72126 |
| 5.422362 | 8 | 12.19905 | 13 | 32.79264 | 28 | 87.29483 | 47 | SMARCC1 | G-401 | Kidney | GSM18358 | 2617 | 72130 |
| 135.3088 | 90 | 168.9303 | 112 | 166.1843 | 152 | 211.5232 | 181 | TP63 | JHU-029 | None | GSM23503 | 14348 | 72220 |
| 520.1415 | 80 | 945.2103 | 125 | 351.7361 | 87 | 357.7527 | 93 | MYC | None | None | GSM23048 | 2331 | 72337 |
| 447.1883 | 59 | 482.9314 | 67 | 257.0688 | 57 | 255.1412 | 60 | BRD4 | Gen2.2 | None | GSM19750 | 1315 | 72345 |
| 691.4634 | 93 | 628.3369 | 95 | 406.2594 | 91 | 263.8435 | 76 | BRD4 | Gen2.2 | None | GSM19750 | 2052 | 72347 |
| 826.6961 | 109 | 1031.273 | 135 | 566.6873 | 116 | 407.3468 | 102 | BRD4 | CAL-1 | None | GSM19750 | 2305 | 72351 |
| 982.4395 | 612 | 991.6902 | 747 | 684.9388 | 605 | 658.2284 | 647 | RUNX1 | Kasumi-1 | None | GSM20260 | 40570 | 72352 |
| 834.2259 | 458 | 823.7816 | 543 | 598.9338 | 497 | 558.7983 | 511 | HDAC2 | Kasumi-1 | None | GSM20260 | 41729 | 72353 |
| 935.9663 | 470 | 950.5091 | 588 | 680.8713 | 486 | 658.4726 | 523 | HDAC1 | Kasumi-1 | None | GSM20260 | 30767 | 72354 |
| 885.2373 | 403 | 891.6675 | 494 | 630.4956 | 408 | 607.9649 | 437 | EP300 | Kasumi-1 | None | GSM20260 | 29926 | 72355 |
| 1056.192 | 417 | 1089.441 | 543 | 780.9491 | 400 | 760.7694 | 429 | TAL1 | Kasumi-1 | None | GSM20260 | 16883 | 72356 |
| 67.49415 | 24 | 73.75782 | 28 | 34.44195 | 26 | 78.68468 | 41 | GATA2 | Kasumi-1 | None | GSM20260 | 2194 | 72357 |
| 893.9677 | 342 | 949.2322 | 477 | 687.3981 | 387 | 639.7837 | 388 | RUNX1 | Kasumi-1 | None | GSM20260 | 23178 | 72358 |
| 913.0986 | 647 | 938.2978 | 814 | 636.4466 | 684 | 617.0202 | 749 | ERG | Kasumi-1 | None | GSM20260 | 56372 | 72359 |
| 1027.467 | 634 | 1064.559 | 814 | 729.851 | 638 | 688.6745 | 657 | FLI1 | Kasumi-1 | None | GSM20260 | 37850 | 72360 |
| 115.4121 | 65 | 206.4197 | 96 | 86.57538 | 89 | 131.727 | 114 | CTCF | U2OS | Bone | GSM23416 | 9104 | 72362 |
| 883.6912 | 757 | 923.1186 | 960 | 657.7505 | 946 | 604.732 | 979 | TCF4 | Gen2.2 | None | GSM19750 | 92124 | 72366 |
| 830.2874 | 597 | 869.1963 | 774 | 628.2101 | 745 | 588.4399 | 779 | TCF4 | CAL-1 | None | GSM19750 | 69071 | 72367 |
| 1031.193 | 323 | 1018.51 | 380 | 721.8438 | 288 | 732.5462 | 322 | SIX1 | None | Embryonic | GSM19046 | 14183 | 72382 |
| 152.4429 | 39 | 228.6125 | 54 | 77.35474 | 40 | 224.3813 | 73 | SIX1 | None | Embryonic | GSM19045 | 2489 | 72383 |
| 648.9782 | 120 | 863.8643 | 155 | 389.9416 | 128 | 440.8429 | 145 | BRD9 | NOMO-1 | None | GSM20928 | 5128 | 72398 |
| 978.8641 | 337 | 968.1772 | 399 | 741.7869 | 354 | 714.6885 | 374 | ORC2 | K562 | Bone Marr | GSM17178 | 17467 | 72426 |
| 108.2588 | 24 | 176.4014 | 35 | 39.89631 | 20 | 115.7258 | 37 | GUCY1B3 | U87 | None | GSM22026 | 1154 | 72432 |
| 436.5775 | 57 | 584.3084 | 74 | 251.0501 | 55 | 305.5301 | 65 | AFF4 | SEM | None | GSM23737 | 1253 | 72467 |
| 371.8314 | 84 | 355.2418 | 90 | 122.2129 | 68 | 276.7151 | 108 | ESR1 | None | Breast | GSM24440 | 4248 | 72498 |
| 148.4 | 30 | 141.6047 | 32 | 43.24564 | 22 | 226.2969 | 56 | ESR1 | None | Breast | GSM24439 | 1328 | 72500 |
| 343.9109 | 80 | 402.9665 | 96 | 144.3425 | 73 | 285.7987 | 109 | ESR1 | None | Breast | GSM24439 | 4157 | 72502 |
| 269.2999 | 65 | 344.6296 | 82 | 102.5142 | 57 | 275.9191 | 99 | ESR1 | None | Breast | GSM24439 | 3556 | 72504 |
| 380.1608 | 80 | 446.0309 | 96 | 148.2199 | 69 | 365.3139 | 116 | ESR1 | None | Breast | GSM24439 | 3648 | 72506 |
| 240.7198 | 65 | 281.0519 | 78 | 93.44329 | 59 | 285.8658 | 108 | ESR1 | None | Breast | GSM24439 | 4114 | 72508 |
| 364.538 | 134 | 481.8317 | 171 | 202.1154 | 151 | 160.1185 | 148 | TRIM28 | None | None | GSM22304 | 11714 | 72552 |
| 542.1646 | 88 | 737.7714 | 115 | 203.2744 | 70 | 238.7724 | 81 | ZNF143 | U937 | None | GSM15877 | 2652 | 72553 |
| 1152.348 | 232 | 1166.156 | 297 | 887.5987 | 222 | 807.5624 | 224 | HEXIM1 | A375 | None | GSM16617 | 6510 | 72603 |
| 1052.578 | 574 | 1070.302 | 714 | 758.2912 | 569 | 733.8014 | 611 | HEXIM1 | A375 | None | GSM16617 | 31168 | 72604 |
| 762.1407 | 131 | 983.3474 | 166 | 448.4918 | 136 | 610.8841 | 170 | CDK9 | A375 | None | GSM16617 | 4868 | 72605 |
| 1010.552 | 373 | 1023.881 | 472 | 747.946 | 373 | 714.5116 | 389 | CDK9 | A375 | None | GSM16617 | 18802 | 72606 |
| 934.692 | 231 | 950.836 | 296 | 527.7626 | 237 | 446.3896 | 235 | HEXIM1 | A375 | None | GSM16615 | 12979 | 72608 |
| 663.8581 | 156 | 967.1597 | 223 | 541.9712 | 200 | 479.1669 | 201 | HEXIM1 | A375 | None | GSM16615 | 8914 | 72610 |
| 1294.072 | 293 | 1340.157 | 392 | 1014.489 | 265 | 1024.448 | 313 | SP1 | A375 | None | GSM16615 | 5268 | 72611 |

| | | | | | | | | | | | | | |
|---|---|---|---|---|---|---|---|---|---|---|---|---|---|
| 1272.934 | 512 | 1315.194 | 677 | 954.3991 | 461 | 933.0985 | 495 | SP1 | A375 | None | GSM16615 | 12153 | 72612 |
| 914.1241 | 187 | 984.5159 | 240 | 361.2279 | 165 | 403.4133 | 186 | ETS1 | None | Umbilical \ | GSM24427 | 8960 | 72623 |
| 219.3737 | 34 | 392.439 | 53 | 83.40393 | 27 | 54.54521 | 23 | ETS1 | None | Umbilical \ | GSM24427 | 1045 | 72624 |
| 994.2608 | 485 | 1008.725 | 609 | 726.7191 | 495 | 685.3191 | 506 | ETS1 | None | Umbilical \ | GSM24427 | 28243 | 72625 |
| 985.2819 | 401 | 1010.722 | 519 | 719.5022 | 401 | 704.792 | 437 | ETS1 | None | Umbilical \ | GSM24427 | 22022 | 72626 |
| 1078.279 | 282 | 1072.956 | 341 | 806.7755 | 265 | 731.9213 | 266 | ETS1 | None | Umbilical \ | GSM24427 | 10220 | 72627 |
| 1063.54 | 265 | 1079.432 | 337 | 783.2348 | 253 | 750.5559 | 264 | ETS1 | None | Umbilical \ | GSM24427 | 9775 | 72628 |
| 908.5209 | 147 | 1076.296 | 181 | 566.2167 | 155 | 450.3397 | 146 | BRD4 | None | Umbilical \ | GSM24427 | 4778 | 72629 |
| 635.2335 | 105 | 650.1639 | 116 | 490.2592 | 124 | 388.0776 | 116 | BRD4 | None | Umbilical \ | GSM24427 | 3408 | 72630 |
| 14.23277 | 20 | 14.0552 | 23 | 15.8463 | 35 | 74.30375 | 68 | HOXA9 | HEK293FT | Kidney | GSM15296 | 5684 | 72702 |
| 31.97548 | 22 | 26.68043 | 23 | 18.83006 | 28 | 50.34809 | 45 | ASCL1 | K562 | None | GSM19750 | 3584 | 72768 |
| 393.2202 | 208 | 619.5729 | 286 | 239.8254 | 260 | 287.9095 | 302 | CTCF | HCT-116 | Colon | GSM24392 | 27281 | 72779 |
| 441.9588 | 212 | 670.7408 | 287 | 295.4453 | 270 | 335.0318 | 307 | CTCF | K562 | Bone | GSM24392 | 25021 | 72780 |
| 440.8308 | 221 | 655.6811 | 296 | 302.2313 | 286 | 336.676 | 323 | CTCF | K562 | Bone | GSM24392 | 27238 | 72781 |
| -0.000224 | 71 | 0.045662 | 97 | 0.029189 | 165 | 0.09808 | 194 | LMNA | HeLa | Cervix | GSM21660 | 85560 | 72793 |
| 1050.789 | 626 | 1094.017 | 806 | 736.8849 | 587 | 711.0961 | 626 | TRAF7 | None | meningiom | GSM24219 | 30052 | 72854 |
| 1222.916 | 480 | 1254 | 614 | 937.4555 | 455 | 888.3806 | 456 | TRAF7 | None | meningiom | GSM24219 | 12002 | 72855 |
| 1144.25 | 405 | 1179.482 | 535 | 860.8461 | 374 | 816.7826 | 374 | TRAF7 | None | meningiom | GSM24219 | 11433 | 72856 |
| 1066.687 | 429 | 1076.69 | 527 | 783.5691 | 396 | 754.0487 | 418 | TRAF7 | None | meningiom | GSM24219 | 16035 | 72857 |
| 921.8134 | 372 | 936.6553 | 472 | 667.0279 | 376 | 659.1794 | 419 | MYF5 | Rh18 | None | GSM22431 | 24004 | 72878 |
| 48.13426 | 47 | 46.77105 | 53 | 34.14608 | 67 | 46.1325 | 81 | TBX5 | None | None | GSM22800 | 9761 | 72907 |
| 298.8811 | 49 | 286.9455 | 52 | 201.7109 | 53 | 200.357 | 56 | MED1 | MHH-MED· | None | GSM22800 | 1437 | 72908 |
| 81.5756 | 46 | 92.98536 | 55 | 37.73964 | 52 | 78.25568 | 75 | GATA4 | None | None | GSM22800 | 6338 | 72912 |
| 1103.143 | 291 | 1161.345 | 407 | 876.4538 | 302 | 816.5825 | 289 | HDAC2 | None | None | GSM22800 | 9138 | 72913 |
| 28.33767 | 35 | 29.87013 | 41 | 21.93267 | 53 | 58.07776 | 81 | TBX5 | None | None | GSM22800 | 8693 | 72917 |
| 17.54691 | 11 | 29.32706 | 16 | 18.86171 | 18 | 46.19111 | 29 | GATA4 | None | None | GSM22800 | 1860 | 72922 |
| 48.13426 | 47 | 46.77105 | 53 | 34.14608 | 67 | 46.1325 | 81 | TBX5 | None | None | GSM22800 | 9761 | 72923 |
| 298.8811 | 49 | 286.9455 | 52 | 201.7109 | 53 | 200.357 | 56 | MED1 | MHH-MED· | None | GSM22800 | 1437 | 72924 |
| 81.5756 | 46 | 92.98536 | 55 | 37.73964 | 52 | 78.25568 | 75 | GATA4 | None | None | GSM22800 | 6338 | 72928 |
| 117.1434 | 76 | 96.51952 | 79 | 95.80521 | 109 | 113.4328 | 126 | GATA4 | None | None | GSM22800 | 11738 | 72934 |
| 301.487 | 42 | 420.3172 | 56 | 237.1948 | 49 | 289.2201 | 58 | HDAC2 | None | None | GSM22800 | 1040 | 72935 |
| 23.06044 | 28 | 26.67189 | 34 | 16.28759 | 41 | 38.26421 | 60 | TBX5 | None | None | GSM22799 | 6823 | 72942 |
| 194.8731 | 65 | 148.696 | 63 | 112.9033 | 74 | 171.2279 | 96 | GATA4 | None | None | GSM22799 | 5436 | 72947 |
| 64.71691 | 51 | 59.94401 | 56 | 23.51743 | 56 | 55.31755 | 82 | TBX5 | None | None | GSM22799 | 8828 | 72948 |
| 472.5868 | 67 | 578.504 | 82 | 380.6512 | 78 | 314.3493 | 74 | MED1 | MHH-MED· | None | GSM22799 | 1456 | 72949 |
| 60.58311 | 31 | 54.10948 | 33 | 31.94197 | 36 | 76.00401 | 56 | GATA4 | None | None | GSM22799 | 3832 | 72953 |
| 97.35757 | 69 | 80.24903 | 72 | 93.06947 | 106 | 99.89404 | 118 | GATA4 | None | None | GSM22799 | 11392 | 72959 |
| 214.5946 | 48 | 300.2111 | 64 | 145.4894 | 56 | 204.4221 | 71 | AR | None | Prostate | GSM24279 | 2680 | 72970 |
| 416.8892 | 96 | 500.867 | 117 | 215.3437 | 98 | 404.8731 | 143 | AR | None | Prostate | GSM24279 | 5362 | 72971 |
| 401.8831 | 49 | 487.8969 | 60 | 261.9307 | 51 | 440.4401 | 73 | SRC | None | Prostate | GSM24279 | 1074 | 72972 |
| 405.097 | 61 | 451.9055 | 71 | 282.0287 | 68 | 531.9208 | 103 | SRC | None | Prostate | GSM24279 | 1886 | 72973 |
| 49.31225 | 14 | 129.3939 | 27 | 61.21356 | 23 | 166.0618 | 42 | AR | None | Prostate | GSM24279 | 1080 | 72974 |
| 228.6011 | 48 | 246.0262 | 55 | 117.9877 | 48 | 272.2996 | 79 | AR | None | Prostate | GSM24279 | 2385 | 72975 |
| 525.7744 | 321 | 688.8872 | 450 | 384.3841 | 437 | 363.5301 | 467 | CTCF | COLO-829 | None | GSM21782 | 48213 | 72977 |
| 506.3967 | 306 | 678.0729 | 418 | 324.4593 | 398 | 345.3216 | 443 | CTCF | COLO-829 | None | GSM21782 | 46005 | 72978 |
| 376.7919 | 240 | 420.1495 | 283 | 524.3754 | 415 | 604.413 | 521 | ESR1 | MCF-7 | Breast | GSM22578 | 35436 | 72992 |
| 614.1619 | 314 | 586.9087 | 346 | 600.2711 | 497 | 629.2421 | 608 | ESR1 | MCF-7 | Breast | GSM22578 | 39711 | 72993 |
| 134.5095 | 111 | 115.5424 | 119 | 103.3048 | 162 | 169.3528 | 210 | FOXA1 | MCF-7 | Breast | GSM22578 | 22357 | 72994 |
| 19.08059 | 19 | 13.53515 | 19 | 5.765417 | 21 | 48.77093 | 49 | FOXA1 | MCF-7 | Breast | GSM22578 | 4504 | 72995 |
| 168.6955 | 69 | 146.0499 | 72 | 121.5695 | 90 | 276.9267 | 140 | FOXA1 | MCF-7 | Breast | GSM22578 | 7560 | 72996 |
| 6.275823 | 30 | 5.9503 | 35 | 0.753598 | 38 | 12.16866 | 71 | ESR1 | MCF-7 | Breast | GSM22578 | 14717 | 72997 |
| 1.557427 | 106 | 2.622199 | 137 | 1.908212 | 219 | 10.91568 | 301 | ESR1 | MCF-7 | Breast | GSM22578 | 92286 | 72998 |
| 1.481043 | 108 | 1.837447 | 135 | 0.303722 | 197 | 3.496494 | 268 | ESR1 | MCF-7 | Breast | GSM22578 | 95383 | 72999 |
| 1.273986 | 20 | 0.229675 | 19 | 0.553646 | 35 | 7.860893 | 62 | CTCF | MCF-7 | Breast | GSM22578 | 13983 | 73000 |
| 56.44767 | 162 | 114.7602 | 233 | 57.39723 | 284 | 54.77972 | 310 | CTCF | MCF-7 | Breast | GSM22578 | 64502 | 73001 |
| 92.13908 | 149 | 105.433 | 180 | 57.48355 | 220 | 49.57283 | 233 | CTCF | MCF-7 | Breast | GSM22578 | 45547 | 73002 |
| 1114.46 | 212 | 1070.57 | 231 | 828.3422 | 215 | 759.2526 | 218 | ATF3 | HepG2 | Liver | GSM80350 | 6452 | 7301 |
| 554.5375 | 128 | 763.6274 | 168 | 449.0757 | 163 | 465.9775 | 177 | BHLHE40 | HepG2 | Liver | GSM80348 | 7093 | 7302 |
| 650.8561 | 351 | 652.3636 | 413 | 585.6638 | 560 | 612.5238 | 684 | GRHL2 | MCF-7 | Breast | GSM21718 | 52501 | 73183 |
| 647.872 | 290 | 667.0469 | 329 | 624.0726 | 447 | 618.7113 | 504 | GRHL2 | MCF-7 | Breast | GSM21718 | 33191 | 73184 |
| 658.8462 | 301 | 699.9656 | 362 | 628.0652 | 478 | 621.5517 | 538 | GRHL2 | MCF-7 | Breast | GSM21718 | 36441 | 73185 |
| 762.8065 | 401 | 682.2925 | 401 | 640.8947 | 577 | 675.0226 | 712 | FOXA1 | MCF-7 | Breast | GSM21718 | 44415 | 73186 |
| 805.116 | 415 | 747.1165 | 442 | 684.9302 | 601 | 708.2614 | 728 | FOXA1 | MCF-7 | Breast | GSM21718 | 41163 | 73187 |
| 784.5233 | 391 | 735.6187 | 420 | 690.2473 | 592 | 703.752 | 698 | FOXA1 | MCF-7 | Breast | GSM21718 | 37738 | 73188 |
| 779.8021 | 353 | 723.7572 | 371 | 675.1847 | 520 | 702.2734 | 634 | FOXA1 | MCF-7 | Breast | GSM21718 | 33540 | 73189 |
| 778.3957 | 378 | 704.2824 | 383 | 660.5312 | 541 | 696.8374 | 678 | FOXA1 | MCF-7 | Breast | GSM21718 | 37233 | 73190 |
| 413.3166 | 79 | 443.8534 | 90 | 182.6273 | 71 | 203.7147 | 80 | EZH2 | H9 | Embryo | GSM20293 | 3134 | 73191 |
| 803.8109 | 617 | 814.9585 | 760 | 613.4682 | 793 | 634.8785 | 949 | FOXA1 | HepG2 | Liver | GSM80343 | 79218 | 7322 |
| 759.6098 | 618 | 773.3012 | 766 | 578.2069 | 811 | 613.8234 | 997 | FOXA1 | HepG2 | Liver | GSM80346 | 91505 | 7323 |
| 308.6116 | 132 | 440.9921 | 175 | 215.8679 | 169 | 203.5207 | 178 | IKZF1 | None | None | GSM14201 | 14116 | 73234 |
| 80.91898 | 38 | 121.5149 | 52 | 69.78158 | 54 | 99.74901 | 68 | IKZF1 | None | None | GSM14201 | 4520 | 73235 |
| 209.5463 | 87 | 260.3397 | 108 | 152.731 | 113 | 197.8132 | 136 | IKZF1 | None | None | GSM14201 | 9065 | 73236 |

| | | | | | | | | | | | | | |
|---|---|---|---|---|---|---|---|---|---|---|---|---|---|
| 94.77946 | 39 | 144.9365 | 54 | 79.68164 | 54 | 95.66371 | 63 | IKZF1 | None | None | GSM14201 | 4084 | 73237 |
| 282.7391 | 155 | 502.4492 | 226 | 254.2799 | 226 | 204.928 | 225 | IKZF1 | None | None | GSM14201 | 20814 | 73238 |
| 80.47404 | 52 | 170.8261 | 83 | 93.55232 | 86 | 114.7984 | 101 | IKZF1 | None | None | GSM14201 | 8063 | 73239 |
| 364.7068 | 273 | 507.3455 | 353 | 190.331 | 346 | 208.2938 | 389 | LARP7 | HeLa | Cervix | GSM23323 | 51973 | 73297 |
| 13.87822 | 19 | 38.12485 | 33 | 19.44994 | 36 | 92.69633 | 72 | LARP7 | HeLa | None | GSM23323 | 5345 | 73298 |
| 8.04706 | 6 | 11.04908 | 8 | 4.903606 | 8 | 84.72196 | 32 | PARP1 | None | Kidney | GSM19389 | 1254 | 73342 |
| 18.59616 | 12 | 41.14748 | 20 | 34.73648 | 25 | 85.30597 | 41 | NOTCH1 | None | None | GSM19229 | 2030 | 73405 |
| 325.2829 | 79 | 332.2003 | 88 | 120.9966 | 69 | 233.2049 | 101 | RBPJ | None | None | GSM19229 | 4451 | 73406 |
| 965.3311 | 450 | 1009.461 | 600 | 719.9731 | 476 | 695.5209 | 511 | RBPJ | None | None | GSM19229 | 27066 | 73407 |
| 1008.755 | 375 | 1019.152 | 469 | 731.1406 | 354 | 731.093 | 401 | RBPJ | None | None | GSM19229 | 18149 | 73408 |
| 1210.341 | 743 | 1272.803 | 965 | 872.3877 | 715 | 837.24 | 759 | HEY1 | HepG2 | Liver | GSM80344 | 27614 | 7341 |
| 833.9562 | 529 | 863.3169 | 680 | 627.2371 | 643 | 634.9394 | 745 | HNF4A | HepG2 | Liver | GSM80346 | 55625 | 7343 |
| 496.1308 | 144 | 625.1746 | 180 | 301.4984 | 166 | 332.1063 | 186 | MED1 | None | None | GSM24648 | 11001 | 73444 |
| 845.0742 | 265 | 911.6132 | 383 | 507.0224 | 294 | 509.4285 | 317 | RUNX1 | THP-1 | None | GSM21080 | 20270 | 73446 |
| 876.1056 | 447 | 897.3586 | 565 | 628.879 | 474 | 614.6989 | 517 | RUNX1 | MV4-11 | None | GSM21080 | 36504 | 73449 |
| 996.2546 | 704 | 1015.638 | 875 | 714.7124 | 764 | 662.5746 | 783 | T | None | None | GSM21769 | 53718 | 73453 |
| 1003.273 | 753 | 1036.606 | 948 | 715.1364 | 828 | 679.9652 | 880 | T | None | None | GSM21769 | 61272 | 73454 |
| 297.6987 | 86 | 488.3949 | 124 | 154.6398 | 91 | 200.6419 | 110 | MLLT3 | THP-1 | None | GSM21080 | 6086 | 73462 |
| 963.8497 | 345 | 974.6363 | 429 | 764.6368 | 388 | 749.1844 | 423 | BRD4 | SUM229 | Breast | GSM23306 | 16423 | 73500 |
| 984.8135 | 316 | 963.7199 | 365 | 736.3475 | 310 | 738.8749 | 356 | BRD4 | SUM229 | Breast | GSM23306 | 13939 | 73501 |
| 990.492 | 405 | 990.6758 | 488 | 809.5897 | 491 | 793.958 | 540 | BRD4 | SUM229 | Breast | GSM23306 | 19522 | 73502 |
| 1004.347 | 514 | 1023.127 | 638 | 808.3326 | 610 | 789.3786 | 668 | BRD4 | SUM229 | Breast | GSM23306 | 25801 | 73503 |
| 895.8983 | 522 | 915.6617 | 654 | 653.7827 | 553 | 626.3146 | 590 | BRD4 | SUM159PT | Breast | GSM23306 | 35925 | 73504 |
| 886.5751 | 384 | 901.102 | 479 | 684.3685 | 432 | 652.2638 | 451 | BRD4 | SUM159PT | Breast | GSM23306 | 24336 | 73505 |
| 918.2394 | 481 | 931.1323 | 594 | 679.4496 | 508 | 642.85 | 525 | BRD4 | SUM159PT | Breast | GSM23306 | 29436 | 73506 |
| 928.3614 | 549 | 948.0046 | 684 | 686.7621 | 591 | 649.1757 | 613 | BRD4 | SUM159PT | Breast | GSM23306 | 35889 | 73507 |
| 1001.878 | 385 | 999.7242 | 463 | 755.7104 | 382 | 721.2952 | 396 | BRD4 | SUM159PT | Breast | GSM23305 | 16225 | 73508 |
| 933.5172 | 617 | 949.9406 | 764 | 674.9336 | 659 | 658.3499 | 727 | BRD4 | SUM159PT | Breast | GSM23305 | 43237 | 73509 |
| 868.1887 | 639 | 901.406 | 809 | 621.7385 | 711 | 605.4804 | 785 | BRD4 | SUM159PT | Breast | GSM23305 | 58781 | 73510 |
| 952.2061 | 587 | 970.3304 | 724 | 692.7683 | 610 | 665.4884 | 653 | BRD4 | SUM159PT | Breast | GSM23305 | 36893 | 73511 |
| 948.5667 | 590 | 969.9938 | 736 | 707.7347 | 647 | 675.9178 | 689 | BRD4 | SUM159PT | Breast | GSM23305 | 38382 | 73512 |
| 909.8235 | 584 | 919.065 | 717 | 672.7684 | 656 | 653.9598 | 714 | BRD4 | SUM159PT | Breast | GSM23305 | 44741 | 73513 |
| 960.7043 | 574 | 981.9682 | 720 | 709.7724 | 615 | 677.5137 | 650 | BRD4 | SUM159PT | Breast | GSM23305 | 35232 | 73514 |
| 862.7056 | 532 | 866.6493 | 648 | 622.2009 | 568 | 613.8422 | 636 | BRD4 | SUM159PT | Breast | GSM23305 | 40390 | 73515 |
| 846.6627 | 543 | 846.9217 | 655 | 604.2431 | 579 | 591.1125 | 640 | BRD4 | SUM159PT | Breast | GSM23305 | 44641 | 73516 |
| 936.8529 | 508 | 957.1262 | 636 | 688.4033 | 523 | 660.9003 | 555 | BRD4 | SUM159PT | Breast | GSM23305 | 29442 | 73517 |
| 943.5629 | 479 | 955.254 | 592 | 684.2287 | 476 | 663.5498 | 514 | BRD4 | SUM159PT | Breast | GSM23305 | 25938 | 73518 |
| 327.6255 | 205 | 351.0985 | 238 | 308.3738 | 307 | 418.8646 | 374 | EP300 | SUM159PT | Breast | GSM23305 | 30671 | 73519 |
| 246.7302 | 149 | 268.2705 | 174 | 263.7686 | 234 | 343.0635 | 281 | EP300 | SUM159PT | Breast | GSM23305 | 21300 | 73520 |
| 757.1037 | 545 | 773.0072 | 677 | 585.7343 | 709 | 580.1595 | 797 | EP300 | SUM159PT | Breast | GSM23305 | 68862 | 73521 |
| 738.0882 | 387 | 738.8633 | 474 | 486.0189 | 452 | 466.8399 | 484 | EP300 | SUM159PT | Breast | GSM23305 | 44773 | 73522 |
| 928.1299 | 669 | 934.9088 | 812 | 665.1355 | 725 | 664.6013 | 830 | BRD4 | SUM159PT | Breast | GSM23305 | 53438 | 73523 |
| 948.4903 | 687 | 968.7993 | 851 | 679.4904 | 749 | 674.0303 | 848 | BRD4 | SUM159PT | Breast | GSM23305 | 53686 | 73524 |
| 918.5227 | 688 | 936.5465 | 851 | 661.0978 | 770 | 649.8597 | 860 | BRD4 | SUM159PT | Breast | GSM23305 | 58926 | 73525 |
| 958.1491 | 680 | 967.0717 | 826 | 683.8895 | 728 | 669.6964 | 806 | BRD4 | SUM159PT | Breast | GSM23305 | 50686 | 73526 |
| 624.74 | 130 | 779.1963 | 161 | 490.5772 | 158 | 529.701 | 175 | BRD4 | HCC1806 | Breast | GSM23305 | 5477 | 73527 |
| 363.116 | 72 | 450.0587 | 89 | 317.8552 | 92 | 325.6564 | 99 | BRD4 | HCC1806 | Breast | GSM23305 | 2802 | 73528 |
| 981.7318 | 277 | 1003.392 | 356 | 772.2897 | 299 | 776.4769 | 343 | BRD4 | HCC1806 | Breast | GSM23305 | 11982 | 73529 |
| 564.2113 | 107 | 871.6779 | 150 | 349.775 | 113 | 376.7194 | 125 | BRD4 | HCC1806 | Breast | GSM23305 | 3804 | 73530 |
| 368.7327 | 376 | 466.5976 | 466 | 227.489 | 522 | 253.0474 | 591 | CEBPB | SUM159PT | Breast | GSM23305 | 88416 | 73533 |
| 300.2863 | 319 | 310.3103 | 369 | 163.3694 | 428 | 183.842 | 486 | CEBPB | SUM159PT | Breast | GSM23305 | 74981 | 73534 |
| 249.8124 | 282 | 225.735 | 312 | 141.2907 | 385 | 169.5557 | 445 | CEBPB | SUM159PT | Breast | GSM23305 | 68295 | 73535 |
| 345.7011 | 299 | 328.2963 | 334 | 148.7098 | 363 | 217.1252 | 445 | CEBPB | SUM159PT | Breast | GSM23305 | 60506 | 73536 |
| 237.9558 | 69 | 125.7823 | 55 | 127.0191 | 73 | 144.0179 | 83 | EP300 | SUM159PT | Breast | GSM23305 | 4535 | 73537 |
| 39.51853 | 16 | 14.71259 | 11 | 36.91157 | 23 | 28.51591 | 22 | EP300 | SUM159PT | Breast | GSM23305 | 1568 | 73538 |
| 119.9978 | 59 | 67.83821 | 51 | 92.11244 | 79 | 131.2292 | 99 | EP300 | SUM159PT | Breast | GSM23305 | 6691 | 73540 |
| 635.2671 | 101 | 477.7336 | 93 | 322.6996 | 94 | 275.9529 | 92 | MED1 | SUM159PT | Breast | GSM23305 | 2944 | 73541 |
| 302.577 | 67 | 306.9081 | 74 | 115.4203 | 57 | 127.3767 | 64 | MED1 | SUM159PT | Breast | GSM23305 | 3004 | 73542 |
| 765.051 | 164 | 773.0246 | 181 | 462.4039 | 177 | 437.4992 | 184 | MED1 | SUM159PT | Breast | GSM23305 | 7685 | 73543 |
| 1037.721 | 249 | 1013.728 | 288 | 779.9068 | 252 | 735.2504 | 261 | MED1 | SUM159PT | Breast | GSM23305 | 9066 | 73544 |
| 986.6451 | 336 | 991.311 | 413 | 749.3727 | 342 | 738.6086 | 377 | BRD4 | SUM159PT | Breast | GSM23305 | 15334 | 73545 |
| 1025.262 | 220 | 1042.566 | 280 | 726.7834 | 229 | 691.1819 | 238 | BRD4 | SUM159PT | Breast | GSM23305 | 7838 | 73546 |
| 961.7092 | 424 | 953.6279 | 507 | 718.9423 | 439 | 694.6226 | 471 | BRD4 | SUM159PT | Breast | GSM23305 | 22868 | 73547 |
| 1027.797 | 310 | 1020.823 | 368 | 764.5382 | 289 | 760.8948 | 324 | BRD4 | SUM159PT | Breast | GSM23305 | 11628 | 73548 |
| 930.9683 | 387 | 959.3242 | 502 | 695.3153 | 401 | 670.4798 | 427 | LEO1 | SEM | None | GSM22122 | 22259 | 73696 |
| 1030.26 | 262 | 1059.171 | 345 | 810.388 | 273 | 716.0978 | 267 | PAF1 | SEM | None | GSM22122 | 9533 | 73697 |
| 1129.425 | 300 | 1153.506 | 383 | 853.4813 | 264 | 812.1183 | 265 | MED1 | SEM | None | GSM22122 | 7316 | 73698 |
| 947.2552 | 544 | 970.1848 | 683 | 691.9821 | 560 | 637.5698 | 558 | MED1 | SEM | None | GSM22122 | 30841 | 73699 |
| 176.5162 | 33 | 325.7845 | 52 | 174.0619 | 45 | 120.011 | 39 | KDM2B | SEM | None | GSM22122 | 1238 | 73700 |
| 1038.085 | 616 | 1077.006 | 787 | 770.5004 | 634 | 712.9433 | 628 | KDM2B | SEM | None | GSM22122 | 29039 | 73701 |
| 853.0433 | 614 | 888.0228 | 781 | 599.943 | 669 | 566.5046 | 706 | KMT2A | None | Bone Marr | GSM23645 | 59193 | 73707 |
| 1034.778 | 589 | 1037.591 | 715 | 741.8041 | 576 | 716.6708 | 621 | KMT2A | None | Cord blood | GSM23645 | 30343 | 73708 |

| | | | | | | | | | | | | | |
|---|---|---|---|---|---|---|---|---|---|---|---|---|---|
| 295.4672 | 54 | 305.055 | 60 | 184.5923 | 57 | 413.465 | 94 | AFF1 | None | Spleen | GSM23645 | 1879 | 73709 |
| 898.6562 | 493 | 916.8873 | 616 | 639.5912 | 500 | 625.1123 | 550 | KMT2A | None | Spleen | GSM23645 | 34642 | 73710 |
| 71.80892 | 24 | 107.0636 | 33 | 26.37452 | 22 | 81.74835 | 40 | AFF1 | SEM | None | GSM23645 | 1954 | 73711 |
| 609.1611 | 337 | 681.2773 | 433 | 258.7233 | 375 | 300.0276 | 430 | AFF1 | SEM | None | GSM23645 | 46184 | 73712 |
| 1004.928 | 447 | 1020.858 | 555 | 725.0359 | 416 | 708.971 | 456 | AFF1 | SEM | None | GSM23645 | 20677 | 73713 |
| 745.982 | 373 | 760.3719 | 468 | 422.2179 | 407 | 411.2514 | 438 | AFF1 | SEM | None | GSM23645 | 39778 | 73714 |
| 899.5831 | 579 | 941.7019 | 753 | 659.889 | 636 | 637.2769 | 691 | KMT2A | SEM | None | GSM23645 | 43181 | 73715 |
| 1103.562 | 514 | 1138.343 | 661 | 819.5572 | 485 | 747.9993 | 459 | KMT2A | SEM | None | GSM23645 | 17293 | 73716 |
| 1028.726 | 386 | 1048.193 | 485 | 773.8531 | 381 | 703.5702 | 353 | BRD4 | SEM | None | GSM22122 | 15847 | 73719 |
| 857.2148 | 299 | 865.9871 | 374 | 585.5689 | 329 | 509.8722 | 334 | CREBBP | None | Tonsil | GSM23867 | 21061 | 73720 |
| 847.9649 | 289 | 853.2276 | 358 | 501.161 | 306 | 470.046 | 321 | CREBBP | None | Tonsil | GSM23867 | 20916 | 73721 |
| 26.09546 | 17 | 28.27217 | 20 | 15.88441 | 22 | 9.751195 | 20 | NR3C1 | HepG2 | Liver | GSM22654 | 2853 | 73725 |
| 52.47894 | 66 | 94.61897 | 95 | 104.8642 | 139 | 84.79899 | 140 | TAL1 | None | Blood | GSM24521 | 17048 | 73728 |
| 769.4099 | 597 | 796.5777 | 758 | 589.8791 | 788 | 531.6621 | 794 | GATA1 | None | Blood | GSM24521 | 86248 | 73729 |
| 91.2839 | 20 | 172.0529 | 32 | 76.61523 | 26 | 201.6288 | 47 | MEN1 | T47D | Breast | GSM24667 | 1106 | 73757 |
| 267.4782 | 42 | 162.4292 | 34 | 298.2286 | 61 | 353.5634 | 71 | FOXA1 | MCF-7 | Breast | GSM24666 | 1306 | 73758 |
| 697.8317 | 182 | 703.0675 | 202 | 790.656 | 287 | 788.0072 | 326 | FOXA1 | MCF-7 | Breast | GSM24666 | 12061 | 73759 |
| 1212.753 | 179 | 1208.098 | 213 | 812.8302 | 168 | 597.8676 | 150 | KMT2D | MCF-7 | Breast | GSM22646 | 3481 | 73846 |
| 1097.938 | 173 | 1063.353 | 193 | 724.9475 | 185 | 807.8925 | 208 | MEN1 | MCF-7 | Breast | GSM22646 | 5163 | 73848 |
| 995.4628 | 148 | 1112.087 | 180 | 821.3824 | 179 | 609.1269 | 161 | MEN1 | MCF-7 | Breast | GSM22646 | 4072 | 73849 |
| 929.6084 | 467 | 958.8946 | 606 | 684.2707 | 500 | 629.091 | 494 | LMO1 | Jurkat | Blood | GSM24745 | 32226 | 73854 |
| 22.83596 | 36 | 20.33611 | 40 | 5.229582 | 41 | 93.62565 | 109 | ZNF84 | 293T | None | GSM24666 | 10912 | 73925 |
| 159.7068 | 50 | 336.6084 | 83 | 116.0567 | 63 | 315.2205 | 110 | ZNF778 | 293T | None | GSM24666 | 4078 | 73928 |
| 29.81082 | 37 | 23.48576 | 39 | 10.64559 | 45 | 63.85032 | 88 | ZNF766 | 293T | None | GSM24666 | 9825 | 73929 |
| 96.09583 | 25 | 138.886 | 34 | 75.64409 | 32 | 248.8907 | 64 | ZNF708 | 293T | None | GSM24666 | 1672 | 73930 |
| 41.63199 | 23 | 56.43147 | 30 | 19.14942 | 26 | 181.9078 | 76 | ZNF707 | 293T | None | GSM24666 | 3326 | 73931 |
| 10.02239 | 33 | 31.63376 | 56 | 7.990919 | 56 | 61.58432 | 111 | ZNF680 | 293T | None | GSM24666 | 14138 | 73932 |
| 142.7928 | 109 | 323.5921 | 176 | 118.2007 | 161 | 199.1584 | 212 | ZNF680 | 293T | None | GSM24666 | 20064 | 73933 |
| 620.8211 | 105 | 861.1951 | 139 | 312.9827 | 100 | 526.8384 | 140 | ZNF610 | 293T | None | GSM24666 | 3777 | 73937 |
| 29.98242 | 13 | 31.5703 | 15 | 21.16916 | 17 | 201.8175 | 56 | ZNF558 | 293T | None | GSM24665 | 1598 | 73940 |
| 125.0772 | 29 | 169.4514 | 38 | 117.3977 | 40 | 309.6358 | 72 | ZNF549 | 293T | None | GSM24665 | 1642 | 73941 |
| 170.5509 | 47 | 362.7393 | 79 | 119.4244 | 57 | 233.6077 | 85 | ZNF547 | 293T | None | GSM24665 | 3252 | 73943 |
| 28.91093 | 12 | 78.51862 | 23 | 82.89953 | 31 | 167.848 | 48 | ZNF528 | 293T | None | GSM24665 | 1422 | 73944 |
| 27.31273 | 40 | 32.21538 | 49 | 9.666836 | 50 | 38.38422 | 82 | ZNF382 | 293T | None | GSM24665 | 11320 | 73951 |
| 4.948337 | 16 | 8.029319 | 22 | 2.150854 | 24 | 30.99578 | 56 | ZNF382 | 293T | None | GSM24665 | 7253 | 73952 |
| 60.50863 | 17 | 79.79606 | 22 | 43.38638 | 21 | 191.4726 | 49 | ZNF354A | 293T | None | GSM24665 | 1273 | 73953 |
| 11.61604 | 7 | 26.11993 | 12 | 25.01307 | 16 | 213.9007 | 52 | ZNF33A | 293T | None | GSM24665 | 1258 | 73954 |
| 36.43428 | 13 | 38.70429 | 15 | 21.01678 | 15 | 194.6675 | 50 | ZNF33A | 293T | None | GSM24665 | 1293 | 73955 |
| 13.31742 | 7 | 34.26676 | 13 | 8.343478 | 9 | 219.3439 | 50 | ZNF331 | 293T | None | GSM24665 | 1116 | 73956 |
| 90.16301 | 37 | 180.116 | 59 | 90.75656 | 56 | 247.5598 | 97 | ZNF320 | 293T | None | GSM24665 | 4041 | 73958 |
| 31.42673 | 29 | 43.51699 | 38 | 10.89476 | 33 | 62.90099 | 67 | ZNF317 | 293T | None | GSM24665 | 6382 | 73959 |
| 135.7862 | 44 | 164.1389 | 54 | 44.89414 | 39 | 222.2201 | 88 | ZNF2 | 293T | None | GSM24665 | 3659 | 73961 |
| 130.6591 | 29 | 236.0061 | 45 | 166.7203 | 47 | 255.3765 | 63 | ZNF266 | 293T | None | GSM24665 | 1550 | 73963 |
| 1009.708 | 255 | 1042.497 | 340 | 759.5476 | 266 | 743.5489 | 281 | ZNF263 | 293T | None | GSM24665 | 11261 | 73965 |
| 638.9213 | 235 | 817.1011 | 333 | 354.2947 | 272 | 506.313 | 340 | ZNF257 | 293T | None | GSM24665 | 23922 | 73966 |
| 1170.086 | 403 | 1217.637 | 544 | 889.6041 | 376 | 879.9992 | 420 | ZNF202 | 293T | None | GSM24664 | 11278 | 73970 |
| 1.560721 | 9 | 2.293039 | 12 | 3.661657 | 21 | 57.73028 | 58 | ZNF136 | 293T | None | GSM24664 | 5246 | 73974 |
| 8.017635 | 8 | 30.25216 | 17 | 14.42902 | 17 | 148.7246 | 54 | ZNF101 | 293T | None | GSM24664 | 2069 | 73976 |
| 8.440019 | 50 | 3.226171 | 48 | 4.060664 | 80 | 19.16118 | 122 | ZIM3 | 293T | None | GSM24664 | 25979 | 73977 |
| 86.05084 | 64 | 162.6415 | 96 | 82.77282 | 100 | 137.6559 | 132 | ZFP57 | 293T | None | GSM24664 | 11577 | 73979 |
| 20.33535 | 27 | 40.96572 | 41 | 17.2796 | 43 | 115.9093 | 95 | ZFP57 | 293T | None | GSM24664 | 7448 | 73980 |
| 0.557672 | 33 | 0.078725 | 34 | 0.713694 | 68 | 1.198668 | 81 | TRIM28 | H1 | Embryo | GSM20673 | 27219 | 73982 |
| 1206.854 | 444 | 1226.113 | 559 | 912.1798 | 401 | 859.114 | 399 | BCL11B | None | Thymus | GSM22474 | 10832 | 73984 |
| 1187.832 | 169 | 1243.058 | 240 | 735.0936 | 161 | 878.0093 | 188 | BCL11B | None | Thymus | GSM22474 | 3698 | 73985 |
| 1171.52 | 329 | 1207.005 | 436 | 915.1221 | 320 | 847.2755 | 298 | BCL11B | None | Thymus | GSM22474 | 8374 | 73986 |
| 1071.38 | 153 | 1160.783 | 206 | 609.2456 | 150 | 546.6629 | 150 | BCL11B | None | Thymus | GSM22474 | 3967 | 73987 |
| 185.4998 | 48 | 205.4411 | 56 | 182.3758 | 68 | 378.3062 | 106 | ESR1 | None | endometri | GSM24673 | 2948 | 73996 |
| 198.905 | 33 | 340.0183 | 50 | 164.0738 | 41 | 445.2155 | 77 | ESR1 | None | endometri | GSM24673 | 1203 | 73998 |
| 1033.013 | 307 | 1028.562 | 371 | 786.208 | 311 | 778.3878 | 343 | ESR1 | None | endometri | GSM24673 | 13088 | 73999 |
| 348.5693 | 181 | 374.5004 | 210 | 390.3403 | 288 | 485.505 | 340 | ESR1 | None | endometri | GSM24673 | 23936 | 74000 |
| 26.88478 | 12 | 37.47961 | 16 | 59.4735 | 27 | 158.9907 | 48 | ESR1 | None | endometri | GSM24672 | 1481 | 74003 |
| 84.1354 | 57 | 76.89594 | 62 | 129.54 | 106 | 166.4297 | 127 | ESR1 | None | endometri | GSM24672 | 9111 | 74007 |
| 118.6136 | 34 | 244.0145 | 56 | 150.4608 | 55 | 206.3757 | 69 | SPDEF | A549 | Lung | GSM23110 | 2242 | 74047 |
| 215.1975 | 58 | 423.8775 | 93 | 154.7034 | 70 | 172.8963 | 79 | SPDEF | A549 | Lung | GSM23110 | 3536 | 74048 |
| 311.3152 | 114 | 353.2367 | 135 | 179.5277 | 131 | 290.7252 | 174 | NKX2-1 | A549 | Lung | GSM23109 | 10181 | 74050 |
| 577.2294 | 179 | 767.8812 | 230 | 335.151 | 206 | 474.1852 | 258 | NKX2-1 | A549 | Lung | GSM23109 | 15464 | 74051 |
| 937.7735 | 930 | 947.1732 | 1122 | 664.5351 | 1163 | 654.727 | 1316 | NKX2-1 | A549 | Lung | GSM23109 | 137545 | 74052 |
| 696.2573 | 210 | 807.8883 | 251 | 781.4951 | 368 | 832.252 | 483 | ESR1 | None | None | GSM23053 | 16327 | 74115 |
| 954.4627 | 517 | 974.0058 | 654 | 750.4077 | 620 | 744.4639 | 695 | RUNX2 | MCF-7 | None | GSM23053 | 34155 | 74116 |
| 974.5368 | 533 | 1004.174 | 686 | 779.4155 | 657 | 757.7842 | 715 | RUNX2 | MCF-7 | None | GSM23053 | 34189 | 74117 |
| 100.1155 | 36 | 99.77751 | 40 | 170.9388 | 69 | 323.7316 | 102 | ESR1 | MCF7-LTED | None | GSM23053 | 3226 | 74119 |
| 126.8123 | 62 | 109.5938 | 65 | 132.8284 | 96 | 197.7353 | 123 | ESR1 | MCF-7 | None | GSM23053 | 7585 | 74120 |

| | | | | | | | | | | | | | |
|---|---|---|---|---|---|---|---|---|---|---|---|---|---|
| 364.1867 | 159 | 451.6986 | 197 | 529.3989 | 282 | 702.592 | 364 | ESR1 | MCF-7 | None | GSM23053 | 18469 | 74121 |
| 897.1169 | 540 | 830.1266 | 570 | 706.3933 | 677 | 714.2009 | 784 | FOXA1 | T47D | Breast | GSM22424 | 45300 | 74138 |
| 905.4456 | 622 | 856.8025 | 684 | 706.3009 | 777 | 718.6657 | 913 | FOXA1 | T47D | Breast | GSM22424 | 55816 | 74139 |
| 513.8258 | 124 | 664.2357 | 157 | 688.9819 | 204 | 790.7647 | 233 | ESR1 | T47D | Breast | GSM22424 | 6986 | 74140 |
| 642.1901 | 151 | 736.4824 | 179 | 719.974 | 226 | 824.5699 | 265 | ESR1 | T47D | Breast | GSM22424 | 8323 | 74141 |
| 851.3574 | 688 | 803.7394 | 764 | 671.1794 | 911 | 684.5296 | 1068 | FOXA1 | T47D | Breast | GSM22424 | 79995 | 74142 |
| 850.4178 | 702 | 796.0527 | 772 | 662.3866 | 917 | 654.6438 | 1035 | FOXA1 | T47D | Breast | GSM22424 | 83061 | 74143 |
| 650.7735 | 167 | 717.4642 | 194 | 727.033 | 251 | 813.2923 | 311 | ESR1 | T47D | Breast | GSM22424 | 10288 | 74144 |
| 641.5386 | 164 | 648.5145 | 182 | 714.1059 | 246 | 800.5699 | 296 | ESR1 | T47D | Breast | GSM22424 | 10050 | 74145 |
| 330.9353 | 125 | 319.0412 | 137 | 594.8566 | 244 | 745.4279 | 293 | GATA3 | MCF-7 | Breast | GSM21549 | 12371 | 74146 |
| 377.4656 | 161 | 422.498 | 190 | 583.1232 | 293 | 717.3131 | 379 | GATA3 | MCF-7 | Breast | GSM21549 | 18251 | 74147 |
| 125.8234 | 44 | 101.7794 | 44 | 315.446 | 103 | 537.5274 | 145 | GATA3 | MCF-7 | Breast | GSM21549 | 3980 | 74148 |
| 971.6575 | 297 | 946.6205 | 337 | 855.8037 | 431 | 867.2263 | 504 | NR3C1 | MCF-7 | Breast | GSM21549 | 15378 | 74149 |
| 997.7478 | 390 | 955.0554 | 430 | 859.6376 | 537 | 865.2553 | 618 | NR3C1 | MCF-7 | Breast | GSM21549 | 20303 | 74151 |
| 843.5277 | 413 | 818.5543 | 474 | 711.641 | 579 | 748.4589 | 721 | NR3C1 | MCF-7 | Breast | GSM21549 | 35766 | 74153 |
| 840.1548 | 354 | 821.9615 | 413 | 727.9769 | 518 | 760.4097 | 639 | NR3C1 | MCF-7 | Breast | GSM21549 | 29497 | 74155 |
| 227.9723 | 85 | 206.1376 | 90 | 365.9214 | 156 | 631.5175 | 218 | ESR1 | MCF-7 | Breast | GSM21549 | 7599 | 74157 |
| 252.2561 | 86 | 282.3383 | 101 | 385.7712 | 153 | 585.5864 | 201 | ESR1 | MCF-7 | Breast | GSM21549 | 6795 | 74158 |
| 584.7041 | 120 | 722.2488 | 148 | 826.4559 | 200 | 932.6235 | 245 | ESR1 | MCF-7 | Breast | GSM21549 | 5188 | 74159 |
| 321.1505 | 53 | 342.8972 | 60 | 501.6646 | 93 | 583.1549 | 107 | ESR1 | MCF-7 | Breast | GSM21549 | 1656 | 74160 |
| 1252.014 | 634 | 1308.591 | 833 | 919.8931 | 572 | 876.505 | 591 | BRD2 | None | Pontine | GSM22656 | 16747 | 74202 |
| 1091.992 | 138 | 1214.758 | 182 | 633.5344 | 134 | 550.2936 | 131 | BRD4 | SF8628 | Pontine | GSM22656 | 2910 | 74206 |
| 1202.831 | 322 | 1238.029 | 427 | 924.6875 | 293 | 897.1327 | 307 | BRD4 | SF8628 | Pontine | GSM22656 | 7172 | 74207 |
| 675.7241 | 90 | 786.5948 | 107 | 347.7448 | 82 | 386.3009 | 92 | BRD2 | SF8628 | Pontine | GSM22656 | 2033 | 74208 |
| 1199.75 | 456 | 1217.726 | 574 | 893.5138 | 401 | 847.9243 | 399 | BRD2 | SF8628 | Pontine | GSM22656 | 10973 | 74209 |
| 253.1515 | 42 | 449.3975 | 65 | 187.0534 | 49 | 343.8822 | 73 | MLLT1 | MV4-11 | None | GSM23657 | 1508 | 74291 |
| 831.6149 | 456 | 824.9064 | 547 | 576.8501 | 465 | 567.681 | 519 | CDK9 | MV4;11 | None | GSM24059 | 39016 | 74304 |
| 932.8689 | 472 | 945.4722 | 586 | 658.5109 | 459 | 624.4557 | 479 | CDK9 | MV4;11 | None | GSM24059 | 29396 | 74308 |
| 893.6993 | 218 | 890.411 | 252 | 367.1509 | 199 | 315.4468 | 200 | MLLT1 | MV4;11 | None | GSM21837 | 11163 | 74315 |
| 805.8178 | 104 | 1033.524 | 131 | 363.8221 | 87 | 300.427 | 83 | MLLT1 | MV4;11 | None | GSM21837 | 2088 | 74318 |
| 445.1481 | 249 | 485.2095 | 291 | 185.9941 | 277 | 205.1834 | 312 | NEUROD1 | D283 Med | None | GSM24329 | 36002 | 74341 |
| 887.7672 | 629 | 895.4428 | 769 | 594.6911 | 637 | 569.1641 | 685 | NEUROD1 | D283 Med | None | GSM24329 | 59043 | 74342 |
| 797.6601 | 594 | 788.5259 | 706 | 517.9224 | 615 | 517.8866 | 704 | NEUROD1 | D283 Med | None | GSM24329 | 72505 | 74343 |
| 388.8482 | 411 | 483.2171 | 506 | 190.6745 | 542 | 209.6177 | 612 | OTX2 | D283 Med | None | GSM24329 | 102087 | 74344 |
| 185.7679 | 379 | 178.7213 | 435 | 99.52373 | 553 | 87.17585 | 597 | OTX2 | D283 Med | None | GSM24329 | 131713 | 74345 |
| 1034.364 | 506 | 1054.964 | 642 | 731.1588 | 473 | 728.4874 | 535 | NEUROD1 | D-341 | None | GSM24329 | 26169 | 74346 |
| 1113.326 | 495 | 1139.791 | 631 | 804.1135 | 452 | 801.2255 | 512 | NEUROD1 | D-341 | None | GSM24329 | 19359 | 74347 |
| 881.4237 | 517 | 932.9906 | 692 | 627.2071 | 551 | 630.3716 | 633 | NEUROD1 | D-341 | None | GSM24329 | 44508 | 74348 |
| 811.6862 | 507 | 812.1119 | 616 | 568.3851 | 557 | 556.6507 | 619 | OTX2 | D-341 | None | GSM24329 | 55046 | 74349 |
| 831.0399 | 545 | 848.7247 | 685 | 585.9267 | 607 | 580.8165 | 684 | OTX2 | D-341 | None | GSM24329 | 58426 | 74350 |
| 699.8051 | 512 | 713.0826 | 640 | 510.18 | 642 | 489.2294 | 698 | OTX2 | D-341 | None | GSM24329 | 79425 | 74351 |
| 722.8043 | 529 | 733.0052 | 653 | 510.9832 | 631 | 500.9886 | 704 | OTX2 | D-341 | None | GSM24329 | 77484 | 74352 |
| 329.7653 | 266 | 322.7086 | 300 | 185.0877 | 347 | 235.0564 | 409 | OTX2 | D-341 | None | GSM24329 | 52073 | 74353 |
| 440.379 | 358 | 471.3452 | 418 | 241.6849 | 469 | 259.6082 | 525 | OTX2 | D-341 | None | GSM24329 | 74460 | 74354 |
| 263.7067 | 209 | 307.041 | 252 | 150.1542 | 272 | 213.1446 | 333 | OTX2 | D-341 | None | GSM24329 | 39417 | 74355 |
| 257.0914 | 273 | 256.4406 | 312 | 139.4791 | 366 | 184.339 | 435 | OTX2 | D-341 | None | GSM24329 | 64162 | 74356 |
| 937.6685 | 627 | 980.5735 | 809 | 661.2447 | 658 | 631.7469 | 701 | NEUROD1 | D-341 | None | GSM24329 | 50199 | 74357 |
| 407.6783 | 360 | 445.9454 | 424 | 226.6482 | 478 | 227.3615 | 525 | OTX2 | D-341 | None | GSM24329 | 78503 | 74358 |
| 642.2552 | 427 | 613.7015 | 487 | 433.5391 | 561 | 390.479 | 593 | OTX2 | D-341 | None | GSM24329 | 72876 | 74359 |
| 755.6141 | 462 | 765.3524 | 572 | 517.6666 | 515 | 495.5224 | 552 | NEUROD1 | D283 Med | None | GSM24329 | 56023 | 74360 |
| 145.2928 | 309 | 105.9064 | 329 | 79.40205 | 454 | 69.62987 | 489 | OTX2 | D283 Med | None | GSM24329 | 107589 | 74361 |
| 264.0417 | 352 | 334.82 | 437 | 145.5367 | 492 | 148.049 | 545 | OTX2 | D283 Med | None | GSM24329 | 98960 | 74362 |
| 457.4927 | 120 | 590.7799 | 152 | 527.5957 | 185 | 722.3112 | 231 | ESR1 | MCF-7 | Breast | GSM16440 | 7843 | 74381 |
| 21.82351 | 41 | 26.75754 | 51 | 20.39703 | 69 | 29.00101 | 84 | RELA | MCF-7 | Breast | GSM16440 | 13448 | 74382 |
| 651.0804 | 160 | 784.3711 | 195 | 829.2989 | 268 | 882.4889 | 355 | ESR1 | MCF-7 | Breast | GSM16439 | 9734 | 74383 |
| 227.941 | 68 | 331.8885 | 92 | 436.2754 | 137 | 687.9868 | 185 | ESR1 | MCF-7 | Breast | GSM16439 | 5033 | 74384 |
| 274.0088 | 81 | 436.9168 | 115 | 401.9007 | 142 | 477.244 | 165 | RELA | MCF-7 | Breast | GSM16439 | 5930 | 74385 |
| 284.76 | 86 | 342.9802 | 105 | 382.1233 | 144 | 553.6456 | 185 | RELA | MCF-7 | Breast | GSM16439 | 6377 | 74386 |
| 932.5855 | 251 | 971.2083 | 341 | 818.98 | 368 | 810.2051 | 409 | RELA | MCF-7 | Breast | GSM16439 | 14596 | 74387 |
| 88.80106 | 24 | 160.1133 | 37 | 84.822 | 34 | 128.9893 | 45 | RELA | MCF-7 | Breast | GSM16439 | 1664 | 74391 |
| 261.0154 | 138 | 271.9902 | 158 | 447.4205 | 264 | 684.2517 | 351 | RELA | MCF-7 | Breast | GSM16439 | 18286 | 74392 |
| 3.489492 | 4 | 21.10568 | 11 | 10.49998 | 11 | 111.6224 | 37 | ESR1 | MCF-7 | Breast | GSM16439 | 1272 | 74393 |
| 365.1179 | 80 | 451.1367 | 99 | 443.8726 | 125 | 688.6911 | 168 | ESR1 | MCF-7 | Breast | GSM16439 | 4052 | 74394 |
| 437.9307 | 92 | 422.3645 | 99 | 543.6354 | 145 | 655.2906 | 170 | ESR1 | MCF-7 | Breast | GSM16439 | 4448 | 74395 |
| 437.1979 | 170 | 438.5967 | 190 | 692.2111 | 310 | 737.6281 | 384 | ESR1 | MCF-7 | Breast | GSM16439 | 17045 | 74396 |
| 969.8066 | 226 | 992.6062 | 295 | 839.1573 | 314 | 862.5759 | 382 | ESR1 | MCF-7 | Breast | GSM16439 | 11433 | 74397 |
| 612.8384 | 191 | 647.8249 | 218 | 776.5715 | 346 | 798.294 | 417 | ESR1 | MCF-7 | Breast | GSM16439 | 15352 | 74398 |
| 869.2251 | 364 | 883.2731 | 457 | 756.4322 | 537 | 782.9346 | 653 | ESR1 | MCF-7 | Breast | GSM16439 | 28832 | 74399 |
| 721.9788 | 355 | 698.8757 | 412 | 659.38 | 596 | 677.618 | 710 | ESR1 | MCF-7 | Breast | GSM16439 | 44399 | 74400 |
| 1062.755 | 480 | 1069.355 | 593 | 878.4903 | 604 | 876.3874 | 683 | ESR1 | MCF-7 | Breast | GSM16439 | 22378 | 74401 |
| 17.58048 | 9 | 81.9785 | 23 | 62.36398 | 26 | 131.7507 | 41 | ESR1 | MCF-7 | Breast | GSM16439 | 1324 | 74402 |
| 974.5155 | 207 | 949.2947 | 238 | 738.0961 | 249 | 898.7977 | 389 | AR | LNCaP | Prostate | GSM20928 | 10540 | 74410 |

| | | | | | | | | | | | | | |
|---|---|---|---|---|---|---|---|---|---|---|---|---|---|
| 961.634 | 218 | 944.8538 | 255 | 793.5313 | 274 | 920.2623 | 445 | AR | LNCaP | Prostate | GSM20928 | 11533 | 74411 |
| 70.43907 | 17 | 36.51659 | 13 | 23.99158 | 14 | 104.8345 | 32 | ZSCAN5A | BeWo | None | GSM22558 | 1002 | 74427 |
| 1136.895 | 426 | 1149.805 | 529 | 849.9293 | 396 | 805.2813 | 399 | ELF1 | None | None | GSM22245 | 13906 | 74429 |
| 1177.2 | 615 | 1221.403 | 795 | 900.4471 | 646 | 859.5599 | 669 | ELF1 | None | None | GSM22245 | 22661 | 74430 |
| 894.2014 | 690 | 927.0407 | 873 | 627.4395 | 765 | 587.274 | 803 | TFAP4 | COLO-320 | None | GSM22245 | 69158 | 74431 |
| 1176.574 | 906 | 1256.662 | 1164 | 896.4268 | 1083 | 848.2145 | 1151 | TFAP4 | SNU175 | None | GSM22245 | 54721 | 74432 |
| 113.4871 | 42 | 91.51544 | 42 | 109.329 | 61 | 142.3359 | 74 | PROX1 | None | Umbilical V | GSM18310 | 3865 | 74485 |
| 349.9927 | 184 | 320.5972 | 199 | 199.3031 | 224 | 266.4676 | 271 | EP300 | None | Umbilical V | GSM18310 | 24045 | 74486 |
| 23.38329 | 18 | 7.751404 | 13 | 22.2321 | 28 | 43.81378 | 40 | EP300 | None | Umbilical V | GSM18310 | 3243 | 74487 |
| 199.4009 | 49 | 109.0077 | 39 | 91.30285 | 47 | 131.3959 | 60 | PROX1 | None | Umbilical V | GSM18310 | 2759 | 74488 |
| 1205.118 | 171 | 1211.546 | 213 | 877.0084 | 177 | 852.054 | 184 | YY1 | HeLa | Cervix | GSM17083 | 3869 | 74510 |
| 172.5875 | 107 | 341.5572 | 164 | 224.1982 | 183 | 233.8543 | 201 | CBFB | Saos-2 | None | GSM20411 | 15717 | 74545 |
| 212.4395 | 79 | 306.7263 | 106 | 123.1229 | 92 | 220.5551 | 128 | RUNX2 | Saos-2 | None | GSM20411 | 7612 | 74546 |
| 875.1782 | 523 | 898.1502 | 663 | 594.6938 | 517 | 558.0573 | 539 | MAZ | T778 | None | GSM14995 | 43687 | 74571 |
| 861.2144 | 109 | 1151.446 | 141 | 592.2909 | 116 | 427.7311 | 102 | SPIN1 | T778 | None | GSM13838 | 2221 | 74573 |
| 323.5913 | 63 | 463.3592 | 85 | 218.0035 | 71 | 237.8316 | 79 | BCL3 | K562 | Bone Marr | GSM80351 | 2690 | 7461 |
| 358.5157 | 60 | 421.1687 | 72 | 405.1873 | 88 | 359.5941 | 87 | ZMYM3 | MCF-7 | None | GSM25748 | 2018 | 74636 |
| 1148.594 | 200 | 1234.3 | 313 | 749.7129 | 189 | 916.8583 | 233 | NRF1 | MCF-7 | None | GSM25748 | 5471 | 74637 |
| 1154.451 | 380 | 1194.55 | 508 | 916.9841 | 400 | 861.6837 | 393 | NRF1 | MCF-7 | None | GSM25748 | 11668 | 74638 |
| 1276.959 | 303 | 1312.093 | 400 | 1008.933 | 286 | 976.0412 | 296 | KLF10 | MCF-7 | None | GSM25748 | 5863 | 74639 |
| 338.3802 | 53 | 370.5969 | 61 | 131.4017 | 43 | 218.2692 | 60 | ATF2 | HepG2 | Liver | GSM25748 | 1681 | 74640 |
| 158.1158 | 44 | 166.3534 | 50 | 242.3894 | 79 | 247.9029 | 85 | SMAD4 | HepG2 | Liver | GSM25748 | 2979 | 74641 |
| 258.1096 | 89 | 340.9789 | 114 | 336.1658 | 148 | 335.0137 | 158 | TSC22D4 | HepG2 | Liver | GSM25748 | 7659 | 74642 |
| 680.8563 | 167 | 717.8313 | 189 | 295.6654 | 157 | 259.8548 | 159 | NFATC3 | HepG2 | Liver | GSM25748 | 9039 | 74643 |
| 230.6104 | 51 | 245.1142 | 58 | 288.5246 | 81 | 381.9973 | 100 | HES1 | HepG2 | Liver | GSM25748 | 2576 | 74644 |
| 842.7179 | 505 | 864.8152 | 639 | 591.8178 | 542 | 553.9141 | 561 | USF2 | GM12878 | Blood | GSM25747 | 49082 | 74645 |
| 35.99016 | 17 | 39.55189 | 20 | 22.45717 | 21 | 39.34252 | 29 | TSC22D4 | GM12878 | Blood | GSM25747 | 2059 | 74646 |
| 350.1133 | 184 | 444.4404 | 230 | 194.62 | 223 | 183.7768 | 237 | ETV6 | GM12878 | Blood | GSM25747 | 24686 | 74647 |
| 213.2881 | 107 | 362.2392 | 154 | 196.6351 | 157 | 133.0839 | 145 | ETV6 | GM12878 | Blood | GSM25747 | 13475 | 74648 |
| 24.37086 | 12 | 52.27295 | 20 | 45.74582 | 25 | 53.89168 | 29 | TP73 | GM12878 | Blood | GSM25747 | 1649 | 74649 |
| 692.9826 | 195 | 877.0967 | 259 | 393.4223 | 217 | 341.6349 | 219 | DRAP1 | GM12878 | Blood | GSM25747 | 14335 | 74650 |
| 12.01721 | 13 | 2.226498 | 8 | -0.040965 | 4 | 0.018684 | 7 | BATF3 | GM12878 | Blood | GSM25747 | 3039 | 74651 |
| 115.574 | 64 | 122.8545 | 74 | 81.94836 | 85 | 86.59555 | 94 | ZMYM3 | GM12878 | Blood | GSM25747 | 8543 | 74652 |
| 857.0081 | 133 | 839.2292 | 143 | 499.8924 | 134 | 434.3496 | 132 | ZNF18 | GM12878 | Blood | GSM25747 | 3961 | 74653 |
| 98.83521 | 87 | 117.0843 | 106 | 105.2756 | 142 | 76.46086 | 138 | ZMYM3 | K562 | Bone Marr | GSM25747 | 17189 | 74654 |
| 888.4555 | 334 | 887.0286 | 407 | 682.9897 | 382 | 642.721 | 390 | USF2 | HepG2 | Liver | GSM25747 | 23853 | 74655 |
| 160.8664 | 62 | 165.0407 | 70 | 153.5638 | 90 | 151.4734 | 96 | ZMYM3 | HepG2 | Liver | GSM25747 | 5969 | 74656 |
| 195.2008 | 83 | 242.2301 | 103 | 198.1884 | 125 | 214.1466 | 139 | ARID3A | HepG2 | Liver | GSM25747 | 8868 | 74657 |
| 1092.218 | 314 | 1138.937 | 428 | 840.9157 | 312 | 828.6257 | 343 | NRF1 | HepG2 | Liver | GSM25747 | 11017 | 74658 |
| 431.1793 | 282 | 659.022 | 392 | 330.1993 | 401 | 374.8835 | 456 | ZBTB2 | HepG2 | Liver | GSM25747 | 49962 | 74659 |
| 110.0933 | 115 | 125.3655 | 138 | 105.4914 | 183 | 95.28257 | 193 | ZMYM3 | K562 | Bone Marr | GSM25747 | 25576 | 74660 |
| 540.8551 | 147 | 663.714 | 181 | 448.2636 | 193 | 354.527 | 185 | ZMYM3 | HepG2 | Liver | GSM25747 | 9999 | 74661 |
| 75.13963 | 25 | 71.14434 | 27 | 39.32576 | 27 | 90.50109 | 43 | ZMYM3 | GM12878 | Blood | GSM25747 | 2088 | 74662 |
| 1034.921 | 297 | 1054.552 | 380 | 799.5084 | 310 | 739.7156 | 296 | ZNF18 | K562 | Bone Marr | GSM25747 | 12638 | 74663 |
| 1161.005 | 544 | 1189.092 | 692 | 876.185 | 544 | 828.5701 | 550 | NRF1 | K562 | Bone Marr | GSM25747 | 19549 | 74664 |
| 1165.537 | 392 | 1202.502 | 512 | 888.8175 | 374 | 850.0171 | 384 | NRF1 | K562 | Bone Marr | GSM25747 | 12026 | 74665 |
| 1153.879 | 587 | 1187.44 | 760 | 865.9394 | 602 | 811.432 | 601 | NRF1 | K562 | Bone Marr | GSM25747 | 23469 | 74666 |
| 1143.46 | 208 | 1148.175 | 257 | 693.0541 | 186 | 744.3821 | 205 | MED1 | K562 | Bone Marr | GSM25747 | 5907 | 74667 |
| 902.8217 | 238 | 931.5882 | 314 | 555.5527 | 263 | 632.027 | 299 | BATF | GM12878 | Blood | GSM25747 | 15865 | 74668 |
| 283.9117 | 108 | 321.7466 | 128 | 242.9184 | 149 | 235.6195 | 158 | ZMYM3 | HepG2 | Liver | GSM25747 | 10371 | 74669 |
| 1083.198 | 358 | 1057.471 | 407 | 826.0095 | 361 | 789.7389 | 369 | ZNF639 | K562 | Bone Marr | GSM25747 | 14204 | 74670 |
| 980.0964 | 161 | 1072.037 | 200 | 613.0981 | 172 | 515.9979 | 167 | ZNF639 | K562 | Bone Marr | GSM25747 | 5706 | 74671 |
| 77.99645 | 23 | 124.5207 | 33 | 99.48675 | 38 | 97.04326 | 40 | ZMYM3 | HepG2 | Liver | GSM25747 | 1738 | 74672 |
| 1157.08 | 214 | 1180.418 | 277 | 929.0273 | 223 | 822.8926 | 215 | RUVBL1 | HepG2 | Liver | GSM25747 | 5716 | 74673 |
| 303.3564 | 67 | 374.9132 | 83 | 257.9826 | 87 | 305.4715 | 101 | SMAD4 | HepG2 | Liver | GSM25747 | 3477 | 74674 |
| 634.2264 | 133 | 698.0947 | 154 | 502.2612 | 165 | 476.1075 | 171 | SMAD4 | HepG2 | Liver | GSM25747 | 6353 | 74675 |
| 303.7824 | 42 | 245.5732 | 40 | 145.4373 | 37 | 144.155 | 39 | ZMYM3 | HepG2 | Liver | GSM25747 | 1057 | 74676 |
| 128.1907 | 33 | 89.85011 | 30 | 55.49537 | 31 | 51.08334 | 32 | ZMYM3 | HepG2 | Liver | GSM25747 | 1980 | 74677 |
| 707.0065 | 110 | 679.337 | 117 | 358.2805 | 103 | 400.4432 | 116 | STAT2 | GM12878 | Blood | GSM25747 | 3364 | 74678 |
| 560.836 | 123 | 589.8565 | 139 | 424.6022 | 150 | 343.7445 | 144 | FOXO3 | K562 | Bone Marr | GSM25747 | 6241 | 74680 |
| 1148.855 | 742 | 1215.386 | 977 | 832.2038 | 765 | 796.8055 | 809 | SMAD4 | HepG2 | Liver | GSM25747 | 37237 | 74681 |
| 1107.679 | 688 | 1135.207 | 869 | 805.2976 | 706 | 781.9568 | 769 | FOXO3 | HepG2 | Liver | GSM25747 | 35213 | 74682 |
| 693.7561 | 402 | 713.9158 | 513 | 483.0759 | 510 | 462.9983 | 547 | ZBTB2 | GM12878 | Blood | GSM25747 | 60946 | 74683 |
| 905.499 | 121 | 1151.841 | 152 | 418.2492 | 105 | 437.5955 | 114 | EZH2 | K562 | Bone Marr | GSM25747 | 2935 | 74684 |
| 123.8445 | 62 | 192.5885 | 86 | 169.8095 | 109 | 237.2704 | 136 | NFIL3 | HepG2 | Liver | GSM25747 | 8076 | 74685 |
| 1189.672 | 426 | 1192.822 | 526 | 920.7832 | 426 | 894.9531 | 451 | KLF10 | HepG2 | Liver | GSM25747 | 12373 | 74686 |
| 101.7125 | 23 | 86.54259 | 23 | 11.75526 | 11 | 22.52913 | 16 | GRHL3 | NHEK | Skin | GSM24985 | 1041 | 74687 |
| 974.482 | 399 | 1006.216 | 527 | 725.9476 | 415 | 690.6592 | 429 | GRHL3 | NHEK | Skin | GSM24985 | 22998 | 74688 |
| 166.9192 | 29 | 244.8365 | 40 | 122.4627 | 34 | 285.0354 | 58 | KMT2D | T47D | Breast | GSM25080 | 1147 | 74823 |
| 519.911 | 170 | 504.2655 | 186 | 650.8375 | 275 | 722.9955 | 314 | T | T47D | None | GSM24768 | 14036 | 74830 |
| 459.458 | 167 | 439.0684 | 182 | 533.2042 | 263 | 683.3413 | 316 | T | T47D | None | GSM24768 | 15548 | 74831 |
| 517.4277 | 153 | 485.0796 | 164 | 719.5807 | 259 | 777.4094 | 303 | T | T47D | None | GSM24768 | 11182 | 74832 |

| | | | | | | | | | | | | | |
|---|---|---|---|---|---|---|---|---|---|---|---|---|---|
| 1060.927 | 262 | 1094.295 | 348 | 762.7019 | 248 | 802.2207 | 283 | JMJD6 | None | None | GSM23609 | 9591 | 74836 |
| 1212.314 | 579 | 1277.835 | 787 | 908.7373 | 548 | 880.5944 | 586 | JMJD6 | None | None | GSM23609 | 16159 | 74838 |
| 267.0628 | 50 | 366.9805 | 66 | 202.6724 | 60 | 477.1725 | 102 | AR | LNCaP | Prostate | GSM21228 | 2098 | 74839 |
| 280.4898 | 136 | 273.8897 | 151 | 166.6706 | 167 | 235.2991 | 207 | FOS | K562 | Bone Marr | GSM77764 | 17012 | 7484 |
| 333.1887 | 57 | 396.1038 | 69 | 222.2382 | 63 | 571.8474 | 113 | AR | LNCaP | Prostate | GSM21228 | 2094 | 74840 |
| 387.5701 | 102 | 464.4021 | 124 | 640.0936 | 188 | 743.0151 | 216 | GRHL2 | LNCaP | Prostate | GSM21228 | 6338 | 74841 |
| 485.042 | 120 | 524.7427 | 138 | 668.7501 | 201 | 751.1264 | 227 | GRHL2 | LNCaP | Prostate | GSM21228 | 7067 | 74842 |
| 927.1423 | 324 | 890.577 | 360 | 653.5249 | 325 | 615.59 | 340 | JMJD6 | None | None | GSM19220 | 19787 | 74843 |
| 556.9607 | 135 | 750.8426 | 175 | 365.1124 | 156 | 317.3215 | 156 | JMJD6 | None | None | GSM19220 | 7992 | 74844 |
| 76.50248 | 78 | 121.9265 | 107 | 136.6355 | 154 | 136.6285 | 167 | GATA2 | K562 | Bone Marr | GSM77764 | 16049 | 7485 |
| 178.7882 | 33 | 259.0109 | 45 | 49.01077 | 23 | 54.73052 | 26 | HDAC8 | K562 | Bone Marr | GSM77764 | 1298 | 7486 |
| 376.8321 | 183 | 429.9265 | 218 | 278.0686 | 245 | 328.6156 | 283 | JUNB | K562 | Bone Marr | GSM77763 | 23098 | 7487 |
| 795.2686 | 525 | 803.1406 | 647 | 614.9916 | 669 | 592.9834 | 727 | JUND | K562 | Bone Marr | GSM77763 | 59302 | 7488 |
| 200.0808 | 34 | 275.4662 | 45 | 83.9057 | 29 | 74.39847 | 29 | NR4A1 | K562 | Bone Marr | GSM77763 | 1196 | 7489 |
| 790.9785 | 381 | 819.2299 | 494 | 597.5458 | 450 | 587.6019 | 500 | CTCF | THP-1 | None | GSM25442 | 35853 | 74902 |
| 813.6665 | 379 | 847.0125 | 500 | 619.9715 | 446 | 608.0053 | 491 | CTCF | THP-1 | None | GSM25442 | 32756 | 74903 |
| 797.2201 | 433 | 842.4636 | 578 | 595.0428 | 505 | 579.5453 | 554 | CTCF | THP-1 | None | GSM25442 | 41941 | 74904 |
| 755.9978 | 350 | 806.8565 | 482 | 579.3677 | 435 | 573.0727 | 485 | CTCF | THP-1 | None | GSM25442 | 36610 | 74905 |
| 892.5402 | 492 | 930.5446 | 643 | 657.352 | 533 | 626.447 | 560 | POLR3D | THP-1 | None | GSM25442 | 35951 | 74906 |
| 785.3711 | 610 | 832.1044 | 798 | 571.8644 | 735 | 566.1795 | 831 | POLR3D | THP-1 | None | GSM25442 | 75616 | 74907 |
| 1042.482 | 264 | 1044.126 | 325 | 726.101 | 250 | 766.5181 | 274 | POLR3D | THP-1 | None | GSM25442 | 10090 | 74908 |
| 946.1638 | 602 | 982.9717 | 775 | 690.9476 | 652 | 659.4176 | 695 | POLR3D | THP-1 | None | GSM25442 | 43541 | 74909 |
| 53.95731 | 24 | 106.283 | 38 | 77.04234 | 43 | 66.22439 | 43 | CUX1 | K562 | None | GSM24392 | 2809 | 74934 |
| 92.95473 | 27 | 111.2499 | 33 | 53.54048 | 30 | 102.0159 | 44 | CUX1 | K562 | None | GSM24392 | 2007 | 74937 |
| 124.4896 | 38 | 158.6304 | 48 | 130.8917 | 57 | 144.9431 | 64 | CUX1 | K562 | None | GSM24392 | 2982 | 74939 |
| 355.318 | 54 | 495.7194 | 72 | 226.4003 | 57 | 197.4476 | 56 | NOTCH1 | MO1043 | None | GSM24355 | 1621 | 75003 |
| 8.24393 | 32 | 17.31446 | 47 | 1.811383 | 43 | 22.30034 | 83 | SMARCE1 | None | None | GSM25462 | 14769 | 75037 |
| 859.9057 | 595 | 875.3375 | 738 | 608.4591 | 653 | 603.5197 | 741 | TP53 | MDA-MB-2 | Breast | GSM25015 | 58267 | 75042 |
| 1012.391 | 681 | 1054.61 | 867 | 704.1511 | 672 | 700.1612 | 765 | E2F1 | MDA-MB-2 | Breast | GSM25015 | 43103 | 75043 |
| 1078.177 | 189 | 1080.828 | 233 | 671.4959 | 190 | 639.966 | 197 | MYC | MDA-MB-2 | Breast | GSM25015 | 5934 | 75044 |
| 37.3893 | 57 | 24.46097 | 57 | 29.37293 | 89 | 86.61727 | 138 | FOSL1 | MDA-MB-2 | Breast | GSM25015 | 16408 | 75045 |
| 771.4557 | 396 | 811.0977 | 527 | 585.999 | 481 | 482.9993 | 467 | MYB | DU-528 | Blood | GSM24666 | 40764 | 75136 |
| 878.9587 | 562 | 922.406 | 732 | 645.6528 | 633 | 599.209 | 646 | MYB | PF382 | Blood | GSM24666 | 49360 | 75137 |
| 799.3902 | 457 | 824.2394 | 584 | 580.1339 | 515 | 530.9583 | 516 | MYB | LOUCY | Blood | GSM24666 | 44808 | 75138 |
| 1095.434 | 304 | 1112.135 | 388 | 845.79 | 302 | 794.188 | 294 | RELA | Huh7 | Liver | GSM23609 | 10114 | 75139 |
| 1162.334 | 193 | 1191.307 | 255 | 956.0784 | 209 | 780.7803 | 195 | RELA | Huh7 | Liver | GSM23609 | 4780 | 75140 |
| 1231.802 | 681 | 1285.88 | 893 | 879.6567 | 620 | 850.7615 | 662 | HEY1 | K562 | Bone Marr | GSM80338 | 22145 | 7527 |
| 919.4504 | 433 | 933.2215 | 539 | 641.9001 | 408 | 607.844 | 426 | ZBTB48 | U2OS | None | GSM25431 | 27283 | 75273 |
| 793.7001 | 377 | 801.9811 | 469 | 398.8895 | 370 | 389.3906 | 398 | ZBTB48 | U2OS | None | GSM25431 | 34718 | 75274 |
| 885.9871 | 478 | 901.4543 | 597 | 617.0744 | 475 | 569.7677 | 478 | ZBTB48 | U2OS | None | GSM25431 | 36119 | 75275 |
| 822.2262 | 380 | 831.6686 | 472 | 430.1707 | 362 | 415.2294 | 387 | ZBTB48 | U2OS | None | GSM25431 | 32153 | 75276 |
| 38.03809 | 134 | 11.70969 | 119 | 5.505741 | 162 | 4.059985 | 174 | ZBTB48 | U2OS | None | GSM25431 | 52115 | 75277 |
| 36.7459 | 128 | 32.6577 | 145 | 14.6938 | 183 | 9.940291 | 190 | ZBTB48 | U2OS | None | GSM25431 | 49463 | 75278 |
| 68.20851 | 117 | 81.45492 | 143 | 14.28146 | 131 | 41.60729 | 186 | ZBTB48 | U2OS | None | GSM25431 | 32880 | 75279 |
| 12.59999 | 85 | 14.18504 | 103 | 0.927259 | 106 | 4.530085 | 144 | ZBTB48 | U2OS | None | GSM25431 | 41306 | 75280 |
| 857.5469 | 435 | 861.3316 | 527 | 617.5978 | 458 | 596.794 | 495 | ZBTB48 | HeLa | None | GSM25431 | 34516 | 75281 |
| 856.5149 | 397 | 857.3888 | 478 | 638.4519 | 435 | 606.23 | 455 | ZBTB48 | HeLa | None | GSM25431 | 30217 | 75282 |
| 836.3623 | 388 | 838.6878 | 472 | 530.181 | 389 | 553.089 | 428 | ZBTB48 | HeLa | None | GSM25431 | 31445 | 75283 |
| 887.7068 | 424 | 894.4794 | 522 | 651.8352 | 450 | 631.3209 | 486 | ZBTB48 | HeLa | None | GSM25431 | 30051 | 75284 |
| 731.1333 | 322 | 741.8807 | 406 | 352.1307 | 355 | 288.8259 | 359 | ZBTB48 | HeLa | None | GSM25431 | 35926 | 75285 |
| 254.9907 | 192 | 265.6334 | 221 | 105.1265 | 221 | 126.2398 | 256 | ZBTB48 | HeLa | None | GSM25431 | 32819 | 75286 |
| 931.139 | 276 | 955.796 | 363 | 661.142 | 293 | 633.0943 | 308 | ZBTB48 | HeLa | None | GSM25431 | 15083 | 75287 |
| 584.1727 | 247 | 645.9008 | 289 | 255.8934 | 263 | 255.1997 | 285 | ZBTB48 | HeLa | None | GSM25431 | 26302 | 75288 |
| 145.4977 | 53 | 135.6631 | 57 | 250.2174 | 102 | 390.5254 | 136 | CDX2 | LS174T | None | GSM25604 | 4927 | 75366 |
| 199.6772 | 35 | 281.3614 | 47 | 167.8158 | 44 | 218.8113 | 54 | STAT3 | None | None | GSM16413 | 1332 | 75512 |
| 603.1947 | 209 | 827.7644 | 293 | 328.7789 | 236 | 359.2427 | 264 | CTCF | None | Retina | GSM23163 | 19105 | 75553 |
| 767.7005 | 262 | 817.897 | 342 | 370.1433 | 282 | 399.2612 | 314 | CTCF | None | Retina | GSM23163 | 24128 | 75554 |
| 781.3753 | 298 | 829.1097 | 413 | 516.3813 | 355 | 469.1545 | 369 | CTCF | None | Retina | GSM23163 | 27267 | 75555 |
| 670.6923 | 233 | 839.5829 | 332 | 394.9582 | 271 | 381.0928 | 288 | CTCF | None | Retina | GSM23163 | 20861 | 75556 |
| 772.8296 | 340 | 804.1827 | 448 | 537.1263 | 399 | 496.8969 | 419 | CTCF | None | Retina | GSM23163 | 32380 | 75557 |
| 685.8833 | 277 | 790.0456 | 385 | 429.9857 | 336 | 387.1024 | 349 | CTCF | None | Retina | GSM23163 | 27745 | 75558 |
| 1121.944 | 547 | 1155.844 | 709 | 825.8546 | 523 | 778.5026 | 527 | BRD4 | None | Retina | GSM23163 | 19714 | 75559 |
| 1147.772 | 351 | 1169.253 | 444 | 880.6192 | 326 | 814.4826 | 306 | BRD4 | None | Retina | GSM23163 | 8621 | 75560 |
| 1144.292 | 455 | 1152.417 | 555 | 858.2693 | 414 | 791.949 | 397 | BRD4 | None | Retina | GSM23163 | 12692 | 75561 |
| 1122.094 | 520 | 1155.533 | 672 | 827.934 | 485 | 779.9649 | 486 | BRD4 | None | Retina | GSM23162 | 16894 | 75562 |
| 924.8623 | 129 | 1167.047 | 170 | 487.5426 | 120 | 522.0618 | 132 | BRD4 | None | Retina | GSM23162 | 3067 | 75563 |
| 396.5324 | 177 | 483.4727 | 217 | 173.0755 | 189 | 216.2577 | 223 | SMC1A | HAP-1 | Embryo | GSM24938 | 19565 | 75568 |
| 262.7643 | 95 | 419.5957 | 134 | 132.3243 | 103 | 121.0616 | 107 | SMC1A | HAP-1 | Embryo | GSM24938 | 8510 | 75569 |
| 785.7432 | 269 | 806.5092 | 341 | 383.0828 | 289 | 400.8921 | 318 | SMC1A | HAP-1 | Embryo | GSM24938 | 23740 | 75570 |
| 384.6464 | 143 | 538.518 | 188 | 209.8579 | 162 | 207.787 | 174 | SMC1A | HAP-1 | Embryo | GSM24938 | 13464 | 75571 |
| 434.662 | 268 | 664.9467 | 365 | 269.7818 | 347 | 272.1608 | 379 | CTCF | HAP-1 | Embryo | GSM24938 | 41870 | 75572 |
| 434.4026 | 241 | 696.1637 | 336 | 281.2958 | 312 | 281.5131 | 339 | CTCF | HAP-1 | Embryo | GSM24938 | 34233 | 75573 |

| | | | | | | | | | | | | | |
|---|---|---|---|---|---|---|---|---|---|---|---|---|---|
| 456.585 | 270 | 679.7143 | 370 | 277.8793 | 346 | 266.086 | 371 | CTCF | HAP-1 | Embryo | GSM24938 | 41315 | 75574 |
| 445.1365 | 259 | 688.6997 | 363 | 312.1783 | 348 | 302.4389 | 374 | CTCF | HAP-1 | Embryo | GSM24938 | 38824 | 75575 |
| 877.899 | 622 | 915.8782 | 799 | 663.3874 | 753 | 638.7605 | 818 | RAD21 | Rh4 | None | GSM22141 | 61297 | 75836 |
| 512.5073 | 236 | 673.4682 | 299 | 351.3391 | 303 | 345.6561 | 326 | EP300 | Rh4 | None | GSM22141 | 27464 | 75837 |
| 9.655639 | 21 | 68.59706 | 52 | 37.57822 | 57 | 81.07869 | 83 | MYOG | Rh4 | None | GSM22141 | 7335 | 75838 |
| 306.8736 | 140 | 355.6674 | 168 | 313.9911 | 213 | 315.9061 | 230 | MYOD1 | Rh4 | None | GSM22141 | 16454 | 75839 |
| 987.1943 | 219 | 998.8988 | 277 | 518.0057 | 209 | 477.7281 | 215 | MYCN | Rh4 | None | GSM22141 | 9775 | 75840 |
| 176.7082 | 44 | 251.2008 | 59 | 127.6303 | 53 | 163.222 | 64 | MED1 | Rh4 | None | GSM22141 | 2554 | 75841 |
| 884.8062 | 623 | 904.6566 | 781 | 647.9132 | 725 | 616.1018 | 772 | CTCF | Rh4 | None | GSM22141 | 61424 | 75847 |
| 813.1941 | 544 | 867.2462 | 726 | 624.2228 | 686 | 599.8151 | 737 | CTCF | Rh4 | None | GSM22140 | 60553 | 75848 |
| 81.8697 | 39 | 91.0491 | 46 | 40.99873 | 44 | 43.98646 | 49 | BRD4 | Rh4 | None | GSM22140 | 4571 | 75849 |
| 893.4216 | 273 | 915.7859 | 352 | 623.2248 | 300 | 588.7819 | 314 | BRD4 | Rh4 | None | GSM22140 | 16863 | 75850 |
| 13.717 | 20 | 9.147957 | 20 | 7.420569 | 28 | 46.09747 | 57 | FOXO1 | 7250 | None | GSM22140 | 5865 | 75851 |
| 8.829226 | 6 | 2.754579 | 4 | 1.630678 | 5 | 7.601698 | 10 | FOXO1 | 7250 | None | GSM22140 | 1208 | 75852 |
| 300.1491 | 99 | 398.9518 | 127 | 189.6937 | 116 | 180.6292 | 122 | BRD4 | 7250 | None | GSM22140 | 7685 | 75854 |
| 1219.593 | 406 | 1245.387 | 517 | 926.3265 | 365 | 891.5126 | 374 | E2F1 | MM1.S | None | GSM21325 | 9053 | 75924 |
| 1314.035 | 600 | 1359.185 | 789 | 962.6887 | 532 | 946.3475 | 574 | KDM1A | PC-3M-luc | Prostate | GSM13741 | 14124 | 75942 |
| 815.0656 | 249 | 786.2415 | 273 | 724.8063 | 372 | 749.4265 | 451 | ESR1 | MCF-7 | Mammary | GSM24672 | 19262 | 76100 |
| 500.3359 | 103 | 409.4083 | 101 | 491.4855 | 142 | 685.7762 | 180 | ESR1 | MCF-7 | Mammary | GSM24672 | 4611 | 76101 |
| 754.3223 | 282 | 728.8077 | 314 | 689.4624 | 460 | 721.6786 | 571 | ESR1 | MCF-7 | Mammary | GSM24672 | 29233 | 76102 |
| 574.5726 | 198 | 509.1539 | 208 | 721.576 | 336 | 738.5141 | 402 | ESR1 | MCF-7 | Mammary | GSM24672 | 17989 | 76103 |
| 702.0572 | 315 | 640.9576 | 340 | 654.3715 | 531 | 688.1561 | 660 | ESR1 | MCF-7 | Mammary | GSM24672 | 39363 | 76104 |
| 700.4506 | 355 | 655.5304 | 388 | 639.0216 | 597 | 667.5695 | 731 | ESR1 | MCF-7 | Mammary | GSM24672 | 48152 | 76105 |
| 569.996 | 306 | 610.4604 | 355 | 628.884 | 552 | 657.0513 | 674 | ESR1 | MCF-7 | Mammary | GSM24672 | 44828 | 76106 |
| 670.3219 | 323 | 666.4293 | 367 | 631.3338 | 540 | 669.2784 | 677 | ESR1 | MCF-7 | Mammary | GSM24672 | 43484 | 76107 |
| 696.1627 | 364 | 661.898 | 409 | 628.1782 | 604 | 666.5777 | 752 | ESR1 | MCF-7 | Mammary | GSM24672 | 50718 | 76108 |
| 6.748681 | 8 | 3.342477 | 7 | 1.742645 | 9 | 23.27392 | 24 | H2AFX | HeLa | Cervix | GSM14816 | 2240 | 76140 |
| 1348.998 | 861 | 1401.454 | 1087 | 1011.703 | 917 | 1000.215 | 1022 | MYC | LNCaP | Prostate | GSM19072 | 27634 | 76149 |
| 1280.318 | 625 | 1328.44 | 820 | 999.3078 | 659 | 987.6997 | 729 | MYC | LNCaP | Prostate | GSM19072 | 17157 | 76151 |
| 1206.268 | 244 | 1214.438 | 302 | 968.9968 | 250 | 970.0578 | 284 | MYC | LNCaP | Prostate | GSM19072 | 5722 | 76152 |
| 883.9551 | 360 | 836.5693 | 391 | 677.2539 | 420 | 776.8552 | 620 | AR | LNCaP | Prostate | GSM19072 | 28487 | 76154 |
| 834.8804 | 548 | 756.9339 | 564 | 641.5889 | 692 | 734.23 | 961 | AR | LNCaP | Prostate | GSM19072 | 61248 | 76155 |
| 883.9751 | 283 | 840.2577 | 309 | 662.3773 | 326 | 788.7032 | 497 | AR | LNCaP | Prostate | GSM19071 | 20582 | 76156 |
| 275.1778 | 42 | 363.0071 | 54 | 130.3932 | 37 | 73.45156 | 29 | MYC | HCT-116 | Colon | GSM25767 | 1123 | 76186 |
| 580.6894 | 108 | 862.1137 | 148 | 429.8687 | 125 | 370.5274 | 123 | AR | None | None | GSM24122 | 3769 | 76204 |
| 463.4363 | 79 | 731.9941 | 113 | 273.7678 | 81 | 279.6337 | 87 | AR | None | Prostate | GSM24122 | 2676 | 76205 |
| 1059.361 | 242 | 1062.182 | 298 | 746.0137 | 235 | 580.4343 | 221 | AR | None | Prostate | GSM24122 | 8425 | 76206 |
| 225.3911 | 204 | 324.1868 | 267 | 154.9334 | 285 | 182.2476 | 328 | CTCF | MCF10CA1 | Breast | GSM25990 | 40686 | 76247 |
| 224.97 | 192 | 302.3046 | 245 | 149.6457 | 264 | 171.0531 | 301 | CTCF | MCF10CA1 | Breast | GSM25990 | 37060 | 76248 |
| 266.1802 | 246 | 401.0613 | 327 | 182.7679 | 345 | 201.4407 | 389 | CTCF | None | Breast | GSM25990 | 50552 | 76249 |
| 193.1091 | 167 | 261.7374 | 214 | 122.4738 | 226 | 148.9948 | 263 | CTCF | None | Breast | GSM25990 | 32113 | 76250 |
| 445.4912 | 305 | 633.9717 | 402 | 286.2928 | 407 | 352.8268 | 476 | CTCF | MCF-10A | None | GSM25990 | 53246 | 76251 |
| 265.6846 | 186 | 364.406 | 240 | 171.8815 | 248 | 226.1094 | 297 | CTCF | MCF-10A | None | GSM25990 | 31627 | 76252 |
| 152.3458 | 66 | 170.461 | 78 | 147.1853 | 98 | 259.4232 | 136 | JUND | Calu-3 | Lung | GSM22662 | 7411 | 76277 |
| 80.34234 | 51 | 123.1897 | 70 | 90.75605 | 84 | 211.339 | 130 | JUN | Calu-3 | Lung | GSM22662 | 8195 | 76278 |
| 115.8316 | 80 | 126.1989 | 94 | 85.23343 | 112 | 220.3019 | 176 | JUN | Calu-3 | Lung | GSM22662 | 13624 | 76279 |
| 890.337 | 228 | 895.7948 | 286 | 591.9977 | 270 | 631.4163 | 298 | EHF | None | None | GSM22662 | 15385 | 76282 |
| 85.60664 | 29 | 128.1177 | 40 | 58.67639 | 36 | 146.5967 | 60 | EHF | None | None | GSM22662 | 2632 | 76283 |
| 397.3055 | 51 | 504.2157 | 64 | 238.2837 | 50 | 323.5501 | 63 | YY1 | LCL | Blood | GSM25972 | 1046 | 76288 |
| 594.6247 | 78 | 821.0927 | 103 | 342.4382 | 75 | 324.9007 | 77 | YY1 | LCL | Blood | GSM25972 | 1633 | 76289 |
| 708.5359 | 91 | 899.4453 | 114 | 380.8339 | 84 | 374.4677 | 88 | YY1 | LCL | Blood | GSM25972 | 1851 | 76290 |
| 956.4376 | 129 | 1197.355 | 175 | 624.0122 | 134 | 518.7785 | 128 | YY1 | LCL | Blood | GSM25972 | 2832 | 76291 |
| 928.1119 | 135 | 1167.936 | 188 | 559.2983 | 136 | 481.7641 | 133 | YY1 | LCL | Blood | GSM25972 | 3411 | 76292 |
| 540.5277 | 69 | 679.4055 | 86 | 351.9152 | 71 | 319.8913 | 71 | YY1 | LCL | Blood | GSM25972 | 1406 | 76293 |
| 886.7866 | 135 | 1138.329 | 185 | 504.7257 | 133 | 426.022 | 129 | YY1 | LCL | Blood | GSM25972 | 3715 | 76294 |
| 1087.179 | 187 | 1143.39 | 265 | 702.9541 | 190 | 606.968 | 187 | YY1 | LCL | Blood | GSM25972 | 5443 | 76295 |
| 1098.571 | 160 | 1171.622 | 225 | 645.5686 | 160 | 558.458 | 157 | YY1 | LCL | Blood | GSM25972 | 4229 | 76296 |
| 827.3557 | 108 | 1086.271 | 138 | 502.797 | 107 | 366.7156 | 95 | YY1 | LCL | Blood | GSM25972 | 2213 | 76297 |
| 682.159 | 91 | 953.6979 | 121 | 362.9199 | 84 | 334.5702 | 85 | YY1 | LCL | Blood | GSM25972 | 1963 | 76299 |
| 1150.473 | 163 | 1193.994 | 225 | 701.0339 | 161 | 594.8271 | 156 | YY1 | LCL | Blood | GSM25972 | 3766 | 76300 |
| 1131.671 | 156 | 1172.702 | 202 | 601.7589 | 146 | 421.7883 | 128 | YY1 | LCL | Blood | GSM25972 | 3695 | 76301 |
| 1072.694 | 221 | 1089.745 | 282 | 743.0497 | 217 | 565.1754 | 201 | YY1 | LCL | Blood | GSM25972 | 6901 | 76302 |
| 1122.158 | 168 | 1136.24 | 217 | 695.6247 | 171 | 523.8939 | 156 | YY1 | LCL | Blood | GSM25972 | 4357 | 76303 |
| 956.7775 | 133 | 1181.016 | 182 | 621.8618 | 139 | 510.1095 | 132 | YY1 | LCL | Blood | GSM25972 | 3169 | 76304 |
| 1151.3 | 208 | 1176.68 | 267 | 841.0519 | 201 | 631.3446 | 183 | YY1 | LCL | Blood | GSM25972 | 5074 | 76305 |
| 99.48652 | 26 | 120.8253 | 32 | 60.38037 | 29 | 72.6931 | 34 | STAT3 | None | Umbilical C | GSM25458 | 1661 | 76389 |
| 193.0219 | 32 | 232.0348 | 39 | 171.5527 | 41 | 190.3966 | 46 | STAT3 | None | Umbilical C | GSM25458 | 1074 | 76390 |
| 330.58 | 107 | 492.7112 | 146 | 206.1915 | 126 | 168.6963 | 124 | NFKB1 | GM12878 | None | GSM26280 | 8868 | 76400 |
| 330.7991 | 109 | 457.6751 | 143 | 186.4265 | 123 | 198.7858 | 136 | NFKB1 | GM12878 | None | GSM26280 | 9191 | 76401 |
| 566.9881 | 237 | 773.862 | 331 | 365.8274 | 294 | 301.8552 | 295 | RELB | GM12878 | None | GSM26280 | 25582 | 76402 |
| 517.2471 | 219 | 768.1334 | 305 | 368.6625 | 286 | 328.1675 | 295 | RELB | GM12878 | None | GSM26280 | 25370 | 76403 |
| 665.6679 | 211 | 836.7375 | 279 | 440.9953 | 254 | 296.8516 | 231 | RELA | GM12878 | None | GSM26280 | 16816 | 76405 |

| | | | | | | | | | | | | | |
|---|---|---|---|---|---|---|---|---|---|---|---|---|---|
| 609.5262 | 219 | 657.3477 | 253 | 278.2056 | 231 | 270.8964 | 247 | ESCO2 | vu1199-F S | None | GSM21398 | 20561 | 76412 |
| 607.0519 | 78 | 710.3588 | 93 | 234.5345 | 60 | 286.5114 | 71 | PMEL | None | None | GSM14230 | 1639 | 76413 |
| 879.2039 | 503 | 879.3528 | 607 | 603.5144 | 489 | 591.9459 | 542 | PMEL | None | None | GSM14230 | 36319 | 76415 |
| 11.32173 | 21 | 16.76913 | 28 | 74.56178 | 70 | 75.35225 | 76 | MAML3 | SK-N-SH | Brain | GSM16931 | 6378 | 76419 |
| 9.436583 | 21 | 15.5123 | 29 | 71.49714 | 73 | 88.41618 | 86 | MAML3 | SK-N-SH | Brain | GSM16930 | 7147 | 76420 |
| 11.40508 | 23 | 24.24528 | 35 | 62.58638 | 71 | 93.13278 | 90 | MAML3 | SK-N-SH | Brain | GSM16930 | 7557 | 76421 |
| 7.196186 | 10 | 29.28765 | 21 | 55.03439 | 38 | 89.28018 | 51 | MAML3 | SK-N-SH | Brain | GSM16930 | 2814 | 76422 |
| 836.4519 | 127 | 1177.884 | 196 | 636.5003 | 147 | 502.9488 | 137 | PHF2 | None | None | GSM19367 | 3566 | 76468 |
| 1134.938 | 206 | 1178.512 | 280 | 744.0408 | 192 | 697.7004 | 197 | PHF2 | None | None | GSM19367 | 5365 | 76469 |
| 1119.365 | 347 | 1133.473 | 435 | 854.9756 | 321 | 786.3271 | 297 | BRPF3 | RKO | Colon | GSM15561 | 9127 | 76494 |
| 1150.244 | 257 | 1169.65 | 329 | 863.4148 | 225 | 827.8999 | 234 | BRPF3 | RKO | Colon | GSM15561 | 5741 | 76495 |
| 1165.876 | 430 | 1188.416 | 537 | 880.7233 | 384 | 836.3006 | 389 | BRPF3 | RKO | Colon | GSM15561 | 10482 | 76496 |
| 933.5656 | 251 | 925.2418 | 298 | 473.1082 | 236 | 514.2748 | 263 | ZMYND8 | ZR-75-30 | None | GSM20295 | 13601 | 76498 |
| 1006.781 | 292 | 1025.13 | 375 | 860.7431 | 389 | 832.7827 | 408 | ZMYND8 | ZR-75-30 | None | GSM20295 | 13055 | 76499 |
| 1361.212 | 220 | 1340.374 | 252 | 1074.005 | 193 | 1021.672 | 185 | CREB1 | KG1 | None | GSM19379 | 2905 | 76508 |
| 1357.319 | 211 | 1340.882 | 244 | 1069.103 | 183 | 1039.835 | 192 | CREB1 | KG1 | None | GSM19379 | 2835 | 76509 |
| 1208.536 | 186 | 1224.04 | 232 | 667.5107 | 156 | 734.2735 | 174 | ZBTB16 | None | None | GSM19435 | 4078 | 76518 |
| 1208.759 | 732 | 1246.033 | 931 | 853.2466 | 699 | 791.4997 | 694 | SETD1A | None | None | GSM16983 | 31877 | 76519 |
| 1324.429 | 876 | 1350.483 | 1078 | 915.7844 | 827 | 875.6439 | 873 | BPTF | None | None | GSM16983 | 33716 | 76520 |
| 432.6307 | 150 | 613.7696 | 199 | 198.5338 | 158 | 209.0061 | 174 | BRD9 | NOMO-1 | None | GSM20928 | 14066 | 76526 |
| 1158.988 | 183 | 1203.865 | 259 | 875.189 | 194 | 818.4288 | 198 | EPC1 | K562 | None | GSM17052 | 4640 | 76527 |
| 518.4165 | 69 | 762.073 | 95 | 361.4441 | 75 | 368.0041 | 80 | EPC1 | K562 | None | GSM17052 | 1630 | 76528 |
| 1163.315 | 456 | 1188.582 | 582 | 887.7036 | 448 | 817.4016 | 424 | MBTD1 | K562 | None | GSM17052 | 14173 | 76529 |
| 1.049914 | 7 | 15.19184 | 20 | 0.740992 | 13 | 16.88489 | 33 | CHAT | SH-SY5Y | None | GSM18981 | 4605 | 76547 |
| 2.51922 | 21 | 1.712014 | 23 | 0.372706 | 30 | 5.988382 | 53 | CHAT | SH-SY5Y | None | GSM18981 | 12805 | 76548 |
| 82.88709 | 23 | 98.75349 | 28 | 77.83553 | 32 | 127.2895 | 44 | ZNF618 | 293T | None | GSM21114 | 1520 | 76567 |
| 1016.542 | 293 | 1059.507 | 400 | 764.5933 | 291 | 730.522 | 301 | UHRF2 | 293T | None | GSM21114 | 13034 | 76568 |
| 7.638093 | 7 | 5.567305 | 7 | 0 | 3 | 115.7761 | 43 | CENPC | HuRef | None | GSM17098 | 1585 | 76574 |
| 1031.823 | 323 | 1022.244 | 383 | 722.4196 | 288 | 732.0822 | 321 | SIX1 | None | Kidney | GSM19705 | 14172 | 76585 |
| 160.7884 | 40 | 230.4719 | 54 | 78.25693 | 40 | 226.5186 | 73 | SIX1 | None | Kidney | GSM19705 | 2463 | 76587 |
| 992.0082 | 393 | 999.3628 | 487 | 734.6424 | 393 | 704.1309 | 411 | ZMYND8 | DU145 | Prostate | GSM21872 | 19448 | 76673 |
| 1377.651 | 901 | 1510.17 | 1212 | 968.9534 | 867 | 938.5762 | 933 | ASXL1 | HEK293 | None | GSM12503 | 29226 | 76682 |
| 940.3568 | 100 | 1042.306 | 115 | 493.8647 | 89 | 589.6259 | 104 | CCND2 | MM1.S | Blood | GSM23028 | 1558 | 76683 |
| 854.123 | 157 | 1036.117 | 204 | 354.9426 | 138 | 339.0546 | 144 | CCND2 | MM1.S | Blood | GSM23028 | 6322 | 76686 |
| 773.8482 | 145 | 797.6007 | 161 | 470.7063 | 152 | 274.8788 | 124 | MGEA5 | BJAB | None | GSM22959 | 5183 | 76709 |
| 349.0066 | 182 | 476.0644 | 235 | 243.76 | 240 | 266.5875 | 269 | NEUROG2 | None | Fetal Lung | GSM19701 | 24112 | 76809 |
| 584.8667 | 274 | 739.9893 | 373 | 448.16 | 371 | 403.4033 | 386 | NEUROG2 | None | Fetal Lung | GSM19701 | 34134 | 76810 |
| 610.0959 | 276 | 705.6098 | 337 | 405.8584 | 352 | 356.1358 | 363 | NEUROG2 | None | Fetal Lung | GSM19701 | 33165 | 76811 |
| 716.7038 | 330 | 727.9343 | 411 | 447.946 | 406 | 454.841 | 443 | NEUROG2 | None | Fetal Lung | GSM19701 | 40087 | 76812 |
| 131.6516 | 75 | 151.5503 | 90 | 128.0476 | 114 | 127.7578 | 123 | NEUROG2 | None | Fetal Lung | GSM19701 | 10521 | 76813 |
| 504.9356 | 232 | 734.5997 | 313 | 355.9616 | 303 | 381.9509 | 337 | NEUROG2 | None | Fetal Lung | GSM19701 | 28033 | 76814 |
| 78.5376 | 74 | 134.1454 | 105 | 97.97358 | 129 | 131.2087 | 156 | NEUROG2 | None | Fetal Lung | GSM19701 | 15490 | 76815 |
| 117.6175 | 105 | 168.033 | 138 | 94.26602 | 155 | 148.7369 | 198 | NEUROG2 | None | Fetal Lung | GSM19701 | 20988 | 76816 |
| 153.8388 | 134 | 226.446 | 178 | 110.773 | 190 | 145.3469 | 227 | NEUROG2 | None | Fetal Lung | GSM19701 | 26321 | 76817 |
| 182.1012 | 152 | 252.6845 | 197 | 126.6015 | 212 | 171.3299 | 256 | NEUROG2 | None | Fetal Lung | GSM19701 | 29098 | 76818 |
| 47.49689 | 59 | 88.19395 | 86 | 47.21079 | 97 | 103.8538 | 139 | NEUROG2 | None | Fetal Lung | GSM19701 | 14775 | 76819 |
| 87.51927 | 75 | 86.68074 | 85 | 74.68958 | 113 | 103.2161 | 138 | NEUROG2 | None | Fetal Lung | GSM19701 | 14658 | 76820 |
| 292.8825 | 39 | 393.7826 | 51 | 239.7937 | 47 | 313.9595 | 58 | SETX | A549 | Lung | GSM12783 | 1006 | 76830 |
| 365.3117 | 56 | 436.7939 | 68 | 195.1659 | 54 | 226.0029 | 62 | SETX | A549 | Lung | GSM12783 | 1770 | 76831 |
| 172.5074 | 80 | 182.5063 | 92 | 148.7443 | 113 | 75.86264 | 93 | EBF3 | SK-N-SH | Brain | GSM24103 | 9203 | 76847 |
| 56.42545 | 22 | 41.38757 | 21 | 40.50761 | 28 | 43.09463 | 31 | EBF3 | SK-N-SH | Brain | GSM24103 | 2123 | 76848 |
| 666.0665 | 361 | 672.5163 | 445 | 421.161 | 464 | 509.1703 | 540 | EBF3 | SK-N-SH | Brain | GSM24103 | 51863 | 76849 |
| 1109.666 | 203 | 1121.244 | 255 | 657.2697 | 185 | 820.8166 | 221 | ZMYND8 | HEK293 | Embryonic | GSM23225 | 5720 | 76851 |
| 230.6905 | 121 | 252.3329 | 141 | 81.13054 | 120 | 170.1131 | 172 | CDCA2 | HeLa | Cervix | GSM22261 | 13613 | 76854 |
| 1137.107 | 261 | 1152.008 | 329 | 856.0437 | 232 | 879.8818 | 276 | ZMYND8 | HEK293 | Kidney | GSM21714 | 6805 | 76856 |
| 1060.67 | 287 | 1084.3 | 366 | 776.3165 | 261 | 816.5831 | 320 | ZMYND8 | HEK293 | Kidney | GSM21714 | 9892 | 76857 |
| 615.2719 | 93 | 764.3308 | 115 | 263.3036 | 78 | 313.9197 | 91 | IRAK4 | SEM | None | GSM23737 | 2462 | 76858 |
| 1119.508 | 156 | 1137.346 | 184 | 493.679 | 132 | 519.717 | 144 | IRAK1 | HEK293 | Embryonic | GSM23736 | 3727 | 76859 |
| 10.10978 | 7 | 12.97325 | 9 | 22.24427 | 16 | 52.81854 | 26 | CD74 | None | None | GSM23563 | 1316 | 76925 |
| 40.12675 | 19 | 35.16152 | 20 | 57.09613 | 34 | 110.2977 | 50 | CD74 | None | None | GSM23563 | 2300 | 76926 |
| 55.52085 | 26 | 58.86854 | 30 | 74.30003 | 45 | 87.15738 | 52 | CD74 | None | None | GSM23563 | 3067 | 76927 |
| 28.78784 | 11 | 9.44605 | 7 | 20.3015 | 14 | 29.53814 | 18 | CD74 | None | None | GSM23563 | 1145 | 76928 |
| 657.9267 | 113 | 868.154 | 145 | 391.4192 | 117 | 377.5031 | 122 | CD74 | None | None | GSM23563 | 3965 | 76930 |
| 21.18992 | 11 | 35.08532 | 16 | 9.32796 | 12 | 34.4985 | 23 | CD74 | None | None | GSM23563 | 1555 | 76931 |
| 949.5041 | 572 | 1017.304 | 782 | 714.2334 | 646 | 666.1667 | 657 | CNOT3 | HCT-116 | None | GSM16173 | 39715 | 76964 |
| 942.5014 | 702 | 971.7479 | 881 | 667.0392 | 768 | 658.7472 | 863 | CNOT3 | None | Snap-froze | GSM16173 | 60573 | 76965 |
| 385.8947 | 90 | 482.8942 | 112 | 208.7759 | 93 | 198.2787 | 97 | MBOAT4 | HeLa | Cervix | GSM15874 | 4711 | 76967 |
| 57.35334 | 15 | 79.55153 | 20 | 48.78365 | 20 | 66.14626 | 25 | LRWD1 | U2OS | None | GSM21440 | 1027 | 76970 |
| 67.87942 | 20 | 61.24455 | 21 | 37.87089 | 22 | 32.534 | 22 | LRWD1 | U2OS | None | GSM21440 | 1564 | 76971 |
| 794.2983 | 169 | 986.3682 | 225 | 309.5618 | 147 | 359.8262 | 169 | GLI2 | THP-1 | None | GSM22122 | 7706 | 76995 |
| 818.0014 | 320 | 832.5668 | 402 | 612.5602 | 371 | 578.7261 | 391 | PITX3 | SH-SY5Y | Midbrain | GSM24499 | 24874 | 76999 |
| 133.518 | 50 | 129.7729 | 55 | 48.8157 | 48 | 196.9866 | 96 | ZNF93 | 293T | None | GSM24666 | 4936 | 77018 |

| | | | | | | | | | | | | | |
|---|---|---|---|---|---|---|---|---|---|---|---|---|---|
| 21.74007 | 11 | 27.57663 | 14 | 18.46001 | 16 | 188.6352 | 54 | ZNF879 | 293T | None | GSM24666 | 1607 | 77022 |
| 218.2492 | 36 | 263.2985 | 44 | 123.05 | 36 | 464.048 | 81 | ZNF846 | 293T | None | GSM24666 | 1250 | 77024 |
| 20.70158 | 18 | 26.80034 | 23 | 13.47534 | 25 | 157.4356 | 76 | ZNF823 | 293T | None | GSM24666 | 3843 | 77025 |
| 15.54622 | 9 | 33.41432 | 15 | 17.19821 | 15 | 152.908 | 47 | ZNF81 | 293T | None | GSM24666 | 1522 | 77026 |
| 35.33338 | 22 | 68.22819 | 34 | 29.26481 | 32 | 100.5401 | 59 | ZNF808 | 293T | None | GSM24666 | 3535 | 77027 |
| 76.76881 | 45 | 95.00782 | 56 | 45.90893 | 57 | 141.7704 | 98 | ZNF793 | 293T | None | GSM24666 | 6747 | 77032 |
| 57.16726 | 70 | 98.49946 | 99 | 39.73384 | 103 | 74.86869 | 138 | ZNF793 | 293T | None | GSM24666 | 17863 | 77033 |
| 365.287 | 92 | 488.7021 | 119 | 178.8323 | 93 | 317.2993 | 131 | ZNF786 | 293T | None | GSM24666 | 5743 | 77037 |
| 982.3006 | 141 | 1149.709 | 184 | 415.7104 | 118 | 647.7473 | 159 | ZNF783 | 293T | None | GSM24666 | 3717 | 77039 |
| 81.80761 | 25 | 62.1488 | 24 | 27.77061 | 22 | 244.1821 | 69 | ZNF780A | 293T | None | GSM24666 | 1994 | 77041 |
| 132.2244 | 26 | 160.7019 | 32 | 50.99274 | 22 | 313.6241 | 63 | ZNF777 | 293T | None | GSM24666 | 1192 | 77043 |
| 108.7645 | 45 | 116.1729 | 52 | 76.67346 | 58 | 168.6165 | 89 | ZNF765 | 293T | None | GSM24666 | 4892 | 77045 |
| 34.3087 | 30 | 49.56069 | 40 | 31.71862 | 47 | 84.85539 | 75 | ZNF75D | 293T | None | GSM24666 | 6132 | 77047 |
| 51.26124 | 41 | 78.48378 | 56 | 40.91506 | 60 | 99.88037 | 92 | ZNF75D | 293T | None | GSM24666 | 7675 | 77048 |
| 2.12881 | 4 | 3.834402 | 6 | 1.022764 | 6 | 48.88206 | 28 | ZNF736 | 293T | None | GSM24666 | 1550 | 77052 |
| 81.80739 | 22 | 98.50018 | 27 | 25.39879 | 18 | 202.5948 | 55 | ZNF730 | 293T | None | GSM24666 | 1524 | 77053 |
| 54.62102 | 17 | 72.06874 | 22 | 41.7814 | 22 | 193.0507 | 52 | ZNF716 | 293T | None | GSM24666 | 1437 | 77054 |
| 721.7907 | 148 | 841.171 | 177 | 365.0913 | 147 | 654.5219 | 210 | ZNF695 | 293T | None | GSM24666 | 7149 | 77059 |
| 242.9837 | 62 | 414.1381 | 92 | 154.35 | 71 | 285.9 | 103 | ZNF682 | 293T | None | GSM24666 | 3921 | 77061 |
| 100.4803 | 30 | 280.8502 | 59 | 71.02175 | 37 | 232.4466 | 72 | ZNF682 | 293T | None | GSM24666 | 2318 | 77062 |
| 39.18015 | 29 | 54.06223 | 38 | 18.50946 | 35 | 103.9731 | 75 | ZNF675 | 293T | None | GSM24666 | 5353 | 77064 |
| 317.3886 | 77 | 442.3396 | 102 | 194.4281 | 86 | 317.0574 | 117 | ZNF674 | 293T | None | GSM24666 | 4568 | 77065 |
| 238.1186 | 64 | 415.6068 | 96 | 188.558 | 82 | 267.0921 | 104 | ZNF671 | 293T | None | GSM24666 | 4231 | 77066 |
| 10.98886 | 15 | 10.40872 | 17 | 5.16884 | 20 | 74.83638 | 58 | ZNF649 | 293T | None | GSM24666 | 4339 | 77068 |
| 12.08363 | 8 | 37.35614 | 16 | 14.48958 | 14 | 163.6952 | 49 | ZNF627 | 293T | None | GSM24666 | 1528 | 77069 |
| 35.72128 | 14 | 85.30521 | 25 | 25.21379 | 18 | 236.0439 | 60 | ZNF616 | 293T | None | GSM24666 | 1529 | 77074 |
| 81.26323 | 36 | 172.6581 | 59 | 37.56683 | 39 | 186.922 | 86 | ZNF611 | 293T | None | GSM24666 | 4065 | 77078 |
| 53.46015 | 21 | 85.51114 | 30 | 31.81221 | 25 | 153.5395 | 57 | ZNF605 | 293T | None | GSM24666 | 2254 | 77079 |
| 81.69722 | 40 | 93.74323 | 48 | 34.70515 | 43 | 133.7435 | 82 | ZNF585A | 293T | None | GSM24666 | 5124 | 77081 |
| 55.61796 | 20 | 121.6917 | 34 | 52.16399 | 29 | 192.8462 | 60 | ZNF573 | 293T | None | GSM24666 | 1946 | 77082 |
| 23.36833 | 11 | 43.10986 | 17 | 36.46351 | 21 | 168.0215 | 49 | ZNF567 | 293T | None | GSM24666 | 1473 | 77085 |
| 59.8222 | 17 | 21.67107 | 11 | 15.62962 | 13 | 175.7883 | 47 | ZNF565 | 293T | None | GSM24665 | 1286 | 77087 |
| 191.9956 | 37 | 301.9564 | 53 | 148.2618 | 45 | 352.5428 | 77 | ZNF561 | 293T | None | GSM24665 | 1629 | 77090 |
| 158.5644 | 29 | 112.7349 | 26 | 155.6792 | 40 | 305.2511 | 62 | ZNF557 | 293T | None | GSM24665 | 1188 | 77091 |
| 139.8701 | 25 | 163.114 | 30 | 94.65676 | 28 | 256.2439 | 52 | ZNF555 | 293T | None | GSM24665 | 1000 | 77092 |
| 229.4537 | 43 | 415.3089 | 67 | 196.0326 | 55 | 340.1664 | 79 | ZNF543 | 293T | None | GSM24665 | 1803 | 77096 |
| 800.2899 | 365 | 792.6414 | 438 | 405.7426 | 365 | 511.3634 | 432 | ZNF534 | 293T | None | GSM24665 | 36797 | 77098 |
| 80.71977 | 22 | 131.324 | 32 | 53.66955 | 26 | 248.2566 | 62 | ZNF530 | 293T | None | GSM24665 | 1549 | 77099 |
| 945.3085 | 289 | 972.1786 | 381 | 580.3854 | 281 | 644.2322 | 316 | ZNF519 | 293T | None | GSM24665 | 16964 | 77103 |
| 42.35644 | 45 | 80.7021 | 67 | 47.51791 | 77 | 73.46298 | 98 | ZNF506 | 293T | None | GSM24665 | 10670 | 77104 |
| 22.95949 | 30 | 31.46491 | 39 | 12.05732 | 41 | 79.73233 | 86 | ZNF485 | 293T | None | GSM24665 | 8153 | 77108 |
| 27.04 | 12 | 47.09324 | 18 | 22.65316 | 17 | 260.4278 | 63 | ZNF480 | 293T | None | GSM24665 | 1508 | 77111 |
| 531.7617 | 261 | 731.3364 | 365 | 298.9071 | 319 | 270.6009 | 334 | ZNF479 | 293T | None | GSM24665 | 35751 | 77112 |
| 453.8731 | 117 | 929.1002 | 191 | 305.4363 | 138 | 419.9191 | 172 | ZNF468 | 293T | None | GSM24665 | 7432 | 77113 |
| 167.8833 | 37 | 207.9862 | 46 | 76.71442 | 35 | 250.1241 | 69 | ZNF460 | 293T | None | GSM24665 | 1945 | 77114 |
| 152.5458 | 42 | 161.8617 | 48 | 124.4291 | 55 | 252.581 | 84 | ZNF44 | 293T | None | GSM24665 | 2943 | 77116 |
| 85.95254 | 26 | 81.27676 | 28 | 72.72733 | 35 | 150.858 | 54 | ZNF445 | 293T | None | GSM24665 | 2022 | 77117 |
| 313.8178 | 142 | 436.6956 | 186 | 106.2103 | 140 | 229.9624 | 204 | ZNF441 | 293T | None | GSM24665 | 17277 | 77120 |
| 18.81386 | 13 | 43.28883 | 22 | 67.85075 | 37 | 109.7995 | 50 | ZNF440 | 293T | None | GSM24665 | 2427 | 77121 |
| 8.580989 | 6 | 14.88772 | 9 | 6.939013 | 9 | 165.2932 | 45 | ZNF433 | 293T | None | GSM24665 | 1255 | 77123 |
| 28.41514 | 14 | 80.53366 | 27 | 56.3695 | 30 | 187.9356 | 59 | ZNF430 | 293T | None | GSM24665 | 1960 | 77126 |
| 71.12581 | 45 | 163.5974 | 75 | 85.31485 | 76 | 130.4017 | 98 | ZNF429 | 293T | None | GSM24665 | 7206 | 77127 |
| 130.4526 | 57 | 162.2451 | 71 | 55.29169 | 60 | 176.3961 | 106 | ZNF425 | 293T | None | GSM24665 | 6524 | 77128 |
| 148.1426 | 28 | 167.5534 | 33 | 54.83478 | 23 | 390.4114 | 72 | ZNF383 | 293T | None | GSM24665 | 1207 | 77131 |
| 962.1463 | 145 | 1095.531 | 173 | 454.006 | 131 | 497.7038 | 146 | ZNF343 | 293T | None | GSM24665 | 4412 | 77133 |
| 28.5251 | 13 | 42.77475 | 18 | 22.48992 | 18 | 199.8531 | 57 | ZNF337 | 293T | None | GSM24665 | 1684 | 77135 |
| 7.99181 | 7 | 18.85888 | 12 | 4.23083 | 9 | 43.12024 | 26 | ZNF334 | 293T | None | GSM24665 | 1568 | 77136 |
| 127.9793 | 29 | 181.2362 | 39 | 78.58118 | 32 | 324.438 | 73 | ZNF333 | 293T | None | GSM24665 | 1601 | 77137 |
| 59.27458 | 21 | 106.2721 | 32 | 54.04766 | 30 | 211.7565 | 64 | ZNF304 | 293T | None | GSM24665 | 2013 | 77140 |
| 207.8792 | 97 | 292.9482 | 128 | 93.38237 | 106 | 188.899 | 152 | ZNF28 | 293T | None | GSM24665 | 11707 | 77143 |
| 17.02205 | 12 | 34.17944 | 19 | 18.50485 | 20 | 130.9577 | 53 | ZNF287 | 293T | None | GSM24665 | 2273 | 77144 |
| 31.32845 | 59 | 27.89983 | 66 | 8.410715 | 73 | 40.05535 | 120 | ZNF284 | 293T | None | GSM24665 | 20075 | 77146 |
| 706.8955 | 137 | 917.6121 | 174 | 277.1587 | 118 | 427.0969 | 156 | ZNF282 | 293T | None | GSM24665 | 5892 | 77148 |
| 800.4292 | 158 | 1023.809 | 216 | 393.3845 | 153 | 545.1977 | 192 | ZNF282 | 293T | None | GSM24665 | 7014 | 77149 |
| 736.6761 | 161 | 994.3397 | 230 | 422.8254 | 172 | 483.217 | 196 | ZNF273 | 293T | None | GSM24665 | 8438 | 77150 |
| 4.402437 | 7 | 7.250591 | 10 | 10.65937 | 17 | 106.4071 | 50 | ZNF26 | 293T | None | GSM24665 | 2478 | 77151 |
| 56.97571 | 19 | 33.21271 | 16 | 27.54536 | 20 | 222.044 | 61 | ZNF267 | 293T | None | GSM24665 | 1721 | 77152 |
| 30.26213 | 11 | 38.44668 | 14 | 24.49879 | 15 | 262.2995 | 56 | ZNF254 | 293T | None | GSM24665 | 1151 | 77154 |
| 1.563221 | 13 | 6.873313 | 23 | 0.733047 | 22 | 23.91353 | 56 | ZNF248 | 293T | None | GSM24665 | 8300 | 77155 |
| 108.5607 | 27 | 130.0649 | 33 | 61.23101 | 29 | 288.9249 | 70 | ZNF222 | 293T | None | GSM24664 | 1690 | 77158 |
| 596.5332 | 121 | 1045.181 | 186 | 372.3395 | 133 | 607.3693 | 182 | ZNF189 | 293T | None | GSM24664 | 5722 | 77164 |
| 79.31664 | 52 | 76.72462 | 58 | 40.16852 | 63 | 100.4456 | 97 | ZNF182 | 293T | None | GSM24664 | 8523 | 77165 |

| | | | | | | | | | | | | | |
|---|---|---|---|---|---|---|---|---|---|---|---|---|---|
| 118.289 | 24 | 182.0051 | 34 | 100.6852 | 31 | 303.7303 | 61 | ZNF180 | 293T | None | GSM24664 | 1154 | 77167 |
| 15.49184 | 13 | 59.93062 | 28 | 10.02855 | 18 | 127.2272 | 58 | ZNF17 | 293T | None | GSM24664 | 2788 | 77168 |
| 40.81556 | 17 | 68.92992 | 25 | 31.85601 | 23 | 205.4271 | 62 | ZNF169 | 293T | None | GSM24664 | 1923 | 77169 |
| 8.963411 | 10 | 45.52478 | 24 | 12.48283 | 19 | 161.6732 | 64 | ZNF169 | 293T | None | GSM24664 | 2616 | 77170 |
| 23.2671 | 11 | 38.23628 | 16 | 23.07774 | 17 | 206.3497 | 55 | ZNF157 | 293T | None | GSM24664 | 1492 | 77171 |
| 176.9671 | 71 | 208.5351 | 86 | 130.9854 | 93 | 194.2536 | 119 | ZNF141 | 293T | None | GSM24664 | 7417 | 77173 |
| 30.07676 | 14 | 57.66931 | 22 | 37.79639 | 24 | 197.9877 | 59 | ZNF135 | 293T | None | GSM24664 | 1824 | 77174 |
| 21.76399 | 29 | 20.98997 | 33 | 24.88632 | 51 | 55.66758 | 74 | ZNF133 | 293T | None | GSM24664 | 8031 | 77175 |
| 38.24272 | 14 | 66.10741 | 21 | 41.74064 | 22 | 206.5556 | 54 | ZNF124 | 293T | None | GSM24664 | 1434 | 77177 |
| 5.662781 | 5 | 25.26713 | 12 | 6.568341 | 9 | 135.4569 | 41 | ZNF10 | 293T | None | GSM24664 | 1295 | 77179 |
| 12.37186 | 7 | 52.71911 | 17 | 26.50148 | 16 | 245.2951 | 55 | ZNF100 | 293T | None | GSM24664 | 1216 | 77180 |
| 270.1736 | 65 | 390.8195 | 88 | 182.5585 | 76 | 293.7706 | 103 | ZNF100 | 293T | None | GSM24664 | 3794 | 77181 |
| 126.7693 | 34 | 189.4236 | 47 | 71.20488 | 37 | 239.0632 | 73 | ZNF100 | 293T | None | GSM24664 | 2340 | 77182 |
| 439.8288 | 107 | 609.5576 | 141 | 189.1733 | 101 | 411.2498 | 157 | ZKSCAN5 | 293T | None | GSM24664 | 6309 | 77183 |
| 397.1689 | 52 | 359.4843 | 53 | 131.0571 | 37 | 478.8227 | 82 | ZKSCAN2 | 293T | None | GSM24664 | 1216 | 77185 |
| 18.13949 | 10 | 31.80511 | 15 | 16.15309 | 15 | 141.1131 | 46 | ZFP69 | 293T | None | GSM24664 | 1580 | 77189 |
| 300.3377 | 59 | 388.2854 | 75 | 240.7024 | 73 | 407.9855 | 103 | ZSCAN2 | MCF-7 | None | GSM25748 | 2557 | 77254 |
| 765.2877 | 142 | 876.5899 | 168 | 514.5279 | 160 | 507.6921 | 169 | DPF1 | MCF-7 | None | GSM25748 | 5913 | 77255 |
| 1004.731 | 493 | 999.6984 | 586 | 828.4712 | 628 | 802.9679 | 678 | ZXDC | MCF-7 | None | GSM25748 | 28106 | 77256 |
| 1236.29 | 191 | 1248.869 | 241 | 868.3174 | 177 | 715.2328 | 168 | ZNF75A | MCF-7 | None | GSM25748 | 3816 | 77257 |
| 968.3811 | 381 | 971.4424 | 468 | 831.9493 | 525 | 833.6977 | 603 | ZNF701 | MCF-7 | None | GSM25748 | 21900 | 77258 |
| 1075.355 | 334 | 1068.348 | 401 | 879.3914 | 398 | 881.2437 | 456 | WDHD1 | MCF-7 | None | GSM25748 | 13212 | 77259 |
| 1055.15 | 310 | 1056.911 | 379 | 878.6566 | 386 | 883.9649 | 445 | WDHD1 | MCF-7 | None | GSM25748 | 12746 | 77260 |
| 392.5348 | 144 | 468.4984 | 175 | 206.7972 | 161 | 202.1272 | 172 | ZNF597 | GM12878 | Blood | GSM25747 | 13675 | 77261 |
| 112.6265 | 61 | 93.34314 | 63 | 49.13056 | 67 | 57.22344 | 77 | HMG20A | GM12878 | Blood | GSM25747 | 8051 | 77262 |
| 386.2511 | 199 | 465.5235 | 243 | 338.2013 | 287 | 534.6424 | 373 | NFIA | K562 | Bone Marr | GSM25747 | 26588 | 77263 |
| 227.0139 | 56 | 131.2458 | 46 | 88.14865 | 50 | 62.78468 | 46 | DPF1 | K562 | Bone Marr | GSM25747 | 3262 | 77264 |
| 152.6923 | 34 | 150.1968 | 37 | 84.08362 | 35 | 159.6555 | 52 | ZSCAN2 | HepG2 | Liver | GSM25747 | 1692 | 77266 |
| 104.6518 | 36 | 104.8135 | 40 | 72.06407 | 44 | 96.11169 | 54 | NFIA | HepG2 | Liver | GSM25747 | 2925 | 77267 |
| 1054.47 | 324 | 1016.074 | 360 | 787.229 | 313 | 761.2599 | 332 | L3MBTL4 | HepG2 | Liver | GSM25747 | 13425 | 77271 |
| 574.9084 | 211 | 795.6944 | 281 | 296.9396 | 233 | 327.8872 | 262 | TRIM25 | None | Breast | GSM20987 | 19271 | 77298 |
| 922.0303 | 501 | 980.1362 | 679 | 701.0336 | 561 | 680.8128 | 610 | TRIM25 | BT-549 | Breast | GSM20987 | 33594 | 77299 |
| 974.8845 | 510 | 1036.946 | 696 | 730.201 | 548 | 683.0718 | 555 | TRIM25 | BT-549 | Breast | GSM20987 | 30197 | 77300 |
| 240.0277 | 177 | 299.087 | 219 | 144.6115 | 229 | 157.5814 | 257 | SP140 | None | None | GSM23989 | 29159 | 77331 |
| 122.345 | 38 | 139.0634 | 45 | 7.760206 | 17 | 13.96406 | 23 | SP140 | None | None | GSM23989 | 2682 | 77332 |
| 197.9163 | 42 | 294.8699 | 58 | 72.48485 | 35 | 334.957 | 83 | TAZ | SW480 | Colon | GSM24028 | 1959 | 77366 |
| 567.0484 | 124 | 689.9512 | 152 | 365.843 | 140 | 244.5414 | 123 | H2AFX | IMR90 | Lung | GSM20679 | 6429 | 77448 |
| 548.4441 | 233 | 796.9589 | 349 | 432.2205 | 318 | 379.4218 | 326 | NFKB2 | L1236 | None | GSM15563 | 27619 | 77449 |
| 339.3936 | 182 | 702.0726 | 285 | 259.2036 | 252 | 287.939 | 284 | NFKB2 | L1236 | None | GSM15563 | 26328 | 77450 |
| 172.7402 | 42 | 314.5334 | 65 | 107.3836 | 47 | 266.4946 | 80 | SON | K562 | None | GSM20405 | 2448 | 77457 |
| 468.7055 | 73 | 720.7355 | 103 | 327.7094 | 81 | 329.248 | 86 | TP53 | None | Breast | GSM16247 | 2171 | 77458 |
| 926.8147 | 643 | 940.4802 | 793 | 653.0778 | 687 | 616.1404 | 721 | KMT2A | MV4-11 | None | GSM18973 | 53321 | 77486 |
| 391.193 | 68 | 463.482 | 82 | 231.0859 | 70 | 323.4335 | 89 | HIF1A | U2OS | Bone | GSM22576 | 2399 | 77518 |
| 162.6772 | 68 | 305.2819 | 104 | 97.43704 | 81 | 101.8253 | 89 | HIF1A | U2OS | Bone | GSM22576 | 6983 | 77519 |
| 1124.009 | 249 | 1162.03 | 334 | 846.7957 | 231 | 796.7759 | 238 | TFDP1 | U266B1 | None | GSM21325 | 6779 | 77534 |
| 1210.751 | 372 | 1235.786 | 477 | 913.6907 | 325 | 862.6786 | 319 | TFDP1 | MM1.S | Blood | GSM21325 | 8088 | 77535 |
| 1074.208 | 478 | 1112.468 | 624 | 804.5883 | 482 | 758.2625 | 486 | REL | GM12878 | None | GSM26280 | 20837 | 77536 |
| 1197.727 | 214 | 1204.189 | 265 | 917.858 | 202 | 864.6298 | 207 | REL | GM12878 | None | GSM26280 | 4764 | 77537 |
| 713.5918 | 463 | 764.2887 | 626 | 548.1334 | 615 | 533.2418 | 675 | CTCF | H1 | Embryo | GSM62407 | 68304 | 8086 |
| 714.0821 | 510 | 752.8287 | 671 | 536.4481 | 661 | 518.2803 | 725 | CTCF | HeLa | Cervix | GSM62407 | 78170 | 8087 |
| 453.8263 | 299 | 650.5847 | 406 | 314.285 | 410 | 374.2364 | 474 | CTCF | None | Umbilical V | GSM62407 | 52681 | 8088 |
| 640.521 | 406 | 702.6733 | 570 | 494.0226 | 569 | 491.7907 | 633 | CTCF | K562 | Bone Marr | GSM62408 | 71615 | 8089 |
| 852.5296 | 687 | 939.2112 | 940 | 653.9117 | 881 | 615.6505 | 936 | CTCF | NHEK | Skin | GSM62408 | 82596 | 8090 |
| 1115.512 | 152 | 1195.62 | 209 | 666.3131 | 153 | 759.2133 | 174 | MYC | GM12878 | Blood | GSM74852 | 3857 | 8091 |
| 1203.543 | 140 | 1284.143 | 194 | 686.3678 | 134 | 752.3765 | 149 | MYC | GM12878 | Blood | GSM74852 | 2743 | 8092 |
| 153.1976 | 34 | 203.2946 | 44 | 82.60954 | 35 | 163.0052 | 53 | MYC | H1 | Embryo | GSM74853 | 1814 | 8096 |
| 492.8889 | 86 | 746.219 | 120 | 494.4448 | 119 | 494.6105 | 126 | MYC | HeLa | Cervix | GSM74853 | 3221 | 8098 |
| 681.4227 | 133 | 829.5915 | 163 | 622.0997 | 177 | 747.1453 | 207 | MYC | HeLa | Cervix | GSM74853 | 5947 | 8099 |
| 343.3582 | 245 | 555.7516 | 338 | 250.7093 | 345 | 321.1974 | 409 | CTCF | HepG2 | Liver | GSM74853 | 45202 | 8103 |
| 479.3657 | 316 | 644.9882 | 419 | 311.2703 | 425 | 376.9144 | 494 | CTCF | HepG2 | Liver | GSM74853 | 56560 | 8104 |
| 1062.396 | 315 | 1081.568 | 409 | 843.4522 | 348 | 828.341 | 379 | MYC | HepG2 | Liver | GSM74854 | 12658 | 8105 |
| 959.7345 | 145 | 1088.88 | 171 | 554.2789 | 146 | 555.3461 | 155 | MYC | HepG2 | Liver | GSM74854 | 4424 | 8106 |
| 1129.719 | 295 | 1142.658 | 376 | 879.4029 | 299 | 856.0031 | 316 | MYC | HepG2 | Liver | GSM74854 | 9310 | 8107 |
| 1099.519 | 595 | 1109.846 | 733 | 795.0956 | 578 | 773.8567 | 629 | MYC | None | Umbilical V | GSM74854 | 28038 | 8112 |
| 1126.711 | 160 | 1173.616 | 218 | 547.9322 | 143 | 477.016 | 141 | AUTS2 | HeLa | Cervix | GSM28601 | 3881 | 81145 |
| 1108.476 | 229 | 1165.365 | 316 | 894.6704 | 232 | 849.4608 | 235 | ZFX | HCT-116 | Colon | GSM27417 | 5475 | 81159 |
| 1063.631 | 344 | 1086.279 | 440 | 824.1087 | 358 | 790.6049 | 374 | MYC | K562 | Bone Marr | GSM74855 | 13970 | 8116 |
| 1061.749 | 367 | 1081.853 | 466 | 810.865 | 352 | 779.2535 | 369 | ZFX | HCT-116 | Colon | GSM27417 | 11606 | 81160 |
| 1053.305 | 311 | 1077.07 | 395 | 812.449 | 297 | 799.1332 | 325 | ZFX | None | Kidney | GSM27417 | 9142 | 81161 |
| 879.0466 | 308 | 887.8818 | 382 | 490.4829 | 293 | 523.9857 | 325 | ZFX | None | Kidney | GSM27417 | 17355 | 81162 |
| 1092.558 | 463 | 1126.494 | 601 | 818.6163 | 465 | 794.9955 | 497 | MYC | K562 | Bone Marr | GSM74855 | 19356 | 8117 |
| 922.651 | 577 | 859.5912 | 612 | 746.249 | 745 | 770.3697 | 895 | FOXA1 | 22RV1 | Prostate | GSM22771 | 45274 | 81170 |

| | | | | | | | | | | | | | |
|---|---|---|---|---|---|---|---|---|---|---|---|---|---|
| 1152.019 | 523 | 1187.351 | 676 | 881.8159 | 528 | 840.6457 | 541 | MYCN | SHEP-21N | Bone Marr | GSM21135 | 17140 | 81174 |
| 1127.99 | 361 | 1165.27 | 474 | 887.1527 | 379 | 831.0321 | 366 | MYC | K562 | Bone Marr | GSM74855 | 11955 | 8118 |
| 999.1822 | 341 | 984.6369 | 400 | 798.5694 | 387 | 798.0652 | 443 | FXR2 | AGS | None | GSM27528 | 15713 | 81190 |
| 1010.126 | 254 | 1061.998 | 357 | 808.1673 | 290 | 779.8614 | 304 | MYC | NHEK | Skin | GSM74855 | 11540 | 8122 |
| 772.3655 | 634 | 781.4451 | 775 | 547.3487 | 754 | 514.1237 | 801 | RUNX1 | None | Blood | GSM26875 | 87449 | 81223 |
| 821.9276 | 560 | 843.9123 | 705 | 593.1012 | 645 | 550.8173 | 668 | TET2 | None | Blood | GSM26875 | 61270 | 81224 |
| 126.5753 | 25 | 173.297 | 33 | 78.65053 | 27 | 435.3282 | 75 | CTCF | 226LDM | None | GSM27315 | 1111 | 81228 |
| 614.1228 | 86 | 689.6058 | 100 | 198.092 | 61 | 300.0788 | 81 | SMARCA4 | HepG2 | Liver | GSM27409 | 2030 | 81230 |
| 90.19667 | 158 | 99.32389 | 188 | 54.28065 | 229 | 79.8538 | 279 | EP300 | None | None | GSM25502 | 46128 | 81281 |
| 160.0963 | 30 | 262.58 | 44 | 101.6037 | 32 | 124.7507 | 38 | BRD4 | A549 | Lung | GSM23594 | 1095 | 81291 |
| 779.7354 | 145 | 803.3823 | 161 | 301.485 | 121 | 334.3726 | 136 | BRD4 | A375 | Skin | GSM23594 | 5219 | 81292 |
| 874.0955 | 375 | 877.2594 | 457 | 639.1721 | 387 | 617.1907 | 415 | BRD4 | A375 | Skin | GSM23594 | 24974 | 81293 |
| 952.511 | 328 | 957.4166 | 402 | 710.4159 | 330 | 695.5587 | 359 | BRD4 | A375 | Skin | GSM23594 | 16349 | 81294 |
| 985.5245 | 263 | 978.194 | 314 | 611.5368 | 247 | 679.2136 | 278 | BRD4 | A375 | Skin | GSM23594 | 10661 | 81295 |
| 414.7724 | 55 | 525.4217 | 69 | 268.9222 | 57 | 267.286 | 60 | MORC2 | HeLa | Cervix | GSM25144 | 1245 | 81315 |
| 415.9842 | 54 | 555.4388 | 70 | 223.5363 | 50 | 232.1145 | 54 | MORC2 | HeLa | Cervix | GSM25144 | 1164 | 81316 |
| 0.093063 | 8 | 3.40053 | 19 | 2.380732 | 28 | 26.63212 | 59 | MAF | None | Blood | GSM25928 | 8804 | 81326 |
| 423.4351 | 239 | 602.2764 | 313 | 253.4319 | 300 | 255.2697 | 327 | RAD21 | HCT-116 | Colon | GSM28096 | 33507 | 81366 |
| 169.908 | 30 | 297.1908 | 46 | 118.2342 | 34 | 100.1772 | 33 | NUP153 | K562 | Bone Marr | GSM26352 | 1156 | 81390 |
| 307.3231 | 42 | 416.1673 | 55 | 121.0648 | 33 | 161.1062 | 41 | NUP98 | K562 | Bone Marr | GSM26352 | 1012 | 81393 |
| 1134.214 | 189 | 1107.383 | 215 | 691.561 | 178 | 611.7571 | 177 | BRD4 | K562 | Bone Marr | GSM26352 | 5055 | 81395 |
| 1123.941 | 298 | 1130.016 | 375 | 872.7823 | 302 | 817.8036 | 292 | BRD4 | K562 | Bone Marr | GSM26352 | 9655 | 81397 |
| 1190.215 | 192 | 1196.825 | 236 | 1055.985 | 261 | 1051.086 | 291 | ESR1 | MCF-7 | Breast | GSM27476 | 4425 | 81409 |
| 493.7785 | 147 | 629.4678 | 185 | 276.94 | 165 | 225.3956 | 162 | RELA | None | Blood | GSM26799 | 11949 | 81421 |
| 30.51719 | 21 | 36.97634 | 26 | 27.52327 | 32 | 92.41437 | 58 | RELA | SW480 | Colon | GSM27465 | 3686 | 81526 |
| 803.6946 | 428 | 821.7431 | 537 | 613.8032 | 509 | 582.0722 | 532 | TP53 | SW480 | Colon | GSM27465 | 38347 | 81528 |
| 815.0176 | 621 | 855.1382 | 805 | 586.252 | 728 | 561.5637 | 789 | TET2 | Jurkat | Blood | GSM24670 | 72727 | 81536 |
| 869.8812 | 554 | 914.3932 | 731 | 655.1685 | 654 | 600.7014 | 657 | TET2 | Jurkat | Blood | GSM24670 | 49705 | 81543 |
| 409.0415 | 127 | 657.3639 | 180 | 241.1932 | 144 | 305.6172 | 172 | BMI1 | C4-2 | Prostate | GSM25791 | 9637 | 81557 |
| 264.2633 | 236 | 318.0493 | 288 | 115.3202 | 287 | 141.224 | 334 | EP300 | None | None | GSM25502 | 47352 | 81576 |
| 333.0584 | 207 | 304.1975 | 225 | 194.282 | 261 | 132.7964 | 250 | EP300 | None | None | GSM25502 | 31442 | 81582 |
| 105.3091 | 80 | 164.8574 | 110 | 74.94477 | 112 | 118.888 | 144 | RAD21 | None | Umbilical V | GSM24868 | 14705 | 81584 |
| 260.6933 | 133 | 332.3351 | 167 | 167.5754 | 170 | 237.6241 | 211 | RAD21 | None | Umbilical V | GSM24868 | 17622 | 81585 |
| 173.3269 | 116 | 225.4948 | 147 | 134.1859 | 165 | 200.2583 | 208 | RAD21 | None | Umbilical V | GSM24868 | 19457 | 81586 |
| 318.7662 | 171 | 421.6807 | 218 | 199.0524 | 218 | 264.1592 | 263 | RAD21 | None | Umbilical V | GSM24868 | 23903 | 81587 |
| 201.2949 | 36 | 392.9448 | 59 | 132.9113 | 40 | 272.3011 | 63 | PSMD1 | None | None | GSM23338 | 1473 | 81596 |
| 1007.156 | 566 | 1007.917 | 684 | 779.7218 | 633 | 753.3581 | 684 | BRD4 | RT112 | Urinary Bla | GSM23594 | 29381 | 81604 |
| 891.345 | 486 | 879.0496 | 573 | 683.9709 | 556 | 670.1285 | 615 | BRD4 | PC-9 | Lung | GSM23594 | 32657 | 81607 |
| 976.4766 | 247 | 956.1414 | 285 | 768.0273 | 273 | 684.8109 | 276 | BRD4 | PC-9 | Lung | GSM23594 | 10855 | 81608 |
| 544.6804 | 124 | 558.0226 | 138 | 518.74 | 169 | 535.8969 | 183 | BRD4 | RT112 | Urinary Bla | GSM23594 | 6144 | 81609 |
| 898.9946 | 530 | 914.9296 | 660 | 693.6992 | 619 | 669.1045 | 664 | BRD4 | PC-9 | Lung | GSM23594 | 36333 | 81611 |
| 1039.289 | 371 | 1017.836 | 424 | 765.4508 | 345 | 752.6649 | 376 | BRD4 | N87 | Stomach | GSM23594 | 14615 | 81612 |
| 938.0237 | 444 | 946.7093 | 547 | 727.7693 | 499 | 709.6999 | 541 | BRD4 | PC-9 | Lung | GSM23594 | 24898 | 81613 |
| 1000.31 | 280 | 1008.594 | 348 | 781.0464 | 298 | 754.386 | 315 | BRD4 | PC-9 | Lung | GSM23594 | 12346 | 81614 |
| 1137.262 | 531 | 1185.668 | 706 | 857.8887 | 521 | 804.7779 | 516 | NCAPH2 | HEK293 | Embryonic | GSM25717 | 17421 | 81619 |
| 1140.754 | 511 | 1204.95 | 694 | 864.8277 | 497 | 816.4706 | 498 | NCAPH2 | HEK293 | Embryonic | GSM25717 | 16013 | 81621 |
| 1037.314 | 787 | 1094.87 | 1010 | 740.291 | 857 | 706.4439 | 918 | MAX | MKL-1 | None | GSM17118 | 57489 | 81646 |
| 506.0266 | 246 | 732.4816 | 341 | 324.8744 | 307 | 332.6757 | 336 | CTCF | ID00015 | None | GSM20407 | 28188 | 81660 |
| 509.4887 | 280 | 712.4751 | 396 | 341.3689 | 362 | 397.6397 | 416 | CTCF | GM12878 | None | GSM20407 | 36536 | 81662 |
| 801.8156 | 469 | 833.9415 | 610 | 601.9827 | 555 | 577.1867 | 596 | CTCF | GM17942 | None | GSM20407 | 45770 | 81664 |
| 222.5846 | 117 | 198.2189 | 125 | 184.2359 | 166 | 173.0796 | 175 | ATF3 | None | Skin | GSM21519 | 15926 | 81700 |
| 371.0192 | 158 | 433.3975 | 190 | 196.9207 | 181 | 226.8411 | 207 | MED1 | hMSC-TER1 | None | GSM28029 | 17007 | 81711 |
| 130.4931 | 107 | 96.96937 | 108 | 75.77617 | 139 | 158.1242 | 196 | FOSL1 | MG-63 | None | GSM26353 | 19311 | 81732 |
| 899.3655 | 203 | 942.6493 | 253 | 513.8573 | 215 | 435.9613 | 213 | FOS | MG-63 | None | GSM26353 | 10121 | 81733 |
| 90.1737 | 65 | 85.7403 | 72 | 69.68941 | 92 | 96.83696 | 113 | FOSL1 | 143B | None | GSM26353 | 10688 | 81736 |
| 991.7369 | 122 | 779.2085 | 114 | 1019.143 | 165 | 1113.519 | 212 | AR | LNCaP | Prostate | GSM23938 | 2563 | 81774 |
| 988.7388 | 221 | 946.112 | 241 | 697.9096 | 243 | 781.0777 | 278 | KLF5 | BICR31 | Head and r | GSM23566 | 10274 | 81775 |
| 510.365 | 201 | 571.0421 | 237 | 690.2366 | 355 | 747.4921 | 473 | ESR1 | MIN | Breast | GSM21251 | 21821 | 81802 |
| 871.0871 | 167 | 1015.591 | 213 | 491.0331 | 171 | 742.9175 | 225 | ESR1 | MIN | Breast | GSM21251 | 6713 | 81804 |
| 43.77801 | 41 | 91.64528 | 64 | 93.26872 | 89 | 223.0485 | 139 | TFAP2C | H9 | Embryo | GSM28176 | 8810 | 81808 |
| 67.13885 | 21 | 36.80087 | 17 | 44.80226 | 25 | 332.3932 | 76 | FOXA2 | BJ | Skin | GSM24306 | 1585 | 81822 |
| 4.27956 | 6 | 1.801022 | 5 | 4.861562 | 11 | 33.47235 | 26 | NOTCH3 | None | None | GSM27938 | 1999 | 81877 |
| 48.06044 | 14 | 34.65814 | 13 | 50.37891 | 21 | 117.2986 | 35 | NOTCH3 | None | None | GSM27938 | 1078 | 81879 |
| 801.3931 | 109 | 787.6801 | 117 | 699.7061 | 135 | 923.1203 | 167 | ERF | VCaP | Prostate | GSM26124 | 2631 | 81882 |
| 1053.933 | 151 | 864.2566 | 146 | 985.7084 | 196 | 1016.598 | 247 | ERF | VCaP | Prostate | GSM26124 | 4066 | 81883 |
| 945.4196 | 328 | 986.608 | 442 | 707.3601 | 346 | 628.2714 | 338 | HIRA | IMR90 | Lung | GSM19367 | 19376 | 81884 |
| 1015.809 | 375 | 1045.952 | 486 | 787.6912 | 398 | 722.8824 | 380 | SMARCC1 | None | epithelioid | GSM24096 | 16536 | 81895 |
| 994.2802 | 339 | 1009.791 | 426 | 769.428 | 358 | 724.4357 | 358 | SMARCA4 | None | epithelioid | GSM24096 | 14685 | 81896 |
| 947.9863 | 237 | 925.5365 | 272 | 531.2482 | 232 | 574.8962 | 258 | MKL2 | A673 | Muscle | GSM24366 | 10772 | 81943 |
| 879.9087 | 279 | 867.0904 | 325 | 519.4127 | 280 | 565.8922 | 313 | MKL2 | A673 | Muscle | GSM24366 | 15112 | 81944 |
| 353.4653 | 59 | 467.9972 | 76 | 221.3443 | 62 | 241.7956 | 69 | MKL1 | A673 | Muscle | GSM24366 | 1904 | 81946 |
| 182.8007 | 202 | 149.5236 | 214 | 69.8299 | 236 | 89.24103 | 278 | HCFC1R1 | None | Cartilage | GSM26776 | 38251 | 81963 |

| | | | | | | | | | | | | | |
|---|---|---|---|---|---|---|---|---|---|---|---|---|---|
| 788.0289 | 625 | 814.4508 | 792 | 660.6017 | 918 | 631.9928 | 994 | CTCF | MCF-7 | Breast | GSM27477 | 87089 | 82007 |
| 347.4388 | 85 | 391.6581 | 100 | 278.0366 | 108 | 338.1294 | 127 | RXRA | JMSU-1 | None | GSM28773 | 4939 | 82015 |
| 933.4945 | 284 | 966.1207 | 379 | 724.818 | 317 | 712.5575 | 350 | RXRA | JMSU-1 | None | GSM28773 | 16447 | 82016 |
| 34.94956 | 43 | 55.21171 | 59 | 28.81459 | 67 | 46.89383 | 86 | HNF1A | None | Epididymis | GSM19824 | 11106 | 82017 |
| 23.55636 | 44 | 27.8914 | 54 | 24.61058 | 76 | 28.45665 | 87 | HNF1A | None | Epididymis | GSM19824 | 13978 | 82019 |
| 204.8269 | 71 | 210.421 | 80 | 204.3961 | 104 | 283.4168 | 130 | RXRA | JMSU-1 | None | GSM28773 | 6075 | 82022 |
| 315.4713 | 105 | 323.2001 | 118 | 285.454 | 146 | 271.6942 | 153 | RXRA | JMSU-1 | None | GSM28773 | 8556 | 82023 |
| 3.498564 | 10 | 1.881472 | 10 | 14.40456 | 28 | 36.19667 | 43 | DNMT3B | TKO | None | GSM26425 | 4085 | 82035 |
| 1171.022 | 244 | 1192.137 | 315 | 906.3267 | 227 | 869.111 | 233 | TET1 | HUES8 hES | None | GSM26425 | 5998 | 82036 |
| 4.83309 | 6 | 2.127539 | 5 | 0 | 3 | 44.46717 | 28 | DNMT3B | HUES8 hES | None | GSM26425 | 1720 | 82037 |
| 35.00524 | 136 | 43.53227 | 169 | 107.7767 | 320 | 196.0166 | 417 | GATA3 | H9 | Embryo | GSM28176 | 20272 | 82055 |
| 709.3555 | 452 | 722.9911 | 567 | 534.3416 | 574 | 518.6928 | 629 | NR3C1 | None | Muscle | GSM25199 | 61582 | 82060 |
| 643.0834 | 388 | 658.7209 | 491 | 419.0166 | 510 | 440.6376 | 566 | NR3C1 | None | Muscle | GSM25199 | 60880 | 82061 |
| 708.7971 | 440 | 717.1314 | 546 | 531.0933 | 552 | 502.135 | 587 | NR3C1 | None | Muscle | GSM25199 | 58923 | 82062 |
| 647.1269 | 427 | 647.599 | 521 | 485.0203 | 574 | 486.5563 | 640 | NR3C1 | None | Muscle | GSM25199 | 69077 | 82063 |
| 492.8279 | 339 | 634.4685 | 444 | 369.9772 | 476 | 402.1341 | 534 | NR3C1 | None | Muscle | GSM25199 | 59133 | 82064 |
| 548.5621 | 366 | 629.8211 | 462 | 434.0093 | 522 | 414.3814 | 561 | NR3C1 | None | Muscle | GSM25199 | 63230 | 82065 |
| 1169.327 | 399 | 1142.954 | 456 | 921.4013 | 414 | 889.5807 | 430 | ESR1 | None | Breast | GSM27970 | 11310 | 82072 |
| 1134.916 | 252 | 1124.722 | 300 | 952.6866 | 304 | 960.995 | 350 | ESR1 | None | Breast | GSM27970 | 7320 | 82073 |
| 926.3083 | 285 | 863.019 | 295 | 772.5606 | 371 | 817.2069 | 476 | FOXA1 | None | Breast | GSM27970 | 16523 | 82074 |
| 901.601 | 89 | 1198.532 | 115 | 618.0987 | 92 | 801.1364 | 113 | ESR1 | None | Breast | GSM27970 | 1235 | 82075 |
| 837.6216 | 371 | 758.6789 | 370 | 668.2663 | 468 | 697.8665 | 575 | FOXA1 | None | Breast | GSM27970 | 30960 | 82081 |
| 993.0508 | 160 | 1095.9 | 210 | 571.7149 | 160 | 571.5398 | 170 | BRD4 | C4-2 | Prostate | GSM23504 | 4752 | 82112 |
| 909.9319 | 122 | 952.2511 | 136 | 767.3883 | 148 | 1012.017 | 183 | BRD4 | C4-2 | Prostate | GSM23504 | 2896 | 82113 |
| 250.9477 | 65 | 379.4239 | 90 | 246.5007 | 92 | 439.493 | 132 | ESR1 | T47D | Breast | GSM21251 | 3998 | 82124 |
| 177.0884 | 58 | 326.4136 | 89 | 128.8477 | 73 | 161.827 | 87 | KDM3B | HCT-116 | HCT116 | GSM18471 | 4735 | 82126 |
| 628.2909 | 175 | 728.4523 | 209 | 385.5331 | 200 | 403.4356 | 219 | KDM3B | HCT-116 | HCT116 | GSM18471 | 12170 | 82127 |
| 212.4634 | 51 | 322.6709 | 71 | 221.9767 | 74 | 268.9956 | 87 | ESR1 | T47D | Breast | GSM21251 | 2844 | 82132 |
| 1182 | 439 | 1218.886 | 568 | 903.1644 | 407 | 865.9121 | 420 | GLI2 | THP-1 | None | GSM27790 | 11302 | 82146 |
| 1184.094 | 753 | 1261.399 | 1001 | 855.5404 | 749 | 818.7742 | 793 | MAPK1 | HeLa | Cervix | GSM22575 | 31284 | 82186 |
| 1118.595 | 345 | 1104.136 | 406 | 843.8587 | 327 | 801.6838 | 331 | MYCN | NGP | None | GSM24828 | 11537 | 82208 |
| 163.5924 | 93 | 116.026 | 90 | 204.297 | 155 | 347.8651 | 210 | FOXA2 | BJ | Skin | GSM29775 | 12244 | 82250 |
| 957.5751 | 132 | 1182.331 | 177 | 498.3043 | 120 | 503.9211 | 128 | BRD4 | MutuI | None | GSM26115 | 2723 | 82316 |
| 767.0951 | 97 | 1070.209 | 129 | 388.7875 | 87 | 337.0873 | 85 | BRD4 | MutuI | None | GSM26115 | 2031 | 82317 |
| 1178.36 | 227 | 1175.352 | 269 | 908.9395 | 210 | 751.4383 | 201 | BRD4 | MutuI | None | GSM26115 | 5063 | 82318 |
| 1244.509 | 135 | 1282.782 | 166 | 1096.803 | 164 | 1102.236 | 182 | ESR1 | MCF-7 | Breast | GSM27476 | 2203 | 82325 |
| 766.5761 | 177 | 857.028 | 207 | 844.6953 | 287 | 875.4541 | 352 | ESR1 | MCF-7 | Breast | GSM27476 | 9870 | 82326 |
| 1179.584 | 237 | 1180.483 | 290 | 1031.736 | 313 | 1029.846 | 355 | ESR1 | MCF-7 | Breast | GSM27476 | 6010 | 82327 |
| 939.9862 | 266 | 937.8239 | 320 | 834.8255 | 394 | 863.3387 | 487 | ESR1 | MCF-7 | Breast | GSM27476 | 15059 | 82328 |
| 824.1411 | 188 | 917.7916 | 225 | 855.772 | 308 | 877.664 | 370 | ESR1 | MCF-7 | Breast | GSM27476 | 10436 | 82330 |
| 529.4332 | 63 | 554.2865 | 70 | 498.4443 | 80 | 665.3786 | 100 | ESR1 | MCF-7 | Breast | GSM27476 | 1157 | 82331 |
| 1148.528 | 432 | 1182.589 | 559 | 889.9936 | 419 | 834.9248 | 411 | KLF15 | None | Muscle | GSM25199 | 11348 | 82343 |
| 1180.93 | 405 | 1194.931 | 507 | 911.046 | 379 | 852.6564 | 369 | KLF15 | None | Muscle | GSM25199 | 9436 | 82344 |
| 1128.664 | 204 | 1176.007 | 278 | 661.257 | 179 | 686.5321 | 194 | ZFX | PrEC | Prostate | GSM29095 | 5002 | 82348 |
| 659.8717 | 128 | 956.4761 | 173 | 394.8491 | 135 | 676.7284 | 190 | BCL11A | V5 | None | GSM27715 | 5182 | 82369 |
| 136.2163 | 27 | 226.4975 | 40 | 63.33274 | 25 | 55.72461 | 25 | STAT3 | TMD8 | None | GSM28554 | 1184 | 82372 |
| 109.7034 | 70 | 100.3712 | 76 | 82.93674 | 97 | 97.85545 | 112 | BACH2 | None | None | GSM27380 | 10525 | 82378 |
| 928.6424 | 388 | 911.7627 | 448 | 781.6833 | 509 | 777.271 | 575 | ESR1 | LTED | Breast | GSM26708 | 22762 | 82380 |
| 1027.721 | 716 | 1012.557 | 837 | 829.4419 | 897 | 823.5105 | 1015 | ESR1 | LTED | Breast | GSM26708 | 43892 | 82381 |
| 102.9603 | 23 | 234.8053 | 41 | 90.07503 | 30 | 205.2632 | 50 | HDAC2 | None | None | GSM17066 | 1125 | 82391 |
| 328.3958 | 57 | 545.4029 | 84 | 234.0187 | 64 | 191.3793 | 61 | SMARCA4 | HepG2 | Liver | GSM27409 | 1808 | 82422 |
| 928.9671 | 145 | 1085.568 | 176 | 434.6234 | 129 | 374.5679 | 127 | SMARCA4 | HepG2 | Liver | GSM27409 | 4073 | 82423 |
| 177.3736 | 68 | 233.6898 | 87 | 456.8702 | 159 | 602.9079 | 195 | ESR1 | MCF-7 | Breast | GSM26489 | 6446 | 82425 |
| 101.9912 | 37 | 148.0975 | 50 | 236.1791 | 83 | 424.3844 | 120 | ESR1 | MCF-7 | Breast | GSM26489 | 3401 | 82427 |
| 903.5251 | 111 | 1174.627 | 141 | 521.7443 | 106 | 538.4695 | 114 | NELFA | HeLa | Adherent I | GSM24651 | 2077 | 82509 |
| 880.5702 | 123 | 1181.179 | 169 | 535.3653 | 123 | 578.9779 | 136 | NELFA | HeLa | Adherent I | GSM24651 | 2775 | 82510 |
| 1124.704 | 150 | 1203.286 | 200 | 657.3936 | 146 | 637.6242 | 152 | NELFA | HeLa | Adherent I | GSM24651 | 3157 | 82512 |
| 1208.797 | 173 | 1230.418 | 220 | 783.5229 | 162 | 764.472 | 169 | SUPT5H | HeLa | Adherent I | GSM24651 | 3250 | 82513 |
| 275.1221 | 42 | 835.4062 | 91 | 232.2401 | 51 | 402.4472 | 74 | SUPT5H | HeLa | Adherent I | GSM24651 | 1097 | 82516 |
| 1240.694 | 229 | 1147.671 | 214 | 1017.465 | 242 | 1086.043 | 334 | AR | LNCaP | Prostate | GSM23938 | 4647 | 82536 |
| 543.2498 | 148 | 571.5746 | 168 | 349.8895 | 172 | 702.6152 | 257 | AR | LNCaP | Prostate | GSM23938 | 9895 | 82537 |
| 64.24673 | 46 | 61.34968 | 51 | 27.04553 | 52 | 23.27382 | 54 | TP53 | HCT-116 | HCT116 | GSM22962 | 7352 | 82544 |
| 94.88964 | 22 | 80.07622 | 22 | 32.41593 | 18 | 54.74947 | 25 | TP53 | HCT-116 | HCT116 | GSM22962 | 1199 | 82545 |
| 85.39075 | 75 | 70.48855 | 79 | 31.69145 | 85 | 41.22567 | 101 | TP53 | Osteosarcc | None | GSM22962 | 14111 | 82549 |
| 941.3751 | 527 | 976.8725 | 680 | 662.3512 | 525 | 619.4332 | 535 | ZBTB7A | HUDEP-2 | None | GSM27715 | 35954 | 82591 |
| 1002.057 | 523 | 1026.369 | 661 | 732.4556 | 527 | 689.6155 | 539 | ZBTB7A | HUDEP-2 | None | GSM27715 | 27228 | 82592 |
| 32.60857 | 13 | 89.05774 | 25 | 37.46456 | 21 | 67.22404 | 30 | ZBTB7A | K562 | Bone Marr | GSM27715 | 1413 | 82593 |
| 995.1839 | 488 | 989.0993 | 587 | 724.4908 | 482 | 685.6259 | 496 | ZBTB7A | HUDEP-2 | None | GSM27715 | 25158 | 82594 |
| 1061.141 | 182 | 1108.191 | 254 | 680.6754 | 192 | 635.4655 | 197 | CHD1 | None | Bone Marr | GSM23599 | 5823 | 82606 |
| 1049.746 | 181 | 1097.37 | 254 | 544.5616 | 173 | 480.8251 | 173 | CHD1 | None | Bone Marr | GSM23599 | 5993 | 82607 |
| 439.0137 | 513 | 569.6259 | 701 | 218.9806 | 703 | 314.1555 | 847 | SPI1 | THP-1 | Blood | GSM23599 | 150206 | 82608 |
| 423.0025 | 487 | 559.1402 | 656 | 233.0107 | 680 | 307.6279 | 804 | SPI1 | THP-1 | Blood | GSM23599 | 139015 | 82609 |

| | | | | | | | | | | | | | |
|---|---|---|---|---|---|---|---|---|---|---|---|---|---|
| 384.7591 | 456 | 532.727 | 595 | 208.1092 | 635 | 259.4825 | 740 | SPI1 | THP-1 | Blood | GSM23599 | 130866 | 82610 |
| 348.6027 | 439 | 536.05 | 594 | 225.267 | 643 | 250.1134 | 728 | SPI1 | THP-1 | Blood | GSM23599 | 129615 | 82611 |
| 175.0764 | 280 | 265.7579 | 369 | 123.5024 | 423 | 113.7835 | 457 | SPI1 | THP-1 | Blood | GSM23599 | 83727 | 82612 |
| 152.6352 | 233 | 209.8997 | 298 | 116.2816 | 357 | 103.6515 | 381 | SPI1 | THP-1 | Blood | GSM23599 | 66651 | 82613 |
| 58.18739 | 18 | 81.59901 | 24 | 45.26051 | 23 | 81.25858 | 33 | PRDM1 | U266B1 | None | GSM27354 | 1329 | 82633 |
| 189.082 | 44 | 310.2276 | 64 | 287.6218 | 77 | 439.1124 | 103 | NR0B1 | H460 cells | None | GSM23839 | 2157 | 82635 |
| 874.7614 | 282 | 853.5126 | 321 | 692.2351 | 327 | 695.4617 | 377 | NFIC | HepG2 | Liver | GSM29026 | 17145 | 82639 |
| 1110.107 | 261 | 1117.675 | 326 | 832.2661 | 239 | 819.3784 | 254 | SATB1 | HepG2 | Liver | GSM29026 | 7237 | 82642 |
| 50.7817 | 50 | 28.96688 | 46 | 11.82525 | 50 | 13.09222 | 57 | FOXA2 | BJ | Skin | GSM24014 | 9843 | 82643 |
| 376.531 | 362 | 313.655 | 388 | 218.1763 | 486 | 216.6702 | 533 | FOXA2 | BJ | Skin | GSM24014 | 78037 | 82644 |
| 851.9292 | 492 | 854.5273 | 600 | 624.4638 | 555 | 597.9022 | 590 | T | None | Blood | GSM26799 | 44419 | 82662 |
| 14.22674 | 19 | 14.37538 | 22 | 6.681219 | 25 | 31.46155 | 46 | GRIP1 | THP-1 | Blood | GSM26617 | 5140 | 82702 |
| 1217.981 | 332 | 1254.291 | 429 | 952.9352 | 307 | 903.9764 | 307 | OGT | None | Colon | GSM24599 | 6876 | 82732 |
| 693.5815 | 655 | 718.3527 | 825 | 507.365 | 858 | 482.2239 | 932 | SPI1 | None | Blood | GSM25928 | 120284 | 82735 |
| 521.3201 | 233 | 642.5143 | 287 | 199.5034 | 240 | 240.4832 | 279 | EP300 | None | None | GSM25502 | 27295 | 82759 |
| 137.3155 | 111 | 187.1627 | 143 | 98.85691 | 155 | 134.5384 | 188 | MED1 | hMSC-TERT | None | GSM28029 | 19879 | 82793 |
| 73.11931 | 38 | 66.98607 | 41 | 33.16421 | 42 | 51.14019 | 54 | MED1 | hMSC-TERT | None | GSM28029 | 4911 | 82794 |
| 995.6932 | 417 | 961.6889 | 462 | 758.2894 | 430 | 727.2738 | 449 | BRD4 | N87 | Stomach | GSM23594 | 18719 | 82804 |
| 775.9973 | 246 | 847.4847 | 328 | 459.0466 | 281 | 327.3194 | 263 | CDK9 | MOLT-4 | None | GSM20909 | 19139 | 82835 |
| 822.3843 | 269 | 836.2277 | 339 | 501.7106 | 302 | 310.2174 | 267 | CDK9 | MOLT-4 | None | GSM20909 | 19771 | 82838 |
| 283.9502 | 46 | 311.8808 | 53 | 126.1951 | 40 | 59.69466 | 29 | BRD4 | MOLT-4 | None | GSM20909 | 1449 | 82839 |
| 1149.944 | 868 | 1144.621 | 1028 | 847.1829 | 975 | 862.3105 | 1143 | ESR1 | None | Breast | GSM27970 | 50990 | 82844 |
| 1182.067 | 242 | 1188.445 | 300 | 1002.659 | 290 | 991.9833 | 321 | ESR1 | None | Breast | GSM27970 | 5962 | 82845 |
| 1140.298 | 407 | 1151.944 | 507 | 883.2792 | 410 | 871.6483 | 450 | ESR1 | None | Breast | GSM27970 | 12618 | 82847 |
| 1239.178 | 314 | 1259.296 | 406 | 1002.759 | 326 | 989.7534 | 357 | ESR1 | None | Breast | GSM27970 | 6628 | 82848 |
| 1107.518 | 143 | 1178.285 | 176 | 890.9519 | 169 | 1025.824 | 208 | ESR1 | None | Breast | GSM27970 | 3331 | 82849 |
| 1238.745 | 349 | 1248.801 | 437 | 986.5508 | 351 | 963.2611 | 373 | ESR1 | None | Breast | GSM27970 | 7556 | 82850 |
| 702.8697 | 142 | 579.0351 | 140 | 596.1797 | 182 | 763.1472 | 220 | ESR1 | None | Breast | GSM27970 | 6529 | 82851 |
| 1129.444 | 534 | 1118.548 | 627 | 936.2298 | 658 | 912.8842 | 711 | ESR1 | None | Breast | GSM27970 | 20343 | 82852 |
| 451.3135 | 86 | 467.2615 | 96 | 289.2659 | 94 | 442.5279 | 125 | ESR1 | None | Breast | GSM27970 | 3506 | 82853 |
| 948.4448 | 264 | 950.3998 | 325 | 826.9947 | 378 | 830.7333 | 436 | ESR1 | None | Breast | GSM27970 | 14567 | 82854 |
| 999.1448 | 613 | 976.4637 | 714 | 793.788 | 765 | 765.7705 | 824 | FOXA2 | A549 | Lung | GSM24014 | 42454 | 82934 |
| 925.3938 | 559 | 905.2758 | 651 | 733.7796 | 708 | 734.7467 | 809 | FOXA2 | A549 | Lung | GSM24014 | 46247 | 82935 |
| 197.5234 | 180 | 126.5153 | 174 | 114.6762 | 239 | 111.3031 | 259 | FOXA2 | BJ | Skin | GSM24014 | 36383 | 82936 |
| 205.2232 | 192 | 147.829 | 194 | 126.7738 | 261 | 169.928 | 313 | FOXA2 | BJ | Skin | GSM24014 | 39656 | 82937 |
| 137.0523 | 155 | 96.87106 | 157 | 87.23697 | 218 | 95.11802 | 245 | FOXA2 | BJ | Skin | GSM24014 | 36349 | 82944 |
| 167.6091 | 175 | 111.5108 | 173 | 83.91154 | 227 | 91.24226 | 255 | FOXA2 | BJ | Skin | GSM24014 | 39230 | 82945 |
| 836.8968 | 146 | 994.0887 | 176 | 433.0319 | 141 | 426.8643 | 149 | ARID5B | Jurkat | Blood | GSM25705 | 5211 | 82963 |
| 732.2378 | 333 | 771.7101 | 448 | 511.833 | 422 | 550.1001 | 470 | CTCF | None | None | GSM22472 | 40059 | 82974 |
| 646.2374 | 323 | 733.2793 | 445 | 467.023 | 434 | 463.9322 | 470 | CTCF | JHU-06 | None | GSM22472 | 44899 | 82975 |
| 681.2574 | 412 | 725.2229 | 554 | 500.1096 | 534 | 518.6805 | 618 | CTCF | None | None | GSM22472 | 62211 | 82982 |
| 216.0328 | 66 | 220.8367 | 74 | 172.2049 | 86 | 171.9391 | 92 | PRMT5 | MDA-MB-2 | Breast | GSM19648 | 5001 | 83016 |
| 688.2444 | 253 | 770.2773 | 312 | 657.8532 | 369 | 629.0714 | 388 | RELA | Detroit 562 | None | GSM24198 | 23440 | 83024 |
| 31.47768 | 17 | 63.12867 | 27 | 31.12909 | 26 | 28.39339 | 27 | RELA | Detroit 562 | None | GSM24198 | 2292 | 83025 |
| 69.2381 | 48 | 87.30373 | 60 | 47.3568 | 64 | 39.28076 | 65 | RELA | Detroit 562 | None | GSM24198 | 7272 | 83026 |
| 240.6342 | 60 | 339.9411 | 80 | 168.5668 | 71 | 207.3006 | 84 | RELA | Detroit 562 | None | GSM24198 | 3407 | 83028 |
| 111.8353 | 86 | 165.5707 | 115 | 174.9802 | 160 | 331.3504 | 225 | RELA | Detroit 562 | None | GSM24198 | 14614 | 83029 |
| 56.30937 | 23 | 62.25214 | 27 | 86.94217 | 42 | 35.94698 | 30 | RELA | Detroit 562 | None | GSM24198 | 2252 | 83030 |
| 109.2613 | 29 | 163.2164 | 40 | 88.31864 | 37 | 68.98509 | 35 | RELA | Detroit 562 | None | GSM24198 | 1745 | 83031 |
| 357.682 | 62 | 435.2672 | 76 | 205.8554 | 63 | 280.8087 | 79 | ZNF483 | BG01V - AT | None | GSM25639 | 2208 | 83048 |
| 446.7678 | 62 | 781.3968 | 95 | 286.0907 | 64 | 164.9222 | 50 | DDX21 | HeLa | Cervix | GSM23713 | 1464 | 83099 |
| 1197.246 | 263 | 1200.465 | 320 | 912.0954 | 224 | 875.0958 | 230 | DDX21 | HeLa | Cervix | GSM23713 | 5507 | 83100 |
| 658.2633 | 94 | 826.1049 | 117 | 321.2051 | 84 | 314.1201 | 88 | TCOF1 | HeLa | Cervix | GSM23713 | 2330 | 83101 |
| 264.946 | 44 | 362.8586 | 58 | 131.4809 | 41 | 446.6753 | 86 | AR | LNCaP | Prostate | GSM23029 | 1498 | 83108 |
| 773.4127 | 389 | 807.288 | 514 | 606.7515 | 503 | 572.9321 | 528 | SMC3 | HeLa | None | GSM27477 | 43453 | 83136 |
| 224.593 | 112 | 437.6196 | 172 | 153.4519 | 144 | 169.4898 | 162 | SMC3 | HeLa | None | GSM27477 | 13298 | 83137 |
| 767.5772 | 351 | 784.2748 | 447 | 532.1527 | 418 | 571.5046 | 478 | SMC3 | HeLa | None | GSM27477 | 38676 | 83138 |
| 418.1042 | 183 | 679.4011 | 257 | 227.5938 | 212 | 236.7241 | 233 | SMC3 | HeLa | None | GSM27477 | 19752 | 83139 |
| 953.0224 | 466 | 858.8108 | 459 | 776.3417 | 596 | 857.2032 | 819 | AR | LNCaP | Prostate | GSM27169 | 30966 | 83143 |
| 1051.042 | 272 | 1047.95 | 330 | 784.3299 | 261 | 671.2092 | 258 | RNF2 | None | None | GSM28058 | 9852 | 83146 |
| 166.9587 | 58 | 159.4616 | 63 | 77.05984 | 60 | 135.9225 | 83 | KDM2B | None | None | GSM28058 | 5007 | 83147 |
| 1067.413 | 431 | 1104.72 | 561 | 810.7269 | 427 | 763.5518 | 426 | RYBP | None | None | GSM28058 | 15671 | 83150 |
| 407.7581 | 168 | 337.7242 | 172 | 647.1303 | 307 | 700.7543 | 361 | ESR1 | MCF-7 | Breast | GSM29704 | 17695 | 83180 |
| 257.1608 | 116 | 279.2999 | 135 | 450.9009 | 224 | 717.1368 | 298 | ESR1 | MCF-7 | Breast | GSM29704 | 13257 | 83182 |
| 340.3829 | 145 | 305.231 | 154 | 562.53 | 270 | 713.9829 | 334 | ESR1 | MCF-7 | Breast | GSM29704 | 15347 | 83183 |
| 346.0476 | 136 | 333.1728 | 149 | 583.4515 | 256 | 730.932 | 311 | ESR1 | MCF-7 | Breast | GSM29704 | 13523 | 83186 |
| 70.1836 | 43 | 59.25693 | 45 | 13.00935 | 36 | 50.14346 | 63 | EP300 | None | None | GSM25501 | 6334 | 83198 |
| 615.7518 | 404 | 665.6867 | 543 | 383.6734 | 539 | 417.2442 | 605 | CTCF | H1 | Embryo | GSM51837 | 75212 | 832 |
| 45.84865 | 24 | 16.42331 | 17 | 31.09279 | 31 | 68.24736 | 47 | SMARCC1 | TTC1240 | Malignant | GSM24096 | 3056 | 83222 |
| 756.4009 | 551 | 737.9608 | 644 | 554.3369 | 666 | 544.9507 | 744 | SMARCB1 | TTC1240 | Malignant | GSM24096 | 67445 | 83223 |
| 1035.989 | 338 | 1066.326 | 443 | 796.0925 | 351 | 729.5817 | 328 | SMARCC1 | TTC1240 | Malignant | GSM24096 | 14228 | 83224 |
| 1011.992 | 264 | 1013.486 | 323 | 780.5305 | 271 | 576.3623 | 250 | SMARCA4 | TTC1240 | Malignant | GSM24096 | 11045 | 83225 |

| | | | | | | | | | | | | | |
|---|---|---|---|---|---|---|---|---|---|---|---|---|---|
| 724.1674 | 583 | 763.771 | 759 | 618.7483 | 884 | 607.5359 | 988 | CTCF | MCF-7 | Breast | GSM28779 | 90527 | 83249 |
| 659.0817 | 461 | 712.8423 | 627 | 506.661 | 625 | 488.6766 | 681 | CTCF | MCF-7 | Breast | GSM28779 | 77580 | 83251 |
| 671.4429 | 383 | 681.5413 | 477 | 546.0928 | 553 | 560.9157 | 654 | CTCF | MCF-7 | Breast | GSM28779 | 59134 | 83252 |
| 654.1792 | 462 | 715.2457 | 640 | 497.7945 | 622 | 479.2451 | 679 | CTCF | MCF-7 | Breast | GSM28779 | 78997 | 83253 |
| 514.2452 | 123 | 525.5716 | 137 | 638.1029 | 194 | 833.3631 | 237 | GATA3 | T47D | Mammary | GSM26445 | 6776 | 83258 |
| 874.5261 | 476 | 824.8046 | 516 | 716.0469 | 627 | 721.476 | 729 | FOXA1 | T47D | Mammary | GSM26445 | 39575 | 83259 |
| 716.2585 | 268 | 740.5508 | 307 | 649.4329 | 393 | 674.7019 | 477 | GATA3 | T47D | Mammary | GSM26445 | 27119 | 83264 |
| 538.3281 | 182 | 547.6257 | 204 | 492.8703 | 255 | 723.3665 | 344 | GATA3 | T47D | Mammary | GSM26445 | 15472 | 83265 |
| 551.791 | 211 | 468.2457 | 218 | 495.9885 | 297 | 697.4598 | 413 | GATA3 | T47D | Mammary | GSM26445 | 20639 | 83266 |
| 893.9976 | 487 | 819.6171 | 504 | 720.0795 | 629 | 739.1597 | 748 | FOXA1 | T47D | Mammary | GSM26445 | 39298 | 83268 |
| 501.132 | 76 | 445.1402 | 77 | 617.6442 | 116 | 799.3986 | 142 | FOXA1 | T47D | Mammary | GSM26445 | 2173 | 83269 |
| 533.1255 | 76 | 405.4945 | 70 | 548.4395 | 104 | 716.3076 | 128 | FOXA1 | T47D | Mammary | GSM26445 | 1977 | 83270 |
| 255.3911 | 40 | 203.3445 | 38 | 160.4059 | 42 | 331.3663 | 67 | ESR1 | None | Breast | GSM27971 | 1255 | 83325 |
| 993.6666 | 339 | 949.6243 | 372 | 844.6858 | 451 | 862.2686 | 536 | ESR1 | None | Breast | GSM27971 | 16910 | 83326 |
| 1162.294 | 883 | 1170.677 | 1062 | 871.0459 | 1021 | 887.2 | 1200 | ESR1 | None | Breast | GSM27971 | 51000 | 83328 |
| 1050.129 | 519 | 1054.959 | 633 | 848.5722 | 621 | 828.3864 | 680 | ESR1 | None | Breast | GSM27971 | 24584 | 83331 |
| 512.4764 | 80 | 531.5405 | 89 | 336.8434 | 86 | 336.0365 | 91 | ESR1 | None | Breast | GSM27971 | 2422 | 83332 |
| -0.000465 | 37 | 0.035035 | 51 | 0.799827 | 104 | 1.216601 | 122 | PHOX2B | CLB-Ga | None | GSM26643 | 40894 | 83392 |
| 303.8444 | 86 | 196.982 | 76 | 489.8933 | 157 | 774.3752 | 212 | ESR1 | MCF-7 | Breast | GSM27477 | 5757 | 83427 |
| 781.0934 | 322 | 769.1364 | 381 | 708.182 | 516 | 738.5885 | 639 | ESR1 | MCF-7 | Breast | GSM27477 | 31038 | 83428 |
| 989.5236 | 273 | 937.4414 | 289 | 863.053 | 381 | 894.7027 | 468 | ESR1 | MCF-7 | Breast | GSM27477 | 12597 | 83429 |
| 785.6845 | 285 | 756.5389 | 321 | 721.6555 | 471 | 752.6979 | 583 | ESR1 | MCF-7 | Breast | GSM27477 | 26457 | 83430 |
| 765.2301 | 336 | 737.697 | 381 | 693.3318 | 542 | 719.2178 | 662 | ESR1 | MCF-7 | Breast | GSM27477 | 34791 | 83431 |
| 1037.932 | 417 | 1016.508 | 488 | 876.4422 | 547 | 884.0111 | 630 | ESR1 | MCF-7 | Breast | GSM27477 | 19283 | 83432 |
| 961.4684 | 241 | 948.0977 | 281 | 866.0596 | 364 | 881.4698 | 431 | ESR1 | MCF-7 | Breast | GSM27477 | 11764 | 83433 |
| 724.1847 | 147 | 873.5638 | 179 | 880.2117 | 227 | 908.6188 | 280 | ESR1 | MCF-7 | Breast | GSM27477 | 6651 | 83434 |
| 956.325 | 332 | 976.5579 | 427 | 702.0135 | 340 | 697.3402 | 381 | GTF3C5 | IMR-5 | None | GSM20820 | 20362 | 83437 |
| 551.4645 | 100 | 609.4252 | 116 | 209.991 | 84 | 288.2099 | 105 | GTF3C5 | IMR-5 | None | GSM20820 | 4110 | 83438 |
| 33.48958 | 18 | 23.00078 | 17 | 31.67674 | 27 | 57.68447 | 38 | NR3C1 | MCF-10A | Breast | GSM27353 | 2423 | 83501 |
| 103.4201 | 99 | 205.596 | 149 | 137.8714 | 175 | 216.9375 | 225 | NR3C1 | MCF-10A | Breast | GSM27353 | 20013 | 83502 |
| 968.6035 | 181 | 979.4745 | 202 | 804.7667 | 228 | 885.7099 | 292 | ESR1 | None | Breast | GSM27971 | 7640 | 83530 |
| 137.1378 | 28 | 132.2927 | 30 | 55.89484 | 24 | 196.7924 | 50 | MORC2 | K562 | None | GSM25094 | 1146 | 83532 |
| 152.5913 | 31 | 102.4332 | 27 | 46.74161 | 23 | 210.0714 | 54 | MORC2 | K562 | None | GSM25094 | 1250 | 83534 |
| 1105.711 | 417 | 1153.471 | 561 | 853.3328 | 418 | 804.2132 | 416 | FOXA2 | BJ | Skin | GSM29775 | 13766 | 83588 |
| 14.40592 | 122 | 10.04831 | 135 | 11.51508 | 216 | 14.11529 | 249 | FOXA2 | BJ | Skin | GSM29775 | 66245 | 83589 |
| 364.9318 | 135 | 602.9532 | 193 | 285.3031 | 178 | 407.0061 | 224 | CTCF | SHEP-21N | Bone Marr | GSM25723 | 12482 | 83591 |
| 378.7079 | 181 | 591.0818 | 249 | 292.2459 | 245 | 391.7603 | 298 | CTCF | SHEP-21N | Bone Marr | GSM25723 | 21839 | 83592 |
| 427.4094 | 190 | 734.533 | 274 | 314.2353 | 250 | 426.8384 | 306 | CTCF | SHEP-21N | Bone Marr | GSM25723 | 21445 | 83595 |
| 62.09879 | 16 | 105.5975 | 24 | 46.65834 | 20 | 217.603 | 49 | RBPJ | None | None | GSM27461 | 1083 | 83600 |
| 49.22181 | 24 | 170.4053 | 51 | 106.9271 | 53 | 116.1852 | 59 | IRF1 | None | Blood | GSM26799 | 3171 | 83604 |
| 198.1973 | 32 | 199.4907 | 35 | 208.7377 | 45 | 294.8326 | 58 | NR3C1 | None | Breast | GSM27971 | 1022 | 83616 |
| 1022.854 | 180 | 830.396 | 175 | 665.9437 | 197 | 876.595 | 257 | FOXA1 | None | Breast | GSM27971 | 6679 | 83619 |
| 898.815 | 281 | 818.0775 | 276 | 726.2674 | 348 | 748.702 | 420 | FOXA1 | None | Breast | GSM27971 | 17846 | 83620 |
| 660.7298 | 104 | 480.6034 | 94 | 327.0614 | 96 | 521.6469 | 131 | FOXA1 | None | Breast | GSM27971 | 3155 | 83621 |
| 982.7966 | 291 | 927.7587 | 307 | 765.3508 | 318 | 815.4685 | 418 | FOXA1 | None | Breast | GSM27971 | 14471 | 83622 |
| 969.7578 | 433 | 888.0872 | 434 | 751.0502 | 488 | 772.542 | 591 | FOXA1 | None | Breast | GSM27971 | 25146 | 83623 |
| 953.7604 | 261 | 891.2457 | 265 | 779.8747 | 313 | 806.4603 | 388 | FOXA1 | None | Breast | GSM27971 | 12990 | 83624 |
| 353.81 | 57 | 517.4398 | 78 | 274.5948 | 67 | 340.8687 | 80 | NR3C1 | None | Breast | GSM27971 | 1727 | 83625 |
| 46.91543 | 87 | 58.02568 | 108 | 38.31913 | 139 | 30.65556 | 144 | MYOG | H9 | Embryo | GSM26295 | 27144 | 83689 |
| 237.6846 | 140 | 303.4726 | 175 | 85.29593 | 144 | 105.6843 | 169 | DNMT3A | MCF-7 | None | GSM27359 | 17847 | 83708 |
| 2.296287 | 6 | 11.16794 | 13 | 6.34035 | 15 | 3.845077 | 14 | SMARCA2 | SK-N-MC | Brain | GSM24721 | 2459 | 83717 |
| 826.7926 | 755 | 903.0721 | 1002 | 679.4926 | 1077 | 672.4923 | 1220 | CTCF | MCF-7 | Breast | GSM27477 | 108645 | 83736 |
| 690.3491 | 356 | 738.8915 | 487 | 550.8468 | 495 | 535.8434 | 543 | CTCF | MCF-7 | Breast | GSM27477 | 52294 | 83737 |
| 716.1172 | 398 | 757.6257 | 536 | 597.0849 | 585 | 578.099 | 636 | CTCF | MCF-7 | Breast | GSM27477 | 53846 | 83738 |
| 710.1267 | 467 | 756.5624 | 627 | 575.7198 | 665 | 556.2108 | 726 | CTCF | MCF-7 | Breast | GSM27477 | 68539 | 83739 |
| 354.9274 | 222 | 559.0082 | 304 | 276.4762 | 313 | 274.1704 | 339 | CTCF | MCF-7 | Breast | GSM27477 | 35407 | 83740 |
| 746.0758 | 567 | 813.5105 | 777 | 560.0559 | 731 | 545.4694 | 808 | CTCF | MCF-7 | Breast | GSM27477 | 82986 | 83741 |
| 695.5401 | 423 | 754.0042 | 585 | 522.1164 | 542 | 510.8834 | 602 | CTCF | MCF-7 | Breast | GSM27477 | 61797 | 83742 |
| 1181.384 | 436 | 1203.801 | 555 | 885.9526 | 391 | 866.1644 | 423 | FAM208A | K562 | None | GSM27898 | 10945 | 83747 |
| 1153.136 | 483 | 1177.263 | 613 | 862.6163 | 445 | 823.5705 | 454 | FAM208A | K562 | None | GSM27898 | 13346 | 83748 |
| 137.4674 | 27 | 124.9585 | 28 | 41.11471 | 20 | 35.73468 | 20 | GRIP1 | THP-1 | Blood | GSM26618 | 1199 | 83760 |
| 105.119 | 34 | 76.94071 | 32 | 220.7443 | 71 | 765.0751 | 149 | ESR1 | MCF-7 | Breast | GSM29153 | 2360 | 83810 |
| 38.97549 | 13 | 83.23618 | 22 | 81.48164 | 28 | 620.4789 | 94 | ESR1 | MCF-7 | Breast | GSM29153 | 1108 | 83812 |
| 93.60509 | 31 | 97.02145 | 35 | 237.0314 | 72 | 765.7237 | 146 | ESR1 | MCF-7 | Breast | GSM29153 | 2318 | 83813 |
| 277.2346 | 122 | 273.0399 | 135 | 521.8687 | 238 | 766.2433 | 331 | ESR1 | MCF-7 | Breast | GSM29153 | 11714 | 83816 |
| 173.0503 | 71 | 199.747 | 85 | 340.6611 | 145 | 721.3828 | 225 | ESR1 | MCF-7 | Breast | GSM29153 | 7156 | 83817 |
| 144.9237 | 62 | 148.0558 | 70 | 289.8236 | 128 | 678.9942 | 209 | ESR1 | MCF-7 | Breast | GSM29153 | 6552 | 83818 |
| 84.71487 | 20 | 32.64239 | 13 | 140.1676 | 37 | 762.6138 | 105 | ESR1 | MCF-7 | Breast | GSM29153 | 1030 | 83819 |
| 885.5537 | 338 | 887.7802 | 416 | 589.4562 | 343 | 584.0179 | 368 | RUNX1 | SKH1 | None | GSM23268 | 23759 | 83832 |
| 1014.466 | 564 | 1025.204 | 694 | 723.4341 | 562 | 706.7811 | 613 | RUNX1 | SKH1 | None | GSM23268 | 32974 | 83833 |
| 1231.702 | 834 | 1287.375 | 1058 | 933.99 | 903 | 937.1605 | 1038 | HNF4G | LNCaP | Prostate | GSM22771 | 30100 | 83855 |
| 876.6065 | 620 | 791.0409 | 636 | 712.7041 | 837 | 780.7459 | 1094 | FOXA1 | LNCaP | Prostate | GSM22771 | 62283 | 83865 |

| | | | | | | | | | | | | | |
|---|---|---|---|---|---|---|---|---|---|---|---|---|---|
| 47.59472 | 91 | 47.05121 | 105 | 53.8005 | 160 | 83.68213 | 200 | SIRT1 | HCT-116 | HCT116 | GSM24748 | 29012 | 83885 |
| 1.527326 | 68 | 6.363698 | 102 | 8.981128 | 171 | 6.947119 | 183 | MCM2 | HCT-116 | HCT116 | GSM24748 | 53495 | 83886 |
| 0.431016 | 4 | 0.150182 | 4 | 4.034539 | 14 | 6.499972 | 18 | SIRT1 | K562 | None | GSM24748 | 2789 | 83889 |
| 961.4206 | 377 | 977.1885 | 470 | 705.6173 | 367 | 629.1589 | 353 | BRD4 | MOLT-4 | None | GSM20909 | 18842 | 83903 |
| 96.42905 | 22 | 124.2256 | 28 | 90.18804 | 30 | 243.2887 | 55 | HOXA11 | K562 | None | GSM24105 | 1159 | 83917 |
| 140.1608 | 29 | 157.3817 | 34 | 65.3774 | 27 | 388.752 | 76 | CTCF | K562 | None | GSM24105 | 1301 | 83918 |
| 45.95236 | 15 | 110.1104 | 27 | 70.59985 | 27 | 411.4039 | 76 | BRD4 | J-Lat A72 | None | GSM26767 | 1119 | 83932 |
| 1000.328 | 172 | 1028.917 | 201 | 471.0657 | 156 | 509.4581 | 173 | PHIP | HEK293 | Embryonic | GSM27114 | 5418 | 83936 |
| 54.76905 | 16 | 131.5656 | 29 | 60.69372 | 24 | 82.19215 | 30 | PHIP | HEK293 | Embryonic | GSM27114 | 1012 | 83937 |
| 1149.806 | 592 | 1129.391 | 684 | 915.4966 | 669 | 917.4641 | 768 | ESR1 | None | Breast | GSM27971 | 22623 | 83942 |
| 82.23068 | 20 | 31.30765 | 13 | 153.1507 | 40 | 766.8472 | 108 | ESR1 | MCF-7 | Breast | GSM29153 | 1163 | 84007 |
| 104.4326 | 22 | 81.7814 | 21 | 50.0575 | 21 | 183.7204 | 45 | EP300 | BICR31 | Head and r | GSM23566 | 1088 | 84010 |
| 164.1055 | 28 | 254.6343 | 40 | 156.778 | 38 | 271.7173 | 55 | SOX2 | None | None | GSM19739 | 1084 | 84035 |
| 7.795832 | 36 | 45.62286 | 73 | 44.79689 | 105 | 46.34391 | 116 | GATA1 | None | Blood | GSM72239 | 17199 | 8409 |
| 9.759851 | 34 | 41.89013 | 64 | 59.52989 | 104 | 56.02167 | 111 | GATA1 | None | Blood | GSM72239 | 14727 | 8410 |
| 1249.925 | 220 | 1238.793 | 255 | 771.6262 | 173 | 856.9521 | 194 | RPE | None | Retina | GSM28650 | 4163 | 84107 |
| 12.64911 | 41 | 51.63696 | 76 | 58.44956 | 114 | 44.47464 | 114 | GATA1 | None | Blood | GSM72239 | 17091 | 8411 |
| 1030.629 | 520 | 1024.605 | 622 | 755.0049 | 526 | 719.5943 | 546 | GATA2 | None | Blood | GSM72239 | 27341 | 8412 |
| 64.41224 | 39 | 77.77289 | 48 | 55.02905 | 57 | 58.09251 | 63 | SMAD1 | None | Blood | GSM72239 | 5955 | 8413 |
| 66.21557 | 42 | 81.02282 | 52 | 55.04605 | 61 | 78.34346 | 76 | SMAD1 | None | Blood | GSM72239 | 6650 | 8414 |
| 710.5951 | 321 | 746.2406 | 426 | 505.9562 | 424 | 484.1675 | 452 | SMAD1 | None | Blood | GSM72239 | 40942 | 8415 |
| 271.1699 | 157 | 253.4134 | 172 | 166.4503 | 201 | 201.5174 | 234 | SMAD1 | None | Blood | GSM72240 | 23395 | 8416 |
| 592.1862 | 165 | 895.7768 | 230 | 325.2876 | 180 | 387.3823 | 209 | TCF7L2 | None | Blood | GSM72240 | 11787 | 8417 |
| 757.1183 | 211 | 860.1989 | 263 | 379.8016 | 222 | 396.038 | 243 | TCF7L2 | None | Blood | GSM72240 | 15366 | 8418 |
| 758.4455 | 111 | 914.7822 | 135 | 373.317 | 101 | 548.2169 | 132 | MYCN | SHEP-21N | Bone Marr | GSM21135 | 3038 | 84181 |
| 1145.148 | 513 | 1050.288 | 509 | 956.9003 | 639 | 1004.806 | 809 | AR | LNCaP-abl | Prostate | GSM21225 | 18646 | 84184 |
| 716.2088 | 569 | 756.2228 | 741 | 607.9105 | 855 | 592.6809 | 949 | CTCF | MCF-7 | Breast | GSM28779 | 89334 | 84200 |
| 1214.553 | 207 | 1155.31 | 211 | 1091.027 | 287 | 1123.116 | 355 | HOXB13 | 22RV1 | Prostate | GSM25372 | 4369 | 84204 |
| 1254.157 | 184 | 1119.762 | 158 | 1114.834 | 239 | 1153.445 | 301 | HOXB13 | 22RV1 | Prostate | GSM25372 | 3309 | 84205 |
| 1215.113 | 198 | 1120.967 | 186 | 1013.037 | 222 | 1063.461 | 290 | FOXA1 | 22RV1 | Prostate | GSM25372 | 4083 | 84210 |
| 27.759 | 26 | 30.76307 | 31 | 41.00872 | 49 | 76.01271 | 68 | TCF4 | SH-SY5Y | None | GSM25458 | 5595 | 84213 |
| 742.698 | 118 | 795.7323 | 134 | 336.0105 | 105 | 298.5539 | 105 | AR | 22RV1 | Prostate | GSM22771 | 3857 | 84224 |
| 563.9742 | 134 | 684.0366 | 164 | 333.355 | 147 | 421.9187 | 176 | AR | 22RV1 | Prostate | GSM22771 | 7792 | 84225 |
| 1013.024 | 451 | 903.5636 | 424 | 799.414 | 518 | 858.7944 | 686 | FOXA1 | LNCaP | Prostate | GSM22771 | 23735 | 84230 |
| 288.945 | 81 | 395.7178 | 106 | 223.6147 | 103 | 310.0571 | 129 | HNF4G | LNCaP | Prostate | GSM22771 | 5571 | 84232 |
| 44.54362 | 62 | 113.317 | 102 | 119.2741 | 145 | 122.8324 | 159 | GATA1 | K562 | Bone Marr | GSM28771 | 16605 | 84241 |
| 57.65496 | 93 | 76.10126 | 118 | 95.53719 | 182 | 141.0005 | 226 | TAL1 | K562 | Bone Marr | GSM28771 | 26625 | 84242 |
| 206.2974 | 185 | 270.3944 | 234 | 229.2747 | 306 | 203.7532 | 320 | GATA1 | K562 | Bone Marr | GSM28771 | 38279 | 84244 |
| 25.06674 | 65 | 69.9436 | 106 | 68.51109 | 151 | 105.9901 | 190 | TAL1 | K562 | Bone Marr | GSM28771 | 23715 | 84246 |
| 33.19698 | 63 | 42.04475 | 79 | 22.18112 | 96 | 51.41945 | 134 | CEBPB | None | Blood | GSM25927 | 20186 | 84251 |
| 1.496281 | 4 | 0.27322 | 3 | 2.498876 | 9 | 8.226297 | 15 | CEBPB | None | Blood | GSM25927 | 2104 | 84252 |
| 9.723159 | 39 | 35.16896 | 68 | 61.99667 | 119 | 82.81744 | 143 | GATA1 | K562 | Bone Marr | GSM72241 | 17626 | 8428 |
| 1140.707 | 344 | 1154.46 | 425 | 885.7961 | 336 | 854.4325 | 348 | ESR1 | Ishikawa | Uterus | GSM26640 | 9903 | 84285 |
| 100.0472 | 27 | 107.477 | 31 | 87.41521 | 36 | 114.0847 | 44 | ESR1 | Ishikawa | Uterus | GSM26640 | 1690 | 84286 |
| 4.012044 | 30 | 23.87355 | 58 | 69.34862 | 120 | 79.61929 | 137 | GATA1 | K562 | Bone Marr | GSM72241 | 16848 | 8429 |
| 153.3469 | 261 | 127.0334 | 284 | 84.80929 | 367 | 81.64884 | 402 | KDM4A | breast tripl | Breast | GSM24984 | 73380 | 84293 |
| 38.59386 | 60 | 85.79733 | 93 | 88.49569 | 133 | 134.2546 | 168 | GATA2 | K562 | Bone Marr | GSM72241 | 17442 | 8430 |
| 26.35764 | 44 | 72.03582 | 74 | 100.2043 | 118 | 98.32718 | 127 | GATA2 | K562 | Bone Marr | GSM72241 | 13321 | 8431 |
| 101.8161 | 54 | 176.8908 | 79 | 122.7513 | 90 | 103.0539 | 90 | SMAD1 | K562 | Bone Marr | GSM72241 | 7348 | 8432 |
| 51.61616 | 15 | 98.80111 | 24 | 30.9085 | 17 | 93.71811 | 32 | SMAD1 | K562 | Bone Marr | GSM72241 | 1163 | 8433 |
| 876.2657 | 556 | 762.8313 | 528 | 705.0252 | 728 | 761.8243 | 937 | FOXA1 | LNCaP | Prostate | GSM24808 | 51338 | 84338 |
| 863.5228 | 565 | 748.8267 | 540 | 675.1889 | 721 | 741.8584 | 947 | FOXA1 | LNCaP | Prostate | GSM24808 | 55528 | 84339 |
| 23.22901 | 18 | 125.7157 | 46 | 94.14797 | 53 | 117.3965 | 63 | TCF7L2 | K562 | Bone Marr | GSM72241 | 3476 | 8434 |
| 482.4001 | 81 | 506.4485 | 91 | 214.8074 | 72 | 551.236 | 127 | SMARCA4 | LNCaP | Prostate | GSM24808 | 2867 | 84341 |
| 941.6243 | 412 | 823.2687 | 378 | 802.5035 | 574 | 842.8617 | 721 | HOXB13 | LNCaP | Prostate | GSM24808 | 26681 | 84342 |
| 837.8821 | 588 | 718.4658 | 556 | 674.1313 | 788 | 720.5401 | 992 | FOXA1 | LNCaP | Prostate | GSM24808 | 63918 | 84344 |
| 873.4849 | 82 | 651.7222 | 73 | 494.006 | 75 | 528.0487 | 82 | FOXA1 | LNCaP | Prostate | GSM24808 | 1050 | 84345 |
| 999.4876 | 339 | 884.8047 | 303 | 852.8239 | 453 | 878.0856 | 554 | HOXB13 | LNCaP | Prostate | GSM24808 | 16799 | 84346 |
| 1062.095 | 311 | 963.8571 | 296 | 894.0452 | 397 | 937.1478 | 505 | HOXB13 | LNCaP | Prostate | GSM24808 | 12445 | 84347 |
| 47.34376 | 36 | 99.57894 | 57 | 80.85537 | 71 | 92.45956 | 81 | TCF7L2 | K562 | Bone Marr | GSM72241 | 6558 | 8435 |
| 745.6646 | 651 | 814.9644 | 876 | 577.9126 | 875 | 540.4894 | 929 | RUNX1 | None | Kidney | GSM24670 | 99785 | 84398 |
| 520.2266 | 385 | 599.3932 | 478 | 330.6268 | 524 | 335.5246 | 576 | CEBPA | U937 | Pleura | GSM72242 | 77521 | 8444 |
| 940.3598 | 115 | 909.5632 | 122 | 690.3979 | 128 | 1048.361 | 172 | FOXA1 | VCaP | Prostate | GSM25372 | 2434 | 84447 |
| 917.874 | 935 | 826.1103 | 1008 | 758.4702 | 1343 | 772.7878 | 1567 | FOXA1 | VCaP | Prostate | GSM25372 | 138653 | 84448 |
| 884.3624 | 837 | 804.2586 | 899 | 738.127 | 1202 | 765.6383 | 1432 | FOXA1 | VCaP | Prostate | GSM25372 | 113051 | 84450 |
| 882.3936 | 853 | 806.8889 | 925 | 728.7181 | 1216 | 755.0117 | 1448 | FOXA1 | VCaP | Prostate | GSM25372 | 118796 | 84451 |
| 892.8552 | 865 | 805.3088 | 925 | 732.7805 | 1226 | 766.2744 | 1469 | FOXA1 | VCaP | Prostate | GSM25372 | 119019 | 84452 |
| 1139.544 | 461 | 1171.868 | 595 | 864.602 | 436 | 811.6377 | 432 | SUPT5H | MOLT-4 | None | GSM23677 | 13170 | 84456 |
| 116.4518 | 158 | 191.7551 | 217 | 93.89512 | 245 | 148.4496 | 308 | CEBPA | U937 | Pleura | GSM72242 | 42981 | 8446 |
| 822.7932 | 112 | 1076.439 | 143 | 448.0632 | 106 | 735.1505 | 148 | MYCN | SHEP-21N | Bone Marr | GSM21135 | 2652 | 84465 |
| 1126.439 | 444 | 1150.767 | 569 | 830.9527 | 405 | 788.4066 | 411 | BRD4 | BE2-C | None | GSM21135 | 14361 | 84467 |
| 569.3382 | 330 | 676.0273 | 435 | 411.3451 | 452 | 404.4289 | 489 | SMAD1 | U937 | Pleura | GSM72242 | 52609 | 8447 |

| | | | | | | | | | | | | | |
|---|---|---|---|---|---|---|---|---|---|---|---|---|---|
| 470.5113 | 128 | 460.3611 | 140 | 201.862 | 124 | 233.1736 | 142 | TCF7L2 | U937 | Pleura | GSM72242 | 8817 | 8448 |
| 94.52342 | 88 | 130.1471 | 114 | 122.5203 | 155 | 140.7228 | 177 | GATA1 | K562 | Bone Marr | GSM28771 | 18134 | 84533 |
| 12.68105 | 19 | 21.69078 | 27 | 49.60942 | 53 | 51.01108 | 58 | GATA1 | K562 | Bone Marr | GSM28771 | 5602 | 84534 |
| 18.37009 | 37 | 27.12051 | 49 | 40.74471 | 81 | 58.63138 | 100 | TAL1 | K562 | Bone Marr | GSM28771 | 11832 | 84535 |
| 14.12668 | 40 | 21.19164 | 53 | 33.91981 | 90 | 72.61721 | 126 | TAL1 | K562 | Bone Marr | GSM28771 | 15280 | 84536 |
| 3.050284 | 20 | 37.28438 | 51 | 31.07981 | 69 | 72.72033 | 101 | CEBPA | K562 | Bone Marr | GSM78836 | 11056 | 8455 |
| 1326.463 | 131 | 1304.797 | 147 | 582.88 | 104 | 505.9823 | 101 | EZH2 | KG1 | Bone Marr | GSM29473 | 1795 | 84554 |
| 1304.481 | 131 | 1312.947 | 155 | 598.8209 | 107 | 546.8419 | 107 | EZH2 | KG1 | Bone Marr | GSM29473 | 1849 | 84555 |
| 211.941 | 207 | 359.3396 | 289 | 276.4392 | 364 | 320.5326 | 417 | CEBPA | K562 | Bone Marr | GSM78836 | 46178 | 8456 |
| 64.63268 | 18 | 77.4938 | 22 | 65.63664 | 26 | 589.1062 | 94 | ESR1 | MCF-7 | Breast | GSM29153 | 1171 | 84569 |
| 1300.17 | 829 | 1369.831 | 1062 | 906.5603 | 747 | 869.6218 | 794 | GATA1 | K562 | Bone Marr | GSM78836 | 26444 | 8458 |
| 144.6138 | 51 | 195.0388 | 66 | 66.89097 | 52 | 84.32949 | 62 | SMC1A | LCL | None | GSM24434 | 4025 | 84580 |
| 107.9651 | 33 | 146.5226 | 43 | 44.46807 | 31 | 80.94941 | 44 | SMC1A | LCL | None | GSM24434 | 2252 | 84581 |
| 976.3501 | 221 | 952.2652 | 255 | 464.1175 | 203 | 514.1922 | 228 | NIPBL | LCL | None | GSM24434 | 10500 | 84582 |
| 411.1609 | 126 | 473.0368 | 150 | 229.5093 | 139 | 204.4658 | 142 | NIPBL | LCL | None | GSM24434 | 9377 | 84583 |
| 982.7021 | 503 | 988.0731 | 617 | 702.1525 | 495 | 686.8882 | 539 | MED1 | LCL | None | GSM24434 | 29078 | 84585 |
| 22.27439 | 19 | 33.7155 | 26 | 30.74057 | 35 | 78.13301 | 56 | GATA1 | K562 | Bone Marr | GSM78836 | 3966 | 8459 |
| 107.9002 | 33 | 184.477 | 49 | 97.68014 | 46 | 171.2045 | 65 | IRF1 | JHU-06 | None | GSM23944 | 2600 | 84604 |
| 44.76418 | 17 | 131.3065 | 34 | 70.57634 | 32 | 126.7761 | 46 | IRF1 | JHU-06 | None | GSM23944 | 1773 | 84605 |
| 194.9197 | 159 | 161.4804 | 168 | 151.9368 | 231 | 215.6891 | 284 | JUNB | JHU-06 | None | GSM23944 | 30716 | 84608 |
| 384.7195 | 290 | 460.0557 | 353 | 283.1155 | 411 | 406.594 | 506 | JUN | JHU-06 | None | GSM23944 | 55485 | 84610 |
| 935.1024 | 751 | 972.6152 | 949 | 677.378 | 879 | 656.8186 | 966 | JUN | JHU-06 | None | GSM23944 | 76806 | 84611 |
| 69.68059 | 28 | 63.97086 | 30 | 37.77351 | 32 | 94.15151 | 52 | NFE2L2 | JHU-06 | None | GSM23944 | 2971 | 84612 |
| 951.2714 | 475 | 903.9335 | 516 | 808.2743 | 646 | 841.2774 | 797 | FOXA1 | 22RV1 | Prostate | GSM25372 | 30258 | 84613 |
| 433.1776 | 188 | 555.0277 | 236 | 364.6732 | 261 | 319.8936 | 267 | STAT3 | MDA-MB-1 | None | GSM22780 | 20822 | 84616 |
| 650.0423 | 267 | 670.7771 | 303 | 451.1109 | 342 | 499.3558 | 385 | STAT3 | MDA-MB-1 | None | GSM22780 | 29314 | 84617 |
| 809.682 | 271 | 787.124 | 312 | 483.6087 | 311 | 458.7659 | 328 | STAT3 | MDA-MB-2 | None | GSM22780 | 23377 | 84618 |
| 734.64 | 227 | 642.7484 | 236 | 474.2832 | 270 | 499.4833 | 297 | STAT3 | MDA-MB-2 | None | GSM22780 | 17931 | 84619 |
| 1006.291 | 833 | 1023.603 | 1017 | 795.4708 | 1084 | 810.5919 | 1276 | STAT3 | HCC70 | Breast | GSM22780 | 75470 | 84620 |
| 899.9662 | 497 | 888.4419 | 584 | 771.5646 | 700 | 783.6691 | 824 | STAT3 | HCC70 | Breast | GSM22780 | 38313 | 84621 |
| 737.891 | 194 | 898.1285 | 255 | 648.9881 | 260 | 590.4675 | 266 | STAT3 | None | None | GSM22780 | 12428 | 84622 |
| 972.9343 | 321 | 972.2098 | 387 | 780.258 | 376 | 740.9831 | 382 | STAT3 | None | None | GSM22780 | 16737 | 84623 |
| 695.9606 | 520 | 690.5768 | 624 | 452.5399 | 600 | 467.9723 | 671 | FLI1 | A673 | Muscle | GSM24366 | 75155 | 84628 |
| 762.6973 | 578 | 765.0981 | 703 | 529.6889 | 659 | 502.8466 | 706 | FLI1 | A673 | Muscle | GSM24366 | 73486 | 84629 |
| 793.9291 | 474 | 832.0983 | 621 | 573.2398 | 530 | 545.9855 | 565 | SRF | A673 | Muscle | GSM24366 | 45827 | 84630 |
| 748.7778 | 379 | 781.1373 | 498 | 506.5884 | 441 | 451.9006 | 459 | SRF | A673 | Muscle | GSM24366 | 40966 | 84631 |
| 738.9344 | 447 | 733.204 | 537 | 536.5789 | 533 | 474.3739 | 548 | CEBPB | MV4;11 | Blood | GSM23450 | 57210 | 84640 |
| 96.82389 | 38 | 73.48877 | 37 | 34.47643 | 36 | 68.56937 | 52 | MED1 | hMSC-TER1 | None | GSM28029 | 3739 | 84651 |
| 33.21835 | 70 | 55.08431 | 96 | 49.04484 | 134 | 95.88856 | 181 | TAL1 | K562 | Bone Marr | GSM28771 | 23216 | 84685 |
| 45.99179 | 45 | 72.77924 | 62 | 63.77256 | 82 | 84.61171 | 99 | GATA1 | K562 | Bone Marr | GSM28771 | 9595 | 84686 |
| 95.68388 | 28 | 73.60155 | 27 | 50.67146 | 30 | 43.43852 | 30 | SMC3 | HeLa | Cervix | GSM28138 | 2124 | 84733 |
| 264.0674 | 136 | 346.8148 | 173 | 202.6868 | 186 | 237.1586 | 214 | SMC3 | HeLa | Cervix | GSM28138 | 17814 | 84734 |
| 184.5441 | 120 | 270.138 | 160 | 163.9569 | 178 | 187.6422 | 203 | SMC3 | HeLa | Cervix | GSM28138 | 18952 | 84735 |
| 806.9735 | 120 | 995.8344 | 148 | 411.9585 | 112 | 413.1684 | 119 | MYC | RPM-MC | None | GSM23019 | 3427 | 84760 |
| 219.497 | 82 | 328.5784 | 112 | 151.9691 | 103 | 225.6638 | 132 | RUNX1 | Kasumi-1 | Blood | GSM72270 | 7816 | 8478 |
| 169.5684 | 200 | 145.04 | 217 | 129.2197 | 298 | 146.6371 | 339 | EP300 | None | None | GSM25502 | 48244 | 84782 |
| 451.8773 | 191 | 632.3953 | 250 | 301.1662 | 240 | 303.3224 | 260 | GATA2 | SKH1 | None | GSM24949 | 20592 | 84784 |
| 325.0156 | 155 | 555 | 223 | 263.382 | 214 | 274.4771 | 235 | GATA2 | SKH1 | None | GSM24949 | 18601 | 84785 |
| 115.1779 | 54 | 208.6784 | 81 | 149.5388 | 92 | 197.3599 | 112 | RUNX1 | Kasumi-1 | Blood | GSM72270 | 6457 | 8479 |
| 815.3013 | 518 | 818.6595 | 630 | 610.2443 | 626 | 577.942 | 661 | RUNX1 | None | Blood | GSM72270 | 56309 | 8481 |
| 1210.886 | 483 | 1274.877 | 659 | 927.0904 | 452 | 887.772 | 468 | KAT7 | RKO | Colon | GSM81828 | 11740 | 8482 |
| 141.3579 | 80 | 319.5336 | 132 | 95.196 | 105 | 178.1095 | 146 | VDR | THP-1 | Blood | GSM23714 | 11226 | 84824 |
| 24.00736 | 12 | 84.17717 | 26 | 19.70609 | 17 | 87.44989 | 37 | VDR | THP-1 | Blood | GSM23714 | 1639 | 84828 |
| 28.23879 | 13 | 84.55943 | 26 | 17.1954 | 16 | 87.56313 | 37 | VDR | THP-1 | Blood | GSM23714 | 1657 | 84830 |
| 1129.773 | 626 | 1175.547 | 809 | 820.4618 | 589 | 782.8332 | 615 | PCGF1 | None | None | GSM28058 | 22450 | 84831 |
| 584.0574 | 107 | 686.5684 | 128 | 215.0565 | 86 | 240.0526 | 97 | PAF1 | HCT-116 | HCT116 | GSM27690 | 3596 | 84915 |
| 264.0424 | 80 | 272.5135 | 90 | 134.692 | 84 | 272.1934 | 125 | ZEB1 | H1975 | Lung | GSM28572 | 5704 | 84926 |
| 1160.759 | 215 | 1176.732 | 273 | 817.6833 | 197 | 915.4121 | 231 | ZFX | MCF-7 | Breast | GSM27417 | 4657 | 84927 |
| 899.6786 | 275 | 918.9218 | 351 | 503.8001 | 267 | 551.4526 | 299 | ZFX | MCF-7 | Breast | GSM27417 | 14336 | 84928 |
| 579.7364 | 79 | 585.1683 | 86 | 308.3836 | 73 | 313.5342 | 78 | ZNF711 | MCF-7 | Breast | GSM27417 | 1743 | 84929 |
| 292.9693 | 124 | 518.2887 | 183 | 197.1516 | 157 | 168.7402 | 159 | RUNX1 | Kasumi-1 | Blood | GSM85082 | 13638 | 8493 |
| 858.5528 | 120 | 1164.656 | 157 | 426.4382 | 107 | 402.6359 | 110 | ZNF711 | MCF-7 | Breast | GSM27417 | 2641 | 84930 |
| 1192.056 | 241 | 1189.491 | 292 | 852.4604 | 205 | 826.697 | 214 | ZFX | C4-2B | Prostate | GSM27417 | 4856 | 84931 |
| 376.129 | 169 | 622.7752 | 240 | 202.8442 | 198 | 182.8933 | 206 | RUNX1 | Kasumi-1 | Blood | GSM85082 | 19971 | 8494 |
| 1086.283 | 507 | 1107.226 | 633 | 811.3162 | 495 | 792.2666 | 535 | NIPBL | HCT-116 | Colon | GSM28096 | 18943 | 84953 |
| 31.88776 | 25 | 91.34895 | 46 | 31.92159 | 40 | 147.8147 | 83 | CEBPB | HL-60 | None | GSM28711 | 4731 | 84984 |
| 151.6729 | 81 | 236.3551 | 112 | 120.6234 | 113 | 133.1006 | 127 | RUNX1 | HL-60 | None | GSM28711 | 10942 | 84986 |
| 1170.947 | 531 | 1187.506 | 658 | 865.7643 | 476 | 804.7634 | 464 | SUPT5H | MOLT-4 | None | GSM23677 | 14142 | 85013 |
| 351.2228 | 62 | 623.568 | 95 | 300.3573 | 77 | 269.5588 | 77 | INO80 | HepG2 | Liver | GSM28772 | 1994 | 85027 |
| 566.72 | 242 | 618.5232 | 282 | 652.8885 | 402 | 638.0057 | 438 | GRHL2 | MCF-7 | Breast | GSM29704 | 27720 | 85047 |
| 20.61474 | 12 | 10.67043 | 10 | 8.577165 | 13 | 33.42413 | 25 | SPI1 | None | Blood | GSM26956 | 1794 | 85055 |
| 37.4801 | 30 | 66.88481 | 44 | 33.94976 | 46 | 66.62676 | 65 | SPI1 | None | Blood | GSM26956 | 5592 | 85056 |

| | | | | | | | | | | | | | |
|---|---|---|---|---|---|---|---|---|---|---|---|---|---|
| 85.72959 | 37 | 57.02344 | 34 | 59.54226 | 47 | 47.27023 | 46 | SPI1 | None | Blood | GSM26956 | 3879 | 85057 |
| 451.6614 | 104 | 684.6754 | 144 | 332.6199 | 125 | 259.0753 | 118 | RXRG | None | Blood | GSM26956 | 5355 | 85058 |
| 551.0063 | 152 | 803.8482 | 205 | 350.009 | 176 | 363.7295 | 192 | RXRG | None | Blood | GSM26956 | 10374 | 85059 |
| 264.9414 | 50 | 568.3101 | 86 | 301.6 | 74 | 177.9341 | 59 | RXRG | None | Blood | GSM26956 | 1926 | 85060 |
| 1281.526 | 765 | 1338.888 | 981 | 889.2961 | 666 | 871.2778 | 723 | USF2 | None | None | GSM25502 | 23746 | 85079 |
| 861.8461 | 459 | 876.3196 | 576 | 617.4248 | 486 | 588.239 | 512 | EP300 | None | None | GSM25502 | 37469 | 85080 |
| 1059.29 | 268 | 1069.764 | 332 | 795.6367 | 254 | 829.1994 | 307 | BRD4 | J-Lat A72 | None | GSM26768 | 8730 | 85084 |
| 2.659574 | 11 | 6.14001 | 17 | 0.50311 | 15 | 34.55933 | 50 | CEBPB | None | Blood | GSM25928 | 5723 | 85110 |
| 0.112592 | 8 | -0.053441 | 6 | 0.122167 | 17 | 9.17372 | 41 | MAF | None | Blood | GSM25928 | 7826 | 85111 |
| 109.4682 | 112 | 130.7448 | 137 | 96.71795 | 174 | 86.5159 | 183 | SPI1 | None | Blood | GSM25928 | 25202 | 85115 |
| 3.074531 | 7 | 10.15518 | 13 | 0.250026 | 7 | 4.98864 | 16 | SETDB1 | HeLa | Cervix | GSM25144 | 2690 | 85129 |
| 100.3403 | 21 | 174.6492 | 32 | 58.09698 | 22 | 142.607 | 38 | HDAC2 | 5637 | None | GSM17067 | 1005 | 85150 |
| 657.8512 | 202 | 760.2094 | 241 | 291.2501 | 204 | 273.079 | 214 | EWSR1 | A673 | None | GSM26664 | 15980 | 85158 |
| 303.092 | 74 | 400.8297 | 95 | 155.9244 | 75 | 146.7361 | 78 | EWSR1 | A673 | None | GSM26664 | 4056 | 85159 |
| 492.2197 | 160 | 647.8169 | 204 | 267.083 | 177 | 242.8991 | 183 | EWSR1 | A673 | None | GSM26664 | 13084 | 85160 |
| 141.2388 | 80 | 148.3354 | 92 | 49.68884 | 82 | 50.83209 | 90 | EWSR1 | A673 | None | GSM26664 | 10860 | 85161 |
| 1031.571 | 868 | 1093.53 | 1107 | 720.9142 | 965 | 681.0673 | 1028 | MAX | MKL-1 | None | GSM17118 | 74630 | 85164 |
| 1131.862 | 511 | 1176.991 | 675 | 839.1435 | 497 | 834.5642 | 554 | EP400 | MKL-1 | None | GSM17118 | 19709 | 85166 |
| 141.6147 | 300 | 214.0638 | 392 | 152.5656 | 512 | 188.8866 | 596 | TET2 | Jurkat | Blood | GSM22746 | 99856 | 85217 |
| 926.8143 | 484 | 956.2042 | 627 | 681.4508 | 523 | 655.3189 | 555 | RUNX1 | Jurkat | Blood | GSM22746 | 34454 | 85218 |
| 528.7725 | 280 | 703.0528 | 378 | 298.7412 | 342 | 329.2144 | 385 | CTCF | HCT-116 | Colon | GSM28096 | 37710 | 85284 |
| 483.8535 | 310 | 687.3574 | 448 | 341.9289 | 421 | 366.9243 | 470 | CTCF | HCT-116 | Colon | GSM28096 | 49478 | 85285 |
| 349.9395 | 170 | 464.3259 | 217 | 222.2835 | 214 | 384.1552 | 288 | CTCF | HCT-116 | Colon | GSM28096 | 21103 | 85286 |
| 217.1168 | 70 | 332.5057 | 97 | 106.6658 | 73 | 320.4286 | 131 | SMC1A | HCT-116 | Colon | GSM28096 | 5344 | 85288 |
| 424.8868 | 247 | 688.4851 | 346 | 267.5135 | 318 | 397.1251 | 399 | SMC1A | HCT-116 | Colon | GSM28096 | 36148 | 85289 |
| 203.4691 | 70 | 230.8905 | 83 | 98.51563 | 74 | 114.4332 | 85 | RAD21 | HCT-116 | Colon | GSM28096 | 6031 | 85290 |
| 492.8701 | 71 | 694.4314 | 95 | 387.6022 | 83 | 238.117 | 67 | MYCN | KELLY | None | GSM24828 | 1813 | 85291 |
| 1169.185 | 267 | 1163.177 | 318 | 907.726 | 252 | 851.617 | 243 | VEZF1 | HeLa-S3 | Cervix | GSM86619 | 6790 | 8532 |
| 584.1813 | 91 | 884.1941 | 127 | 344.0414 | 92 | 380.4189 | 103 | BRD4 | None | None | GSM28071 | 2715 | 85341 |
| 872.5781 | 410 | 881.8741 | 507 | 621.2896 | 423 | 633.1328 | 498 | EP300 | None | None | GSM25502 | 32680 | 85378 |
| 714.5227 | 484 | 701.1978 | 573 | 436.0352 | 549 | 426.3978 | 595 | ATF3 | None | None | GSM23338 | 72829 | 85390 |
| 653.5091 | 87 | 775.5805 | 105 | 509.3549 | 102 | 572.2368 | 115 | ATF3 | None | None | GSM23338 | 2242 | 85391 |
| 886.9621 | 609 | 915.603 | 769 | 641.0247 | 676 | 596.1826 | 698 | ATF3 | GM00739 | None | GSM23338 | 53676 | 85392 |
| 1090.185 | 380 | 1048.388 | 417 | 820.3409 | 356 | 769.8406 | 352 | BRD4 | N87 | Stomach | GSM23594 | 12279 | 85397 |
| 960.3295 | 425 | 972.7086 | 530 | 738.1129 | 460 | 719.4002 | 502 | BRD4 | N87 | Stomach | GSM23594 | 21752 | 85398 |
| 1022.174 | 430 | 1006.843 | 504 | 747.1755 | 407 | 749.8011 | 468 | BRD4 | H3122 | Lung | GSM23594 | 18875 | 85399 |
| 1084.748 | 369 | 1067.331 | 429 | 811.2393 | 344 | 778.7215 | 357 | BRD4 | H3122 | Lung | GSM23594 | 12132 | 85401 |
| 712.3212 | 81 | 465.3955 | 67 | 217.6972 | 53 | 372.4074 | 76 | BRD4 | H3122 | Lung | GSM23594 | 1329 | 85402 |
| 535.2342 | 84 | 512.6037 | 89 | 260.0563 | 76 | 286.5373 | 85 | BRD4 | A549 | Lung | GSM23594 | 2386 | 85403 |
| 1082.676 | 209 | 1083.111 | 250 | 585.1423 | 182 | 702.858 | 213 | BRD4 | H3122 | Lung | GSM23594 | 5878 | 85404 |
| 444.0766 | 61 | 455.1372 | 67 | 305.6788 | 65 | 225.8664 | 58 | BRD4 | A549 | Lung | GSM23594 | 1325 | 85405 |
| 1081.145 | 306 | 1086.581 | 381 | 838.0426 | 310 | 814.806 | 326 | BRD4 | A549 | Lung | GSM23594 | 10260 | 85406 |
| 190.7551 | 141 | 320.6909 | 199 | 257.7825 | 249 | 311.5692 | 290 | TFAP2A | None | None | GSM28176 | 25406 | 85423 |
| 690.0259 | 240 | 798.5828 | 305 | 472.0315 | 300 | 640.1029 | 368 | TFAP2A | None | None | GSM28176 | 22735 | 85427 |
| 5.894997 | 14 | 13.47894 | 22 | 14.37374 | 33 | 111.0265 | 79 | GATA2 | None | None | GSM28176 | 5564 | 85433 |
| 318.4655 | 63 | 546.2398 | 94 | 133.2455 | 54 | 116.4553 | 54 | PHIP | HCT-116 | HCT116 | GSM27113 | 2180 | 85459 |
| 547.1411 | 100 | 666.8926 | 122 | 245.9411 | 88 | 226.1642 | 90 | PHIP | HCT-116 | HCT116 | GSM27113 | 3113 | 85463 |
| 1113.213 | 301 | 1108.423 | 362 | 827.311 | 272 | 782.0185 | 271 | RNF2 | HEK293T | Embryonic | GSM85500 | 9826 | 8549 |
| 345.4309 | 62 | 610.3722 | 95 | 277.9615 | 76 | 242.9513 | 75 | RYBP | HEK293T | Embryonic | GSM85500 | 2355 | 8550 |
| 656.107 | 107 | 818.5357 | 133 | 380.6164 | 109 | 487.3258 | 132 | BMI1 | HEK293T | Embryonic | GSM85501 | 3624 | 8553 |
| 1057.125 | 352 | 1046.852 | 420 | 797.6629 | 343 | 766.4044 | 354 | BRD4 | K562 | Bone Marr | GSM26352 | 13019 | 85536 |
| 82.31586 | 42 | 89.32303 | 49 | 82.79301 | 64 | 151.3852 | 90 | FOSL1 | MNNG | None | GSM26353 | 5233 | 85560 |
| 1006.846 | 318 | 990.3038 | 374 | 735.9163 | 303 | 586.4555 | 289 | HIRA | IMR90 | Lung | GSM19367 | 15086 | 85563 |
| 274.8803 | 41 | 469.5201 | 62 | 162.4938 | 41 | 133.5823 | 39 | STAT1 | AGS | None | GSM27528 | 1127 | 85567 |
| 217.6691 | 152 | 311.0297 | 200 | 157.399 | 211 | 195.9373 | 248 | CEBPD | JHU-06 | None | GSM23943 | 25889 | 85611 |
| 105.383 | 44 | 265.356 | 79 | 101.9502 | 65 | 74.80029 | 61 | RAD21 | IMR-5 | None | GSM27593 | 4674 | 85620 |
| 716.2948 | 168 | 929.7295 | 214 | 252.1826 | 145 | 399.1235 | 192 | GPS2 | HEK293T | Embryonic | GSM86574 | 9800 | 8563 |
| 257.2553 | 217 | 426.0575 | 301 | 205.4858 | 318 | 231.899 | 361 | CTCF | HeLa | None | GSM27477 | 42810 | 85639 |
| 1071.381 | 560 | 1123.35 | 747 | 776.8797 | 558 | 748.705 | 591 | TBL1X | HEK293T | Embryonic | GSM86574 | 28238 | 8564 |
| 340.4979 | 276 | 517.9406 | 369 | 227.2542 | 381 | 260.8808 | 435 | CTCF | HeLa | None | GSM27477 | 54580 | 85640 |
| 311.4776 | 261 | 477.0139 | 350 | 222.9886 | 370 | 272.371 | 431 | CTCF | HeLa | None | GSM27477 | 52765 | 85641 |
| 403.5948 | 65 | 772.8456 | 105 | 242.6001 | 67 | 274.7533 | 76 | NCOR1 | HEK293T | Embryonic | GSM86574 | 2111 | 8565 |
| 924.2784 | 146 | 818.2285 | 148 | 473.7102 | 137 | 501.408 | 150 | ESR1 | MIN | Breast | GSM21251 | 4275 | 85653 |
| 165.0906 | 47 | 152.9062 | 50 | 240.7253 | 83 | 536.8758 | 135 | ESR1 | MIN | Breast | GSM21251 | 3473 | 85654 |
| 463.6071 | 162 | 477.3429 | 183 | 307.8395 | 199 | 367.0042 | 231 | TP53 | SW480 | Colon | GSM27465 | 14870 | 85656 |
| 536.0654 | 80 | 958.4051 | 124 | 290.1975 | 77 | 466.504 | 106 | NCOR1 | HEK293T | Embryonic | GSM86574 | 2311 | 8566 |
| 338.1558 | 148 | 555.1686 | 210 | 214.8745 | 185 | 326.1904 | 237 | CTCF | None | Cord blood | GSM28617 | 17425 | 85680 |
| 1184.971 | 622 | 1239.36 | 817 | 851.2584 | 561 | 808.607 | 575 | USF2 | None | None | GSM25502 | 19974 | 85688 |
| 294.4956 | 185 | 369.6955 | 230 | 115.0653 | 204 | 153.564 | 245 | EP300 | None | None | GSM25502 | 28673 | 85689 |
| 980.2098 | 535 | 984.5912 | 647 | 785.501 | 642 | 765.173 | 703 | BRD4 | RT112 | Urinary Bla | GSM23594 | 30544 | 85706 |
| 896.3641 | 505 | 909.8636 | 626 | 672.5676 | 566 | 645.8615 | 605 | SMARCC1 | HS-ES-2M | epithelioid | GSM24096 | 37925 | 85732 |
| 1067.319 | 342 | 1069.07 | 417 | 832.8149 | 354 | 774.8782 | 340 | SMARCA4 | HS-ES-2M | epithelioid | GSM24096 | 12410 | 85737 |

| | | | | | | | | | | | | | |
|---|---|---|---|---|---|---|---|---|---|---|---|---|---|
| 1055.317 | 547 | 1086.776 | 696 | 782.6364 | 552 | 758.1581 | 590 | SMARCC1 | HS-ES-2M | epithelioid | GSM24096 | 24862 | 85738 |
| 22.34197 | 10 | 43.73818 | 16 | 10.71733 | 11 | 44.73116 | 23 | ARID1A | HCT-116 | HCT116 | GSM27197 | 1248 | 85741 |
| 704.3572 | 379 | 761.2186 | 526 | 505.3597 | 476 | 523.0038 | 524 | CTCF | ID00016 | None | GSM20407 | 47317 | 85767 |
| 1016.284 | 465 | 1010.873 | 556 | 739.5219 | 436 | 718.9695 | 474 | BRD4 | LCL | Blood | GSM26435 | 19655 | 85771 |
| 965.8509 | 395 | 958.7807 | 469 | 691.6151 | 368 | 682.3912 | 409 | BRD4 | LCL | Blood | GSM26435 | 19274 | 85772 |
| 609.089 | 346 | 710.9379 | 487 | 431.3332 | 468 | 432.6695 | 510 | CTCF | JHU-011 | Fetal Liver | GSM27300 | 53371 | 85815 |
| 639.8611 | 361 | 701.5465 | 490 | 440.96 | 484 | 448.1024 | 530 | CTCF | None | Blood | GSM27300 | 56005 | 85819 |
| 660.5503 | 364 | 711.4437 | 499 | 500.5885 | 503 | 468.911 | 535 | CTCF | JHU-011 | Blood | GSM27300 | 55230 | 85826 |
| 179.2645 | 44 | 184.4753 | 49 | 62.62824 | 36 | 169.9784 | 63 | MKL2 | A673 | Muscle | GSM24366 | 2028 | 85837 |
| 1004.656 | 322 | 936.686 | 332 | 809.2239 | 379 | 879.8775 | 520 | AR | LNCaP | Prostate | GSM23938 | 15396 | 85854 |
| 101.4974 | 23 | 67.12628 | 20 | 63.19851 | 25 | 95.03564 | 33 | AR | LNCaP | Prostate | GSM23938 | 1105 | 85855 |
| 1279.558 | 177 | 1198.634 | 172 | 1158.535 | 252 | 1212.871 | 327 | AR | LNCaP | Prostate | GSM23938 | 3098 | 85856 |
| 999.9004 | 293 | 957.3803 | 323 | 827.5578 | 365 | 921.3671 | 533 | AR | LNCaP | Prostate | GSM23938 | 13805 | 85857 |
| 780.8108 | 345 | 831.3556 | 474 | 616.8086 | 442 | 583.8731 | 462 | CTCF | None | Fetal Liver | GSM27300 | 34616 | 85870 |
| 438.0729 | 87 | 618.6725 | 116 | 231.7724 | 86 | 208.5787 | 87 | ESR1 | MIN | Breast | GSM21251 | 3537 | 85907 |
| 437.6723 | 87 | 511.3559 | 104 | 242.0865 | 88 | 366.1859 | 116 | ESR1 | MIN | Breast | GSM21251 | 3546 | 85908 |
| 696.6833 | 442 | 704.9892 | 549 | 513.7294 | 550 | 494.1569 | 597 | KLF15 | None | Muscle | GSM25199 | 61713 | 85934 |
| 607.5264 | 357 | 658.7212 | 448 | 411.495 | 474 | 445.8528 | 531 | KLF15 | None | Muscle | GSM25199 | 54228 | 85935 |
| 1240.545 | 394 | 1266.847 | 507 | 949.824 | 357 | 911.8855 | 364 | KLF15 | None | Muscle | GSM25199 | 8397 | 85941 |
| 1172.269 | 427 | 1196.911 | 546 | 903.7303 | 407 | 848.6145 | 400 | KLF15 | None | Muscle | GSM25199 | 10801 | 85942 |
| 254.5935 | 97 | 208.3861 | 98 | 473.2456 | 191 | 610.2793 | 231 | ESR1 | MCF-7 | Breast | GSM29704 | 8894 | 85954 |
| 150.5006 | 32 | 204.3937 | 42 | 82.61655 | 33 | 116.527 | 42 | KLF4 | TIME | Brain | GSM25737 | 1620 | 85956 |
| 531.3087 | 65 | 629.5151 | 78 | 344.297 | 66 | 382.4516 | 74 | SUPT5H | HeLa | Adherent I | GSM24650 | 1187 | 85966 |
| 1212.052 | 384 | 1213.359 | 468 | 917.8918 | 339 | 902.0861 | 370 | NELFA | HeLa | Adherent I | GSM24650 | 8344 | 85971 |
| 1228.712 | 349 | 1242.463 | 437 | 945.4105 | 314 | 906.654 | 320 | NELFA | HeLa | Adherent I | GSM24650 | 6899 | 85972 |
| 4.997408 | 7 | 6.356663 | 9 | 10.42614 | 16 | 165.1217 | 60 | GATA3 | None | None | GSM28176 | 2281 | 85975 |
| 1.406109 | 6 | 3.838353 | 10 | 4.653586 | 16 | 77.09728 | 51 | GATA3 | None | None | GSM28176 | 3201 | 85977 |
| 5.183263 | 9 | 14.86998 | 16 | 22.38943 | 27 | 203.9167 | 79 | GATA2 | None | None | GSM28176 | 3245 | 85979 |
| 10.12053 | 29 | 22.44195 | 44 | 8.945863 | 50 | 50.4832 | 92 | CEBPB | None | Blood | GSM28044 | 11785 | 85984 |
| 27.37926 | 68 | 42.52934 | 91 | 26.43607 | 117 | 55.31245 | 158 | CEBPB | None | Blood | GSM28044 | 24863 | 85985 |
| 18.28092 | 22 | 11.0668 | 21 | 13.23548 | 33 | 47.84003 | 57 | SPI1 | None | Blood | GSM28044 | 5800 | 85986 |
| 1290.196 | 138 | 1239.414 | 145 | 631.7989 | 117 | 691.8991 | 130 | ZNF131 | HeLa | Cervix | GSM19590 | 2158 | 85994 |
| 598.5928 | 109 | 520.7274 | 110 | 352.5372 | 113 | 362.7748 | 122 | GATAD1 | HeLa | Cervix | GSM19590 | 4086 | 85995 |
| 57.83202 | 49 | 80.48109 | 64 | 66.26612 | 82 | 74.58383 | 93 | MED1 | hMSC-TERT | None | GSM28029 | 9256 | 86002 |
| 532.3958 | 99 | 628.7474 | 119 | 403.4288 | 117 | 511.6735 | 141 | ERF | VCaP | Prostate | GSM26124 | 3698 | 86004 |
| 348.1167 | 46 | 444.218 | 58 | 316.6446 | 58 | 418.32 | 72 | ERF | VCaP | Prostate | GSM26124 | 1033 | 86005 |
| 976.2753 | 212 | 952.4421 | 243 | 617.6901 | 228 | 695.4051 | 258 | ERG | VCaP | Prostate | GSM26124 | 9868 | 86006 |
| 333.5519 | 72 | 505.6962 | 100 | 301.5582 | 95 | 485.4912 | 130 | AR | VCaP | Prostate | GSM26124 | 3292 | 86007 |
| 787.321 | 295 | 778.4816 | 354 | 708.9561 | 472 | 758.7619 | 616 | AHR | MCF-7 | Breast | GSM24067 | 28025 | 86157 |
| 164.0624 | 31 | 207.5886 | 39 | 177.2775 | 45 | 449.3732 | 81 | AHRR | MCF-7 | Breast | GSM24067 | 1278 | 86159 |
| 818.489 | 193 | 892.4074 | 223 | 422.6443 | 200 | 409.1791 | 211 | ZBTB7A | K562 | Bone Marr | GSM27715 | 11623 | 86195 |
| 1040.562 | 500 | 1083.982 | 659 | 775.5261 | 514 | 752.8407 | 549 | ZBTB7A | K562 | Bone Marr | GSM27715 | 24897 | 86196 |
| 1052.036 | 516 | 1103.68 | 687 | 797.6785 | 540 | 732.1975 | 523 | ZBTB7A | K562 | Bone Marr | GSM27715 | 23929 | 86198 |
| 769.6117 | 158 | 1005.746 | 214 | 513.0042 | 178 | 755.9002 | 231 | BCL11A | HUDEP-2 | None | GSM27715 | 7015 | 86200 |
| 564.0007 | 83 | 888.9638 | 119 | 755.7632 | 133 | 1052.228 | 170 | BRD4 | C4-2 | Prostate | GSM23504 | 2310 | 86205 |
| 1238.998 | 692 | 1240.001 | 839 | 1017.178 | 839 | 1013.696 | 956 | BRD4 | C4-2 | Prostate | GSM23504 | 22587 | 86207 |
| 979.5125 | 330 | 963.2911 | 380 | 836.8202 | 436 | 857.7133 | 523 | ESR1 | SUM44 | Breast | GSM26708 | 15763 | 86209 |
| 921.1604 | 466 | 909.1371 | 544 | 791.5825 | 647 | 798.488 | 747 | ESR1 | SUM44 | Breast | GSM26708 | 29993 | 86210 |
| 266.5894 | 86 | 138.7336 | 69 | 250.5922 | 121 | 409.5133 | 164 | ESR1 | MCF-7 | Breast | GSM26708 | 6606 | 86212 |
| 622.3303 | 125 | 640.1278 | 139 | 827.7995 | 201 | 928.2165 | 247 | ESR1 | MCF-7 | Breast | GSM26708 | 5191 | 86213 |
| 775.0594 | 465 | 779.1586 | 567 | 688.2971 | 723 | 709.1826 | 865 | ESR1 | MCF-7 | Breast | GSM26489 | 52541 | 86282 |
| 737.8898 | 347 | 764.8033 | 453 | 689.1554 | 596 | 722.2085 | 736 | ESR1 | MCF-7 | Breast | GSM26489 | 40384 | 86283 |
| 445.171 | 119 | 436.3607 | 130 | 528.4626 | 185 | 729.1724 | 232 | GATA3 | T47D | Mammary | GSM26445 | 7537 | 86315 |
| 589.5397 | 136 | 535.4814 | 142 | 631.6399 | 198 | 860.8064 | 260 | GATA3 | T47D | Mammary | GSM26445 | 7112 | 86316 |
| 428.1304 | 104 | 344.8306 | 102 | 559.034 | 169 | 622.4342 | 190 | GATA3 | T47D | Mammary | GSM26445 | 5876 | 86317 |
| 62.84304 | 16 | 78.13501 | 20 | 57.94668 | 22 | 156.0066 | 40 | ESR1 | T47D | Mammary | GSM26445 | 1002 | 86340 |
| 35.66069 | 12 | 49.46172 | 16 | 8.199694 | 9 | 70.15295 | 27 | POU5F1 | iPS cells | None | GSM21837 | 1117 | 86398 |
| 44.61711 | 18 | 43.91997 | 20 | 15.39961 | 17 | 7.604852 | 14 | CTCF | IMR-5 | None | GSM27593 | 1991 | 86416 |
| 1179.609 | 391 | 1180.191 | 478 | 884.6799 | 351 | 841.1828 | 354 | BRD4 | Akata | None | GSM26115 | 10121 | 86429 |
| 1008.455 | 510 | 1031.587 | 647 | 734.7002 | 497 | 666.3853 | 474 | BRD4 | MOLT-4 | None | GSM20909 | 23452 | 86433 |
| 282.0524 | 248 | 467.9888 | 343 | 180.2942 | 340 | 234.7837 | 404 | CTCF | HeLa | None | GSM27477 | 50845 | 86441 |
| 1534.839 | 1133 | 1558.544 | 1350 | 1074.585 | 1261 | 1119.715 | 1531 | BRD4 | C4-2 | Prostate | GSM23504 | 38644 | 86456 |
| 993.4261 | 349 | 1060.419 | 499 | 764.1007 | 365 | 729.0656 | 377 | CHD1 | None | Bone Marr | GSM23599 | 15758 | 86486 |
| 433.7744 | 93 | 568.8183 | 119 | 272.5026 | 103 | 887.671 | 204 | AR | VCaP | Prostate | GSM71739 | 4628 | 8651 |
| 754.9632 | 527 | 686.4889 | 553 | 645.0578 | 791 | 701.8164 | 1014 | AR | VCaP | Prostate | GSM71739 | 70786 | 8652 |
| 489.8003 | 107 | 638.2245 | 136 | 222.9674 | 100 | 236.3729 | 110 | BRD4 | SKMEL5 | None | GSM19682 | 4941 | 86522 |
| 653.3324 | 78 | 778.155 | 94 | 364.3793 | 73 | 432.2835 | 85 | ESR1 | None | Breast | GSM27971 | 1467 | 86526 |
| 257.1322 | 49 | 338.4475 | 63 | 406.369 | 88 | 729.6914 | 130 | ESR1 | None | Breast | GSM27971 | 2041 | 86527 |
| 727.2233 | 314 | 715.6424 | 372 | 619.3747 | 469 | 702.4979 | 662 | AR | VCaP | Prostate | GSM71739 | 37973 | 8653 |
| 11.464 | 15 | 14.50711 | 19 | 1.019644 | 13 | 13.21942 | 29 | POU5F1 | None | None | GSM26489 | 4138 | 86531 |
| 15.64656 | 12 | 46.89586 | 23 | 11.21043 | 17 | 36.61951 | 30 | POU5F1 | H9 | Embryo | GSM26489 | 2375 | 86532 |
| 77.86766 | 102 | 76.0725 | 116 | 21.74206 | 116 | 32.88997 | 142 | NANOG | None | None | GSM26489 | 25109 | 86533 |

| | | | | | | | | | | | | | |
|---|---|---|---|---|---|---|---|---|---|---|---|---|---|
| 32.13946 | 66 | 30.27894 | 75 | 16.07554 | 93 | 19.66637 | 108 | NANOG | H9 | Embryo | GSM26489 | 20758 | 86534 |
| 1163.777 | 620 | 1169.598 | 760 | 958.9845 | 772 | 932.7606 | 833 | ERG | VCaP | Prostate | GSM71739 | 24788 | 8654 |
| 1054.572 | 253 | 1045.144 | 300 | 890.6833 | 317 | 920.9189 | 392 | ESR1 | None | Breast | GSM27971 | 9599 | 86546 |
| 1118.858 | 642 | 1165.073 | 827 | 927.0695 | 814 | 921.0964 | 924 | ERG | VCaP | Prostate | GSM71739 | 30026 | 8655 |
| 1136.288 | 785 | 1170.027 | 984 | 894.1704 | 953 | 900.0454 | 1104 | ERG | VCaP | Prostate | GSM71739 | 43624 | 8656 |
| 1027.366 | 546 | 1087.688 | 738 | 779.9165 | 589 | 721.7784 | 581 | ESR1 | Ishikawa | Uterus | GSM26640 | 28732 | 86560 |
| 693.3763 | 107 | 1035.412 | 148 | 486.1428 | 119 | 543.7098 | 134 | ESR1 | Ishikawa | Uterus | GSM26640 | 3165 | 86561 |
| 1231.581 | 360 | 1283.838 | 492 | 942.9001 | 333 | 925.7497 | 356 | HDAC1 | VCaP | Prostate | GSM71739 | 8324 | 8657 |
| 1237.627 | 391 | 1272.23 | 517 | 936.462 | 349 | 906.7605 | 364 | HDAC1 | VCaP | Prostate | GSM71739 | 9061 | 8658 |
| 1209.9 | 425 | 1255.449 | 570 | 984.4624 | 475 | 965.6399 | 517 | HDAC2 | VCaP | Prostate | GSM71740 | 11578 | 8659 |
| 1169.748 | 517 | 1197.735 | 657 | 940.1009 | 582 | 927.7843 | 643 | HDAC2 | VCaP | Prostate | GSM71740 | 17288 | 8660 |
| 1294.647 | 209 | 1212.026 | 200 | 1181.736 | 297 | 1192.551 | 346 | HDAC3 | VCaP | Prostate | GSM71740 | 3551 | 8661 |
| 1036.996 | 291 | 983.9105 | 306 | 891.5579 | 389 | 956.1413 | 523 | HDAC3 | VCaP | Prostate | GSM71740 | 12208 | 8662 |
| 61.98604 | 23 | 118.3533 | 36 | 63.56686 | 34 | 141.103 | 54 | EZH2 | VCaP | Prostate | GSM71740 | 1952 | 8663 |
| 229.2994 | 74 | 297.8198 | 94 | 176.4285 | 95 | 319.6276 | 135 | EZH2 | VCaP | Prostate | GSM71740 | 5775 | 8664 |
| 548.6662 | 66 | 767.0867 | 88 | 243.452 | 54 | 289.6057 | 63 | OGT | None | Colon | GSM24599 | 1225 | 86643 |
| 25.69429 | 52 | 25.28254 | 60 | 14.51776 | 76 | 38.89686 | 110 | GLI2 | THP-1 | Blood | GSM26617 | 16903 | 86683 |
| 20.91619 | 33 | 30.91369 | 44 | 12.65267 | 48 | 67.41281 | 91 | GLI2 | THP-1 | Blood | GSM26617 | 9784 | 86686 |
| 942.1209 | 201 | 937.8133 | 235 | 841.3459 | 293 | 865.6124 | 354 | ESR1 | None | Breast | GSM27970 | 10208 | 86729 |
| 45.43945 | 20 | 9.708758 | 11 | 6.822177 | 14 | 52.4755 | 35 | STAG1 | None | Embryo | GSM80062 | 2382 | 8673 |
| 369.2972 | 80 | 385.4842 | 90 | 391.9761 | 116 | 676.3639 | 165 | AR | None | Breast | GSM27970 | 3945 | 86730 |
| 945.8369 | 232 | 882.8464 | 236 | 772.3389 | 282 | 806.7815 | 354 | FOXA1 | None | Breast | GSM27970 | 12131 | 86731 |
| 356.6776 | 73 | 372.2231 | 82 | 363.8055 | 103 | 481.1891 | 127 | ESR1 | None | Breast | GSM27970 | 3303 | 86733 |
| 654.0457 | 131 | 429.4425 | 114 | 342.0755 | 131 | 511.3907 | 171 | FOXA1 | None | Breast | GSM27970 | 5929 | 86734 |
| 447.9564 | 69 | 306.8186 | 60 | 678.473 | 119 | 880.5799 | 146 | AR | MCF-7 | Breast | GSM27970 | 2032 | 86735 |
| 392.3279 | 125 | 334.3203 | 128 | 627.0786 | 228 | 800.4118 | 288 | NR3C1 | MCF-7 | Breast | GSM27970 | 9903 | 86736 |
| 1165.06 | 518 | 1197.578 | 666 | 880.2872 | 494 | 830.0053 | 495 | INTS12 | None | None | GSM21255 | 15180 | 86783 |
| 1024.206 | 533 | 1052.758 | 682 | 728.7924 | 515 | 714.6805 | 569 | RUNX1 | SKH1 | None | GSM23268 | 29580 | 86788 |
| 827.493 | 184 | 938.8878 | 225 | 480.2964 | 199 | 421.7564 | 200 | CEBPA | SKH1 | None | GSM23268 | 10099 | 86793 |
| 384.083 | 84 | 366.5788 | 90 | 233.962 | 92 | 288.7983 | 109 | CEBPA | SKH1 | None | GSM23268 | 4338 | 86794 |
| 90.93819 | 32 | 118.7899 | 41 | 100.561 | 50 | 118.982 | 58 | JUN | JHU-06 | None | GSM25761 | 3000 | 86803 |
| 847.7825 | 462 | 862.6283 | 581 | 637.3326 | 547 | 637.3723 | 623 | JUN | JHU-06 | None | GSM25761 | 44044 | 86804 |
| 888.6473 | 591 | 912.5528 | 748 | 659.4542 | 693 | 616.5732 | 719 | JUN | JHU-06 | None | GSM25761 | 56610 | 86805 |
| 1050.958 | 293 | 1050.355 | 360 | 787.9739 | 286 | 766.2858 | 303 | ERG | JHU-06 | None | GSM25761 | 12107 | 86806 |
| 3.683695 | 68 | 1.273524 | 70 | 15.97227 | 167 | 47.75792 | 236 | GATA4 | BJ | Skin | GSM24014 | 45189 | 86817 |
| 531.7587 | 162 | 547.2068 | 182 | 494.3962 | 225 | 664.4128 | 277 | STAT3 | MDA-MB-4 | None | GSM22780 | 11744 | 86819 |
| 672.1316 | 474 | 725.2349 | 642 | 525.8072 | 652 | 535.4827 | 762 | CTCF | MCF-7 | Breast | GSM28779 | 77377 | 86828 |
| 925.6964 | 507 | 918.8552 | 604 | 759.2744 | 663 | 748.6068 | 742 | STAT3 | MDA-MB-4 | None | GSM22780 | 37254 | 86835 |
| 59.93788 | 25 | 126.5961 | 41 | 83.64526 | 43 | 72.09723 | 43 | SUZ12 | NT2-D1 | None | GSM27058 | 2314 | 86837 |
| 1252.023 | 239 | 1225.598 | 275 | 908.8029 | 197 | 896.4559 | 207 | U2AF1 | HEK293T | Embryonic | GSM25594 | 4390 | 86849 |
| 1245.981 | 148 | 1234.14 | 178 | 631.9602 | 129 | 588.8043 | 131 | U2AF1 | HEK293T | Embryonic | GSM25594 | 2626 | 86850 |
| 1109.91 | 125 | 1219.851 | 143 | 537.4594 | 107 | 448.3397 | 102 | U2AF1 | HEK293T | Embryonic | GSM25594 | 2098 | 86856 |
| 1030.27 | 119 | 1170.029 | 139 | 505.3104 | 103 | 427.2447 | 99 | U2AF1 | HEK293T | Embryonic | GSM25594 | 2084 | 86857 |
| 337.3053 | 47 | 216.9391 | 39 | 266.0956 | 55 | 413.7622 | 75 | FOXA1 | None | Breast | GSM27970 | 1175 | 86889 |
| 861.0903 | 297 | 835.8978 | 338 | 748.728 | 426 | 755.0456 | 492 | GATA3 | None | Breast | GSM27971 | 20367 | 86894 |
| 513.1817 | 115 | 526.3009 | 128 | 504.1972 | 159 | 709.2355 | 202 | NR3C1 | None | Breast | GSM27971 | 5561 | 86896 |
| 169.1725 | 33 | 176.1467 | 37 | 169.7772 | 46 | 269.476 | 63 | NR3C1 | None | Breast | GSM27971 | 1398 | 86897 |
| 248.9633 | 37 | 207.4117 | 36 | 139.3981 | 36 | 291.8279 | 58 | NR3C1 | None | Breast | GSM27971 | 1045 | 86899 |
| 245.9779 | 58 | 290.5855 | 70 | 365.2698 | 101 | 605.1419 | 141 | GATA3 | None | Breast | GSM27971 | 3105 | 86902 |
| 108.4982 | 52 | 132.0281 | 64 | 162.2355 | 94 | 188.4485 | 108 | RXRA | JMSU-1 | None | GSM28773 | 6094 | 86915 |
| 216.1474 | 73 | 180.1152 | 74 | 159.9501 | 93 | 230.1741 | 118 | RXRA | JMSU-1 | None | GSM28773 | 6166 | 86919 |
| 354.4342 | 150 | 574.2736 | 211 | 329.7387 | 215 | 477.8131 | 272 | RELA | Detroit 562 | None | GSM24198 | 15247 | 86943 |
| 215.5711 | 284 | 141.7647 | 285 | 125.2199 | 392 | 160.0766 | 461 | FOXA2 | BJ | Skin | GSM29774 | 69308 | 86987 |
| 519.8778 | 271 | 706.3926 | 365 | 376.1004 | 359 | 351.1821 | 380 | SMC3 | HeLa | None | GSM27477 | 34295 | 87039 |
| 614.7999 | 314 | 719.7063 | 427 | 462.9532 | 431 | 488.2668 | 477 | SMC3 | HeLa | None | GSM27477 | 45375 | 87040 |
| 412.1045 | 214 | 569.1171 | 277 | 305.9629 | 286 | 285.5197 | 302 | SMC3 | HeLa | None | GSM27477 | 26715 | 87042 |
| 709.7498 | 339 | 736.0674 | 446 | 539.1184 | 456 | 542.2404 | 500 | SMC3 | HeLa | None | GSM27477 | 45526 | 87043 |
| 45.53294 | 37 | 31.34115 | 36 | 39.52787 | 55 | 28.18577 | 53 | SMAD2 | H1 | Embryo | GSM26356 | 6380 | 87046 |
| 26.67892 | 28 | 30.88274 | 34 | 9.939388 | 33 | 46.43394 | 61 | TEAD4 | H1 | Embryo | GSM26356 | 6390 | 87047 |
| 60.98514 | 34 | 60.09183 | 38 | 42.85522 | 45 | 82.67445 | 64 | SMAD2 | H1 | Embryo | GSM26356 | 4575 | 87048 |
| 1211.415 | 270 | 1109.302 | 251 | 989.6844 | 292 | 1061.956 | 401 | AR | LNCaP | Prostate | GSM27169 | 6334 | 87051 |
| 958.5424 | 460 | 869.021 | 454 | 773.2506 | 570 | 844.4529 | 771 | AR | LNCaP | Prostate | GSM27169 | 29541 | 87052 |
| 93.08529 | 22 | 98.31054 | 25 | 24.57387 | 16 | 125.2412 | 39 | RNF2 | None | None | GSM28058 | 1239 | 87070 |
| 994.8984 | 348 | 1021.815 | 448 | 753.6175 | 357 | 708.07 | 357 | BCOR | None | None | GSM28058 | 16425 | 87073 |
| 1141.428 | 724 | 1193.389 | 929 | 813.1128 | 693 | 772.6133 | 723 | KDM2B | None | None | GSM28058 | 29865 | 87074 |
| 534.8084 | 59 | 690.7498 | 75 | 356.3241 | 61 | 483.6941 | 77 | BRD4 | LREX' | Prostate | GSM27715 | 1034 | 87081 |
| 588.0172 | 275 | 722.7126 | 359 | 378.5693 | 349 | 354.0553 | 369 | SMC1A | None | Foreskin | GSM22746 | 34745 | 87092 |
| 664.4353 | 301 | 739.3164 | 397 | 444.1258 | 388 | 494.6262 | 438 | CTCF | SH-SY5Y | Bone Marr | GSM28709 | 38269 | 87100 |
| 709.5979 | 429 | 743.6142 | 563 | 530.206 | 539 | 519.8754 | 601 | CTCF | SK-N-AS | Brain | GSM28709 | 59454 | 87102 |
| 81.2175 | 31 | 88.3764 | 36 | 163.5681 | 64 | 166.9371 | 69 | GRHL2 | MCF-7 | Breast | GSM29704 | 2729 | 87110 |
| 492.9977 | 221 | 544.2309 | 259 | 648.5801 | 377 | 654.0861 | 439 | GRHL2 | MCF-7 | Breast | GSM29704 | 26081 | 87111 |
| 500.2962 | 220 | 581.9325 | 264 | 612.2218 | 359 | 641.8173 | 413 | GRHL2 | MCF-7 | Breast | GSM29704 | 25460 | 87112 |

| | | | | | | | | | | | | | |
|---|---|---|---|---|---|---|---|---|---|---|---|---|---|
| 397.7586 | 155 | 315.6992 | 155 | 663.8758 | 289 | 722.9177 | 337 | ESR1 | MCF-7 | Breast | GSM29704 | 15096 | 87114 |
| 150.1292 | 48 | 65.19023 | 35 | 311.1376 | 100 | 436.3007 | 127 | ESR1 | MCF-7 | Breast | GSM29704 | 3500 | 87115 |
| 224.3658 | 82 | 228.2156 | 92 | 361.0239 | 151 | 472.7094 | 184 | ESR1 | MCF-7 | Breast | GSM29704 | 7301 | 87116 |
| 728.9005 | 305 | 713.6219 | 352 | 656.2907 | 487 | 640.9783 | 535 | GRHL2 | MCF-7 | Breast | GSM29704 | 33782 | 87118 |
| 384.5515 | 150 | 369.0075 | 164 | 489.8209 | 247 | 527.0761 | 274 | GRHL2 | MCF-7 | Breast | GSM29704 | 14611 | 87119 |
| 45.3957 | 36 | 59.7599 | 46 | 15.64711 | 39 | 38.28472 | 58 | EWSR1 | A673 | Muscle | GSM24720 | 6465 | 87143 |
| 509.966 | 210 | 570.9867 | 247 | 264.2098 | 236 | 244.7768 | 248 | SMARCB1 | G-401 | Malignant | GSM24096 | 21141 | 87159 |
| 401.169 | 212 | 489.5716 | 260 | 276.6916 | 278 | 265.8111 | 297 | SMARCC1 | G-401 | Malignant | GSM24096 | 28075 | 87160 |
| 323.1275 | 195 | 305.3575 | 215 | 317.1625 | 299 | 346.5387 | 335 | GATA2 | None | None | GSM28978 | 30126 | 87168 |
| 824.5804 | 214 | 868.528 | 253 | 776.4764 | 314 | 811.4688 | 394 | ESR1 | T47D | Mammary | GSM26445 | 13269 | 87200 |
| 146.014 | 69 | 182.1689 | 86 | 107.322 | 92 | 122.9827 | 105 | SMC3 | HeLa | Cervix | GSM28138 | 8431 | 87201 |
| 581.7584 | 224 | 683.6885 | 269 | 312.8006 | 251 | 261.6036 | 253 | SMC1A | JHU-06 | None | GSM22472 | 20109 | 87229 |
| 449.3406 | 115 | 728.4655 | 165 | 259.8043 | 126 | 372.0694 | 160 | SMC1A | None | None | GSM22472 | 7232 | 87230 |
| 260.1501 | 74 | 378.9531 | 100 | 182.3139 | 89 | 153.6766 | 88 | SMC1A | None | None | GSM22472 | 4847 | 87231 |
| 648.9909 | 224 | 823.8741 | 305 | 387.8778 | 261 | 455.2527 | 301 | SMC1A | JHU-06 | None | GSM22472 | 19491 | 87232 |
| 875.8266 | 371 | 898.8446 | 476 | 651.471 | 401 | 641.8867 | 445 | SMC1A | None | None | GSM22472 | 25950 | 87233 |
| 201.6345 | 32 | 305.0989 | 45 | 120.911 | 33 | 331.1117 | 62 | SMC1A | None | None | GSM22472 | 1078 | 87234 |
| 1008.025 | 195 | 1004.191 | 237 | 480.204 | 180 | 504.6781 | 197 | EZH1 | None | None | GSM23713 | 7262 | 87343 |
| 1115.688 | 517 | 1146.707 | 664 | 829.7303 | 502 | 786.6015 | 511 | EZH1 | None | None | GSM23713 | 18942 | 87346 |
| 1.137531 | 90 | 5.095055 | 132 | 19.43727 | 258 | 30.06245 | 311 | GATA6 | None | Endoderm | GSM20507 | 73187 | 87381 |
| 1101.553 | 737 | 1125.404 | 915 | 784.4 | 733 | 737.7128 | 757 | KDM2B | None | None | GSM28058 | 35475 | 87390 |
| 1126.697 | 727 | 1184.362 | 940 | 810.3147 | 719 | 767.6286 | 748 | KDM2B | None | None | GSM28058 | 31993 | 87391 |
| 195.196 | 52 | 145.0879 | 49 | 158.1292 | 67 | 183.5085 | 77 | FOXA2 | M.M. | None | GSM24306 | 3302 | 87392 |
| 114.3342 | 62 | 72.66877 | 57 | 45.68842 | 66 | 97.93104 | 96 | FOXA2 | M.M. | None | GSM24306 | 8233 | 87393 |
| 200.6732 | 249 | 122.0649 | 243 | 107.2515 | 338 | 116.4037 | 380 | FOXA2 | BJ | Skin | GSM24306 | 62944 | 87396 |
| 8.815971 | 52 | 6.378129 | 57 | 15.27419 | 105 | 41.05803 | 149 | FOXA2 | BJ | Skin | GSM24306 | 24070 | 87397 |
| 229.4293 | 64 | 152.0646 | 57 | 262.7898 | 98 | 265.3482 | 105 | FOXA2 | BJ | Skin | GSM24306 | 4138 | 87398 |
| 5.533849 | 120 | 0.613734 | 112 | 10.01972 | 249 | 39.1182 | 351 | GATA4 | BJ | Skin | GSM24306 | 80713 | 87399 |
| 279.3184 | 211 | 270.0315 | 236 | 164.2227 | 274 | 183.3185 | 310 | EP300 | None | None | GSM25501 | 37163 | 87438 |
| 240.465 | 40 | 216.9704 | 41 | 228.2103 | 53 | 389.2371 | 76 | ESR1 | None | Breast | GSM27971 | 1307 | 87458 |
| 956.2491 | 189 | 953.1148 | 211 | 806.2765 | 241 | 861.6365 | 300 | ESR1 | None | Breast | GSM27971 | 8520 | 87459 |
| 342.498 | 71 | 367.003 | 81 | 448.8325 | 115 | 632.3756 | 147 | ESR1 | None | Breast | GSM27971 | 3299 | 87466 |
| 1100.329 | 238 | 1108.268 | 300 | 914.9271 | 282 | 921.4756 | 330 | ESR1 | None | Breast | GSM27971 | 7764 | 87467 |
| 1162.147 | 633 | 1162.177 | 769 | 903.0272 | 694 | 880.3508 | 756 | ESR1 | None | Breast | GSM27971 | 24731 | 87472 |
| 1068.264 | 222 | 1038.564 | 253 | 889.4813 | 270 | 913.9486 | 328 | ESR1 | None | Breast | GSM27971 | 7947 | 87473 |
| 1301.623 | 633 | 1344.6 | 813 | 965.9725 | 577 | 927.4116 | 599 | MAPK1 | HeLa | Cervix | GSM22575 | 15485 | 87476 |
| 605.1727 | 459 | 673.0601 | 642 | 475.3509 | 648 | 460.0361 | 713 | CTCF | MCF-7 | Breast | GSM28779 | 85032 | 87487 |
| 652.1217 | 496 | 712.5051 | 682 | 501.5858 | 671 | 476.5922 | 725 | CTCF | MCF-7 | Breast | GSM28779 | 82398 | 87488 |
| 1197.255 | 550 | 1202.289 | 667 | 956.8989 | 607 | 952.1247 | 685 | ESR1 | None | Breast | GSM27971 | 17150 | 87528 |
| 1078.994 | 378 | 1068.062 | 448 | 884.4288 | 448 | 894.2827 | 519 | ESR1 | None | Breast | GSM27971 | 14539 | 87529 |
| 24.54137 | 16 | 9.606505 | 12 | 22.93985 | 24 | 10.13195 | 19 | HAND2 | CLB-Ga | None | GSM26643 | 2341 | 87612 |
| 3.978979 | 22 | 8.45155 | 32 | 29.02061 | 69 | 33.48877 | 79 | GATA3 | CLB-Ga | None | GSM26643 | 11009 | 87613 |
| 179.5985 | 39 | 277.3995 | 55 | 135.1302 | 47 | 274.7045 | 73 | MORC2 | hESC | None | GSM27898 | 1872 | 87623 |
| 296.337 | 68 | 305.1406 | 76 | 192.9321 | 77 | 241.91 | 92 | MORC2 | hESC | None | GSM27898 | 3573 | 87624 |
| 750.569 | 570 | 797.1838 | 754 | 629.6032 | 845 | 623.7185 | 953 | CTCF | MCF-7 | Breast | GSM27477 | 84982 | 87644 |
| 691.9887 | 425 | 721.8397 | 554 | 557.78 | 600 | 553.5442 | 677 | CTCF | MCF-7 | Breast | GSM27477 | 63111 | 87645 |
| 682.8561 | 369 | 729.2857 | 502 | 483.6751 | 487 | 479.4399 | 528 | CTCF | MCF-7 | Breast | GSM27477 | 54704 | 87646 |
| 734.4895 | 531 | 773.2746 | 696 | 550.0552 | 679 | 543.3867 | 765 | CTCF | MCF-7 | Breast | GSM27477 | 77969 | 87647 |
| 26.70403 | 16 | 20.51166 | 16 | 24.67071 | 24 | 65.82416 | 40 | NR3C1 | MCF-10A | Breast | GSM27353 | 2357 | 87649 |
| 810.8928 | 688 | 858.5725 | 891 | 646.7451 | 944 | 641.6482 | 1068 | CTCF | MCF-7 | Breast | GSM27477 | 92864 | 87650 |
| 782.9402 | 571 | 828.6932 | 751 | 656.7377 | 829 | 630.2493 | 898 | CTCF | MCF-7 | Breast | GSM27477 | 73300 | 87651 |
| 179.5108 | 133 | 311.138 | 190 | 173.8572 | 205 | 303.9503 | 275 | NR3C1 | MCF-10A | Breast | GSM27353 | 22554 | 87652 |
| 1156.25 | 418 | 1178.366 | 539 | 877.0133 | 408 | 822.5053 | 402 | MYC | IMR-5 | None | GSM20820 | 14194 | 87653 |
| 1153.449 | 437 | 1159.129 | 541 | 863.2637 | 422 | 806.3896 | 412 | MYC | IMR-5 | None | GSM20820 | 15587 | 87654 |
| 61.58018 | 40 | 91.32863 | 54 | 85.26601 | 71 | 165.0523 | 102 | NR3C1 | MCF-10A | Breast | GSM27353 | 6079 | 87678 |
| 36.21146 | 13 | 21.26862 | 11 | 34.93921 | 19 | 107.4182 | 36 | NR3C1 | MCF-10A | Breast | GSM27353 | 1226 | 87679 |
| 12.23692 | 11 | 5.442424 | 9 | 9.106749 | 16 | 30.03265 | 28 | NR3C1 | MCF-10A | Breast | GSM27353 | 2357 | 87680 |
| 197.9346 | 145 | 320.7021 | 201 | 193.9709 | 224 | 257.0705 | 270 | NR3C1 | MCF-10A | Breast | GSM27353 | 24331 | 87683 |
| 40.92819 | 19 | 32.34931 | 19 | 41.67839 | 29 | 67.66775 | 39 | NR3C1 | MCF-10A | Breast | GSM27353 | 2217 | 87684 |
| 226.4761 | 82 | 226.0465 | 91 | 497.7726 | 175 | 849.6613 | 253 | ESR1 | MCF-7 | Breast | GSM29153 | 6885 | 87688 |
| 85.45884 | 33 | 121.4291 | 44 | 262.7078 | 85 | 789.3203 | 164 | ESR1 | MCF-7 | Breast | GSM29153 | 3030 | 87689 |
| 579.4646 | 86 | 958.2153 | 127 | 322.1064 | 84 | 442.9316 | 106 | CHD1 | K562 | Bone Marr | GSM83098 | 2476 | 8769 |
| 1211.184 | 375 | 1238.557 | 480 | 924.3025 | 345 | 877.2214 | 343 | FGFR1 | None | None | GSM24391 | 9120 | 87695 |
| 1163.191 | 313 | 1203.693 | 420 | 892.8157 | 291 | 865.6674 | 308 | CHD1 | K562 | Bone Marr | GSM83098 | 8281 | 8770 |
| 386.718 | 59 | 404.8139 | 66 | 184.1526 | 53 | 176.6162 | 55 | CHD7 | K562 | Bone Marr | GSM83099 | 1752 | 8771 |
| 895.3049 | 141 | 1108.89 | 185 | 596.5006 | 154 | 635.9093 | 169 | SETDB1 | K562 | Bone Marr | GSM83099 | 4516 | 8772 |
| 1233.469 | 556 | 1291.64 | 741 | 924.8749 | 502 | 885.8154 | 517 | ARID2 | TTC1240 | Malignant | GSM24959 | 13117 | 87728 |
| 1150.869 | 611 | 1190.538 | 782 | 836.7769 | 568 | 810.0798 | 605 | ARID2 | TTC1240 | Malignant | GSM24959 | 21025 | 87729 |
| 1127.722 | 179 | 1130.04 | 221 | 744.2669 | 183 | 541.8747 | 164 | SMARCA4 | G-401 | Malignant | GSM24959 | 4955 | 87730 |
| 18.03653 | 99 | 21.42264 | 122 | 8.010396 | 157 | 9.216766 | 179 | NKX2-1 | hESC | None | GSM26655 | 51889 | 87737 |
| -0.045375 | 0 | 0 | 1 | 0.080798 | 3 | 0.20505 | 4 | NKX2-1 | hESC | None | GSM26655 | 1454 | 87738 |
| 696.0831 | 101 | 1088.426 | 144 | 399.0779 | 100 | 508.5823 | 121 | HDAC1 | K562 | Bone Marr | GSM83099 | 2795 | 8774 |

| | | | | | | | | | | | | | |
|---|---|---|---|---|---|---|---|---|---|---|---|---|---|
| 227.4759 | 69 | 219.35 | 75 | 419.5131 | 136 | 791.9718 | 202 | AHR | MCF-7 | Breast | GSM24067 | 5103 | 87747 |
| 1156.904 | 584 | 1194.584 | 752 | 857.0298 | 563 | 828.8464 | 596 | HDAC1 | K562 | Bone Marr | GSM83099 | 20901 | 8775 |
| 440.7122 | 78 | 774.3924 | 119 | 272.1341 | 83 | 364.7427 | 103 | HDAC2 | K562 | Bone Marr | GSM83099 | 2953 | 8776 |
| 1066.487 | 215 | 1116.051 | 297 | 838.9756 | 233 | 838.8665 | 254 | HDAC2 | K562 | Bone Marr | GSM83099 | 7638 | 8777 |
| 412.6295 | 57 | 539.3877 | 73 | 160.8052 | 45 | 251.0679 | 61 | HDAC6 | K562 | Bone Marr | GSM83099 | 1450 | 8778 |
| 962.0011 | 199 | 847.7611 | 204 | 797.9107 | 252 | 827.0373 | 299 | FOXA1 | None | Breast | GSM27971 | 9296 | 87788 |
| 275.4098 | 58 | 255.4055 | 61 | 134.1156 | 56 | 369.2018 | 101 | AR | None | Breast | GSM27971 | 2750 | 87789 |
| 411.5162 | 54 | 487.0023 | 65 | 180.3588 | 45 | 287.9079 | 62 | HDAC6 | K562 | Bone Marr | GSM83099 | 1262 | 8779 |
| 769.972 | 157 | 533.0801 | 141 | 872.5403 | 238 | 883.7638 | 281 | AR | None | Breast | GSM27971 | 7263 | 87790 |
| 1018.628 | 191 | 1019.412 | 237 | 891.0097 | 266 | 923.5138 | 336 | AR | None | Breast | GSM27971 | 7790 | 87791 |
| 253.4785 | 46 | 234.0778 | 48 | 234.9734 | 61 | 484.7784 | 97 | AR | None | Breast | GSM27971 | 1774 | 87793 |
| 1012.772 | 386 | 1016.401 | 477 | 861.4408 | 514 | 852.6045 | 572 | AR | None | Breast | GSM27971 | 18682 | 87795 |
| 175.1013 | 33 | 166.0084 | 35 | 156.2815 | 43 | 303.4115 | 66 | AR | None | Breast | GSM27971 | 1339 | 87796 |
| 351.2187 | 56 | 304.3705 | 56 | 357.1693 | 77 | 426.8235 | 90 | AR | None | Breast | GSM27971 | 1744 | 87797 |
| 80.21107 | 26 | 139.6952 | 39 | 38.40976 | 27 | 54.20425 | 34 | CBX3 | K562 | Bone Marr | GSM83099 | 2182 | 8780 |
| 928.5727 | 618 | 861.5911 | 657 | 753.2294 | 811 | 775.7694 | 972 | FOXA1 | 22RV1 | Prostate | GSM22771 | 51790 | 87801 |
| 943.3587 | 348 | 889.6699 | 374 | 773.2915 | 443 | 848.3467 | 612 | HNF4G | 22RV1 | Prostate | GSM22771 | 21019 | 87802 |
| 1080.798 | 150 | 1152.712 | 186 | 412.9696 | 119 | 591.0535 | 153 | KDM5C | K562 | Bone Marr | GSM83100 | 3961 | 8781 |
| 29.27328 | 27 | 34.41997 | 33 | 12.92562 | 33 | 24.84504 | 45 | CREBBP | K562 | Bone Marr | GSM83100 | 5809 | 8782 |
| 185.6831 | 112 | 238.4168 | 141 | 183.9777 | 172 | 199.4433 | 192 | KDM1A | K562 | Bone Marr | GSM83100 | 16503 | 8783 |
| 23.09363 | 9 | 101.3427 | 23 | 35.66585 | 17 | 194.4578 | 45 | HDAC2 | hESC | None | GSM17066 | 1015 | 87832 |
| 594.749 | 213 | 788.7369 | 272 | 320.5973 | 236 | 318.0198 | 254 | ZEB1 | MDA-231 | Mammary | GSM23606 | 17717 | 87839 |
| 177.8399 | 61 | 227.3668 | 77 | 96.68107 | 68 | 157.5292 | 91 | EP300 | K562 | Bone Marr | GSM83100 | 5353 | 8787 |
| 483.7087 | 64 | 734.3611 | 90 | 222.4339 | 55 | 382.3684 | 79 | KAT2B | K562 | Bone Marr | GSM83100 | 1525 | 8788 |
| 451.26 | 68 | 671.1468 | 94 | 232.0352 | 63 | 252.7704 | 70 | PHIP | HCT-116 | HCT116 | GSM27113 | 1839 | 87884 |
| 198.5708 | 35 | 364.0398 | 55 | 38.3426 | 20 | 154.4419 | 44 | PHIP | HCT-116 | HCT116 | GSM27113 | 1186 | 87886 |
| 1289.696 | 772 | 1395.944 | 1044 | 932.9223 | 725 | 884.7869 | 744 | PHF8 | K562 | Bone Marr | GSM83100 | 22999 | 8789 |
| 1260.202 | 456 | 1273.532 | 570 | 957.9339 | 411 | 914.2125 | 420 | RPE | None | Retina | GSM28650 | 10190 | 87891 |
| 1234.835 | 333 | 1275.862 | 436 | 973.2396 | 321 | 925.3522 | 315 | RPE | None | Retina | GSM28650 | 7132 | 87892 |
| 1095.926 | 534 | 1175.837 | 751 | 833.5751 | 569 | 784.5859 | 571 | ZBTB7A | K562 | Bone Marr | GSM27715 | 23925 | 87893 |
| 1079.718 | 595 | 1120.092 | 772 | 792.7106 | 607 | 750.8006 | 623 | ZBTB7A | K562 | Bone Marr | GSM27715 | 29880 | 87897 |
| 1285.955 | 796 | 1373.275 | 1058 | 919.6048 | 750 | 873.3099 | 769 | PHF8 | K562 | Bone Marr | GSM83100 | 25078 | 8790 |
| 1202.308 | 636 | 1278.011 | 862 | 897.0666 | 617 | 859.6216 | 648 | KDM5B | K562 | Bone Marr | GSM83101 | 20649 | 8791 |
| 400.2292 | 177 | 422.0812 | 203 | 346.0708 | 249 | 476.1799 | 307 | STAT3 | H358 | Lung | GSM27528 | 19914 | 87919 |
| 1151.302 | 220 | 1155.246 | 271 | 775.1803 | 200 | 719.2145 | 204 | KDM5B | K562 | Bone Marr | GSM83101 | 5898 | 8792 |
| 284.5658 | 58 | 353.0533 | 72 | 149.8067 | 58 | 229.8992 | 77 | STAT1 | H358 | Lung | GSM27528 | 2660 | 87922 |
| 947.2259 | 136 | 1125.992 | 164 | 619.228 | 145 | 593.5527 | 150 | RBBP5 | K562 | Bone Marr | GSM83101 | 3761 | 8793 |
| 1220.122 | 193 | 1206.607 | 228 | 902.9825 | 186 | 987.4089 | 223 | ZIC2 | None | Peritoneal | GSM27380 | 4031 | 87938 |
| 1234.962 | 397 | 1254.346 | 507 | 945.5272 | 363 | 911.779 | 376 | GLI2 | THP-1 | None | GSM27790 | 8983 | 87939 |
| 1153.863 | 174 | 1178.056 | 226 | 753.5769 | 175 | 677.9826 | 175 | RBBP5 | K562 | Bone Marr | GSM83101 | 4526 | 8794 |
| 913.1596 | 104 | 1169.791 | 131 | 487.001 | 94 | 443.1911 | 94 | GLI2 | THP-1 | None | GSM27790 | 1754 | 87941 |
| 1229.554 | 433 | 1258.589 | 550 | 931.6669 | 390 | 898.2024 | 404 | GLI2 | THP-1 | None | GSM27790 | 9987 | 87942 |
| 91.38345 | 27 | 103.5104 | 32 | 124.9446 | 46 | 606.6777 | 116 | ESR1 | MCF-7 | Breast | GSM29153 | 1981 | 87947 |
| 212.3982 | 70 | 241.5455 | 83 | 485.3331 | 154 | 897.2542 | 231 | ESR1 | MCF-7 | Breast | GSM29153 | 5621 | 87948 |
| 1116.23 | 551 | 1148.851 | 697 | 818.5983 | 520 | 799.2476 | 564 | RBBP5 | K562 | Bone Marr | GSM83101 | 20291 | 8795 |
| 118.2287 | 56 | 195.6672 | 80 | 48.85712 | 58 | 81.39692 | 77 | REST | K562 | Bone Marr | GSM83101 | 6125 | 8796 |
| 143.0667 | 34 | 182.6658 | 43 | 37.64567 | 25 | 179.3852 | 58 | RNF2 | K562 | Bone Marr | GSM83101 | 1934 | 8798 |
| 1205.913 | 151 | 1217.117 | 193 | 719.6938 | 149 | 765.6819 | 163 | SAP30 | K562 | Bone Marr | GSM83101 | 3283 | 8799 |
| 1247.352 | 538 | 1317.129 | 735 | 948.8905 | 516 | 900.0109 | 513 | SAP30 | K562 | Bone Marr | GSM83101 | 13811 | 8800 |
| 1.929053 | 5 | 0.475059 | 4 | 4.497719 | 12 | 3.344134 | 12 | SIRT1 | K562 | None | GSM24748 | 2165 | 88010 |
| 2.93568 | 8 | 1.509224 | 8 | 5.871575 | 18 | 11.88536 | 25 | SIRT1 | K562 | None | GSM24748 | 3504 | 88011 |
| 1149.76 | 601 | 1197.846 | 783 | 848.8235 | 582 | 794.5512 | 578 | FGFR1 | None | None | GSM24391 | 21771 | 88045 |
| 874.9617 | 398 | 836.9445 | 441 | 707.5806 | 509 | 800.8181 | 731 | AR | LNCaP | Prostate | GSM22771 | 31026 | 88049 |
| 211.0328 | 120 | 190.151 | 129 | 126.571 | 150 | 203.2173 | 195 | MED1 | hMSC-TERT | None | GSM28029 | 16404 | 88055 |
| 52.15505 | 34 | 53.87323 | 39 | 42.7611 | 49 | 78.84861 | 68 | MED1 | hMSC-TERT | None | GSM28029 | 5349 | 88056 |
| 18.63349 | 15 | 18.45191 | 17 | 13.35113 | 21 | 15.13896 | 24 | MED1 | hMSC-TERT | None | GSM28029 | 2788 | 88057 |
| 909.0226 | 163 | 1064.681 | 223 | 530.709 | 169 | 566.6429 | 186 | CHD1 | H1 | Embryo | GSM83102 | 6211 | 8806 |
| 17.6995 | 22 | 31.78621 | 32 | 47.89538 | 53 | 78.34382 | 70 | TCF4 | SH-SY5Y | None | GSM25458 | 5719 | 88062 |
| 14.86219 | 207 | 2.937869 | 198 | 3.015799 | 320 | 4.999291 | 378 | FOXA2 | BJ | Skin | GSM29775 | 124404 | 88065 |
| 71.75166 | 194 | 38.27889 | 192 | 45.45 | 299 | 57.10986 | 348 | FOXA2 | BJ | Skin | GSM29775 | 70852 | 88066 |
| 1095.91 | 378 | 1137.235 | 509 | 846.8513 | 385 | 801.0936 | 386 | CHD1 | H1 | Embryo | GSM83102 | 13314 | 8807 |
| 220.7742 | 93 | 342.9602 | 129 | 110.5818 | 104 | 163.9531 | 132 | CHD7 | H1 | Embryo | GSM83102 | 10102 | 8808 |
| 13.18615 | 7 | 67.87256 | 19 | 45.31427 | 20 | 122.3454 | 36 | EZH2 | H1 | Embryo | GSM83102 | 1068 | 8809 |
| 1083.975 | 547 | 1114.27 | 699 | 798.9458 | 547 | 756.6233 | 557 | HDAC2 | H1 | Embryo | GSM83102 | 24975 | 8810 |
| 1230.87 | 185 | 1209.807 | 218 | 825.6548 | 171 | 802.9815 | 178 | KDM5A | H1 | Embryo | GSM83103 | 3700 | 8812 |
| 123.9817 | 40 | 151.0943 | 49 | 46.95664 | 36 | 86.07852 | 51 | MKL2 | A673 | Muscle | GSM24366 | 2557 | 88131 |
| 696.979 | 380 | 739.3719 | 512 | 559.3805 | 535 | 551.7129 | 597 | CTCF | MCF-7 | Breast | GSM27477 | 55825 | 88132 |
| 0.246411 | 1 | 0.699419 | 2 | 3.370906 | 6 | 3.9135 | 7 | SMARCC1 | SK-N-MC | Brain | GSM24721 | 1028 | 88136 |
| 14.63481 | 9 | 8.698659 | 8 | 0.359004 | 4 | 6.386294 | 11 | SMARCC1 | SK-N-MC | Brain | GSM24721 | 1470 | 88137 |
| 4.741041 | 6 | 4.79988 | 7 | 2.37127 | 8 | 44.0299 | 28 | SMARCA2 | SK-N-MC | Brain | GSM24721 | 1716 | 88138 |
| 53.20457 | 15 | 94.34741 | 23 | 36.00872 | 18 | 80.28834 | 29 | KDM5B | H1 | Embryo | GSM83103 | 1131 | 8814 |
| 35.12215 | 12 | 48.92111 | 16 | 58.689 | 23 | 114.9211 | 35 | EP300 | 786-O | Kidney | GSM22934 | 1050 | 88140 |

| | | | | | | | | | | | | | |
|---|---|---|---|---|---|---|---|---|---|---|---|---|---|
| 1204.735 | 748 | 1282.074 | 992 | 840.6965 | 683 | 803.4308 | 717 | KDM4A | H1 | Embryo | GSM83103 | 28179 | 8816 |
| 397.8128 | 96 | 618.6638 | 135 | 216.2674 | 101 | 216.2023 | 108 | EP300 | H1 | Embryo | GSM83103 | 5591 | 8817 |
| 1307.608 | 716 | 1385.597 | 955 | 941.8912 | 638 | 915.1109 | 686 | PHF8 | H1 | Embryo | GSM83103 | 19149 | 8818 |
| 1106.598 | 387 | 1096.86 | 456 | 809.5424 | 347 | 794.3744 | 378 | RBBP5 | H1 | Embryo | GSM83103 | 13637 | 8819 |
| 1110.644 | 668 | 1137.082 | 841 | 782.1885 | 638 | 747.3106 | 667 | RBBP5 | H1 | Embryo | GSM83103 | 31727 | 8820 |
| 1141.296 | 495 | 1196.138 | 667 | 865.9607 | 493 | 832.6675 | 516 | SAP30 | H1 | Embryo | GSM83104 | 17660 | 8821 |
| 791.0347 | 143 | 1057.817 | 197 | 378.7232 | 134 | 379.8379 | 143 | SUZ12 | H1 | Embryo | GSM83104 | 5506 | 8823 |
| 688.5966 | 488 | 740.7117 | 661 | 551.6755 | 689 | 528.3057 | 746 | CTCF | MCF-7 | Breast | GSM28779 | 76385 | 88275 |
| 1008.942 | 374 | 898.9164 | 344 | 826.9782 | 452 | 868.2447 | 576 | HOXB13 | LNCaP | Prostate | GSM25372 | 17452 | 88278 |
| 1091.764 | 284 | 1004.492 | 272 | 920.925 | 346 | 969.0359 | 449 | AR | 22RV1 | Prostate | GSM25372 | 9465 | 88282 |
| 1076.191 | 287 | 977.6776 | 270 | 901.6771 | 355 | 943.2518 | 450 | AR | 22RV1 | Prostate | GSM25372 | 10256 | 88283 |
| 1131.536 | 277 | 1176.117 | 376 | 882.4368 | 272 | 863.3886 | 293 | MYCN | T.T | None | GSM25723 | 8020 | 88296 |
| 967.7727 | 251 | 990.6923 | 324 | 616.3322 | 252 | 647.347 | 276 | MYCN | NGP | None | GSM21135 | 12046 | 88298 |
| 958.0799 | 249 | 987.6617 | 330 | 546.133 | 243 | 504.8171 | 251 | MYCN | KELLY | None | GSM21135 | 12799 | 88303 |
| 801.9136 | 496 | 807.8446 | 613 | 693.2188 | 740 | 711.4235 | 882 | ESR1 | MCF-7 | Breast | GSM26489 | 53601 | 88335 |
| 840.6305 | 253 | 859.638 | 326 | 780.3725 | 427 | 785.8809 | 495 | ESR1 | MCF-7 | Breast | GSM26489 | 19484 | 88336 |
| 268.9787 | 135 | 367.0803 | 175 | 182.1815 | 176 | 212.0307 | 202 | SMC3 | HeLa | Cervix | GSM28138 | 17779 | 88343 |
| 589.1104 | 245 | 692.3622 | 295 | 683.0193 | 427 | 715.3625 | 532 | ESR1 | MCF-7 | Breast | GSM26489 | 26465 | 88345 |
| 191.2458 | 101 | 295.8381 | 139 | 139.1518 | 136 | 129.4663 | 143 | SMC3 | HeLa | Cervix | GSM28138 | 13670 | 88347 |
| 468.6384 | 186 | 509.492 | 216 | 710.4789 | 345 | 744.4697 | 432 | ESR1 | MCF-7 | Breast | GSM26489 | 18950 | 88349 |
| 1098.084 | 319 | 1115.618 | 401 | 866.0582 | 315 | 861.7852 | 354 | ZFX | C4-2B | Prostate | GSM27417 | 8556 | 88351 |
| 6.293889 | 37 | 41.69697 | 76 | 56.25351 | 121 | 77.97589 | 147 | TAL1 | K562 | Bone Marr | GSM28771 | 18599 | 88353 |
| 41.36287 | 50 | 69.03429 | 70 | 97.22113 | 112 | 118.7571 | 131 | GATA1 | K562 | Bone Marr | GSM28771 | 12372 | 88355 |
| 890.1519 | 212 | 891.6191 | 242 | 820.9355 | 329 | 851.2509 | 407 | ESR1 | MCF-7 | Breast | GSM27476 | 12459 | 88372 |
| 960.1391 | 375 | 908.6801 | 402 | 818.6892 | 509 | 913.098 | 732 | AR | LNCaP | Prostate | GSM24808 | 21517 | 88420 |
| 1044.414 | 448 | 937.2269 | 421 | 872.9239 | 569 | 922.6 | 730 | AR | LNCaP | Prostate | GSM24808 | 20990 | 88423 |
| 1019.58 | 347 | 927.9971 | 334 | 835.7355 | 422 | 892.717 | 559 | ARID1A | LNCaP | Prostate | GSM24808 | 16106 | 88426 |
| 1117.037 | 272 | 990.7685 | 235 | 941.0604 | 337 | 976.0434 | 417 | ARID1A | LNCaP | Prostate | GSM24808 | 8634 | 88427 |
| 22.1342 | 32 | 22.22043 | 37 | 52.80315 | 73 | 120.2783 | 109 | GATA2 | None | None | GSM28176 | 9150 | 88445 |
| 1118.507 | 423 | 1125.362 | 517 | 843.0675 | 400 | 804.6031 | 408 | NIPBL | HCT-116 | Colon | GSM28096 | 13164 | 88450 |
| 282.7582 | 58 | 494.8058 | 88 | 262.5559 | 78 | 234.1989 | 78 | BRD4 | MV4;11 | Blood | GSM23450 | 2664 | 88453 |
| 934.0485 | 316 | 890.3325 | 342 | 746.8642 | 377 | 828.992 | 534 | HNF4G | 22RV1 | Prostate | GSM22771 | 18881 | 88463 |
| 972.0292 | 515 | 917.7156 | 553 | 875.6194 | 783 | 904.7159 | 944 | HOXB13 | VCaP | Prostate | GSM25372 | 32612 | 88495 |
| 966.2684 | 517 | 908.8965 | 555 | 884.3478 | 813 | 905.9767 | 969 | HOXB13 | VCaP | Prostate | GSM25372 | 33791 | 88496 |
| 985.8433 | 560 | 931.8318 | 609 | 889.0932 | 853 | 905.4043 | 1007 | HOXB13 | VCaP | Prostate | GSM25372 | 36258 | 88497 |
| 901.6024 | 880 | 791.7444 | 916 | 734.6213 | 1242 | 765.5124 | 1487 | FOXA1 | VCaP | Prostate | GSM25372 | 123343 | 88498 |
| 991.9087 | 576 | 917.6526 | 605 | 878.424 | 851 | 891.6657 | 1003 | HOXB13 | VCaP | Prostate | GSM25372 | 37392 | 88500 |
| 953.2476 | 606 | 905.387 | 665 | 842.3525 | 898 | 867.2483 | 1080 | HOXB13 | VCaP | Prostate | GSM25372 | 45976 | 88501 |
| 124.9576 | 48 | 166.7392 | 62 | 183.4347 | 86 | 91.80149 | 67 | SKI | HL-60 | None | GSM28711 | 4810 | 88503 |
| 1173.688 | 506 | 1200.552 | 647 | 874.7051 | 467 | 849.0905 | 495 | PAF1 | DLD-1 | None | GSM27690 | 13955 | 88509 |
| 1199.211 | 555 | 1212.957 | 686 | 885.8123 | 501 | 857.4914 | 532 | PAF1 | DLD-1 | None | GSM27690 | 15170 | 88511 |
| 1174.07 | 499 | 1209.996 | 648 | 883.0143 | 464 | 854.0221 | 492 | PAF1 | DLD-1 | None | GSM27690 | 13667 | 88512 |
| 37.33358 | 31 | 34.32201 | 34 | 37.2332 | 49 | 40.62865 | 55 | T | MCF-7 | Breast | GSM26712 | 5636 | 88587 |
| 32.15472 | 20 | 16.92667 | 17 | 11.90944 | 21 | 43.15286 | 38 | T | MCF-7 | Breast | GSM26712 | 2996 | 88588 |
| 32.94352 | 23 | 35.71427 | 27 | 22.53073 | 31 | 37.26649 | 41 | T | MCF-7 | Breast | GSM26712 | 3749 | 88590 |
| 157.4649 | 92 | 195.7586 | 114 | 152.6452 | 138 | 137.0769 | 143 | T | MCF-7 | Breast | GSM26712 | 12189 | 88591 |
| 61.27603 | 23 | 41.33329 | 21 | 23.60538 | 22 | 34.41938 | 28 | T | MCF-7 | Breast | GSM26712 | 2087 | 88593 |
| 9.024154 | 19 | 7.739357 | 21 | 8.436939 | 32 | 9.975112 | 37 | BCL6 | OCI-Ly1 | Bone Marr | GSM27542 | 6288 | 88598 |
| 9.584779 | 18 | 11.09646 | 22 | 18.44685 | 38 | 17.74498 | 41 | BCL6 | OCI-Ly1 | Bone Marr | GSM27542 | 5647 | 88599 |
| 714.3334 | 103 | 802.0807 | 120 | 355.0623 | 94 | 397.9563 | 106 | MYC | IMEC | Breast | GSM23019 | 2772 | 88605 |
| 1186.81 | 192 | 1223.373 | 262 | 712.2022 | 169 | 646.6012 | 170 | YAP1 | OVCAR5 | ovarian | GSM13776 | 4219 | 88621 |
| 828.8594 | 604 | 856.242 | 765 | 590.5716 | 686 | 557.9488 | 728 | EP300 | None | None | GSM25502 | 66266 | 88622 |
| 65.46503 | 41 | 95.48484 | 55 | 37.14249 | 51 | 32.90131 | 53 | ZBTB16 | KG1 | Bone Marr | GSM29473 | 6239 | 88633 |
| 56.42436 | 36 | 67.12979 | 44 | 38.05084 | 48 | 54.7947 | 60 | ZBTB16 | KG1 | Bone Marr | GSM29473 | 5601 | 88634 |
| 753.0796 | 567 | 747.9695 | 681 | 585.5042 | 748 | 575.0201 | 835 | SS18 | TTC1240 | Malignant | GSM24097 | 74469 | 88639 |
| 1031.969 | 264 | 1061.073 | 344 | 776.1795 | 262 | 627.4096 | 252 | SS18 | TTC1240 | Malignant | GSM24097 | 10282 | 88643 |
| 844.121 | 361 | 835.3602 | 425 | 547.5083 | 365 | 405.7182 | 350 | SMARCB1 | None | epithelioid | GSM24097 | 25576 | 88645 |
| 828.5223 | 322 | 835.2655 | 402 | 564.0965 | 363 | 622.0885 | 423 | ERG | JHU-06 | None | GSM23944 | 28324 | 88664 |
| 586.7465 | 216 | 666.8679 | 256 | 465.2374 | 288 | 519.2159 | 325 | STAT3 | HCC1143 | None | GSM22780 | 20875 | 88680 |
| 71.15333 | 19 | 198.6259 | 38 | 72.52406 | 28 | 190.7661 | 50 | TRIM28 | LNCaP | Prostate | GSM24808 | 1361 | 88682 |
| 931.1224 | 505 | 795.6598 | 453 | 768.4717 | 667 | 834.1452 | 881 | TLE3 | LNCaP | Prostate | GSM24808 | 36803 | 88683 |
| 871.655 | 514 | 758.5915 | 488 | 736.9518 | 727 | 789.5266 | 932 | TLE3 | LNCaP | Prostate | GSM24808 | 46313 | 88684 |
| 99.3286 | 23 | 140.6909 | 31 | 77.68978 | 29 | 296.0712 | 64 | TRIM28 | LNCaP | Prostate | GSM24808 | 1360 | 88688 |
| 922.445 | 509 | 813.1604 | 485 | 778.4891 | 705 | 833.5611 | 906 | TLE3 | LNCaP | Prostate | GSM24808 | 38208 | 88689 |
| 901.8044 | 513 | 772.5566 | 464 | 759.5763 | 710 | 815.1607 | 916 | TLE3 | LNCaP | Prostate | GSM24808 | 41603 | 88690 |
| 1059.22 | 463 | 1075.803 | 579 | 779.7061 | 439 | 729.9734 | 438 | BRD4 | LCL | Blood | GSM26435 | 18089 | 88724 |
| 275.1352 | 186 | 287.2917 | 214 | 172.7807 | 243 | 199.3733 | 278 | MED1 | hMSC-TERT | None | GSM28029 | 29857 | 88736 |
| 995.848 | 398 | 1023.347 | 520 | 774.1711 | 442 | 780.8905 | 513 | GATA4 | A549 | Lung | GSM22557 | 20687 | 88777 |
| 549.0846 | 76 | 681.6542 | 94 | 500.7319 | 96 | 390.8504 | 88 | INO80 | HepG2 | Liver | GSM28772 | 1773 | 88784 |
| 158.5027 | 38 | 234.8509 | 52 | 117.9618 | 45 | 90.10938 | 42 | INO80 | HepG2 | Liver | GSM28772 | 1729 | 88796 |
| 1087.21 | 379 | 1122.38 | 501 | 830.9428 | 381 | 795.4835 | 394 | XCL1 | C6661 | None | GSM25231 | 14733 | 88801 |
| 583.8341 | 338 | 691.4629 | 459 | 405.2267 | 457 | 480.7979 | 528 | CTCF | NB69 | None | GSM28710 | 54311 | 88815 |

| | | | | | | | | | | | | | |
|---|---|---|---|---|---|---|---|---|---|---|---|---|---|
| 116.3608 | 52 | 236.6597 | 83 | 120.3793 | 80 | 79.41469 | 72 | PRMT1 | None | None | GSM29770 | 6065 | 88884 |
| 501.0186 | 245 | 591.9405 | 296 | 315.0874 | 308 | 295.1352 | 326 | PRMT1 | None | None | GSM29770 | 30919 | 88886 |
| 500.931 | 107 | 568.7701 | 126 | 186.9821 | 91 | 328.3665 | 128 | CHD4 | SCC9 | None | GSM25791 | 4999 | 88895 |
| 500.9934 | 99 | 514.6056 | 110 | 203.6536 | 86 | 242.0048 | 100 | CHD4 | SCC9 | None | GSM25791 | 4082 | 88896 |
| 553.6044 | 104 | 679.9661 | 128 | 278.1586 | 100 | 275.304 | 106 | CHD4 | SCC9 | None | GSM25791 | 4125 | 88897 |
| 719.4856 | 140 | 813.9233 | 164 | 283.5682 | 119 | 375.5903 | 146 | CHD4 | SCC9 | None | GSM25791 | 5365 | 88898 |
| 283.4305 | 53 | 259.9481 | 55 | 111.2131 | 44 | 125.8521 | 50 | PAF1 | HCT-116 | HCT116 | GSM25710 | 1888 | 88906 |
| 768.3653 | 604 | 815.5957 | 791 | 649.706 | 898 | 637.9879 | 1004 | CTCF | MCF-7 | Breast | GSM27477 | 88037 | 88917 |
| 373.0097 | 138 | 381.8922 | 155 | 187.6959 | 148 | 214.8588 | 169 | SMARCA4 | J-Lat A72 | None | GSM26768 | 11387 | 88920 |
| 100.9747 | 29 | 138.3607 | 38 | 81.87224 | 37 | 301.4118 | 78 | BRD4 | J-Lat A72 | None | GSM26768 | 1775 | 88924 |
| 427.1243 | 239 | 548.0315 | 300 | 307.1188 | 325 | 303.0375 | 351 | SMC1A | None | Blood | GSM25928 | 36158 | 88934 |
| 225.4654 | 114 | 290.9842 | 144 | 107.0737 | 129 | 199.515 | 178 | SMC1A | None | Blood | GSM25928 | 14663 | 88935 |
| -0.213649 | 0 | 0.10878 | 2 | 5.006928 | 10 | 3.868715 | 10 | SPI1 | None | Blood | GSM25928 | 1752 | 88936 |
| 961.8958 | 459 | 978.9859 | 575 | 711.2331 | 480 | 684.7992 | 511 | RELA | JHU-06 | None | GSM23944 | 28410 | 88945 |
| 1060.256 | 656 | 1100.3 | 844 | 762.3136 | 677 | 745.758 | 743 | RELA | JHU-06 | None | GSM23944 | 38980 | 88946 |
| 962.598 | 527 | 989.7318 | 670 | 704.8409 | 557 | 689.5946 | 609 | RELA | JHU-06 | None | GSM23944 | 34708 | 88947 |
| 1033.502 | 719 | 1122.019 | 977 | 748.7594 | 785 | 728.6757 | 860 | RELA | JHU-06 | None | GSM23944 | 50806 | 88948 |
| 48.40249 | 30 | 163.3916 | 61 | 37.16173 | 42 | 131.9134 | 78 | VDR | THP-1 | Blood | GSM23714 | 4606 | 88952 |
| 15.08607 | 10 | 38.78434 | 18 | 7.724807 | 12 | 49.41762 | 29 | VDR | THP-1 | Blood | GSM23714 | 1670 | 88954 |
| 375.7272 | 147 | 344.9344 | 157 | 178.7888 | 156 | 169.1516 | 165 | ARID1A | HCT-116 | HCT116 | GSM27197 | 13042 | 88976 |
| 124.3564 | 61 | 151.8716 | 75 | 97.40089 | 82 | 130.1124 | 100 | ARID1A | HCT-116 | HCT116 | GSM27197 | 6723 | 88981 |
| 150.5975 | 40 | 131.3716 | 41 | 134.4556 | 54 | 155.5803 | 62 | STAT3 | AGS | None | GSM27529 | 2503 | 89056 |
| 800.1882 | 287 | 761.3496 | 318 | 724.7412 | 468 | 767.6043 | 595 | ESR1 | MCF-7 | Breast | GSM27361 | 26159 | 89061 |
| 777.8476 | 267 | 694.4871 | 282 | 734.5138 | 456 | 764.4397 | 562 | ESR1 | MCF-7 | Breast | GSM27361 | 24598 | 89062 |
| 382.6623 | 90 | 396.457 | 101 | 796.3946 | 188 | 916.5868 | 215 | ESR1 | MCF-7 | Breast | GSM27361 | 4933 | 89063 |
| 768.961 | 240 | 753.5414 | 264 | 702.4041 | 348 | 738.3767 | 435 | JUN | MCF-7 | Breast | GSM27361 | 19650 | 89064 |
| 543.3423 | 187 | 532.2268 | 206 | 596.9679 | 286 | 713.9407 | 354 | JUN | MCF-7 | Breast | GSM27361 | 16588 | 89065 |
| 181.1643 | 58 | 224.4975 | 72 | 200.6546 | 89 | 295.4063 | 115 | JUN | MCF-7 | Breast | GSM27361 | 4626 | 89066 |
| 946.2345 | 213 | 932.5653 | 250 | 464.0793 | 208 | 339.6733 | 193 | EGR1 | HL-60 | Blood | GSM28364 | 10942 | 89075 |
| 982.8614 | 236 | 979.4289 | 286 | 584.7625 | 234 | 406.3816 | 211 | EGR1 | HL-60 | Blood | GSM28364 | 11271 | 89076 |
| 776.9027 | 465 | 725.1711 | 504 | 662.4641 | 683 | 662.2782 | 780 | FOXA1 | WS8 | None | GSM30964 | 52329 | 89243 |
| 788.6553 | 415 | 742.8818 | 454 | 681.7764 | 617 | 692.3777 | 725 | FOXA1 | WS8 | None | GSM30964 | 42523 | 89244 |
| 799.3126 | 376 | 759.0447 | 416 | 694.337 | 562 | 710.55 | 669 | FOXA1 | WS8 | None | GSM30964 | 35743 | 89245 |
| 797.7125 | 155 | 1001.064 | 195 | 495.0995 | 166 | 512.349 | 180 | TFAP2C | hESC | None | GSM29017 | 6080 | 89256 |
| 370.2394 | 72 | 388.5966 | 81 | 179.0838 | 68 | 152.8324 | 67 | BRD4 | ST1 | None | GSM24816 | 2949 | 89275 |
| 768.4744 | 142 | 1036.641 | 189 | 426.1949 | 143 | 375.8845 | 143 | BRD4 | ST1 | None | GSM24816 | 5366 | 89279 |
| 1095.873 | 208 | 1132.822 | 280 | 897.5375 | 226 | 678.1688 | 205 | BRD4 | KK1 | None | GSM24816 | 5645 | 89283 |
| 111.5405 | 98 | 174.7059 | 134 | 59.03664 | 126 | 120.6713 | 175 | CEBPB | None | Epididymis | GSM29290 | 19619 | 89292 |
| 402.7661 | 386 | 516.1877 | 481 | 311.5156 | 572 | 327.9445 | 637 | CEBPB | None | Epididymis | GSM29290 | 91120 | 89293 |
| 202.7785 | 282 | 235.5193 | 341 | 172.4946 | 445 | 171.2173 | 488 | CEBPB | None | Epididymis | GSM29290 | 80983 | 89295 |
| 491.7607 | 316 | 492.1818 | 358 | 323.7873 | 424 | 269.0983 | 435 | CEBPB | None | Epididymis | GSM29290 | 54256 | 89297 |
| 506.7287 | 159 | 442.5534 | 165 | 772.886 | 292 | 825.971 | 382 | ESR1 | MCF-7 | Breast | GSM27525 | 12809 | 89298 |
| 404.8541 | 139 | 380.5626 | 150 | 751.7862 | 274 | 815.1505 | 359 | ESR1 | MCF-7 | Breast | GSM27525 | 12296 | 89299 |
| 598.4329 | 212 | 535.1118 | 224 | 733.5373 | 379 | 774.4106 | 484 | ESR1 | MCF-7 | Breast | GSM27525 | 19923 | 89300 |
| 608.9079 | 187 | 605.3864 | 207 | 778.508 | 338 | 828.8265 | 439 | ESR1 | MCF-7 | Breast | GSM27525 | 14968 | 89301 |
| 945.9211 | 304 | 956.2704 | 385 | 566.8155 | 285 | 596.2364 | 313 | LDB1 | None | None | GSM33948 | 17225 | 89356 |
| 940.4772 | 375 | 951.8102 | 468 | 681.0866 | 374 | 645.2072 | 382 | LDB1 | None | None | GSM33948 | 23073 | 89357 |
| 866.9898 | 519 | 880.5849 | 644 | 630.879 | 576 | 584.8605 | 586 | BRD9 | EOL1 | None | GSM30943 | 45814 | 89358 |
| 700.1675 | 201 | 858.6169 | 255 | 383.5721 | 220 | 447.1825 | 253 | BRD9 | EOL1 | None | GSM30943 | 14730 | 89359 |
| 686.4753 | 566 | 693.3193 | 693 | 493.7141 | 709 | 480.6769 | 787 | SMARCA4 | EOL1 | None | GSM30943 | 91753 | 89360 |
| 1205.258 | 232 | 1225.216 | 298 | 827.756 | 196 | 920.6785 | 229 | RBP2 | THP-1 | Blood | GSM31119 | 4812 | 89369 |
| 779.4566 | 468 | 803.1897 | 596 | 557.5489 | 533 | 566.6347 | 622 | RAD21 | THP-1 | Blood | GSM31119 | 53634 | 89370 |
| 1228.923 | 398 | 1229.692 | 486 | 935.6552 | 358 | 887.9968 | 357 | RBP2 | THP-1 | Blood | GSM31119 | 8725 | 89371 |
| 1142.963 | 447 | 1157.912 | 562 | 847.4133 | 401 | 830.8035 | 435 | RBP2 | THP-1 | Blood | GSM31119 | 12630 | 89372 |
| 1208.619 | 496 | 1242.923 | 633 | 895.668 | 436 | 874.8466 | 470 | RBP2 | THP-1 | Blood | GSM31119 | 12595 | 89373 |
| 709.7524 | 390 | 759.9189 | 534 | 544.1798 | 503 | 478.9152 | 520 | CTCF | GM04025 | B-lymphoc | GSM32117 | 52261 | 89383 |
| 912.4611 | 183 | 1042.386 | 268 | 427.2883 | 173 | 502.4562 | 200 | EZH2 | LNCaP | Prostate | GSM28790 | 8037 | 89386 |
| 6.274139 | 55 | 2.761511 | 56 | 8.435227 | 107 | 27.05767 | 153 | TSC22D3 | MCF-10A | Breast | GSM26755 | 29253 | 89394 |
| 1078.195 | 536 | 1118.137 | 700 | 799.4655 | 520 | 743.3866 | 514 | BRD4 | K562 | Bone Marr | GSM30315 | 19885 | 89419 |
| 1164.815 | 495 | 1187.423 | 619 | 872.6285 | 458 | 816.1141 | 451 | BRD4 | K562 | Bone Marr | GSM30315 | 14066 | 89420 |
| 1088.594 | 510 | 1102.489 | 632 | 813.0362 | 490 | 760.6801 | 485 | BRD4 | K562 | Bone Marr | GSM30315 | 17732 | 89421 |
| 1156.004 | 569 | 1212.175 | 756 | 867.4129 | 549 | 822.1164 | 559 | BRD4 | K562 | Bone Marr | GSM30315 | 17966 | 89423 |
| 1205.256 | 537 | 1236.328 | 680 | 898.8129 | 486 | 861.01 | 504 | BRD4 | K562 | Bone Marr | GSM30315 | 14255 | 89428 |
| 811.6366 | 562 | 798.9877 | 663 | 670.9685 | 781 | 674.9147 | 899 | SMAD3 | HCC1954 | Breast | GSM28073 | 59999 | 89465 |
| 701.3882 | 154 | 809.8154 | 183 | 486.5385 | 179 | 687.3683 | 227 | SOX4 | HCC1954 | Breast | GSM28073 | 7587 | 89466 |
| 613.3952 | 413 | 647.9536 | 540 | 399.292 | 555 | 470.4974 | 641 | SMAD3 | MDA-MB-2 | Breast | GSM28073 | 73914 | 89467 |
| 689.6275 | 443 | 719.3689 | 575 | 525.9071 | 577 | 507.6891 | 628 | SMAD3 | MDA-MB-2 | Breast | GSM28073 | 63392 | 89468 |
| 835.1302 | 169 | 998.6633 | 221 | 388.5326 | 160 | 477.6279 | 189 | SOX4 | MDA-MB-2 | Breast | GSM28073 | 7716 | 89469 |
| 993.5662 | 175 | 1046.666 | 220 | 483.5432 | 165 | 446.8169 | 169 | SOX4 | MDA-MB-2 | Breast | GSM28073 | 6533 | 89470 |
| 827.5792 | 526 | 823.0706 | 629 | 698.1646 | 747 | 705.2749 | 864 | SMAD3 | HCC1954 | Breast | GSM28073 | 52684 | 89471 |
| 858.0737 | 553 | 840.9519 | 645 | 694.8229 | 727 | 707.3103 | 853 | SMAD3 | HCC1954 | Breast | GSM28073 | 49871 | 89472 |
| 779.2718 | 548 | 768.6189 | 648 | 683.0084 | 832 | 685.9793 | 959 | SMAD3 | HCC1954 | Breast | GSM28073 | 65368 | 89473 |

| | | | | | | | | | | | | | |
|---|---|---|---|---|---|---|---|---|---|---|---|---|---|
| 1253.965 | 166 | 1262.837 | 210 | 742.5911 | 149 | 842.8501 | 169 | RBP2 | MCF-7 | Breast | GSM70075 | 3227 | 89543 |
| 830.7805 | 607 | 815.1524 | 716 | 665.7241 | 820 | 674.0638 | 953 | FOXA1 | RT-4 | transitiona | GSM23103 | 69611 | 89554 |
| 554.671 | 166 | 542.3309 | 182 | 418.1569 | 211 | 579.5038 | 263 | FOXA1 | RT-4 | transitiona | GSM23103 | 12704 | 89563 |
| 1259.816 | 170 | 1292.817 | 230 | 793.8688 | 155 | 930.0728 | 179 | RBP2 | MCF-7 | Breast | GSM70075 | 3239 | 89574 |
| 693.8037 | 555 | 740.7348 | 732 | 524.5428 | 733 | 496.6992 | 787 | CTCF | MOLM-13 | None | GSM30943 | 90546 | 89590 |
| 648.3242 | 399 | 642.2316 | 479 | 403.378 | 512 | 416.6912 | 565 | DPF2 | MOLM-13 | None | GSM30943 | 63572 | 89592 |
| 1090.45 | 311 | 1089.202 | 378 | 814.9929 | 282 | 743.7619 | 274 | SS18 | SYO1 | None | GSM30943 | 9666 | 89594 |
| 1031.299 | 346 | 1031.484 | 425 | 754.051 | 324 | 726.5784 | 340 | BRD9 | SYO1 | None | GSM30943 | 15259 | 89595 |
| 808.6279 | 640 | 898.371 | 888 | 584.791 | 764 | 573.8529 | 852 | CTCF | SYO1 | None | GSM30943 | 78764 | 89597 |
| 1170.773 | 335 | 1165.93 | 394 | 891.9023 | 299 | 858.146 | 312 | RBP2 | M.M. | None | GSM31119 | 8343 | 89602 |
| 490.0225 | 258 | 707.296 | 355 | 303.9424 | 327 | 363.1282 | 379 | CTCF | THP-1 | Blood | GSM31120 | 35414 | 89609 |
| 1257.58 | 613 | 1278.317 | 766 | 918.9268 | 547 | 873.6824 | 556 | CTRL | HL-60 | Blood | GSM28365 | 15986 | 89610 |
| 746.6124 | 385 | 766.9252 | 493 | 507.462 | 453 | 480.39 | 482 | RAD21 | THP-1 | Blood | GSM31120 | 44996 | 89611 |
| 507.5272 | 243 | 705.7822 | 316 | 315.5934 | 304 | 296.5988 | 322 | RAD21 | THP-1 | Blood | GSM31120 | 30524 | 89613 |
| 520.5682 | 289 | 716.9974 | 417 | 364.1063 | 387 | 452.5229 | 455 | CTCF | THP-1 | Blood | GSM31120 | 42663 | 89615 |
| 519.1852 | 288 | 695.1966 | 394 | 322.3565 | 369 | 366.5642 | 420 | CTCF | THP-1 | Blood | GSM31120 | 42181 | 89616 |
| 502.8686 | 292 | 701.3074 | 417 | 382.2239 | 405 | 421.6804 | 456 | CTCF | THP-1 | Blood | GSM31120 | 44710 | 89617 |
| 1244.824 | 571 | 1295.462 | 752 | 940.6839 | 551 | 900.51 | 569 | NELFA | HEK293 | Embryonic | GSM30207 | 15800 | 89652 |
| 1303.537 | 722 | 1374.239 | 947 | 954.1302 | 677 | 921.7326 | 715 | NELFA | HEK293 | Embryonic | GSM30207 | 19918 | 89655 |
| 277.2639 | 110 | 269.5333 | 121 | 329.5508 | 176 | 592.1763 | 248 | TET2 | MCF-7 | Breast | GSM34147 | 10758 | 89703 |
| 38.32791 | 13 | 69.1335 | 20 | 26.34954 | 16 | 107.8248 | 35 | TET2 | MCF-7 | Breast | GSM34147 | 1151 | 89704 |
| 335.7203 | 68 | 513.6596 | 95 | 221.3854 | 75 | 427.0638 | 113 | ZEB1 | RKO | Colon | GSM23449 | 2721 | 89715 |
| 424.5708 | 81 | 404.6529 | 86 | 214.7339 | 77 | 291.2822 | 96 | ZEB1 | MIA PaCa-2 | Pancreatic | GSM23449 | 2919 | 89716 |
| 245.0681 | 46 | 295.252 | 56 | 125.99 | 44 | 213.9387 | 62 | ZEB1 | MIA PaCa-2 | Pancreatic | GSM23449 | 1679 | 89717 |
| 182.1705 | 36 | 237.6234 | 46 | 116.0972 | 39 | 266.142 | 65 | ZEB1 | MIA PaCa-2 | Pancreatic | GSM23449 | 1461 | 89718 |
| 1165.195 | 530 | 1220.078 | 706 | 866.7941 | 495 | 834.7287 | 522 | RBP2 | THP-1 | Blood | GSM31120 | 16507 | 89799 |
| 1177.311 | 572 | 1220.745 | 741 | 870.9598 | 534 | 848.5118 | 578 | RBP2 | THP-1 | Blood | GSM31120 | 18081 | 89803 |
| 1181.349 | 537 | 1228.573 | 701 | 878.6217 | 495 | 849.5222 | 524 | RBP2 | THP-1 | Blood | GSM31120 | 15958 | 89804 |
| 1160.568 | 514 | 1194.74 | 668 | 861.0993 | 480 | 833.0362 | 510 | RBP2 | THP-1 | Blood | GSM31120 | 16288 | 89805 |
| 1188.601 | 505 | 1218.879 | 640 | 876.4377 | 453 | 860.3563 | 494 | RBP2 | THP-1 | Blood | GSM31120 | 14345 | 89806 |
| 1150.315 | 500 | 1184.659 | 640 | 851.1115 | 458 | 830.3568 | 494 | RBP2 | THP-1 | Blood | GSM31120 | 15985 | 89807 |
| 1179.198 | 490 | 1204.368 | 624 | 875.4472 | 449 | 852.9867 | 481 | RBP2 | THP-1 | Blood | GSM31120 | 14109 | 89808 |
| 727.8939 | 257 | 766.7655 | 304 | 343.6396 | 277 | 381.3946 | 312 | PDX1 | None | Pancreas | GSM28580 | 25131 | 89815 |
| 635.2113 | 106 | 641.348 | 116 | 602.8844 | 139 | 1039.356 | 225 | HOXB13 | None | Prostate | GSM26432 | 3216 | 89854 |
| 1041.242 | 183 | 968.5166 | 191 | 846.8405 | 223 | 980.0082 | 343 | HOXB13 | None | Prostate | GSM26432 | 6482 | 89856 |
| 758.878 | 116 | 698.5027 | 120 | 770.4396 | 157 | 1070.523 | 238 | HOXB13 | None | Prostate | GSM26432 | 3140 | 89857 |
| 761.3691 | 421 | 780.9177 | 537 | 583.4899 | 529 | 530.6868 | 526 | MYOD1 | CTR | None | GSM22591 | 47295 | 89894 |
| 792.417 | 465 | 809.0065 | 587 | 601.1419 | 570 | 561.5938 | 589 | MYOD1 | CTR | None | GSM22591 | 49933 | 89895 |
| 214.2937 | 135 | 367.1002 | 193 | 150.1592 | 182 | 170.5456 | 207 | MYOG | CTR | None | GSM22591 | 20532 | 89897 |
| 56.45717 | 26 | 116.3038 | 42 | 45.97902 | 36 | 132.2811 | 63 | MYOG | CTR | None | GSM22591 | 3066 | 89898 |
| 835.8763 | 450 | 859.3274 | 578 | 624.578 | 523 | 565.6007 | 511 | MYOD1 | CTR | None | GSM22591 | 40682 | 89899 |
| 252.6922 | 315 | 353.9949 | 405 | 142.3283 | 438 | 149.0156 | 488 | GLI2 | THP-1 | Blood | GSM31121 | 84017 | 89908 |
| 166.9459 | 234 | 198.0942 | 285 | 106.7805 | 340 | 87.15417 | 355 | GLI2 | THP-1 | Blood | GSM31121 | 64882 | 89909 |
| 97.81745 | 27 | 86.9027 | 28 | 221.0348 | 60 | 284.3024 | 73 | RARG | RWPE1 | Prostate | GSM32730 | 1827 | 89951 |
| 61.31786 | 16 | 63.17523 | 18 | 99.49477 | 30 | 193.4968 | 46 | RARG | RWPE1 | Prostate | GSM32730 | 1072 | 89953 |
| 1154.792 | 228 | 1156.379 | 277 | 717.1561 | 192 | 751.2707 | 209 | ARID2 | Aska | None | GSM28875 | 5567 | 89955 |
| 248.2326 | 56 | 378.2196 | 78 | 103.3166 | 50 | 198.787 | 74 | SMARCA4 | Aska | None | GSM28875 | 2647 | 89956 |
| 385.3495 | 71 | 536.5854 | 94 | 117.1996 | 52 | 71.59904 | 44 | SMARCA4 | Aska | None | GSM28875 | 2521 | 89957 |
| 203.6025 | 78 | 249.7385 | 96 | 85.44578 | 78 | 89.71697 | 86 | SMARCC1 | Aska | None | GSM28875 | 6818 | 89958 |
| 1097.181 | 449 | 1100.667 | 550 | 798.9816 | 407 | 767.9382 | 422 | SMARCC1 | Aska | None | GSM28875 | 15953 | 89959 |
| 86.6992 | 42 | 51.61197 | 37 | 58.07675 | 53 | 99.56595 | 72 | SS18 | Aska | None | GSM28875 | 4703 | 89960 |
| 432.8177 | 69 | 597.6797 | 91 | 240.8837 | 67 | 227.6353 | 69 | SS18 | Aska | None | GSM28875 | 1944 | 89961 |
| 780.4307 | 107 | 908.9142 | 127 | 282.0734 | 80 | 320.6439 | 91 | ARID2 | Aska | None | GSM28875 | 2316 | 89963 |
| 132.619 | 66 | 204.254 | 91 | 119.3317 | 96 | 178.8694 | 123 | ESR2 | MDA-MB-2 | Breast | GSM29182 | 8265 | 89980 |
| 53.32076 | 25 | 65.15935 | 31 | 40.64675 | 34 | 85.37084 | 51 | ESR2 | MDA-MB-2 | Breast | GSM29182 | 3152 | 89981 |
| 69.2148 | 28 | 86.29769 | 35 | 62.18051 | 40 | 105.7461 | 55 | ESR2 | MDA-MB-2 | Breast | GSM29182 | 2939 | 89982 |
| 381.1575 | 217 | 471.9465 | 268 | 244.6597 | 281 | 276.4161 | 319 | RAD21 | M.M. | None | GSM31119 | 31467 | 89992 |
| 54.57406 | 23 | 51.36911 | 25 | 58.58303 | 36 | 176.6016 | 66 | TCF4 | LS174T | Colon | GSM46012 | 2591 | 9 |
| 673.0473 | 407 | 727.1926 | 561 | 495.5694 | 536 | 506.5979 | 604 | CTCF | THP-1 | Blood | GSM31120 | 62446 | 90018 |
| 668.7752 | 387 | 722.3984 | 533 | 442.8289 | 498 | 485.5479 | 560 | CTCF | THP-1 | Blood | GSM31120 | 58638 | 90019 |
| 630.743 | 370 | 699.7772 | 512 | 411.081 | 489 | 436.4676 | 544 | CTCF | THP-1 | Blood | GSM31120 | 59328 | 90020 |
| 586.0819 | 351 | 697.7092 | 493 | 380.0802 | 463 | 443.7776 | 532 | CTCF | THP-1 | Blood | GSM31120 | 56172 | 90021 |
| 579.3753 | 363 | 690.8074 | 510 | 421.8125 | 502 | 446.5754 | 558 | CTCF | THP-1 | Blood | GSM31120 | 61199 | 90022 |
| 610.4204 | 357 | 702.078 | 496 | 400.515 | 473 | 454.1364 | 538 | CTCF | THP-1 | Blood | GSM31120 | 57250 | 90023 |
| 539.7311 | 352 | 686.012 | 503 | 401.0627 | 492 | 386.2759 | 530 | CTCF | THP-1 | Blood | GSM31120 | 60818 | 90024 |
| 630.5504 | 363 | 702.2473 | 499 | 389.6069 | 470 | 447.565 | 537 | CTCF | THP-1 | Blood | GSM31120 | 57347 | 90025 |
| 670.8001 | 384 | 719.3251 | 526 | 437.8162 | 493 | 496.8885 | 561 | CTCF | THP-1 | Blood | GSM31120 | 57974 | 90026 |
| 663.799 | 382 | 711.5728 | 520 | 407.5215 | 485 | 445.7678 | 545 | CTCF | THP-1 | Blood | GSM31120 | 58847 | 90027 |
| 923.5845 | 597 | 839.9664 | 613 | 813.6757 | 883 | 849.8731 | 1082 | HOXB13 | LN95 | Prostate | GSM26432 | 46235 | 90052 |
| 898.5207 | 496 | 805.0567 | 486 | 743.3564 | 649 | 782.5407 | 812 | HOXB13 | 22RV1 | Prostate | GSM26432 | 35631 | 90057 |
| 917.1087 | 471 | 828.1366 | 465 | 759.7331 | 614 | 808.9801 | 791 | HOXB13 | 22RV1 | Prostate | GSM26432 | 32338 | 90058 |
| 73.11931 | 38 | 66.98607 | 41 | 33.16421 | 42 | 51.14019 | 54 | MED1 | hMSC-TER1 | None | GSM31009 | 4911 | 90063 |

| | | | | | | | | | | | | | |
|---|---|---|---|---|---|---|---|---|---|---|---|---|---|
| 137.3155 | 111 | 187.1627 | 143 | 98.85691 | 155 | 134.5384 | 188 | MED1 | hMSC-TER1 | None | GSM31009 | 19879 | 90064 |
| 166.8852 | 122 | 178.4454 | 142 | 105.086 | 161 | 124.702 | 186 | MED1 | hMSC-TER1 | None | GSM31009 | 20610 | 90065 |
| 529.8913 | 213 | 604.5087 | 253 | 261.8462 | 236 | 279.68 | 262 | MED1 | hMSC-TER1 | None | GSM31009 | 22078 | 90066 |
| 129.6363 | 78 | 89.7249 | 75 | 73.00077 | 96 | 127.7792 | 129 | MED1 | hMSC-TER1 | None | GSM31009 | 11225 | 90067 |
| 275.1352 | 186 | 287.2917 | 214 | 172.7807 | 243 | 199.3733 | 278 | MED1 | hMSC-TER1 | None | GSM31009 | 29857 | 90068 |
| 81.35961 | 85 | 70.71945 | 92 | 65.74933 | 127 | 106.1439 | 163 | MED1 | hMSC-TER1 | None | GSM31009 | 18139 | 90069 |
| 369.1057 | 202 | 414.7558 | 239 | 227.6402 | 255 | 268.0447 | 294 | MED1 | hMSC-TER1 | None | GSM31009 | 27439 | 90070 |
| 94.61208 | 44 | 89.46934 | 48 | 39.27124 | 46 | 78.46228 | 66 | MED1 | hMSC-TER1 | None | GSM31009 | 5103 | 90071 |
| 272.2203 | 173 | 319.8996 | 209 | 179.7551 | 228 | 251.4992 | 280 | MED1 | hMSC-TER1 | None | GSM31009 | 26271 | 90072 |
| 120.0005 | 40 | 250.8553 | 66 | 88.31726 | 51 | 190.2901 | 79 | MLLT3 | MLL-AF9 | None | GSM27868 | 3454 | 90077 |
| 362.7685 | 221 | 549.9368 | 297 | 243.3787 | 289 | 225.5665 | 306 | CTCF | None | None | GSM29487 | 30963 | 90083 |
| 420.7092 | 240 | 625.9828 | 320 | 276.0767 | 309 | 253.8262 | 326 | CTCF | None | None | GSM29487 | 32027 | 90084 |
| 174.5428 | 79 | 244.1725 | 104 | 138.8502 | 108 | 134.8997 | 115 | MAPK1 | CTR | None | GSM22591 | 9115 | 90089 |
| 944.8116 | 529 | 991.9289 | 696 | 684.8109 | 548 | 646.242 | 569 | EZH2 | CTR | None | GSM22591 | 35487 | 90091 |
| 92.95654 | 60 | 103.1487 | 71 | 39.70693 | 68 | 47.78061 | 79 | MAPK1 | CTR | None | GSM22591 | 9510 | 90092 |
| 1196.134 | 495 | 1225.606 | 633 | 934.7948 | 505 | 900.968 | 523 | TET2 | MCF-7 | Breast | GSM34147 | 13964 | 90143 |
| 1435.618 | 168 | 1430.323 | 195 | 855.036 | 127 | 860.7957 | 134 | TET2 | MCF-7 | Breast | GSM34147 | 1672 | 90144 |
| 235.5966 | 45 | 217.7514 | 47 | 146.4588 | 48 | 236.9534 | 66 | HIF1A | HKC8 | Kidney | GSM34178 | 1762 | 90165 |
| 111.3596 | 55 | 83.70227 | 54 | 142.0129 | 92 | 238.0591 | 125 | EPAS1 | HKC8 | Kidney | GSM34178 | 6349 | 90167 |
| 162.624 | 43 | 129.8948 | 42 | 108.8997 | 50 | 230.7114 | 78 | HIF1A | HKC8 | Kidney | GSM34178 | 2582 | 90168 |
| 91.07927 | 56 | 69.3788 | 56 | 140.6454 | 103 | 395.3336 | 177 | EPAS1 | HKC8 | Kidney | GSM34178 | 7843 | 90169 |
| 863.779 | 176 | 969.715 | 210 | 505.0299 | 183 | 508.1276 | 196 | TFAP2C | hESC | None | GSM29017 | 6949 | 90175 |
| 838.338 | 286 | 861.343 | 369 | 481.2517 | 305 | 424.2762 | 312 | GFI1 | Kasumi-1 | None | GSM27592 | 22175 | 90189 |
| 1152.497 | 585 | 1214.254 | 784 | 846.6024 | 553 | 812.8804 | 579 | RBP2 | THP-1 | Blood | GSM31120 | 20337 | 90190 |
| 430.9188 | 275 | 620.26 | 361 | 299.604 | 373 | 325.7358 | 418 | RAD21 | THP-1 | Blood | GSM31120 | 44850 | 90191 |
| 1193.695 | 440 | 1214.732 | 561 | 893.6887 | 400 | 859.3475 | 415 | RBP2 | THP-1 | Blood | GSM31120 | 11581 | 90192 |
| 1203.111 | 424 | 1233.289 | 545 | 888.0175 | 366 | 870.1832 | 393 | RBP2 | THP-1 | Blood | GSM31120 | 10506 | 90193 |
| 1192.945 | 400 | 1224.673 | 512 | 893.4412 | 348 | 865.9687 | 366 | RBP2 | THP-1 | Blood | GSM31120 | 9764 | 90194 |
| 1204.563 | 424 | 1232.701 | 542 | 896.6321 | 372 | 874.277 | 394 | RBP2 | THP-1 | Blood | GSM31120 | 10451 | 90195 |
| 1195.805 | 462 | 1224.997 | 591 | 898.2084 | 420 | 862.5495 | 431 | RBP2 | THP-1 | Blood | GSM31120 | 12057 | 90196 |
| 1189.459 | 517 | 1221.843 | 670 | 883.1108 | 475 | 847.4117 | 495 | RBP2 | THP-1 | Blood | GSM31120 | 14775 | 90197 |
| 1188.812 | 512 | 1224.562 | 667 | 888.9693 | 474 | 842.8712 | 480 | RBP2 | THP-1 | Blood | GSM31120 | 14568 | 90198 |
| 1182.199 | 554 | 1216.956 | 711 | 866.6299 | 502 | 827.9707 | 522 | RBP2 | THP-1 | Blood | GSM31120 | 16896 | 90199 |
| 1201.745 | 433 | 1222.306 | 551 | 891.2941 | 385 | 894.7198 | 440 | SETD7 | HES-3 | Embryo | GSM28791 | 11621 | 90215 |
| 1199.911 | 322 | 1205.581 | 392 | 904.4293 | 281 | 889.5961 | 305 | SETD7 | HES-3 | Embryo | GSM28791 | 7727 | 90217 |
| 433.4513 | 62 | 420.0922 | 66 | 202.225 | 54 | 123.36 | 44 | SETD7 | HES-3 | Embryo | GSM28791 | 1565 | 90218 |
| 340.8018 | 45 | 237.2605 | 39 | 89.2575 | 28 | 139.7852 | 38 | SETD7 | HES-3 | Embryo | GSM28791 | 1008 | 90219 |
| 355.6676 | 46 | 392.9409 | 53 | 163.3477 | 39 | 253.5828 | 53 | SETD7 | HES-3 | Embryo | GSM28791 | 1011 | 90220 |
| 1194.337 | 500 | 1224.95 | 644 | 877.7531 | 448 | 862.577 | 488 | SETD7 | HES-3 | Embryo | GSM28791 | 14518 | 90222 |
| 180.659 | 50 | 133.1115 | 47 | 96.82725 | 53 | 76.48023 | 51 | TP53 | U2OS | Bone | GSM30169 | 3318 | 90253 |
| 751.5945 | 337 | 768.1492 | 428 | 428.5975 | 382 | 542.5821 | 453 | RAD21 | THP-1 | Blood | GSM31119 | 37576 | 90270 |
| 147.7114 | 117 | 217.42 | 156 | 72.56855 | 145 | 132.8357 | 194 | IRF4 | KK1 | None | GSM24816 | 22079 | 90298 |
| 299.1205 | 124 | 441.1047 | 167 | 250.6191 | 170 | 263.4295 | 187 | TP53 | SW480 | Colon | GSM32043 | 12497 | 90300 |
| 915.5843 | 350 | 915.1668 | 420 | 694.2868 | 375 | 684.045 | 415 | KMT2B | SW480 | Colon | GSM32043 | 20731 | 90302 |
| 922.9522 | 636 | 930.0398 | 778 | 671.5802 | 705 | 662.8822 | 794 | TP53 | SW480 | Colon | GSM32043 | 50749 | 90303 |
| 1385.938 | 198 | 1430.896 | 263 | 1044.793 | 163 | 1083.938 | 181 | TET2 | MCF-7 | Breast | GSM34147 | 2333 | 90315 |
| 1293.486 | 291 | 1339.64 | 398 | 1026.555 | 273 | 985.6695 | 279 | TET2 | MCF-7 | Breast | GSM34147 | 5093 | 90316 |
| 991.7089 | 475 | 978.2968 | 559 | 806.9088 | 593 | 805.694 | 672 | ESR1 | MCF-7 | Breast | GSM34147 | 26895 | 90319 |
| 661.663 | 388 | 667.1037 | 480 | 377.0648 | 479 | 341.6902 | 506 | FLI1 | A673 | Muscle | GSM28575 | 59834 | 90348 |
| 667.3853 | 382 | 683.0582 | 485 | 344.2834 | 455 | 316.4825 | 483 | NKX2-2 | A673 | Muscle | GSM28575 | 57685 | 90349 |
| 217.1909 | 50 | 291.6438 | 65 | 246.8561 | 76 | 322.3002 | 93 | MYC | DLD-1 | Colon | GSM32897 | 2748 | 90357 |
| 954.842 | 206 | 973.0405 | 262 | 758.8813 | 255 | 654.8294 | 253 | MYC | DLD-1 | Colon | GSM32897 | 10197 | 90358 |
| 838.0926 | 403 | 837.2761 | 485 | 610.0204 | 432 | 584.8182 | 462 | TRIM24 | MOLM-13 | None | GSM26873 | 32764 | 90382 |
| 881.377 | 323 | 853.0236 | 367 | 561.8259 | 325 | 488.447 | 330 | TRIM24 | MOLM-13 | None | GSM26873 | 21148 | 90383 |
| 444.2475 | 227 | 716.0028 | 315 | 334.7164 | 305 | 408.8244 | 357 | CTCF | LCL | Blood | GSM32129 | 28262 | 90390 |
| 543.7434 | 288 | 676.5606 | 360 | 302.4023 | 345 | 394.0408 | 413 | CTCF | LCL | Blood | GSM32129 | 35522 | 90393 |
| 543.6797 | 252 | 741.3393 | 347 | 378.2078 | 326 | 432.8108 | 372 | CTCF | LCL | Blood | GSM32129 | 29723 | 90399 |
| 529.5678 | 249 | 752.932 | 357 | 378.0511 | 325 | 481.9524 | 387 | CTCF | LCL | Blood | GSM32129 | 29187 | 90400 |
| 905.4199 | 545 | 925.4823 | 684 | 645.5608 | 564 | 605.2814 | 585 | SET | SET2 | None | GSM26868 | 40269 | 90521 |
| 734.9472 | 146 | 852.8975 | 174 | 468.0981 | 161 | 309.5834 | 140 | STAT1 | SET2 | None | GSM26868 | 6507 | 90522 |
| 627.866 | 148 | 731.9425 | 177 | 375.3208 | 162 | 233.3869 | 139 | STAT1 | SET2 | None | GSM26868 | 8140 | 90523 |
| 639.6392 | 117 | 776.0432 | 143 | 352.9213 | 118 | 261.2873 | 108 | STAT1 | SET2 | None | GSM26868 | 4586 | 90524 |
| 516.2027 | 112 | 610.3957 | 135 | 385.3838 | 135 | 222.5677 | 110 | STAT1 | SET2 | None | GSM26868 | 5476 | 90525 |
| 1114.868 | 148 | 1164.572 | 177 | 573.6196 | 135 | 460.7595 | 127 | ETS1 | HEYA8 | ovarian | GSM27169 | 3275 | 90553 |
| 1192.549 | 254 | 1183.649 | 301 | 917.0333 | 229 | 733.2962 | 209 | ETS1 | Ovcar8 | ovarian | GSM27169 | 5743 | 90554 |
| 654.6255 | 376 | 690.0833 | 499 | 381.9375 | 477 | 462.9296 | 555 | CTCF | HeLa | None | GSM28722 | 59665 | 90566 |
| 662.5312 | 381 | 723.7821 | 535 | 428.3015 | 493 | 502.7873 | 569 | CTCF | HeLa | None | GSM28722 | 59167 | 90567 |
| 583.8747 | 322 | 708.3705 | 444 | 399.1513 | 426 | 458.9722 | 487 | CTCF | HeLa | None | GSM28722 | 46561 | 90568 |
| 1199.277 | 468 | 1217.485 | 588 | 888.1845 | 417 | 870.1221 | 450 | RBP2 | THP-1 | Blood | GSM31120 | 12581 | 90569 |
| 1200.968 | 461 | 1227.493 | 589 | 901.4814 | 423 | 878.7888 | 456 | RBP2 | THP-1 | Blood | GSM31120 | 12502 | 90570 |
| 1165.352 | 489 | 1215.844 | 650 | 875.2659 | 466 | 853.5653 | 497 | RBP2 | THP-1 | Blood | GSM31120 | 14954 | 90573 |
| 811.2187 | 384 | 786.3557 | 441 | 707.1654 | 575 | 711.7096 | 664 | FOXA1 | WS8 | None | GSM30964 | 35695 | 90574 |

| | | | | | | | | | | | | | |
|---|---|---|---|---|---|---|---|---|---|---|---|---|---|
| 784.1046 | 410 | 746.4638 | 458 | 685.7665 | 623 | 698.0844 | 735 | FOXA1 | WS8 | None | GSM30964 | 43100 | 90576 |
| 765.2718 | 438 | 730.5669 | 493 | 659.9481 | 658 | 671.2329 | 771 | FOXA1 | WS8 | None | GSM30964 | 49653 | 90577 |
| 26.52459 | 16 | 43.74254 | 23 | 19.38598 | 22 | 78.71609 | 44 | TP53 | 116 | None | GSM29889 | 2502 | 90649 |
| 252.3035 | 85 | 209.4133 | 86 | 102.5174 | 83 | 176.0268 | 113 | TP53 | 116 | None | GSM29889 | 7095 | 90650 |
| 1062.968 | 245 | 1085.724 | 321 | 733.2155 | 236 | 777.625 | 259 | ETS1 | None | Umbilical V | GSM29474 | 9038 | 90690 |
| 771.8632 | 413 | 712.3353 | 435 | 657.4504 | 598 | 665.5693 | 698 | FOXA1 | TAM_R | Mammary | GSM30964 | 43646 | 90759 |
| 768.0086 | 400 | 758.2515 | 473 | 666.6344 | 598 | 643.1718 | 648 | GRHL2 | TAM_R | Mammary | GSM30964 | 42609 | 90760 |
| 743.9305 | 358 | 736.4322 | 430 | 654.2106 | 556 | 635.0484 | 607 | GRHL2 | TAM_R | Mammary | GSM30964 | 40445 | 90761 |
| 764.0917 | 401 | 758.4215 | 481 | 658.3454 | 598 | 635.5388 | 647 | GRHL2 | TAM_R | Mammary | GSM30964 | 43635 | 90762 |
| 758.0842 | 500 | 713.6445 | 551 | 634.061 | 719 | 625.8368 | 808 | FOXA1 | TAM_R | Mammary | GSM30964 | 61040 | 90763 |
| 769.2237 | 492 | 710.0723 | 527 | 624.7139 | 677 | 631.6988 | 785 | FOXA1 | TAM_R | Mammary | GSM30964 | 57614 | 90764 |
| 1197.818 | 328 | 1192.302 | 397 | 897.636 | 285 | 870.2656 | 300 | RBP2 | THP-1 | Blood | GSM31119 | 7644 | 90832 |
| 739.55 | 340 | 764.8311 | 446 | 495.2857 | 418 | 554.0547 | 475 | CTCF | THP-1 | Blood | GSM31119 | 40240 | 90833 |
| 704.1439 | 323 | 756.1815 | 443 | 481.2875 | 420 | 468.6955 | 451 | CTCF | THP-1 | Blood | GSM31119 | 41254 | 90836 |
| 40.67121 | 25 | 131.9135 | 50 | 25.0419 | 32 | 58.48297 | 49 | TP53 | None | None | GSM29889 | 3801 | 90842 |
| 232.8444 | 36 | 381.1534 | 53 | 187.2443 | 43 | 253.2103 | 54 | TP53 | None | None | GSM29889 | 1041 | 90843 |
| 303.5633 | 83 | 424.9109 | 110 | 137.3341 | 82 | 197.6045 | 104 | TP53 | None | None | GSM29889 | 5679 | 90844 |
| 542.0702 | 144 | 634.233 | 173 | 289.9786 | 154 | 285.7408 | 164 | TP53 | None | None | GSM29889 | 9780 | 90845 |
| 193.8062 | 39 | 172.4746 | 40 | 41.92573 | 25 | 135.8873 | 48 | TP53 | None | None | GSM29889 | 1772 | 90846 |
| 522.5894 | 342 | 688.1229 | 496 | 397.5427 | 487 | 453.85 | 555 | CTCF | None | None | GSM28166 | 62824 | 90862 |
| 624.9186 | 384 | 705.2422 | 548 | 436.7073 | 529 | 489.732 | 603 | CTCF | None | None | GSM28166 | 68830 | 90863 |
| 520.1391 | 363 | 644.8232 | 491 | 324.469 | 489 | 386.7575 | 565 | RAD21 | None | None | GSM28166 | 71066 | 90864 |
| 655.0701 | 406 | 708.2654 | 560 | 479.8143 | 552 | 419.915 | 576 | RAD21 | None | None | GSM28166 | 70381 | 90865 |
| 439.8134 | 77 | 424.0668 | 82 | 195.824 | 67 | 193.7415 | 71 | NIPBL | None | None | GSM28166 | 2332 | 90868 |
| 245.7638 | 52 | 335.7009 | 68 | 128.8394 | 51 | 151.0462 | 59 | NIPBL | None | None | GSM28166 | 2128 | 90869 |
| 41.90336 | 46 | 32.71412 | 48 | 25.56565 | 64 | 37.22805 | 79 | TP63 | None | None | GSM34558 | 10892 | 90881 |
| 118.2864 | 103 | 135.8764 | 124 | 54.42372 | 127 | 63.93694 | 146 | TP63 | None | Ectoderm | GSM34558 | 20924 | 90882 |
| 34.29586 | 75 | 36.54437 | 89 | 11.27462 | 99 | 30.92191 | 139 | TP63 | None | Ectoderm | GSM34558 | 26529 | 90883 |
| 20.89768 | 26 | 75.17763 | 51 | 16.02886 | 40 | 45.932 | 63 | TP63 | None | None | GSM34558 | 7073 | 90884 |
| 536.4263 | 182 | 827.6439 | 253 | 291.9247 | 205 | 387.2033 | 249 | STAG2 | MCF-10A | Breast | GSM29422 | 16495 | 90908 |
| 276.9431 | 89 | 294.2511 | 102 | 125.3669 | 91 | 177.5441 | 114 | STAG1 | MCF-10A | Breast | GSM29422 | 7425 | 90910 |
| 495.036 | 187 | 689.7552 | 245 | 302.4196 | 225 | 394.3453 | 271 | STAG2 | MCF-10A | Breast | GSM29422 | 19000 | 90911 |
| 211.902 | 70 | 245.7799 | 84 | 105.9458 | 75 | 135.9135 | 90 | STAG1 | MCF-10A | Breast | GSM29422 | 6007 | 90912 |
| 222.847 | 55 | 304.0864 | 72 | 98.19243 | 52 | 196.2345 | 78 | EZH2 | LNCaP | Prostate | GSM28790 | 3156 | 90916 |
| 776.4878 | 429 | 733.7145 | 471 | 660.4464 | 623 | 664.0681 | 719 | FOXA1 | TAM_R | Mammary | GSM30964 | 45691 | 90994 |
| 748.5114 | 455 | 698.2363 | 496 | 633.3427 | 670 | 632.937 | 763 | FOXA1 | TAM_R | Mammary | GSM30964 | 55579 | 90995 |
| 589.7144 | 289 | 729.6189 | 393 | 367.7953 | 362 | 335.2951 | 380 | CTCF | T47D | Mammary | GSM30451 | 36356 | 90998 |
| 705.8243 | 406 | 742.8997 | 538 | 500.889 | 502 | 468.9934 | 534 | CTCF | T47D | Mammary | GSM30451 | 53399 | 90999 |
| 701.1763 | 310 | 769.3035 | 430 | 424.9973 | 379 | 392.3762 | 399 | CTCF | T47D | Mammary | GSM30451 | 35675 | 91000 |
| 567.4277 | 265 | 734.0479 | 357 | 345.8084 | 326 | 347.0133 | 354 | CTCF | T47D | Mammary | GSM30451 | 31416 | 91001 |
| 95.61127 | 55 | 178.5152 | 83 | 30.76145 | 55 | 72.29743 | 82 | ZNF366 | SCLC-24H | None | GSM20812 | 7867 | 91038 |
| 137.8899 | 58 | 143.0427 | 66 | 40.22126 | 52 | 80.05784 | 74 | ZNF366 | None | None | GSM20812 | 6185 | 91039 |
| 146.5496 | 61 | 150.516 | 69 | 44.20562 | 55 | 79.00465 | 75 | ZNF366 | None | None | GSM20812 | 6252 | 91040 |
| 207.0087 | 151 | 294.2613 | 198 | 78.58725 | 169 | 162.6233 | 235 | ZNF366 | None | None | GSM20812 | 26013 | 91041 |
| 163.046 | 116 | 296.8323 | 170 | 71.58923 | 135 | 130.5668 | 181 | ZNF366 | SCLC-24H | None | GSM20812 | 19401 | 91043 |
| 107.0603 | 123 | 168.6329 | 167 | 49.19981 | 157 | 92.84473 | 208 | ZNF366 | SCLC-24H | None | GSM20812 | 28632 | 91044 |
| 159.5491 | 49 | 194.6789 | 60 | 61.47881 | 44 | 57.23773 | 46 | SUZ12 | CRL7250 (fi | None | GSM28876 | 2920 | 91046 |
| 1054.208 | 540 | 1077.579 | 680 | 761.1961 | 510 | 727.0853 | 535 | SS18 | None | None | GSM28876 | 23011 | 91047 |
| 712.565 | 575 | 701.3931 | 680 | 511.8477 | 706 | 497.3259 | 780 | SS18 | None | None | GSM28876 | 86714 | 91048 |
| 156.0866 | 93 | 140.6019 | 100 | 97.10048 | 118 | 111.8564 | 135 | SS18 | V5 | None | GSM28876 | 12840 | 91051 |
| 442.6214 | 96 | 569.2408 | 121 | 236.5261 | 96 | 215.4318 | 98 | SUZ12 | None | None | GSM28876 | 4109 | 91052 |
| 517.8516 | 261 | 588.4786 | 310 | 282.5805 | 313 | 298.9443 | 347 | SS18 | V5 | None | GSM28876 | 34080 | 91054 |
| 173.8619 | 192 | 210.7528 | 235 | 59.16079 | 223 | 89.91032 | 275 | NANOG | None | None | GSM28166 | 41906 | 91059 |
| 287.5358 | 271 | 375.7456 | 340 | 105.7406 | 313 | 135.7159 | 369 | NANOG | None | None | GSM28166 | 52453 | 91062 |
| 958.1805 | 244 | 969.8615 | 304 | 524.0854 | 231 | 563.5411 | 256 | ZNF143 | None | None | GSM28166 | 11485 | 91063 |
| 232.1213 | 121 | 326.7448 | 159 | 120.3349 | 142 | 256.3262 | 207 | CTCF | None | None | GSM28166 | 15803 | 91065 |
| 960.3843 | 218 | 971.8109 | 277 | 445.0255 | 201 | 461.4766 | 219 | ZNF143 | None | None | GSM28166 | 10083 | 91066 |
| 312.2066 | 85 | 406.3372 | 108 | 186.2482 | 94 | 207.7417 | 106 | KLF4 | None | None | GSM28166 | 5240 | 91067 |
| 67.75498 | 67 | 30.71964 | 57 | 8.167332 | 57 | 25.55448 | 85 | POU5F1 | None | None | GSM28166 | 13284 | 91068 |
| 217.3102 | 266 | 387.9632 | 373 | 181.3592 | 405 | 219.6395 | 470 | RAD21 | HUES64 | Embryo | GSM28945 | 63620 | 91096 |
| 212.6269 | 169 | 329.7417 | 229 | 158.1556 | 238 | 279.672 | 317 | CTCF | HUES64 | Embryo | GSM28945 | 30003 | 91097 |
| 256.9996 | 303 | 316.5313 | 373 | 134.866 | 410 | 142.8014 | 458 | GLI2 | THP-1 | Blood | GSM31121 | 77685 | 91098 |
| 8.918468 | 16 | 5.767602 | 16 | 1.053267 | 16 | 29.31153 | 45 | CEBPB | THP-1 | Blood | GSM31121 | 5112 | 91099 |
| 109.0497 | 176 | 137.4616 | 219 | 79.86663 | 268 | 97.47737 | 311 | CEBPB | THP-1 | Blood | GSM31121 | 50940 | 91100 |
| 247.6643 | 336 | 329.4725 | 424 | 132.0783 | 467 | 132.0233 | 515 | GLI2 | THP-1 | Blood | GSM31121 | 95491 | 91101 |
| 89.47537 | 131 | 110.2684 | 162 | 54.91202 | 188 | 98.31491 | 244 | CEBPB | THP-1 | Blood | GSM31121 | 35681 | 91102 |
| 249.8219 | 159 | 360.3295 | 210 | 149.1517 | 202 | 259.562 | 269 | CTCF | HUES64 | Embryo | GSM28945 | 24077 | 91103 |
| 499.516 | 92 | 468.8869 | 97 | 261.4145 | 90 | 251.7613 | 94 | EGR1 | A375 | None | GSM32127 | 3564 | 91126 |
| 720.881 | 116 | 707.3805 | 125 | 345.9849 | 106 | 347.9519 | 113 | AR | A375 | Skin | GSM32127 | 3678 | 91127 |
| 573.2389 | 94 | 812.1577 | 126 | 280.2885 | 87 | 249.2418 | 87 | AR | A375 | Skin | GSM32127 | 3090 | 91128 |
| 1174.631 | 296 | 1161.345 | 352 | 888.0686 | 266 | 852.1093 | 271 | SMARCA4 | TTC1240 | None | GSM33260 | 7818 | 91191 |
| 183.6332 | 34 | 310.5204 | 51 | 207.5412 | 51 | 250.9785 | 60 | T | None | None | GSM30998 | 1458 | 91202 |

| | | | | | | | | | | | | | |
|---|---|---|---|---|---|---|---|---|---|---|---|---|---|
| 701.6652 | 140 | 950.3913 | 182 | 324.0426 | 131 | 265.0695 | 127 | ZNF341 | None | None | GSM30998 | 6109 | 91203 |
| 733.0493 | 138 | 1017.1 | 182 | 351.0667 | 130 | 265.8756 | 121 | ZNF341 | None | None | GSM30998 | 5543 | 91204 |
| 760.0867 | 491 | 805.3479 | 657 | 613.5551 | 677 | 591.326 | 733 | CTCF | MCF-7 | Breast | GSM33571 | 61168 | 91207 |
| 37.38052 | 17 | 59.10585 | 24 | 17.07775 | 18 | 42.43077 | 29 | CTCF | MCF-7 | Breast | GSM33571 | 1822 | 91208 |
| 806.8317 | 547 | 841.8669 | 708 | 661.741 | 755 | 636.9081 | 821 | CTCF | MCF-7 | Breast | GSM33571 | 62319 | 91209 |
| 860.6121 | 620 | 894.5233 | 794 | 661.3183 | 768 | 632.4007 | 824 | CTCF | MCF-7 | Breast | GSM33571 | 61048 | 91210 |
| 762.279 | 480 | 795.6827 | 627 | 590.8167 | 622 | 582.3468 | 694 | CTCF | MCF-7 | Breast | GSM33571 | 57396 | 91211 |
| 776.5067 | 549 | 824.8056 | 727 | 623.6947 | 751 | 610.412 | 833 | CTCF | MCF-7 | Breast | GSM33571 | 69842 | 91212 |
| 743.8548 | 489 | 787.0263 | 647 | 612.0518 | 692 | 583.9452 | 744 | CTCF | MCF-7 | Breast | GSM33571 | 63547 | 91213 |
| 764.7919 | 502 | 798.2651 | 652 | 602.6266 | 667 | 583.9861 | 729 | CTCF | MCF-7 | Breast | GSM33571 | 62242 | 91214 |
| 760.5663 | 543 | 800.9571 | 708 | 614.5671 | 750 | 595.25 | 825 | CTCF | MCF-7 | Breast | GSM33571 | 71196 | 91215 |
| 738.7162 | 490 | 768.1133 | 633 | 605.7538 | 694 | 576.2918 | 742 | CTCF | MCF-7 | Breast | GSM33571 | 65139 | 91216 |
| 1029.965 | 314 | 1051.354 | 403 | 786.5421 | 313 | 798.4882 | 367 | RAD21 | T47D | Mammary | GSM30451 | 12376 | 91222 |
| 1019.789 | 418 | 1058.623 | 553 | 773.6961 | 433 | 755.2884 | 469 | RAD21 | T47D | Mammary | GSM30451 | 18936 | 91223 |
| 1032.854 | 265 | 1037.736 | 330 | 754.0191 | 259 | 745.8995 | 275 | RAD21 | T47D | Mammary | GSM30451 | 10238 | 91224 |
| 911.1336 | 187 | 999.8872 | 246 | 562.244 | 203 | 617.9532 | 227 | RAD21 | T47D | Mammary | GSM30451 | 8364 | 91225 |
| 994.3029 | 432 | 987.1747 | 520 | 734.5825 | 435 | 722.602 | 478 | RAD21 | T47D | Mammary | GSM30451 | 21971 | 91226 |
| 216.0886 | 45 | 304.9152 | 60 | 186.536 | 57 | 212.7874 | 65 | SS18 | HS-SY-2 | None | GSM29161 | 1780 | 91236 |
| 38.89015 | 14 | 31.8174 | 14 | 20.56239 | 15 | 88.58474 | 33 | KDM2B | HS-SY-2 | None | GSM29161 | 1130 | 91237 |
| 534.661 | 164 | 477.553 | 172 | 781.8275 | 302 | 835.8208 | 396 | ESR1 | MCF-7 | Breast | GSM27525 | 12993 | 91252 |
| 711.3063 | 255 | 657.0769 | 274 | 723.4337 | 442 | 751.4534 | 542 | ESR1 | MCF-7 | Breast | GSM27525 | 24533 | 91253 |
| 353.6655 | 215 | 507.2779 | 282 | 249.441 | 285 | 363.4833 | 356 | CTCF | LCL | Blood | GSM32129 | 29249 | 91256 |
| 221.3619 | 42 | 359.0049 | 61 | 82.62551 | 34 | 115.0868 | 43 | SMARCC1 | None | None | GSM28875 | 1527 | 91279 |
| 75.801 | 44 | 49.15678 | 41 | 43.98401 | 55 | 130.826 | 93 | JUN | None | Umbilical V | GSM29474 | 6541 | 91320 |
| 52.38477 | 33 | 71.30929 | 43 | 50.19864 | 51 | 117.5497 | 79 | JUN | None | Umbilical V | GSM29474 | 5352 | 91321 |
| 66.45145 | 45 | 63.56521 | 50 | 49.06841 | 63 | 104.3497 | 92 | JUN | None | Umbilical V | GSM29474 | 7580 | 91322 |
| 44.99726 | 29 | 69.16335 | 40 | 109.3429 | 67 | 134.3933 | 79 | GATA2 | None | Umbilical V | GSM29474 | 4730 | 91327 |
| 45.09909 | 29 | 26.38035 | 26 | 85.91069 | 60 | 84.27223 | 64 | GATA2 | None | Umbilical V | GSM29474 | 4705 | 91328 |
| 32.32571 | 26 | 33.33207 | 30 | 72.32289 | 58 | 91.07265 | 69 | GATA2 | None | Umbilical V | GSM29474 | 5062 | 91329 |
| 1088.472 | 405 | 1118.155 | 524 | 831.5337 | 410 | 787.4557 | 414 | SMARCA4 | TTC1240 | None | GSM33260 | 15644 | 91331 |
| 928.8368 | 354 | 909.241 | 412 | 681.0536 | 358 | 585.5092 | 352 | BRD9 | Aska | None | GSM33260 | 21760 | 91332 |
| 1082.273 | 294 | 1103.158 | 382 | 814.5416 | 281 | 784.0693 | 291 | SMARCA4 | TTC1240 | None | GSM33260 | 10621 | 91333 |
| 1139.915 | 347 | 1150.043 | 438 | 876.475 | 340 | 817.8527 | 327 | SMARCA4 | TTC1240 | None | GSM33260 | 10820 | 91334 |
| 507.0922 | 267 | 718.0837 | 375 | 345.819 | 353 | 347.9799 | 384 | CTCF | TTC1240 | None | GSM33260 | 38285 | 91337 |
| 1295.188 | 188 | 1312.79 | 242 | 880.0881 | 160 | 1037.598 | 193 | CDK9 | T47D | Mammary | GSM31249 | 2420 | 91351 |
| 1138.496 | 324 | 1201.031 | 457 | 880.0748 | 313 | 867.1993 | 344 | ELL2 | 293T | Embryonic | GSM33605 | 9549 | 91384 |
| 529.8734 | 76 | 847.6862 | 110 | 271.507 | 70 | 408.5753 | 93 | SETD7 | hESC | None | GSM28791 | 1960 | 91429 |
| 1246.794 | 775 | 1259.124 | 944 | 918.7769 | 777 | 899.6526 | 855 | E2F1 | LNCaP | Prostate | GSM24924 | 26683 | 91432 |
| 1264.834 | 442 | 1292.974 | 566 | 994.5128 | 432 | 948.9164 | 437 | E2F1 | LNCaP | Prostate | GSM24924 | 9586 | 91433 |
| 193.9566 | 40 | 351.4109 | 62 | 154.104 | 49 | 101.4359 | 42 | ZNF341 | None | None | GSM28770 | 1715 | 91448 |
| 103.0291 | 24 | 247.202 | 44 | 76.63187 | 29 | 76.97382 | 31 | ZNF341 | None | None | GSM28770 | 1275 | 91449 |
| 1132.149 | 436 | 1158.649 | 558 | 848.857 | 410 | 802.5512 | 410 | RBP2 | M.M. | None | GSM31119 | 13644 | 91459 |
| 1143.97 | 334 | 1165.138 | 425 | 871.3979 | 313 | 826.1502 | 310 | RBP2 | M.M. | None | GSM31119 | 9208 | 91460 |
| 1147.415 | 384 | 1160.291 | 478 | 860.1561 | 346 | 849.1035 | 380 | RBP2 | M.M. | None | GSM31119 | 10624 | 91461 |
| 1185.573 | 223 | 1234.506 | 314 | 943.6966 | 222 | 914.4257 | 230 | MYB | None | None | GSM25853 | 5389 | 91489 |
| 1049.332 | 233 | 971.3679 | 231 | 753.9466 | 238 | 899.5551 | 350 | AR | 22RV1 | Prostate | GSM24956 | 9238 | 91524 |
| 1097.384 | 474 | 1027.189 | 491 | 876.4175 | 546 | 928.91 | 699 | AR | 22RV1 | Prostate | GSM24956 | 19763 | 91525 |
| 911.3807 | 590 | 847.7269 | 635 | 736.1137 | 782 | 800.422 | 1016 | AR | 22RV1 | Prostate | GSM24956 | 52934 | 91526 |
| 894.442 | 586 | 819.5893 | 609 | 721.0464 | 772 | 769.0732 | 977 | AR | 22RV1 | Prostate | GSM24956 | 54628 | 91527 |
| 1145.298 | 714 | 1094.51 | 793 | 869.8902 | 792 | 929.3293 | 1017 | AR | 22RV1 | Prostate | GSM24956 | 34627 | 91528 |
| 1220.326 | 695 | 1227.348 | 843 | 921.6233 | 707 | 929.7727 | 822 | BRD4 | 22RV1 | Prostate | GSM24956 | 22591 | 91529 |
| 1067.906 | 644 | 1033.229 | 730 | 839.2254 | 762 | 885.9084 | 962 | AR | 22RV1 | Prostate | GSM24956 | 36394 | 91530 |
| 1073.032 | 587 | 1002.469 | 620 | 821.7316 | 651 | 872.9034 | 838 | AR | 22RV1 | Prostate | GSM24956 | 30389 | 91531 |
| 1250.623 | 899 | 1247.252 | 1064 | 895.4863 | 924 | 897.7009 | 1066 | BRD4 | 22RV1 | Prostate | GSM24956 | 37218 | 91532 |
| 1288.941 | 903 | 1286.501 | 1076 | 933.9354 | 944 | 947.1435 | 1112 | BRD4 | 22RV1 | Prostate | GSM24956 | 35940 | 91533 |
| 809.4376 | 522 | 837.2806 | 670 | 590.0385 | 613 | 577.2224 | 678 | RAD21 | THP-1 | Blood | GSM31119 | 57819 | 91543 |
| 699.1206 | 390 | 749.5935 | 534 | 514.1029 | 501 | 503.9169 | 541 | CTCF | GM09236 | B-lymphoc | GSM32117 | 54707 | 91552 |
| 684.8524 | 363 | 748.463 | 514 | 486.449 | 477 | 479.5826 | 516 | CTCF | GM09236 | B-lymphoc | GSM32117 | 51469 | 91553 |
| 700.3713 | 383 | 749.5129 | 524 | 514.9602 | 492 | 509.8113 | 533 | CTCF | GM09237 | B-lymphoc | GSM32117 | 52156 | 91554 |
| 711.202 | 430 | 742.3088 | 565 | 537.0498 | 551 | 499.7984 | 571 | CTCF | GM09237 | B-lymphoc | GSM32117 | 59436 | 91555 |
| 569.9152 | 265 | 715.5688 | 339 | 348.5928 | 329 | 384.2915 | 370 | RAD21 | THP-1 | Blood | GSM31119 | 32963 | 91558 |
| 1209.815 | 435 | 1219.034 | 536 | 909.7673 | 395 | 864.2652 | 394 | BRD4 | None | None | GSM29425 | 10884 | 91582 |
| 1180.973 | 454 | 1188.478 | 550 | 912.1916 | 427 | 842.2265 | 406 | BRD4 | None | None | GSM29425 | 10769 | 91583 |
| 1150.006 | 392 | 1171.807 | 496 | 888.6061 | 376 | 841.133 | 374 | BRD4 | None | None | GSM29425 | 10207 | 91585 |
| 1203.043 | 509 | 1217.707 | 626 | 898.2698 | 457 | 841.8354 | 446 | BRD4 | None | None | GSM29425 | 12568 | 91586 |
| 1145.463 | 404 | 1164.546 | 513 | 880.4432 | 390 | 822.5092 | 375 | BRD4 | None | None | GSM29425 | 11055 | 91587 |
| 1198.934 | 433 | 1208.1 | 531 | 931.2715 | 414 | 851.6971 | 377 | BRD4 | None | None | GSM29425 | 10393 | 91588 |
| 1190.978 | 179 | 1198.952 | 222 | 765.2673 | 167 | 667.3854 | 164 | BRD4 | None | None | GSM29425 | 3639 | 91589 |
| 595.4321 | 125 | 872.069 | 170 | 363.8535 | 136 | 448.4497 | 161 | AFF1 | H2171 | Lung | GSM30739 | 5984 | 91608 |
| 759.9084 | 525 | 798.8345 | 688 | 613.2567 | 726 | 595.3698 | 799 | CTCF | MCF-7 | Breast | GSM33571 | 67866 | 91661 |
| 782.1007 | 517 | 833.6375 | 692 | 623.4624 | 694 | 594.2335 | 740 | CTCF | MCF-7 | Breast | GSM33571 | 61026 | 91662 |
| 762.4195 | 536 | 795.761 | 692 | 603.6142 | 722 | 591.0573 | 802 | CTCF | MCF-7 | Breast | GSM33571 | 69801 | 91663 |

| | | | | | | | | | | | | | |
|---|---|---|---|---|---|---|---|---|---|---|---|---|---|
| 810.5466 | 587 | 864.3458 | 785 | 611.2273 | 732 | 569.6059 | 762 | CTCF | MCF-7 | Breast | GSM33571 | 69760 | 91664 |
| 713.0747 | 466 | 755.2901 | 619 | 589.767 | 680 | 575.9154 | 751 | CTCF | MCF-7 | Breast | GSM33571 | 67885 | 91665 |
| 652.4084 | 363 | 691.8075 | 479 | 540.8856 | 532 | 540.4142 | 603 | CTCF | MCF-7 | Breast | GSM33571 | 57142 | 91666 |
| 758.2556 | 469 | 808.9045 | 633 | 619.4214 | 650 | 593.1067 | 701 | CTCF | MCF-7 | Breast | GSM33571 | 56099 | 91667 |
| 772.0161 | 555 | 812.098 | 728 | 608.7695 | 744 | 598.395 | 835 | CTCF | MCF-7 | Breast | GSM33571 | 72335 | 91668 |
| 813.8643 | 610 | 875.8566 | 821 | 646.7388 | 819 | 617.8869 | 885 | CTCF | MCF-7 | Breast | GSM33571 | 74409 | 91669 |
| 766.1604 | 542 | 842.3262 | 758 | 635.913 | 779 | 613.1837 | 850 | CTCF | MCF-7 | Breast | GSM33571 | 71176 | 91670 |
| 741.5225 | 172 | 914.4186 | 212 | 380.2106 | 175 | 292.3346 | 166 | BMI1 | LCL | Blood | GSM33844 | 9405 | 91707 |
| 1171.274 | 248 | 1198.396 | 326 | 966.5378 | 273 | 943.2157 | 293 | ESR1 | MCF-7 | Breast | GSM34147 | 6149 | 91709 |
| 1220.591 | 474 | 1223.093 | 575 | 957.0882 | 484 | 930.0708 | 515 | KDM6A | MCF-7 | Breast | GSM34147 | 12630 | 91736 |
| 1291.226 | 288 | 1316.142 | 371 | 997.8055 | 245 | 970.8126 | 260 | TET2 | MCF-7 | Breast | GSM34147 | 4759 | 91737 |
| 101.7744 | 51 | 83.01979 | 52 | 121.7269 | 83 | 221.5581 | 117 | EPAS1 | HKC8 | Kidney | GSM34178 | 5955 | 91772 |
| 55.04584 | 22 | 52.59295 | 24 | 79.19546 | 39 | 309.587 | 84 | EPAS1 | HKC8 | Kidney | GSM34178 | 2187 | 91773 |
| 1106.098 | 121 | 1284.704 | 157 | 596.2099 | 111 | 641.0317 | 122 | RBP2 | MCF-7 | Breast | GSM70075 | 2142 | 91774 |
| 1269.972 | 278 | 1287.498 | 359 | 970.6631 | 244 | 968.7794 | 274 | RBP2 | MCF-7 | Breast | GSM70075 | 5563 | 91801 |
| 735.4314 | 325 | 788.7717 | 456 | 482.415 | 405 | 558.7031 | 464 | CTCF | None | Breast | GSM30172 | 39980 | 91802 |
| 504.3419 | 281 | 688.9174 | 379 | 320.0921 | 368 | 392.3429 | 430 | CTCF | None | Breast | GSM30172 | 44637 | 91803 |
| 717.2156 | 521 | 778.9568 | 713 | 525.5248 | 657 | 530.8867 | 757 | CTCF | None | Breast | GSM30172 | 79885 | 91804 |
| 726.984 | 461 | 765.7777 | 609 | 584.8109 | 640 | 571.7537 | 708 | CTCF | MCF-7 | Breast | GSM33571 | 62433 | 91919 |
| 766.7267 | 554 | 812.16 | 731 | 629.0872 | 786 | 603.1433 | 852 | CTCF | MCF-7 | Breast | GSM33571 | 73251 | 91920 |
| 765.0486 | 556 | 813.144 | 737 | 597.2413 | 738 | 591.5451 | 831 | CTCF | MCF-7 | Breast | GSM33571 | 73140 | 91922 |
| 729.8346 | 503 | 772.1486 | 665 | 596.5895 | 715 | 580.0934 | 784 | CTCF | MCF-7 | Breast | GSM33571 | 70539 | 91923 |
| 699.0414 | 390 | 727.0496 | 507 | 581.3514 | 569 | 559.6456 | 616 | CTCF | MCF-7 | Breast | GSM33571 | 54203 | 91924 |
| 721.3252 | 440 | 767.4237 | 590 | 600.6016 | 636 | 570.2204 | 677 | CTCF | MCF-7 | Breast | GSM33571 | 58118 | 91925 |
| 710.5834 | 458 | 754.3245 | 613 | 573.1627 | 647 | 561.5316 | 716 | CTCF | MCF-7 | Breast | GSM33571 | 66077 | 91926 |
| 762.6903 | 479 | 804.8093 | 635 | 614.1149 | 646 | 584.3419 | 688 | CTCF | MCF-7 | Breast | GSM33571 | 56281 | 91927 |
| 544.8084 | 344 | 659.9195 | 455 | 515.8276 | 522 | 515.8401 | 583 | CTCF | MCF-7 | Breast | GSM33571 | 58412 | 91928 |
| 746.2948 | 485 | 766.3628 | 615 | 618.3314 | 693 | 599.3599 | 759 | CTCF | MCF-7 | Breast | GSM33571 | 62463 | 91929 |
| 79.33083 | 49 | 152.9442 | 75 | 48.45487 | 63 | 87.55233 | 86 | TCF7L1 | RPMI-8402 | Blood | GSM95760 | 7628 | 9193 |
| 56.34601 | 74 | 94.13948 | 103 | 90.25124 | 143 | 79.30379 | 149 | TCF7L1 | RPMI-8402 | Blood | GSM95760 | 18991 | 9194 |
| 88.25496 | 50 | 144.5912 | 71 | 50.7268 | 62 | 74.40109 | 78 | GATA3 | RPMI-8402 | Blood | GSM95760 | 7235 | 9195 |
| 61.63312 | 52 | 58.70008 | 58 | 27.97968 | 63 | 45.53628 | 81 | GATA3 | RPMI-8402 | Blood | GSM95760 | 10104 | 9196 |
| 785.7337 | 235 | 887.3309 | 332 | 476.7811 | 273 | 358.5268 | 260 | TCF12 | RPMI-8402 | Blood | GSM95761 | 18914 | 9197 |
| 905.4101 | 543 | 936.6163 | 698 | 660.909 | 600 | 605.1634 | 597 | TCF12 | RPMI-8402 | Blood | GSM95761 | 45045 | 9198 |
| 132.1741 | 35 | 187.9916 | 47 | 79.50237 | 39 | 41.8818 | 31 | CTCF | HepG2 | Liver | GSM30199 | 2261 | 91983 |
| 207.2399 | 167 | 286.0406 | 216 | 183.6659 | 253 | 109.3903 | 232 | TAL1 | RPMI-8402 | Blood | GSM95761 | 31792 | 9199 |
| 311.4265 | 109 | 531.396 | 159 | 182.0053 | 125 | 270.0569 | 160 | CTCF | HepG2 | Liver | GSM30199 | 9249 | 91993 |
| 874.3262 | 387 | 895.2408 | 492 | 674.6425 | 446 | 636.1207 | 457 | RAD21 | T47D | Mammary | GSM34155 | 27604 | 92003 |
| 1160.042 | 588 | 1192.927 | 741 | 856.0054 | 544 | 796.5598 | 537 | T | None | None | GSM27429 | 17960 | 92005 |
| 1128.123 | 664 | 1179.799 | 859 | 811.5692 | 615 | 763.9525 | 626 | T | None | None | GSM27429 | 22511 | 92006 |
| 210.4279 | 108 | 215.1688 | 122 | 100.1863 | 119 | 87.64039 | 123 | T | None | None | GSM27429 | 11815 | 92007 |
| 1086.572 | 491 | 1128.959 | 647 | 805.6588 | 461 | 760.9279 | 467 | T | None | None | GSM27429 | 16146 | 92008 |
| 244.9946 | 150 | 323.276 | 191 | 202.5667 | 216 | 138.5875 | 204 | TAL1 | RPMI-8402 | Blood | GSM95761 | 23268 | 9201 |
| 1201.865 | 502 | 1236.123 | 647 | 888.9962 | 452 | 878.8165 | 498 | RBP2 | THP-1 | Blood | GSM31120 | 13719 | 92039 |
| 1251.277 | 272 | 1270.743 | 344 | 970.0152 | 247 | 926.5315 | 246 | HCFC1 | HeLa | Cervix | GSM77942 | 5502 | 9205 |
| 1155.66 | 204 | 1189.722 | 270 | 751.2713 | 188 | 784.3323 | 204 | HCFC1 | HeLa | Cervix | GSM77942 | 5371 | 9206 |
| 520.2937 | 91 | 580.7839 | 106 | 509.5925 | 124 | 729.1326 | 160 | CBX8 | PC-3 | Prostate | GSM25722 | 3364 | 92062 |
| 908.7876 | 673 | 893.7548 | 790 | 673.2202 | 794 | 677.1859 | 914 | CBX8 | PC-3 | Prostate | GSM25722 | 65321 | 92064 |
| 992.332 | 628 | 1023.786 | 800 | 714.9883 | 663 | 689.847 | 716 | PWWP2A | HeLa | None | GSM29829 | 42754 | 92097 |
| 1067.889 | 737 | 1095.864 | 924 | 752.8289 | 759 | 717.9353 | 803 | PWWP2A | HeLa | None | GSM29829 | 45893 | 92098 |
| 747.4972 | 103 | 910.2898 | 126 | 314.0493 | 85 | 256.9228 | 81 | ZNF143 | HeLa | Cervix | GSM77943 | 2568 | 9210 |
| 210.0682 | 39 | 107.242 | 29 | 79.41088 | 32 | 190.079 | 54 | TP53 | A549 | Lung | GSM26773 | 1505 | 92109 |
| 789.5896 | 388 | 756.5158 | 439 | 693.9574 | 595 | 704.8489 | 696 | FOXA1 | WS8 | None | GSM30964 | 39000 | 92112 |
| 781.7386 | 397 | 750.7666 | 447 | 687.3059 | 609 | 688.4248 | 696 | FOXA1 | WS8 | None | GSM30964 | 41035 | 92113 |
| 803.7102 | 376 | 777.5661 | 430 | 697.0384 | 561 | 710.4458 | 657 | FOXA1 | WS8 | None | GSM30964 | 35383 | 92114 |
| 739.6966 | 239 | 777.0157 | 272 | 717.4866 | 369 | 721.8366 | 424 | GRHL2 | WS8 | None | GSM30964 | 19817 | 92115 |
| 809.5918 | 314 | 809.7162 | 379 | 720.8906 | 483 | 717.4955 | 545 | GRHL2 | WS8 | None | GSM30964 | 26807 | 92116 |
| 819.2067 | 302 | 817.2317 | 368 | 717.5296 | 455 | 703.5256 | 499 | GRHL2 | WS8 | None | GSM30964 | 25394 | 92117 |
| 1236.001 | 279 | 1256.455 | 357 | 943.6761 | 243 | 915.159 | 255 | GABPA | HeLa | Cervix | GSM77943 | 5913 | 9212 |
| 249.0399 | 65 | 355.3801 | 87 | 160.5169 | 74 | 204.3578 | 89 | TFAP2C | hESC | None | GSM29017 | 3788 | 92121 |
| 251.6042 | 75 | 478.2445 | 117 | 226.1521 | 102 | 247.8129 | 114 | TFAP2C | hESC | None | GSM29017 | 5098 | 92125 |
| 1030.076 | 147 | 1168.631 | 199 | 622.486 | 150 | 478.7676 | 138 | YY1 | HeLa | Cervix | GSM77943 | 4026 | 9213 |
| 1160.247 | 485 | 1167.913 | 604 | 864.4272 | 457 | 813.0962 | 456 | CTRL | HL-60 | Blood | GSM28364 | 15694 | 92149 |
| 1115.275 | 475 | 1136.12 | 599 | 826.8775 | 450 | 771.5105 | 444 | CTRL | HL-60 | Blood | GSM28364 | 17161 | 92150 |
| 1297.868 | 506 | 1345.85 | 667 | 997.7504 | 472 | 962.1648 | 489 | POLR2B | HeLa | Cervix | GSM77943 | 10925 | 9216 |
| 127.2545 | 69 | 147.5628 | 83 | 62.04464 | 79 | 148.3624 | 121 | TBX5 | None | Blood | GSM21839 | 9047 | 92169 |
| 702.7499 | 377 | 762.4945 | 531 | 540.3267 | 494 | 483.9105 | 516 | CTCF | GM04025 | B-lymphoc | GSM32117 | 50871 | 92197 |
| 1190.007 | 393 | 1208.126 | 494 | 901.8536 | 350 | 874.4224 | 368 | TOP1 | HEK293 | Embryonic | GSM27383 | 9138 | 92223 |
| 935.1677 | 524 | 944.0498 | 648 | 672.2572 | 542 | 627.2915 | 551 | SET | SET2 | None | GSM26868 | 35411 | 92238 |
| 239.6335 | 391 | 334.9317 | 499 | 160.0857 | 586 | 129.2158 | 617 | SPI1 | M.M. | None | GSM31119 | 123024 | 92247 |
| 307.7328 | 471 | 446.9141 | 606 | 194.5043 | 694 | 187.2096 | 762 | SPI1 | M.M. | None | GSM31119 | 147358 | 92248 |
| 274.985 | 445 | 397.2156 | 572 | 172.0002 | 657 | 169.3176 | 725 | SPI1 | M.M. | None | GSM31119 | 141567 | 92249 |

| | | | | | | | | | | | | | |
|---|---|---|---|---|---|---|---|---|---|---|---|---|---|
| 54.73724 | 41 | 65.08766 | 50 | 36.86394 | 55 | 48.11782 | 66 | NFATC2 | None | Blood | GSM28100 | 6682 | 92277 |
| 227.5625 | 79 | 265.188 | 95 | 125.6513 | 89 | 129.4584 | 97 | CBX3 | HCT-116 | Colon | GSM69972 | 7044 | 9228 |
| 596.5738 | 64 | 504.2615 | 62 | 313.4347 | 57 | 278.0413 | 56 | KMT2D | NT2-D1 | None | GSM32734 | 1030 | 92283 |
| 289.5129 | 136 | 283.0598 | 151 | 155.4412 | 160 | 152.6695 | 172 | CBX3 | HCT-116 | Colon | GSM69972 | 16688 | 9230 |
| 1085.807 | 640 | 1141.849 | 842 | 798.9602 | 648 | 785.0047 | 717 | MYC | SW1271 | Lung | GSM30739 | 29795 | 92301 |
| 965.3517 | 710 | 991.3549 | 886 | 688.9974 | 768 | 645.0312 | 796 | MYC | SW1271 | Lung | GSM30739 | 54081 | 92302 |
| 749.2541 | 113 | 1192.733 | 180 | 352.5812 | 101 | 422.0057 | 118 | ELL2 | SW1271 | Lung | GSM30739 | 3259 | 92303 |
| 43.10483 | 13 | 40.65875 | 14 | 4.936782 | 7 | 101.2495 | 32 | CBX3 | HCT-116 | Colon | GSM69973 | 1070 | 9231 |
| 5.785782 | 7 | 19.76112 | 14 | 4.464024 | 11 | 50.1913 | 32 | CBX3 | HCT-116 | Colon | GSM69973 | 2118 | 9232 |
| 480.8895 | 263 | 726.5789 | 387 | 337.4861 | 351 | 346.9589 | 385 | CTCF | THP-1 | Blood | GSM31120 | 37583 | 92326 |
| 275.0959 | 75 | 308.3267 | 88 | 193.0603 | 90 | 298.4102 | 119 | SOX4 | HCC1954 | Breast | GSM28073 | 4762 | 92329 |
| 651.0599 | 297 | 725.7851 | 382 | 412.6817 | 373 | 397.9485 | 399 | RAD21 | THP-1 | Blood | GSM31120 | 36443 | 92330 |
| 653.9916 | 313 | 748.0293 | 435 | 400.4228 | 391 | 351.4581 | 405 | RAD21 | THP-1 | Blood | GSM31120 | 40368 | 92335 |
| 1221.676 | 437 | 1283.031 | 600 | 931.5408 | 404 | 894.4957 | 417 | AFF4 | 293T | Embryonic | GSM33605 | 10899 | 92336 |
| 1185.588 | 262 | 1185.541 | 312 | 911.7561 | 233 | 887.3705 | 249 | AFF4 | 293T | Embryonic | GSM33605 | 6036 | 92337 |
| 1215.596 | 649 | 1256.456 | 837 | 905.4178 | 624 | 835.0917 | 605 | MYC | 293T | Embryonic | GSM33605 | 19315 | 92338 |
| 1206.559 | 386 | 1214.99 | 474 | 918.0472 | 349 | 883.7288 | 362 | AFF1 | 293T | Embryonic | GSM33605 | 9246 | 92340 |
| 1177.929 | 383 | 1218.039 | 503 | 907.0433 | 363 | 869.206 | 371 | AFF1 | 293T | Embryonic | GSM33605 | 10192 | 92341 |
| 553.9821 | 304 | 713.241 | 428 | 351.4059 | 392 | 409.0935 | 450 | CTCF | THP-1 | Blood | GSM31120 | 44198 | 92343 |
| 46.37699 | 121 | 54.98846 | 148 | 21.12668 | 173 | 25.69285 | 200 | BATF3 | KK1 | None | GSM24816 | 43332 | 92360 |
| 1220.039 | 218 | 1268.026 | 299 | 967.3375 | 209 | 938.1415 | 218 | KDM5A | HCT-116 | HCT116 | GSM28629 | 4458 | 92366 |
| 1217.817 | 237 | 1251.374 | 316 | 965.9079 | 228 | 934.0878 | 234 | KDM5A | HCT-116 | HCT116 | GSM28629 | 4922 | 92367 |
| 13.07184 | 7 | 29.04921 | 12 | 10.39519 | 10 | 114.23 | 35 | CBX3 | HCT-116 | Colon | GSM89822 | 1119 | 9237 |
| 79.88394 | 89 | 71.89478 | 98 | 73.14354 | 141 | 104.0302 | 173 | AR | None | Epididymis | GSM29290 | 20993 | 92373 |
| 805.6953 | 295 | 832.6631 | 384 | 621.4035 | 367 | 515.4714 | 366 | RUNX1 | None | Epididymis | GSM29290 | 25932 | 92374 |
| 64.30712 | 29 | 87.93325 | 38 | 40.26952 | 36 | 122.9894 | 64 | AR | None | Epididymis | GSM29290 | 3484 | 92375 |
| 419.8672 | 138 | 463.0749 | 161 | 293.713 | 171 | 291.2534 | 183 | RUNX1 | None | Epididymis | GSM29290 | 11423 | 92377 |
| 787.8438 | 277 | 746.2094 | 304 | 741.4148 | 486 | 757.3646 | 576 | ESR1 | MCF-7 | Breast | GSM27525 | 26012 | 92378 |
| 794.4421 | 380 | 769.4395 | 435 | 727.5722 | 622 | 748.482 | 745 | ESR1 | MCF-7 | Breast | GSM27525 | 37219 | 92379 |
| 662.0938 | 230 | 612.9447 | 247 | 738.6343 | 407 | 788.4075 | 530 | ESR1 | MCF-7 | Breast | GSM27525 | 21316 | 92380 |
| 789.3849 | 325 | 735.3737 | 345 | 718.5158 | 531 | 761.9861 | 672 | ESR1 | MCF-7 | Breast | GSM27525 | 31670 | 92381 |
| 601.3121 | 223 | 592.408 | 247 | 734.2928 | 413 | 764.7414 | 508 | ESR1 | MCF-7 | Breast | GSM27525 | 21787 | 92382 |
| 615.5515 | 208 | 580.442 | 225 | 769.2604 | 393 | 789.1029 | 475 | ESR1 | MCF-7 | Breast | GSM27525 | 18479 | 92383 |
| 446.0158 | 261 | 679.6167 | 355 | 305.3493 | 344 | 273.7116 | 360 | CTCF | None | None | GSM29487 | 36705 | 92420 |
| 784.4639 | 440 | 786.5261 | 536 | 568.9539 | 497 | 431.7395 | 475 | BRD7 | MOLM-13 | None | GSM30943 | 45745 | 92423 |
| 874.6366 | 384 | 871.2079 | 464 | 648.1374 | 421 | 537.3489 | 397 | BRD9 | MOLM-13 | None | GSM30943 | 29322 | 92424 |
| 704.1794 | 501 | 690.2783 | 591 | 518.8384 | 625 | 490.4829 | 668 | SMARCA4 | MOLM-13 | None | GSM30943 | 71039 | 92425 |
| 732.4571 | 577 | 771.2156 | 749 | 542.2407 | 732 | 512.8069 | 783 | CTCF | EOL1 | None | GSM30943 | 86664 | 92426 |
| 815.6551 | 573 | 795.5839 | 666 | 570.3108 | 635 | 550.4763 | 692 | DPF2 | EOL1 | None | GSM30943 | 64000 | 92428 |
| 836.8204 | 476 | 825.3018 | 563 | 602.1231 | 523 | 556.0385 | 527 | BRD7 | EOL1 | None | GSM30943 | 44268 | 92429 |
| 756.6068 | 638 | 753.346 | 766 | 533.9724 | 763 | 509.8919 | 829 | SMARCC1 | EOL1 | None | GSM30943 | 90323 | 92430 |
| 1182.934 | 352 | 1184.247 | 432 | 892.5067 | 316 | 863.2915 | 331 | RBP2 | M.M. | None | GSM31119 | 8811 | 92432 |
| 1152.472 | 467 | 1168.182 | 584 | 849.4648 | 419 | 826.0761 | 448 | RBP2 | M.M. | None | GSM31119 | 13869 | 92438 |
| 1145.641 | 447 | 1144.754 | 536 | 832.6132 | 386 | 808.6397 | 409 | RBP2 | M.M. | None | GSM31119 | 13157 | 92443 |
| 1135.626 | 364 | 1167.446 | 478 | 870.5544 | 355 | 846.7047 | 380 | MYB | None | None | GSM25853 | 11488 | 92444 |
| 1110.979 | 542 | 1149.83 | 698 | 827.4254 | 522 | 788.4519 | 538 | BRD4 | K562 | Bone Marr | GSM30315 | 18772 | 92478 |
| 1135.143 | 541 | 1171.401 | 696 | 845.9953 | 519 | 808.8209 | 538 | BRD4 | K562 | Bone Marr | GSM30315 | 17793 | 92479 |
| 1117.65 | 561 | 1166.892 | 739 | 830.9849 | 544 | 789.24 | 555 | BRD4 | K562 | Bone Marr | GSM30315 | 19732 | 92480 |
| 1146.64 | 280 | 1164.489 | 354 | 861.8546 | 248 | 841.9315 | 265 | SIRT6 | L-C-B | None | GSM27467 | 7935 | 92514 |
| 958.4574 | 203 | 958.7567 | 247 | 504.3098 | 204 | 393.1963 | 194 | RAD21 | T47D | Mammary | GSM34155 | 9153 | 92516 |
| 745.6977 | 524 | 770.8274 | 666 | 540.0774 | 627 | 528.3149 | 698 | SMAD3 | MDA-MB-2 | Breast | GSM28073 | 67396 | 92518 |
| 464.9323 | 298 | 656.3389 | 401 | 306.1307 | 396 | 362.0453 | 457 | SMAD3 | MDA-MB-2 | Breast | GSM28073 | 48247 | 92519 |
| 967.508 | 303 | 1006.87 | 403 | 742.2053 | 316 | 694.736 | 317 | RAD21 | T47D | Mammary | GSM34155 | 14255 | 92550 |
| 945.8067 | 122 | 1194.067 | 154 | 644.7755 | 130 | 560.9659 | 127 | BRD2 | H23 | Lung | GSM28011 | 2641 | 92552 |
| 940.7267 | 489 | 953.9864 | 611 | 699.3053 | 526 | 656.2158 | 536 | RAD21 | T47D | Mammary | GSM34155 | 30840 | 92585 |
| 482.7859 | 246 | 729.5648 | 351 | 347.1422 | 323 | 444.3082 | 385 | GLIS3 | hESC | None | GSM29465 | 29690 | 92597 |
| 334.0916 | 50 | 413.8721 | 62 | 200.9794 | 51 | 600.3455 | 101 | AR | LNCaP | Prostate | GSM77536 | 1471 | 9261 |
| 1137.074 | 483 | 1177.135 | 626 | 860.5556 | 470 | 834.384 | 499 | RBP2 | THP-1 | Blood | GSM31120 | 15890 | 92643 |
| 682.4021 | 356 | 699.4759 | 454 | 458.3362 | 462 | 454.7359 | 501 | RAD21 | THP-1 | Blood | GSM31120 | 50111 | 92647 |
| 221.1229 | 103 | 257.797 | 124 | 104.1021 | 114 | 123.7788 | 132 | RAD21 | THP-1 | Blood | GSM31120 | 12038 | 92657 |
| 697.4373 | 373 | 713.3768 | 471 | 495.7982 | 479 | 525.1633 | 539 | RAD21 | THP-1 | Blood | GSM31120 | 50870 | 92660 |
| -0.235052 | 0 | 0 | 1 | -0.277664 | 1 | -0.475033 | 1 | FOXA1 | None | Prostate | GSM31408 | 1601 | 92669 |
| 990.427 | 572 | 1007.954 | 714 | 723.6761 | 597 | 676.2913 | 607 | BRD9 | SYO1 | None | GSM30943 | 35314 | 92671 |
| 854.3921 | 553 | 856.213 | 668 | 625.6099 | 621 | 597.5137 | 663 | SS18 | SYO1 | None | GSM30943 | 48083 | 92672 |
| 1010.989 | 219 | 1004.281 | 267 | 580.5115 | 214 | 464.0163 | 205 | BRD9 | Jurkat | None | GSM30943 | 9507 | 92673 |
| 1122.367 | 326 | 1111.329 | 381 | 834.5989 | 292 | 789.8649 | 290 | BRD9 | TTC1240 | None | GSM30943 | 10319 | 92674 |
| 711.3171 | 499 | 800.481 | 718 | 551.6292 | 667 | 523.0322 | 711 | CTCF | Jurkat | None | GSM30943 | 73505 | 92675 |
| 1108.645 | 432 | 1104.647 | 518 | 802.271 | 386 | 772.4924 | 406 | SMARCB1 | TTC1240 | None | GSM30943 | 15696 | 92676 |
| 1182.614 | 436 | 1205.951 | 560 | 884.5677 | 394 | 848.8806 | 408 | RBP2 | THP-1 | Blood | GSM31120 | 11749 | 92690 |
| 1136.106 | 521 | 1164.447 | 670 | 835.6381 | 491 | 812.2887 | 523 | RBP2 | THP-1 | Blood | GSM31120 | 18204 | 92691 |
| 532.4004 | 282 | 721.8764 | 398 | 356.7417 | 369 | 349.9139 | 398 | CTCF | THP-1 | Blood | GSM31120 | 39597 | 92695 |
| 492.362 | 270 | 709.6152 | 381 | 341.2929 | 359 | 378.8457 | 405 | CTCF | THP-1 | Blood | GSM31120 | 38815 | 92699 |

| | | | | | | | | | | | | | |
|---|---|---|---|---|---|---|---|---|---|---|---|---|---|
| 366.7434 | 53 | 381.7092 | 59 | 129.8956 | 40 | 208.9912 | 55 | USF2 | K562 | Bone Marr | GSM30317 | 1428 | 92713 |
| 509.8281 | 76 | 423.9526 | 74 | 180.7659 | 58 | 223.6861 | 69 | USF2 | K562 | Bone Marr | GSM30317 | 2138 | 92714 |
| 267.519 | 132 | 404.8091 | 180 | 218.6874 | 187 | 330.2978 | 239 | CTCF | HCC1806 | Breast | GSM33175 | 18100 | 92754 |
| 861.9364 | 426 | 866.8139 | 525 | 621.8382 | 449 | 584.3437 | 465 | BRD4 | MV4-11 | Blood | GSM27166 | 33116 | 92766 |
| 904.081 | 371 | 895.6787 | 445 | 646.3556 | 375 | 633.4225 | 411 | BRD4 | MV4-11 | Blood | GSM27166 | 26115 | 92767 |
| 880.8202 | 493 | 888.3403 | 604 | 625.2882 | 510 | 592.176 | 532 | BRD4 | MV4-11 | Blood | GSM27166 | 37184 | 92768 |
| 949.5385 | 733 | 992.0396 | 938 | 678.5764 | 834 | 641.8302 | 885 | BRD4 | MV4-11 | Blood | GSM27166 | 70192 | 92769 |
| 877.0577 | 312 | 878.539 | 384 | 592.6598 | 331 | 538.5481 | 342 | BRD4 | MV4-11 | Blood | GSM27166 | 21822 | 92770 |
| 916.019 | 762 | 953.2107 | 965 | 647.7291 | 881 | 618.794 | 953 | BRD4 | MV4-11 | Blood | GSM27166 | 84279 | 92771 |
| 432.3619 | 235 | 704.2108 | 331 | 309.9948 | 314 | 356.7218 | 359 | CTCF | MIA PaCa- | Pancreatic | GSM26536 | 32093 | 92837 |
| 568.2425 | 105 | 607.4107 | 119 | 261.8578 | 95 | 183.5312 | 85 | SUZ12 | Aska | None | GSM28875 | 3672 | 92860 |
| 96.96633 | 33 | 201.7592 | 54 | 62.1333 | 38 | 69.36002 | 43 | SUZ12 | Aska | None | GSM28875 | 2247 | 92861 |
| 318.5406 | 169 | 458.8863 | 224 | 221.2006 | 224 | 241.904 | 251 | CTCF | M.M. | None | GSM31119 | 23390 | 92905 |
| 333.5491 | 188 | 535.3935 | 261 | 229.3223 | 250 | 274.6728 | 290 | CTCF | M.M. | None | GSM31119 | 27356 | 92906 |
| 336.3834 | 184 | 509.0144 | 249 | 214.4171 | 237 | 237.5813 | 267 | CTCF | M.M. | None | GSM31119 | 26149 | 92907 |
| 322.4398 | 185 | 514.1 | 256 | 229.9709 | 250 | 258.093 | 283 | CTCF | M.M. | None | GSM31119 | 27334 | 92908 |
| 323.4106 | 172 | 448.5125 | 224 | 212.0639 | 223 | 237.0235 | 252 | CTCF | M.M. | None | GSM31119 | 23800 | 92909 |
| 343.2724 | 188 | 531.6159 | 257 | 251.0749 | 255 | 247.805 | 275 | CTCF | M.M. | None | GSM31119 | 26697 | 92910 |
| 341.599 | 186 | 478.1736 | 243 | 239.1599 | 248 | 244.4258 | 271 | CTCF | M.M. | None | GSM31119 | 26305 | 92911 |
| 523.9619 | 264 | 659.5074 | 328 | 315.3497 | 329 | 344.6961 | 369 | RAD21 | M.M. | None | GSM31119 | 34955 | 92912 |
| 498.0112 | 261 | 670.5183 | 334 | 316.7869 | 334 | 372.8911 | 385 | RAD21 | M.M. | None | GSM31119 | 35893 | 92913 |
| 478.2931 | 255 | 606.8535 | 318 | 304.3879 | 327 | 353.3968 | 375 | RAD21 | M.M. | None | GSM31119 | 35494 | 92914 |
| 573.2899 | 294 | 713.8881 | 398 | 381.9218 | 382 | 350.0587 | 402 | RAD21 | THP-1 | Blood | GSM31120 | 39995 | 92934 |
| 1180.81 | 525 | 1221.186 | 686 | 873.7121 | 487 | 836.6143 | 503 | RBP2 | THP-1 | Blood | GSM31120 | 15766 | 92935 |
| 1192.016 | 543 | 1233.335 | 711 | 875.5082 | 496 | 849.3061 | 529 | RBP2 | THP-1 | Blood | GSM31120 | 16109 | 92936 |
| 555.0594 | 269 | 726.731 | 364 | 363.6364 | 344 | 368.7021 | 375 | RAD21 | THP-1 | Blood | GSM31120 | 34285 | 92938 |
| 703.027 | 319 | 738.2723 | 417 | 425.6425 | 394 | 410.9106 | 422 | RAD21 | THP-1 | Blood | GSM31120 | 39282 | 92939 |
| 667.2458 | 313 | 747.9593 | 431 | 439.3045 | 401 | 399.2039 | 420 | RAD21 | THP-1 | Blood | GSM31120 | 39860 | 92940 |
| 562.3971 | 248 | 736.2904 | 324 | 310.5228 | 293 | 330.321 | 325 | RAD21 | THP-1 | Blood | GSM31120 | 28948 | 92942 |
| 473.6741 | 229 | 583.076 | 282 | 276.5594 | 280 | 318.1524 | 320 | RAD21 | THP-1 | Blood | GSM31120 | 28961 | 92943 |
| 662.9121 | 238 | 686.5067 | 270 | 307.8054 | 255 | 323.3963 | 281 | PDX1 | None | Pancreas | GSM28580 | 23451 | 92949 |
| 583.7295 | 214 | 659.0882 | 253 | 276.3571 | 231 | 288.8416 | 254 | PDX1 | None | Pancreas | GSM28580 | 21125 | 92950 |
| 288.0467 | 161 | 287.6988 | 181 | 189.0906 | 208 | 232.9047 | 244 | MED1 | hMSC-TERT | None | GSM31009 | 21847 | 92968 |
| 124.4969 | 69 | 102.0957 | 71 | 62.27412 | 80 | 114.7591 | 110 | MED1 | hMSC-TERT | None | GSM31009 | 9179 | 92969 |
| 399.888 | 200 | 366.0976 | 216 | 191.5477 | 227 | 290.9047 | 288 | MED1 | hMSC-TERT | None | GSM31009 | 25266 | 92970 |
| 125.1589 | 93 | 141.8849 | 111 | 92.27704 | 130 | 143.1762 | 166 | MED1 | hMSC-TERT | None | GSM31009 | 15818 | 92971 |
| 569.9572 | 286 | 577.2511 | 323 | 341.8084 | 354 | 393.812 | 405 | MED1 | hMSC-TERT | None | GSM31009 | 36588 | 92974 |
| 284.4842 | 179 | 351.5858 | 221 | 178.8233 | 232 | 194.8008 | 260 | MED1 | hMSC-TERT | None | GSM31009 | 27341 | 92976 |
| 92.05528 | 44 | 61.87513 | 41 | 37.75584 | 46 | 44.13979 | 53 | MED1 | hMSC-TERT | None | GSM31009 | 5215 | 92977 |
| 1316.817 | 557 | 1386.362 | 767 | 999.4328 | 518 | 971.4825 | 548 | PHF8 | HEK293T | Embryonic | GSM52038 | 12263 | 93 |
| 612.5634 | 317 | 728.8227 | 443 | 380.3892 | 403 | 451.6679 | 466 | CTCF | THP-1 | Blood | GSM31120 | 44216 | 93012 |
| 564.6111 | 307 | 706.7476 | 421 | 370.4237 | 401 | 414.9702 | 454 | CTCF | THP-1 | Blood | GSM31120 | 44839 | 93017 |
| 528.0995 | 288 | 716.2986 | 411 | 365.3262 | 383 | 410.3931 | 434 | CTCF | THP-1 | Blood | GSM31120 | 41694 | 93019 |
| 18.07829 | 9 | 11.18394 | 8 | 10.84344 | 11 | 33.82047 | 20 | TP53 | MCF-7 | Breast | GSM27400 | 1184 | 93105 |
| 77.06406 | 19 | 19.51359 | 10 | 30.69055 | 17 | 82.43236 | 30 | ESR2 | MDA-MB-2 | Breast | GSM29182 | 1144 | 93123 |
| 149.3828 | 86 | 217.7449 | 115 | 109.202 | 117 | 205.0138 | 163 | ESR2 | MDA-MB-2 | Breast | GSM29182 | 12364 | 93126 |
| 288.0764 | 137 | 409.5998 | 181 | 231.1512 | 189 | 278.0351 | 220 | ESR2 | MDA-MB-2 | Breast | GSM29182 | 16696 | 93127 |
| 909.7786 | 632 | 940.8806 | 805 | 652.5181 | 705 | 621.4443 | 749 | CTCF | BCBL1 | None | GSM94171 | 58394 | 9316 |
| 652.7225 | 410 | 710.5978 | 569 | 450.8742 | 541 | 490.1793 | 615 | CTCF | THP-1 | Blood | GSM31120 | 67482 | 93161 |
| 1002.794 | 731 | 1049.974 | 936 | 725.0798 | 811 | 691.4583 | 860 | SMC1A | BCBL1 | None | GSM94171 | 56416 | 9317 |
| 1027.569 | 514 | 1068.414 | 677 | 751.9108 | 521 | 729.778 | 564 | SMC1A | LCL | Blood | GSM94121 | 28297 | 9319 |
| 1106.989 | 485 | 1130.95 | 615 | 825.3444 | 473 | 798.4756 | 502 | NIPBL | LCL | Blood | GSM94121 | 19310 | 9320 |
| 934.6138 | 563 | 843.3291 | 563 | 819.9491 | 820 | 867.5851 | 1031 | HOXB13 | LN95 | Prostate | GSM26432 | 40977 | 93209 |
| 26.7304 | 19 | 17.59453 | 18 | 10.19443 | 21 | 51.53584 | 43 | HOXA9 | SEM | None | GSM94032 | 3409 | 9321 |
| 603.8607 | 406 | 621.6843 | 498 | 404.6223 | 548 | 374.8794 | 585 | SMARCA4 | MOLM-13 | None | GSM33260 | 70465 | 93210 |
| 94.60854 | 67 | 184.1854 | 102 | 65.85955 | 92 | 64.18027 | 99 | MEIS1 | SEM | None | GSM94032 | 11506 | 9322 |
| 52.15505 | 34 | 53.87323 | 39 | 42.7611 | 49 | 78.84861 | 68 | MED1 | hMSC-TERT | None | GSM31009 | 5349 | 93227 |
| 57.83202 | 49 | 80.48109 | 64 | 66.26612 | 82 | 74.58383 | 93 | MED1 | hMSC-TERT | None | GSM31009 | 9256 | 93228 |
| 96.82389 | 38 | 73.48877 | 37 | 34.47643 | 36 | 68.56937 | 52 | MED1 | hMSC-TERT | None | GSM31009 | 3739 | 93229 |
| 2.491896 | 6 | 9.691937 | 12 | 4.165131 | 13 | 28.56627 | 29 | MEIS1 | SEM | None | GSM94032 | 2697 | 9324 |
| 536.0536 | 331 | 663.8585 | 439 | 353.2126 | 441 | 347.3535 | 478 | RAD21 | THP-1 | Blood | GSM31120 | 54418 | 93325 |
| 470.542 | 307 | 650.9204 | 411 | 345.4149 | 427 | 380.9402 | 481 | RAD21 | THP-1 | Blood | GSM31120 | 52347 | 93326 |
| 526.1302 | 304 | 672.3011 | 398 | 327.6363 | 393 | 323.6137 | 426 | RAD21 | THP-1 | Blood | GSM31120 | 46738 | 93327 |
| 600.1099 | 323 | 667.2246 | 398 | 343.3387 | 402 | 360.5058 | 445 | RAD21 | THP-1 | Blood | GSM31120 | 47273 | 93328 |
| 608.1848 | 350 | 670.0619 | 450 | 359.056 | 446 | 411.2534 | 509 | RAD21 | THP-1 | Blood | GSM31120 | 55156 | 93329 |
| 429.0587 | 253 | 600.6152 | 329 | 272.6757 | 329 | 277.4597 | 360 | RAD21 | THP-1 | Blood | GSM31120 | 38539 | 93330 |
| 433.7587 | 246 | 610.6984 | 321 | 272.5349 | 317 | 310.5513 | 361 | RAD21 | THP-1 | Blood | GSM31120 | 36320 | 93331 |
| 398.6554 | 255 | 602.1902 | 342 | 257.2794 | 337 | 315.9688 | 394 | RAD21 | THP-1 | Blood | GSM31120 | 41491 | 93332 |
| 412.8355 | 262 | 525.6268 | 327 | 236.7224 | 332 | 257.0815 | 372 | RAD21 | THP-1 | Blood | GSM31120 | 42314 | 93333 |
| 495.771 | 310 | 657.446 | 412 | 332.2786 | 416 | 347.1221 | 460 | RAD21 | THP-1 | Blood | GSM31120 | 51302 | 93334 |
| 1054.989 | 242 | 1077.211 | 317 | 735.654 | 237 | 709.4965 | 248 | ETS1 | None | Umbilical V | GSM29474 | 9106 | 93394 |
| 976.8988 | 372 | 1004.392 | 486 | 717.3049 | 371 | 698.8181 | 400 | ETS1 | None | Umbilical V | GSM29474 | 20340 | 93396 |

| | | | | | | | | | | | | | |
|---|---|---|---|---|---|---|---|---|---|---|---|---|---|
| 942.4791 | 590 | 969.383 | 750 | 698.7946 | 655 | 662.5106 | 684 | RAD21 | T47D | Mammary | GSM34155 | 41598 | 93397 |
| 972.2313 | 300 | 984.1187 | 372 | 740.6497 | 309 | 676.0098 | 310 | RAD21 | T47D | Mammary | GSM34155 | 14304 | 93399 |
| 17.72343 | 49 | 25.59825 | 64 | 11.24162 | 76 | 29.96387 | 108 | BATF3 | KK1 | None | GSM24816 | 18006 | 93421 |
| 5.161478 | 129 | 13.51308 | 181 | 2.778534 | 230 | 6.306477 | 283 | BATF3 | ST1 | None | GSM24816 | 87672 | 93422 |
| 6.888543 | 28 | 9.398049 | 36 | 2.260103 | 40 | 16.99217 | 70 | IRF4 | ST1 | None | GSM24816 | 12331 | 93423 |
| 407.9144 | 255 | 543.0986 | 324 | 174.6375 | 290 | 203.7896 | 333 | BATF3 | ST1 | None | GSM24816 | 38973 | 93424 |
| 619.6891 | 364 | 673.8375 | 474 | 299.2616 | 434 | 301.3234 | 475 | NKX2-2 | A673 | Muscle | GSM28575 | 57449 | 93479 |
| 446.5979 | 206 | 584.4708 | 261 | 218.2603 | 233 | 223.7473 | 255 | FLI1 | A673 | Muscle | GSM28575 | 24565 | 93480 |
| 547.9598 | 246 | 750.8109 | 339 | 298.0589 | 290 | 352.5966 | 335 | SOX2 | A673 | Muscle | GSM28575 | 29333 | 93484 |
| 450.0501 | 213 | 709.5437 | 294 | 226.1044 | 245 | 243.1497 | 273 | SOX2 | A673 | Muscle | GSM28575 | 26291 | 93485 |
| 230.9439 | 56 | 206.8776 | 58 | 134.1065 | 60 | 425.6836 | 116 | HIF1A | HKC8 | Kidney | GSM34178 | 3028 | 93506 |
| 134.4058 | 34 | 137.9881 | 38 | 93.33324 | 40 | 261.7511 | 73 | HIF1A | HKC8 | Kidney | GSM34178 | 1945 | 93510 |
| 595.2084 | 203 | 648.1403 | 235 | 270.7697 | 209 | 271.7381 | 226 | TRIM24 | MOLM-13 | None | GSM26873 | 16679 | 93511 |
| 425.5294 | 200 | 599.7974 | 262 | 286.3043 | 255 | 384.2034 | 310 | CTCF | LCL | Blood | GSM32129 | 23342 | 93518 |
| 1151.061 | 531 | 1198.952 | 704 | 851.3287 | 499 | 817.522 | 521 | RBP2 | THP-1 | Blood | GSM31120 | 17585 | 93550 |
| 363.5245 | 68 | 403.2346 | 79 | 149.6934 | 59 | 256.9415 | 83 | BRD4 | H2171 | Lung | GSM27167 | 2733 | 93570 |
| 914.5795 | 527 | 922.346 | 645 | 648.4528 | 538 | 609.4751 | 557 | BRD4 | MV4-11 | Blood | GSM27167 | 37626 | 93576 |
| 609.6578 | 93 | 1120.328 | 146 | 264.5567 | 79 | 430.1309 | 109 | NRF1 | HeLa | None | GSM28723 | 2567 | 93597 |
| 590.5508 | 85 | 1048.3 | 131 | 228.1652 | 67 | 396.1395 | 96 | NRF1 | HeLa | None | GSM28723 | 2144 | 93598 |
| 620.7849 | 94 | 1157.398 | 149 | 314.9001 | 87 | 372.1677 | 101 | NRF1 | HeLa | None | GSM28723 | 2628 | 93599 |
| 694.9239 | 102 | 1187.939 | 155 | 299.029 | 86 | 436.9638 | 112 | NRF1 | HeLa | None | GSM28723 | 2729 | 93600 |
| 819.6284 | 125 | 1193.108 | 200 | 404.3155 | 114 | 617.3487 | 152 | NRF1 | HeLa | None | GSM28723 | 3449 | 93601 |
| 1190.01 | 348 | 1204.176 | 440 | 910.6727 | 324 | 892.8595 | 349 | BRD2 | H23 | Lung | GSM28011 | 8819 | 93605 |
| 1062.691 | 146 | 1157.612 | 180 | 470.1207 | 124 | 507.1903 | 137 | ELL2 | 293T | Embryonic | GSM33605 | 3563 | 93636 |
| 30.99742 | 61 | 66.3899 | 92 | 20.77619 | 93 | 68.6146 | 145 | CEBPB | THP-1 | Blood | GSM31120 | 19599 | 93666 |
| 1209.161 | 447 | 1223.29 | 559 | 901.6665 | 400 | 877.1757 | 425 | RBP2 | THP-1 | Blood | GSM31120 | 11259 | 93667 |
| 503.9793 | 274 | 702.6335 | 378 | 367.2326 | 371 | 441.3506 | 431 | CTCF | THP-1 | Blood | GSM31120 | 39326 | 93668 |
| 474.1174 | 247 | 709.5786 | 340 | 309.1701 | 319 | 375.3654 | 372 | CTCF | THP-1 | Blood | GSM31120 | 33927 | 93669 |
| 687.0682 | 379 | 713.8911 | 492 | 460.84 | 482 | 511.2171 | 544 | RAD21 | THP-1 | Blood | GSM31120 | 54186 | 93670 |
| 1181.167 | 476 | 1235.977 | 639 | 889.6043 | 446 | 865.7434 | 473 | RBP2 | THP-1 | Blood | GSM31120 | 13301 | 93671 |
| 1167.104 | 372 | 1180.616 | 466 | 879.7804 | 334 | 853.5576 | 352 | RBP2 | THP-1 | Blood | GSM31120 | 9621 | 93672 |
| 1181.168 | 408 | 1199.082 | 513 | 876.8931 | 359 | 864.4339 | 392 | RBP2 | THP-1 | Blood | GSM31120 | 10759 | 93673 |
| 467.1984 | 58 | 555.9048 | 70 | 376.3355 | 68 | 523.4543 | 87 | TRIM24 | LNCaP | Prostate | GSM28911 | 1228 | 93733 |
| 665.1332 | 92 | 744.7933 | 107 | 313.3072 | 81 | 624.1319 | 126 | TRIM24 | LNCaP | Prostate | GSM28911 | 2351 | 93734 |
| 1032.449 | 569 | 988.02 | 631 | 783.8561 | 628 | 787.8842 | 722 | TRIM28 | LNCaP | Prostate | GSM28911 | 31885 | 93735 |
| 970.4634 | 297 | 920.4875 | 322 | 731.5413 | 310 | 787.5927 | 413 | TRIM28 | LNCaP | Prostate | GSM28911 | 15940 | 93736 |
| 635.383 | 99 | 667.0018 | 111 | 376.1625 | 101 | 995.7131 | 184 | AR | LNCaP | Prostate | GSM28911 | 3087 | 93737 |
| 658.9328 | 96 | 654.7883 | 104 | 376.5632 | 95 | 861.4065 | 160 | AR | LNCaP | Prostate | GSM28911 | 2691 | 93738 |
| 80.02126 | 37 | 146.4672 | 56 | 79.89109 | 56 | 70.28753 | 57 | TP53 | None | None | GSM29889 | 4341 | 93743 |
| 80.98324 | 26 | 92.60429 | 31 | 42.72002 | 28 | 137.7842 | 53 | TP53 | None | None | GSM29889 | 2080 | 93744 |
| 78.25166 | 248 | 75.56537 | 288 | 85.6604 | 446 | 88.95211 | 497 | CUL4A | U2OS | None | GSM31477 | 108184 | 93778 |
| 0.355594 | 120 | 1.616712 | 166 | 9.4105 | 331 | 33.19474 | 446 | SKP2 | HCT-116 | Colon | GSM31477 | 119409 | 93779 |
| 3.180489 | 133 | 4.979068 | 171 | 9.880595 | 302 | 22.80651 | 382 | DTL | U2OS | None | GSM31477 | 106548 | 93780 |
| 2.982949 | 136 | 3.620241 | 169 | 19.42025 | 343 | 43.50338 | 439 | SKP2 | HCT-116 | Colon | GSM31477 | 109335 | 93781 |
| 730.0298 | 628 | 689.5194 | 703 | 599.4633 | 899 | 594.8302 | 1021 | FOXA1 | TAM_R | Mammary | GSM30964 | 94736 | 93862 |
| 762.0129 | 510 | 704.4051 | 548 | 625.5384 | 713 | 620.5411 | 805 | FOXA1 | TAM_R | Mammary | GSM30964 | 62046 | 93863 |
| 736.5075 | 331 | 782.1649 | 452 | 459.7351 | 397 | 424.9592 | 418 | CTCF | T47D | Mammary | GSM30451 | 36960 | 93878 |
| 743.2424 | 332 | 793.2107 | 461 | 467.7913 | 395 | 450.1816 | 422 | CTCF | T47D | Mammary | GSM30451 | 36304 | 93879 |
| 707.4278 | 447 | 756.5824 | 604 | 526.521 | 558 | 507.1699 | 607 | CTCF | T47D | Mammary | GSM30451 | 60965 | 93880 |
| 364.5901 | 48 | 376.6487 | 53 | 148.8388 | 38 | 203.7475 | 48 | BRD4 | MDA-MB-2 | Breast | GSM27361 | 1036 | 93920 |
| 1201.258 | 448 | 1246.044 | 584 | 908.4546 | 400 | 872.7869 | 413 | TOP1 | HEK293 | Embryonic | GSM27383 | 10602 | 93994 |
| 621.2511 | 352 | 715.0975 | 499 | 408.5127 | 465 | 513.5307 | 552 | CTCF | HeLa | None | GSM28723 | 55090 | 94009 |
| 659.2309 | 376 | 707.6633 | 517 | 415.7744 | 488 | 498.8308 | 566 | CTCF | HeLa | None | GSM28723 | 58949 | 94010 |
| 639.7034 | 357 | 698.5364 | 480 | 413.8956 | 468 | 515.5249 | 556 | CTCF | HeLa | None | GSM28723 | 55165 | 94011 |
| 661.7055 | 366 | 716.6083 | 508 | 415.3898 | 475 | 503.6859 | 553 | CTCF | HeLa | None | GSM28723 | 56713 | 94012 |
| 653.9878 | 348 | 726.0817 | 485 | 421.4963 | 453 | 509.4573 | 527 | CTCF | HeLa | None | GSM28723 | 51808 | 94013 |
| 10.47798 | 15 | 1.347219 | 9 | 8.70266 | 24 | 63.84564 | 55 | FOS | THP-1 | Blood | GSM31121 | 4425 | 94016 |
| 873.743 | 254 | 865.9654 | 308 | 424.0835 | 253 | 534.0571 | 301 | GLI2 | THP-1 | Blood | GSM31121 | 17406 | 94017 |
| 229.2287 | 278 | 399.9175 | 386 | 194.6641 | 423 | 208.9942 | 474 | RAD21 | HUES64 | Embryo | GSM28945 | 64508 | 94021 |
| 254.9702 | 47 | 428.6368 | 70 | 156.0613 | 50 | 324.7061 | 79 | STAG2 | MCF-10A | Breast | GSM29422 | 1891 | 94041 |
| 302.6407 | 118 | 368.2392 | 145 | 152.0009 | 131 | 227.0831 | 167 | STAG1 | MCF-10A | Breast | GSM29422 | 12147 | 94042 |
| 434.8861 | 89 | 707.0643 | 129 | 290.1855 | 101 | 287.4033 | 107 | STAG2 | MCF-10A | Breast | GSM29422 | 4180 | 94043 |
| 537.9248 | 215 | 771.648 | 289 | 317.5515 | 258 | 397.2257 | 305 | STAG2 | MCF-10A | Breast | GSM29422 | 23496 | 94044 |
| 302.9159 | 112 | 344.112 | 133 | 160.8313 | 126 | 213.3361 | 153 | STAG1 | MCF-10A | Breast | GSM29422 | 10890 | 94046 |
| 523.4282 | 203 | 736.5843 | 267 | 304.9912 | 241 | 384.7166 | 286 | STAG2 | MCF-10A | Breast | GSM29422 | 21436 | 94047 |
| 1041.508 | 160 | 1147.872 | 228 | 539.1317 | 152 | 648.6556 | 178 | EZH2 | LNCaP | Prostate | GSM28790 | 4897 | 94049 |
| 1168.482 | 384 | 1204.71 | 505 | 897.2657 | 369 | 863.2255 | 383 | EZH2 | LNCaP | Prostate | GSM28790 | 11019 | 94056 |
| 53.93744 | 27 | 134.3243 | 48 | 73.13545 | 48 | 133.1606 | 68 | TSC22D3 | MCF-10A | Breast | GSM26755 | 3746 | 94061 |
| 1194.379 | 419 | 1216.083 | 533 | 882.424 | 368 | 880.3884 | 413 | SETD7 | HES-3 | Embryo | GSM28791 | 11270 | 94087 |
| 1085.942 | 238 | 1014.404 | 241 | 834.5916 | 239 | 833.2606 | 262 | FOXA2 | None | Endometri | GSM34397 | 7841 | 94090 |
| 1109.451 | 299 | 1089.258 | 349 | 872.4101 | 303 | 804.6603 | 284 | FOXA2 | P388/NVB- | Endometri | GSM34397 | 8977 | 94091 |
| 1135.805 | 197 | 1089.617 | 207 | 757.0461 | 189 | 657.2905 | 186 | FOXA2 | None | Endometri | GSM34397 | 5117 | 94093 |

| | | | | | | | | | | | | | |
|---|---|---|---|---|---|---|---|---|---|---|---|---|---|
| 1026.833 | 365 | 1013.833 | 429 | 785.4491 | 374 | 768.2192 | 404 | FOXA2 | None | Endometri | GSM34397 | 15479 | 94094 |
| 131.3812 | 30 | 52.64157 | 20 | 39.1267 | 23 | 89.48416 | 37 | TP53 | UO31 | None | GSM26773 | 1589 | 94111 |
| 779.4233 | 417 | 715.5947 | 435 | 648.8626 | 586 | 661.4409 | 691 | FOXA1 | TAM_R | Mammary | GSM30964 | 43663 | 94131 |
| 958.744 | 373 | 995.6038 | 492 | 739.2763 | 405 | 714.224 | 432 | RAD21 | T47D | Mammary | GSM30451 | 20200 | 94135 |
| 15.7939 | 35 | 27.69712 | 49 | 34.6532 | 75 | 62.29387 | 100 | NR3C1 | SUP-B15 | None | GSM28717 | 10631 | 94142 |
| 532.7155 | 273 | 705.6098 | 362 | 350.0194 | 352 | 347.216 | 381 | RAD21 | THP-1 | Blood | GSM31120 | 36511 | 94192 |
| 1115.537 | 568 | 1151.328 | 732 | 820.8268 | 540 | 776.2595 | 549 | SS18 | None | None | GSM28876 | 20022 | 94195 |
| 29.39363 | 11 | 32.71718 | 13 | 12.66349 | 11 | 69.33291 | 27 | SMARCA4 | None | None | GSM28166 | 1059 | 94197 |
| 31.22739 | 19 | 23.98734 | 19 | 12.0382 | 20 | 66.6031 | 44 | SMARCA4 | None | None | GSM28166 | 2694 | 94200 |
| 552.7445 | 155 | 659.5282 | 187 | 241.3289 | 147 | 322.2748 | 180 | RNF2 | None | None | GSM28166 | 8611 | 94201 |
| 22.95199 | 35 | 16.59094 | 36 | 9.652889 | 45 | 28.4239 | 68 | POU5F1 | None | None | GSM28166 | 9313 | 94202 |
| 863.7555 | 190 | 963.9538 | 246 | 339.1715 | 165 | 527.3925 | 218 | RNF2 | None | None | GSM28166 | 8163 | 94204 |
| 36.79594 | 26 | 32.9116 | 28 | 21.87874 | 33 | 160.6534 | 83 | HSF1 | None | None | GSM28166 | 4067 | 94207 |
| 28.26593 | 14 | 33.06318 | 17 | 40.03836 | 25 | 78.2152 | 37 | EZH2 | None | Umbilical \ | GSM29474 | 1722 | 94229 |
| 141.192 | 45 | 187.1101 | 58 | 85.19469 | 52 | 173.2848 | 78 | EP300 | None | Umbilical \ | GSM29474 | 3696 | 94230 |
| 119.7892 | 31 | 242.1437 | 51 | 73.96535 | 35 | 116.9159 | 47 | EP300 | None | Umbilical \ | GSM29474 | 2006 | 94233 |
| 24.90137 | 17 | 43.35469 | 25 | 45.39178 | 35 | 70.96732 | 46 | GATA2 | None | Umbilical \ | GSM29474 | 3032 | 94234 |
| 106.1721 | 27 | 107.3468 | 30 | 51.71095 | 27 | 170.5537 | 53 | EP300 | None | Umbilical \ | GSM29474 | 1703 | 94235 |
| 204.3828 | 64 | 333.1473 | 92 | 128.3027 | 75 | 171.6107 | 92 | EP300 | None | Umbilical \ | GSM29474 | 5065 | 94236 |
| 680.6972 | 451 | 702.2746 | 577 | 568.4558 | 674 | 553.4464 | 745 | CTCF | MCF-7 | Breast | GSM33571 | 72094 | 94308 |
| 768.8222 | 515 | 807.3715 | 675 | 590.7154 | 656 | 582.8782 | 738 | CTCF | MCF-7 | Breast | GSM33571 | 61458 | 94309 |
| 754.8301 | 498 | 800.3859 | 661 | 607.4662 | 684 | 579.4368 | 734 | CTCF | MCF-7 | Breast | GSM33571 | 64035 | 94311 |
| 803.828 | 611 | 831.5246 | 776 | 639.4425 | 826 | 622.2938 | 911 | CTCF | MCF-7 | Breast | GSM33571 | 76506 | 94312 |
| 755.9171 | 534 | 791.0041 | 694 | 610.855 | 741 | 598.4816 | 826 | CTCF | MCF-7 | Breast | GSM33571 | 70995 | 94313 |
| 755.9333 | 515 | 815.3649 | 701 | 625.2208 | 736 | 597.9 | 791 | CTCF | MCF-7 | Breast | GSM33571 | 67226 | 94314 |
| 709.9103 | 431 | 749.1063 | 573 | 594.4015 | 635 | 575.4362 | 692 | CTCF | MCF-7 | Breast | GSM33571 | 60501 | 94315 |
| 793.0176 | 412 | 852.2942 | 573 | 597.9371 | 494 | 562.7067 | 514 | CTCF | MCF-7 | Breast | GSM33571 | 42618 | 94316 |
| 751.6648 | 468 | 799.1428 | 630 | 602.7859 | 635 | 586.1519 | 698 | CTCF | MCF-7 | Breast | GSM33571 | 57532 | 94317 |
| 775.1613 | 528 | 813.3299 | 689 | 617.1481 | 707 | 597.9703 | 774 | CTCF | MCF-7 | Breast | GSM33571 | 64125 | 94318 |
| 247.1287 | 43 | 275.6721 | 50 | 106.4807 | 37 | 131.3943 | 44 | SMARCA4 | HS-SY-2 | None | GSM29161 | 1426 | 94347 |
| 233.4266 | 71 | 201.972 | 73 | 214.9968 | 99 | 198.7408 | 102 | SMAD3 | MDA-MB-2 | Breast | GSM24298 | 5319 | 94376 |
| 450.2693 | 65 | 646.999 | 88 | 248.4996 | 62 | 155.4371 | 51 | SMARCC1 | SYO1 | None | GSM28875 | 1612 | 94383 |
| 94.04875 | 66 | 83.77221 | 71 | 59.45454 | 86 | 56.77597 | 92 | SS18 | SYO1 | None | GSM28875 | 10327 | 94384 |
| 932.1723 | 171 | 1014.39 | 209 | 426.0096 | 156 | 455.1377 | 172 | SS18 | Aska | None | GSM28875 | 6347 | 94391 |
| 566.7445 | 104 | 544.558 | 111 | 257.6597 | 94 | 282.661 | 105 | SS18 | Aska | None | GSM28875 | 3805 | 94393 |
| 562.7019 | 75 | 527.1484 | 78 | 267.4562 | 65 | 153.4618 | 51 | SS18 | SYO1 | None | GSM28875 | 1637 | 94395 |
| 157.0934 | 80 | 309.1748 | 124 | 100.2466 | 101 | 212.838 | 149 | CTCF | None | None | GSM28166 | 10054 | 94411 |
| 295.5936 | 262 | 447.8142 | 349 | 192.332 | 360 | 236.3263 | 420 | RAD21 | None | None | GSM28166 | 52815 | 94412 |
| 503.2649 | 328 | 661.2344 | 448 | 364.7533 | 454 | 349.5872 | 488 | RAD21 | None | None | GSM28166 | 55972 | 94413 |
| 279.408 | 288 | 420.5426 | 381 | 190.1614 | 409 | 218.1829 | 467 | RAD21 | None | None | GSM28166 | 64980 | 94414 |
| 268.6078 | 241 | 451.778 | 335 | 189.5475 | 341 | 242.2719 | 403 | RAD21 | None | None | GSM28166 | 49495 | 94415 |
| 172.156 | 43 | 284.9177 | 63 | 122.2344 | 52 | 238.5361 | 78 | ERG | None | Umbilical \ | GSM29474 | 2713 | 94437 |
| 684.9769 | 141 | 886.4456 | 179 | 394.2728 | 150 | 435.4294 | 168 | ERG | None | Umbilical \ | GSM29474 | 7046 | 94438 |
| 123.7599 | 24 | 163.0954 | 31 | 66.3793 | 24 | 138.7679 | 38 | ERG | None | Umbilical \ | GSM29474 | 1073 | 94440 |
| 991.0005 | 456 | 997.8623 | 564 | 731.9487 | 472 | 689.9106 | 480 | ETS1 | None | Umbilical \ | GSM29474 | 26011 | 94441 |
| 84.02971 | 34 | 92.99 | 40 | 130.4628 | 63 | 167.3973 | 76 | JUN | None | Umbilical \ | GSM29474 | 3631 | 94442 |
| 620.7215 | 147 | 816.3814 | 188 | 332.4473 | 154 | 218.6145 | 136 | BRD9 | TTC1240 | None | GSM33260 | 8585 | 94459 |
| 37.03721 | 16 | 45.9286 | 20 | 5.683019 | 11 | 3.335915 | 10 | BRD9 | TTC1240 | None | GSM33260 | 1789 | 94460 |
| 535.0302 | 100 | 731.0384 | 131 | 372.3971 | 114 | 413.6659 | 128 | RARA | None | None | GSM25853 | 4003 | 94496 |
| 1159.517 | 173 | 1174.225 | 218 | 718.6567 | 168 | 690.0048 | 174 | ELL2 | H2171 | Lung | GSM30739 | 4418 | 94505 |
| 894.6617 | 164 | 1051.114 | 223 | 428.8972 | 155 | 438.1186 | 167 | ELL2 | SW1271 | Lung | GSM30739 | 6635 | 94506 |
| 1171.693 | 259 | 1169.749 | 315 | 893.1151 | 231 | 863.745 | 240 | RBP2 | M.M. | None | GSM31119 | 6090 | 94507 |
| 521.6946 | 71 | 770.7383 | 98 | 360.8628 | 77 | 334.9596 | 78 | AFF1 | SW1271 | Lung | GSM30739 | 1728 | 94508 |
| 404.9148 | 51 | 463.5382 | 60 | 129.7986 | 35 | 205.8241 | 48 | AFF4 | H2171 | Lung | GSM30739 | 1029 | 94509 |
| 867.8489 | 133 | 1112.868 | 171 | 472.4678 | 129 | 595.7077 | 155 | AFF1 | H2171 | Lung | GSM30739 | 3919 | 94510 |
| 862.0566 | 124 | 1148.848 | 161 | 496.9155 | 123 | 466.9917 | 126 | AFF1 | SW1271 | Lung | GSM30739 | 3365 | 94511 |
| 909.501 | 120 | 1198.593 | 158 | 479.5315 | 111 | 498.3286 | 120 | AFF4 | SW1271 | Lung | GSM30739 | 2696 | 94512 |
| 612.8299 | 96 | 951.1783 | 136 | 280.7638 | 85 | 227.7508 | 81 | ELL2 | H2171 | Lung | GSM30739 | 2914 | 94513 |
| 888.8303 | 120 | 1051.778 | 144 | 352.0038 | 95 | 553.4026 | 129 | AFF4 | H2171 | Lung | GSM30739 | 2742 | 94514 |
| 588.8108 | 93 | 894.1197 | 130 | 286.1807 | 85 | 359.9743 | 102 | AFF4 | SW1271 | Lung | GSM30739 | 2837 | 94515 |
| 762.5243 | 572 | 809.2448 | 754 | 629.846 | 824 | 611.3394 | 905 | CTCF | MCF-7 | Breast | GSM33571 | 78900 | 94542 |
| 743.8493 | 482 | 777.146 | 629 | 593.4617 | 659 | 574.5453 | 717 | CTCF | MCF-7 | Breast | GSM33571 | 63181 | 94543 |
| 793.3193 | 512 | 827.8392 | 667 | 613.6132 | 651 | 606.341 | 732 | CTCF | MCF-7 | Breast | GSM33571 | 57522 | 94544 |
| 733.156 | 484 | 782.1658 | 653 | 587.6258 | 671 | 587.1855 | 763 | CTCF | MCF-7 | Breast | GSM33571 | 67094 | 94545 |
| 753.6174 | 487 | 780.1644 | 625 | 592.2822 | 649 | 571.4791 | 705 | CTCF | MCF-7 | Breast | GSM33571 | 62389 | 94546 |
| 794.7908 | 553 | 836.5926 | 720 | 625.3152 | 725 | 607.7564 | 798 | CTCF | MCF-7 | Breast | GSM33571 | 64827 | 94547 |
| 635.2355 | 163 | 783.926 | 201 | 362.4845 | 177 | 288.5243 | 171 | BRD9 | Aska | None | GSM33260 | 9975 | 94624 |
| 715.5379 | 115 | 700.954 | 124 | 348.938 | 107 | 893.0924 | 189 | AR | 22RV1 | Prostate | GSM24956 | 3912 | 94663 |
| 118.5316 | 28 | 94.11509 | 27 | 67.40748 | 29 | 105.6958 | 39 | SS18 | HS-SY-2 | None | GSM29160 | 1320 | 94682 |
| 18.72448 | 13 | 32.07126 | 19 | 8.355697 | 15 | 34.77538 | 29 | NFATC1 | None | Blood | GSM28100 | 2259 | 94683 |
| 1013.572 | 343 | 1030.536 | 432 | 734.7272 | 313 | 743.3621 | 366 | IRF4 | None | Blood | GSM28100 | 14978 | 94684 |
| 1087.757 | 315 | 1101.007 | 397 | 833.6248 | 302 | 786.088 | 300 | KDM2B | HS-SY-2 | None | GSM29160 | 9626 | 94685 |

| | | | | | | | | | | | | | |
|---|---|---|---|---|---|---|---|---|---|---|---|---|---|
| 790.6852 | 462 | 810.9057 | 584 | 567.3967 | 526 | 566.6626 | 598 | RAD21 | THP-1 | Blood | GSM31119 | 51088 | 94770 |
| 678.6407 | 415 | 736.011 | 575 | 524.4079 | 563 | 501.3653 | 607 | CTCF | MCF-7 | Breast | GSM33571 | 66268 | 94779 |
| 771.9135 | 536 | 830.2786 | 725 | 618.4826 | 727 | 605.3077 | 807 | CTCF | MCF-7 | Breast | GSM33571 | 66789 | 94780 |
| 155.4487 | 44 | 82.62303 | 35 | 46.79815 | 36 | 39.28119 | 36 | TP53 | U2OS | Bone | GSM30170 | 3021 | 94788 |
| 258.1377 | 93 | 265.9026 | 105 | 280.3909 | 142 | 535.1631 | 207 | TET2 | MCF-7 | Breast | GSM34147 | 8276 | 94814 |
| 351.6609 | 72 | 255.3458 | 66 | 422.3818 | 111 | 722.8047 | 158 | ESR1 | MDA-MB-1 | Mammary | GSM29316 | 3265 | 94818 |
| 797.0537 | 296 | 768.3695 | 335 | 747.4127 | 508 | 785.518 | 640 | ESR1 | MDA-MB-1 | Mammary | GSM29316 | 26960 | 94820 |
| 212.4946 | 45 | 173.1228 | 44 | 327.105 | 80 | 564.7059 | 115 | ESR1 | MDA-MB-1 | Mammary | GSM29316 | 2159 | 94821 |
| 172.0808 | 37 | 112.4556 | 32 | 273.0625 | 67 | 639.6053 | 115 | ESR1 | MDA-MB-1 | Mammary | GSM29316 | 1807 | 94822 |
| 826.0974 | 338 | 798.9904 | 385 | 732.4442 | 520 | 787.6081 | 683 | ESR1 | MDA-MB-1 | Mammary | GSM29316 | 29132 | 94829 |
| 707.2577 | 219 | 858.2531 | 295 | 407.083 | 245 | 397.2341 | 261 | CTCF | T47D | Mammary | GSM34155 | 16070 | 94832 |
| 809.0389 | 242 | 871.7933 | 327 | 519.1372 | 283 | 456.8843 | 288 | CTCF | T47D | Mammary | GSM34155 | 17225 | 94833 |
| 804.7537 | 387 | 853.4117 | 527 | 611.6917 | 453 | 579.8661 | 476 | CTCF | T47D | Mammary | GSM34155 | 34264 | 94834 |
| 844.7414 | 383 | 879.6693 | 504 | 629.2789 | 420 | 599.1736 | 441 | CTCF | T47D | Mammary | GSM34155 | 29647 | 94835 |
| 829.0292 | 343 | 854.4172 | 448 | 603.1467 | 383 | 573.1959 | 405 | CTCF | T47D | Mammary | GSM34155 | 27596 | 94836 |
| 812.6433 | 417 | 860.8516 | 567 | 600.9025 | 474 | 580.7344 | 513 | CTCF | T47D | Mammary | GSM34155 | 37450 | 94837 |
| 0.11257 | 5 | 0.600396 | 8 | 1.515796 | 16 | 0.105761 | 12 | CTCF | HepG2 | Liver | GSM30199 | 4670 | 94850 |
| 7.235523 | 7 | 5.211491 | 7 | 0 | 3 | 6.130521 | 12 | CTCF | HepG2 | Liver | GSM30199 | 1727 | 94853 |
| 1338.774 | 200 | 1360.613 | 250 | 863.6821 | 153 | 1013.093 | 177 | CDK9 | T47D | Mammary | GSM31250 | 2293 | 94867 |
| 1255.308 | 851 | 1227.358 | 979 | 915.7263 | 870 | 908.4324 | 976 | BRD4 | 22RV1 | Prostate | GSM24956 | 31322 | 94880 |
| 1251.299 | 513 | 1287.063 | 661 | 930.6073 | 449 | 913.6528 | 491 | ZFX | 22RV1 | Prostate | GSM24956 | 12222 | 94881 |
| 1237.481 | 417 | 1266.136 | 538 | 940.1485 | 376 | 921.4085 | 408 | ZFX | 22RV1 | Prostate | GSM24956 | 9872 | 94882 |
| 621.3277 | 293 | 754.0473 | 415 | 382.0276 | 370 | 512.4692 | 448 | CTCF | None | Breast | GSM30172 | 39916 | 94885 |
| 1177.238 | 193 | 1183.877 | 239 | 625.8091 | 161 | 755.8658 | 189 | PKNOX1 | HeLa | Cervix | GSM27153 | 4670 | 94906 |
| 1177.867 | 470 | 1179.925 | 570 | 884.8277 | 424 | 821.4464 | 408 | BRD4 | None | None | GSM29425 | 12028 | 94941 |
| 1183.789 | 484 | 1200.368 | 596 | 907.0435 | 459 | 840.7833 | 435 | BRD4 | None | None | GSM29425 | 12137 | 94942 |
| 1196.063 | 303 | 1197.382 | 367 | 937.797 | 286 | 856.0261 | 258 | BRD4 | None | None | GSM29425 | 6712 | 94943 |
| 1175.767 | 471 | 1194.308 | 592 | 891.9966 | 437 | 822.9062 | 415 | BRD4 | None | None | GSM29425 | 12111 | 94944 |
| 1190.981 | 246 | 1224.674 | 324 | 941.4354 | 237 | 866.5267 | 224 | BRD4 | None | None | GSM29425 | 5437 | 94948 |
| 154.8195 | 53 | 227.4691 | 72 | 71.94235 | 55 | 77.00159 | 61 | ZNF366 | None | None | GSM20812 | 4602 | 94951 |
| 129.6438 | 34 | 263.8758 | 56 | 87.83019 | 40 | 171.8416 | 60 | HIF1A | None | Umbilical V | GSM95597 | 2134 | 9496 |
| 904.8358 | 113 | 800.4462 | 113 | 387.3149 | 90 | 445.1865 | 103 | ONECUT2 | 22RV1 | Prostate | GSM25719 | 1889 | 94983 |
| 793.5157 | 90 | 694.1158 | 89 | 354.4835 | 72 | 370.2496 | 78 | ONECUT2 | 22RV1 | Prostate | GSM25719 | 1276 | 94984 |
| 847.1426 | 529 | 912.7566 | 725 | 646.1925 | 644 | 596.1568 | 653 | CTCF | MCF-7 | Breast | GSM33571 | 51294 | 95016 |
| 757.7726 | 507 | 786.2269 | 652 | 605.3001 | 689 | 586.3095 | 754 | CTCF | MCF-7 | Breast | GSM33571 | 65125 | 95017 |
| 743.6246 | 451 | 771.2207 | 584 | 587.9839 | 599 | 562.38 | 641 | CTCF | MCF-7 | Breast | GSM33571 | 54350 | 95018 |
| 760.9602 | 497 | 786.3505 | 633 | 610.124 | 673 | 580.7165 | 717 | CTCF | MCF-7 | Breast | GSM33571 | 59885 | 95019 |
| 734.4187 | 492 | 773.6711 | 646 | 553.7573 | 623 | 522.0338 | 661 | CTCF | MCF-7 | Breast | GSM33571 | 66772 | 95020 |
| 749.1843 | 522 | 798.5144 | 696 | 595.1149 | 710 | 583.0095 | 790 | CTCF | MCF-7 | Breast | GSM33571 | 69171 | 95021 |
| 756.6002 | 515 | 806.3563 | 689 | 606.2281 | 709 | 596.3393 | 792 | CTCF | MCF-7 | Breast | GSM33571 | 67738 | 95022 |
| 749.1163 | 534 | 782.4912 | 691 | 606.553 | 749 | 593.9568 | 834 | CTCF | MCF-7 | Breast | GSM33571 | 73392 | 95023 |
| 782.3675 | 547 | 815.2943 | 707 | 624.7945 | 742 | 593.2021 | 792 | CTCF | MCF-7 | Breast | GSM33571 | 68610 | 95024 |
| 822.2709 | 625 | 858.0784 | 800 | 666.944 | 862 | 647.0772 | 948 | CTCF | MCF-7 | Breast | GSM33571 | 74679 | 95025 |
| 411.3064 | 297 | 489.4261 | 361 | 347.5404 | 441 | 357.1376 | 485 | GATA2 | HDFa | None | GSM26989 | 56692 | 95036 |
| 267.1333 | 179 | 268.1398 | 203 | 205.2696 | 254 | 313.1354 | 322 | GATA2 | HDFa | None | GSM26989 | 30494 | 95037 |
| 44.18056 | 23 | 81.50853 | 35 | 89.31788 | 49 | 139.7149 | 65 | GATA2 | HDFa | None | GSM26989 | 3174 | 95039 |
| 43.39248 | 15 | 49.84198 | 18 | 11.39633 | 12 | 40.23621 | 23 | FOXK1 | HeLa | Cervix | GSM95668 | 1367 | 9505 |
| 46.36432 | 20 | 62.28224 | 26 | 10.45308 | 16 | 54.94826 | 35 | FOXK1 | HeLa | Cervix | GSM95669 | 2194 | 9506 |
| 831.8371 | 592 | 881.3552 | 779 | 720.4006 | 884 | 721.9118 | 1016 | TFAP2C | MCF-7 | Breast | GSM88942 | 67442 | 9507 |
| 1263.232 | 1081 | 1332.874 | 1329 | 960.4403 | 1392 | 907.0255 | 1510 | TFAP2C | BT-474 | Mammary | GSM88942 | 82383 | 9508 |
| 1074.784 | 744 | 1135.789 | 967 | 869.0673 | 946 | 854.8903 | 1060 | TFAP2C | MDA-MB-4 | Mammary | GSM88942 | 46587 | 9509 |
| 1071.597 | 936 | 1096.65 | 1144 | 830.1493 | 1209 | 815.3061 | 1363 | TFAP2C | SKBR-3 | Mammary | GSM88942 | 86559 | 9510 |
| 794.1925 | 521 | 816.3861 | 664 | 545.3044 | 573 | 534.5071 | 637 | NANOG | H1 | Embryo | GSM51837 | 62474 | 955 |
| 354.2653 | 141 | 572.6528 | 199 | 198.9196 | 164 | 264.2543 | 199 | PRDM14 | H1 | Embryo | GSM56308 | 14522 | 966 |
| 4.794209 | 33 | 17.5452 | 55 | 5.773254 | 64 | 31.69938 | 107 | CEBPA | THP-1 | Blood | GSM35190 | 18157 | 99670 |
| 9.448138 | 53 | 26.8996 | 83 | 14.0996 | 106 | 39.64049 | 151 | CEBPA | THP-1 | Blood | GSM35190 | 26443 | 99675 |
| 1.126369 | 16 | 14.68457 | 37 | 3.593063 | 39 | 26.17245 | 72 | CEBPA | THP-1 | Blood | GSM35190 | 11278 | 99678 |
| 6.921473 | 31 | 25.99434 | 54 | 6.747769 | 56 | 35.90285 | 96 | CEBPA | THP-1 | Blood | GSM35190 | 14730 | 99680 |
| 5.364328 | 36 | 17.44923 | 58 | 5.056547 | 66 | 28.09286 | 109 | CEBPA | THP-1 | Blood | GSM35190 | 19354 | 99681 |
| 11.36721 | 59 | 30.61124 | 91 | 14.05322 | 112 | 46.05488 | 166 | CEBPA | THP-1 | Blood | GSM35190 | 28361 | 99682 |
| 9.968279 | 46 | 22.60366 | 68 | 10.06998 | 83 | 36.79206 | 127 | CEBPA | THP-1 | Blood | GSM35190 | 21054 | 99684 |
| 20.44877 | 34 | 45.83197 | 53 | 25.00952 | 61 | 33.04347 | 73 | IRF8 | THP-1 | Blood | GSM35149 | 9913 | 99718 |
| 22.48782 | 94 | 51.35441 | 138 | 28.09185 | 172 | 29.95545 | 193 | CEBPA | THP-1 | Blood | GSM35149 | 37326 | 99720 |
| 585.8671 | 84 | 521.6784 | 85 | 277.7622 | 74 | 455.1412 | 103 | FOXA1 | 22RV1 | Prostate | GSM35081 | 2123 | 99854 |
| 960.5669 | 427 | 883.5231 | 431 | 761.1808 | 500 | 779.3447 | 595 | FOXA1 | 22RV1 | Prostate | GSM35081 | 24470 | 99855 |
| 650.3277 | 89 | 569.2159 | 89 | 407.7158 | 91 | 629.1081 | 123 | FOXA1 | 22RV1 | Prostate | GSM35081 | 2105 | 99856 |
| 117.5601 | 139 | 75.19576 | 136 | 35.29994 | 157 | 44.7065 | 184 | FOXA1 | HEK293 | Embryonic | GSM35081 | 30937 | 99859 |
| 227.2534 | 58 | 154.4902 | 52 | 310.1943 | 97 | 501.2504 | 133 | FOXA1 | HEK293 | Embryonic | GSM35081 | 3440 | 99861 |
| 1304.012 | 185 | 1215.52 | 175 | 1148.625 | 236 | 1232.888 | 332 | TLE3 | C4-2B | Prostate | GSM35081 | 2892 | 99863 |
| 638.3439 | 75 | 470.6465 | 67 | 272.5844 | 60 | 858.3682 | 124 | FOXA1 | 22RV1 | Prostate | GSM35081 | 1367 | 99865 |
| 943.5759 | 524 | 876.0017 | 552 | 750.998 | 644 | 768.7632 | 769 | FOXA1 | 22RV1 | Prostate | GSM35080 | 35937 | 99872 |
| 1078.319 | 174 | 796.0833 | 158 | 644.4454 | 178 | 900.1989 | 233 | FOXA1 | 22RV1 | Prostate | GSM35080 | 5612 | 99877 |

| | | | | | | | | | | | | | |
|---|---|---|---|---|---|---|---|---|---|---|---|---|---|
| 804.4403 | 379 | 737.521 | 393 | 620.7396 | 465 | 644.4021 | 565 | FOXA1 | 22RV1 | Prostate | GSM35080 | 36740 | 99878 |
| 984.2432 | 224 | 887.6816 | 216 | 825.2413 | 280 | 873.0296 | 368 | FOXA1 | LAPC4 | None | GSM35080 | 10131 | 99881 |
| 635.38 | 399 | 367.463 | 368 | 431.6144 | 539 | 516.6892 | 666 | FOXA1 | LAPC4 | None | GSM35080 | 68171 | 99882 |
| 1081.897 | 400 | 997.232 | 394 | 928.6293 | 530 | 1000.377 | 717 | FOXA1 | C4-2B | Prostate | GSM35080 | 15740 | 99885 |
| 985.4139 | 571 | 889.7344 | 565 | 801.6147 | 724 | 868.565 | 955 | FOXA1 | LNCaP | Prostate | GSM35080 | 36575 | 99888 |
| 1089.837 | 155 | 1040.823 | 164 | 1006.428 | 214 | 1091.493 | 308 | FOXA1 | LAPC4 | None | GSM35080 | 4219 | 99890 |
| 544.968 | 306 | 696.0675 | 417 | 382.3779 | 408 | 385.404 | 445 | CTCF | A549 | Lung | GSM24378 | 43814 | 99898 |
| 964.091 | 277 | 921.9404 | 300 | 752.3862 | 302 | 733.4936 | 326 | JUN | A549 | Lung | GSM24378 | 13610 | 99903 |
| 151.2255 | 103 | 118.2611 | 105 | 129.4679 | 150 | 246.4504 | 209 | JUNB | A549 | Lung | GSM24378 | 15958 | 99904 |
| 104.8578 | 69 | 108.38 | 79 | 115.8876 | 111 | 222.0016 | 157 | JUNB | A549 | Lung | GSM24378 | 10436 | 99905 |
| 95.77625 | 67 | 84.98277 | 72 | 95.31949 | 104 | 249.3802 | 167 | JUNB | A549 | Lung | GSM24378 | 10688 | 99906 |
| 95.49971 | 69 | 73.63289 | 70 | 88.54894 | 104 | 209.2792 | 159 | JUNB | A549 | Lung | GSM24378 | 10980 | 99907 |
| 1001.183 | 209 | 990.7505 | 249 | 632.4052 | 218 | 656.991 | 237 | JUN | A549 | Lung | GSM24378 | 8567 | 99908 |
| 1022.049 | 215 | 1015.182 | 257 | 609.6174 | 210 | 753.023 | 249 | JUN | A549 | Lung | GSM24378 | 8128 | 99909 |
| 1057.889 | 182 | 1049.501 | 213 | 697.4539 | 195 | 741.5825 | 214 | JUN | A549 | Lung | GSM24378 | 6062 | 99910 |
| 776.2993 | 194 | 813.342 | 219 | 783.1794 | 279 | 820.817 | 352 | ZNF843 | HEK293 | Embryonic | GSM28271 | 10976 | 99914 |
| 902.1148 | 481 | 953.6307 | 647 | 689.4667 | 562 | 681.9239 | 627 | CTCF | None | Pancreas | GSM28275 | 37424 | 99926 |
| 176.6397 | 135 | 162.8462 | 148 | 157.2732 | 202 | 212.6279 | 245 | CBFA2T2 | K562 | None | GSM28275 | 23237 | 99930 |
| 645.9219 | 292 | 747.7225 | 393 | 481.6243 | 390 | 486.5156 | 424 | RAD21 | A549 | Lung | GSM31064 | 35558 | 99945 |
| 645.6307 | 284 | 726.578 | 357 | 425.0316 | 359 | 457.4374 | 400 | RAD21 | A549 | Lung | GSM31064 | 33025 | 99946 |
| 593.7751 | 264 | 735.8262 | 345 | 374.0043 | 328 | 454.2029 | 383 | RAD21 | A549 | Lung | GSM31064 | 30692 | 99947 |
| 745.265 | 354 | 774.225 | 463 | 586.1163 | 460 | 581.1582 | 518 | RAD21 | A549 | Lung | GSM31064 | 39431 | 99948 |
| 797.2194 | 467 | 809.7655 | 580 | 611.797 | 574 | 607.315 | 646 | RAD21 | A549 | Lung | GSM31064 | 47049 | 99949 |
| 943.0734 | 688 | 949.436 | 835 | 683.3595 | 773 | 673.734 | 866 | RAD21 | A549 | Lung | GSM31064 | 58083 | 99950 |
| 638.6597 | 292 | 738.4544 | 387 | 440.3621 | 377 | 515.4174 | 434 | SMC3 | A549 | Lung | GSM31064 | 34807 | 99951 |
| 643.006 | 280 | 755.9149 | 377 | 462.1123 | 366 | 559.33 | 427 | SMC3 | A549 | Lung | GSM31064 | 32335 | 99952 |
| 581.1524 | 266 | 754.8717 | 372 | 439.8833 | 358 | 511.5565 | 411 | SMC3 | A549 | Lung | GSM31064 | 32692 | 99953 |
| 713.6619 | 297 | 760.2001 | 389 | 466.9943 | 373 | 568.1918 | 436 | SMC3 | A549 | Lung | GSM31064 | 33475 | 99954 |
| 491.1542 | 224 | 748.6081 | 312 | 316.2018 | 281 | 395.5071 | 332 | SMC3 | A549 | Lung | GSM31064 | 26267 | 99955 |
| 569.6345 | 251 | 743.2965 | 335 | 377.9234 | 318 | 427.0892 | 361 | SMC3 | A549 | Lung | GSM31064 | 29015 | 99956 |
| 522.3347 | 248 | 713.1604 | 321 | 367.524 | 323 | 436.6369 | 374 | RAD21 | A549 | Lung | GSM31064 | 29760 | 99957 |
| 484.3481 | 229 | 704.234 | 304 | 285.5831 | 279 | 390.5231 | 341 | RAD21 | A549 | Lung | GSM31064 | 27700 | 99958 |
| 593.5125 | 266 | 736.1 | 350 | 396.0903 | 339 | 433.1779 | 380 | RAD21 | A549 | Lung | GSM31064 | 31527 | 99959 |
| 725.1366 | 337 | 755.4579 | 441 | 563.9647 | 445 | 569.4359 | 505 | SMC3 | A549 | Lung | GSM31064 | 41119 | 99960 |
| 793.98 | 438 | 817.2732 | 562 | 602.7514 | 529 | 604.5215 | 609 | SMC3 | A549 | Lung | GSM31064 | 44921 | 99961 |
| 599.2387 | 274 | 726.0155 | 353 | 431.5123 | 360 | 452.2349 | 397 | SMC3 | A549 | Lung | GSM31064 | 32749 | 99962 |
| 564.2673 | 259 | 728.1685 | 341 | 399.2412 | 339 | 433.7791 | 379 | RAD21 | A549 | Lung | GSM31064 | 31526 | 99963 |
| 436.2218 | 220 | 617.0051 | 288 | 307.6358 | 288 | 358.6218 | 331 | RAD21 | A549 | Lung | GSM31064 | 27237 | 99964 |
| 736.1482 | 309 | 765.7566 | 404 | 459.6884 | 376 | 567.1649 | 442 | RAD21 | A549 | Lung | GSM31063 | 33293 | 99965 |
| 733.5525 | 368 | 763.0388 | 483 | 577.0909 | 480 | 585.1707 | 559 | RAD21 | A549 | Lung | GSM31064 | 42282 | 99966 |
| 733.4813 | 338 | 758.9109 | 441 | 560.0992 | 436 | 569.3555 | 492 | RAD21 | A549 | Lung | GSM31064 | 37942 | 99967 |
| 602.8245 | 286 | 731.0023 | 385 | 478.6657 | 393 | 482.3891 | 427 | RAD21 | A549 | Lung | GSM31064 | 36006 | 99968 |
| 475.0673 | 215 | 671.7237 | 282 | 263.4689 | 253 | 355.8724 | 308 | SMC3 | A549 | Lung | GSM31063 | 24312 | 99969 |
| 775.3884 | 316 | 795.3491 | 406 | 544.5094 | 384 | 606.3636 | 454 | SMC3 | A549 | Lung | GSM31063 | 30863 | 99970 |
| 518.0146 | 219 | 690.6856 | 280 | 301.4383 | 262 | 381.2932 | 311 | SMC3 | A549 | Lung | GSM31063 | 24398 | 99971 |
| 762.7139 | 393 | 798.1658 | 517 | 608.9921 | 519 | 591.6927 | 569 | SMC3 | A549 | Lung | GSM31063 | 42914 | 99972 |
| 500.4084 | 248 | 718.9391 | 336 | 363.0263 | 329 | 388.82 | 366 | SMC3 | A549 | Lung | GSM31063 | 31195 | 99974 |
| 125.5138 | 34 | 167.3163 | 44 | 88.77141 | 41 | 130.1303 | 53 | KDM1A | A549 | Lung | GSM27005 | 2202 | 99976 |
| 353.5521 | 63 | 630.5243 | 97 | 214.8137 | 66 | 288.108 | 82 | HDAC2 | A549 | Lung | GSM27005 | 2287 | 99981 |

Table S10

| Cell.Rx | Active_Enhancer | Bivalent_Promoter | Poised_Enhancer | Polycomb | Promoter | Transcribed | Outside_ChromHMM |
|---|---|---|---|---|---|---|---|
| LNCaP.EtOH | 10805 | 1608 | 10181 | 2629 | 14381 | 1253 | 4368 |
| LNCaP.DHT | 10351 | 1702 | 9962 | 2576 | 16543 | 1820 | 2936 |
| C42.EtOH | 9919 | 1212 | 10714 | 2552 | 10798 | 2641 | 18667 |
| C42.DHT | 10960 | 1084 | 11846 | 2510 | 11468 | 2699 | 18059 |

Table S11

| Cell.Rx | ChromHMM | TAD.region | NumberGenes |
|---|---|---|---|
| C42.EtOH | Outside_ChromHMM | Outside.TAD | 19450 |
| C42.DHT | Outside_ChromHMM | Outside.TAD | 19001 |
| LNCaP.DHT | Promoter | Outside.TAD | 13136 |
| LNCaP.EtOH | Promoter | Outside.TAD | 11140 |
| LNCaP.EtOH | Active_Enhancer | Outside.TAD | 8303 |
| C42.DHT | Promoter | Outside.TAD | 8111 |
| C42.DHT | Active_Enhancer | Outside.TAD | 7954 |
| C42.EtOH | Promoter | Outside.TAD | 7821 |
| LNCaP.DHT | Active_Enhancer | Outside.TAD | 7811 |
| LNCaP.EtOH | Poised_Enhancer | Outside.TAD | 7785 |
| LNCaP.DHT | Poised_Enhancer | Outside.TAD | 7683 |
| C42.EtOH | Active_Enhancer | Outside.TAD | 6834 |
| LNCaP.DHT | Promoter | TAD | 5758 |
| LNCaP.EtOH | Promoter | TAD | 5087 |
| LNCaP.EtOH | Active_Enhancer | TAD | 3791 |
| C42.DHT | Transcribed | Outside.TAD | 3680 |
| LNCaP.EtOH | Outside_ChromHMM | Outside.TAD | 3579 |
| LNCaP.DHT | Active_Enhancer | TAD | 3375 |
| LNCaP.EtOH | Poised_Enhancer | TAD | 3345 |
| LNCaP.DHT | Poised_Enhancer | TAD | 3208 |
| C42.EtOH | Transcribed | Outside.TAD | 3179 |
| C42.DHT | Promoter | TAD | 3057 |
| C42.EtOH | Promoter | TAD | 3020 |
| C42.DHT | Active_Enhancer | TAD | 2840 |
| C42.DHT | Poised_Enhancer | Outside.TAD | 2813 |
| C42.EtOH | Active_Enhancer | TAD | 2717 |
| C42.EtOH | Outside_ChromHMM | TAD | 2361 |
| LNCaP.DHT | Outside_ChromHMM | Outside.TAD | 2356 |
| C42.EtOH | Poised_Enhancer | Outside.TAD | 2328 |
| C42.DHT | Outside_ChromHMM | TAD | 2119 |
| LNCaP.EtOH | Polycomb | Outside.TAD | 1951 |
| LNCaP.DHT | Polycomb | Outside.TAD | 1627 |
| LNCaP.DHT | Transcribed | Outside.TAD | 1440 |
| LNCaP.EtOH | Bivalent_Promoter | Outside.TAD | 1199 |
| LNCaP.DHT | Bivalent_Promoter | Outside.TAD | 1187 |
| LNCaP.EtOH | Outside_ChromHMM | TAD | 1167 |
| C42.DHT | Transcribed | TAD | 1149 |
| LNCaP.DHT | Polycomb | TAD | 1079 |
| C42.EtOH | Transcribed | TAD | 977 |
| LNCaP.EtOH | Transcribed | Outside.TAD | 918 |
| C42.DHT | Poised_Enhancer | TAD | 804 |
| C42.EtOH | Poised_Enhancer | TAD | 779 |
| LNCaP.EtOH | Polycomb | TAD | 762 |
| LNCaP.DHT | Outside_ChromHMM | TAD | 620 |
| LNCaP.DHT | Bivalent_Promoter | TAD | 525 |
| C42.EtOH | Polycomb | Outside.TAD | 509 |
| C42.EtOH | Bivalent_Promoter | Outside.TAD | 473 |
| LNCaP.EtOH | Bivalent_Promoter | TAD | 444 |
| LNCaP.DHT | Transcribed | TAD | 383 |
| LNCaP.EtOH | Transcribed | TAD | 369 |
| C42.DHT | Polycomb | Outside.TAD | 268 |
| C42.DHT | Bivalent_Promoter | Outside.TAD | 228 |
| C42.EtOH | Polycomb | TAD | 125 |
| C42.EtOH | Bivalent_Promoter | TAD | 123 |
| C42.DHT | Bivalent_Promoter | TAD | 47 |
| C42.DHT | Polycomb | TAD | 8 |

Table S12

| Cell.Rx | ChromHMM | Methyl.region | NumberGenes |
|---|---|---|---|
| C42.DHT | Outside_ChromHMM | Outside.CpG | 13903 |
| C42.EtOH | Outside_ChromHMM | Outside.CpG | 13870 |
| C42.EtOH | Outside_ChromHMM | CpG.region | 13057 |
| LNCaP.DHT | Promoter | Outside.CpG | 12905 |
| C42.DHT | Outside_ChromHMM | CpG.region | 11760 |
| LNCaP.EtOH | Promoter | Outside.CpG | 11045 |
| LNCaP.EtOH | Active_Enhancer | Outside.CpG | 8749 |
| LNCaP.DHT | Active_Enhancer | Outside.CpG | 8254 |
| LNCaP.EtOH | Poised_Enhancer | Outside.CpG | 7979 |
| LNCaP.DHT | Poised_Enhancer | Outside.CpG | 7416 |
| C42.EtOH | Promoter | Outside.CpG | 7348 |
| C42.DHT | Promoter | CpG.region | 7092 |
| C42.DHT | Active_Enhancer | Outside.CpG | 6863 |
| C42.DHT | Promoter | Outside.CpG | 6746 |
| C42.EtOH | Active_Enhancer | Outside.CpG | 6268 |
| LNCaP.DHT | Promoter | CpG.region | 6238 |
| C42.DHT | Active_Enhancer | CpG.region | 6196 |
| LNCaP.EtOH | Promoter | CpG.region | 5682 |
| C42.EtOH | Active_Enhancer | CpG.region | 4667 |
| C42.EtOH | Promoter | CpG.region | 4663 |
| LNCaP.EtOH | Outside_ChromHMM | Outside.CpG | 3896 |
| LNCaP.DHT | Poised_Enhancer | CpG.region | 3626 |
| C42.DHT | Transcribed | Outside.CpG | 3118 |
| LNCaP.DHT | Active_Enhancer | CpG.region | 3054 |
| LNCaP.EtOH | Poised_Enhancer | CpG.region | 2991 |
| LNCaP.EtOH | Active_Enhancer | CpG.region | 2943 |
| C42.EtOH | Transcribed | Outside.CpG | 2783 |
| LNCaP.DHT | Outside_ChromHMM | Outside.CpG | 2699 |
| C42.DHT | Transcribed | CpG.region | 2617 |
| C42.DHT | Poised_Enhancer | Outside.CpG | 2562 |
| C42.EtOH | Poised_Enhancer | Outside.CpG | 2142 |
| LNCaP.EtOH | Polycomb | Outside.CpG | 2076 |
| C42.EtOH | Transcribed | CpG.region | 2030 |
| LNCaP.DHT | Polycomb | Outside.CpG | 2014 |
| C42.DHT | Poised_Enhancer | CpG.region | 1926 |
| LNCaP.DHT | Transcribed | Outside.CpG | 1399 |
| C42.EtOH | Poised_Enhancer | CpG.region | 1368 |
| LNCaP.DHT | Bivalent_Promoter | Outside.CpG | 1347 |
| LNCaP.EtOH | Bivalent_Promoter | Outside.CpG | 1346 |
| LNCaP.EtOH | Transcribed | Outside.CpG | 844 |
| LNCaP.EtOH | Outside_ChromHMM | CpG.region | 799 |
| LNCaP.EtOH | Polycomb | CpG.region | 636 |
| LNCaP.DHT | Polycomb | CpG.region | 572 |
| C42.EtOH | Polycomb | Outside.CpG | 468 |
| LNCaP.DHT | Transcribed | CpG.region | 457 |
| LNCaP.EtOH | Transcribed | CpG.region | 411 |
| C42.EtOH | Bivalent_Promoter | Outside.CpG | 408 |
| LNCaP.DHT | Bivalent_Promoter | CpG.region | 375 |
| C42.EtOH | Polycomb | CpG.region | 324 |
| C42.EtOH | Bivalent_Promoter | CpG.region | 320 |
| LNCaP.EtOH | Bivalent_Promoter | CpG.region | 281 |
| LNCaP.DHT | Outside_ChromHMM | CpG.region | 251 |
| C42.DHT | Bivalent_Promoter | Outside.CpG | 247 |
| C42.DHT | Polycomb | Outside.CpG | 223 |
| C42.DHT | Polycomb | CpG.region | 85 |
| C42.DHT | Bivalent_Promoter | CpG.region | 35 |
